# Excessive assimilation of ammonium by plastidic glutamine synthetase is a major cause of ammonium toxicity in *Arabidopsis thaliana*

**DOI:** 10.1101/764324

**Authors:** Takushi Hachiya, Jun Inaba, Mayumi Wakazaki, Mayuko Sato, Kiminori Toyooka, Atsuko Miyagi, Maki Kawai-Yamada, Takatoshi Kiba, Alain Gojon, Hitoshi Sakakibara

## Abstract

Plants use nitrate and ammonium in the soil as their main nitrogen sources. Recently, ammonium has attracted attention due to evidence suggesting that, in C_3_ species, an elevated CO_2_ environment inhibits nitrate assimilation. However, high concentrations of ammonium as the sole nitrogen source for plants causes impaired growth, i.e. ammonium toxicity. Although ammonium toxicity has been studied for a long time, the primary cause remains to be elucidated. Here, we show that ammonium assimilation in plastids rather than ammonium accumulation is a primary cause for toxicity. Our genetic screen of ammonium-tolerant Arabidopsis lines with enhanced shoot growth identified plastidic *GLUTAMINE SYNTHETASE 2* (*GLN2*) as the causal gene. Our reciprocal grafting of wild-type and *GLN2* or *GLN1;2*-deficient lines suggested that shoot GLN2 activity results in ammonium toxicity, whilst root GLN1;2 activity prevents it. With exposure to toxic levels of ammonium, the shoot GLN2 reaction produced an abundance of protons within cells, thereby elevating shoot acidity and stimulating expression of acidic stress-responsive genes. Application of an alkaline ammonia solution to the toxic ammonium medium efficiently alleviated the ammonium toxicity with a concomitant reduction in shoot acidity. Consequently, we conclude that a primary cause of ammonium toxicity is acidic stress in the shoot. This fundamental insight provides a framework for enhanced understanding of ammonium toxicity in plants.

## Introduction

Nitrate and ammonium are the main sources of nitrogen (N) for most plants. Recent studies suggest that elevated CO_2_ inhibits nitrate reduction in C_3_ species, such as wheat and *Arabidopsis*, whereas ammonium utilization does not decrease (1). One estimate predicts that only a 1% drop in nitrogen use efficiency could increase worldwide cultivation costs for crops by about $1 billion annually (2). Therefore, increasing ammonium use by crops is an important goal for agriculture as CO_2_ levels rise in the world; however, millimolar concentrations of ammonium as the sole N source causes growth suppression and chlorosis in plants, compared with nitrate (3, 4, 5). This phenomenon is widely known as ammonium toxicity, but the primary cause of impaired growth remains to be identified.

Plants grown in high ammonium conditions show several distinct characteristics from those grown in nitrate (3, 4, 5). These toxic symptoms have evoked several hypotheses about the toxic causes, including futile transmembrane ammonium cycling, deficiencies in inorganic cations and organic acids, impaired hormonal homeostasis, disordered pH regulation, and the uncoupling of photophosphorylation; however, some of the symptoms are not directly associated with growth suppression by ammonium toxicity (6), making it difficult to determine the toxic cause. Several efforts have isolated ammonium-sensitive mutants in *Arabidopsis thaliana* and determined their causative genes (4, 5). *GMP1* is a causal gene whose deficiency causes stunted growth of primary roots under high ammonium conditions (7). Given that GMP1 is crucial for synthesizing GDP-mannose as a substrate for *N*-glycosylation, lack of *N*-glycoproteins could be involved in ammonium hypersensitivity. In accordance with this hypothesis, the ammonium-dependent inhibition of primary root growth was shown to be partly attenuated by the lack of a GDP-mannose pyrophosphohydrolase that hydrolyses GDP-mannose to mannose 1-phosphate and GMP (8). In another study, a genetic screen focusing on severely chlorotic Arabidopsis leaves identified *AMOS1*, a gene encoding a plastid metalloprotease, as a factor for improving ammonium tolerance (9). Transcriptome analysis revealed that an AMOS1-dependent mechanism regulates more than half of the transcriptional changes triggered by toxic levels of ammonium. On the other hand, recent studies found that ammonium toxicity was partly alleviated by deficiencies in EIN2 and EIN3, regulators of ethylene responses, or by the application of ethylene biosynthesis and action inhibitors (10, 11). This suggests that ammonium toxicity would be mediated via the ethylene signaling pathway.

The above-described genetic studies have succeeded in determining molecular components closely associated with ammonium toxicity. Nevertheless, the initial event that triggers ammonium toxicity remains to be identified and characterized. To address this question, we screened ammonium-insensitive Arabidopsis lines that were expected to attenuate toxicity and isolated *ami2*. Interestingly, the defect in *ami2* was downregulation of the *GLUTAMINE SYNTHETASE 2* (*GLN2*) gene encoding an ammonium assimilatory enzyme. We identified that in the presence of toxic levels of ammonium, large levels of proton production, due to excessive primary assimilation of ammonium by GLN2, aggravate the acidic burden and lead to plant toxicity.

## Results

### A Genetic Screen Isolated an Ammonium-Insensitive Mutant

To find ammonium-insensitive lines, a gain-of-function population of the Arabidopsis FOX (full-length cDNA overexpressing) lines (12) was used. An apparent ammonium-insensitive mutant was identified that shows enhanced growth of cotyledons that are greener than wild-type (Col) when grown on 10 mM ammonium as the sole N source; the mutant was named *ammonium-insensitive 2* (*ami2*) (Fig. 1*A*). The fresh weights of *ami2* 11-d-old shoots were approximately double those of Col when grown on ammonium (Fig. 1*B*). In contrast, in media containing 10 mM nitrate or 5 mM ammonium plus 5 mM nitrate, the shoot fresh weights of *ami2* were less than those of Col. In media containing 10 mM ammonium, the percentage increase in fresh weight of *ami2* relative to Col was much larger for shoots (by ca. 110%) than for roots (by ca. 50%) (Fig. 1*C*). The greater shoot growth in *ami2* was reduced in media with lower concentrations of ammonium (0.4, 2 mM), in which the shoot growth of Col was greater than that when grown on media containing 10 mM ammonium (Fig. 1*D*). Moreover, nitrate addition in the presence of 10 mM ammonium attenuated the deficiency in shoot growth more effectively in Col than in *ami2*, decreasing the growth difference in a concentration-dependent manner (*SI Appendix*, Fig. S1 *A* and *B*). A time-course analysis of shoot growth revealed that increased ammonium tolerance of the *ami2* plants compared to Col was significant as soon as 5 d after culture initiation (*SI Appendix*, Fig. S1*C*). These results indicate that ammonium tolerance in *ami2* is manifested specifically under harsh ammonium conditions.

**Fig. 1.**
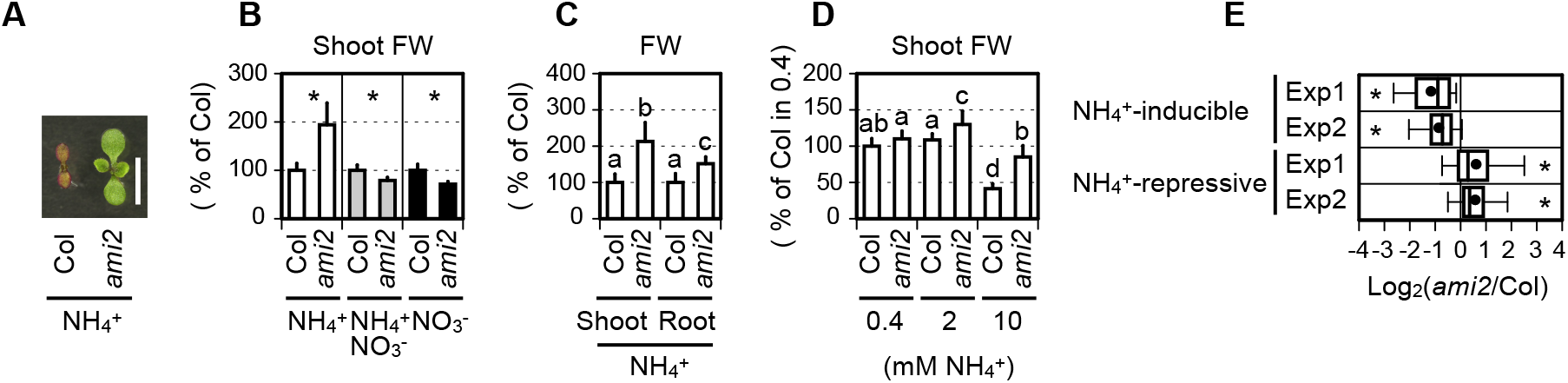
Enhanced shoot growth of *ami2* in the presence of 10 mM ammonium. (*A*) A representative photograph of shoots from the wild-type (Col) and *ami2* grown on media containing 10 mM ammonium for 11 d. The scale bar represents 5 mm. (*B*) Fresh weights (FW) of shoots from Col and *ami2* grown on media containing 10 mM ammonium (mean ± SD; n = 10), 5 mM ammonium nitrate (mean ± SD; n = 5), or 10 mM nitrate (mean ± SD; n = 5) for 11 d. (*C*) FW of shoots and roots from Col and *ami2* grown on media containing 10 mM ammonium for 11 d (mean ± SD; n = 26). (*D*) FW of shoots grown on media containing 0.4, 2, or 10 mM ammonium for 11 d (mean ± SD, n = 5). (*E*) Box plots of the differences in the expression of the ammonium stress-responsive genes between the Col and *ami2* shoots 3 d after transfer to media containing 10 mM ammonium. The gene list was obtained from (9) (For further details, see *Datasets*, Table S1). Two independent experiments (Exp1 and Exp2) were performed. Nine shoots from three plates constituted a single biological replicate. An individual box plot shows the median (heavy vertical line), the 25^th^ to 75^th^ percentiles (right and left sides of the box), the 10^th^ to 90^th^ percentiles (whiskers), and the mean (closed circle). (*B*–*D*) Six shoots from one plate constituted a single biological replicate. (*B*, *E*) Welch’s *t*-test was run at α = 0.05; \**p* < 0.05. (*C*, *D*) Tukey-Kramer’s multiple comparison test was conducted at a significance level of *P* < 0.05 only when a one-way ANOVA was significant at *P* < 0.05. Different letters denote significant differences.

To corroborate this enhanced ammonium tolerance in *ami2*, we performed microarray experiments and compared the expression of genes responsive to toxic levels of ammonium (9) between the Col and *ami2* shoots growing in media containing 10 mM ammonium (Fig. 1*E* and *Datasets*, Table S1). The transcript levels of ammonium-inducible genes were significantly reduced in *ami2* shoots compared with Col shoots, whereas those of ammonium-repressive genes showed the opposite trend. A reverse transcription-quantitative PCR (RT-qPCR) analysis confirmed that expression of *MIOX2* and *PDH2*, two representative ammonium-inducible genes, was more upregulated in the presence of ammonium than in nitrate-containing media in Col, but not in *ami2* (*SI Appendix*, Fig. S2*A*). The expression of a house-keeping gene *TIP41* was less changed (*SI Appendix*, Fig. S2*B*). Collectively, these results indicate that ammonium toxicity is attenuated in *ami2* shoots.

### *GLUTAMINE SYNTHETASE 2* is a Major Causative Gene for Ammonium Toxicity

Next, to identify the causative gene in *ami2*, we recovered the transgene in a vector using specific primers and sequenced the construct. The gene was identified as *GLUTAMINE SYNTHETASE 2* (*GLN2*), the sole plastidic isoform in *A. thaliana* (Fig. 2*A* and *SI Appendix*, Fig. S2*C*). Because the transgene was driven by the cauliflower mosaic virus *35S* promoter, we expected that overexpression of *GLN2* would enhance ammonium tolerance; however, in media containing 10 mM ammonium, the transcript levels of *GLN2* in *ami2* shoots were downregulated to about 5% of those in Col (Fig. 2*B*). In contrast, among the major cytosolic *GLUTAMINE SYNTHETASE* genes (*GLN1s*), *GLN1;1* was upregulated in *ami2* shoots, but *GLN1;2* and *GLN1;3* were slightly downregulated (*SI Appendix*, Fig. S2*D*). Also, an immunoblot analysis using anti-GLN antibodies (13) confirmed that the protein levels of GLN2 were remarkably lower in *ami2* shoots compared with Col, whereas the signal intensities corresponding to GLN1s were comparable between the mutant and wild type (Fig. 2*C*). These findings suggested that overexpression of *GLN2* cDNA would result in a co-suppression event (14), which would make it difficult to test for phenotypic complementation by introducing the *GLN2* transgene. To ensure that reduced expression of *GLN2* enhances ammonium tolerance, we obtained another *GLN2*-deficient line having a T-DNA insertion at the 3′-UTR region of *GLN2* (SALK_051953, designated as *gln2*, *SI Appendix*, Fig. S2*C*). As expected, *gln2* phenocopied *ami2* in terms of the reduced *GLN2* and GLN2 expression (Fig. 2 *B* and *C*), the enhanced ammonium tolerance (Fig. 2 *D* and *E*), and the lowered induction of ammonium-inducible genes when grown on ammonium (*SI Appendix*, Fig. S2*A*). Thus, we concluded that *GLN2* is a causative gene for ammonium toxicity.

**Fig. 2.**
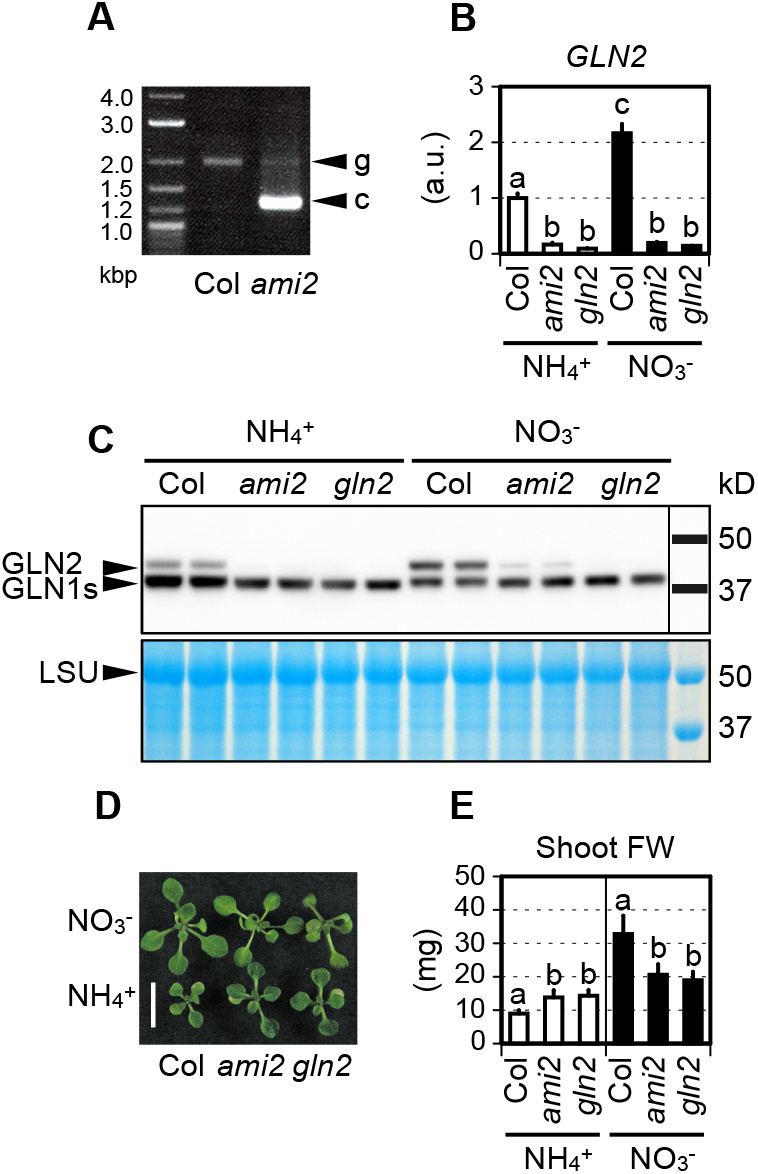
Downregulation of *GLN2* enhances ammonium tolerance. (*A*) Genomic PCR using *GLN2*-specific primers. g and c denote the PCR fragments derived from genomic DNA and cDNA sequences corresponding to *GLN2*, respectively. (*B*) Relative transcript levels of *GLN2* in the shoots of Col, *ami2*, and *gln2* 3 d after transfer to media containing 10 mM ammonium or 10 mM nitrate (mean ± SD; n = 3). Six shoots from two plates constituted a single biological replicate. (*C*) Immunodetection of GLN1s and GLN2 isoproteins using specific antisera raised against maize GLN following SDS-PAGE and immunoblotting of total proteins from the shoots of Col, *ami2*, and *gln2* 5 d after transfer to media containing 10 mM ammonium or 10 mM nitrate. LSU denotes large subunits of RuBisCO. (*D*) A representative photograph of shoots from Col, *ami2*, and *gln2* 7 d after transfer to media containing 10 mM ammonium or 10 mM nitrate. The scale bar represents 10 mm. (*E*) FW of shoots from Col, *ami2*, and *gln2* 7 d after transfer to media containing 10 mM ammonium (mean ± SD; n = 8) or 10 mM nitrate (mean ± SD; n = 5). Mean values of three shoots from one plate constituted a single biological replicate. (*B*, *E*) Tukey-Kramer’s multiple comparison test was conducted at a significance level of *P* < 0.05 only when a one-way ANOVA was significant at *P* < 0.05. Different letters denote significant differences.

### Shoot GLN2 Causes Ammonium Toxicity; Root GLN1;2 Attenuates Ammonium Toxicity

Previous studies had reported that mutants deficient in *AtGLN1;2* were hypersensitive to millimolar concentrations of ammonium (15–17). We also confirmed the ammonium hypersensitivity of *gln1;2-1* and *gln1;2-2* (*SI Appendix*, Fig. S3 *A* and *B*). *GLN2* and *GLN1;2*, therefore, have opposite effects on ammonium toxicity. To discover how GLN2 and GLN1;2 are involved in the toxicity, we evaluated the distribution of *GLN2* and *GLN1;2* expression between shoots and roots in Col plants. In the presence of ammonium or nitrate, the steady-state levels of *GLN2* expression were consistently higher in the shoots than in the roots, whereas expression of *GLN1;2* was much higher in the roots (Fig. 3 *A* and *B* and *SI Appendix*, Fig. S4 *A* and *B*), implying that both shoot GLN2 and root GLN1;2 could affect ammonium toxicity. To support this hypothesis, we performed a growth analysis using reciprocally-grafted plants between Col and *ami2* (Fig. 3*C* and *SI Appendix*, Fig. S4*C*) and between Col and *gln1;2-1* (Fig. 3*D* and *SI Appendix*, Fig. S4*D*). Prior to the analysis, we confirmed that shoot expression of *GLN2* was lower in the *ami2*-derived shoots irrespective of root-genotype (*SI Appendix*, Fig. S5), because *GLN2* mRNA is suggested to be root-to-shoot mobile (18). Only when the scion was derived from *ami2* was shoot growth significantly enhanced in the presence of 10 mM ammonium (Fig. 3*C*). On the other hand, deficiency in root *GLN1;2* content was sufficient to decrease shoot growth in ammonium (Fig. 3*D*). Further, we observed that in ammonium-grown plants, the total enzymatic activities of GLNs were significantly reduced by ca. 30-40% in 5-d-old shoots of *ami2* and *gln2* and by ca. 40-60% in 5-d-old roots of *gln1;2-1* and *gln1;2-2* compared with Col (*SI Appendix*, Fig. S6 *A* and *B*). Additionally, partially compensatory inductions of other *GLNs* were found in the mutants (*SI Appendix*, Fig. S6 *C* and *D*). Our findings demonstrate that although shoot GLN2 causes ammonium toxicity in the shoot, root GLN1;2 attenuates ammonium toxicity.

**Fig. 3.**
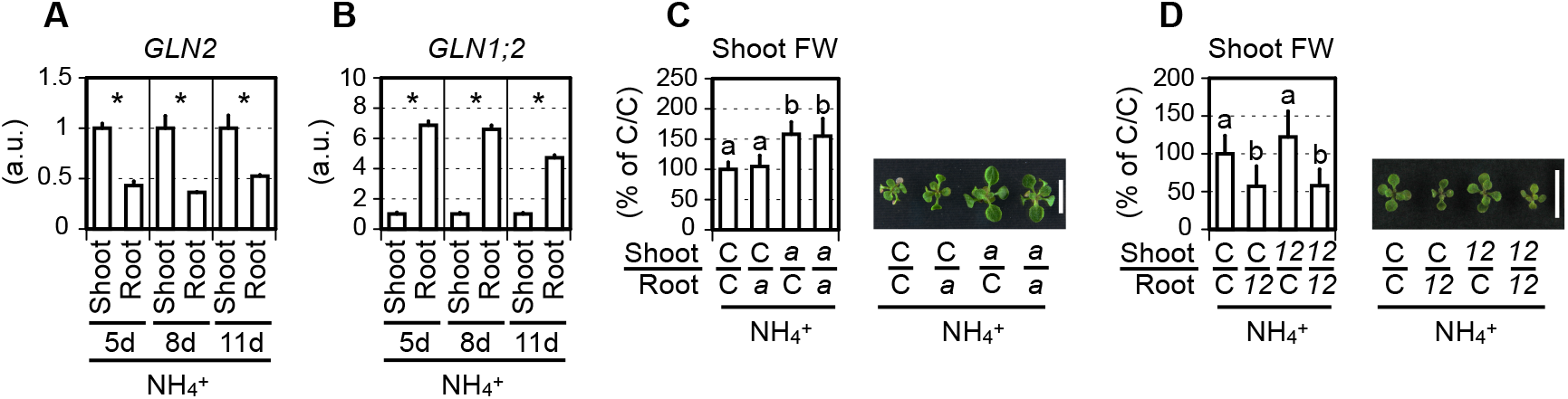
Shoot GLN2 causes ammonium toxicity, whilst root GLN1;2 attenuates ammonium toxicity. (*A*) Relative transcript levels of *GLN2* in the shoots and roots of Col grown on media containing 10 mM ammonium for 5, 8, or 11 d (mean ± SD; n = 3). (*B*) Relative transcript levels of *GLN1;2* in the shoots and roots of Col grown on media containing 10 mM ammonium for 5, 8, or 11 d (mean ± SD; n = 3). (*A*, *B*) Twelve shoots and roots from one plate constituted a single biological replicate. Welch’s *t*-test was run at α = 0.05; \**p* < 0.05. (*C*) FW of shoots from reciprocally-grafted plants between Col (C) and *ami2* (*a*) 7 d after transfer to media containing 10 mM ammonium (mean ± SD; n = 8). (*D*) FW of shoots from reciprocally-grafted plants between Col (C) and *gln1.2-1* 7 d after transfer to media containing 10 mM ammonium (mean ± SD; n = 10). (*C*, *D*) One shoot from one plate constituted a single biological replicate. Tukey-Kramer’s multiple comparison test was conducted at a significance level of *P* < 0.05 only when a one-way ANOVA was significant at *P* < 0.05. Different letters denote significant differences. Representative photograph of shoots 7 d after transfer to media containing 10 mM ammonium are shown. The scale bar represents 10 mm.

### Decreased GLN2 Activity Reduces the Conversion of Ammonium to Amino Acids in Shoots

It is generally held that ammonium *per se* is a toxic compound (19). On the other hand, a deficiency in *GLN2* content should lead to ammonium accumulation in the shoot. Our determination of shoot ammonium content revealed that *ami2* and *gln2* shoots grown on 10 mM ammonium both accumulated more than 100 μmol g^−1^ fresh weight of ammonium (Fig. 4*A*), albeit the two mutants accumulated more fresh weight than Col (*SI Appendix*, Fig. S7). This result indicates that ammonium assimilation by GLN2, rather than ammonium accumulation, triggers ammonium toxicity in the shoot.

**Fig. 4.**
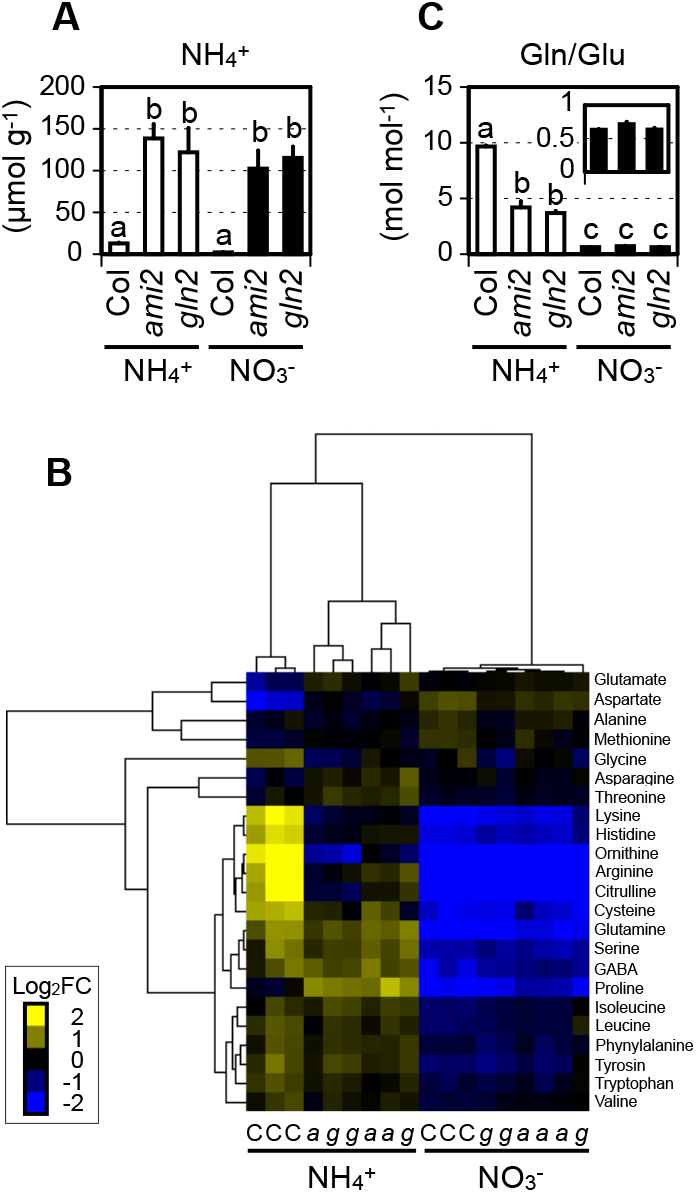
Decreased activity of GLN2 reduces the conversion of ammonium to amino acids in shoots. (*A*) The shoot ammonium content of Col, *ami2*, and *gln2* 5 d after transfer to media containing 10 mM ammonium or 10 mM nitrate (mean ± SD; n = 3). Three shoots from one plate constituted a single biological replicate. (*B*) Hierarchical clustering of the shoot amino acid content of Col (C), *ami2* (*a*), and *gln2* (*g*) 5 d after transfer to media containing 10 mM ammonium or 10 mM nitrate. The color spectrum from yellow to blue corresponds to the relative content of each amino acid. (*C*) The molar ratio of Gln to Glu in the shoots of Col, *ami2*, and *gln2* 5 d after transfer to media containing 10 mM ammonium or 10 mM nitrate (mean ± SE; n = 3). (*A*, *C*) Tukey-Kramer’s multiple comparison test was conducted at a significance level of *P* < 0.05 only when a one-way ANOVA was significant at *P* < 0.05. Different letters denote significant differences. (*B*, *C*) Six shoots from two plates constituted a single biological replicate. Three biological replicates were sampled separately three times.

An ample supply of ammonium increases the concentrations of amino acids compared with nitrate supply alone (6, 20). In particular, the molar ratios of Gln to Glu are elevated at higher ammonium levels, suggesting that Gln synthesis by glutamine synthetase (GLN) overflows glutamate synthase (GOGAT) capacity. Our hierarchical cluster analysis of amino acid content in shoots clearly demonstrated that the type of N source, i.e. 10 mM ammonium or nitrate, was the strongest determinant for plant amino acid composition (Fig. 4*B* and *SI Appendix*, S8*A*). In this analysis, Col and the *GLN2*-deficient lines categorized into separate clusters depending on the N source. The molar ratio of Gln to Glu (Fig. 4*C*), total amino acid-N content per amino acid (*SI Appendix*, Fig. S8*B*), total amino acid-N content per fresh weight (*SI Appendix*, Fig. S8*C*), and the molar ratios of N to C in total amino acids (*SI Appendix*, Fig. S8*D*) were consistently larger in ammonium-grown shoots than nitrate-grown shoots, and this large ammonium-N input was partly but significantly attenuated by *GLN2* deficiency. These findings suggest that the GLN2 reaction leads to excessive incorporation of ammonium-N into amino acids in shoots when toxic levels of ammonium are present.

### Ammonium Assimilation by GLN2 Causes Acidic Stress

The amino acid profiles suggested that metabolic imbalances due to excessive ammonium assimilation by GLN2 could be a cause of ammonium toxicity. We have previously demonstrated that nitrate addition at adequate concentrations mitigates ammonium toxicity without reducing amino acid accumulation (6). Therefore, a phenomenon triggered by some GLN2-mediated process other than amino acid accumulation should be a cause of ammonium toxicity. Notably, the GLN reaction is a proton-producing process (21). The stoichiometry of this reaction is two protons per each glutamine produced, one proton of which is derived from ATP hydrolysis and the other is from deprotonation of NH_4_^+^. Conversely, the subsequent ferredoxin-dependent glutamate synthase (Fd-GOGAT) reaction consumes two protons per one glutamine incorporated. Given that the molar ratio of Gln to Glu was about 10 in the ammonium condition but close to 1 in the nitrate condition (Fig. 4*C*), proton production in the ammonium condition could proceed beyond its consumption. Strikingly, a previous study found that 43% of ammonium-inducible genes correspond to acidic stress-inducible genes in Arabidopsis roots (22, 23). Thus, we hypothesized that excessive ammonium assimilation by GLN2 causes acidic stress to the plants growing in ammonium.

We re-surveyed our microarray data by focusing on previously identified acidic stress-responsive genes (23) (Fig. 5*A* and *Datasets*, Table S2). All acidic stress-inducible genes were entirely downregulated in *ami2* shoots compared with Col, whereas the acidic stress-repressive genes showed the opposite trend. The transcript levels of *ALMT1*, a typical acidic stress-inducible gene, were determined in the shoots of Col and *ami2* plants incubated in 10 mM ammonium or nitrate with or without methionine sulfoximine (MSX), an inhibitor of the GLN reaction (Fig. 5*B*). *ALMT1* expression was much higher in the ammonium-treated Col shoots than the nitrate-treated samples. This ammonium-dependent induction was significantly diminished in the *ami2* shoots and was mimicked by MSX treatment. Also, other proton-inducible genes such as *GABA-T*, *GAD1*, *GDH2*, *PGIP1*, and *PGIP2* (24) were ammonium-inducible, and their inductions were suppressed or attenuated by *GLN2* deficiency (*SI Appendix*, Fig. S9*A*). These results support our hypothesis associating ammonium assimilation with acidic stress. Moreover, in Col and *ami2* reciprocally-grafted plants growing in the ammonium condition, *ALMT1* expression was significantly lower in the *ami2*-derived shoots than the Col-derived shoots (*SI Appendix*, Fig. S9*B*), indicating that shoot GLN2 locally causes acidic stress to the shoot. Furthermore, *ALMT1* expression was analyzed using grafted plants between Col and a mutant lacking the STOP1 transcription factor (*stop1-KO*) that induces *ALMT1* to respond to acidic stress (24) (*SI Appendix*, Fig. S9*C*). In the *stop1-KO*-derived shoots, the ammonium-dependent induction of *ALMT1* disappeared, reconfirming the notion that acidic stress occurs in plants growing in ammonium.

**Fig. 5.**
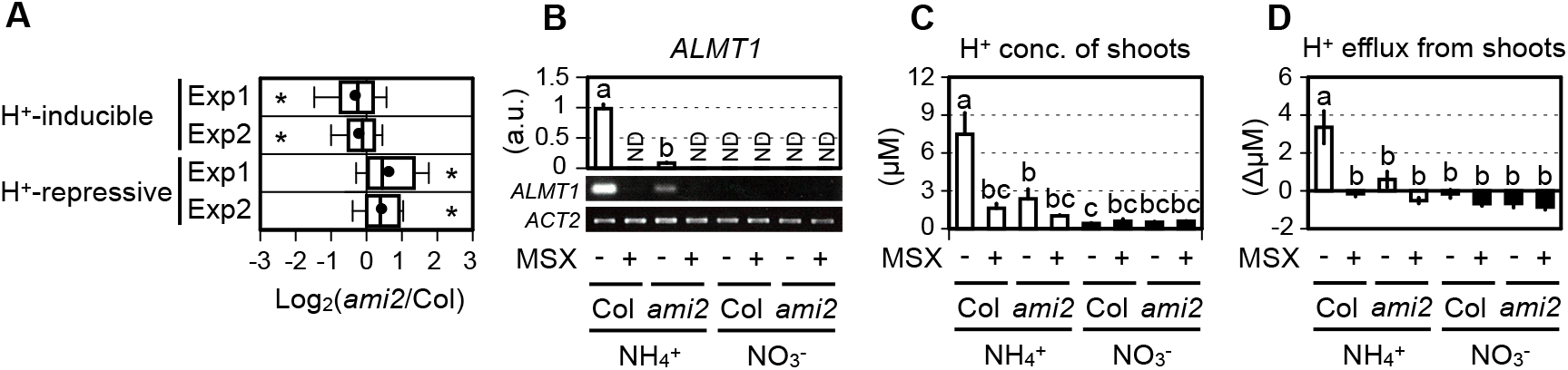
Ammonium assimilation by GLN2 causes acidic stress. (*A*) Box plots of the differences in expression of the acidic stress-responsive genes between the Col and *ami2* shoots 3 d after transfer to media containing 10 mM ammonium. The gene list was obtained from (23) (For further details, see *Datasets*, Table S2). Two independent experiments (Exp1 and Exp2) were performed. Nine shoots from three plates constituted a single biological replicate. An individual box plot shows the median (heavy vertical line), the 25^th^ to 75^th^ percentiles (right and left sides of the box), the 10^th^ to 90^th^ percentiles (whiskers), and the mean (closed circle). Welch’s *t*-test was run at α = 0.05; \**p* < 0.05. (*B*) Effects of MSX treatment on the relative transcript level of *ALMT1* in the Col and *ami2* shoots 3 d after transfer to media containing 10 mM ammonium or 10 mM nitrate. The transcript levels were evaluated both by RT-qPCR (mean ± SD; n = 3) and semi-quantitative RT-PCR with agarose gel electrophoresis. *ACTIN2* (*ACT2*) was the internal standard. Three shoots from one plate constituted a single biological replicate. (*C*) Effects of MSX treatment on proton concentrations in water extracts from the Col and *ami2* shoots 3 d after transfer to media containing 10 mM ammonium or 10 mM nitrate (mean ± SD; n = 3). Three shoots from one plate constituted a single biological replicate. (*D*) Effects of MSX treatment on proton efflux rates from the Col and *ami2* shoots 3 d after transfer to media containing 10 mM ammonium or 10 mM nitrate (mean ± SE; n = 3). Three shoots from one plate constituted a single biological replicate. (*B*-*D*) Tukey-Kramer’s multiple comparison test was conducted at a significance level of *P* < 0.05 only when a one-way ANOVA was significant at *P* < 0.05. Different letters denote significant differences.

It is widely accepted that the reduction from nitrate to ammonium consumes a proton, suggesting that nitrate reduction could attenuate acidic stress caused by excess ammonium and might explain why nitrate addition alleviates ammonium toxicity. To verify this hypothesis, we analyzed shoot expression of *ALMT1* using grafted plants between Col and the *NITRATE REDUCTASE*-null mutant (designated as NR-null) (25) (*SI Appendix*, Fig. S9*D*). Addition of 2.5 mM nitrate diminished the ammonium-dependent *ALMT1* induction in the Col-derived shoots but not in the NR-null-derived shoots, thereby supporting the above hypothesis.

To obtain direct evidence for ammonium-dependent proton production, we measured the proton concentrations of water extracts from the Col and *ami2* shoots incubated in media containing 10 mM ammonium or nitrate with or without MSX (Fig. 5*C*). The ammonium-treated Col shoots contained the highest concentrations of protons; proton content was significantly decreased by *GLN2* deficiency and by MSX treatment to levels comparable to those in nitrate-treated shoots. A similar trend was observed among the Col, *ami2*, and *gln2* shoots grown on ammonium- or nitrate-containing media (*SI Appendix*, Fig. S9*E*).

The presence of ammonium in cultures generally acidifies the external media (22). Thus, we quantified the proton efflux from the Col and *ami2* shoots incubated in media containing 10 mM ammonium or nitrate with or without MSX (Fig. 5*D*). Incubation of the Col shoots in the presence of ammonium strongly acidified the external media, which was alleviated by *GLN2* deficiency and by MSX treatment. A similar tendency was observed by qualitative measurements with a pH indicator of proton effluxes from mesophyll cells where GLN2 is predominantly expressed (*SI Appendix*, Fig. S9*F*). Thus, we conclude that ammonium assimilation by GLN2 without nitrate increases shoot acidity.

### Ammonium Toxicity is Closely Associated with Acidic Stress

If acidic stress rather than ammonium accumulation has a dominant effect on ammonium toxicity, an application of alkaline ammonia should reduce the toxicity. Given that the GLN2 reaction is a primary cause of increased acidic stress, an elevation in medium pH may increase the shoot growth of Col more effectively than that of the *GLN2*-deficient mutants. As expected, addition of a 25% ammonia solution to media containing 10 mM ammonium elevated the pH from 5.7 to 6.7 and significantly improved shoot growth with a concomitant decrease in acidity (Fig. 6*A*). Fresh weights of Col shoots grown at pH 6.7 increased by ca. 180% compared with those grown at pH 5.7, whereas fresh weights of *ami2* and *gln2* shoots only increased by ca. 30% and 60%, respectively (Fig. 6*B*). In addition, the acid-sensitive *STOP1*-deficient mutants had slightly but significantly lower shoot growth when grown in 10 mM ammonium (Fig. 6*C* and *SI Appendix*, Fig. S9*G*), although their acid-hypersensitivity has been described only in roots to date (24, 26). Moreover, the NR-null-derived shoots that lack a proton-consuming nitrate reduction capacity failed to attenuate ammonium toxicity by nitrate addition (Fig. 6*D*). Collectively, our results lead to the conclusion that acidic stress is one of the primary causes of ammonium toxicity.

**Fig. 6.**
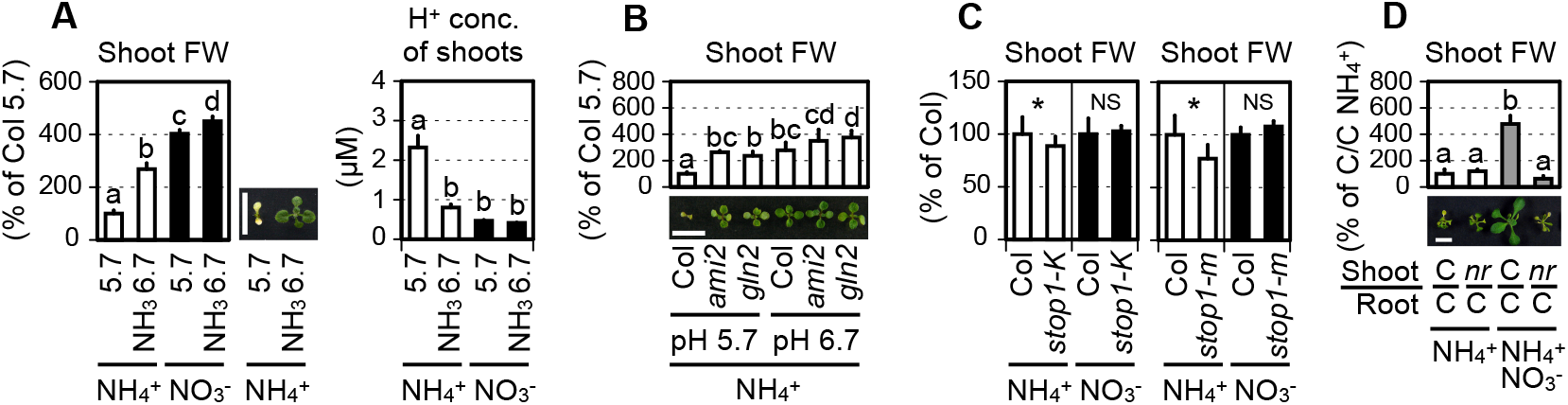
Ammonium toxicity is closely linked with acidic stress. (*A*) Effects of NH_3_ application on shoot FW and proton concentrations in water extracts of Col grown on media containing 10 mM ammonium or 10 mM nitrate for 5 d (mean ± SD; n = 3). Thirty-seven shoots from one plate constituted a single biological replicate. The pH was adjusted to pH 5.7 with 1N KOH; subsequently, 25% (v/v) ammonia was added to adjust the pH from 5.7 to 6.7. A representative photograph of 11-d-old shoots grown on 10 mM ammonium (12 plants per plate) is shown. (*B*) Effects of intermediate pH on the FW of shoots from Col, *ami2*, and *gln2* grown on media containing 10 mM ammonium for 11 d (mean ± SD; n = 6). Three shoots from one plate constituted a single biological replicate. The pH was adjusted to pH 5.7 with 1N KOH; subsequently, 1N NaOH was used to adjust the pH from 5.7 to 6.7 to maintain the potassium concentration constant among all samples. A representative photograph of 11-d-old shoots is shown. (*C*) FW of shoots from Col, *stop1*-*KO* (*stop1-k*), and the *stop1* mutant (*stop1-m*) grown on media containing 10 mM ammonium (mean ± SD; n = 20) or 10 mM nitrate (mean ± SD; n = 5) for 11 d. Six shoots from one plate constituted a single biological replicate. Welch’s *t*-test was run at α = 0.05; \**p* < 0.05. NS denotes not significant. (*D*) FW of shoots from plants grafted between Col (C) and the NR-null mutant (*nr*) 7 d after transfer to media containing 10 mM ammonium (NH_4_^+^) or 2.5 mM nitrate and 10 mM ammonium (NH_4_^+^ NO_3_^−^) conditions (mean ± SD; n = 3). One shoot from one plate constituted a single biological replicate. A representative photograph of shoots 7 d after transfer to media is shown. (*A*, *B*, *D*) Tukey-Kramer’s multiple comparison test was conducted at a significance level of *P* < 0.05 only when a one-way ANOVA was significant at *P* < 0.05. Different letters denote significant differences. The scale bar represents 10 mm.

### GLN2 Causes Ammonium Toxicity Independently of NRT1.1

We have already reported that Arabidopsis NITRATE TRANSPORTER 1.1 (NRT1.1), acting as a nitrate transceptor, also aggravates ammonium toxicity (27). This result was recently confirmed by another group (11). Thus, we investigated whether NRT1.1 and GLN2 increase the sensitivity to ammonium through a common mechanism. A RT-qPCR analysis revealed that deficiency of either *NRT1.1* or *GLN2* did not downregulate the expression of the other gene (*SI Appendix*, Fig. S10*A*), and that *GLN2* expression was almost 3-times higher in *nrt1.1* than in Col. Therefore, the enhanced ammonium tolerance of *nrt1.1* cannot be explained by reduced *GLN2* expression as in *gln2*. Moreover, a homozygous double mutant of *NRT1.1* and *GLN2* showed slightly but significantly larger shoot fresh weight, leaf number, shoot diameter, and chlorophyll content compared with any of the single mutants (*SI Appendix*, Fig. S10 *B*-*D*). These findings suggest that *NRT1.1* and *GLN2* are implicated in ammonium-sensitivity independently.

## Discussion

Although ammonium is a toxic compound for plant growth, our results demonstrate that ammonium assimilation by shoot GLN2 rather than ammonium accumulation is a major cause of ammonium toxicity (Fig. 2 and Fig. 4). In plants growing in toxic levels of ammonium as a sole N source, assimilation of ammonium by GLN2 would occur largely due to bypassing nitrate reduction as the rate-limiting step for N assimilation. The resultant increase in the ratio of Gln to Glu content (Fig. 4*C*) corresponds to the preferential enhancement of the proton-producing GLN reaction over the proton-consuming GOGAT reaction. This metabolic imbalance exerted by the GLN2 reaction leads to the production of large amounts of protons in shoot cells that stimulate proton effluxes to the apoplasm; however, the volume of the shoot apoplasm is probably too small to accommodate such a large proton efflux. Thus, in the presence of toxic levels of ammonium, the GLN2 reaction causes acidic stress inside and outside the cells and triggers acidic stress responses that modulate gene expression (Fig. 5 and *SI Appendix*, Fig. S9 *A*-*F*). Given that the wild-type when grown at a higher pH phenocopies the ammonium-insensitive lines at lower pH, the acidic stress-sensitive mutants show ammonium-hypersensitivity, and proton-consuming nitrate reduction alleviates ammonium toxicity (Fig. 6 and *SI Appendix*, Fig. S9G), we conclude that acidic stress is one of the primary causes for ammonium toxicity. In this framework, upregulation of Arabidopsis *NIA1* and *NIA2* genes encoding nitrate reductase (NR) (23, 24) and activation of spinach NR (28) responding to acidic stress are understandable regulatory responses in the context of maintaining cellular pH homeostasis.

The present study does not address how ammonium-dependent acidification triggers growth deficiency at the cellular and subcellular scales. The chloroplastic localization of GLN2 indicates that proton production must occur within chloroplasts in the elevated ammonium condition. A previous study reported abnormal chloroplast membrane structure including swollen compartments at late stages of ammonium toxicity (29); however, we did not find any similar structural changes in the shoots of ammonium-grown plants (*SI Appendix*, Fig. S11*A*), where the intermediates of the Calvin-Benson cycle were not depleted compared with nitrate-grown shoots (*SI Appendix*, Fig. S11*B*). These observations do not support a deficiency in chloroplast function as a primary cause of ammonium toxicity. Apoplastic pH in sunflower leaves and cytosolic pH in carrot cell suspensions decrease after application of millimolar levels of ammonium (30, 31). The ammonium-inducible genes whose expression is downregulated by *GLN2* deficiency, *PGIP1* and *PGIP2* (*SI Appendix*, Fig. S9*A*), contribute to cell wall stabilization under acidic stress (26), implying apoplastic acidification as a target of ammonium toxicity. On the other hand, in the presence of toxic levels of ammonium, the GABA shunt-related genes (*SI Appendix*, Fig. S9*A*) and oxygen uptake rates (32) are induced as biochemical pH-stats (33) that may represent an intracellular acidic burden. Given that changes in pH environments influence a wide spectrum of physiological processes, elucidating the relationship between ammonium-dependent acidification and growth deficiency awaits future study.

At the whole-plant scale, our grafting work demonstrated that root GLN1;2 activity attenuates ammonium toxicity in the shoots, whilst shoot GLN2 activity causes the condition (Fig. 3). Considering that GLN1;2 is the ammonium-inducible low-affinity enzyme expressed in the epidermis and cortex of roots, and its deficiency elevates ammonium levels in xylem sap when ammonium is supplied (17), root GLN1;2 could act as a barrier to prevent the shoot-to-root transport of ammonium, thus avoiding ammonium assimilation by shoot GLN2. In oilseed rape plants, replacing 3 mM nitrate in a nutrient solution with 10 mM ammonium increased the ammonium levels in xylem sap linearly with time, attaining concentrations greater than 5 mM (34), which could indicate breaking through the barrier. On the other hand, we did not determine whether shoot GLN1 isozymes attenuate or deteriorate the toxicity. With ammonium nitrate nutrition, Arabidopsis shoot GLN1;2 activity promotes shoot growth (35). We observed a larger protein signal corresponding to shoot GLN1s when plants received ammonium rather than nitrate nutrition (*SI Appendix*, Fig. S12 *A* and *B*), implying a barrier function of GLN1 in the shoot. Further grafting work using several combinations of multiple mutants on *GLN1s* are required to confirm this hypothesis.

The present study demonstrated that *GLN2* and *NRT1.1* reduce ammonium tolerance via separate mechanisms when plants experience high ammonium conditions (*SI Appendix*, Fig. S10). On the other hand, these genes are nitrate-inducible genes that are crucial for plant adaptation to nitrate-dominant environments (36, 37). This observation suggests that the adaptive traits to nitrate and ammonium could be exclusive, and therefore, breeding elevated CO_2_-adapted crops in terms of their mode of N utilization, i.e. ammonium-tolerant crops, might sacrifice their adaptability to nitrate.

## Materials and Methods

Detailed information on plant materials, their growth conditions, isolation of ammonium-insensitive lines, expression analyses for mRNAs and proteins, the grafting procedure, the activity assay, metabolite analysis by mass spectrometry, physiological analyses, TEM observations, and statistical analyses is provided in *SI Appendix*, *SI Materials and Methods*.

## ACKNOWLEDGMENTS

Seeds of the *A thaliana* NR-null mutant were kindly provided by Prof. Nigel M. Crawford, Division of Biological Sciences, University of California, San Diego. This work was supported by the Building of Consortia for the Development of Human Resources in Science and Technology, by the Japan Society for the Promotion of Science KAKENHI Grant No. JP17K15237, by the Inamori Foundation, by the Agropolis Foundation No. 1502-405, and by a Grant-in-Aid for Young Scientists from Shimane University.

## References

1. Rubio-Asensio JS, Bloom AJ (2016) Inorganic nitrogen form: a major player in wheat and Arabidopsis responses to elevated CO_2_. J Exp Bot 68: 2611–2625.

2. Kant S, Bi YM, Rothstein SJ (2011) Understanding plant response to nitrogen limitation for the improvement of crop nitrogen use efficiency. J Exp Bot 62: 1499–1509.

3. Britto DT, Kronzucker HJ (2002) NH_4_^+^ toxicity in higher plants: a critical review. J Plant Physiol 159: 567–584.

4. Li B, Li G, Kronzucker HJ, Baluška F, Shi W (2014) Ammonium stress in *Arabidopsis* signaling, genetic loci, and physiological targets. Trends Plant Sci 19: 107–114.

5. Esteban R, Ariz I, Cruz C, Moran JF (2016) Mechanisms of ammonium toxicity and the quest for tolerance. Plant Sci 248: 92–101.

6. Hachiya, T et al. (2012) Nitrate addition alleviates ammonium toxicity without lessening ammonium accumulation, organic acid depletion and inorganic cation depletion in Arabidopsis thaliana shoots. Plant Cell Physiol 53: 577–591.

7. Qin C, et al. (2008) GDP-mannose pyrophosphorylase is a genetic determinant of ammonium sensitivity in *Arabidopsis thaliana*. Proc Natl Acad Sci USA 105: 18308–18313.

8. Tanaka H, et al. (2015) Identi cation and characterization of *Arabidopsis* AtNUDX9 as a GDP-d-mannose pyrophosphohydrolase: its involvement in root growth inhibition in response to ammonium. J Exp Bot 66: 5797–5808.

9. Li B, et al. (2012) Arabidopsis plastid metalloprotease AMOS1/EGY1 integrates with ABA signaling to regulate global gene expression in response to ammonium stress. Plant Physiol 160: 2040–2051.

10. Li G, et al. (2019) The Arabidopsis *AMOT1/EIN3* gene plays a important role in the amelioration of ammonium toxicity. J Exp Bot 70: 1375–1388

11. Jian S, et al. (2018) NRT1.1-related NH_4_^+^ toxicity is associated with a disturbed balance between NH_4_^+^ uptake and assimilation. Plant Physiol 178: 1473–1488.

12. Ichikawa T, et al. (2006) The FOX hunting system: an alternative gain-of-function gene hunting technique. Plant J 48: 974–985.

13. Sakakibara H, Kawabata S, Hase T, Sugiyama T (1992) Differential effects of nitrate and light on the expression of glutamine synthetases and ferredoxin-dependent glutamate synthase in maize. Plant Cell Physiol 33: 1193–1198.

14. Cheng XF, Wang ZY (2005) Overexpression of *COL9*, a *CONSTANS-LIKE* gene, delays flowering by reducing expression of *CO* and *FT* in *Arabidopsis thaliana*. Plant J 43: 758–768.

15. Lothier J, et al. (2011) The cytosolic glutamine synthetase GLN1;2 plays a role in the control of plant growth and ammonium homeostasis in *Arabidopsis* rosettes when nitrate supply is not limiting. J Exp Bot 62: 1375–1390.

16. Guan M, Møller IS, Schjoerring JK (2015) Two cytosolic glutamine synthetase isoforms play specific roles for seed germination and seed yield structure in *Arabidopsis*. J Exp Bot 66: 203–212.

17. Konishi N, et al. (2017) Contributions of two cytosolic glutamine synthetase isozymes to ammonium assimilation in *Arabidopsis* roots. J Exp Bot 68: 613–625.

18. Thieme CJ, et al. (2015) Endogenous *Arabidopsis* messenger RNAs transported to distant tissues. Nat Plants 1: 15025.

19. Bittsánszky A, Pilinszky K, Gyulai G, Komives T (2015) Overcoming ammonium toxicity. Plant Sci 231: 184–190.

20. Sato S, Soga T, Nishioka T, Tomita M (2004) Simultaneous determination of the main metabolites in rice leaves using capillary electrophoresis mass spectrometry and capillary electrophoresis diode array detection. Plant J 40: 151–163.

21. Britto DT, Kronzucker HJ (2005) Nitrogen acquisition, PEP carboxylase, and cellular pH homeostasis: new views on old paradigms. Plant Cell Environ 28: 1396–1409.

22. Patterson K, et al. (2010) Distinct signalling pathways and transcriptome response signatures differentiate ammonium- and nitrate-supplied plants. Plant Cell Environ 33: 1486–1501.

23. Lager I, et al. (2010) Changes in external pH rapidly alter plant gene expression and modulate auxin and elicitor responses. Plant Cell Environ 33: 1513–1528.

24. Sawaki Y, et al. (2009) STOP1 regulates multiple genes that protect Arabidopsis from proton and aluminum toxicities. Plant Physiol 150: 281–294.

25. Wang R, Xing X, Crawford N (2007) Nitrite acts as a transcriptome signal at micromolar concentrations in Arabidopsis roots. Plant Physiol 145: 1735–1745.

26. Kobayashi Y, et al. (2014) STOP2 activates transcription of several genes for Al- and Low pH-tolerance that are regulated by STOP1 in *Arabidopsis*. Mol Plant 7: 311–322.

27. Hachiya T, et al. (2011) Evidence for a nitrate-independent function of the nitrate sensor NRT1.1 in Arabidopsis thaliana. J Plant Res 124: 425–430.

28. Kaiser WM, Brendle-Behnisch E (1995) Acid-base-modulation of nitrate reductase in leaf tissues. Planta 196: 1–6.

29. Puritch GS, Barker AV (1967) Structure and function of tomato leaf chloroplasts during ammonium toxicity. Plant Physiol 42: 1229–1238.

30. Carroll AD, et al. (1994) Ammonium assimilation and the role of γ-aminobutyric acid in pH homeostasis in carrot cell suspensions. Plant Physiol 106: 513–520.

31. Hoffmann B, Plänker R, Mengel K (1992) Measurement of pH in the apoplast of sunflower leaves by means of fluorescence. Physiol Plant 84: 146–153.

32. Hachiya T, et al. (2010) Ammonium-dependent respiratory increase is dependent on the cytochrome pathway in *Arabidopsis thaliana* shoots. Plant Cell Environ 33: 1888–1897.

33. Sakano K (1998) Revision of Biochemical pH-Stat: Involvement of Alternative Pathway Metabolisms. Plant Cell Physiol 39: 467–473.

34. Schjoerring JK, Husted S, Mäck G, Mattsson M (2002) The regulation of ammonium translocation in plants. J Exp Bot 53: 883–890.

35. Guan M, Schjoerring JK (2016) Peering into the separate roles of root and shoot cytosolic glutamine synthetase 1;2 by use of grafting experiments in Arabidopsis. Plant Signal Behav 11: e1245253.

36. Hachiya T, Sakakibara H (2017) Interactions between nitrate and ammonium in their uptake, allocation, assimilation, and signaling in plants. J Exp Bot 68: 2501–2512.

37. Marchive C, et al. (2013) Nuclear retention of the transcription factor NLP7 orchestrates the early response to nitrate in plants. Nat Commun 4: 1713.

