## Supplementary material for "Excessive assimilation of ammonium by plastidic glutamine synthetase is a major cause of ammonium toxicity in *Arabidopsis thaliana*": Supple MM & Figs

**Author affiliations: :** [a]Department of Molecular and Functional Genomics, Interdisciplinary Center for Science Research, Shimane University, Matsue 690-8504, Japan, [b]Department of Biological Mechanisms and Functions, Graduate School of Bioagricultural Sciences, Nagoya University, Furo-cho, Chikusa-ku, Nagoya, Aichi 464-8601, Japan, [c]Institute for Advanced Research, Nagoya University, Furo-cho, Chikusa-ku, Nagoya, Aichi 464-8602, Japan, [d]RIKEN Center for Sustainable Resource Science, 1-7-22 Suehiro-cho, Tsurumi-ku, Yokohama, Kanagawa 230-0045, Japan, [e]Graduate School of Science and Engineering, Saitama University, 255 Shimo-Okubo, Sakura-ku, Saitama-city, Saitama 338-8570, Japan, [f]Biochimie et Physiologie Moléculaire des Plantes, CNRS/INRA/SupAgro-M/Montpellier University, Montpellier 34060, France.

**Corresponding Author:** Takushi Hachiya (http://orcid.org/0000-0002-9239-6529)
Departement of Molecular and Functional Genomics, Interdisciplinary Center for Science Research, Shimane University, Matsue 690-8504, Japan

**This PDF file includes:**
SI Materials and Methods, Figs. S1 to S12, References for SI reference citations

**Other supplementary materials for this manuscript include the following:**
Tables S1 to S4

**SI Materials and Methods**

**Plant materials and growth conditions.** *A. thaliana*, accession Columbia (Col) was the control line used in this study. The T-DNA insertion mutants *gln2* (SALK_051953), *gln1;2-1* (SALK_145235) (1), *gln1;2-2* (SALK_102291) (2), *stop1-KO* (SALK_114108) (3), and *nrt1.1* (SALK_097431) (4) were purchased from the European Arabidopsis Stock Centre. The *Arabidopsis* FOX lines (5) and the ethyl methanesulfonate-mutagenized s*top1* mutant (psi00011) (3) were obtained from the RIKEN BioResource Research Center. The NR-null mutant (6) was acquired from Professor Nigel M. Crawford (University of California, San Diego).

Seeds were surface-sterilized and sown in plastic Petri dishes (diameter 90 mm, depth 20 mm, Iwaki, Tokyo, Japan) containing about 30-mL N-modified Murashige and Skoog medium, supplemented with 10 mM MES, 2% (w/v) sucrose, and 0.25% (w/v) gellan gum (pH 5.7). The N and K sources were 10 mM $KNO_3$ (10 mM nitrate condition), 5 mM $NH_4NO_3$ with 10 mM KCl (5 mM ammonium nitrate condition), or 5 mM $(NH_4)_2SO_4$ with 10 mM KCl (10 mM ammonium condition). The seeds were placed in the dark at 4˚C for 3 d. Plants were grown in a horizontal position under a photosynthetic photon flux density of 100–130 μmol $m^{-2}$ $s^{-1}$ (16 h light/8 h dark cycle) at 23°C. For the transfer experiments, surface-sterilized seeds were sown in plastic Petri dishes (length, 140 mm; width, 100 mm; depth, 20 mm; Eiken Chemical Co. Ltd., Taito-ku, Tokyo, Japan) containing 50-mL half-strength modified Murashige and Skoog medium containing 2.5 mM ammonium as the sole N source at pH 6.7 (7). The seeds were placed in the dark at 4˚C for 3 d. The plants were grown in a vertical position for 7 d under a photosynthetic photon flux density of 100–130 μmol $m^{-2}$ $s^{-1}$ (16 h light/8 h dark cycle) at 23 °C. The plants were transferred with sterilized tweezers to varying N conditions and grown in a horizontal position for further experiments. Other details of plantlet cultivation are presented in the Results section and the figure legends.

**Isolation of ammonium-insensitive lines.** Forty-five seedlings per plate of *Arabidopsis* FOX lines (5) were grown on media containing 10 mM ammonium for 11 d in a horizontal position. The insensitivity to ammonium was visually evaluated based on size and color of the cotyledons or true leaves. The next generation was obtained to refresh the seeds for the more detailed second screen. The ammonium insensitivity of the line was evaluated quantitatively by determining their fresh weights in comparison with the wild types. In total, seed mixtures that included 15800 lines (pss10001-10016) were used for a series of screens, resulting in the isolation of three lines.

**Extraction of total RNA.** Shoots and roots were harvested, immediately frozen with liquid $N_2$, and stored at −80°C until use. Frozen samples were ground with a Multi-Beads Shocker (Yasui Kikai Corp., Osaka Prefecture, Osaka, Japan) using zirconia beads (diameter, 5 mm). Total RNA was extracted using the RNeasy Plant Mini Kit (Qiagen) and on-column DNase digestion according to the manufacturer's instructions.

**Reverse transcription and quantitative (real-time) PCR.** Reverse transcription was performed using a ReverTra Ace qPCR RT Master Mix with gDNA Remover (Toyobo Co. Ltd., Tokyo, Japan) according to the manufacturer's instructions. The synthesized cDNA was diluted 10-fold with distilled water and used in quantitative PCR (qPCR). Transcript levels were measured using a StepOnePlus Real-Time PCR System (Thermo Fisher Scientific, Waltham, MA, USA). The obtained cDNA (2 μL) was amplified in the presence of 10-μL KAPA SYBR FAST qPCR Kit (Nippon Genetics Co. Ltd., Tokyo, Japan), 0.4-μL specific primers (0.2 μM final concentration), and 7.2-μL sterile water. Transcript levels were quantified using a relative standard curve with *ACTIN3* as the internal standard. A dilution series of total cDNAs were used as templates to generate the standard curves. The primer sequences used in the experiments are shown in *Datasets*, Table S3.

**Immunoblot analysis of GLN isoproteins.** Immunoblot analysis of GLN proteins was performed based on the method reported by (7). Shoots were harvested, frozen with liquid $N_2$, and stored at −80°C until use. Frozen samples were ground with a Multi-Beads Shocker (Yasui Kikai Corp.) using zirconia beads (diameter, 5 mm). Total proteins were extracted with 10 volumes of sample buffer [2% (w/v) SDS, 62.5 mM Tris–HCl (pH 6.8), 7.5% (v/v) glycerol, 50 mM DTT and 0.01% (w/v) bromophenol blue] containing a protease inhibitor tablet (Roche Diagnostics, Basel, Switzerland). The extracts were incubated at 95°C for 5 min followed by cooling on ice and centrifugation at 20,400 ×*g* at room temperature (20°C–25°C) for 10 min. A 10-µL aliquot of the supernatant (equivalent to approximately 1-mg fresh sample) was subjected to SDS-PAGE in a 12% (w/v) gel (TGX FastCast Acrylamide Kit, Bio-Rad Laboratories, Hercules, CA, USA) and transferred to a PVDF membrane (Trans-Blot Turbo Mini PVDF Transfer Packs, Bio-Rad Laboratories, Hercules, CA, USA) using HIGH MW with a semi-dry blotting system (Trans-Blot Turbo Transfer System, Bio-Rad Laboratories). The membrane was incubated overnight in blocking buffer containing 5% (w/v) ECL Prime Blocking Agent (GE Healthcare, Little Chalfont, UK), 0.1% (v/v) Tween-20, 50 mM Tris-HCl, and 150 mM NaCl (pH 7.6). The blocked membrane was then incubated for 1 hour with a 1/10,000 dilution of polyclonal antibodies raised against maize cytosolic glutamine synthetase (8). After rinsing with a buffer containing 0.1% Tween-20, 50 mM Tris-HCl, and 150 mM NaCl (pH 7.6), the antigen–antibody complex was detected using a 1/100,000 dilution of horseradish peroxidase conjugated to goat anti-rabbit IgG (NA935, GE Healthcare) and visualized by chemiluminescent detection (ECL Prime, GE Healthcare) using ImageQuant LAS 3000 mini (Fujifilm, Tokyo, Japan). The signal intensities for each band corresponding to GLN1s and GLN2 isoproteins were quantified using Image J software, version 10.2. After detection, the membrane was stained with the GelCode Blue Stain Reagent (ThermoFisher Scientific).

**Microarray analysis.** Total RNA was extracted as described above. RNA quality was assessed using an Agilent 2100 bioanalyzer (Agilent Technologies). RNA amplification, labelling, hybridization and scanning with the 3′ IVT Express Kit (Affymetrix) and the GeneChip Arabidopsis Genome ATH1 Array (Affymetrix) were conducted according to the manufacturer's instructions. The data set from the microarray chips was normalized by the Microarray Suite 5.0 (MAS5) method (Affymetrix). When the signal detection of a transcript was labelled "Absent" or "Marginal", the sequence was omitted from the subsequent quantitative analysis. The raw data are shown in *Datasets*, Table S4. Changes in respective gene expression levels between Col and *ami2* were represented as logarithms to base 2 of the ratios in signal intensities.

**Grafting between shoots and roots.** *Arabidopsis* seedlings were grown for 4 d on media containing 2.5 mM ammonium as the sole N source at pH 6.7 (7) before grafting. Each seedling was perpendicularly cut at the hypocotyl with the tip of an injection needle (NN-2613S, TERUMO, Tokyo, Japan) on a mixed cellulose membrane (HAWP09000, MERCK MILLIPORE, Darmstadt, Germany). The scion was kept in touch with the rootstock through a section of 0.4 mm diameter silicon tubing as described (9). The grafted plants were grown for another 4 d at 27˚C, and then, for 2 d at 23˚C. Plants without adventitious roots were transferred to the fresh media and further grown and used for analyses of growth and gene expression.

**Determination of glutamine synthetase activity.** GLN activity was determined as the ADP-dependent conversion rate of L-glutamine to γ-glutamylhydroxamate. Shoots and roots were harvested, immediately frozen with liquid $N_2$ and stored at −80°C until use. Frozen samples were ground with a TissueLyser II (QIAGEN) using 5 mm zirconia beads. The powder was mixed with 10 volumes of extraction buffer (100 mM Tris-HCl, pH7.5, 1% (w/v) PVP-40, 1 mM EDTA, 1 mM $MnCl_2$, 0.5% (v/v) β-mercaptoethanol, 0.1 mM 4-APMSF). The extracts were centrifuged at 12,000 ×*g* at 4°C for 10 min. The reaction was started by adding 45 µl of pre-incubated assay buffer (40 mM imidazole-HCl, pH 7.0, 20 mM sodium arsenate, 0.5 mM ADP, 3 mM $MnCl_2$, 60 mM $NH_2OH$, 30 mM L-glutamine) to 5 µl of the supernatant. The mixture was incubated at 30°C for 15 min. The reaction was stopped by adding 30 µl of $FeCl_3$-TCA-HCl solution (2.6% $FeCl_3 \cdot 6H_2O$, 4% trichloroacetic acid in 1N HCl). Ferric γ-glutamylhydroxamic acid was measured by spectrophotometric absorbance at 540 nm.

**Determination of amino acids and photosynthetic intermediates.** Amino acids and photosynthetic intermediates were extracted and their concentrations were determined based on the method reported by (10) with minor modifications. Shoots were harvested, frozen with liquid $N_2$, and stored at −80°C until use. Frozen samples were ground with a Multi-Beads Shocker (Yasui Kikai Corp.) using zirconia beads (diameter, 5 mm). Metabolites were extracted with 50% (v/v) methanol containing 50 µM PIPES and 50 µM methionine sulfone (internal standard). After the first centrifugation (21,500 ×*g*, 5 min,

4°C), the supernatant was transferred to a 3 kDa cut-off filter (UFC500324, Millipore, Billerica, MA, USA) and centrifuged again (13,700 ×*g*, 30 min, 4 °C). The filtered extract was used for capillary electrophoresis-mass spectrometry (CE-MS) analysis.

Measurements of metabolites were performed using an Agilent 6400 series triple quadrupole CE-MS system (CE; 7100 Capillary Electrophoresis, MS; 6420 Triple Quad LC/MS, Agilent Technologies, Santa Clara, CA, USA) in multi reaction monitoring (MRM) mode (10, 11). A DB-WAX capillary (polyethylene glycol-coated, 100 cm × 50 μm i.d., Agilent Technology) with 20 mM ammonium acetate (pH 8.5) as the running buffer was used for measuring anionic compounds (e.g. organic acids and phosphorylated compounds), and an uncoated fused silica capillary (100 cm × 50 μm i.d., GL Sciences, Tokyo, Japan) with 1 M formic acetate was used for cationic compounds (e.g. amino acids). MS analysis at the applied −25 kV for anions was carried out in negative ion mode. Cations were determined in positive ion mode (applied voltage 25 kV). For MS stabilization, 5 mM ammonium acetate (for anions) or 0.1% formic acid (v/v) (for cations) in 50% (v/v) methanol was used as sheath solution, applied to the capillary at 10 μl $min^{-1}$ using an isocratic HPLC pump (Agilent 1200 series) equipped with a 1:100 splitter. The capillary voltage (± 3500 V) and the drying nitrogen gas (at 320 °C) flow (8 l $min^{-1}$) were held constant for approximately 30 min during each electrophoresis run. Quantitative accuracy was determined using known concentrations of standard reference compounds using Agilent MassHunter Software, version B.07.00.

**Determination of ammonium.** Ammonium was extracted and its concentration was determined with slight modifications to the method reported by (12). Shoots were harvested, immediately frozen with liquid $N_2$ and stored at −80°C until use. Frozen samples were ground with a Multi-Beads Shocker (Yasui Kikai Corp.) using zirconia beads (diameter, 5 mm). One ml of 0.1 N HCl and 500 μl of chloroform were added to the frozen powder, followed by vortexing for 15 min. The mixture was centrifuged at 12,000 ×*g* at 8°C for 10 min. The aqueous phase was transferred to a microtube containing 50 mg of acid-washed activated charcoal (No. 035-18081; Wako, Osaka, Japan). The mixture was vortexed and centrifuged at 20,400 ×*g* at 8°C for 10 min. The ammonium content of the supernatant was spectroscopically determined using an Ammonia Test Kit (No. 277-14401, Wako) according to the manufacturer's instructions.

**Determination of $H^+$ concentration in a water extract of shoots.** For Fig. 5*C*, plants grown on media containing 10 mM ammonium or 10 mM nitrate were submerged in 5 mL of the corresponding liquid medium in the presence or absence of 1 mM methionine sulfoximine (MSX). The plants were returned to their original positions on each medium and incubated for 5 h in the light. Shoots were harvested, immediately frozen with liquid $N_2$ and stored at −80°C until use. For Fig. 6*A* and *SI Appendix*, Fig. S9*E*, the shoots were harvested without prior incubation in liquid media, immediately frozen with liquid $N_2$ and stored at −80°C until use. Ten volumes of $H_2O$ for Fig. 5*C* and *SI Appendix*, Fig. S9*E* and 40 volumes of $H_2O$ for Fig. 6*A* were added to the frozen powder followed by centrifugation at 10,000 ×*g* at room temperature for 10 min. The pH of the supernatant was measured with a portable pH meter (B212, Horiba, Ltd.). The values were converted to proton concentrations.

**Determination of $H^+$ efflux from shoots.** Excised shoots from the plants grown on media containing 10 mM ammonium or 10 mM nitrate were submerged in 100 volumes of the corresponding liquid medium without MES buffer in the presence or absence of 1 mM methionine sulfoximine (MSX). The submerged shoots were incubated in liquid media for 5 h in the light. The pH changes in the incubation media were measured with a portable pH meter (B212, Horiba, Ltd.). The changes in pH were converted to changes in proton concentrations.

**Determination of $H^+$ efflux from mesophyll protoplasts**. Mesophyll protoplasts were prepared with slight modifications to the method reported by (13). In one dish, 20 plants were grown for 7 d on media containing 2.5 mM ammonium as the sole N source at pH 6.7 (7) in a vertical position. Ten plants were transferred to fresh medium with the same composition and grown for another 7 d. Two plants were further transferred to a medium containing 10 mM ammonium and grown for 3 d in a horizontal position. The adaxial epidermal surface was affixed to plastic tape, whereas the abaxial epidermal surface was attached to a strip of 15-mm wide Scotch transparent tape (3 M, St. Paul, MN, USA). The Scotch tape was then carefully pulled away from the plastic tape, peeling away the abaxial epidermal surface cell layer. The plastic tape samples with adhering peeled leaves were transferred to 1.5 mL microtubes containing 1 mL of an enzyme solution [0.75% (w/v) cellulase 'Onozuka' R10 (Yakult, Tokyo, Japan), 0.25% (w/v) macerozyme 'Onozuka' R10 (Yakult), 0.4 M mannitol, 8 mM $CaCl_2$, and 5 mM MES, pH 5.6]. The tube was slowly rotated for 20 min at room temperature. The liberated protoplasts were filtered through a 30-μm nylon filter (NY30-HD, SEMITEC, Osaka, Japan), and collected by centrifugation at 100 ×*g* at 4˚C for 5 min. The protoplast suspensions of leaf samples from 16 plants were pooled. The supernatant was removed, and the protoplasts were re-suspended in 1 mL of a wash solution [0.4 M mannitol and 8 mM $CaCl_2$, pH 6.5] followed by centrifugation at 100 ×*g* at 4˚C for 5 min. This washing step was repeated once more, afterwhich the protoplasts were suspended in 100 μl of the wash solution. Ten μl of protoplast suspension was mixed with 90 μl of an assay solution [0.4 M mannitol, 8 mM $CaCl_2$, 0.02% bromocresol purple (BCP) pH 6.5] supplemented with 5 mM $(NH_4)_2SO_4$ and 10 mM KCl in the presence or absence of 1 mM MSX, 10 mM KCl, or 10 mM $KNO_3$. The mixture was incubated at 23˚C in the light until color changes were observed.

**Observation of chloroplast ultrastructure in mesophyll cells with TEM.** For ultrastructural analyses, true leaves from Col, *ami2*, and *gln2* 5 d after the transfer to media containing 10 mM ammonium or 10 mM nitrate were analyzed by transmission electron microscopy according to a previously described method (14) with modification. Leaves were fixed with 4% (w/v) paraformaldehyde and 2% (v/v) glutaraldehyde in 50 mM sodium cacodylate buffer (pH 7.4) overnight at 4ºC. After the samples were post fixed with 1% (w/v) osmium tetroxide for 3 h and dehydrated in a graded methanol series, the samples were embedded in EPON812 resin. Micrographs were recorded using a JEM-1400 (JEOL) transmission electron microscope.

**Statistical and cluster analyses.** All statistical analyses were conducted using R software, version 2.15.3. Hierarchical clustering was performed using Gene Cluster software, version 3.0, with “Correlation (uncentered)” as the similarity metric and “Average linkage” as the clustering method. The results were visualized using Java TreeView software, version 1.1.6r4. Other details of analyses are provided in the Results and in the table and figure legends.

**Additional data table S1-4**

**Table S1.** The list of ammonium-inducible genes and ammonium-repressive genes in *A. thaliana* shoots.

**Table S2.** The list of acidic stress-inducible genes and acidic stress-repressive genes in *A. thaliana* shoots.

**Table S3.** Primer sequences.

**Table S4.** Raw data of microarray analysis.

**References**


1. Guan M, Møller IS, Schjoerring JK (2015) Two cytosolic glutamine synthetase isoforms play specific roles for seed germination and seed yield structure in *Arabidopsis*. *J Exp Bot* 66: 203-212.
2. Lothier J, et al. (2011) The cytosolic glutamine synthetase GLN1;2 plays a role in the control of plant growth and ammonium homeostasis in *Arabidopsis* rosettes when nitrate supply is not limiting. *J Exp Bot* 62: 1375-1390.
3. Sawaki Y, et al. (2009) STOP1 regulates multiple genes that protect Arabidopsis from proton and aluminum toxicities. *Plant Physiol* 150: 281-294.
4. Hachiya T, et al. (2011) Evidence for a nitrate-independent function of the nitrate sensor NRT1.1 in Arabidopsis thaliana. *J Plant Res* 124: 425-430.
5. Ichikawa T, et al. (2006) The FOX hunting system: an alternative gain-of-function gene hunting technique. *Plant J* 48 : 974-985.
6. Wang R, Xing X, Crawford N (2007) Nitrite acts as a transcriptome signal at micromolar concentrations in Arabidopsis roots. *Plant Physiol* 145: 1735-1745.
7. Okamoto Y, et al. (2019) Shoot nitrate underlies a perception of nitrogen satiety to trigger local and systemic signaling cascades in *Arabidopsis thaliana*. *Soil Sci Plant Nutr* 65: 56-64.
8. Sakakibara H, Kawabata S, Hase T, Sugiyama T (1992) Differential effects of nitrate and light on the expression of glutamine synthetases and ferredoxin-dependent glutamate synthase in maize. *Plant Cell Physiol* 33: 1193-1198.
9. Tabata R, et al. (2014) Perception of root-derived peptides by shoot LRR-RKs mediates systemic N-demand signaling. *Science* 346: 343-346.
10. Miyagi A, et al. (2010) Principal component and hierarchical clustering analysis of metabolites in destructive weeds; polygonaceous plants. *Metabolomic* 6: 146-155.
11. Miyagi K, et al. (2019) Oxalate contents in leaves of two rice cultivars grown at a free-air $CO_2$ enrichment (FACE) site. *Plant Prod Sci* 22: 407-411.
12. Hachiya T, et al. (2016) Arabidopsis root-type ferredoxin: NADP (H) oxidoreductase 2 is involved in detoxification of nitrite in roots. *Plant Cell Physiol* 57: 2440-2450.
13. Endo M, Shimizu H, Araki T (2016) Rapid and simple isolation of vascular, epidermal and mesophyll cells from plant leaf tissue. *Nat protoc* 11: 1388.

14. Toyooka K, Okamoto T, Minamikawa T (2000) Mass transport of proform of a KDEL-tailed cysteine proteinase (SH-EP) to protein storage vacuoles by endoplasmic reticulum–derived vesicle is involved in protein mobilization in germinating seeds. *J Cell Biol* 148: 453-464.

### Supplementary Figure S1

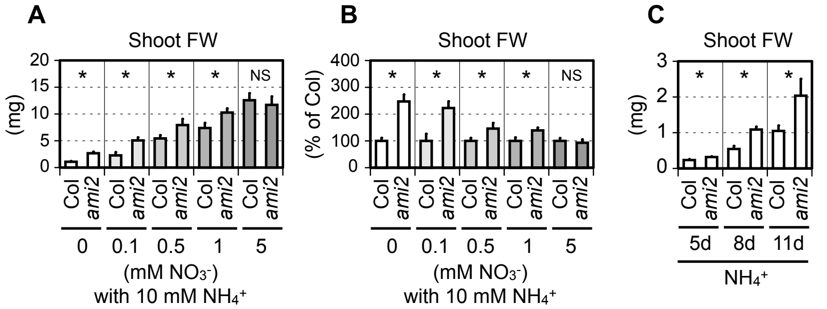


**Fig. S1** The *ami2* mutant exhibited enhanced ammonium tolerance during early stages of development. (*A*) FW of shoots and (*B*) relative FW (relative to Col as one) from Col and *ami2* grown on media containing 10 mM ammonium with varying concentrations of nitrate (mean ± SD; n = 10). Six shoots from one plate constituted a single biological replicate. (*C*) FW of shoots from Col and *ami2* grown on media containing 10 mM ammonium for 5 (mean ± SD; n = 5), 8 (mean ± SD; n = 5), or 11 d (mean ± SD; n = 10). Six shoots or roots from one plate constituted a single biological replicate. (*A*-*C*) Welch's *t*-test was run at α = 0.05; *$p < 0.05$.

### Supplementary Figure S2

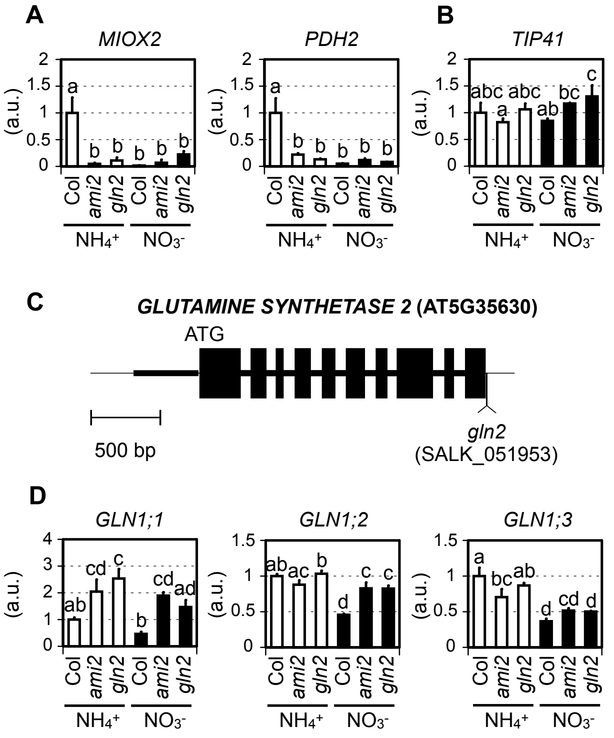


**Fig. S2** *GLN2* is a causative gene for ammonium toxicity. (*A*, *B*, *D*) Relative transcript levels of ammonium-inducible genes (13) (*A*), a house-keeping gene *TIP41* (*B*), and the *GLN1* gene (*D*) in the shoots of Col, *ami2*, and *gln2 3* d after transfer to media containing 10 mM ammonium (mean ± SD; n = 3). Six shoots from two plates constituted a single biological replicate. Tukey-Kramer's multiple comparison test was conducted at a significance level of $P < 0.05$ only when a one-way ANOVA was significant at $P < 0.05$. Different letters denote significant differences. (*C*) Genomic structure of the *GLN2* gene (AT5G35630). Black boxes, bold lines, and thin lines indicate exons, introns, and UTRs, respectively. T-DNA in *gln2* (SALK_051953) is inserted within the 3-UTR.

### Supplementary Figure S3

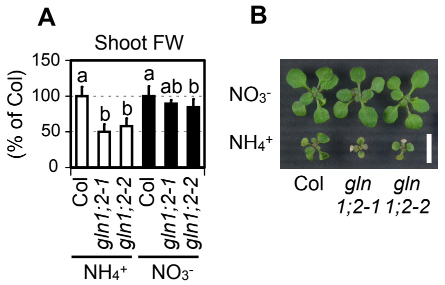


**Fig. S3** A deficiency in GLN1;2 causes ammonium hypersensitivity. (*A*) FW of shoots from Col, *gln1;2-1*, and *gln1;2-2* 7 d after transfer to media containing 10 mM ammonium or 10 mM nitrate (mean ± SD; n = 10). One shoot from one plate constituted a single biological replicate. Tukey-Kramer's multiple comparison test was conducted at a significance level of $P < 0.05$ only when a one-way ANOVA was significant at $P < 0.05$. Different letters denote significant differences. (*B*) A representative photograph of 14-d-old shoots is shown. The scale bar represents 10 mm.

### Supplementary Figure S4

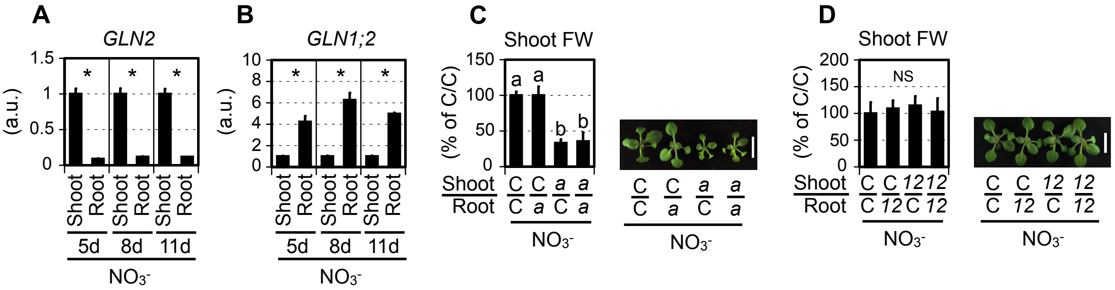


**Fig. S4** Shoot *GLN2* is crucial for shoot growth when nitrate is the N-source. (*A*) Relative transcript levels of *GLN2* in the shoots and roots of Col grown on 10 mM nitrate for 5, 8, or 11 d (mean ± SD; n = 3). (*B*) Relative transcript levels of *GLN1;2* in the shoots and roots of Col grown on media containing 10 mM nitrate for 5, 8, or 11 d (mean ± SD; n = 3). (*A*, *B*) Twelve shoots and roots from one plate constituted a single biological replicate. Welch's *t*-test was run at α = 0.05; $*p < 0.05$. (*C*) FW of shoots from reciprocally-grafted plants between Col (C) and *ami2* (a) 7 d after transfer to media containing 10 mM nitrate (mean ± SD; n = 3). (*D*) FW of shoots from reciprocally-grafted plants between Col (C) and *gln1.2-1* (12) 7 d after transfer to media containing 10 mM nitrate (mean ± SD; n = 8). (*C*, *D*) One shoot from one plate constituted a single biological replicate. Tukey-Kramer's multiple comparison test was conducted at a significance level of $P < 0.05$ only when a one-way ANOVA was significant at $P < 0.05$. Different letters denote significant differences. Representative photograph of shoots 7 d after transfer to media containing 10 mM nitrate are shown. The scale bar represents 10 mm.

### Supplementary Figure S5

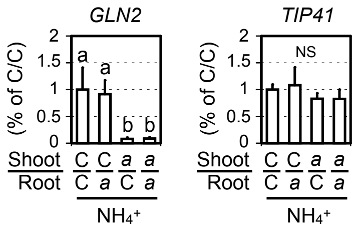


**Fig. S5** Root expression of *GLN2* does not affect the transcript levels of shoot *GLN2*. Relative transcript levels of *GLN2* and *TIP41*, a house-keeping gene, in the shoots of reciprocally-grafted plants between Col (C) and *ami2* (a) 3 d after transfer to media containing 10 mM ammonium (mean ± SD; n = 3). Three shoots from three plates constituted a single biological replicate. Tukey-Kramer's multiple comparison test was conducted at a significance level of $P < 0.05$ only when a one-way ANOVA was significant at $P < 0.05$. NS denotes not significant.

### Supplementary Figure S6

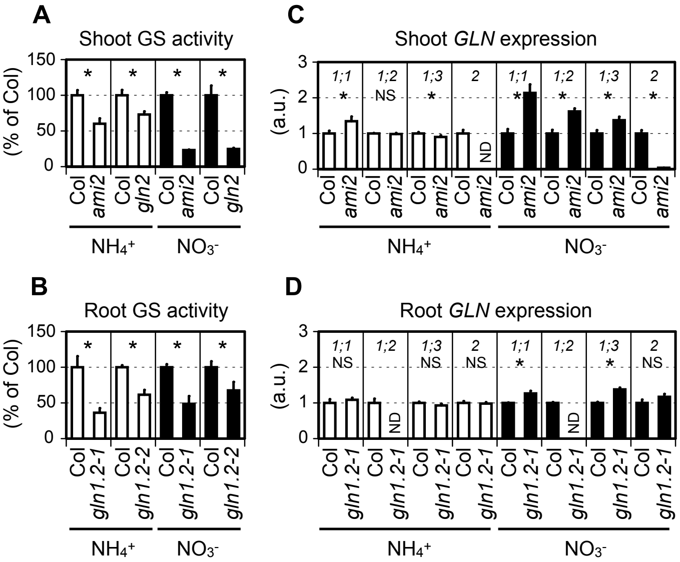


**Fig. S6** Deficiencies in *GLN2* and *GLN1;2* reduce GS activities in shoots and roots, respectively. (*A*) Shoot GS activities in Col, *ami2*, and *gln2* grown on media containing 10 mM ammonium or 10 mM nitrate for 5 d (mean ± SD; n = 3). Thirty-seven shoots from one plate constituted a single biological replicate. (*B*) Root GS activities in Col, *gln1;2-1*, and *gln1;2-2* grown on media containing 10 mM ammonium or 10 mM nitrate for 5 d (mean ± SD; n = 3). Thirty-seven shoots from one plate constituted a single biological replicate. (*C*) Relative transcript levels of *GLN1;1*, *GLN1;2*, *GLN1;3*, and *GLN2* in the shoots of Col and *ami2* grown on 10 mM ammonium or 10 mM nitrate for 5 d (mean ± SD; n = 3). Thirty-seven shoots from one plate constituted a single biological replicate. (*D*) Relative transcript levels of *GLN1;1*, *GLN1;2*, *GLN1;3*, and *GLN2* in the roots of Col and *gln1;2-1* grown on media containing 10 mM ammonium or 10 mM nitrate for 5 d (mean ± SD; n = 3). Thirty-seven shoots from one plate constituted a single biological replicate. (*A-D*) Welch's *t*-test was run at α = 0.05; *$p < 0.05$. NS denotes not significant.

### Supplementary Figure S7

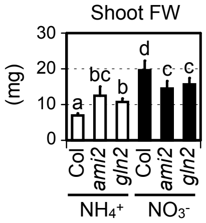


**Fig. S7** *GLN2* deficiency enhances ammonium insensitivity. FW of shoots from Col, *ami2*, and *gln2* 5 d after transfer to media containing 10 mM ammonium or 10 mM nitrate (mean ± SD; n = 6). Three shoots from one plate constituted a single biological replicate. Tukey-Kramer's multiple comparison test was conducted at a significance level of $P < 0.05$ only when a one-way ANOVA was significant at $P < 0.05$. Different letters denote significant differences.

### Supplementary Figure S8

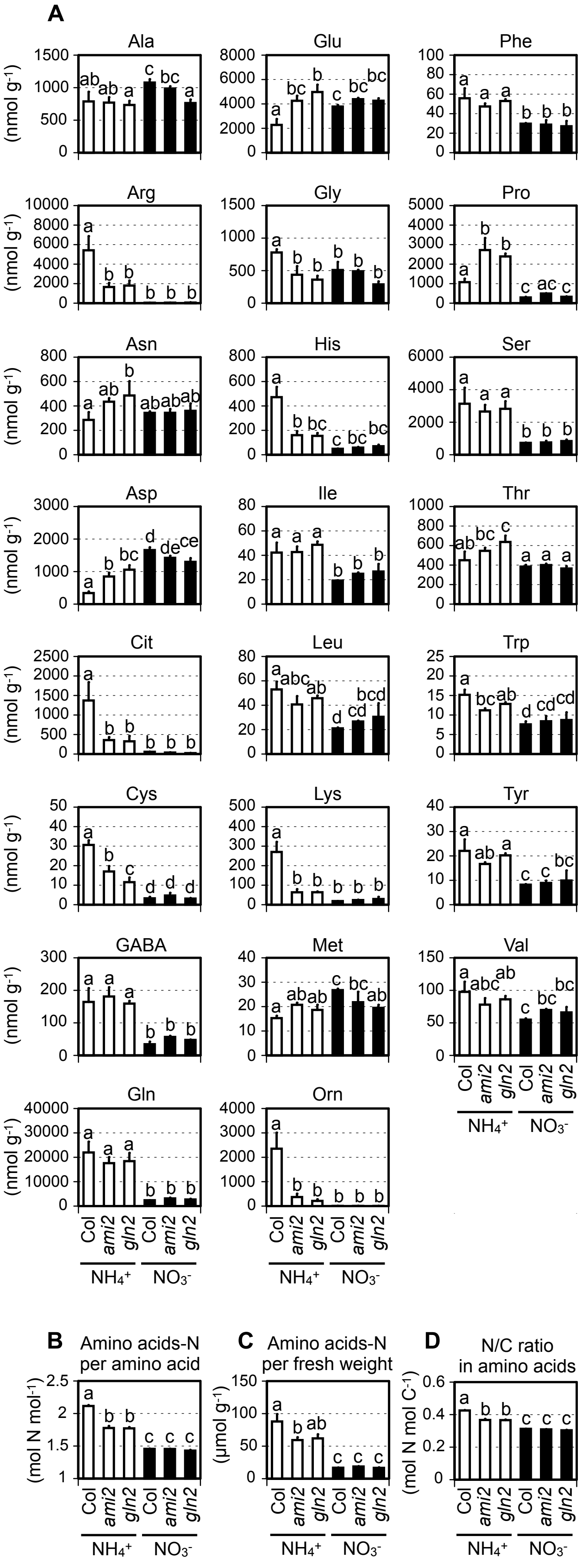


**Fig. S8** *GLN2* deficiency reduces ammonium-N incorporation into amino acids when plants are grown on ammonium. (*A*-*D*) Amino acid content (*A*), total amino acid-N concentration per amino acid (mol N $mol^{-1}$) (*B*), total amino acid-N concentration per fresh weight (μmol $g^{-1}$) (*C*), and molar ratios of N to C in total amino acids (mol N mol $C^{-1}$) (*D*) in the shoots of Col, *ami2*, and *gln2* 5 d after transfer to media containing 10 mM ammonium or 10 mM nitrate. Six shoots from two independent plates constitute one biological replicate. Three biological replicates were sampled separately three times (mean ± SE; n = 3). Tukey-Kramer's multiple comparison test was conducted at a significance level of $P < 0.05$ only when a one-way ANOVA was significant at $P < 0.05$. Different letters denote significant differences.

### Supplementary Figure S9

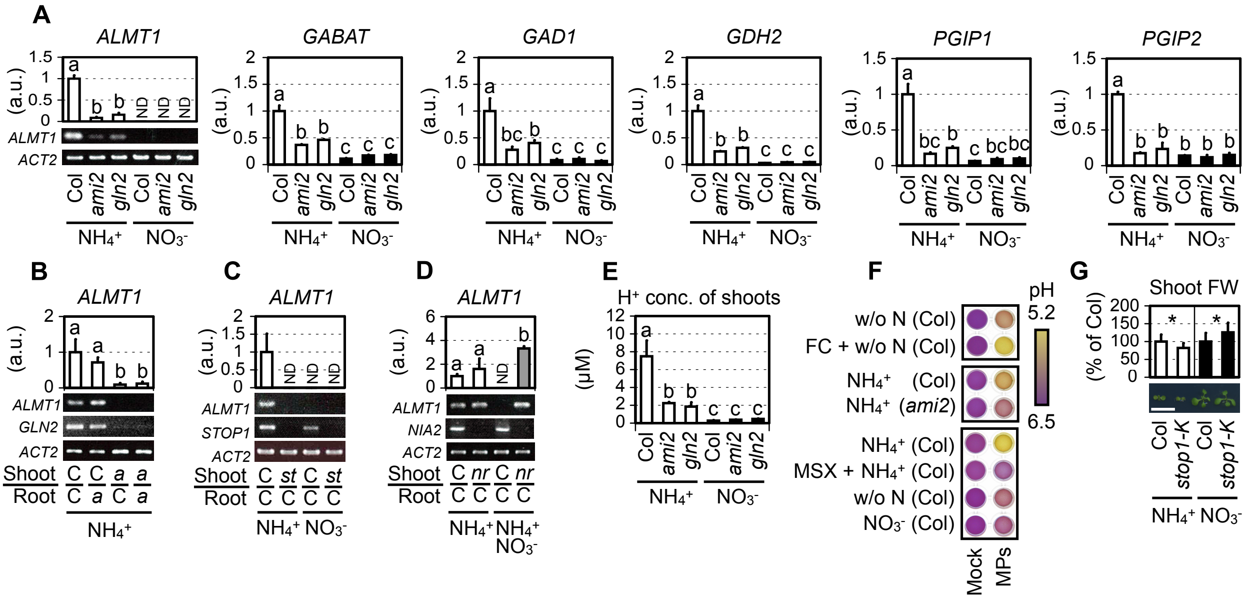


**Fig. S9** Ammonium assimilation by *GLN2* causes acidic stress. (*A*) Relative transcript levels of acidic stress-inducible genes (26, 27) in the shoots of Col, *ami2*, and *gln2* 3 d after transfer to media containing 10 mM ammonium or 10 mM nitrate (mean ± SD; n = 3). The transcript levels of *ALMT1* were evaluated both by reverse transcription-qPCR and semi-quantitative RT-PCR with agarose gel electrophoresis. *ACTIN2* (*ACT2*) was the internal standard. Three shoots from one plate constituted a single biological replicate. (*B*) Relative transcript levels of *ALMT1* in the shoots from reciprocally-grafted plants between Col (C) and *ami2* (a) 3 d after transfer to media containing 10 mM ammonium (mean ± SD; n = 3). Three shoots from three plates constituted a single biological replicate. The transcript levels of *ALMT1*, *GLN2*, and *ACTIN2* were evaluated by semi-quantitative RT-PCR with agarose gel electrophoresis. (*C*) Relative transcript levels of *ALMT1* in the shoots from grafted plants between Col (C) and *stop1-KO* (st) 3 d after transfer to media containing 10 mM ammonium or 10 mM nitrate (mean ± SD; n = 7). Two shoots from two plates constituted a single biological replicate. The transcript levels of *ALMT1*, *STOP1*, and *ACTIN2* were evaluated by semi-quantitative RT-PCR with agarose gel electrophoresis. (*D*) Relative transcript levels of *ALMT1* in the shoots from grafted plants between Col (C) and the *NR*-null mutant (nr) 3 d after transfer to media containing 10 mM ammonium ($NH_4^+$) or 2.5 mM nitrate and 10 mM ammonium ($NH_4^+$ $NO_3^-$) (mean ± SD; n = 3). One shoot from one plate constituted a single biological replicate. The transcript levels of *ALMT1*, *NIA2*, and *ACTIN2* were evaluated by semi-quantitative RT-PCR with agarose gel electrophoresis. (*E*) Proton concentrations of water extracts of shoots from Col, *ami2*, and *gln2* 5 d after transfer to media containing 10 mM ammonium or 10 mM nitrate (mean ± SD; n = 4). Six shoots from two plates constituted a single biological replicate. (*F*) Qualitative evaluation of proton efflux from mesophyll protoplasts (MPs) prepared from Col and *ami2* to liquid media containing 10 mM ammonium ($NH_4^+$), no N (w/o N), or 10 mM nitrate ($NO_3^-$) in the presence of 0.02% (w/v) bromocresol purple adjusted to pH 6.7. Fusicoccin (FC), an irreversible activator of plasma-membrane $H^+$-ATPase, was added at a final concentration of 1 M as a control. A representative photograph is shown. (*G*) FW of shoots from Col and *stop1-KO* (*stop1-k*) (mean ± SD; n = 5). In one dish, five seedlings of each line grown on media containing 2.5 mM ammonium at pH 6.7 for 5 d were transferred to media containing 10 mM ammonium or 10 mM nitrate and further grown for 6 d. Welch's *t*-test was run at α = 0.05; **p* < 0.05. A representative photograph of 11-d-old shoots is shown. The scale bar represents 10 mm. (*A-E*) Tukey-Kramer's multiple comparison test was conducted at a significance level of *P* < 0.05 only when a one-way ANOVA was significant at *P* < 0.05. Different letters denote significant differences.

### Supplementary Figure S10

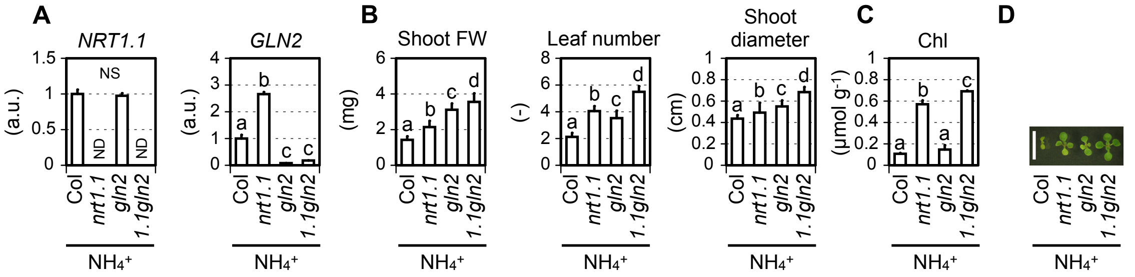


**Fig. S10** Deficiencies in *NRT1.1* and *GLN2* additively enhance ammonium insensitivity. (*A*) Relative transcript levels of *NRT1.1* and *GLN2* in seedlings of Col, *nrt1.1*, *gln2*, and *nrt1.1gln2* grown on media containing 10 mM ammonium for 5 d (mean ± SD; n = 3). Thirty-seven seedlings from one plate constituted a single biological replicate. (*B*) Shoot FW, leaf number, and shoot diameter of Col, *nrt1.1*, *gln2*, and *nrt1.1gln2* (mean ± SD; n = 19-20). In one dish, three seeds of each line of Col, *nrt1.1*, *gln2*, and *nrt1.1gln2* were grown on media containing 10 mM ammonium for 11 d. (*C*) Chlorophyll content in the shoots of Col, *nrt1.1*, *gln2*, and *nrt1.1gln2* grown on media containing 10 mM ammonium for 5 d (mean ± SD; n = 3). Thirty-seven shoots constituted a single biological replicate. (*D*) A representative photograph of 11-d-old shoots is shown. The scale bar represents 10 mm. (*A*-*C*) Tukey-Kramer's multiple comparison test was conducted at a significance level of $P < 0.05$ only when a one-way ANOVA was significant at $P < 0.05$. Different letters denote significant differences.

### Supplementary Figure S11

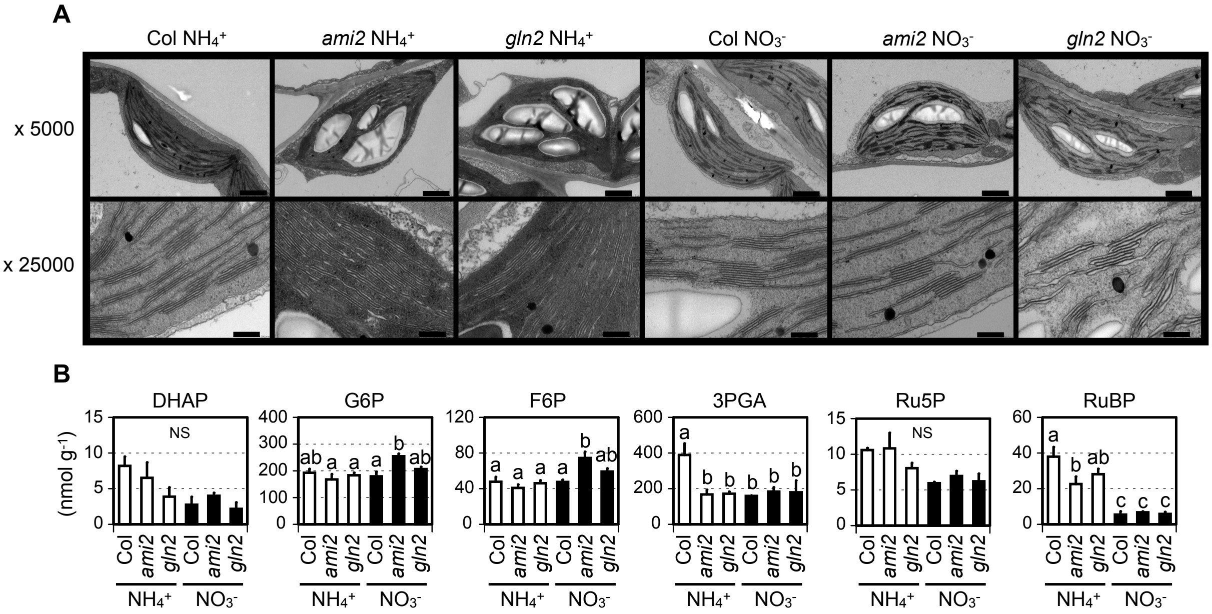


**Fig. S11** Abnormal chloroplast membrane structure and depletion of photosynthetic intermediates were not observed under conditions of ammonium toxicity as previously reported (28). (*A*) Representative micrographs of chloroplast ultrastructure of mesophyll cells in true leaves of Col, *ami2*, and *gln2* 5 d after transfer to media containing 10 mM ammonium or 10 mM nitrate. Upper and lower panels represent 5,000- and 25,000-fold magnifications, respectively. The scale bars at upper and lower panels represent 1 µm and 200 nm, respectively. (*B*) Contents of Calvin-Benson cycle intermediates in shoots of Col, *ami2*, and *gln2* 5 d after transfer to media containing 10 mM ammonium or 10 mM nitrate. Six shoots from two independent plates constitute one biological replicate. Three biological replicates were sampled separately three times (mean ± SE; n = 3). Tukey-Kramer's multiple comparison test was conducted at a significance level of $P < 0.05$ only when a one-way ANOVA was significant at $P < 0.05$. Different letters denote significant differences.

### Supplementary Figure S12

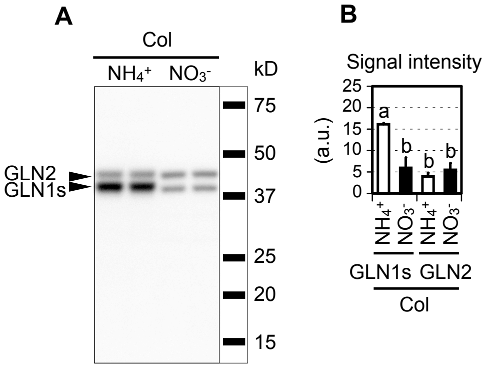


**Fig. S12** The GLN1 protein content in the shoot was higher in plants grown on media containing ammonium than those grown on nitrate-containing media. (*A*) Immunodetection of GLN1s and GLN2 isoproteins with specific antisera raised against maize GS following SDS-PAGE and immunoblotting of total protein extracts of Col shoots 5 d after transfer to media containing 10 mM ammonium or 10 mM nitrate. The positions of the molecular weight markers are shown on the right. (*B*) Signal intensities corresponding to GLN1s and GLN2 isoproteins (mean ± SD; n = 3). Three shoots from one plate constituted a single biological replicate. The intensity was quantified by ImageJ software, version 10.2. Tukey-Kramer's multiple comparison test was conducted at a significance level of $P < 0.05$ only when a one-way ANOVA was significant at $P < 0.05$. Different letters denote significant differences.
