## Supplementary material for "Excessive assimilation of ammonium by plastidic glutamine synthetase is a major cause of ammonium toxicity in *Arabidopsis thaliana*": Supple Tables

**Supplementary Table S1** The list of ammonium-inducible genes and ammonium-repressive genes in *A. thaliana* shoots (Log2 |FC| ≧ 1, Li et al. 2012 Plant Physiol.). In our experiments, the Log2 (ami2/Col) on gene expression was calculated, only when the calls were positive in two independent experiments.

**Ammonium-inducible genes in shoots**

| AGI No. | Log2 (NH4+ treatment/Control) (Li et al. 2012) | Log2 (ami2/Col) (Experiment 1 in the present paper) | Log2 (ami2/Col) (Experiment 2 in the present paper) |
|---|---|---|---|
| AT1G02470 | 2.530907744 | -1.26121941 | -0.668723361 |
| AT1G05680 | 3.221190204 | -1.7431746 | -0.615990443 |
| AT1G09500 | 3.060624649 | -2.144746868 | 0.223459899 |
| AT1G17180 | 2.420312664 | -1.555091971 | -1.040902186 |
| AT1G22500 | 2.315795701 | -1.880496789 | -1.174334221 |
| AT1G31750 | 1.80620102 | -0.457303226 | -0.632665926 |
| AT1G49150 | 3.964264542 | - | - |
| AT1G67810 | 2.033973449 | -0.656193143 | -0.584625172 |
| AT1G73120 | 4.156507954 | -0.495647877 | -1.034314726 |
| AT2G02930 | 1.217810797 | -0.302966081 | -0.371165347 |
| AT2G03130 | 4.426149973 | - | - |
| AT2G04050 | 2.245196305 | -4.070779614 | -2.530819565 |
| AT2G04070 | 3.151802137 | -1.569481512 | -0.927451917 |
| AT2G21640 | 2.156931318 | -1.774351898 | -1.693480448 |
| AT2G27830 | 1.762050977 | -1.345320912 | -0.762671307 |
| AT2G29490 | 1.414340834 | -0.864913818 | -0.196707603 |
| AT2G29500 | 1.599976909 | -1.546386358 | -0.910774311 |
| AT2G38210 | 2.144255394 | -0.366675069 | -0.507224206 |
| AT2G41730 | 2.087300691 | -1.463208979 | -1.238750033 |
| AT2G47520 | 3.607931191 | - | - |
| AT3G04000 | 1.991087193 | -1.84433859 | -0.848483211 |
| AT3G06700 | 2.025120022 | -0.095281543 | 0.225941489 |
| AT3G13080 | 1.375396088 | -0.574869023 | -0.203038987 |
| AT3G20340 | 1.952278888 | - | - |
| AT3G28740 | 1.912624188 | -0.924188799 | -0.933480552 |
| AT3G46230 | 3.076493905 | 1.340340132 | 0.465110095 |
| AT3G61620 | 4.977044846 | -0.44366406 | -0.544603053 |
| AT3G61630 | 2.285280248 | -1.894888165 | -1.650513396 |
| AT4G21830 | 2.14667779 | -3.508618131 | -2.62083789 |
| AT4G23750 | 1.283811398 | - | - |
| AT4G28020 | 1.451308313 | -0.419153607 | 0.056734407 |
| AT4G29740 | 1.575449681 | -0.118510455 | -0.521368735 |
| AT4G30670 | 2.233346681 | -1.394732514 | -0.780967137 |
| AT5G09570 | 3.096250614 | -0.718858919 | -0.59612186 |
| AT5G19120 | 1.986372598 | -1.031337968 | -1.25743612 |
| AT5G23730 | 1.280229635 | 0.16801995 | 0.046088926 |
| AT5G27920 | 1.876631167 | -0.481946043 | -0.624071538 |
| AT5G40690 | 2.089252045 | -0.380992291 | -0.049238008 |
| AT5G49560 | 1.952859871 | -0.8420823 | -0.908129948 |
| AT5G50760 | 1.905885499 | -3.027037419 | -2.420482521 |
| AT5G54530 | 1.731044898 | -0.452705755 | -0.34044242 |
| AT5G57980 | 1.753085052 | -0.402546555 | -1.155855419 |
| AT5G66650 | 2.119369256 | -2.848520671 | -2.182046498 |

**Ammonium-repressive genes in shoots**

| AGI code | Log2 (NH4+ treatment/Control) (Li et al. 2012) | Log2 (ami2/Col) (Experiment 1 in the present paper) | Log2 (ami2/Col) (Experiment 2 in the present paper) |
|---|---|---|---|
| AT1G04770 | -1.655520841 | 0.682163086 | 0.102702952 |
| AT1G14360 | -2.12651421 | 0.009272223 | 0.208501095 |
| AT1G21910 | -1.117544191 | 0.110252477 | 0.082532066 |
| AT1G22160 | -2.161460162 | 1.086254915 | 0.777909095 |
| AT1G68600 | -2.950932158 | -0.63842075 | 0.542862749 |
| AT1G75380 | -1.569049136 | 0.291294901 | 0.348994483 |
| AT1G80130 | -2.19923552 | 0.904346006 | 1.490029083 |
| AT2G02810 | -2.143545317 | 0.331967667 | -0.241421893 |
| AT2G14560 | -1.737198118 | 1.226727801 | 1.543477237 |
| AT3G08970 | -1.729261846 | -0.084841788 | 0.366127899 |
| AT3G23250 | -2.134164331 | -0.86697458 | -0.88502052 |
| AT3G24090 | -1.576735165 | 0.585620215 | 0.302009589 |
| AT3G26210 | -1.871473522 | 1.50577577 | 2.063805109 |
| AT3G45970 | -2.174803168 | -0.084366003 | 0.625197474 |
| AT3G53668 | -2.115082045 | 0.823321671 | 0.187278568 |
| AT3G57240 | -1.593190272 | 1.536587727 | 0.94753258 |
| AT4G02330 | -2.989247529 | -0.246475129 | 0.269147705 |
| AT4G15210 | -3.232677217 | -0.630381625 | 0.548966714 |
| AT4G21760 | -1.779471891 | 1.902758162 | 0.566896316 |
| AT4G23140 | -2.345261074 | 4.023975273 | 3.825475764 |
| AT4G23160 | -2.045229484 | 4.023975273 | 3.825475764 |
| AT4G23210 | -1.702240695 | 0.279673543 | -0.048136522 |
| AT4G23810 | -2.013759706 | 3.420728089 | 1.704664304 |
| AT4G25810 | -3.486855567 | -1.512796307 | -1.375792122 |
| AT4G39210 | -1.859046258 | 0.272907939 | 0.629223873 |
| AT5G03160 | -2.188264274 | -0.265036759 | 0.244738887 |
| AT5G24660 | -1.349765292 | 0.649869446 | 0.224545162 |
| AT5G25440 | -1.833581643 | 0.978800834 | 0.933445205 |
| AT5G26220 | -3.496802685 | 0.206534509 | -1.47518128 |
| AT5G44380 | -1.641589185 | - | - |
| AT5G48370 | -1.796039105 | 0.103950964 | 0.355190549 |
| AT5G49520 | -2.279332586 | - | - |
| AT5G55050 | -1.966109801 | -1.625436041 | -0.15467767 |

**Supplementary Table S2** The list of acidic stress-inducible genes and acidic stress-repressive genes in *A. thaliana* shoots (Log2 |FC| ≧ 1, Lager et al. 2010 Plant Cell Environ.). In our experiments, the Log2 (ami2/Col) on each gene expression was calculated, only when the calls were positive in two independent experiments.

**Early acidic stress-inducible genes**

| AGI No. | Log2 (pH4.5 treatment/pH 6.0 Control) (Lager et al. 2010) | Log2 (ami2/Col) (Experiment 1 in the present paper) | Log2 (ami2/Col) (Experiment 2 in the present paper) |
|---|---|---|---|
| AT5G57560 | 5.527097871 | -1.671621478 | -1.466712848 |
| AT1G35140 | 4.763262797 | -2.451161746 | -1.010239185 |
| AT5G52760 | 4.175273261 | 0.474936201 | -0.259275542 |
| AT4G38400 | 3.658613589 | -0.479204178 | -0.014555836 |
| AT4G25810 | 3.592046004 | -1.512796307 | -1.375792122 |
| AT4G08950 | 3.496917844 | -1.407234886 | -0.898631283 |
| AT5G54490 | 3.350534519 | 0.242284758 | 0.317981479 |
| AT4G38420 | 3.139316992 | -0.200907305 | -0.124498542 |
| AT5G66650 | 3.039246678 | -2.848520666 | -2.182046496 |
| AT4G37240 | 2.822147002 | 0.133926803 | -0.097526349 |
| AT4G27280 | 2.809344174 | -0.280407681 | 0.062755973 |
| AT4G24570 | 2.678358836 | 0.046960012 | -0.142263905 |
| AT5G63790 | 2.617050269 | 0.045685372 | 0.45046784 |
| AT1G69930 | 2.590582018 | - | - |
| AT3G50930 | 2.566591488 | -0.156588929 | -0.116087968 |
| AT5G52750 | 2.493993848 | -0.040947761 | 0.388992704 |
| AT1G59660 | 2.491241379 | 0.72156301 | 0.388087529 |
| AT1G80840 | 2.447669755 | 0.646505806 | 0.35011764 |
| AT2G41640 | 2.429509609 | -0.370680273 | -0.485210547 |
| AT1G61340 | 2.418881471 | -0.294752242 | 0.256825918 |
| AT3G55840 | 2.415158996 | - | - |
| AT5G60350 | 2.339686527 | - | - |
| AT1G76640 | 2.307465859 | - | - |
| AT2G32190 | 2.286472699 | -0.486360745 | -0.242515038 |
| AT5G37770 | 2.255420961 | 0.184847711 | -0.002497176 |
| AT3G45970 | 2.233589913 | -0.084366003 | 0.625197473 |
| AT4G38410 | 2.189062274 | - | - |
| AT4G30290 | 2.178772512 | -0.081650289 | -0.186748732 |
| AT2G32020 | 2.177665556 | - | - |
| AT3G45960 | 2.150892241 | - | - |
| AT1G76650 | 2.140970509 | -0.627157418 | -0.356029088 |
| AT4G24110 | 2.118572368 | -1.686011502 | -0.843409637 |
| AT3G62720 | 2.11187186 | -0.512693127 | 0.156504486 |
| AT4G02330 | 2.069322953 | -0.246475129 | 0.269147705 |
| AT2G28400 | 2.063871361 | -1.555709774 | -0.363841737 |
| AT5G14730 | 2.052536442 | -0.17953735 | 0.049456387 |
| AT1G01720 | 2.041089664 | -0.369701517 | 0.216868375 |
| AT5G06860 | 2.034453144 | -2.266804489 | -1.591122045 |
| AT5G09980 | 2.010952788 | - | - |
| AT1G02810 | 2.000145655 | -0.601450624 | -0.63900234 |
| AT4G30280 | 1.994611159 | -1.280605572 | -0.949434083 |
| AT3G51680 | 1.969594953 | - | - |
| AT5G22530 | 1.932422358 | -1.377157616 | -1.677681513 |
| AT3G28180 | 1.929785527 | -0.561285587 | -0.032410959 |
| AT2G37430 | 1.924571958 | - | - |
| AT3G56400 | 1.922501118 | 0.917760155 | 0.963977794 |
| AT2G30130 | 1.918376647 | - | - |
| AT1G68410 | 1.89887547 | -0.737685682 | 0.366815093 |
| AT4G31800 | 1.873107745 | -0.869575431 | -0.96348513 |
| AT1G32920 | 1.866893015 | -1.065897785 | -0.11131988 |
| AT5G64870 | 1.865259112 | 0.513732196 | 0.06619849 |
| AT4G17490 | 1.829223064 | -0.462372228 | -0.112410914 |
| AT2G14960 | 1.828261898 | - | - |
| AT3G44735 | 1.814725058 | 0.547071901 | 0.472752997 |
| AT1G57560 | 1.813899082 | 0.499274036 | 0.343301746 |
| AT4G36500 | 1.798746008 | -0.306965223 | -0.069544195 |
| AT1G25400 | 1.754529859 | -0.353636955 | -0.14575869 |
| AT4G30430 | 1.733672206 | - | - |
| AT5G47220 | 1.728499799 | 0.029404011 | -0.548038083 |
| AT3G17690 | 1.723482155 | - | - |
| AT4G29780 | 1.722382394 | 0.122328759 | -0.335153033 |
| AT4G01870 | 1.717600431 | -0.498541264 | 0.281839037 |
| AT2G01300 | 1.710203768 | - | - |
| AT5G53980 | 1.695805372 | - | - |
| AT3G09280 | 1.667050621 | - | - |
| AT4G24160 | 1.665447599 | 0.12900758 | 0.498619207 |
| AT4G13180 | 1.657863402 | -0.46998569 | -0.037571917 |
| AT1G44830 | 1.654587335 | 2.962455631 | 0.675273272 |
| AT2G24600 | 1.651173854 | -0.450603559 | -0.15475735 |
| AT3G57450 | 1.635568462 | -0.591910166 | -0.242678684 |
| AT3G16150 | 1.635517969 | -0.251877438 | 0.560335529 |
| AT4G36010 | 1.631449736 | -0.673704434 | 0.134626338 |
| AT2G16900 | 1.630417209 | 0.132679876 | -0.082064537 |
| AT2G47550 | 1.621831034 | -1.166157188 | -0.394910799 |
| AT4G00080 | 1.618957391 | - | - |
| AT5G22570 | 1.613577145 | - | - |
| AT2G15490 | 1.61159448 | -1.795790988 | -1.280233391 |
| AT2G04500 | 1.595361509 | - | - |
| AT1G66180 | 1.594096978 | -1.382721029 | -0.957745165 |
| AT5G41080 | 1.588956395 | - | - |
| AT4G34710 | 1.572598142 | -1.226798339 | -1.081414305 |
| AT1G69920 | 1.565017488 | - | - |
| AT1G65310 | 1.562744592 | - | - |
| AT1G80240 | 1.561924522 | - | - |
| AT1G77450 | 1.540369449 | -0.859396143 | -0.256048317 |
| AT4G31940 | 1.534174698 | - | - |
| AT2G17230 | 1.532040305 | 0.348694484 | 0.005686626 |
| AT3G56880 | 1.512729875 | -0.078895292 | -0.173740672 |
| AT3G50060 | 1.494266092 | 0.296551532 | -0.066255073 |
| AT1G14550 | 1.491646252 | - | - |

| | | | |
|---|---|---|---|
| AT3G54150 | 1.49087154 | -1.165722566 | -0.94640932 |
| AT5G64310 | 1.490050922 | 0.088113828 | 0.463369509 |
| AT1G73010 | 1.48703199 | -0.543719518 | -0.478750399 |
| AT5G45110 | 1.481954449 | -0.248288473 | 0.974313356 |
| AT2G41100 | 1.476901409 | 0.146153215 | -0.19242011 |
| AT3G25790 | 1.463326493 | - | - |
| AT5G42380 | 1.450689518 | - | - |
| AT2G34930 | 1.449624167 | -0.946351336 | -0.486213251 |
| AT1G78340 | 1.441873147 | - | - |
| AT3G25780 | 1.433575882 | -1.216378212 | -1.236816129 |
| AT2G17880 | 1.429653466 | - | - |
| AT5G10830 | 1.426290992 | -0.27424188 | -0.111235241 |
| AT2G45400 | 1.424091649 | -0.615519968 | -0.414356501 |
| AT1G76520 | 1.413280169 | -0.837628857 | -0.423431459 |
| AT1G13600 | 1.407762328 | - | - |
| AT4G29610 | 1.403566424 | -0.207018437 | 0.440527846 |
| AT5G37260 | 1.392737302 | 0.311594381 | -0.060012671 |
| AT1G43910 | 1.392552297 | -0.522760765 | -0.933461881 |
| AT3G05320 | 1.39041628 | 0.411505808 | 0.117066264 |
| AT2G19800 | 1.389961252 | -2.855102827 | -2.53379774 |
| AT3G25730 | 1.383593126 | 0.328187085 | -0.51904519 |
| AT5G26200 | 1.378710616 | - | - |
| AT1G14480 | 1.375904045 | -0.257239547 | -0.032207014 |
| AT1G66160 | 1.374969337 | -0.771990723 | -0.677575935 |
| AT3G53400 | 1.372561093 | 0.170694234 | -0.025283743 |
| AT5G14760 | 1.363473726 | -0.036037528 | -0.38175964 |
| AT1G27730 | 1.362403784 | 0.457252831 | 0.037352142 |
| AT1G55920 | 1.353808388 | -0.137324231 | 0.138117727 |
| AT2G39980 | 1.353166077 | 1.030435094 | 0.487730839 |
| AT2G01180 | 1.351607877 | -0.674229839 | -0.307123625 |
| AT5G51190 | 1.348507799 | -0.154956411 | -0.601547413 |
| AT2G22880 | 1.346486183 | - | - |
| AT2G22470 | 1.332873794 | - | - |
| AT3G28580 | 1.328253838 | - | - |
| AT2G21210 | 1.319759534 | - | - |
| AT5G66210 | 1.304259922 | 0.331674733 | 0.010448533 |
| AT2G44840 | 1.298915212 | -1.855456064 | -1.327261539 |
| AT1G27200 | 1.284308606 | -0.912624741 | -0.827583383 |
| AT5G10380 | 1.275130716 | 0.225409123 | -0.148580176 |
| AT5G54380 | 1.2585512 | -0.296409231 | 0.254156964 |
| AT5G08350 | 1.249032053 | -0.309560496 | -0.339828202 |
| AT3G54810 | 1.248119262 | 0.465420447 | 0.024683365 |
| AT4G27652 | 1.234740277 | -0.31931508 | -0.158384219 |
| AT1G67810 | 1.232909722 | -0.656193143 | -0.584625171 |
| AT4G13340 | 1.231582058 | -0.7194127 | -0.58068882 |
| AT2G36220 | 1.225655652 | 0.65838357 | 0.431836788 |
| AT5G38700 | 1.224822364 | - | - |
| AT1G77640 | 1.223118185 | - | - |
| AT2G32140 | 1.217174713 | -0.736965594 | -0.689280796 |
| AT2G43290 | 1.217041638 | 0.628771186 | 0.431031899 |
| AT2G46400 | 1.213375877 | 0.093011232 | 0.017685624 |
| AT4G14680 | 1.2112142 | -0.895488824 | -0.743355383 |
| AT2G38470 | 1.210477508 | -0.066061053 | 0.231145827 |
| AT3G25600 | 1.210281286 | 0.431531008 | 0.757121122 |
| AT3G59080 | 1.209667724 | -0.916598249 | -0.153288225 |
| AT4G30470 | 1.20625076 | -0.5642115 | 0.052569218 |
| AT3G23170 | 1.190104987 | 0.203489604 | 0.730001646 |
| AT3G61190 | 1.188984113 | -0.093197889 | -1.163605405 |
| AT2G46600 | 1.168467877 | -0.356533388 | -0.155624812 |
| AT2G45080 | 1.162127845 | 1.946576963 | -0.18602146 |
| AT2G15480 | 1.161985901 | -0.243029777 | 0.15273524 |
| AT3G15540 | 1.157743939 | - | - |
| AT5G18470 | 1.155103084 | 0.831496793 | 0.518498514 |
| AT1G21910 | 1.154764206 | 0.110252477 | 0.082532067 |
| AT4G37400 | 1.154524903 | - | - |
| AT4G34150 | 1.152828546 | 0.568845724 | 0.09393277 |
| AT3G44630 | 1.146369224 | 0.288753947 | 0.295783657 |
| AT2G06850 | 1.144606451 | 1.123003311 | 0.257360171 |
| AT2G29120 | 1.141535124 | 0.060744191 | -0.50891931 |
| AT1G56060 | 1.139637736 | 0.40746428 | -0.120012979 |
| AT4G17460 | 1.136119759 | 1.032736786 | 0.622973225 |
| AT3G27360 | 1.135804302 | 0.344950104 | 0.267357739 |
| AT3G59220 | 1.134463931 | -1.985091109 | -0.732813951 |
| AT5G48570 | 1.131192085 | 1.339559455 | 0.931425003 |
| AT1G72940 | 1.128435674 | -0.922627388 | -0.46153787 |
| AT1G17180 | 1.127002904 | -1.555091971 | -1.040902187 |
| AT1G17620 | 1.126103195 | -0.287974931 | 0.285246512 |
| AT1G72920 | 1.125975654 | -0.849499871 | -0.281344815 |
| AT4G25390 | 1.124849513 | - | - |
| AT4G11780 | 1.123933202 | - | - |
| AT5G62020 | 1.123389928 | -0.488769625 | -0.448942839 |
| AT2G27080 | 1.121345197 | -0.302656723 | 0.308885057 |
| AT5G39670 | 1.118775537 | 0.520575113 | -0.081577589 |
| AT1G37130 | 1.118497479 | -0.756000943 | -0.535422857 |
| AT3G30775 | 1.10793146 | -2.544420874 | -1.949893863 |
| AT2G26440 | 1.103643874 | 0.31072215 | 0.149178943 |
| AT3G59350 | 1.097966915 | -0.489110086 | -0.198264522 |
| AT1G05000 | 1.095732214 | -1.491639062 | -0.927500478 |
| AT2G48080 | 1.090996946 | - | - |
| AT1G13540 | 1.084179394 | - | - |
| AT2G17070 | 1.079622507 | - | - |
| AT2G24610 | 1.079119486 | - | - |
| AT2G16060 | 1.077158528 | - | - |
| AT1G66400 | 1.076350523 | -0.042924326 | -0.031866454 |
| AT2G47160 | 1.06948429 | 0.92631887 | 0.446184824 |

| | | | |
|---|---|---|---|
| AT3G24530 | 1.069115827 | 0.1901409 | 0.268295311 |
| AT2G32030 | 1.067517925 | -0.75808502 | -0.656347134 |
| AT2G41380 | 1.064469649 | -1.82066817 | -1.134195148 |
| AT5G49280 | 1.062379762 | -1.699586145 | -0.650641734 |
| AT5G12940 | 1.061787101 | -0.588461642 | -0.870137248 |
| AT5G22890 | 1.061716475 | - | - |
| AT2G22500 | 1.058666426 | -0.061667776 | -0.070244486 |
| AT5G57340 | 1.057457674 | 0.734499257 | 1.045738663 |
| AT1G69680 | 1.056639047 | -0.143924861 | 0.29757469 |
| AT3G26980 | 1.053980621 | 0.045592577 | 0.039287567 |
| AT4G12720 | 1.05172895 | 0.172192687 | 0.072772185 |
| AT3G55980 | 1.047093757 | -0.015247932 | 0.198646362 |
| AT1G13590 | 1.046649736 | - | - |
| AT5G02220 | 1.045560324 | - | - |
| AT3G19680 | 1.042550888 | 0.123002703 | -0.203854591 |
| AT5G13190 | 1.034121009 | 0.041698851 | 0.010606117 |
| AT1G62500 | 1.031096748 | 0.885964757 | 0.242684654 |
| AT1G69840 | 1.026999613 | 0.189371956 | -0.000733222 |
| AT1G12740 | 1.026095534 | - | - |
| AT4G28085 | 1.015692456 | -0.802484112 | -1.394729773 |
| AT4G08040 | 1.01385316 | - | - |
| AT3G61640 | 1.011642333 | -0.181149439 | -0.218006682 |
| AT5G56080 | 1.010902656 | - | - |
| AT1G12950 | 1.005934295 | - | - |
| AT5G65300 | 1.005891603 | -0.125208355 | -0.001656366 |
| AT5G64300 | 1.003717501 | -0.215072127 | -0.231981824 |

**Early acidic stress-repressive genes**

| AGI No. | Log2 (pH4.5 treatment/pH 6.0 Control) (Lager et al. 2010) | Log2 (ami2/Col) (Experiment 1 in the present paper) | Log2 (ami2/Col) (Experiment 2 in the present paper) |
|---|---|---|---|
| AT5G02260 | -3.401703007 | 0.535131345 | 0.582237863 |
| AT3G26740 | -2.13975885 | 0.090057003 | 0.005919573 |
| AT5G45340 | -2.114755268 | - | - |
| AT1G16510 | -1.842964828 | 1.267421319 | 1.006734235 |
| AT2G45050 | -1.777881598 | - | - |
| AT1G80450 | -1.474034689 | - | - |
| AT3G07010 | -1.315597567 | - | - |
| AT5G03960 | -1.30911748 | - | - |
| AT2G44740 | -1.299345675 | - | - |
| AT1G71030 | -1.267371341 | 0.073571967 | -0.520347435 |
| AT2G25680 | -1.250333956 | 0.058569451 | 0.917259569 |
| AT1G20190 | -1.237813876 | 0.047379701 | 0.334253044 |
| AT4G13510 | -1.212370035 | 1.179105555 | 0.595781651 |
| AT2G30520 | -1.212041543 | 0.856382414 | 0.519887777 |
| AT1G13110 | -1.192682735 | 0.251925401 | 0.166990222 |
| AT5G15230 | -1.143807707 | 2.495380534 | 2.366497796 |
| AT5G44910 | -1.13081523 | - | - |
| AT5G57785 | -1.122729635 | 1.340822407 | 0.957166705 |
| AT4G10270 | -1.110506151 | 1.404024507 | 0.336126411 |
| AT4G33790 | -1.096815297 | - | - |
| AT5G42830 | -1.086240508 | - | - |
| AT1G67750 | -1.08336485 | 1.727287277 | 1.010414229 |
| AT2G02950 | -1.058169068 | -0.066572672 | -0.312277925 |
| AT5G15120 | -1.032909602 | - | - |
| AT1G48330 | -1.029177 | 1.803568319 | 1.096765255 |
| AT5G06270 | -1.02877219 | 0.35374132 | 0.384634077 |
| AT1G70710 | -1.026915882 | 0.862894956 | 0.23769881 |
| AT5G13930 | -1.026659554 | -0.82163532 | -0.473066749 |
| AT1G05300 | -1.009640463 | 0.006490516 | 0.558979946 |
| AT3G30180 | -1.008736062 | -0.018654051 | -0.24941075 |
| AT3G51240 | -1.005477082 | -0.946246397 | -0.356544305 |
| AT1G24530 | -1.001620433 | 1.741806857 | 0.394748879 |

**Supplementary Table S3** Primer sequences

| AGI No. | Gene name | Description | Primer 1 (5'->3') | Primer 2 (5'->3') | Reference |
|---|---|---|---|---|---|
| AT3G18780 | *ACT2* | RT-PCR | TGTCCTCCTCACTTTCATCAGC | CATCAATTCGATCACTCAGAGC | Kiba et al. (2013) |
| AT3G53750 | *ACT3* | Q-PCR | GGCTAACCGTGAGAAGATGA | CGACCTGCAAGATCAAGACG | Hachiya et al. (2016) |
| AT1G08430 | *ALMT1* | Q-PCR | GGCAGTGTGCCTACAGGATT | AGATTTGCATGCAGCTTGTG | The present study |
| AT1G08430 | *ALMT1* | RT-PCR | GGCCGACCGTGCTATACGAG | CATGAGTCCTGTGAACTCCC | Sawaki et al. (2009) |
| AT3G22200 | *GABAT* | Q-PCR | TGTTAGTCCGTGTTGCAGGTGA | CACTTCTTGTGCTGAGCCTTGA | Sawaki et al. (2009) |
| AT5G17330 | *GAD1* | Q-PCR | ACAGGAAGAAGACGAGTGGTAT | GAAACTTGGTTCAGTAATAAAGGGA | Sawaki et al. (2009) |
| AT5G07440 | *GDH2* | Q-PCR | CACTGGAAAGCCCATTGATCT | TGTCGCTTACTGCAACCACT | Patterson et al. (2010) |
| AT5G37600 | *GLN1;1* | Q-PCR | CATCAACCTTAACCTCTCAGACTCCACT | ACTTCAGCTGCAACATCAGGGTTGCTA | Ishiyama et al. (2004) |
| AT1G66200 | *GLN1;2* | Q-PCR | AGCCAAGCTTCTCGATCGCC | TGGAATGGAGCTGGTGCTCA | Hachiya et al. (2016) |
| AT3G17820 | *GLN1;3* | Q-PCR | GAAAGCACGAAACCGCAGA | GCGACTCCCCAAGAGAATG | The present study |
| AT5G35630 | *GLN2* | Q-PCR | TTGACCAGTTCTCATGGGGC | TTAGATGCTGGACGGCGATC | Hachiya et al. (2016) |
| AT5G35630 | *GLN2* | RT-PCR/gPCR | ATGGCTCAGATCTTAGCAGC | TTAAACATTCAAAGAAAGCTT | The present study |
| AT2G19800 | *MIOX2* | Q-PCR | GCTGTCGTTGGCGATACATTTC | AGGGTCGTGCCATTCTTCTTAG | Endres and Tenhaken (2009) |
| AT1G37130 | *NIA2* | Q-PCR | AAGGGAGGAACTGGATGGTT | CGGTACTGTATGCCCAACCT | Konishi and Yanagisawa (2011) |
| AT1G37130 | *NIA2* | RT-PCR | CTTACAAACCTCCCGTTCCA | GCATAATCGCCTCGGAAATA | The present study |
| AT1G12110 | *NRT1.1* | Q-PCR | AAACAAAAGCTGCCACACAC | GCACGATTTGTTTCACTTCCTC | Lager et al. (2010) |
| AT5G38710 | *PDH2* | Q-PCR | TCCACAGGTGCTTTAGACCG | TCCTGTGTCTTCACCAAGCC | The present study |
| AT5G06860 | *PGIP1* | Q-PCR | TAAACCAAGCTTATCTCTAGGATTA | CCATCAAATAAAACATTTTGAAAATGTGC | Sawaki et al. (2009) |
| AT5G06870 | *PGIP2* | Q-PCR | CCTAATCATACAAGACGGCGAGA | TTGGCGATAGTGGGTTGGA | Di et al. (2010) |
| AT1G34370 | *STOP1* | Q-PCR/RT-PCR | GAAGGTCATGTTGTGGAGGAAC | GAGTATGCGGTGCGAGAATC | Sawaki et al. (2009) |
| AT4G34270 | *TIP41* | Q-PCR | CATTTCAGTCTCTATCTGCGAAAGGGTATCC | CACCACAATAAGTCAGTGGAGTAACTCCTTAC | Hong et al. (2010) |

**Supplementary Table S4** Raw data of microarray analysis

| | Col Exp1 | ina1c1_Detection | ami2 Exp1 | ina3a1_Detection | Col Exp2 | ina2c2_Detection | ami2 Exp2 | ina4a2_Detection | Gene Symbol |
|---|---|---|---|---|---|---|---|---|---|
| 244901_at | 32.9 | P | 57.1 | P | 75 | P | 52.6 | P | ORF25 |
| 244902_at | 91.7 | P | 64.9 | P | 46.2 | A | 81.5 | P | NAD4L |
| 244903_at | 215.1 | P | 182.8 | P | 149.8 | P | 199.4 | P | ArthMp059 |
| 244904_at | 79.2 | P | 45.8 | P | 79.9 | P | 70.1 | P | ArthMp060 |
| 244905_at | 29.9 | A | 26.3 | A | 54.1 | A | 22.5 | A | ArthMp061 |
| 244906_at | 88.1 | P | 142.3 | P | 121.4 | P | 116.1 | P | ORF240A |
| 244907_at | 11.1 | A | 3.5 | A | 4.9 | A | 10 | A | ArthMp063 |
| 244908_at | 2.7 | A | 1.3 | A | 1.8 | A | 1.6 | A | ArthMp064 |
| 244909_at | 31.1 | A | 5.3 | A | 26.8 | A | 6.5 | A | ArthMp066 |
| 244910_s_at | 8.4 | A | 35.6 | P | 44.8 | P | 33.7 | P | ArthMp067 |
| 244911_at | 5.3 | A | 1.2 | A | 3.5 | A | 10.5 | A | ArthMp071 |
| 244912_at | 1705.6 | P | 778.1 | P | 1681.6 | P | 928.7 | P | --- |
| 244913_at | 3.6 | A | 3.3 | A | 10 | A | 2.7 | A | ArthMp073 |
| 244914_at | 1.8 | A | 2.2 | A | 1.1 | A | 10.4 | A | ArthMp074 |
| 244915_s_at | 16.7 | A | 46.2 | A | 3.5 | A | 15.7 | A | ArthMp075 |
| 244916_at | 23 | A | 15.1 | A | 23.5 | A | 3.2 | A | ORF187 |
| 244917_at | 3.5 | A | 2.3 | A | 5 | A | 3.1 | A | ArthMp076 |
| 244918_at | 4.5 | A | 3 | A | 31.6 | A | 9.2 | A | ArthMp078 |
| 244919_at | 1.6 | A | 10.5 | A | 28.5 | A | 8.7 | A | --- |
| 244920_s_at | 202.6 | P | 117.3 | P | 145.1 | P | 123.4 | P | --- |
| 244921_s_at | 167.9 | P | 124.2 | P | 125.5 | P | 104.7 | P | --- |
| 244922_s_at | 45.3 | P | 54.2 | P | 74.4 | P | 42.4 | P | --- |
| 244923_s_at | 41.4 | A | 44.1 | A | 56.9 | A | 43.7 | A | --- |
| 244924_at | 52.7 | A | 38.9 | A | 50.9 | A | 55.4 | A | ArthMp091 |
| 244925_at | 32.9 | A | 26.9 | P | 35.6 | A | 15.5 | A | nad7 |
| 244926_s_at | 7.8 | A | 3.7 | A | 4.7 | A | 4.3 | A | --- |
| 244927_at | 50.4 | A | 32.4 | A | 98.2 | A | 70.5 | A | --- |
| 244928_s_at | 183.9 | P | 143.3 | P | 185.3 | P | 151.4 | P | --- |
| 244929_at | 484.4 | P | 330.1 | P | 372.6 | P | 310.2 | P | nad4 |
| 244930_at | 29.1 | A | 12 | A | 22.8 | A | 1.8 | A | --- |
| 244931_at | 42.9 | M | 31.4 | P | 41.7 | A | 28.7 | P | ArthMp056 |
| 244932_at | 834.7 | P | 744.1 | P | 777.4 | P | 696.6 | P | psaC |
| 244933_at | 1051.7 | P | 521.3 | P | 666.2 | P | 559.4 | P | ndhE |
| 244934_at | 709.2 | P | 511.4 | P | 921.4 | P | 591.2 | P | ndhG |
| 244935_at | 16537.8 | P | 17068.3 | P | 21518.9 | P | 17883.3 | P | ndhI |
| 244936_at | 8469.3 | P | 8045.7 | P | 8560 | P | 8644.6 | P | ndhA |
| 244937_at | 721.7 | P | 830.5 | P | 818.1 | P | 779.1 | P | ndhH |
| 244938_at | 784.6 | P | 407.4 | P | 515.1 | P | 421.2 | P | rps15 |
| 244939_at | 1017 | P | 426.8 | P | 678.4 | P | 439.5 | P | rps12 |
| 244940_at | 6537.1 | P | 3224.5 | P | 5493.4 | P | 3549 | P | rps12 |
| 244941_at | 28.8 | A | 39.5 | A | 56.8 | P | 30.9 | A | ArthMp002 |
| 244942_at | 23.2 | A | 6.9 | A | 6.1 | A | 6.2 | A | ArthMp005 |
| 244943_at | 115.1 | P | 93.9 | P | 112.9 | P | 116 | P | NAD9 |
| 244944_s_at | 520.9 | P | 268.6 | P | 358.3 | P | 288 | P | rpl16 /// rps3 |
| 244945_at | 94.6 | P | 76.1 | P | 128.8 | P | 113.3 | P | ccb206 |
| 244946_at | 5.6 | A | 10.7 | A | 10.7 | A | 2.7 | A | ArthMp011 |
| 244947_at | 19.3 | A | 36.8 | A | 29 | A | 29 | P | ArthMp012 |
| 244948_at | 1.5 | A | 2.7 | A | 1.7 | A | 7.7 | A | ArthMp013 |
| 244949_at | 45.8 | A | 20.7 | A | 32.2 | A | 11.2 | A | ArthMp014 |
| 244950_at | 126.6 | P | 100.9 | P | 132.6 | P | 149.4 | P | cox2 |
| 244951_s_at | 186.3 | P | 161.6 | P | 224.1 | P | 197.1 | P | --- |
| 244952_at | 15.2 | A | 31.7 | P | 32.3 | P | 19.9 | A | ArthMp023 |
| 244953_s_at | 39.6 | P | 44.3 | P | 57.3 | P | 47.3 | P | --- |
| 244954_s_at | 10.8 | A | 6.1 | A | 10.8 | A | 2.5 | A | --- |
| 244955_at | 2.1 | A | 10.8 | A | 33.1 | A | 16 | A | ArthMp030 |
| 244956_s_at | 47.4 | M | 36.1 | P | 42.4 | P | 66.6 | A | ArthMp029 |
| 244957_at | 57.2 | P | 23.4 | A | 46.1 | P | 32 | A | ArthMp034 |
| 244958_at | 27.8 | A | 17.5 | A | 22.8 | A | 34.6 | A | ArthMp038 |
| 244959_s_at | 278.1 | P | 182.5 | P | 242.3 | P | 178.8 | P | --- |
| 244960_at | 4693.3 | P | 5200.2 | P | 5203.9 | P | 4850.8 | P | rpl32 |
| 244961_at | 2091 | P | 1855.1 | P | 1587.4 | P | 1464.5 | P | ccsA |
| 244962_at | 3712.2 | P | 3234.9 | P | 3672.2 | P | 2416.5 | P | ndhD |
| 244963_at | 9795.7 | P | 9411.9 | P | 11301.4 | P | 8941.5 | P | psbF |
| 244964_at | 5717.1 | P | 6291.8 | P | 6143.6 | P | 6267.3 | P | psbE |
| 244965_at | 870.3 | P | 512.3 | P | 633.8 | P | 451.7 | P | petL |
| 244966_at | 332.8 | P | 293.8 | P | 377.5 | P | 257.2 | P | petG |
| 244967_at | 3602.3 | P | 4374.8 | P | 3663.1 | P | 3771.8 | P | psaJ |
| 244968_at | 4220 | P | 4803.9 | P | 4426.5 | P | 4508.7 | P | rpl33 |
| 244969_at | 1242.2 | P | 1219 | P | 1237.7 | P | 856.7 | P | rps18 |
| 244970_at | 1105.4 | P | 417.2 | P | 787.1 | P | 471.3 | P | rpl20 |
| 244971_at | 6556.4 | P | 4510 | P | 5116.5 | P | 4201.1 | P | clpP |
| 244972_at | 729.2 | P | 618.9 | P | 544.2 | P | 596.9 | P | psbB |
| 244973_at | 448.7 | P | 388.2 | P | 321.7 | P | 283.3 | P | psbT |
| 244974_at | 419.5 | P | 296.7 | P | 241 | P | 355.9 | P | psbN |
| 244975_at | 2214.5 | P | 2089.9 | P | 1772.5 | P | 1856 | P | psbH |
| 244976_at | 7329.4 | P | 7984.3 | P | 8319.7 | P | 8335.2 | P | petB |
| 244977_at | 99.1 | P | 108.5 | P | 69.6 | P | 75.6 | P | petD |
| 244978_at | 7534.8 | P | 7370.9 | P | 9831.1 | P | 6837.2 | P | rpoA |
| 244979_at | 12587.7 | P | 11640.8 | P | 13160.3 | P | 13322 | P | rps11 |
| 244980_at | 9993.4 | P | 7639.9 | P | 7063.1 | P | 8582.5 | P | rpl36 |
| 244981_at | 3488.8 | P | 3276 | P | 4092.6 | P | 2963.5 | P | rps8 |
| 244982_at | 2740.8 | P | 1739.2 | P | 2727.3 | P | 1917.6 | P | rpl14 |
| 244983_at | 4042.9 | P | 2279.9 | P | 2420.6 | P | 2501.2 | P | rpl16 |
| 244984_at | 7399.3 | P | 5966.7 | P | 7037.3 | P | 5571.3 | P | rps3 |
| 244985_at | 9219.3 | P | 6986.1 | P | 7930.5 | P | 5997.9 | P | rpl22 |
| 244986_at | 4858.6 | P | 4719.5 | P | 5068.8 | P | 3444.1 | P | rps19 |
| 244987_s_at | 16646 | P | 13838.6 | P | 17092.3 | P | 14414.7 | P | rpl2 |
| 244988_s_at | 9378.5 | P | 8777.3 | P | 11204.9 | P | 6935.6 | P | rpl23 |
| 244989_s_at | 145.5 | P | 57.7 | P | 97.3 | P | 70.6 | P | ycf2 |
| 244990_s_at | 165.6 | P | 88.2 | P | 56.4 | P | 74.6 | P | --- |
| 244991_s_at | 97.9 | P | 44.5 | P | 100.5 | P | 68.3 | P | ndhB |
| 244992_s_at | 14890.1 | P | 8045.9 | P | 8965.3 | P | 8196.8 | P | rps7 |
| 244993_s_at | 4104.9 | P | 2390.9 | P | 2407.8 | P | 2334.5 | P | ycf1 |
| 244994_at | 22 | P | 19.1 | A | 35 | A | 20.4 | A | ndhF |
| 244995_at | 194.1 | P | 161 | P | 189.7 | P | 149.5 | P | atpI |
| 244996_at | 1147.5 | P | 738.8 | P | 978.5 | P | 583.3 | P | rps2 |
| 244997_at | 475.4 | P | 219.4 | P | 283.3 | P | 185.7 | P | rpoC2 |
| 244998_at | 1462.9 | P | 939 | P | 1382.9 | P | 937.5 | P | rpoC1 |
| 244999_at | 95.9 | P | 66.5 | P | 97.2 | P | 58 | P | rpoB |
| 245000_at | 121.6 | P | 132.5 | P | 182.1 | P | 141.8 | P | petN |
| 245001_at | 15826.4 | P | 20296.1 | P | 20701.7 | P | 13465.3 | P | psbM |
| 245002_at | 1064.4 | P | 783 | P | 1093.6 | P | 821.5 | P | psbD |
| 245003_at | 2748.4 | P | 1953.1 | P | 2140.5 | P | 1982.6 | P | psbC |
| 245004_at | 4307.1 | P | 2688.8 | P | 2922.7 | P | 2517.3 | P | psbZ |
| 245005_at | 50.5 | P | 42.4 | P | 78.8 | P | 58 | P | rps14 |

| | | | | | | | | | |
|---|---|---|---|---|---|---|---|---|---|
| 245006_at | 10822.3 | P | 7459.4 | P | 10462.7 | P | 7477.4 | P | psaB |
| 245007_at | 1403.6 | P | 1267.4 | P | 1406.1 | P | 1248.8 | P | psaA |
| 245008_at | 1367.1 | P | 611.7 | P | 755.9 | P | 578.2 | P | ycf3 |
| 245009_at | 4255.7 | P | 2339.2 | P | 3408.5 | P | 2247.4 | P | rps4 |
| 245010_at | 577.6 | P | 356.6 | P | 461.8 | P | 308 | P | ndhJ |
| 245011_at | 514.7 | P | 402 | P | 538.3 | P | 444.5 | P | ndhK |
| 245012_at | 4760.7 | P | 6080 | P | 5672.4 | P | 6099.5 | P | ndhC |
| 245013_at | 13348.2 | P | 11440.7 | P | 15171.5 | P | 11502.7 | P | atpE |
| 245014_at | 14228 | P | 11404.7 | P | 14569.2 | P | 11333.5 | P | atpB |
| 245015_at | 1250.9 | P | 872.8 | P | 1483.8 | P | 998 | P | rbcL |
| 245016_at | 1540.3 | P | 771.6 | P | 1149.8 | P | 789 | P | accD |
| 245017_at | 4023.2 | P | 2587 | P | 4290.8 | P | 2506.8 | P | psaI |
| 245018_at | 2347.6 | P | 1507.3 | P | 2334.5 | P | 1319.4 | P | ycf4 |
| 245019_at | 3344.4 | P | 1739.3 | P | 2701.3 | P | 1693.4 | P | cemA |
| 245020_at | 2932.2 | P | 1668.3 | P | 2333.2 | P | 1627.8 | P | petA |
| 245021_at | 4687.1 | P | 4482.2 | P | 4036.7 | P | 3902.3 | P | psbJ |
| 245022_at | 1833.5 | P | 1837.5 | P | 2290.4 | P | 1602 | P | psbL |
| 245023_at | 5401.1 | P | 6157 | P | 6554.9 | P | 5966.6 | P | psbI |
| 245024_at | 2474.5 | P | 1774.9 | P | 2242.5 | P | 1672.1 | P | atpA |
| 245025_at | 2530.8 | P | 1185.8 | P | 1743.9 | P | 1253.2 | P | atpF |
| 245026_at | 2048.7 | P | 1080.4 | P | 1407 | P | 1319.4 | P | atpH |
| 245027_at | 335.3 | P | 403.5 | P | 333.1 | P | 387.3 | P | HO2 |
| 245028_at | 253.1 | P | 363.8 | P | 290.5 | P | 299.2 | P | --- |
| 245029_at | 342.3 | P | 393.2 | P | 443.1 | P | 397.4 | P | YAB5 |
| 245030_at | 15.9 | A | 2.5 | A | 1.5 | A | 1.1 | A | --- |
| 245031_at | 31.5 | A | 37.6 | P | 31.2 | M | 30.2 | P | --- |
| 245032_at | 97.8 | P | 31.4 | P | 112.9 | P | 24.3 | P | --- |
| 245033_at | 26.5 | A | 3.3 | A | 19.3 | A | 8.8 | A | --- |
| 245034_at | 37.7 | A | 37.9 | A | 52.2 | A | 73.1 | A | --- |
| 245035_at | 96.9 | P | 139 | P | 151.8 | P | 88.3 | P | ATARD3 |
| 245036_at | 4.7 | A | 15.2 | A | 25.4 | A | 4.6 | A | Iqd4 |
| 245037_at | 7.6 | A | 46.6 | P | 38.2 | A | 25.2 | A | PIP5K3 |
| 245038_at | 978.1 | P | 619.2 | P | 620.6 | P | 740 | P | PLA2A |
| 245039_at | 387.2 | P | 498.8 | P | 402.4 | P | 467.3 | P | --- |
| 245040_at | 27.5 | A | 54.6 | A | 10.7 | A | 55.1 | M | --- |
| 245041_at | 514 | P | 208.9 | P | 418.2 | P | 248.3 | P | AR781 |
| 245042_at | 581.9 | P | 769.7 | P | 702.5 | P | 669.9 | P | HEMD |
| 245043_at | 569.9 | P | 494.1 | P | 608.3 | P | 521.7 | P | RCY1 |
| 245044_at | 16928.9 | P | 15355.8 | P | 17368.7 | P | 15969.5 | P | --- |
| 245045_at | 1127.3 | P | 991.8 | P | 903 | P | 799.5 | P | --- |
| 245046_at | 951.2 | P | 1141.2 | P | 1185.7 | P | 1106.7 | P | PDE135 |
| 245047_at | 2541 | P | 2174.7 | P | 2821.7 | P | 2361.7 | P | psbA |
| 245048_at | 5537.1 | P | 5888.7 | P | 5333.4 | P | 4424.2 | P | matK |
| 245049_at | 206.3 | P | 217.6 | P | 214 | P | 155.6 | P | rps16 |
| 245050_at | 6646.1 | P | 8171.2 | P | 8887.7 | P | 7589.3 | P | psbK |
| 245051_at | 523.2 | P | 437 | P | 478.6 | P | 315.3 | P | WRKY15 |
| 245052_at | 803.9 | P | 997.1 | P | 761.9 | P | 844.9 | P | --- |
| 245053_at | 20.2 | A | 16 | A | 8.1 | A | 1.2 | A | --- |
| 245054_at | 253.5 | P | 210.2 | P | 207.6 | P | 222.5 | P | --- |
| 245055_at | 89.5 | P | 90.9 | P | 77.2 | P | 86.7 | P | --- |
| 245056_at | 52.7 | P | 60.6 | A | 53.1 | P | 52.3 | P | UGT76D1 |
| 245057_at | 3.2 | A | 7 | A | 6.6 | A | 5.6 | A | --- |
| 245058_at | 18.3 | A | 3.4 | A | 4.7 | A | 6.6 | A | --- |
| 245059_at | 419.4 | P | 545.7 | P | 551.4 | P | 574.4 | P | RHC2A |
| 245060_at | 4124.2 | P | 3925.8 | P | 4572.1 | P | 4585.3 | P | CYT1 |
| 245061_at | 17865.8 | P | 19076.6 | P | 17305.6 | P | 19451.1 | P | RCA |
| 245062_at | 671.2 | P | 803.3 | P | 731.2 | P | 779.8 | P | BPM3 |
| 245063_at | 476.7 | P | 510.6 | P | 588.7 | P | 477.9 | P | --- |
| 245064_at | 393.7 | P | 267.6 | P | 265.6 | P | 357.5 | P | --- |
| 245065_at | 794.4 | P | 787.7 | P | 739 | P | 806.2 | P | --- |
| 245066_at | 412.3 | P | 447.1 | P | 422.9 | P | 426.7 | P | TOPP4 |
| 245067_at | 1871.6 | P | 2286.7 | P | 1909.4 | P | 2720.3 | P | RNS2 |
| 245068_at | 21.9 | A | 33.7 | A | 44.5 | A | 9.8 | A | UGT84B1 |
| 245069_at | 1.8 | A | 12.5 | A | 9.2 | A | 1.3 | A | UGT84B2 |
| 245070_at | 1.8 | A | 3.1 | A | 6.8 | A | 3.1 | A | --- |
| 245071_at | 6.6 | A | 7.9 | A | 2.8 | A | 31.8 | A | --- |
| 245072_s_at | 8 | A | 19.7 | A | 25.2 | P | 25 | A | CYP81D7 |
| 245073_at | 2.3 | A | 16.6 | A | 9.8 | A | 4.8 | A | --- |
| 245074_at | 174.2 | P | 220.8 | P | 137.2 | P | 202.7 | P | --- |
| 245075_at | 229.3 | P | 159.2 | P | 214.9 | P | 185 | P | CYP96A1 |
| 245076_at | 270.9 | P | 148.1 | P | 115 | A | 137.3 | P | GH3.3 |
| 245077_at | 17.8 | A | 21.2 | A | 11.1 | A | 18.2 | A | --- |
| 245078_at | 145.9 | P | 227.9 | P | 155.8 | P | 364.9 | P | --- |
| 245079_at | 232.3 | P | 186.3 | P | 226 | P | 156.4 | P | --- |
| 245080_at | 41.6 | A | 65.5 | P | 72.2 | A | 67.1 | P | --- |
| 245081_at | 42.7 | A | 51.2 | P | 44.3 | A | 42.3 | P | --- |
| 245082_at | 8.2 | A | 29 | A | 4.9 | A | 11.4 | A | --- |
| 245083_at | 662.2 | P | 762.9 | P | 674.2 | P | 795.5 | P | ATRER1C1 |
| 245084_at | 147 | P | 183.6 | P | 160.8 | P | 146.4 | P | AtMYB70 |
| 245085_at | 4629.2 | P | 4308 | P | 5819.2 | P | 4508.3 | P | PAB4 |
| 245086_at | 20.6 | A | 4.6 | A | 28.3 | A | 21.7 | A | --- |
| 245087_at | 98 | P | 147.5 | P | 144.1 | P | 169.7 | P | DAR2 |
| 245088_at | 258 | P | 244.6 | P | 343.3 | P | 271.2 | P | --- |
| 245089_at | 344.2 | P | 214.8 | P | 259.6 | P | 215.6 | P | --- |
| 245090_at | 109.4 | P | 168.9 | P | 190.8 | P | 171.2 | P | --- |
| 245091_at | 5.1 | A | 3.8 | A | 27.5 | A | 3.8 | A | --- |
| 245092_at | 291.3 | P | 421 | P | 342.3 | P | 420.2 | P | BZIP17 |
| 245093_at | 203 | P | 298.3 | P | 248.1 | P | 235.8 | P | --- |
| 245094_at | 4801.7 | P | 5316 | P | 4964.7 | P | 5143.1 | P | DPE2 |
| 245095_at | 107 | P | 116.1 | P | 99.4 | P | 97.2 | P | --- |
| 245096_at | 2394.8 | P | 3481.4 | P | 2413.3 | P | 2960.8 | P | ATCYSA |
| 245097_at | 194.3 | P | 257.7 | P | 253.1 | P | 287.9 | P | --- |
| 245098_at | 910.6 | P | 1331.9 | P | 1037.1 | P | 1277.9 | P | ERS1 |
| 245099_at | 550.2 | P | 733.5 | P | 498.7 | P | 688.8 | P | RHC1A |
| 245100_at | 254.2 | P | 328.6 | P | 272.1 | P | 249 | P | UBP5 |
| 245101_at | 1273.7 | P | 1410.2 | P | 1519.5 | P | 1371.9 | P | CYP98A3 |
| 245102_at | 28.9 | A | 5 | A | 34.2 | A | 14.1 | A | --- |
| 245103_at | 18.5 | A | 18.5 | A | 36.3 | A | 10.8 | A | --- |
| 245104_at | 495.8 | P | 562.4 | P | 569.3 | P | 572.5 | P | --- |
| 245105_at | 59 | A | 43.7 | A | 33.7 | A | 36.1 | P | --- |
| 245106_at | 572.2 | P | 442.1 | P | 521.2 | P | 529 | P | --- |
| 245107_at | 2.8 | A | 8 | A | 5.5 | A | 1.9 | A | AT-HSFB3 |
| 245108_at | 17.6 | A | 8 | A | 17.3 | A | 3.4 | A | CKX1 |
| 245109_at | 187.9 | P | 225.3 | P | 207.6 | P | 173.1 | P | --- |
| 245110_at | 60.6 | P | 72.2 | P | 111.4 | M | 60.1 | A | --- |
| 245111_at | 1.4 | A | 4.9 | A | 8.2 | A | 2.3 | A | --- |
| 245112_at | 393.6 | P | 450.9 | P | 431.3 | P | 452.2 | P | GPDHC1 |
| 245113_at | 63.5 | A | 60.7 | A | 40.7 | A | 103 | A | MIZ1 |
| 245114_at | 1052 | P | 877.5 | P | 974.9 | P | 925.1 | P | TFIIB |

| | | | | | | | | | |
|---|---|---|---|---|---|---|---|---|---|
| 245115_at | 3120.7 | P | 3290.3 | P | 3562.5 | P | 2964.5 | P | SFGH |
| 245116_at | 663.4 | P | 883.1 | P | 813.3 | P | 670.2 | P | --- |
| 245117_at | 238.5 | P | 554.2 | P | 230.8 | P | 430.3 | P | ACA4 |
| 245118_at | 1800.2 | P | 1780.9 | P | 1699.8 | P | 1918.9 | P | NTRC |
| 245119_at | 158 | P | 122.2 | P | 133.4 | P | 95.3 | P | --- |
| 245120_at | 315.9 | P | 388.2 | P | 525.3 | P | 380.6 | P | HOS1 |
| 245121_at | 6386.5 | P | 5609.5 | P | 6399.1 | P | 5818.8 | P | --- |
| 245122_at | 848.7 | P | 514 | P | 798.9 | P | 584.6 | P | --- |
| 245123_at | 2969.5 | P | 3138.2 | P | 2428.4 | P | 2321.7 | P | CAO |
| 245124_at | 1091.6 | P | 1020 | P | 1129.3 | P | 1154.2 | P | U1A |
| 245125_at | 3.9 | A | 21.7 | A | 21.8 | A | 7.7 | A | CKI1 |
| 245126_at | 40.6 | A | 28.5 | A | 47.2 | A | 44.3 | A | MYB12 |
| 245127_at | 1402.2 | P | 878.5 | P | 891 | P | 868.7 | P | ATMHX |
| 245128_at | 25.8 | A | 17 | A | 8.5 | A | 1.9 | A | --- |
| 245129_at | 30.2 | A | 32.6 | A | 49.9 | A | 52 | M | CRR4 |
| 245130_at | 330 | P | 440 | P | 304.7 | P | 448.1 | P | --- |
| 245131_s_at | 579.3 | P | 561.3 | P | 483.3 | P | 510.5 | P | emb1067 |
| 245132_at | 136.9 | P | 166.1 | P | 263.2 | P | 210.7 | P | --- |
| 245133_at | 89.6 | P | 107.2 | P | 149.5 | P | 108.7 | P | GAE4 |
| 245134_s_at | 575.2 | P | 606.2 | P | 434.9 | P | 694 | P | --- |
| 245135_at | 7.1 | A | 31.9 | A | 4.6 | A | 30.7 | A | --- |
| 245136_at | 624.6 | P | 523 | P | 387.1 | P | 388.6 | P | --- |
| 245137_at | 132.8 | P | 134.5 | P | 119.5 | P | 105.9 | P | --- |
| 245138_at | 314.1 | P | 342.1 | P | 350.5 | P | 404.8 | P | AFO |
| 245139_at | 56.4 | P | 53.9 | A | 66.3 | P | 61.7 | P | --- |
| 245140_at | 14 | A | 28.8 | A | 9.2 | A | 1.9 | A | LBD18 |
| 245141_at | 267.2 | P | 174.4 | P | 282.4 | P | 211.9 | P | BEN1 |
| 245142_at | 299.7 | P | 286.4 | P | 301.6 | P | 329.1 | P | --- |
| 245143_at | 85.5 | P | 173.7 | P | 132.9 | P | 134 | P | ZPR1 |
| 245144_at | 1124 | P | 1144.5 | P | 1529.1 | P | 1179.3 | P | MAP1A |
| 245145_at | 1614.1 | P | 1204 | P | 1426.6 | P | 1335.8 | P | DHDPS2 |
| 245146_at | 648.3 | P | 637.8 | P | 439 | P | 514.2 | P | GOS12 |
| 245147_at | 191.3 | P | 88.9 | P | 178.4 | P | 124.7 | P | ATRAD51C |
| 245148_at | 8.1 | A | 8.8 | A | 15 | A | 5.3 | A | --- |
| 245149_at | 123.8 | P | 69.9 | P | 132.9 | P | 106.6 | P | ATSWI3A |
| 245150_at | 1930.5 | P | 2855 | P | 2065.9 | P | 3009.3 | P | PHR2 |
| 245151_at | 47.8 | P | 21.3 | P | 52.2 | P | 39.7 | P | --- |
| 245152_at | 301.4 | P | 627.3 | P | 579.7 | P | 558.6 | P | --- |
| 245153_at | 9.8 | A | 11.5 | P | 17.4 | A | 21.7 | A | --- |
| 245154_at | 4.1 | A | 7.7 | A | 30.2 | A | 6.6 | A | --- |
| 245155_at | 1640.7 | P | 2080 | P | 1763.2 | P | 2017.3 | P | --- |
| 245156_at | 227 | P | 231.4 | P | 300.9 | P | 273.9 | P | CPK7 |
| 245157_at | 9 | A | 18.2 | A | 20.4 | A | 15.5 | A | --- |
| 245158_at | 3.7 | A | 18.1 | A | 5.4 | A | 11.8 | A | RALFL18 |
| 245159_at | 22.1 | A | 27.7 | A | 3.9 | A | 29.1 | A | ATCSLD1 |
| 245160_at | 14.9 | P | 34.9 | A | 13.2 | A | 14.7 | A | AtRLP28 |
| 245161_at | 97.8 | A | 86.4 | P | 131.9 | A | 79.8 | A | NSP2 |
| 245162_at | 39.4 | A | 47.3 | A | 47.3 | P | 31.3 | A | XID |
| 245163_at | 21.3 | A | 2.6 | A | 3.1 | A | 8.1 | A | --- |
| 245164_at | 2475.5 | P | 2179.8 | P | 2918.3 | P | 2193 | P | HSP60-2 |
| 245165_at | 1581.7 | P | 1566.1 | P | 1833.3 | P | 1467.5 | P | --- |
| 245166_at | 190.5 | P | 244.8 | P | 268.1 | P | 210.6 | P | --- |
| 245167_s_at | 1014.1 | P | 1362.6 | P | 1066.4 | P | 1153.7 | P | ATVAMP723 |
| 245168_at | 9925.4 | P | 7671 | P | 6349.3 | P | 7538.4 | P | PKT3 |
| 245169_at | 2151.3 | P | 1946.3 | P | 2195.1 | P | 1929.9 | P | --- |
| 245170_at | 4.3 | A | 23.4 | A | 61.4 | A | 17.3 | A | --- |
| 245171_at | 2.4 | A | 16 | A | 6.8 | A | 5.2 | A | --- |
| 245172_at | 44.3 | P | 29.2 | A | 58.7 | A | 41 | P | --- |
| 245173_at | 55 | P | 7.9 | A | 85.6 | P | 25.1 | P | --- |
| 245174_at | 70.6 | P | 89.3 | P | 86.6 | P | 101.8 | P | --- |
| 245175_at | 2495.3 | P | 3591.7 | P | 2646.2 | P | 3854.1 | P | UNE5 |
| 245176_at | 579.6 | P | 603.8 | P | 688.9 | P | 620.7 | P | --- |
| 245177_at | 22.2 | A | 24.9 | A | 26.8 | A | 19.8 | A | --- |
| 245178_at | 671.9 | P | 484.9 | P | 588.7 | P | 493.5 | P | FIS1B |
| 245179_at | 183.2 | P | 180.3 | P | 132.8 | P | 129.6 | P | --- |
| 245180_at | 622.6 | P | 628.5 | P | 751.5 | P | 713.7 | P | --- |
| 245181_at | 162.7 | P | 154.5 | P | 128.3 | P | 155 | P | --- |
| 245182_at | 260.5 | P | 235.5 | P | 198.6 | P | 248.7 | P | --- |
| 245183_at | 124.3 | P | 182.7 | P | 198.5 | P | 183 | P | --- |
| 245184_at | 1.8 | A | 12.4 | A | 31.7 | A | 23.8 | A | --- |
| 245185_at | 85.9 | P | 80.2 | P | 92.5 | M | 102.9 | P | --- |
| 245186_at | 8.7 | A | 3.3 | A | 7.4 | A | 23.8 | A | ARR11 |
| 245187_s_at | 1188.4 | P | 841.6 | P | 1071.9 | P | 913 | P | --- |
| 245188_at | 645.8 | P | 971.7 | P | 719.1 | P | 910.6 | P | --- |
| 245189_at | 0.9 | A | 1 | A | 1.4 | A | 1.4 | A | --- |
| 245190_at | 102.4 | P | 102.7 | P | 155.1 | P | 105.6 | P | --- |
| 245191_at | 21.7 | A | 6.3 | A | 14.6 | A | 22.2 | A | TEL2 |
| 245192_at | 0.9 | A | 1.9 | A | 6.6 | A | 3.9 | A | --- |
| 245193_at | 2963.2 | P | 1880.3 | P | 2779.6 | P | 1853.5 | P | SUFE2 |
| 245194_at | 138.4 | P | 130.1 | P | 229.6 | P | 142.8 | P | --- |
| 245195_at | 9860.5 | P | 12248.7 | P | 11237.2 | P | 11614.6 | P | PSBY |
| 245196_at | 275.2 | P | 911.2 | P | 434.8 | P | 875.9 | P | --- |
| 245197_at | 286.9 | P | 311.5 | P | 283.4 | P | 294.9 | P | --- |
| 245198_at | 2851.7 | P | 2948.5 | P | 2743.3 | P | 2836 | P | --- |
| 245199_at | 2097 | P | 1966.9 | P | 2027.8 | P | 2145.3 | P | KCR1 |
| 245200_at | 290.7 | P | 379.5 | P | 358.1 | P | 337.9 | P | --- |
| 245201_at | 949.9 | P | 1024.1 | P | 1015.5 | P | 1092.7 | P | CSK |
| 245202_at | 283.8 | P | 322.8 | P | 350.8 | P | 311.7 | P | --- |
| 245203_at | 0.7 | A | 1.7 | A | 0.9 | A | 0.6 | A | --- |
| 245204_at | 5.8 | A | 7.2 | A | 3.9 | A | 2.8 | A | --- |
| 245205_at | 42.4 | A | 40.1 | A | 45.4 | A | 45.2 | A | --- |
| 245206_at | 286 | P | 360.6 | P | 356 | P | 358.2 | P | DGS1 |
| 245207_at | 665.2 | P | 693.5 | P | 776.1 | P | 864.1 | P | --- |
| 245208_at | 123 | P | 42.9 | A | 100.7 | P | 53.9 | P | LRP1 |
| 245209_at | 27.4 | A | 33 | A | 25.4 | A | 4.9 | A | --- |
| 245210_at | 177.1 | P | 214.8 | P | 229.5 | P | 207.5 | P | --- |
| 245211_at | 666.2 | P | 822.2 | P | 790.1 | P | 818.3 | P | SEC10 |
| 245212_at | 3.4 | A | 15.3 | A | 5.5 | A | 3.4 | A | AT1G44224 |
| 245213_at | 8871.7 | P | 11549.4 | P | 10015.3 | P | 12585 | P | NPQ4 |
| 245214_at | 2.8 | A | 19.5 | A | 36.3 | A | 17.9 | A | --- |
| 245215_at | 581.7 | P | 686.1 | P | 696.5 | P | 786.1 | P | ATFXG1 |
| 245216_at | 1625.3 | P | 1633.2 | P | 1477.2 | P | 1658.5 | P | IMPA-2 |
| 245217_at | 1.9 | A | 3 | A | 4.8 | A | 1.5 | A | --- |
| 245218_s_at | 839.2 | P | 1016.4 | P | 1084.6 | P | 1036.5 | P | --- |
| 245219_at | 2072.6 | P | 2118.8 | P | 2138 | P | 2089.6 | P | --- |
| 245220_at | 0.3 | A | 1 | A | 1.4 | A | 0.8 | A | --- |
| 245221_s_at | 17.8 | A | 3.2 | A | 12.3 | A | 16.1 | A | --- |
| 245222_at | 3.6 | A | 0.8 | A | 2.6 | A | 1.8 | A | --- |
| 245223_at | 1.1 | A | 2.3 | A | 7.6 | A | 1.1 | A | --- |

| | | | | | | | | | |
|---|---|---|---|---|---|---|---|---|---|
| 245224_at | 16.6 | A | 26.1 | A | 22.5 | A | 12.3 | A | --- |
| 245225_at | 15.1 | A | 6.6 | A | 5.1 | A | 19.1 | A | --- |
| 245226_at | 41.7 | A | 40.6 | A | 23.6 | A | 32.4 | A | --- |
| 245227_s_at | 229.3 | P | 203.2 | P | 240.5 | P | 235.1 | P | --- |
| 245228_at | 319.4 | P | 257 | P | 370.4 | P | 277 | P | COBL2 |
| 245229_at | 403.6 | P | 396.5 | P | 330.8 | P | 418.9 | P | --- |
| 245230_at | 82.7 | P | 83.6 | P | 85.9 | P | 81.9 | P | --- |
| 245231_at | 119.4 | P | 83.6 | P | 104.8 | P | 86.1 | P | --- |
| 245232_at | 2.5 | A | 10.1 | A | 7.5 | A | 11.7 | A | ADF7 |
| 245233_at | 5.8 | A | 5.8 | A | 16 | A | 2 | A | --- |
| 245234_at | 34.4 | A | 3.6 | A | 21.1 | A | 7 | A | AtMYB18 |
| 245235_at | 1660.9 | P | 1778 | P | 1574.1 | P | 1649.3 | P | --- |
| 245236_at | 73 | P | 81.6 | P | 101.1 | P | 76.9 | P | MSH3 |
| 245237_at | 117.6 | P | 125.3 | P | 126 | P | 104.8 | P | SLK1 |
| 245238_at | 1615.4 | P | 2186.9 | P | 1576.3 | P | 1918.2 | P | ACYB-2 |
| 245239_at | 50.4 | P | 29.7 | A | 46.3 | A | 24.5 | A | FWA |
| 245240_at | 3.7 | A | 7.2 | A | 9 | A | 4.1 | A | --- |
| 245241_at | 2.9 | A | 4.1 | A | 2.2 | A | 3.9 | A | --- |
| 245242_at | 547.1 | P | 844.5 | P | 738.6 | P | 742.9 | P | CH1 |
| 245243_at | 15.1 | A | 46.2 | P | 45.7 | A | 48.2 | A | --- |
| 245244_at | 1383.7 | P | 394.8 | P | 1337 | P | 542.4 | P | ILL6 |
| 245245_at | 22 | A | 14.6 | A | 52.2 | A | 20.6 | A | hemb2 |
| 245246_at | 18.5 | A | 7.3 | A | 4.1 | A | 7 | A | --- |
| 245247_at | 728.4 | P | 529.5 | P | 656.8 | P | 495.5 | P | SCL13 |
| 245248_at | 257.1 | P | 325.7 | P | 290.2 | P | 342.6 | P | ATB' GAMMA |
| 245249_at | 4504.4 | P | 2353 | P | 3627.1 | P | 2510.7 | P | ACX1 |
| 245250_at | 331.5 | P | 240.6 | P | 305.5 | P | 282.6 | P | ATERF6 |
| 245251_at | 410 | P | 323 | P | 364.5 | P | 336.5 | P | CBL1 |
| 245252_at | 444 | P | 243.1 | P | 445.6 | P | 216.9 | P | ATERF-1 |
| 245253_at | 802.5 | P | 280.9 | P | 550.6 | P | 543.5 | P | HPL1 |
| 245254_at | 1518.7 | P | 816.4 | P | 1606.6 | P | 959.7 | P | APS3 |
| 245255_at | 832.3 | P | 873.3 | P | 821.5 | P | 828.2 | P | COP9 |
| 245256_at | 201.8 | P | 242.2 | P | 228.2 | P | 167 | P | FAR1 |
| 245257_at | 8.9 | A | 6.6 | A | 0.8 | A | 2.9 | A | CAM8 |
| 245258_at | 0.9 | A | 11.6 | A | 10.1 | A | 10.2 | A | ATPEN1 |
| 245259_at | 49.1 | P | 77.3 | P | 50.1 | M | 69.9 | P | PAKRP1 |
| 245260_at | 540.8 | P | 671.2 | P | 646.7 | P | 769.8 | P | --- |
| 245261_at | 363.3 | P | 344.7 | P | 429.7 | P | 303.2 | P | --- |
| 245262_at | 506.9 | P | 250.2 | P | 316.6 | P | 428.2 | P | --- |
| 245263_at | 1065.6 | P | 733.2 | P | 1199.6 | P | 803.9 | P | --- |
| 245264_at | 705.8 | P | 460.2 | P | 639.7 | P | 392.3 | P | --- |
| 245265_at | 180.1 | P | 636.8 | P | 256.1 | P | 609.6 | P | ACD6 |
| 245266_at | 341 | P | 475.3 | P | 366.8 | P | 367.1 | P | --- |
| 245267_at | 2.3 | A | 56.2 | P | 32.3 | A | 21.4 | A | --- |
| 245268_at | 132.8 | P | 170.4 | P | 238.6 | P | 142.6 | P | PRMT7 |
| 245269_at | 896.4 | P | 973.8 | P | 832.8 | P | 941.4 | P | --- |
| 245270_at | 3972.9 | P | 6392.7 | P | 5829 | P | 6773.7 | P | TUA6 |
| 245271_at | 298.4 | P | 358.5 | P | 251.4 | P | 298.1 | P | --- |
| 245272_at | 138.5 | P | 132.6 | P | 141.7 | P | 170.4 | P | --- |
| 245273_at | 325.2 | P | 369.6 | P | 344.6 | P | 343.7 | P | --- |
| 245274_at | 936.9 | P | 1125.2 | P | 1266.6 | P | 1164.4 | P | --- |
| 245275_at | 1306.8 | P | 844.2 | P | 795.4 | P | 1163.7 | P | BAM5 |
| 245276_at | 109.7 | A | 172.8 | P | 139.7 | A | 108.2 | A | ATHB-2 |
| 245277_at | 774.7 | P | 826.2 | P | 734.4 | P | 1010.3 | P | IAGLU |
| 245278_at | 626 | P | 561.3 | P | 715.3 | P | 600.9 | P | SYP23 |
| 245279_at | 705.1 | P | 744.6 | P | 566.2 | P | 706.7 | P | --- |
| 245280_at | 524.1 | P | 508.2 | P | 476.2 | P | 526.1 | P | VRN2 |
| 245281_at | 2530.3 | P | 3503.1 | P | 3220.6 | P | 3265.6 | P | CLA1 |
| 245282_at | 449.8 | P | 581.7 | P | 413.2 | P | 563.5 | P | --- |
| 245283_at | 59.1 | P | 157.5 | P | 87.3 | P | 110.5 | P | AT-HSFA5 |
| 245284_at | 1557.5 | P | 1857.6 | P | 1578.5 | P | 1827.9 | P | PDS3 |
| 245285_s_at | 3493.9 | P | 2055.7 | P | 2984.1 | P | 2226.8 | P | SBP1 |
| 245286_at | 10342.6 | P | 7704.4 | P | 7592.2 | P | 7279.4 | P | OASA1 |
| 245287_at | 931.4 | P | 821 | P | 779.2 | P | 838.4 | P | --- |
| 245288_at | 451.4 | P | 477.1 | P | 317.9 | P | 519.1 | P | ADA2B |
| 245289_at | 2347.3 | P | 2810.5 | P | 2143.1 | P | 2671 | P | --- |
| 245290_at | 634.6 | P | 872.5 | P | 641.5 | P | 774.6 | P | --- |
| 245291_at | 1355.8 | P | 1126.8 | P | 1720.9 | P | 1192.6 | P | --- |
| 245292_at | 317.4 | P | 324.4 | P | 387.5 | P | 308.2 | P | --- |
| 245293_at | 1158.7 | P | 1079 | P | 1008 | P | 1158.9 | P | --- |
| 245294_at | 324.2 | P | 393 | P | 385 | P | 375.7 | P | CKI1 |
| 245295_at | 279.4 | P | 277.1 | P | 249.9 | P | 325.4 | P | --- |
| 245296_at | 1099.7 | P | 3187.1 | P | 2169.4 | P | 2704.3 | P | ATOPT3 |
| 245297_at | 211.4 | P | 168.8 | P | 161.6 | P | 168.4 | P | --- |
| 245298_at | 408.6 | P | 339.1 | P | 425.3 | P | 423.1 | P | --- |
| 245299_at | 2225.1 | P | 1819.7 | P | 1901.6 | P | 2072.3 | P | ATRAB1C |
| 245300_at | 28.5 | A | 28.7 | A | 37.7 | A | 21.7 | A | CBL6 |
| 245301_at | 435.7 | P | 483.8 | P | 341.8 | P | 459.2 | P | FPS2 |
| 245302_at | 162.2 | P | 205.9 | P | 121.1 | P | 223.5 | P | KAN3 |
| 245303_at | 239.2 | P | 246.8 | P | 259.3 | P | 258.1 | P | --- |
| 245304_at | 1424.5 | P | 1085.4 | P | 1041.9 | P | 1082.6 | P | --- |
| 245305_at | 31.6 | P | 45.3 | A | 58.1 | A | 44.1 | A | --- |
| 245306_at | 1500.3 | P | 630.4 | P | 3907.6 | P | 1108.2 | P | ELIP2 |
| 245307_at | 420.7 | P | 459.7 | P | 530.3 | P | 463.7 | P | --- |
| 245308_at | 468.8 | P | 535.3 | P | 546.6 | P | 562.3 | P | --- |
| 245309_at | 137.8 | P | 203.7 | P | 136.5 | P | 200 | P | --- |
| 245310_at | 298.8 | P | 351.7 | P | 314.7 | P | 360.7 | P | RALFL31 |
| 245311_at | 6461.2 | P | 5913.8 | P | 7614 | P | 6671.8 | P | --- |
| 245312_at | 77.6 | P | 122.7 | P | 114.8 | P | 123.3 | P | --- |
| 245313_at | 554.6 | P | 432.6 | P | 365.6 | P | 377.2 | P | --- |
| 245314_at | 121.9 | P | 83.9 | P | 121 | P | 64.5 | A | --- |
| 245315_at | 1684.9 | P | 1886.6 | P | 1985 | P | 1932.5 | P | PBD2 |
| 245316_at | 616 | P | 588.5 | P | 628 | P | 581 | P | --- |
| 245317_at | 1118.7 | P | 638.5 | P | 520.2 | P | 702.6 | P | --- |
| 245318_at | 2403.1 | P | 5456 | P | 3933.3 | P | 5424.4 | P | --- |
| 245319_at | 1971.3 | P | 1633.5 | P | 1734.7 | P | 1876.5 | P | --- |
| 245320_at | 554.1 | P | 517.5 | P | 458 | P | 512.2 | P | --- |
| 245321_at | 4967.8 | P | 6346.4 | P | 4592.2 | P | 5547.2 | P | --- |
| 245322_at | 15.5 | A | 41.3 | A | 30.2 | A | 19.4 | A | --- |
| 245323_at | 1132.7 | P | 1473.5 | P | 985.1 | P | 1445.1 | P | --- |
| 245324_at | 518.2 | P | 616.3 | P | 518.5 | P | 616.4 | P | --- |
| 245325_at | 1.9 | A | 54.9 | A | 33.2 | A | 33.8 | A | XTR7 |
| 245326_at | 559.9 | P | 762.1 | P | 768.1 | P | 732.5 | P | --- |
| 245327_at | 176.9 | P | 253.5 | P | 182.9 | P | 171 | P | --- |
| 245328_at | 34.7 | A | 97 | P | 76 | P | 111.7 | P | --- |
| 245329_at | 660.4 | P | 1501.4 | P | 544.5 | P | 903.9 | P | --- |
| 245330_at | 444.4 | P | 745.7 | P | 476.2 | P | 645.2 | P | --- |
| 245331_at | 982.9 | P | 1068 | P | 1327 | P | 1016.1 | P | --- |
| 245332_at | 1650.3 | P | 1290.9 | P | 1424.1 | P | 1297.7 | P | --- |

| | | | | | | | | | |
|---|---|---|---|---|---|---|---|---|---|
| 245333_at | 1719.7 | P | 1511.1 | P | 1587.1 | P | 1550.6 | P | --- |
| 245334_at | 1657.8 | P | 1313 | P | 1169.6 | P | 1553.2 | P | RALFL33 |
| 245335_at | 5.8 | A | 4.4 | A | 3.9 | A | 4.7 | A | ATOEP16-2 |
| 245336_at | 100 | P | 231.2 | P | 126.1 | A | 184.8 | P | --- |
| 245337_at | 384.6 | P | 331.4 | P | 381.4 | P | 322.6 | P | --- |
| 245338_at | 392.1 | P | 460.1 | P | 379.9 | P | 417.7 | P | --- |
| 245339_at | 171.3 | P | 190 | P | 249.4 | P | 192.1 | P | COBL7 |
| 245340_at | 1479.2 | P | 2021 | P | 1691.4 | P | 1993.1 | P | --- |
| 245341_at | 265.7 | P | 214.8 | P | 334.1 | P | 199.6 | P | --- |
| 245342_at | 8445.4 | P | 7969.6 | P | 9088.2 | P | 8102.8 | P | --- |
| 245343_at | 248.3 | P | 497.6 | P | 316.3 | P | 498.7 | P | --- |
| 245344_at | 529.8 | P | 538 | P | 536.6 | P | 531.9 | P | --- |
| 245345_at | 6.2 | A | 27.6 | A | 24.2 | A | 16.8 | A | --- |
| 245346_at | 2168.7 | P | 4514.1 | P | 2705.1 | P | 5949.1 | P | CT-BMY |
| 245347_at | 2906.8 | P | 3248.3 | P | 2391.5 | P | 3325.5 | P | --- |
| 245348_at | 861.3 | P | 1210.6 | P | 899.2 | P | 1306.5 | P | ATTPS5 |
| 245349_at | 642.6 | P | 327.3 | P | 443.8 | P | 276.3 | P | MES16 |
| 245350_at | 1626.9 | P | 1922.9 | P | 1574.2 | P | 1556 | P | --- |
| 245351_at | 1248.7 | P | 1379 | P | 1189.5 | P | 1315.6 | P | CKB2 |
| 245352_at | 3472.5 | P | 1849.1 | P | 3402.4 | P | 2456.8 | P | UGT84A3 |
| 245353_at | 12.5 | A | 33.5 | A | 19.4 | A | 36.8 | A | --- |
| 245354_at | 3564.7 | P | 4370.7 | P | 3821.3 | P | 4213.8 | P | LIL3:1 |
| 245355_at | 7033.5 | P | 6743.7 | P | 8729.8 | P | 7534.7 | P | --- |
| 245356_at | 13553.8 | P | 11836 | P | 14480.3 | P | 12097.6 | P | MEE58 |
| 245357_at | 2192.5 | P | 3033 | P | 2666.2 | P | 3085.1 | P | --- |
| 245358_at | 756.9 | P | 731.4 | P | 908.7 | P | 758 | P | PRL1 |
| 245359_at | 1204.8 | P | 915.2 | P | 1029.3 | P | 971.6 | P | IBR10 |
| 245360_at | 1752.8 | P | 1445.9 | P | 1585.4 | P | 1403.4 | P | --- |
| 245361_at | 397.5 | P | 455.1 | P | 363.4 | P | 398.7 | P | --- |
| 245362_at | 61 | P | 124.8 | P | 87.4 | P | 134.6 | P | HAT1 |
| 245363_at | 42.2 | A | 54.6 | A | 54.8 | A | 59.8 | A | --- |
| 245364_at | 270.2 | P | 396.6 | P | 255.1 | P | 486.7 | P | --- |
| 245365_at | 563.6 | P | 729.5 | P | 500.5 | P | 623.3 | P | --- |
| 245366_at | 413.8 | P | 413.9 | P | 434.7 | P | 477.8 | P | AtMAPR4 |
| 245367_at | 124.5 | P | 161.9 | P | 138.2 | P | 186.3 | P | NRPB9B |
| 245368_at | 340 | P | 520.4 | P | 403.3 | P | 453.3 | P | --- |
| 245369_at | 41.8 | A | 42.6 | P | 78.8 | A | 47 | M | --- |
| 245370_at | 187.1 | P | 255.6 | P | 228.4 | P | 263 | P | --- |
| 245371_at | 24.8 | P | 52.9 | A | 54.5 | A | 34.7 | A | --- |
| 245372_at | 5776.8 | P | 5219.1 | P | 5296 | P | 5436.4 | P | --- |
| 245373_s_at | 5971.5 | P | 6146.9 | P | 4564.5 | P | 5991.4 | P | ATARD2 |
| 245374_at | 669.6 | P | 599.1 | P | 681.1 | P | 563.5 | P | --- |
| 245375_at | 178.9 | P | 144.2 | P | 72.2 | P | 108.3 | P | --- |
| 245376_at | 10.7 | A | 37.9 | P | 38.8 | A | 28.7 | A | --- |
| 245377_at | 14.3 | A | 13.3 | A | 24 | A | 18.3 | A | --- |
| 245378_at | 46.8 | A | 59.7 | A | 123 | A | 56.4 | A | HDG4 |
| 245379_at | 96 | P | 104.7 | P | 88.2 | P | 86.9 | P | HSF1 |
| 245380_at | 40.4 | A | 73.1 | P | 69.8 | P | 79.4 | P | --- |
| 245381_at | 2.7 | A | 21.8 | A | 55.6 | A | 10.2 | A | MYB39 |
| 245382_at | 11.9 | A | 23 | A | 40.6 | A | 19.2 | A | --- |
| 245383_at | 224 | P | 268.2 | P | 377.9 | P | 268.4 | P | --- |
| 245384_at | 125.8 | P | 90.1 | P | 99.7 | P | 87.7 | P | --- |
| 245385_at | 1101.4 | P | 821 | P | 1048.2 | P | 785.9 | P | --- |
| 245386_at | 936 | P | 610 | P | 985.1 | P | 806.8 | P | RALFL32 |
| 245387_at | 1520.3 | P | 1618.3 | P | 1821.4 | P | 1701.5 | P | --- |
| 245388_at | 3834.6 | P | 3650.9 | P | 3977.9 | P | 3929.1 | P | --- |
| 245389_at | 83.9 | A | 46 | A | 67.1 | A | 94.8 | A | --- |
| 245390_at | 794.1 | P | 628.4 | P | 676.9 | P | 616.7 | P | --- |
| 245391_at | 3234 | P | 3182.4 | P | 2500.5 | P | 2880.6 | P | ATG8F |
| 245392_at | 38.4 | P | 123.4 | P | 66.3 | A | 89.9 | P | --- |
| 245393_at | 175.2 | P | 107.1 | P | 169.6 | P | 95.3 | P | --- |
| 245394_at | 4910.2 | P | 5612.7 | P | 4577.7 | P | 5765.2 | P | --- |
| 245395_at | 346.9 | P | 413.3 | P | 363.9 | P | 364.9 | P | --- |
| 245396_at | 623.9 | P | 963.4 | P | 675.4 | P | 991.4 | P | --- |
| 245397_at | 104.8 | A | 155.8 | P | 110.1 | A | 154.1 | A | IAA1 |
| 245398_at | 374.1 | P | 354.2 | P | 411.3 | P | 295.3 | P | --- |
| 245399_at | 526.9 | P | 1061.3 | P | 547 | P | 898.9 | P | TIP2;2 |
| 245400_at | 6360.4 | P | 6803 | P | 6889.3 | P | 6692.9 | P | --- |
| 245401_at | 403.5 | P | 759.9 | P | 332.5 | P | 571.5 | P | --- |
| 245402_at | 9.5 | A | 3 | A | 3.6 | A | 3.2 | A | --- |
| 245403_at | 6.2 | A | 9.8 | A | 2.3 | A | 18.2 | A | --- |
| 245404_at | 263.4 | P | 200.2 | P | 229.4 | P | 207.5 | P | --- |
| 245405_at | 164 | P | 181.3 | P | 173 | P | 234.2 | P | --- |
| 245406_at | 45.1 | A | 33.3 | A | 64.4 | A | 55.6 | A | ATRABB1A |
| 245407_at | 1392.3 | P | 1189.2 | P | 1115.3 | P | 1277.8 | P | RABB1C |
| 245408_at | 153.6 | P | 204.8 | P | 205.6 | P | 219.6 | P | --- |
| 245409_at | 20.6 | A | 8.4 | A | 28.8 | A | 19 | A | --- |
| 245410_at | 24.4 | P | 15.7 | A | 30.7 | A | 23.4 | P | ATMAP70-5 |
| 245411_at | 125.8 | P | 144.8 | P | 144.3 | P | 131.4 | P | --- |
| 245412_at | 43.1 | A | 65.4 | P | 63.1 | A | 57.3 | P | --- |
| 245413_at | 1167.3 | P | 750.6 | P | 897 | P | 757.4 | P | NS1 |
| 245414_at | 189.5 | P | 183.8 | P | 206.9 | P | 150.2 | P | --- |
| 245415_at | 1039.1 | P | 1083.7 | P | 1103.3 | P | 1210.9 | P | ATG2484-1 |
| 245416_at | 72.3 | P | 70.2 | P | 53.1 | P | 105.7 | P | --- |
| 245417_at | 274.6 | P | 277.4 | P | 308.1 | P | 278.4 | P | --- |
| 245418_at | 753.1 | P | 825.2 | P | 708 | P | 805.4 | P | --- |
| 245419_at | 3 | A | 14.5 | A | 7.9 | A | 5.1 | A | MSH4 |
| 245420_at | 102.7 | A | 134.1 | A | 130.8 | A | 143.9 | A | --- |
| 245421_at | 266.9 | P | 228.3 | P | 229 | P | 247.1 | P | --- |
| 245422_at | 2799.1 | P | 345.5 | P | 1948.4 | P | 579.1 | P | --- |
| 245423_at | 83.7 | P | 53.9 | P | 90.6 | A | 70 | P | --- |
| 245424_at | 1.2 | A | 5.4 | A | 5 | A | 2.1 | A | --- |
| 245425_at | 757.8 | P | 803.8 | P | 768.7 | P | 808.5 | P | UCH3 |
| 245426_at | 319.3 | P | 406.4 | P | 335 | P | 384.7 | P | --- |
| 245427_at | 418 | P | 276.3 | P | 257.9 | P | 326.7 | P | --- |
| 245428_at | 346.8 | P | 434 | P | 429.9 | P | 316.5 | P | --- |
| 245429_at | 42.7 | A | 27.3 | A | 95.2 | A | 10.3 | A | --- |
| 245430_at | 404.5 | P | 337.9 | P | 301.6 | P | 361.8 | P | FIP2 |
| 245431_at | 396.3 | P | 343.1 | P | 373 | P | 360.1 | P | --- |
| 245432_at | 989.8 | P | 1250.9 | P | 891.9 | P | 1004.9 | P | --- |
| 245433_at | 1217.3 | P | 1108.8 | P | 920.1 | P | 1075.5 | P | --- |
| 245434_at | 299.6 | P | 238.7 | P | 267.9 | P | 294.3 | P | --- |
| 245435_at | 27.5 | P | 58 | P | 52.8 | P | 42.5 | P | AT4G17140 |
| 245436_at | 22.7 | A | 27.8 | A | 5 | A | 29.6 | A | --- |
| 245437_at | 273.2 | P | 130.5 | P | 244.4 | P | 160.4 | P | --- |
| 245438_at | 239.2 | P | 252.7 | P | 258.2 | P | 304 | P | --- |
| 245439_at | 180.5 | P | 187.4 | P | 131.6 | P | 155.8 | P | --- |
| 245440_at | 18.5 | A | 9.4 | A | 35.8 | A | 18.9 | A | --- |
| 245441_at | 112.8 | P | 117.3 | P | 192.3 | P | 98 | P | PSD1 |

| | | | | | | | | | |
|---|---|---|---|---|---|---|---|---|---|
| 245442_at | 464.3 | P | 492.8 | P | 417.5 | P | 484.1 | P | --- |
| 245443_at | 17.7 | A | 20.5 | A | 1.7 | A | 15.6 | A | --- |
| 245444_at | 44.7 | A | 20.6 | A | 4.8 | A | 17.7 | A | ATTPS03 |
| 245445_at | 122.5 | P | 94.6 | P | 102.1 | P | 135.3 | P | --- |
| 245446_at | 466.3 | P | 611.1 | P | 510.1 | P | 511.5 | P | --- |
| 245447_at | 9.8 | A | 3.4 | A | 27.4 | A | 6.3 | A | --- |
| 245448_at | 380.2 | P | 610 | P | 273.4 | P | 492.5 | P | RPP4 |
| 245449_at | 291.9 | P | 292.3 | P | 286.3 | P | 266.1 | P | --- |
| 245450_at | 152.2 | P | 290.4 | P | 158.2 | P | 371.3 | P | --- |
| 245451_at | 281.6 | P | 789.7 | P | 555.4 | P | 651.2 | P | SNC1 |
| 245452_at | 2.7 | A | 3.3 | A | 0.9 | A | 2.4 | A | --- |
| 245453_at | 109.2 | P | 66.8 | P | 56.6 | A | 66.5 | A | --- |
| 245454_at | 56.5 | P | 31.2 | A | 39.4 | A | 35.6 | P | --- |
| 245455_at | 1.7 | A | 1.6 | A | 2.9 | A | 1.4 | A | --- |
| 245456_at | 1260.5 | P | 1781.1 | P | 984.7 | P | 1212.3 | P | RPP5 |
| 245457_s_at | 76.6 | P | 116 | P | 60.7 | A | 69.6 | A | --- |
| 245458_at | 60 | A | 109.8 | A | 57.1 | A | 90.3 | A | --- |
| 245459_at | 43.6 | P | 48.1 | P | 45.2 | A | 51.3 | P | --- |
| 245460_at | 635.5 | P | 915.5 | P | 843.8 | P | 976.2 | P | RLM3 |
| 245461_at | 71.3 | P | 84.2 | P | 89.2 | M | 69.9 | P | --- |
| 245462_at | 169.3 | P | 188.9 | P | 186.2 | P | 196.2 | P | --- |
| 245463_at | 239.4 | P | 373.3 | P | 365.5 | P | 436.3 | P | ATEXLB1 |
| 245464_at | 56.1 | A | 137.7 | P | 105.3 | P | 94.8 | P | UGLYAH |
| 245465_at | 51.3 | A | 11.8 | A | 47.2 | A | 232.2 | P | ATCSLA01 |
| 245466_at | 29.1 | A | 33.1 | P | 48.7 | P | 40.4 | P | --- |
| 245467_at | 103.1 | P | 87.8 | P | 117.1 | A | 87.2 | P | --- |
| 245468_at | 32.4 | A | 20.8 | A | 35.7 | A | 39.9 | A | --- |
| 245469_at | 6.3 | A | 8.3 | A | 5 | A | 2.6 | A | --- |
| 245470_at | 1.5 | A | 1.8 | A | 2.2 | A | 1.7 | A | --- |
| 245471_at | 1.4 | A | 0.9 | A | 12 | A | 2.6 | A | --- |
| 245472_at | 1.7 | A | 3.6 | A | 3.6 | A | 1 | A | --- |
| 245473_at | 3 | A | 2.9 | A | 4.7 | A | 0.6 | A | --- |
| 245474_at | 537.5 | P | 810.1 | P | 531.5 | P | 767.3 | P | --- |
| 245475_at | 44.9 | A | 15.7 | A | 70.8 | P | 3.6 | A | --- |
| 245476_at | 10.6 | A | 17.5 | A | 32.9 | A | 4.4 | A | --- |
| 245477_at | 159.7 | P | 156.7 | P | 163.5 | P | 188.8 | P | ARR2 |
| 245478_at | 498.4 | P | 481.3 | P | 398 | P | 451.6 | P | ARA1 |
| 245479_at | 607.3 | P | 564.6 | P | 845.4 | P | 682.6 | P | --- |
| 245480_at | 317.1 | P | 309.7 | P | 421 | P | 266.1 | P | --- |
| 245481_at | 760.1 | P | 724.5 | P | 727.3 | P | 606.8 | P | --- |
| 245482_at | 482.6 | P | 594.5 | P | 516.6 | P | 537.6 | P | --- |
| 245483_at | 11856 | P | 12590.1 | P | 11400.9 | P | 13469.2 | P | --- |
| 245484_at | 699.3 | P | 763.1 | P | 672.7 | P | 691.2 | P | ECHIA |
| 245485_at | 19.8 | A | 43.3 | A | 46.3 | A | 30.4 | A | --- |
| 245486_x_at | 284.5 | P | 236.8 | A | 215 | A | 233.4 | M | --- |
| 245487_at | 443.8 | P | 400.9 | P | 439.8 | P | 412 | P | PHYD |
| 245488_at | 3.6 | A | 24.9 | A | 20.2 | A | 12.6 | A | --- |
| 245489_at | 166.8 | P | 162.5 | P | 145.6 | P | 135.4 | P | FCA |
| 245490_at | 30.7 | A | 37.3 | A | 16.9 | A | 9.3 | A | LDL3 |
| 245491_at | 92.6 | P | 71.4 | P | 97.7 | P | 45.4 | P | LDL3 |
| 245492_at | 274.8 | P | 235.5 | P | 245 | P | 276.3 | P | SPK1 |
| 245493_at | 27.3 | P | 50.8 | P | 84 | P | 75.2 | P | --- |
| 245494_at | 1231.4 | P | 1472.3 | P | 1246.6 | P | 1507.7 | P | --- |
| 245495_at | 39.7 | A | 67.7 | M | 65.7 | P | 41.7 | A | --- |
| 245496_at | 355.4 | P | 276.3 | P | 335.6 | P | 269.6 | P | --- |
| 245497_at | 9.5 | A | 13 | A | 6.3 | A | 7.5 | A | --- |
| 245498_at | 81.5 | P | 53.9 | P | 72.5 | P | 77.4 | P | --- |
| 245499_at | 119.3 | P | 141.6 | P | 128.9 | P | 115.6 | P | INT4 |
| 245500_at | 2 | A | 2.5 | A | 3.7 | A | 2.9 | A | --- |
| 245501_at | 754.9 | P | 494.7 | P | 654.1 | P | 735 | P | --- |
| 245502_at | 817.4 | P | 596.4 | P | 1158.7 | P | 588 | P | --- |
| 245503_at | 3.1 | A | 2.3 | A | 28.5 | A | 14 | A | --- |
| 245504_at | 14.5 | P | 27.8 | A | 8.8 | A | 33.6 | P | --- |
| 245505_at | 12.6 | A | 52.4 | P | 29.8 | A | 2.6 | A | --- |
| 245506_at | 28.5 | A | 64.9 | P | 27.2 | A | 42.6 | P | --- |
| 245507_at | 3.2 | A | 3 | A | 3.8 | A | 6.2 | A | --- |
| 245508_at | 20.1 | A | 5.9 | A | 5.6 | A | 26.7 | A | --- |
| 245509_at | 64.5 | P | 59.1 | P | 59.3 | P | 51.7 | P | --- |
| 245510_at | 23.9 | A | 30.5 | A | 23.8 | A | 29.2 | A | --- |
| 245511_at | 8.9 | A | 3.1 | A | 7.2 | A | 2 | A | --- |
| 245512_at | 1520.8 | P | 1080 | P | 1659.8 | P | 1281.6 | P | --- |
| 245513_at | 937.1 | P | 995.6 | P | 919.9 | P | 951.4 | P | ATVAMP724 |
| 245514_at | 160 | P | 162.3 | P | 155.4 | P | 167.6 | P | --- |
| 245515_at | 50 | P | 43.4 | P | 57.6 | P | 48.9 | P | --- |
| 245516_at | 255.3 | P | 247.9 | P | 173 | P | 231.5 | P | --- |
| 245517_at | 323.7 | P | 352.7 | P | 189.7 | P | 276.8 | P | --- |
| 245518_at | 152.2 | P | 133.2 | P | 173.4 | P | 168.6 | P | ATRH1 |
| 245519_at | 30.4 | A | 3.9 | A | 27.3 | A | 8.5 | A | --- |
| 245520_at | 12 | A | 2.2 | A | 3.4 | A | 1.9 | A | ATTS1 |
| 245521_at | 316.3 | P | 394.2 | P | 370.4 | P | 428.4 | P | ESD4 |
| 245522_at | 107.8 | P | 123.4 | P | 159 | P | 115.5 | P | --- |
| 245523_at | 1577 | P | 811.5 | P | 2196.6 | P | 1172.1 | P | ATDI21 |
| 245524_at | 661.5 | P | 775.9 | P | 597.2 | P | 616.5 | P | --- |
| 245525_at | 1381.2 | P | 1514.9 | P | 1511.5 | P | 1387 | P | --- |
| 245526_at | 148.7 | P | 132 | P | 176.7 | P | 170.4 | P | NRPD4 |
| 245527_at | 96.4 | P | 107.3 | P | 92.9 | P | 124.5 | P | --- |
| 245528_at | 1895.4 | P | 1303.3 | P | 1770.6 | P | 1462.7 | P | PPDK |
| 245529_at | 232.7 | P | 260.4 | P | 289.1 | P | 214.4 | P | MAA3 |
| 245530_at | 22.8 | A | 29.1 | A | 53 | P | 12.5 | A | --- |
| 245531_at | 5.2 | A | 4 | A | 78.3 | A | 11.1 | A | scpl30 |
| 245532_at | 1287 | P | 1337 | P | 925.1 | P | 1279.9 | P | CYP97B3 |
| 245533_at | 285.4 | P | 282.4 | P | 242.9 | P | 272.1 | P | CCT2 |
| 245534_at | 9.1 | A | 2.2 | A | 9.5 | A | 2.5 | A | --- |
| 245535_at | 58.1 | P | 86.1 | P | 144 | P | 97.9 | P | --- |
| 245536_at | 148.3 | P | 135.4 | P | 179.1 | P | 141.1 | P | SDG2 |
| 245537_at | 15.8 | M | 24.2 | P | 25.1 | A | 38.4 | P | --- |
| 245538_at | 9.3 | A | 29.2 | A | 39.7 | A | 2.6 | A | --- |
| 245539_at | 1.6 | A | 4 | A | 8 | A | 2.5 | A | PDR2 |
| 245540_at | 113.5 | P | 117.1 | P | 144.9 | P | 147.5 | P | PDR2 |
| 245541_at | 101.1 | P | 103.8 | P | 103.4 | P | 97.3 | P | --- |
| 245542_at | 43.1 | A | 14.9 | A | 22.2 | A | 13.6 | A | --- |
| 245543_at | 462.8 | P | 256.8 | P | 360.1 | P | 258.5 | P | --- |
| 245544_at | 41 | A | 33.7 | A | 37.2 | A | 35.5 | A | --- |
| 245545_at | 81.6 | A | 33.9 | A | 61.2 | A | 41.9 | A | UGT71B5 |
| 245546_at | 35.6 | A | 49.2 | M | 95.3 | P | 52.4 | P | ATCSLB05 |
| 245547_at | 19.9 | A | 2.4 | A | 7.1 | A | 14.6 | A | CYP702A2 |
| 245548_at | 36.5 | M | 5.6 | A | 44.7 | M | 20.1 | A | CYP702A3 |
| 245549_at | 16.3 | A | 1.8 | A | 2.3 | A | 4.6 | A | ATCSLB06 |
| 245550_at | 9.9 | A | 15.8 | A | 38 | A | 18.6 | A | CYP705A1 |

| | | | | | | | | | |
|---|---|---|---|---|---|---|---|---|---|
| 245551_at | 0.9 | A | 1.6 | A | 7.8 | A | 1.5 | A | CYP705A2 |
| 245552_at | 9.5 | A | 13.8 | A | 27.5 | A | 15.1 | A | CYP705A3 |
| 245553_at | 7.9 | A | 17.3 | A | 35 | A | 2.6 | A | BARS1 |
| 245554_at | 10.4 | A | 23.9 | A | 21.2 | A | 23.7 | A | CYP705A4 |
| 245555_at | 4.1 | A | 75.3 | A | 23.3 | A | 100.6 | A | --- |
| 245556_at | 8.4 | A | 0.6 | A | 14.9 | A | 0.8 | A | --- |
| 245557_at | 400.3 | P | 468.1 | P | 407.6 | P | 442.6 | P | PUX5 |
| 245558_at | 49 | P | 79.3 | P | 74.2 | M | 77.3 | P | --- |
| 245559_at | 31.7 | A | 16.5 | A | 13.5 | A | 4 | A | --- |
| 245560_at | 12.4 | A | 20.7 | A | 45.5 | A | 50.3 | A | UGT84A1 |
| 245561_at | 177.2 | P | 101.1 | P | 159.5 | P | 121.6 | P | UGT84A4 |
| 245562_at | 423.7 | P | 426.4 | P | 453.9 | P | 521.6 | P | --- |
| 245563_at | 134.5 | P | 162.8 | P | 101.2 | P | 183 | P | CIPK4 |
| 245564_at | 76.8 | A | 75.2 | P | 97.4 | A | 78.6 | A | emb2739 |
| 245565_at | 184.9 | P | 280.7 | P | 254.9 | P | 242.5 | P | --- |
| 245566_at | 47.5 | A | 97.4 | P | 24.5 | A | 40.2 | A | --- |
| 245567_at | 2.2 | A | 6.6 | A | 17.1 | A | 39 | A | GLP9 |
| 245568_at | 31 | A | 67.6 | A | 55.3 | A | 63.1 | A | --- |
| 245569_at | 492.7 | P | 366.2 | P | 395.4 | P | 378.1 | P | NRPE7 |
| 245570_at | 6.9 | A | 5.7 | A | 4.8 | A | 7.5 | A | CLPB2 |
| 245571_at | 23.8 | A | 3.6 | A | 3.5 | A | 17.3 | A | --- |
| 245572_at | 199.8 | P | 162.3 | P | 218.1 | P | 202.5 | P | PPD2 |
| 245573_at | 41.7 | P | 12.5 | A | 47.7 | P | 31.3 | A | --- |
| 245574_at | 58.8 | P | 103.2 | P | 44.9 | A | 71.6 | P | IQD19 |
| 245575_at | 217.5 | P | 367 | P | 268.5 | P | 406.2 | P | --- |
| 245576_at | 151.1 | A | 267.7 | P | 178 | A | 244.2 | P | TCX2 |
| 245577_at | 31.9 | A | 21.2 | A | 27.9 | A | 17.8 | A | --- |
| 245578_at | 156.6 | P | 149.3 | P | 246.2 | P | 172.6 | P | ATSUV3 |
| 245579_at | 33.6 | P | 14.1 | A | 8.5 | A | 22 | A | --- |
| 245580_at | 96.9 | P | 55.3 | P | 91 | P | 68.9 | P | --- |
| 245581_at | 93.6 | P | 62.9 | P | 83.5 | P | 69 | P | --- |
| 245582_at | 75.9 | P | 70.1 | P | 83 | P | 64.4 | P | LOI1 |
| 245583_at | 202.1 | P | 172 | P | 179.7 | P | 230.2 | P | --- |
| 245584_at | 67.3 | P | 85.6 | P | 66.2 | P | 68.2 | P | ATAO1 |
| 245585_at | 17 | A | 38.6 | P | 40.2 | P | 48.8 | P | --- |
| 245586_at | 3.6 | A | 3.3 | A | 3.2 | A | 5.4 | A | --- |
| 245587_at | 157.3 | P | 186.1 | P | 157.1 | P | 190.9 | P | --- |
| 245588_at | 140.2 | P | 97.5 | P | 107.2 | P | 112.8 | P | --- |
| 245589_at | 7.5 | A | 3 | A | 31 | A | 2.8 | A | --- |
| 245590_at | 13.7 | A | 14.6 | A | 34.7 | P | 15.9 | M | --- |
| 245591_at | 0.8 | A | 1.1 | A | 0.9 | A | 15.2 | A | --- |
| 245592_at | 657.2 | P | 628.9 | P | 579.3 | P | 615.8 | P | NF-YB3 |
| 245593_at | 20 | A | 57.4 | P | 30.5 | A | 69.7 | P | IAA14 |
| 245594_at | 644.6 | P | 624.3 | P | 689.4 | P | 646.9 | P | --- |
| 245595_at | 73.9 | A | 94.1 | P | 95 | A | 66.6 | P | --- |
| 245596_at | 1.8 | A | 10.1 | A | 40.9 | A | 3.8 | A | AtPRD1 |
| 245597_at | 360.1 | P | 238.4 | P | 317.1 | P | 309.5 | P | --- |
| 245598_at | 48 | P | 101.9 | P | 61 | P | 118.5 | P | --- |
| 245599_at | 399.2 | P | 417.6 | P | 337.9 | P | 327.2 | P | RHF1A |
| 245600_at | 697.9 | P | 770.7 | P | 631.7 | P | 725 | P | --- |
| 245601_at | 396.7 | P | 560.5 | P | 348.7 | P | 433.9 | P | --- |
| 245602_at | 1006.5 | P | 1871.3 | P | 762.2 | P | 1589.4 | P | --- |
| 245603_at | 36 | A | 23.8 | A | 42.2 | A | 6.2 | A | --- |
| 245604_at | 22.9 | A | 36.6 | A | 26.6 | A | 6.8 | A | --- |
| 245605_at | 1013.9 | P | 977 | P | 1035.8 | P | 966.3 | P | --- |
| 245606_at | 105.1 | P | 103.6 | P | 137.1 | P | 125.2 | P | --- |
| 245607_at | 58.6 | P | 102.9 | P | 67 | A | 57.4 | P | --- |
| 245608_at | 243.8 | P | 306.8 | P | 259 | P | 264.9 | P | --- |
| 245609_at | 2.8 | A | 9.7 | A | 15.2 | P | 10.7 | A | --- |
| 245610_at | 29.1 | P | 39.6 | A | 40.4 | A | 78.7 | P | --- |
| 245611_at | 47.3 | P | 56.6 | P | 80.4 | A | 73.6 | M | --- |
| 245612_at | 1036 | P | 1472.9 | P | 1061.7 | P | 1549.8 | P | HCD1 |
| 245613_at | 26.3 | A | 6.6 | A | 4 | A | 7.2 | A | AT4G14450 /// ATBET12 |
| 245614_at | 21.6 | A | 2 | A | 15.3 | A | 22.1 | A | --- |
| 245615_at | 39.8 | P | 25.6 | P | 79.8 | P | 36.4 | A | --- |
| 245616_at | 84.6 | A | 101.5 | P | 149.3 | P | 114.7 | P | --- |
| 245617_at | 25.2 | A | 28.6 | P | 24 | P | 31.9 | P | --- |
| 245618_at | 290.6 | P | 275.9 | P | 344.3 | P | 258.4 | P | --- |
| 245619_at | 55 | A | 75.3 | P | 57.4 | P | 62.3 | P | --- |
| 245620_at | 49.9 | P | 40.1 | A | 42.6 | A | 40.9 | A | --- |
| 245621_at | 697.1 | P | 1019.2 | P | 738.8 | P | 959.4 | P | AAE15 |
| 245622_at | 5.5 | A | 2.3 | A | 2 | A | 1 | A | MEE48 |
| 245623_s_at | 70.8 | P | 110.9 | P | 121.6 | P | 68.9 | P | --- |
| 245624_at | 1549 | P | 331.6 | P | 2411.4 | P | 814.7 | P | --- |
| 245625_at | 1402.7 | P | 1454.5 | P | 1287.2 | P | 1301.6 | P | --- |
| 245626_at | 1625.6 | P | 1603.4 | P | 1351.7 | P | 1519.5 | P | --- |
| 245627_at | 2.9 | A | 2.9 | A | 1.8 | A | 4.6 | A | AtGolS2 |
| 245628_at | 926.1 | P | 288.5 | P | 884.6 | P | 525.1 | P | PAP1 |
| 245629_at | 3726.7 | P | 4031.3 | P | 3558.2 | P | 3777.3 | P | --- |
| 245630_at | 132 | P | 66.5 | P | 89.7 | P | 78 | P | --- |
| 245631_at | 1173.5 | P | 998.2 | P | 1382 | P | 1066 | P | OVA9 |
| 245632_at | 281.9 | P | 315.5 | P | 384.9 | P | 313.3 | P | ATRBL10 |
| 245633_at | 708.9 | P | 634.1 | P | 655 | P | 619.6 | P | AtTLP10 |
| 245634_at | 4 | A | 1.1 | A | 3.7 | A | 2.2 | A | --- |
| 245635_at | 80.4 | P | 115.9 | P | 115.6 | P | 138.7 | P | AtIDD16 |
| 245636_at | 22.8 | A | 31.7 | A | 22.4 | A | 31.5 | A | --- |
| 245637_at | 160.2 | P | 411.9 | P | 211.5 | P | 396.1 | P | --- |
| 245638_s_at | 360.4 | P | 404.7 | P | 256.1 | P | 320.5 | P | --- |
| 245639_at | 2507.1 | P | 1815.7 | P | 3057 | P | 2054.8 | P | --- |
| 245640_at | 10.8 | A | 13.4 | A | 2.5 | A | 2.5 | A | --- |
| 245641_at | 249.1 | P | 173.5 | P | 194.2 | P | 147.3 | P | --- |
| 245642_at | 8675.8 | P | 7829.3 | P | 8417.6 | P | 7625.1 | P | --- |
| 245643_at | 17.7 | A | 26.2 | A | 16.2 | A | 32 | A | MYB116 |
| 245644_at | 217.7 | P | 207.2 | P | 244.6 | P | 247.7 | P | --- |
| 245645_at | 155.2 | P | 205.4 | P | 130 | P | 191.3 | P | ATMAP70-2 |
| 245646_x_at | 22.6 | A | 2.3 | A | 32.5 | A | 16.6 | A | --- |
| 245647_at | 57.1 | A | 37.8 | A | 51.6 | A | 56 | A | --- |
| 245648_at | 9.7 | A | 4.3 | A | 8.9 | A | 2.2 | A | --- |
| 245649_at | 312.6 | P | 295.5 | P | 329.6 | P | 326.7 | P | --- |
| 245650_at | 62.9 | A | 61.6 | P | 46.9 | A | 51.2 | A | --- |
| 245651_s_at | 2144.8 | P | 2387.9 | P | 2614.2 | P | 2380 | P | --- |
| 245652_at | 18.5 | A | 1.5 | A | 14.8 | A | 3.1 | A | --- |
| 245653_at | 2 | A | 1.4 | A | 3.4 | A | 1.5 | A | FBD1 |
| 245654_at | 25.5 | P | 25.6 | P | 32.5 | P | 39.3 | P | --- |
| 245655_at | 13.3 | A | 0.9 | A | 0.8 | A | 0.1 | A | --- |
| 245656_at | 4.9 | A | 1 | A | 2.8 | A | 3 | A | --- |
| 245657_at | 70.9 | P | 161.5 | P | 112 | P | 134.1 | P | --- |
| 245658_at | 2.8 | A | 33.1 | A | 2.1 | A | 5.6 | A | RALFL4 |
| 245659_at | 378.4 | P | 205.2 | P | 335.1 | P | 207.5 | P | --- |

| | | | | | | | | | |
|---|---|---|---|---|---|---|---|---|---|
| 245660_at | 322 | P | 282.3 | P | 296.6 | P | 266.2 | P | --- |
| 245661_at | 94.9 | P | 79.8 | P | 97.8 | P | 96.2 | P | ATPUP3 |
| 245662_at | 262.6 | P | 157.2 | P | 177.4 | P | 162.6 | P | --- |
| 245663_at | 1.1 | A | 1.2 | A | 4.2 | A | 3 | A | --- |
| 245664_at | 14 | A | 10.6 | A | 16.7 | A | 1.5 | A | --- |
| 245665_at | 280 | P | 379 | P | 274 | P | 280.5 | P | --- |
| 245666_at | 490.9 | P | 387.5 | P | 379.3 | P | 382.7 | P | --- |
| 245667_at | 973.6 | P | 809.5 | P | 602.9 | P | 773.8 | P | FIP1 |
| 245668_at | 1024.7 | P | 354.1 | P | 818.1 | P | 467.1 | P | DYL1 |
| 245669_at | 16.3 | A | 17.7 | A | 28.4 | A | 32.2 | A | LEC2 |
| 245670_at | 148.7 | P | 118.3 | P | 123 | P | 132.2 | P | ATJ1 |
| 245671_at | 1070.8 | P | 546 | P | 943.7 | P | 657 | P | PUP1 |
| 245672_at | 96.5 | P | 124.9 | P | 161.9 | P | 146.1 | P | --- |
| 245673_at | 66.1 | P | 53.9 | M | 69 | P | 57 | P | --- |
| 245674_at | 3.4 | A | 29.4 | A | 43 | A | 5.4 | A | --- |
| 245675_at | 21.9 | A | 22.5 | A | 63.1 | A | 11.5 | A | --- |
| 245676_at | 24.3 | A | 134.8 | P | 77.8 | P | 110.1 | P | --- |
| 245677_at | 52.5 | P | 100.3 | P | 46.2 | P | 61.8 | P | --- |
| 245678_at | 201.6 | P | 186.1 | P | 193.7 | P | 192.5 | P | --- |
| 245679_at | 280 | P | 350.2 | P | 202.4 | P | 281.3 | P | --- |
| 245680_at | 68.3 | P | 58.3 | P | 125.9 | P | 62.7 | P | --- |
| 245681_at | 391.5 | P | 402.8 | P | 383.4 | P | 343.6 | P | --- |
| 245682_at | 390.5 | P | 508 | P | 387.9 | P | 420 | P | --- |
| 245683_at | 436.2 | P | 390.8 | P | 394.7 | P | 397.8 | P | UBP8 |
| 245684_at | 1274.2 | P | 1185.9 | P | 859.8 | P | 1013.9 | P | RHF2A |
| 245685_at | 110.6 | P | 97.5 | P | 100.2 | P | 113.4 | P | E2F1 |
| 245686_at | 2215.2 | P | 2146 | P | 2004.9 | P | 1986.1 | P | J2 |
| 245687_at | 521.9 | P | 489.2 | P | 401.6 | P | 482.6 | P | DEG15 |
| 245688_at | 781.7 | P | 1795 | P | 1158 | P | 1981.9 | P | AGP31 |
| 245689_at | 17.1 | A | 26 | A | 30.9 | A | 4.3 | A | --- |
| 245690_at | 191.9 | P | 243.7 | P | 386.7 | P | 258.2 | P | PAL3 |
| 245691_at | 241.4 | P | 220.7 | P | 230.5 | P | 210.6 | P | ELF6 |
| 245692_at | 24.4 | A | 148.2 | P | 692 | P | 241.7 | P | BHLH101 |
| 245693_at | 858.7 | P | 1016.8 | P | 699.2 | P | 1026.2 | P | WCRKC2 |
| 245694_at | 682 | P | 818 | P | 711.3 | P | 764.9 | P | --- |
| 245695_at | 142.9 | P | 201.6 | P | 191 | P | 187.3 | P | --- |
| 245696_at | 17 | A | 26.2 | A | 11.3 | A | 4.1 | A | PKS4 |
| 245697_at | 48.4 | P | 81.6 | P | 67 | M | 94.2 | P | AtMC9 |
| 245698_at | 284.2 | P | 498.8 | P | 304 | P | 445.9 | P | --- |
| 245699_at | 273 | P | 177.1 | P | 196.9 | P | 153.6 | P | --- |
| 245700_at | 31.7 | A | 49.3 | A | 37.6 | A | 58.2 | A | ACA3 |
| 245701_at | 4934.1 | P | 4225.7 | P | 4803.3 | P | 4650.4 | P | GLU1 |
| 245702_at | 131.2 | P | 133 | P | 110.2 | P | 86.7 | P | SYTC |
| 245703_at | 17.5 | A | 17.4 | A | 10.2 | A | 10 | A | --- |
| 245704_at | 249.1 | P | 283.2 | P | 326.3 | P | 271.6 | P | KTF1 |
| 245705_at | 49.3 | P | 15.9 | A | 40.9 | A | 20.4 | A | --- |
| 245706_at | 83.2 | P | 55.2 | A | 74.9 | A | 46 | A | --- |
| 245707_at | 34.3 | A | 40.5 | P | 71.9 | P | 39.4 | P | anac077 |
| 245708_at | 41.2 | A | 6.6 | A | 61 | A | 13.6 | A | --- |
| 245709_at | 224.1 | P | 168.5 | P | 197.5 | P | 171.8 | P | --- |
| 245710_at | 150.5 | P | 73.2 | P | 147.4 | P | 93.8 | M | --- |
| 245711_at | 303.3 | P | 216.8 | P | 246.3 | P | 204.7 | P | ZAT6 |
| 245712_at | 472.6 | P | 480.1 | P | 439.2 | P | 473.3 | P | ATLDA |
| 245713_at | 18.6 | A | 4.4 | A | 6.9 | A | 25.2 | A | NAMT1 |
| 245714_at | 1081.8 | P | 832.3 | P | 1066.9 | P | 986.5 | P | --- |
| 245715_s_at | 9387.7 | P | 7982.9 | P | 7189.1 | P | 8482.4 | P | --- |
| 245716_at | 632 | P | 760.8 | P | 723.7 | P | 785 | P | NDC1 |
| 245717_at | 93.6 | P | 71.7 | P | 90.3 | P | 69.5 | P | --- |
| 245718_at | 169.7 | P | 288.7 | P | 271.6 | P | 243.7 | P | --- |
| 245719_at | 44.4 | A | 83.2 | P | 32.6 | A | 67 | P | --- |
| 245720_at | 6 | A | 5.4 | A | 5.7 | A | 5.6 | A | --- |
| 245721_s_at | 354.2 | P | 327.7 | P | 339 | P | 381.3 | P | --- |
| 245722_at | 247.1 | P | 273.4 | P | 297.7 | P | 268 | P | --- |
| 245723_at | 130.1 | P | 100.4 | P | 114.3 | M | 95.8 | P | --- |
| 245724_at | 468.7 | P | 427.1 | P | 493.1 | P | 503.7 | P | --- |
| 245725_at | 76.3 | P | 145.6 | P | 80.7 | P | 123.3 | P | SUS6 |
| 245726_at | 99.7 | P | 96.1 | P | 86.7 | P | 97.9 | P | HDG11 |
| 245727_at | 169.5 | P | 253.1 | P | 213.5 | P | 223 | P | --- |
| 245728_at | 3.3 | A | 6 | A | 22.8 | A | 6.7 | A | --- |
| 245729_at | 1700.7 | P | 1434.5 | P | 2009 | P | 1473.1 | P | --- |
| 245730_at | 391.4 | P | 718.2 | P | 423.5 | P | 574.9 | P | --- |
| 245731_at | 709.5 | P | 763.3 | P | 796.6 | P | 731.3 | P | MKK9 |
| 245732_at | 300.6 | P | 308.1 | P | 242.1 | P | 291 | P | --- |
| 245733_at | 244.5 | P | 272.9 | P | 257.4 | P | 232.3 | P | --- |
| 245734_at | 384.4 | P | 351.4 | P | 323.8 | P | 283.1 | P | --- |
| 245735_at | 2.1 | A | 28.5 | M | 21.8 | A | 4.1 | A | MYB54 |
| 245736_at | 1272.2 | P | 2319 | P | 2412.4 | P | 2339.7 | P | ATDR4 |
| 245737_at | 27 | A | 43.6 | P | 22.5 | A | 38.3 | A | --- |
| 245738_at | 67.5 | A | 88 | P | 37.9 | A | 50.9 | A | --- |
| 245739_at | 153.6 | P | 279 | P | 277 | P | 330.5 | P | CYCA1;1 |
| 245740_at | 806.9 | P | 972.9 | P | 821.8 | P | 814.1 | P | AAP5 |
| 245741_at | 4.5 | A | 10 | A | 28 | A | 19.3 | A | --- |
| 245742_at | 930.8 | P | 1216.5 | P | 916.1 | P | 1098.2 | P | ALDH3H1 |
| 245743_at | 154 | P | 183.9 | P | 153.4 | P | 205.3 | P | --- |
| 245744_at | 1062.9 | P | 1483.1 | P | 1195 | P | 1428.3 | P | --- |
| 245745_at | 1572.7 | P | 1995.3 | P | 2041.2 | P | 1884.8 | P | --- |
| 245746_at | 1431.3 | P | 1327.4 | P | 1544.9 | P | 1367 | P | --- |
| 245747_at | 434 | P | 499.5 | P | 295.6 | P | 410.2 | P | --- |
| 245748_at | 516.3 | P | 420.3 | P | 407.8 | P | 499.6 | P | --- |
| 245749_at | 214 | P | 192.8 | P | 179 | P | 292.2 | P | --- |
| 245750_at | 1532.7 | P | 3018.8 | P | 1774.4 | P | 3011.2 | P | HTA10 |
| 245751_s_at | 381.4 | P | 344.1 | P | 246.2 | P | 311.6 | P | --- |
| 245752_at | 3 | A | 2.3 | A | 37.5 | A | 6.4 | A | --- |
| 245753_at | 1.4 | A | 6.2 | A | 13.7 | A | 9.2 | A | --- |
| 245754_at | 10.3 | A | 0.3 | A | 0.8 | A | 0.3 | A | --- |
| 245755_at | 56.6 | P | 65.7 | P | 41 | P | 82.5 | P | --- |
| 245756_at | 1297.1 | P | 814.9 | P | 1167.6 | P | 757.3 | P | --- |
| 245757_at | 365.3 | P | 66.8 | P | 212.1 | P | 105.3 | P | PHI-1 |
| 245758_at | 121.8 | P | 114.2 | P | 113.5 | P | 132.5 | P | KAN2 |
| 245759_at | 411.7 | P | 477.3 | P | 423.1 | P | 469 | P | --- |
| 245760_s_at | 580 | P | 505.6 | P | 578.4 | P | 448.4 | P | --- |
| 245761_at | 548 | P | 459.6 | P | 448.3 | P | 410.3 | P | --- |
| 245762_at | 56.3 | P | 103.1 | P | 86.3 | P | 78.2 | P | --- |
| 245763_at | 302.3 | P | 303.1 | P | 235.1 | P | 309.3 | P | --- |
| 245764_s_at | 211.2 | P | 265 | P | 223.9 | P | 309.6 | P | ATCSA-1 |
| 245765_at | 1215.3 | P | 1109.4 | P | 1068.8 | P | 1039.6 | P | --- |
| 245766_at | 14.6 | A | 24.9 | A | 44.7 | A | 16.4 | A | --- |
| 245767_at | 26.1 | A | 11.1 | A | 60.2 | P | 40.3 | A | --- |
| 245768_at | 3452.9 | P | 2700.4 | P | 3024.9 | P | 2701.6 | P | --- |

| | | | | | | | | | |
|---|---|---|---|---|---|---|---|---|---|
| 245769_at | 181 | P | 112.4 | P | 131.8 | P | 101 | P | INT2 |
| 245770_at | 227.2 | P | 154.3 | P | 237.4 | P | 173.4 | P | --- |
| 245771_at | 43 | P | 13.2 | A | 32.5 | A | 30.7 | A | --- |
| 245772_at | 229.9 | P | 254 | P | 286.7 | P | 228 | P | --- |
| 245773_at | 246.6 | P | 254.1 | P | 307.2 | P | 259.7 | P | --- |
| 245774_at | 326.7 | P | 430 | P | 436 | P | 455.3 | P | TCP24 |
| 245775_at | 738.1 | P | 564.2 | P | 780.5 | P | 483.2 | P | CIPK23 |
| 245776_at | 339 | P | 246.7 | P | 281 | P | 285.8 | P | --- |
| 245777_at | 108.7 | P | 233.5 | P | 130.6 | P | 130.6 | P | atnudt21 |
| 245778_at | 920.6 | P | 1219.3 | P | 827.4 | P | 1064.9 | P | --- |
| 245779_at | 2.7 | A | 1.3 | A | 21.1 | A | 1.7 | A | --- |
| 245780_at | 1485.6 | P | 1709 | P | 1525.2 | P | 1785 | P | --- |
| 245781_at | 536 | P | 517.3 | P | 476.9 | P | 513.6 | P | SBP1 |
| 245782_at | 1 | A | 1 | A | 1.8 | A | 16.4 | A | --- |
| 245783_s_at | 109.1 | P | 155.6 | P | 173.6 | M | 163.2 | P | --- |
| 245784_at | 89.6 | P | 106.7 | P | 88.3 | P | 154.6 | P | --- |
| 245785_at | 3.9 | A | 2.5 | A | 2.3 | A | 2.5 | A | ATCSLD6 |
| 245786_at | 209.4 | P | 272.4 | P | 245.2 | P | 275.5 | P | --- |
| 245787_at | 758 | P | 750.3 | P | 626.3 | P | 617.7 | P | --- |
| 245788_at | 103.9 | P | 90.2 | P | 113.6 | P | 83.4 | P | --- |
| 245789_at | 326.5 | P | 397.7 | P | 428.6 | P | 376.1 | P | --- |
| 245790_at | 1506.9 | P | 1449 | P | 1583.2 | P | 1299.6 | P | ATS1 |
| 245791_at | 1741.8 | P | 2155 | P | 1375.1 | P | 2057.3 | P | ATDAD1 |
| 245792_at | 189.9 | P | 226.4 | P | 203.8 | P | 250.5 | P | PRR1 |
| 245793_at | 1630.1 | P | 1502.5 | P | 1243 | P | 1636.1 | P | --- |
| 245794_at | 201.6 | P | 144.9 | P | 199.9 | P | 135.7 | P | XTR4 |
| 245795_at | 1477.9 | P | 1764.8 | P | 1359.3 | P | 1621.4 | P | --- |
| 245796_at | 2211.1 | P | 2002.4 | P | 2244.8 | P | 1830 | P | RCD1 |
| 245797_at | 659.4 | P | 830.3 | P | 817.9 | P | 669.3 | P | --- |
| 245798_at | 31.2 | P | 12.8 | P | 29.7 | A | 29.5 | P | --- |
| 245799_at | 387.7 | P | 108.4 | P | 238.7 | P | 125.4 | P | AtRLP6 |
| 245800_at | 200.7 | P | 193.4 | P | 250.5 | P | 225.4 | P | HSFB4 |
| 245801_at | 0.6 | A | 0.8 | A | 2.4 | A | 1.1 | A | --- |
| 245802_at | 29.4 | P | 2.5 | A | 30.3 | A | 14.4 | A | --- |
| 245803_at | 6476.3 | P | 7255.6 | P | 5691.9 | P | 6549.1 | P | RD21 |
| 245804_at | 2 | A | 15.3 | A | 1.4 | A | 3.4 | A | --- |
| 245805_at | 8.3 | A | 1.1 | A | 1.2 | A | 11.3 | A | --- |
| 245806_at | 1809.4 | P | 2543.2 | P | 2600.7 | P | 2588.2 | P | LHCA5 |
| 245807_at | 58.5 | A | 68.8 | P | 93.4 | A | 62.9 | M | RAP2.1 |
| 245808_at | 129.3 | P | 102.1 | P | 168.3 | P | 104.8 | P | ATRBP1 |
| 245809_at | 469.2 | P | 509.2 | P | 549.7 | P | 505.3 | P | XF1 |
| 245810_at | 114.5 | P | 213.8 | P | 154.8 | P | 186.6 | P | --- |
| 245811_at | 1.1 | A | 4.8 | A | 2.4 | A | 12.3 | A | --- |
| 245812_at | 3.4 | A | 0.7 | A | 0.6 | A | 2.7 | A | --- |
| 245813_at | 4.5 | A | 27.4 | A | 6.5 | A | 3.1 | A | --- |
| 245814_at | 30 | A | 80.5 | A | 17.5 | A | 82.9 | P | --- |
| 245815_at | 352.8 | P | 278.9 | P | 330.9 | P | 278.4 | P | --- |
| 245816_at | 122.8 | P | 109.4 | P | 101 | P | 140.8 | P | --- |
| 245817_at | 1002.5 | P | 877.8 | P | 1007.1 | P | 961.2 | P | --- |
| 245818_at | 152.9 | P | 265.6 | P | 158.8 | P | 275.2 | P | --- |
| 245819_at | 33 | A | 50.3 | A | 33.2 | A | 22.2 | A | CAL |
| 245820_at | 34.2 | A | 7.8 | A | 30.3 | A | 57.8 | A | --- |
| 245821_at | 414.9 | P | 505.5 | P | 489.6 | P | 465.7 | P | --- |
| 245822_at | 1255.8 | P | 1331.9 | P | 1487.5 | P | 1248.1 | P | --- |
| 245823_at | 2.6 | A | 3.6 | A | 6.7 | A | 2.3 | A | --- |
| 245824_at | 12.5 | A | 20.1 | A | 3.8 | A | 4.5 | A | ATPUP17 |
| 245825_at | 306.1 | P | 430.6 | P | 367.4 | P | 452.4 | P | ATSK42 |
| 245826_at | 11.1 | A | 22.1 | A | 30.7 | A | 20.1 | A | --- |
| 245827_at | 24.8 | P | 38.2 | P | 32.6 | A | 24 | A | --- |
| 245828_at | 358.4 | P | 462.3 | P | 440.7 | P | 455.7 | P | VIM1 |
| 245829_at | 13.9 | A | 16 | A | 21.8 | A | 13.8 | A | --- |
| 245830_at | 165.1 | P | 164.6 | P | 175.3 | P | 175.3 | P | --- |
| 245831_at | 167.2 | P | 175.2 | P | 166.4 | P | 145.3 | P | --- |
| 245832_at | 2429.5 | P | 2043.9 | P | 1980.6 | P | 2245.2 | P | EMB1144 |
| 245833_at | 49.2 | M | 35.2 | A | 53.9 | A | 24 | A | --- |
| 245834_at | 6.8 | A | 5.1 | A | 8.9 | A | 2.3 | A | --- |
| 245835_at | 4.9 | A | 14.5 | P | 31.9 | A | 17.2 | A | --- |
| 245836_at | 0.8 | A | 0.7 | A | 0.8 | A | 0.3 | A | --- |
| 245837_at | 12.4 | A | 10.2 | A | 15.2 | A | 3.1 | A | --- |
| 245838_at | 36.6 | P | 39.7 | P | 70.2 | P | 60.9 | P | --- |
| 245839_at | 13 | A | 25.2 | M | 36.5 | A | 24.6 | A | --- |
| 245840_at | 78.3 | P | 198.9 | P | 122.2 | P | 116.9 | P | --- |
| 245841_s_at | 12326.1 | P | 10821.1 | P | 12985 | P | 11628 | P | XW6 |
| 245842_at | 29.7 | A | 21.1 | A | 23.1 | A | 22.6 | A | RXF26 |
| 245843_at | 416.9 | P | 546.4 | P | 466.8 | P | 483.7 | P | --- |
| 245844_at | 316.5 | P | 376.2 | P | 491.6 | P | 276.7 | P | --- |
| 245845_at | 414.3 | P | 413.3 | P | 504.7 | P | 433.7 | P | PERK10 |
| 245846_at | 133.8 | P | 193.1 | P | 144.4 | P | 173.2 | P | --- |
| 245847_at | 24.9 | A | 41.1 | A | 68.9 | A | 46.5 | P | MYB64 |
| 245848_at | 234.2 | P | 238.5 | P | 337.2 | P | 261.3 | P | FY |
| 245849_at | 176.9 | P | 285.1 | P | 257.5 | P | 239.7 | P | --- |
| 245850_at | 45.7 | P | 69.9 | P | 80.8 | P | 68.8 | P | KEG |
| 245851_at | 250 | P | 234.7 | P | 228.2 | P | 208.7 | P | KEG |
| 245852_at | 4712.8 | P | 6096.3 | P | 5129.6 | P | 5970.5 | P | --- |
| 245853_at | 379.6 | P | 507.8 | P | 378.1 | P | 465.3 | P | --- |
| 245854_at | 4228.1 | P | 1991.9 | P | 3558.8 | P | 2644.1 | P | AAC2 |
| 245855_at | 786.9 | P | 510 | P | 751.1 | P | 407.6 | P | SULTR4;1 |
| 245856_at | 3.5 | A | 8 | A | 3.7 | A | 14.7 | A | --- |
| 245857_at | 16.4 | A | 1.8 | A | 12.5 | A | 3.6 | A | --- |
| 245858_at | 15.8 | A | 21.9 | A | 15.9 | A | 15.7 | A | --- |
| 245859_at | 169.9 | P | 343.8 | P | 273.9 | P | 322.5 | P | ATNEK3 |
| 245860_at | 51.3 | A | 93.3 | A | 72 | A | 7.7 | A | --- |
| 245861_at | 699.3 | P | 496.4 | P | 666.7 | P | 345.8 | P | --- |
| 245862_at | 1266 | P | 1205.2 | P | 1476 | P | 1171.5 | P | --- |
| 245863_s_at | 341.1 | P | 300.8 | P | 253.7 | P | 234.6 | P | --- |
| 245864_at | 102.7 | M | 107.3 | P | 98.9 | A | 155 | P | --- |
| 245865_at | 3 | A | 2.2 | A | 3.2 | A | 2.6 | A | --- |
| 245866_s_at | 590.3 | P | 488.8 | P | 351.9 | P | 471.9 | P | --- |
| 245867_at | 1190.2 | P | 1276 | P | 1212.3 | P | 1286.7 | P | ATATP-PRT1 |
| 245868_at | 1271.5 | P | 1394 | P | 1371.7 | P | 1128.5 | P | CAT2 |
| 245869_at | 49.5 | A | 46.8 | A | 82 | A | 76.3 | A | --- |
| 245870_at | 204.9 | P | 217.2 | P | 167.1 | P | 215.9 | P | --- |
| 245871_at | 21.8 | A | 13.4 | A | 27.8 | A | 20.8 | P | --- |
| 245872_at | 110.2 | P | 101.4 | P | 101.6 | P | 85.8 | P | --- |
| 245873_at | 39.6 | P | 71.3 | P | 61.9 | P | 71 | P | CIB5 |
| 245874_at | 26.6 | A | 30.9 | M | 21.3 | A | 27.4 | A | --- |
| 245875_at | 27.4 | A | 27.3 | A | 27 | A | 14.5 | A | --- |
| 245876_at | 652.2 | P | 668.5 | P | 855 | P | 711.1 | P | --- |
| 245877_at | 301 | P | 441.2 | P | 361.5 | P | 334.5 | P | --- |

| | | | | | | | | | |
|---|---|---|---|---|---|---|---|---|---|
| 245878_at | 117.3 | P | 144.7 | P | 226.5 | P | 135.2 | P | --- |
| 245879_at | 177.7 | P | 145.6 | P | 193.2 | P | 147.7 | P | ATTOC64-V |
| 245880_at | 11.2 | A | 11.3 | A | 3.3 | A | 12.6 | A | --- |
| 245881_at | 394.2 | P | 445 | P | 535.6 | P | 524.4 | P | --- |
| 245882_at | 175.6 | P | 68.1 | A | 77.2 | A | 73.3 | A | DIC3 |
| 245883_at | 31.3 | A | 16.1 | P | 23 | A | 33.3 | A | --- |
| 245884_at | 83.7 | P | 223.4 | P | 88.6 | P | 241.2 | P | --- |
| 245885_at | 1719.8 | P | 707.4 | P | 840.4 | P | 776.5 | P | EXL4 |
| 245886_at | 1908.6 | P | 1691.5 | P | 2256.7 | P | 1804.2 | P | --- |
| 245887_at | 385.1 | P | 471.3 | P | 406 | P | 441.1 | P | --- |
| 245888_at | 524.3 | P | 476.3 | P | 457 | P | 508.3 | P | --- |
| 245889_at | 1.3 | A | 96 | P | 69 | P | 97 | P | --- |
| 245890_at | 22.7 | A | 1.5 | A | 2 | A | 11.2 | A | --- |
| 245891_at | 982.5 | P | 3304.8 | P | 1685.2 | P | 3041.7 | P | AAP2 |
| 245892_at | 5.4 | A | 3.3 | A | 57.4 | A | 27.8 | A | --- |
| 245893_at | 461 | P | 377.2 | P | 412.6 | P | 425.3 | P | --- |
| 245894_at | 512 | P | 590.5 | P | 488.9 | P | 501.5 | P | --- |
| 245895_at | 396.3 | P | 475.8 | P | 543.6 | P | 507.1 | P | SRT2 |
| 245896_at | 260.4 | P | 339.1 | P | 243.5 | P | 339.2 | P | KIWI |
| 245897_at | 171.8 | P | 258.5 | P | 236.3 | P | 168.5 | P | KUP7 |
| 245898_at | 165.2 | P | 160.3 | P | 158.8 | P | 164.2 | P | --- |
| 245899_at | 257.5 | P | 257.2 | P | 236.4 | P | 187.1 | P | ALF4 |
| 245900_at | 402.6 | P | 442.1 | P | 376.1 | P | 410.8 | P | --- |
| 245901_at | 523.6 | P | 600.9 | P | 390 | P | 536.1 | P | KNAT4 |
| 245902_at | 63.4 | P | 39.3 | A | 62.9 | A | 48 | P | --- |
| 245903_at | 189.5 | P | 121.9 | P | 189.5 | P | 117.2 | P | SYTD |
| 245904_at | 245.5 | P | 221.9 | P | 202.9 | P | 221.5 | P | ATSPS2F |
| 245905_at | 1379.6 | P | 926.8 | P | 1253 | P | 826.6 | P | --- |
| 245906_at | 992.6 | P | 1040.5 | P | 1184.9 | P | 967.4 | P | --- |
| 245907_at | 14.6 | A | 8.4 | A | 7.5 | A | 2.7 | A | --- |
| 245908_at | 6.9 | A | 2.2 | A | 8.3 | A | 6.1 | A | LAC14 |
| 245909_at | 243.3 | P | 141.2 | P | 244.7 | P | 177.7 | P | --- |
| 245910_at | 671.9 | P | 632.3 | P | 659.5 | P | 650.6 | P | EICBP.B |
| 245911_at | 1452.2 | P | 1761.6 | P | 1652.9 | P | 1715.9 | P | STT3A |
| 245912_at | 119.6 | P | 89.4 | P | 218.6 | P | 141.9 | P | SULTR3;5 |
| 245913_at | 2219 | P | 2510.2 | P | 1855.3 | P | 2041.5 | P | --- |
| 245914_at | 422.4 | P | 471.2 | P | 619.5 | P | 496.2 | P | OEP80 |
| 245915_s_at | 4949.2 | P | 5944.4 | P | 5007.1 | P | 6049 | P | TUA3 |
| 245916_at | 16.3 | A | 19.6 | P | 4.8 | A | 8.5 | A | --- |
| 245917_at | 493.3 | P | 442.8 | P | 539.3 | P | 514 | P | --- |
| 245918_at | 1.6 | A | 1.4 | A | 2.7 | A | 8.4 | A | --- |
| 245919_at | 16.1 | A | 13.3 | A | 2.1 | A | 5.8 | A | --- |
| 245920_at | 32.1 | P | 18.1 | P | 40.2 | P | 20.1 | A | --- |
| 245921_at | 5.1 | A | 38.3 | A | 22.5 | A | 26.7 | A | --- |
| 245922_at | 0.4 | A | 2.3 | A | 1.9 | A | 0.6 | A | --- |
| 245923_at | 1 | A | 14.5 | A | 19 | A | 2.2 | A | --- |
| 245924_at | 2624 | P | 3360.4 | P | 2488.6 | P | 3286.7 | P | --- |
| 245925_at | 165 | P | 220.4 | P | 272.6 | P | 199.9 | P | BZO2H3 |
| 245926_at | 171.3 | P | 139.8 | P | 125.9 | P | 115.8 | P | --- |
| 245927_at | 76 | A | 85.7 | A | 101.5 | A | 60.1 | A | --- |
| 245928_s_at | 7702 | P | 1978.2 | P | 10459.1 | P | 2632.4 | P | VSP1 |
| 245929_at | 320.2 | P | 319.2 | P | 396.5 | P | 327.8 | P | --- |
| 245930_at | 313.9 | P | 330.8 | P | 357.3 | P | 330.9 | P | --- |
| 245931_at | 13.5 | A | 15.4 | A | 35.7 | A | 27.3 | A | --- |
| 245932_at | 80.8 | A | 115 | P | 76.1 | A | 88.9 | P | --- |
| 245933_at | 171.5 | P | 166.9 | P | 202.7 | P | 139.8 | P | VPS9B |
| 245934_at | 1038.2 | P | 818.1 | P | 1153 | P | 1009.7 | P | anac082 |
| 245935_at | 105.5 | P | 115.6 | P | 130.6 | P | 108.8 | P | --- |
| 245936_at | 597.1 | P | 743.3 | P | 691.9 | P | 810.7 | P | --- |
| 245937_at | 481.4 | P | 905.2 | P | 674 | P | 788.3 | P | --- |
| 245938_at | 598 | P | 740.5 | P | 730.4 | P | 741.8 | P | --- |
| 245939_at | 6901.6 | P | 7231.7 | P | 6793.9 | P | 7434.6 | P | --- |
| 245940_at | 462.6 | P | 426.2 | P | 362.5 | P | 477.8 | P | --- |
| 245941_at | 1156.6 | P | 1063.5 | P | 948.7 | P | 1223 | P | emb2734 |
| 245942_at | 39.2 | P | 33.9 | A | 53.4 | A | 36 | P | --- |
| 245943_at | 205.7 | P | 275.7 | P | 261.3 | P | 299.9 | P | --- |
| 245944_at | 17 | A | 3.8 | A | 35 | A | 20.8 | A | MSL9 |
| 245945_at | 13.4 | A | 2.4 | A | 29.1 | A | 2.2 | A | ROPGEF10 |
| 245946_at | 76.8 | A | 89.6 | A | 55.4 | A | 74.3 | A | --- |
| 245947_at | 117.3 | P | 190.2 | P | 199.4 | P | 189.6 | P | ACL5 |
| 245948_at | 2361.9 | P | 2955.2 | P | 1832.3 | P | 2538.8 | P | --- |
| 245949_at | 7433.6 | P | 7314.6 | P | 8408.1 | P | 7452.5 | P | --- |
| 245950_at | 220.7 | P | 257.5 | P | 275.6 | P | 243.5 | P | --- |
| 245951_at | 3798.9 | P | 2508.1 | P | 3403.5 | P | 2356.9 | P | ASP2 |
| 245952_at | 3957.1 | P | 4355.4 | P | 3788.1 | P | 4706.7 | P | --- |
| 245953_at | 8 | A | 5.1 | A | 8.5 | A | 4.7 | A | --- |
| 245954_at | 130 | P | 197.6 | P | 116 | P | 116.8 | P | FRS10 |
| 245955_at | 16 | A | 9.2 | A | 8.2 | A | 2.9 | A | BGLU24 |
| 245956_s_at | 7063.8 | P | 8208.8 | P | 9225 | P | 9295.5 | P | BIP2 |
| 245957_at | 1856.8 | P | 2214.7 | P | 2012.1 | P | 1970.1 | P | --- |
| 245958_at | 3.5 | A | 2.9 | A | 1.6 | A | 4.9 | A | --- |
| 245959_at | 8.6 | A | 20.2 | A | 1.7 | A | 8.4 | A | --- |
| 245960_at | 238.1 | P | 228.8 | P | 200.9 | P | 209.7 | P | S1P |
| 245961_at | 84.5 | P | 74.3 | M | 100.9 | A | 107 | P | --- |
| 245962_at | 18.9 | A | 13.6 | A | 7.4 | A | 5.6 | A | --- |
| 245963_at | 4.8 | A | 27.5 | P | 20.7 | A | 22.8 | A | --- |
| 245964_at | 74.5 | P | 47.1 | P | 29.5 | A | 46.5 | A | --- |
| 245965_at | 249.1 | P | 269.4 | P | 221.9 | P | 288.5 | P | --- |
| 245966_at | 3.8 | A | 2.5 | A | 3.3 | A | 1.8 | A | RAP2.11 |
| 245967_at | 5.4 | A | 12.3 | A | 13.1 | A | 6.7 | A | --- |
| 245968_at | 20 | A | 6.4 | A | 5.4 | A | 5.6 | A | --- |
| 245969_at | 195.1 | P | 157 | P | 169.8 | P | 116.6 | P | --- |
| 245970_at | 6.7 | A | 6.7 | A | 74.6 | A | 12.1 | A | BGAL7 |
| 245971_at | 625.9 | P | 593.8 | P | 529.2 | P | 632.6 | P | NPH4 |
| 245972_at | 193 | P | 225.5 | P | 236 | P | 197.4 | P | --- |
| 245973_at | 1.1 | A | 0.9 | A | 0.8 | A | 0.5 | A | --- |
| 245974_at | 11.1 | A | 59.3 | A | 58.5 | A | 35.2 | A | --- |
| 245975_at | 715.8 | P | 659.4 | P | 661.9 | P | 740.5 | P | --- |
| 245976_at | 90.3 | P | 30.4 | A | 43 | P | 26.3 | A | WRKY75 |
| 245977_at | 855 | P | 569 | P | 602.7 | P | 627 | P | G6PD2 |
| 245978_at | 1.1 | A | 15.2 | A | 2.6 | A | 1.3 | A | --- |
| 245979_at | 2.6 | A | 3.8 | A | 3.9 | A | 2.5 | A | ATEXO70C1 |
| 245980_at | 893 | P | 1385.9 | P | 1075.4 | P | 1256.3 | P | --- |
| 245981_at | 402.7 | P | 620.2 | P | 430.8 | P | 594.4 | P | --- |
| 245982_at | 76.9 | P | 45.5 | A | 54.5 | A | 60.2 | A | SAG29 |
| 245983_at | 423 | P | 445.5 | P | 436.3 | P | 439.4 | P | 5-FCL |
| 245984_at | 430.9 | P | 465.6 | P | 396.3 | P | 428.1 | P | --- |
| 245985_at | 3309 | P | 3343.4 | P | 3927.2 | P | 3824.6 | P | --- |
| 245986_at | 462 | P | 446.5 | P | 537.3 | P | 443.5 | P | PBS1 |

| | | | | | | | | | |
|---|---|---|---|---|---|---|---|---|---|
| 245987_at | 1725.1 | P | 1775.8 | P | 1364.5 | P | 1541.4 | P | ANAC083 |
| 245988_at | 270.8 | P | 291.8 | P | 292.3 | P | 272.2 | P | --- |
| 245989_s_at | 1696.8 | P | 1400.7 | P | 1604.6 | P | 1535.7 | P | UBQ4 |
| 245990_at | 36.6 | A | 44.8 | A | 43.1 | A | 60.7 | A | --- |
| 245991_at | 1185.9 | P | 1095.8 | P | 1107.3 | P | 951.2 | P | --- |
| 245992_at | 55.7 | P | 44.3 | P | 29.6 | M | 26.1 | P | --- |
| 245993_at | 6737.1 | P | 4615.2 | P | 6522.6 | P | 5283.7 | P | --- |
| 245994_at | 44.5 | P | 45.5 | A | 59.2 | A | 32.6 | A | --- |
| 245995_at | 17.6 | A | 5.5 | A | 39.5 | A | 15.1 | A | --- |
| 245996_at | 4 | A | 3.4 | A | 5 | A | 1.6 | A | --- |
| 245997_at | 22.8 | A | 11.3 | A | 8.1 | A | 20.9 | A | --- |
| 245998_at | 1050.9 | P | 1145.8 | P | 816.6 | P | 1185.6 | P | SUS1 |
| 245999_at | 3012.7 | P | 3307.4 | P | 2293.7 | P | 3269.7 | P | COPT5 |
| 246000_at | 29.3 | A | 39.5 | A | 23.7 | A | 29.1 | A | --- |
| 246001_at | 134.7 | P | 106 | P | 131.3 | P | 103.8 | P | --- |
| 246002_at | 230.3 | P | 391.8 | P | 324.8 | P | 422.3 | P | --- |
| 246003_at | 11906.4 | P | 11162.9 | P | 11269 | P | 10829.2 | P | CPN20 |
| 246004_at | 9801.9 | P | 12774.8 | P | 13229.6 | P | 13932.9 | P | GER3 |
| 246005_at | 459 | P | 381.8 | P | 491.8 | P | 415.2 | P | --- |
| 246006_at | 2979.8 | P | 3005.5 | P | 2566 | P | 2928 | P | YLS8 |
| 246007_at | 3367.4 | P | 3256.3 | P | 2420.3 | P | 3239 | P | FTRA2 |
| 246008_at | 256 | P | 311.5 | P | 324.3 | P | 274.1 | P | --- |
| 246009_at | 274.7 | P | 272.4 | P | 281.3 | P | 238.9 | P | ATSTE14B |
| 246010_at | 109.9 | P | 139.7 | P | 117.9 | P | 153.3 | P | --- |
| 246011_at | 281.2 | P | 337.5 | P | 265.5 | P | 359.6 | P | --- |
| 246012_at | 206.9 | P | 184.2 | P | 143.3 | P | 173.2 | P | --- |
| 246013_at | 15.8 | A | 4.7 | A | 46.8 | P | 20.4 | A | --- |
| 246014_at | 2.9 | A | 11.4 | A | 4.1 | A | 2 | A | --- |
| 246015_at | 279.4 | P | 289.9 | P | 291.7 | P | 234.7 | P | --- |
| 246016_at | 31.1 | A | 7.1 | A | 4.8 | A | 6.2 | A | AHK5 |
| 246017_at | 922.3 | P | 757.9 | P | 783.3 | P | 787.5 | P | --- |
| 246018_at | 269.1 | P | 620.4 | P | 156.9 | P | 332.4 | P | --- |
| 246019_at | 451.1 | P | 751.9 | P | 534.9 | P | 717.8 | P | --- |
| 246020_at | 436.6 | P | 452.7 | P | 475.6 | P | 429.3 | P | --- |
| 246021_at | 604.6 | P | 990.8 | P | 650.2 | P | 1030.3 | P | --- |
| 246022_at | 1.8 | A | 4.1 | A | 19.6 | A | 13 | A | --- |
| 246023_at | 42.2 | P | 33.9 | M | 65.7 | P | 30.4 | P | EIL2 |
| 246024_at | 3.4 | A | 3.6 | A | 24.8 | A | 6.1 | A | --- |
| 246025_at | 20.5 | A | 14.3 | A | 23.9 | A | 26.7 | A | --- |
| 246026_at | 472.7 | P | 535.2 | P | 476.4 | P | 551.1 | P | --- |
| 246027_at | 599.9 | P | 504.3 | P | 519 | P | 516.8 | P | --- |
| 246028_at | 330 | P | 404.9 | P | 368.1 | P | 340.3 | P | --- |
| 246029_at | 6181.7 | P | 6405.4 | P | 5811.5 | P | 5544.3 | P | --- |
| 246030_at | 2696.5 | P | 1895.7 | P | 2630.1 | P | 2071.2 | P | --- |
| 246031_at | 644.1 | P | 599.1 | P | 795.4 | P | 633.1 | P | --- |
| 246032_s_at | 2.7 | A | 4.7 | A | 26.9 | A | 2.1 | A | --- |
| 246033_at | 4424.2 | P | 5027.3 | P | 4542 | P | 5250.7 | P | HEMC |
| 246034_at | 229.4 | P | 185.1 | P | 206.8 | P | 163.4 | P | --- |
| 246035_at | 3584 | P | 2836.4 | P | 3375.6 | P | 2930.8 | P | --- |
| 246036_at | 713.8 | P | 585.9 | P | 813.8 | P | 693.9 | P | AtAGAL2 |
| 246037_at | 161.9 | P | 184.7 | P | 202.8 | P | 179 | P | --- |
| 246038_s_at | 127 | P | 129.2 | P | 129.1 | P | 172.2 | P | atnudt20 |
| 246039_at | 535.3 | P | 315.3 | P | 354.2 | P | 304.6 | P | --- |
| 246040_at | 1870.4 | P | 2104.5 | P | 2165.2 | P | 2169.6 | P | --- |
| 246041_at | 1144.9 | P | 1396.7 | P | 1136.7 | P | 1391 | P | --- |
| 246042_at | 12451.4 | P | 5501.3 | P | 11844.4 | P | 6973.2 | P | --- |
| 246043_at | 220 | P | 233 | P | 213.9 | P | 229.7 | P | --- |
| 246044_at | 728.4 | P | 794.1 | P | 650.2 | P | 713.9 | P | CDPK19 |
| 246045_at | 340.4 | P | 324.7 | P | 321.8 | P | 347.9 | P | --- |
| 246046_at | 14.7 | A | 28.3 | A | 38.4 | A | 35.9 | A | --- |
| 246047_at | 25.4 | P | 36.7 | A | 55 | A | 49.3 | A | --- |
| 246048_at | 1.3 | A | 13.7 | A | 1.9 | A | 0.6 | A | --- |
| 246049_at | 21.8 | A | 3.7 | A | 6.2 | A | 2.1 | A | --- |
| 246050_s_at | 1179.3 | P | 1254.8 | P | 1218.8 | P | 1261.3 | P | --- |
| 246051_at | 3288.2 | P | 4113.7 | P | 4516.8 | P | 4548.6 | P | GME |
| 246052_at | 99.9 | P | 83.6 | P | 119.2 | P | 112.2 | P | --- |
| 246053_at | 277 | P | 334.7 | P | 229 | P | 255.3 | P | --- |
| 246054_at | 9.5 | A | 17.7 | A | 14.9 | A | 24.5 | A | --- |
| 246055_at | 1160.5 | P | 1227.6 | P | 1060.2 | P | 1212.4 | P | AtAGAL1 |
| 246056_at | 232.4 | P | 320.4 | P | 378.4 | P | 324.9 | P | --- |
| 246057_at | 802.7 | P | 940.5 | P | 791.1 | P | 944.9 | P | --- |
| 246058_at | 292.1 | P | 314.3 | P | 297 | P | 320.7 | P | --- |
| 246059_at | 320.8 | P | 386.9 | P | 322.2 | P | 256.1 | P | --- |
| 246060_at | 734.3 | P | 669.8 | P | 747.9 | P | 714.8 | P | --- |
| 246061_at | 331.8 | P | 310.4 | P | 372.3 | P | 317.5 | P | RANGAP2 |
| 246062_at | 1295.2 | P | 1262.6 | P | 1260.1 | P | 1120.6 | P | --- |
| 246063_at | 78.1 | P | 106.2 | P | 126.3 | P | 118.1 | P | --- |
| 246064_at | 957.8 | P | 806.2 | P | 976.5 | P | 806.6 | P | --- |
| 246065_at | 8.3 | A | 3.6 | A | 25 | A | 9.6 | A | CPK34 |
| 246066_at | 502.6 | P | 519.8 | P | 381.9 | P | 447.3 | P | SMG7 |
| 246067_at | 6.1 | A | 8.7 | A | 5.2 | A | 16.8 | A | --- |
| 246068_at | 15395.4 | P | 14635.4 | P | 13522.8 | P | 14622 | P | --- |
| 246069_at | 268 | P | 336.7 | P | 378.9 | P | 372.3 | P | --- |
| 246070_at | 3561.5 | P | 2651.3 | P | 3689.5 | P | 3258.5 | P | --- |
| 246071_at | 472.1 | P | 531.2 | P | 277.3 | P | 420.2 | P | SPX1 |
| 246072_at | 42.8 | M | 43.1 | P | 67.1 | P | 38.8 | A | PI |
| 246073_at | 1712.7 | P | 1784.9 | P | 1634.1 | P | 1575.9 | P | --- |
| 246074_at | 582.4 | P | 627.5 | P | 733.2 | P | 607.4 | P | --- |
| 246075_at | 88.4 | P | 136.5 | P | 92.4 | P | 125.9 | P | MGD2 |
| 246076_at | 2586.2 | P | 3023.9 | P | 3040.5 | P | 3110.2 | P | ATSPS1F |
| 246077_at | 13 | A | 5.4 | A | 24.1 | A | 20.4 | A | CHR42 |
| 246078_s_at | 21.1 | M | 33.5 | P | 70.6 | P | 28.6 | A | --- |
| 246079_at | 123.3 | A | 130.2 | P | 105.8 | A | 117.1 | P | --- |
| 246080_at | 32.3 | P | 9.5 | A | 29.9 | A | 5.5 | A | --- |
| 246081_s_at | 536.9 | P | 543.1 | P | 321.2 | P | 463.2 | P | --- |
| 246082_at | 236.8 | P | 180.4 | P | 230.3 | P | 156.6 | P | EFR |
| 246083_at | 81.4 | P | 119.6 | P | 135.8 | P | 106.5 | P | AL5 |
| 246084_at | 79.5 | P | 82.8 | P | 104.1 | P | 82.1 | P | ATBRXL4 |
| 246085_at | 109.5 | P | 113.4 | P | 138.2 | P | 109.4 | P | ATBRXL4 |
| 246086_at | 21.2 | A | 9.3 | A | 60.5 | A | 48.5 | A | --- |
| 246087_at | 180.9 | P | 195.4 | P | 225.6 | P | 231.1 | P | --- |
| 246088_at | 536.1 | P | 400.6 | P | 506.7 | P | 439.4 | P | --- |
| 246089_at | 2103.5 | P | 1859.4 | P | 1958.2 | P | 2060.3 | P | RBX1 |
| 246090_at | 1159.7 | P | 1075 | P | 1086.4 | P | 1097.8 | P | WAV2 |
| 246091_at | 424.8 | P | 412 | P | 517.4 | P | 479.7 | P | --- |
| 246092_at | 3710.5 | P | 3832.5 | P | 3302.7 | P | 4044.5 | P | --- |
| 246093_at | 6.6 | A | 27.9 | A | 27.2 | A | 4.9 | A | --- |
| 246094_at | 295.1 | P | 184.5 | P | 300.8 | P | 242.3 | P | --- |
| 246095_at | 171.3 | P | 180.1 | P | 126.9 | P | 162.9 | P | --- |

| | | | | | | | | | |
|---|---|---|---|---|---|---|---|---|---|
| 246096_at | 11.8 | A | 4 | A | 5 | A | 2.3 | A | BETAG4 |
| 246097_at | 417.2 | P | 507.6 | P | 494.5 | P | 565.1 | P | HHP1 |
| 246098_at | 596.3 | P | 487.7 | P | 470.9 | P | 511.8 | P | --- |
| 246099_at | 6393.6 | P | 3612.6 | P | 3488 | P | 2352.1 | P | ATBCB |
| 246100_at | 15.8 | A | 2.2 | A | 35.7 | A | 14.3 | A | --- |
| 246101_at | 2.6 | A | 50.3 | A | 6.1 | A | 17.2 | A | --- |
| 246102_at | 15.1 | A | 1.3 | A | 2.8 | A | 17.1 | P | --- |
| 246103_at | 55.4 | P | 110.1 | P | 88.5 | P | 113.6 | P | AN3 |
| 246104_at | 24.2 | A | 58.8 | A | 11 | A | 32.7 | A | WRKY74 |
| 246105_at | 19.3 | A | 1.2 | A | 1.9 | A | 7 | A | --- |
| 246106_at | 0.9 | A | 1.4 | A | 2.2 | A | 2.5 | A | --- |
| 246107_at | 31.4 | M | 23.6 | A | 30.5 | A | 31.8 | A | --- |
| 246108_at | 181.3 | P | 154 | P | 111.5 | P | 143.2 | P | --- |
| 246109_at | 271.5 | P | 324.8 | P | 295.5 | P | 313.4 | P | --- |
| 246110_at | 931.5 | P | 910.9 | P | 944.4 | P | 822.3 | P | --- |
| 246111_at | 235 | P | 310 | P | 256.2 | P | 285.9 | P | --- |
| 246112_at | 258.9 | P | 340.9 | P | 381.6 | P | 304.7 | P | --- |
| 246113_at | 33.9 | P | 24.1 | A | 47.2 | A | 31.3 | P | --- |
| 246114_at | 2489.2 | P | 901.4 | P | 2266.5 | P | 830.3 | P | DIN10 |
| 246115_at | 371.4 | P | 413.9 | P | 324.3 | P | 376.1 | P | --- |
| 246116_at | 23.8 | A | 22.3 | P | 33.3 | P | 26.1 | P | --- |
| 246117_at | 135.8 | P | 224.1 | P | 251.3 | P | 228 | P | DCL4 |
| 246118_at | 19.5 | A | 8.5 | A | 20.2 | A | 22 | A | BG5 |
| 246119_at | 1104.2 | P | 1187.5 | P | 980.5 | P | 1143.8 | P | TIP1 |
| 246120_at | 364 | P | 368.3 | P | 389.1 | P | 418.5 | P | --- |
| 246121_at | 16.1 | A | 13.6 | A | 23.8 | A | 20.2 | A | --- |
| 246122_at | 824.1 | P | 789.7 | P | 820.7 | P | 835.7 | P | PHT4;5 |
| 246123_at | 3.2 | A | 7.9 | A | 40.5 | A | 0.9 | A | --- |
| 246124_at | 1524.1 | P | 1694.7 | P | 1864 | P | 1832.8 | P | --- |
| 246125_at | 282.5 | P | 371.9 | P | 312.7 | P | 344.1 | P | --- |
| 246126_at | 853.8 | P | 1162.9 | P | 964.3 | P | 1100.8 | P | ATNUDX19 |
| 246127_s_at | 26 | A | 15.1 | A | 40.3 | A | 26.5 | A | --- |
| 246128_at | 3.9 | A | 24.9 | A | 34 | A | 8.7 | A | --- |
| 246129_at | 10.9 | A | 9.9 | A | 8.8 | A | 1.5 | A | --- |
| 246130_at | 24.7 | A | 22.9 | A | 29.6 | A | 17.8 | A | --- |
| 246131_at | 809 | P | 1019.4 | P | 908.6 | P | 912.2 | P | B73 |
| 246132_at | 52 | A | 121.4 | P | 42 | A | 107.8 | P | ATRAD51 |
| 246133_at | 483.6 | P | 429.6 | P | 489.4 | P | 311 | P | AAO1 |
| 246134_at | 217.9 | P | 236.2 | P | 238.2 | P | 269.6 | P | TSL |
| 246135_at | 352.4 | P | 416.3 | P | 395 | P | 445.3 | P | --- |
| 246136_at | 51.3 | A | 28.3 | A | 44.1 | A | 35.7 | A | --- |
| 246137_at | 76.5 | P | 139.5 | P | 112.3 | P | 148 | P | LSH1 |
| 246138_at | 28.9 | P | 37.5 | P | 71.8 | P | 30.7 | P | --- |
| 246139_at | 434.3 | P | 420.5 | P | 500.2 | P | 409.9 | P | --- |
| 246140_at | 251.4 | P | 255.4 | P | 286.8 | P | 263.1 | P | --- |
| 246141_at | 98.7 | P | 164 | P | 128.9 | P | 127.5 | P | --- |
| 246142_at | 11 | A | 24.5 | M | 20.9 | A | 7.5 | A | --- |
| 246143_at | 575.2 | P | 583.5 | P | 570.4 | P | 638.4 | P | --- |
| 246144_at | 161.8 | P | 84.1 | P | 112.4 | P | 90.3 | P | --- |
| 246145_at | 2 | A | 14.6 | A | 4.2 | A | 1.6 | A | --- |
| 246146_at | 457.6 | P | 655.6 | P | 457.8 | P | 461.2 | P | --- |
| 246147_s_at | 1619.6 | P | 1648.5 | P | 1615.7 | P | 1593.4 | P | RPT6A |
| 246148_at | 246.2 | P | 262.2 | P | 228.5 | P | 260.4 | P | --- |
| 246149_at | 22.9 | A | 10.2 | A | 37.1 | A | 20.9 | A | --- |
| 246150_at | 175.1 | P | 316 | P | 192.3 | P | 258.8 | P | --- |
| 246151_at | 92.2 | P | 72.5 | P | 48.6 | P | 81.5 | P | --- |
| 246152_at | 484.3 | P | 359.3 | P | 594.6 | P | 404.7 | P | IPT9 |
| 246153_s_at | 4676.2 | P | 5272.9 | P | 4268.8 | P | 4474.5 | P | RAN-1 |
| 246154_at | 3595.7 | P | 4663.6 | P | 4576.6 | P | 4075.5 | P | --- |
| 246155_at | 98.6 | P | 82.5 | P | 131 | P | 92.9 | P | --- |
| 246156_at | 2402.8 | P | 2628.2 | P | 2515.2 | P | 2680 | P | --- |
| 246157_at | 3181.5 | P | 2776.3 | P | 3260 | P | 3032.5 | P | --- |
| 246158_at | 1737.5 | P | 1428.8 | P | 1576.9 | P | 1485.3 | P | --- |
| 246159_at | 506.5 | P | 599.8 | P | 611.8 | P | 641.6 | P | --- |
| 246160_at | 365.2 | P | 491.3 | P | 419.2 | P | 476.2 | P | FBX2 |
| 246161_at | 1858.9 | P | 1869 | P | 2432 | P | 1862.5 | P | JAZ12 |
| 246162_at | 460 | P | 589.2 | P | 575.7 | P | 481.7 | P | --- |
| 246163_at | 13 | A | 4.9 | A | 5.1 | A | 22.3 | A | --- |
| 246164_at | 20 | A | 14.4 | A | 0.8 | A | 1.3 | A | --- |
| 246165_at | 39.1 | A | 2.7 | A | 18.5 | A | 15.6 | A | --- |
| 246166_at | 19.8 | A | 4.6 | A | 18.4 | A | 4.2 | A | --- |
| 246167_at | 5.3 | A | 36.8 | A | 79 | A | 21.1 | A | --- |
| 246168_at | 1.2 | A | 1 | A | 4.5 | A | 0.9 | A | --- |
| 246169_at | 383.1 | P | 404.7 | P | 393.6 | P | 359.5 | P | --- |
| 246170_at | 1508.3 | P | 1787 | P | 1683.2 | P | 1758.5 | P | --- |
| 246171_at | 1144.7 | P | 1123.1 | P | 964.8 | P | 918.7 | P | --- |
| 246172_s_at | 5.4 | A | 3.4 | A | 26 | A | 27.1 | A | ACS1 |
| 246173_s_at | 738.5 | P | 755.3 | P | 826.2 | P | 847.1 | P | --- |
| 246174_s_at | 12.3 | A | 3.5 | A | 3.8 | A | 9.2 | A | --- |
| 246175_s_at | 79.8 | P | 79.7 | P | 63.6 | P | 55.9 | A | --- |
| 246176_at | 5.8 | A | 3.2 | A | 1.8 | A | 1.9 | A | --- |
| 246177_at | 5.2 | A | 16.5 | A | 1.9 | A | 6.7 | A | --- |
| 246178_s_at | 31.5 | P | 33.9 | P | 10.2 | A | 22 | P | --- |
| 246179_at | 1.6 | A | 21.3 | A | 31 | A | 25.8 | A | --- |
| 246180_at | 394.9 | P | 416.2 | P | 426.1 | P | 404.9 | P | --- |
| 246181_at | 2.8 | A | 3.1 | A | 31.3 | A | 6.2 | A | --- |
| 246182_at | 25.6 | A | 31.8 | P | 68 | P | 42.2 | P | --- |
| 246183_at | 96.5 | A | 76.5 | A | 45.4 | A | 44.9 | A | --- |
| 246184_at | 2000.8 | P | 2161.1 | P | 1691.3 | P | 2030.2 | P | --- |
| 246185_at | 340 | P | 441.5 | P | 441.6 | P | 501.2 | P | ATMS3 |
| 246186_at | 27.4 | A | 12.4 | A | 29.1 | A | 1.4 | A | --- |
| 246187_at | 76.7 | P | 59.9 | P | 88.3 | P | 53.8 | A | AT5G21020 |
| 246188_at | 53.1 | P | 58.3 | P | 29.4 | A | 55.9 | M | --- |
| 246189_at | 386.1 | P | 414.5 | P | 335.1 | P | 384 | P | --- |
| 246190_at | 171.1 | P | 219.1 | P | 239.9 | P | 173.7 | P | ATBPM5 |
| 246191_at | 3528.8 | P | 3042.7 | P | 3671.8 | P | 3516 | P | --- |
| 246192_at | 3193.6 | P | 2949.7 | P | 2750 | P | 2685 | P | EIF2 BETA |
| 246193_at | 105.4 | P | 108.6 | P | 146.2 | P | 116.1 | P | --- |
| 246194_at | 2117 | P | 1883.1 | P | 1958.1 | P | 2274.5 | P | ACD2 |
| 246195_at | 290.2 | P | 286.3 | P | 356.1 | P | 283.6 | P | UBC17 |
| 246196_at | 346.9 | P | 244.7 | P | 309.8 | P | 256.2 | P | --- |
| 246197_at | 12.9 | A | 11.5 | P | 16.6 | A | 16.9 | A | CEN2 |
| 246198_at | 1009.2 | P | 1590.3 | P | 1092.3 | P | 1473.8 | P | GGPS1 |
| 246199_at | 1404.1 | P | 1504.9 | P | 1573.3 | P | 1510.7 | P | --- |
| 246200_at | 179.9 | P | 197.4 | P | 205 | P | 191.6 | P | --- |
| 246201_at | 729.5 | P | 784.2 | P | 588.2 | P | 765.4 | P | --- |
| 246202_at | 1095.2 | P | 1206 | P | 1374.4 | P | 1222.7 | P | MAP1D |
| 246203_at | 53.7 | A | 101.4 | P | 68.9 | M | 100.7 | P | --- |
| 246204_at | 331.3 | P | 352.8 | P | 476.8 | P | 400 | P | --- |

| | | | | | | | | | |
|---|---|---|---|---|---|---|---|---|---|
| 246205_at | 608.5 | P | 542.6 | P | 733.7 | P | 519.2 | P | --- |
| 246206_at | 32 | A | 29.7 | A | 32.8 | A | 26.1 | A | PBRP |
| 246207_at | 380 | P | 387.1 | P | 446.4 | P | 444 | P | --- |
| 246208_at | 11.9 | A | 30.1 | A | 8.8 | A | 9.1 | A | SFH12 |
| 246209_at | 188.9 | P | 199.1 | P | 196.2 | P | 153 | P | BLH2 |
| 246210_at | 409.7 | P | 432.5 | P | 468 | P | 534.9 | P | --- |
| 246211_at | 895 | P | 764.6 | P | 616.3 | P | 686.6 | P | GBF1 |
| 246212_at | 151.3 | P | 173.9 | P | 170.9 | P | 123.2 | P | SPT |
| 246213_at | 328.2 | P | 339.9 | P | 417.9 | P | 338.7 | P | ATLCB1 |
| 246214_at | 985 | P | 568.8 | P | 534.3 | P | 502.9 | P | HSF4 |
| 246215_at | 598.4 | P | 408.4 | P | 476.1 | P | 394.1 | P | --- |
| 246216_at | 124.7 | P | 206.6 | P | 129.2 | P | 130 | P | ROT3 |
| 246217_at | 266.2 | P | 268.2 | P | 222.8 | P | 271.2 | P | AP2 |
| 246218_at | 811 | P | 857.9 | P | 704.3 | P | 762.1 | P | RCE1 |
| 246219_at | 2480.2 | P | 2143.4 | P | 2226.9 | P | 2068.8 | P | APP1 |
| 246220_at | 425.2 | P | 480.8 | P | 386.6 | P | 467 | P | --- |
| 246221_at | 1130.3 | P | 768.9 | P | 829.5 | P | 816 | P | SMP2 |
| 246222_at | 146.5 | P | 119.4 | P | 122.3 | P | 132 | P | RAP2.10 |
| 246223_at | 294.7 | P | 317.2 | P | 310.9 | P | 314.5 | P | IRX14 |
| 246224_at | 31 | A | 30 | A | 59.4 | M | 32.4 | A | --- |
| 246225_at | 734.1 | P | 643.5 | P | 777 | P | 697.8 | P | LEJ2 |
| 246226_at | 1360.3 | P | 1656.4 | P | 1149.9 | P | 1712.9 | P | HCF164 |
| 246227_at | 11.9 | A | 5.2 | A | 25.5 | A | 4 | A | --- |
| 246228_at | 62.9 | M | 169.5 | P | 139.5 | P | 189.2 | P | --- |
| 246229_at | 3.9 | A | 28.6 | A | 8.1 | A | 19.1 | A | sks15 |
| 246230_at | 142.5 | P | 289.4 | P | 195.7 | P | 224.6 | P | --- |
| 246231_at | 388.7 | P | 375.9 | P | 446.6 | P | 445 | P | --- |
| 246232_at | 638.9 | P | 507 | P | 525.9 | P | 495 | P | EMB2754 |
| 246233_at | 263 | P | 309.7 | P | 240.3 | P | 226 | P | --- |
| 246234_at | 301.2 | P | 413.5 | P | 440.6 | P | 442.2 | P | --- |
| 246235_at | 194.6 | P | 184.6 | P | 125.7 | P | 177.4 | P | --- |
| 246236_at | 92 | A | 101.5 | A | 125.1 | A | 106.9 | P | --- |
| 246237_at | 1345.1 | P | 1435.8 | P | 1531.2 | P | 1356.3 | P | --- |
| 246238_at | 207.5 | P | 232.4 | P | 103.4 | P | 151.7 | P | --- |
| 246239_at | 271.4 | P | 209.2 | P | 191.9 | P | 214 | P | --- |
| 246240_at | 302.5 | P | 287.4 | P | 353.5 | P | 304.4 | P | --- |
| 246241_at | 4.8 | A | 20.7 | A | 31.2 | A | 5.3 | A | PLP4 |
| 246242_at | 23.9 | A | 17 | A | 29.4 | A | 25.6 | A | --- |
| 246243_at | 186.8 | P | 174 | P | 212.3 | P | 149.9 | P | --- |
| 246244_at | 242.8 | P | 178.3 | P | 242.9 | P | 199.9 | P | --- |
| 246245_at | 715.1 | P | 712 | P | 692.4 | P | 689.3 | P | --- |
| 246246_at | 120.9 | P | 54 | P | 110.9 | P | 71.9 | P | --- |
| 246247_at | 1061 | P | 661.4 | P | 1037.8 | P | 768.9 | P | --- |
| 246248_at | 1.4 | A | 11.6 | A | 1.7 | A | 1 | A | --- |
| 246249_at | 626.6 | P | 431.8 | P | 686.9 | P | 436.8 | P | --- |
| 246250_at | 42.1 | P | 29 | P | 48.2 | P | 42 | P | CP1 |
| 246251_at | 33.4 | A | 15.5 | A | 47.4 | A | 12.1 | A | --- |
| 246252_s_at | 0.5 | A | 4.5 | A | 5.8 | A | 3.2 | A | PLP1 |
| 246253_at | 504.1 | P | 377.6 | P | 536.3 | P | 379 | P | MYB73 |
| 246254_at | 3.9 | A | 14 | A | 2.3 | A | 25.9 | A | ATMPK14 |
| 246255_at | 192.1 | P | 182.7 | P | 164.6 | P | 174.9 | P | ATPBRP |
| 246256_at | 22.1 | A | 6.2 | A | 2 | A | 11.4 | A | --- |
| 246257_at | 1333 | P | 1445.3 | P | 1367.7 | P | 1254.4 | P | ATU2AF65A |
| 246258_at | 60 | P | 49.4 | P | 54.5 | P | 57.2 | P | --- |
| 246259_at | 469.7 | P | 340.4 | P | 469.9 | P | 365.2 | P | --- |
| 246260_at | 701.9 | P | 414 | P | 535.5 | P | 392.8 | P | --- |
| 246261_at | 161.5 | P | 131 | P | 136.7 | P | 119.3 | P | --- |
| 246262_at | 198.7 | P | 211.2 | P | 174.9 | P | 203.6 | P | --- |
| 246263_at | 716.5 | P | 837.2 | P | 659.9 | P | 756.2 | P | --- |
| 246264_at | 118.6 | P | 141.7 | P | 144.5 | P | 85.6 | P | FRL2 |
| 246265_at | 657.1 | P | 621.2 | P | 852.1 | P | 658.2 | P | AT-IE |
| 246266_at | 377 | P | 487 | P | 350.1 | P | 510.2 | P | NFD3 |
| 246267_at | 10497.8 | P | 10732.3 | P | 10838.5 | P | 10956.9 | P | ACBP6 |
| 246268_at | 1813.8 | P | 1948.2 | P | 1453.3 | P | 1894.8 | P | CYP97A3 |
| 246269_at | 90.8 | P | 192.1 | P | 114 | P | 204.1 | P | --- |
| 246270_at | 1424.4 | P | 1151.4 | P | 1092.2 | P | 1040.8 | P | --- |
| 246271_at | 23.9 | A | 36.1 | A | 50.1 | A | 33.8 | A | --- |
| 246272_at | 317.9 | P | 179.9 | P | 260.3 | P | 166.2 | P | MES9 |
| 246273_at | 2.6 | A | 164.9 | P | 296.4 | P | 271.7 | P | --- |
| 246274_at | 23 | A | 25.1 | A | 45.3 | A | 50.1 | A | --- |
| 246275_at | 223.6 | P | 488.6 | P | 371.6 | P | 386.9 | P | BEE2 |
| 246276_at | 943.3 | P | 901.3 | P | 1181.9 | P | 837 | P | HMA1 |
| 246277_at | 15.1 | A | 3.2 | A | 17.2 | A | 4.5 | A | --- |
| 246278_at | 306.5 | P | 283.2 | P | 266.7 | P | 248.9 | P | --- |
| 246279_at | 4.1 | A | 3.5 | A | 1.8 | A | 1.6 | A | ATHB40 |
| 246280_at | 595.9 | P | 848.6 | P | 611.6 | P | 793.7 | P | --- |
| 246281_at | 407.3 | P | 481.1 | P | 344.4 | P | 595.8 | P | NAPRT1 |
| 246282_at | 180.9 | P | 122.9 | P | 181.8 | P | 122.3 | P | --- |
| 246283_at | 809.2 | P | 604.2 | P | 591.7 | P | 455.4 | P | --- |
| 246284_at | 329.4 | P | 508.4 | P | 395 | P | 409.8 | P | --- |
| 246285_at | 353.1 | P | 338.9 | P | 345 | P | 332.8 | P | --- |
| 246286_at | 4410.7 | P | 3538.4 | P | 3553.6 | P | 3249.8 | P | --- |
| 246287_at | 296.7 | P | 289.3 | P | 243.1 | P | 265.5 | P | --- |
| 246288_at | 805.4 | P | 1261.6 | P | 745.2 | P | 1117 | P | --- |
| 246289_at | 2236.1 | P | 2117.1 | P | 2011.3 | P | 1783.1 | P | --- |
| 246290_at | 2255.8 | P | 3458.8 | P | 2191.9 | P | 3215.8 | P | CAM3 |
| 246291_at | 201.8 | P | 196.7 | P | 302.4 | P | 209.2 | P | --- |
| 246292_at | 1257.4 | P | 1348.4 | P | 1213.4 | P | 1284.9 | P | UBA2A |
| 246293_at | 241.6 | P | 535.8 | P | 259.6 | P | 367.6 | P | SIB1 |
| 246294_at | 9000.5 | P | 8932.2 | P | 7504.1 | P | 8346.5 | P | PSRP5 |
| 246295_at | 207.7 | P | 178 | P | 218.4 | P | 175.5 | P | CIP111 |
| 246296_at | 284.6 | P | 240.3 | P | 323.7 | P | 321.8 | P | --- |
| 246297_at | 70.3 | P | 42.5 | A | 24.6 | A | 59.1 | A | --- |
| 246298_at | 231.7 | P | 324.4 | P | 322.5 | P | 322.3 | P | ETO1 |
| 246299_at | 1.8 | A | 3.6 | A | 14.1 | A | 0.8 | A | EM1 |
| 246300_at | 315.9 | P | 335.4 | P | 267.3 | P | 338.7 | P | SAC8 |
| 246301_at | 604.5 | P | 813.7 | P | 555.3 | P | 788.8 | P | CPK13 |
| 246302_at | 2103 | P | 1749.6 | P | 1083.5 | P | 1301.7 | P | CAX3 |
| 246303_at | 615 | P | 1256.7 | P | 1039.1 | P | 1092.8 | P | --- |
| 246304_at | 2362.7 | P | 2097.9 | P | 2515.4 | P | 1998 | P | ACX4 |
| 246305_at | 324.3 | P | 401.2 | P | 259 | P | 307.9 | P | --- |
| 246306_at | 567.8 | P | 404.7 | P | 525.9 | P | 506.3 | P | ATBAG4 |
| 246307_at | 4234.9 | P | 3051.7 | P | 4905.8 | P | 3573.2 | P | ATG2 |
| 246308_at | 2895.9 | P | 3443.8 | P | 3391.1 | P | 3547.8 | P | G4 |
| 246309_at | 465.7 | P | 303.4 | P | 452 | P | 323.4 | P | ATG1 |
| 246310_at | 424.2 | P | 319.5 | P | 656.7 | P | 405 | P | SULTR3;1 |
| 246311_at | 637.6 | P | 706.7 | P | 745.7 | P | 647.9 | P | HMGB1 |
| 246312_at | 336.9 | P | 322.7 | P | 263.3 | P | 347.1 | P | XLG3 |
| 246313_at | 188 | P | 224.7 | P | 167.5 | P | 208.5 | P | --- |

| | | | | | | | | | |
|---|---|---|---|---|---|---|---|---|---|
| 246314_at | 363 | P | 361.2 | P | 361.5 | P | 345.4 | P | AREB3 |
| 246315_at | 87.8 | P | 97.8 | P | 141.8 | P | 101.5 | P | --- |
| 246316_at | 62.2 | A | 14 | A | 49 | A | 13.3 | A | --- |
| 246317_at | 259.1 | P | 268.8 | P | 365.4 | P | 244.6 | P | --- |
| 246318_at | 262.9 | P | 363.5 | P | 267.9 | P | 313.3 | P | --- |
| 246319_at | 1980.6 | P | 2155.2 | P | 2001.6 | P | 2394.3 | P | --- |
| 246320_at | 872.5 | P | 1081.9 | P | 914.2 | P | 1114.8 | P | --- |
| 246321_at | 34.1 | A | 50.3 | M | 71.6 | P | 46.4 | P | --- |
| 246322_at | 277.8 | P | 316.3 | P | 311.7 | P | 339.5 | P | --- |
| 246323_at | 112 | A | 127.7 | A | 146.3 | P | 120.7 | P | --- |
| 246324_at | 491.8 | P | 506 | P | 566.3 | P | 552.8 | P | SR45 |
| 246325_at | 171.5 | P | 159.8 | P | 175.4 | P | 163.6 | P | ABA3 |
| 246326_at | 69.2 | P | 82 | P | 95.8 | P | 89.6 | P | REV7 |
| 246327_at | 248.1 | P | 274.8 | P | 160.5 | P | 249.4 | P | --- |
| 246328_at | 455.5 | P | 405.2 | P | 470.4 | P | 389.6 | P | --- |
| 246329_at | 219.9 | P | 305.7 | P | 333.9 | P | 388.9 | P | --- |
| 246330_at | 373.2 | P | 546.4 | P | 376.8 | P | 553.1 | P | AAO2 |
| 246331_at | 113.2 | P | 97.7 | P | 98.2 | A | 97 | M | --- |
| 246332_at | 32.4 | A | 33.6 | A | 28.7 | A | 20.5 | A | --- |
| 246333_at | 4.8 | A | 11.5 | A | 5.1 | A | 7.4 | A | --- |
| 246334_at | 218.5 | P | 233.1 | P | 275.2 | P | 235.2 | P | --- |
| 246335_at | 3034.3 | P | 1798.2 | P | 2303.5 | P | 1626.9 | P | ACD1 |
| 246336_at | 12.4 | A | 19.2 | A | 4.1 | A | 6.2 | A | ATCHX4 |
| 246337_at | 2 | A | 3.9 | A | 10.9 | A | 17.2 | A | CHX12 |
| 246338_s_at | 5.2 | A | 1.3 | A | 2.5 | A | 2.3 | A | ATCHX10 |
| 246339_at | 3696.1 | P | 4688.5 | P | 3698.7 | P | 4452.5 | P | RPL9 |
| 246340_s_at | 3383.3 | P | 741.7 | P | 1416.6 | P | 989.8 | P | --- |
| 246341_x_at | 1.9 | A | 1.2 | A | 2.1 | A | 5.7 | A | --- |
| 246342_at | 8.1 | A | 2.8 | A | 45 | A | 4.8 | A | FAR6 |
| 246343_at | 552.4 | P | 559.4 | P | 455.6 | P | 560 | P | --- |
| 246344_at | 13.2 | A | 18.9 | A | 31.3 | A | 32.6 | P | --- |
| 246345_at | 289.4 | P | 322.3 | P | 239.6 | P | 246 | P | --- |
| 246346_at | 86.2 | P | 76.6 | P | 138.8 | P | 122 | P | --- |
| 246347_at | 191.2 | P | 185.6 | P | 183.3 | P | 206.4 | P | --- |
| 246348_at | 602.2 | P | 629.3 | P | 586.5 | P | 555.8 | P | --- |
| 246349_at | 3.1 | A | 2 | A | 2 | A | 6.1 | A | --- |
| 246350_at | 173.5 | P | 115.8 | P | 202 | P | 123.5 | P | --- |
| 246351_at | 493.9 | P | 545.3 | P | 634.5 | P | 549.4 | P | --- |
| 246352_at | 4.6 | A | 1.1 | A | 2.5 | A | 1.7 | A | --- |
| 246353_s_at | 12.5 | A | 18 | A | 2.3 | A | 6.8 | A | --- |
| 246354_at | 19 | A | 26.7 | A | 29.9 | A | 3.2 | A | --- |
| 246355_at | 33.5 | A | 43.1 | A | 43.3 | A | 23.7 | A | --- |
| 246356_x_at | 6.9 | A | 15.2 | A | 4.3 | A | 19.8 | A | --- |
| 246357_x_at | 1.9 | A | 11.8 | A | 2.1 | A | 1.1 | A | --- |
| 246358_x_at | 1.2 | A | 12.5 | A | 2.2 | A | 1.7 | A | --- |
| 246359_x_at | 34.7 | M | 4.1 | A | 21 | A | 21.1 | A | --- |
| 246360_at | 12.3 | A | 1.4 | A | 1.4 | A | 0.7 | A | --- |
| 246361_s_at | 16.6 | A | 10.9 | A | 8 | A | 7.2 | A | --- |
| 246362_at | 3.7 | A | 0.5 | A | 20.3 | A | 12.1 | A | --- |
| 246363_at | 15.3 | A | 3.2 | A | 34.5 | A | 19.4 | A | --- |
| 246364_at | 25.6 | P | 8.5 | A | 11.6 | A | 17.1 | A | --- |
| 246365_at | 24.9 | A | 3 | A | 1.6 | A | 3.4 | A | --- |
| 246366_at | 22.5 | A | 45.5 | A | 16.5 | A | 35.5 | P | --- |
| 246367_at | 2.8 | A | 14.4 | A | 12.6 | A | 3.6 | A | --- |
| 246368_at | 49.2 | P | 95.2 | P | 53.8 | A | 55.3 | P | --- |
| 246369_at | 42 | P | 31.5 | P | 55.9 | P | 51.5 | P | --- |
| 246370_at | 39.9 | P | 29.1 | A | 38.3 | A | 16.9 | A | --- |
| 246371_at | 486.3 | P | 659.5 | P | 587.1 | P | 682.7 | P | --- |
| 246372_at | 13.2 | A | 11.3 | A | 22.8 | A | 1.2 | A | --- |
| 246373_at | 1.6 | A | 26.1 | A | 1.5 | A | 4.5 | A | --- |
| 246374_at | 22.8 | A | 17.8 | A | 28.7 | A | 6.2 | A | --- |
| 246375_at | 37.2 | P | 15.3 | A | 3.7 | A | 20.4 | P | --- |
| 246376_at | 805.8 | P | 630.9 | P | 706.3 | P | 532.9 | P | IAA18 |
| 246377_at | 13.8 | A | 49.5 | A | 99.4 | A | 37.8 | A | --- |
| 246378_at | 399.6 | P | 596.6 | P | 377.7 | P | 592.1 | P | --- |
| 246379_s_at | 4337.8 | P | 3308 | P | 4154.7 | P | 3528.2 | P | --- |
| 246380_at | 19 | A | 19.6 | A | 17.7 | A | 22.2 | A | CYP96A15 |
| 246381_at | 1.4 | A | 3.1 | A | 7.8 | A | 12.2 | A | --- |
| 246382_at | 25.9 | A | 7.3 | A | 24.6 | A | 3.4 | A | --- |
| 246383_at | 96.2 | P | 73.2 | P | 107.3 | P | 105.3 | P | --- |
| 246384_at | 1318.8 | P | 1134.4 | P | 1374.7 | P | 1148.4 | P | --- |
| 246385_at | 34.5 | A | 47.7 | A | 38.9 | A | 41.1 | P | CYCA1;2 |
| 246386_at | 76.5 | P | 87.6 | P | 86.8 | P | 95.7 | P | BGAL16 |
| 246387_at | 50.1 | A | 42 | A | 42.4 | A | 19.6 | A | --- |
| 246388_at | 26.9 | A | 41.3 | A | 41.4 | A | 15.9 | A | --- |
| 246389_at | 83.2 | A | 79 | A | 87.3 | M | 34.2 | A | AAP3 |
| 246390_at | 35 | A | 57.6 | A | 87.4 | A | 59.1 | A | --- |
| 246391_at | 817.2 | P | 1011.1 | P | 764.2 | P | 1121.7 | P | --- |
| 246392_at | 11.2 | A | 20.9 | P | 22.3 | A | 5.8 | A | --- |
| 246393_at | 189.9 | P | 138.2 | P | 270.9 | P | 179.8 | P | --- |
| 246394_at | 78.4 | P | 39.7 | P | 50.3 | P | 66.7 | P | --- |
| 246395_at | 32.6 | P | 83.8 | P | 67 | P | 106.7 | P | --- |
| 246396_at | 908.8 | P | 803.3 | P | 765.4 | P | 791.2 | P | BCA6 |
| 246397_at | 2.4 | A | 1.7 | A | 8.6 | A | 1.8 | A | AtRLP9 |
| 246398_at | 319.2 | P | 391 | P | 336.3 | P | 335 | P | --- |
| 246399_at | 359.9 | P | 535.3 | P | 452 | P | 516.9 | P | --- |
| 246400_at | 413.5 | P | 447.1 | P | 508.3 | P | 437.9 | P | --- |
| 246401_at | 47.4 | P | 67 | P | 33.5 | P | 42.5 | P | AtMYB50 |
| 246402_at | 20.9 | A | 11.9 | A | 24.2 | A | 5 | A | --- |
| 246403_at | 571.4 | P | 1331.7 | P | 268.8 | P | 1041.1 | P | --- |
| 246404_at | 282.5 | P | 333 | P | 321.1 | P | 320.3 | P | --- |
| 246405_at | 182.7 | P | 477 | P | 96.8 | P | 196.3 | P | --- |
| 246406_at | 8.1 | A | 19.4 | A | 3.6 | A | 4.9 | A | --- |
| 246407_at | 18 | A | 6.8 | A | 54.6 | A | 19.7 | A | --- |
| 246408_at | 1236.3 | P | 1386.4 | P | 1291.3 | P | 1373.8 | P | --- |
| 246409_at | 94.2 | P | 127.8 | P | 119.6 | P | 140.9 | P | --- |
| 246410_at | 127.2 | P | 130.2 | P | 201.7 | P | 172.3 | P | --- |
| 246411_at | 600.4 | P | 552.9 | P | 679.1 | P | 663.9 | P | --- |
| 246412_at | 520.8 | P | 425.1 | P | 458.1 | P | 512.7 | P | --- |
| 246413_at | 64.3 | P | 64.1 | P | 70.9 | P | 45 | P | --- |
| 246414_at | 8.3 | A | 0.6 | A | 2.8 | A | 1.3 | A | --- |
| 246415_at | 142.7 | P | 229.6 | P | 162.2 | P | 249.3 | P | --- |
| 246416_at | 11.4 | A | 2.1 | A | 2.3 | A | 8.4 | A | --- |
| 246417_at | 949.8 | P | 732 | P | 699.3 | P | 821.6 | P | --- |
| 246418_at | 37.6 | A | 70.9 | A | 55.8 | A | 82.6 | A | --- |
| 246419_at | 14.6 | A | 5.5 | A | 20.2 | A | 23.1 | A | UGT78D3 |
| 246420_at | 221.7 | P | 206.9 | P | 183.3 | P | 243.5 | P | --- |
| 246421_at | 947.5 | P | 1110.2 | P | 999 | P | 922.3 | P | --- |
| 246422_at | 300.7 | P | 375.1 | P | 342.6 | P | 340.8 | P | ATARFB1B |

| | | | | | | | | | |
|---|---|---|---|---|---|---|---|---|---|
| 246423_at | 161.6 | P | 134.3 | P | 172.7 | P | 130.2 | P | --- |
| 246424_at | 860.9 | P | 977.3 | P | 830.1 | P | 989 | P | XPO1A |
| 246425_at | 90.2 | P | 103 | P | 97.6 | P | 139 | P | IRX3 |
| 246426_at | 1.5 | A | 3.7 | A | 4.3 | A | 3.5 | A | BBM |
| 246427_at | 335 | P | 367.7 | P | 383 | P | 341.5 | P | ER-ANT1 |
| 246428_at | 93 | P | 112.9 | P | 124.9 | P | 167.8 | P | --- |
| 246429_at | 150.3 | P | 200.8 | P | 191.8 | P | 225.9 | P | --- |
| 246430_at | 4.1 | A | 10.5 | A | 38 | A | 9.2 | A | --- |
| 246431_at | 0.2 | A | 1.5 | A | 1.5 | A | 1.7 | A | APC1 |
| 246432_at | 9 | A | 2 | A | 13.8 | A | 22.8 | A | RGL3 |
| 246433_at | 164.9 | P | 182.9 | P | 204.6 | P | 194.2 | P | --- |
| 246434_at | 682.3 | P | 1142.6 | P | 602.9 | P | 1135.9 | P | RCP1 |
| 246435_at | 70.4 | P | 91.2 | P | 106.9 | P | 73.7 | P | --- |
| 246436_at | 463.8 | P | 351.8 | P | 465.9 | P | 348.2 | P | --- |
| 246437_at | 5.4 | A | 3 | A | 2 | A | 6.1 | A | --- |
| 246438_at | 30.3 | A | 7.1 | A | 37.7 | A | 41.9 | A | --- |
| 246439_at | 299.2 | P | 294.3 | P | 384.2 | P | 295.6 | P | --- |
| 246440_at | 772.5 | P | 455 | P | 619.1 | P | 474.1 | P | --- |
| 246441_at | 2823.6 | P | 2543.8 | P | 2695.9 | P | 3041 | P | --- |
| 246442_at | 391.5 | P | 314.9 | P | 376.4 | P | 269.9 | P | --- |
| 246443_at | 215.6 | P | 294.7 | P | 297.7 | P | 311.4 | P | --- |
| 246444_at | 187.2 | P | 151.3 | P | 226.6 | P | 187 | P | --- |
| 246445_at | 727.2 | P | 1029.7 | P | 875.3 | P | 982.6 | P | --- |
| 246446_at | 259.4 | P | 208.9 | P | 260.5 | P | 212.7 | P | --- |
| 246447_at | 378.7 | P | 442.7 | P | 438 | P | 389.1 | P | DOT2 |
| 246448_at | 362.7 | P | 411.8 | P | 458.5 | P | 389.8 | P | --- |
| 246449_at | 405.6 | P | 394 | P | 409.8 | P | 320.5 | P | --- |
| 246450_at | 178.3 | P | 203.1 | P | 146.4 | P | 180.4 | P | HSF3 |
| 246451_at | 1064.2 | P | 1090.2 | P | 1304.3 | P | 1090.5 | P | --- |
| 246452_at | 1339.3 | P | 1353.1 | P | 1537.8 | P | 1242.4 | P | BPA1 |
| 246453_at | 801.2 | P | 756.6 | P | 817 | P | 662.7 | P | SYP21 |
| 246454_at | 1824 | P | 2996.6 | P | 1915.1 | P | 2345.8 | P | DHAR3 |
| 246455_at | 31.1 | A | 4.3 | A | 6.1 | A | 22.5 | A | --- |
| 246456_at | 10.9 | A | 7.8 | A | 40 | A | 6.5 | A | ATTERT |
| 246457_at | 396.8 | P | 230 | P | 459.1 | P | 233.6 | P | TOZ |
| 246458_at | 73 | P | 47.7 | P | 82.2 | P | 45.7 | P | --- |
| 246459_at | 41.5 | A | 37.1 | P | 37.6 | A | 12.4 | A | --- |
| 246460_at | 538.7 | P | 453.9 | P | 444.2 | P | 405.6 | P | CSLD2 |
| 246461_at | 136 | P | 95 | P | 150.3 | P | 109.2 | P | --- |
| 246462_at | 116.6 | P | 232.2 | P | 120.1 | P | 191.2 | P | --- |
| 246463_at | 2672.4 | P | 2098.2 | P | 1878.8 | P | 1946.1 | P | AT-AER |
| 246464_at | 790.6 | P | 684.9 | P | 452.7 | P | 951.1 | P | --- |
| 246465_at | 180.2 | P | 211.7 | P | 203.1 | P | 178.6 | P | --- |
| 246466_at | 605.7 | P | 721.9 | P | 835.3 | P | 878.1 | P | --- |
| 246467_at | 41.4 | A | 34.7 | A | 7.7 | A | 49.8 | A | --- |
| 246468_at | 1698.4 | P | 867.6 | P | 2266.1 | P | 1532.2 | P | UGT78D2 |
| 246469_at | 306.4 | P | 344.2 | P | 207.4 | P | 327.4 | P | --- |
| 246470_at | 3.5 | A | 1.4 | A | 11.8 | A | 17 | A | --- |
| 246471_s_at | 2.6 | A | 18 | A | 22.7 | A | 3.2 | A | --- |
| 246472_at | 1.8 | A | 9.2 | A | 35.5 | A | 12.9 | A | --- |
| 246473_at | 13.7 | A | 32.8 | A | 32 | A | 20.6 | A | --- |
| 246474_at | 18.2 | A | 18.3 | P | 22.5 | A | 5.2 | A | --- |
| 246475_at | 33.6 | M | 58.3 | P | 79.7 | P | 64.2 | P | --- |
| 246476_at | 262.4 | P | 297.1 | P | 263.3 | P | 290.2 | P | --- |
| 246477_at | 1.2 | A | 16.2 | A | 15.4 | A | 10.8 | A | AtMYB9 |
| 246478_at | 303.2 | P | 295.1 | P | 363.2 | P | 290.4 | P | --- |
| 246479_at | 1866.3 | P | 1031.5 | P | 1757 | P | 1080.9 | P | --- |
| 246480_at | 1070.1 | P | 1115.5 | P | 1108.6 | P | 1137.2 | P | --- |
| 246481_s_at | 3151.9 | P | 5566.2 | P | 2632.4 | P | 7674 | P | KIN1 |
| 246482_at | 484.6 | P | 601.6 | P | 538.8 | P | 620.1 | P | PAM1 |
| 246483_at | 312.9 | P | 323.2 | P | 361.5 | P | 390.2 | P | NIK1 |
| 246484_at | 457.7 | P | 471.5 | P | 389.4 | P | 433.5 | P | --- |
| 246485_at | 8.2 | A | 79.3 | M | 49.9 | A | 65.1 | M | AtCXE17 |
| 246486_at | 776.1 | P | 836.1 | P | 819 | P | 867.5 | P | --- |
| 246487_at | 601.9 | P | 824.1 | P | 512.8 | P | 685 | P | --- |
| 246488_at | 4842 | P | 4529.9 | P | 4519.3 | P | 4340.4 | P | --- |
| 246489_at | 1502.4 | P | 2128 | P | 1594 | P | 2123.8 | P | GRF5 |
| 246490_at | 363.5 | P | 711.6 | P | 496.1 | P | 665.7 | P | CPuORF10 |
| 246491_at | 38.2 | P | 38.7 | P | 42.2 | A | 40.5 | P | --- |
| 246492_at | 283.8 | P | 367.2 | P | 372.2 | P | 396.2 | P | --- |
| 246493_at | 169.2 | P | 188.8 | P | 150.8 | P | 170 | P | CRS1 |
| 246494_at | 239.3 | P | 161 | P | 331.8 | P | 185.1 | P | CSLA11 |
| 246495_at | 307.3 | P | 197.9 | P | 256.9 | P | 187.6 | P | --- |
| 246496_at | 336 | P | 350.7 | P | 291 | P | 359.9 | P | --- |
| 246497_at | 632.4 | P | 644.4 | P | 585.4 | P | 568.4 | P | --- |
| 246498_at | 35.7 | A | 58.6 | P | 40.8 | A | 54.8 | P | --- |
| 246499_at | 528.1 | P | 460.9 | P | 441.5 | P | 563 | P | --- |
| 246500_at | 311.5 | P | 429.2 | P | 535 | P | 383.4 | P | SYN4 |
| 246501_at | 238.2 | P | 272.1 | P | 305 | P | 268.8 | P | --- |
| 246502_at | 177.4 | P | 153.3 | P | 186.1 | P | 144.4 | P | --- |
| 246503_at | 5663.6 | P | 5277.3 | P | 6942.9 | P | 6277.3 | P | --- |
| 246504_at | 39.1 | P | 33.9 | A | 55.7 | P | 30.9 | A | --- |
| 246505_at | 602 | P | 1103.6 | P | 1014.2 | P | 1276.6 | P | --- |
| 246506_at | 891.3 | P | 941.8 | P | 972.2 | P | 1006.5 | P | --- |
| 246507_at | 802.5 | P | 569.6 | P | 687 | P | 570.8 | P | --- |
| 246508_at | 2547.5 | P | 2582.1 | P | 2547.6 | P | 2254.6 | P | PGLCT |
| 246509_at | 750.3 | P | 922.6 | P | 842.8 | P | 857.3 | P | EMB2247 |
| 246510_at | 619.5 | P | 1585.7 | P | 738.5 | P | 1480.9 | P | DND1 |
| 246511_at | 1312.7 | P | 1142.6 | P | 1245.4 | P | 1161.5 | P | --- |
| 246512_at | 69.9 | P | 95.9 | P | 82.8 | P | 110.3 | P | IRX6 |
| 246513_at | 259.5 | P | 265.6 | P | 266.1 | P | 259.8 | P | --- |
| 246514_at | 184.9 | P | 119.3 | M | 141.8 | A | 149.6 | P | --- |
| 246515_at | 135.7 | P | 141.4 | P | 109.8 | P | 162.9 | P | --- |
| 246516_at | 451.6 | P | 518.8 | P | 336.6 | P | 527.9 | P | --- |
| 246517_at | 131.4 | P | 121.2 | P | 168.5 | P | 159.8 | P | --- |
| 246518_at | 255.5 | P | 238.7 | P | 202.3 | P | 266.6 | P | AtGNA1 |
| 246519_at | 13.8 | A | 36.5 | A | 3.4 | A | 3.2 | A | --- |
| 246520_at | 82.5 | P | 115.7 | P | 106.6 | P | 161.7 | P | --- |
| 246521_at | 1148.3 | P | 956.1 | P | 1421.4 | P | 951.7 | P | --- |
| 246522_at | 21.5 | A | 41.5 | A | 6.9 | A | 21.5 | A | AtbZIP3 |
| 246523_at | 811.8 | P | 646.4 | P | 901.1 | P | 671.3 | P | COL1 |
| 246524_at | 348.8 | P | 317.7 | P | 298 | P | 293.6 | P | ATPCME |
| 246525_at | 76 | P | 78.9 | P | 142.3 | P | 90.8 | P | CO |
| 246526_at | 20.3 | A | 11.9 | A | 4 | A | 17 | A | GLIP7 |
| 246527_at | 1408.1 | P | 1004.9 | P | 1386.7 | P | 1007.4 | P | --- |
| 246528_at | 423.9 | P | 518.2 | P | 471.1 | P | 491.6 | P | --- |
| 246529_at | 354.2 | P | 349.7 | P | 246.8 | P | 273 | P | --- |
| 246530_at | 10.7 | A | 19.2 | A | 35.4 | A | 6.9 | A | --- |
| 246531_at | 5.9 | A | 7.6 | A | 7.3 | A | 23.5 | A | 41518 |

| | | | | | | | | | |
|---|---|---|---|---|---|---|---|---|---|
| 246532_at | 481.6 | P | 390.7 | P | 479.8 | P | 310.9 | P | --- |
| 246533_at | 502.6 | P | 574.6 | P | 399.3 | P | 542.6 | P | --- |
| 246534_at | 0.9 | A | 4.2 | A | 4.9 | A | 6.1 | A | --- |
| 246535_at | 29.4 | A | 46.7 | A | 7.3 | A | 48.1 | A | --- |
| 246536_at | 134.4 | P | 137 | P | 142.8 | P | 102.2 | P | --- |
| 246537_at | 53 | M | 57.1 | A | 89.1 | A | 51 | P | --- |
| 246538_at | 594.5 | P | 619.1 | P | 757.3 | P | 661.9 | P | --- |
| 246539_at | 661.8 | P | 647 | P | 577.9 | P | 650.4 | P | MUB2 |
| 246540_at | 47.8 | P | 78.4 | M | 69.8 | A | 58.9 | P | SP1L4 |
| 246541_at | 560.6 | P | 624.7 | P | 568.2 | P | 668.3 | P | --- |
| 246542_at | 332.9 | P | 294.1 | P | 328.3 | P | 319.1 | P | SNL2 |
| 246543_at | 33.7 | A | 5 | A | 56.5 | A | 27.6 | A | --- |
| 246544_at | 17.2 | A | 10.2 | A | 16.6 | A | 20.1 | A | PIN8 |
| 246545_at | 2.7 | A | 12.7 | A | 4.2 | A | 1.4 | A | --- |
| 246546_at | 3743.3 | P | 2959 | P | 3415.5 | P | 3270 | P | VDAC3 |
| 246547_at | 1199.8 | P | 1346.8 | P | 1267.9 | P | 1400.3 | P | --- |
| 246548_at | 3023 | P | 4666.8 | P | 3009.6 | P | 4484.4 | P | --- |
| 246549_at | 369.6 | P | 677.2 | P | 375.1 | P | 686.8 | P | --- |
| 246550_at | 1997.2 | P | 4786.6 | P | 1537.1 | P | 4574.8 | P | --- |
| 246551_at | 21.1 | A | 34.5 | A | 13.2 | A | 38.7 | A | --- |
| 246552_at | 10 | A | 2.5 | A | 28.1 | A | 6.7 | A | --- |
| 246553_at | 129.9 | P | 222.3 | P | 195.1 | P | 167.3 | P | EDL1 |
| 246554_at | 1989.4 | P | 2448.2 | P | 2328.2 | P | 2991.3 | P | CLPB3 |
| 246555_at | 189.2 | P | 316.7 | P | 164 | P | 274.4 | P | GAUT14 |
| 246556_at | 5.8 | A | 21.9 | M | 23.3 | A | 6.3 | A | --- |
| 246557_at | 58.9 | P | 99.7 | P | 91.8 | P | 83.6 | P | --- |
| 246558_at | 238.2 | P | 228.4 | P | 293.1 | P | 249.4 | P | EMB2773 |
| 246559_at | 603.3 | P | 476.3 | P | 687.8 | P | 512.9 | P | --- |
| 246560_at | 19.8 | A | 3.6 | A | 46.7 | A | 27.9 | A | --- |
| 246561_at | 176.3 | P | 210.2 | P | 223.2 | P | 217.1 | P | --- |
| 246562_at | 70.5 | P | 180.8 | P | 79.6 | P | 139.4 | P | LNG1 |
| 246563_at | 23 | A | 16.4 | A | 17.2 | A | 12.7 | A | --- |
| 246564_at | 1092.3 | P | 603.1 | P | 781.6 | P | 846.1 | P | --- |
| 246565_at | 236.8 | P | 380.9 | P | 386.7 | P | 398.4 | P | BCCP2 |
| 246566_at | 290.5 | P | 307.9 | P | 297.5 | P | 303.7 | P | --- |
| 246567_at | 371.4 | P | 406.5 | P | 414.5 | P | 386 | P | GMII |
| 246568_at | 69.4 | P | 114.5 | P | 84.4 | P | 70.4 | P | DEL2 |
| 246569_at | 11 | A | 1.2 | A | 16.6 | A | 1.2 | A | --- |
| 246570_at | 8.9 | A | 9.3 | A | 5.3 | A | 5.5 | A | --- |
| 246571_at | 1.4 | A | 1.4 | A | 3.7 | A | 14.6 | A | --- |
| 246572_at | 64.2 | A | 60.9 | A | 111.8 | A | 49.9 | A | --- |
| 246573_at | 255.4 | P | 325.7 | P | 260 | P | 446.2 | P | --- |
| 246574_at | 8.2 | A | 7.8 | A | 4.8 | A | 4 | A | --- |
| 246575_at | 714.9 | P | 635.4 | P | 831.7 | P | 664.7 | P | --- |
| 246576_at | 448.1 | P | 481 | P | 475.9 | P | 463.7 | P | ROPGEF14 |
| 246577_at | 19.5 | A | 15.9 | A | 21.3 | A | 2.8 | A | AGL92 |
| 246578_at | 0.9 | A | 1.4 | A | 2.4 | A | 12.6 | A | AGL86 |
| 246579_at | 3 | A | 1.8 | A | 9.1 | A | 3.1 | A | --- |
| 246580_at | 116 | P | 256.9 | P | 284.6 | P | 285.7 | P | --- |
| 246581_at | 63.8 | P | 69.5 | A | 94.2 | A | 65.9 | A | --- |
| 246582_at | 76.2 | P | 55.5 | P | 90.7 | P | 58.5 | P | --- |
| 246583_at | 357.4 | P | 429.4 | P | 409.1 | P | 440.3 | P | --- |
| 246584_at | 288 | P | 254.3 | P | 263.8 | P | 273 | P | --- |
| 246585_at | 12.2 | A | 23.9 | A | 3.4 | A | 12.2 | A | ATMYB66 |
| 246586_at | 32.6 | P | 60.5 | P | 63.8 | P | 56.3 | P | --- |
| 246587_at | 36.1 | A | 8.6 | A | 3.9 | A | 28.4 | A | --- |
| 246588_at | 16.8 | A | 14.4 | A | 21.2 | A | 19.6 | A | --- |
| 246589_at | 39.4 | A | 34.7 | A | 24.2 | A | 2.7 | A | --- |
| 246590_at | 30.5 | A | 35.3 | P | 42.5 | A | 22.8 | A | CNGC18 |
| 246591_at | 282.2 | P | 440.8 | P | 318.7 | P | 382.3 | P | --- |
| 246592_at | 4.2 | A | 12.7 | A | 8.2 | A | 3.1 | A | --- |
| 246593_at | 241.5 | P | 361.4 | P | 326.4 | P | 315.1 | P | --- |
| 246594_at | 2552.9 | P | 2466.1 | P | 2740.4 | P | 2370.4 | P | P5CR |
| 246595_at | 14235.2 | P | 8163.1 | P | 13609.9 | P | 9029.6 | P | FDH |
| 246596_at | 9314.4 | P | 10904.3 | P | 10223.1 | P | 9272.6 | P | CA2 |
| 246597_at | 287.8 | P | 280.7 | P | 275.7 | P | 211.6 | P | AO |
| 246598_at | 344.6 | P | 278 | P | 340.3 | P | 288.8 | P | --- |
| 246599_at | 8 | A | 18.5 | A | 5.2 | A | 1.9 | A | --- |
| 246600_at | 173.8 | P | 205.7 | P | 167.6 | P | 164.9 | P | SAG101 |
| 246601_at | 542.8 | P | 867.4 | P | 877.1 | P | 821.6 | P | --- |
| 246602_at | 171.7 | P | 297.1 | P | 340.9 | P | 269.8 | P | --- |
| 246603_at | 575.1 | P | 694 | P | 513.2 | P | 809.2 | P | --- |
| 246604_at | 3.3 | A | 4.6 | A | 20.4 | A | 27.5 | A | --- |
| 246605_at | 16.2 | A | 23.2 | A | 6.9 | A | 1.2 | A | --- |
| 246606_at | 3.2 | A | 34.9 | A | 47.3 | P | 31.7 | P | --- |
| 246607_at | 240.2 | P | 185.9 | P | 144.8 | P | 183.1 | P | --- |
| 246608_at | 5.3 | A | 6.4 | A | 18.9 | A | 2.4 | A | --- |
| 246609_at | 12.9 | A | 7.7 | A | 3.1 | A | 1.3 | A | --- |
| 246610_at | 67.3 | P | 86.5 | P | 114.5 | P | 57 | P | --- |
| 246611_at | 770.3 | P | 781.2 | P | 831.7 | P | 854.8 | P | MBD02 |
| 246612_at | 266.5 | P | 360.6 | P | 281 | P | 282.3 | P | --- |
| 246613_at | 4089.8 | P | 3334.7 | P | 4240.3 | P | 3230.5 | P | CAC2 |
| 246614_at | 93.2 | P | 54.6 | P | 81.3 | P | 76.1 | P | SOS2 |
| 246615_at | 14.3 | A | 20.1 | A | 30.8 | A | 24.3 | A | --- |
| 246616_at | 3.7 | A | 3.5 | A | 6.5 | A | 2.4 | A | --- |
| 246617_at | 30.6 | A | 25.5 | A | 72.5 | P | 60.3 | P | --- |
| 246618_at | 24.2 | A | 24.8 | A | 74.6 | P | 2.8 | A | --- |
| 246619_at | 1017.9 | P | 1344.2 | P | 968.7 | P | 1222.8 | P | --- |
| 246620_at | 633.1 | P | 331.8 | P | 434.5 | P | 364.2 | P | CYP81D1 |
| 246621_at | 3099.3 | P | 3324.3 | P | 2630.4 | P | 3345.1 | P | --- |
| 246622_at | 270.7 | P | 280.1 | P | 225 | P | 275 | P | --- |
| 246623_at | 6483.4 | P | 5379.4 | P | 6040 | P | 5802.9 | P | ATNUC-L1 |
| 246624_at | 21.1 | A | 40.3 | A | 38 | A | 21 | A | YUC10 |
| 246625_at | 70.2 | A | 82 | A | 65.6 | A | 106.5 | A | --- |
| 246626_at | 48.5 | A | 46.1 | A | 76.8 | A | 48.4 | A | --- |
| 246627_s_at | 2980.5 | P | 2292.8 | P | 2897.7 | P | 2566.5 | P | --- |
| 246628_at | 1338 | P | 1226.4 | P | 1317.5 | P | 1313.8 | P | --- |
| 246629_at | 1.8 | A | 3.2 | A | 2.5 | A | 1.8 | A | --- |
| 246630_at | 301 | P | 427.9 | P | 336.3 | P | 283.9 | P | --- |
| 246631_at | 837.8 | P | 1068.4 | P | 782.2 | P | 920.2 | P | --- |
| 246632_at | 152.5 | P | 84.8 | P | 108.9 | P | 131.7 | P | --- |
| 246633_at | 46.8 | P | 110.7 | P | 48 | A | 88 | P | --- |
| 246634_at | 1356.9 | P | 1188.8 | P | 1010.4 | P | 1105 | P | --- |
| 246635_at | 17.4 | A | 28.6 | A | 37.2 | A | 35.4 | A | --- |
| 246636_at | 5431.2 | P | 4779.4 | P | 4949.2 | P | 5587.8 | P | PAP26 |
| 246637_at | 1.6 | A | 2 | A | 2.4 | A | 1.7 | A | --- |
| 246638_at | 20.7 | A | 16.9 | A | 5.2 | A | 41.3 | A | --- |
| 246639_x_at | 2 | A | 0.9 | A | 5.8 | A | 8.7 | A | --- |
| 246640_x_at | 100.1 | P | 118.2 | P | 143.1 | A | 90.5 | A | --- |

| | | | | | | | | | |
|---|---|---|---|---|---|---|---|---|---|
| 246641_s_at | 2 | A | 1.7 | A | 2.3 | A | 2.1 | A | --- |
| 246642_s_at | 5.1 | A | 1.9 | A | 2.6 | A | 1.1 | A | --- |
| 246643_s_at | 0.6 | A | 0.3 | A | 1.8 | A | 0.5 | A | --- |
| 246644_at | 720.2 | P | 1160.2 | P | 806.4 | P | 894.7 | P | --- |
| 246645_at | 405 | P | 481.8 | P | 363.1 | P | 408.3 | P | --- |
| 246646_at | 1.1 | A | 1.3 | A | 2 | A | 1.3 | A | --- |
| 246647_at | 19 | A | 13.4 | A | 16.7 | A | 20.2 | A | --- |
| 246648_at | 1.1 | A | 2.6 | A | 2.8 | A | 1.4 | A | --- |
| 246649_at | 8.5 | A | 24.8 | A | 27.1 | A | 5.8 | A | --- |
| 246650_at | 912.2 | P | 1157.4 | P | 894.9 | P | 1182.2 | P | --- |
| 246651_at | 1429.3 | P | 1558.8 | P | 1335.1 | P | 1536.7 | P | --- |
| 246652_at | 1.3 | A | 1 | A | 3.5 | A | 18.2 | M | --- |
| 246653_at | 381 | P | 444.9 | P | 356.9 | P | 479.7 | P | --- |
| 246654_s_at | 1368.8 | P | 1636 | P | 1566.8 | P | 1804.4 | P | EGY1 |
| 246655_at | 11.9 | A | 16.7 | A | 24.3 | A | 10.5 | A | --- |
| 246656_at | 4.4 | A | 10.8 | A | 39 | A | 24.7 | A | --- |
| 246657_at | 20.9 | A | 1.8 | A | 2.6 | A | 9.6 | A | --- |
| 246658_at | 0.6 | A | 11.1 | P | 0.9 | A | 2.5 | A | --- |
| 246659_at | 17.3 | A | 2.8 | A | 26.4 | A | 9.2 | A | --- |
| 246660_at | 872.6 | P | 1127 | P | 968.5 | P | 1007.5 | P | --- |
| 246661_at | 0.4 | A | 0.1 | A | 0.6 | A | 7.5 | A | --- |
| 246662_at | 12.1 | A | 15.5 | A | 34.4 | A | 2.6 | A | --- |
| 246663_at | 5.9 | A | 4.2 | A | 16.7 | A | 4.2 | A | --- |
| 246664_at | 3.2 | A | 6.2 | A | 19 | A | 6.4 | A | --- |
| 246665_s_at | 69.6 | A | 51.9 | A | 75 | A | 36.7 | A | --- |
| 246666_at | 39.1 | P | 21.2 | A | 35.1 | P | 37 | A | --- |
| 246667_at | 3.9 | A | 7.1 | A | 9.3 | A | 3.8 | A | --- |
| 246668_at | 1.5 | A | 2.9 | A | 7.1 | A | 2.2 | A | --- |
| 246669_at | 12.3 | A | 17.9 | A | 30.7 | A | 2.9 | A | --- |
| 246670_at | 15.9 | A | 3.8 | A | 7.1 | A | 0.3 | A | --- |
| 246671_at | 9.4 | A | 22.1 | A | 4.2 | A | 25.1 | A | --- |
| 246672_at | 12.1 | A | 22.4 | M | 12.8 | A | 16.3 | A | --- |
| 246673_at | 7938.3 | P | 8713.3 | P | 8403.1 | P | 9065 | P | RPS1 |
| 246674_at | 179.6 | P | 161.5 | P | 166.8 | P | 188.1 | P | --- |
| 246675_at | 23.4 | A | 20.9 | A | 10.7 | A | 29.3 | P | --- |
| 246676_at | 16.9 | A | 1.5 | A | 7.9 | A | 8.5 | A | --- |
| 246677_at | 2.1 | A | 3.7 | A | 17.5 | A | 3.5 | A | --- |
| 246678_at | 17.4 | A | 4.4 | A | 37.6 | A | 25.6 | A | --- |
| 246679_at | 1.4 | A | 1.9 | A | 4.2 | A | 5.8 | A | --- |
| 246680_at | 23.2 | A | 40.7 | A | 65.6 | A | 15.6 | A | --- |
| 246681_at | 278 | P | 434.4 | P | 330.8 | P | 368.7 | P | --- |
| 246682_at | 1657.1 | P | 1249.8 | P | 1245.3 | P | 1052.6 | P | XGD1 |
| 246683_at | 77.9 | P | 66.8 | P | 93.6 | P | 90.2 | P | --- |
| 246684_at | 23.2 | A | 30.3 | A | 45.9 | A | 29.4 | A | CDR1 |
| 246685_at | 16.2 | A | 6.6 | A | 23.2 | A | 8.8 | A | --- |
| 246686_at | 17.4 | A | 37.2 | A | 5 | A | 27.8 | A | --- |
| 246687_at | 329.3 | P | 281.7 | P | 194.5 | P | 293.7 | P | --- |
| 246688_at | 18 | P | 1.8 | A | 15 | A | 9.1 | A | --- |
| 246689_at | 7.2 | A | 5.7 | A | 37.9 | P | 1 | A | --- |
| 246690_at | 4.5 | A | 24.4 | A | 17.1 | A | 8.3 | A | --- |
| 246691_at | 37.3 | A | 29.8 | A | 62.8 | A | 24.8 | A | --- |
| 246692_at | 19.2 | A | 8.6 | A | 19.2 | A | 12.8 | A | --- |
| 246693_at | 1.7 | A | 1.5 | A | 2.9 | A | 2.3 | A | --- |
| 246694_at | 16.1 | A | 8.4 | A | 22.9 | A | 1 | A | --- |
| 246695_at | 6.5 | A | 6.2 | A | 11.7 | A | 7.5 | A | --- |
| 246696_at | 17.9 | A | 22.7 | A | 3.7 | A | 2.2 | A | --- |
| 246697_at | 2.4 | A | 4.5 | A | 5.4 | A | 6.9 | A | --- |
| 246698_at | 8.1 | A | 15.1 | A | 23.9 | A | 6 | A | --- |
| 246699_at | 698.1 | P | 605.5 | P | 608.1 | P | 606.5 | P | --- |
| 246700_at | 38.9 | A | 68.1 | P | 80.9 | P | 86.5 | P | --- |
| 246701_at | 350 | P | 478.8 | P | 327.8 | P | 404.2 | P | CYSD2 |
| 246702_at | 2772.3 | P | 2564.5 | P | 2823.4 | P | 2741.8 | P | --- |
| 246703_at | 39.2 | A | 25.3 | A | 36.2 | A | 29.5 | A | WNK9 |
| 246704_at | 15.4 | A | 23.8 | P | 41.9 | P | 37.9 | M | --- |
| 246705_x_at | 1.1 | A | 2.9 | A | 1.9 | A | 21.3 | A | --- |
| 246706_at | 10.5 | A | 10 | A | 2.1 | A | 10.5 | A | --- |
| 246707_at | 6 | A | 7.5 | A | 8.1 | A | 14 | A | --- |
| 246708_at | 394.9 | P | 357.9 | P | 617.3 | P | 355.8 | P | --- |
| 246709_s_at | 0.7 | A | 0.4 | A | 3 | A | 4.9 | A | --- |
| 246710_at | 5 | A | 9.9 | A | 6.8 | A | 19.1 | A | --- |
| 246711_at | 1.1 | A | 1.7 | A | 1.4 | A | 1.1 | A | --- |
| 246712_at | 11.8 | A | 14.6 | A | 1.4 | A | 16.4 | A | --- |
| 246713_at | 6.5 | A | 17.2 | A | 58.9 | P | 27.5 | A | --- |
| 246714_at | 314.6 | P | 291.3 | P | 240.2 | P | 336.6 | P | --- |
| 246715_at | 37.4 | A | 23.4 | A | 7.3 | A | 40.7 | A | --- |
| 246716_s_at | 26.8 | P | 14.9 | A | 4.4 | A | 36.4 | P | --- |
| 246717_at | 40.6 | A | 34.9 | A | 49.6 | A | 16.8 | A | --- |
| 246718_at | 4.4 | A | 9.2 | A | 9.3 | A | 16.9 | A | --- |
| 246719_at | 1.2 | A | 3.7 | A | 15.3 | A | 0.6 | A | --- |
| 246720_at | 15.8 | A | 0.5 | A | 9.7 | A | 3.9 | A | --- |
| 246721_at | 14.2 | A | 11.3 | A | 40.6 | A | 32.6 | A | --- |
| 246722_at | 1.7 | A | 5.6 | A | 6.5 | A | 6.7 | A | --- |
| 246723_at | 2.1 | A | 3.8 | A | 7.8 | A | 2.9 | A | --- |
| 246724_at | 206.6 | P | 109.3 | P | 190 | P | 163.9 | P | --- |
| 246725_at | 20.9 | A | 34.1 | A | 7 | A | 43.8 | A | --- |
| 246726_at | 2.4 | A | 1.7 | A | 2.7 | A | 3.3 | A | --- |
| 246727_at | 141.9 | P | 70.2 | P | 113.4 | P | 84.8 | P | --- |
| 246728_at | 367.1 | P | 389.9 | P | 310.7 | P | 462.4 | P | --- |
| 246729_at | 3.4 | A | 0.8 | A | 1.1 | A | 9.1 | A | --- |
| 246730_at | 4052 | P | 3576.6 | P | 5442.1 | P | 3919.8 | P | --- |
| 246731_at | 57.1 | P | 38.7 | P | 52.3 | P | 77.3 | P | ACBP5 |
| 246732_at | 169.9 | P | 142.7 | P | 213.2 | P | 165.1 | P | --- |
| 246733_at | 131.5 | P | 50.4 | P | 164.4 | P | 97.3 | P | --- |
| 246734_at | 35.8 | A | 29.9 | P | 39.2 | A | 26.9 | A | RECQSIM |
| 246735_at | 1160.5 | P | 1281.8 | P | 1168.3 | P | 1336.5 | P | HTA7 |
| 246736_at | 913.6 | P | 1066.5 | P | 1219.3 | P | 1026.5 | P | --- |
| 246737_at | 240.4 | P | 273.2 | P | 260.3 | P | 267.7 | P | --- |
| 246738_at | 484.2 | P | 543.6 | P | 633.8 | P | 533.3 | P | EMB2775 |
| 246739_at | 19.8 | P | 23.5 | A | 49.4 | P | 24.2 | A | --- |
| 246740_at | 2 | A | 10 | A | 6.7 | A | 1.1 | A | --- |
| 246741_at | 878.1 | P | 1039.3 | P | 852.6 | P | 1099.6 | P | emb1644 |
| 246742_at | 826.4 | P | 766.1 | P | 752.3 | P | 653.4 | P | TOPP8 |
| 246743_at | 185 | P | 199.8 | P | 132.8 | P | 241.6 | P | --- |
| 246744_at | 5543 | P | 3443.9 | P | 4812.8 | P | 4200.3 | P | --- |
| 246745_at | 4623.1 | P | 3970.5 | P | 4486.5 | P | 4300.3 | P | --- |
| 246746_at | 823 | P | 894.9 | P | 756.5 | P | 995.2 | P | --- |
| 246747_at | 7757.7 | P | 7988.1 | P | 9244.5 | P | 7525.9 | P | --- |
| 246748_at | 367 | P | 476.6 | P | 420.1 | P | 495.1 | P | --- |
| 246749_at | 552.7 | P | 538.3 | P | 414.3 | P | 514.7 | P | --- |

| | | | | | | | | | |
|---|---|---|---|---|---|---|---|---|---|
| 246750_at | 167.6 | P | 180.2 | P | 150.2 | P | 154.4 | P | AT5G27860 |
| 246751_at | 14.2 | A | 1.4 | A | 6.2 | A | 5.4 | A | --- |
| 246752_at | 19.3 | P | 26.9 | P | 50.8 | P | 16.5 | A | --- |
| 246753_at | 3.1 | A | 3.8 | A | 25.4 | A | 10.3 | A | --- |
| 246754_at | 30.2 | A | 27.2 | A | 32.7 | A | 7.8 | A | NF-YC8 |
| 246755_at | 322.9 | P | 231.2 | P | 344 | P | 223.2 | P | --- |
| 246756_at | 677.4 | P | 546.4 | P | 657 | P | 590 | P | --- |
| 246757_at | 8.2 | A | 36.4 | A | 46.7 | A | 39.3 | A | WPP3 |
| 246758_at | 7278.4 | P | 6768.6 | P | 8348 | P | 7583.2 | P | --- |
| 246759_at | 305.9 | P | 387.5 | P | 314.5 | P | 335.5 | P | --- |
| 246760_at | 307.9 | P | 335.3 | P | 306.6 | P | 298.1 | P | --- |
| 246761_at | 3.7 | A | 12.2 | A | 30 | A | 26.8 | A | --- |
| 246762_at | 248.3 | P | 208.4 | P | 288.9 | P | 245.1 | P | CYCH;1 |
| 246763_at | 774.2 | P | 670.3 | P | 536.1 | P | 675.4 | P | NHX1 |
| 246764_s_at | 2.4 | A | 1.9 | A | 0.8 | A | 1.1 | A | --- |
| 246765_at | 209.8 | P | 188.5 | P | 299.5 | P | 161.7 | P | --- |
| 246766_at | 37.1 | A | 35.2 | A | 72.5 | A | 44.1 | A | --- |
| 246767_at | 697.2 | P | 584 | P | 763.9 | P | 676.1 | P | --- |
| 246768_at | 407.1 | P | 430.2 | P | 519.5 | P | 464.8 | P | --- |
| 246769_at | 113.1 | P | 150.1 | P | 102.5 | P | 111.5 | P | --- |
| 246770_at | 264.9 | P | 188.4 | P | 165.9 | P | 153.8 | P | --- |
| 246771_at | 0.7 | A | 0.9 | A | 1 | A | 3.8 | A | --- |
| 246772_at | 280 | P | 445.2 | P | 424.9 | P | 395.4 | P | --- |
| 246773_at | 20.5 | A | 30.9 | A | 21.3 | A | 32.7 | A | --- |
| 246774_at | 7.1 | A | 34.4 | A | 43 | A | 43.2 | A | --- |
| 246775_at | 836.8 | P | 796.7 | P | 925.6 | P | 770.1 | P | MIRO1 |
| 246776_at | 92.9 | P | 143.4 | P | 92.8 | P | 162 | P | --- |
| 246777_at | 149.2 | P | 186.4 | P | 106.5 | P | 154.4 | P | --- |
| 246778_at | 359.3 | P | 375 | P | 371.3 | P | 437.9 | P | MK |
| 246779_at | 711.8 | P | 380.1 | P | 551.8 | P | 366.4 | P | PNC2 |
| 246780_at | 2629.7 | P | 2614.2 | P | 2979.6 | P | 2401 | P | --- |
| 246781_at | 530.7 | P | 709.1 | P | 549.9 | P | 509.1 | P | SFP1 |
| 246782_at | 83.1 | P | 121.1 | P | 97.1 | P | 84.1 | P | GID1C |
| 246783_at | 96.3 | P | 168.6 | P | 110.6 | P | 100.2 | P | SFP2 |
| 246784_at | 491.7 | P | 639.1 | P | 516.5 | P | 671.6 | P | --- |
| 246785_at | 4565.4 | P | 2312.1 | P | 3916.1 | P | 2678.6 | P | GSH2 |
| 246786_at | 170.9 | P | 139.4 | P | 106.9 | P | 142.9 | P | --- |
| 246787_s_at | 1.8 | A | 1.4 | A | 2.1 | A | 20.7 | A | --- |
| 246788_at | 17.7 | A | 14.3 | A | 19.2 | A | 13.3 | A | AGL89 |
| 246789_at | 1132.4 | P | 869.4 | P | 957 | P | 820.3 | P | LACS7 |
| 246790_at | 41.8 | A | 38.4 | A | 60.4 | A | 27.6 | A | --- |
| 246791_at | 2588.3 | P | 2655.3 | P | 1613.9 | P | 2036.1 | P | --- |
| 246792_at | 402.4 | P | 543.3 | P | 441 | P | 548.8 | P | --- |
| 246793_at | 177 | P | 238.3 | P | 159.1 | P | 195.1 | P | --- |
| 246794_s_at | 105.3 | P | 125.9 | P | 138.1 | P | 131.7 | P | --- |
| 246795_at | 3.9 | A | 2.9 | A | 15.3 | A | 12.6 | A | --- |
| 246796_at | 101.8 | P | 134.4 | P | 79.5 | P | 84 | P | --- |
| 246797_at | 33.4 | P | 87.6 | P | 77.9 | P | 103.9 | P | --- |
| 246798_at | 27.7 | A | 35.6 | A | 14.4 | A | 41.5 | A | --- |
| 246799_at | 270.9 | P | 329.3 | P | 326 | P | 307.9 | P | --- |
| 246800_at | 691.6 | P | 953.1 | P | 734.3 | P | 953.4 | P | SHM2 |
| 246801_at | 2413.9 | P | 2027.1 | P | 2821.6 | P | 2113.3 | P | --- |
| 246802_at | 27.9 | A | 48 | P | 67.6 | P | 75.2 | P | ATK4 |
| 246803_at | 103.9 | P | 70.1 | P | 97.2 | P | 96 | P | SYP41 |
| 246804_at | 1.5 | A | 1.4 | A | 1 | A | 1.8 | A | --- |
| 246805_at | 19.4 | A | 21.2 | A | 26.6 | A | 22.6 | A | AGL101 |
| 246806_at | 2.6 | A | 0.5 | A | 7 | A | 16.8 | A | --- |
| 246807_at | 55.6 | A | 45.5 | A | 58.8 | M | 43.9 | A | GLR2.1 |
| 246808_at | 143.5 | P | 143.7 | P | 161.4 | P | 176.9 | P | --- |
| 246809_s_at | 663.4 | P | 412.2 | P | 719.7 | P | 472 | P | --- |
| 246810_at | 11.6 | A | 14.3 | A | 0.6 | A | 0.9 | A | --- |
| 246811_at | 3.1 | A | 3.3 | A | 9.3 | A | 4.2 | A | --- |
| 246812_at | 1 | A | 0.4 | A | 7.6 | A | 0.4 | A | --- |
| 246813_at | 3.2 | A | 0.8 | A | 2 | A | 1.4 | A | --- |
| 246814_at | 39 | A | 8.1 | A | 50.4 | M | 12.5 | A | ACP5 |
| 246815_at | 87.2 | P | 50.9 | P | 108.6 | P | 61 | P | --- |
| 246816_at | 39.2 | P | 32.6 | A | 65.3 | A | 25.5 | A | --- |
| 246817_at | 50.8 | P | 61.5 | P | 77.5 | P | 87.2 | P | --- |
| 246818_at | 320.1 | P | 299.1 | P | 314.9 | P | 312.3 | P | EMB976 |
| 246819_at | 1.3 | A | 0.1 | A | 0.8 | A | 8.5 | A | --- |
| 246820_at | 192.7 | P | 222.4 | P | 211.3 | P | 218 | P | --- |
| 246821_at | 199.9 | P | 534.5 | P | 209.1 | P | 286.8 | P | CBP60G |
| 246822_at | 182.6 | P | 201 | P | 217.5 | P | 198.2 | P | --- |
| 246823_at | 8.2 | A | 2.2 | A | 6 | A | 1.9 | A | --- |
| 246824_at | 553.6 | P | 575 | P | 534 | P | 578.9 | P | --- |
| 246825_at | 311.7 | P | 125.4 | P | 238.1 | P | 96 | P | --- |
| 246826_at | 4.8 | A | 8.7 | A | 5.3 | A | 9.5 | A | UGT72E3 |
| 246827_at | 58.1 | P | 87.5 | P | 78.9 | P | 93.2 | P | --- |
| 246828_at | 3.4 | A | 3.3 | A | 11.8 | A | 5 | A | --- |
| 246829_at | 4108.2 | P | 3672.1 | P | 3918.4 | P | 4075.9 | P | ATGWD3 |
| 246830_at | 2004.2 | P | 2176.3 | P | 2218.9 | P | 1942.6 | P | --- |
| 246831_at | 1119.8 | P | 1290.9 | P | 618.6 | P | 925.6 | P | MSS1 |
| 246832_at | 1258.2 | P | 793.8 | P | 1338.6 | P | 837.8 | P | --- |
| 246833_at | 1.1 | A | 1.7 | A | 1.3 | A | 1.1 | A | --- |
| 246834_at | 21.8 | A | 17.3 | A | 21.9 | A | 6.9 | A | --- |
| 246835_at | 16.8 | A | 36 | A | 51.6 | A | 25.3 | A | --- |
| 246836_s_at | 1.8 | A | 12.3 | A | 12.1 | A | 8.4 | A | --- |
| 246837_at | 92.7 | P | 219.9 | P | 82.1 | P | 172.9 | P | --- |
| 246838_at | 187.7 | P | 113.4 | P | 196 | P | 137.7 | P | --- |
| 246839_at | 111.5 | P | 79 | P | 77.3 | A | 141.4 | P | --- |
| 246840_at | 478.2 | P | 424.5 | P | 414.6 | P | 365.4 | P | --- |
| 246841_at | 20.9 | A | 12 | A | 9.8 | A | 3.8 | A | --- |
| 246842_at | 106 | P | 77.9 | P | 61.5 | P | 93.4 | P | --- |
| 246843_at | 473.5 | P | 510.7 | P | 437 | P | 457.2 | P | --- |
| 246844_at | 45.8 | A | 54.5 | P | 57.6 | A | 44.7 | A | ATMYB86 |
| 246845_at | 1604.6 | P | 1698.3 | P | 1809.6 | P | 1578.8 | P | --- |
| 246846_at | 541.6 | P | 567.2 | P | 501.3 | P | 609.9 | P | --- |
| 246847_at | 766.3 | P | 854.1 | P | 878.6 | P | 772.1 | P | ATIREG3 |
| 246848_at | 5.4 | A | 9.2 | A | 7.5 | A | 7.4 | A | --- |
| 246849_at | 303.1 | P | 493.3 | P | 485.4 | P | 474.2 | P | --- |
| 246850_at | 488 | P | 332.7 | P | 480.9 | P | 373.1 | P | LON1 |
| 246851_at | 0.3 | A | 2.3 | A | 2 | A | 1.3 | A | --- |
| 246852_at | 2419.5 | P | 2132 | P | 2660.5 | P | 2008.1 | P | --- |
| 246853_at | 35.7 | P | 24.1 | A | 31.9 | A | 11.7 | A | --- |
| 246854_at | 3.3 | A | 3.8 | A | 11.6 | A | 2.1 | A | --- |
| 246855_at | 290.3 | P | 122.7 | P | 389.7 | P | 157.8 | P | --- |
| 246856_at | 504.9 | P | 462.4 | P | 508.3 | P | 450.7 | P | AL4 |
| 246857_at | 6.1 | A | 6.7 | A | 8.7 | A | 1.4 | A | --- |
| 246858_at | 355.2 | P | 367.8 | P | 341.5 | P | 301.2 | P | --- |

| | | | | | | | | | |
|---|---|---|---|---|---|---|---|---|---|
| 246859_at | 1.3 | A | 1.7 | A | 3.7 | A | 2.4 | A | --- |
| 246860_at | 527.5 | P | 585.9 | P | 620.6 | P | 573.9 | P | --- |
| 246861_at | 95.9 | P | 147.4 | P | 138.9 | P | 139.7 | P | IAA28 |
| 246862_at | 1301.6 | P | 1313.6 | P | 1068.5 | P | 1319.3 | P | PEX4 |
| 246863_at | 1650.3 | P | 1298.9 | P | 1430.2 | P | 1137.8 | P | --- |
| 246864_at | 631.4 | P | 574.4 | P | 654 | P | 476.3 | P | GA3 |
| 246865_s_at | 1153.7 | P | 911.1 | P | 1114.4 | P | 1060.5 | P | TIF3B1 |
| 246866_at | 25.6 | A | 6.4 | A | 22.8 | A | 17.6 | A | --- |
| 246867_at | 14.9 | A | 1.7 | A | 4.8 | A | 12.2 | A | --- |
| 246868_at | 27.1 | A | 28.7 | A | 18.6 | A | 22.8 | A | --- |
| 246869_at | 29.5 | A | 15.8 | A | 38.7 | A | 36 | A | --- |
| 246870_at | 2235.6 | P | 2034.4 | P | 2090.7 | P | 1870.7 | P | FC1 |
| 246871_at | 10.2 | A | 21.6 | P | 22.6 | A | 23 | P | --- |
| 246872_at | 23.2 | A | 3.8 | A | 17.8 | A | 11.4 | A | --- |
| 246873_at | 36.2 | A | 24.9 | A | 17.6 | A | 7.1 | A | --- |
| 246874_at | 2.9 | A | 4.3 | A | 7.7 | A | 7.3 | A | ASD2 |
| 246875_at | 20.3 | A | 29.7 | A | 70.2 | P | 15.6 | A | --- |
| 246876_at | 2.5 | A | 1.2 | A | 21.1 | A | 3.7 | A | --- |
| 246877_at | 1.7 | A | 1.1 | A | 4.4 | A | 2.3 | A | --- |
| 246878_at | 19.5 | A | 12.3 | A | 3.6 | A | 3.4 | A | --- |
| 246879_at | 575.7 | P | 361.6 | P | 402.4 | P | 389.6 | P | --- |
| 246880_s_at | 11213.3 | P | 12520.3 | P | 15255.8 | P | 10403.8 | P | TGG2 |
| 246881_at | 177.6 | P | 176.9 | P | 116.4 | P | 148.9 | P | HDA2 |
| 246882_at | 334.7 | P | 177 | P | 233.2 | P | 250.2 | P | --- |
| 246883_at | 61.1 | P | 54.7 | P | 33.9 | P | 58.5 | P | --- |
| 246884_at | 230 | P | 265.4 | P | 718.4 | P | 258.4 | P | --- |
| 246885_at | 79.2 | P | 145.9 | P | 84.7 | P | 149.2 | P | --- |
| 246886_at | 445.1 | P | 436.3 | P | 496.8 | P | 410.5 | P | CLC-D |
| 246887_at | 25.2 | A | 27.2 | A | 33 | A | 11.1 | A | --- |
| 246888_at | 567.8 | P | 264 | P | 366.4 | P | 238.4 | P | --- |
| 246889_at | 16.3 | A | 7.9 | A | 26.5 | A | 15.7 | A | --- |
| 246890_at | 48.5 | A | 17.1 | A | 38.3 | A | 28.3 | A | --- |
| 246891_at | 154 | P | 168 | P | 198.5 | P | 201.1 | P | --- |
| 246892_at | 150.4 | P | 145 | P | 136.2 | P | 155.3 | P | --- |
| 246893_at | 470.4 | P | 661.6 | P | 520.3 | P | 577.7 | P | --- |
| 246894_at | 18.9 | A | 1.7 | A | 1.8 | A | 13.3 | A | --- |
| 246895_at | 1053.8 | P | 1198.7 | P | 1009.5 | P | 1144.9 | P | CID6 |
| 246896_at | 4.5 | A | 3.6 | A | 5.6 | A | 12.7 | A | --- |
| 246897_at | 1100.1 | P | 1032.9 | P | 937.4 | P | 1024.3 | P | --- |
| 246898_at | 38 | A | 57.9 | P | 52.9 | P | 48.4 | P | --- |
| 246899_at | 131.8 | P | 152.5 | P | 183.8 | P | 157.5 | P | --- |
| 246900_at | 20.2 | A | 52.9 | A | 37.6 | A | 70 | M | YUC6 |
| 246901_at | 707.5 | P | 880.5 | P | 658.1 | P | 807.1 | P | --- |
| 246902_at | 32.6 | P | 40.5 | P | 11.3 | A | 23.3 | A | --- |
| 246903_at | 2.4 | A | 8.6 | A | 1 | A | 1.3 | A | --- |
| 246904_at | 105.1 | P | 88.9 | P | 108.7 | P | 98.3 | P | DNMT2 |
| 246905_at | 147.1 | P | 120.7 | P | 206.9 | P | 158.1 | P | --- |
| 246906_at | 125 | P | 134.8 | P | 209.3 | P | 166.5 | P | --- |
| 246907_at | 273.2 | P | 503 | P | 392.3 | P | 482.6 | P | --- |
| 246908_at | 3636.8 | P | 5460.3 | P | 2731.2 | P | 5159.9 | P | RD22 |
| 246909_at | 583.7 | P | 453.8 | P | 347.5 | P | 409.7 | P | --- |
| 246910_at | 215.6 | P | 175.6 | P | 243.9 | P | 171.5 | P | --- |
| 246911_at | 43.8 | A | 64.1 | P | 81.7 | A | 64.2 | P | tny |
| 246912_at | 45.1 | P | 29.5 | P | 2.8 | A | 38.2 | P | --- |
| 246913_at | 18.4 | A | 62.7 | A | 94.4 | P | 113.8 | P | --- |
| 246914_at | 63.6 | A | 57.5 | A | 56.4 | A | 53.4 | A | --- |
| 246915_at | 41 | P | 33.3 | A | 11.9 | A | 34.7 | A | ATNADP-ME3 |
| 246916_at | 13.8 | A | 14.6 | P | 0.9 | A | 20 | A | AtRLP52 |
| 246917_at | 1356.8 | P | 1444.7 | P | 1231 | P | 1177.6 | P | --- |
| 246918_at | 25.5 | A | 28.9 | A | 33 | A | 34.3 | A | --- |
| 246919_at | 3365.6 | P | 3414.9 | P | 4842.3 | P | 3308.6 | P | --- |
| 246920_at | 172.9 | P | 354.9 | P | 189.1 | P | 335.4 | P | --- |
| 246921_at | 403.4 | P | 413.6 | P | 387.9 | P | 371.7 | P | --- |
| 246922_at | 46.8 | A | 66 | A | 9.8 | A | 47 | A | CIPK25 |
| 246923_at | 1132.3 | P | 1337.3 | P | 1278.8 | P | 1362.6 | P | --- |
| 246924_at | 359.1 | P | 330.7 | P | 356.4 | P | 341.1 | P | --- |
| 246925_at | 45.2 | A | 37.3 | A | 32.7 | A | 19.2 | A | CYP71B14 |
| 246926_at | 102.5 | P | 122.7 | P | 101.4 | P | 143.3 | P | --- |
| 246927_s_at | 68 | P | 182.6 | P | 11.6 | A | 118.3 | P | --- |
| 246928_at | 392.7 | P | 420.9 | P | 361.3 | P | 404 | P | --- |
| 246929_at | 113 | P | 108.4 | P | 89.2 | P | 93.3 | P | --- |
| 246930_at | 488.5 | P | 359.5 | P | 339 | P | 315.1 | P | KNAT3 |
| 246931_at | 361.5 | P | 418.1 | P | 371.1 | P | 365.6 | P | --- |
| 246932_at | 102.4 | P | 197.5 | P | 115.4 | P | 116.9 | P | --- |
| 246933_at | 2.2 | A | 12.9 | A | 5.5 | A | 15.2 | A | ZFP3 |
| 246934_at | 16.1 | A | 18.5 | A | 13.9 | A | 17.1 | A | --- |
| 246935_at | 599.7 | P | 949.4 | P | 598.8 | P | 791.4 | P | EBF2 |
| 246936_at | 449.3 | P | 700.2 | P | 408 | P | 603.9 | P | --- |
| 246937_at | 12.3 | A | 23.5 | A | 24.4 | P | 32.3 | A | PLDALPHA3 |
| 246938_at | 24.2 | P | 43.3 | A | 15 | A | 32 | P | CYCA2;1 |
| 246939_at | 19.9 | A | 156.7 | P | 48.3 | P | 133.2 | P | SHN2 |
| 246940_at | 12.2 | A | 16.3 | A | 34.9 | A | 16.9 | A | --- |
| 246941_at | 37.3 | A | 4.3 | A | 4.2 | A | 27.9 | A | --- |
| 246942_at | 9.5 | A | 4.2 | A | 27.2 | A | 15.9 | A | --- |
| 246943_at | 308.5 | P | 608 | P | 171.9 | P | 328.3 | P | --- |
| 246944_at | 472 | P | 351.3 | P | 361.2 | P | 263.2 | P | --- |
| 246945_s_at | 513.2 | P | 531.5 | P | 375.4 | P | 562.4 | P | --- |
| 246946_at | 277.2 | P | 273.4 | P | 271.8 | P | 300.8 | P | --- |
| 246947_at | 51.2 | P | 112.2 | P | 59.1 | A | 103.4 | P | CYP71B11 |
| 246948_at | 8.5 | A | 27.2 | P | 12.4 | A | 24.7 | P | CYP71B12 |
| 246949_at | 17.9 | P | 32.9 | P | 14.9 | A | 17.8 | P | CYP71B13 |
| 246950_at | 187.9 | P | 208.3 | P | 194.8 | P | 222.8 | P | TAF5 |
| 246951_at | 129.7 | P | 102.4 | P | 117.4 | P | 108.1 | P | --- |
| 246952_at | 181.7 | P | 171.4 | P | 216.8 | P | 176 | P | OFP13 |
| 246953_at | 480.3 | P | 452.1 | P | 537.3 | P | 422.3 | P | VPS60.2 |
| 246954_at | 2870.7 | P | 2745.5 | P | 2645.6 | P | 2868 | P | --- |
| 246955_at | 328.1 | P | 316 | P | 332.9 | P | 286.2 | P | CPK1 |
| 246956_at | 65.5 | A | 93.4 | P | 112 | A | 95.7 | P | RTM2 |
| 246957_at | 815.1 | P | 763.1 | P | 948.4 | P | 799.2 | P | --- |
| 246958_at | 3162.8 | P | 3591.8 | P | 4358.6 | P | 3646.3 | P | --- |
| 246959_at | 1466.6 | P | 1763 | P | 1515.5 | P | 1799.5 | P | --- |
| 246960_at | 397.7 | P | 391.3 | P | 446.8 | P | 398.4 | P | --- |
| 246961_at | 52.6 | P | 54.4 | P | 79.6 | P | 60.7 | P | --- |
| 246962_s_at | 726.8 | P | 736.6 | P | 757.1 | P | 686.7 | P | BZIP9 |
| 246963_at | 9.1 | A | 28.3 | A | 41.6 | A | 3.8 | A | --- |
| 246964_at | 162.1 | P | 197.7 | P | 192.3 | P | 187.2 | P | --- |
| 246965_at | 546.4 | P | 407.6 | P | 400.8 | P | 396.2 | P | --- |
| 246966_at | 160.8 | P | 234.6 | P | 324.6 | P | 267.1 | P | CRY3 |
| 246967_at | 19.3 | A | 18.8 | A | 28.9 | A | 21.8 | P | FPF1 |

| | | | | | | | | | |
|---|---|---|---|---|---|---|---|---|---|
| 246968_at | 242.6 | P | 243.6 | P | 207.1 | P | 266.6 | P | --- |
| 246969_at | 28.2 | A | 8.9 | A | 62.5 | P | 13.1 | A | --- |
| 246970_at | 44.8 | A | 40.3 | A | 9.2 | A | 41.3 | A | CYP714A2 |
| 246971_at | 3.7 | A | 7 | A | 57.2 | A | 26.5 | A | --- |
| 246972_s_at | 44.6 | P | 64.4 | P | 66.9 | P | 82.3 | A | CYP71A14 |
| 246973_at | 340.1 | P | 288.4 | P | 249.2 | P | 292.9 | P | --- |
| 246974_at | 122.7 | P | 81.3 | P | 126.1 | P | 73.7 | P | --- |
| 246975_at | 462.1 | P | 567.7 | P | 442.7 | P | 552.1 | P | --- |
| 246976_s_at | 940.8 | P | 1261.3 | P | 893.5 | P | 1075.8 | P | --- |
| 246977_at | 3427.6 | P | 3228.3 | P | 2309 | P | 3640.3 | P | COL4 |
| 246978_at | 59.4 | P | 8.2 | A | 78 | A | 18.2 | A | CYP714A1 |
| 246979_s_at | 172.2 | P | 91.7 | P | 152.9 | P | 136.9 | P | --- |
| 246980_at | 286.2 | P | 316.8 | P | 289.9 | P | 328.4 | P | --- |
| 246981_at | 74.6 | A | 67.9 | P | 78.9 | A | 79 | A | --- |
| 246982_s_at | 55.8 | P | 88.3 | P | 40.8 | P | 83.9 | P | --- |
| 246983_at | 165.6 | P | 163.3 | P | 268.6 | P | 166.6 | P | --- |
| 246984_at | 64.9 | A | 54.5 | A | 37.9 | A | 50 | A | CYP81G1 |
| 246985_at | 659.2 | P | 698.4 | P | 566.8 | P | 724.8 | P | --- |
| 246986_at | 858 | P | 824.1 | P | 624.1 | P | 930.8 | P | RLK |
| 246987_at | 1544.3 | P | 1215.7 | P | 1388.7 | P | 1353 | P | MYBR1 |
| 246988_at | 54.9 | A | 106.9 | P | 50.9 | A | 75.9 | A | --- |
| 246989_at | 120.7 | P | 130.3 | P | 109.3 | P | 73.8 | P | --- |
| 246990_at | 3694.9 | P | 3455.3 | P | 3047.1 | P | 3653.4 | P | ARA12 |
| 246991_at | 2.5 | A | 2.4 | A | 4.5 | A | 1.6 | A | --- |
| 246992_at | 19 | A | 3.2 | A | 5.7 | A | 10.9 | A | --- |
| 246993_at | 36.3 | P | 59.7 | P | 49.2 | P | 27.9 | A | AZF1 |
| 246994_at | 108.6 | P | 118 | P | 96.2 | P | 123.7 | P | --- |
| 246995_at | 430.6 | P | 324.5 | P | 355.2 | P | 345.3 | P | --- |
| 246996_at | 177.6 | P | 240.1 | P | 208.4 | P | 224.8 | P | LBD37 |
| 246997_at | 87.5 | P | 93 | P | 61.4 | P | 118.9 | P | --- |
| 246998_at | 663.5 | P | 1873.2 | P | 2584.1 | P | 2308.3 | P | --- |
| 246999_at | 281.1 | P | 222.6 | P | 267.7 | P | 222.5 | P | NPY3 |
| 247000_at | 1112.6 | P | 841.3 | P | 1191.4 | P | 874.8 | P | CKA1 |
| 247001_at | 1786.2 | P | 2390.7 | P | 2244.2 | P | 2399.6 | P | NRAMP4 |
| 247002_at | 525.9 | P | 614.6 | P | 705.4 | P | 688.9 | P | HOS15 |
| 247003_at | 5.6 | A | 14.9 | A | 6.6 | A | 11.4 | A | --- |
| 247004_at | 116.9 | P | 114.4 | P | 109.4 | P | 116.1 | P | DG1 |
| 247005_at | 69.2 | P | 52.2 | P | 54.8 | A | 87.1 | P | --- |
| 247006_at | 2416.2 | P | 1636.2 | P | 2141.3 | P | 2125.9 | P | --- |
| 247007_at | 2751.2 | P | 2337.6 | P | 3431.7 | P | 2539.9 | P | VDAC2 |
| 247008_at | 773.2 | P | 1015 | P | 635.9 | P | 861.8 | P | ATARLA1D |
| 247009_at | 7834.8 | P | 3981.7 | P | 5109.9 | P | 4287.9 | P | --- |
| 247010_at | 3282.1 | P | 3012.6 | P | 4013.9 | P | 3331.5 | P | --- |
| 247011_at | 4353.5 | P | 4612.5 | P | 4854.3 | P | 4821.3 | P | FRO1 |
| 247012_at | 478.2 | P | 501.9 | P | 488.2 | P | 551.9 | P | TRB2 |
| 247013_at | 3310.8 | P | 2751.2 | P | 3379.6 | P | 3363.8 | P | BT4 |
| 247014_at | 447 | P | 368.6 | P | 475 | P | 355.8 | P | --- |
| 247015_at | 88 | P | 70.4 | P | 102.8 | P | 89.8 | P | --- |
| 247016_at | 17.9 | A | 1.4 | A | 10.2 | A | 6.1 | A | --- |
| 247017_at | 2.9 | A | 7.9 | A | 15.1 | A | 2.4 | A | --- |
| 247018_at | 16.1 | A | 32.5 | A | 23.2 | A | 4 | A | --- |
| 247019_at | 17.4 | A | 25.6 | A | 44.8 | A | 12.7 | A | --- |
| 247020_at | 73 | M | 62.7 | P | 27.2 | A | 52.4 | P | --- |
| 247021_at | 2.4 | A | 1.7 | A | 4 | A | 4.9 | A | --- |
| 247022_at | 3.4 | A | 49.9 | P | 60.9 | P | 42.1 | A | --- |
| 247023_at | 45.9 | P | 127.2 | P | 92.1 | P | 58.2 | P | HEC1 |
| 247024_at | 15 | A | 27 | P | 37.4 | A | 19.3 | A | --- |
| 247025_at | 2200.7 | P | 3052.2 | P | 2976.3 | P | 2871 | P | ABA1 |
| 247026_at | 2.5 | A | 2.6 | A | 40.4 | A | 4.4 | A | MAPKKK19 |
| 247027_at | 31.7 | A | 56.9 | A | 32.8 | A | 21.4 | A | --- |
| 247028_at | 93.4 | P | 134.7 | P | 113.3 | P | 135.2 | P | ICU2 |
| 247029_at | 98.6 | A | 115.9 | A | 115.9 | A | 85 | A | --- |
| 247030_at | 88.5 | P | 101.9 | P | 56 | P | 71.1 | P | --- |
| 247031_at | 100.4 | P | 126.8 | P | 135.6 | P | 178.8 | P | --- |
| 247032_at | 605 | P | 413.1 | P | 570.9 | P | 473.9 | P | SDN3 |
| 247033_at | 552.2 | P | 534.1 | P | 453.4 | P | 554.8 | P | SKIP2 |
| 247034_at | 1299.6 | P | 1656.4 | P | 1403.9 | P | 1835.7 | P | CYCD3;2 |
| 247035_at | 126.6 | P | 137.8 | P | 169.3 | P | 147.6 | P | ALC |
| 247036_at | 651.9 | P | 772.8 | P | 823.2 | P | 739.8 | P | --- |
| 247037_at | 1960 | P | 3204.8 | P | 2001 | P | 3164.7 | P | RALFL34 |
| 247038_at | 221.6 | P | 147.7 | P | 173.4 | P | 124.9 | P | EPS1 |
| 247039_at | 109.1 | P | 133.5 | P | 126.7 | P | 180.2 | P | ATEB1C |
| 247040_at | 710.2 | P | 451.6 | P | 1026.2 | P | 564 | P | --- |
| 247041_at | 184.4 | P | 67 | P | 142.9 | P | 87.2 | P | TOE3 |
| 247042_at | 1570 | P | 1475.2 | P | 1982.1 | P | 1553.1 | P | --- |
| 247043_at | 908.4 | P | 809.7 | P | 895 | P | 812.2 | P | SNRK2.3 |
| 247044_at | 135.9 | P | 262.1 | P | 172.3 | P | 224.5 | P | MAPKKK5 |
| 247045_at | 269.1 | P | 310.2 | P | 227.8 | P | 301 | P | --- |
| 247046_at | 421.4 | P | 257.7 | P | 402.3 | P | 324.7 | P | --- |
| 247047_at | 1653 | P | 229.5 | P | 1768.9 | P | 389.8 | P | --- |
| 247048_at | 208.9 | P | 219.7 | P | 202.9 | P | 231.9 | P | --- |
| 247049_at | 448 | P | 545.5 | P | 400.1 | P | 475.3 | P | --- |
| 247050_at | 34.2 | A | 7.5 | A | 46.5 | A | 9.4 | A | --- |
| 247051_at | 21.2 | A | 7.4 | A | 16.3 | A | 39.2 | A | --- |
| 247052_at | 34.9 | P | 30.9 | P | 62.8 | P | 31.5 | P | HB53 |
| 247053_at | 47.6 | A | 37 | A | 28.2 | A | 24.9 | A | --- |
| 247054_at | 335.6 | P | 300.9 | P | 313.5 | P | 300.7 | P | --- |
| 247055_at | 19.7 | A | 24 | A | 54.3 | P | 28.9 | A | --- |
| 247056_at | 119.2 | P | 177.7 | P | 140 | P | 187.5 | P | CHR1 |
| 247057_at | 97.6 | P | 175.3 | P | 188.6 | P | 135.2 | P | --- |
| 247058_at | 2824.4 | P | 3169.9 | P | 3095.5 | P | 2935.6 | P | DGL1 |
| 247059_at | 77 | P | 124.4 | P | 139.2 | P | 128.6 | P | UGT72E2 |
| 247060_at | 3701 | P | 2629.4 | P | 3878 | P | 3132.3 | P | SDH1-1 |
| 247061_at | 1.3 | A | 13.7 | A | 3.3 | A | 29.4 | A | --- |
| 247062_at | 280.8 | P | 272.6 | P | 251.6 | P | 276.6 | P | --- |
| 247063_at | 383.4 | P | 336.4 | P | 485.4 | P | 391.7 | P | --- |
| 247064_at | 8.1 | A | 30.8 | P | 19.4 | A | 26.1 | P | --- |
| 247065_s_at | 484 | P | 436 | P | 404.4 | P | 369.5 | P | --- |
| 247066_at | 70.8 | P | 70.3 | P | 56.5 | A | 97.2 | P | --- |
| 247067_at | 406.4 | P | 477.8 | P | 497.4 | P | 468.1 | P | --- |
| 247068_at | 51.1 | A | 80.9 | P | 64.7 | P | 108.2 | P | --- |
| 247069_at | 160.6 | P | 300.9 | P | 348 | P | 270.3 | P | sks17 |
| 247070_at | 8.3 | A | 9.5 | A | 4.1 | A | 3 | A | --- |
| 247071_at | 39.8 | P | 58.6 | P | 58.9 | P | 27.4 | P | DAR3 |
| 247072_at | 388.7 | P | 414.6 | P | 366.7 | P | 354.3 | P | --- |
| 247073_at | 16194 | P | 15688.1 | P | 15459.1 | P | 15042.9 | P | PSBO1 |
| 247074_at | 133.2 | P | 171.9 | P | 160.5 | P | 194.9 | P | --- |
| 247075_at | 862.9 | P | 633.8 | P | 732.3 | P | 676.5 | P | PLP3b |
| 247076_at | 3017.8 | P | 2589.5 | P | 2623.4 | P | 2490.1 | P | GAMMA CA3 |

| | | | | | | | | | |
|---|---|---|---|---|---|---|---|---|---|
| 247077_at | 669.4 | P | 857.4 | P | 779.2 | P | 847.2 | P | --- |
| 247078_at | 130.6 | P | 137.6 | P | 191.6 | M | 140.1 | P | --- |
| 247079_at | 858.6 | P | 841 | P | 953.3 | P | 837.8 | P | AKRP |
| 247080_at | 1400 | P | 1504.5 | P | 1240 | P | 1657.7 | P | PAD2 |
| 247081_at | 148.3 | P | 125.3 | P | 152.2 | P | 142.5 | P | ATRAD17 |
| 247082_at | 674.9 | P | 554.6 | P | 552.6 | P | 547.6 | P | ATGRIP |
| 247083_at | 252.9 | P | 283.9 | P | 268.4 | P | 245.6 | P | --- |
| 247084_at | 2.7 | A | 31.7 | A | 4 | A | 33.2 | A | NAC105 |
| 247085_at | 64.4 | P | 201.1 | P | 58.3 | P | 183.7 | P | --- |
| 247086_at | 328.1 | P | 334.3 | P | 342 | P | 324.7 | P | --- |
| 247087_at | 59.6 | P | 25.7 | A | 26.3 | A | 64.8 | P | --- |
| 247088_at | 39.7 | P | 40.6 | A | 57.1 | A | 32.2 | A | --- |
| 247089_at | 244.1 | P | 280.4 | P | 182 | P | 234.7 | P | --- |
| 247090_at | 15.9 | A | 8.4 | A | 5.2 | A | 10.1 | A | --- |
| 247091_at | 1.4 | A | 25.2 | A | 17.8 | A | 13.9 | A | --- |
| 247092_at | 326.8 | P | 304.6 | P | 379.3 | P | 307.2 | P | ATFOLT1 |
| 247093_at | 79.6 | P | 127.1 | P | 135 | P | 88.2 | P | SHI |
| 247094_at | 26.1 | A | 19.8 | A | 30.9 | A | 17.5 | A | GMD1 |
| 247095_at | 182.2 | P | 136.7 | P | 123.5 | P | 216.6 | P | RAB18 |
| 247096_at | 19.8 | A | 9.7 | A | 5.6 | A | 2 | A | --- |
| 247097_at | 117.7 | P | 233.7 | P | 155.5 | P | 218 | P | --- |
| 247098_at | 546.9 | P | 752.6 | P | 771.7 | P | 750.4 | P | --- |
| 247099_at | 85.6 | P | 42.6 | P | 49.2 | M | 34.6 | P | --- |
| 247100_at | 140 | P | 99 | P | 187.6 | P | 88.9 | P | --- |
| 247101_at | 1009.3 | P | 1734.9 | P | 1126 | P | 1708.1 | P | --- |
| 247102_at | 188.4 | P | 244.5 | P | 145.8 | P | 178.2 | P | --- |
| 247103_at | 58.6 | P | 89.1 | P | 44.4 | A | 86.7 | P | DAR7 |
| 247104_at | 73.5 | P | 35.8 | M | 73.1 | P | 44.6 | P | DAR6 |
| 247105_at | 67.8 | P | 53.9 | P | 33.9 | A | 63.1 | P | DAR5 |
| 247106_at | 1173.9 | P | 1112.6 | P | 1617.9 | P | 1014.9 | P | --- |
| 247107_at | 5504.3 | P | 5369 | P | 4595.7 | P | 5480.2 | P | STR16 |
| 247108_at | 257.7 | P | 279.9 | P | 299.5 | P | 231.1 | P | RMR1 |
| 247109_at | 1810.1 | P | 905.9 | P | 1161.7 | P | 1095.1 | P | ATPSK5 |
| 247110_at | 167.4 | P | 152.9 | P | 195 | P | 176.9 | P | --- |
| 247111_at | 771.3 | P | 592.6 | P | 662.5 | P | 587.4 | P | --- |
| 247112_at | 1706.6 | P | 1441.1 | P | 1628.5 | P | 1461 | P | --- |
| 247113_at | 773.7 | P | 757.6 | P | 909.6 | P | 652.8 | P | --- |
| 247114_at | 287.6 | P | 391.4 | P | 308.3 | P | 371.9 | P | --- |
| 247115_at | 237.7 | P | 242.2 | P | 241.1 | P | 273 | P | ZWI |
| 247116_at | 62.2 | P | 118.7 | P | 138.8 | P | 92 | P | MLO10 |
| 247117_at | 1610.7 | P | 1487.2 | P | 1292.5 | P | 1381.1 | P | CHY1 |
| 247118_at | 149.4 | P | 203.8 | P | 89.6 | P | 202.8 | P | ACR1 |
| 247119_at | 295.8 | P | 260.8 | P | 342.8 | P | 269.7 | P | --- |
| 247120_at | 1288.5 | P | 1034.2 | P | 977.5 | P | 1138.9 | P | --- |
| 247121_at | 289.5 | P | 299.1 | P | 274.5 | P | 289.2 | P | --- |
| 247122_at | 32.1 | A | 33.1 | A | 6.8 | A | 44.2 | A | ATSAC1B |
| 247123_at | 354.2 | P | 332.6 | P | 446.8 | P | 350.2 | P | --- |
| 247124_at | 368.1 | P | 407.9 | P | 238.3 | P | 358.2 | P | --- |
| 247125_at | 360.1 | P | 387.3 | P | 332.6 | P | 303.6 | P | --- |
| 247126_at | 213.6 | P | 68.8 | P | 115.1 | P | 72 | P | --- |
| 247127_at | 334.1 | P | 268.8 | P | 404.6 | P | 322.9 | P | --- |
| 247128_at | 17.4 | A | 25.7 | A | 53.8 | A | 31.3 | A | --- |
| 247129_at | 25 | A | 28.9 | A | 5.4 | A | 4.4 | A | --- |
| 247130_at | 586.4 | P | 454.4 | P | 541.6 | P | 476.3 | P | --- |
| 247131_at | 3552.2 | P | 7787.9 | P | 4166.3 | P | 6980.7 | P | FNR1 |
| 247132_at | 651 | P | 630 | P | 660 | P | 601.1 | P | ARO2 |
| 247133_at | 14.6 | A | 20.7 | A | 69 | A | 24.4 | A | --- |
| 247134_at | 234.4 | P | 301 | P | 249.7 | P | 343.9 | P | --- |
| 247135_at | 12.7 | A | 13.4 | A | 23.9 | A | 19.9 | A | --- |
| 247136_at | 627.1 | P | 239.2 | P | 284.6 | P | 217.3 | P | --- |
| 247137_at | 362.9 | P | 456.7 | P | 357.7 | P | 360.3 | P | CPK28 |
| 247138_at | 1884.1 | P | 1546.6 | P | 1674.2 | P | 1543.2 | P | --- |
| 247139_at | 902.1 | P | 794.1 | P | 811.2 | P | 747.9 | P | --- |
| 247140_at | 217.9 | P | 282 | P | 186.5 | P | 207.5 | P | --- |
| 247141_at | 155.9 | P | 140.2 | P | 144.4 | P | 133.8 | P | --- |
| 247142_at | 69.2 | P | 67.8 | P | 92.4 | P | 73.1 | P | --- |
| 247143_at | 22.1 | A | 30.2 | A | 37.2 | A | 18.4 | A | --- |
| 247144_at | 62.4 | A | 60.8 | A | 13.4 | A | 8.9 | A | --- |
| 247145_at | 38.6 | P | 30.1 | A | 25.5 | A | 2.8 | A | --- |
| 247146_at | 27.9 | A | 26.2 | A | 43.1 | A | 75.4 | A | --- |
| 247147_at | 1683.5 | P | 1550.9 | P | 1765.4 | P | 1809.4 | P | GTE7 |
| 247148_at | 2026.5 | P | 2315.3 | P | 1903.8 | P | 2419.6 | P | IAA9 |
| 247149_at | 2479.3 | P | 1014.5 | P | 2267.5 | P | 1317.5 | P | --- |
| 247150_at | 386.5 | P | 400.8 | P | 390.9 | P | 399.8 | P | --- |
| 247151_at | 502.1 | P | 531.8 | P | 486.3 | P | 509 | P | bHLH093 |
| 247152_at | 1483.6 | P | 1606.3 | P | 1656.6 | P | 1500.2 | P | --- |
| 247153_at | 734.9 | P | 861 | P | 1002.6 | P | 961.5 | P | BAM1 |
| 247154_at | 200.9 | P | 255.6 | P | 244.1 | P | 221 | P | HSL2 |
| 247155_at | 866.4 | P | 547.4 | P | 921.7 | P | 534.6 | P | --- |
| 247156_at | 1005.8 | P | 1034.9 | P | 1031.2 | P | 1114.1 | P | --- |
| 247157_at | 360 | P | 435.4 | P | 508.1 | P | 417.7 | P | LINC4 |
| 247158_at | 933.2 | P | 1051.9 | P | 951.7 | P | 971.4 | P | ATBCAT-5 |
| 247159_at | 1.6 | A | 3.7 | A | 2.6 | A | 2.7 | A | ACS5 |
| 247160_at | 17 | A | 12.1 | A | 31.1 | A | 31.2 | P | --- |
| 247161_at | 337 | P | 672.5 | P | 428.6 | P | 680.5 | P | --- |
| 247162_at | 975.1 | P | 957.4 | P | 841.7 | P | 878.9 | P | --- |
| 247163_at | 814.3 | P | 935.9 | P | 922.1 | P | 887.9 | P | --- |
| 247164_at | 1301.1 | P | 902.5 | P | 1164.1 | P | 1028.8 | P | NFS1 |
| 247165_at | 10.8 | A | 35.2 | A | 23.7 | A | 5.4 | A | MYB68 |
| 247166_at | 1171.2 | P | 1607.5 | P | 1447.9 | P | 1662.3 | P | --- |
| 247167_at | 89.7 | P | 58 | P | 149.4 | P | 60 | P | --- |
| 247168_at | 616 | P | 539.6 | P | 777.1 | P | 665.7 | P | --- |
| 247169_at | 153.9 | P | 166.4 | P | 182 | P | 137.6 | P | --- |
| 247170_at | 27.2 | A | 44.2 | A | 6.8 | A | 35.3 | A | --- |
| 247171_at | 169.5 | P | 180.8 | P | 219.5 | P | 186.4 | P | --- |
| 247172_at | 49.5 | A | 51.2 | P | 60.9 | P | 43.2 | A | --- |
| 247173_at | 1747.3 | P | 1884.7 | P | 1976.4 | P | 1854.7 | P | --- |
| 247174_at | 537.7 | P | 534.6 | P | 556.6 | P | 639 | P | --- |
| 247175_at | 252 | P | 126.4 | P | 189.1 | P | 211.5 | P | GCL1 |
| 247176_at | 1445.6 | P | 979.5 | P | 1046.2 | P | 853.6 | P | ACX2 |
| 247177_at | 203.3 | P | 186.4 | P | 174.3 | P | 174.1 | P | --- |
| 247178_at | 343.5 | P | 243.9 | P | 268.2 | P | 209.9 | P | --- |
| 247179_at | 54.2 | P | 68.1 | P | 67 | A | 36.5 | M | --- |
| 247180_at | 45.9 | A | 45 | A | 6.2 | A | 56.8 | A | --- |
| 247181_at | 15.4 | A | 25.4 | P | 40.5 | P | 27.2 | P | --- |
| 247182_at | 390.2 | P | 368.1 | P | 424.2 | P | 442.8 | P | HB25 |
| 247183_at | 366.7 | P | 429.7 | P | 434.4 | P | 417.1 | P | --- |
| 247184_at | 174 | P | 131.6 | P | 209.2 | P | 207.2 | P | UBP17 |
| 247185_at | 276.8 | P | 264.8 | P | 218.8 | P | 280 | P | --- |

| | | | | | | | | | |
|---|---|---|---|---|---|---|---|---|---|
| 247186_at | 862.8 | P | 842 | P | 638.6 | P | 855.4 | P | --- |
| 247187_at | 330.3 | P | 313.6 | P | 245.5 | P | 275.7 | P | --- |
| 247188_at | 2702.1 | P | 2904.7 | P | 2793.1 | P | 2862.3 | P | GRF8 |
| 247189_at | 296.1 | P | 415.8 | P | 434.2 | P | 384.3 | P | AGP7 |
| 247190_at | 63.7 | P | 88.1 | P | 139.9 | P | 98.6 | P | CYCD4;1 |
| 247191_at | 807.2 | P | 1256.4 | P | 778.9 | P | 1153.4 | P | ATHB5 |
| 247192_at | 2559.3 | P | 2577.9 | P | 1874 | P | 3023.7 | P | --- |
| 247193_at | 2294.2 | P | 2174.2 | P | 3242.2 | P | 2428.4 | P | --- |
| 247194_at | 2005.4 | P | 1889.7 | P | 1745.4 | P | 1712.7 | P | --- |
| 247195_at | 6.4 | A | 7.1 | A | 49.2 | A | 3.3 | A | --- |
| 247196_at | 35.6 | A | 31.2 | P | 44.4 | A | 45 | A | AIL7 |
| 247197_at | 190.7 | P | 167.7 | P | 125.6 | P | 105.4 | P | --- |
| 247198_at | 150.2 | P | 171.7 | P | 156 | P | 150 | P | --- |
| 247199_at | 211.8 | P | 194.7 | P | 209.2 | P | 162.8 | P | TGA1 |
| 247200_at | 58.2 | P | 74.2 | P | 129.1 | P | 101.8 | P | --- |
| 247201_at | 5250.7 | P | 6029.5 | P | 5414.2 | P | 6097.4 | P | --- |
| 247202_at | 862.8 | P | 970.7 | P | 781.6 | P | 1081.4 | P | AtRABA4a |
| 247203_at | 183.2 | P | 149.4 | P | 172.8 | P | 118.5 | P | --- |
| 247204_at | 35.7 | P | 40.9 | P | 45.8 | P | 33.4 | P | AtRABH1a |
| 247205_at | 4.4 | A | 4.8 | A | 9.3 | A | 3.8 | A | PROPEP2 |
| 247206_at | 180.1 | P | 164.8 | P | 269.4 | P | 217.9 | P | --- |
| 247207_at | 488.8 | P | 579.3 | P | 483 | P | 562 | P | --- |
| 247208_at | 85.8 | P | 122.5 | P | 78.8 | P | 82.5 | P | --- |
| 247209_at | 467 | P | 397 | P | 441.6 | P | 352.4 | P | --- |
| 247210_at | 3932.7 | P | 2385.4 | P | 3426.2 | P | 2702.2 | P | ANNAT2 |
| 247211_at | 255.9 | P | 317.5 | P | 266.2 | P | 303.6 | P | --- |
| 247212_at | 465.8 | P | 632.7 | P | 556.2 | P | 635.4 | P | --- |
| 247213_at | 86.6 | P | 92.5 | P | 112.9 | A | 86.7 | P | PROPEP1 |
| 247214_at | 297.6 | P | 492.6 | P | 387 | P | 466.7 | P | --- |
| 247215_at | 26.6 | A | 5 | A | 4.1 | A | 4.3 | A | PROPEP3 |
| 247216_at | 1298.9 | P | 1437.8 | P | 1256.4 | P | 1494.4 | P | DPE1 |
| 247217_s_at | 146.5 | P | 111.4 | P | 63.3 | P | 88.1 | P | MAF3 |
| 247218_at | 523.4 | P | 567.1 | P | 644.3 | P | 543.3 | P | ASN2 |
| 247219_at | 649.9 | P | 550.9 | P | 595.9 | P | 619.6 | P | CIP8 |
| 247220_at | 761.4 | P | 662.8 | P | 748.5 | P | 738.7 | P | CDKC2 |
| 247221_at | 375 | P | 489.7 | P | 451.2 | P | 346 | P | LIP1 |
| 247222_at | 1666.4 | P | 2710.7 | P | 1926 | P | 2329.4 | P | ATGCN5 |
| 247223_at | 23.3 | A | 31.9 | A | 36.3 | A | 23.7 | A | MAF4 |
| 247224_at | 37.9 | P | 71.1 | P | 43.9 | A | 70.4 | P | MAF5 |
| 247225_at | 58 | P | 88 | P | 88 | P | 78.9 | P | BST1 |
| 247226_at | 21.3 | A | 8.2 | A | 27.6 | A | 18.3 | A | --- |
| 247227_at | 4.2 | A | 20.3 | A | 4.7 | A | 2.7 | A | --- |
| 247228_at | 45.6 | P | 75.2 | P | 45.8 | P | 80 | P | --- |
| 247229_at | 33 | A | 5 | A | 37.7 | A | 11.9 | A | --- |
| 247230_at | 57.8 | M | 56.9 | P | 39.7 | P | 62.8 | P | --- |
| 247231_at | 36.7 | A | 55 | A | 45.6 | A | 37.1 | A | AtMYB53 |
| 247232_at | 825.9 | P | 1828 | P | 1068.7 | P | 1669.1 | P | ATH13 |
| 247233_at | 217.7 | P | 198.5 | P | 215 | P | 247.3 | P | --- |
| 247234_at | 75.5 | P | 98.5 | P | 109.5 | P | 95.8 | P | --- |
| 247235_at | 537.2 | P | 642.2 | P | 557.3 | P | 626.7 | P | --- |
| 247236_at | 22.5 | A | 31.6 | A | 3.9 | A | 5.4 | A | --- |
| 247237_at | 181.7 | P | 182.7 | P | 125.8 | P | 172 | P | --- |
| 247238_at | 406.2 | P | 423.2 | P | 405.2 | P | 382.7 | P | HAM1 |
| 247239_at | 145.5 | P | 156.9 | P | 198.3 | P | 195.8 | P | --- |
| 247240_at | 126 | P | 139.2 | P | 130.5 | P | 121.1 | P | CMPG2 |
| 247241_at | 572.6 | P | 335.1 | P | 518.2 | P | 387.5 | P | --- |
| 247242_at | 12.9 | A | 2.1 | A | 3.5 | A | 6.4 | A | --- |
| 247243_at | 137.2 | P | 101.6 | P | 127.8 | P | 90.5 | P | --- |
| 247244_at | 69.7 | P | 54.5 | P | 79.6 | P | 66.9 | P | --- |
| 247245_at | 11.8 | A | 29.7 | A | 4.8 | A | 3.6 | A | --- |
| 247246_at | 515.4 | P | 867.3 | P | 666.3 | P | 957.7 | P | C/VIF2 |
| 247247_at | 509.4 | P | 505.4 | P | 571.9 | P | 466.6 | P | --- |
| 247248_at | 592.7 | P | 480.6 | P | 457.5 | P | 547.3 | P | --- |
| 247249_at | 596 | P | 666.1 | P | 763.5 | P | 734.4 | P | --- |
| 247250_at | 84.3 | P | 132.7 | P | 104.3 | A | 126.5 | M | FAS2 |
| 247251_at | 1344 | P | 1977.8 | P | 1506.2 | P | 1864.8 | P | CESA6 |
| 247252_at | 362.4 | P | 636.1 | P | 405.1 | P | 676.8 | P | --- |
| 247253_at | 25.8 | A | 24.9 | A | 46.1 | A | 29.7 | A | --- |
| 247254_at | 8.4 | A | 42.6 | P | 37.4 | P | 55.3 | P | CLE21 |
| 247255_at | 93.8 | P | 114.2 | P | 105.5 | P | 117 | P | --- |
| 247256_at | 404.9 | P | 319.3 | P | 440.2 | P | 317 | P | --- |
| 247257_at | 363.5 | P | 421.9 | P | 290.9 | P | 413.3 | P | RPN5B |
| 247258_at | 1105 | P | 860.4 | P | 1080.4 | P | 995.4 | P | --- |
| 247259_at | 437.1 | P | 489.7 | P | 374.6 | P | 412.8 | P | CPR5 |
| 247260_at | 594.2 | P | 740.9 | P | 638.7 | P | 763.5 | P | --- |
| 247261_at | 854.1 | P | 1062.4 | P | 1277 | P | 1204.7 | P | --- |
| 247262_at | 709.8 | P | 899.8 | P | 616.6 | P | 740.9 | P | AtFAAH |
| 247263_at | 163 | P | 213.7 | P | 104.2 | P | 167.2 | P | --- |
| 247264_at | 10 | A | 39.1 | A | 22.1 | A | 20.4 | A | XND1 |
| 247265_at | 430.5 | P | 559 | P | 408.3 | P | 808.3 | P | CPuORF22 |
| 247266_at | 972.5 | P | 1563 | P | 731.3 | P | 1089.5 | P | XYL4 |
| 247267_at | 3216 | P | 2875.1 | P | 3140.8 | P | 3087.4 | P | RPS28 |
| 247268_at | 951.4 | P | 721.2 | P | 1077.4 | P | 961.6 | P | --- |
| 247269_at | 3.5 | A | 10.3 | A | 6.9 | A | 7.8 | A | AOX2 |
| 247270_at | 647.7 | P | 493.4 | P | 615.6 | P | 452.5 | P | --- |
| 247271_at | 1551.9 | P | 1444 | P | 1359.7 | P | 1345.2 | P | --- |
| 247272_at | 2218.1 | P | 1910.9 | P | 2294.4 | P | 1953.6 | P | --- |
| 247273_at | 322.1 | P | 421 | P | 328.3 | P | 362.7 | P | CPuORF40 |
| 247274_at | 135.8 | P | 151.8 | P | 143.2 | P | 167.8 | P | --- |
| 247275_at | 993.4 | P | 1061.6 | P | 907.4 | P | 880.3 | P | BETA-UP |
| 247276_at | 207.4 | P | 151 | P | 200.2 | P | 160.7 | P | HEN4 |
| 247277_at | 518.5 | P | 478.7 | P | 555.2 | P | 482.8 | P | --- |
| 247278_at | 275.6 | P | 380 | P | 407.7 | P | 374.9 | P | --- |
| 247279_at | 1564 | P | 1662.5 | P | 1380.3 | P | 1903.1 | P | AGP1 |
| 247280_at | 9686.9 | P | 2891.2 | P | 8427 | P | 3647.9 | P | EXL2 |
| 247281_at | 8085.2 | P | 8464.8 | P | 6601.7 | P | 7887.1 | P | FKBP12 |
| 247282_at | 578.6 | P | 490.8 | P | 467.2 | P | 383.6 | P | AtMC3 |
| 247283_at | 3110.1 | P | 1668.1 | P | 1830.4 | P | 1803.5 | P | --- |
| 247284_at | 135.3 | P | 357.8 | P | 111 | P | 283.5 | P | OPT4 |
| 247285_at | 10231 | P | 10573.8 | P | 6978.5 | P | 10530.4 | P | --- |
| 247286_at | 961.1 | P | 900.7 | P | 1083.1 | P | 938.1 | P | DiT2.2 |
| 247287_at | 398.2 | P | 155.1 | P | 215.9 | P | 235.2 | P | --- |
| 247288_at | 334.4 | P | 616.9 | P | 284 | P | 585.5 | P | NPH3 |
| 247289_at | 2348.6 | P | 2386.8 | P | 2412 | P | 2574.8 | P | DIT2.1 |
| 247290_at | 48.3 | A | 53.7 | A | 31.9 | A | 58.2 | A | --- |
| 247291_at | 300.7 | P | 342.8 | P | 328.2 | P | 348.8 | P | --- |
| 247292_at | 51.4 | P | 59 | P | 52.8 | P | 42.2 | A | --- |
| 247293_at | 164.5 | P | 172.4 | P | 64.7 | A | 101.1 | P | --- |
| 247294_at | 61.6 | A | 58.5 | A | 68.2 | A | 62 | A | XRCC2 |

| | | | | | | | | | |
|---|---|---|---|---|---|---|---|---|---|
| 247295_at | 405.2 | P | 325.1 | P | 279.9 | P | 339.6 | P | --- |
| 247296_at | 284 | P | 325.8 | P | 352.7 | P | 362.2 | P | --- |
| 247297_at | 7.4 | A | 66.3 | A | 78 | A | 46.4 | M | --- |
| 247298_at | 1334.8 | P | 1379.1 | P | 1304.3 | P | 1346.2 | P | RSW3 |
| 247299_at | 21.8 | A | 35.7 | P | 38.9 | A | 34 | A | --- |
| 247300_at | 390.1 | P | 409.9 | P | 304.5 | P | 318.9 | P | FCLY |
| 247301_at | 59.5 | P | 74.4 | P | 71.1 | M | 122.1 | P | --- |
| 247302_at | 495.4 | P | 467.9 | P | 387.4 | P | 382.8 | P | VPS20.1 |
| 247303_at | 1332.3 | P | 1382.6 | P | 1348.3 | P | 1123.9 | P | ATHDH |
| 247304_at | 482.7 | P | 456.7 | P | 399.5 | P | 394 | P | AAP4 |
| 247305_at | 142.4 | P | 118.4 | P | 80.6 | P | 76.2 | P | --- |
| 247306_at | 565.2 | P | 575.6 | P | 580.6 | P | 614.9 | P | PP7 |
| 247307_at | 1832.4 | P | 2264.6 | P | 1829.1 | P | 2306.1 | P | --- |
| 247308_at | 85.3 | P | 72.3 | P | 85.3 | P | 82.7 | P | --- |
| 247309_at | 374.4 | P | 424.3 | P | 441.8 | P | 440.9 | P | --- |
| 247310_at | 66.1 | P | 123.2 | P | 115.8 | P | 135.2 | P | CHR24 |
| 247311_at | 148.9 | P | 140.7 | P | 191.2 | P | 165.7 | P | EMB2780 |
| 247312_at | 206.9 | P | 116.4 | P | 125.8 | P | 114.3 | P | --- |
| 247313_at | 3117.4 | P | 2313.2 | P | 3669.5 | P | 2348.6 | P | SAL1 |
| 247314_at | 67.2 | P | 241.1 | P | 54.5 | P | 132.3 | P | SAL2 |
| 247315_at | 191.6 | P | 177.1 | P | 191.6 | P | 166 | P | --- |
| 247316_at | 1006.4 | P | 1412.2 | P | 1209 | P | 1287.2 | P | --- |
| 247317_at | 89.6 | P | 74.8 | P | 85.9 | P | 78.2 | A | anac103 |
| 247318_at | 157.2 | P | 200.5 | P | 203.4 | P | 150.1 | P | --- |
| 247319_at | 634.5 | P | 478.1 | P | 589.8 | P | 580.5 | P | ERS |
| 247320_at | 9443 | P | 9492.9 | P | 7130.7 | P | 8700.8 | P | PSAN |
| 247321_s_at | 432.5 | P | 492.1 | P | 341.2 | P | 350.4 | P | PI-4KBETA2 |
| 247322_at | 297.2 | P | 339 | P | 332.3 | P | 322.8 | P | --- |
| 247323_at | 277.9 | P | 678.7 | P | 231.8 | P | 496.5 | P | --- |
| 247324_at | 19.7 | A | 1.3 | A | 2.1 | A | 1.7 | A | --- |
| 247325_at | 717.1 | P | 570.3 | P | 527.8 | P | 594.2 | P | SC35 |
| 247326_at | 227.8 | P | 265.5 | P | 235.2 | P | 296.8 | P | --- |
| 247327_at | 1077.4 | P | 1381.6 | P | 742.1 | P | 1228.9 | P | --- |
| 247328_at | 2474.3 | P | 2118.7 | P | 1895.6 | P | 2346.4 | P | --- |
| 247329_at | 169.3 | P | 177.4 | P | 172.4 | P | 181.1 | P | --- |
| 247330_at | 2671.2 | P | 2352.8 | P | 2120.3 | P | 2316.2 | P | GAMMA CAL1 |
| 247331_at | 1029.3 | P | 795.2 | P | 1096.5 | P | 798.6 | P | FP3 |
| 247332_at | 208.6 | P | 277.3 | P | 286.7 | P | 289.1 | P | --- |
| 247333_at | 1.9 | A | 36.3 | P | 18.9 | A | 43.5 | P | FLS5 |
| 247334_at | 324.6 | P | 336.9 | P | 243.1 | P | 318 | P | CDKE;1 |
| 247335_at | 196.6 | P | 166.2 | P | 243 | P | 181.6 | P | --- |
| 247336_at | 307.5 | P | 336.3 | P | 317.5 | P | 328.5 | P | --- |
| 247337_at | 19.6 | A | 37.8 | A | 12.3 | A | 16 | A | PDF2.5 |
| 247338_at | 915.6 | P | 833.8 | P | 916.4 | P | 777.9 | P | --- |
| 247339_at | 149.3 | P | 165 | P | 184.5 | P | 119 | P | --- |
| 247340_at | 169.9 | P | 267.3 | P | 216.4 | P | 255.5 | P | --- |
| 247341_at | 32.2 | A | 3.1 | A | 6.9 | A | 33.3 | A | --- |
| 247342_at | 4.4 | A | 3.3 | A | 7.7 | A | 1.4 | A | --- |
| 247343_at | 18.8 | A | 10.3 | A | 24.3 | A | 5.5 | A | --- |
| 247344_at | 37.6 | A | 34.3 | A | 54.1 | M | 33.2 | A | --- |
| 247345_at | 293.9 | P | 239.3 | P | 316.5 | P | 262 | P | --- |
| 247346_at | 172.7 | P | 122.6 | P | 188.3 | P | 184.8 | P | ATDGK2 |
| 247347_at | 596.2 | P | 1268 | P | 834.1 | P | 1259.5 | P | SHA1 |
| 247348_at | 536.3 | P | 537.5 | P | 439.2 | P | 521.2 | P | BGAL10 |
| 247349_at | 38.3 | A | 55.2 | A | 34.6 | A | 50.7 | A | --- |
| 247350_at | 144.3 | P | 157.9 | P | 192.4 | P | 164.4 | P | --- |
| 247351_at | 1905.3 | P | 1966.6 | P | 1599.8 | P | 2186.1 | P | ANAC102 |
| 247352_at | 49.4 | P | 144.9 | P | 89 | M | 97.7 | P | SNRK2.5 |
| 247353_at | 591.3 | P | 606.5 | P | 559 | P | 546.5 | P | --- |
| 247354_at | 31.3 | A | 81.9 | A | 80.5 | A | 67.5 | A | FLS3 |
| 247355_at | 1280.7 | P | 1352.5 | P | 1169.4 | P | 1323.2 | P | SPT42 |
| 247356_at | 848.1 | P | 760.2 | P | 553.2 | P | 662.4 | P | MUM2 |
| 247357_at | 212.7 | P | 245.2 | P | 297.9 | P | 325.2 | P | --- |
| 247358_at | 21.3 | A | 51.7 | P | 42.2 | A | 41.5 | P | FLS2 |
| 247359_at | 34.4 | A | 91.7 | P | 52.1 | P | 77.8 | P | --- |
| 247360_at | 100.1 | P | 23.8 | A | 64.3 | A | 44.2 | A | CYP94B1 |
| 247361_at | 119.7 | P | 157.6 | P | 122.2 | P | 167.1 | P | --- |
| 247362_at | 104.6 | P | 194.2 | P | 259.4 | P | 230.3 | P | ATPAP29 |
| 247363_at | 259.5 | P | 260.6 | P | 289.9 | P | 273.5 | P | --- |
| 247364_at | 415.1 | P | 364.7 | P | 478.2 | P | 434.2 | P | --- |
| 247365_at | 261.6 | P | 303.8 | P | 283.2 | P | 285.6 | P | --- |
| 247366_at | 252.7 | P | 240.5 | P | 265.9 | P | 261.6 | P | --- |
| 247367_at | 330.6 | P | 202.1 | P | 321.7 | P | 229.1 | P | --- |
| 247368_at | 155.7 | P | 240.5 | P | 239.6 | P | 213.3 | P | --- |
| 247369_at | 15.1 | A | 1.2 | A | 9 | A | 2 | A | --- |
| 247370_at | 98.6 | P | 142.5 | P | 92.3 | P | 102 | P | --- |
| 247371_at | 415.4 | P | 401.4 | P | 467.4 | P | 508.2 | P | HDA6 |
| 247372_at | 237.6 | P | 231.4 | P | 289.2 | P | 214.3 | P | --- |
| 247373_at | 386.6 | P | 441.5 | P | 406.3 | P | 347.9 | P | --- |
| 247374_at | 821.8 | P | 762.6 | P | 652.7 | P | 704.1 | P | --- |
| 247375_at | 175.9 | P | 169.7 | P | 203.1 | P | 181 | P | --- |
| 247376_at | 3981.1 | P | 4758.3 | P | 4575 | P | 4741.1 | P | NDPK2 |
| 247377_at | 475.9 | P | 1507.3 | P | 429.6 | P | 1282.9 | P | --- |
| 247378_at | 369.9 | P | 328 | P | 313.3 | P | 352.5 | P | --- |
| 247379_at | 31 | P | 33.9 | A | 59.1 | P | 35.2 | A | --- |
| 247380_at | 326.6 | P | 357.7 | P | 334.6 | P | 383.8 | P | --- |
| 247381_at | 3 | A | 2.2 | A | 6.4 | A | 2.2 | A | --- |
| 247382_at | 1930.5 | P | 2325.9 | P | 2422.7 | P | 2451.6 | P | ADK1 |
| 247383_at | 405.3 | P | 626.7 | P | 331 | P | 543.6 | P | --- |
| 247384_at | 745.4 | P | 794.2 | P | 850.8 | P | 663.6 | P | --- |
| 247385_at | 453.7 | P | 769 | P | 660 | P | 726 | P | emb2746 |
| 247386_at | 769.3 | P | 982.3 | P | 1025.5 | P | 1023.9 | P | emb2746 |
| 247387_at | 126.4 | A | 184.2 | P | 319.2 | P | 231.2 | P | --- |
| 247388_s_at | 103.3 | P | 135.5 | P | 124.3 | P | 84.4 | P | NF-YC1 |
| 247389_at | 392.6 | P | 256.9 | P | 387.3 | P | 309.5 | P | --- |
| 247390_at | 214.4 | P | 224.4 | P | 195.3 | P | 177.2 | P | --- |
| 247391_at | 4.2 | A | 37.6 | A | 6.1 | A | 50.8 | A | --- |
| 247392_at | 1809.1 | P | 2026.9 | P | 2248.4 | P | 2116.9 | P | GSA1 |
| 247393_at | 124 | M | 134.4 | P | 85 | A | 129.8 | P | --- |
| 247394_at | 143.8 | P | 176.4 | A | 103.1 | P | 144.6 | M | --- |
| 247395_at | 404.7 | P | 356.1 | P | 258.2 | P | 298.3 | P | --- |
| 247396_at | 572.8 | P | 655.6 | P | 499.4 | P | 596 | P | --- |
| 247397_at | 130 | P | 144.9 | P | 173.6 | P | 90.5 | P | --- |
| 247398_at | 284.1 | P | 285.3 | P | 286.7 | P | 242.8 | P | --- |
| 247399_at | 169.1 | P | 112.3 | P | 151.2 | P | 129.2 | P | --- |
| 247400_at | 407 | P | 500.9 | P | 470.5 | P | 526.3 | P | --- |
| 247401_at | 2258.7 | P | 2451.1 | P | 2292 | P | 2691.6 | P | DXR |
| 247402_at | 1.5 | A | 0.9 | A | 0.3 | A | 5.5 | A | --- |
| 247403_at | 1999.8 | P | 1633 | P | 1404.9 | P | 1712.4 | P | --- |

| | | | | | | | | | |
|---|---|---|---|---|---|---|---|---|---|
| 247404_at | 1408.6 | P | 1646.6 | P | 1670.9 | P | 1664.8 | P | --- |
| 247405_at | 552.6 | P | 616.6 | P | 689.8 | P | 642.1 | P | ARAC10 |
| 247406_at | 202.7 | P | 318.2 | P | 218.1 | P | 258 | P | ARR6 |
| 247407_at | 299.2 | P | 392.3 | P | 350.2 | P | 355.1 | P | --- |
| 247408_at | 68.6 | P | 104.6 | P | 98.2 | P | 124.5 | P | --- |
| 247409_at | 809.5 | P | 653.4 | P | 592.4 | P | 613.4 | P | --- |
| 247410_at | 213.9 | P | 160.4 | P | 160.6 | P | 196.5 | P | emb1692 |
| 247411_at | 624 | P | 380.6 | P | 610.8 | P | 451.8 | P | --- |
| 247412_at | 185.2 | P | 148.5 | P | 144.1 | P | 144.4 | P | --- |
| 247413_at | 81.4 | P | 111 | P | 85.9 | P | 89 | P | --- |
| 247414_at | 729.9 | P | 1003.8 | P | 934.9 | P | 1159.8 | P | EMB2759 |
| 247415_at | 264.1 | P | 289.4 | P | 192.4 | P | 269.7 | P | --- |
| 247416_at | 22.1 | A | 19 | A | 5.8 | A | 25.1 | A | --- |
| 247417_at | 241 | P | 281.1 | P | 240.5 | P | 255.1 | P | --- |
| 247418_at | 629.5 | P | 747.2 | P | 475.7 | P | 499.5 | P | --- |
| 247419_at | 153.6 | P | 187.7 | P | 145.8 | P | 175.3 | P | --- |
| 247420_at | 109.1 | P | 154.9 | P | 108.3 | P | 144.3 | P | --- |
| 247421_at | 36 | A | 35.4 | A | 31.4 | A | 16.4 | A | --- |
| 247422_at | 1128.7 | P | 1267.1 | P | 1264.9 | P | 1335.7 | P | PEX14 |
| 247423_at | 21.7 | A | 32.2 | P | 63.9 | P | 35.2 | A | --- |
| 247424_at | 54.7 | M | 32.9 | A | 100.8 | P | 62.1 | A | AtVEX1 |
| 247425_at | 117.4 | P | 201.6 | P | 120.9 | P | 241.7 | P | --- |
| 247426_at | 467.1 | P | 558.3 | P | 351.7 | P | 427.7 | P | --- |
| 247427_at | 270.9 | P | 332 | P | 350 | P | 414.6 | P | --- |
| 247428_at | 416.5 | P | 377.9 | P | 358.6 | P | 500.9 | P | --- |
| 247429_at | 404.2 | P | 386.6 | P | 389.2 | P | 358 | P | --- |
| 247430_at | 697.1 | P | 967.6 | P | 705.2 | P | 894.3 | P | --- |
| 247431_at | 114 | P | 172.7 | P | 65.5 | M | 128.9 | P | SRO5 |
| 247432_at | 123.5 | P | 134.8 | P | 171.8 | P | 144.6 | P | ATEB1B |
| 247433_at | 1395.9 | P | 894.3 | P | 1384.5 | P | 1028.2 | P | UBC3 |
| 247434_at | 1190.8 | P | 1156.4 | P | 995.8 | P | 999.1 | P | --- |
| 247435_at | 45.8 | A | 31.7 | A | 35.5 | A | 6.4 | A | ATGSTU9 |
| 247436_at | 1446.2 | P | 1215.8 | P | 1713.3 | P | 1350.6 | P | ALDH12A1 |
| 247437_at | 6.3 | A | 3.8 | A | 9.9 | A | 1.3 | A | HVA22B |
| 247438_at | 402.9 | P | 433.1 | P | 293.1 | P | 353.4 | P | --- |
| 247439_at | 439.7 | P | 782.7 | P | 467.9 | P | 970.2 | P | AHA11 |
| 247440_at | 813.6 | P | 1377.2 | P | 1072.9 | P | 959.5 | P | --- |
| 247441_at | 92.6 | P | 198.9 | P | 166.8 | P | 177.4 | P | --- |
| 247442_s_at | 3318.8 | P | 4096.7 | P | 3260.1 | P | 4278.1 | P | TUB2 |
| 247443_at | 894.1 | P | 1078.4 | P | 946.4 | P | 1190.9 | P | --- |
| 247444_at | 136.3 | P | 217.8 | P | 167.9 | P | 215.6 | P | HIPL2 |
| 247445_at | 476.3 | P | 417.1 | P | 427.5 | P | 484.5 | P | --- |
| 247446_at | 716.6 | P | 623.7 | P | 722.3 | P | 640.5 | P | --- |
| 247447_at | 33.9 | A | 66.9 | P | 52.2 | P | 42.6 | P | --- |
| 247448_at | 10.5 | A | 69.1 | M | 34.9 | A | 32.6 | A | --- |
| 247449_at | 686 | P | 516 | P | 673.3 | P | 573.3 | P | --- |
| 247450_at | 4123 | P | 5483.6 | P | 4423.8 | P | 5145.3 | P | --- |
| 247451_at | 16 | A | 10.9 | A | 3.3 | A | 5.7 | A | ATMYB99 |
| 247452_at | 131.5 | P | 289.2 | P | 163 | P | 212.2 | P | CDF1 |
| 247453_at | 769.6 | P | 421.3 | P | 740.5 | P | 447.4 | P | --- |
| 247454_at | 1067.3 | P | 593.1 | P | 1151.3 | P | 742.9 | P | --- |
| 247455_at | 104.1 | P | 147.1 | P | 128.9 | P | 155 | P | MYB96 |
| 247456_at | 51.2 | P | 46.1 | P | 40.3 | P | 44.5 | P | AtZIP12 |
| 247457_at | 60.3 | P | 76.3 | P | 89.4 | A | 83.8 | P | --- |
| 247458_at | 10.9 | A | 80.4 | A | 53.1 | A | 66.6 | A | AtCXE20 |
| 247459_at | 1727.6 | P | 1325.8 | P | 1552.5 | P | 1478.7 | P | --- |
| 247460_at | 1052.8 | P | 864.8 | P | 1291.5 | P | 934.9 | P | OXA1 |
| 247461_at | 80.6 | P | 72.6 | A | 122.6 | P | 98.1 | P | ATBAG2 |
| 247462_at | 4 | A | 31.4 | A | 2.7 | A | 4.9 | A | --- |
| 247463_at | 226.6 | P | 311.3 | P | 440 | P | 617 | P | --- |
| 247464_at | 305.3 | P | 340 | P | 248.8 | P | 272.6 | P | IQD23 |
| 247465_at | 3058.7 | P | 2060.7 | P | 3614.2 | P | 2423.3 | P | PRH75 |
| 247466_at | 263 | P | 222.7 | P | 298.9 | P | 209.6 | P | --- |
| 247467_at | 228.9 | P | 236.8 | P | 271.8 | P | 198.9 | P | --- |
| 247468_at | 1309 | P | 1202.9 | P | 1610.3 | P | 1148.2 | P | ARF2 |
| 247469_at | 2.8 | A | 1.5 | A | 16.1 | A | 2.7 | A | AGL42 |
| 247470_at | 383.6 | P | 453.4 | P | 338 | P | 580.1 | P | --- |
| 247471_at | 90.7 | P | 75.1 | P | 126.9 | P | 115 | P | ERL1 |
| 247472_at | 1.5 | A | 2.1 | A | 4.7 | A | 0.4 | A | MAP65-9 |
| 247473_at | 45.1 | P | 49.7 | P | 52.2 | P | 24.9 | A | --- |
| 247474_at | 14.9 | A | 13.5 | A | 3.3 | A | 14.4 | A | --- |
| 247475_at | 33.5 | A | 24.4 | A | 38.5 | P | 39.6 | P | IRE |
| 247476_at | 4.2 | A | 4.1 | A | 5 | A | 7.7 | A | --- |
| 247477_at | 42.3 | A | 22 | A | 51.5 | A | 40.2 | A | --- |
| 247478_at | 97.6 | P | 470.4 | P | 197.9 | P | 599.7 | P | --- |
| 247479_at | 34.4 | A | 34.3 | P | 69.8 | P | 34.3 | P | ANAC101 |
| 247480_at | 2071.2 | P | 2522.5 | P | 2418.8 | P | 2291.1 | P | ATBAG7 |
| 247481_at | 5.7 | A | 9.9 | A | 4.5 | A | 5.5 | A | --- |
| 247482_at | 119.4 | P | 141 | P | 135.6 | P | 104.9 | P | SMC2 |
| 247483_at | 30.1 | A | 33.4 | A | 47.1 | A | 56.4 | A | --- |
| 247484_at | 2.7 | A | 7.8 | A | 0.8 | A | 0.6 | A | --- |
| 247485_at | 1 | A | 1.5 | A | 4.5 | A | 1.2 | A | ARR23 |
| 247486_at | 522.7 | P | 572.8 | P | 613 | P | 636.1 | P | --- |
| 247487_at | 102 | P | 123 | P | 120.2 | P | 84.7 | P | --- |
| 247488_at | 8977.7 | P | 6111 | P | 7861.3 | P | 8218.5 | P | --- |
| 247489_at | 260.2 | P | 330 | P | 295.6 | P | 270.3 | P | --- |
| 247490_at | 17.8 | A | 36.9 | A | 46.6 | A | 16 | A | LFY |
| 247491_at | 464.6 | P | 226.7 | P | 426.8 | P | 300 | P | --- |
| 247492_at | 78.7 | P | 52.4 | A | 54 | A | 35.9 | A | --- |
| 247493_at | 603.4 | P | 714.7 | P | 667.8 | P | 573.5 | P | --- |
| 247494_at | 3901.5 | P | 4166.7 | P | 4199.3 | P | 4485.5 | P | CNX1 |
| 247495_at | 479.2 | P | 344.9 | P | 467.4 | P | 411 | P | IPK2BETA |
| 247496_at | 530.7 | P | 568.9 | P | 624.9 | P | 594.7 | P | GUT1 |
| 247497_at | 602.9 | P | 456.2 | P | 728.3 | P | 447.9 | P | PPAN |
| 247498_at | 460.4 | P | 426.4 | P | 421.7 | P | 485.4 | P | --- |
| 247499_at | 43.5 | P | 27.3 | M | 35.2 | A | 61 | P | --- |
| 247500_at | 175.8 | P | 168.9 | P | 178 | P | 224.3 | P | --- |
| 247501_at | 56.2 | A | 49.9 | P | 51.7 | A | 51.5 | A | --- |
| 247502_at | 66.2 | P | 55.6 | P | 44.7 | P | 64 | P | --- |
| 247503_at | 30.5 | A | 39.6 | P | 32 | A | 46.2 | A | AGD1 |
| 247504_at | 125.8 | P | 187.9 | P | 131.2 | P | 119.1 | P | --- |
| 247505_at | 830.4 | P | 865.5 | P | 850.6 | P | 801.8 | P | --- |
| 247506_at | 769.9 | P | 818.3 | P | 746.6 | P | 864.5 | P | AML1 |
| 247507_at | 224 | P | 235.3 | P | 237.4 | P | 212 | P | APO3 |
| 247508_at | 853.6 | P | 740.2 | P | 919.9 | P | 812.7 | P | ARF2 |
| 247509_at | 311.1 | P | 221.7 | P | 327.2 | P | 239.7 | P | HSFB2A |
| 247510_at | 120 | P | 130.6 | P | 185.5 | P | 139.5 | P | --- |
| 247511_at | 30 | A | 6.3 | A | 7 | A | 19.2 | A | --- |
| 247512_at | 0.9 | A | 1.6 | A | 11.3 | A | 1.2 | A | --- |

| | | | | | | | | | |
|---|---|---|---|---|---|---|---|---|---|
| 247513_at | 959.8 | P | 1036.3 | P | 878.9 | P | 909.2 | P | --- |
| 247514_at | 298.1 | P | 406.5 | P | 300.1 | P | 331 | P | PMSR1 |
| 247515_at | 10.5 | A | 42.2 | P | 11.7 | A | 3.3 | A | ATH14 |
| 247516_at | 5.7 | A | 6.4 | A | 5.9 | A | 5 | A | --- |
| 247517_at | 1831.3 | P | 1875.8 | P | 1518.1 | P | 1799.5 | P | --- |
| 247518_at | 104.4 | P | 93.1 | P | 63.1 | P | 78.9 | P | --- |
| 247519_at | 144.8 | P | 82 | P | 133.6 | P | 79.4 | P | ANAC100 |
| 247520_at | 512.3 | P | 502.7 | P | 467 | P | 526.1 | P | --- |
| 247521_at | 270.9 | P | 232.2 | P | 246.4 | P | 220.2 | P | --- |
| 247522_at | 124.1 | P | 149.9 | P | 119.9 | P | 158.8 | P | --- |
| 247523_at | 7598.5 | P | 8292.8 | P | 7883 | P | 8816.9 | P | RPE |
| 247524_at | 234.1 | P | 128.2 | P | 235.2 | P | 109.3 | P | ACHT5 |
| 247525_at | 1365.7 | P | 1136.2 | P | 1364.1 | P | 1249.9 | P | TOC1 |
| 247526_at | 23.8 | A | 22.1 | P | 1.8 | A | 21.6 | P | --- |
| 247527_at | 132.3 | P | 212.2 | P | 237.7 | P | 297.3 | P | --- |
| 247528_at | 80.2 | P | 85.1 | P | 47.4 | P | 73 | P | MIM |
| 247529_at | 1151.8 | P | 706.6 | P | 965 | P | 785.2 | P | --- |
| 247530_at | 116.8 | P | 111.1 | P | 104.5 | P | 134.6 | P | --- |
| 247531_at | 2.9 | A | 1.1 | A | 1.9 | A | 1.1 | A | --- |
| 247532_at | 89.9 | P | 154 | P | 117.6 | P | 104.7 | P | --- |
| 247533_at | 312.6 | P | 226.7 | P | 200.4 | P | 238.4 | P | --- |
| 247534_at | 933 | P | 860.6 | P | 816.1 | P | 931.7 | P | PFK4 |
| 247535_at | 26.3 | A | 33.6 | A | 12.5 | A | 10.8 | A | --- |
| 247536_at | 3.9 | A | 8 | A | 2.6 | A | 1.6 | A | CYCP4;2 |
| 247537_s_at | 31.6 | P | 15.1 | A | 41.3 | P | 29.4 | P | ATH15 |
| 247538_at | 37.1 | P | 42.2 | P | 47.3 | P | 33.5 | P | ATH16 |
| 247539_at | 0.9 | A | 1.5 | A | 3.3 | A | 3.4 | A | --- |
| 247540_at | 757.1 | P | 1124 | P | 956.3 | P | 919.2 | P | --- |
| 247541_at | 485.1 | P | 818.2 | P | 805.5 | P | 863.9 | P | --- |
| 247542_at | 676.6 | P | 865 | P | 827.5 | P | 788 | P | ATG3 |
| 247543_at | 172.5 | P | 353.8 | P | 206.9 | P | 308.2 | P | --- |
| 247544_at | 1292.5 | P | 1411.4 | P | 1466 | P | 1451.5 | P | --- |
| 247545_at | 1045.1 | P | 1118.9 | P | 947.4 | P | 1031.1 | P | --- |
| 247546_at | 1.6 | A | 17.8 | A | 33.6 | A | 6.7 | A | --- |
| 247547_at | 101.6 | P | 125.2 | P | 157.3 | P | 105.1 | P | --- |
| 247548_at | 59.3 | A | 13.3 | A | 72.1 | A | 42.2 | M | --- |
| 247549_at | 777.7 | P | 938.6 | P | 1143.5 | P | 936.8 | P | MYB28 |
| 247550_at | 181.1 | P | 177.8 | P | 220.3 | P | 156.4 | P | --- |
| 247551_at | 479.9 | P | 563.8 | P | 551.4 | P | 586.2 | P | --- |
| 247552_at | 2076.7 | P | 3092.9 | P | 2155.6 | P | 2956.4 | P | COB |
| 247553_at | 7.2 | A | 45.5 | A | 17.3 | A | 30.7 | A | AGL8 |
| 247554_at | 293.2 | P | 567 | P | 260.4 | P | 494.6 | P | ATEXO70E2 |
| 247555_at | 2711.3 | P | 2409.4 | P | 3340.7 | P | 2573.2 | P | ECT3 |
| 247556_at | 101.7 | P | 101.1 | P | 102.1 | P | 126.5 | P | --- |
| 247557_at | 0.5 | A | 3 | A | 2.1 | A | 2.4 | A | --- |
| 247558_at | 361.8 | P | 459 | P | 385.5 | P | 387.6 | P | HDA05 |
| 247559_at | 37.8 | A | 40.6 | A | 3.3 | A | 34.5 | A | HDA18 |
| 247560_at | 37.8 | A | 59 | A | 47.1 | A | 48.6 | A | --- |
| 247561_at | 15.6 | A | 20.2 | A | 35 | A | 15.9 | A | --- |
| 247562_at | 13.2 | A | 16.4 | A | 4.6 | A | 15.9 | A | --- |
| 247563_at | 788.8 | P | 1067.1 | P | 1175.2 | P | 1309.8 | P | PDCB1 |
| 247564_at | 782 | P | 606.6 | P | 558.8 | P | 606.8 | P | --- |
| 247565_at | 418.8 | P | 497.3 | P | 322.5 | P | 487.3 | P | VIP4 |
| 247566_at | 5702.4 | P | 5208.7 | P | 5711.9 | P | 5052 | P | --- |
| 247567_at | 93.8 | P | 100.2 | P | 88.2 | P | 103.5 | P | --- |
| 247568_at | 30.9 | A | 21.5 | A | 19.3 | A | 15.1 | A | --- |
| 247569_at | 1243.7 | P | 1345.5 | P | 1031.3 | P | 1360.6 | P | --- |
| 247570_at | 92.7 | P | 88.9 | P | 78.6 | P | 64.5 | P | AtGUS1 |
| 247571_at | 1239.3 | P | 1320.7 | P | 820.3 | P | 1101.9 | P | SNAP33 |
| 247572_at | 887.9 | P | 665.4 | P | 915.5 | P | 809.4 | P | --- |
| 247573_at | 5.3 | A | 87.8 | P | 30.8 | A | 89.5 | P | AACT1 |
| 247574_at | 620.7 | P | 883.6 | P | 618.1 | P | 729.4 | P | --- |
| 247575_at | 819 | P | 628.5 | P | 866.7 | P | 724.9 | P | GR-RBP3 |
| 247576_at | 3.2 | A | 34.4 | A | 36.8 | A | 7.8 | A | --- |
| 247577_at | 186.7 | P | 293.6 | P | 182.4 | P | 342.2 | P | --- |
| 247578_at | 92.9 | P | 125.7 | P | 99.7 | P | 136.2 | P | --- |
| 247579_at | 13.3 | A | 18 | A | 39.1 | A | 24.8 | P | CYP89A3 |
| 247580_at | 266.6 | P | 344.2 | P | 227 | P | 261.4 | P | --- |
| 247581_at | 39 | A | 42.3 | A | 47.8 | A | 37.3 | A | --- |
| 247582_at | 32.1 | A | 19.2 | A | 22.6 | A | 5.6 | A | --- |
| 247583_at | 747.4 | P | 785 | P | 686.4 | P | 819.8 | P | --- |
| 247584_at | 4685.4 | P | 4024.4 | P | 3838.1 | P | 4389.2 | P | --- |
| 247585_at | 367.7 | P | 619.9 | P | 366 | P | 431.2 | P | --- |
| 247586_at | 62.6 | A | 59.1 | A | 79.6 | A | 48.9 | A | PIP2;4 |
| 247587_at | 1335.1 | P | 1352.9 | P | 1387.2 | P | 1272.6 | P | --- |
| 247588_at | 2230.9 | P | 2185.8 | P | 1920.6 | P | 1947.5 | P | ATPDIL1-4 |
| 247589_at | 360.5 | P | 538.2 | P | 537 | P | 528.4 | P | REV |
| 247590_at | 44 | P | 50.8 | P | 62 | A | 70.4 | P | --- |
| 247591_at | 5.9 | A | 6.2 | A | 9.2 | A | 12.1 | A | NRT2.4 |
| 247592_at | 43.4 | A | 5.1 | A | 23.1 | A | 16.3 | A | NRT2.3 |
| 247593_at | 3904.1 | P | 2957.2 | P | 3492.5 | P | 3784.6 | P | GCN1 |
| 247594_at | 569.4 | P | 437.9 | P | 487.1 | P | 358.3 | P | --- |
| 247595_at | 164.3 | P | 217.4 | P | 213.7 | P | 210 | P | --- |
| 247596_at | 130 | A | 258.3 | P | 206.7 | P | 246.3 | P | --- |
| 247597_at | 325.8 | P | 355.6 | P | 208.7 | P | 347.1 | P | AtRABA1f |
| 247598_at | 190.9 | P | 196.8 | P | 305.3 | P | 203.8 | P | --- |
| 247599_at | 158 | P | 148.7 | P | 163.1 | P | 122.2 | P | --- |
| 247600_at | 882 | P | 470.5 | P | 823 | P | 555.9 | P | MYB34 |
| 247601_at | 591.1 | P | 757.9 | P | 468.7 | P | 608.2 | P | OBP4 |
| 247602_at | 68.3 | P | 204.8 | P | 60.4 | P | 188.7 | P | RLK1 |
| 247603_at | 94.4 | P | 128.6 | P | 117.7 | P | 108.4 | P | --- |
| 247604_at | 199.7 | P | 294.6 | P | 119.7 | P | 227.3 | P | COBL5 |
| 247605_at | 141.9 | P | 147 | P | 185.5 | P | 169.5 | P | TCP5 |
| 247606_at | 343 | P | 341.4 | P | 330.3 | P | 344.6 | P | --- |
| 247607_at | 771.1 | P | 671.7 | P | 850.2 | P | 737.7 | P | --- |
| 247608_at | 814.3 | P | 739.2 | P | 869.6 | P | 743.2 | P | --- |
| 247609_at | 213.1 | P | 251.2 | P | 241.6 | P | 298.5 | P | --- |
| 247610_at | 99.3 | P | 95.5 | P | 64.7 | P | 89.5 | P | --- |
| 247611_at | 518.4 | P | 658.9 | P | 483.6 | P | 699.5 | P | --- |
| 247612_at | 127.3 | P | 109.5 | P | 134.3 | P | 124.7 | P | --- |
| 247613_at | 10.1 | A | 17 | A | 10.2 | A | 13.3 | A | --- |
| 247614_at | 2.8 | A | 3.3 | A | 4.8 | A | 12.4 | A | --- |
| 247615_at | 146.2 | P | 63.7 | P | 156.1 | P | 65.6 | P | --- |
| 247616_at | 60.4 | P | 18.8 | A | 69 | M | 51.6 | M | --- |
| 247617_at | 126.5 | P | 121.2 | P | 129.3 | P | 102.5 | P | --- |
| 247618_at | 12.9 | A | 34.2 | A | 3.5 | A | 5.9 | A | --- |
| 247619_at | 6.3 | A | 19.8 | A | 21.5 | A | 49.4 | A | --- |
| 247620_at | 8.7 | A | 7.3 | A | 6.8 | A | 9 | A | --- |
| 247621_at | 87.5 | P | 73.1 | P | 109.1 | P | 112.5 | P | --- |

| | | | | | | | | | |
|---|---|---|---|---|---|---|---|---|---|
| 247622_at | 30.1 | A | 4.3 | A | 8.3 | A | 20.8 | A | --- |
| 247623_at | 25.1 | A | 20.6 | A | 36.1 | A | 24.7 | A | --- |
| 247624_at | 1396 | P | 1232.6 | P | 1019.6 | P | 1132.1 | P | --- |
| 247625_at | 93.8 | P | 169.4 | P | 157.4 | P | 123.9 | P | --- |
| 247626_at | 346 | P | 158.7 | P | 368.6 | P | 165.6 | P | --- |
| 247627_at | 16357.9 | P | 12838.8 | P | 13186.3 | P | 15731.5 | P | AALP |
| 247628_at | 122.7 | P | 169.1 | P | 81.4 | P | 112.1 | P | --- |
| 247629_at | 315.6 | P | 205.5 | P | 175.8 | P | 196.9 | P | SIZ1 |
| 247630_at | 880.7 | P | 837.7 | P | 858.1 | P | 884.9 | P | SIZ1 |
| 247631_at | 18.2 | P | 24 | P | 26.2 | A | 21.7 | A | AGL62 |
| 247632_at | 1221.9 | P | 853.2 | P | 905.1 | P | 842.5 | P | --- |
| 247633_at | 42 | A | 58.9 | P | 69.3 | A | 42.2 | A | --- |
| 247634_at | 18.7 | A | 16.8 | A | 1.8 | A | 17.6 | A | --- |
| 247635_at | 484.6 | P | 544.5 | P | 510.1 | P | 600.9 | P | GRIK2 |
| 247636_at | 587.2 | P | 717.6 | P | 699.5 | P | 786.3 | P | --- |
| 247637_at | 4998.9 | P | 4753.7 | P | 4472.2 | P | 5077.7 | P | HDS |
| 247638_at | 106 | P | 218.3 | P | 148.4 | P | 221.2 | P | FLA12 |
| 247639_s_at | 0.3 | A | 0.7 | A | 0.6 | A | 0.5 | A | --- |
| 247640_at | 42.6 | A | 52.9 | A | 38.4 | A | 40.9 | A | --- |
| 247641_at | 1695.5 | P | 1342.8 | P | 1932.3 | P | 1726 | P | PDX2 |
| 247642_at | 348.3 | P | 279 | P | 286.1 | P | 270.3 | P | --- |
| 247643_at | 337.8 | P | 284.7 | P | 371.7 | P | 298.9 | P | ARF4 |
| 247644_s_at | 23332.3 | P | 23387.5 | P | 28838.2 | P | 25630.8 | P | --- |
| 247645_at | 30 | A | 7.2 | A | 28.9 | A | 18.3 | A | --- |
| 247646_at | 6.1 | A | 22.1 | A | 1.9 | A | 24.8 | A | --- |
| 247647_at | 28.1 | A | 34.2 | A | 41.5 | P | 40.5 | A | --- |
| 247648_at | 140 | M | 164.6 | P | 169.4 | P | 132.8 | P | LAC17 |
| 247649_at | 90.7 | P | 31.6 | P | 78.1 | P | 48.2 | P | --- |
| 247650_at | 594 | P | 671.4 | P | 469.7 | P | 657.5 | P | --- |
| 247651_at | 1769.2 | P | 2839 | P | 2102.4 | P | 2686.7 | P | HTA6 |
| 247652_at | 2949.6 | P | 2925.9 | P | 3204.5 | P | 2236.7 | P | HTB4 |
| 247653_at | 997 | P | 1069.3 | P | 896.9 | P | 1080 | P | --- |
| 247654_at | 4177.9 | P | 4009.2 | P | 5207.7 | P | 4525 | P | --- |
| 247655_at | 334.5 | P | 356.5 | P | 231.1 | P | 244.5 | P | RHL41 |
| 247656_at | 1855.2 | P | 2383.1 | P | 1905.5 | P | 2376 | P | ADF4 |
| 247657_at | 44.1 | P | 85.3 | P | 84.3 | P | 82.4 | P | --- |
| 247658_at | 4676.8 | P | 5562.1 | P | 4578.3 | P | 5368.2 | P | ADF3 |
| 247659_at | 204.6 | P | 160 | P | 207.7 | P | 128.3 | P | NRPC1 |
| 247660_at | 2.4 | A | 2 | A | 3.8 | A | 1.2 | A | --- |
| 247661_at | 1.3 | A | 3.3 | A | 15.8 | A | 18.4 | A | --- |
| 247662_at | 23 | A | 29.9 | A | 11.9 | A | 17.8 | A | --- |
| 247663_at | 0.8 | A | 1.4 | A | 2.6 | A | 0.9 | A | APUM18 |
| 247664_at | 255.3 | P | 279.3 | P | 249.7 | P | 297.3 | P | TOE2 |
| 247665_at | 2 | A | 21.9 | A | 3.1 | A | 2.9 | A | --- |
| 247666_at | 14.2 | A | 18.4 | A | 29.1 | A | 1.9 | A | --- |
| 247667_at | 32.2 | M | 85.2 | P | 92.4 | P | 68.8 | P | --- |
| 247668_at | 678.8 | P | 629.1 | P | 510.1 | P | 671 | P | APRR3 |
| 247669_at | 506.8 | P | 496.4 | P | 567.5 | P | 549.1 | P | --- |
| 247670_at | 197.3 | P | 158.8 | P | 172.7 | P | 158.4 | P | --- |
| 247671_at | 248.6 | P | 278.9 | P | 228.2 | P | 302.8 | P | --- |
| 247672_at | 21.1 | A | 48.1 | A | 39.4 | A | 30.1 | A | TET4 |
| 247673_at | 107.8 | P | 141.2 | P | 110.8 | P | 138 | P | SEN2 |
| 247674_at | 3 | A | 12.3 | A | 50.1 | A | 8.7 | A | --- |
| 247675_at | 33 | A | 50.7 | A | 138.7 | P | 75.3 | P | --- |
| 247676_at | 71.2 | P | 61.1 | P | 108.4 | P | 59.3 | P | --- |
| 247677_at | 1112.9 | P | 1258.9 | P | 1230.3 | P | 1177.8 | P | ORP3C |
| 247678_at | 32.5 | P | 46.5 | P | 66.7 | P | 50.1 | P | ZIP2 |
| 247679_at | 3262.4 | P | 3246 | P | 3178.7 | P | 3200.8 | P | --- |
| 247680_s_at | 27.1 | A | 15.6 | A | 16.8 | A | 9.5 | A | --- |
| 247681_at | 5.7 | A | 8.7 | A | 2.4 | A | 9.8 | A | --- |
| 247682_at | 48.2 | P | 25.2 | P | 6.2 | A | 56.9 | P | --- |
| 247683_at | 52.6 | A | 37.4 | A | 10.6 | A | 40.5 | A | --- |
| 247684_at | 29.6 | A | 28.8 | P | 7 | A | 38.7 | P | --- |
| 247685_at | 44.8 | A | 35.2 | A | 76.7 | A | 29 | A | --- |
| 247686_at | 90.4 | P | 66.6 | P | 63.7 | A | 58.9 | A | --- |
| 247687_at | 111.8 | P | 109.6 | P | 122.5 | P | 137.4 | P | --- |
| 247688_at | 9.2 | A | 17.7 | A | 6.5 | A | 12.3 | A | --- |
| 247689_at | 925.3 | P | 809.5 | P | 999.8 | P | 923.8 | P | --- |
| 247690_at | 269.7 | P | 214.2 | P | 281.2 | P | 257.3 | P | MBD7 |
| 247691_at | 68.4 | P | 81.8 | P | 25.1 | A | 78.1 | P | HSP18.2 |
| 247692_s_at | 2644.9 | P | 4415.6 | P | 3082.4 | P | 4620.8 | P | --- |
| 247693_at | 354 | P | 427.8 | P | 324 | P | 323.6 | P | ATEXO70H7 |
| 247694_at | 450 | P | 661.9 | P | 538.5 | P | 624.3 | P | --- |
| 247695_at | 501.1 | P | 439.4 | P | 274.8 | P | 449.5 | P | VIP2 |
| 247696_at | 203.1 | P | 721.8 | P | 264.8 | P | 434.2 | P | MYB59 |
| 247697_at | 26.5 | A | 18.7 | A | 37.5 | A | 38.5 | A | SBT5.4 |
| 247698_at | 536.9 | P | 405.6 | P | 655.7 | P | 474.8 | P | --- |
| 247699_at | 498.8 | P | 359.1 | P | 374 | P | 354 | P | --- |
| 247700_at | 21 | A | 6.6 | A | 1.9 | A | 3.4 | A | --- |
| 247701_at | 60.7 | P | 99.8 | P | 79.7 | P | 71.3 | P | --- |
| 247702_at | 815.9 | P | 1123.8 | P | 956.3 | P | 1157.2 | P | --- |
| 247703_at | 292.6 | P | 323.6 | P | 351.3 | P | 353.1 | P | SRR1 |
| 247704_at | 190.4 | P | 40.5 | P | 128.3 | P | 60.8 | P | RTFL5 |
| 247705_at | 312.4 | P | 280.9 | P | 394.9 | P | 307.1 | P | --- |
| 247706_at | 598.4 | P | 609.7 | P | 729.2 | P | 672.3 | P | --- |
| 247707_at | 242.4 | P | 283.7 | P | 216.9 | P | 231.6 | P | --- |
| 247708_at | 413.4 | P | 358.9 | P | 359.6 | P | 424.6 | P | --- |
| 247709_at | 1054.4 | P | 1616.4 | P | 1213.1 | P | 1545.2 | P | --- |
| 247710_at | 15.7 | A | 30.6 | A | 59.4 | P | 41.2 | A | --- |
| 247711_at | 3.6 | A | 3 | A | 3.8 | A | 2.4 | A | --- |
| 247712_at | 928.8 | P | 869 | P | 881.3 | P | 855.7 | P | UBC7 |
| 247713_at | 94.2 | P | 179.9 | P | 50.7 | A | 206.5 | P | --- |
| 247714_at | 42.5 | P | 44.6 | P | 60.7 | P | 31.1 | A | WOX2 |
| 247715_at | 64.9 | P | 68.3 | A | 93.7 | A | 64.7 | A | --- |
| 247716_at | 264.3 | P | 323.6 | P | 290.5 | P | 301.4 | P | --- |
| 247717_at | 3403.8 | P | 702 | P | 125.1 | P | 2026.1 | P | LTP3 |
| 247718_at | 4218.8 | P | 188.1 | P | 42.6 | A | 695 | P | LTP4 |
| 247719_at | 2.4 | A | 3.9 | A | 3 | A | 15.9 | A | --- |
| 247720_at | 844.1 | P | 880.2 | P | 696.9 | P | 927.6 | P | UXS3 |
| 247721_at | 1080.3 | P | 1236.1 | P | 1258.6 | P | 1183.1 | P | --- |
| 247722_at | 523.9 | P | 437.9 | P | 495.1 | P | 509 | P | ATRABA2D |
| 247723_at | 625.2 | P | 182.9 | P | 220.1 | P | 292.3 | P | --- |
| 247724_at | 194.9 | P | 205.1 | P | 226 | P | 234.7 | P | MBD6 |
| 247725_at | 364.2 | P | 415 | P | 376.3 | P | 409 | P | --- |
| 247726_at | 267.6 | P | 258.5 | P | 285 | P | 262.8 | P | TRP1 |
| 247727_at | 36.3 | A | 50.3 | A | 35.6 | A | 49.9 | A | --- |
| 247728_at | 19.3 | A | 0.7 | A | 2.3 | A | 7.4 | A | --- |
| 247729_at | 56.5 | P | 15.1 | A | 43.1 | A | 14.9 | A | --- |
| 247730_at | 14.4 | A | 1.3 | A | 6.6 | A | 3.2 | A | UGT76E1 |

| | | | | | | | | | |
|---|---|---|---|---|---|---|---|---|---|
| 247731_at | 39.1 | A | 36.3 | A | 48.7 | A | 13.5 | A | UGT76E2 |
| 247732_at | 82.4 | P | 55.9 | P | 77.8 | P | 62.7 | P | --- |
| 247733_at | 439.3 | P | 448.4 | P | 397 | P | 516.6 | P | --- |
| 247734_at | 401.4 | P | 283.8 | P | 320.1 | P | 346.5 | P | --- |
| 247735_at | 1268.2 | P | 768.6 | P | 1283.1 | P | 683.8 | P | ZEU1 |
| 247736_at | 22.1 | A | 12.6 | A | 40.3 | A | 6.5 | A | ACT4 |
| 247737_at | 4.6 | A | 16.7 | A | 8.6 | A | 17.4 | A | --- |
| 247738_at | 469.8 | P | 445.4 | P | 383.2 | P | 420.5 | P | --- |
| 247739_at | 113.4 | P | 64.2 | P | 117.6 | P | 68.1 | P | --- |
| 247740_at | 46.4 | P | 99.2 | P | 26.8 | A | 64.9 | M | CRCK1 |
| 247741_at | 341.9 | P | 544.5 | P | 371.6 | P | 605 | P | GIL1 |
| 247742_at | 216.4 | P | 228.5 | P | 173.8 | P | 225.9 | P | --- |
| 247743_at | 220.3 | P | 216.8 | P | 229.2 | P | 236.5 | P | --- |
| 247744_at | 70.7 | P | 86.5 | P | 96.3 | P | 93.9 | P | --- |
| 247745_at | 1310.1 | P | 2028 | P | 1392.6 | P | 2021.5 | P | COPT1 |
| 247746_at | 377.2 | P | 505.9 | P | 505.1 | P | 521.8 | P | ATUCP2 |
| 247747_at | 229 | P | 265.9 | P | 243.9 | P | 250.4 | P | --- |
| 247748_at | 756.1 | P | 768.9 | P | 832.9 | P | 810.6 | P | --- |
| 247749_at | 4.8 | A | 3.2 | A | 5.1 | A | 2.3 | A | MYB119 |
| 247750_at | 1141.5 | P | 1282.7 | P | 931.3 | P | 1407.5 | P | --- |
| 247751_at | 91.4 | P | 242.3 | P | 59.8 | P | 221.8 | P | --- |
| 247752_at | 37.8 | A | 2 | A | 8.4 | A | 21.5 | A | --- |
| 247753_at | 5.1 | A | 34.2 | A | 15.5 | A | 4.7 | A | --- |
| 247754_at | 63.9 | P | 137.9 | P | 91.5 | P | 129.3 | P | --- |
| 247755_at | 17.5 | A | 23.4 | A | 47.2 | A | 3.6 | A | ATSBT4.12 |
| 247756_at | 22 | A | 40.3 | A | 25.1 | A | 35.6 | A | --- |
| 247757_at | 11.1 | A | 7.4 | A | 16.9 | A | 3.6 | A | --- |
| 247758_at | 1.5 | A | 7.2 | A | 14.8 | A | 16.2 | A | ATSBT4.13 |
| 247759_at | 5.1 | A | 29 | A | 24.4 | A | 3.2 | A | COPT3 |
| 247760_at | 364.3 | P | 277.6 | P | 310.8 | P | 393 | P | --- |
| 247761_at | 1453.1 | P | 1442.9 | P | 1473.6 | P | 1376 | P | TOPP2 |
| 247762_at | 123.9 | A | 103.8 | A | 145.9 | A | 51.3 | A | --- |
| 247763_at | 445.8 | P | 449.1 | P | 463.7 | P | 437 | P | NRPB7 |
| 247764_at | 36.8 | A | 17.2 | A | 18.7 | A | 25 | A | --- |
| 247765_at | 57.5 | A | 94 | P | 80.6 | A | 108.9 | P | CYP86A1 |
| 247766_at | 1089.6 | P | 1297.5 | P | 1312.2 | P | 1173.9 | P | ftsh9 |
| 247767_at | 14.3 | A | 17.9 | P | 38.5 | P | 4.7 | A | AGL82 |
| 247768_at | 316.5 | P | 467.4 | P | 386.3 | P | 472 | P | --- |
| 247769_at | 0.9 | A | 2 | A | 13.5 | A | 8.2 | A | LAC16 |
| 247770_at | 687.8 | P | 835.4 | P | 1071.2 | P | 1186.4 | P | --- |
| 247771_at | 1071.2 | P | 654.8 | P | 682.1 | P | 858.6 | P | RANBP1 |
| 247772_at | 88.6 | P | 13.4 | A | 90.6 | P | 39.7 | A | --- |
| 247773_at | 36.5 | A | 24.3 | A | 27.5 | A | 28.9 | A | --- |
| 247774_at | 49.3 | P | 54.3 | P | 71.4 | A | 62.7 | P | --- |
| 247775_at | 90.2 | P | 72.3 | P | 71.4 | A | 47.6 | P | PLC5 |
| 247776_at | 197 | P | 218.9 | P | 187.3 | P | 243.2 | P | PLC4 |
| 247777_at | 1343.5 | P | 1591.8 | P | 1402.1 | P | 1615.4 | P | --- |
| 247778_at | 1.8 | A | 5.2 | A | 3.3 | A | 2 | A | --- |
| 247779_at | 118.3 | P | 126.3 | P | 99.3 | P | 154.7 | P | DDB2 |
| 247780_at | 469.3 | P | 737.6 | P | 753.6 | P | 767.1 | P | --- |
| 247781_at | 30 | A | 14.9 | A | 21.2 | A | 9.5 | A | --- |
| 247782_at | 39.7 | P | 49.9 | P | 46.6 | P | 34.7 | P | --- |
| 247783_at | 1515.2 | P | 1097.3 | P | 909.6 | P | 1099.6 | P | --- |
| 247784_at | 32.6 | A | 16.9 | A | 4.2 | A | 17.9 | A | --- |
| 247785_at | 4.2 | A | 1.3 | A | 6.1 | A | 1.3 | A | --- |
| 247786_at | 492.9 | P | 421 | P | 419.6 | P | 459.3 | P | PMR5 |
| 247787_at | 592.4 | P | 665.4 | P | 710.6 | P | 662.9 | P | --- |
| 247788_at | 234.6 | P | 207.2 | P | 208.9 | P | 242.7 | P | --- |
| 247789_at | 35.9 | A | 3.5 | A | 6.7 | A | 10.1 | A | --- |
| 247790_at | 335.2 | P | 396.7 | P | 306.2 | P | 297.8 | P | --- |
| 247791_at | 3483.9 | P | 3656.6 | P | 3184 | P | 3531.6 | P | ROC7 |
| 247792_at | 315.4 | P | 370.2 | P | 239.7 | P | 320.5 | P | --- |
| 247793_at | 628.7 | P | 588.4 | P | 658.3 | P | 572.6 | P | PSY1 |
| 247794_at | 291.5 | P | 153.2 | P | 118.2 | P | 150.7 | P | PLC1 |
| 247795_at | 225 | P | 175.9 | P | 182.8 | P | 169 | P | --- |
| 247796_at | 4.4 | A | 19.6 | A | 3.4 | A | 17.7 | A | --- |
| 247797_at | 2.8 | A | 14.7 | A | 4.4 | A | 5.9 | A | --- |
| 247798_at | 33.6 | A | 7.6 | A | 12.2 | A | 19 | P | --- |
| 247799_at | 79.9 | P | 4.6 | A | 45.8 | A | 7.4 | A | --- |
| 247800_at | 639.5 | P | 463.3 | P | 406 | P | 327.5 | P | --- |
| 247801_at | 526.6 | P | 545.4 | P | 528.4 | P | 538.1 | P | --- |
| 247802_at | 70 | A | 47.1 | A | 80.6 | A | 69.7 | A | ATL63 |
| 247803_at | 42.8 | P | 71.6 | P | 63.8 | M | 78.3 | A | --- |
| 247804_at | 29.2 | P | 49.3 | P | 73.7 | P | 37.9 | P | SVL5 |
| 247805_at | 21.1 | A | 3.4 | A | 5.8 | A | 5 | A | ATYKT62 |
| 247806_at | 1160.2 | P | 1090.3 | P | 1046.1 | P | 1165.4 | P | STA1 |
| 247807_at | 13.2 | A | 15.1 | A | 7.9 | A | 3 | A | OFP3 |
| 247808_at | 269.5 | P | 223.7 | P | 213.4 | P | 221 | P | ECT10 |
| 247809_at | 31 | P | 44.7 | P | 52.9 | P | 52.1 | P | --- |
| 247810_at | 4679.4 | P | 4093.8 | P | 4038 | P | 4311.2 | P | RPT3 |
| 247811_at | 640.4 | P | 680.2 | P | 601.8 | P | 589.8 | P | ATEXO70B1 |
| 247812_at | 209.2 | P | 608.3 | P | 298 | P | 478.7 | P | --- |
| 247813_at | 6706.6 | P | 6349.4 | P | 7533.3 | P | 7489.1 | P | --- |
| 247814_at | 29.7 | A | 106.3 | P | 46.4 | A | 100.4 | P | MES18 |
| 247815_at | 3249.5 | P | 2956.4 | P | 4154.1 | P | 2929 | P | --- |
| 247816_at | 1290.3 | P | 1528.1 | P | 1287.4 | P | 1598.7 | P | --- |
| 247817_at | 1466 | P | 1391.1 | P | 1283.7 | P | 1312.4 | P | --- |
| 247818_at | 261.8 | P | 208.5 | P | 273.4 | P | 241.7 | P | --- |
| 247819_at | 1183.4 | P | 427 | P | 959.6 | P | 440.4 | P | WNK4 |
| 247820_at | 324.4 | P | 374.7 | P | 241.4 | P | 391.1 | P | SIP1 |
| 247821_at | 845.7 | P | 898.4 | P | 1010.1 | P | 908.8 | P | MSI1 |
| 247822_at | 358.6 | P | 325.1 | P | 365.2 | P | 307.2 | P | SNX2a |
| 247823_at | 305.3 | P | 337.2 | P | 374.4 | P | 300.5 | P | --- |
| 247824_at | 33.4 | P | 32.2 | A | 44.2 | P | 22.4 | A | CHX25 |
| 247825_at | 650.8 | P | 527 | P | 646.9 | P | 376.9 | P | TAF15b |
| 247826_at | 310.9 | P | 314.5 | P | 414.2 | P | 371.7 | P | --- |
| 247827_at | 60.4 | P | 63.4 | P | 46.1 | A | 86.1 | P | LSH5 |
| 247828_at | 238 | P | 213.6 | P | 143.6 | P | 202.1 | P | --- |
| 247829_at | 142.5 | P | 116.7 | P | 154.8 | A | 148.9 | P | --- |
| 247830_at | 166.7 | P | 201.7 | P | 158.2 | P | 181.7 | P | --- |
| 247831_at | 23.9 | P | 35.8 | A | 35.3 | P | 46.1 | P | --- |
| 247832_at | 45.7 | A | 39.7 | A | 6.8 | A | 27.3 | A | EOL2 |
| 247833_at | 209.8 | P | 145.7 | P | 175.2 | P | 154.4 | P | --- |
| 247834_at | 1666 | P | 1482.4 | P | 1886.3 | P | 1573.1 | P | --- |
| 247835_at | 294.2 | P | 172.2 | P | 226.9 | P | 167.4 | P | --- |
| 247836_at | 587.1 | P | 465.5 | P | 545.4 | P | 507 | P | --- |
| 247837_at | 41.1 | A | 28.2 | A | 104.5 | A | 66.6 | A | --- |
| 247838_at | 1422.5 | P | 1112.6 | P | 1390.2 | P | 1159.6 | P | UBP23 |
| 247839_at | 2185 | P | 2296.4 | P | 2870.6 | P | 2600.6 | P | --- |

| | | | | | | | | | |
|---|---|---|---|---|---|---|---|---|---|
| 247840_at | 1984.1 | P | 1555.6 | P | 2216.4 | P | 1670.8 | P | --- |
| 247841_at | 407 | P | 501.7 | P | 457 | P | 452.4 | P | ATFIP1[V] |
| 247842_at | 860.4 | P | 987.2 | P | 970.1 | P | 935.3 | P | --- |
| 247843_at | 4.5 | A | 3.4 | A | 4.6 | A | 3.6 | A | SVL4 |
| 247844_at | 17 | A | 26.3 | A | 2.9 | A | 30 | A | ARR18 |
| 247845_at | 742.5 | P | 601.9 | P | 776.6 | P | 604.1 | P | --- |
| 247846_at | 404.2 | P | 388.1 | P | 421.1 | P | 451 | P | --- |
| 247847_at | 1468.9 | P | 1540.1 | P | 1110.1 | P | 1460.5 | P | --- |
| 247848_at | 230.3 | P | 345.3 | P | 269 | P | 265.9 | P | --- |
| 247849_at | 408.9 | P | 291.5 | P | 470.7 | P | 363.1 | P | --- |
| 247850_at | 206.6 | P | 201.4 | P | 244.5 | P | 188.5 | P | --- |
| 247851_at | 5707.1 | P | 5455.3 | P | 4594.4 | P | 5435.3 | P | TIL |
| 247852_at | 2026.2 | P | 2085.9 | P | 1743 | P | 2108.8 | P | YKT61 |
| 247853_at | 1625.3 | P | 1839.4 | P | 1637.5 | P | 1635.1 | P | PHOT2 |
| 247854_at | 815.6 | P | 828.8 | P | 808.8 | P | 736.1 | P | --- |
| 247855_at | 239.5 | P | 163.8 | P | 195 | P | 183.3 | P | --- |
| 247856_at | 158.7 | P | 257.1 | P | 237 | P | 290.9 | P | --- |
| 247857_at | 11.4 | A | 28.5 | A | 33.3 | A | 23 | A | --- |
| 247858_at | 577.7 | P | 740.4 | P | 512.6 | P | 668.4 | P | TTL |
| 247859_at | 413.8 | P | 329.8 | P | 394.3 | P | 324.9 | P | --- |
| 247860_at | 1039.3 | P | 803.6 | P | 1013.1 | P | 799.4 | P | --- |
| 247861_at | 46.1 | P | 55 | A | 26 | A | 90.8 | P | --- |
| 247862_at | 3173.4 | P | 4192.1 | P | 3046.9 | P | 3267.2 | P | --- |
| 247863_at | 1278.1 | P | 872.8 | P | 980.9 | P | 754.1 | P | --- |
| 247864_s_at | 2283.9 | P | 1278.3 | P | 1562.9 | P | 1186.4 | P | --- |
| 247865_at | 1147.2 | P | 1406.1 | P | 1084 | P | 1072.2 | P | --- |
| 247866_at | 279.7 | P | 37.3 | A | 285.5 | P | 52.4 | P | XTR3 |
| 247867_at | 344 | P | 227.6 | P | 263.8 | P | 314.2 | P | CIPK21 |
| 247868_at | 0.7 | A | 5.4 | A | 7.5 | A | 1.4 | A | MYB36 |
| 247869_at | 9.3 | A | 4.5 | A | 20.3 | A | 25.3 | A | ZFP2 |
| 247870_at | 328.3 | P | 408.5 | P | 322.6 | P | 424.7 | P | --- |
| 247871_at | 2.8 | A | 6.9 | A | 3.6 | A | 22.9 | A | --- |
| 247872_at | 20.9 | A | 35.1 | A | 28.9 | A | 33.3 | P | --- |
| 247873_at | 3.6 | A | 18.5 | A | 43.9 | A | 42.5 | P | --- |
| 247874_at | 1402.2 | P | 1249 | P | 896.1 | P | 1324.4 | P | --- |
| 247875_at | 48.7 | A | 48.8 | A | 72.7 | P | 44.5 | A | --- |
| 247876_at | 12.4 | A | 9.6 | A | 1.1 | A | 2.9 | A | --- |
| 247877_at | 126.6 | P | 126.2 | P | 114.1 | P | 122.9 | P | XBAT32 |
| 247878_at | 32.3 | A | 184.5 | P | 21.7 | A | 118.5 | P | --- |
| 247879_at | 17.1 | A | 13.7 | A | 4.8 | A | 6.8 | A | --- |
| 247880_at | 94.4 | P | 219.2 | P | 70 | P | 192.4 | P | --- |
| 247881_at | 208.1 | P | 393.7 | P | 152.9 | P | 302.1 | P | --- |
| 247882_at | 318.6 | P | 807 | P | 408.5 | P | 793.1 | P | --- |
| 247883_at | 2.8 | A | 8.7 | A | 32.3 | A | 20.5 | A | --- |
| 247884_at | 568 | P | 846.7 | P | 573.6 | P | 955.4 | P | CER3 |
| 247885_at | 48.6 | P | 63.9 | P | 7.5 | A | 49.5 | P | --- |
| 247886_at | 769.8 | P | 989.1 | P | 770 | P | 915.4 | P | --- |
| 247887_at | 50.1 | P | 52 | P | 80 | P | 53.6 | P | --- |
| 247888_at | 60.5 | A | 42.9 | P | 61.9 | A | 60.5 | P | --- |
| 247889_at | 1137.6 | P | 1384.1 | P | 1317.4 | P | 1413.4 | P | APO2 |
| 247890_at | 673.9 | P | 695.9 | P | 747.3 | P | 687.3 | P | ATCNGC5 |
| 247891_at | 290.7 | P | 361.3 | P | 321.1 | P | 358.7 | P | --- |
| 247892_at | 71.3 | A | 96.4 | P | 77.3 | A | 96.2 | P | --- |
| 247893_at | 80.5 | P | 60.9 | P | 125 | P | 56.1 | P | RPB5C |
| 247894_at | 239.7 | P | 199.2 | P | 178.1 | P | 171.2 | P | CPL4 |
| 247895_at | 5.4 | A | 28.5 | A | 35 | A | 36.2 | A | --- |
| 247896_at | 366.7 | P | 296.2 | P | 382.8 | P | 298.9 | P | --- |
| 247897_at | 2.1 | A | 1.6 | A | 3.3 | A | 2.1 | A | TET15 |
| 247898_at | 244.2 | P | 376.7 | P | 292.6 | P | 367.5 | P | ZTL |
| 247899_at | 1603.9 | P | 1847.6 | P | 998 | P | 1729.2 | P | --- |
| 247900_at | 2419.7 | P | 2248.2 | P | 2359 | P | 2518.5 | P | --- |
| 247901_at | 430.5 | P | 342.9 | P | 432.2 | P | 367.9 | P | --- |
| 247902_at | 3979.2 | P | 2869.4 | P | 4728 | P | 3414.8 | P | AHA3 |
| 247903_at | 128.8 | P | 214.3 | P | 104 | P | 214.7 | P | --- |
| 247904_at | 63.6 | P | 102.4 | P | 102 | P | 65.7 | P | AIL5 |
| 247905_at | 3 | A | 8.8 | A | 2.9 | A | 2.7 | A | --- |
| 247906_at | 20.1 | A | 24.2 | A | 29.1 | A | 29.9 | A | IAA33 |
| 247907_at | 491 | P | 606 | P | 514.7 | P | 494.5 | P | --- |
| 247908_at | 329.3 | P | 510.2 | P | 305.6 | P | 425.1 | P | GS1 |
| 247909_at | 263.1 | P | 294.9 | P | 291.3 | P | 363.1 | P | --- |
| 247910_at | 210.2 | P | 262.9 | P | 217.1 | P | 264.3 | P | --- |
| 247911_at | 21.8 | A | 47.7 | A | 15.6 | A | 28.2 | A | XRCC3 |
| 247912_at | 61.6 | P | 39.7 | P | 55.7 | P | 32 | P | --- |
| 247913_at | 21.6 | A | 15 | A | 43.9 | M | 11.3 | A | --- |
| 247914_at | 17.3 | A | 4.1 | A | 2.2 | A | 29.1 | A | --- |
| 247915_at | 37.6 | P | 27.2 | P | 71 | P | 30 | M | --- |
| 247916_at | 289.8 | P | 297.9 | P | 337.8 | P | 350.9 | P | BIO1 |
| 247917_at | 25 | A | 40.5 | A | 45.5 | A | 45.9 | A | BIO1 |
| 247918_at | 374.1 | P | 451.4 | P | 359.6 | P | 387.9 | P | --- |
| 247919_at | 35.1 | A | 50.6 | P | 31 | A | 14 | A | --- |
| 247920_at | 9.9 | A | 18 | A | 2.7 | A | 17.1 | A | --- |
| 247921_at | 394.6 | P | 581.3 | P | 264.7 | P | 374.5 | P | COL5 |
| 247922_at | 37.9 | P | 33.3 | P | 16.9 | A | 45.5 | P | --- |
| 247923_at | 726.8 | P | 794.3 | P | 955.9 | P | 822.1 | P | VDAC4 |
| 247924_at | 1050.1 | P | 695.6 | P | 612.8 | P | 725.4 | P | --- |
| 247925_at | 2691.3 | P | 844.8 | P | 2423.4 | P | 876.8 | P | TCH4 |
| 247926_at | 582.5 | P | 497.3 | P | 875.8 | P | 501.7 | P | --- |
| 247927_at | 3.8 | A | 22.1 | A | 39.9 | A | 33.2 | A | --- |
| 247928_at | 60.2 | A | 57.7 | A | 51.2 | A | 69.2 | A | --- |
| 247929_at | 2269.9 | P | 2006.8 | P | 2199.8 | P | 2091.5 | P | --- |
| 247930_at | 659.8 | P | 471 | P | 358.2 | P | 474.6 | P | --- |
| 247931_at | 1657.2 | P | 2218.6 | P | 1777.3 | P | 2343.7 | P | --- |
| 247932_at | 11.2 | A | 22.5 | A | 42.9 | A | 21.3 | A | --- |
| 247933_at | 114.7 | P | 53.3 | A | 136.2 | P | 71 | A | --- |
| 247934_at | 206.6 | P | 212.5 | P | 209.2 | P | 201.9 | P | --- |
| 247935_at | 1531.1 | P | 1085 | P | 1368.4 | P | 1219.9 | P | --- |
| 247936_at | 815.5 | P | 1115.7 | P | 1162.1 | P | 1069.7 | P | LUT2 |
| 247937_at | 739.1 | P | 810.1 | P | 920.6 | P | 887.1 | P | ACA8 |
| 247938_at | 100.2 | P | 102.5 | P | 98.2 | P | 110.6 | P | --- |
| 247939_at | 209 | P | 154.4 | P | 118 | P | 188.7 | P | ATLIG4 |
| 247940_at | 36.7 | P | 24.6 | M | 54.6 | P | 33.7 | P | PSD2 |
| 247941_at | 8.6 | A | 16.4 | A | 14 | A | 9.5 | A | --- |
| 247942_at | 849 | P | 509.1 | P | 686.7 | P | 541.8 | P | --- |
| 247943_at | 441.8 | P | 638.7 | P | 472.8 | P | 636.2 | P | --- |
| 247944_at | 179.5 | P | 197.8 | P | 235 | P | 200.8 | P | --- |
| 247945_at | 529.1 | P | 329.8 | P | 294 | P | 286.4 | P | --- |
| 247946_at | 615.8 | P | 584.7 | P | 748.7 | P | 623.1 | P | CIA2 |
| 247947_at | 33.2 | A | 9.8 | A | 6.9 | A | 39.4 | A | EIR1 |
| 247948_at | 116.4 | P | 142 | P | 110.8 | P | 144.5 | P | --- |

| | | | | | | | | | |
|---|---|---|---|---|---|---|---|---|---|
| 247949_at | 194.4 | P | 122.4 | P | 86.9 | P | 139.2 | P | CYP81F2 |
| 247950_at | 306.5 | P | 341.4 | P | 426.4 | P | 331.6 | P | --- |
| 247951_at | 53.3 | P | 51 | P | 21.3 | P | 57.2 | P | ORP4C |
| 247952_at | 97.6 | P | 75.8 | P | 40.8 | A | 95 | P | CYP71B10 |
| 247953_at | 14.4 | A | 18.2 | A | 1.6 | A | 2.4 | A | CYP71B10 |
| 247954_at | 83.8 | P | 253.2 | P | 81 | A | 123.2 | P | BGAL4 |
| 247955_at | 783.1 | P | 657.1 | P | 916.1 | P | 764.1 | P | NAP1;3 |
| 247956_at | 30.9 | P | 27.3 | A | 40.5 | A | 30.2 | M | CKX3 |
| 247957_at | 306.4 | P | 150.4 | A | 213 | A | 189.4 | M | ABI2 |
| 247958_at | 36.2 | P | 74.4 | P | 58.8 | P | 54.6 | P | --- |
| 247959_at | 132.7 | P | 107.6 | P | 95.3 | P | 105.8 | P | --- |
| 247960_at | 790.3 | P | 1145.8 | P | 839.1 | P | 863.7 | P | NMT1 |
| 247961_at | 12.4 | A | 3.2 | A | 3.6 | A | 34.7 | A | --- |
| 247962_at | 108.4 | P | 152.1 | P | 140.3 | P | 127.7 | P | MKK6 |
| 247963_at | 152.2 | P | 178.7 | P | 152.3 | P | 155.4 | P | --- |
| 247964_at | 1283.2 | P | 1332.7 | P | 1413.9 | P | 1468.1 | P | PRF3 |
| 247965_at | 85.1 | P | 115.3 | P | 88.4 | P | 141.1 | P | AGP14 |
| 247966_at | 92.2 | P | 131.9 | P | 87.9 | P | 125.6 | P | --- |
| 247967_at | 8.1 | A | 14.9 | A | 20.4 | A | 24.1 | A | anac099 |
| 247968_at | 6205.4 | P | 5718.7 | P | 6476.7 | P | 6844.2 | P | --- |
| 247969_at | 7.7 | A | 17.5 | A | 2.7 | A | 4.1 | A | --- |
| 247970_at | 44.9 | A | 67.7 | P | 31.2 | A | 52.8 | P | --- |
| 247971_at | 207.6 | P | 228.6 | P | 242 | P | 215.4 | P | --- |
| 247972_at | 156.9 | P | 299 | P | 269.9 | P | 285.1 | P | HAG2 |
| 247973_at | 27.7 | M | 40.6 | A | 40.1 | A | 27.2 | A | --- |
| 247974_at | 74.7 | P | 70.7 | P | 61.4 | A | 57.3 | A | --- |
| 247975_at | 22 | A | 18.4 | A | 3.6 | A | 22.1 | A | --- |
| 247976_at | 21.9 | A | 8.5 | A | 21.4 | A | 34.8 | A | --- |
| 247977_at | 294.1 | P | 443.6 | P | 262.9 | P | 451.3 | P | --- |
| 247978_at | 4841.5 | P | 4884.9 | P | 6357.6 | P | 5390.9 | P | --- |
| 247979_at | 335.2 | P | 315.9 | P | 342.7 | P | 299.6 | P | --- |
| 247980_at | 276.9 | P | 265.1 | P | 472.4 | P | 244.3 | P | GNC |
| 247981_at | 33.6 | A | 3.9 | A | 1.9 | A | 1.1 | A | MIOX5 |
| 247982_at | 1756.2 | P | 1278.8 | P | 1559.5 | P | 1316.3 | P | ATSERAT1;1 |
| 247983_at | 1565.6 | P | 1315.9 | P | 1244 | P | 1479.9 | P | PFK7 |
| 247984_s_at | 163.5 | P | 233.7 | P | 169.8 | P | 235.1 | P | ILL1 |
| 247985_at | 374.3 | P | 385.8 | P | 457.1 | P | 352.5 | P | --- |
| 247986_at | 29.6 | A | 18.5 | A | 68.5 | A | 32.4 | A | --- |
| 247987_at | 425.3 | P | 491.7 | P | 478.5 | P | 382.7 | P | --- |
| 247988_at | 451 | P | 418.7 | P | 461.6 | P | 386.8 | P | --- |
| 247989_at | 1578.2 | P | 1288 | P | 1531.9 | P | 1404.9 | P | --- |
| 247990_at | 1146.2 | P | 1185.4 | P | 1116.8 | P | 1214 | P | --- |
| 247991_at | 2.7 | A | 36.4 | A | 3.6 | A | 13.6 | A | ATEXPA14 |
| 247992_at | 226.2 | P | 243.9 | P | 232.1 | P | 242.8 | P | --- |
| 247993_at | 356.2 | P | 431.9 | P | 390.9 | P | 417.7 | P | --- |
| 247994_at | 824.6 | P | 855.6 | P | 847.1 | P | 819.9 | P | --- |
| 247995_at | 32.2 | P | 53.6 | P | 63.6 | P | 58.5 | P | --- |
| 247996_at | 1475.5 | P | 1418.1 | P | 888.4 | P | 1317.2 | P | --- |
| 247997_at | 452.8 | P | 461.7 | P | 398.5 | P | 349.6 | P | ATARP8 |
| 247998_at | 40.2 | A | 33.1 | P | 30.4 | P | 31.6 | P | --- |
| 247999_at | 1788.2 | P | 1553.8 | P | 1500.1 | P | 1478 | P | UBC30 |
| 248000_at | 509.6 | P | 639.5 | P | 458.9 | P | 509 | P | --- |
| 248001_at | 755.6 | P | 916.6 | P | 816.1 | P | 921.7 | P | CBL2 |
| 248002_at | 216.6 | P | 218.8 | P | 144.8 | P | 163.2 | P | ATARP8 |
| 248003_at | 127.3 | P | 160.9 | P | 126.7 | P | 146.7 | P | --- |
| 248004_at | 52.6 | P | 62.1 | P | 73.4 | P | 13.7 | P | PRA1.G2 |
| 248005_at | 340 | P | 385.5 | P | 404.1 | P | 360.5 | P | --- |
| 248006_at | 168.5 | P | 253.5 | P | 135.3 | P | 186.9 | P | HAP8 |
| 248007_at | 600.5 | P | 1029.9 | P | 570.7 | P | 676.2 | P | --- |
| 248008_at | 201.5 | P | 253.9 | P | 168.2 | P | 236.9 | P | WRKY2 |
| 248009_at | 656.5 | P | 903.4 | P | 950.8 | P | 953.2 | P | CSN6A |
| 248010_at | 1032.3 | P | 1071.1 | P | 947.8 | P | 1086 | P | PEX5 |
| 248011_at | 29.9 | A | 31.1 | A | 20.9 | A | 19.2 | P | GAMT2 |
| 248012_at | 52.5 | P | 38.9 | A | 52.1 | M | 45 | M | --- |
| 248013_at | 2.7 | A | 1.9 | A | 5.4 | A | 2.9 | A | ACA8 |
| 248014_at | 251 | P | 297.1 | P | 220.8 | P | 220.5 | P | --- |
| 248015_at | 18.6 | A | 56.4 | M | 46.3 | A | 57.2 | P | --- |
| 248016_at | 181.5 | P | 179.5 | P | 223.7 | P | 155 | P | --- |
| 248017_at | 144.9 | P | 123.5 | P | 159.9 | P | 162.6 | P | --- |
| 248018_at | 1.6 | A | 16.9 | A | 12.1 | A | 10.8 | A | --- |
| 248019_at | 6.6 | A | 2.9 | A | 2 | A | 3 | A | --- |
| 248020_at | 1.7 | A | 7.2 | A | 30 | A | 39.1 | A | --- |
| 248021_at | 849 | P | 685.3 | P | 1021.2 | P | 832.8 | P | --- |
| 248022_at | 10.2 | A | 22.9 | A | 5.5 | A | 3.7 | A | APUM12 |
| 248023_at | 166.1 | P | 201.8 | P | 175 | P | 181.3 | P | --- |
| 248024_at | 66 | P | 76.5 | P | 85.8 | P | 75.8 | P | --- |
| 248025_at | 1573.4 | P | 1379 | P | 1316.9 | P | 1398.4 | P | NOI |
| 248026_at | 1038.1 | P | 1243 | P | 999.3 | P | 1097.1 | P | --- |
| 248027_at | 572 | P | 616.9 | P | 579.3 | P | 565.6 | P | ATKCO1 |
| 248028_at | 1235.5 | P | 1203.1 | P | 1935.4 | P | 1307.9 | P | --- |
| 248029_at | 225.5 | P | 250.4 | P | 244.1 | P | 259.1 | P | BAM4 |
| 248030_at | 203.3 | P | 222.5 | P | 192.6 | P | 211.9 | P | SRT1 |
| 248031_at | 310.9 | P | 267.9 | P | 287.4 | P | 294.7 | P | --- |
| 248032_at | 153.1 | P | 189.9 | P | 151.8 | P | 175.6 | P | --- |
| 248033_s_at | 6.7 | A | 5.3 | A | 4.9 | A | 22.4 | A | --- |
| 248034_at | 909.9 | P | 1111.4 | P | 892.4 | P | 1012.8 | P | D6PK |
| 248035_at | 113.8 | M | 105.7 | P | 78.1 | A | 69.6 | P | --- |
| 248036_at | 817.3 | P | 543.8 | P | 801.8 | P | 606.3 | P | OLI2 |
| 248037_at | 958.2 | P | 1192.1 | P | 683.9 | P | 1090.5 | P | OPT1 |
| 248038_at | 32.6 | A | 27 | A | 28.6 | A | 8.2 | A | --- |
| 248039_at | 10.9 | A | 61.8 | A | 53.3 | A | 37.9 | A | --- |
| 248040_at | 51.8 | P | 48 | P | 86.9 | A | 60.7 | P | --- |
| 248041_at | 923.8 | P | 1092.1 | P | 924.5 | P | 939.2 | P | emb2731 |
| 248042_at | 492.9 | P | 552 | P | 473.8 | P | 545.1 | P | --- |
| 248043_s_at | 8139.3 | P | 9957 | P | 11077 | P | 12310.4 | P | --- |
| 248044_at | 275.2 | P | 279.6 | P | 264.8 | P | 266.2 | P | --- |
| 248045_at | 4081.8 | P | 5007.3 | P | 4972.3 | P | 5261.3 | P | HSP81-2 |
| 248046_at | 554 | P | 582.8 | P | 697.9 | P | 557.7 | P | --- |
| 248047_at | 13.3 | A | 13.3 | P | 6.9 | A | 5 | A | --- |
| 248048_at | 17.4 | A | 5.3 | A | 2.2 | A | 12.5 | A | NAS2 |
| 248049_at | 361.3 | P | 268.6 | P | 350.7 | P | 237 | P | COX15 |
| 248050_at | 110.3 | P | 197.1 | P | 112.9 | P | 165.8 | P | --- |
| 248051_at | 21.6 | A | 56 | P | 56.3 | A | 60.2 | A | ATMYB103 |
| 248052_at | 9.7 | A | 10 | A | 8.6 | A | 13.4 | A | --- |
| 248053_at | 119.9 | P | 104.7 | P | 102.5 | P | 141.7 | P | AtNMNAT |
| 248054_at | 28.1 | P | 57.4 | P | 47.4 | M | 60.8 | P | --- |
| 248055_at | 47.1 | P | 70 | P | 64 | P | 58.7 | P | --- |
| 248056_at | 492 | P | 512.3 | P | 424.4 | P | 517.9 | P | XYLT |
| 248057_at | 81.7 | P | 140.9 | P | 91.7 | P | 125.9 | P | --- |

| | | | | | | | | | |
|---|---|---|---|---|---|---|---|---|---|
| 248058_at | 935.9 | P | 899.8 | P | 868 | P | 998.5 | P | --- |
| 248059_at | 217.8 | P | 261.3 | P | 315.5 | P | 274.8 | P | TRN1 |
| 248060_at | 72 | P | 78.9 | P | 58.4 | M | 74.7 | P | --- |
| 248061_at | 63.9 | A | 105.3 | P | 93.2 | A | 90.1 | P | --- |
| 248062_at | 251.9 | P | 1140.1 | P | 255.4 | P | 926.2 | P | --- |
| 248063_at | 314.6 | P | 270.4 | P | 380 | P | 270.5 | P | --- |
| 248064_at | 69.9 | P | 42.9 | A | 30.4 | A | 22.8 | A | --- |
| 248065_at | 272.2 | P | 318.1 | P | 314.9 | P | 272.6 | P | --- |
| 248066_at | 21.1 | A | 5.5 | A | 34.3 | A | 16.1 | A | QRT1 |
| 248067_at | 294.8 | P | 303.3 | P | 264.7 | P | 275.6 | P | --- |
| 248068_at | 929.8 | P | 991.3 | P | 1094.9 | P | 1093.3 | P | --- |
| 248069_at | 27.5 | A | 67.8 | P | 58.2 | A | 51.3 | M | --- |
| 248070_at | 471.7 | P | 493.3 | P | 399.3 | P | 393.7 | P | --- |
| 248071_at | 334.8 | P | 248.8 | P | 322.2 | P | 318.2 | P | --- |
| 248072_at | 6.6 | A | 4 | A | 7.3 | A | 5.6 | A | --- |
| 248073_at | 59 | P | 40.4 | P | 60.4 | M | 62.5 | P | --- |
| 248074_at | 410.3 | P | 855.3 | P | 621.6 | P | 934.8 | P | FLA1 |
| 248075_at | 256.9 | P | 584.3 | P | 364 | P | 414.8 | P | CRR21 |
| 248076_at | 3.7 | A | 6.9 | A | 33.3 | A | 20.9 | A | --- |
| 248077_at | 41.8 | M | 29 | A | 35.4 | A | 33 | A | --- |
| 248078_at | 25.6 | A | 28.8 | A | 50.5 | A | 6 | A | --- |
| 248079_at | 344.7 | P | 351.3 | P | 282.8 | P | 306.5 | P | --- |
| 248080_at | 168.3 | P | 165.8 | P | 210.1 | P | 167.2 | P | --- |
| 248081_at | 251.6 | P | 262.5 | P | 273.9 | P | 268.7 | P | EDM2 |
| 248082_at | 161 | P | 198.1 | P | 125.3 | P | 210.6 | P | --- |
| 248083_at | 7.5 | A | 11.7 | A | 8.3 | A | 7.1 | A | --- |
| 248084_at | 100 | P | 74.6 | P | 87.5 | A | 66.4 | A | NHX3 |
| 248085_at | 380.7 | P | 338.9 | P | 379.2 | P | 368.4 | P | SVL1 |
| 248086_at | 39.3 | A | 39.9 | P | 28.8 | A | 31.6 | P | GEX1 |
| 248087_at | 198.4 | P | 181.1 | P | 193 | P | 205.6 | P | --- |
| 248088_at | 2123.1 | P | 2004.4 | P | 2817.8 | P | 1980.1 | P | --- |
| 248089_at | 18.7 | A | 26.1 | P | 6 | A | 19.7 | A | AtRAN4 |
| 248090_at | 4 | A | 14.5 | A | 7.1 | A | 14.1 | A | MAPKKK15 |
| 248091_at | 1189.9 | P | 868.2 | P | 1117.3 | P | 846.5 | P | VTC5 |
| 248092_at | 61.9 | A | 126.4 | P | 91.6 | A | 109.5 | P | SUMO3 |
| 248093_at | 483.5 | P | 508.6 | P | 470.3 | P | 493.6 | P | --- |
| 248094_at | 2049.2 | P | 2068.9 | P | 1962.3 | P | 2142.2 | P | --- |
| 248095_at | 577.9 | P | 857.1 | P | 653.9 | P | 860.1 | P | ATMAP65-1 |
| 248096_at | 28.4 | A | 25 | A | 3.6 | A | 23.6 | A | --- |
| 248097_at | 168.3 | P | 282.9 | P | 218.2 | P | 291.5 | P | PPX2 |
| 248098_at | 2063.1 | P | 2172.4 | P | 1796.5 | P | 2319.4 | P | --- |
| 248099_at | 607.5 | P | 624.3 | P | 582.2 | P | 580.5 | P | TOP1ALPHA |
| 248100_at | 752.7 | P | 1227.5 | P | 796.8 | P | 1314.3 | P | --- |
| 248101_at | 655 | P | 452.7 | P | 618.3 | P | 516.1 | P | --- |
| 248102_at | 578.8 | P | 505.1 | P | 674.1 | P | 548 | P | --- |
| 248103_at | 2040.3 | P | 2144.6 | P | 2046.5 | P | 2105.6 | P | SUMO2 |
| 248104_at | 95.9 | P | 59.7 | A | 109 | P | 67.1 | A | IAMT1 |
| 248105_at | 1537.8 | P | 1306.7 | P | 1556.7 | P | 1426 | P | FTSZ1-1 |
| 248106_at | 215.4 | P | 301.7 | P | 292.1 | P | 279.6 | P | --- |
| 248107_at | 3390.5 | P | 2911.4 | P | 3214.1 | P | 2967.9 | P | RAN3 |
| 248108_at | 541.1 | P | 542.7 | P | 482.9 | P | 533.8 | P | CNX5 |
| 248109_at | 726.1 | P | 693 | P | 588.2 | P | 676.3 | P | TOP1BETA |
| 248110_at | 25.7 | A | 16.9 | A | 45.8 | A | 39.6 | A | --- |
| 248111_at | 28.9 | A | 30.9 | A | 28.1 | A | 19.5 | A | --- |
| 248112_at | 7 | A | 12.8 | A | 7.4 | A | 2.3 | A | --- |
| 248113_at | 34.4 | A | 26.4 | A | 62.3 | A | 48.4 | P | --- |
| 248114_at | 33.5 | P | 34.2 | P | 56.9 | P | 50.4 | P | --- |
| 248115_at | 1403.9 | P | 1211.8 | P | 1549.5 | P | 1127 | P | --- |
| 248116_at | 6 | A | 19 | A | 3.1 | A | 1.7 | A | MYB120 |
| 248117_at | 440.4 | P | 402.5 | P | 411.2 | P | 360.8 | P | FIP2 |
| 248118_at | 1590.5 | P | 515.5 | P | 605.9 | P | 544.3 | P | --- |
| 248119_at | 140.2 | P | 132.5 | P | 207.7 | P | 114.3 | P | --- |
| 248120_at | 2217.8 | P | 2876.5 | P | 1711.1 | P | 2487.6 | P | --- |
| 248121_at | 68.6 | P | 72.5 | P | 81.4 | P | 101.8 | P | GAUT12 |
| 248122_at | 7.9 | A | 6.8 | A | 2.8 | A | 8.8 | A | --- |
| 248123_at | 59.4 | P | 111.5 | P | 107.2 | P | 103.5 | P | --- |
| 248124_at | 298.9 | P | 312.9 | P | 263.7 | P | 250.9 | P | ATG18F |
| 248125_at | 31.1 | A | 34.9 | A | 94.2 | P | 35.9 | P | --- |
| 248126_at | 4784.7 | P | 4965.3 | P | 4089.1 | P | 5150 | P | ROC5 |
| 248127_at | 1446 | P | 1490.2 | P | 1448.1 | P | 1435.5 | P | --- |
| 248128_at | 22364.6 | P | 21868 | P | 27394.6 | P | 24261.3 | P | THI1 |
| 248129_at | 156.7 | P | 157.6 | P | 123.9 | P | 140.5 | P | --- |
| 248130_at | 23.5 | P | 22.7 | P | 37.2 | A | 9.3 | A | --- |
| 248131_at | 345 | P | 373.9 | P | 295.6 | P | 411.7 | P | --- |
| 248132_at | 596.1 | P | 463.9 | P | 507.7 | P | 444.9 | P | SGP1 |
| 248133_at | 81.8 | P | 90.2 | P | 109.6 | P | 76.3 | P | --- |
| 248134_at | 486.6 | P | 566.3 | P | 368.5 | P | 456.2 | P | --- |
| 248135_at | 90.1 | P | 103.8 | P | 96.6 | A | 114.7 | P | --- |
| 248136_at | 210 | P | 204.2 | P | 261.5 | P | 221.6 | P | --- |
| 248137_at | 42.1 | A | 26.4 | A | 66.4 | P | 60.3 | M | --- |
| 248138_at | 2986.6 | P | 2510.6 | P | 2526.1 | P | 2795.1 | P | PDC2 |
| 248139_at | 755.8 | P | 751.1 | P | 914.4 | P | 828.7 | P | --- |
| 248140_at | 694.6 | P | 565.1 | P | 614.4 | P | 475 | P | --- |
| 248141_at | 6.4 | A | 17.2 | A | 2.3 | A | 0.8 | A | --- |
| 248142_at | 284.8 | P | 255.6 | P | 213.1 | P | 251.2 | P | --- |
| 248143_at | 35.6 | P | 35.9 | M | 3.6 | A | 38.7 | A | --- |
| 248144_at | 667.7 | P | 704.6 | P | 680.6 | P | 648.4 | P | GPT1 |
| 248145_at | 162.3 | P | 103 | P | 153.6 | P | 171.3 | P | --- |
| 248146_at | 5717.2 | P | 6118.8 | P | 5098.8 | P | 5906.2 | P | --- |
| 248147_at | 1801.4 | P | 1604 | P | 1837.3 | P | 1623.6 | P | ATRBP45A |
| 248148_at | 331.7 | P | 346.5 | P | 421.9 | P | 341.6 | P | --- |
| 248149_at | 813.6 | P | 803 | P | 628 | P | 833.7 | P | --- |
| 248150_at | 92.3 | P | 183 | P | 123.1 | P | 201.3 | P | ATK3 |
| 248151_at | 9966.6 | P | 12584.3 | P | 9847.2 | P | 11327.2 | P | LHCB3 |
| 248152_at | 92.8 | P | 62 | M | 116.5 | P | 77.4 | P | MRE11 |
| 248153_at | 841.6 | P | 487.6 | P | 575.9 | P | 549.7 | P | ATCNGC4 |
| 248154_at | 38.6 | P | 57.1 | P | 66.6 | P | 70.5 | P | --- |
| 248155_at | 4052.8 | P | 2849.2 | P | 4268.2 | P | 2956.9 | P | AHL |
| 248156_at | 8.9 | A | 0.6 | A | 2.2 | A | 0.4 | A | --- |
| 248157_at | 1.2 | A | 18.7 | A | 41.1 | M | 12.5 | A | --- |
| 248158_at | 543.4 | P | 525.5 | P | 455.9 | P | 564.4 | P | --- |
| 248159_at | 10.2 | A | 21.2 | A | 25.9 | A | 23.9 | A | --- |
| 248160_at | 20.4 | A | 64.7 | P | 5.1 | A | 34.2 | A | --- |
| 248161_at | 8.9 | A | 6.3 | A | 2.7 | A | 11.5 | A | --- |
| 248162_at | 2968.4 | P | 3115.7 | P | 2054.1 | P | 2966.8 | P | FQR1 |
| 248163_at | 164.3 | P | 522.8 | P | 178.5 | P | 568.5 | P | DFL1 |
| 248164_at | 156.4 | P | 185 | P | 139.1 | P | 173.4 | P | PBP1 |
| 248165_at | 1345.9 | P | 1418.9 | P | 1305.1 | P | 1346.1 | P | PHOS32 |
| 248166_at | 113.1 | P | 122.4 | P | 130.2 | P | 154.7 | P | --- |

| | | | | | | | | | |
|---|---|---|---|---|---|---|---|---|---|
| 248167_at | 117.7 | P | 86 | P | 129.4 | P | 102.2 | P | --- |
| 248168_at | 29.3 | A | 25.9 | M | 31.4 | P | 41 | A | BGLU41 |
| 248169_at | 42.2 | P | 196.6 | P | 38.1 | P | 177.1 | P | ANK |
| 248170_at | 0.6 | A | 0.6 | A | 1.8 | A | 0.6 | A | --- |
| 248171_at | 1789.4 | P | 1727.2 | P | 1482.1 | P | 1626.1 | P | ILR3 |
| 248172_at | 122.3 | P | 130.5 | P | 113.4 | P | 124.7 | P | --- |
| 248173_at | 1401.3 | P | 1145.8 | P | 961.5 | P | 1067.1 | P | --- |
| 248174_at | 7562.9 | P | 8439.8 | P | 6997.4 | P | 7513.3 | P | --- |
| 248175_at | 366.1 | P | 614 | P | 257.4 | P | 477.4 | P | RAT5 |
| 248176_at | 92.1 | P | 93.4 | P | 82.9 | A | 92.7 | P | Fh5 |
| 248177_at | 309.7 | P | 368.8 | P | 354.3 | P | 315.2 | P | --- |
| 248178_at | 3 | A | 9.6 | A | 43.2 | A | 27.7 | A | --- |
| 248179_at | 467.9 | P | 381 | P | 418.4 | P | 499 | P | THE1 |
| 248180_at | 914.6 | P | 867.2 | P | 690.1 | P | 897 | P | AGD5 |
| 248181_at | 1453.3 | P | 1527.3 | P | 1411.9 | P | 1457.1 | P | --- |
| 248182_at | 9.5 | A | 32.8 | P | 13.1 | A | 18 | A | --- |
| 248183_at | 8.9 | A | 41 | A | 45.6 | A | 6 | A | --- |
| 248184_at | 16.9 | A | 12.2 | A | 2.1 | A | 19.5 | A | --- |
| 248185_at | 1880.2 | P | 293.2 | P | 2471.9 | P | 744.7 | P | UF3GT |
| 248186_at | 1207.4 | P | 1273.1 | P | 1349.9 | P | 1271.1 | P | --- |
| 248187_at | 850.7 | P | 1078.3 | P | 872.9 | P | 1042.5 | P | --- |
| 248188_at | 24.4 | A | 29.6 | A | 46.3 | A | 43.2 | A | HSFA9 |
| 248189_at | 90.2 | P | 151.2 | P | 141.1 | P | 159.6 | P | --- |
| 248190_at | 52.8 | A | 83.2 | M | 108.3 | A | 90.1 | P | --- |
| 248191_at | 58.9 | P | 98.2 | P | 136.4 | P | 101.2 | P | --- |
| 248192_at | 62.8 | P | 107.8 | P | 82 | P | 59.4 | P | ILL3 |
| 248193_at | 860.3 | P | 779.8 | P | 602.5 | P | 712.4 | P | HGO |
| 248194_at | 4.7 | A | 3.3 | A | 1 | A | 1.6 | A | --- |
| 248195_at | 622.1 | P | 628.3 | P | 358.4 | P | 426.9 | P | ATMAMI |
| 248196_at | 2.5 | A | 3.9 | A | 9.2 | A | 4.9 | A | --- |
| 248197_at | 58.2 | P | 123.4 | P | 57.7 | A | 94.5 | P | PORA |
| 248198_at | 61.6 | P | 140.6 | P | 128.7 | P | 131.2 | P | --- |
| 248199_at | 1715.9 | P | 829 | P | 1433.2 | P | 788.2 | P | --- |
| 248200_at | 4361.1 | P | 5152.3 | P | 3665.5 | P | 4618.4 | P | ATOMT1 |
| 248201_at | 289.2 | P | 403.5 | P | 381.3 | P | 375.4 | P | PTAC15 |
| 248202_at | 5.2 | A | 13.6 | A | 33.5 | A | 12.2 | A | --- |
| 248203_at | 4.9 | A | 15.7 | A | 36.1 | A | 4.5 | A | MYB49 |
| 248204_at | 78.3 | P | 117.8 | P | 126.3 | P | 109.7 | P | ATM2 |
| 248205_at | 135.3 | P | 100.7 | P | 96.3 | P | 93.4 | P | --- |
| 248206_at | 16.2 | A | 38 | A | 34.2 | A | 35.8 | A | --- |
| 248207_at | 584.4 | P | 617.7 | P | 566.2 | P | 645.1 | P | --- |
| 248208_at | 1.4 | A | 2 | A | 4.2 | A | 1.9 | A | ATHB52 |
| 248209_at | 1.5 | A | 3.3 | A | 2.1 | A | 1.4 | A | --- |
| 248210_at | 1.2 | A | 0.8 | A | 2.2 | A | 0.9 | A | --- |
| 248211_at | 1.3 | A | 6 | A | 6.6 | A | 1.7 | A | --- |
| 248212_at | 9.3 | A | 31.7 | A | 40.8 | A | 2.9 | A | --- |
| 248213_at | 63 | P | 61.2 | A | 93.2 | A | 76.5 | A | AtGRF7 |
| 248214_at | 2.6 | A | 1.7 | A | 5.8 | A | 1.6 | A | --- |
| 248215_at | 22 | A | 15.3 | A | 9.8 | A | 5.4 | A | --- |
| 248216_at | 31.9 | A | 15.2 | A | 5.7 | A | 5.6 | A | --- |
| 248217_at | 10061.3 | P | 8277 | P | 8296.7 | P | 9050.3 | P | CB5-E |
| 248218_at | 26.3 | A | 52.4 | A | 78.2 | A | 36.4 | A | --- |
| 248219_at | 751.7 | P | 795.7 | P | 819.4 | P | 923.6 | P | --- |
| 248220_at | 1137.9 | P | 920.8 | P | 914.3 | P | 973.9 | P | --- |
| 248221_at | 632.2 | P | 731.8 | P | 505.6 | P | 651.1 | P | VPS26A |
| 248222_at | 468.1 | P | 437.6 | P | 448.8 | P | 400.6 | P | --- |
| 248223_at | 613.1 | P | 713.2 | P | 665.8 | P | 681.7 | P | --- |
| 248224_at | 1867 | P | 2542.1 | P | 1467.2 | P | 2285.5 | P | --- |
| 248225_at | 36.4 | A | 28.2 | M | 75.3 | A | 16.1 | A | --- |
| 248226_at | 177.2 | P | 175.3 | P | 209.1 | P | 191.9 | P | --- |
| 248227_at | 42.3 | A | 99.9 | P | 37.9 | A | 94.1 | P | --- |
| 248228_at | 270.2 | P | 259 | P | 311.8 | P | 305.8 | P | --- |
| 248229_at | 2.6 | A | 5.5 | A | 8.2 | A | 1.6 | A | --- |
| 248230_at | 106.1 | P | 148.7 | P | 163.4 | P | 192.1 | P | --- |
| 248231_at | 161.4 | P | 181.8 | A | 217.2 | P | 151.9 | A | --- |
| 248232_at | 171.5 | P | 197.9 | P | 210.8 | P | 211.5 | P | MLO11 |
| 248233_at | 24.7 | P | 6.2 | A | 34.8 | A | 24.4 | A | --- |
| 248234_at | 1749.4 | P | 1542.6 | P | 1828.4 | P | 1426 | P | --- |
| 248235_at | 1100.7 | P | 1346.8 | P | 1446.1 | P | 1313.2 | P | emb2737 |
| 248236_at | 192.4 | P | 116.8 | P | 89.7 | P | 108.4 | P | --- |
| 248237_at | 301.9 | P | 246.2 | P | 289 | P | 267.4 | P | --- |
| 248238_at | 238 | P | 247.5 | P | 291.1 | P | 217.5 | P | --- |
| 248239_at | 294.2 | P | 272.1 | P | 302.7 | P | 296.4 | P | --- |
| 248240_at | 35.5 | A | 50.6 | M | 64.5 | A | 30.8 | A | CUC2 |
| 248241_at | 3.3 | A | 2.3 | A | 5 | A | 4.7 | A | --- |
| 248242_at | 1146.2 | P | 1383 | P | 1349.9 | P | 1288.6 | P | --- |
| 248243_at | 769.8 | P | 282.3 | P | 492.6 | P | 467.2 | P | CPuORF50 |
| 248244_at | 6.7 | A | 24.3 | A | 27.1 | A | 38.4 | A | --- |
| 248245_at | 29.7 | A | 47.5 | A | 19 | A | 29.6 | A | --- |
| 248246_at | 47.9 | A | 46.7 | P | 61.7 | P | 49.5 | A | TRY |
| 248247_at | 101.4 | P | 123.3 | P | 156.6 | M | 132.7 | P | SPCH |
| 248248_at | 1166.7 | P | 1172.2 | P | 768.9 | P | 1211.9 | P | SPDS3 |
| 248249_at | 1148.4 | P | 1260.1 | P | 1207.1 | P | 1380.8 | P | --- |
| 248250_at | 601.3 | P | 694.4 | P | 533.6 | P | 706.6 | P | CNGC1 |
| 248251_at | 97.8 | P | 78 | P | 98.6 | P | 48.9 | P | --- |
| 248252_at | 31.1 | A | 14.1 | A | 5.1 | A | 2.5 | A | AGP22 |
| 248253_at | 80.3 | P | 62.3 | A | 82.3 | P | 26.2 | A | CRF3 |
| 248254_at | 30.8 | A | 39.3 | A | 47.4 | A | 31.1 | A | --- |
| 248255_at | 2597 | P | 2172.4 | P | 2468.6 | P | 2075.1 | P | CLPX |
| 248256_at | 310.5 | P | 251.6 | P | 211.3 | P | 209.8 | P | --- |
| 248257_at | 27.2 | A | 38.1 | A | 20 | A | 43.8 | A | --- |
| 248258_at | 938 | P | 867.9 | P | 902.6 | P | 836.5 | P | --- |
| 248259_at | 877.8 | P | 882.5 | P | 866.7 | P | 765.8 | P | --- |
| 248260_at | 1 | A | 3.1 | A | 2.1 | A | 3.3 | A | --- |
| 248261_at | 193.6 | P | 215.7 | P | 176.1 | P | 228.3 | P | PDV1 |
| 248262_at | 473.2 | P | 587.3 | P | 516.9 | P | 676.3 | P | --- |
| 248263_at | 786.7 | P | 1308.4 | P | 745 | P | 1117.6 | P | PMEPCRF |
| 248264_at | 711.6 | P | 986.8 | P | 593.9 | P | 775.9 | P | --- |
| 248265_at | 130.2 | P | 147 | P | 104.1 | P | 116.3 | P | SDG29 |
| 248266_at | 422.8 | P | 380.3 | P | 422.5 | P | 427.7 | P | --- |
| 248267_at | 1282.6 | P | 891.5 | P | 1128 | P | 907.3 | P | GLT1 |
| 248268_at | 1055.8 | P | 1008 | P | 1107.5 | P | 982.9 | P | --- |
| 248269_at | 273.3 | P | 336.8 | P | 338.6 | P | 316.6 | P | ACBP1 |
| 248270_at | 863.9 | P | 3624.9 | P | 3322.9 | P | 3363.6 | P | ORG1 |
| 248271_at | 1659.9 | P | 1305.2 | P | 1655.1 | P | 1527 | P | --- |
| 248272_at | 890.1 | P | 885.8 | P | 818.3 | P | 786.7 | P | --- |
| 248273_at | 113.4 | P | 204.7 | P | 124.5 | P | 221.2 | P | --- |
| 248274_at | 25.3 | A | 38.6 | P | 31.4 | A | 44.4 | A | ATOPT9 |
| 248275_at | 1.4 | A | 0.9 | A | 2.1 | A | 2.2 | A | ATOPT8 |

| | | | | | | | | | |
|---|---|---|---|---|---|---|---|---|---|
| 248276_at | 767.2 | P | 1103.5 | P | 838.6 | P | 669.5 | P | YSL3 |
| 248277_at | 41.4 | A | 114.1 | P | 53.6 | A | 82.8 | P | --- |
| 248278_at | 44.8 | A | 40.8 | A | 44.8 | A | 35 | A | --- |
| 248279_at | 141.4 | P | 125.8 | A | 104 | A | 123.7 | P | ATIM |
| 248280_at | 65.8 | A | 67.4 | A | 123.7 | M | 62.9 | P | --- |
| 248281_at | 316.4 | P | 304.6 | P | 361.9 | P | 307.2 | P | --- |
| 248282_at | 165.1 | P | 149.5 | P | 155.6 | P | 114 | P | --- |
| 248283_at | 2489 | P | 1796.3 | P | 2796.5 | P | 1938.1 | P | PKP-BETA1 |
| 248284_at | 55.8 | P | 75.2 | M | 124.2 | P | 54.3 | A | --- |
| 248285_at | 1021.4 | P | 1374.1 | P | 953.8 | P | 1272.6 | P | --- |
| 248286_at | 91.2 | P | 100.1 | P | 93.6 | P | 86.1 | P | --- |
| 248287_at | 1417.1 | P | 1964.9 | P | 1499.7 | P | 1727 | P | --- |
| 248288_at | 3205.5 | P | 3373.4 | P | 3465.1 | P | 3557.5 | P | --- |
| 248289_at | 244.1 | P | 284.7 | P | 211 | P | 303.6 | P | --- |
| 248290_at | 144.7 | P | 131.3 | P | 92.4 | P | 99.9 | P | --- |
| 248291_at | 167.6 | P | 208 | P | 161 | P | 175.1 | P | --- |
| 248292_at | 191.1 | P | 188 | P | 196.1 | P | 160.6 | P | --- |
| 248293_at | 946.2 | P | 647.6 | P | 806.6 | P | 669.6 | P | --- |
| 248294_at | 154.9 | P | 135.6 | P | 118.7 | P | 94 | P | --- |
| 248295_at | 602.3 | P | 572.3 | P | 724.8 | P | 616.3 | P | --- |
| 248296_at | 78.2 | P | 132.3 | P | 110.7 | P | 119.8 | P | --- |
| 248297_at | 16 | A | 28.6 | M | 58.6 | A | 21 | A | --- |
| 248298_at | 43 | P | 84.2 | P | 65 | A | 94.1 | A | --- |
| 248299_at | 135.7 | P | 99.1 | P | 122.3 | P | 83.5 | P | --- |
| 248300_at | 702.4 | P | 480.6 | P | 697.5 | P | 519.2 | P | TAP46 |
| 248301_at | 170.5 | P | 252.6 | P | 232 | P | 183.8 | P | --- |
| 248302_at | 1209.5 | P | 1017 | P | 1105.8 | P | 946.7 | P | --- |
| 248303_at | 834.8 | P | 1107.5 | P | 906.9 | P | 1094.2 | P | FTSH11 |
| 248304_at | 673.4 | P | 732.2 | P | 847.6 | P | 714.5 | P | PTB2 |
| 248305_at | 618.2 | P | 798 | P | 741.5 | P | 759.9 | P | ATUSP |
| 248306_at | 63.5 | P | 49.4 | P | 98.6 | P | 60.3 | P | WRKY27 |
| 248307_at | 168.3 | P | 121 | P | 157.3 | P | 113.2 | P | --- |
| 248308_at | 273.6 | P | 397.8 | P | 297.5 | P | 345.6 | P | --- |
| 248309_at | 337.8 | P | 768 | P | 452.8 | P | 671.6 | P | --- |
| 248310_at | 219.2 | P | 292.4 | P | 174.7 | P | 283.8 | P | CPuORF14 |
| 248311_at | 944.9 | P | 567.1 | P | 675.6 | P | 746 | P | BETA-OHASE 2 |
| 248312_at | 299.6 | P | 354.8 | P | 234.1 | P | 313.1 | P | --- |
| 248313_at | 600.2 | P | 708.9 | P | 546.2 | P | 633.6 | P | --- |
| 248314_at | 1.1 | A | 2.5 | A | 3.1 | A | 19 | A | --- |
| 248315_at | 99.6 | P | 102.1 | P | 107.6 | P | 121.9 | P | MEF1 |
| 248316_at | 2.3 | A | 6.9 | A | 23.3 | A | 1 | A | --- |
| 248317_at | 10.3 | A | 1.6 | A | 4.9 | A | 10.7 | A | --- |
| 248318_at | 1.7 | A | 2.6 | A | 1.8 | A | 1.3 | A | --- |
| 248319_at | 14.4 | A | 16 | A | 45 | P | 37.2 | A | --- |
| 248320_at | 3.2 | A | 8.4 | A | 0.7 | A | 3.7 | A | --- |
| 248321_at | 41.1 | P | 30.3 | P | 47.9 | P | 29.2 | P | --- |
| 248322_at | 516.6 | P | 718 | P | 590.3 | P | 493.2 | P | --- |
| 248323_at | 3.5 | A | 0.9 | A | 2.1 | A | 1.6 | A | --- |
| 248324_at | 15.9 | A | 34.9 | A | 31.6 | A | 24.1 | A | --- |
| 248325_at | 33.8 | A | 37.1 | A | 43.9 | A | 55.4 | P | --- |
| 248326_at | 1206.7 | P | 712.9 | P | 1120.1 | P | 751.3 | P | --- |
| 248327_at | 593.2 | P | 576.6 | P | 370.3 | P | 484.9 | P | --- |
| 248328_at | 184.1 | P | 188 | P | 240.9 | P | 230.9 | P | --- |
| 248329_at | 1319 | P | 1960 | P | 1390.6 | P | 1738.5 | P | --- |
| 248330_at | 1402.8 | P | 909.5 | P | 1122.6 | P | 892.5 | P | --- |
| 248331_at | 6079.7 | P | 4986.7 | P | 6560.5 | P | 5611.4 | P | --- |
| 248332_at | 748.8 | P | 1351.3 | P | 628.3 | P | 996.4 | P | ATHSP90.1 |
| 248333_at | 115.6 | P | 51 | A | 73.5 | A | 115.9 | P | --- |
| 248334_s_at | 507.6 | P | 489.9 | P | 596.8 | P | 509.3 | P | --- |
| 248335_at | 2195.7 | P | 1448.8 | P | 2130.2 | P | 1702.7 | P | --- |
| 248336_at | 1053.5 | P | 829.8 | P | 1059.8 | P | 905.6 | P | --- |
| 248337_at | 1478.1 | P | 2626.3 | P | 442.4 | P | 2867.5 | P | LTI78 |
| 248338_at | 1344 | P | 1884.5 | P | 1640.3 | P | 1810.6 | P | HCF106 |
| 248339_at | 1023.9 | P | 1248 | P | 1225 | P | 1166.8 | P | OVA6 |
| 248340_at | 457.9 | P | 548.4 | P | 461.8 | P | 556.7 | P | --- |
| 248341_at | 55.5 | P | 99.9 | P | 66.5 | P | 103.1 | P | --- |
| 248342_at | 124.8 | P | 134.8 | P | 149.9 | P | 143.1 | P | MBD13 |
| 248343_at | 13.9 | A | 16.3 | A | 14.7 | A | 0.8 | A | AtMYB19 |
| 248344_at | 183.8 | P | 149.9 | P | 123.4 | P | 145.8 | P | --- |
| 248345_at | 22.4 | A | 14.2 | A | 9.8 | A | 24.1 | A | SHOC1 |
| 248346_at | 393.5 | P | 470.3 | P | 386.7 | P | 448.8 | P | ATGB1 |
| 248347_at | 121.5 | P | 94.1 | P | 105.3 | P | 82.1 | P | --- |
| 248348_at | 697.3 | P | 762 | P | 703.4 | P | 807.6 | P | --- |
| 248349_at | 1834.8 | P | 2293.1 | P | 2055.2 | P | 2264.5 | P | MSBP1 |
| 248350_at | 11 | A | 5.4 | A | 38.8 | A | 26.4 | A | --- |
| 248351_at | 645.1 | P | 870.7 | P | 618.9 | P | 571.9 | P | --- |
| 248352_at | 24.4 | A | 41.9 | A | 13.5 | A | 31.9 | A | LTI65 |
| 248353_at | 737.2 | P | 420 | P | 613.2 | P | 557.4 | P | CYP96A4 |
| 248354_at | 5.9 | A | 5.7 | A | 5.3 | A | 26.7 | P | --- |
| 248355_at | 3.8 | A | 16.2 | A | 1.1 | A | 10.2 | A | ATEXO70A2 |
| 248356_at | 3.7 | A | 13.2 | A | 4.3 | A | 17.5 | A | ATEXO70A3 |
| 248357_at | 216.3 | P | 207.5 | P | 277.3 | P | 204.4 | P | --- |
| 248358_at | 52.9 | P | 32.9 | A | 28.8 | A | 31.7 | A | CYP715A1 |
| 248359_at | 72.1 | P | 95.2 | P | 69.8 | P | 46.9 | P | --- |
| 248360_at | 364.8 | P | 264.5 | P | 212.3 | P | 269 | P | --- |
| 248361_at | 12.7 | A | 28.3 | P | 22.5 | A | 21.5 | P | EDA41 |
| 248362_at | 5930.9 | P | 4352.6 | P | 5797.1 | P | 4449.9 | P | FIB1 |
| 248363_at | 1.5 | A | 2.1 | A | 9.1 | A | 8.9 | A | --- |
| 248364_at | 10.6 | A | 6.3 | A | 7.6 | A | 3.6 | A | --- |
| 248365_at | 1.4 | A | 1.6 | A | 2.6 | A | 1.2 | A | --- |
| 248366_at | 478.6 | P | 467.8 | P | 450.3 | P | 487.1 | P | --- |
| 248367_at | 1.6 | A | 1.7 | A | 5.9 | A | 1 | A | ADF10 |
| 248368_at | 9.9 | A | 9.9 | A | 21.4 | A | 27.4 | A | --- |
| 248369_at | 1481.5 | P | 1417.1 | P | 1515.2 | P | 1487.2 | P | ATRSP41 |
| 248370_at | 25.2 | A | 6.8 | A | 35 | A | 6.9 | A | HDG7 |
| 248371_at | 29.1 | A | 3.8 | A | 6.1 | A | 18.3 | A | GA20OX2 |
| 248372_at | 48 | P | 76 | P | 70.3 | A | 58.5 | P | --- |
| 248373_at | 3 | A | 16.3 | A | 4.2 | A | 15.2 | A | --- |
| 248374_at | 4.3 | A | 2.2 | A | 6.6 | A | 1.1 | A | AGL71 |
| 248375_at | 224.4 | P | 194.5 | P | 214.3 | P | 267 | P | KEA5 |
| 248376_at | 1198.6 | P | 1336.8 | P | 1303.2 | P | 1325.4 | P | --- |
| 248377_at | 1342.9 | P | 802.4 | P | 680.3 | P | 1626.2 | P | --- |
| 248378_at | 271.9 | P | 406.7 | P | 348.1 | P | 278 | P | --- |
| 248379_at | 329.8 | P | 379.4 | P | 355.7 | P | 405.1 | P | PBS2 |
| 248380_at | 1946.6 | P | 1729.1 | P | 1850.7 | P | 1935.4 | P | PGM |
| 248381_at | 983.8 | P | 690.9 | P | 815.4 | P | 697.3 | P | --- |
| 248382_at | 559.9 | P | 805.8 | P | 697.8 | P | 742 | P | --- |
| 248383_at | 8.5 | A | 7.5 | A | 24.7 | A | 15.1 | A | --- |
| 248384_at | 11.9 | A | 12.1 | A | 7 | A | 5.5 | A | --- |

| | | | | | | | | | |
|---|---|---|---|---|---|---|---|---|---|
| 248385_at | 27.9 | A | 94.2 | P | 59.7 | A | 87 | P | --- |
| 248386_at | 1269.7 | P | 1130.3 | P | 1147.8 | P | 1163.9 | P | NRPB6A |
| 248387_at | 620.5 | P | 697.8 | P | 844.4 | P | 648.1 | P | --- |
| 248388_at | 522.2 | P | 527.6 | P | 461.2 | P | 646.4 | P | --- |
| 248389_at | 3.4 | A | 15.1 | A | 10.8 | A | 20.4 | A | CBF4 |
| 248390_at | 13.2 | A | 5.9 | A | 2.4 | A | 2.2 | A | IMPA-8 |
| 248391_at | 246.8 | P | 252.2 | P | 289.1 | P | 264.7 | P | --- |
| 248392_at | 61.7 | A | 13.4 | A | 37.1 | A | 16.4 | A | --- |
| 248393_at | 1059 | P | 1314.3 | P | 1217.8 | P | 1500.7 | P | ATBAG1 |
| 248394_at | 199.3 | P | 137.1 | P | 152.5 | P | 116 | P | --- |
| 248395_at | 283.9 | P | 118.6 | P | 246.9 | P | 142.5 | P | AtPP2-A14 |
| 248396_at | 24.4 | A | 4.3 | A | 4.6 | A | 3.8 | A | --- |
| 248397_at | 3.3 | A | 4.6 | A | 4.4 | A | 1.7 | A | --- |
| 248398_at | 3072.2 | P | 2957.1 | P | 3114.2 | P | 2547.9 | P | --- |
| 248399_at | 612.7 | P | 626.4 | P | 898.6 | P | 679 | P | --- |
| 248400_at | 2.9 | A | 3.1 | A | 5.8 | A | 2.7 | A | --- |
| 248401_at | 464.5 | P | 499.2 | P | 446.1 | P | 446.4 | P | HCF208 |
| 248402_at | 348.8 | P | 528.2 | P | 398.2 | P | 479.6 | P | crr1 |
| 248403_at | 549 | P | 541.6 | P | 639.1 | P | 506.8 | P | --- |
| 248404_at | 232 | P | 212.1 | P | 282.7 | P | 233.6 | P | ATTPPA |
| 248405_at | 25.4 | A | 19.4 | A | 63.1 | A | 33 | A | SKS2 |
| 248406_at | 6.2 | A | 3 | A | 6.4 | A | 4.6 | A | --- |
| 248407_at | 14.4 | A | 1.6 | A | 55.7 | A | 10.3 | A | --- |
| 248408_at | 2.3 | A | 13.5 | A | 4.7 | A | 2.8 | A | --- |
| 248409_at | 1044.6 | P | 1132.2 | P | 1075.3 | P | 1088.9 | P | LPA2 |
| 248410_at | 1478.4 | P | 1334.2 | P | 1285.6 | P | 1516 | P | --- |
| 248411_at | 10.1 | A | 20.1 | A | 31.4 | A | 10.6 | A | --- |
| 248412_at | 104.8 | P | 127.2 | P | 160.2 | P | 139.2 | P | --- |
| 248413_at | 63.6 | P | 130.6 | P | 90.3 | P | 167.9 | P | PLE |
| 248414_at | 22.3 | A | 21.9 | A | 13.2 | A | 12.3 | A | --- |
| 248415_at | 590.3 | P | 434.5 | P | 471.7 | P | 334.4 | P | --- |
| 248416_at | 62.8 | A | 133.3 | P | 44.7 | A | 77.8 | A | --- |
| 248417_at | 2.4 | A | 7.3 | A | 7.3 | A | 18.1 | A | --- |
| 248418_at | 287.3 | P | 330.9 | P | 314 | P | 333.3 | P | CPSF160 |
| 248419_at | 1245.5 | P | 1746.8 | P | 1755.8 | P | 1785.2 | P | EXL3 |
| 248420_at | 217.4 | P | 531.5 | P | 415.5 | P | 536.9 | P | --- |
| 248421_at | 908 | P | 871.7 | P | 813.9 | P | 786.7 | P | --- |
| 248422_at | 340.8 | P | 320.1 | P | 266.8 | P | 255 | P | YLS7 |
| 248423_at | 17.2 | A | 43.5 | A | 49 | A | 31.4 | A | --- |
| 248424_at | 38.4 | M | 43.2 | P | 82 | P | 51.7 | P | --- |
| 248425_at | 124.1 | P | 156.1 | P | 123.8 | P | 138.5 | P | ACS12 |
| 248426_at | 583.1 | P | 533.1 | P | 514.8 | P | 602.4 | P | --- |
| 248427_at | 1740 | P | 1637.3 | P | 1948.1 | P | 1545.8 | P | ATSBT1.3 |
| 248428_at | 35.3 | A | 25.5 | A | 9.6 | A | 38.7 | A | AHG1 |
| 248429_at | 299.4 | P | 249.6 | P | 275.3 | P | 280.7 | P | --- |
| 248430_at | 14.6 | A | 1.4 | A | 1.4 | A | 1.7 | A | --- |
| 248431_at | 3.6 | A | 21.6 | A | 8.6 | A | 11.2 | A | --- |
| 248432_at | 222.4 | P | 75.5 | P | 135.5 | P | 109.3 | P | --- |
| 248433_at | 1022.5 | P | 1325.2 | P | 1070.5 | P | 1337 | P | --- |
| 248434_at | 605.6 | P | 174.8 | P | 482.2 | P | 204.3 | P | --- |
| 248435_at | 20.4 | A | 35.9 | P | 70.4 | P | 45.4 | P | OLEO3 |
| 248436_at | 283.5 | P | 314.5 | P | 448.4 | P | 359 | P | --- |
| 248437_at | 85.3 | P | 131.4 | P | 163.4 | P | 135.9 | P | EMF2 |
| 248438_at | 279.2 | P | 271.9 | P | 373.5 | P | 321.7 | P | EMF2 |
| 248439_at | 25.7 | A | 33.2 | A | 11 | A | 7.1 | A | --- |
| 248440_at | 166.9 | P | 192.6 | P | 161.7 | P | 175.3 | P | --- |
| 248441_at | 26.1 | A | 3.9 | A | 4.2 | A | 20.2 | A | --- |
| 248442_at | 552.2 | P | 337.3 | P | 440.1 | P | 421.3 | P | --- |
| 248443_at | 26.5 | A | 33.4 | A | 38.2 | A | 38 | A | --- |
| 248444_at | 4.4 | A | 0.2 | A | 9.6 | A | 0.7 | A | --- |
| 248445_at | 305.3 | P | 292.7 | P | 283.9 | P | 233.7 | P | --- |
| 248446_at | 770.7 | P | 760.5 | P | 776.1 | P | 675.2 | P | --- |
| 248447_at | 444 | P | 558.5 | P | 524.1 | P | 553.9 | P | PABN1 |
| 248448_at | 163 | P | 146.4 | P | 205.6 | P | 135.5 | P | --- |
| 248449_at | 4838.8 | P | 5071.2 | P | 5041 | P | 5208.8 | P | --- |
| 248450_at | 153.4 | P | 127.6 | P | 162.3 | P | 79.5 | P | --- |
| 248451_at | 481.7 | P | 593 | P | 585.9 | P | 544.8 | P | --- |
| 248452_at | 653.8 | P | 578.3 | P | 709.9 | P | 646.5 | P | --- |
| 248453_at | 371.9 | P | 389.8 | P | 295.3 | P | 341.9 | P | --- |
| 248454_at | 159.2 | P | 231.8 | P | 241.4 | P | 264 | P | --- |
| 248455_at | 25.9 | A | 1.2 | A | 48.1 | A | 31 | A | --- |
| 248456_at | 39.7 | P | 41.7 | P | 20.8 | A | 65.7 | P | --- |
| 248457_at | 7.7 | A | 27.3 | A | 11.1 | A | 4.3 | A | --- |
| 248458_at | 541.6 | P | 593.3 | P | 520.4 | P | 593.5 | P | EYE |
| 248459_at | 1420.2 | P | 1476.3 | P | 1445.5 | P | 1372.5 | P | CRL |
| 248460_at | 984.9 | P | 1267.7 | P | 1104.1 | P | 1111.8 | P | --- |
| 248461_s_at | 3026.2 | P | 2829.8 | P | 3268.8 | P | 3173.9 | P | FUM1 |
| 248462_at | 1010.6 | P | 792.7 | P | 766.8 | P | 859.7 | P | NBP35 |
| 248463_at | 185.3 | P | 136.5 | P | 177.4 | P | 127.4 | P | --- |
| 248464_at | 23.2 | A | 31.9 | A | 41.9 | A | 32 | A | --- |
| 248465_at | 2098.7 | P | 789.2 | P | 1808.4 | P | 1102.3 | P | --- |
| 248466_at | 170.1 | P | 191.6 | P | 197.2 | P | 111.7 | P | HVA22E |
| 248467_at | 332.6 | P | 470.1 | P | 471.5 | P | 410.8 | P | --- |
| 248468_at | 81.1 | A | 43.5 | A | 154.1 | A | 50 | A | RGP4 |
| 248469_at | 8.3 | A | 9.1 | A | 51.5 | A | 30.1 | A | anac097 |
| 248470_at | 28.7 | A | 3.4 | A | 29.5 | A | 2.6 | A | --- |
| 248471_at | 435.6 | P | 395.9 | P | 421.1 | P | 431.5 | P | --- |
| 248472_at | 335.4 | P | 387.3 | P | 345.7 | P | 350.2 | P | --- |
| 248473_at | 1552.8 | P | 1279.7 | P | 1806.5 | P | 1345.7 | P | TIM8 |
| 248474_at | 2472.6 | P | 2467.6 | P | 2592.6 | P | 2446.7 | P | MAB1 |
| 248475_at | 0.6 | A | 1.6 | A | 1.8 | A | 3 | A | --- |
| 248476_at | 98.8 | P | 104.2 | P | 107.4 | P | 110.3 | P | --- |
| 248477_at | 470.5 | P | 489.1 | P | 482.8 | P | 570.2 | P | --- |
| 248478_at | 1239.1 | P | 1305.4 | P | 1285.1 | P | 1406 | P | UBC27 |
| 248479_at | 0.7 | A | 1.3 | A | 19.8 | A | 2.4 | A | --- |
| 248480_at | 9053.8 | P | 7790.6 | P | 6883 | P | 7751.4 | P | CLPC1 |
| 248481_at | 117.8 | P | 114.2 | P | 127.2 | P | 100.6 | P | --- |
| 248482_at | 65.2 | P | 49.3 | P | 65.2 | P | 50 | P | --- |
| 248483_at | 51.4 | P | 66.7 | P | 62.6 | P | 72.9 | P | --- |
| 248484_at | 45.9 | A | 41.9 | A | 68.3 | A | 50.1 | P | --- |
| 248485_at | 429.7 | P | 508.8 | P | 471.4 | P | 470.9 | P | --- |
| 248486_at | 161.6 | P | 7.5 | A | 137.8 | A | 47.1 | A | RHD2 |
| 248487_at | 3956.9 | P | 2373.5 | P | 2354.1 | P | 1904.8 | P | ERD1 |
| 248488_at | 44.8 | A | 45.6 | P | 79.7 | A | 52.1 | P | --- |
| 248489_at | 1.9 | A | 1.7 | A | 7.1 | A | 20.5 | A | --- |
| 248490_at | 10.5 | A | 22.2 | A | 24.4 | M | 21.9 | A | --- |
| 248491_at | 3395.3 | P | 2041.1 | P | 2551.9 | P | 2196.6 | P | --- |
| 248492_at | 1708.2 | P | 1565.5 | P | 2017.9 | P | 1706.9 | P | --- |
| 248493_at | 929 | P | 1064 | P | 931.4 | P | 1193.3 | P | FSD2 |

| | | | | | | | | | |
|---|---|---|---|---|---|---|---|---|---|
| 248494_at | 38 | A | 51.9 | P | 33.1 | A | 13.4 | A | AtHSD6 |
| 248495_at | 169.5 | P | 101.6 | P | 181.6 | P | 115.4 | P | --- |
| 248496_at | 148.1 | P | 357.6 | P | 168.8 | P | 273.8 | P | --- |
| 248497_at | 539.1 | P | 527.4 | P | 453.9 | P | 441.2 | P | ATEXO70F1 |
| 248498_at | 62.5 | P | 72 | P | 61.6 | M | 45.6 | P | --- |
| 248499_at | 580.5 | P | 628.2 | P | 775.3 | P | 711.2 | P | PAP27 |
| 248500_at | 326.8 | P | 390.3 | P | 414 | P | 318.5 | P | --- |
| 248501_at | 211.8 | P | 136.3 | P | 182.6 | P | 145.6 | P | MEMB12 |
| 248502_at | 175.5 | P | 282.8 | P | 149.7 | P | 299.6 | P | --- |
| 248503_at | 1.6 | A | 3.9 | A | 7.8 | A | 2.9 | A | NF-YC7 |
| 248504_at | 35.9 | A | 59.2 | M | 78.1 | M | 21.4 | A | NF-YC6 |
| 248505_at | 61.1 | A | 34.5 | A | 6.1 | A | 9.3 | A | --- |
| 248506_at | 948.7 | P | 636.9 | P | 1145.5 | P | 781.2 | P | --- |
| 248507_at | 373.9 | P | 450.2 | P | 404.5 | P | 490.8 | P | --- |
| 248508_at | 808.3 | P | 840.5 | P | 727.1 | P | 808.6 | P | ELO3 |
| 248509_at | 131.3 | P | 251.7 | P | 72.7 | P | 106.8 | P | --- |
| 248510_at | 250.8 | P | 274.1 | P | 197.8 | M | 267.5 | P | --- |
| 248511_at | 307.7 | P | 605.6 | P | 378.1 | P | 518.6 | P | CPI1 |
| 248512_at | 3426.9 | P | 3908.5 | P | 3563.7 | P | 4029.7 | P | SIG2 |
| 248513_at | 1 | A | 2.3 | A | 3 | A | 0.8 | A | --- |
| 248514_s_at | 7.8 | A | 2.9 | A | 32.9 | A | 3.6 | A | --- |
| 248515_at | 486.9 | P | 446.2 | P | 369.5 | P | 314.5 | P | --- |
| 248516_at | 28.7 | A | 39.2 | P | 50.9 | P | 67.8 | P | --- |
| 248517_at | 433.6 | P | 359.9 | P | 453.8 | P | 315.1 | P | --- |
| 248518_at | 37.4 | A | 44.3 | A | 63.7 | A | 31.7 | A | --- |
| 248519_at | 3 | A | 28.3 | A | 21.4 | A | 2.8 | A | AtHSD7 |
| 248520_at | 46.1 | P | 8.1 | A | 30.3 | A | 25.8 | P | AtHSD1 |
| 248521_s_at | 478 | P | 531.8 | P | 513.3 | P | 436.1 | P | --- |
| 248522_at | 210.1 | P | 184.9 | P | 207.9 | P | 200.7 | P | --- |
| 248523_s_at | 829.5 | P | 704.4 | P | 736.4 | P | 761.2 | P | SAE1B |
| 248524_s_at | 1379.1 | P | 785.1 | P | 1148.6 | P | 607.5 | P | --- |
| 248525_s_at | 2.9 | A | 18.7 | A | 26.8 | A | 29.1 | A | --- |
| 248526_at | 227.7 | P | 380.2 | P | 267.5 | P | 427.9 | P | --- |
| 248527_at | 200.5 | P | 209.1 | P | 218.4 | P | 244.8 | P | --- |
| 248528_at | 851 | P | 104.4 | P | 619.4 | P | 115.7 | P | --- |
| 248529_at | 679.7 | P | 890.2 | P | 715.7 | P | 834.9 | P | --- |
| 248530_at | 258.3 | P | 349.7 | P | 386.6 | P | 468.1 | P | --- |
| 248531_at | 640.8 | P | 551.7 | P | 645.8 | P | 545.3 | P | --- |
| 248532_at | 2850.6 | P | 2514.3 | P | 2363.8 | P | 2415 | P | NFU2 |
| 248533_at | 225.4 | P | 236.6 | P | 226.6 | P | 261.8 | P | --- |
| 248534_at | 56.7 | A | 49.9 | A | 59.7 | A | 64.5 | A | --- |
| 248535_at | 29.2 | A | 43.9 | A | 66.5 | A | 39.7 | A | --- |
| 248536_at | 30 | A | 36.8 | A | 59.9 | A | 34.3 | M | --- |
| 248537_at | 843.3 | P | 611.6 | P | 633.6 | P | 637.1 | P | --- |
| 248538_at | 577.7 | P | 491.2 | P | 527.5 | P | 521.3 | P | --- |
| 248539_at | 564.4 | P | 202 | P | 539.7 | P | 202.8 | P | --- |
| 248540_at | 1042.4 | P | 950.9 | P | 1231.6 | P | 1078 | P | FRO8 |
| 248541_at | 487.5 | P | 609.6 | P | 531.9 | P | 576.8 | P | --- |
| 248542_at | 19.3 | A | 6.2 | A | 29.6 | A | 6.9 | A | --- |
| 248543_at | 489.9 | P | 411.5 | P | 486.9 | P | 435.4 | P | --- |
| 248544_at | 182.9 | P | 222.4 | P | 348.7 | P | 204.7 | P | PIMT2 |
| 248545_at | 21.1 | A | 26.2 | A | 46.2 | A | 11.8 | A | --- |
| 248546_at | 36.9 | P | 28 | P | 54.4 | P | 25.8 | P | --- |
| 248547_at | 329 | P | 429.5 | P | 376.7 | P | 391.6 | P | EMB1006 |
| 248548_at | 24 | A | 33 | P | 27.9 | A | 34.8 | P | --- |
| 248549_at | 600.7 | P | 710.2 | P | 921.2 | P | 674.5 | P | --- |
| 248550_at | 587.7 | P | 785.7 | P | 638.8 | P | 817.6 | P | QS |
| 248551_at | 124.2 | P | 593.8 | P | 137.5 | P | 437.9 | P | WR3 |
| 248552_at | 2025.2 | P | 2396.2 | P | 1513.1 | P | 2334.3 | P | --- |
| 248553_at | 114.5 | P | 127.6 | P | 138.1 | P | 118.3 | P | --- |
| 248554_at | 158.5 | P | 119.7 | P | 132.2 | P | 108.4 | P | --- |
| 248555_at | 239.7 | P | 334.4 | P | 247.2 | P | 348.1 | P | --- |
| 248556_at | 187.7 | P | 214.4 | P | 228.3 | P | 191.7 | P | --- |
| 248557_at | 1263.4 | P | 1578.1 | P | 1372.3 | P | 1660.9 | P | AFB5 |
| 248558_at | 173.6 | P | 230 | P | 253.4 | P | 216 | P | --- |
| 248559_at | 26.9 | A | 92.9 | P | 31.6 | A | 60.1 | P | --- |
| 248560_at | 1427.5 | P | 1451.1 | P | 1307.7 | P | 1251.9 | P | ATPPOX |
| 248561_at | 21.1 | A | 21.3 | A | 53.4 | A | 29.8 | A | --- |
| 248562_at | 50.6 | P | 80.3 | P | 102.5 | P | 46 | P | --- |
| 248563_at | 26.5 | A | 4.1 | A | 58.9 | A | 12.1 | A | --- |
| 248564_at | 90.1 | P | 96.3 | P | 63.2 | A | 80.9 | P | --- |
| 248565_at | 263.3 | P | 270.1 | P | 198.4 | P | 255 | P | --- |
| 248566_s_at | 754.2 | P | 1268.9 | P | 1083.3 | P | 932.2 | P | ATFRO7 |
| 248567_at | 27.7 | A | 26.6 | A | 51.9 | A | 45.3 | A | --- |
| 248568_at | 708.3 | P | 917.7 | P | 495.3 | P | 729.5 | P | --- |
| 248569_at | 3.6 | A | 2.4 | A | 2.5 | A | 2.8 | A | --- |
| 248570_at | 22.4 | A | 20.3 | A | 10.5 | A | 16.9 | A | --- |
| 248571_at | 21.8 | P | 6.7 | A | 1.8 | A | 3.7 | A | AT5G49780 |
| 248572_at | 27.4 | A | 70.6 | A | 36.2 | A | 64.3 | A | --- |
| 248573_at | 3051.2 | P | 3303.4 | P | 3023 | P | 3350.1 | P | ATGH9A1 |
| 248574_at | 591.1 | P | 673.9 | P | 551.4 | P | 638.7 | P | --- |
| 248575_at | 720 | P | 580.2 | P | 653.4 | P | 463.1 | P | --- |
| 248576_at | 1250.3 | P | 1463.3 | P | 1064.7 | P | 1496.1 | P | MMT |
| 248577_at | 2.2 | A | 2.6 | A | 13.9 | A | 12.7 | A | --- |
| 248578_at | 619.4 | P | 510.6 | P | 647.6 | P | 453.7 | P | emb1879 |
| 248579_at | 367 | P | 307 | P | 391.6 | P | 321 | P | --- |
| 248580_at | 2351.8 | P | 1229.1 | P | 1953.3 | P | 1402.6 | P | CLC-C |
| 248581_at | 144.1 | P | 215.3 | P | 159.5 | P | 213.7 | P | --- |
| 248582_at | 2014.8 | P | 2822.7 | P | 2640.8 | P | 3221.9 | P | CPHSC70-2EAT SHOCK PROTEIN 70-2 |
| 248583_at | 505 | P | 429.5 | P | 422.7 | P | 399.2 | P | emb1441 |
| 248584_at | 82.6 | P | 134.1 | P | 143.4 | P | 102.2 | P | --- |
| 248585_at | 40 | P | 34.8 | M | 44.8 | P | 29.9 | A | --- |
| 248586_at | 233 | P | 174.7 | P | 146.3 | P | 172.9 | P | --- |
| 248587_at | 768.5 | P | 651.2 | P | 736.6 | P | 700 | P | --- |
| 248588_at | 1391 | P | 1730.8 | P | 1415.3 | P | 1738.1 | P | --- |
| 248589_at | 1247.5 | P | 1210.7 | P | 1042.1 | P | 1257.3 | P | PDF3 |
| 248590_at | 391.1 | P | 426.2 | P | 525.2 | P | 438.2 | P | --- |
| 248591_at | 969.1 | P | 985.8 | P | 1043.4 | P | 905.9 | P | --- |
| 248592_at | 802.6 | P | 247.1 | P | 646 | P | 411.5 | P | --- |
| 248593_at | 27.6 | A | 15.7 | A | 5.6 | A | 32.6 | A | --- |
| 248594_at | 669.7 | P | 459.8 | P | 392.4 | P | 589 | P | --- |
| 248595_at | 578.5 | P | 553.5 | P | 600.9 | P | 535.5 | P | HRB1 |
| 248596_at | 224.8 | P | 248.6 | P | 273.6 | P | 363.5 | P | AtMYB111 |
| 248597_at | 400.7 | P | 385.3 | P | 550.3 | P | 425.5 | P | MET1 |
| 248598_at | 2.3 | A | 2.8 | A | 11.8 | A | 9.5 | A | --- |
| 248599_at | 6 | A | 55.7 | A | 5.7 | A | 10.7 | A | --- |
| 248600_at | 85.3 | P | 55.7 | P | 60 | P | 45.9 | P | --- |
| 248601_at | 366.9 | P | 372.6 | P | 363.4 | P | 303.4 | P | --- |
| 248602_s_at | 8.1 | A | 0.6 | A | 6.3 | A | 1.1 | A | --- |

| | | | | | | | | | |
|---|---|---|---|---|---|---|---|---|---|
| 248603_at | 117.3 | P | 139.1 | P | 128 | P | 131.3 | P | --- |
| 248604_at | 92 | P | 102.5 | P | 77.7 | A | 117.6 | P | --- |
| 248605_at | 144 | P | 150.7 | P | 180.6 | P | 152.6 | P | --- |
| 248606_at | 232.3 | P | 228.4 | P | 167 | P | 194.9 | P | CPuORF4 |
| 248607_at | 2467.7 | P | 2956.5 | P | 2534.7 | P | 3887.4 | P | ATCP1 |
| 248608_at | 728.9 | P | 999.1 | P | 878.6 | P | 1081.6 | P | ACLB-2 |
| 248609_at | 60.4 | P | 69.9 | P | 90 | P | 76.3 | P | --- |
| 248610_at | 102.9 | P | 148.5 | P | 136.7 | P | 132.5 | P | --- |
| 248611_at | 107.9 | P | 117.2 | P | 88.2 | M | 99 | P | WRKY48 |
| 248612_at | 493.1 | P | 491.5 | P | 536.9 | P | 515.1 | P | --- |
| 248613_at | 442.4 | P | 630.5 | P | 521 | P | 656.2 | P | --- |
| 248614_at | 185 | P | 103.2 | P | 203.8 | P | 108.6 | P | --- |
| 248615_at | 420.6 | P | 505.8 | P | 314.9 | P | 413.1 | P | AtPNG1 |
| 248616_at | 488.1 | P | 451.6 | P | 454.7 | P | 375.7 | P | --- |
| 248617_at | 5.7 | A | 0.1 | A | 0.3 | A | 1.8 | A | --- |
| 248618_at | 16.7 | A | 15.4 | A | 3 | A | 10.5 | A | AtMYB78 |
| 248619_at | 306.3 | P | 664.8 | P | 287.1 | P | 550.2 | P | AAP6 |
| 248620_at | 57.9 | P | 77 | P | 84.7 | A | 55.7 | P | --- |
| 248621_at | 16.4 | A | 17.5 | A | 4.9 | A | 4.2 | A | --- |
| 248622_at | 228 | P | 543.1 | P | 229.5 | P | 262.9 | P | --- |
| 248623_at | 37 | A | 74 | P | 69.6 | A | 58 | P | --- |
| 248624_at | 1633.6 | P | 1690.4 | P | 1598.3 | P | 1471.9 | P | --- |
| 248625_at | 1619.4 | P | 1183.2 | P | 1613.6 | P | 1670.9 | P | PKT2 |
| 248626_at | 8.4 | A | 22.6 | A | 39.3 | A | 28.3 | A | --- |
| 248627_at | 79.7 | P | 82.7 | P | 72.2 | P | 70.1 | P | --- |
| 248628_at | 613.7 | P | 589.6 | P | 580.6 | P | 599.7 | P | --- |
| 248629_at | 380.8 | P | 380.7 | P | 386.2 | P | 391.7 | P | --- |
| 248630_at | 158.8 | P | 156.2 | P | 224.2 | P | 163.7 | P | --- |
| 248631_at | 232.3 | P | 185.2 | P | 256.5 | P | 220.4 | P | --- |
| 248632_at | 191.9 | P | 241.9 | P | 215.2 | P | 247.4 | P | SLD5 |
| 248633_at | 190.5 | P | 249.7 | P | 256.3 | P | 220.1 | P | PRMT4A |
| 248634_at | 848.9 | P | 1113.3 | P | 910.4 | P | 1058.8 | P | OVA2 |
| 248635_at | 20.3 | A | 29.9 | A | 20.9 | A | 16.6 | A | --- |
| 248636_at | 2.4 | A | 4.4 | A | 24.1 | A | 2.3 | A | --- |
| 248637_at | 124.9 | P | 106.4 | P | 144.7 | P | 132 | P | --- |
| 248638_at | 18.6 | A | 17.5 | A | 37 | A | 22.3 | M | KCS21 |
| 248639_at | 2564.2 | P | 2583.6 | P | 2876.3 | P | 2606.1 | P | HCT |
| 248640_at | 270.4 | P | 284 | P | 244.7 | P | 332.2 | P | --- |
| 248641_at | 38.6 | P | 36.4 | A | 30.6 | A | 21.8 | A | --- |
| 248642_at | 78.9 | P | 51.6 | P | 85.7 | P | 55 | P | --- |
| 248643_at | 44.4 | A | 37.5 | P | 57.5 | A | 39 | M | --- |
| 248644_at | 31.8 | A | 28 | A | 6.8 | A | 1.3 | A | --- |
| 248645_at | 19.8 | A | 20.8 | A | 2.9 | A | 7 | A | GEX2 |
| 248646_at | 229.2 | P | 190.2 | P | 239.5 | P | 226.9 | P | --- |
| 248647_at | 45.4 | A | 35.4 | A | 72.1 | A | 45.4 | A | SUS2 |
| 248648_at | 191.8 | P | 212.9 | P | 256.4 | P | 321.2 | P | --- |
| 248649_at | 1.4 | A | 5.5 | A | 3 | A | 5.5 | A | APRR4 |
| 248650_at | 12.3 | A | 3.2 | A | 6.3 | A | 2.5 | A | --- |
| 248651_at | 32 | A | 25.7 | A | 24.7 | A | 20.2 | A | --- |
| 248652_at | 50.9 | P | 44 | A | 53.4 | A | 41.4 | A | SHV2 |
| 248653_at | 13.4 | A | 41.9 | A | 41.8 | A | 5.7 | A | --- |
| 248654_at | 69.6 | A | 47.4 | A | 17.3 | A | 11.3 | A | IMPA-5 |
| 248655_at | 1745.6 | P | 1744.3 | P | 1912.8 | P | 1850.3 | P | --- |
| 248656_at | 96.4 | P | 216.5 | P | 124.7 | P | 179.8 | P | --- |
| 248657_at | 159.4 | P | 403.4 | P | 178.8 | P | 341 | P | ROF2 |
| 248658_at | 98.5 | P | 161.3 | P | 139.5 | P | 186.9 | P | ATSMC3 |
| 248659_at | 327.5 | P | 446.6 | P | 351.8 | P | 366.3 | P | --- |
| 248660_at | 5.7 | A | 26.5 | A | 19.2 | A | 24.5 | A | --- |
| 248661_at | 93.2 | A | 78.7 | A | 104.2 | A | 101.2 | A | AGL80 |
| 248662_at | 1.7 | A | 1.8 | A | 3.7 | A | 20.4 | A | --- |
| 248663_at | 819.4 | P | 909 | P | 942.2 | P | 855.8 | P | --- |
| 248664_at | 1282.7 | P | 1875.4 | P | 1519.9 | P | 1845.8 | P | FKBP15-2 |
| 248665_at | 968 | P | 886.6 | P | 903 | P | 956.1 | P | --- |
| 248666_at | 773.2 | P | 774.5 | P | 922.4 | P | 713.8 | P | --- |
| 248667_at | 22.6 | A | 12.4 | A | 11.9 | A | 5.2 | A | --- |
| 248668_at | 203.7 | P | 208.8 | P | 203.8 | P | 201.2 | P | XRI1 |
| 248669_at | 375.3 | P | 398.2 | P | 361.3 | P | 527.5 | P | --- |
| 248670_at | 57.1 | A | 29.2 | A | 47.9 | P | 37.8 | P | --- |
| 248671_at | 1.3 | A | 32.4 | A | 4.7 | A | 2.7 | A | --- |
| 248672_at | 2.7 | A | 1.2 | A | 2 | A | 2.2 | A | --- |
| 248673_at | 27.5 | P | 3.8 | A | 7 | A | 4.1 | A | --- |
| 248674_at | 89.9 | P | 101.8 | P | 111.4 | P | 97.7 | P | --- |
| 248675_at | 56.6 | P | 92.7 | P | 136.4 | P | 86.7 | P | ICK6 |
| 248676_at | 175.1 | P | 21.3 | A | 403.6 | P | 36.4 | P | ATSDI1 |
| 248677_at | 1.1 | A | 11 | A | 6.3 | A | 22.6 | A | --- |
| 248678_at | 615.7 | P | 655.6 | P | 657.6 | P | 748.7 | P | SAD1 |
| 248679_at | 416.5 | P | 543.9 | P | 489.5 | P | 572.3 | P | --- |
| 248680_at | 419.6 | P | 304.4 | P | 391 | P | 440.4 | P | PANC |
| 248681_at | 276.7 | P | 940.2 | P | 249.5 | P | 769.4 | P | --- |
| 248682_at | 3236.4 | P | 3972.5 | P | 3255 | P | 3608.8 | P | CB5-D |
| 248683_at | 83.6 | P | 653 | P | 252.7 | P | 507.8 | P | --- |
| 248684_at | 553.4 | P | 1398.8 | P | 639.1 | P | 1396.1 | P | DIR1 |
| 248685_at | 144.9 | P | 202.7 | P | 228.6 | P | 198.7 | P | --- |
| 248686_at | 105.1 | P | 220.9 | P | 84.5 | P | 170.2 | P | --- |
| 248687_at | 3534.2 | P | 4353.7 | P | 4149.7 | P | 4755.3 | P | ADG1 |
| 248688_at | 422.6 | P | 872.8 | P | 443 | P | 633.4 | P | --- |
| 248689_at | 822.8 | P | 741.4 | P | 888.6 | P | 749.1 | P | PAT1 |
| 248690_at | 936.9 | P | 884.1 | P | 996.3 | P | 932.1 | P | ACAT2 |
| 248691_at | 138 | P | 233.3 | P | 172.1 | P | 222.7 | P | --- |
| 248692_s_at | 16.5 | A | 31.8 | P | 27.1 | P | 27.5 | P | --- |
| 248693_at | 196.5 | P | 251 | P | 223.3 | P | 298.3 | P | --- |
| 248694_at | 378.9 | P | 300.6 | P | 337.7 | P | 318.4 | P | --- |
| 248695_at | 2.9 | A | 3.5 | A | 7.8 | A | 1.1 | A | --- |
| 248696_at | 186.6 | P | 164.1 | P | 254.7 | P | 205.4 | P | --- |
| 248697_at | 348 | P | 374 | P | 274.4 | P | 351 | P | --- |
| 248698_at | 1811 | P | 2128.4 | P | 1906.2 | P | 1612.2 | P | --- |
| 248699_at | 2.8 | A | 4.3 | A | 3.6 | A | 3.2 | A | ATZIP4 |
| 248700_at | 37.6 | P | 49.4 | P | 66.2 | P | 28.9 | P | ATGLR1.2 |
| 248701_at | 181.6 | P | 153.8 | P | 105.6 | P | 143.3 | P | GLR1.3 |
| 248702_at | 44.5 | A | 34.1 | A | 82.9 | A | 29.9 | A | --- |
| 248703_at | 12.6 | A | 37.6 | M | 24.9 | A | 25.2 | A | --- |
| 248704_at | 282.2 | P | 422.2 | P | 410 | P | 391.2 | P | sks3 |
| 248705_at | 157.8 | P | 245.6 | P | 203.7 | P | 181.9 | P | --- |
| 248706_at | 56 | A | 34.8 | A | 34.7 | A | 29.3 | A | --- |
| 248707_at | 13.1 | A | 35.5 | A | 25.3 | A | 8.1 | A | --- |
| 248708_at | 3.2 | A | 21.6 | P | 1.7 | A | 4.3 | A | --- |
| 248709_at | 635 | P | 672.3 | P | 803 | P | 724.2 | P | --- |
| 248710_at | 588.4 | P | 900.1 | P | 780.3 | P | 958.7 | P | --- |
| 248711_at | 18.5 | A | 7.4 | A | 31.2 | A | 13.7 | A | --- |

| | | | | | | | | | |
|---|---|---|---|---|---|---|---|---|---|
| 248712_at | 3.5 | A | 2.4 | A | 5.3 | A | 1.9 | A | --- |
| 248713_at | 3217.2 | P | 2015.4 | P | 2143.8 | P | 2480.2 | P | NSP5 |
| 248714_at | 11.8 | P | 9 | A | 12.2 | A | 3 | A | --- |
| 248715_at | 21.9 | A | 85.5 | P | 7.8 | A | 64.3 | P | --- |
| 248716_at | 41.3 | A | 28.3 | A | 37.7 | A | 33.2 | M | --- |
| 248717_at | 21.6 | A | 32.4 | A | 28.8 | A | 18.4 | A | --- |
| 248718_at | 818.7 | P | 1152.5 | P | 942 | P | 1164.7 | P | FPS1 |
| 248719_at | 1486 | P | 1170.3 | P | 1553.4 | P | 1140.7 | P | RBOHD |
| 248720_at | 399.4 | P | 437.5 | P | 471.8 | P | 459.1 | P | D6PKL2 |
| 248721_at | 1852.5 | P | 1849.2 | P | 1824.7 | P | 1885.1 | P | GAUT4 |
| 248722_at | 444.4 | P | 166.7 | P | 261.7 | P | 203.8 | P | PFK2 |
| 248723_at | 42.8 | A | 36.3 | A | 41.3 | A | 7.2 | A | --- |
| 248724_at | 240.1 | P | 263.1 | P | 296.5 | P | 251.4 | P | --- |
| 248725_at | 1.9 | A | 32.5 | A | 6.2 | A | 12.8 | A | --- |
| 248726_at | 109.1 | P | 92.3 | P | 103.1 | P | 65 | P | ATRABA4C |
| 248727_at | 32.3 | P | 45 | P | 33.6 | A | 42.7 | P | CYP705A5 |
| 248728_at | 8 | A | 47.7 | A | 67.5 | A | 53 | A | CYP708A2 |
| 248729_at | 60.6 | A | 44.3 | P | 77 | A | 48.4 | A | THAS1 |
| 248730_at | 20.2 | A | 20.5 | A | 34.3 | A | 44.3 | A | --- |
| 248731_at | 29.8 | A | 82.5 | P | 68.1 | A | 74.4 | P | --- |
| 248732_at | 23.5 | M | 34.2 | P | 24.7 | A | 35.7 | P | XTH20 |
| 248733_at | 11.4 | A | 23.9 | A | 48.7 | A | 13.2 | A | ELP1 |
| 248734_at | 38.8 | A | 24.1 | A | 40 | P | 48.8 | A | ELP1 |
| 248735_at | 52.8 | M | 17.4 | A | 61.5 | A | 17.3 | A | TT10 |
| 248736_at | 17 | A | 15.9 | A | 12.2 | A | 3.3 | A | --- |
| 248737_at | 609 | P | 329.1 | P | 593.2 | P | 414.8 | P | --- |
| 248738_at | 814.5 | P | 831.6 | P | 757.8 | P | 741.5 | P | --- |
| 248739_at | 1197.7 | P | 895.6 | P | 1233.6 | P | 1048.5 | P | GFA2 |
| 248740_at | 36.3 | A | 26.8 | A | 44.6 | A | 56.4 | A | --- |
| 248741_at | 72 | P | 48.8 | P | 105.9 | P | 61.8 | P | SLY2 |
| 248742_at | 2.7 | A | 1.6 | A | 28.6 | A | 20.2 | A | --- |
| 248743_at | 683.8 | P | 414.5 | P | 454.6 | P | 563.8 | P | --- |
| 248744_at | 737.1 | P | 609 | P | 684.3 | P | 672.9 | P | --- |
| 248745_at | 154.9 | P | 129.3 | P | 153.4 | P | 198.8 | P | --- |
| 248746_at | 1932.3 | P | 1788.8 | P | 1980.1 | P | 2031.5 | P | --- |
| 248747_at | 5395.5 | P | 5271.7 | P | 7616.3 | P | 5320.6 | P | --- |
| 248748_at | 2617.5 | P | 3009.3 | P | 3173.1 | P | 3040.4 | P | AMK2 |
| 248749_at | 1681.2 | P | 992.3 | P | 1071.1 | P | 1099.9 | P | ERF1-1 |
| 248750_at | 28.3 | A | 49.1 | P | 44.4 | P | 44.8 | P | --- |
| 248751_at | 728.4 | P | 469.3 | P | 574.5 | P | 457.5 | P | --- |
| 248752_at | 31.3 | A | 4.5 | A | 19.6 | A | 25 | A | --- |
| 248753_at | 186 | P | 181.8 | P | 232.4 | P | 244 | P | mtACP3 |
| 248754_at | 2 | A | 9.1 | A | 2.8 | A | 4.4 | A | NF-YB6 |
| 248755_at | 194.6 | P | 227.8 | P | 282.3 | P | 207.7 | P | --- |
| 248756_at | 3744.8 | P | 1910.5 | P | 2778.6 | P | 2065.7 | P | TDT |
| 248757_at | 1072.9 | P | 799 | P | 999 | P | 828.4 | P | --- |
| 248758_at | 1208.5 | P | 1085.2 | P | 1068.9 | P | 1301.4 | P | --- |
| 248759_at | 511.1 | P | 273.1 | P | 438.1 | P | 248.3 | P | --- |
| 248760_at | 6511.8 | P | 5321.9 | P | 5454 | P | 5702.3 | P | --- |
| 248761_at | 37.6 | A | 40.1 | A | 36.3 | A | 40.3 | A | --- |
| 248762_at | 359.2 | P | 386.8 | P | 386.2 | P | 404.7 | P | --- |
| 248763_at | 1442.1 | P | 474 | P | 1182.6 | P | 710.3 | P | --- |
| 248764_at | 450.4 | P | 449.6 | P | 544 | P | 542.1 | P | NF-YB2 |
| 248765_at | 474.7 | P | 323 | P | 394.2 | P | 399.8 | P | ATNUDT2 |
| 248766_at | 209.9 | P | 281.2 | P | 204.5 | P | 215.8 | P | --- |
| 248767_at | 422.6 | P | 439.2 | P | 472.5 | P | 416.2 | P | --- |
| 248768_at | 6418.2 | P | 5519.1 | P | 6935.9 | P | 6092.8 | P | --- |
| 248769_at | 384.3 | P | 356.3 | P | 318.7 | P | 307.6 | P | --- |
| 248770_at | 3.6 | A | 14.3 | A | 5 | A | 3.3 | A | --- |
| 248771_at | 186.6 | P | 158.2 | P | 167 | P | 197.2 | P | --- |
| 248772_at | 138.5 | P | 130.9 | P | 122.5 | P | 152.4 | P | --- |
| 248773_at | 291.8 | P | 415.1 | P | 379.2 | P | 402.7 | P | FRA1 |
| 248774_at | 330.2 | P | 417.3 | P | 206.8 | P | 314.8 | P | --- |
| 248775_at | 7.2 | A | 5.7 | A | 5.7 | A | 4.9 | A | CCR4 |
| 248776_at | 44.2 | A | 98.4 | P | 99.6 | P | 87 | A | --- |
| 248777_at | 47.9 | P | 61.6 | P | 51.4 | A | 60 | P | --- |
| 248778_at | 120.2 | P | 123.5 | P | 128.1 | P | 144.4 | P | --- |
| 248779_at | 459.6 | P | 505.6 | P | 772.2 | P | 495.6 | P | --- |
| 248780_at | 641.8 | P | 581.2 | P | 700.1 | P | 634.9 | P | ATPGLP2 |
| 248781_at | 725.7 | P | 996.8 | P | 933.6 | P | 1074.5 | P | --- |
| 248782_at | 32.3 | P | 28.8 | P | 47.7 | P | 40.7 | P | --- |
| 248783_at | 134.2 | P | 99.5 | P | 94.5 | P | 135.7 | P | --- |
| 248784_at | 98 | P | 179.4 | P | 109.6 | P | 150 | P | --- |
| 248785_at | 133.4 | P | 133.1 | P | 159.2 | P | 144.8 | P | --- |
| 248786_at | 230.8 | P | 176.8 | P | 184 | P | 175.3 | P | --- |
| 248787_at | 451.4 | P | 446.3 | P | 449.6 | P | 466.8 | P | --- |
| 248788_at | 182.6 | P | 192.8 | P | 205.8 | P | 175.1 | P | --- |
| 248789_at | 63.5 | A | 54.9 | A | 76.9 | A | 37.8 | A | --- |
| 248790_at | 29.1 | A | 28 | A | 21.7 | A | 34.2 | A | TIP2;3 |
| 248791_at | 17.6 | A | 22.3 | P | 45.8 | P | 17.4 | A | --- |
| 248792_at | 3123.9 | P | 2512.7 | P | 2809.3 | P | 2456.1 | P | ATRAB1A |
| 248793_at | 1343.3 | P | 986.6 | P | 852 | P | 885.7 | P | atnudt8 |
| 248794_at | 403.1 | P | 411.4 | P | 484.1 | P | 331.1 | P | ERF2 |
| 248795_at | 410.6 | P | 591.8 | P | 432.8 | P | 542.3 | P | --- |
| 248796_at | 581 | P | 499.8 | P | 370.7 | P | 452.1 | P | --- |
| 248797_at | 4545 | P | 3757.8 | P | 4040.5 | P | 4256.6 | P | --- |
| 248798_at | 2947.3 | P | 3735 | P | 3399.2 | P | 3265.8 | P | --- |
| 248799_at | 92 | P | 118.3 | P | 68.1 | M | 101 | P | ERF5 |
| 248800_at | 1415.9 | P | 1310.1 | P | 1241.7 | P | 1207.8 | P | RPS19 |
| 248801_at | 120 | P | 168.7 | P | 118.1 | P | 194.7 | P | HAT2 |
| 248802_at | 4225.1 | P | 3248.4 | P | 3694.3 | P | 3111.7 | P | --- |
| 248803_at | 97 | P | 88.3 | P | 137.6 | P | 89.9 | P | --- |
| 248804_at | 30 | A | 23.4 | A | 5.6 | A | 24.2 | A | --- |
| 248805_at | 467 | P | 485 | P | 412.2 | P | 477.3 | P | --- |
| 248806_at | 190.3 | P | 219 | P | 192.6 | P | 232.5 | P | --- |
| 248807_at | 393 | P | 725.6 | P | 337.9 | P | 838 | P | --- |
| 248808_at | 75.5 | P | 83.9 | P | 54.9 | A | 71.7 | P | --- |
| 248809_at | 208.9 | P | 229.7 | P | 206.1 | P | 250.7 | P | AtRABA5a |
| 248810_at | 11.8 | A | 8.9 | A | 14.1 | A | 3.3 | A | ADR1-L3 |
| 248811_at | 584.5 | P | 803.1 | P | 628.7 | P | 706.6 | P | --- |
| 248812_at | 1116.7 | P | 965.4 | P | 586 | P | 836.2 | P | --- |
| 248813_at | 1166.4 | P | 1193.5 | P | 1587.3 | P | 1170.4 | P | VAM3 |
| 248814_at | 86.8 | P | 71.1 | P | 98 | P | 108.3 | P | --- |
| 248815_at | 320.9 | P | 347.7 | P | 443.3 | P | 349.8 | P | --- |
| 248816_at | 506.9 | P | 475 | P | 545.6 | P | 455.7 | P | LBA1 |
| 248817_at | 787 | P | 745 | P | 668 | P | 714.8 | P | --- |
| 248818_at | 640.7 | P | 806 | P | 573 | P | 657.6 | P | LON2 |
| 248819_at | 448.1 | P | 281.7 | P | 505.9 | P | 333.2 | P | --- |
| 248820_at | 981.9 | P | 977.9 | P | 1092.2 | P | 926.5 | P | --- |

| | | | | | | | | | |
|---|---|---|---|---|---|---|---|---|---|
| 248821_at | 729 | P | 770 | P | 530.5 | P | 606.1 | P | --- |
| 248822_at | 3.7 | A | 26.6 | A | 19.7 | A | 15.9 | A | --- |
| 248823_s_at | 6.5 | A | 2.1 | A | 10.8 | A | 10.7 | A | --- |
| 248824_at | 17.8 | A | 4.2 | A | 25.7 | A | 10.1 | A | --- |
| 248825_at | 4882.4 | P | 4987.2 | P | 5114.1 | P | 5124.2 | P | --- |
| 248826_at | 731.3 | P | 696.7 | P | 807.2 | P | 641.1 | P | CKB1 |
| 248827_at | 219.3 | P | 297.8 | P | 235 | P | 281.9 | P | CBL9 |
| 248828_at | 3176.8 | P | 3316.2 | P | 2517 | P | 3117.4 | P | --- |
| 248829_at | 249.8 | P | 236 | P | 268.4 | P | 254.7 | P | --- |
| 248830_at | 11.3 | A | 2.9 | A | 15.5 | A | 19.8 | A | --- |
| 248831_at | 3.8 | A | 2.7 | A | 4.4 | A | 17.3 | A | --- |
| 248832_at | 16.5 | A | 10.9 | A | 36.3 | A | 28.5 | A | --- |
| 248833_at | 1659.8 | P | 2801.2 | P | 1546.9 | P | 2344.4 | P | ATBI1 |
| 248834_at | 247.1 | P | 221.1 | P | 304.1 | P | 239.3 | P | --- |
| 248835_at | 106.8 | P | 120.2 | P | 102.6 | A | 124 | P | --- |
| 248836_at | 16.2 | A | 29.3 | A | 38.3 | A | 37.6 | A | --- |
| 248837_at | 2.4 | A | 1.9 | A | 8.4 | A | 1.3 | A | --- |
| 248838_at | 1861.5 | P | 2079.7 | P | 2462.1 | P | 2155.5 | P | BOU |
| 248839_at | 156.4 | P | 182.8 | P | 192.1 | P | 194.9 | P | bHLH071 |
| 248840_at | 9.2 | A | 13.7 | A | 8.6 | A | 6.4 | A | --- |
| 248841_at | 126.6 | A | 189.4 | A | 173.6 | A | 205 | P | UBP21 |
| 248842_at | 182.3 | P | 193.1 | P | 251.1 | P | 217.2 | P | --- |
| 248843_at | 67.5 | A | 107.8 | A | 96.7 | A | 80.5 | M | HB-7 |
| 248844_s_at | 24.9 | A | 30.7 | A | 43.3 | A | 45.2 | A | --- |
| 248845_at | 233.2 | P | 185 | P | 163.8 | P | 154.4 | P | --- |
| 248846_at | 21.6 | M | 25.6 | P | 46.7 | P | 30.3 | A | --- |
| 248847_at | 72.8 | P | 114.2 | P | 70.1 | P | 107.5 | P | --- |
| 248848_at | 58.2 | P | 76.1 | P | 49.8 | A | 54.7 | P | --- |
| 248849_at | 4.6 | A | 3 | A | 4.3 | A | 25.5 | A | PGP7 |
| 248850_at | 276.8 | P | 265.5 | P | 366.5 | P | 262.5 | P | --- |
| 248851_s_at | 95.1 | P | 114.4 | P | 103.1 | P | 130.3 | P | --- |
| 248852_at | 294.3 | P | 361.3 | P | 229.4 | P | 424.7 | P | --- |
| 248853_at | 162.9 | P | 326.2 | P | 194.6 | P | 309.6 | P | BSK2 |
| 248854_at | 1178.8 | P | 1205.3 | P | 850.6 | P | 984.7 | P | --- |
| 248855_at | 98.3 | P | 46.6 | A | 49.1 | A | 42.9 | A | anac096 |
| 248856_at | 388.2 | P | 393.9 | P | 368.4 | P | 439.4 | P | --- |
| 248857_at | 81.5 | P | 42.4 | P | 104 | P | 37.3 | P | --- |
| 248858_at | 1225.1 | P | 1400.4 | P | 1298.6 | P | 1374.1 | P | --- |
| 248859_at | 15.8 | A | 9.2 | A | 36.8 | A | 2.9 | A | --- |
| 248860_at | 36.6 | A | 69.5 | A | 12.7 | A | 37.9 | A | --- |
| 248861_at | 427.2 | P | 1207.2 | P | 686 | P | 1159.7 | P | TRN2 |
| 248862_at | 802.3 | P | 404.8 | P | 916.2 | P | 491.7 | P | --- |
| 248863_at | 834.3 | P | 923.9 | P | 761.4 | P | 882.9 | P | AGD9 |
| 248864_at | 390.8 | P | 322.1 | P | 404.1 | P | 316.1 | P | --- |
| 248865_at | 521.5 | P | 589.8 | P | 642.7 | P | 553.4 | P | --- |
| 248866_s_at | 21 | A | 0.6 | A | 1.4 | A | 2.2 | A | --- |
| 248867_at | 10.4 | A | 2.1 | A | 20 | A | 6.4 | A | --- |
| 248868_at | 622.4 | P | 445.2 | P | 595.7 | P | 406.7 | P | --- |
| 248869_at | 369.3 | P | 321.7 | P | 470.5 | P | 376.2 | P | --- |
| 248870_at | 645.2 | P | 329 | P | 436.4 | P | 300.1 | P | --- |
| 248871_at | 133 | P | 117.9 | P | 118.1 | P | 134.5 | P | --- |
| 248872_at | 5.2 | A | 26.6 | A | 2.4 | A | 14 | A | MSP2 |
| 248873_at | 420 | P | 451.7 | P | 383.1 | P | 429.2 | P | --- |
| 248874_at | 138.4 | P | 117.4 | P | 123.1 | P | 99.9 | P | --- |
| 248875_at | 124.4 | P | 164.9 | P | 110.8 | P | 87 | P | --- |
| 248876_at | 5.2 | A | 5.2 | A | 3.7 | A | 12.5 | A | --- |
| 248877_at | 66.7 | P | 78.5 | P | 102.8 | P | 57.3 | P | --- |
| 248878_at | 1285.8 | P | 1000.5 | P | 1674.6 | P | 976.8 | P | --- |
| 248879_at | 1530 | P | 970.6 | P | 1045.7 | P | 878.8 | P | DELTA-OAT |
| 248880_at | 3.3 | A | 19.2 | A | 2.5 | A | 23.7 | A | --- |
| 248881_at | 3296.7 | P | 3536.8 | P | 2809.7 | P | 2867.1 | P | --- |
| 248882_at | 1716.4 | P | 1756.2 | P | 1926.4 | P | 1705.9 | P | CUL4 |
| 248883_at | 308.7 | P | 317.5 | P | 293.3 | P | 268.9 | P | --- |
| 248884_at | 2027.1 | P | 1686.9 | P | 1075.7 | P | 1752.1 | P | --- |
| 248885_at | 285.2 | P | 372.1 | P | 315.6 | P | 333.7 | P | --- |
| 248886_at | 12420 | P | 13508.7 | P | 12673.3 | P | 13918 | P | APE2 |
| 248887_at | 31.5 | A | 64 | A | 3.1 | A | 44 | A | --- |
| 248888_at | 292.6 | P | 212.8 | P | 290.2 | P | 247.5 | P | KAT1 |
| 248889_at | 205.3 | P | 253 | P | 129.1 | P | 231.1 | P | --- |
| 248890_at | 184.9 | P | 210.7 | P | 185.3 | P | 217.1 | P | --- |
| 248891_at | 288.9 | P | 319.5 | P | 284.8 | P | 303.4 | P | MCM3 |
| 248892_at | 1 | A | 5.1 | A | 22 | A | 0.9 | A | --- |
| 248893_at | 50.4 | A | 46 | A | 74.9 | A | 52.1 | A | --- |
| 248894_at | 14.6 | A | 6.9 | A | 53.2 | A | 25.3 | A | --- |
| 248895_at | 168.6 | P | 298.4 | P | 143 | P | 186.8 | P | FLS2 |
| 248896_at | 171 | P | 87.9 | P | 116.6 | P | 91.1 | P | WRKY8 |
| 248897_at | 36.7 | M | 11.2 | A | 27 | A | 29.5 | A | KCO3 |
| 248898_at | 38.7 | A | 9.1 | A | 57.6 | A | 17.2 | A | KCO2 |
| 248899_at | 510.3 | P | 458.1 | P | 534.3 | P | 472.3 | P | --- |
| 248900_at | 117.3 | P | 108.7 | P | 131.9 | P | 86.8 | P | PRP39-2 |
| 248901_at | 252.1 | P | 263.6 | P | 258.3 | P | 258.5 | P | --- |
| 248902_at | 19.7 | A | 16.9 | A | 24.9 | A | 33.3 | A | --- |
| 248903_at | 2593.6 | P | 2880.6 | P | 3232.9 | P | 2884.7 | P | KAS I |
| 248904_at | 21.7 | A | 10.1 | A | 15.7 | A | 10.1 | A | --- |
| 248905_at | 2101.6 | P | 1920.5 | P | 2198.2 | P | 1880.6 | P | --- |
| 248906_at | 1478.5 | P | 1810.6 | P | 1632 | P | 1755.9 | P | --- |
| 248907_at | 343 | P | 476.9 | P | 410.9 | P | 380.4 | P | --- |
| 248908_at | 941.1 | P | 1342.1 | P | 1050.4 | P | 1219.6 | P | MEE62 |
| 248909_at | 13.3 | A | 4.8 | A | 4.5 | A | 8.6 | A | CIPK19 |
| 248910_at | 98.1 | P | 495.7 | P | 122.2 | P | 332.7 | P | CIPK20 |
| 248911_at | 3.5 | A | 78.8 | A | 3.3 | A | 71.1 | A | DOG1 |
| 248912_at | 281.8 | P | 725 | P | 314.5 | P | 599.5 | P | --- |
| 248913_at | 330.7 | P | 369.6 | P | 450.8 | P | 309.6 | P | --- |
| 248914_at | 924 | P | 1263.3 | P | 953.3 | P | 1161.6 | P | AtRABA1c |
| 248915_at | 14.2 | A | 2.2 | A | 3.6 | A | 2.8 | A | --- |
| 248916_at | 153.5 | P | 164.3 | P | 108.8 | P | 127.7 | P | --- |
| 248917_at | 23.8 | A | 36.7 | A | 37.9 | A | 11.3 | A | --- |
| 248918_at | 1.9 | A | 10.4 | A | 2 | A | 10.7 | A | SAG12 |
| 248919_at | 73.8 | P | 71.2 | P | 70.2 | P | 64.1 | P | --- |
| 248920_at | 626.1 | P | 912.5 | P | 797.6 | P | 940.9 | P | CHLI2 |
| 248921_at | 1027.7 | P | 1055.6 | P | 1233.3 | P | 1120.6 | P | --- |
| 248922_at | 10.8 | A | 6.5 | A | 4.1 | A | 8.5 | A | WOX8 |
| 248923_at | 206.9 | P | 261.4 | P | 214.2 | P | 257.4 | P | atnudt11 |
| 248924_at | 15.6 | A | 17.9 | A | 56.7 | A | 13.9 | A | --- |
| 248925_at | 26.1 | A | 21.1 | A | 22.9 | A | 37 | P | --- |
| 248926_at | 5 | A | 3.1 | A | 4.3 | A | 3.1 | A | --- |
| 248927_at | 269.1 | P | 292.8 | P | 292.2 | P | 280.6 | P | APG7 |
| 248928_at | 21.6 | A | 33.8 | A | 8.4 | A | 27.3 | A | ARAC2 |
| 248929_at | 8.3 | A | 1.4 | A | 3.2 | A | 0.9 | A | --- |

| | | | | | | | | | |
|---|---|---|---|---|---|---|---|---|---|
| 248930_at | 2 | A | 16.6 | A | 28.6 | A | 7.2 | A | --- |
| 248931_at | 24.6 | A | 10.5 | A | 8.2 | A | 1.1 | A | --- |
| 248932_at | 123.7 | P | 219.7 | P | 71.1 | P | 112.2 | P | PTR3 |
| 248933_at | 527.9 | P | 546 | P | 568.5 | P | 561.6 | P | --- |
| 248934_at | 69.4 | P | 95.4 | P | 54.7 | P | 53.9 | P | --- |
| 248935_at | 94.1 | P | 117.2 | P | 110.4 | P | 73.1 | P | --- |
| 248936_at | 249.7 | P | 185.1 | P | 226.2 | P | 179.2 | P | RHA1 |
| 248937_at | 33.2 | A | 24.9 | A | 51.9 | A | 28.1 | A | AtRLP55 |
| 248938_at | 139.9 | A | 140.2 | A | 122.1 | A | 112.6 | M | --- |
| 248939_at | 43.7 | P | 31.2 | P | 43.2 | P | 27.2 | P | --- |
| 248940_at | 58.5 | P | 90.3 | P | 80.3 | P | 111.5 | P | --- |
| 248941_s_at | 163.2 | P | 219.2 | P | 136 | P | 248.1 | P | --- |
| 248942_at | 277.5 | P | 340.1 | P | 248 | P | 239.2 | P | --- |
| 248943_s_at | 282.1 | P | 377.3 | P | 294.8 | P | 343.8 | P | --- |
| 248944_at | 591.5 | P | 657.9 | P | 536.3 | P | 520.8 | P | --- |
| 248945_at | 1077 | P | 1310.6 | P | 920.4 | P | 1214.2 | P | --- |
| 248946_at | 19.8 | P | 35.5 | P | 17.4 | A | 32.9 | P | --- |
| 248947_at | 33 | P | 49.7 | P | 43.6 | A | 44.4 | P | --- |
| 248948_at | 57 | P | 101.5 | P | 69.3 | M | 122.4 | P | --- |
| 248949_at | 0.6 | A | 0.5 | A | 18.5 | A | 1.2 | A | --- |
| 248950_at | 5047 | P | 5521.3 | P | 4841.1 | P | 5070.6 | P | CLPP4 |
| 248951_at | 1089.3 | P | 1373.6 | P | 1291.8 | P | 1380.9 | P | --- |
| 248952_at | 2549.2 | P | 1515.2 | P | 2149 | P | 1594.5 | P | --- |
| 248953_at | 611.8 | P | 978.2 | P | 581.7 | P | 755.1 | P | --- |
| 248954_at | 329.7 | P | 449.3 | P | 400.6 | P | 415.8 | P | --- |
| 248955_at | 402.4 | P | 401.8 | P | 356.1 | P | 406 | P | GAS41 |
| 248956_at | 150.6 | P | 137.8 | P | 198.3 | P | 118.1 | P | --- |
| 248957_at | 1112.7 | P | 1195.4 | P | 1338.8 | P | 1148.1 | P | --- |
| 248958_at | 340.5 | P | 450.5 | P | 436.9 | P | 507.6 | P | --- |
| 248959_at | 20.7 | P | 48.8 | P | 49.9 | P | 52.6 | P | --- |
| 248960_at | 14.7 | A | 21.9 | A | 29.3 | A | 18.9 | A | --- |
| 248961_at | 142.1 | P | 840.5 | P | 186.4 | P | 624.1 | P | --- |
| 248962_at | 1149 | P | 2125.8 | P | 1254.1 | P | 1702.9 | P | FKBP13 |
| 248963_at | 70.1 | P | 149.8 | P | 96.6 | P | 185.7 | P | --- |
| 248964_at | 21.2 | A | 93.5 | P | 73.9 | P | 72.6 | P | CYP707A3 |
| 248965_at | 126.3 | P | 159 | P | 147.3 | P | 154.5 | P | --- |
| 248966_at | 628.9 | P | 664.5 | P | 521.6 | P | 832.9 | P | --- |
| 248967_at | 2794.4 | P | 3493.1 | P | 3458.3 | P | 3314.6 | P | --- |
| 248968_at | 1324.7 | P | 1158.5 | P | 698.8 | P | 852.8 | P | --- |
| 248969_at | 31.1 | A | 25.8 | A | 22.1 | A | 28.1 | A | --- |
| 248970_at | 378.8 | P | 733.2 | P | 326.3 | P | 622 | P | --- |
| 248971_at | 22.7 | A | 53.6 | P | 15.7 | A | 40.5 | P | --- |
| 248972_at | 1046.1 | P | 1043.8 | P | 784.4 | P | 1103.5 | P | ATDSS1(V) |
| 248973_at | 262.1 | P | 258.8 | P | 218.5 | P | 233.4 | P | TTR1 |
| 248974_at | 188.6 | P | 252.3 | P | 246 | P | 190.7 | P | --- |
| 248975_at | 366.8 | P | 414.9 | P | 438.9 | P | 488.8 | P | --- |
| 248976_at | 188.5 | P | 149 | P | 133.8 | P | 146.9 | P | ARAD2 |
| 248977_at | 153.4 | P | 198.7 | P | 186.3 | P | 160.8 | P | --- |
| 248978_at | 22.9 | A | 24.8 | A | 28.5 | A | 22.9 | A | AtPP2-A8 |
| 248979_at | 39 | A | 9.9 | A | 3.9 | A | 23.7 | A | AtPP2-A6 |
| 248980_at | 60.7 | A | 42.3 | A | 87.3 | A | 47.2 | P | AtPP2-A7 |
| 248981_at | 449.7 | P | 378.6 | P | 238 | P | 467.6 | P | NPR3 |
| 248982_at | 42.9 | P | 18.9 | P | 35.9 | P | 3.1 | A | --- |
| 248983_at | 2345.8 | P | 2244 | P | 1920.4 | P | 1717.1 | P | RHA1 |
| 248984_at | 471.3 | P | 290.7 | P | 578.3 | P | 330.4 | P | NRPC2 |
| 248985_at | 281.1 | P | 326.2 | P | 282.3 | P | 275 | P | --- |
| 248986_at | 1492.2 | P | 1494.6 | P | 1489.8 | P | 1394.1 | P | --- |
| 248987_at | 4 | A | 1.7 | A | 2.6 | A | 4.8 | A | --- |
| 248988_at | 376 | P | 437.9 | P | 447.4 | P | 467.3 | P | --- |
| 248989_at | 20.8 | A | 12.5 | A | 3.3 | A | 1.3 | A | --- |
| 248990_at | 1.1 | A | 2.1 | A | 2.8 | A | 16 | A | --- |
| 248991_at | 30 | A | 38.3 | M | 50 | P | 37.1 | A | --- |
| 248992_at | 5.2 | A | 42.1 | A | 5.5 | A | 4.4 | A | --- |
| 248993_at | 1.5 | A | 14.3 | A | 2.3 | A | 4.2 | A | --- |
| 248994_at | 648.4 | P | 557.8 | P | 434.7 | P | 533.2 | P | RPS4 |
| 248995_at | 183 | P | 177.3 | P | 185.2 | P | 190.6 | P | RRS1 |
| 248996_at | 159.8 | P | 155.1 | P | 99.2 | P | 167.1 | P | RRS1 |
| 248997_at | 259.7 | P | 223.9 | P | 288.5 | P | 248.3 | P | BMY2 |
| 248998_at | 26.9 | A | 46.5 | A | 63.4 | A | 43.4 | A | --- |
| 248999_at | 3.7 | A | 30 | A | 3.7 | A | 2.6 | A | --- |
| 249000_at | 47 | P | 57 | P | 60.1 | P | 49.1 | P | --- |
| 249001_at | 7.3 | A | 3.6 | A | 2.9 | A | 25.4 | A | --- |
| 249002_at | 207.7 | P | 597.9 | P | 340.3 | P | 461.7 | P | --- |
| 249003_at | 100.4 | P | 147 | P | 153.8 | P | 132.4 | P | --- |
| 249004_at | 159.1 | P | 113.9 | P | 126 | P | 113.8 | P | --- |
| 249005_at | 2.4 | A | 12.6 | A | 2.3 | A | 12.2 | A | --- |
| 249006_at | 208.4 | P | 189.6 | P | 263.2 | P | 223.4 | P | --- |
| 249007_at | 1123.2 | P | 2022.1 | P | 1427.9 | P | 2059.7 | P | --- |
| 249008_at | 425.9 | P | 449.1 | P | 754.8 | P | 467.3 | P | --- |
| 249009_at | 45.2 | A | 28.8 | A | 60.5 | P | 10.8 | A | MAP18 |
| 249010_at | 1302 | P | 5116.8 | P | 1385.8 | P | 3894.7 | P | --- |
| 249011_at | 261 | P | 242.7 | P | 293.3 | P | 273.9 | P | --- |
| 249012_at | 3.2 | A | 4.3 | A | 49.2 | A | 4.3 | A | CYP706A3 |
| 249013_at | 2.3 | A | 3.1 | A | 15.5 | A | 3.8 | A | GSO2 |
| 249014_at | 587.8 | P | 678 | P | 648.1 | P | 632.5 | P | --- |
| 249015_at | 464.6 | P | 500.8 | P | 514.3 | P | 549 | P | --- |
| 249016_at | 252.4 | P | 292.9 | P | 325.5 | P | 315.5 | P | REV1 |
| 249017_at | 26.2 | A | 11.4 | A | 17.7 | A | 20.6 | A | --- |
| 249018_at | 2.8 | A | 13.1 | A | 18.7 | A | 23.1 | A | --- |
| 249019_at | 147.5 | P | 145.2 | P | 202.3 | P | 152.5 | P | --- |
| 249020_at | 185.4 | P | 189.8 | P | 176.6 | P | 203.3 | P | CHR4 |
| 249021_at | 109.5 | P | 118.8 | P | 100.7 | P | 122.6 | P | --- |
| 249022_s_at | 0.5 | A | 0.7 | A | 1.6 | A | 0.7 | A | --- |
| 249023_at | 54.8 | A | 53.1 | A | 74 | P | 47.9 | M | --- |
| 249024_at | 556 | P | 564.6 | P | 582.6 | P | 555.1 | P | --- |
| 249025_at | 2422.5 | P | 1560.7 | P | 2106.4 | P | 1636.1 | P | --- |
| 249026_at | 741 | P | 650.9 | P | 792 | P | 697.2 | P | OSB3 |
| 249027_at | 542.4 | P | 640.9 | P | 473.2 | P | 578.5 | P | RAN1 |
| 249028_at | 51.3 | M | 98 | P | 62.8 | A | 83.6 | A | POLH |
| 249029_at | 376.5 | P | 410.7 | P | 425.2 | P | 362.2 | P | --- |
| 249030_at | 37.7 | P | 14.8 | A | 41 | A | 38.4 | M | --- |
| 249031_at | 4.8 | A | 14.1 | A | 15.3 | A | 12.5 | A | --- |
| 249032_at | 28.6 | P | 51 | P | 29.2 | A | 49.4 | M | --- |
| 249033_at | 36.7 | P | 57.5 | P | 71.1 | P | 45.1 | P | --- |
| 249034_at | 2.9 | A | 2.5 | A | 4.6 | A | 3.2 | A | NUC |
| 249035_at | 451 | P | 821.8 | P | 736.3 | P | 681.1 | P | GLK2 |
| 249036_at | 584.5 | P | 546.2 | P | 682.4 | P | 543.5 | P | CBP20 |
| 249037_at | 1087.5 | P | 1134.6 | P | 1172.2 | P | 1309.5 | P | FLA13 |
| 249038_at | 63.7 | P | 75.9 | P | 94.7 | P | 58.7 | P | RING1A |

| | | | | | | | | | |
|---|---|---|---|---|---|---|---|---|---|
| 249039_at | 6.2 | A | 3.2 | A | 12.6 | A | 14.4 | A | --- |
| 249040_at | 1721 | P | 1407.1 | P | 1840.1 | P | 1576.4 | P | --- |
| 249041_at | 3.8 | A | 1.3 | A | 2.3 | A | 1.3 | A | --- |
| 249042_at | 63.9 | P | 58.7 | P | 96.4 | M | 58.2 | A | --- |
| 249043_at | 2.5 | A | 1.6 | A | 13.6 | A | 4.4 | A | --- |
| 249044_at | 241.3 | P | 339.9 | P | 294.1 | P | 337.2 | P | PHT4;6 |
| 249045_at | 96.8 | P | 90.7 | P | 54.1 | A | 82.6 | P | --- |
| 249046_at | 262.4 | P | 686.5 | P | 460.8 | P | 612.4 | P | --- |
| 249047_at | 164.9 | P | 209.5 | P | 206.4 | P | 191.6 | P | --- |
| 249048_at | 5 | A | 4.5 | A | 31.9 | A | 36.6 | A | --- |
| 249049_at | 4348 | P | 4546.7 | P | 5599.3 | P | 5016.5 | P | TUB4 |
| 249050_at | 256.7 | P | 342.7 | P | 278.3 | P | 255.6 | P | --- |
| 249051_at | 118.8 | P | 92.9 | P | 75.9 | P | 91.1 | P | --- |
| 249052_at | 777.7 | P | 1013.8 | P | 440.9 | P | 868.2 | P | PDF1.2 |
| 249053_at | 1.2 | A | 3.6 | A | 1.9 | A | 5 | A | --- |
| 249054_at | 337.8 | P | 282.3 | P | 469.8 | P | 317.9 | P | --- |
| 249055_at | 48.5 | A | 41.7 | A | 63.6 | A | 33.9 | A | --- |
| 249056_at | 0.9 | A | 1.1 | A | 2 | A | 2.9 | A | --- |
| 249057_at | 203.1 | P | 155.9 | P | 108.7 | P | 119.2 | P | DUR |
| 249058_at | 180.3 | P | 81.7 | A | 116 | A | 155.9 | P | TAO1 |
| 249059_at | 80.7 | P | 148 | P | 147.5 | P | 122.5 | P | --- |
| 249060_at | 228.6 | P | 416.6 | P | 249.7 | P | 325.4 | P | VPS2.2 |
| 249061_at | 31.5 | A | 146.6 | P | 82.6 | P | 145.3 | P | --- |
| 249062_at | 26.8 | A | 30.2 | A | 23.7 | A | 27 | A | --- |
| 249063_at | 409.7 | P | 637.8 | P | 536.9 | P | 648.7 | P | POP1 |
| 249064_at | 456.3 | P | 537.1 | P | 387.1 | P | 524.9 | P | --- |
| 249065_at | 70.8 | P | 153.9 | P | 11.4 | A | 47.7 | P | --- |
| 249066_at | 131.8 | P | 173.9 | P | 211.6 | P | 160.7 | P | --- |
| 249067_at | 1351.9 | P | 1479.7 | P | 1250 | P | 1451 | P | --- |
| 249068_at | 435 | P | 159.8 | P | 232.5 | P | 165.8 | P | PDLP1 |
| 249069_at | 107.9 | P | 71.4 | P | 121.5 | P | 61.4 | P | --- |
| 249070_at | 134.2 | P | 192 | P | 109.9 | P | 187.3 | P | CESA4 |
| 249071_at | 249.5 | P | 151.1 | P | 292.9 | P | 163.7 | P | --- |
| 249072_at | 250.2 | P | 237 | P | 184.3 | P | 190.9 | P | --- |
| 249073_at | 3587.3 | P | 2971.1 | P | 3978.2 | P | 3266.6 | P | --- |
| 249074_at | 275.1 | P | 198.3 | P | 253.5 | P | 196 | P | --- |
| 249075_at | 1011.6 | P | 796.8 | P | 835 | P | 849.9 | P | --- |
| 249076_at | 1671.5 | P | 1143 | P | 1341.8 | P | 1367.6 | P | TOM22-V |
| 249077_at | 8214.7 | P | 7361.4 | P | 7841.9 | P | 7359.9 | P | HOT5 |
| 249078_at | 778.2 | P | 1138.6 | P | 779.7 | P | 1045.9 | P | CAD1 |
| 249079_at | 244 | P | 260.7 | P | 277.7 | P | 229.8 | P | --- |
| 249080_at | 62.3 | P | 93.8 | P | 71.5 | P | 139 | P | SUVR2 |
| 249081_at | 910.2 | P | 794.5 | P | 534.8 | P | 646.2 | P | --- |
| 249082_at | 33.1 | A | 12.5 | A | 5.9 | A | 17.4 | A | CRA1 |
| 249083_at | 15.4 | A | 2.3 | A | 28.6 | A | 1.1 | A | ATPHB7 |
| 249084_at | 107.8 | P | 92.3 | P | 124.8 | P | 110.8 | P | --- |
| 249085_at | 391.8 | P | 453.6 | P | 377.5 | P | 404.1 | P | --- |
| 249086_at | 290.2 | P | 307.4 | P | 231.5 | P | 320.9 | P | --- |
| 249087_at | 34.9 | A | 50 | P | 26.6 | A | 70.4 | P | ERF9 |
| 249088_at | 53.2 | A | 58.2 | P | 92 | M | 87.8 | P | --- |
| 249089_at | 494.1 | P | 482.2 | P | 495.4 | P | 413.6 | P | --- |
| 249090_at | 358.5 | P | 221.4 | P | 295.3 | P | 228.7 | P | --- |
| 249091_at | 228.9 | P | 190.3 | P | 164.5 | P | 211 | P | CLH2 |
| 249092_at | 188 | P | 202.3 | P | 196 | P | 204 | P | --- |
| 249093_at | 319 | P | 312.7 | P | 290 | P | 347.2 | P | --- |
| 249094_at | 21.4 | A | 12 | A | 66.3 | A | 39.3 | A | YUC5 |
| 249095_at | 394.5 | P | 517.3 | P | 423.7 | P | 446 | P | MYA2 |
| 249096_at | 78.3 | P | 234.9 | P | 76.2 | P | 111 | P | --- |
| 249097_at | 14.1 | A | 5 | A | 15.9 | A | 4.4 | A | --- |
| 249098_at | 17.3 | A | 18.6 | A | 2 | A | 15.1 | A | --- |
| 249099_at | 3.2 | A | 6.6 | A | 5.3 | A | 7.6 | A | --- |
| 249100_at | 692.3 | P | 667.7 | P | 629.8 | P | 654 | P | --- |
| 249101_at | 35.6 | P | 64.3 | P | 53.7 | P | 70.2 | P | --- |
| 249102_at | 9.3 | A | 14.9 | A | 3.8 | A | 3.2 | A | --- |
| 249103_at | 375.1 | P | 397.2 | P | 353 | P | 353 | P | UAH |
| 249104_at | 16.1 | A | 28.9 | P | 27.2 | A | 26.2 | A | ATSUC6 |
| 249105_at | 28.9 | A | 0.9 | A | 27.8 | A | 24.9 | P | --- |
| 249106_at | 255.1 | P | 306.9 | P | 343.1 | P | 300.3 | P | --- |
| 249107_at | 199.4 | P | 209.8 | P | 191.3 | P | 187.1 | P | --- |
| 249108_at | 22 | A | 12.8 | A | 5.8 | A | 19.7 | A | --- |
| 249109_at | 299.1 | P | 397.2 | P | 360.5 | P | 333.2 | P | ATAUX2-11 |
| 249110_at | 44.1 | P | 45.6 | P | 63.7 | P | 27 | P | --- |
| 249111_at | 5.3 | A | 31.7 | A | 62.7 | A | 28.2 | A | --- |
| 249112_at | 1762.7 | P | 1804.6 | P | 1688.3 | P | 1627.3 | P | APS4 |
| 249113_at | 96.2 | P | 134.8 | P | 124.3 | P | 140 | P | --- |
| 249114_at | 6.2 | A | 36.1 | A | 84.9 | M | 7.3 | A | --- |
| 249115_at | 532.4 | P | 486.7 | P | 467.3 | P | 537 | P | ZLL |
| 249116_at | 119.7 | P | 131.3 | P | 153.7 | P | 159 | P | --- |
| 249117_at | 16 | A | 1.5 | A | 9.9 | A | 0.7 | A | AT-HSFA6A |
| 249118_at | 85.2 | P | 82.5 | P | 110.8 | P | 79.9 | P | --- |
| 249119_at | 358.4 | P | 369.2 | P | 398.5 | P | 397.7 | P | --- |
| 249120_at | 1052.1 | P | 1639 | P | 1244 | P | 1532.5 | P | --- |
| 249121_at | 2385.1 | P | 3280 | P | 2247.7 | P | 3280.1 | P | --- |
| 249122_at | 1620.9 | P | 2121.6 | P | 1804.8 | P | 2078 | P | ARD4 |
| 249123_at | 1024.4 | P | 1086.8 | P | 923.2 | P | 1061.6 | P | KCS20 |
| 249124_at | 153 | P | 174.4 | P | 182.8 | P | 202 | P | --- |
| 249125_at | 1770.6 | P | 1209.7 | P | 962 | P | 1169.9 | P | --- |
| 249126_at | 173 | P | 149.2 | P | 153.8 | P | 178.6 | P | TOPP6 |
| 249127_at | 395.8 | P | 251.6 | P | 359.9 | P | 259.6 | P | ATARP9 |
| 249128_at | 435.6 | P | 535.2 | P | 283 | P | 479.4 | P | --- |
| 249129_at | 117.8 | P | 142.3 | P | 96.5 | P | 142.7 | P | CYCA3;1 |
| 249130_at | 403 | P | 319.8 | P | 299.6 | P | 291.6 | P | --- |
| 249131_at | 2.9 | A | 1.7 | A | 8.2 | A | 3.4 | A | APUM14 |
| 249132_at | 20.3 | A | 3.8 | A | 4.4 | A | 28.9 | A | --- |
| 249133_at | 820.2 | P | 946.3 | P | 1028.6 | P | 807.2 | P | TAF4 |
| 249134_at | 1017.5 | P | 1070.8 | P | 662.1 | P | 971.4 | P | --- |
| 249135_at | 84.9 | P | 64.1 | P | 42 | A | 107.6 | P | --- |
| 249136_at | 81.9 | P | 71.2 | P | 113.3 | P | 69.2 | P | --- |
| 249137_at | 669.8 | P | 426.7 | P | 535.1 | P | 476.4 | P | --- |
| 249138_at | 215.5 | P | 276 | P | 217.4 | P | 267.2 | P | WPP1 |
| 249139_at | 256.4 | P | 198.3 | P | 230.1 | P | 194 | P | AZF3 |
| 249140_at | 431.5 | P | 385.4 | P | 344.9 | P | 307.6 | P | --- |
| 249141_at | 17.9 | A | 12.8 | A | 13.9 | A | 20 | A | --- |
| 249142_at | 1.8 | A | 3.9 | A | 3.6 | A | 1.9 | A | --- |
| 249143_at | 27.8 | P | 13.4 | A | 43 | P | 20.7 | A | --- |
| 249144_at | 456.7 | P | 437 | P | 385 | P | 374.3 | P | SPL2 |
| 249145_at | 607.1 | P | 720.8 | P | 596.7 | P | 664.4 | P | ATDCI1 |
| 249146_at | 283.6 | P | 238.6 | P | 243.3 | A | 243.8 | P | --- |
| 249147_at | 1279.2 | P | 1066.1 | P | 1191.5 | P | 1024.9 | P | --- |

| | | | | | | | | | |
|---|---|---|---|---|---|---|---|---|---|
| 249148_at | 2427.4 | P | 1950.5 | P | 2096.1 | P | 2013.3 | P | --- |
| 249149_at | 963.3 | P | 967.3 | P | 933.3 | P | 984.5 | P | ckl8 |
| 249150_at | 2.1 | A | 2.1 | A | 2.4 | A | 2 | A | PHT6 |
| 249151_at | 9.9 | A | 32 | A | 21.8 | A | 19.6 | A | PHT3 |
| 249152_s_at | 49.8 | P | 101.7 | P | 110.9 | P | 91.4 | P | APT1 |
| 249153_s_at | 106 | P | 134.5 | P | 84.1 | A | 81.5 | P | --- |
| 249154_at | 2.3 | A | 4.5 | A | 2.5 | A | 17.3 | A | --- |
| 249155_at | 4.2 | A | 2.7 | A | 3 | A | 5 | A | --- |
| 249156_s_at | 144.4 | P | 138.5 | P | 183 | P | 120.8 | P | --- |
| 249157_at | 2.7 | A | 1.5 | A | 0.7 | A | 14.1 | A | --- |
| 249158_at | 784.5 | P | 703.5 | P | 653.9 | P | 636.2 | P | ETFBETA |
| 249159_at | 3929.1 | P | 3280.3 | P | 3363.3 | P | 3391 | P | --- |
| 249160_at | 113.6 | P | 108.2 | P | 103.7 | P | 155.6 | P | --- |
| 249161_at | 2467.2 | P | 2160.4 | P | 2366.4 | P | 2289.9 | P | PAF1 |
| 249162_at | 622.1 | P | 817.4 | P | 592.4 | P | 845.6 | P | --- |
| 249163_at | 1087.9 | P | 1167.2 | P | 1347.1 | P | 1114.4 | P | --- |
| 249164_at | 898.4 | P | 634.9 | P | 836.5 | P | 725.6 | P | U2AF35B |
| 249165_at | 551.3 | P | 351.7 | P | 446.1 | P | 419.5 | P | ATIPK1 |
| 249166_at | 37 | A | 31.7 | A | 52.5 | A | 35.4 | A | --- |
| 249167_at | 75.5 | P | 97.3 | P | 92.5 | P | 125.5 | P | --- |
| 249168_at | 331.7 | P | 340.8 | P | 284.2 | P | 260.1 | P | --- |
| 249169_at | 29.9 | M | 32.3 | P | 61.3 | P | 61.7 | P | --- |
| 249170_at | 29.8 | A | 3.2 | A | 26.2 | A | 16.2 | A | --- |
| 249171_at | 636.2 | P | 617.6 | P | 645.9 | P | 609.8 | P | --- |
| 249172_at | 615 | P | 474.2 | P | 517.6 | P | 429.9 | P | --- |
| 249173_at | 2 | A | 1.7 | A | 4.9 | A | 1.6 | A | --- |
| 249174_at | 1175.2 | P | 784.7 | P | 1166.3 | P | 1032.2 | P | --- |
| 249175_at | 704 | P | 767.9 | P | 794.9 | P | 745.1 | P | COP8 |
| 249176_at | 12416.8 | P | 11451.1 | P | 12317.5 | P | 11177.1 | P | ATTRX3 |
| 249177_at | 1316.7 | P | 1114.3 | P | 1116.7 | P | 985.2 | P | --- |
| 249178_at | 2471.9 | P | 2461.4 | P | 2430.1 | P | 2384.8 | P | SCP2 |
| 249179_at | 1.9 | A | 15.7 | A | 4.5 | A | 2.7 | A | --- |
| 249180_at | 1326.6 | P | 1197.3 | P | 1214.2 | P | 1155.2 | P | RPT4A |
| 249181_at | 512.2 | P | 648.6 | P | 589.8 | P | 628.1 | P | --- |
| 249182_at | 832.6 | P | 994.1 | P | 1221.9 | P | 989.4 | P | --- |
| 249183_at | 569.8 | P | 685.7 | P | 579.6 | P | 653.9 | P | UBC18 |
| 249184_at | 145.5 | P | 249.7 | P | 267.3 | P | 258 | P | --- |
| 249185_at | 1.7 | A | 1.3 | A | 3.8 | A | 2.4 | A | --- |
| 249186_at | 57.5 | P | 47.4 | P | 79.7 | P | 40.1 | P | --- |
| 249187_at | 1279.2 | P | 1455.2 | P | 858 | P | 1316.4 | P | --- |
| 249188_at | 103.2 | P | 78.6 | P | 46.4 | A | 41.6 | A | --- |
| 249189_at | 141.6 | P | 112.3 | P | 173 | P | 129.2 | P | AtHB27 |
| 249190_at | 398.4 | P | 265.8 | P | 319.5 | P | 241.8 | P | BKI1 |
| 249191_at | 158.3 | P | 369.3 | P | 384.3 | P | 407.4 | P | --- |
| 249192_at | 386.8 | P | 461.4 | P | 439.3 | P | 468.5 | P | --- |
| 249193_at | 767.7 | P | 814.6 | P | 972.6 | P | 900.3 | P | ARC6 |
| 249194_at | 4.3 | A | 1.2 | A | 11.6 | A | 12.9 | A | --- |
| 249195_s_at | 39.5 | A | 51.3 | A | 12.2 | A | 20 | A | --- |
| 249196_at | 338.1 | P | 410.5 | P | 393.2 | P | 382 | P | --- |
| 249197_at | 10.4 | A | 42.8 | P | 30.9 | P | 8.8 | A | CML37 |
| 249198_s_at | 252.7 | P | 288.2 | P | 213.3 | P | 273.4 | P | --- |
| 249199_at | 683.7 | P | 869.8 | P | 850.8 | P | 685.6 | P | BPC6 |
| 249200_at | 658.9 | P | 546.5 | P | 402.7 | P | 517.9 | P | XRN2 |
| 249201_at | 214.5 | P | 193.7 | P | 280.8 | P | 290.5 | P | --- |
| 249202_at | 12.2 | A | 4 | A | 9.7 | A | 10.5 | A | CYP705A12 |
| 249203_at | 19.7 | A | 22.9 | A | 23.5 | A | 30.5 | A | CYP71A16 |
| 249204_at | 632.6 | P | 723.2 | P | 545 | P | 779 | P | --- |
| 249205_at | 3.5 | A | 4.8 | A | 2.5 | A | 3 | A | MRN1 |
| 249206_at | 13.9 | A | 15.1 | A | 31.7 | P | 32.3 | A | ATS |
| 249207_at | 16 | A | 8.3 | A | 5 | A | 20.1 | A | --- |
| 249208_at | 4217.9 | P | 1320.5 | P | 4015 | P | 1863.8 | P | AOS |
| 249209_at | 173.7 | P | 215 | P | 193.7 | P | 210.1 | P | --- |
| 249210_at | 272.8 | P | 334.1 | P | 305 | P | 372.1 | P | --- |
| 249211_at | 197.5 | P | 321.2 | P | 306 | P | 391 | P | --- |
| 249212_at | 33.1 | A | 48 | P | 59.2 | P | 41.3 | P | --- |
| 249213_at | 41.3 | M | 37 | P | 29.5 | A | 26.3 | P | --- |
| 249214_at | 254.5 | P | 446.4 | P | 332.5 | P | 399.8 | P | --- |
| 249215_at | 1836.2 | P | 400.5 | P | 3547.2 | P | 1148.9 | P | DFR |
| 249216_at | 508.7 | P | 673 | P | 760.6 | P | 667 | P | scpl42 |
| 249217_at | 5649.1 | P | 4896.2 | P | 4905.9 | P | 4984.5 | P | UBL5 |
| 249218_at | 1640.5 | P | 1566.7 | P | 1647.4 | P | 1469.9 | P | --- |
| 249219_at | 148.2 | P | 195.4 | P | 194.2 | P | 181.8 | P | SDG25 |
| 249220_at | 612.4 | P | 929.3 | P | 781 | P | 820 | P | --- |
| 249221_at | 49.6 | A | 54.2 | P | 60.2 | P | 61.7 | P | --- |
| 249222_at | 37 | P | 45.2 | P | 31.6 | P | 49.3 | P | --- |
| 249223_at | 2.1 | A | 1.9 | A | 3.6 | A | 1.8 | A | --- |
| 249224_at | 857.4 | P | 721.2 | P | 818.5 | P | 760.7 | P | --- |
| 249225_at | 112.3 | P | 113.4 | P | 101 | A | 132.9 | P | --- |
| 249226_at | 11.3 | A | 9.8 | A | 8.3 | A | 18.6 | M | --- |
| 249227_at | 15.1 | A | 9.8 | A | 50.1 | A | 2.8 | A | --- |
| 249228_at | 1916.8 | P | 2118.6 | P | 2091.7 | P | 2185.2 | P | ASK2 |
| 249229_at | 2100.9 | P | 2213.3 | P | 2104.1 | P | 2174.9 | P | --- |
| 249230_at | 460.7 | P | 832.6 | P | 597 | P | 789.3 | P | --- |
| 249231_at | 1542.5 | P | 1522.8 | P | 1493.2 | P | 1415.7 | P | ABIL4 |
| 249232_at | 1635.6 | P | 1959.5 | P | 1582.5 | P | 1918.9 | P | ADL1 |
| 249233_at | 915.4 | P | 683.1 | P | 1017.8 | P | 775.7 | P | --- |
| 249234_at | 137.8 | P | 110.4 | P | 140.8 | P | 91.5 | P | --- |
| 249235_at | 1403.3 | P | 1737.8 | P | 1946.5 | P | 1809 | P | BG_PPAP |
| 249236_at | 869.5 | P | 1106.9 | P | 778.1 | P | 1159 | P | --- |
| 249237_at | 2374.3 | P | 2982.2 | P | 2251 | P | 3012.9 | P | --- |
| 249238_at | 922.5 | P | 935.8 | P | 1266.2 | P | 950.9 | P | --- |
| 249239_at | 891.7 | P | 980.9 | P | 1001.6 | P | 923.9 | P | CPuORF58 |
| 249240_at | 596.2 | P | 666.5 | P | 571.7 | P | 627 | P | --- |
| 249241_at | 2.8 | A | 1.2 | A | 2.6 | A | 15.7 | A | scpl41 |
| 249242_at | 345 | P | 462.5 | P | 239 | P | 415.9 | P | --- |
| 249243_s_at | 1.6 | A | 7.7 | A | 1.5 | A | 0.5 | A | BGLU13 |
| 249244_at | 6799.2 | P | 5068.6 | P | 5566.9 | P | 5391.4 | P | VAR1 |
| 249245_at | 70.3 | P | 57.2 | M | 99.3 | P | 54.8 | P | --- |
| 249246_at | 15.4 | A | 11.3 | A | 6.6 | A | 5.9 | A | --- |
| 249247_at | 762.6 | P | 904.5 | P | 1061.8 | P | 874.3 | P | --- |
| 249248_at | 453.8 | P | 548.2 | P | 425.5 | P | 521.5 | P | --- |
| 249249_at | 59 | A | 68.5 | A | 68.5 | A | 82.5 | A | --- |
| 249250_at | 11.8 | A | 3.9 | A | 2.7 | A | 3.4 | A | --- |
| 249251_at | 428.2 | P | 488 | P | 437.4 | P | 459.4 | P | --- |
| 249252_at | 67.6 | P | 82.7 | P | 32.1 | A | 55.1 | P | --- |
| 249253_at | 231.2 | P | 134.4 | P | 256.5 | P | 166.7 | P | --- |
| 249254_at | 17.3 | P | 51.3 | P | 43.6 | P | 13.7 | M | --- |
| 249255_at | 166.1 | P | 60.1 | P | 109.4 | P | 58.8 | P | CHX18 |
| 249256_at | 28.6 | A | 44.3 | A | 30.8 | A | 69.5 | P | --- |

| | | | | | | | | | |
|---|---|---|---|---|---|---|---|---|---|
| 249257_at | 22.4 | A | 24.2 | A | 30.9 | A | 9.3 | A | --- |
| 249258_at | 207.1 | P | 225.9 | P | 276.6 | P | 269.5 | P | --- |
| 249259_at | 10.4 | A | 28.3 | A | 33.8 | A | 26.5 | A | --- |
| 249260_at | 6.1 | A | 15.8 | A | 47.3 | A | 37.5 | A | --- |
| 249261_at | 22.1 | A | 49.5 | A | 53.2 | A | 64.8 | A | --- |
| 249262_at | 42.8 | A | 35.4 | A | 29.4 | A | 39.7 | A | --- |
| 249263_at | 6.3 | A | 6 | A | 4.5 | A | 9.2 | A | --- |
| 249264_s_at | 239.9 | P | 335.5 | P | 264.2 | P | 178.5 | P | --- |
| 249265_at | 8502.1 | P | 9331 | P | 10225.1 | P | 9736.4 | P | --- |
| 249266_at | 1038.8 | P | 983.2 | P | 1314.4 | P | 1020.3 | P | --- |
| 249267_at | 1142.8 | P | 1084 | P | 1323 | P | 1247.6 | P | BTI3 |
| 249268_at | 1301.9 | P | 1120.3 | P | 807.7 | P | 1125.1 | P | --- |
| 249269_at | 274.4 | P | 312.6 | P | 318.5 | P | 323 | P | --- |
| 249270_at | 16.5 | A | 3.4 | A | 9.5 | A | 16.3 | A | --- |
| 249271_at | 934.6 | P | 895.5 | P | 751.7 | P | 863.5 | P | CIP1 |
| 249272_at | 5.2 | A | 8.3 | A | 11.7 | A | 23.8 | A | --- |
| 249273_at | 135.4 | P | 152.4 | P | 155.3 | P | 145.6 | P | --- |
| 249274_at | 50.9 | A | 51 | A | 53.2 | A | 41.8 | A | --- |
| 249275_at | 191.6 | P | 238.1 | P | 211.6 | P | 228.5 | P | --- |
| 249276_at | 187 | P | 276.3 | P | 265.8 | P | 329.2 | P | POLA3 |
| 249277_at | 12.2 | A | 15.6 | A | 19.8 | A | 22.2 | A | --- |
| 249278_at | 47.3 | M | 101 | P | 75.7 | A | 79.2 | P | --- |
| 249279_at | 174.8 | P | 225.1 | P | 189 | P | 226.8 | P | --- |
| 249280_at | 164.7 | P | 236.2 | P | 128.1 | P | 248.5 | P | --- |
| 249281_at | 629.8 | P | 676.8 | P | 667 | P | 644.2 | P | --- |
| 249282_at | 468 | P | 521.4 | P | 508.5 | P | 473.6 | P | --- |
| 249283_at | 317.2 | P | 247.7 | P | 233.8 | P | 230.2 | P | --- |
| 249284_at | 336.6 | P | 223.9 | P | 299.3 | P | 196 | P | --- |
| 249285_at | 842.9 | P | 693.7 | P | 769.6 | P | 701.6 | P | --- |
| 249286_at | 109.1 | P | 137.5 | P | 112.2 | P | 110 | P | --- |
| 249287_at | 18 | A | 5.5 | A | 3.8 | A | 1.6 | A | --- |
| 249288_at | 436.2 | P | 533.9 | P | 743.9 | P | 576.7 | P | --- |
| 249289_at | 56.7 | P | 131.7 | P | 118.8 | P | 120.5 | P | --- |
| 249290_at | 209.8 | P | 288.4 | P | 364 | P | 335.4 | P | --- |
| 249291_at | 2874.4 | P | 2567.5 | P | 2679.3 | P | 2580.2 | P | ATGSTT1 |
| 249292_at | 139.4 | P | 156.9 | P | 198.3 | P | 146.4 | P | UVH1 |
| 249293_at | 435.1 | P | 472.1 | P | 445 | P | 377.2 | P | --- |
| 249294_at | 185.2 | P | 125.3 | P | 157.9 | P | 123 | P | --- |
| 249295_at | 3.6 | A | 8.4 | A | 41 | A | 35.5 | A | --- |
| 249296_at | 1.8 | A | 2 | A | 1.8 | A | 1.2 | A | --- |
| 249297_at | 58.9 | P | 50.6 | A | 61.7 | P | 54.8 | P | --- |
| 249298_at | 170.6 | P | 153.8 | P | 191.6 | P | 164.3 | P | --- |
| 249299_at | 964.9 | P | 1082.7 | P | 840 | P | 1184.5 | P | UBC4 |
| 249300_at | 1.8 | A | 17.9 | A | 14.4 | A | 21.9 | A | --- |
| 249301_at | 3.4 | A | 11.8 | A | 3.4 | A | 0.7 | A | --- |
| 249302_at | 17.4 | A | 11 | A | 17.8 | A | 22.8 | A | --- |
| 249303_at | 323.9 | P | 315.2 | P | 271.4 | P | 389 | P | --- |
| 249304_at | 232.1 | P | 200.1 | P | 172.5 | P | 158.5 | P | GLA1 |
| 249305_at | 2.5 | A | 1.4 | A | 5.3 | A | 1.2 | A | --- |
| 249306_at | 499.9 | P | 495.8 | P | 482.1 | P | 588.7 | P | --- |
| 249307_s_at | 303.8 | P | 297.5 | P | 421.9 | P | 290.1 | P | XPB1 |
| 249308_at | 448.8 | P | 481.5 | P | 420.6 | P | 418 | P | --- |
| 249309_at | 492.2 | P | 426.8 | P | 380.3 | P | 415.8 | P | BEL1 |
| 249310_at | 6316.4 | P | 4937.3 | P | 7010.2 | P | 5125.4 | P | --- |
| 249311_at | 22.4 | A | 5.3 | A | 7.1 | A | 3.3 | A | --- |
| 249312_at | 30.7 | A | 52.9 | M | 53 | A | 53.8 | P | --- |
| 249313_at | 438.3 | P | 421.6 | P | 455.4 | P | 428.7 | P | --- |
| 249314_at | 101.2 | P | 147 | P | 159.9 | P | 124.2 | P | --- |
| 249315_at | 1133.7 | P | 775.2 | P | 1079 | P | 963.5 | P | --- |
| 249316_s_at | 294 | P | 192.8 | P | 210.6 | P | 255.7 | P | ATGSTT2 |
| 249317_at | 63.8 | P | 81 | P | 62.4 | P | 33.7 | P | ATGSTT2 |
| 249318_at | 389.2 | P | 501.3 | P | 445.2 | P | 486.9 | P | AtUK/UPRT1 |
| 249319_s_at | 55 | P | 65.7 | P | 90.6 | P | 58.5 | P | --- |
| 249320_at | 227.1 | P | 253.5 | P | 229.4 | P | 248.7 | P | --- |
| 249321_at | 102.9 | P | 128.6 | P | 110.7 | P | 123.2 | P | --- |
| 249322_at | 1021.7 | P | 798.8 | P | 909.7 | P | 806.9 | P | TOM20-4 |
| 249323_at | 2.3 | A | 6 | A | 2.5 | A | 1.5 | A | FLA20 |
| 249324_at | 2836.3 | P | 2927.2 | P | 1890.2 | P | 2803.9 | P | --- |
| 249325_at | 677 | P | 329.7 | P | 746.4 | P | 374.8 | P | UPM1 |
| 249326_at | 55.6 | P | 61.4 | P | 104.7 | P | 69.1 | P | SYN2 |
| 249327_at | 562.1 | P | 463.9 | P | 711.6 | P | 472.1 | P | ATCLC-A |
| 249328_at | 50.4 | A | 66.9 | A | 81.4 | A | 82 | M | ATR |
| 249329_at | 214.4 | P | 215.1 | P | 249.3 | P | 180.4 | P | --- |
| 249330_at | 140.5 | P | 155.9 | P | 173.9 | P | 156.5 | P | --- |
| 249331_at | 7680.7 | P | 9244.7 | P | 7924.4 | P | 8537.3 | P | RPL27 |
| 249332_at | 104.7 | P | 179.9 | P | 168.1 | P | 159.5 | P | --- |
| 249333_at | 18.8 | A | 0.3 | A | 2.8 | A | 21.5 | A | GLIP1 |
| 249334_at | 221.3 | P | 183.6 | P | 232.9 | M | 202.9 | P | YSL4 |
| 249335_at | 1187.5 | P | 1036.7 | P | 1174.5 | P | 1124.3 | P | NRPB12 |
| 249336_at | 406.8 | P | 228.7 | P | 439 | P | 307 | P | DRB5 |
| 249337_at | 62.6 | P | 54 | P | 68.1 | P | 11.3 | A | --- |
| 249338_at | 25.7 | P | 16.5 | A | 33.1 | A | 11.2 | A | anac095 |
| 249339_at | 162.9 | P | 180.7 | P | 173.9 | P | 152 | P | --- |
| 249340_at | 87.8 | P | 237.3 | P | 133.4 | P | 180.7 | P | --- |
| 249341_at | 34.3 | P | 33 | P | 43.8 | P | 31.9 | P | --- |
| 249342_at | 671.6 | P | 681.3 | P | 734.4 | P | 700.1 | P | --- |
| 249343_at | 1230.5 | P | 1197.1 | P | 1439.9 | P | 1240.3 | P | SDH2-2 |
| 249344_at | 2623 | P | 2310.4 | P | 3150.4 | P | 2382 | P | ATPHB3 |
| 249345_at | 116.4 | P | 170.8 | P | 130.8 | P | 150 | P | --- |
| 249346_at | 210.9 | P | 1265.5 | P | 118.2 | P | 671.9 | P | LTH1 |
| 249347_at | 630.8 | P | 716.3 | P | 610.3 | P | 832 | P | --- |
| 249348_at | 7.1 | A | 8.4 | A | 39.5 | A | 3.4 | A | --- |
| 249349_at | 11.5 | A | 3.4 | A | 13.2 | A | 0.8 | A | MYB24 |
| 249350_at | 11.9 | A | 3 | A | 34.5 | A | 5.9 | A | MYB115 |
| 249351_at | 133.2 | P | 121 | P | 119.1 | P | 128.3 | P | ATMKK3 |
| 249352_at | 4.7 | A | 2.4 | A | 3.1 | A | 12.7 | A | MYB22 |
| 249353_at | 22.1 | A | 1.7 | A | 26.3 | A | 17.2 | A | OLEO2 |
| 249354_at | 635.8 | P | 522.9 | P | 700.6 | P | 644.5 | P | EMB3012 |
| 249355_at | 592.6 | P | 859.8 | P | 508 | P | 812.5 | P | --- |
| 249356_at | 173.6 | P | 256.9 | P | 248 | P | 201.6 | P | --- |
| 249357_at | 366.9 | P | 344.6 | P | 405.1 | P | 298.8 | P | --- |
| 249358_at | 24.5 | A | 46.9 | P | 63.1 | P | 52.7 | P | --- |
| 249359_at | 279.2 | P | 257.5 | P | 235.1 | P | 215.8 | P | --- |
| 249360_at | 372.4 | P | 364.3 | P | 400 | P | 385.9 | P | --- |
| 249361_at | 301.1 | P | 401.3 | P | 253.2 | P | 356.3 | P | --- |
| 249362_at | 122.6 | P | 161.8 | P | 120.3 | P | 136.6 | P | --- |
| 249363_at | 1.1 | A | 3.4 | A | 1.3 | A | 2.3 | A | DegP13 |
| 249364_at | 19.8 | A | 27 | A | 40.1 | A | 2.8 | A | --- |
| 249365_at | 151.4 | P | 137.6 | P | 113.9 | P | 163.2 | P | --- |

| | | | | | | | | | |
|---|---|---|---|---|---|---|---|---|---|
| 249366_at | 427.6 | P | 445.9 | P | 443.5 | P | 439 | P | --- |
| 249367_at | 2.7 | A | 24.1 | A | 5.5 | A | 0.9 | A | --- |
| 249368_at | 36.2 | P | 82.7 | P | 78.1 | P | 92.2 | P | --- |
| 249369_at | 33 | A | 35.6 | A | 29.9 | A | 42.6 | A | --- |
| 249370_at | 322.4 | P | 322 | P | 350.1 | P | 329.3 | P | --- |
| 249371_at | 9.4 | A | 3.4 | A | 14.7 | A | 0.8 | A | --- |
| 249372_at | 639.4 | P | 847.2 | P | 502.1 | P | 782.7 | P | G6PD6 |
| 249373_at | 1094.1 | P | 1176.5 | P | 933.4 | P | 1022.4 | P | --- |
| 249374_at | 1078.5 | P | 1107.3 | P | 968.6 | P | 1228.6 | P | PBB2 |
| 249375_at | 513.1 | P | 415.2 | P | 485.4 | P | 413.7 | P | AGP24 |
| 249376_at | 1 | A | 0.4 | A | 0.4 | A | 1.2 | A | --- |
| 249377_at | 308.5 | P | 236.9 | P | 181.8 | P | 175.7 | P | --- |
| 249378_at | 873.5 | P | 897.1 | P | 484.8 | P | 656.5 | P | --- |
| 249379_at | 63.9 | P | 42.5 | P | 41.6 | P | 46.8 | P | --- |
| 249380_at | 9911.5 | P | 8581.8 | P | 10434.6 | P | 8339.9 | P | --- |
| 249381_at | 34.2 | A | 8.3 | A | 22.6 | A | 18.2 | A | --- |
| 249382_at | 31.4 | A | 17 | A | 23.7 | A | 3.7 | A | --- |
| 249383_at | 30.2 | P | 49.4 | P | 39.5 | P | 16.6 | A | PRE1 |
| 249384_at | 53.1 | P | 86.9 | P | 107.6 | P | 77.3 | P | --- |
| 249385_at | 2093 | P | 2443.3 | P | 1937.8 | P | 2428.4 | P | ATTRX2 |
| 249386_at | 189.4 | P | 230.3 | P | 194.3 | P | 170.5 | P | --- |
| 249387_at | 26.7 | A | 21.6 | A | 60.5 | A | 29 | A | --- |
| 249388_at | 29.1 | A | 34.7 | P | 32.8 | A | 2.5 | A | --- |
| 249389_at | 4 | A | 3.9 | A | 4.4 | A | 2.3 | A | --- |
| 249390_at | 41.8 | P | 24.6 | P | 41.5 | A | 22 | A | --- |
| 249391_at | 171 | P | 214.7 | P | 193.8 | P | 194.8 | P | --- |
| 249392_at | 332 | P | 317.1 | P | 353.1 | P | 367.2 | P | --- |
| 249393_at | 1052.3 | P | 825 | P | 736 | P | 898.3 | P | AtRLP54 |
| 249394_at | 21.2 | A | 0.8 | A | 21.1 | A | 23.1 | A | --- |
| 249395_at | 963.7 | P | 809.4 | P | 882 | P | 738.9 | P | --- |
| 249396_at | 840.9 | P | 699.9 | P | 808.7 | P | 731.5 | P | DegP9 |
| 249397_at | 51 | A | 86.5 | P | 77.7 | A | 64.1 | A | --- |
| 249398_at | 321.7 | P | 362.4 | P | 385.2 | P | 379.3 | P | --- |
| 249399_at | 0.6 | A | 1.7 | A | 8 | A | 3.2 | A | --- |
| 249400_at | 420.5 | P | 672.9 | P | 494 | P | 620.8 | P | --- |
| 249401_at | 23.9 | A | 17.4 | A | 36.9 | A | 9.5 | A | --- |
| 249402_at | 1.5 | A | 1.1 | A | 2.2 | A | 1.8 | A | --- |
| 249403_at | 91.7 | P | 128.4 | P | 134.2 | P | 134.5 | P | --- |
| 249404_at | 1078.6 | P | 1181.5 | P | 1007.1 | P | 1172.3 | P | EMB506 |
| 249405_at | 139.3 | P | 171.6 | P | 159.4 | P | 198.3 | P | ERA1 |
| 249406_at | 138.4 | P | 64.4 | P | 162.6 | P | 74 | P | --- |
| 249407_at | 3.6 | A | 3.8 | A | 12 | A | 3 | A | --- |
| 249408_at | 112.9 | P | 122.5 | P | 89.9 | P | 139.1 | P | MYB23 |
| 249409_at | 351.1 | P | 288.7 | P | 256.4 | P | 248.3 | P | --- |
| 249410_at | 162.9 | A | 153.1 | A | 159.1 | A | 132.3 | P | --- |
| 249411_at | 301.6 | P | 537.8 | P | 449.5 | P | 635 | P | SIP1 |
| 249412_at | 42.4 | A | 39.1 | A | 39.2 | A | 16.2 | A | --- |
| 249413_at | 48.9 | A | 12.9 | A | 25.6 | A | 17.7 | A | AtRABG1 |
| 249414_at | 4.7 | A | 4 | A | 2.3 | A | 3.1 | A | MYB89 |
| 249415_at | 51.5 | P | 97.8 | P | 67 | A | 64.8 | P | CDF2 |
| 249416_at | 5262 | P | 5573 | P | 5077.3 | P | 5595.9 | P | --- |
| 249417_at | 374.2 | P | 536.8 | P | 316.5 | P | 299.1 | P | --- |
| 249418_at | 14.2 | A | 34.8 | P | 58.6 | P | 21.9 | P | --- |
| 249419_s_at | 13.4 | A | 3.5 | A | 6 | A | 23.5 | A | AGL98 |
| 249420_at | 1.4 | A | 1.2 | A | 2.3 | A | 2 | A | anac094 |
| 249421_at | 737.6 | P | 972.2 | P | 1052.3 | P | 1019.8 | P | DEG8 |
| 249422_at | 291.9 | P | 300 | P | 320.9 | P | 353.6 | P | AtHB23 |
| 249423_at | 111.4 | P | 134.2 | P | 93.7 | P | 141.1 | P | --- |
| 249424_s_at | 338.1 | P | 338.4 | P | 273.1 | P | 359.9 | P | --- |
| 249425_at | 643.5 | P | 559 | P | 664.2 | P | 606.8 | P | --- |
| 249426_at | 96.3 | P | 119.1 | P | 164.8 | P | 115.7 | P | --- |
| 249427_at | 1648.6 | P | 843.5 | P | 1639.6 | P | 794.9 | P | --- |
| 249428_at | 13.5 | A | 15.8 | A | 30.7 | A | 2.2 | A | --- |
| 249429_at | 8.2 | A | 0.5 | A | 2 | A | 10 | A | --- |
| 249430_at | 97.7 | P | 107.5 | P | 122.4 | P | 110 | P | --- |
| 249431_at | 25.4 | A | 5.9 | A | 4.7 | A | 34.9 | A | --- |
| 249432_at | 31.3 | A | 26.8 | A | 6.2 | A | 16 | A | CLPS5 |
| 249433_at | 282.5 | P | 293.9 | P | 291.2 | P | 272.7 | P | --- |
| 249434_at | 153.5 | P | 176.5 | P | 187.2 | P | 181.8 | P | --- |
| 249435_at | 6.2 | A | 24.2 | A | 2.6 | A | 7.1 | A | --- |
| 249436_at | 247.3 | P | 243.4 | P | 229.3 | P | 268.7 | P | --- |
| 249437_at | 345.4 | P | 499.1 | P | 328.1 | P | 539.3 | P | --- |
| 249438_at | 2.3 | A | 41.4 | P | 28.1 | A | 39.4 | P | AATP1 |
| 249439_at | 38.6 | P | 126.3 | P | 83.2 | P | 94.7 | P | --- |
| 249440_at | 36.3 | A | 48.5 | A | 26.2 | A | 24.4 | A | --- |
| 249441_at | 1907.4 | P | 1802.1 | P | 1524 | P | 1893.1 | P | --- |
| 249442_at | 1680 | P | 1078.5 | P | 1453.4 | P | 1061.9 | P | --- |
| 249443_at | 484.9 | P | 499.4 | P | 491.5 | P | 500.1 | P | --- |
| 249444_at | 9.2 | A | 12.6 | A | 40 | A | 1.4 | A | --- |
| 249445_at | 36.4 | P | 30.6 | P | 24.3 | A | 23 | A | --- |
| 249446_at | 35 | A | 18.3 | A | 50.4 | A | 4.3 | A | --- |
| 249447_at | 1.5 | A | 0.8 | A | 28.2 | A | 12.4 | A | PTEN1 |
| 249448_at | 1.4 | A | 1.1 | A | 1.7 | A | 2.1 | A | cdc2cAt |
| 249449_at | 3.4 | A | 4.6 | A | 4.5 | A | 1 | A | --- |
| 249450_at | 22.7 | A | 16.3 | A | 22.1 | A | 24.3 | A | SnRK1.3 |
| 249451_s_at | 7.7 | A | 1.9 | A | 0.8 | A | 0.5 | A | --- |
| 249452_at | 316.9 | P | 416 | P | 340.7 | P | 374.9 | P | --- |
| 249453_at | 2095.7 | P | 1951 | P | 1840 | P | 2048.3 | P | SGR4 |
| 249454_at | 547.9 | P | 530.5 | P | 138.6 | P | 303.8 | P | --- |
| 249455_s_at | 9.6 | A | 13.1 | A | 5.1 | A | 0.5 | A | --- |
| 249456_at | 1405.4 | P | 1304.3 | P | 1237.2 | P | 1297.4 | P | --- |
| 249457_s_at | 70.1 | P | 74.6 | P | 62.2 | P | 62.6 | P | VIM4 |
| 249458_at | 27.2 | A | 34.1 | P | 55.2 | A | 41.2 | M | --- |
| 249459_at | 110.7 | P | 38.9 | P | 84.6 | P | 29.9 | P | --- |
| 249460_at | 0.6 | A | 2.4 | A | 18 | A | 5.7 | A | --- |
| 249461_at | 29.8 | A | 11.9 | A | 31.2 | A | 3.5 | A | --- |
| 249462_at | 76 | P | 111.5 | P | 95.7 | A | 90.2 | P | EMB2744 |
| 249463_s_at | 2.1 | A | 1.1 | A | 27.1 | P | 4.5 | A | anac064 |
| 249464_at | 169.1 | P | 246.8 | P | 243 | P | 233.5 | P | EMB2745 |
| 249465_at | 2.2 | A | 1.5 | A | 2.8 | A | 2.6 | A | AIG2L |
| 249466_at | 5641.8 | P | 5011.8 | P | 5431.4 | P | 5695.7 | P | RPL5B |
| 249467_at | 187.7 | P | 193 | P | 54.5 | P | 112.5 | P | ATNAC6 |
| 249468_at | 18.7 | A | 1.8 | A | 6.3 | A | 6.7 | A | --- |
| 249469_at | 333.3 | P | 592.6 | P | 357.8 | P | 511.6 | P | --- |
| 249470_at | 181.7 | P | 172.6 | P | 202 | P | 206.3 | P | --- |
| 249471_at | 336 | P | 481.5 | P | 440 | P | 381.7 | P | EDL2 |
| 249472_at | 632.5 | P | 667.1 | P | 629.8 | P | 674.4 | P | CRR7 |
| 249473_at | 575.3 | P | 711.7 | P | 705.6 | P | 814.3 | P | AHP3 |
| 249474_s_at | 32.1 | A | 49.1 | P | 33.4 | A | 27.7 | P | GER2 |

| | | | | | | | | | |
|---|---|---|---|---|---|---|---|---|---|
| 249475_at | 651.5 | P | 498.5 | P | 845.7 | P | 569.1 | P | --- |
| 249476_at | 6.9 | A | 6.3 | A | 9.5 | A | 36.4 | A | --- |
| 249477_s_at | 194.4 | P | 289.3 | P | 176.6 | P | 285.5 | P | --- |
| 249478_at | 9.1 | A | 5.4 | A | 10.2 | A | 9 | A | --- |
| 249479_at | 30.5 | A | 30.1 | A | 62.6 | A | 22.2 | A | --- |
| 249480_s_at | 402.3 | P | 531.2 | P | 534.2 | P | 468.2 | P | --- |
| 249481_at | 142.6 | P | 147.5 | P | 112.2 | P | 99 | P | --- |
| 249482_at | 5292.5 | P | 7738.2 | P | 6754.5 | P | 7767.4 | P | --- |
| 249483_at | 422.8 | P | 457.3 | P | 431.1 | P | 429.7 | P | --- |
| 249484_at | 1.9 | A | 10.7 | A | 4.4 | A | 30 | A | BR6OX1 |
| 249485_at | 106.1 | P | 193.8 | P | 92.9 | P | 128.9 | P | --- |
| 249486_at | 325.5 | P | 189.8 | P | 249.4 | P | 202.4 | P | --- |
| 249487_at | 418.7 | P | 447 | P | 405 | P | 452.2 | P | TAP2 |
| 249488_at | 7.6 | A | 1.9 | A | 13.2 | A | 2.4 | A | --- |
| 249489_at | 59.9 | P | 77.2 | P | 74.4 | A | 48.9 | A | --- |
| 249490_s_at | 2.7 | A | 38 | A | 23.5 | A | 17.2 | A | --- |
| 249491_at | 0.6 | A | 4.5 | A | 3.4 | A | 0.6 | A | --- |
| 249492_at | 13.3 | A | 18 | A | 16.9 | A | 18 | A | GER2 |
| 249493_at | 476.3 | P | 608 | P | 619.9 | P | 575.8 | P | --- |
| 249494_at | 3811.1 | P | 1774.8 | P | 2857.8 | P | 2125.3 | P | --- |
| 249495_at | 9.3 | A | 17.5 | A | 1.6 | A | 2.1 | A | GLP6 |
| 249496_s_at | 20.3 | A | 27.3 | A | 25.6 | A | 12.2 | A | --- |
| 249497_at | 68.3 | P | 56.6 | P | 129 | P | 45.4 | P | --- |
| 249498_at | 22 | A | 28.7 | A | 4.8 | A | 24.4 | A | --- |
| 249499_at | 218.3 | P | 253 | P | 186.6 | P | 217.2 | P | --- |
| 249500_at | 3.6 | A | 5.1 | A | 13.4 | A | 3.1 | A | ATEXPA21 |
| 249501_s_at | 10.9 | A | 1.2 | A | 23.8 | A | 20.2 | A | ATEXP26 |
| 249502_s_at | 3.8 | A | 2.4 | A | 5.3 | A | 2 | A | ATEXPA23 |
| 249503_at | 2.9 | A | 1.8 | A | 13 | A | 14.6 | A | ATEXPA24 |
| 249504_at | 76.8 | P | 146.8 | P | 140.8 | P | 81.7 | P | --- |
| 249505_at | 25 | A | 34.9 | A | 26.7 | A | 26.1 | A | --- |
| 249506_at | 298.2 | P | 229.5 | P | 370.6 | P | 210.9 | P | --- |
| 249507_at | 23.8 | A | 27.5 | A | 27.8 | A | 19.7 | P | --- |
| 249508_at | 289.7 | P | 350.4 | P | 231.1 | P | 282 | P | --- |
| 249509_at | 22.1 | A | 19.5 | A | 27.4 | P | 5.3 | A | --- |
| 249510_at | 645.7 | P | 702.7 | P | 558.6 | P | 760.2 | P | --- |
| 249511_at | 3.5 | A | 7.4 | A | 4 | A | 10.2 | A | --- |
| 249512_at | 2010.3 | P | 2274 | P | 2101.1 | P | 1963.5 | P | --- |
| 249513_at | 153.2 | P | 227.2 | P | 231.7 | P | 184.3 | P | --- |
| 249514_at | 3253.3 | P | 4243 | P | 3208.4 | P | 3877.9 | P | GRF3 |
| 249515_at | 3852.3 | P | 1579.2 | P | 3232.9 | P | 1948.3 | P | --- |
| 249516_s_at | 22.3 | P | 24.8 | A | 2.3 | A | 1.8 | A | --- |
| 249517_at | 3.6 | A | 3.9 | A | 3.1 | A | 12.6 | A | --- |
| 249518_at | 108.9 | P | 88.6 | P | 111.2 | P | 81.6 | A | --- |
| 249519_at | 2191.6 | P | 2849.2 | P | 2381.6 | P | 2594.5 | P | APE1 |
| 249520_at | 3.8 | A | 5.5 | A | 5.7 | A | 1.8 | A | --- |
| 249521_at | 185.1 | P | 214.7 | P | 199.1 | P | 212.8 | P | --- |
| 249522_at | 20.1 | A | 32.5 | P | 25 | A | 18 | A | --- |
| 249523_at | 715.2 | P | 868.9 | P | 825 | P | 775.4 | P | ACYB-1 |
| 249524_at | 1101.1 | P | 1220 | P | 1082.3 | P | 1178.3 | P | --- |
| 249525_at | 1660.1 | P | 1508.4 | P | 1277.9 | P | 1431.5 | P | APE1 |
| 249526_at | 214.7 | P | 279 | P | 186 | P | 225.3 | P | --- |
| 249527_at | 502.4 | P | 98.9 | P | 405.6 | P | 77.8 | P | --- |
| 249528_at | 301.6 | P | 241.9 | P | 273.3 | P | 225.2 | P | --- |
| 249529_at | 184.8 | P | 127.5 | P | 106.5 | P | 120.3 | P | --- |
| 249530_at | 24.6 | A | 27.3 | A | 49 | A | 3.5 | A | --- |
| 249531_at | 3.6 | A | 49.9 | A | 37 | A | 21 | A | AtGDU7 |
| 249532_at | 92.9 | P | 71.5 | P | 103.2 | P | 50.5 | P | --- |
| 249533_at | 11.6 | A | 4 | A | 45.4 | A | 7.9 | A | --- |
| 249534_at | 5.3 | A | 1.6 | A | 24 | A | 9.7 | A | AtbZIP43 |
| 249535_at | 88.2 | P | 90.7 | P | 70.6 | P | 90 | P | --- |
| 249536_at | 1.5 | A | 0.8 | A | 6.2 | A | 1.9 | A | --- |
| 249537_at | 1237.9 | P | 1065.1 | P | 990.5 | P | 1173 | P | --- |
| 249538_at | 478.1 | P | 386.1 | P | 456.9 | P | 391.8 | P | --- |
| 249539_at | 13.2 | A | 17.6 | A | 32.3 | A | 5.9 | A | --- |
| 249540_at | 74.1 | P | 38.4 | M | 105.2 | P | 38.1 | P | --- |
| 249541_at | 34.7 | P | 16.4 | P | 25.2 | P | 15 | A | --- |
| 249542_at | 171.5 | P | 213.6 | P | 132.8 | P | 223.4 | P | NF-YC12 |
| 249543_at | 3.9 | A | 3.1 | A | 5 | A | 3.4 | A | --- |
| 249544_at | 185.8 | P | 330.9 | P | 207.7 | P | 298.6 | P | ASF1B |
| 249545_at | 85.9 | P | 64.7 | P | 76.7 | P | 75.4 | P | --- |
| 249546_at | 63.4 | P | 86.1 | P | 80.2 | P | 118.8 | P | PMI15 |
| 249547_at | 16.8 | A | 6.9 | A | 6.3 | A | 30.2 | A | --- |
| 249548_at | 3.9 | A | 3.1 | A | 5.6 | A | 2.8 | A | --- |
| 249549_at | 17.6 | A | 1.2 | A | 2 | A | 1.3 | A | --- |
| 249550_at | 80 | P | 130 | P | 66 | M | 75.1 | P | --- |
| 249551_at | 263.3 | P | 300.7 | P | 239 | P | 348.6 | P | --- |
| 249552_s_at | 60.8 | P | 88.4 | P | 81.7 | A | 81.4 | P | --- |
| 249553_at | 14.9 | A | 12.5 | A | 30.2 | A | 19 | A | --- |
| 249554_at | 268.8 | P | 570.6 | P | 396.4 | P | 501.3 | P | --- |
| 249555_at | 117.1 | A | 155.1 | P | 76.2 | A | 171.3 | P | --- |
| 249556_at | 24.3 | A | 1.7 | A | 24.8 | A | 2.8 | A | --- |
| 249557_at | 27.2 | A | 35.7 | A | 66.4 | P | 31.7 | A | PR5K |
| 249558_at | 41.5 | P | 38.3 | P | 48.5 | P | 49.6 | P | --- |
| 249559_at | 7.3 | A | 23.7 | A | 15.8 | A | 46.5 | A | --- |
| 249560_at | 2.9 | A | 3.4 | A | 6.5 | A | 1.6 | A | LCR80 |
| 249561_at | 97.1 | P | 79.8 | P | 104.7 | P | 83.4 | P | --- |
| 249562_at | 20.8 | A | 30.3 | A | 45.3 | P | 19.3 | A | --- |
| 249563_at | 237.2 | P | 278.7 | P | 173.9 | P | 274.9 | P | --- |
| 249564_at | 2.5 | A | 2.6 | A | 4.5 | A | 3 | A | --- |
| 249565_at | 10.6 | A | 20.9 | P | 27.4 | M | 6 | A | --- |
| 249566_at | 53.4 | A | 45.3 | A | 59.4 | A | 18.8 | A | CYP735A1 |
| 249567_at | 276.1 | P | 129.8 | P | 269.7 | P | 185.9 | P | --- |
| 249568_at | 7.9 | A | 14.9 | A | 21.9 | A | 23.3 | A | --- |
| 249569_at | 56.5 | P | 68.1 | P | 48 | A | 78.4 | P | --- |
| 249570_at | 16.7 | A | 3.3 | A | 47.2 | A | 7.2 | A | --- |
| 249571_at | 2.7 | A | 4 | A | 21.3 | A | 19.2 | A | --- |
| 249572_at | 94.8 | A | 104 | P | 133.9 | A | 106.1 | P | EMB2656 |
| 249573_at | 26 | A | 9.5 | A | 4.3 | A | 19.8 | A | UBQ9 |
| 249574_at | 36.3 | A | 82 | A | 99 | A | 83.5 | A | PDLP7 |
| 249575_at | 133.4 | P | 182.1 | P | 131.4 | P | 157.8 | P | --- |
| 249576_at | 30.8 | P | 107.1 | P | 80.2 | P | 133.3 | P | --- |
| 249577_at | 135.6 | P | 123.2 | P | 179.6 | P | 101.3 | P | --- |
| 249578_at | 25.9 | A | 11.6 | A | 20.4 | A | 2.6 | A | --- |
| 249579_at | 490.2 | P | 470.2 | P | 451 | P | 515.4 | P | ATARLA1A |
| 249580_at | 1362.5 | P | 697.4 | P | 965.7 | P | 730.5 | P | --- |
| 249581_at | 4951.4 | P | 11749.3 | P | 4933.3 | P | 11004.9 | P | GSR 1 |
| 249582_at | 3735.6 | P | 3994.5 | P | 3249.3 | P | 3892 | P | CAM1 |
| 249583_at | 681.8 | P | 775 | P | 751.7 | P | 750.4 | P | TCH2 |

| | | | | | | | | | |
|---|---|---|---|---|---|---|---|---|---|
| 249584_s_at | 19 | A | 29.7 | A | 8.5 | A | 26.9 | A | NIP4;1 |
| 249585_at | 957.3 | P | 848.3 | P | 673 | P | 833.3 | P | OXP1 |
| 249586_at | 3.4 | A | 6.4 | A | 5.5 | A | 5.3 | A | --- |
| 249587_at | 1.5 | A | 1 | A | 3 | A | 17.9 | A | --- |
| 249588_at | 67.5 | P | 99.5 | P | 61.4 | A | 78.9 | P | --- |
| 249589_at | 978.5 | P | 937 | P | 1114.2 | P | 877.5 | P | SOS4 |
| 249590_at | 2.8 | A | 9.1 | A | 7.3 | A | 7.5 | A | --- |
| 249591_at | 1.6 | A | 1.7 | A | 5.2 | A | 3.2 | A | --- |
| 249592_at | 229.5 | P | 333.3 | P | 403.3 | P | 351.8 | P | --- |
| 249593_at | 15.8 | A | 1.2 | A | 28.1 | A | 15.7 | A | --- |
| 249594_at | 21.1 | A | 2.9 | A | 24.9 | A | 15.5 | A | --- |
| 249595_at | 313.7 | P | 464 | P | 215.7 | P | 314 | P | --- |
| 249596_s_at | 31.6 | P | 57.6 | P | 26.8 | A | 40.2 | P | --- |
| 249597_at | 1.1 | A | 7.7 | A | 1.4 | A | 1.2 | A | --- |
| 249598_at | 17 | P | 19.7 | P | 9.9 | A | 14.7 | A | --- |
| 249599_at | 49.4 | P | 140.3 | P | 111.3 | P | 146.2 | P | --- |
| 249600_s_at | 74.7 | P | 84.2 | P | 86.7 | P | 94.7 | P | --- |
| 249601_at | 50.3 | A | 34.7 | A | 59.4 | P | 21.4 | A | --- |
| 249602_at | 29 | A | 20.7 | A | 20.6 | A | 12.3 | A | --- |
| 249603_at | 18.5 | A | 30.8 | A | 42.9 | A | 29.3 | M | --- |
| 249604_x_at | 27.6 | A | 40.6 | P | 33.6 | P | 20 | P | --- |
| 249605_at | 2 | A | 5 | A | 6.2 | A | 3.8 | A | --- |
| 249606_at | 252.2 | P | 313 | P | 311.7 | P | 299 | P | RVE2 |
| 249607_at | 7.2 | A | 23.3 | A | 49.5 | A | 4.6 | A | --- |
| 249608_at | 769.8 | P | 970.6 | P | 987.2 | P | 946.4 | P | --- |
| 249609_at | 11.3 | A | 10.4 | A | 14.5 | A | 0.7 | A | --- |
| 249610_at | 2450.7 | P | 2620.4 | P | 2193 | P | 2552 | P | --- |
| 249611_at | 408 | P | 468.7 | P | 460.5 | P | 470.9 | P | ATSRL1 |
| 249612_at | 250.8 | P | 264.3 | P | 255.5 | P | 270.9 | P | --- |
| 249613_at | 352 | P | 444.7 | P | 347.5 | P | 407 | P | --- |
| 249614_at | 29.5 | A | 34.5 | P | 36.2 | A | 104.6 | P | WSD1 |
| 249615_x_at | 55.4 | A | 4.6 | A | 7 | A | 15.4 | A | AGL105 |
| 249616_s_at | 107.2 | P | 77.3 | P | 76.2 | P | 56.4 | P | --- |
| 249617_at | 4.7 | A | 30.7 | A | 15.5 | A | 6.7 | A | --- |
| 249618_at | 68.2 | A | 54.8 | A | 66.9 | A | 40.9 | P | --- |
| 249619_at | 93.2 | P | 87.8 | P | 98.1 | P | 93.9 | P | GORK |
| 249620_at | 3.7 | A | 7.1 | A | 11.9 | A | 9.1 | A | --- |
| 249621_at | 125.8 | P | 141.5 | P | 184.9 | P | 176.7 | P | --- |
| 249622_at | 107.2 | P | 115.1 | P | 106.1 | P | 96.2 | P | --- |
| 249623_at | 1.2 | A | 1.9 | A | 3.2 | A | 1.6 | A | --- |
| 249624_at | 70.1 | P | 48.4 | P | 62.3 | P | 68.8 | P | --- |
| 249625_at | 441.6 | P | 479.8 | P | 394.5 | P | 445.5 | P | --- |
| 249626_at | 253.2 | P | 248.4 | P | 153.8 | P | 175.4 | P | --- |
| 249627_at | 3161.1 | P | 3374.3 | P | 3592.4 | P | 3264.8 | P | EMB1467 |
| 249628_at | 137.6 | P | 271.6 | P | 117.1 | P | 217.5 | P | --- |
| 249629_at | 65.7 | P | 139 | P | 123 | M | 95 | P | --- |
| 249630_s_at | 3.6 | A | 10.4 | A | 18.1 | A | 21.9 | A | --- |
| 249631_at | 16.6 | A | 33.5 | M | 42.6 | M | 19.6 | A | --- |
| 249632_at | 26.1 | A | 48.7 | A | 53.6 | A | 23.6 | A | --- |
| 249633_at | 65.1 | P | 79.6 | P | 73.9 | P | 81.3 | P | SUS5 |
| 249634_at | 0.9 | A | 0.8 | A | 1 | A | 1.1 | A | --- |
| 249635_at | 26.5 | A | 42.4 | A | 44 | P | 59.7 | A | ATGSL09 |
| 249636_at | 614.7 | P | 690.9 | P | 736.5 | P | 896.5 | P | BGLU42 |
| 249637_at | 21.8 | A | 27.6 | A | 13.9 | A | 9.2 | A | --- |
| 249638_at | 3018.1 | P | 2691.8 | P | 3218.2 | P | 2633.9 | P | --- |
| 249639_at | 111.4 | P | 161.1 | P | 128.8 | P | 146.2 | P | --- |
| 249640_at | 233.4 | P | 293.3 | P | 221.7 | P | 280.7 | P | CAT3 |
| 249641_at | 273.9 | P | 234.5 | P | 308.7 | P | 262.1 | P | DegP10 |
| 249642_at | 23.7 | A | 6.1 | A | 22.4 | A | 10.6 | A | --- |
| 249643_at | 12.1 | A | 16.3 | A | 3 | A | 22.1 | A | --- |
| 249644_at | 114.3 | P | 205.5 | P | 160.3 | P | 173.1 | P | --- |
| 249645_at | 151.1 | P | 129.3 | P | 266.6 | P | 224.5 | P | THI2.2 |
| 249646_at | 5.7 | A | 5.2 | A | 1.4 | A | 12.6 | A | --- |
| 249647_at | 32.7 | A | 56.8 | A | 69.2 | P | 23 | A | --- |
| 249648_at | 4.5 | A | 10.5 | A | 11.5 | A | 25 | M | --- |
| 249649_at | 17.1 | A | 3 | A | 20.3 | A | 11.3 | A | CHX24 |
| 249650_at | 132.8 | P | 137.9 | P | 177.5 | P | 160.7 | P | SEF |
| 249651_at | 130.2 | P | 280.1 | P | 224 | P | 243 | P | ARF8 |
| 249652_at | 523.1 | P | 684 | P | 577.5 | P | 574.5 | P | --- |
| 249653_at | 21.8 | A | 2.3 | A | 4.7 | A | 5.9 | A | --- |
| 249654_at | 1.1 | A | 3.8 | A | 1.8 | A | 0.9 | A | --- |
| 249655_at | 0.7 | A | 15.3 | A | 21.2 | A | 11.3 | A | --- |
| 249656_at | 125.5 | P | 125.2 | P | 148.1 | P | 111.7 | P | --- |
| 249657_at | 3.1 | A | 6.7 | A | 17.4 | A | 5.1 | A | --- |
| 249658_s_at | 4503.1 | P | 5905.5 | P | 4724.8 | P | 5089.4 | P | --- |
| 249659_s_at | 318.7 | P | 591.7 | P | 341.9 | P | 636.3 | P | --- |
| 249660_at | 0.9 | A | 10.6 | A | 2.4 | A | 2 | A | --- |
| 249661_x_at | 21.3 | A | 1.2 | A | 1.2 | A | 1.6 | A | --- |
| 249662_s_at | 21 | P | 33.9 | P | 33 | P | 22 | M | --- |
| 249663_s_at | 1.9 | A | 13.5 | A | 6.4 | A | 3.3 | A | --- |
| 249664_at | 21.9 | A | 1.1 | A | 10.3 | A | 17.9 | A | --- |
| 249665_at | 48.4 | P | 29.7 | P | 46.7 | A | 40 | P | --- |
| 249666_at | 377.2 | P | 374.5 | P | 412.2 | P | 394.1 | P | PHYC |
| 249667_at | 19.4 | A | 8.3 | A | 12 | A | 4.6 | A | --- |
| 249668_at | 10 | A | 29.5 | A | 49.9 | A | 5.7 | A | --- |
| 249669_at | 19.6 | A | 2.6 | A | 8.8 | A | 0.8 | A | --- |
| 249670_at | 6 | A | 12.2 | A | 3.8 | A | 1.2 | A | --- |
| 249671_at | 8.1 | A | 21.6 | A | 3.6 | A | 8 | A | LBD35 |
| 249672_at | 580.4 | P | 558.2 | P | 818.3 | P | 487.7 | P | --- |
| 249673_at | 1.1 | A | 17.9 | A | 16.6 | A | 14.6 | A | CYP79A4P |
| 249674_at | 143 | P | 143.2 | P | 178.9 | P | 168.9 | P | --- |
| 249675_at | 35.6 | P | 7.9 | A | 47.1 | P | 11.9 | A | --- |
| 249676_at | 47.2 | P | 41.1 | A | 48.4 | A | 33.6 | A | --- |
| 249677_at | 608.3 | P | 1092.4 | P | 551.6 | P | 963.8 | P | --- |
| 249678_at | 660.1 | P | 698 | P | 526.4 | P | 753.6 | P | --- |
| 249679_at | 981.2 | P | 942.9 | P | 1018.8 | P | 977.6 | P | --- |
| 249680_s_at | 17.2 | A | 26.6 | P | 29.1 | A | 22.8 | A | --- |
| 249681_s_at | 2 | A | 1 | A | 2.5 | A | 2.1 | A | --- |
| 249682_at | 3.1 | A | 0.5 | A | 6.3 | A | 0.7 | A | --- |
| 249683_x_at | 1.3 | A | 6 | A | 8.3 | A | 9.7 | A | --- |
| 249684_s_at | 10.2 | A | 2.1 | A | 0.7 | A | 1.3 | A | --- |
| 249685_at | 260.7 | P | 340.9 | P | 286 | P | 385.8 | P | CCB3 |
| 249686_at | 16 | A | 29 | A | 47.7 | A | 3.8 | A | CYP716A2 |
| 249687_at | 19 | A | 29.5 | P | 38.9 | A | 27.7 | A | ATPEN3 |
| 249688_at | 2945.8 | P | 2493.1 | P | 2742.1 | P | 2556.9 | P | --- |
| 249689_at | 0.8 | A | 6.6 | A | 1.2 | A | 3.2 | A | --- |
| 249690_at | 1029.2 | P | 1049.5 | P | 1086.1 | P | 994.9 | P | --- |
| 249691_at | 1473.3 | P | 1395.6 | P | 1324.4 | P | 1386.5 | P | HCF109 |
| 249692_s_at | 11.6 | A | 23.6 | A | 18 | A | 36.2 | A | --- |

| | | | | | | | | | |
|---|---|---|---|---|---|---|---|---|---|
| 249693_at | 182.6 | P | 281.3 | P | 230.5 | P | 254.1 | P | AHK2 |
| 249694_at | 831.2 | P | 1143.7 | P | 952.5 | P | 968.2 | P | G6PD1 |
| 249695_at | 19.7 | A | 21.1 | A | 14.2 | A | 15.6 | A | --- |
| 249696_at | 1.5 | A | 1.1 | A | 2.2 | A | 1.2 | A | --- |
| 249697_at | 0.5 | A | 0.2 | A | 1.1 | A | 0.3 | A | --- |
| 249698_at | 72.8 | A | 78.7 | P | 122.3 | P | 112.2 | P | MIS12 |
| 249699_at | 11.2 | A | 12 | A | 38.4 | A | 19.4 | A | --- |
| 249700_at | 6843.9 | P | 5782.5 | P | 7850.8 | P | 6661.4 | P | --- |
| 249701_at | 886.6 | P | 1102.1 | P | 927.3 | P | 1033.6 | P | --- |
| 249702_at | 175.3 | P | 169.1 | P | 154.3 | P | 127.8 | P | --- |
| 249703_at | 406.7 | P | 485.5 | P | 392 | P | 443.1 | P | --- |
| 249704_at | 21.8 | A | 33.7 | A | 32.5 | A | 17.8 | A | TT2 |
| 249705_at | 2.1 | A | 4.5 | A | 7.3 | A | 4.6 | A | --- |
| 249706_at | 4.1 | A | 23.1 | A | 20.2 | A | 24.1 | A | HDA7 |
| 249707_at | 22.9 | A | 18.6 | A | 15.9 | A | 9.7 | A | --- |
| 249708_at | 12.6 | A | 2.7 | A | 34.3 | A | 4.8 | A | --- |
| 249709_at | 53.7 | A | 71.5 | P | 27.4 | A | 94.3 | P | iqd33 |
| 249710_at | 10824 | P | 1772 | P | 11440 | P | 2251.8 | P | GS2 |
| 249711_at | 4570.6 | P | 3986.6 | P | 3059.2 | P | 4427.3 | P | --- |
| 249712_at | 1374.4 | P | 1408.3 | P | 1599.3 | P | 1285.6 | P | LSP1 |
| 249713_at | 40.7 | A | 47.9 | A | 52.1 | A | 25.5 | P | --- |
| 249714_at | 27.3 | A | 42.8 | A | 44.7 | A | 29.5 | P | --- |
| 249715_at | 3.8 | A | 3.6 | A | 18.1 | A | 19.6 | A | --- |
| 249716_at | 36 | P | 26.6 | P | 41.9 | A | 49.8 | P | SAP |
| 249717_at | 565 | P | 705.8 | P | 521.4 | P | 752.1 | P | --- |
| 249718_at | 176.1 | P | 443.1 | P | 408.2 | P | 436.3 | P | --- |
| 249719_at | 2912.5 | P | 2428.3 | P | 1858.2 | P | 2096 | P | --- |
| 249720_at | 413.9 | P | 354.9 | P | 433.6 | P | 347.5 | P | FIM2 |
| 249721_at | 46.3 | A | 66.9 | A | 64 | A | 104.6 | A | LSU4 |
| 249722_at | 201.1 | P | 191.1 | P | 203.7 | P | 188.8 | P | --- |
| 249723_at | 559.5 | P | 576.9 | P | 568.7 | P | 532.2 | P | --- |
| 249724_at | 117.2 | P | 148.9 | P | 168.4 | P | 140.2 | P | --- |
| 249725_at | 12.2 | A | 2.3 | A | 2.4 | A | 7.8 | A | --- |
| 249726_at | 0.7 | A | 15.4 | A | 20.6 | P | 0.8 | A | --- |
| 249727_at | 109.2 | P | 113.2 | P | 117.8 | P | 100.2 | P | MRU1 |
| 249728_at | 32.9 | P | 60.5 | P | 96.9 | P | 58.9 | P | --- |
| 249729_at | 3.7 | A | 5.4 | A | 3 | A | 4.4 | A | --- |
| 249730_at | 591.3 | P | 447.3 | P | 343.4 | P | 373.8 | P | --- |
| 249731_at | 47.1 | A | 10.3 | A | 6 | A | 13.4 | A | CID13 |
| 249732_at | 2517.7 | P | 129 | P | 2594.8 | P | 223.3 | P | --- |
| 249733_at | 1078.2 | P | 1156.3 | P | 1319.4 | P | 1156.2 | P | emb2024 |
| 249734_at | 320 | P | 353.2 | P | 245.1 | P | 307.8 | P | IRE1-1 |
| 249735_at | 372.6 | P | 449.1 | P | 519.4 | P | 499 | P | --- |
| 249736_at | 854.9 | P | 904.2 | P | 1016.3 | P | 979.6 | P | --- |
| 249737_at | 44.9 | A | 25 | P | 58.4 | M | 31.9 | A | --- |
| 249738_at | 27.7 | A | 15 | A | 29.1 | A | 5.7 | A | --- |
| 249739_at | 535.8 | P | 697.6 | P | 498.2 | P | 545.9 | P | TTG1 |
| 249740_at | 613.1 | P | 555.4 | P | 564.6 | P | 591.8 | P | --- |
| 249741_at | 409.6 | P | 372.6 | P | 387.9 | P | 427.7 | P | APRR5 |
| 249742_at | 3643.4 | P | 3944.9 | P | 3664.5 | P | 3377.4 | P | --- |
| 249743_at | 3.6 | A | 1.1 | A | 13.3 | A | 3.9 | A | BGLU31 |
| 249744_at | 10.9 | A | 1.3 | A | 1.7 | A | 12 | A | BGLU32 |
| 249745_at | 31.9 | A | 21.9 | A | 64.1 | M | 48.8 | P | AtPP2-B12 |
| 249746_at | 251.7 | P | 356.8 | P | 296.3 | P | 282.9 | P | TIP |
| 249747_at | 145.6 | P | 82.1 | P | 201.5 | P | 91.6 | P | --- |
| 249748_at | 391.9 | P | 632.6 | P | 405.3 | P | 575.6 | P | --- |
| 249749_at | 75 | P | 41 | P | 79 | P | 51.1 | P | BIN4 |
| 249750_at | 1650.6 | P | 1387.6 | P | 1422.2 | P | 1215 | P | --- |
| 249751_at | 1689.5 | P | 1468.6 | P | 1563.3 | P | 1466.1 | P | --- |
| 249752_at | 255.7 | P | 401.2 | P | 344.4 | P | 402.4 | P | LSU2 |
| 249753_at | 1073.6 | P | 1500.2 | P | 842.1 | P | 1388.4 | P | --- |
| 249754_at | 463.9 | P | 1826 | P | 491.3 | P | 1324.5 | P | DMR6 |
| 249755_at | 51.6 | P | 102.3 | P | 69.9 | P | 100.1 | P | --- |
| 249756_at | 7.3 | A | 2.9 | A | 5.4 | A | 3.2 | A | --- |
| 249757_at | 14.2 | A | 13.1 | A | 10.2 | A | 12.7 | A | --- |
| 249758_at | 228.4 | P | 234.4 | P | 214.1 | P | 175.3 | P | --- |
| 249759_at | 445.7 | P | 612 | P | 724.8 | P | 599.2 | P | YSL2 |
| 249760_at | 1.2 | A | 1.5 | A | 27.4 | A | 1.7 | A | TPS21 |
| 249761_at | 26 | P | 25.5 | A | 40.2 | M | 29.8 | A | --- |
| 249762_at | 212.6 | P | 182 | P | 232.2 | P | 198.4 | P | --- |
| 249763_at | 187.4 | P | 156.8 | P | 213.2 | P | 178.2 | P | --- |
| 249764_at | 530.6 | P | 568.5 | P | 676.4 | P | 578.4 | P | MIND |
| 249765_at | 331.1 | P | 401.4 | P | 266.8 | P | 326.5 | P | SLAH3 |
| 249766_at | 1.1 | A | 2.4 | A | 3.5 | A | 14.3 | A | --- |
| 249767_at | 107.8 | P | 61.1 | P | 111.4 | P | 71 | P | --- |
| 249768_at | 33.4 | P | 12.7 | A | 49.3 | M | 12.4 | A | --- |
| 249769_at | 191.9 | P | 540.9 | P | 240 | P | 442.8 | P | SIGE |
| 249770_at | 1.7 | A | 2.3 | A | 2 | A | 1.6 | A | WRKY30 |
| 249771_at | 29.1 | A | 4.3 | A | 33.7 | A | 20 | A | --- |
| 249772_at | 30.5 | A | 42.5 | A | 37.2 | A | 58.4 | A | --- |
| 249773_at | 12.6 | A | 30.4 | A | 4.6 | A | 16.9 | A | SQP2 |
| 249774_at | 762.3 | P | 814.7 | P | 925 | P | 933.6 | P | SQP1 |
| 249775_at | 1317.8 | P | 905.5 | P | 1366.5 | P | 816.2 | P | SQE6 |
| 249776_at | 324.1 | P | 352.9 | P | 326.4 | P | 316.3 | P | --- |
| 249777_at | 962.3 | P | 1608.9 | P | 973.6 | P | 1022.8 | P | --- |
| 249778_at | 2689.3 | P | 2884.1 | P | 2644.4 | P | 2799.7 | P | --- |
| 249779_at | 51.7 | A | 71.9 | A | 33.5 | A | 33.9 | A | --- |
| 249780_at | 2.8 | A | 2.7 | A | 5 | A | 2.4 | A | --- |
| 249781_at | 1.6 | A | 16.6 | A | 34.2 | A | 20.3 | A | --- |
| 249782_at | 360.5 | P | 370.1 | P | 417.3 | A | 349.6 | P | --- |
| 249783_at | 214 | P | 137.1 | P | 131.4 | P | 126.5 | P | SOS3 |
| 249784_at | 299.1 | P | 102.6 | P | 285 | P | 120.6 | P | --- |
| 249785_at | 1357.7 | P | 1261.5 | P | 1409.7 | P | 1334 | P | SSI1 |
| 249786_at | 275.8 | P | 213.3 | P | 231.2 | P | 179.6 | P | ABIL3 |
| 249787_at | 303.1 | P | 297.5 | P | 261.3 | P | 308.4 | P | --- |
| 249788_at | 26.6 | A | 106.8 | A | 46.8 | A | 88.1 | P | ATXR6 |
| 249789_at | 252 | P | 214.3 | P | 210.4 | P | 206.7 | P | --- |
| 249790_at | 218 | P | 66.3 | P | 157.4 | P | 94.1 | P | --- |
| 249791_at | 288.6 | P | 264.1 | P | 248.3 | P | 209 | P | AAP7 |
| 249792_at | 150.6 | P | 133.4 | P | 186.4 | P | 174.3 | P | PHS1 |
| 249793_at | 115.8 | P | 54.7 | P | 88 | P | 81 | P | --- |
| 249794_at | 265.8 | P | 168.5 | P | 224 | P | 207.1 | P | AtCXE18 |
| 249795_at | 5367.6 | P | 5349.8 | P | 6297.7 | P | 5411.7 | P | RPS11-BETA |
| 249796_at | 1853.3 | P | 1821.1 | P | 1441.7 | P | 1894.3 | P | --- |
| 249797_at | 86.5 | P | 104.4 | P | 82.5 | P | 108.6 | P | --- |
| 249798_at | 294.7 | P | 331.1 | P | 363.5 | P | 375.3 | P | --- |
| 249799_at | 937 | P | 1273.6 | P | 870.8 | P | 962 | P | LCB2 |
| 249800_at | 2292.2 | P | 2153.1 | P | 2751.2 | P | 2220.7 | P | MTN3 |
| 249801_at | 239.3 | P | 223.8 | P | 263.4 | P | 229.9 | P | CDPK9 |

| | | | | | | | | | |
|---|---|---|---|---|---|---|---|---|---|
| 249802_at | 40.1 | P | 47.5 | P | 49.9 | P | 35.3 | P | --- |
| 249803_at | 14.4 | A | 10.2 | A | 43.4 | A | 19.8 | A | --- |
| 249804_at | 1.4 | A | 2.6 | A | 1.3 | A | 7.7 | A | AtGolS5 |
| 249805_at | 55 | P | 72.1 | P | 89.2 | P | 88.6 | P | --- |
| 249806_at | 403.4 | P | 282.8 | P | 394.8 | P | 307.4 | P | --- |
| 249807_at | 112.5 | P | 171.1 | P | 155.1 | P | 130.1 | P | --- |
| 249808_at | 608.7 | P | 579.7 | P | 558.2 | P | 647.5 | P | --- |
| 249809_at | 51.1 | M | 72 | P | 42.5 | A | 70.6 | P | --- |
| 249810_at | 1078.4 | P | 1310.4 | P | 1471.3 | P | 1262.7 | P | --- |
| 249811_at | 378.7 | P | 447.2 | P | 446 | P | 551.1 | P | --- |
| 249812_at | 167.6 | P | 81 | P | 184.2 | P | 107.6 | P | --- |
| 249813_at | 281.2 | P | 433.1 | P | 270.8 | P | 358.4 | P | EMB3009 |
| 249814_at | 2 | A | 3.1 | A | 5.4 | A | 3.1 | A | --- |
| 249815_at | 3830.7 | P | 2487.4 | P | 4832.5 | P | 3589.9 | P | --- |
| 249816_at | 409.6 | P | 315 | P | 354.8 | P | 348 | P | CPSF100 |
| 249817_at | 3950.5 | P | 1844.8 | P | 3952.4 | P | 1933.2 | P | --- |
| 249818_at | 285.3 | P | 548.1 | P | 401.9 | P | 526.9 | P | TUB8 |
| 249819_at | 1.7 | A | 2.2 | A | 3.1 | A | 3 | A | --- |
| 249820_at | 4.7 | A | 6.8 | A | 6.6 | A | 4.8 | A | --- |
| 249821_at | 157 | P | 136.1 | P | 191.8 | P | 112.8 | P | --- |
| 249822_at | 310.5 | P | 199 | P | 267.4 | P | 241 | P | --- |
| 249823_s_at | 18.3 | P | 52.7 | P | 11.9 | A | 34.6 | P | --- |
| 249824_at | 108 | M | 124.2 | P | 110.7 | M | 108.6 | P | --- |
| 249825_at | 492.4 | P | 401.9 | P | 352.3 | P | 428.3 | P | --- |
| 249826_at | 988.5 | P | 1528.1 | P | 841.2 | P | 1437 | P | FSD3 |
| 249827_at | 182.7 | P | 263.5 | P | 192.3 | P | 185.1 | P | --- |
| 249828_at | 999 | P | 786.3 | P | 1048.1 | P | 878.2 | P | --- |
| 249829_at | 791.6 | P | 820.6 | P | 757.7 | P | 726.9 | P | PDF5 |
| 249830_at | 340.4 | P | 329.6 | P | 404 | P | 302.8 | P | PYRD |
| 249831_at | 342.4 | P | 416.4 | P | 288.5 | P | 441.9 | P | --- |
| 249832_at | 73.7 | P | 91.3 | P | 87.7 | P | 86.3 | P | --- |
| 249833_at | 121.9 | P | 154.4 | P | 123.4 | P | 155.2 | P | --- |
| 249834_at | 1094.5 | P | 1068.9 | P | 1111 | P | 1202.5 | P | FTRA1 |
| 249835_s_at | 828.9 | P | 672.4 | P | 541.6 | P | 629.9 | P | --- |
| 249836_at | 231.9 | P | 266.1 | P | 313.5 | P | 303.6 | P | HMGB6 |
| 249837_at | 40.1 | P | 64.9 | P | 67.2 | P | 58.8 | P | --- |
| 249838_at | 191.1 | P | 140.6 | P | 161.5 | P | 197.8 | P | --- |
| 249839_at | 200.6 | P | 190.5 | P | 227.5 | P | 201.4 | P | --- |
| 249840_at | 1051.3 | P | 1182.1 | P | 1042.2 | P | 1053.4 | P | ATLCBK1 |
| 249841_at | 146 | P | 183.5 | P | 184.1 | P | 156.2 | P | --- |
| 249842_at | 366.7 | P | 442.4 | P | 383.5 | P | 429.5 | P | --- |
| 249843_at | 141.3 | P | 148.2 | P | 136.4 | P | 110.4 | P | SGS3 |
| 249844_at | 197 | P | 124.3 | P | 213.4 | P | 161.2 | P | --- |
| 249845_at | 495.3 | P | 552.7 | P | 519.2 | P | 553.9 | P | --- |
| 249846_at | 1020.9 | P | 910.5 | P | 834.6 | P | 882.5 | P | --- |
| 249847_at | 537.4 | P | 549.3 | P | 755.7 | P | 447 | P | SCPL34 |
| 249848_at | 2.5 | A | 13.3 | A | 4.9 | A | 9.3 | A | NIC3 |
| 249849_at | 132.7 | P | 92.5 | P | 94.2 | P | 108.9 | P | NIC2 |
| 249850_at | 4377.7 | P | 3781.3 | P | 3852.6 | P | 4218.6 | P | --- |
| 249851_at | 27.3 | A | 38 | A | 57.1 | A | 33.9 | A | TT16 |
| 249852_at | 35.7 | P | 32.8 | P | 48.4 | P | 49.4 | P | STP11 |
| 249853_at | 93.2 | P | 63.2 | P | 76.8 | P | 45.4 | A | ATSTE14A |
| 249854_at | 34.7 | P | 36.2 | A | 34.9 | A | 24.8 | A | --- |
| 249855_at | 12.8 | A | 8.2 | A | 37.7 | A | 29.7 | A | --- |
| 249856_at | 30 | A | 31.5 | A | 7.8 | A | 42.2 | P | scpl47 |
| 249857_at | 5 | A | 7.9 | A | 1.4 | A | 2.2 | A | --- |
| 249858_at | 18.9 | A | 30.5 | A | 51.3 | A | 44.2 | A | MYB37 |
| 249859_at | 210.2 | P | 266.4 | P | 284.3 | P | 273.5 | P | --- |
| 249860_at | 1190.3 | P | 436.2 | P | 436.7 | P | 641.2 | P | --- |
| 249861_at | 304 | P | 321.7 | P | 313.8 | P | 277.1 | P | --- |
| 249862_at | 485.9 | P | 527.7 | P | 586.2 | P | 510.5 | P | --- |
| 249863_at | 682 | P | 674.7 | P | 647.4 | P | 654.7 | P | VPS24.1 |
| 249864_at | 1266.8 | P | 1746.4 | P | 1427.3 | P | 1519.8 | P | ATMGT10 |
| 249865_at | 67.6 | P | 72.3 | P | 68.3 | P | 70.3 | P | --- |
| 249866_at | 12382.5 | P | 7600.1 | P | 13633.3 | P | 7645.3 | P | MAM1 |
| 249867_at | 2459.1 | P | 2245.2 | P | 3381.9 | P | 2083.5 | P | IMS2 |
| 249868_at | 20.2 | A | 1 | A | 28.6 | A | 13.9 | A | TET12 |
| 249869_at | 669.5 | P | 443.6 | P | 717 | P | 613.2 | P | AAE17 |
| 249870_at | 185 | P | 245.9 | P | 218 | P | 241.6 | P | TGH |
| 249871_at | 194.3 | P | 202.5 | P | 201 | P | 188.7 | P | --- |
| 249872_at | 99.2 | P | 101.7 | P | 95.9 | P | 109.5 | P | --- |
| 249873_at | 1283.2 | P | 1191.2 | P | 1415.9 | P | 1164 | P | NCLPP7 |
| 249874_at | 588.5 | P | 536.3 | P | 665.1 | P | 645.2 | P | --- |
| 249875_at | 3658.6 | P | 3384.4 | P | 3788.9 | P | 3590.4 | P | HCF136 |
| 249876_at | 4675 | P | 5024.7 | P | 5007 | P | 4879.6 | P | CaS |
| 249877_at | 114.8 | P | 104.8 | P | 132.1 | P | 74.3 | P | HUA2 |
| 249878_at | 146.9 | P | 150.7 | P | 123 | P | 127.7 | P | NF-YB13 |
| 249879_at | 89.9 | P | 101.1 | P | 87.6 | P | 111.1 | P | --- |
| 249880_at | 35.3 | A | 32.7 | A | 26.6 | M | 23.7 | A | --- |
| 249881_at | 13.4 | A | 24 | A | 30 | A | 31.4 | A | CYP86B1 |
| 249882_at | 125.1 | P | 33.5 | A | 134.5 | P | 75.8 | P | --- |
| 249883_at | 1.1 | A | 19.9 | A | 21.6 | A | 0.6 | A | CHX3 |
| 249884_at | 1 | A | 5.3 | A | 2.5 | A | 4.5 | A | CHX9 |
| 249885_at | 82.5 | P | 108.4 | P | 95.3 | P | 120.1 | P | F8H |
| 249886_at | 318.6 | P | 208.9 | P | 192.6 | P | 218.7 | P | --- |
| 249887_at | 32.4 | A | 84.1 | P | 66 | P | 74.1 | P | --- |
| 249888_s_at | 554.9 | P | 581.5 | P | 709.8 | P | 542.1 | P | --- |
| 249889_at | 68.7 | P | 84 | P | 107.5 | P | 88.8 | P | --- |
| 249890_at | 11.1 | A | 43.9 | P | 28.8 | A | 3.5 | A | WRKY38 |
| 249891_at | 16.1 | A | 10.1 | A | 6.7 | A | 36.8 | A | --- |
| 249892_at | 5.9 | A | 48 | A | 52.4 | A | 62 | A | --- |
| 249893_at | 1.7 | A | 1.2 | A | 2.1 | A | 1.6 | A | --- |
| 249894_at | 392.7 | P | 2089.1 | P | 517.7 | P | 1891.5 | P | --- |
| 249895_at | 177.9 | P | 262.9 | P | 101.4 | P | 139.8 | P | FAR1 |
| 249896_at | 106.5 | P | 41 | P | 73.9 | P | 23.1 | P | --- |
| 249897_at | 12.5 | M | 7.4 | A | 27.7 | A | 26.9 | A | --- |
| 249898_at | 835.3 | P | 916.8 | P | 844.9 | P | 938.8 | P | --- |
| 249899_at | 789.9 | P | 776 | P | 779.5 | P | 859.9 | P | --- |
| 249900_at | 971.6 | P | 1357 | P | 1094.7 | P | 1053.1 | P | emb1211 |
| 249901_at | 3189.9 | P | 2585.3 | P | 3705.3 | P | 2786.6 | P | HD2B |
| 249902_at | 21.7 | A | 35.4 | A | 33.9 | A | 29.5 | A | --- |
| 249903_at | 104.4 | P | 138.1 | P | 123.5 | P | 119.4 | P | --- |
| 249904_at | 172.9 | P | 287 | P | 250.4 | P | 264.3 | P | --- |
| 249905_at | 127.5 | P | 166.1 | P | 157.1 | P | 135.8 | P | --- |
| 249906_at | 143.7 | P | 171.7 | P | 224.4 | P | 166.3 | P | --- |
| 249907_at | 48.2 | A | 54.2 | A | 55.1 | A | 68.8 | P | RAD5 |
| 249908_at | 301.4 | P | 397.8 | P | 360.4 | P | 299.8 | P | --- |
| 249909_s_at | 597.2 | P | 715.9 | P | 667.3 | P | 719.1 | P | --- |
| 249910_at | 543.5 | P | 328.8 | P | 410.1 | P | 327.1 | P | ADT5 |

| | | | | | | | | | |
|---|---|---|---|---|---|---|---|---|---|
| 249911_at | 908.3 | P | 1321.8 | P | 1253.9 | P | 1383.7 | P | CSLA02 |
| 249912_at | 360.9 | P | 336.9 | P | 498.4 | P | 366.1 | P | RER1 |
| 249913_at | 11.9 | A | 28.7 | A | 27.6 | A | 8 | A | --- |
| 249914_at | 312.5 | P | 367.9 | P | 287.4 | P | 333 | P | --- |
| 249915_at | 32.4 | P | 32 | P | 57.2 | P | 28.4 | P | --- |
| 249916_at | 668 | P | 818.2 | P | 1193.5 | P | 989.1 | P | HTB2 |
| 249917_at | 101.3 | P | 245.3 | P | 29 | A | 299 | P | --- |
| 249918_at | 1198.4 | P | 2156.7 | P | 927.1 | P | 1486.4 | P | --- |
| 249919_at | 2048.5 | P | 3742.8 | P | 1448.4 | P | 3157.6 | P | --- |
| 249920_at | 155.8 | P | 118.3 | P | 227.7 | P | 123.6 | P | --- |
| 249921_at | 42.7 | A | 38.7 | A | 59.1 | A | 30 | A | --- |
| 249922_at | 5760.1 | P | 7531.7 | P | 6074.9 | P | 6037.1 | P | AILP1 |
| 249923_at | 847 | P | 414.4 | P | 607 | P | 253.9 | P | --- |
| 249924_at | 302.3 | P | 259.1 | P | 285.8 | P | 284 | P | KAPP |
| 249925_at | 496.6 | P | 502.3 | P | 601.9 | P | 443.5 | P | --- |
| 249926_at | 748.2 | P | 713.4 | P | 867.3 | P | 797.5 | P | ECR1 |
| 249927_at | 1117.6 | P | 1419.5 | P | 1064.6 | P | 1383 | P | APL1 |
| 249928_at | 230.5 | P | 159.4 | P | 174.9 | P | 212.7 | P | --- |
| 249929_at | 1280.7 | P | 1477.9 | P | 1774.1 | P | 1519.1 | P | --- |
| 249930_at | 689.1 | P | 816.1 | P | 736.6 | P | 793.5 | P | ATVAMP714 |
| 249931_at | 274.3 | P | 294.8 | P | 307.5 | P | 283.1 | P | QQT1 |
| 249932_at | 47.1 | P | 45.2 | P | 91.6 | P | 55.8 | P | --- |
| 249933_at | 158 | P | 193.7 | P | 110.4 | P | 207.8 | P | --- |
| 249934_at | 10.4 | A | 1.1 | A | 31.3 | A | 6.3 | A | --- |
| 249935_at | 3.4 | A | 11.2 | A | 28.6 | A | 23.4 | A | FAR7 |
| 249936_at | 436.3 | P | 459.1 | P | 443 | P | 396.4 | P | --- |
| 249937_at | 37.9 | A | 28.9 | A | 64.8 | A | 43 | A | --- |
| 249938_at | 378.8 | P | 358.6 | P | 447.4 | P | 378.4 | P | RIN1 |
| 249939_at | 3.6 | A | 12.4 | A | 5.7 | A | 26.7 | A | --- |
| 249940_at | 4.8 | A | 15.6 | A | 38.7 | A | 41.3 | A | anac090 |
| 249941_at | 603.1 | P | 564.6 | P | 463.9 | P | 450.3 | P | --- |
| 249942_at | 1674.5 | P | 1919.1 | P | 1426.7 | P | 2167.4 | P | GAPB |
| 249943_at | 338.2 | P | 366.4 | P | 286.7 | P | 341.1 | P | --- |
| 249944_at | 376.4 | P | 360 | P | 307.3 | P | 367.8 | P | anac089 |
| 249945_at | 2430.5 | P | 2320.2 | P | 3130.9 | P | 2943.5 | P | --- |
| 249946_at | 38.8 | A | 67.5 | P | 90.9 | P | 77.2 | P | --- |
| 249947_at | 145.1 | P | 169.4 | P | 109.2 | P | 122.8 | P | --- |
| 249948_at | 191.9 | P | 198.9 | P | 237.7 | P | 188.4 | P | --- |
| 249949_at | 25.9 | A | 3.6 | A | 37 | A | 9.4 | A | --- |
| 249950_at | 1.5 | A | 3 | A | 8.8 | A | 1.7 | A | --- |
| 249951_at | 33.4 | A | 46.5 | A | 80.9 | M | 74.1 | A | BUD2 |
| 249952_at | 79.2 | P | 80.9 | P | 122.2 | P | 77.8 | P | --- |
| 249953_at | 305.6 | P | 342 | P | 382.9 | P | 363.3 | P | FRS12 |
| 249954_at | 116.7 | P | 97.6 | P | 78.2 | P | 81.5 | P | --- |
| 249955_at | 60.1 | P | 39.7 | P | 67.9 | A | 60.6 | P | --- |
| 249956_at | 403.1 | P | 547.1 | P | 514.3 | P | 470 | P | --- |
| 249957_at | 969.7 | P | 818.8 | P | 713.3 | P | 839.5 | P | --- |
| 249958_at | 124.2 | P | 187.1 | P | 172.7 | P | 215.5 | P | --- |
| 249959_at | 2063.8 | P | 1856.4 | P | 1968.8 | P | 2042.8 | P | --- |
| 249960_at | 364.5 | P | 455.2 | P | 392.3 | P | 506.5 | P | SPL7 |
| 249961_at | 63.5 | P | 88.2 | P | 96.9 | P | 87.6 | P | --- |
| 249962_at | 5.3 | A | 1.6 | A | 3.2 | A | 1.7 | A | --- |
| 249963_at | 434.6 | P | 340.7 | P | 306.7 | P | 263.1 | P | ATBPM1 |
| 249964_at | 382.5 | P | 668.2 | P | 427.3 | P | 606.9 | P | MPK16 |
| 249965_at | 81 | P | 82.8 | P | 62 | P | 90.3 | P | --- |
| 249966_at | 348 | P | 382.7 | P | 401.1 | P | 379.2 | P | --- |
| 249967_at | 317.2 | P | 351.5 | P | 270.4 | P | 267.6 | P | --- |
| 249968_at | 283.9 | P | 321.8 | P | 318.7 | P | 324.9 | P | --- |
| 249969_at | 366.2 | P | 387.3 | P | 470.5 | P | 392.2 | P | --- |
| 249970_at | 95.3 | A | 23.8 | A | 119.2 | A | 72.1 | A | --- |
| 249971_at | 128.6 | P | 67.6 | A | 110.3 | P | 32.9 | A | --- |
| 249972_at | 2.9 | A | 7.9 | A | 7.7 | A | 3.6 | A | IPT5 |
| 249973_at | 369.6 | P | 336.2 | P | 293.4 | P | 297.3 | P | --- |
| 249974_at | 157.8 | P | 160.1 | P | 108.1 | P | 137.7 | P | --- |
| 249975_s_at | 1251.2 | P | 941 | P | 1361.9 | P | 946 | P | --- |
| 249976_at | 1.6 | A | 3.2 | A | 34 | A | 15.8 | A | SCL28 |
| 249977_at | 277 | P | 131.2 | P | 178.8 | P | 119.7 | P | EMB3007 |
| 249978_at | 316.8 | P | 290 | P | 224.4 | P | 321.6 | P | --- |
| 249979_s_at | 72.4 | P | 34.1 | P | 69.3 | P | 70.5 | P | --- |
| 249980_at | 3.7 | A | 12.3 | A | 3.5 | A | 1.7 | A | --- |
| 249981_at | 2.2 | A | 2.3 | A | 3.6 | A | 2.5 | A | --- |
| 249982_at | 1517.6 | P | 1743.1 | P | 1112.2 | P | 1559.3 | P | --- |
| 249983_at | 507.1 | P | 902.4 | P | 433.1 | P | 620.4 | P | --- |
| 249984_at | 791.2 | P | 580.1 | P | 782.2 | P | 569 | P | --- |
| 249985_at | 801.8 | P | 1051.6 | P | 788.1 | P | 941.8 | P | --- |
| 249986_at | 266.1 | P | 396.6 | P | 327.6 | P | 408.4 | P | --- |
| 249987_at | 329.1 | P | 237.7 | P | 260.2 | P | 230.2 | P | --- |
| 249988_at | 1942.4 | P | 1360.8 | P | 1375.6 | P | 1341.1 | P | --- |
| 249989_at | 224.3 | P | 192 | P | 243.9 | P | 277.6 | P | --- |
| 249990_at | 78.9 | P | 90.4 | P | 103.6 | P | 97.3 | P | --- |
| 249991_at | 124.8 | P | 197.3 | P | 112.2 | P | 174.3 | P | --- |
| 249992_at | 36.9 | A | 22.5 | A | 3.3 | A | 25.3 | A | PUCHI |
| 249993_at | 654.1 | P | 558.3 | P | 564 | P | 678.9 | P | --- |
| 249994_at | 644.4 | P | 635.7 | P | 592.9 | P | 607.2 | P | --- |
| 249995_at | 1.5 | A | 4.4 | A | 2.6 | A | 32.4 | A | --- |
| 249996_at | 291.8 | P | 466.3 | P | 165.1 | P | 246.7 | P | --- |
| 249997_at | 357.5 | P | 438.8 | P | 355.5 | P | 423.7 | P | CHR17 |
| 249998_at | 1094.1 | P | 1189.8 | P | 1224.2 | P | 1257.9 | P | FASS |
| 249999_at | 279.1 | P | 344.3 | P | 261.6 | P | 299.7 | P | --- |
| 250000_at | 822.5 | P | 910.2 | P | 896 | P | 752.1 | P | --- |
| 250001_at | 646.7 | P | 711.8 | P | 696.7 | P | 641.4 | P | AtTLP11 |
| 250002_at | 77.2 | P | 191.8 | P | 163.2 | P | 184.1 | P | AGP25 |
| 250003_at | 23.3 | A | 5.2 | A | 25.3 | A | 3.3 | A | --- |
| 250004_at | 105 | P | 142.4 | P | 168.6 | P | 139.3 | P | --- |
| 250005_at | 287.2 | P | 447.5 | P | 364.9 | P | 408.3 | P | --- |
| 250006_at | 1360.9 | P | 1583.4 | P | 1224.5 | P | 1728.6 | P | PCB2 |
| 250007_at | 104.1 | A | 137.7 | P | 113.8 | M | 120.3 | P | BMY3 |
| 250008_at | 242.1 | P | 261.2 | P | 206.5 | P | 219.6 | P | --- |
| 250009_at | 270.4 | P | 187.5 | P | 324 | P | 265.7 | P | --- |
| 250010_at | 34.1 | A | 20.1 | A | 42.3 | A | 43.4 | A | --- |
| 250011_at | 54.8 | P | 55.5 | P | 94.3 | P | 69.4 | P | --- |
| 250012_x_at | 48.1 | A | 87.5 | P | 74.8 | A | 80.8 | M | --- |
| 250013_at | 283.5 | P | 257 | P | 282.3 | P | 280.3 | P | --- |
| 250014_at | 3587.1 | P | 2935.9 | P | 3148.1 | P | 2795.5 | P | TRP1 |
| 250015_at | 269.3 | P | 200.6 | P | 197.6 | P | 206.2 | P | DRT101 |
| 250016_at | 878.3 | P | 1040.4 | P | 911.9 | P | 929.7 | P | CSD3 |
| 250017_at | 539.8 | P | 704.2 | P | 649.4 | P | 694.4 | P | --- |
| 250018_at | 802.5 | P | 919.8 | P | 536.2 | P | 736.1 | P | --- |
| 250019_at | 8.9 | A | 1.3 | A | 5.5 | A | 7.3 | A | --- |

| | | | | | | | | | |
|---|---|---|---|---|---|---|---|---|---|
| 250020_at | 53 | A | 67.2 | P | 72.7 | A | 36.5 | A | --- |
| 250021_at | 658 | P | 751.6 | P | 744.5 | P | 689.1 | P | --- |
| 250022_at | 110.7 | P | 145.7 | P | 131.6 | P | 155.5 | P | --- |
| 250023_at | 1.5 | A | 26 | A | 20.7 | A | 4.1 | A | --- |
| 250024_at | 140.9 | P | 66.6 | P | 115.9 | P | 60.5 | P | ANAC087 |
| 250025_at | 128.7 | P | 67.2 | A | 192.8 | P | 84.9 | A | SIP1;2 |
| 250026_at | 23.1 | A | 29.1 | P | 57.5 | A | 16.6 | A | --- |
| 250027_at | 428.1 | P | 350.1 | P | 341.2 | P | 366.5 | P | --- |
| 250028_at | 1518.5 | P | 1141.5 | P | 968.2 | P | 1116.3 | P | --- |
| 250029_at | 803.4 | P | 585 | P | 781.7 | P | 630.1 | P | --- |
| 250030_at | 418.4 | P | 403.6 | P | 295 | P | 457.9 | P | ATAPRL7 |
| 250031_at | 142.7 | P | 118.4 | P | 163.5 | P | 160.5 | P | MYR1 |
| 250032_at | 1605.2 | P | 818.7 | P | 1491.3 | P | 810.6 | P | GDH1 |
| 250033_at | 1069.3 | P | 1033.2 | P | 1030.2 | P | 1006.8 | P | NCBP |
| 250034_at | 406.1 | P | 435.6 | P | 481.9 | P | 490.8 | P | ATAPY2 |
| 250035_at | 472.5 | P | 622.6 | P | 529 | P | 542.8 | P | --- |
| 250036_at | 37.2 | P | 28.7 | P | 52.2 | A | 50.8 | P | --- |
| 250037_at | 5 | A | 3.8 | A | 5.2 | A | 3.4 | A | --- |
| 250038_at | 37.4 | M | 71.3 | P | 23.8 | A | 63.3 | P | --- |
| 250039_at | 43.6 | A | 54.9 | P | 32.2 | A | 61.6 | P | --- |
| 250040_at | 219.2 | P | 243.9 | P | 253.8 | P | 238.7 | P | --- |
| 250041_at | 305.1 | P | 378.3 | P | 333.6 | P | 397.6 | P | PIR121 |
| 250042_at | 600.2 | P | 631.1 | P | 590 | P | 718.7 | P | --- |
| 250043_at | 88.1 | P | 121.4 | P | 91.7 | P | 113.6 | P | --- |
| 250044_at | 69.6 | P | 53.1 | P | 79.5 | P | 72.6 | P | --- |
| 250045_at | 1104.2 | P | 370 | P | 1254.3 | P | 682.8 | P | --- |
| 250046_at | 1.9 | A | 3.6 | A | 7 | A | 3.2 | A | --- |
| 250047_at | 29 | A | 29 | A | 65.3 | A | 5.9 | A | --- |
| 250048_s_at | 1.3 | A | 2.4 | A | 3.8 | A | 8.1 | A | --- |
| 250049_at | 109.5 | P | 123 | P | 183.5 | P | 137.8 | P | --- |
| 250050_at | 224.9 | P | 263.5 | P | 306.3 | P | 233.7 | P | VAR3 |
| 250051_at | 48.2 | P | 50.3 | P | 77.8 | M | 54.2 | P | AtMYB56 |
| 250052_at | 24.4 | A | 25.5 | P | 42.6 | A | 25.5 | P | --- |
| 250053_at | 292.9 | P | 339.8 | P | 278 | P | 285.1 | P | --- |
| 250054_at | 2814 | P | 1650.4 | P | 2633.1 | P | 1924 | P | CAX7 |
| 250055_at | 3547.9 | P | 3120 | P | 3777.5 | P | 3170.2 | P | ATCBR |
| 250056_at | 285.9 | P | 350.1 | P | 293.9 | P | 320.4 | P | --- |
| 250057_at | 1584 | P | 1447.7 | P | 1492.1 | P | 1541.7 | P | --- |
| 250058_at | 7986.2 | P | 8466.2 | P | 8301.6 | P | 9083.8 | P | PSRP6 |
| 250059_at | 3.8 | A | 41.5 | A | 9.2 | A | 2.1 | A | --- |
| 250060_at | 195.7 | P | 196.5 | P | 223.2 | P | 220.7 | P | TFL2 |
| 250061_at | 1919.6 | P | 1805.6 | P | 2027.4 | P | 1726.1 | P | EMB1241 |
| 250062_at | 2710.4 | P | 1473.1 | P | 2200.8 | P | 1607 | P | --- |
| 250063_at | 325.2 | P | 280 | P | 346.9 | P | 248.2 | P | CSA1 |
| 250064_at | 263.8 | P | 263 | P | 314.8 | P | 294.6 | P | DAR4 |
| 250065_at | 228.7 | P | 220.6 | P | 168.2 | P | 182.4 | P | --- |
| 250066_at | 237 | P | 241 | P | 269.8 | P | 286.6 | P | --- |
| 250067_at | 19.8 | A | 2.6 | A | 22.8 | A | 10.4 | A | --- |
| 250068_at | 18.5 | A | 22.9 | A | 49.1 | A | 41.6 | A | --- |
| 250069_at | 26.9 | A | 30.2 | A | 34.1 | A | 21.7 | A | --- |
| 250070_at | 139.5 | P | 114.9 | P | 143.3 | P | 121.2 | P | --- |
| 250071_at | 13.3 | A | 1.8 | A | 1.4 | A | 2.2 | A | --- |
| 250072_at | 117.3 | P | 120.5 | P | 181.5 | P | 129.1 | P | --- |
| 250073_at | 4849.2 | P | 2810.6 | P | 4023.9 | P | 3811.9 | P | ENH1 |
| 250074_at | 513 | P | 984.4 | P | 745.4 | P | 1016.9 | P | --- |
| 250075_at | 478.5 | P | 827 | P | 547.7 | P | 941.3 | P | --- |
| 250076_at | 2469.8 | P | 2677.7 | P | 2566 | P | 2350.8 | P | --- |
| 250077_at | 337.5 | P | 415.4 | P | 386.2 | P | 407 | P | --- |
| 250078_at | 68.1 | M | 80 | P | 67.8 | A | 83 | P | ORC3 |
| 250079_at | 686.2 | P | 826.5 | P | 849.3 | P | 823 | P | --- |
| 250080_at | 1441.6 | P | 1676.4 | P | 1832.4 | P | 1612.5 | P | TIC40 |
| 250081_at | 1 | A | 0.4 | A | 2.4 | A | 0.4 | A | --- |
| 250082_at | 6.8 | A | 0.9 | A | 28.6 | A | 21.8 | A | --- |
| 250083_at | 2386.9 | P | 460.4 | P | 2805.2 | P | 1244.3 | P | ATGSTF12 |
| 250084_at | 98.9 | P | 130.1 | P | 174.4 | P | 162.1 | P | SDG40 |
| 250085_at | 184.9 | P | 164.6 | P | 203.3 | P | 180.4 | P | --- |
| 250086_at | 29.1 | A | 20.8 | A | 38.6 | A | 31 | A | anac086 |
| 250087_at | 104.3 | P | 86.4 | P | 125.8 | P | 109 | P | --- |
| 250088_at | 405.2 | P | 442.2 | P | 449.6 | P | 306.6 | P | APG5 |
| 250089_at | 38.9 | A | 18.2 | A | 47.8 | A | 29.3 | A | HDG9 |
| 250090_at | 829.4 | P | 301.7 | P | 732.3 | P | 365.8 | P | GAD |
| 250091_at | 12.8 | A | 9 | A | 16.9 | A | 2.8 | A | --- |
| 250092_at | 25.5 | A | 25.5 | P | 40.6 | P | 42.7 | A | --- |
| 250093_at | 91.4 | P | 76.9 | P | 86.2 | M | 76.4 | P | --- |
| 250094_at | 6734.3 | P | 2948.5 | P | 4105.2 | P | 3604.4 | P | --- |
| 250095_at | 2672.3 | P | 2654.4 | P | 2867.9 | P | 2752.6 | P | --- |
| 250096_at | 1553 | P | 2062.3 | P | 1169.2 | P | 1732.8 | P | --- |
| 250097_at | 1203.9 | P | 933.9 | P | 1154.6 | P | 1138.9 | P | --- |
| 250098_at | 96.3 | P | 105.7 | P | 43.1 | A | 73.5 | P | --- |
| 250099_at | 23.9 | A | 56.4 | P | 42.9 | A | 51.9 | M | --- |
| 250100_at | 48.3 | P | 220.4 | P | 56.7 | A | 247.1 | P | GLN1;4 |
| 250101_at | 7.7 | A | 31.3 | A | 11.2 | A | 8.6 | A | BGLU2 |
| 250102_at | 377.8 | P | 470.1 | P | 463.4 | P | 485.8 | P | LRR1 |
| 250103_at | 39.5 | P | 76.9 | P | 61 | A | 46.8 | P | MYB43 |
| 250104_at | 168.8 | P | 215.7 | P | 186.1 | P | 179.2 | P | --- |
| 250105_at | 310.6 | P | 207.8 | P | 304.1 | P | 236.4 | P | RAD4 |
| 250106_at | 17.9 | A | 41.9 | P | 32.7 | A | 39.2 | P | --- |
| 250107_at | 103.1 | P | 138 | P | 99.6 | M | 166.3 | P | SPX4 |
| 250108_at | 33.9 | P | 61.8 | P | 53.8 | P | 50.8 | P | ATHB-3 |
| 250109_at | 1216.4 | P | 6859 | P | 1190.1 | P | 6137.2 | P | GASA4 |
| 250110_at | 1294.2 | P | 2267.6 | P | 1543.3 | P | 2059.2 | P | --- |
| 250111_at | 1031.1 | P | 927.6 | P | 1136.1 | P | 899.4 | P | --- |
| 250112_at | 393.8 | P | 495.9 | P | 476.9 | P | 503.5 | P | --- |
| 250113_at | 156.3 | P | 162.5 | P | 146.2 | P | 167.6 | P | FRL1 |
| 250114_s_at | 1145.9 | P | 909.4 | P | 907.6 | P | 808.8 | P | AAE5 |
| 250115_at | 147.8 | P | 113.2 | P | 117.2 | P | 122.6 | P | --- |
| 250116_at | 19.2 | A | 3.3 | A | 25.7 | A | 11.2 | A | --- |
| 250117_at | 626.6 | P | 584.1 | P | 690.7 | P | 585.9 | P | IPP1 |
| 250118_at | 6.7 | A | 1.6 | A | 14.8 | A | 1.7 | A | --- |
| 250119_at | 2534 | P | 2478.2 | P | 2921.2 | P | 2468.2 | P | --- |
| 250120_at | 96.3 | P | 124 | P | 146.5 | P | 157.4 | P | RIC4 |
| 250121_at | 3 | A | 0.9 | A | 3.5 | A | 9.8 | A | --- |
| 250122_at | 130.7 | P | 131 | P | 138.9 | P | 128.6 | P | --- |
| 250123_at | 58.2 | P | 103.8 | P | 73.5 | P | 85.7 | P | PIN5 |
| 250124_at | 101.4 | P | 151.5 | P | 184 | P | 128.9 | P | --- |
| 250125_at | 1959.1 | P | 2542.2 | P | 2313.8 | P | 2268 | P | CAC1 |
| 250126_at | 208.1 | P | 283.4 | P | 201.5 | P | 281.4 | P | UCH1 |
| 250127_at | 312.3 | P | 341.7 | P | 301.9 | P | 377.2 | P | --- |
| 250128_at | 305.2 | P | 282.8 | P | 351.2 | P | 308.5 | P | ZFN3 |

| | | | | | | | | | |
|---|---|---|---|---|---|---|---|---|---|
| 250129_at | 1916.7 | P | 935.5 | P | 1370.2 | P | 1052.2 | P | --- |
| 250130_at | 654.2 | P | 891.5 | P | 666.6 | P | 775.3 | P | --- |
| 250131_at | 710.5 | P | 798 | P | 656.8 | P | 819.4 | P | --- |
| 250132_at | 134.9 | P | 117.7 | A | 143.4 | A | 137.9 | M | KAN |
| 250133_at | 778.1 | P | 1345.2 | P | 760.8 | P | 918.3 | P | TRXF2 |
| 250134_at | 564.1 | P | 461 | P | 634.8 | P | 559.4 | P | --- |
| 250135_at | 73.4 | A | 53.4 | A | 146.9 | A | 36.6 | A | --- |
| 250136_at | 23.3 | A | 40.1 | A | 82 | A | 25.2 | A | DRM1 |
| 250137_at | 445.3 | P | 481.7 | P | 347.4 | P | 396.2 | P | --- |
| 250138_at | 123.5 | P | 279.1 | P | 132.2 | P | 255.7 | P | --- |
| 250139_at | 224.7 | P | 237.4 | P | 221.8 | P | 230.2 | P | DRM2 |
| 250140_at | 64.8 | P | 116.5 | P | 135.7 | P | 112.2 | P | DRM2 |
| 250141_at | 214.9 | P | 440.1 | P | 318.3 | P | 478.7 | P | SK13 |
| 250142_at | 9.4 | A | 2.1 | A | 37.1 | A | 33.2 | P | --- |
| 250143_at | 53.3 | P | 49 | A | 52.3 | A | 58.6 | A | ATARFA1B |
| 250144_at | 643.1 | P | 597.3 | P | 672 | P | 576.4 | P | --- |
| 250145_at | 32.5 | A | 61.1 | P | 41.6 | A | 54.3 | P | --- |
| 250146_at | 670 | P | 862.4 | P | 800.6 | P | 974.6 | P | PDF1B |
| 250147_at | 191.9 | P | 230.7 | P | 263.4 | P | 158.9 | P | --- |
| 250148_at | 255.9 | P | 247.8 | P | 304.2 | P | 230.2 | P | --- |
| 250149_at | 219.5 | P | 160.7 | P | 343.5 | P | 227.8 | P | --- |
| 250150_at | 340.2 | P | 314.5 | P | 377.5 | P | 321.8 | P | --- |
| 250151_at | 858.3 | P | 363.7 | P | 759.7 | P | 579 | P | ATNRT2.7 |
| 250152_at | 23.7 | A | 244.3 | P | 104.4 | P | 95.1 | P | --- |
| 250153_at | 5.5 | A | 11.9 | A | 19 | A | 1.7 | A | WRKY72 |
| 250154_at | 32 | A | 20.9 | A | 10.5 | A | 2.4 | A | --- |
| 250155_at | 84.9 | P | 204.3 | P | 96.3 | P | 173.4 | P | --- |
| 250156_at | 177.7 | P | 148.1 | P | 169.4 | P | 147.3 | P | --- |
| 250157_at | 40.8 | P | 43.1 | P | 58.9 | P | 30.3 | A | --- |
| 250158_at | 177.1 | P | 156.6 | P | 161 | P | 159 | P | --- |
| 250159_at | 12260.1 | P | 10636.5 | P | 13507.4 | P | 11366.1 | P | --- |
| 250160_at | 208.7 | P | 372.1 | P | 347.5 | P | 403.6 | P | ATHB30 |
| 250161_at | 140.1 | P | 65.5 | P | 64.1 | A | 55.2 | A | --- |
| 250162_at | 49 | A | 11.3 | A | 37 | A | 5.2 | A | FTSH6 |
| 250163_at | 163.4 | P | 190.1 | P | 137.1 | P | 191.5 | P | --- |
| 250164_at | 154.2 | P | 139.7 | P | 162.7 | P | 168.1 | P | --- |
| 250165_at | 36.3 | P | 30.5 | A | 22.9 | A | 35.8 | A | --- |
| 250166_at | 63.3 | P | 45.6 | P | 54.7 | A | 58 | P | --- |
| 250167_at | 202.1 | P | 140.1 | P | 191.7 | P | 183.2 | P | ATMYB16 |
| 250168_at | 1216.3 | P | 957.3 | P | 1124.2 | P | 848.8 | P | --- |
| 250169_at | 55.1 | P | 67.1 | P | 63.2 | P | 62.6 | P | --- |
| 250170_at | 1579.3 | P | 2275.6 | P | 1681.4 | P | 2003.6 | P | --- |
| 250171_at | 4.4 | A | 5.2 | A | 8.8 | A | 13.8 | A | ATPHB5 |
| 250172_at | 21.4 | A | 34.8 | A | 28.2 | A | 12.2 | A | --- |
| 250173_at | 12.9 | A | 26.9 | A | 1.8 | A | 25.5 | A | AtMYB40 |
| 250174_at | 75.8 | P | 91.8 | P | 47.8 | A | 89.2 | P | AGP6 |
| 250175_at | 337.6 | P | 222.1 | P | 359.4 | P | 259.8 | P | --- |
| 250176_at | 1.4 | A | 1.2 | A | 5.4 | A | 1.1 | A | CYP724A1 |
| 250177_at | 156.8 | P | 227.3 | P | 228.3 | P | 176.3 | P | RGLG2 |
| 250178_at | 1007.5 | P | 1188.7 | P | 1240.7 | P | 1140.5 | P | --- |
| 250179_at | 345.8 | P | 261.5 | P | 393.1 | P | 257.7 | P | --- |
| 250180_at | 451.9 | P | 469.4 | P | 399 | P | 433.1 | P | --- |
| 250181_at | 1073.1 | P | 1215.9 | P | 897.7 | P | 998.2 | P | --- |
| 250182_at | 22.4 | A | 26.3 | A | 7.9 | A | 5.3 | A | --- |
| 250183_at | 48.7 | P | 64 | P | 70.4 | P | 41.5 | P | --- |
| 250184_at | 606.2 | P | 750.1 | P | 837 | P | 657.2 | P | --- |
| 250185_at | 334 | P | 320.2 | P | 303 | P | 296.8 | P | AtCXE16 |
| 250186_at | 249.5 | P | 311.6 | P | 288.5 | P | 326.1 | P | --- |
| 250187_at | 296.5 | P | 324.8 | P | 397 | P | 258.4 | P | --- |
| 250188_at | 925.2 | P | 866.3 | P | 918.9 | P | 952.3 | P | COP13 |
| 250189_at | 208.2 | P | 213.9 | P | 205.8 | P | 214 | P | --- |
| 250190_at | 6526 | P | 7352.7 | P | 5875.2 | P | 6965.8 | P | --- |
| 250191_at | 538.3 | P | 517.4 | P | 482.8 | P | 501.3 | P | ATBET9 |
| 250192_at | 2300.5 | P | 1828.5 | P | 2270.3 | P | 2005.8 | P | --- |
| 250193_at | 148.9 | P | 140.6 | P | 186.5 | P | 179.4 | P | --- |
| 250194_at | 310.8 | P | 354.2 | P | 285.3 | P | 330.4 | P | --- |
| 250195_at | 1.3 | A | 1.5 | A | 10.8 | A | 1.3 | A | --- |
| 250196_at | 204.2 | P | 154.4 | P | 277.4 | P | 162.5 | P | --- |
| 250197_at | 1759.3 | P | 1566.2 | P | 2053.7 | P | 1479.5 | P | --- |
| 250198_at | 168.9 | P | 245.2 | P | 253.1 | P | 278.5 | P | NAP14 |
| 250199_at | 129.4 | P | 7.5 | A | 9.1 | A | 4.7 | A | MPL1 |
| 250200_at | 11.9 | A | 13.9 | A | 8 | A | 3.9 | A | --- |
| 250201_at | 93.7 | P | 222.7 | P | 115.6 | P | 159.9 | P | --- |
| 250202_at | 373.7 | P | 401.8 | P | 412 | P | 392 | P | --- |
| 250203_at | 897.8 | P | 1104.9 | P | 855.1 | P | 991.9 | P | --- |
| 250204_at | 3.4 | A | 4.4 | A | 10.5 | A | 29.4 | A | ATEXO70C2 |
| 250205_at | 30.5 | A | 4.4 | A | 9.9 | A | 14.8 | A | --- |
| 250206_at | 5373.3 | P | 5464.5 | P | 6583.5 | P | 5408 | P | --- |
| 250207_at | 5624.6 | P | 3182.4 | P | 8249.5 | P | 5943.2 | P | TT4 |
| 250208_at | 87.9 | A | 53.4 | A | 84.2 | A | 29.2 | A | anac084 |
| 250209_at | 3756.9 | P | 4154 | P | 3292.5 | P | 4030.7 | P | --- |
| 250210_at | 484.9 | P | 530 | P | 548.5 | P | 466.2 | P | --- |
| 250211_at | 42.8 | A | 79.2 | P | 52 | P | 78.6 | P | --- |
| 250212_at | 278.8 | P | 220.8 | P | 222 | P | 217 | P | SUVH4 |
| 250213_at | 165.8 | P | 92.4 | M | 142.7 | P | 107.6 | P | TBP1 |
| 250214_at | 54 | P | 80 | P | 102.3 | P | 95.8 | P | EXGT-A4 |
| 250215_at | 176.5 | P | 176.5 | P | 136.8 | P | 180.7 | P | --- |
| 250216_at | 72.1 | P | 154.8 | P | 214.8 | P | 113.1 | P | --- |
| 250217_at | 2135.2 | P | 1798.5 | P | 2180.6 | P | 2152.8 | P | --- |
| 250218_at | 378.2 | P | 406.7 | P | 464.2 | P | 321.4 | P | CHC1 |
| 250219_at | 65.6 | P | 25.1 | A | 38.8 | A | 18.2 | A | MPL1 |
| 250220_at | 281.3 | P | 359 | P | 451 | P | 493.7 | P | --- |
| 250221_at | 487.3 | P | 535.2 | P | 630.4 | P | 459.4 | P | HEMG2 |
| 250222_at | 758.8 | P | 565 | P | 813.6 | P | 612.1 | P | --- |
| 250223_at | 5.4 | A | 5.7 | A | 18.2 | A | 31.8 | A | ROXY2 |
| 250224_at | 8 | A | 3.5 | A | 4.7 | A | 4.4 | A | --- |
| 250225_at | 509 | P | 459.6 | P | 485 | P | 415.5 | P | --- |
| 250226_at | 1587.4 | P | 1345.3 | P | 1879.5 | P | 1493.4 | P | --- |
| 250227_at | 202.5 | P | 135.3 | A | 261.6 | P | 94.4 | P | --- |
| 250228_at | 149.5 | P | 252.5 | P | 213.2 | P | 313.5 | P | FZR3 |
| 250229_at | 1194.6 | P | 1558.3 | P | 1061 | P | 1207 | P | NACA3 |
| 250230_at | 2.5 | A | 49.5 | A | 12.3 | A | 24.9 | A | --- |
| 250231_at | 120.2 | M | 121.1 | P | 125.4 | P | 102.9 | P | LEP |
| 250232_at | 333.4 | P | 416.3 | P | 263.6 | P | 304.6 | P | --- |
| 250233_at | 180.1 | P | 277.5 | P | 188.2 | P | 262.7 | P | IQD11 |
| 250234_at | 2032.4 | P | 1489.7 | P | 1585.6 | P | 1464.7 | P | --- |
| 250235_s_at | 4455 | P | 3937.6 | P | 4913.6 | P | 4365.6 | P | --- |
| 250236_at | 5233.8 | P | 5051.7 | P | 5691.9 | P | 4865.5 | P | --- |
| 250237_at | 502.9 | P | 583.6 | P | 516.7 | P | 526.1 | P | --- |

| | | | | | | | | | |
|---|---|---|---|---|---|---|---|---|---|
| 250238_at | 252.8 | P | 300.2 | P | 253 | P | 245.6 | P | DCP2 |
| 250239_at | 26.2 | P | 57.2 | P | 97.5 | P | 81.1 | P | --- |
| 250240_at | 93.9 | P | 101.2 | P | 78.5 | P | 58.3 | P | --- |
| 250241_at | 48.2 | A | 44.8 | A | 67.4 | A | 29.7 | A | --- |
| 250242_at | 22 | A | 15.3 | A | 8.4 | A | 1.1 | A | --- |
| 250243_at | 6970.3 | P | 5382.3 | P | 4987.4 | P | 5646.6 | P | GUN5 |
| 250244_at | 264.9 | P | 235.5 | P | 249.9 | P | 181.9 | P | ABO1 |
| 250245_at | 277.1 | P | 351 | P | 292 | P | 318.1 | P | --- |
| 250246_at | 24.3 | A | 9.7 | A | 10.9 | A | 35.8 | A | ATPAO1 |
| 250247_at | 835.5 | P | 796.8 | P | 853.8 | P | 738.7 | P | --- |
| 250248_at | 806.1 | P | 2001 | P | 2156.6 | P | 2001.2 | P | ZIF1 |
| 250249_at | 261.9 | P | 259.7 | P | 264.2 | P | 244.5 | P | --- |
| 250250_at | 387.3 | P | 336.3 | P | 412.1 | P | 343.7 | P | --- |
| 250251_at | 15.7 | A | 6.1 | A | 34.9 | A | 38.8 | A | --- |
| 250252_at | 2868.1 | P | 1365.5 | P | 2525.6 | P | 1729.8 | P | ZIFL1 |
| 250253_at | 697 | P | 816.9 | P | 641.1 | P | 824.4 | P | PDAT |
| 250254_at | 2779 | P | 2532.7 | P | 2748.5 | P | 2685.8 | P | SMT1 |
| 250255_at | 1152.6 | P | 1239.3 | P | 1186.1 | P | 1034.9 | P | SIG4 |
| 250256_at | 2934.1 | P | 3304.2 | P | 3716.9 | P | 3344.9 | P | --- |
| 250257_at | 1228.9 | P | 993.6 | P | 1236.4 | P | 1140.7 | P | --- |
| 250258_at | 15.3 | A | 36.6 | A | 18.4 | A | 22.3 | A | AGL15 |
| 250259_at | 1694.9 | P | 1151.7 | P | 1396.6 | P | 1275.6 | P | --- |
| 250260_at | 937.1 | P | 1005.8 | P | 769.6 | P | 920.3 | P | --- |
| 250261_at | 296.4 | P | 294 | P | 382.3 | P | 330.5 | P | --- |
| 250262_at | 1007.3 | P | 970.1 | P | 906.2 | P | 1016.7 | P | --- |
| 250263_at | 204.7 | P | 186.6 | P | 205.6 | P | 197.3 | P | --- |
| 250264_at | 960.7 | P | 551.2 | P | 662.7 | P | 544.8 | P | --- |
| 250265_at | 135.6 | P | 157.9 | P | 160.8 | P | 130.3 | P | --- |
| 250266_at | 2.4 | A | 11.5 | A | 5 | A | 22.7 | A | --- |
| 250267_at | 562.9 | P | 131.9 | P | 380.3 | P | 168.3 | P | --- |
| 250268_s_at | 706.4 | P | 692.9 | P | 771.7 | P | 608.4 | P | --- |
| 250269_at | 7.7 | A | 32.1 | A | 27.7 | A | 29.1 | P | --- |
| 250270_at | 260.2 | P | 182.5 | P | 252.1 | P | 216.9 | P | --- |
| 250271_at | 18.4 | P | 8.1 | A | 30.1 | P | 21.2 | P | CLE40 |
| 250272_at | 1526.7 | P | 1751.4 | P | 1521.9 | P | 1870.1 | P | ATGSL12 |
| 250273_at | 683.9 | P | 589.3 | P | 647.4 | P | 559.8 | P | EMB3011 |
| 250274_at | 540.1 | P | 501.2 | P | 472.6 | P | 529.3 | P | --- |
| 250275_at | 1255.4 | P | 1628 | P | 1401.8 | P | 1558.2 | P | --- |
| 250276_at | 2018.9 | P | 2311.4 | P | 1892.8 | P | 2025.6 | P | --- |
| 250277_at | 309.6 | P | 205.9 | P | 327 | P | 178.9 | P | --- |
| 250278_at | 9054.2 | P | 8060.2 | P | 8850.3 | P | 8364.2 | P | DiT1 |
| 250279_at | 207.3 | P | 278.6 | P | 214 | P | 251.2 | P | --- |
| 250280_at | 7.7 | A | 9.8 | A | 33.9 | A | 8.2 | A | --- |
| 250281_at | 195.9 | P | 462.7 | P | 230.1 | P | 448.8 | P | --- |
| 250282_at | 268.3 | P | 286.7 | P | 331.6 | P | 295.4 | P | --- |
| 250283_at | 173.2 | P | 142.3 | P | 190.9 | P | 164.5 | P | --- |
| 250284_at | 114.1 | P | 165.1 | P | 184 | P | 169.2 | P | CRN |
| 250285_at | 469.6 | P | 567.9 | P | 506.5 | P | 567.7 | P | SFC |
| 250286_at | 210.6 | P | 297 | P | 148.8 | P | 154.8 | P | PBS3 |
| 250287_at | 804.3 | P | 455.5 | P | 556.7 | P | 340.3 | P | Rap2.6L |
| 250288_at | 46.7 | A | 51.9 | A | 11.4 | A | 51.9 | A | --- |
| 250289_at | 1868.7 | P | 1923.5 | P | 1599.2 | P | 1611 | P | --- |
| 250290_at | 220.8 | P | 249.1 | P | 229.6 | P | 228.9 | P | --- |
| 250291_at | 1554.8 | P | 990.4 | P | 1366.4 | P | 1200 | P | AK-LYS1 |
| 250292_at | 834.6 | P | 234.4 | P | 584.2 | P | 224.7 | P | JAZ10 |
| 250293_s_at | 1763.7 | P | 688.9 | P | 1370.3 | P | 785 | P | --- |
| 250294_at | 56.1 | P | 39.6 | P | 48.7 | M | 39.4 | A | --- |
| 250295_at | 784 | P | 868.8 | P | 864.6 | P | 921.1 | P | NEF1 |
| 250296_at | 24.5 | A | 50.4 | P | 28.5 | A | 50 | P | HSP17.6II |
| 250297_at | 511.5 | P | 616.9 | P | 672.5 | P | 595.9 | P | --- |
| 250298_at | 613.7 | P | 682.1 | P | 744.3 | P | 799 | P | --- |
| 250299_at | 552.7 | P | 645.3 | P | 675.9 | P | 576.6 | P | --- |
| 250300_at | 518.6 | P | 484.3 | P | 547.1 | P | 469.7 | P | --- |
| 250301_at | 424.2 | P | 396 | P | 355.5 | P | 435.4 | P | --- |
| 250302_at | 27.1 | A | 78.7 | P | 43 | P | 45 | A | AtcwINV6 |
| 250303_at | 86.9 | P | 59.3 | P | 77.7 | P | 73.7 | P | --- |
| 250304_at | 753.1 | P | 778.5 | P | 1038.1 | P | 879 | P | --- |
| 250305_at | 1312.4 | P | 1496.3 | P | 1393.5 | P | 1212.8 | P | --- |
| 250306_at | 440.5 | P | 380 | P | 494.1 | P | 436.7 | P | --- |
| 250307_at | 1357.5 | P | 1167.1 | P | 1808.9 | P | 1249.9 | P | --- |
| 250308_at | 20.6 | A | 3.5 | A | 1.4 | A | 24.1 | A | CPK17 |
| 250309_at | 165.9 | P | 156 | P | 190.2 | P | 147.7 | P | --- |
| 250310_at | 346.9 | P | 309.9 | P | 342.7 | P | 363.5 | P | --- |
| 250311_at | 37.9 | A | 38.7 | M | 67 | P | 45.6 | P | --- |
| 250312_at | 175.7 | P | 228.6 | P | 186.6 | P | 233.6 | P | --- |
| 250313_at | 1192 | P | 1149.6 | P | 1143.9 | P | 1247 | P | --- |
| 250314_at | 702.6 | P | 701.4 | P | 601.6 | P | 781.1 | P | --- |
| 250315_at | 611.5 | P | 851.5 | P | 818.6 | P | 756.8 | P | PDE149 |
| 250316_at | 3222.6 | P | 3580.6 | P | 2542.4 | P | 3306.1 | P | ATCYS1 |
| 250317_at | 1409.6 | P | 2248 | P | 1611.9 | P | 2191.8 | P | TUB6 |
| 250318_at | 684.6 | P | 929.7 | P | 787.1 | P | 777.6 | P | --- |
| 250319_at | 1063.9 | P | 1004.3 | P | 1097.1 | P | 958.2 | P | --- |
| 250320_at | 304 | P | 341.2 | P | 275.9 | P | 341.2 | P | NF-YA1 |
| 250321_at | 285.6 | P | 406.9 | P | 295.6 | P | 366.7 | P | --- |
| 250322_at | 45.9 | A | 73 | A | 49.5 | A | 39.3 | A | MYB46 |
| 250323_at | 113.2 | P | 58.9 | P | 101.8 | P | 76.5 | A | --- |
| 250324_at | 1833.4 | P | 1651.7 | P | 1291.2 | P | 1679 | P | --- |
| 250325_s_at | 4.4 | A | 19.1 | A | 5.3 | A | 22.7 | A | --- |
| 250326_at | 376.4 | P | 321.7 | P | 401.5 | P | 357.1 | P | MSL10 |
| 250327_at | 93 | P | 194.2 | P | 115.8 | P | 128.1 | P | --- |
| 250328_at | 55.2 | P | 55.6 | P | 55.8 | P | 57.1 | P | --- |
| 250329_at | 255.7 | P | 259 | P | 263 | P | 242.2 | P | KEA6 |
| 250330_at | 941.3 | P | 879.2 | P | 887.2 | P | 760.4 | P | --- |
| 250331_at | 10.9 | A | 1.9 | A | 13 | A | 5.2 | A | --- |
| 250332_at | 796.1 | P | 990.7 | P | 933.1 | P | 785.8 | P | --- |
| 250333_at | 353.1 | P | 388.4 | P | 421.8 | P | 397.2 | P | TIM17-3 |
| 250334_at | 6586.6 | P | 5450.7 | P | 7348.7 | P | 5945.4 | P | --- |
| 250335_at | 261.2 | P | 250.3 | P | 219.4 | P | 160.8 | P | --- |
| 250336_at | 949.5 | P | 807.9 | P | 886.4 | P | 729.3 | P | --- |
| 250337_at | 445.3 | P | 789.1 | P | 513.9 | P | 659.3 | P | --- |
| 250338_at | 864.6 | P | 912.9 | P | 986.2 | P | 807.4 | P | --- |
| 250339_at | 7108.9 | P | 6141.9 | P | 5618.9 | P | 5836.1 | P | ATNADP-ME2 |
| 250340_at | 821.6 | P | 834.4 | P | 653.8 | P | 660.8 | P | --- |
| 250341_at | 296.3 | P | 275.6 | P | 278.7 | P | 230.7 | P | --- |
| 250342_at | 21.7 | A | 37.2 | A | 43.1 | A | 28.8 | A | --- |
| 250343_at | 748 | P | 670.6 | P | 678.6 | P | 676.5 | P | --- |
| 250344_at | 219.6 | P | 236.1 | P | 308.1 | P | 252.5 | P | --- |
| 250345_at | 4.3 | A | 24.9 | A | 40.5 | A | 3.9 | A | --- |
| 250346_at | 1352.2 | P | 1166.2 | P | 1433.5 | P | 1226.5 | P | --- |

| | | | | | | | | | |
|---|---|---|---|---|---|---|---|---|---|
| 250347_at | 518.4 | P | 510.6 | P | 564 | P | 546.8 | P | --- |
| 250348_at | 29.6 | A | 37.4 | A | 33.9 | A | 33 | A | --- |
| 250349_at | 36.2 | A | 36.5 | A | 40.4 | A | 45.7 | A | --- |
| 250350_at | 555.5 | P | 537.3 | P | 420.4 | P | 538.3 | P | --- |
| 250351_at | 38.8 | P | 126.5 | P | 24.1 | A | 92.9 | P | AT-HSP17.6A |
| 250352_at | 710.1 | P | 750.5 | P | 729 | P | 807.4 | P | --- |
| 250353_at | 548.8 | P | 580.6 | P | 667.5 | P | 563.2 | P | --- |
| 250354_at | 405.3 | P | 548.5 | P | 341.9 | P | 489.1 | P | --- |
| 250355_at | 673.5 | P | 641.9 | P | 655.9 | P | 593.5 | P | --- |
| 250356_at | 863.2 | P | 996.4 | P | 897.2 | P | 874.5 | P | --- |
| 250357_at | 331.5 | P | 378.1 | P | 349 | P | 373.3 | P | --- |
| 250358_at | 8872.3 | P | 10493.6 | P | 9870.6 | P | 10977.8 | P | AGP15 |
| 250359_at | 232.2 | P | 277.4 | P | 270.2 | P | 266.1 | P | --- |
| 250360_at | 3.7 | A | 49 | A | 30.2 | A | 47.3 | A | --- |
| 250361_at | 28.9 | A | 51.5 | P | 55.5 | A | 14.9 | A | --- |
| 250362_at | 193.8 | P | 265.7 | P | 232 | P | 236.3 | P | DXPS3 |
| 250363_at | 351.3 | P | 226.6 | P | 322.6 | P | 248.1 | P | --- |
| 250364_at | 4.4 | A | 2.6 | A | 5.2 | A | 1.9 | A | --- |
| 250365_at | 275.9 | P | 218.9 | P | 201.6 | P | 230 | P | --- |
| 250366_at | 798.7 | P | 2273.1 | P | 945.4 | P | 1972.7 | P | --- |
| 250367_s_at | 3420.7 | P | 2902.3 | P | 3574.3 | P | 2894.3 | P | --- |
| 250368_at | 822.8 | P | 972.4 | P | 935.3 | P | 885.3 | P | --- |
| 250369_at | 63.7 | P | 92.5 | P | 96.4 | P | 87.4 | P | CYC3B |
| 250370_at | 135.5 | P | 175.8 | P | 130.2 | P | 168.5 | P | --- |
| 250371_at | 2718.5 | P | 3319.4 | P | 2637.9 | P | 2979.6 | P | --- |
| 250372_at | 163.8 | P | 160.4 | P | 146.7 | P | 200.3 | P | --- |
| 250373_at | 24.2 | A | 54.7 | P | 34 | A | 41.2 | P | --- |
| 250374_at | 141.2 | P | 123.5 | M | 166.6 | M | 141.8 | P | EMF1 |
| 250375_at | 13.7 | A | 36.1 | A | 15.9 | A | 13.7 | A | --- |
| 250376_at | 181.2 | P | 185.1 | P | 163.4 | P | 191.5 | P | --- |
| 250377_at | 1610.3 | P | 2153.7 | P | 1677.6 | P | 2013 | P | --- |
| 250378_at | 4.3 | A | 23.3 | A | 7 | A | 9.6 | A | --- |
| 250379_at | 45.2 | P | 76.8 | P | 65.2 | P | 86 | P | TINY2 |
| 250380_at | 196.5 | P | 150.4 | P | 121.3 | P | 112.1 | P | --- |
| 250381_at | 60.6 | P | 103.9 | P | 47.1 | P | 107.3 | P | --- |
| 250382_at | 922.1 | P | 1191.3 | P | 884.6 | P | 917.4 | P | --- |
| 250383_at | 819.2 | P | 1278 | P | 1096 | P | 1200.5 | P | --- |
| 250384_at | 2814.5 | P | 2756.7 | P | 2463.3 | P | 3099.6 | P | --- |
| 250385_at | 4878 | P | 2697.5 | P | 3786.4 | P | 2825.4 | P | ASP3 |
| 250386_at | 74.5 | A | 154.7 | P | 88 | A | 173.4 | P | MYB3R-4 |
| 250387_at | 4 | A | 13.3 | A | 29.9 | A | 17.8 | A | --- |
| 250388_at | 91.6 | P | 83.4 | P | 92.3 | P | 68.5 | P | --- |
| 250389_at | 29.4 | A | 30.4 | A | 67.2 | A | 41.3 | A | YUC4 |
| 250390_at | 890.2 | P | 1048.3 | P | 786.6 | P | 1067.4 | P | --- |
| 250391_at | 49.1 | A | 45.5 | A | 77.9 | A | 49 | A | --- |
| 250392_at | 17.1 | A | 14 | A | 32.1 | A | 6.5 | A | --- |
| 250393_at | 58.3 | P | 78.6 | P | 95.3 | P | 78.5 | P | --- |
| 250394_at | 334 | P | 497.1 | P | 359.6 | P | 491 | P | --- |
| 250395_at | 182.8 | P | 189.2 | P | 165.6 | P | 187.5 | P | --- |
| 250396_at | 74.8 | P | 47.1 | A | 75.3 | P | 66.3 | A | --- |
| 250397_at | 4566 | P | 4366 | P | 3927.5 | P | 4229 | P | --- |
| 250398_at | 438.5 | P | 400.9 | P | 444.2 | P | 427 | P | --- |
| 250399_at | 391.3 | P | 397.8 | P | 360.4 | P | 390.7 | P | --- |
| 250400_at | 179.6 | P | 264.3 | P | 142.2 | P | 207.7 | P | --- |
| 250401_at | 2024.3 | P | 2293.5 | P | 2166.8 | P | 2082 | P | --- |
| 250402_at | 481.3 | P | 349.2 | P | 387.7 | P | 387.1 | P | --- |
| 250403_at | 1152.1 | P | 1018.2 | P | 1263.4 | P | 1062.3 | P | --- |
| 250404_at | 2596.3 | P | 3211.2 | P | 2589.9 | P | 3051.3 | P | --- |
| 250405_at | 301.2 | P | 275.9 | P | 229.2 | P | 311.4 | P | UBP22 |
| 250406_at | 617.7 | P | 679.8 | P | 597.4 | P | 743.6 | P | ER |
| 250407_at | 285.8 | P | 341.4 | P | 296.8 | P | 322.9 | P | ATCM2 |
| 250408_at | 140.5 | P | 119.1 | P | 113.6 | P | 136 | P | CIPK5 |
| 250409_at | 4302.4 | P | 5121.2 | P | 4850.9 | P | 4973 | P | --- |
| 250410_at | 297.9 | P | 273 | P | 293.9 | P | 278.6 | P | --- |
| 250411_at | 29.7 | A | 16.6 | A | 21.1 | A | 24.3 | P | --- |
| 250412_at | 499.6 | P | 772.1 | P | 420.9 | P | 682.7 | P | ATVAMP713 |
| 250413_at | 216.2 | P | 185.4 | P | 217.7 | P | 223.1 | P | APT5 |
| 250414_at | 56.8 | A | 60.4 | A | 67.5 | A | 54.5 | A | GLR2.6 |
| 250415_at | 15.9 | A | 11.8 | A | 5.4 | A | 20.8 | A | GLR2.5 |
| 250416_at | 1.8 | A | 1.7 | A | 55.5 | A | 9.8 | A | --- |
| 250417_at | 378.2 | P | 190.8 | P | 199.7 | P | 138.9 | P | --- |
| 250418_at | 368.7 | P | 358 | P | 519 | P | 345.2 | P | --- |
| 250419_at | 111.7 | P | 216.3 | P | 148.3 | P | 164.6 | P | --- |
| 250420_at | 290.6 | P | 286.4 | P | 258.5 | P | 312.7 | P | HY5 |
| 250421_at | 525.2 | P | 559.3 | P | 534.2 | P | 521.4 | P | OCP3 |
| 250422_at | 1384.9 | P | 1512.8 | P | 1262.5 | P | 1571.9 | P | --- |
| 250423_s_at | 282.9 | P | 261.7 | P | 299.8 | P | 284 | P | CYP81K2 |
| 250424_at | 1145.5 | P | 1511.1 | P | 1088.4 | P | 1271.1 | P | GTE2 |
| 250425_at | 15.8 | A | 47 | A | 9.6 | A | 47.1 | A | --- |
| 250426_at | 62.7 | P | 82 | P | 81.3 | P | 74.1 | P | AIL6 |
| 250427_at | 2.1 | A | 4.1 | A | 19.5 | A | 1.1 | A | --- |
| 250428_at | 1319.2 | P | 1673.5 | P | 1438.1 | P | 1725.8 | P | PAS2 |
| 250429_at | 621.7 | P | 886.2 | P | 689.4 | P | 905.2 | P | --- |
| 250430_at | 628.6 | P | 712 | P | 620.1 | P | 720.5 | P | --- |
| 250431_at | 66 | A | 55.9 | A | 34.7 | A | 31.8 | A | CYCD4;2 |
| 250432_at | 22.3 | A | 18.6 | A | 13.4 | A | 32.1 | A | --- |
| 250433_at | 505.3 | P | 806.4 | P | 620.5 | P | 770.1 | P | --- |
| 250434_at | 826.2 | P | 921.5 | P | 817.3 | P | 977.5 | P | --- |
| 250435_at | 2559.3 | P | 2992.1 | P | 2502.5 | P | 2257.6 | P | RING1 |
| 250436_at | 976.5 | P | 1037.9 | P | 1098.2 | P | 1252.5 | P | --- |
| 250437_at | 119.5 | P | 235.5 | P | 87.3 | P | 224.8 | P | AGP4 |
| 250438_at | 57.1 | A | 62.4 | M | 66.3 | A | 37.5 | A | --- |
| 250439_at | 5077.9 | P | 6032.6 | P | 5245.4 | P | 5895 | P | GRF6 |
| 250440_at | 7739.9 | P | 6458.2 | P | 7057.5 | P | 6222.5 | P | EMB3010 |
| 250441_at | 1007.2 | P | 872.6 | P | 1181.8 | P | 858 | P | --- |
| 250442_at | 549.5 | P | 539.5 | P | 507.9 | P | 533.2 | P | MSL2 |
| 250443_at | 44.8 | A | 59.3 | A | 78 | A | 37.2 | A | RBK1 |
| 250444_at | 604.4 | P | 930 | P | 775.7 | P | 950.4 | P | --- |
| 250445_at | 613.4 | P | 3662.1 | P | 464.4 | P | 2775.2 | P | --- |
| 250446_at | 95.9 | P | 235.4 | P | 144.4 | P | 220.9 | P | --- |
| 250447_at | 28.6 | A | 40.7 | A | 32.4 | A | 37.5 | A | --- |
| 250448_at | 316 | P | 216.8 | P | 356.2 | P | 233 | P | --- |
| 250449_at | 250.7 | P | 207.3 | P | 283 | P | 262 | P | --- |
| 250450_at | 32.3 | P | 54.8 | P | 66.5 | P | 46.5 | P | ATMYB92 |
| 250451_at | 218 | P | 218.8 | P | 216 | P | 231.4 | P | CDKC;1 |
| 250452_at | 296.6 | P | 335.9 | P | 343.9 | P | 366.7 | P | --- |
| 250453_at | 193.7 | P | 177.7 | P | 208.4 | P | 198.4 | P | --- |
| 250454_at | 503.9 | P | 528 | P | 611.7 | P | 426.5 | P | --- |
| 250455_at | 100.7 | P | 48.3 | P | 99.4 | P | 25.9 | M | PROPEP4 |

| | | | | | | | | | |
|---|---|---|---|---|---|---|---|---|---|
| 250456_at | 1750.5 | P | 1933.7 | P | 1857.9 | P | 2095.2 | P | EMB2107 |
| 250457_at | 764.6 | P | 772.3 | P | 667.8 | P | 722.4 | P | --- |
| 250458_s_at | 8187.6 | P | 9344.4 | P | 7647.2 | P | 9738.7 | P | ACT7 |
| 250459_at | 397.8 | P | 318.1 | P | 400.3 | P | 349.6 | P | --- |
| 250460_at | 848.5 | P | 891.3 | P | 889.5 | P | 818.2 | P | --- |
| 250461_at | 493.9 | P | 551.1 | P | 753.4 | P | 625.1 | P | --- |
| 250462_at | 258 | P | 312 | P | 257.5 | P | 329.5 | P | --- |
| 250463_at | 158.8 | P | 230.9 | P | 163.8 | P | 177.5 | P | TGA4 |
| 250464_at | 4.5 | A | 14.2 | A | 29.8 | A | 13.8 | A | --- |
| 250465_at | 634.8 | P | 417.8 | P | 611.1 | P | 507 | P | --- |
| 250466_at | 11.8 | A | 4.8 | A | 2.6 | A | 1.8 | A | --- |
| 250467_at | 1.2 | A | 3.7 | A | 1.9 | A | 6.8 | A | --- |
| 250468_at | 19.6 | A | 11.3 | A | 13.5 | A | 1.3 | A | --- |
| 250469_at | 5 | A | 6.7 | A | 4 | A | 2.5 | A | --- |
| 250470_at | 1774 | P | 1728 | P | 1809.8 | P | 1707 | P | --- |
| 250471_at | 164.4 | P | 136.6 | P | 151.1 | P | 199.4 | P | MIPS3 |
| 250472_at | 2.3 | A | 3.9 | A | 3.5 | A | 9.7 | A | --- |
| 250473_at | 10.2 | A | 0.9 | A | 23 | A | 2.4 | A | ANN6 |
| 250474_at | 0.6 | A | 3.1 | A | 28.9 | A | 22.9 | A | ANNAT7 |
| 250475_at | 1253.4 | P | 1232.5 | P | 1233.2 | P | 1072.5 | P | SULTR2;1 |
| 250476_at | 31.3 | A | 83.3 | A | 102.7 | A | 73.6 | A | FLC |
| 250477_at | 320.6 | P | 359.7 | P | 386.9 | P | 304.5 | P | --- |
| 250478_at | 24.9 | A | 39.3 | A | 35.7 | A | 4 | A | DOT3 |
| 250479_at | 31.7 | A | 51.3 | P | 25.2 | A | 74.7 | M | AtRABH1e |
| 250480_at | 532.2 | P | 459 | P | 406.2 | P | 475.7 | P | --- |
| 250481_at | 52.5 | A | 67.9 | P | 81.5 | P | 77.1 | P | --- |
| 250482_at | 128.4 | P | 136.7 | P | 170.3 | P | 131.9 | P | --- |
| 250483_at | 468.7 | P | 218.6 | P | 459.7 | P | 189.4 | P | MES5 |
| 250484_at | 541.8 | P | 580.7 | P | 624.4 | P | 652.8 | P | ASN3 |
| 250485_at | 53.9 | P | 66.6 | P | 55.2 | P | 70.9 | P | PROPEP5 |
| 250486_at | 316 | P | 350.7 | P | 320 | P | 387.7 | P | --- |
| 250487_at | 88.6 | P | 97.8 | P | 100.7 | P | 60.9 | P | --- |
| 250488_at | 2 | A | 1.5 | A | 2.2 | A | 1.9 | A | --- |
| 250489_s_at | 31.5 | A | 22.1 | A | 23.5 | A | 51.1 | P | --- |
| 250490_at | 389 | P | 409.4 | P | 564.8 | P | 480.9 | P | --- |
| 250491_at | 31.6 | A | 27.5 | A | 45 | A | 42 | A | --- |
| 250492_at | 132.4 | P | 137.3 | P | 122.8 | M | 113.2 | P | ATXR5 |
| 250493_at | 50.8 | P | 29.2 | A | 34.2 | A | 23.6 | A | --- |
| 250494_at | 330.3 | P | 329.4 | P | 273.6 | P | 277.7 | P | HAM2 |
| 250495_at | 727.6 | P | 644.9 | P | 812.4 | P | 705.6 | P | --- |
| 250496_at | 5817.4 | P | 5082.8 | P | 6422.5 | P | 6094.3 | P | AtPPa6 |
| 250497_at | 139.7 | P | 128.3 | P | 140.5 | P | 129.1 | P | --- |
| 250498_at | 6888.8 | P | 7185.6 | P | 7048.1 | P | 6911.1 | P | PMDH2 |
| 250499_at | 14.8 | A | 1.3 | A | 5.9 | A | 17.6 | A | BXL3 |
| 250500_at | 4.9 | A | 132.6 | P | 136.2 | P | 177.6 | P | --- |
| 250501_at | 5.6 | A | 28 | A | 5.7 | A | 9.7 | A | SCPL19 |
| 250502_at | 1924.8 | P | 1409.7 | P | 2119.2 | P | 1867.6 | P | MTHSC70-2 |
| 250503_at | 537.2 | P | 644.4 | P | 559.8 | P | 719.3 | P | --- |
| 250504_at | 86.5 | P | 64.9 | P | 95.6 | P | 94.1 | P | --- |
| 250505_at | 1084.6 | P | 1313.2 | P | 1226.7 | P | 1495.2 | P | CESA5 |
| 250506_at | 2.6 | A | 15.9 | A | 7.6 | A | 2.2 | A | GCN2 |
| 250507_at | 3.4 | A | 1.4 | A | 5.4 | A | 27.4 | A | --- |
| 250508_at | 34.9 | A | 23.5 | A | 31.8 | A | 35.8 | A | --- |
| 250509_at | 42.6 | A | 32.7 | A | 12.9 | A | 33.9 | P | CYP78A7 |
| 250510_at | 731.7 | P | 753.3 | P | 714.7 | P | 775.3 | P | NRPB4 |
| 250511_at | 446.9 | P | 531.5 | P | 555.5 | P | 597.3 | P | --- |
| 250512_at | 233.2 | P | 373.1 | P | 276.1 | P | 315.1 | P | --- |
| 250513_at | 639.1 | P | 934.6 | P | 790.6 | P | 957.4 | P | --- |
| 250514_at | 25.5 | A | 28.4 | P | 47.4 | A | 30.2 | P | --- |
| 250515_at | 295.6 | P | 179.6 | P | 356.9 | P | 236.1 | P | --- |
| 250516_at | 2255.7 | P | 1776.1 | P | 1694.7 | P | 1848.4 | P | --- |
| 250517_at | 3556.1 | P | 3347.9 | P | 3823.9 | P | 3600.9 | P | scpl35 |
| 250518_at | 258.7 | P | 354.7 | P | 248 | P | 254.3 | P | --- |
| 250519_at | 16.8 | A | 30.3 | A | 34.1 | A | 36.1 | A | --- |
| 250520_at | 210.8 | P | 240 | P | 239.7 | P | 189.4 | P | PEX1 |
| 250521_at | 19.8 | A | 20.2 | A | 30.7 | A | 47.3 | A | --- |
| 250522_at | 740.9 | P | 829 | P | 799.4 | P | 799.9 | P | --- |
| 250523_at | 139.1 | P | 112.8 | P | 180.6 | P | 98.2 | P | --- |
| 250524_at | 239.1 | P | 420.6 | P | 374.2 | P | 358 | P | --- |
| 250525_at | 221.6 | P | 283.2 | P | 284.2 | P | 227 | P | ILP1 |
| 250526_at | 2214 | P | 1683.7 | P | 2402.8 | P | 1930.5 | P | --- |
| 250527_at | 878.2 | P | 980.8 | P | 803.7 | P | 1127.1 | P | SNRK2.1 |
| 250528_at | 154.9 | P | 88.8 | P | 186.4 | P | 102.5 | P | --- |
| 250529_at | 512.4 | P | 617 | P | 732.8 | P | 691 | P | --- |
| 250530_at | 129.1 | P | 126.1 | P | 85.1 | P | 132.7 | P | --- |
| 250531_at | 1083.6 | P | 1167.9 | P | 1036.9 | P | 1149.3 | P | --- |
| 250532_at | 1371.4 | P | 1177.8 | P | 1150.1 | P | 1147.6 | P | --- |
| 250533_at | 744.2 | P | 849.5 | P | 1395.9 | P | 1254.5 | P | FLS |
| 250534_at | 3591.8 | P | 3360.7 | P | 3380.4 | P | 3830.8 | P | CI51 |
| 250535_at | 1 | A | 4 | A | 2.9 | A | 11.1 | A | --- |
| 250536_at | 410 | P | 388.6 | P | 386.5 | P | 385.7 | P | --- |
| 250537_at | 70.4 | P | 117.9 | P | 116.9 | P | 114.2 | P | --- |
| 250538_at | 659.7 | P | 397 | P | 646.3 | P | 444.7 | P | STRS2 |
| 250539_at | 839.3 | P | 845 | P | 968 | P | 1056.6 | P | --- |
| 250540_at | 316.3 | P | 413.5 | P | 370.3 | P | 437.8 | P | --- |
| 250541_at | 35.1 | P | 58.3 | P | 64.1 | P | 44.9 | P | --- |
| 250542_at | 108.9 | P | 168.7 | P | 133.7 | P | 124.9 | P | --- |
| 250543_at | 312.3 | P | 268.9 | P | 241.7 | P | 290.2 | P | NF-YB12 |
| 250544_at | 793.4 | P | 697.9 | P | 682.2 | P | 691.7 | P | SYP132 |
| 250545_at | 1002.1 | P | 1284.4 | P | 957.6 | P | 1281.3 | P | ATPK3 |
| 250546_at | 858.6 | P | 681.6 | P | 483.9 | P | 831.4 | P | --- |
| 250547_at | 864.1 | P | 871.5 | P | 808.7 | P | 879.8 | P | --- |
| 250548_at | 231 | P | 338.5 | P | 236.9 | P | 364.7 | P | MBP2C |
| 250549_at | 540.8 | P | 492.8 | P | 673.7 | P | 608.6 | P | --- |
| 250550_at | 115 | P | 166.7 | P | 157.9 | P | 164.6 | P | --- |
| 250551_at | 24.1 | A | 14.3 | A | 40 | A | 48.9 | A | SNAP29 |
| 250552_at | 202.4 | P | 243.3 | P | 200 | P | 246.9 | P | --- |
| 250553_at | 733.2 | P | 1008.6 | P | 905.7 | P | 912.2 | P | --- |
| 250554_at | 452.9 | P | 452.1 | P | 582.5 | P | 437.2 | P | --- |
| 250555_at | 192.5 | P | 188.1 | P | 224 | P | 159.3 | P | --- |
| 250556_at | 215.3 | P | 254.4 | P | 149.8 | P | 191.8 | P | DGK1 |
| 250557_at | 172 | P | 272.1 | P | 178.3 | P | 244.8 | P | CPuORF16 |
| 250558_at | 666.1 | P | 197.9 | P | 1058.9 | P | 382.5 | P | TT7 |
| 250559_at | 167.4 | P | 248.2 | P | 222.9 | P | 222.3 | P | --- |
| 250560_at | 136 | P | 185.2 | P | 151.5 | P | 221.5 | P | RPA70B |
| 250561_at | 1.2 | A | 3.4 | A | 3.6 | A | 1 | A | --- |
| 250562_at | 1796.3 | P | 1576.2 | P | 1864.5 | P | 1553.7 | P | TOM5 |
| 250563_at | 3130.9 | P | 3308.7 | P | 3691.8 | P | 3209.2 | P | --- |
| 250564_at | 2024.6 | P | 1908.2 | P | 2090.8 | P | 1716.6 | P | --- |

| | | | | | | | | | |
|---|---|---|---|---|---|---|---|---|---|
| 250565_at | 1182.1 | P | 1535.9 | P | 1388.3 | P | 1518.5 | P | E13L3 |
| 250566_at | 89.9 | P | 85.1 | P | 129.1 | P | 105.3 | P | TCP17 |
| 250567_at | 26.7 | A | 12.1 | A | 35.3 | A | 55.6 | A | --- |
| 250568_at | 79.7 | P | 73.6 | A | 117.3 | P | 93.7 | P | --- |
| 250569_at | 490.1 | P | 435.7 | P | 568.8 | P | 510.1 | P | BIM1 |
| 250570_at | 540 | P | 662.7 | P | 631.8 | P | 815.8 | P | EMB1873 |
| 250571_at | 261.2 | P | 297 | P | 254.6 | P | 260.5 | P | --- |
| 250572_at | 107 | P | 78.6 | P | 78.7 | P | 107.1 | P | AT5G08230 |
| 250573_at | 31.8 | A | 11.7 | A | 46.8 | A | 4.4 | A | --- |
| 250574_at | 41.8 | A | 60.7 | M | 66.7 | A | 61.5 | A | --- |
| 250575_at | 112.8 | P | 100.8 | P | 21.9 | A | 44.8 | P | --- |
| 250576_at | 7.1 | A | 18.2 | A | 23.5 | A | 28.8 | A | --- |
| 250577_at | 334.7 | P | 404.1 | P | 272.2 | P | 272.5 | P | --- |
| 250578_at | 1.6 | A | 3.2 | A | 2.2 | A | 2.4 | A | --- |
| 250579_at | 10.1 | A | 2.4 | A | 4.4 | A | 4.3 | A | MCT2 |
| 250580_at | 11861 | P | 3637.3 | P | 10249.4 | P | 4624.1 | P | GDH2 |
| 250581_at | 370.7 | P | 518.6 | P | 515.7 | P | 494.4 | P | BON2 |
| 250582_at | 804.9 | P | 1118.2 | P | 854.7 | P | 1052.3 | P | --- |
| 250583_at | 29.6 | A | 35.4 | A | 6.3 | A | 16.3 | A | PEI1 |
| 250584_at | 175.4 | P | 180.5 | P | 203.6 | P | 221.9 | P | --- |
| 250585_at | 56.2 | P | 74.6 | P | 81.7 | P | 73.6 | P | --- |
| 250586_at | 159.2 | P | 220.4 | P | 142.3 | P | 213.8 | P | --- |
| 250587_at | 3.4 | A | 2.8 | A | 11.3 | A | 22.8 | A | --- |
| 250588_at | 56.1 | A | 50.9 | A | 67.4 | A | 44 | A | --- |
| 250589_at | 128.9 | P | 78.2 | A | 161.5 | P | 71.1 | M | MYB76 |
| 250590_at | 178.1 | P | 185.4 | P | 243.5 | P | 169.8 | P | --- |
| 250591_at | 54.2 | P | 65.9 | P | 55.1 | A | 65.8 | P | --- |
| 250592_at | 53.2 | A | 32.3 | A | 13.3 | A | 66.4 | P | --- |
| 250593_at | 60.3 | A | 58.6 | A | 76.6 | A | 83.5 | A | --- |
| 250594_at | 26.5 | A | 2.6 | A | 15.4 | A | 19.4 | A | --- |
| 250595_at | 35.2 | A | 73.1 | P | 42.5 | A | 34.1 | M | --- |
| 250596_at | 21.4 | A | 2.6 | A | 3.6 | A | 2.3 | A | --- |
| 250597_at | 56.9 | A | 45.8 | A | 17.6 | A | 49.2 | A | ANAC080 |
| 250598_at | 271.3 | P | 470.6 | P | 273.4 | P | 390.7 | P | ATMYB29 |
| 250599_at | 196.1 | P | 152.2 | P | 229.9 | P | 175.8 | P | --- |
| 250600_at | 69.3 | A | 92.3 | P | 84.9 | A | 87.4 | P | --- |
| 250601_at | 27.5 | A | 63.5 | P | 39 | A | 59.1 | P | --- |
| 250602_s_at | 305.9 | P | 258.2 | P | 240.2 | P | 244.3 | P | --- |
| 250603_at | 105.9 | P | 119 | P | 52.7 | P | 79.6 | P | --- |
| 250604_at | 1379.4 | P | 1565.7 | P | 1146.9 | P | 1225.4 | P | PIP1B |
| 250605_at | 23.6 | A | 33.8 | A | 35.4 | A | 29.3 | A | --- |
| 250606_s_at | 5.4 | A | 1.6 | A | 8.9 | A | 1.7 | A | PPME1 |
| 250607_at | 1058.3 | P | 988.4 | P | 957.9 | P | 940.2 | P | IPK2a |
| 250608_at | 2.2 | A | 1.4 | A | 2.7 | A | 1.2 | A | --- |
| 250609_at | 1304.1 | P | 1355.9 | P | 1336.1 | P | 1178.3 | P | PMSR3 |
| 250610_at | 17.6 | A | 17.4 | A | 36.7 | A | 8.2 | A | GRP19 |
| 250611_at | 40.1 | A | 18.9 | A | 36.3 | A | 42.3 | A | GA20OX3 |
| 250612_s_at | 2.2 | A | 2.3 | A | 2.1 | A | 2.5 | A | ARR13 |
| 250613_at | 498.4 | P | 448.2 | P | 436.4 | P | 472.5 | P | IQD24 |
| 250614_at | 22.2 | A | 15.1 | A | 18.3 | A | 3.4 | A | --- |
| 250615_at | 343.2 | P | 341.1 | P | 304.6 | P | 461.8 | P | --- |
| 250616_at | 77.1 | M | 120.2 | P | 126.8 | P | 104.8 | P | EMS1 |
| 250617_at | 340.4 | P | 434.8 | P | 350 | P | 424.4 | P | AML4 |
| 250618_at | 866.9 | P | 775.6 | P | 757 | P | 749.9 | P | ATBAG3 |
| 250619_at | 1.8 | A | 4.1 | A | 2.3 | A | 2.1 | A | --- |
| 250620_at | 17.4 | A | 6.7 | A | 44.1 | A | 26.5 | A | ATS3 |
| 250621_at | 533.3 | P | 562.6 | P | 494.3 | P | 597.5 | P | ATRBL3 |
| 250622_at | 29.1 | A | 21.6 | A | 3.1 | A | 1 | A | --- |
| 250623_at | 373.3 | P | 368 | P | 391.6 | P | 409.6 | P | --- |
| 250624_at | 4.7 | A | 3 | A | 8.9 | A | 35.7 | A | --- |
| 250625_at | 627.6 | P | 776.3 | P | 543.8 | P | 639.1 | P | --- |
| 250626_at | 1559.5 | P | 1400.6 | P | 1582.5 | P | 1380.8 | P | --- |
| 250627_at | 287.5 | P | 277.5 | P | 215.8 | P | 259.3 | P | --- |
| 250628_at | 15.6 | A | 11.1 | A | 11.2 | A | 32.8 | A | --- |
| 250629_at | 4 | A | 5.1 | A | 20.3 | A | 3.7 | A | ATRBOHA |
| 250630_at | 82.3 | A | 55.9 | A | 53.5 | A | 45.6 | A | --- |
| 250631_at | 2.4 | A | 2.3 | A | 4.6 | A | 1.9 | A | --- |
| 250632_at | 17.9 | A | 14.7 | A | 28.2 | A | 15.1 | A | CYCP4;3 |
| 250633_at | 1716.6 | P | 1045.3 | P | 1900.8 | P | 1001.6 | P | PMSR2 |
| 250634_at | 5.2 | A | 3.6 | A | 8.2 | A | 6.7 | A | --- |
| 250635_at | 3.8 | A | 7 | A | 8.9 | A | 9.6 | A | GRP14 |
| 250636_at | 12.7 | A | 21.5 | A | 4.8 | A | 2 | A | GRP18 |
| 250637_at | 62 | A | 30.1 | A | 32.5 | A | 37.6 | A | GRP17 |
| 250638_at | 32.4 | A | 27.2 | A | 34.6 | A | 18.2 | A | GRP16 |
| 250639_at | 33.9 | A | 36.2 | A | 69.5 | A | 21 | A | GRP20 |
| 250640_at | 12.1 | A | 43.6 | A | 33.1 | A | 17.2 | A | --- |
| 250641_at | 1.6 | A | 2 | A | 3.1 | A | 1 | A | --- |
| 250642_at | 152.7 | P | 141.8 | P | 188.5 | P | 182.1 | P | ERL2 |
| 250643_at | 256.2 | P | 263 | P | 270.4 | P | 184 | P | --- |
| 250644_at | 470.8 | P | 448.2 | P | 298.3 | P | 376.2 | P | --- |
| 250645_at | 800.5 | P | 998.1 | P | 770.5 | P | 951.7 | P | --- |
| 250646_at | 104.9 | P | 74.3 | P | 100.2 | P | 92.5 | P | --- |
| 250647_at | 155 | P | 245.9 | P | 224.9 | P | 272 | P | --- |
| 250648_at | 81.9 | P | 47 | A | 3.5 | A | 54 | A | --- |
| 250649_at | 1106.3 | P | 1133.8 | P | 1133.4 | P | 1098.7 | P | WCRKC1 |
| 250650_at | 80.6 | A | 67.6 | A | 112.4 | A | 51.5 | A | --- |
| 250651_at | 0.7 | A | 2.6 | A | 6.3 | A | 0.8 | A | CYP93D1 |
| 250652_at | 41.2 | A | 32.9 | A | 28.1 | A | 6.8 | A | FLA21 |
| 250653_at | 58.1 | A | 126.2 | P | 64.9 | P | 95.5 | P | --- |
| 250654_at | 7.7 | A | 85 | P | 41.3 | A | 73 | P | --- |
| 250655_at | 330.3 | P | 507.9 | P | 375.3 | P | 380.5 | P | OBF5 |
| 250656_at | 338.9 | P | 479.2 | P | 302.9 | P | 446 | P | --- |
| 250657_at | 120.3 | P | 125.1 | P | 88.2 | P | 138.1 | P | ST2B |
| 250658_at | 61 | P | 55.9 | P | 46.5 | A | 47.5 | M | --- |
| 250659_at | 14.4 | A | 38.8 | A | 71.1 | A | 17.5 | A | --- |
| 250660_at | 52.7 | P | 28.3 | P | 20.9 | A | 47.7 | P | --- |
| 250661_at | 333 | P | 1350.3 | P | 998.5 | P | 1281.7 | P | --- |
| 250662_at | 176.2 | P | 61 | P | 143.4 | P | 59.1 | P | ST2A |
| 250663_at | 209.6 | P | 178.5 | P | 179 | P | 249.1 | P | PRA1.B6 |
| 250664_at | 96.5 | P | 115.4 | P | 96.9 | P | 111.3 | P | --- |
| 250665_at | 147.4 | P | 476 | P | 141.2 | P | 328 | P | --- |
| 250666_at | 118.7 | P | 135.9 | P | 105.8 | P | 106.1 | P | WRKY26 |
| 250667_at | 3944.6 | P | 3643.2 | P | 4550.9 | P | 3757.3 | P | --- |
| 250668_at | 2551.6 | P | 3284.3 | P | 2969.9 | P | 3058.7 | P | --- |
| 250669_at | 4493.8 | P | 894.3 | P | 2805.8 | P | 1031.5 | P | PGIP2 |
| 250670_at | 4521.4 | P | 939.5 | P | 3286.1 | P | 1090.7 | P | PGIP1 |
| 250671_at | 251.9 | P | 293 | P | 198.1 | P | 289.7 | P | AHBP-1B |
| 250672_at | 403.5 | P | 419.2 | P | 320.8 | P | 399.9 | P | ATJ6 |
| 250673_at | 84.7 | A | 61.7 | A | 57.8 | A | 89.5 | A | CIPK2 |

| | | | | | | | | | |
|---|---|---|---|---|---|---|---|---|---|
| 250674_at | 54.6 | P | 60.9 | P | 73.2 | P | 76.5 | P | LAC13 |
| 250675_at | 895.1 | P | 939.4 | P | 1051.9 | P | 904.5 | P | --- |
| 250676_at | 1865.7 | P | 2905.7 | P | 1262.2 | P | 1885.4 | P | NHL3 |
| 250677_at | 17.4 | A | 18.2 | A | 20.6 | A | 27.7 | A | --- |
| 250678_at | 42.6 | A | 32.7 | M | 12.8 | A | 31.6 | A | --- |
| 250679_at | 346.4 | P | 201.3 | P | 393.1 | P | 244.4 | P | --- |
| 250680_at | 29.5 | A | 26.5 | A | 41.1 | A | 32.7 | A | --- |
| 250681_at | 99.4 | P | 128.9 | P | 129.4 | P | 118.6 | P | --- |
| 250682_x_at | 82.6 | P | 99.9 | P | 120.2 | P | 95.5 | P | --- |
| 250683_x_at | 17.6 | A | 66 | A | 73.1 | P | 37.3 | A | --- |
| 250684_at | 0.9 | A | 19.3 | A | 36.7 | A | 9.8 | A | GIS2 |
| 250685_at | 47 | P | 57 | P | 53.4 | A | 69.8 | P | --- |
| 250686_at | 101.5 | P | 131.3 | P | 109.2 | P | 140.2 | P | SPC98 |
| 250687_at | 631.2 | P | 895.6 | P | 737.1 | P | 860.8 | P | --- |
| 250688_at | 291 | P | 418.7 | P | 276.8 | P | 485.5 | P | NF-YA10 |
| 250689_at | 147 | P | 188.9 | P | 144.6 | P | 166.1 | P | --- |
| 250690_at | 326.2 | P | 456.1 | P | 389.3 | P | 514.9 | P | --- |
| 250691_at | 553.4 | P | 637.2 | P | 501.8 | P | 461.4 | P | --- |
| 250692_at | 279.6 | P | 310.6 | P | 290.2 | P | 269 | P | --- |
| 250693_at | 907 | P | 821.2 | P | 870.3 | P | 956.4 | P | UBP12 |
| 250694_at | 26.6 | A | 80.5 | A | 51.6 | A | 55.3 | A | HAT14 |
| 250695_at | 2.9 | A | 4.1 | A | 2.8 | A | 4 | A | --- |
| 250696_at | 98.9 | P | 121.4 | P | 197 | P | 130.3 | P | --- |
| 250697_at | 29.1 | A | 39.1 | A | 57.4 | A | 41.2 | A | --- |
| 250698_at | 163.7 | P | 181.9 | P | 193.3 | P | 168 | P | --- |
| 250699_at | 3.6 | A | 2.7 | A | 5.7 | A | 3.8 | A | SRF2 |
| 250700_at | 234.9 | P | 287.6 | P | 238.3 | P | 208.6 | P | --- |
| 250701_at | 38.3 | A | 57.6 | A | 45.9 | A | 43 | A | --- |
| 250702_at | 26 | A | 35.1 | A | 10.2 | A | 32.8 | A | --- |
| 250703_at | 2520.9 | P | 1854.1 | P | 2469.2 | P | 1732.5 | P | --- |
| 250704_at | 297.2 | P | 341.4 | P | 299.5 | P | 299.9 | P | --- |
| 250705_at | 3001.2 | P | 2824.3 | P | 3022.3 | P | 2667.1 | P | ATNUDX27 |
| 250706_at | 289.3 | P | 333.1 | P | 310.4 | P | 307.4 | P | DHS |
| 250707_at | 469.8 | P | 576.9 | P | 582.7 | P | 566.2 | P | MEE60 |
| 250708_at | 6.1 | A | 35.6 | A | 12.5 | A | 17.9 | A | RBE |
| 250709_at | 0.7 | A | 18.5 | A | 44 | A | 29.1 | A | LBD33 |
| 250710_at | 130.6 | P | 154.8 | P | 175.5 | P | 114.3 | P | MYB33 |
| 250711_at | 843.8 | P | 484.9 | P | 780.1 | P | 587.4 | P | --- |
| 250712_at | 467.8 | P | 448.1 | P | 405.8 | P | 449.4 | P | --- |
| 250713_at | 2130.4 | P | 2018.7 | P | 1801.3 | P | 1802.5 | P | --- |
| 250714_at | 918.6 | P | 905.5 | P | 857.4 | P | 827.3 | P | SNX1 |
| 250715_at | 261.5 | P | 303.2 | P | 211.1 | P | 258.6 | P | ATO |
| 250716_at | 5.1 | A | 47.7 | A | 23.9 | A | 42.6 | A | ATSUC9 |
| 250717_at | 4.4 | A | 67 | A | 34.6 | A | 22 | A | --- |
| 250718_at | 404.2 | P | 374.6 | P | 368.8 | P | 375.7 | P | emb2735 |
| 250719_at | 88.6 | P | 77.5 | P | 129.1 | P | 92.5 | P | --- |
| 250720_at | 152.4 | P | 167.7 | P | 138.5 | P | 132.9 | P | --- |
| 250721_at | 667.6 | P | 533.1 | P | 786.3 | P | 506.3 | P | --- |
| 250722_at | 144 | P | 58.4 | P | 80.7 | P | 115 | P | --- |
| 250723_at | 118 | P | 107 | P | 135 | P | 98.2 | P | --- |
| 250724_at | 60.2 | A | 116.8 | A | 65 | A | 96.2 | P | --- |
| 250725_at | 67.4 | P | 75.7 | P | 94 | P | 58.4 | P | --- |
| 250726_at | 313.8 | P | 255 | P | 367.2 | P | 295.7 | P | --- |
| 250727_at | 120.8 | P | 150.8 | P | 154.6 | P | 108 | P | --- |
| 250728_at | 340.7 | P | 275.1 | P | 219.6 | P | 242.3 | P | --- |
| 250729_at | 941 | P | 796.3 | P | 795.9 | P | 860.2 | P | UBA 2 |
| 250730_at | 20.5 | A | 13.6 | A | 21.7 | A | 16.3 | A | --- |
| 250731_at | 6 | A | 20.2 | A | 5.5 | A | 19.1 | A | AGL96 |
| 250732_at | 128 | P | 150.9 | P | 140.3 | P | 155.8 | P | --- |
| 250733_at | 3205.2 | P | 5122.9 | P | 4139 | P | 4946.7 | P | 2-Cys Prx B |
| 250734_at | 180.3 | P | 230.4 | P | 206.2 | P | 269.2 | P | --- |
| 250735_at | 606 | P | 641.8 | P | 673.9 | P | 547.7 | P | --- |
| 250736_s_at | 547.6 | P | 374 | P | 470.3 | P | 520.6 | P | --- |
| 250737_at | 936.1 | P | 894.7 | P | 893.1 | P | 769 | P | --- |
| 250738_at | 3328.5 | P | 1463.2 | P | 2256.2 | P | 1284.9 | P | ASA1 |
| 250739_at | 1044.6 | P | 1292.5 | P | 987.7 | P | 1183.2 | P | EGY2 |
| 250740_at | 141.7 | P | 170.6 | P | 127.2 | P | 120.3 | P | SYP31 |
| 250741_at | 119.6 | P | 165.4 | P | 103.4 | P | 170.2 | P | --- |
| 250742_at | 307.2 | P | 311.2 | P | 326.4 | P | 272.5 | P | --- |
| 250743_at | 400.7 | P | 504.6 | P | 383.4 | P | 458.1 | P | --- |
| 250744_at | 18.1 | A | 25.6 | A | 16.6 | A | 52.3 | P | --- |
| 250745_at | 152 | P | 146 | P | 137.5 | P | 146.8 | P | --- |
| 250746_at | 22.5 | A | 55 | A | 8.9 | A | 45.1 | A | --- |
| 250747_at | 1.6 | A | 1.1 | A | 5.4 | A | 1.3 | A | --- |
| 250748_at | 251.9 | P | 320.9 | P | 346.1 | P | 293.7 | P | --- |
| 250749_at | 1496.5 | P | 1647.9 | P | 1304.8 | P | 1762.3 | P | RPN8A |
| 250750_at | 260.5 | P | 249 | P | 273.8 | P | 195.6 | P | UGT76C2 |
| 250751_at | 80.4 | P | 93.4 | P | 81.6 | A | 112.2 | P | --- |
| 250752_at | 1949.6 | P | 1910.6 | P | 1816.5 | P | 1634.9 | P | CPD |
| 250753_at | 318.2 | P | 263.9 | P | 430.7 | P | 279.4 | P | UGT76C2 |
| 250754_at | 493.8 | P | 547.8 | P | 507.9 | P | 481.4 | P | ATE1 |
| 250755_at | 377.4 | P | 434.7 | P | 402.2 | P | 353.1 | P | --- |
| 250756_at | 43.8 | A | 81.1 | P | 100 | P | 92.5 | P | ROPGEF5 |
| 250757_at | 704.2 | P | 563 | P | 790.8 | P | 632.1 | P | ATDFB |
| 250758_at | 175.4 | P | 182.7 | P | 225 | P | 132 | P | EIF3G2 |
| 250759_at | 9.5 | A | 2.4 | A | 20.5 | A | 8.3 | A | --- |
| 250760_at | 3.9 | A | 1.2 | A | 2.9 | A | 3.7 | A | --- |
| 250761_at | 345.5 | P | 395.3 | P | 485.7 | P | 458.7 | P | --- |
| 250762_at | 433.5 | P | 292.1 | P | 540.7 | P | 336.5 | P | --- |
| 250763_at | 955.6 | P | 1121.9 | P | 1178.5 | P | 1009.9 | P | --- |
| 250764_at | 232.4 | P | 413 | P | 432.8 | P | 569.2 | P | --- |
| 250765_at | 528.2 | P | 483.5 | P | 521.7 | P | 497.6 | P | --- |
| 250766_at | 131.2 | P | 126.3 | P | 135.3 | P | 98.7 | P | --- |
| 250767_at | 359.5 | P | 308.1 | P | 471.5 | P | 286.6 | P | ATNFXL2 |
| 250768_at | 274.1 | P | 290.5 | P | 261 | P | 285.8 | P | --- |
| 250769_at | 162.8 | P | 113.1 | P | 225 | P | 141.3 | P | --- |
| 250770_at | 6.4 | A | 26.9 | A | 34.6 | A | 34.1 | A | LAC12 |
| 250771_at | 33.7 | P | 24.9 | A | 8.7 | A | 28.5 | A | --- |
| 250772_at | 57.5 | P | 21.4 | A | 56.4 | P | 30.7 | A | --- |
| 250773_at | 41.8 | P | 52.6 | P | 63.3 | P | 27.6 | A | --- |
| 250774_at | 372.4 | P | 323 | P | 422.1 | P | 273.6 | P | --- |
| 250775_at | 136.7 | P | 187.8 | P | 174.3 | P | 135.2 | P | --- |
| 250776_at | 133.2 | P | 141.2 | P | 89.7 | P | 107.7 | P | --- |
| 250777_at | 1002.6 | P | 907.3 | P | 1054.7 | P | 687.2 | P | --- |
| 250778_at | 12 | A | 28.2 | A | 34.7 | A | 2.6 | A | --- |
| 250779_at | 399.1 | P | 327.8 | P | 501.3 | P | 353.3 | P | EIF2 ALPHA |
| 250780_at | 9.1 | A | 8.8 | A | 24.8 | A | 36.2 | A | ATEXPA2 |
| 250781_at | 267.7 | P | 424.1 | P | 328 | P | 376.3 | P | DREB2A |
| 250782_at | 28.3 | A | 3.4 | A | 3.9 | A | 17 | A | SYN1 |

| | | | | | | | | | |
|---|---|---|---|---|---|---|---|---|---|
| 250783_at | 2.7 | A | 3.1 | A | 2.4 | A | 1.1 | A | CYP79A2 |
| 250784_at | 440.5 | P | 329 | P | 343 | P | 335.2 | P | --- |
| 250785_at | 46.9 | A | 65.4 | A | 43.4 | A | 61.2 | A | --- |
| 250786_at | 299.5 | P | 239 | P | 341.3 | P | 218.3 | P | SDN2 |
| 250787_at | 281.7 | P | 276.7 | P | 256.9 | P | 246.9 | P | EMB2771 |
| 250788_at | 135.9 | P | 186.4 | P | 171.8 | P | 161.1 | P | --- |
| 250789_at | 62.4 | A | 62.5 | A | 78.8 | A | 84.2 | P | --- |
| 250790_at | 189.7 | P | 89.7 | P | 273 | P | 145.6 | P | --- |
| 250791_at | 482.7 | P | 512.9 | P | 469.4 | P | 443.3 | P | AL1 |
| 250792_at | 590.2 | P | 531.6 | P | 661.5 | P | 605.9 | P | --- |
| 250793_at | 2753.1 | P | 404.6 | P | 1712.4 | P | 547.4 | P | --- |
| 250794_at | 481.6 | P | 494.6 | P | 855 | P | 795.5 | P | --- |
| 250795_at | 1.9 | A | 2.6 | A | 4.9 | A | 2 | A | --- |
| 250796_at | 81.9 | A | 67.2 | P | 54.3 | A | 44 | A | --- |
| 250797_at | 168.5 | P | 184 | P | 210.5 | P | 178 | P | --- |
| 250798_at | 89.2 | P | 14.9 | A | 61.9 | A | 35.9 | A | --- |
| 250799_at | 88.1 | A | 84.2 | P | 94.3 | A | 75.8 | A | --- |
| 250800_at | 2614.6 | P | 1835.1 | P | 2052.4 | P | 2085.5 | P | HPR |
| 250801_at | 10.4 | A | 23.5 | A | 21.1 | A | 10.4 | A | --- |
| 250802_at | 1 | A | 1.2 | A | 4 | A | 3.2 | A | --- |
| 250803_at | 57.1 | P | 88.4 | P | 80.8 | P | 67.7 | P | --- |
| 250804_at | 10.1 | A | 8.2 | A | 8.4 | A | 8.7 | A | --- |
| 250805_at | 12.1 | A | 11.3 | A | 5.6 | A | 4.5 | A | --- |
| 250806_at | 2.2 | A | 3 | A | 3.6 | A | 2.4 | A | --- |
| 250807_at | 68.7 | P | 71 | P | 106.7 | P | 60.6 | P | --- |
| 250808_at | 2.6 | A | 3.2 | A | 4.7 | A | 11.8 | A | ATG18E |
| 250809_at | 266 | P | 154.6 | P | 224.5 | P | 173.2 | P | --- |
| 250810_at | 82.1 | P | 150 | P | 102.2 | A | 115 | P | --- |
| 250811_at | 709.4 | P | 636.1 | P | 455.4 | P | 528.8 | P | --- |
| 250812_at | 979.4 | P | 791.7 | P | 804.4 | P | 807.2 | P | --- |
| 250813_at | 658.6 | P | 769.7 | P | 795 | P | 687.1 | P | TOC34 |
| 250814_at | 852.6 | P | 1124.2 | P | 835.8 | P | 873.6 | P | UBC22 |
| 250815_s_at | 1022.4 | P | 1185.1 | P | 1140.7 | P | 1067.3 | P | --- |
| 250816_at | 1864.6 | P | 2063.2 | P | 2071.8 | P | 2072.1 | P | --- |
| 250817_at | 291.4 | P | 362.8 | P | 353.1 | P | 403.3 | P | SUVH1 |
| 250818_at | 557.2 | P | 654.5 | P | 511.9 | P | 522.9 | P | ALA1 |
| 250819_at | 958.4 | P | 899.7 | P | 943.5 | P | 891.8 | P | --- |
| 250820_at | 161.7 | P | 188.2 | P | 218.2 | P | 203.8 | P | --- |
| 250821_at | 122.9 | P | 139.5 | P | 129 | P | 116.7 | P | --- |
| 250822_at | 200.5 | P | 235.2 | P | 197.4 | P | 193.9 | P | --- |
| 250823_at | 81.2 | P | 156 | P | 88.8 | P | 149.8 | P | --- |
| 250824_at | 2627 | P | 3331.8 | P | 2687.2 | P | 3319.9 | P | --- |
| 250825_at | 296.2 | P | 350.5 | P | 451.4 | P | 419.9 | P | --- |
| 250826_at | 24.1 | A | 8.4 | A | 13.1 | A | 6.1 | A | --- |
| 250827_at | 3903.4 | P | 5338.2 | P | 3327.5 | P | 4887 | P | CEV1 |
| 250828_at | 178.7 | P | 1407 | P | 889.3 | P | 917.8 | P | --- |
| 250829_at | 912.6 | P | 1137.7 | P | 849.1 | P | 874.4 | P | ADR1-L2 |
| 250830_at | 546.1 | P | 521.4 | P | 398.4 | P | 467.7 | P | --- |
| 250831_at | 508.6 | P | 560.7 | P | 445.6 | P | 483 | P | --- |
| 250832_at | 44.3 | A | 86 | P | 105.8 | P | 98.4 | P | NAS1 |
| 250833_at | 314.2 | P | 300.7 | P | 265 | P | 294 | P | --- |
| 250834_at | 87.7 | P | 70.5 | P | 74.2 | P | 60.8 | P | DME |
| 250835_at | 83.9 | P | 101 | P | 104.2 | P | 95.7 | P | DME |
| 250836_at | 188.3 | P | 268.4 | P | 196 | P | 177.8 | P | DME |
| 250837_at | 192.1 | P | 167.8 | P | 215.7 | P | 202.9 | P | ATBIOF |
| 250838_at | 1.6 | A | 3.3 | A | 4.3 | A | 4.1 | A | CYP77A9 |
| 250839_at | 3.8 | A | 1.5 | A | 2 | A | 0.8 | A | AGL99 |
| 250840_at | 33.8 | A | 13.3 | A | 7.8 | A | 6.1 | A | --- |
| 250841_at | 93.1 | P | 69 | M | 81.4 | P | 53.3 | A | --- |
| 250842_at | 775.2 | P | 1026 | P | 563.3 | P | 843.6 | P | VTE5 |
| 250843_at | 2509.6 | P | 2598.6 | P | 2564.7 | P | 2716.6 | P | BTR1L |
| 250844_at | 249.2 | P | 231.3 | P | 193.9 | P | 177.9 | P | SIM |
| 250845_at | 1366.4 | P | 1173 | P | 1726.5 | P | 1219.5 | P | --- |
| 250846_at | 4442.8 | P | 2535.9 | P | 3970.4 | P | 2700.6 | P | SIR |
| 250847_at | 182.9 | P | 223 | P | 202.3 | P | 232.1 | P | --- |
| 250848_at | 351.7 | P | 374.2 | P | 431.6 | P | 355.2 | P | PDK1 |
| 250849_at | 1255 | P | 1301.9 | P | 1124.9 | P | 1269.1 | P | NAC2 |
| 250850_at | 377.8 | P | 478.3 | P | 403.5 | P | 449.7 | P | --- |
| 250851_at | 427.7 | P | 477.6 | P | 462.6 | P | 565 | P | --- |
| 250852_at | 235 | P | 179.5 | P | 158.5 | P | 162.9 | P | --- |
| 250853_s_at | 2.1 | A | 3.3 | A | 27.7 | A | 21.1 | A | --- |
| 250854_at | 715.8 | P | 634.5 | P | 723.2 | P | 683.1 | P | --- |
| 250855_at | 10.3 | A | 3.4 | A | 2.3 | A | 3 | A | --- |
| 250856_at | 488 | P | 569.8 | P | 612.9 | P | 599.9 | P | --- |
| 250857_at | 138.5 | P | 142.5 | P | 112.2 | P | 125.2 | P | --- |
| 250858_at | 481.1 | P | 461.8 | P | 431.4 | P | 417.4 | P | --- |
| 250859_at | 83.4 | P | 95.2 | P | 105.5 | P | 110.6 | P | CYP77A4 |
| 250860_at | 10.2 | A | 75.2 | P | 3.9 | A | 82.1 | P | CAT6 |
| 250861_at | 1206.1 | P | 1275 | P | 990.4 | P | 1142.3 | P | --- |
| 250862_s_at | 9490.7 | P | 8650.6 | P | 12690.7 | P | 9251.5 | P | --- |
| 250863_at | 1873.4 | P | 2207 | P | 1671.5 | P | 2319.4 | P | --- |
| 250864_at | 45.1 | P | 71.6 | P | 69.4 | A | 78.3 | P | --- |
| 250865_at | 789.3 | P | 977.7 | P | 800.9 | P | 886.5 | P | --- |
| 250866_at | 703 | P | 636.3 | P | 572 | P | 721.5 | P | --- |
| 250867_at | 3210.7 | P | 2977.4 | P | 3144.6 | P | 2757.2 | P | --- |
| 250868_at | 41 | A | 22 | A | 57.4 | A | 41 | A | MLS |
| 250869_at | 24.3 | A | 5.5 | A | 48.1 | A | 16.6 | A | TFL1 |
| 250870_at | 19.4 | A | 2.5 | A | 17.7 | A | 9.8 | A | --- |
| 250871_at | 30.5 | A | 26.4 | A | 27.9 | A | 18.6 | A | --- |
| 250872_at | 66.9 | P | 97.2 | P | 45 | A | 63.6 | P | IQD12 |
| 250873_at | 2.6 | A | 19.9 | A | 17.8 | A | 6.7 | A | --- |
| 250874_at | 9 | A | 9 | A | 7.6 | A | 15.9 | A | --- |
| 250875_at | 253.7 | P | 165.9 | P | 153 | P | 124.7 | P | --- |
| 250876_at | 19 | A | 20.4 | A | 4 | A | 32 | A | --- |
| 250877_at | 388.7 | P | 385.1 | P | 333.3 | P | 317.6 | P | SDP1 |
| 250878_at | 69.8 | P | 115.1 | P | 82.5 | P | 76.8 | P | --- |
| 250879_at | 235.1 | P | 223.3 | P | 206.5 | P | 187.2 | P | --- |
| 250880_at | 134.6 | P | 128.7 | P | 172.8 | P | 135.9 | P | --- |
| 250881_at | 144.6 | P | 176.2 | P | 163.6 | P | 177.8 | P | --- |
| 250882_at | 53.8 | P | 65.5 | A | 55.7 | A | 43.8 | P | --- |
| 250883_at | 495 | P | 424.8 | P | 568.4 | P | 447.9 | P | --- |
| 250884_at | 2076.6 | P | 2197.2 | P | 1917.7 | P | 2071.7 | P | CPSRP54 |
| 250885_at | 366.7 | P | 458.5 | P | 304.1 | P | 440.7 | P | ATH12 |
| 250886_at | 899.1 | P | 1220.3 | P | 1130.5 | P | 1095.5 | P | --- |
| 250887_at | 67.9 | P | 91.2 | P | 61 | A | 80.5 | P | --- |
| 250888_at | 124.2 | P | 119.7 | P | 131.9 | P | 127.6 | P | --- |
| 250889_at | 17.5 | A | 11.3 | A | 5.1 | A | 6 | A | --- |
| 250890_at | 275.8 | P | 241.9 | P | 252.7 | P | 221.4 | P | --- |
| 250891_at | 166.5 | P | 407.5 | P | 271.9 | P | 310.2 | P | KCS19 |

| | | | | | | | | | |
|---|---|---|---|---|---|---|---|---|---|
| 250892_at | 441.1 | P | 903.5 | P | 389.9 | P | 848 | P | ATCSLA09 |
| 250893_at | 24.7 | A | 41.6 | A | 18.8 | A | 51.2 | A | EMB175 |
| 250894_s_at | 2.9 | A | 2.8 | A | 1.4 | A | 3.5 | A | --- |
| 250895_at | 5207.7 | P | 5250.7 | P | 5878.9 | P | 5421.8 | P | --- |
| 250896_at | 203.8 | P | 230.9 | P | 196.8 | P | 201.4 | P | --- |
| 250897_at | 1295.6 | P | 1408.3 | P | 1525 | P | 1490.7 | P | --- |
| 250898_at | 3416.4 | P | 3561.3 | P | 3979.1 | P | 3976 | P | ADK2 |
| 250899_at | 531.6 | P | 590.2 | P | 429 | P | 573.2 | P | --- |
| 250900_at | 201.7 | P | 265.3 | P | 354.4 | P | 297.4 | P | ATB' ALPHA |
| 250901_at | 115.6 | P | 126.8 | P | 146.4 | P | 153.9 | P | RABC2A |
| 250902_at | 0.9 | A | 1 | A | 16.7 | A | 0.9 | A | --- |
| 250903_at | 24.2 | A | 11.8 | A | 52.2 | A | 23.5 | A | --- |
| 250904_at | 12.3 | A | 24.9 | A | 16.9 | A | 15.5 | A | --- |
| 250905_at | 30 | A | 33.5 | A | 34.7 | A | 31.7 | A | --- |
| 250906_at | 1234.4 | P | 1739.9 | P | 1684.7 | P | 2017.4 | P | SBE2.2 |
| 250907_at | 198.6 | P | 110 | P | 165.4 | P | 146.3 | P | --- |
| 250908_at | 16.6 | A | 48.3 | A | 54 | A | 43 | A | PTL |
| 250909_at | 114.9 | P | 127.1 | P | 131.8 | P | 121.6 | P | --- |
| 250910_at | 60.4 | P | 164.9 | P | 136.3 | P | 114.6 | P | HSFA3 |
| 250911_at | 448.7 | P | 593.9 | P | 501.8 | P | 493.9 | P | CTR1 |
| 250912_at | 797.8 | P | 807.3 | P | 941.2 | P | 801.4 | P | HD2C |
| 250913_at | 159.7 | P | 117.5 | P | 119 | P | 83.5 | P | --- |
| 250914_at | 36.8 | P | 68.8 | P | 53.2 | P | 33.8 | P | TRFL10 |
| 250915_at | 18.2 | A | 21 | M | 32.1 | M | 18.9 | A | HB51 |
| 250916_at | 2226.2 | P | 2081.4 | P | 2150.6 | P | 2109.5 | P | ATMDAR2 |
| 250917_at | 290 | P | 255.2 | P | 310.4 | P | 277.9 | P | --- |
| 250918_at | 1006.1 | P | 775.9 | P | 929.2 | P | 801.4 | P | --- |
| 250919_at | 1021.9 | P | 1036.4 | P | 987.7 | P | 1057.5 | P | --- |
| 250920_at | 81.3 | A | 62.5 | A | 96.1 | A | 88.6 | A | --- |
| 250921_at | 1384.6 | P | 1344.1 | P | 1167.4 | P | 1329.4 | P | --- |
| 250922_at | 1494.4 | P | 1821.7 | P | 1555.2 | P | 1761.5 | P | --- |
| 250923_at | 2774.7 | P | 2620 | P | 2409.5 | P | 2330.5 | P | CDC25 |
| 250924_at | 247.5 | P | 251.5 | P | 215.8 | P | 227.3 | P | --- |
| 250925_at | 412.5 | P | 371 | P | 351.5 | P | 362.9 | P | --- |
| 250926_at | 452.8 | P | 541.4 | P | 577.4 | P | 570.1 | P | --- |
| 250927_at | 39.7 | P | 41.4 | P | 64.6 | P | 61.3 | P | --- |
| 250928_at | 685.8 | P | 734.7 | P | 579.6 | P | 726.9 | P | EIN2 |
| 250929_at | 2880.2 | P | 2365.6 | P | 2323.4 | P | 2829.7 | P | --- |
| 250930_at | 504.7 | P | 420 | P | 386.2 | P | 457.6 | P | ATP58IPK |
| 250931_at | 549.5 | P | 470 | P | 436.8 | P | 338.3 | P | --- |
| 250932_s_at | 529.1 | P | 387.8 | P | 438.6 | P | 351.2 | P | --- |
| 250933_at | 217.9 | P | 312.8 | P | 259.2 | P | 249.7 | P | FLA11 |
| 250934_at | 2065.8 | P | 1560.1 | P | 1643.3 | P | 1593.4 | P | --- |
| 250935_at | 5121.6 | P | 4160.9 | P | 4086.8 | P | 4022.9 | P | UBQ3 |
| 250936_at | 205.3 | P | 298.2 | P | 184.3 | P | 279.5 | P | --- |
| 250937_at | 291.1 | P | 344.9 | P | 148.4 | P | 234.8 | P | --- |
| 250938_at | 133.8 | P | 181.4 | A | 148 | M | 143 | M | --- |
| 250939_at | 736.7 | P | 941.4 | P | 744.1 | P | 898.3 | P | iqd2 |
| 250940_at | 11.2 | A | 16.8 | A | 11.4 | A | 3.9 | A | --- |
| 250941_at | 278.6 | P | 337.3 | P | 335.9 | P | 339.2 | P | --- |
| 250942_at | 294.5 | P | 1033.1 | P | 272.9 | P | 1650.9 | P | --- |
| 250943_at | 20.1 | A | 8.9 | A | 16.5 | A | 3 | A | --- |
| 250944_at | 954.2 | P | 615.8 | P | 681 | P | 638.2 | P | --- |
| 250945_at | 1.8 | A | 6.7 | A | 2.8 | A | 0.8 | A | --- |
| 250946_at | 385 | P | 650.1 | P | 478.8 | P | 692.4 | P | --- |
| 250947_at | 162.4 | P | 173.2 | P | 173.6 | A | 183.6 | P | --- |
| 250948_at | 368.3 | P | 234.7 | P | 240.6 | P | 282.2 | P | --- |
| 250949_at | 47.2 | M | 60.3 | P | 63.1 | P | 71.6 | P | --- |
| 250950_at | 499.2 | P | 469.6 | P | 381.9 | P | 389.8 | P | ATEXO70A1 |
| 250951_at | 12.2 | A | 9.6 | A | 1.2 | A | 13.6 | A | --- |
| 250952_at | 66.9 | P | 51 | P | 110.5 | P | 78.7 | P | ATIREG2 |
| 250953_at | 227.9 | P | 226.7 | P | 197.9 | P | 236.6 | P | DPB |
| 250954_at | 1370.1 | P | 1420.7 | P | 1410.6 | P | 1584.7 | P | ATRAB8C |
| 250955_at | 141.6 | P | 171 | P | 108.1 | P | 208.9 | P | CPUORF47 |
| 250956_at | 53.3 | A | 65.2 | M | 44 | A | 74.5 | P | --- |
| 250957_at | 36.3 | A | 51.4 | P | 42.3 | A | 47.4 | P | --- |
| 250958_at | 107.7 | P | 144.9 | P | 160.2 | P | 137.3 | P | LAC11 |
| 250959_at | 31.2 | A | 17.3 | A | 43.7 | A | 29.6 | A | --- |
| 250960_at | 1981.8 | P | 1628.8 | P | 1686.7 | P | 2151.3 | P | --- |
| 250961_at | 24.4 | P | 33 | A | 40.8 | A | 22.2 | A | --- |
| 250962_at | 6.8 | A | 20.8 | A | 79.4 | A | 13.8 | A | --- |
| 250963_at | 29.6 | P | 22 | P | 36.1 | P | 18.2 | P | --- |
| 250964_at | 2.5 | A | 16.2 | A | 12.1 | A | 19.1 | A | --- |
| 250965_at | 8.2 | A | 45.2 | A | 9 | A | 16.9 | A | --- |
| 250966_at | 174.5 | P | 110.4 | P | 173 | P | 128.4 | P | --- |
| 250967_at | 5061.8 | P | 4986.6 | P | 5683.6 | P | 5683.5 | P | --- |
| 250968_at | 110.1 | P | 200.4 | P | 181.4 | P | 243.8 | P | --- |
| 250969_at | 778.4 | P | 735.9 | P | 732 | P | 729.9 | P | --- |
| 250970_at | 891.4 | P | 776.8 | P | 991.9 | P | 813.7 | P | --- |
| 250971_at | 486.8 | P | 598.3 | P | 390.9 | P | 491.7 | P | PRR7 |
| 250972_at | 799.7 | P | 1217.6 | P | 1049.5 | P | 1180.7 | P | LCL1 |
| 250973_at | 9654.9 | P | 8597.3 | P | 9162.2 | P | 9273.9 | P | --- |
| 250974_at | 197.5 | P | 164.4 | P | 217 | P | 198.9 | P | RHL2 |
| 250975_at | 84.1 | P | 179.7 | P | 78.5 | P | 149.4 | P | --- |
| 250976_at | 19.2 | A | 17.2 | P | 22.6 | A | 23.6 | P | --- |
| 250977_at | 346.4 | P | 235.4 | P | 282.5 | P | 301.9 | P | IMPA-9 |
| 250978_at | 529.6 | P | 687.8 | P | 557.2 | P | 664 | P | --- |
| 250979_at | 58.1 | A | 33 | A | 40.2 | A | 11.2 | A | --- |
| 250980_at | 36.4 | A | 33.6 | A | 54.7 | A | 31.1 | A | --- |
| 250981_at | 139 | P | 204.8 | P | 180.8 | P | 213 | P | --- |
| 250982_at | 7.1 | A | 41.9 | A | 4.3 | A | 39.1 | P | JKD |
| 250983_at | 70.8 | P | 21.8 | A | 46.9 | A | 10 | A | --- |
| 250984_at | 237.6 | P | 267.5 | P | 225.9 | P | 232.9 | P | --- |
| 250985_at | 512.4 | P | 794.2 | P | 488.6 | P | 776 | P | --- |
| 250986_at | 419.2 | P | 406.5 | P | 444.8 | P | 448.6 | P | --- |
| 250987_at | 194.9 | P | 298.6 | P | 270.8 | P | 264.1 | P | --- |
| 250988_at | 380.4 | P | 450.5 | P | 414.1 | P | 381.2 | P | UPL4 |
| 250989_at | 2.5 | A | 13.8 | A | 14 | A | 15.8 | A | CYP96A13 |
| 250990_at | 564.4 | P | 807.3 | P | 518.3 | P | 610.8 | P | NAK |
| 250991_at | 22472.7 | P | 21145.3 | P | 22829.7 | P | 22514.4 | P | MT2B |
| 250992_at | 172.8 | P | 250.4 | P | 212 | P | 317.4 | P | ATEXPA9 |
| 250993_at | 635.8 | P | 678.2 | P | 629.5 | P | 839.1 | P | --- |
| 250994_at | 844.1 | P | 1902.3 | P | 653.2 | A | 1379 | P | --- |
| 250995_at | 17622.7 | P | 18001.8 | P | 22339.2 | P | 19559.1 | P | HSC70-1 |
| 250996_at | 772.1 | P | 775.8 | P | 777 | P | 806.5 | P | --- |
| 250997_at | 27.9 | A | 29.2 | A | 26.8 | A | 44 | P | --- |
| 250998_at | 468.2 | P | 408.4 | P | 326.3 | P | 382.2 | P | --- |
| 250999_at | 301.7 | P | 231.8 | P | 273.5 | P | 260.2 | P | --- |
| 251000_at | 18.4 | A | 56 | P | 25.2 | A | 24.5 | A | --- |

| | | | | | | | | | |
|---|---|---|---|---|---|---|---|---|---|
| 251001_at | 89.1 | P | 86.7 | P | 71.6 | P | 64.6 | P | --- |
| 251002_at | 47.2 | P | 75.8 | P | 92 | P | 85.3 | P | --- |
| 251003_at | 50.1 | P | 81.5 | P | 76.4 | P | 58.9 | P | --- |
| 251004_at | 88.9 | A | 70.4 | A | 117.2 | A | 47.4 | A | --- |
| 251005_at | 623.6 | P | 441.4 | P | 514.4 | P | 493.8 | P | --- |
| 251006_at | 179.4 | P | 192.9 | P | 190.1 | P | 169.9 | P | --- |
| 251007_at | 2704.7 | P | 2200.2 | P | 3065.1 | P | 2355.7 | P | --- |
| 251008_at | 767.3 | P | 819.1 | P | 739.7 | P | 865.6 | P | --- |
| 251009_at | 13.3 | A | 3.3 | A | 27.5 | A | 34.1 | A | --- |
| 251010_at | 24.9 | A | 97.8 | A | 56 | A | 106.8 | A | --- |
| 251011_at | 164.3 | P | 286.5 | P | 111.1 | P | 206 | P | HTA12 |
| 251012_at | 173.3 | P | 150.7 | P | 103.3 | P | 157.3 | P | --- |
| 251013_at | 168.1 | P | 216.2 | P | 196.8 | P | 216.2 | P | --- |
| 251014_at | 18.6 | A | 42.2 | P | 6.8 | A | 54.2 | P | --- |
| 251015_at | 2953.4 | P | 2233.5 | P | 3123 | P | 2529.2 | P | --- |
| 251016_at | 24.6 | A | 14.9 | A | 36.1 | A | 8.3 | A | --- |
| 251017_at | 69.1 | P | 327.6 | P | 61.9 | P | 194.8 | P | --- |
| 251018_at | 4745.1 | P | 3627.9 | P | 4573.8 | P | 4051.8 | P | --- |
| 251019_at | 50.8 | A | 67.4 | P | 62 | A | 86.3 | M | --- |
| 251020_at | 688.3 | P | 894.4 | P | 924.8 | P | 1013.4 | P | NAP9 |
| 251021_at | 8.7 | A | 3.8 | A | 33.8 | A | 3 | A | --- |
| 251022_at | 409.5 | P | 394.5 | P | 437.2 | P | 329.2 | P | --- |
| 251023_at | 22.5 | P | 39.6 | P | 65 | P | 33.6 | P | --- |
| 251024_at | 141 | P | 229.7 | P | 175.6 | P | 239.7 | P | --- |
| 251025_at | 34.7 | A | 46.1 | P | 102.2 | A | 61.4 | P | PCS1 |
| 251026_at | 45.5 | A | 65.4 | P | 51.3 | A | 54.2 | A | FHL |
| 251027_at | 7.1 | A | 26.7 | A | 36.3 | A | 9.7 | A | --- |
| 251028_at | 317.1 | P | 391.2 | P | 214.6 | P | 473.1 | P | --- |
| 251029_at | 843 | P | 706.4 | P | 768.8 | P | 669.5 | P | --- |
| 251030_at | 239.3 | P | 199.1 | P | 278.2 | P | 229.9 | P | NDP1 |
| 251031_at | 2483.8 | P | 3452.5 | P | 2820 | P | 3369.2 | P | OHP |
| 251032_at | 388.2 | P | 376.4 | P | 393.9 | P | 415.4 | P | RPL |
| 251033_at | 335 | P | 333.8 | P | 378.3 | P | 331.3 | P | --- |
| 251034_at | 330.6 | P | 376.5 | P | 353 | P | 318.8 | P | PRA1.A1 |
| 251035_at | 30 | A | 6.4 | A | 6.2 | A | 29.3 | A | --- |
| 251036_at | 1255.6 | P | 2156.1 | P | 1412.5 | P | 1689.4 | P | --- |
| 251037_at | 308.2 | P | 276.4 | P | 301.2 | P | 242.2 | P | UNE18 |
| 251038_at | 5897.3 | P | 6082.8 | P | 5332 | P | 6350.4 | P | --- |
| 251039_at | 31.1 | A | 60.2 | A | 25.9 | A | 56.7 | A | --- |
| 251040_at | 622.6 | P | 713.6 | P | 695.1 | P | 772.7 | P | EMB2730 |
| 251041_at | 56 | A | 66.2 | P | 48.4 | A | 65.3 | P | PRT6 |
| 251042_at | 595.4 | P | 473.5 | P | 539 | P | 404.9 | P | PRT6 |
| 251043_s_at | 1.2 | A | 0.5 | A | 25.8 | A | 10.7 | A | --- |
| 251044_at | 1.4 | A | 2.2 | A | 1.6 | A | 24.8 | A | --- |
| 251045_s_at | 0.2 | A | 0.4 | A | 12.5 | A | 0.4 | A | --- |
| 251046_at | 47 | A | 106.4 | P | 61.5 | P | 78.1 | P | --- |
| 251047_at | 14.2 | A | 48.7 | A | 24.2 | A | 5.4 | A | --- |
| 251048_at | 296.1 | P | 405.7 | P | 405.1 | P | 370.3 | P | --- |
| 251049_at | 52.4 | P | 72.7 | P | 86.6 | P | 54.6 | P | --- |
| 251050_at | 42.6 | P | 73.4 | P | 54.4 | P | 51.1 | P | --- |
| 251051_at | 94.5 | A | 90.2 | A | 112.4 | A | 52 | A | --- |
| 251052_at | 207 | P | 244.7 | P | 243.1 | P | 221.9 | P | DPA |
| 251053_at | 9.2 | A | 28.5 | A | 40.1 | A | 58.1 | A | CAX4 |
| 251054_at | 175.1 | P | 333.8 | P | 176.8 | P | 305.6 | P | LECRKA4.1 |
| 251055_at | 725.2 | P | 721.8 | P | 758.8 | P | 716.8 | P | --- |
| 251056_at | 107.6 | P | 130.1 | P | 133.9 | P | 129.1 | P | RAPTOR2 |
| 251057_at | 249.9 | P | 210.7 | P | 283.5 | P | 264 | P | --- |
| 251058_at | 390 | P | 439.1 | P | 342 | P | 462.6 | P | --- |
| 251059_at | 669.1 | P | 493.4 | P | 557.2 | P | 582.4 | P | CIPK15 |
| 251060_at | 189.2 | P | 443.8 | P | 156.9 | P | 301 | P | ATSR1 |
| 251061_at | 55.5 | A | 31.7 | A | 6.2 | A | 34.5 | A | --- |
| 251062_at | 58.9 | P | 94.4 | P | 65.2 | P | 88.1 | P | OFP1 |
| 251063_at | 432.2 | P | 270.9 | P | 383.7 | P | 243.9 | P | --- |
| 251064_at | 25.6 | A | 18.1 | A | 32.9 | A | 19 | A | --- |
| 251065_at | 185.4 | P | 256 | P | 177.8 | P | 251.1 | P | --- |
| 251066_at | 66.1 | P | 69.1 | A | 57.1 | A | 90.3 | P | --- |
| 251067_at | 105.2 | P | 135.4 | P | 139.5 | P | 154.9 | P | --- |
| 251068_at | 852.5 | P | 949 | P | 984.6 | P | 864 | P | STN8 |
| 251069_at | 85.5 | P | 172.7 | P | 172.4 | P | 161 | P | --- |
| 251070_at | 98.9 | P | 90.3 | P | 117.7 | P | 84.5 | P | --- |
| 251071_at | 198.1 | P | 215.8 | P | 171 | P | 188.9 | P | --- |
| 251072_at | 1418 | P | 538.8 | P | 1353.1 | P | 772.8 | P | --- |
| 251073_at | 3734.2 | P | 3955.3 | P | 4299.5 | P | 3980.8 | P | --- |
| 251074_at | 994.5 | P | 1063.1 | P | 817.3 | P | 937.4 | P | --- |
| 251075_at | 260.7 | P | 302.8 | P | 404.5 | P | 356.2 | P | --- |
| 251076_at | 157.8 | P | 187.7 | P | 186.5 | P | 145.7 | P | --- |
| 251077_at | 463.7 | P | 528.8 | P | 537.3 | P | 519.6 | P | --- |
| 251078_at | 362.8 | P | 311 | P | 214.7 | P | 251.8 | P | --- |
| 251079_at | 40.2 | A | 7.4 | A | 33.4 | A | 4.4 | A | --- |
| 251080_at | 127.9 | P | 145.1 | P | 161.2 | P | 96.4 | P | ROPGEF7 |
| 251081_at | 3 | A | 3.4 | A | 15.7 | A | 1.2 | A | --- |
| 251082_at | 14491.8 | P | 16856.2 | P | 16328.4 | P | 16196.7 | P | --- |
| 251083_at | 585.4 | P | 583.6 | P | 582.4 | P | 572.3 | P | --- |
| 251084_at | 104.3 | P | 161.7 | P | 80.9 | P | 129.5 | P | --- |
| 251085_at | 1.1 | A | 8.4 | A | 3.3 | A | 1.1 | A | --- |
| 251086_at | 202.1 | P | 156.4 | P | 221 | P | 183.1 | P | --- |
| 251087_at | 342.7 | P | 438.9 | P | 439.1 | P | 402.1 | P | --- |
| 251088_at | 4.3 | A | 39.2 | A | 16.1 | A | 21.5 | A | --- |
| 251089_at | 251.2 | P | 288.5 | P | 282.3 | P | 325.6 | P | --- |
| 251090_at | 436.9 | P | 289.5 | P | 458.6 | P | 445.1 | P | --- |
| 251091_at | 3953 | P | 4827 | P | 5674 | P | 5193.1 | P | RSR4 |
| 251092_at | 179.1 | P | 167.6 | P | 177.4 | P | 167.1 | P | --- |
| 251093_at | 44 | A | 78 | P | 100.1 | M | 77.6 | P | --- |
| 251094_at | 1433 | P | 1008.7 | P | 1339.9 | P | 1005.8 | P | --- |
| 251095_at | 210.6 | P | 214.6 | P | 186.5 | P | 200.4 | P | --- |
| 251096_at | 46.9 | A | 42.6 | M | 78.8 | A | 54.5 | A | LECRKA4.2 |
| 251097_at | 1.3 | A | 16.8 | A | 5.4 | A | 16.6 | A | LECRKA4.3 |
| 251098_at | 1657.3 | P | 1772 | P | 1501.6 | P | 1682 | P | --- |
| 251099_at | 44.2 | A | 59.7 | P | 69.9 | P | 60.8 | P | --- |
| 251100_at | 217.9 | P | 182.7 | P | 436.9 | P | 183.6 | P | --- |
| 251101_at | 1.2 | A | 3.9 | A | 22.3 | A | 2.8 | A | CHX26 |
| 251102_at | 45.4 | A | 12.1 | A | 29.4 | A | 18.4 | A | CHX27 |
| 251103_at | 31.9 | A | 34.8 | A | 44.7 | A | 29.2 | A | --- |
| 251104_at | 325.8 | P | 442.7 | P | 247.3 | P | 384.6 | P | --- |
| 251105_at | 69.8 | P | 101.1 | P | 70.3 | P | 91.8 | P | ATSCAR4 |
| 251106_at | 1063.6 | P | 731.6 | P | 1288.2 | P | 828.8 | P | --- |
| 251107_at | 33.8 | P | 75.2 | P | 63.5 | A | 57.2 | P | --- |
| 251108_at | 80 | P | 129 | P | 81.4 | P | 112 | P | --- |
| 251109_at | 4901.9 | P | 3020.7 | P | 5989.7 | P | 4313 | P | FER1 |

| | | | | | | | | | |
|---|---|---|---|---|---|---|---|---|---|
| 251110_at | 321.4 | P | 289.8 | P | 331.2 | P | 323 | P | --- |
| 251111_at | 104 | P | 52.9 | P | 52.4 | P | 54.7 | P | --- |
| 251112_s_at | 179.4 | P | 144 | P | 169.4 | P | 133 | P | PDC3 |
| 251113_at | 127.1 | P | 170.2 | P | 170 | P | 147.5 | P | --- |
| 251114_at | 33 | M | 20.8 | M | 29.1 | A | 28.2 | A | --- |
| 251115_at | 355.8 | P | 331.8 | P | 410.2 | P | 392.9 | P | ESP4 |
| 251116_at | 2.8 | A | 16.7 | A | 8.5 | A | 8.1 | A | scpl40 |
| 251117_at | 519.6 | P | 383.4 | P | 530.7 | P | 399.8 | P | --- |
| 251118_at | 6738.9 | P | 6867.9 | P | 7744.5 | P | 7165.8 | P | APG1 |
| 251119_at | 184.6 | P | 344.3 | P | 313.4 | P | 318 | P | --- |
| 251120_at | 4776.8 | P | 5709.8 | P | 4566.4 | P | 4950.6 | P | --- |
| 251121_at | 463 | P | 528.8 | P | 322.9 | P | 437.4 | P | AGG1 |
| 251122_at | 968.5 | P | 1180.3 | P | 1066.9 | P | 1160.3 | P | --- |
| 251123_at | 279.6 | P | 299.7 | P | 276.6 | P | 297.8 | P | --- |
| 251124_s_at | 128.8 | P | 99.9 | P | 91.7 | P | 115.8 | P | LAC8 |
| 251125_at | 4.5 | A | 43.1 | A | 37.7 | A | 23.2 | A | --- |
| 251126_at | 30.2 | A | 37.1 | A | 34.2 | A | 11.9 | A | --- |
| 251127_at | 11.7 | A | 8.6 | A | 5.4 | A | 4.1 | A | --- |
| 251128_at | 31.1 | A | 35.7 | A | 55.3 | A | 58.9 | A | --- |
| 251129_at | 13.9 | A | 30.3 | A | 7.1 | A | 21.9 | A | --- |
| 251130_at | 75.3 | P | 86.2 | P | 92.5 | P | 78.9 | P | PTR5 |
| 251131_at | 14.8 | A | 40 | P | 29.9 | A | 34.3 | M | LAC10 |
| 251132_at | 72.2 | M | 53.9 | P | 87.7 | P | 82.5 | P | --- |
| 251133_at | 665.8 | P | 649.8 | P | 679.6 | P | 719.6 | P | --- |
| 251134_at | 293.4 | P | 343.9 | P | 360.7 | P | 352.3 | P | AtQSOX2 |
| 251135_at | 2.1 | A | 2.9 | A | 6.5 | A | 2.7 | A | --- |
| 251136_at | 374.5 | P | 285.5 | P | 335.3 | P | 323.1 | P | --- |
| 251137_at | 49.6 | P | 39.5 | A | 25.7 | A | 56.3 | P | --- |
| 251138_at | 227.9 | P | 283.8 | P | 282 | P | 271.9 | P | --- |
| 251139_at | 395.3 | P | 359.9 | P | 381.4 | P | 309.9 | P | --- |
| 251140_at | 815.8 | P | 845 | P | 820.9 | P | 1105.5 | P | --- |
| 251141_at | 141.8 | P | 90.9 | P | 139.6 | P | 113.3 | P | --- |
| 251142_at | 253.1 | P | 621.8 | P | 346.8 | P | 561.3 | P | --- |
| 251143_at | 752.8 | P | 712.3 | P | 766.9 | P | 815 | P | SQD2 |
| 251144_at | 409.8 | P | 669.1 | P | 480.9 | P | 497.5 | P | --- |
| 251145_at | 648.3 | P | 598.4 | P | 665.2 | P | 543.4 | P | --- |
| 251146_at | 3772 | P | 3537.9 | P | 3425.6 | P | 3202.6 | P | CCD1 |
| 251147_at | 81 | P | 105 | P | 58.2 | A | 90.3 | P | --- |
| 251148_at | 302.4 | P | 259.5 | P | 369.2 | P | 240.2 | P | TKL |
| 251149_at | 984.9 | P | 828.4 | P | 691.4 | P | 807.8 | P | --- |
| 251150_at | 299.7 | P | 448.9 | P | 391.4 | P | 464.5 | P | CYCP1;1 |
| 251151_at | 1104.1 | P | 1200 | P | 1142.4 | P | 1227.2 | P | --- |
| 251152_at | 236.2 | P | 264.7 | P | 356.6 | P | 355 | P | RANGAP1 |
| 251153_at | 872.8 | P | 907.4 | P | 769.9 | P | 853 | P | NPL41 |
| 251154_at | 56.1 | A | 143.4 | P | 132 | A | 152.9 | P | ATIPT3 |
| 251155_at | 3858.3 | P | 5649.3 | P | 4302.6 | P | 6964 | P | --- |
| 251156_at | 49.6 | A | 29.5 | A | 48.9 | A | 28.8 | A | EDL3 |
| 251157_at | 3342 | P | 3962.1 | P | 3650.7 | P | 3831.4 | P | CSP41A |
| 251158_at | 696.6 | P | 608.1 | P | 616.2 | P | 616.4 | P | MIRO2 |
| 251159_at | 10.9 | A | 20.2 | A | 14 | A | 4.4 | A | --- |
| 251160_at | 34.9 | A | 57.7 | P | 33.8 | A | 46.2 | A | --- |
| 251161_at | 94.6 | P | 137 | P | 156.7 | P | 158.1 | P | --- |
| 251162_at | 40 | A | 71.4 | P | 65.9 | P | 95.6 | P | --- |
| 251163_at | 1526.1 | P | 1449.8 | P | 1755.2 | P | 1302.2 | P | --- |
| 251164_s_at | 60.1 | A | 45.5 | A | 34.3 | A | 43.3 | A | --- |
| 251165_at | 154 | P | 317.6 | P | 103.2 | P | 287.2 | P | --- |
| 251166_at | 3.8 | A | 6.4 | A | 6 | A | 5.2 | A | AT-HSFA7B |
| 251167_at | 3.4 | A | 2.8 | A | 4.3 | A | 15.2 | A | --- |
| 251168_at | 42.8 | P | 38.1 | P | 63.2 | A | 46.2 | A | --- |
| 251169_at | 634.3 | P | 562.9 | P | 424.9 | P | 516.9 | P | MARD1 |
| 251170_at | 712.7 | P | 700.5 | P | 590.1 | P | 610.9 | P | ATMRK1 |
| 251171_at | 213.4 | P | 212.3 | P | 244.3 | P | 224.9 | P | --- |
| 251172_at | 1950.6 | P | 2226 | P | 2555.4 | P | 2060.8 | P | RRF |
| 251173_at | 110.6 | P | 136.6 | P | 159.2 | P | 146 | P | --- |
| 251174_at | 467.1 | P | 561.5 | P | 599.7 | P | 631.3 | P | PLP9 |
| 251175_at | 246.2 | P | 364.7 | P | 371.4 | P | 365.6 | P | HMT2 |
| 251176_at | 90.3 | P | 143 | P | 29.5 | A | 45 | P | --- |
| 251177_at | 371.4 | P | 412.4 | P | 424.1 | P | 478.6 | P | --- |
| 251178_at | 9.9 | A | 65.5 | A | 73 | A | 76.9 | A | CKX6 |
| 251179_at | 627.6 | P | 683.2 | P | 749.1 | P | 590.1 | P | --- |
| 251180_at | 9.2 | A | 23.6 | A | 3.9 | A | 2 | A | --- |
| 251181_at | 916.6 | P | 944 | P | 859.4 | P | 1036.1 | P | --- |
| 251182_at | 1977.1 | P | 2302.1 | P | 1966.2 | P | 2367 | P | ATERDJ3B |
| 251183_at | 309 | P | 629.7 | P | 312 | P | 668.9 | P | --- |
| 251184_at | 2482.7 | P | 1956.8 | P | 1767.6 | P | 2057.3 | P | --- |
| 251185_at | 10182.9 | P | 8097.9 | P | 11868.3 | P | 9080.6 | P | --- |
| 251186_at | 997 | P | 1160.8 | P | 922.4 | P | 1057.8 | P | --- |
| 251187_at | 455.5 | P | 430.4 | P | 388.3 | P | 362.1 | P | AtATG18a |
| 251188_at | 29.7 | A | 46.3 | P | 46.1 | P | 38.9 | P | --- |
| 251189_at | 1396.7 | P | 1456.6 | P | 1517 | P | 1386.2 | P | --- |
| 251190_at | 55.1 | A | 154.7 | P | 61.9 | A | 98.8 | P | ATL5 |
| 251191_at | 216.6 | P | 141.6 | P | 177.5 | P | 144.6 | P | --- |
| 251192_at | 514.9 | P | 360.9 | P | 384 | P | 428 | P | XT1 |
| 251193_at | 698.4 | P | 790.9 | P | 745.7 | P | 713.9 | P | APG3 |
| 251194_at | 182 | P | 139.3 | P | 130.4 | P | 138.5 | P | --- |
| 251195_at | 18.1 | A | 31.8 | A | 37.9 | A | 31.7 | P | --- |
| 251196_at | 0.8 | A | 1.3 | A | 2 | A | 0.8 | A | --- |
| 251197_at | 39 | A | 181 | P | 95.6 | A | 133.5 | P | --- |
| 251198_at | 670.6 | P | 586 | P | 635.3 | P | 645.9 | P | --- |
| 251199_at | 518.9 | P | 698.9 | P | 434 | P | 748.6 | P | TIR1 |
| 251200_at | 452.6 | P | 303.1 | P | 357.7 | P | 299.9 | P | GID1B |
| 251201_at | 19.6 | A | 1.8 | A | 4.7 | A | 3.4 | A | --- |
| 251202_at | 2.8 | A | 38.3 | A | 25.1 | A | 24.5 | A | --- |
| 251203_at | 245.5 | P | 209.9 | P | 287.5 | P | 243.9 | P | --- |
| 251204_at | 3.6 | A | 4.5 | A | 12 | A | 4 | A | AT3G63095 |
| 251205_at | 823.3 | P | 897.5 | P | 772.1 | P | 764.6 | P | ATGPX5 |
| 251206_at | 189.1 | P | 126.9 | P | 332.8 | P | 243.1 | P | --- |
| 251207_at | 34 | A | 44.7 | A | 39.4 | M | 30.2 | A | --- |
| 251208_at | 318.3 | P | 313.6 | P | 342.2 | P | 299.1 | P | ATOEP16-4 |
| 251209_at | 65 | P | 48 | P | 99.5 | P | 72 | P | --- |
| 251210_at | 568.5 | P | 465.9 | P | 576.4 | P | 446.8 | P | --- |
| 251211_s_at | 251 | P | 191.2 | P | 210.8 | P | 259.1 | P | --- |
| 251212_at | 1.4 | A | 0.8 | A | 1.8 | A | 2.1 | A | --- |
| 251213_at | 1.3 | A | 0.5 | A | 32.2 | A | 10.2 | A | --- |
| 251214_at | 22.3 | A | 31.4 | A | 31.5 | A | 17.5 | A | --- |
| 251215_at | 51.1 | P | 54 | P | 49 | P | 36.5 | P | --- |
| 251216_at | 21.6 | A | 34.1 | A | 39.5 | A | 36.8 | A | --- |
| 251217_at | 1023.2 | P | 1074.7 | P | 1180.5 | P | 1163.9 | P | --- |
| 251218_at | 3892 | P | 5602.2 | P | 4516.2 | P | 5478.4 | P | CP12-2 |

| | | | | | | | | | |
|---|---|---|---|---|---|---|---|---|---|
| 251219_at | 82.2 | M | 145.6 | P | 88.1 | A | 142.2 | P | --- |
| 251220_at | 2072.7 | P | 1840 | P | 1702.6 | P | 1820.2 | P | --- |
| 251221_at | 3865.9 | P | 3895.8 | P | 3912.3 | P | 4070.5 | P | --- |
| 251222_at | 2474.3 | P | 2082.4 | P | 2181.5 | P | 2053 | P | --- |
| 251223_at | 33.8 | A | 46.8 | A | 36.4 | A | 41.6 | A | ATMYB11 |
| 251224_at | 203.7 | P | 204.5 | P | 242.1 | P | 212.6 | P | --- |
| 251225_at | 597.2 | P | 565.2 | P | 462.9 | P | 553.3 | P | GATL7 |
| 251226_at | 2.8 | A | 3.1 | A | 3.1 | A | 2 | A | PRP3 |
| 251227_at | 1818.3 | P | 1785.4 | P | 1654.3 | P | 1946.9 | P | ATMRP10 |
| 251228_at | 5 | A | 11.6 | A | 2.6 | A | 12.3 | A | --- |
| 251229_at | 24.5 | P | 18.6 | A | 46.7 | A | 39.9 | P | BGLU7 |
| 251230_at | 505.2 | P | 642.2 | P | 604.5 | P | 640.8 | P | BGLU8 |
| 251231_at | 32.9 | M | 35.3 | A | 22.3 | A | 19.3 | A | ATGSTF13 |
| 251232_at | 56.7 | A | 67.6 | A | 46.6 | A | 57.2 | A | --- |
| 251233_at | 288.9 | P | 313.6 | P | 282.6 | P | 340.7 | P | DRB4 |
| 251234_s_at | 1820.6 | P | 1130.1 | P | 2081.2 | P | 1165.1 | P | UXS4 |
| 251235_at | 467.2 | P | 301.3 | P | 398.5 | P | 280.1 | P | --- |
| 251236_at | 17.8 | A | 25 | A | 27.8 | A | 18.1 | A | --- |
| 251237_at | 851 | P | 946.8 | P | 897.9 | P | 882.9 | P | CPuORF3 |
| 251238_at | 63 | P | 46.1 | P | 46.8 | P | 41.6 | P | --- |
| 251239_at | 38.4 | A | 3.3 | A | 51.9 | A | 3.1 | A | --- |
| 251240_at | 350.1 | P | 424.4 | P | 305.1 | P | 372.6 | P | --- |
| 251241_s_at | 4671.4 | P | 4975.6 | P | 5212.4 | P | 5007.2 | P | --- |
| 251242_at | 1629.9 | P | 1444.2 | P | 1521.7 | P | 1667.2 | P | RSP31 |
| 251243_at | 3679 | P | 3674.1 | P | 2878.8 | P | 3839 | P | --- |
| 251244_at | 196.2 | P | 281.3 | P | 235.3 | P | 251.3 | P | --- |
| 251245_at | 35.5 | A | 38.2 | A | 20.7 | A | 23.3 | A | PIL2 |
| 251246_at | 40.4 | M | 27.2 | A | 59.4 | A | 23.5 | A | IAA30 |
| 251247_at | 241.1 | P | 214.2 | P | 225.3 | P | 212.8 | P | --- |
| 251248_at | 288 | P | 314.8 | P | 234.5 | P | 277.3 | P | PGP21 |
| 251249_at | 87 | P | 90.3 | P | 88.7 | P | 126.6 | P | --- |
| 251250_at | 9.3 | A | 40.7 | P | 9.9 | A | 21.4 | A | --- |
| 251251_at | 170.7 | P | 220.1 | P | 255.2 | P | 245.7 | P | --- |
| 251252_at | 36.6 | A | 13.1 | A | 16.9 | A | 3.4 | A | --- |
| 251253_at | 491.9 | P | 476.1 | P | 348.6 | P | 491.8 | P | --- |
| 251254_at | 199.5 | P | 269.8 | P | 160.6 | P | 271.5 | P | --- |
| 251255_at | 40.3 | A | 41.5 | M | 43 | A | 62.9 | P | --- |
| 251256_at | 121.3 | P | 140.8 | P | 181.3 | P | 150.2 | P | --- |
| 251257_at | 9796.9 | P | 10339.7 | P | 12671.1 | P | 10168.3 | P | ATARFA1E |
| 251258_at | 4.8 | A | 3.3 | A | 9 | A | 6.9 | A | VGDH2 |
| 251259_at | 137.1 | P | 128.4 | P | 180 | P | 148.1 | P | --- |
| 251260_at | 508.1 | P | 441.7 | P | 620.6 | P | 395.2 | P | --- |
| 251261_at | 466.2 | P | 692.2 | P | 570.8 | P | 716.4 | P | --- |
| 251262_at | 115.6 | P | 105.8 | P | 102 | P | 109.2 | P | --- |
| 251263_at | 828.8 | P | 873.4 | P | 817.8 | P | 685.7 | P | --- |
| 251264_at | 1762.4 | P | 1970.8 | P | 2102 | P | 2010.7 | P | --- |
| 251265_at | 620.4 | P | 516.1 | P | 641.7 | P | 530 | P | --- |
| 251266_at | 4.4 | A | 11.7 | A | 49.6 | A | 16.2 | A | --- |
| 251267_at | 393.7 | P | 343.1 | P | 391.1 | P | 360.6 | P | --- |
| 251268_at | 18.9 | P | 16.4 | A | 21.2 | P | 4.3 | A | --- |
| 251269_at | 832.3 | P | 1012.5 | P | 872 | P | 1054.8 | P | --- |
| 251270_at | 255.4 | P | 355.8 | P | 288.5 | P | 350.8 | P | --- |
| 251271_at | 138.2 | P | 77.7 | A | 160.3 | P | 64.2 | M | --- |
| 251272_at | 327.6 | P | 267.3 | P | 210.4 | P | 258.7 | P | ATHB-12 |
| 251273_at | 613.6 | P | 609.3 | P | 436.7 | P | 581.9 | P | --- |
| 251274_at | 453.7 | P | 425.8 | P | 603.6 | P | 543.9 | P | --- |
| 251275_at | 8.5 | A | 3.1 | A | 4.9 | A | 4.6 | A | --- |
| 251276_at | 79.5 | P | 68.2 | P | 88 | P | 80.2 | P | SDG14 |
| 251277_at | 34.8 | A | 76.1 | P | 27.3 | A | 42.5 | A | ADL1B |
| 251278_at | 222.5 | P | 299.3 | P | 236 | P | 357.6 | P | emb1703 |
| 251279_at | 196.9 | P | 199.3 | P | 225 | P | 172.6 | P | --- |
| 251280_at | 3.6 | A | 36.2 | A | 5.7 | A | 27.4 | A | --- |
| 251281_at | 500 | P | 441 | P | 509.1 | P | 437.7 | P | AGP20 |
| 251282_at | 640.4 | P | 172.2 | P | 448 | P | 142.7 | P | CRF6 |
| 251283_at | 374.8 | P | 408.9 | P | 262.2 | P | 324.8 | P | --- |
| 251284_at | 66.3 | P | 68 | P | 71 | M | 83.9 | P | --- |
| 251285_at | 524.1 | P | 639 | P | 542 | P | 602.7 | P | --- |
| 251286_at | 392.8 | P | 409.5 | P | 374.2 | P | 436.8 | P | --- |
| 251287_at | 918.4 | P | 1534.5 | P | 1001.3 | P | 1581.1 | P | --- |
| 251288_at | 804.2 | P | 591.3 | P | 872.4 | P | 598.1 | P | RRP41 |
| 251289_at | 150.7 | P | 200.2 | P | 260.2 | P | 240.8 | P | ARF18 |
| 251290_at | 82.2 | A | 90 | A | 75.9 | A | 85.1 | P | DAG1 |
| 251291_at | 91.7 | P | 129.8 | P | 87.9 | P | 96.8 | P | --- |
| 251292_at | 58.1 | A | 56.6 | A | 58.7 | P | 47.3 | A | --- |
| 251293_at | 238 | P | 1.3 | A | 90.1 | A | 23.8 | A | --- |
| 251294_at | 8 | A | 9.8 | A | 1.5 | A | 2.5 | A | MTPA1 |
| 251295_at | 264.2 | P | 298.1 | P | 332.8 | P | 332.3 | P | --- |
| 251296_at | 560.6 | P | 643 | P | 611.4 | P | 579.8 | P | --- |
| 251297_at | 145.3 | P | 192.4 | P | 217.4 | P | 209.8 | P | GLP10 |
| 251298_at | 36.7 | A | 90.6 | P | 114.5 | A | 97.1 | A | --- |
| 251299_at | 59.1 | P | 76.1 | P | 64.1 | P | 79.8 | P | --- |
| 251300_at | 405.3 | P | 408.3 | P | 466.4 | P | 430.9 | P | --- |
| 251301_at | 90.6 | A | 55 | A | 44.1 | A | 76.6 | A | CYP78A9 |
| 251302_at | 195.5 | P | 129.9 | P | 191.7 | P | 163.9 | P | NGA2 |
| 251303_at | 27.8 | A | 3.6 | A | 13.2 | A | 26.5 | A | ANAC066 |
| 251304_at | 186.3 | P | 323.7 | P | 256.4 | P | 336.6 | P | --- |
| 251305_at | 10804.4 | P | 11478.3 | P | 11463.3 | P | 11961.4 | P | ROC4 |
| 251306_at | 1146 | P | 1091.7 | P | 913.8 | P | 1041 | P | --- |
| 251307_at | 2412.2 | P | 2124.7 | P | 1952.3 | P | 1760.9 | P | --- |
| 251308_at | 543.2 | P | 669 | P | 470.7 | P | 686.9 | P | GAUT1 |
| 251309_at | 1901.5 | P | 2384.5 | P | 2334.1 | P | 2411.7 | P | --- |
| 251310_at | 409.7 | P | 514.1 | P | 359.4 | P | 437.2 | P | HDG1 |
| 251311_at | 828.7 | P | 843.9 | P | 883.9 | P | 802.1 | P | FUS6 |
| 251312_at | 18.3 | A | 40.2 | P | 42.3 | P | 28.2 | A | AGL13 |
| 251313_at | 104.2 | P | 113.1 | P | 113.5 | P | 107.7 | P | --- |
| 251314_at | 578 | P | 511.3 | P | 523.6 | P | 496.2 | P | --- |
| 251315_at | 2 | A | 1.7 | A | 2 | A | 3.3 | A | --- |
| 251316_s_at | 1036.6 | P | 1068.6 | P | 862 | P | 961.2 | P | --- |
| 251317_at | 88.1 | P | 244.4 | P | 138.8 | P | 232.4 | P | --- |
| 251318_at | 246.1 | P | 326.1 | P | 200.9 | P | 302.3 | P | GDAP1 |
| 251319_at | 126.2 | P | 216.2 | P | 114.7 | P | 203.9 | P | --- |
| 251320_at | 190.9 | P | 125.9 | P | 184.8 | P | 141.9 | P | PANB2 |
| 251321_at | 1406.4 | P | 1636.7 | P | 1312.1 | P | 1413.9 | P | BRH1 |
| 251322_at | 12819.8 | P | 12004.8 | P | 14142.4 | P | 11416 | P | CYSC1 |
| 251323_at | 1917.9 | P | 2337.1 | P | 1495.8 | P | 2028.1 | P | --- |
| 251324_at | 2395.9 | P | 2570.3 | P | 2539.6 | P | 2226.5 | P | PIP1A |
| 251325_s_at | 16376.6 | P | 16854.9 | P | 15207 | P | 16604.4 | P | LHCA2 |
| 251326_at | 158.7 | P | 194.6 | P | 206.8 | P | 184.2 | P | HWS |
| 251327_at | 434.1 | P | 593.7 | P | 543 | P | 536.6 | P | --- |

| | | | | | | | | | |
|---|---|---|---|---|---|---|---|---|---|
| 251328_at | 925.6 | P | 860.6 | P | 780.6 | P | 919.1 | P | ATPOB1 |
| 251329_at | 10.2 | A | 3.6 | A | 6.9 | A | 8.6 | A | SYP73 |
| 251330_at | 280.3 | P | 294.5 | P | 372.4 | P | 373.2 | P | --- |
| 251331_s_at | 502.6 | P | 677.8 | P | 767.3 | P | 649.1 | P | TUBG1 |
| 251332_at | 299.8 | P | 299.8 | P | 310.1 | P | 272.6 | P | --- |
| 251333_at | 442.5 | P | 386 | P | 328.8 | P | 522.3 | P | --- |
| 251334_at | 43.9 | P | 35.2 | P | 77.9 | P | 40 | P | --- |
| 251335_at | 52.4 | M | 16.6 | A | 50.6 | A | 11.4 | A | --- |
| 251336_at | 108.7 | P | 101.9 | P | 108.2 | P | 48.3 | P | BAP1 |
| 251337_at | 3146.2 | P | 3504.4 | P | 3477.1 | P | 2997.2 | P | PBF1 |
| 251338_at | 1930.1 | P | 1984.2 | P | 2006.8 | P | 1851.6 | P | VAP |
| 251339_at | 4.1 | A | 4.2 | A | 7.7 | A | 14.2 | A | --- |
| 251340_at | 351.4 | P | 363.4 | P | 424.7 | P | 365.5 | P | ATARP7 |
| 251341_at | 7150.9 | P | 6331.8 | P | 8262.6 | P | 6857.2 | P | --- |
| 251342_at | 493.4 | P | 630.1 | P | 354.4 | P | 537.6 | P | --- |
| 251343_at | 262.7 | P | 297.6 | P | 260.2 | P | 370.2 | P | --- |
| 251344_at | 0.7 | A | 2.5 | A | 4.8 | A | 1.8 | A | --- |
| 251345_at | 39.4 | P | 23.4 | A | 26.8 | A | 5.3 | A | --- |
| 251346_at | 767.4 | P | 523.9 | P | 854.5 | P | 555.9 | P | --- |
| 251347_at | 243.6 | P | 184.8 | P | 155.5 | P | 221.6 | P | --- |
| 251348_at | 76 | P | 107.1 | P | 104.2 | P | 91.9 | P | --- |
| 251349_s_at | 115.7 | P | 129.3 | P | 110.3 | A | 113.9 | P | --- |
| 251350_at | 23.4 | A | 3.6 | A | 9.5 | A | 14.9 | A | CYP76C7 |
| 251351_s_at | 1165.8 | P | 1471.8 | P | 1229.9 | P | 1308.6 | P | NTMC2T4 |
| 251352_at | 1673.7 | P | 1414.7 | P | 1641 | P | 1493.1 | P | PEX11E |
| 251353_at | 678.2 | P | 638.7 | P | 524.2 | P | 711.4 | P | --- |
| 251354_at | 14.4 | A | 5.3 | A | 31 | A | 13.6 | A | --- |
| 251355_at | 767 | P | 397.9 | P | 733.4 | P | 480.6 | P | --- |
| 251356_at | 265.5 | P | 308 | P | 172.8 | P | 262.5 | P | AtPP2-A13 |
| 251357_at | 10143.1 | P | 8800.9 | P | 8867.3 | P | 9671.6 | P | ARS27A |
| 251358_at | 137.6 | P | 125 | P | 180.7 | P | 138.9 | P | --- |
| 251359_at | 50.8 | P | 29 | M | 134.8 | P | 45.4 | P | --- |
| 251360_at | 209 | P | 268.5 | P | 342.4 | P | 243.6 | P | --- |
| 251361_at | 22.6 | A | 34 | A | 7.3 | A | 51.7 | A | --- |
| 251362_at | 835 | P | 848.7 | P | 925.7 | P | 940.8 | P | --- |
| 251363_at | 99.2 | P | 99.7 | P | 105.8 | P | 145 | P | AtMYB17 |
| 251364_at | 26.2 | A | 5.3 | A | 6.4 | A | 5.4 | A | --- |
| 251365_at | 301.1 | P | 470.5 | P | 510.7 | P | 505.2 | P | --- |
| 251366_at | 19.4 | P | 22.2 | A | 25 | A | 23 | A | --- |
| 251367_at | 127.8 | P | 128.2 | P | 108.2 | P | 109.7 | P | SKIP4 |
| 251368_at | 106.7 | P | 110.8 | P | 141 | P | 116.6 | P | --- |
| 251369_at | 287.6 | P | 324.3 | P | 323.7 | P | 308.5 | P | --- |
| 251370_at | 1731.6 | P | 1290.7 | P | 1426 | P | 1206.6 | P | --- |
| 251371_at | 775.1 | P | 426 | P | 686.4 | P | 500 | P | EDA14 |
| 251372_at | 147.3 | P | 159.6 | P | 126.8 | P | 135.2 | P | --- |
| 251373_at | 504.1 | P | 620.9 | P | 440 | P | 464.6 | P | --- |
| 251374_at | 22.9 | A | 63.5 | M | 53.5 | A | 91.3 | P | HAT3 |
| 251375_at | 135.3 | P | 160.9 | P | 165.6 | P | 164 | P | --- |
| 251376_at | 353.2 | P | 390.8 | P | 422.5 | P | 363.4 | P | --- |
| 251377_at | 7.5 | A | 1.4 | A | 1.6 | A | 8.2 | A | --- |
| 251378_at | 91 | P | 70.5 | P | 70.6 | P | 60.4 | P | --- |
| 251379_at | 77.4 | P | 91.3 | P | 84.2 | P | 136.5 | P | --- |
| 251380_at | 50.6 | P | 56.4 | A | 52.7 | A | 54.6 | P | --- |
| 251381_at | 98.3 | P | 126.3 | P | 120.3 | P | 91.1 | P | PDLP8 |
| 251382_at | 11.3 | A | 10.8 | A | 17.4 | A | 6.1 | A | --- |
| 251383_at | 141 | P | 149.2 | P | 174.4 | P | 184.8 | P | TTN1 |
| 251384_at | 35.5 | P | 3.7 | A | 26.8 | A | 7.5 | A | --- |
| 251385_at | 34.2 | P | 13 | A | 14.5 | A | 2.4 | A | --- |
| 251386_at | 790.7 | P | 860.5 | P | 717.9 | P | 739.3 | P | --- |
| 251387_at | 944 | P | 914 | P | 1036.3 | P | 1030.7 | P | --- |
| 251388_at | 87.9 | P | 116.3 | P | 68.9 | P | 121.8 | P | MAP65-4 |
| 251389_at | 156.9 | P | 158 | P | 129.2 | P | 167.3 | P | --- |
| 251390_at | 955 | P | 949 | P | 761.3 | P | 922.7 | P | --- |
| 251391_at | 683.5 | P | 683.1 | P | 747.1 | P | 720.6 | P | --- |
| 251392_at | 717.7 | P | 472.1 | P | 741.1 | P | 520.9 | P | DHDPS1 |
| 251393_at | 442.8 | P | 324.8 | P | 351.8 | P | 315.9 | P | ATG8G |
| 251394_at | 42.4 | A | 66.8 | P | 56.5 | A | 76.4 | P | FLA10 |
| 251395_at | 1487.6 | P | 2572.3 | P | 1981 | P | 2156.8 | P | FLA8 |
| 251396_at | 13658.2 | P | 15283.5 | P | 16961 | P | 15834.5 | P | --- |
| 251397_at | 26.3 | A | 5.9 | A | 28.8 | A | 17.4 | A | ATEXPB5 |
| 251398_at | 52.4 | A | 40.1 | A | 56.6 | A | 34.2 | A | --- |
| 251399_at | 311.5 | P | 374.2 | P | 363.6 | P | 309.3 | P | --- |
| 251400_at | 780.3 | P | 1640 | P | 602.2 | P | 1000.7 | P | --- |
| 251401_at | 4.8 | A | 3.2 | A | 8.2 | A | 6.7 | A | --- |
| 251402_at | 6.3 | A | 43.8 | A | 17.9 | A | 22.1 | A | --- |
| 251403_at | 1197.7 | P | 1158.2 | P | 1310.7 | P | 1000.5 | P | --- |
| 251404_at | 88.3 | P | 119.2 | P | 80.9 | P | 102.3 | P | --- |
| 251405_at | 83.9 | A | 81.2 | A | 115.8 | P | 61.9 | A | AHA7 |
| 251406_at | 371.9 | P | 372.3 | P | 418.3 | P | 333.9 | P | --- |
| 251407_at | 237.9 | P | 313.7 | P | 247 | P | 275.6 | P | --- |
| 251408_at | 934.7 | P | 968.1 | P | 856.6 | P | 831.1 | P | --- |
| 251409_at | 10376.1 | P | 9326.4 | P | 10544.6 | P | 11167 | P | --- |
| 251410_at | 17.4 | A | 6.2 | A | 6.6 | A | 2.9 | A | UCC3 |
| 251411_at | 692.6 | P | 584.7 | P | 837.6 | P | 720.1 | P | CKB3 |
| 251412_at | 35.3 | P | 49.7 | P | 84.1 | P | 62.8 | P | ATL4 |
| 251413_at | 1057.8 | P | 1239.6 | P | 1196.2 | P | 1275.7 | P | --- |
| 251414_at | 766 | P | 1087.1 | P | 887.6 | P | 957.3 | P | --- |
| 251415_at | 155.8 | P | 233.7 | P | 172.8 | P | 210.3 | P | --- |
| 251416_at | 71.6 | P | 76.9 | P | 56.1 | P | 78.1 | P | --- |
| 251417_at | 12.7 | A | 4.3 | A | 3.6 | A | 5.6 | A | --- |
| 251418_at | 289.9 | P | 257.8 | P | 267.6 | P | 322.3 | P | --- |
| 251419_at | 17.8 | A | 4.8 | A | 4.6 | A | 26 | A | --- |
| 251420_at | 47.2 | A | 56.1 | A | 47.8 | A | 30.5 | A | --- |
| 251421_at | 151.1 | P | 161.6 | P | 213 | P | 171.9 | P | --- |
| 251422_at | 274.3 | P | 309.7 | P | 185.9 | P | 284.7 | P | --- |
| 251423_at | 50.2 | P | 56.8 | P | 98.1 | P | 54.4 | M | CYCP3;2 |
| 251424_at | 1.3 | A | 1.8 | A | 1.2 | A | 1.1 | A | --- |
| 251425_at | 1470.5 | P | 1440.4 | P | 1667.2 | P | 1596.4 | P | --- |
| 251426_at | 526.1 | P | 610.3 | P | 437.5 | P | 458.1 | P | --- |
| 251427_at | 1174.1 | P | 756.6 | P | 807.4 | P | 602.6 | P | BGLU16 |
| 251428_at | 35.6 | P | 0.7 | A | 36.2 | P | 1.4 | A | DIN2 |
| 251429_at | 865.3 | P | 1300.7 | P | 948.7 | P | 1150.5 | P | ADL1E |
| 251430_at | 44.1 | P | 32.8 | P | 33.7 | P | 21.5 | M | --- |
| 251431_at | 1035.3 | P | 988.4 | P | 1079.2 | P | 932.5 | P | EIF4G |
| 251432_at | 710 | P | 423.1 | P | 560.2 | P | 566.1 | P | --- |
| 251433_at | 23.2 | A | 8.3 | A | 28.8 | A | 3.3 | A | --- |
| 251434_at | 2.4 | A | 4.6 | A | 19 | A | 1 | A | --- |
| 251435_at | 7.1 | A | 0.8 | A | 1.1 | A | 1.1 | A | --- |
| 251436_at | 31.5 | A | 135.9 | P | 50.5 | A | 60.3 | P | ARGOS |

| | | | | | | | | | |
|---|---|---|---|---|---|---|---|---|---|
| 251437_at | 466.3 | P | 521 | P | 490.7 | P | 521.1 | P | --- |
| 251438_s_at | 372.2 | P | 78.2 | P | 60 | P | 68.3 | P | --- |
| 251439_at | 329.3 | P | 438 | P | 379.3 | P | 471.6 | P | --- |
| 251440_at | 34.2 | A | 24.7 | A | 63.7 | P | 14.8 | A | ASK13 |
| 251441_at | 3.5 | A | 3.1 | A | 4.1 | A | 3 | A | ASK5 |
| 251442_at | 1002 | P | 1224.8 | P | 1062.2 | P | 1142.5 | P | --- |
| 251443_at | 685.2 | P | 989.4 | P | 524.1 | P | 862.1 | P | --- |
| 251444_at | 1221.7 | P | 1067.2 | P | 1232 | P | 965.6 | P | MAP2B |
| 251445_at | 442.4 | P | 448.2 | P | 388.1 | P | 472.4 | P | --- |
| 251446_at | 931.7 | P | 996.5 | P | 996.3 | P | 1001.9 | P | --- |
| 251447_at | 324.8 | P | 429.8 | P | 371.5 | P | 435 | P | --- |
| 251448_at | 5.3 | A | 8 | A | 28.3 | A | 9 | A | --- |
| 251449_at | 2774.5 | P | 3246.3 | P | 2976.2 | P | 3506.1 | P | ATGDI2 |
| 251450_at | 641 | P | 667.7 | P | 533.3 | P | 615.6 | P | SPL12 |
| 251451_s_at | 111.3 | P | 106.1 | P | 86.1 | P | 117 | P | --- |
| 251452_at | 24.7 | A | 14.7 | A | 31.8 | A | 4.3 | A | --- |
| 251453_at | 123.9 | P | 130.6 | P | 128 | P | 133.4 | P | --- |
| 251454_at | 436.8 | P | 557.5 | P | 502.8 | P | 556.9 | P | --- |
| 251455_at | 67.1 | P | 47.5 | A | 74.6 | A | 44.5 | A | CSY5 |
| 251456_at | 24.5 | P | 4.7 | A | 48.8 | A | 2.9 | A | BGLU27 |
| 251457_s_at | 345.5 | P | 232.7 | P | 173.4 | P | 348.8 | P | ATMRP9 |
| 251458_at | 2.2 | A | 1.6 | A | 2.5 | A | 3.1 | A | --- |
| 251459_at | 101.6 | P | 122.5 | P | 88.7 | A | 123.7 | P | --- |
| 251460_at | 437 | P | 430.2 | P | 376.2 | P | 408 | P | SAC9 |
| 251461_at | 502.2 | P | 651.7 | P | 664.1 | P | 808.5 | P | --- |
| 251462_at | 47.8 | A | 47.6 | A | 48.4 | A | 29.5 | A | ATMPK10 |
| 251463_at | 248.6 | P | 286.6 | P | 310.6 | P | 272.7 | P | --- |
| 251464_at | 624.3 | P | 750.9 | P | 789.6 | P | 573.9 | P | FTA |
| 251465_at | 354.5 | P | 397.4 | P | 384.3 | P | 436.6 | P | UTR6 |
| 251466_at | 17.5 | A | 12.5 | A | 30.1 | A | 4.2 | A | --- |
| 251467_at | 119.1 | P | 118.5 | P | 128.6 | P | 105.6 | P | --- |
| 251468_at | 132.7 | A | 123.8 | A | 174.9 | A | 117.4 | A | --- |
| 251469_at | 49 | A | 35.8 | A | 59.5 | A | 67.3 | A | --- |
| 251470_at | 56.3 | P | 135.3 | P | 118.1 | P | 113.1 | P | --- |
| 251471_at | 26.3 | A | 63.4 | A | 11.9 | A | 20 | A | --- |
| 251472_at | 92.3 | P | 83.5 | P | 65.7 | P | 94.5 | P | --- |
| 251473_at | 16.7 | A | 31.7 | A | 25.4 | A | 31.6 | A | --- |
| 251474_at | 191 | P | 180.7 | P | 199.4 | P | 194.4 | P | --- |
| 251475_at | 712.5 | P | 685 | P | 486.2 | P | 588.4 | P | --- |
| 251476_at | 160.7 | P | 234 | P | 257.8 | P | 232.4 | P | --- |
| 251477_at | 60 | P | 78.2 | P | 83.1 | P | 91 | P | --- |
| 251478_at | 84.2 | P | 104.9 | P | 69.9 | A | 79.5 | P | IQD13 |
| 251479_at | 129.8 | P | 109.9 | P | 62.2 | P | 62.1 | P | ATHLECRK |
| 251480_at | 62.9 | A | 110.8 | P | 49.7 | A | 91.3 | P | --- |
| 251481_at | 0.7 | A | 0.6 | A | 8.1 | A | 5.5 | A | --- |
| 251482_s_at | 22.5 | P | 24.9 | P | 16.6 | P | 21.3 | P | --- |
| 251483_at | 392.9 | P | 422.6 | P | 492.1 | P | 519.4 | P | --- |
| 251484_at | 361 | P | 321.7 | P | 387.8 | P | 303.5 | P | --- |
| 251485_at | 87.1 | P | 118.3 | P | 117.8 | P | 111.3 | P | SYN3 |
| 251486_at | 4813.8 | P | 4440.8 | P | 4288.3 | P | 4795.9 | P | --- |
| 251487_at | 1881.5 | P | 1320.3 | P | 1770.1 | P | 1292.9 | P | OASC |
| 251488_at | 34.5 | A | 25.5 | A | 23.3 | A | 41.1 | A | --- |
| 251489_at | 21.4 | A | 12.9 | A | 9 | A | 32 | A | --- |
| 251490_at | 136.6 | P | 219.3 | P | 187.9 | P | 258.1 | P | --- |
| 251491_at | 29.3 | A | 31.7 | A | 18.1 | A | 22.9 | A | --- |
| 251492_at | 1646 | P | 1361.3 | P | 1451.5 | P | 1276.4 | P | TXR1 |
| 251493_at | 568.1 | P | 545.5 | P | 616.8 | P | 597.4 | P | --- |
| 251494_at | 949.1 | P | 676.2 | P | 757 | P | 659.8 | P | --- |
| 251495_at | 2221.8 | P | 1714.5 | P | 2025.9 | P | 1557.5 | P | --- |
| 251496_at | 700.9 | P | 1065.6 | P | 953 | P | 963.5 | P | --- |
| 251497_at | 290.5 | P | 505.5 | P | 153.3 | P | 360.6 | P | PIL6 |
| 251498_at | 24.4 | A | 20.7 | A | 26.9 | A | 29.9 | A | --- |
| 251499_at | 114.1 | P | 178.8 | P | 138.8 | P | 144.1 | P | ATGSL11 |
| 251500_at | 290.9 | P | 307 | P | 251.9 | P | 297.1 | P | --- |
| 251501_at | 7.2 | A | 5.9 | A | 6.5 | A | 6.6 | A | --- |
| 251502_at | 3.9 | A | 16 | A | 5.6 | A | 21.8 | A | --- |
| 251503_at | 2926 | P | 1136 | P | 3072.2 | P | 1761.2 | P | ATMRP14 |
| 251504_at | 48.2 | A | 94.2 | A | 56.3 | A | 77.5 | A | TT12 |
| 251505_at | 426.9 | P | 360.1 | P | 279.1 | P | 418.9 | P | CPuORF18 |
| 251506_at | 339.2 | P | 327.8 | P | 253.8 | P | 258.6 | P | --- |
| 251507_at | 341.1 | P | 180.7 | P | 237.1 | P | 213.2 | P | --- |
| 251508_at | 369.1 | P | 366.4 | P | 295.8 | P | 385.6 | P | --- |
| 251509_at | 181.4 | P | 315.5 | P | 133.4 | P | 289.9 | P | --- |
| 251510_at | 20.7 | P | 2.1 | A | 2.3 | A | 4.4 | A | --- |
| 251511_at | 8.5 | A | 2.7 | A | 18.5 | A | 8.8 | A | --- |
| 251512_at | 2 | A | 5.8 | A | 3.2 | A | 2.5 | A | --- |
| 251513_at | 394.7 | P | 99.7 | P | 173.5 | P | 104.4 | P | PRN |
| 251514_at | 1.3 | A | 1.3 | A | 3.1 | A | 2.2 | A | --- |
| 251515_at | 23.9 | A | 19.9 | A | 34.1 | A | 1.7 | A | --- |
| 251516_s_at | 534.4 | P | 725.8 | P | 635.2 | P | 614.2 | P | --- |
| 251517_at | 0.8 | A | 1.9 | A | 0.9 | A | 2.2 | A | --- |
| 251518_at | 167.4 | P | 182 | P | 210.7 | P | 171.8 | P | --- |
| 251519_at | 758.6 | P | 1571.1 | P | 1341.9 | P | 1417.3 | P | GUN4 |
| 251520_at | 346.7 | P | 331.9 | P | 390.7 | P | 367.3 | P | --- |
| 251521_at | 84.5 | A | 94.5 | P | 123.3 | P | 102.4 | A | ACR4 |
| 251522_at | 73.9 | P | 110.5 | P | 89.5 | P | 93.2 | P | --- |
| 251523_at | 129 | P | 108.2 | P | 165.5 | P | 108 | P | --- |
| 251524_at | 2414.4 | P | 1903.9 | P | 3874.7 | P | 1958.5 | P | --- |
| 251525_at | 147.7 | P | 165.9 | P | 239 | P | 158 | P | --- |
| 251526_at | 547.8 | P | 548.1 | P | 433.7 | P | 586.3 | P | --- |
| 251527_at | 64.1 | P | 130.1 | P | 118 | P | 132.1 | P | --- |
| 251528_at | 465.3 | P | 515.2 | P | 431.5 | P | 460.2 | P | --- |
| 251529_at | 706.8 | P | 861.7 | P | 643.3 | P | 776.4 | P | --- |
| 251530_at | 109.6 | P | 146.7 | P | 156.6 | P | 142.2 | P | --- |
| 251531_at | 34.1 | A | 163.6 | P | 82.2 | A | 95.5 | P | --- |
| 251532_at | 288.6 | P | 260.7 | P | 262.2 | P | 234.8 | P | --- |
| 251533_at | 457.8 | P | 565 | P | 496.7 | P | 488.6 | P | ATRBL15 |
| 251534_at | 636.7 | P | 677.8 | P | 919.4 | P | 785.1 | P | --- |
| 251535_at | 22.6 | P | 45.9 | P | 24.5 | A | 53.2 | P | --- |
| 251536_at | 8754.8 | P | 7739.3 | P | 9228.9 | P | 8346 | P | --- |
| 251537_at | 467.4 | P | 442.1 | P | 685 | P | 424 | P | --- |
| 251538_at | 750.9 | P | 520.7 | P | 693.9 | P | 589.1 | P | --- |
| 251539_at | 252.7 | P | 286 | P | 343.4 | P | 339.3 | P | --- |
| 251540_at | 3.5 | A | 4 | A | 32.1 | A | 21.8 | A | CSY1 |
| 251541_at | 1135.7 | P | 1114.2 | P | 1073.4 | P | 1102.1 | P | CSY2 |
| 251542_at | 317.5 | P | 306.9 | P | 336.5 | P | 310.5 | P | --- |
| 251543_at | 36.4 | A | 50.3 | A | 21.6 | A | 37.9 | A | --- |
| 251544_at | 172.2 | P | 263.5 | P | 201.3 | P | 210.4 | P | GAUT15 |
| 251545_at | 37.1 | P | 51.9 | P | 63.2 | P | 48.3 | P | MTPA2 |

| | | | | | | | | | |
|---|---|---|---|---|---|---|---|---|---|
| 251546_at | 944.1 | P | 841.1 | P | 1064 | P | 766.9 | P | --- |
| 251547_at | 35 | A | 43.5 | P | 26.1 | A | 35.2 | P | --- |
| 251548_at | 1.7 | A | 1.9 | A | 5.1 | A | 2.1 | A | --- |
| 251549_at | 17.5 | A | 24.8 | A | 54.1 | A | 34 | A | --- |
| 251550_at | 55.2 | P | 89.6 | P | 78.2 | P | 105.7 | P | --- |
| 251551_at | 3175.7 | P | 3133.3 | P | 3245.4 | P | 3189.1 | P | MBF1B |
| 251552_at | 2152.4 | P | 1618.3 | P | 1910.7 | P | 1749.8 | P | --- |
| 251553_at | 182.9 | P | 160.7 | P | 215.6 | P | 165.9 | P | WRKY69 |
| 251554_at | 446.6 | P | 466.7 | P | 483.7 | P | 396.5 | P | --- |
| 251555_at | 25.4 | A | 27.5 | A | 33 | A | 21.9 | A | SHP1 |
| 251556_at | 223.9 | P | 215.4 | P | 229.5 | P | 175.7 | P | --- |
| 251557_at | 4968.9 | P | 4848.3 | P | 4848.5 | P | 4707.7 | P | --- |
| 251558_at | 371.1 | P | 423.4 | P | 433.8 | P | 527.9 | P | --- |
| 251559_at | 496.1 | P | 578.9 | P | 469.7 | P | 506.4 | P | --- |
| 251560_at | 17.5 | A | 15.8 | A | 37.1 | A | 28.2 | P | SPL15 |
| 251561_at | 2648.4 | P | 3580.4 | P | 2933.8 | P | 3564.1 | P | SCE1 |
| 251562_at | 994.6 | P | 947.2 | P | 952.1 | P | 1028.5 | P | --- |
| 251563_at | 541.7 | P | 430.7 | P | 491.9 | P | 396.8 | P | --- |
| 251564_at | 82.3 | P | 71.9 | A | 79.4 | A | 62.2 | A | XIJ |
| 251565_at | 4.6 | A | 3 | A | 12.2 | A | 2.6 | A | LBD29 |
| 251566_at | 2.1 | A | 1.5 | A | 2.2 | A | 2.3 | A | --- |
| 251567_at | 38.3 | A | 41 | A | 48.1 | A | 38.2 | A | --- |
| 251568_at | 36.6 | P | 22.6 | A | 39.9 | A | 36.8 | A | --- |
| 251569_at | 4.1 | A | 3.5 | A | 22.8 | A | 25.9 | A | --- |
| 251570_at | 24.3 | A | 2.9 | A | 7.3 | A | 20.5 | A | --- |
| 251571_at | 25.4 | P | 37.7 | P | 11.7 | A | 19.8 | A | --- |
| 251572_at | 6.3 | A | 43 | A | 4.2 | A | 2.6 | A | --- |
| 251573_at | 1749.9 | P | 1762 | P | 1362.3 | P | 1453.6 | P | --- |
| 251574_at | 149.6 | P | 184.5 | P | 238.2 | P | 184 | P | PDCB5 |
| 251575_at | 52.2 | P | 187.3 | P | 128.3 | P | 184.4 | P | BZIP61 |
| 251576_at | 84 | P | 63.1 | P | 47.4 | A | 57.6 | M | --- |
| 251577_at | 25.7 | A | 29.5 | A | 12.4 | A | 20.8 | A | --- |
| 251578_at | 820.6 | P | 630.8 | P | 881.4 | P | 731.4 | P | --- |
| 251579_at | 1007.4 | P | 1103.7 | P | 877.9 | P | 1061.6 | P | BS14A |
| 251580_at | 3.2 | A | 4.9 | A | 27.8 | A | 10.9 | A | --- |
| 251581_at | 1089.3 | P | 1223.6 | P | 1154.4 | P | 1150.2 | P | --- |
| 251582_at | 381 | P | 499.9 | P | 471.3 | P | 508.1 | P | --- |
| 251583_at | 75.6 | M | 63.4 | P | 44.9 | A | 102 | P | --- |
| 251584_at | 282.3 | P | 306.2 | P | 275.8 | P | 377.8 | P | TTL4 |
| 251585_at | 171.9 | P | 154 | P | 85.7 | P | 152.5 | P | --- |
| 251586_at | 60.8 | P | 110.2 | P | 126.7 | P | 143.8 | P | GIS |
| 251587_at | 17.1 | A | 3 | A | 6.1 | A | 13.4 | A | --- |
| 251588_at | 55.8 | P | 83.3 | P | 62.6 | P | 60.6 | P | --- |
| 251589_at | 493.2 | P | 497.8 | P | 422.3 | P | 474.4 | P | SINAT2 |
| 251590_at | 166.7 | P | 258.3 | P | 243.1 | P | 193.8 | P | AGP23 |
| 251591_at | 153.9 | P | 240.9 | P | 201 | P | 137.7 | P | --- |
| 251592_at | 15.4 | A | 44.3 | A | 26.2 | A | 20.2 | A | NTT |
| 251593_at | 380 | P | 269.6 | P | 397 | P | 286 | P | NRPA1 |
| 251594_at | 224.8 | P | 314.3 | P | 209 | P | 302.1 | P | --- |
| 251595_at | 4 | A | 1.7 | A | 6.5 | A | 2.8 | A | --- |
| 251596_at | 2027.9 | P | 2131 | P | 2413 | P | 2168.2 | P | LPAT2 |
| 251597_at | 202.3 | P | 254.4 | P | 288.1 | P | 243.6 | P | --- |
| 251598_at | 146.2 | P | 162.8 | P | 134.4 | P | 126.6 | P | --- |
| 251599_at | 3255.3 | P | 2843.2 | P | 3610 | P | 3172.1 | P | ADSS |
| 251600_at | 221.3 | P | 212.9 | P | 270.3 | P | 244.2 | P | --- |
| 251601_at | 460.4 | P | 282.4 | P | 321.8 | P | 340.5 | P | --- |
| 251602_at | 1582.9 | P | 1457.1 | P | 1567.1 | P | 1493.5 | P | --- |
| 251603_at | 145.6 | P | 87.9 | P | 101.9 | P | 118.4 | P | --- |
| 251604_at | 37.5 | P | 29.5 | P | 29.2 | P | 19.6 | P | --- |
| 251605_at | 76.2 | P | 73.7 | P | 130.1 | P | 108.3 | P | --- |
| 251606_at | 6 | A | 9.4 | A | 30.8 | A | 6.7 | A | --- |
| 251607_at | 22.3 | A | 0.8 | A | 5.5 | A | 1.1 | A | --- |
| 251608_at | 51.3 | P | 117.2 | P | 86.7 | P | 112.5 | P | UVI4-LIKE |
| 251609_at | 21.4 | M | 9.9 | A | 17.3 | A | 3.4 | A | --- |
| 251610_at | 154.9 | P | 175.6 | P | 155.9 | P | 159.2 | P | --- |
| 251611_at | 204.7 | P | 169 | P | 244.4 | P | 175.4 | P | --- |
| 251612_at | 13.9 | A | 25.2 | A | 5.4 | A | 8 | A | --- |
| 251613_at | 13.7 | A | 37.7 | A | 33.4 | A | 11.8 | A | --- |
| 251614_at | 37.1 | P | 31.8 | M | 65.4 | P | 39.2 | A | --- |
| 251615_at | 58 | P | 48.4 | P | 70.5 | P | 73 | P | --- |
| 251616_at | 792.6 | P | 792.4 | P | 762.6 | P | 839.1 | P | --- |
| 251617_at | 38.5 | A | 50.8 | P | 58.1 | A | 62.7 | A | --- |
| 251618_at | 21.1 | A | 16.2 | A | 27.7 | A | 4.5 | A | --- |
| 251619_at | 366.3 | P | 336.7 | P | 290.7 | P | 338.3 | P | --- |
| 251620_at | 42.3 | A | 6 | A | 35.3 | A | 3.5 | A | --- |
| 251621_at | 73.3 | P | 109.5 | P | 92.1 | P | 100.4 | P | --- |
| 251622_at | 580 | P | 519 | P | 537.3 | P | 483.5 | P | --- |
| 251623_at | 164.3 | P | 139.3 | P | 146.4 | P | 120 | P | AGL18 |
| 251624_at | 1739.7 | P | 1598 | P | 1371 | P | 1344.3 | P | --- |
| 251625_at | 1186.2 | P | 4396.6 | P | 1048.9 | P | 4600 | P | BGL2 |
| 251626_at | 317.3 | P | 391.6 | P | 389.3 | P | 375.5 | P | --- |
| 251627_at | 22.7 | P | 31.2 | A | 2.1 | A | 16.2 | A | BG1 |
| 251628_at | 2284.6 | P | 1990.6 | P | 2651.1 | P | 2159 | P | EIF3E |
| 251629_at | 720.3 | P | 695.8 | P | 520.2 | P | 590.6 | P | VLN3 |
| 251630_at | 318.2 | P | 376.6 | P | 329.3 | P | 326.5 | P | --- |
| 251631_at | 184.7 | P | 185.4 | P | 145.3 | P | 238.9 | P | --- |
| 251632_at | 42.3 | A | 62.8 | P | 78.1 | P | 63.8 | P | --- |
| 251633_at | 58.2 | P | 85 | P | 54 | A | 59.1 | P | --- |
| 251634_at | 254.1 | P | 229.2 | P | 217 | P | 225.1 | P | --- |
| 251635_at | 3.8 | A | 12.5 | A | 14 | A | 3.7 | A | ADPG1 |
| 251636_at | 803.3 | P | 747.2 | P | 610.6 | P | 702.6 | P | CPK32 |
| 251637_at | 197.6 | P | 198.2 | P | 206.7 | P | 193.9 | P | --- |
| 251638_at | 902.3 | P | 795.3 | P | 1441.6 | P | 875 | P | --- |
| 251639_at | 1469.9 | P | 1282.6 | P | 1631.6 | P | 1234.5 | P | --- |
| 251640_at | 2872.5 | P | 1905.8 | P | 1987.4 | P | 1679.7 | P | --- |
| 251641_at | 375.2 | P | 527.4 | P | 397.4 | P | 456.6 | P | --- |
| 251642_at | 3226.1 | P | 605.9 | P | 2770.5 | P | 808.9 | P | AtSIP2 |
| 251643_at | 732.9 | P | 897.6 | P | 707.5 | P | 802.8 | P | AGK2 |
| 251644_at | 42.1 | P | 75.8 | P | 73.7 | P | 98.8 | P | --- |
| 251645_at | 409 | P | 382.9 | P | 350.3 | P | 394.7 | P | --- |
| 251646_at | 327.4 | P | 405.6 | P | 287.9 | P | 394.4 | P | --- |
| 251647_at | 190.5 | P | 178 | P | 179.1 | P | 171.1 | P | --- |
| 251648_at | 40.7 | A | 55.1 | A | 14.8 | A | 34.3 | A | --- |
| 251649_at | 1351.2 | P | 1450.8 | P | 1034 | P | 1261.9 | P | ACA11 |
| 251650_at | 157 | P | 161.7 | P | 125.4 | P | 163.3 | P | --- |
| 251651_at | 16.8 | A | 30.4 | A | 25.3 | A | 29.1 | A | --- |
| 251652_at | 125.4 | P | 60.7 | A | 23.9 | A | 17.5 | A | --- |
| 251653_at | 43.8 | A | 68.9 | A | 53.5 | A | 33.1 | A | BOP1 |
| 251654_at | 33.2 | A | 4.4 | A | 13.4 | A | 3.1 | A | SDP1-LIKE |

| | | | | | | | | | |
|---|---|---|---|---|---|---|---|---|---|
| 251655_at | 6.5 | A | 1.6 | A | 7.4 | A | 5.6 | A | --- |
| 251656_at | 287.1 | P | 280.5 | P | 344.2 | P | 274.2 | P | --- |
| 251657_at | 815.6 | P | 745 | P | 893 | P | 860.3 | P | --- |
| 251658_at | 547 | P | 703.5 | P | 710.7 | P | 796.3 | P | --- |
| 251659_at | 1876.9 | P | 1681.2 | P | 1467.2 | P | 1593.8 | P | BIGYIN |
| 251660_at | 105.8 | P | 123.8 | P | 142.6 | P | 151.2 | P | --- |
| 251661_at | 1196.6 | P | 1088.7 | P | 1325.3 | P | 1152.1 | P | SIP2;1 |
| 251662_at | 541 | P | 585.8 | P | 541.6 | P | 539.4 | P | NRPE5 |
| 251663_at | 425.1 | P | 437.1 | P | 555.3 | P | 399.5 | P | --- |
| 251664_at | 7393.6 | P | 8800.7 | P | 8692.3 | P | 8626.5 | P | CRD1 |
| 251665_at | 282.1 | P | 294.7 | P | 303.8 | P | 293.4 | P | ARR9 |
| 251666_at | 2070.2 | P | 1883.4 | P | 2330 | P | 1972.5 | P | CBL |
| 251667_at | 1751.8 | P | 1114.2 | P | 1641.9 | P | 1227.4 | P | NAP57 |
| 251668_at | 423.3 | P | 100.8 | A | 268.6 | P | 241.4 | P | --- |
| 251669_at | 307.7 | P | 260.5 | P | 258.2 | P | 299.3 | P | --- |
| 251670_at | 424.9 | P | 356.9 | P | 398.3 | P | 377 | P | --- |
| 251671_at | 3.2 | A | 2.1 | A | 25.8 | A | 29 | A | --- |
| 251672_at | 317.2 | P | 438.9 | P | 319.7 | P | 362.9 | P | AGL16 |
| 251673_at | 46.5 | P | 134.9 | P | 74.2 | P | 143.1 | P | BG3 |
| 251674_at | 1.3 | A | 1.2 | A | 2.7 | A | 1.1 | A | --- |
| 251675_at | 293.6 | P | 375.8 | P | 261.7 | P | 300.3 | P | INO80 |
| 251676_at | 348 | P | 329.6 | P | 216.1 | P | 270.1 | P | --- |
| 251677_at | 29.9 | P | 530 | P | 1478.7 | P | 1078.4 | P | BHLH039 |
| 251678_at | 275.5 | P | 208.9 | P | 261.9 | P | 233.5 | P | EDA7 |
| 251679_at | 234.1 | P | 480.4 | P | 361.7 | P | 477.7 | P | --- |
| 251680_at | 42.5 | A | 62.5 | P | 72.6 | A | 53.2 | M | --- |
| 251681_at | 14.2 | A | 5.4 | A | 3.2 | A | 2.6 | A | --- |
| 251682_s_at | 33.7 | A | 25.9 | A | 7.6 | A | 57.3 | A | --- |
| 251683_at | 80.5 | P | 133.8 | P | 98.7 | P | 107 | P | --- |
| 251684_at | 162.6 | P | 125.2 | P | 114.1 | P | 118 | P | --- |
| 251685_at | 474.5 | P | 515 | P | 603.5 | P | 443.3 | P | --- |
| 251686_at | 128.7 | A | 93.2 | A | 99 | A | 87.4 | A | ATATG18D |
| 251687_at | 1894.4 | P | 2057.6 | P | 2077.4 | P | 1956.9 | P | --- |
| 251688_at | 196.5 | P | 398.2 | P | 378.1 | P | 406.3 | P | --- |
| 251689_at | 28.2 | A | 45.3 | A | 40.5 | A | 31.1 | A | --- |
| 251690_at | 575.7 | P | 473 | P | 589.1 | P | 559 | P | --- |
| 251691_at | 2.4 | A | 2.5 | A | 4.5 | A | 2.7 | A | --- |
| 251692_s_at | 8.6 | A | 1.2 | A | 2.1 | A | 1.2 | A | --- |
| 251693_at | 22.7 | A | 56.6 | A | 7 | A | 39.8 | A | --- |
| 251694_s_at | 0.5 | A | 1.9 | A | 15 | A | 1.7 | A | anac063 |
| 251695_at | 122 | P | 103.4 | P | 127.9 | P | 111.2 | P | --- |
| 251696_at | 78.7 | P | 21.4 | A | 46.3 | A | 62.7 | P | --- |
| 251697_at | 39 | A | 7.5 | A | 5 | A | 3.8 | A | --- |
| 251698_at | 5.1 | A | 10.1 | A | 2 | A | 11.6 | A | --- |
| 251699_at | 558.5 | P | 482 | P | 445.8 | P | 453.7 | P | CYP94D2 |
| 251700_at | 117.7 | P | 179 | P | 133.7 | P | 115.9 | P | --- |
| 251701_at | 617.6 | P | 808.2 | P | 569.5 | P | 710.2 | P | --- |
| 251702_at | 67.8 | A | 51.2 | A | 61.8 | A | 39.8 | A | BZIP49 |
| 251703_at | 26.7 | A | 30 | A | 7.3 | A | 18.7 | A | --- |
| 251704_at | 1991.3 | P | 3095.2 | P | 2214.5 | P | 2462.4 | P | --- |
| 251705_at | 381.7 | P | 721.1 | P | 271.7 | P | 530 | P | WRKY70 |
| 251706_at | 201.5 | P | 184.9 | P | 236.6 | P | 170.6 | P | --- |
| 251707_at | 1892.3 | P | 1628.1 | P | 2736.7 | P | 1606.4 | P | --- |
| 251708_at | 220 | P | 215.7 | P | 279.5 | P | 344.2 | P | --- |
| 251709_at | 3.8 | A | 1 | A | 12.1 | A | 13.4 | A | ARR17 |
| 251710_at | 110.8 | P | 98.7 | P | 129.4 | P | 96.4 | P | --- |
| 251711_at | 123 | P | 105.2 | P | 117.1 | P | 85.9 | P | PIP5K4 |
| 251712_at | 299.6 | P | 298.8 | P | 395.6 | P | 330.1 | P | --- |
| 251713_at | 209.4 | P | 198.9 | P | 157.6 | P | 177.9 | P | --- |
| 251714_at | 515 | P | 812.8 | P | 848.6 | P | 1031.5 | P | --- |
| 251715_at | 12.8 | A | 2.2 | A | 4.6 | A | 8.6 | A | --- |
| 251716_at | 3.2 | A | 2.6 | A | 12.6 | A | 5.4 | A | --- |
| 251717_at | 275 | P | 206.3 | P | 225.4 | P | 213.7 | P | LAF3 |
| 251718_at | 63.4 | P | 71.6 | P | 43.2 | P | 40.3 | P | MRLK |
| 251719_at | 840.8 | P | 771.2 | P | 731.9 | P | 825.3 | P | --- |
| 251720_at | 100.9 | P | 124.5 | P | 120.8 | P | 112.9 | P | --- |
| 251721_s_at | 2553.6 | P | 1810.7 | P | 1734.6 | P | 1555.9 | P | ALPHA-SNAP1 |
| 251722_at | 622.9 | P | 289.4 | P | 402.5 | P | 327.1 | P | --- |
| 251723_at | 45.3 | P | 45.3 | P | 69.8 | A | 40 | P | --- |
| 251724_at | 57.4 | M | 39 | P | 26.9 | A | 27 | A | --- |
| 251725_at | 153.8 | P | 158.5 | P | 145.8 | P | 200.8 | P | --- |
| 251726_at | 3.6 | A | 24.2 | A | 23.9 | A | 28.8 | A | --- |
| 251727_at | 545 | P | 1007.5 | P | 897.3 | P | 1046.8 | P | --- |
| 251728_at | 61 | M | 72.9 | P | 50 | A | 47.1 | A | --- |
| 251729_at | 2536 | P | 2525.8 | P | 2573.7 | P | 2525.6 | P | --- |
| 251730_at | 208.5 | P | 250.2 | P | 218.9 | P | 251.7 | P | --- |
| 251731_at | 34.4 | A | 35.6 | A | 40.9 | A | 20.8 | A | --- |
| 251732_at | 801.2 | P | 936.6 | P | 817.5 | P | 1009.2 | P | PRA1.B1 |
| 251733_at | 3578.2 | P | 5045.2 | P | 5323.2 | P | 4721.7 | P | CCH |
| 251734_at | 67.4 | P | 94.3 | P | 64.2 | P | 91 | P | --- |
| 251735_at | 1356.3 | P | 1374.9 | P | 1862.2 | P | 1518 | P | ATFER3 |
| 251736_at | 659.8 | P | 764.9 | P | 778.2 | P | 638 | P | --- |
| 251737_at | 5854.7 | P | 5688.8 | P | 7045.7 | P | 6217.6 | P | --- |
| 251738_at | 2703.9 | P | 2030 | P | 2432.7 | P | 2185.4 | P | EIF3C |
| 251739_at | 524.1 | P | 548.2 | P | 307.4 | P | 504.5 | P | CAN |
| 251740_at | 2315.2 | P | 1438.7 | P | 2256.5 | P | 1446 | P | ROC2 |
| 251741_at | 517 | P | 540.4 | P | 425.2 | P | 514.8 | P | UGP3 |
| 251742_at | 931.3 | P | 1227.2 | P | 883.8 | P | 1225.4 | P | --- |
| 251743_at | 55.1 | P | 54.1 | P | 31 | A | 24.5 | P | --- |
| 251744_at | 1499.9 | P | 1709.9 | P | 1150.1 | P | 1589 | P | --- |
| 251745_at | 570.7 | P | 564.7 | P | 470.8 | P | 540.3 | P | SZF1 |
| 251746_at | 235.9 | P | 179.1 | P | 217 | P | 175 | P | --- |
| 251747_at | 1.8 | A | 1.7 | A | 7.2 | A | 9.2 | A | --- |
| 251748_at | 26.8 | A | 31.6 | A | 63.2 | A | 29.2 | A | --- |
| 251749_at | 105.2 | P | 73.7 | P | 96.1 | P | 74 | P | --- |
| 251750_at | 99.8 | P | 103 | P | 145.1 | P | 157 | P | --- |
| 251751_at | 3.7 | A | 2.7 | A | 1.2 | A | 1.7 | A | --- |
| 251752_at | 294 | P | 183.8 | P | 244.3 | P | 205.8 | P | PROT2 |
| 251753_at | 347 | P | 371.5 | P | 371.2 | P | 408.6 | P | --- |
| 251754_at | 60.9 | A | 47.7 | P | 48.6 | A | 62 | A | --- |
| 251755_at | 1.2 | A | 1.6 | A | 3.2 | A | 1.9 | A | --- |
| 251756_at | 66.6 | P | 80.1 | P | 83 | P | 90.9 | P | --- |
| 251757_at | 538.1 | P | 394.1 | P | 499 | P | 446.3 | P | --- |
| 251758_at | 2724.8 | P | 3046.2 | P | 3115.6 | P | 2817.6 | P | --- |
| 251759_at | 1204.4 | P | 1399.9 | P | 1244 | P | 1248.4 | P | ATDFD |
| 251760_at | 365.4 | P | 345.4 | P | 399.1 | P | 381 | P | --- |
| 251761_at | 55.1 | A | 42.9 | P | 36.6 | M | 37.7 | A | --- |
| 251762_at | 5441.8 | P | 6841.7 | P | 6950.6 | P | 6919.6 | P | SBPASE |
| 251763_at | 125.3 | P | 118.5 | P | 117 | P | 148.1 | P | MYB109 |

| | | | | | | | | | |
|---|---|---|---|---|---|---|---|---|---|
| 251764_at | 1013.6 | P | 1124.7 | P | 992.4 | P | 1116.3 | P | EPC1 |
| 251765_at | 47.3 | A | 47.9 | A | 44.8 | A | 61.7 | P | --- |
| 251766_at | 46.3 | P | 46.6 | P | 58 | P | 34.7 | P | --- |
| 251767_at | 2.8 | A | 32 | A | 40.5 | A | 40.8 | A | --- |
| 251768_at | 20.6 | A | 52.7 | A | 2.7 | A | 57.1 | P | --- |
| 251769_at | 126.3 | P | 119 | P | 82.9 | P | 99.7 | P | CCR3 |
| 251770_at | 613.6 | P | 34.2 | A | 257.9 | P | 39.4 | A | --- |
| 251771_at | 52.5 | P | 53.6 | P | 61.8 | P | 35.1 | P | CSLA14 |
| 251772_at | 248.5 | P | 305.3 | P | 243.7 | P | 342.1 | P | --- |
| 251773_at | 673.3 | P | 949.8 | P | 733.7 | P | 865.6 | P | --- |
| 251774_at | 39.9 | A | 33.8 | A | 33 | A | 19.1 | A | --- |
| 251775_s_at | 2668.6 | P | 3235 | P | 2548.4 | P | 3671.5 | P | P5CS1 |
| 251776_at | 2414.7 | P | 1593.5 | P | 2724.4 | P | 1854.6 | P | emb1624 |
| 251777_s_at | 4.5 | A | 1.4 | A | 5.2 | A | 1.6 | A | --- |
| 251778_at | 80.2 | P | 168.5 | P | 127.4 | P | 138.6 | P | ROPGEF6 |
| 251779_at | 12.8 | A | 10.4 | A | 5 | A | 3 | A | --- |
| 251780_s_at | 369.9 | P | 231.5 | P | 219.5 | P | 279 | P | --- |
| 251781_at | 193.4 | P | 222.2 | P | 254.7 | P | 220.1 | P | PGP20 |
| 251782_at | 566.1 | P | 1035.6 | P | 564.4 | P | 818.1 | P | HEXO1 |
| 251783_at | 2313.1 | P | 2058 | P | 2883.5 | P | 2163.4 | P | RPL23AB |
| 251784_at | 2350.3 | P | 2622.3 | P | 2253.2 | P | 2341.5 | P | PPL1 |
| 251785_at | 677 | P | 522.7 | P | 807.9 | P | 564.5 | P | ATWBC19 |
| 251786_at | 285.6 | P | 325.6 | P | 286.3 | P | 277.2 | P | MKP1 |
| 251787_at | 1655.5 | P | 1425.2 | P | 1803.7 | P | 1590.2 | P | --- |
| 251788_at | 468.9 | P | 324.7 | P | 519.9 | P | 313.1 | P | --- |
| 251789_at | 581.1 | P | 657.8 | P | 550.7 | P | 710.2 | P | --- |
| 251790_at | 242.6 | P | 316.3 | P | 162.5 | P | 242.3 | P | --- |
| 251791_at | 165.4 | P | 136.5 | P | 138.5 | P | 147.8 | P | ATEXPA16 |
| 251792_at | 19 | A | 8.9 | A | 27 | A | 3.7 | A | --- |
| 251793_at | 18.5 | A | 35.7 | A | 34.4 | A | 20.8 | A | --- |
| 251794_at | 4.8 | A | 3.3 | A | 6.2 | A | 4.1 | A | --- |
| 251795_at | 208.9 | P | 191.5 | P | 197.2 | P | 196 | P | --- |
| 251796_at | 1521.6 | P | 2175 | P | 1458.4 | P | 2206.9 | P | --- |
| 251797_at | 163.3 | P | 345.4 | P | 294.1 | P | 231.4 | P | AGF2 |
| 251798_at | 1120.4 | P | 1118 | P | 1312.3 | P | 1148 | P | SCL30 |
| 251799_at | 704.5 | P | 859.1 | P | 608.6 | P | 899.6 | P | --- |
| 251800_at | 429.4 | P | 337 | P | 566 | P | 374.9 | P | RBL |
| 251801_at | 10351 | P | 10183 | P | 11747.8 | P | 10612.8 | P | TPI |
| 251802_at | 793.8 | P | 766.7 | P | 712.8 | P | 714.2 | P | UBC14 |
| 251803_at | 114.4 | P | 158.1 | P | 134 | P | 116.4 | P | --- |
| 251804_at | 2150.2 | P | 1448.2 | P | 1633.7 | P | 1470.9 | P | --- |
| 251805_at | 317.9 | P | 354.7 | P | 445.5 | P | 413 | P | SDIR1 |
| 251806_at | 24.3 | A | 40.3 | P | 46 | A | 53.1 | P | OBP3 |
| 251807_at | 1101.2 | P | 1009.5 | P | 1663.5 | P | 1079.7 | P | OVA1 |
| 251808_at | 468.1 | P | 619.5 | P | 432.1 | P | 469.2 | P | --- |
| 251809_at | 212.4 | P | 189.2 | P | 185.8 | P | 186.4 | P | --- |
| 251810_at | 1664.1 | P | 2538 | P | 2007.8 | P | 2182.2 | P | --- |
| 251811_at | 138.8 | P | 149 | P | 207.9 | P | 204.5 | P | SMZ |
| 251812_at | 170.1 | P | 192.9 | P | 259.4 | P | 180.5 | P | --- |
| 251813_at | 511.2 | P | 762.2 | P | 684.4 | P | 635.5 | P | TON1A |
| 251814_at | 12526.8 | P | 14093.1 | P | 11305.2 | P | 12784.4 | P | LHCA1 |
| 251815_at | 1806.3 | P | 2263.6 | P | 1785.1 | P | 2059.5 | P | CXIP1 |
| 251816_at | 241.5 | P | 279.5 | P | 256.5 | P | 261.4 | P | TON1B |
| 251817_at | 1307.9 | P | 1310.5 | P | 1170.5 | P | 1264.2 | P | ARA6 |
| 251818_at | 533.8 | P | 701.5 | P | 590.7 | P | 563.7 | P | ATVPS33 |
| 251819_at | 630.2 | P | 690.9 | P | 571.4 | P | 714.1 | P | PGPS2 |
| 251820_at | 1053.5 | P | 1319.2 | P | 1156 | P | 1323.3 | P | GSTL2 |
| 251821_at | 541.4 | P | 560.1 | P | 430.7 | P | 531.4 | P | --- |
| 251822_at | 49.6 | P | 44.5 | P | 22.5 | A | 58.4 | P | --- |
| 251823_at | 259.3 | P | 322.1 | P | 421.8 | P | 248.7 | P | --- |
| 251824_at | 11 | A | 10 | A | 40.8 | A | 17.3 | A | --- |
| 251825_at | 114.8 | A | 25.8 | A | 71.8 | A | 17.8 | A | --- |
| 251826_at | 1240.9 | P | 356.9 | P | 1081.5 | P | 513.8 | P | --- |
| 251827_at | 1911.3 | P | 1103 | P | 2159 | P | 1652.7 | P | TT5 |
| 251828_at | 458.1 | P | 458.2 | P | 478.8 | P | 435.9 | P | --- |
| 251829_at | 267 | P | 321.9 | P | 226.7 | P | 343.8 | P | --- |
| 251830_at | 994.5 | P | 750.9 | P | 1143 | P | 840.6 | P | PUR5 |
| 251831_at | 484.8 | P | 527 | P | 454.4 | P | 492.5 | P | --- |
| 251832_at | 10.6 | A | 15.9 | A | 4.7 | A | 24.3 | A | ATEXO70H1 |
| 251833_at | 163.3 | P | 84.2 | P | 114.4 | P | 99 | P | --- |
| 251834_at | 354.3 | P | 367.6 | P | 372.7 | P | 279.3 | P | --- |
| 251835_at | 15 | A | 15 | A | 19.6 | A | 19.5 | A | --- |
| 251836_at | 3.6 | A | 8.7 | A | 20.4 | A | 13 | A | --- |
| 251837_s_at | 1058.1 | P | 881.2 | P | 953.7 | P | 991.5 | P | --- |
| 251838_at | 3.1 | A | 13.9 | A | 19.1 | A | 2 | A | --- |
| 251839_at | 363.5 | P | 332.4 | P | 395.9 | P | 299.9 | P | PLA IIIA |
| 251840_at | 632.5 | P | 988 | P | 629.3 | P | 943.8 | P | ATPDIL1-3 |
| 251841_at | 279.7 | P | 272.2 | P | 387.7 | P | 294.3 | P | --- |
| 251842_at | 32 | A | 3.8 | A | 7.5 | A | 3.3 | A | --- |
| 251843_x_at | 97 | P | 127.1 | P | 119.6 | P | 115.2 | P | ATHRGP1 |
| 251844_at | 56.7 | A | 75.2 | P | 35.8 | A | 65.5 | A | --- |
| 251845_at | 1022.2 | P | 1126 | P | 1148.9 | P | 1049 | P | ATGCN4 |
| 251846_at | 557.9 | P | 837 | P | 597.7 | P | 794 | P | HTA11 |
| 251847_at | 5005.4 | P | 2945.3 | P | 3395.1 | P | 2394.8 | P | TSA1 |
| 251848_at | 797.8 | P | 825.2 | P | 582.3 | P | 802 | P | BZIP25 |
| 251849_at | 277 | P | 306.9 | P | 328.3 | P | 271.3 | P | HAG1 |
| 251850_at | 369.6 | P | 392.3 | P | 484 | P | 288.7 | P | FBL17 |
| 251851_at | 362.1 | P | 411 | P | 304.2 | P | 394.1 | P | TTN8 |
| 251852_at | 407.7 | P | 492.8 | P | 516.9 | P | 593.1 | P | --- |
| 251853_at | 288.9 | P | 395.9 | P | 404.8 | P | 357.8 | P | --- |
| 251854_at | 21.2 | A | 22.8 | A | 26.8 | A | 37.3 | A | --- |
| 251855_at | 732 | P | 804.5 | P | 638.2 | P | 847.1 | P | --- |
| 251856_at | 204.3 | P | 130.1 | P | 221.7 | P | 232.5 | P | AMP1 |
| 251857_at | 17.3 | A | 7.2 | A | 3.4 | A | 9.7 | A | --- |
| 251858_at | 343.6 | P | 521.8 | P | 409.1 | P | 567.1 | P | PIP2;5 |
| 251859_at | 176 | P | 195 | P | 163.7 | P | 222.6 | P | --- |
| 251860_at | 2241.5 | P | 2150.3 | P | 2111.4 | P | 2128.9 | P | GR |
| 251861_at | 450.2 | P | 621.6 | P | 573.7 | P | 583.6 | P | BME3 |
| 251862_at | 227.4 | P | 198.8 | P | 209.3 | P | 214.4 | P | PUB14 |
| 251863_at | 33.3 | P | 37.3 | P | 53.3 | P | 44.7 | P | MRH2 |
| 251864_at | 474.7 | P | 554.4 | P | 644.7 | P | 689.4 | P | PMR6 |
| 251865_at | 73.1 | P | 113.2 | P | 117.2 | P | 120.6 | P | --- |
| 251866_at | 374 | P | 441.8 | P | 444.2 | P | 463 | P | emb1967 |
| 251867_at | 692.4 | P | 966.8 | P | 1005.1 | P | 807.4 | P | SKIP5 |
| 251868_at | 1.6 | A | 3.8 | A | 2.6 | A | 1.6 | A | RPB5E |
| 251869_at | 127.4 | P | 462.7 | P | 126.9 | P | 334.1 | P | --- |
| 251870_at | 30.7 | P | 26.4 | M | 56.7 | P | 49.1 | P | --- |
| 251871_at | 4.5 | A | 0.4 | A | 1.4 | A | 0.5 | A | --- |
| 251872_at | 15.2 | A | 12.1 | A | 33.5 | A | 1.2 | A | --- |

| | | | | | | | | | |
|---|---|---|---|---|---|---|---|---|---|
| 251873_at | 248.6 | P | 235.6 | P | 283.8 | P | 231.4 | P | --- |
| 251874_at | 3.2 | A | 2.7 | A | 4 | A | 2.4 | A | --- |
| 251875_at | 25.1 | A | 45.5 | A | 10.5 | A | 46.3 | A | --- |
| 251876_at | 302.8 | P | 260.3 | P | 331.3 | P | 320.1 | P | RGD3 |
| 251877_at | 1049.1 | P | 1145.3 | P | 894.4 | P | 955.6 | P | ATVAMP727 |
| 251878_at | 5.5 | A | 0.9 | A | 4.5 | A | 1.3 | A | --- |
| 251879_at | 461.1 | P | 719.6 | P | 420.8 | P | 628.9 | P | --- |
| 251880_at | 400.9 | P | 414.4 | P | 329.8 | P | 390.1 | P | --- |
| 251881_at | 174.4 | P | 195.8 | P | 241.7 | P | 203.4 | P | --- |
| 251882_at | 1164.5 | P | 769 | P | 799.9 | P | 660.3 | P | PTR1 |
| 251883_at | 6716 | P | 6797 | P | 5864.7 | P | 6815 | P | --- |
| 251884_at | 257.1 | P | 114.6 | P | 155.9 | P | 80.9 | P | --- |
| 251885_at | 5673.9 | P | 6387.2 | P | 6170.8 | P | 6539.6 | P | --- |
| 251886_at | 137.5 | P | 365.2 | P | 231.7 | P | 342.2 | P | --- |
| 251887_at | 271.1 | P | 320.4 | P | 272.1 | P | 302.7 | P | ATFIP37 |
| 251888_at | 529.9 | P | 623.3 | P | 405.8 | P | 615.7 | P | --- |
| 251889_at | 385.3 | P | 427.1 | P | 457.4 | P | 517.6 | P | --- |
| 251890_at | 464.5 | P | 499.5 | P | 513.9 | P | 483 | P | SCR |
| 251891_at | 102.5 | P | 86 | P | 108 | P | 63.8 | P | WRI1 |
| 251892_at | 179.9 | P | 107.5 | P | 108.2 | P | 113.3 | P | WRI1 |
| 251893_at | 474.8 | P | 494.2 | P | 455.9 | P | 398.3 | P | --- |
| 251894_at | 237.6 | P | 249.6 | P | 249.1 | P | 227 | P | --- |
| 251895_at | 102.7 | P | 225.7 | P | 98.7 | P | 237.8 | P | ATEP3 |
| 251896_at | 180.5 | P | 140.2 | M | 217 | P | 176.5 | P | --- |
| 251897_at | 1948.7 | P | 1872.3 | P | 1810.5 | P | 2002.2 | P | --- |
| 251898_at | 3.9 | A | 44.8 | A | 6.6 | A | 20.5 | A | AP3 |
| 251899_at | 2533.1 | P | 3618.6 | P | 2905.8 | P | 3141.4 | P | --- |
| 251900_at | 187.2 | P | 207.4 | P | 249.8 | P | 192.6 | P | SRS6 |
| 251901_at | 184.2 | P | 162.8 | P | 151.1 | P | 204.7 | P | --- |
| 251902_at | 2567.1 | P | 1869.4 | P | 2536.5 | P | 1989 | P | PUMP1 |
| 251903_at | 185.4 | P | 172.9 | P | 202.3 | P | 195.9 | P | --- |
| 251904_at | 198.2 | P | 257.4 | P | 202.8 | P | 211.9 | P | --- |
| 251905_at | 573.9 | P | 645.5 | P | 547.3 | P | 602 | P | GAPB |
| 251906_at | 229.4 | P | 293.8 | P | 261.4 | P | 309.4 | P | ATCHX20 |
| 251907_at | 119.1 | P | 127.8 | P | 138.6 | P | 153.4 | P | --- |
| 251908_at | 14.5 | A | 16.3 | A | 33.2 | A | 2.8 | A | --- |
| 251909_at | 22.6 | A | 15.9 | A | 7.3 | A | 22.2 | A | TRFL4 |
| 251910_at | 192.6 | P | 223.5 | P | 148 | P | 174.8 | P | --- |
| 251911_at | 3.2 | A | 15.5 | A | 16.8 | A | 13.2 | A | --- |
| 251912_at | 33.5 | A | 25.5 | A | 44.8 | A | 36.9 | A | --- |
| 251913_at | 15.7 | A | 2.3 | A | 5.3 | A | 1.1 | A | --- |
| 251914_at | 197.4 | P | 191.6 | P | 162 | P | 167 | P | --- |
| 251915_at | 119.4 | P | 128 | P | 169.4 | P | 147.7 | P | --- |
| 251916_at | 205.7 | P | 230.4 | P | 199.6 | P | 187.2 | P | --- |
| 251917_at | 553 | P | 601.7 | P | 588.5 | P | 500.5 | P | --- |
| 251918_at | 58.9 | P | 94.9 | P | 50.5 | M | 73 | P | --- |
| 251919_at | 183.8 | P | 458.6 | P | 168.8 | P | 331.6 | P | --- |
| 251920_at | 717.6 | P | 868.9 | P | 839.8 | P | 829.6 | P | --- |
| 251921_at | 5370.4 | P | 5640.2 | P | 6951.7 | P | 5989.7 | P | --- |
| 251922_at | 328.2 | P | 272.5 | P | 190 | P | 201.8 | P | --- |
| 251923_at | 417.6 | P | 345.4 | P | 491.9 | P | 340 | P | --- |
| 251924_at | 2122.5 | P | 2409.2 | P | 1863.3 | P | 2455.8 | P | --- |
| 251925_at | 95 | P | 143.2 | P | 110 | P | 139.2 | P | --- |
| 251926_at | 5799.2 | P | 5501.7 | P | 7118.6 | P | 4698.1 | P | --- |
| 251927_at | 3827.9 | P | 3229.6 | P | 2752.8 | P | 2821 | P | --- |
| 251928_at | 248.8 | P | 266.4 | P | 346.5 | P | 375.2 | P | --- |
| 251929_at | 827.5 | P | 753.8 | P | 908 | P | 767.1 | P | SIGC |
| 251930_at | 473.9 | P | 518.1 | P | 344.4 | P | 413.4 | P | ATRBL4 |
| 251931_at | 426.6 | P | 376.8 | P | 230.1 | P | 326.4 | P | --- |
| 251932_at | 973.6 | P | 871.4 | P | 850.4 | P | 853.2 | P | PAS1 |
| 251933_at | 31.9 | A | 34.5 | A | 34.4 | A | 20.1 | A | --- |
| 251934_at | 21.7 | A | 4.5 | A | 7.1 | A | 10.3 | A | --- |
| 251935_at | 1526.1 | P | 1692.7 | P | 1812.2 | P | 1628.3 | P | --- |
| 251936_at | 402.6 | P | 362.4 | P | 302.7 | P | 352 | P | MEE40 |
| 251937_at | 375 | P | 422.1 | P | 420.2 | P | 412.9 | P | --- |
| 251938_at | 7275.8 | P | 7213.9 | P | 8965 | P | 7617.8 | P | --- |
| 251939_at | 128.4 | P | 112.3 | P | 79 | P | 97.1 | P | --- |
| 251940_at | 16.7 | A | 34.8 | A | 76.8 | P | 28 | A | --- |
| 251941_at | 885.8 | P | 858.9 | P | 797.4 | P | 761.3 | P | --- |
| 251942_at | 200.8 | P | 50 | A | 70.3 | M | 84.5 | P | PDR9 |
| 251943_at | 819.7 | P | 871.2 | P | 1027.5 | P | 936 | P | RSZ32 |
| 251944_at | 14.3 | A | 51.3 | P | 59.2 | A | 19.1 | A | --- |
| 251945_at | 508.3 | P | 630.9 | P | 550 | P | 587.7 | P | UXS1 |
| 251946_at | 252.3 | P | 276.7 | P | 347.1 | P | 297.5 | P | --- |
| 251947_at | 8.1 | A | 8.1 | A | 21.3 | A | 1.6 | A | --- |
| 251948_at | 1477.8 | P | 1353 | P | 1394 | P | 1184.7 | P | --- |
| 251949_at | 23 | A | 10.9 | A | 41.8 | A | 36.7 | A | --- |
| 251950_at | 2.6 | A | 1 | A | 14.8 | A | 3 | A | --- |
| 251951_s_at | 733.7 | P | 548 | P | 763.2 | P | 600.3 | P | --- |
| 251952_at | 55.9 | P | 120.3 | P | 51.4 | P | 75 | P | --- |
| 251953_at | 62.9 | P | 111.3 | P | 40.4 | P | 46 | P | CPuORF51 |
| 251954_at | 395.6 | P | 604.2 | P | 289.3 | P | 596.2 | P | CPuORF51 |
| 251955_at | 99.2 | A | 125.9 | A | 123.2 | A | 109.8 | M | --- |
| 251956_at | 7308.5 | P | 8171.2 | P | 9867.8 | P | 9386.7 | P | CP29 |
| 251957_at | 106.4 | P | 96.2 | P | 112.9 | P | 89.8 | P | --- |
| 251958_at | 2285 | P | 2922.5 | P | 2169.6 | P | 2275.2 | P | --- |
| 251959_at | 148.4 | P | 166.1 | P | 101.7 | P | 151.6 | P | --- |
| 251960_at | 719.3 | P | 662.7 | P | 796.4 | P | 784 | P | HEMB1 |
| 251961_at | 404.9 | P | 506.5 | P | 347.9 | P | 488.8 | P | AtPPa4 |
| 251962_at | 6915.5 | P | 6876 | P | 7759.8 | P | 6929.4 | P | PIP2A |
| 251963_at | 274.5 | P | 251 | P | 243.4 | P | 261.4 | P | AFC1 |
| 251964_at | 293.2 | P | 276.1 | P | 267.1 | P | 223.1 | P | --- |
| 251965_at | 109.9 | P | 103.5 | P | 98.2 | P | 97.2 | P | --- |
| 251966_at | 60.4 | P | 107.6 | P | 123.7 | P | 116.3 | P | --- |
| 251967_at | 326.4 | P | 319.9 | P | 287.5 | P | 286.1 | P | --- |
| 251968_at | 126.4 | P | 198.2 | P | 211.1 | P | 183.3 | P | --- |
| 251969_at | 511.1 | P | 867.1 | P | 702.4 | P | 990.1 | P | LUT1 |
| 251970_at | 24.1 | A | 54.1 | P | 7 | A | 10.4 | A | UGT73D1 |
| 251971_at | 120.5 | P | 50.1 | P | 97.6 | P | 58.5 | P | UGT73C7 |
| 251972_at | 78.4 | P | 114.6 | P | 138.9 | P | 139.7 | P | --- |
| 251973_at | 734.9 | P | 734.1 | P | 787.7 | P | 607.4 | P | --- |
| 251974_at | 23 | P | 7 | A | 3.1 | A | 1.4 | A | AtMYB27 |
| 251975_at | 598.5 | P | 560.5 | P | 374.8 | P | 473.9 | P | --- |
| 251976_at | 123.9 | P | 113.7 | P | 110.9 | P | 132.4 | P | AtRLP45 |
| 251977_at | 10.8 | A | 16.7 | A | 12.4 | A | 8.6 | A | --- |
| 251978_at | 3.1 | A | 3 | A | 5.5 | A | 3 | A | CYP71B30P |
| 251979_at | 94.3 | P | 106.7 | P | 92.9 | A | 104.3 | P | LUT1 |
| 251980_at | 178.8 | P | 197.5 | P | 227.8 | P | 153.4 | P | --- |
| 251981_at | 2.3 | A | 5.1 | A | 5.4 | A | 2.3 | A | --- |

| | | | | | | | | | |
|---|---|---|---|---|---|---|---|---|---|
| 251982_at | 595 | P | 970.2 | P | 856.5 | P | 1065.1 | P | --- |
| 251983_at | 247.7 | P | 154.6 | P | 190.5 | P | 124.1 | P | --- |
| 251984_at | 2820.3 | P | 2277.1 | P | 2849.4 | P | 2682.9 | P | PAL2 |
| 251985_at | 137.2 | P | 139.1 | P | 142.1 | P | 182 | P | --- |
| 251986_at | 78.5 | P | 123.7 | P | 107.6 | P | 99.1 | P | --- |
| 251987_at | 322.4 | P | 325.9 | P | 319.9 | P | 288 | P | CYP71B5 |
| 251988_at | 1.2 | A | 6.7 | A | 2.5 | A | 0.9 | A | CYP71B31 |
| 251989_at | 1635.7 | P | 1599.8 | P | 1530.9 | P | 1580 | P | LOS4 |
| 251990_at | 111.8 | A | 130 | P | 148 | A | 113.4 | A | --- |
| 251991_at | 609 | P | 482.2 | P | 563.8 | P | 448.1 | P | NF-YB10 |
| 251992_at | 82.1 | P | 65 | P | 85.1 | P | 86 | P | --- |
| 251993_at | 4748.7 | P | 4886.8 | P | 5070.6 | P | 5074.1 | P | --- |
| 251994_at | 129.2 | P | 256.3 | P | 174.8 | P | 253.8 | P | KIPK |
| 251995_at | 323.2 | P | 450 | P | 360.1 | P | 411.1 | P | FK |
| 251996_at | 537.7 | P | 861.8 | P | 563.6 | P | 815.2 | P | BGAL2 |
| 251997_at | 7476.1 | P | 6018.6 | P | 7499.1 | P | 7455.7 | P | STV1 |
| 251998_at | 270.5 | P | 330.9 | P | 274.2 | P | 320.3 | P | UPL7 |
| 251999_at | 3.5 | A | 1.7 | A | 2.6 | A | 2.3 | A | --- |
| 252000_at | 238.1 | P | 270 | P | 209.8 | P | 242.7 | P | --- |
| 252001_at | 860.2 | P | 1046.8 | P | 563.1 | P | 1134.8 | P | FTSZ2-2 |
| 252002_at | 295.6 | P | 402.4 | P | 372.3 | P | 328.8 | P | --- |
| 252003_at | 30.5 | P | 14.2 | P | 7.8 | P | 12.9 | A | ZPR3 |
| 252004_at | 1.6 | A | 2.1 | A | 1.6 | A | 4.1 | A | PAP20 |
| 252005_at | 7.1 | A | 7.5 | A | 2 | A | 5.5 | A | PAP21 |
| 252006_at | 23.2 | A | 26 | P | 73.9 | A | 27.7 | A | PAP22 |
| 252007_at | 1.6 | A | 0.6 | A | 3.4 | A | 1.1 | A | --- |
| 252008_at | 477.1 | P | 496.3 | P | 455.8 | P | 497.3 | P | --- |
| 252009_at | 1344.7 | P | 1374.5 | P | 1029.6 | P | 1093.1 | P | --- |
| 252010_at | 401.1 | P | 342.2 | P | 393.4 | P | 353.5 | P | --- |
| 252011_at | 656.5 | P | 499.4 | P | 1058.2 | P | 689.1 | P | ACA1 |
| 252012_at | 4678.6 | P | 4841.7 | P | 4313.1 | P | 4973.5 | P | --- |
| 252013_at | 353.6 | P | 221.3 | P | 366.6 | P | 237 | P | --- |
| 252014_at | 867.3 | P | 601.6 | P | 862.6 | P | 550.5 | P | --- |
| 252015_at | 333 | P | 309 | P | 316.1 | P | 284.4 | P | AtGRF4 |
| 252016_at | 514 | P | 502.4 | P | 567.6 | P | 608.2 | P | --- |
| 252017_at | 4.3 | A | 2.6 | A | 6.6 | A | 3.9 | A | CYP76G1 |
| 252018_at | 200.2 | P | 242.4 | P | 227.2 | P | 226 | P | AtPP2-A15 |
| 252019_at | 3.7 | A | 8 | A | 3.2 | A | 1.9 | A | --- |
| 252020_at | 0.7 | A | 0.6 | A | 0.6 | A | 14.3 | A | ASK6 |
| 252021_at | 41 | A | 22.6 | A | 19.9 | A | 33 | A | --- |
| 252022_at | 9834.8 | P | 9134.6 | P | 10336.9 | P | 10445.9 | P | --- |
| 252023_at | 369.8 | P | 452 | P | 470.2 | P | 543 | P | --- |
| 252024_at | 3642.2 | P | 3719.9 | P | 3070.6 | P | 3258 | P | ATMDAR1 |
| 252025_at | 111.2 | P | 176.9 | P | 190.2 | P | 156.5 | P | --- |
| 252026_at | 256.4 | P | 330.3 | P | 251.6 | P | 292.5 | P | SRPK4 |
| 252027_at | 1784.4 | P | 1412.5 | P | 1502.6 | P | 1303.6 | P | VSR1 |
| 252028_at | 148.6 | P | 232.9 | P | 181.3 | P | 215.5 | P | --- |
| 252029_s_at | 336.6 | P | 297 | P | 262.9 | P | 206.4 | P | --- |
| 252030_at | 3.1 | A | 1.5 | A | 5.1 | A | 3.6 | A | --- |
| 252031_at | 1031.3 | P | 867.7 | P | 1518.4 | P | 936.4 | P | --- |
| 252032_at | 4839.3 | P | 5697.7 | P | 5507.7 | P | 5560 | P | --- |
| 252033_at | 1181.1 | P | 1153.9 | P | 1082.9 | P | 1158.7 | P | --- |
| 252034_at | 1123.2 | P | 904 | P | 1096.1 | P | 945 | P | --- |
| 252035_at | 33.6 | M | 45.9 | A | 3.4 | A | 12.4 | A | KCS15 |
| 252036_at | 554.5 | P | 443.1 | P | 404.6 | P | 370.9 | P | --- |
| 252037_at | 2144.5 | P | 2665.2 | P | 2040.9 | P | 2285.1 | P | CAM9 |
| 252038_at | 252.2 | P | 280.7 | P | 225.6 | P | 227.1 | P | --- |
| 252039_at | 436.2 | P | 594 | P | 367.5 | P | 563.9 | P | --- |
| 252040_at | 711.1 | P | 873.8 | P | 709.2 | P | 783.3 | P | --- |
| 252041_at | 1518.1 | P | 1545.3 | P | 1716.1 | P | 1668.9 | P | NRPB11 |
| 252042_at | 179 | P | 237.8 | P | 185.4 | P | 172.3 | P | --- |
| 252043_at | 600 | P | 589.6 | P | 728.9 | P | 553.9 | P | --- |
| 252044_at | 823.6 | P | 523.1 | P | 534 | P | 476 | P | OEP7 |
| 252045_at | 29.9 | A | 27.9 | A | 9.2 | A | 46.6 | A | PUB22 |
| 252046_at | 21.6 | P | 25.3 | A | 6.7 | A | 42.3 | P | --- |
| 252047_at | 110.5 | P | 119 | P | 130.6 | P | 170.9 | P | --- |
| 252048_at | 1873.6 | P | 2469 | P | 2354.8 | P | 2891 | P | --- |
| 252049_at | 43.6 | A | 43 | P | 52 | A | 38.3 | P | --- |
| 252050_at | 55.6 | P | 81 | P | 67.2 | P | 77 | P | --- |
| 252051_at | 197.9 | P | 174.2 | P | 189.1 | P | 166.1 | P | --- |
| 252052_at | 21.6 | A | 9.5 | A | 1.8 | A | 11.7 | A | AtcwINV2 |
| 252053_at | 477.9 | P | 523.4 | P | 265.2 | P | 381.6 | P | SYP122 |
| 252054_at | 82.4 | A | 59.4 | A | 65 | A | 64.4 | P | OFP18 |
| 252055_at | 3039 | P | 2852.4 | P | 2995.7 | P | 2965.7 | P | --- |
| 252056_at | 9714.7 | P | 8614.9 | P | 11144.1 | P | 9099.3 | P | UBQ1 |
| 252057_at | 37.8 | A | 103.1 | P | 73.5 | A | 68.1 | A | --- |
| 252058_at | 1595.9 | P | 1588.8 | P | 1753.9 | P | 1817.4 | P | --- |
| 252059_at | 2929.4 | P | 2948.7 | P | 2672.9 | P | 3618.6 | P | UEV1D-4 |
| 252060_at | 159.3 | P | 368.7 | P | 120.6 | P | 208.5 | P | PAD4 |
| 252061_at | 2.2 | A | 7.2 | A | 2 | A | 4.8 | A | --- |
| 252062_at | 510.2 | P | 620 | P | 451.8 | P | 570.7 | P | --- |
| 252063_at | 2.1 | A | 4.2 | A | 30.8 | A | 3.9 | A | LTP12 |
| 252064_at | 725.2 | P | 823.4 | P | 726.5 | P | 811.4 | P | --- |
| 252065_at | 843.6 | P | 1114.8 | P | 1039.8 | P | 992 | P | --- |
| 252066_at | 2041.3 | P | 2250.4 | P | 1848.5 | P | 2258.8 | P | FER |
| 252067_at | 897.1 | P | 1027 | P | 1036.7 | P | 1096.6 | P | --- |
| 252068_at | 136.3 | P | 183.1 | P | 55.6 | A | 161 | P | --- |
| 252069_s_at | 450.1 | P | 516.6 | P | 379.1 | P | 566.8 | P | --- |
| 252070_at | 20 | A | 7.1 | A | 12.8 | A | 13.2 | A | --- |
| 252071_at | 23.3 | P | 10.9 | A | 5 | A | 7.2 | A | --- |
| 252072_at | 35.4 | P | 38.3 | P | 66.3 | P | 41.9 | M | --- |
| 252073_at | 39 | P | 64.1 | P | 40.5 | M | 71.6 | P | --- |
| 252074_at | 13.2 | A | 2.3 | A | 4.4 | A | 2.3 | A | --- |
| 252075_at | 4315.9 | P | 4554.6 | P | 3519.5 | P | 3534.5 | P | --- |
| 252076_at | 925.2 | P | 625 | P | 637.7 | P | 505.9 | P | --- |
| 252077_at | 211 | P | 409.2 | P | 215.8 | P | 460.8 | P | --- |
| 252078_at | 69.3 | P | 119.1 | P | 111.9 | P | 135.2 | P | IMK2 |
| 252079_at | 605 | P | 625.7 | P | 790.9 | P | 631.2 | P | CPuORF44 |
| 252080_at | 1474.3 | P | 1675.4 | P | 1948.5 | P | 1727.3 | P | --- |
| 252081_at | 42.8 | P | 60.1 | A | 41.7 | A | 53.2 | P | HSFA7A |
| 252082_at | 124.1 | P | 93.5 | P | 95.1 | P | 50 | P | --- |
| 252083_at | 82.1 | P | 52.1 | P | 57.8 | P | 54.7 | P | --- |
| 252084_at | 91.3 | P | 59.9 | P | 97.5 | P | 62.4 | P | AtSAT1 |
| 252085_s_at | 1.7 | A | 2.1 | A | 16 | A | 1.2 | A | scpl37 |
| 252086_at | 771.2 | P | 642.2 | P | 433.3 | P | 634.5 | P | --- |
| 252087_at | 2.1 | A | 1.4 | A | 3.2 | A | 1.5 | A | chx28 |
| 252088_at | 208.9 | P | 194.1 | P | 195.7 | P | 247 | P | --- |
| 252089_at | 130.9 | P | 240.1 | P | 122.9 | P | 246.7 | P | --- |
| 252090_at | 8 | A | 2.5 | A | 3 | A | 3.9 | A | --- |

| | | | | | | | | | |
|---|---|---|---|---|---|---|---|---|---|
| 252091_at | 497.8 | P | 521.4 | P | 540.3 | P | 520.3 | P | --- |
| 252092_at | 2331.7 | P | 2078.2 | P | 2254.6 | P | 1594.5 | P | SSL4 |
| 252093_at | 238.8 | P | 189.2 | P | 152.6 | P | 165.5 | P | --- |
| 252094_at | 2447.8 | P | 2903.4 | P | 3081.1 | P | 3000.5 | P | PAD1 |
| 252095_at | 1639.8 | P | 1186 | P | 1168.4 | P | 1184.6 | P | --- |
| 252096_at | 66.5 | P | 101.9 | P | 72.9 | P | 78.9 | P | --- |
| 252097_at | 157.4 | P | 122 | P | 167.4 | P | 151.8 | P | --- |
| 252098_at | 4.6 | A | 71.2 | A | 3.9 | A | 41.8 | A | --- |
| 252099_at | 422.2 | P | 513.4 | P | 369.3 | P | 487.4 | P | --- |
| 252100_at | 81.9 | P | 107.9 | P | 85.1 | P | 113.5 | P | --- |
| 252101_at | 75.2 | P | 125.5 | P | 62.1 | P | 118.2 | P | ROP1 |
| 252102_at | 990.9 | P | 530.7 | P | 343.7 | P | 676.8 | P | LTI30 |
| 252103_at | 54.2 | A | 51.8 | M | 75.6 | A | 41.8 | A | --- |
| 252104_at | 521.2 | P | 601.2 | P | 484 | P | 442.6 | P | RHD4 |
| 252105_at | 167.9 | P | 156.6 | P | 154 | P | 186.9 | P | --- |
| 252106_at | 94.6 | P | 135.2 | P | 89.6 | A | 127.6 | P | ATGLR3.6 |
| 252107_at | 38.9 | A | 60.7 | P | 35.2 | A | 45.1 | A | TMT3 |
| 252108_at | 194.2 | P | 161.9 | P | 155 | P | 189.2 | P | --- |
| 252109_at | 2.2 | A | 1.8 | A | 45.8 | A | 5.1 | A | --- |
| 252110_at | 1.6 | A | 7.4 | A | 11.3 | A | 10.4 | A | --- |
| 252111_at | 11.1 | A | 8 | A | 13 | A | 1.9 | A | --- |
| 252112_at | 341.1 | P | 371.6 | P | 427.8 | P | 409.6 | P | --- |
| 252113_at | 258.3 | P | 280.8 | P | 319.8 | P | 351.7 | P | --- |
| 252114_at | 1948 | P | 683.4 | P | 1450.9 | P | 854.2 | P | --- |
| 252115_at | 9157.9 | P | 10179.7 | P | 9422.6 | P | 9439.7 | P | LTP5 |
| 252116_at | 2874 | P | 2963.1 | P | 3105.8 | P | 2893.6 | P | --- |
| 252117_at | 1912.3 | P | 1530.5 | P | 1592.1 | P | 1343.3 | P | YLS2 |
| 252118_at | 96.8 | P | 167.4 | P | 118.7 | P | 89.1 | P | --- |
| 252119_at | 223.7 | P | 193.3 | P | 263.8 | P | 260.2 | P | ATTRX1 |
| 252120_at | 1009.3 | P | 944.4 | P | 831.7 | P | 872.9 | P | --- |
| 252121_at | 1078.7 | P | 1101.5 | P | 1026.6 | P | 997.1 | P | MUR1 |
| 252122_at | 1337.7 | P | 1680.9 | P | 1793.1 | P | 1277.5 | P | --- |
| 252123_at | 1793.9 | P | 931 | P | 2853.4 | P | 2228.6 | P | F3H |
| 252124_at | 1422.8 | P | 1706.7 | P | 1757.7 | P | 1615.6 | P | --- |
| 252125_at | 429.3 | P | 518.6 | P | 422.9 | P | 421.3 | P | RTH |
| 252126_at | 1005.2 | P | 1551.1 | P | 901.8 | P | 1215 | P | --- |
| 252127_at | 136.1 | P | 139.8 | P | 163.7 | P | 144.4 | P | PLP3a |
| 252128_at | 35 | A | 2.2 | A | 31.1 | A | 39.6 | A | MNP |
| 252129_at | 35.2 | A | 51.6 | A | 40.4 | A | 59.2 | A | AtHB28 |
| 252130_at | 6028.3 | P | 6591.8 | P | 7102.6 | P | 6378.6 | P | PSBO2 |
| 252131_at | 640.7 | P | 574.8 | P | 354.4 | P | 327 | P | BCS1 |
| 252132_at | 626.6 | P | 693.4 | P | 690.2 | P | 728.8 | P | --- |
| 252133_at | 266.3 | P | 192.8 | P | 121.3 | P | 104.2 | P | --- |
| 252134_at | 1023.2 | P | 753.6 | P | 941 | P | 809.1 | P | --- |
| 252135_at | 1015.2 | P | 1074.5 | P | 739.5 | P | 828.4 | P | COR413-PM2 |
| 252136_at | 1067 | P | 500.6 | P | 856.2 | P | 377.8 | P | --- |
| 252137_at | 18.2 | A | 26.3 | A | 18.5 | A | 20.3 | A | XERO1 |
| 252138_at | 19.4 | A | 29.1 | A | 27.9 | A | 20 | A | --- |
| 252139_at | 740.7 | P | 956.8 | P | 967.2 | P | 957.3 | P | --- |
| 252140_at | 14.5 | A | 12.8 | A | 32.8 | A | 14.9 | A | --- |
| 252141_at | 875.6 | P | 864.4 | P | 738.8 | P | 760.4 | P | --- |
| 252142_at | 244.6 | P | 239.4 | P | 225.5 | P | 202.6 | P | --- |
| 252143_at | 77 | P | 97.7 | P | 82 | P | 99.8 | P | --- |
| 252144_at | 62.9 | A | 51.2 | A | 73.2 | A | 48.2 | A | --- |
| 252145_at | 9.8 | A | 9.8 | A | 28.4 | A | 16.2 | A | --- |
| 252146_at | 14.7 | A | 1.7 | A | 2.7 | A | 1.2 | A | --- |
| 252147_at | 277 | P | 247.5 | P | 238 | P | 249.4 | P | --- |
| 252148_at | 144.4 | P | 245 | P | 194.1 | P | 242.3 | P | --- |
| 252149_at | 50.9 | P | 87.6 | P | 49.5 | P | 111.4 | P | --- |
| 252150_at | 1022.2 | P | 887.3 | P | 1007.8 | P | 930.4 | P | VPS35C |
| 252151_at | 128.1 | P | 107 | P | 109.2 | P | 193.5 | P | --- |
| 252152_at | 20.8 | A | 22.6 | A | 48.2 | A | 20.3 | A | --- |
| 252153_at | 15.5 | A | 1.3 | A | 3.3 | A | 1.9 | A | --- |
| 252154_at | 615 | P | 419.2 | P | 605.5 | P | 429.3 | P | --- |
| 252155_at | 436.5 | P | 465.4 | P | 508.9 | P | 505.5 | P | --- |
| 252156_at | 28.1 | A | 25.4 | A | 34.1 | A | 34.6 | A | --- |
| 252157_at | 196.5 | P | 218.4 | P | 208.1 | P | 169.3 | P | --- |
| 252158_at | 662.4 | P | 750.8 | P | 754.1 | P | 752.1 | P | CRK |
| 252159_at | 19.4 | A | 34.1 | A | 34.3 | A | 37.1 | A | --- |
| 252160_at | 205.9 | P | 211.5 | P | 163.3 | P | 215 | P | --- |
| 252161_at | 1.7 | A | 1.8 | A | 2.9 | A | 1.9 | A | --- |
| 252162_at | 60 | A | 57 | P | 104.3 | A | 107.6 | P | --- |
| 252163_at | 10.9 | A | 2.5 | A | 32.4 | M | 9.4 | A | --- |
| 252164_at | 192.7 | P | 148.1 | P | 204.2 | P | 155.1 | P | --- |
| 252165_at | 164.3 | P | 199.1 | P | 115.1 | P | 213.2 | P | --- |
| 252166_at | 2300 | P | 2491.6 | P | 2138.3 | P | 2582.3 | P | SNRK2.2 |
| 252167_at | 103 | P | 147.4 | P | 132.7 | P | 157.4 | P | --- |
| 252168_at | 1071.2 | P | 1381.2 | P | 1154.3 | P | 1130.1 | P | MES10 |
| 252169_at | 625.7 | P | 617.2 | P | 583 | P | 552.4 | P | --- |
| 252170_at | 2794.5 | P | 3222.6 | P | 2064.4 | P | 2386.3 | P | HR4 |
| 252171_at | 681.9 | P | 720.6 | P | 776.7 | P | 651.1 | P | --- |
| 252172_at | 22.6 | A | 35.1 | A | 40.1 | A | 34.3 | A | --- |
| 252173_at | 145.8 | P | 164.6 | P | 178 | P | 169.1 | P | --- |
| 252174_at | 383.6 | P | 368.6 | P | 398.7 | P | 381 | P | --- |
| 252175_at | 328.4 | P | 342 | P | 375.9 | P | 286.8 | P | AtIDD2 |
| 252176_at | 6.7 | A | 3.5 | A | 4.2 | A | 4.2 | A | --- |
| 252177_at | 3.9 | A | 1.6 | A | 3.7 | A | 3.8 | A | --- |
| 252178_at | 152.1 | P | 201.3 | P | 124.7 | P | 152.8 | P | --- |
| 252179_at | 852.3 | P | 397.3 | P | 663.7 | P | 360.2 | P | GATL2 |
| 252180_at | 157.4 | P | 216 | P | 239 | P | 195.7 | P | KRP2 |
| 252181_at | 2613.2 | P | 3192.9 | P | 2520.8 | P | 3081.4 | P | --- |
| 252182_at | 1142.5 | P | 977.3 | P | 1428.3 | P | 1111.1 | P | U1-70K |
| 252183_at | 589.1 | P | 808.8 | P | 535.4 | P | 729.5 | P | UGT72E1 |
| 252184_at | 308.7 | P | 216.8 | P | 261.6 | P | 203 | P | DWF4 |
| 252185_at | 288.7 | P | 317.3 | P | 311.2 | P | 432.8 | P | --- |
| 252186_at | 189.3 | P | 206.6 | P | 239 | P | 173.9 | P | --- |
| 252187_at | 102.8 | P | 93.6 | P | 75.1 | M | 106.1 | P | --- |
| 252188_at | 266.8 | P | 269.6 | P | 249 | P | 320.1 | P | --- |
| 252189_at | 488.1 | P | 689 | P | 642 | P | 688.4 | P | CYCD3;3 |
| 252190_at | 14.9 | A | 1.3 | A | 5.7 | A | 2.7 | A | --- |
| 252191_at | 24.6 | P | 27.2 | A | 38.5 | P | 28.2 | P | --- |
| 252192_at | 1244.2 | P | 1287.4 | P | 1067.6 | P | 1330.9 | P | CKA2 |
| 252193_at | 273 | P | 335.3 | P | 271.8 | P | 259.6 | P | MYB77 |
| 252194_at | 127.1 | P | 147.4 | P | 199.2 | P | 139.4 | P | ATPEN3 |
| 252195_at | 0.8 | A | 4.8 | A | 4.5 | A | 1.8 | A | --- |
| 252196_at | 18.7 | A | 2.7 | A | 2.3 | A | 3.2 | A | --- |
| 252197_at | 4.1 | A | 3.3 | A | 6.4 | A | 5.2 | A | --- |
| 252198_x_at | 132.5 | P | 98.7 | M | 178.2 | P | 126.2 | A | --- |
| 252199_at | 562.7 | P | 519.4 | P | 523.9 | P | 501.8 | P | --- |

| | | | | | | | | | |
|---|---|---|---|---|---|---|---|---|---|
| 252200_at | 116.7 | P | 48.4 | P | 74.3 | P | 60 | P | --- |
| 252201_at | 1.5 | A | 10.4 | A | 3.8 | A | 2.4 | A | --- |
| 252202_at | 6.7 | A | 29.4 | A | 39.7 | A | 47.6 | A | --- |
| 252203_at | 12.4 | A | 3.5 | A | 2.1 | A | 23.7 | A | --- |
| 252204_at | 204.1 | P | 258.6 | P | 210.3 | P | 203.2 | P | --- |
| 252205_at | 99.5 | P | 205.9 | P | 116.9 | P | 123.3 | P | --- |
| 252206_at | 1235.9 | P | 1220.4 | P | 1334.8 | P | 1222.8 | P | ATCEN2 |
| 252207_at | 382.9 | P | 305 | P | 338.1 | P | 323.8 | P | --- |
| 252208_at | 387.1 | P | 353.9 | P | 469.1 | P | 351.5 | P | --- |
| 252209_at | 22 | A | 49.3 | P | 4.7 | A | 38.6 | A | --- |
| 252210_at | 225.5 | P | 120.5 | P | 156.8 | P | 173.8 | P | OBP1 |
| 252211_at | 13.7 | A | 7 | A | 8.3 | A | 4 | A | --- |
| 252212_at | 47.8 | A | 47.5 | A | 46.3 | A | 36.1 | A | MAPKKK20 |
| 252213_at | 769.6 | P | 829.8 | P | 731.1 | P | 845.5 | P | --- |
| 252214_at | 129.4 | P | 81.1 | P | 70 | P | 83.7 | P | CEJ1 |
| 252215_at | 110 | P | 137.8 | P | 132.1 | P | 122.3 | P | KICP-02 |
| 252216_at | 67.4 | P | 73.8 | P | 57.7 | M | 58.4 | P | --- |
| 252217_at | 19.8 | A | 28.8 | A | 22.6 | A | 22.2 | M | --- |
| 252218_at | 2.2 | A | 1.2 | A | 42.7 | A | 1.2 | A | --- |
| 252219_at | 21.9 | A | 2.9 | A | 18.4 | A | 5.3 | A | --- |
| 252220_at | 440.3 | P | 223.7 | P | 500.4 | P | 378.2 | P | LBD38 |
| 252221_at | 26.5 | A | 14.9 | A | 3.6 | A | 20.5 | A | --- |
| 252222_at | 3.1 | A | 7 | A | 5.8 | A | 10.7 | A | --- |
| 252223_at | 146 | P | 217.7 | P | 168.1 | P | 171.2 | P | TRB3 |
| 252224_at | 5 | A | 43.3 | A | 27.2 | A | 51.1 | A | ATARLA1B |
| 252225_at | 235.6 | P | 495.4 | P | 363.1 | P | 332.3 | P | ATARLA1C |
| 252226_at | 243.3 | P | 309.8 | P | 212.6 | P | 280 | P | --- |
| 252227_at | 63.3 | P | 65.5 | P | 54.2 | A | 76.1 | P | --- |
| 252228_at | 64.8 | A | 88 | P | 77.2 | A | 71.4 | P | VDAC5 |
| 252229_at | 203.8 | P | 153.9 | P | 206.6 | P | 190 | P | --- |
| 252230_at | 188.8 | P | 219.8 | P | 204.8 | P | 218.9 | P | --- |
| 252231_at | 1954.2 | P | 2373.4 | P | 2277.3 | P | 2627.8 | P | --- |
| 252232_at | 21.4 | A | 8.8 | A | 35.2 | A | 2.6 | A | AtbZIP5 |
| 252233_at | 21.5 | A | 39.2 | P | 62.3 | P | 49.3 | P | MYB84 |
| 252234_at | 286.1 | P | 132.1 | P | 212.2 | P | 125.1 | P | ATPSK4 |
| 252235_at | 13119.6 | P | 11808.2 | P | 15859.3 | P | 13140.7 | P | --- |
| 252236_at | 38.6 | A | 67.3 | P | 102 | M | 54.4 | A | --- |
| 252237_at | 17.5 | A | 30.6 | A | 33.9 | A | 23.9 | A | --- |
| 252238_at | 1.2 | A | 1.8 | A | 1.8 | A | 1.8 | A | --- |
| 252239_at | 481.1 | P | 520.2 | P | 510.9 | P | 499.8 | P | --- |
| 252240_at | 16 | A | 36.2 | P | 64.6 | A | 50.6 | P | --- |
| 252241_at | 62.3 | P | 98.2 | P | 90.4 | P | 113.4 | P | --- |
| 252242_at | 314.8 | P | 315.4 | P | 202.3 | P | 336.6 | P | VFB2 |
| 252243_at | 10.3 | A | 5.9 | A | 19.8 | A | 3.1 | A | --- |
| 252244_at | 11.8 | M | 11.6 | A | 1 | A | 1.4 | A | --- |
| 252245_at | 111 | P | 128.8 | P | 104.1 | P | 124 | P | --- |
| 252246_at | 215.1 | P | 236.6 | P | 211.9 | P | 246.3 | P | --- |
| 252247_at | 94.6 | P | 80.3 | P | 104.3 | P | 95.8 | P | --- |
| 252248_at | 30.9 | P | 93.7 | P | 75.1 | P | 98.5 | P | AtRLP44 |
| 252249_at | 7.5 | A | 23.3 | A | 8.3 | A | 33.2 | A | --- |
| 252250_at | 108.2 | P | 133.6 | P | 101 | P | 91.3 | P | --- |
| 252251_at | 19.6 | A | 31.9 | P | 9.5 | A | 31.1 | P | --- |
| 252252_at | 323.4 | P | 317.4 | P | 290.1 | P | 294.5 | P | RID3 |
| 252253_at | 34.5 | A | 20 | A | 44.7 | A | 27.7 | A | --- |
| 252254_at | 116.3 | P | 200.7 | P | 157.6 | P | 131.2 | P | DMS3 |
| 252255_at | 1546.4 | P | 2315.7 | P | 1624.4 | P | 2401.3 | P | --- |
| 252256_at | 503.7 | P | 592.5 | P | 666.9 | P | 650.8 | P | SRp34a |
| 252257_at | 2.5 | A | 1.5 | A | 5.6 | A | 2.1 | A | --- |
| 252258_at | 7.5 | A | 6.6 | A | 12.3 | A | 2.6 | A | --- |
| 252259_at | 10.8 | A | 3.3 | A | 1.4 | A | 1.8 | A | --- |
| 252260_at | 9.6 | A | 40.2 | A | 22.1 | A | 8 | A | --- |
| 252261_at | 346.6 | P | 370 | P | 344.8 | P | 393 | P | RDR6 |
| 252262_at | 30.6 | A | 23.5 | A | 7.4 | A | 3.8 | A | --- |
| 252263_at | 2.8 | A | 4 | A | 6.7 | A | 4.2 | A | --- |
| 252264_at | 330.9 | P | 270.7 | P | 264 | P | 212.4 | P | --- |
| 252265_at | 13.7 | A | 1.5 | A | 3.9 | A | 1.3 | A | DIN11 |
| 252266_at | 25.3 | A | 27.2 | A | 48.5 | A | 46 | M | --- |
| 252267_at | 233.5 | P | 242.7 | P | 185.2 | P | 247.9 | P | --- |
| 252268_at | 48.3 | P | 114 | P | 85.4 | P | 108.9 | P | --- |
| 252269_at | 45.3 | P | 56.4 | P | 109.9 | P | 52.2 | P | LSU1 |
| 252270_at | 1276.2 | P | 1389.7 | P | 1719 | P | 1350.5 | P | --- |
| 252271_s_at | 760.3 | P | 835 | P | 897.6 | P | 839.8 | P | --- |
| 252272_at | 495.3 | P | 709 | P | 667.1 | P | 726.7 | P | BAM2 |
| 252273_at | 193.3 | P | 167.6 | M | 213.3 | A | 243.9 | M | --- |
| 252274_at | 3575.2 | P | 3102.2 | P | 3807.3 | P | 3087.9 | P | BCAT3 |
| 252275_at | 185.5 | P | 187.2 | P | 179.7 | P | 200.8 | P | UBP26 |
| 252276_at | 872.7 | P | 809.3 | P | 828 | P | 801.5 | P | --- |
| 252277_at | 1539.2 | P | 1829.2 | P | 1849.7 | P | 2012.3 | P | NACA2 |
| 252278_at | 634.1 | P | 687.5 | P | 540.7 | P | 644.3 | P | anac062 |
| 252279_at | 1.3 | A | 27.5 | A | 23.3 | A | 5.6 | A | ACS9 |
| 252280_at | 649.1 | P | 681.5 | P | 753.4 | P | 759.1 | P | iqd21 |
| 252281_at | 237.9 | P | 222.9 | P | 280.8 | P | 280.5 | P | --- |
| 252282_at | 480.9 | P | 673.5 | P | 494.1 | P | 537.7 | P | --- |
| 252283_at | 41.6 | P | 26.9 | M | 27.9 | A | 12.1 | A | --- |
| 252284_at | 10.4 | A | 42.5 | A | 3.9 | A | 25 | A | --- |
| 252285_at | 253.1 | P | 270.6 | P | 212.9 | P | 316.2 | P | --- |
| 252286_at | 53.6 | A | 68.8 | P | 59.6 | A | 47.2 | A | --- |
| 252287_at | 618.8 | P | 718.5 | P | 817.5 | P | 727.2 | P | --- |
| 252288_at | 71.4 | P | 132.9 | P | 72.8 | A | 96.6 | P | --- |
| 252289_at | 13.8 | A | 22.2 | A | 27.8 | A | 3 | A | --- |
| 252290_at | 600.1 | P | 497.5 | P | 447.2 | P | 483.8 | P | --- |
| 252291_s_at | 4801.6 | P | 3658.5 | P | 3707.7 | P | 3989.5 | P | PRXCB |
| 252292_at | 396.8 | P | 371.7 | P | 375.3 | P | 349.5 | P | --- |
| 252293_at | 7485.1 | P | 5122.7 | P | 5177 | P | 4409.8 | P | --- |
| 252294_at | 9862.7 | P | 9737.3 | P | 11997.7 | P | 10455.8 | P | ATBBC1 |
| 252295_at | 548.3 | P | 397.8 | P | 377.6 | P | 415.5 | P | --- |
| 252296_at | 27 | A | 68.7 | P | 72.6 | P | 93.4 | P | --- |
| 252297_at | 5320.8 | P | 5210 | P | 4300.5 | P | 5144.9 | P | EMB1080 |
| 252298_at | 122.6 | P | 188.8 | P | 128 | P | 148.1 | P | --- |
| 252299_at | 36.9 | A | 29.7 | A | 58.4 | A | 24.7 | A | --- |
| 252300_at | 157.5 | P | 74.4 | P | 143.7 | P | 105.8 | P | --- |
| 252301_at | 1.2 | A | 2 | A | 50.9 | A | 4.5 | A | --- |
| 252302_at | 23.4 | A | 9.8 | A | 13 | A | 64.8 | A | --- |
| 252303_at | 245.1 | P | 155.2 | P | 153.6 | P | 121.3 | P | --- |
| 252304_at | 6.9 | A | 38.5 | A | 7.2 | A | 13.5 | A | --- |
| 252305_at | 167 | P | 179.3 | P | 281.2 | P | 236.4 | P | emb1796 |
| 252306_at | 22.6 | A | 10.9 | A | 23.2 | A | 23.3 | M | --- |
| 252307_at | 25.7 | A | 17.5 | A | 43.8 | A | 25.5 | A | --- |
| 252308_at | 744.1 | P | 853.6 | P | 794.3 | P | 882.2 | P | --- |

| | | | | | | | | | |
|---|---|---|---|---|---|---|---|---|---|
| 252309_at | 349 | P | 71.2 | P | 205 | P | 172.4 | P | --- |
| 252310_at | 278.8 | P | 375.5 | P | 380.2 | P | 365 | P | --- |
| 252311_at | 85.5 | P | 126 | P | 128.6 | P | 115.4 | P | --- |
| 252312_at | 28.2 | A | 17.5 | A | 9.2 | A | 24.8 | A | iqd15 |
| 252313_at | 578.1 | P | 688.3 | P | 560.4 | P | 653.5 | P | CID10 |
| 252314_at | 26.1 | A | 21.3 | A | 27.6 | A | 27.1 | A | --- |
| 252315_at | 4029.7 | P | 3201.8 | P | 4092.2 | P | 3697.9 | P | CXE12 |
| 252316_at | 61.6 | P | 87.2 | P | 116 | P | 99.4 | P | ATCXE13 |
| 252317_at | 118.7 | P | 174.3 | P | 140.8 | P | 165 | P | --- |
| 252318_at | 2548.8 | P | 3546.1 | P | 2829.2 | P | 3485.7 | P | GSA2 |
| 252319_at | 108.3 | P | 117.8 | P | 122.3 | P | 149.3 | P | --- |
| 252320_at | 57.1 | A | 26.9 | A | 21.8 | A | 38.5 | A | --- |
| 252321_at | 52.5 | P | 50.8 | P | 62.4 | P | 53.9 | P | --- |
| 252322_at | 58.6 | P | 66 | P | 85 | P | 58.4 | P | --- |
| 252323_at | 1411.9 | P | 1022.4 | P | 876.5 | P | 1058.4 | P | KING1 |
| 252324_at | 576.6 | P | 411 | P | 315 | P | 356.8 | P | --- |
| 252325_at | 4362.1 | P | 4719.6 | P | 5532.6 | P | 4741.7 | P | CSR1 |
| 252326_at | 1698.8 | P | 1676.5 | P | 1657.6 | P | 1537.5 | P | GAMMA CAL2 |
| 252327_at | 5214.4 | P | 4634.7 | P | 6672.8 | P | 5115.5 | P | --- |
| 252328_at | 971.1 | P | 894.2 | P | 940.5 | P | 909.8 | P | --- |
| 252329_at | 678.4 | P | 626.7 | P | 451.3 | P | 632.8 | P | --- |
| 252330_at | 2.4 | A | 11.6 | A | 1.6 | A | 3.7 | A | --- |
| 252331_s_at | 463.6 | P | 371.8 | P | 353.2 | P | 320.1 | P | SPT1 |
| 252332_at | 55.7 | A | 57.1 | P | 54.5 | P | 64.9 | P | --- |
| 252333_at | 41.2 | P | 49.5 | P | 75.8 | P | 41.9 | P | --- |
| 252334_at | 204.6 | P | 36.9 | A | 101.7 | A | 40.8 | A | --- |
| 252335_at | 399.7 | P | 452.5 | P | 388 | P | 347.1 | P | --- |
| 252336_at | 271.5 | P | 322.6 | P | 229.8 | P | 333.2 | P | --- |
| 252337_at | 964.7 | P | 1002.7 | P | 1090 | P | 1030.6 | P | CDC2 |
| 252338_at | 1405.5 | P | 1192.7 | P | 1250.6 | P | 1122.4 | P | MAPR3 |
| 252339_at | 51.4 | A | 51.9 | A | 38.2 | A | 46.1 | A | --- |
| 252340_at | 4.2 | A | 40.7 | A | 41.4 | A | 29.3 | A | AtMYB45 |
| 252341_at | 27.9 | M | 19.6 | A | 65.4 | A | 17.3 | P | --- |
| 252342_at | 27.2 | A | 3.1 | A | 9.2 | A | 23.1 | A | --- |
| 252343_at | 1125.1 | P | 1383.2 | P | 1597.4 | P | 1492.4 | P | --- |
| 252344_s_at | 380.8 | P | 471.9 | P | 492.3 | P | 475 | P | --- |
| 252345_at | 42.2 | A | 86.3 | P | 26.6 | A | 47.1 | P | --- |
| 252346_at | 16.6 | A | 127.1 | P | 23.1 | A | 72.3 | P | --- |
| 252347_at | 1.6 | A | 1.1 | A | 2.1 | A | 1.3 | A | --- |
| 252348_at | 9965.5 | P | 9763.6 | P | 10844 | P | 10708 | P | --- |
| 252349_at | 203.8 | P | 243.6 | P | 263.8 | P | 234.8 | P | APC8 |
| 252350_at | 304.8 | P | 285.2 | P | 265.8 | P | 254 | P | ATM |
| 252351_at | 119.8 | P | 148.3 | P | 179.1 | A | 153.4 | P | --- |
| 252352_at | 92.6 | P | 156.7 | P | 124.5 | P | 174.4 | P | --- |
| 252353_at | 1847.2 | P | 1933.7 | P | 2236.8 | P | 2071.1 | P | --- |
| 252354_at | 1991.6 | P | 1755.2 | P | 1521.8 | P | 1654.9 | P | ALDH10A9 |
| 252355_at | 179.5 | P | 143.4 | P | 212.7 | P | 169.1 | P | --- |
| 252356_at | 14.4 | A | 7.3 | A | 2 | A | 12.6 | A | --- |
| 252357_at | 107.1 | A | 145.6 | P | 120.8 | P | 127.2 | P | --- |
| 252358_at | 101.5 | P | 78.9 | P | 117.3 | P | 117.4 | P | --- |
| 252359_at | 238 | P | 271.5 | P | 319.6 | P | 267.2 | P | --- |
| 252360_at | 5.8 | A | 19.8 | A | 42 | A | 31.8 | A | --- |
| 252361_at | 122 | P | 165.6 | P | 134 | M | 180 | P | --- |
| 252362_at | 1032 | P | 915.3 | P | 781.4 | P | 967.6 | P | PDE312 |
| 252363_at | 332.9 | P | 577.5 | P | 381 | P | 685.9 | P | --- |
| 252364_at | 64.3 | P | 14.8 | A | 53.7 | A | 34.7 | A | --- |
| 252365_at | 744.5 | P | 307.2 | P | 441.6 | P | 408.9 | P | --- |
| 252366_at | 2202.7 | P | 2434.6 | P | 2132.7 | P | 2539 | P | --- |
| 252367_at | 24.3 | A | 1.9 | A | 36.6 | A | 6.5 | A | BT2 |
| 252368_at | 441.1 | P | 77.3 | P | 169.3 | P | 91.2 | P | CYP94B3 |
| 252369_at | 46.8 | A | 33.7 | A | 9.3 | A | 70.6 | P | --- |
| 252370_at | 76.3 | A | 65 | A | 52.9 | A | 58 | A | --- |
| 252371_at | 880.4 | P | 754.1 | P | 798.2 | P | 668.8 | P | --- |
| 252372_at | 6047.4 | P | 4848.3 | P | 5222.8 | P | 4737.8 | P | ALDH2B4 |
| 252373_at | 342.4 | P | 592.3 | P | 302.5 | P | 441.7 | P | EDS1 |
| 252374_at | 159.5 | P | 275.5 | P | 172.3 | P | 197.6 | P | ARR5 |
| 252375_at | 218.8 | P | 247.9 | P | 222.5 | P | 228.3 | P | ROP10 |
| 252376_at | 627.4 | P | 531.8 | P | 539.6 | P | 605.3 | P | ATGLDH |
| 252377_at | 1022.2 | P | 1035.2 | P | 1189 | P | 1072.8 | P | --- |
| 252378_at | 194.1 | P | 117.3 | P | 189 | P | 129.8 | P | --- |
| 252379_at | 615.9 | P | 531.3 | P | 495.3 | P | 489.2 | P | ATH1 |
| 252380_at | 2.2 | A | 11.4 | A | 10.5 | A | 13.8 | A | ATH2 |
| 252381_s_at | 61.4 | P | 69.9 | P | 82.8 | P | 60.6 | P | ABCA4 |
| 252382_at | 8.4 | A | 4.5 | A | 20.7 | A | 8 | A | ATH5 |
| 252383_at | 153.9 | P | 108.2 | P | 106 | P | 90.8 | P | ATH6 |
| 252384_at | 4.4 | A | 4 | A | 31.1 | A | 9.7 | A | ATH7 |
| 252385_at | 2591.5 | P | 2931 | P | 2303.1 | P | 2988.7 | P | MAG1 |
| 252386_at | 92 | P | 57.3 | A | 118.2 | A | 69.9 | M | --- |
| 252387_at | 1771.5 | P | 1294.4 | P | 1399.1 | P | 1386.3 | P | --- |
| 252388_at | 196 | P | 205.7 | P | 210.9 | P | 202.8 | P | --- |
| 252389_at | 2347.4 | P | 2318.1 | P | 2493 | P | 2578.5 | P | --- |
| 252390_at | 1298.9 | P | 1429.7 | P | 1227.8 | P | 1290.8 | P | --- |
| 252391_at | 2298.1 | P | 3007.7 | P | 2261.8 | P | 3206.2 | P | --- |
| 252392_s_at | 341.3 | P | 379.9 | P | 304.5 | P | 348.7 | P | --- |
| 252393_at | 63.5 | P | 78.8 | P | 81.6 | P | 66.7 | P | --- |
| 252394_at | 165 | P | 157.1 | P | 133.5 | P | 173.8 | P | --- |
| 252395_at | 31.3 | A | 76.2 | P | 16.8 | A | 47.2 | P | AHA4 |
| 252396_at | 2.8 | A | 3.2 | A | 8.3 | A | 15.5 | A | --- |
| 252397_at | 7.4 | A | 86.1 | A | 77.8 | A | 72.8 | A | --- |
| 252398_at | 482.4 | P | 410 | P | 371.7 | P | 454 | P | --- |
| 252399_at | 6.8 | A | 5 | A | 2.3 | A | 2.8 | A | CNGC16 |
| 252400_at | 153 | P | 157.4 | P | 113.2 | P | 145.9 | P | --- |
| 252401_at | 20 | A | 2.4 | A | 35.4 | A | 54.1 | P | --- |
| 252402_s_at | 407.5 | P | 471.5 | P | 404.1 | P | 404.4 | P | --- |
| 252403_at | 19.2 | A | 63.8 | P | 18.3 | P | 58.3 | P | --- |
| 252404_at | 747.6 | P | 776.8 | P | 545.8 | P | 769.7 | P | EDD1 |
| 252405_at | 116.4 | P | 131 | P | 120 | A | 113.2 | P | --- |
| 252406_at | 458 | P | 542.6 | P | 375.6 | P | 532.8 | P | --- |
| 252407_at | 2576.3 | P | 2492.5 | P | 2826.9 | P | 2343.4 | P | MDH |
| 252408_at | 139 | P | 146.3 | P | 143 | P | 148.8 | P | ATMYB94 |
| 252409_at | 7431.1 | P | 7170.7 | P | 6660.7 | P | 6460.3 | P | --- |
| 252410_at | 602.7 | P | 625.1 | P | 669.7 | P | 629.4 | P | NOA1 |
| 252411_at | 839.2 | P | 822.8 | P | 758.6 | P | 674.2 | P | PEX11B |
| 252412_at | 390 | P | 372.1 | P | 417.4 | P | 427.4 | P | --- |
| 252413_at | 4447.2 | P | 3796.7 | P | 5206.8 | P | 4135.7 | P | --- |
| 252414_at | 622.2 | P | 587 | P | 498.1 | P | 647 | P | --- |
| 252415_at | 256.6 | P | 73.3 | P | 159.8 | P | 92.2 | P | ASN1 |
| 252416_at | 121.5 | P | 82.1 | P | 101.1 | P | 84 | P | ATSMC2 |
| 252417_at | 420.8 | P | 718.8 | P | 209.6 | P | 396.6 | P | --- |

| | | | | | | | | | |
|---|---|---|---|---|---|---|---|---|---|
| 252418_at | 562.3 | P | 552.4 | P | 453 | P | 456.4 | P | --- |
| 252419_at | 262.3 | P | 250.3 | P | 264.8 | P | 290.4 | P | --- |
| 252420_at | 148.3 | P | 150.1 | P | 136.1 | P | 143.3 | M | --- |
| 252421_at | 609 | P | 386 | P | 558.8 | P | 267.5 | P | --- |
| 252422_at | 864.1 | P | 759.1 | P | 643.2 | P | 772.8 | P | --- |
| 252423_at | 509.3 | P | 550.1 | P | 456.8 | P | 500.7 | P | --- |
| 252424_at | 413 | P | 437.1 | P | 378 | P | 347.7 | P | --- |
| 252425_at | 459.1 | P | 557.6 | P | 433.1 | P | 518.4 | P | AtTCP14 |
| 252426_at | 232.9 | P | 208.1 | P | 287.6 | P | 225.3 | P | --- |
| 252427_at | 151.5 | P | 404 | P | 278.1 | P | 336.4 | P | --- |
| 252428_at | 3.7 | A | 9.9 | A | 10.6 | A | 27 | A | --- |
| 252429_at | 439.1 | P | 881.7 | P | 455.3 | P | 677 | P | CDF3 |
| 252430_at | 13026.3 | P | 15928.3 | P | 14082.3 | P | 13874.4 | P | LHCA4 |
| 252431_at | 182.2 | P | 177.6 | P | 87 | P | 141.5 | P | MAG2 |
| 252432_at | 111 | P | 69.8 | P | 119.5 | P | 104 | P | --- |
| 252433_at | 451.1 | P | 490.6 | P | 328.1 | P | 468.9 | P | --- |
| 252434_s_at | 33.1 | A | 57.3 | M | 115.9 | P | 75 | P | --- |
| 252435_at | 10.4 | A | 1.7 | A | 9.1 | A | 10.8 | A | --- |
| 252436_x_at | 33.8 | A | 18.6 | A | 52.3 | A | 28.6 | A | --- |
| 252437_at | 31.1 | P | 70.7 | P | 69 | P | 71.7 | P | --- |
| 252438_at | 256.6 | P | 276.5 | P | 315.1 | P | 238.5 | P | --- |
| 252439_at | 38.6 | A | 58.9 | A | 8.6 | A | 39.9 | A | --- |
| 252440_at | 4 | A | 22.1 | A | 4.8 | A | 8 | A | TIP5;1 |
| 252441_at | 3099.2 | P | 4081.1 | P | 2541.7 | P | 3862 | P | PTAC16 |
| 252442_at | 680.3 | P | 779.7 | P | 625.1 | P | 956.1 | P | --- |
| 252443_at | 469.6 | P | 434.1 | P | 536.7 | P | 448.1 | P | --- |
| 252444_at | 433 | P | 427.8 | P | 273.9 | P | 409.9 | P | PHT4;3 |
| 252445_at | 566 | P | 774.5 | P | 619.9 | P | 736.8 | P | --- |
| 252446_at | 204.7 | P | 160 | P | 154.2 | P | 152.5 | P | --- |
| 252447_at | 21.7 | A | 3.5 | A | 15 | A | 12.8 | A | --- |
| 252448_at | 2.6 | A | 12.5 | A | 17 | A | 2.3 | A | --- |
| 252449_at | 328.7 | P | 308.8 | P | 248.7 | P | 264.3 | P | ftsh7 |
| 252450_s_at | 32.2 | P | 49.4 | P | 25.6 | A | 22.8 | A | --- |
| 252451_at | 8.8 | A | 1.9 | A | 2.6 | A | 21.9 | A | --- |
| 252452_at | 43.1 | P | 39.3 | P | 44.4 | P | 54.6 | P | --- |
| 252453_at | 282.2 | P | 220 | P | 281.6 | P | 160.7 | P | --- |
| 252454_at | 17.6 | A | 16.5 | A | 26.3 | A | 19.8 | A | --- |
| 252455_at | 11.8 | A | 3 | A | 6.6 | A | 10.5 | A | --- |
| 252456_at | 18.1 | A | 25.9 | A | 33.1 | A | 3 | A | --- |
| 252457_at | 8.2 | A | 54.5 | M | 14 | A | 76.5 | P | --- |
| 252458_at | 25.8 | A | 45.9 | A | 38.9 | A | 17.5 | A | --- |
| 252459_s_at | 451.6 | P | 755.3 | P | 685.8 | P | 717.6 | P | PLC9 |
| 252460_at | 3 | A | 2.3 | A | 2.2 | A | 4.3 | A | --- |
| 252461_at | 14.7 | A | 0.5 | A | 3.9 | A | 2.5 | A | --- |
| 252462_at | 3502.8 | P | 2503 | P | 3109.9 | P | 2382.2 | P | --- |
| 252463_at | 5789.1 | P | 5334.6 | P | 5607.8 | P | 6044.7 | P | --- |
| 252464_at | 1005.8 | P | 1447.4 | P | 688.1 | P | 1073 | P | --- |
| 252465_x_at | 15.9 | A | 3.7 | A | 43.3 | A | 44.6 | A | --- |
| 252466_x_at | 2.4 | A | 3.1 | A | 4.9 | A | 16.7 | A | --- |
| 252467_at | 546.1 | P | 409.3 | P | 488.1 | P | 453 | P | --- |
| 252468_at | 8391.9 | P | 6914.6 | P | 6217.6 | P | 8579.2 | P | PHS2 |
| 252469_at | 210 | P | 139.8 | P | 260.7 | P | 141.7 | P | --- |
| 252470_at | 98.4 | P | 123.1 | P | 101.7 | P | 91.9 | P | --- |
| 252471_at | 579.5 | P | 630.6 | P | 541.8 | P | 689.6 | P | TOPP5 |
| 252472_at | 675.1 | P | 755.7 | P | 611.2 | P | 755.2 | P | ATRABA2C |
| 252473_s_at | 421.8 | P | 397.7 | P | 456.5 | P | 387.8 | P | --- |
| 252474_at | 679.4 | P | 885.5 | P | 660.9 | P | 838.6 | P | --- |
| 252475_s_at | 944.4 | P | 625.1 | P | 824.6 | P | 714.3 | P | PCL1 |
| 252476_at | 2.7 | A | 5.2 | A | 4 | A | 4.1 | A | --- |
| 252477_at | 23.2 | A | 21.5 | A | 35.6 | A | 4.3 | A | --- |
| 252478_at | 885.6 | P | 775.7 | P | 924.4 | P | 771.1 | P | --- |
| 252479_at | 2222.5 | P | 1920.1 | P | 1607 | P | 2091.4 | P | TIM9 |
| 252480_at | 75 | P | 71.4 | P | 153.6 | P | 113.9 | P | MBD5 |
| 252481_at | 979.6 | P | 1096.2 | P | 955.3 | P | 1035.1 | P | --- |
| 252482_at | 1879.7 | P | 833 | P | 1241.8 | P | 1139.6 | P | UGT76E11 |
| 252483_at | 397.9 | P | 318.9 | P | 306.2 | P | 289.1 | P | --- |
| 252484_at | 35.8 | A | 1.7 | A | 5.5 | A | 1.9 | A | --- |
| 252485_at | 949.2 | P | 960.8 | P | 1078.8 | P | 908.1 | P | RPP13 |
| 252486_at | 671.3 | P | 755.9 | P | 753.2 | P | 665.4 | P | PUB13 |
| 252487_at | 3313.4 | P | 189.6 | P | 1910.3 | P | 478.4 | P | UGT76E12 |
| 252488_at | 55 | A | 25.4 | A | 14.4 | A | 4.9 | A | --- |
| 252489_at | 31.2 | A | 28.4 | A | 37.9 | A | 27.3 | A | --- |
| 252490_at | 14.4 | A | 32.6 | P | 24.5 | P | 31.4 | P | --- |
| 252491_at | 1.3 | A | 13.8 | A | 3.9 | A | 15.9 | A | --- |
| 252492_at | 2415 | P | 2525.1 | P | 2247.8 | P | 2505.6 | P | TOC75-III |
| 252493_at | 14.2 | A | 37.9 | A | 44.9 | A | 21.7 | A | --- |
| 252494_at | 23.3 | A | 29.7 | A | 48.6 | P | 34.3 | A | --- |
| 252495_at | 10.4 | A | 17.6 | A | 24.9 | A | 17.3 | A | --- |
| 252496_at | 442.4 | P | 378.7 | P | 350.5 | P | 425.6 | P | CRR2 |
| 252497_at | 5.1 | A | 3.5 | A | 10.8 | A | 5.8 | A | --- |
| 252498_at | 47.2 | A | 3.7 | A | 7.8 | A | 22.6 | P | --- |
| 252499_s_at | 10.6 | A | 10.1 | A | 42.7 | P | 9.8 | A | --- |
| 252500_at | 19.6 | A | 12 | A | 60.9 | A | 3.2 | A | --- |
| 252501_at | 9.7 | A | 24.7 | P | 17.5 | A | 22.5 | A | --- |
| 252502_at | 214.3 | P | 275.5 | P | 422.4 | P | 294.8 | P | COPT2 |
| 252503_at | 37.8 | A | 43.1 | A | 55.2 | A | 32.8 | A | --- |
| 252504_at | 551.6 | P | 609.2 | P | 502.2 | P | 626.7 | P | TRFL1 |
| 252505_at | 0.8 | A | 1 | A | 2.1 | A | 1 | A | --- |
| 252506_at | 95.3 | P | 97.1 | P | 100.8 | P | 90.7 | P | UTR5 |
| 252507_at | 171.6 | P | 253.3 | P | 211.3 | P | 208.9 | P | aTNUDT9 |
| 252508_at | 318.9 | P | 252.7 | P | 313.8 | P | 252.7 | P | --- |
| 252509_at | 24.7 | A | 21.9 | A | 28.7 | A | 33.8 | A | --- |
| 252510_at | 9.9 | A | 14.9 | A | 6.8 | A | 15.7 | A | --- |
| 252511_at | 44.3 | P | 163.1 | P | 45.4 | P | 91 | P | --- |
| 252512_at | 204.5 | P | 180.7 | P | 192.5 | P | 266.5 | P | --- |
| 252513_at | 339.1 | P | 359 | P | 243.4 | P | 275.1 | P | --- |
| 252514_at | 1201.7 | P | 1239.2 | P | 1045.9 | P | 1206.4 | P | ATRAB8A |
| 252515_at | 65.4 | P | 165.6 | P | 89.9 | P | 124.1 | P | HSP17.4 |
| 252516_at | 762.4 | P | 865.6 | P | 854.5 | P | 826.9 | P | ATHRS1 |
| 252517_at | 27.6 | A | 17.4 | A | 13.7 | A | 18.2 | A | --- |
| 252518_at | 9.6 | A | 11.6 | A | 16.1 | A | 16.4 | A | --- |
| 252519_at | 43.4 | M | 38.2 | A | 56.9 | P | 67.1 | P | --- |
| 252520_at | 1.2 | A | 6.7 | A | 1.8 | A | 2.1 | A | --- |
| 252521_at | 7.6 | A | 12.6 | A | 17.1 | A | 12.1 | A | --- |
| 252522_at | 0.7 | A | 0.3 | A | 0.3 | A | 0.7 | A | --- |
| 252523_at | 24 | A | 14.7 | A | 5.3 | A | 9.2 | A | --- |
| 252524_at | 2469.9 | P | 2893.3 | P | 2763.2 | P | 2975.5 | P | --- |
| 252525_at | 594.3 | P | 554.9 | P | 628.7 | P | 685 | P | --- |
| 252526_at | 4.1 | A | 9.5 | A | 24.9 | A | 3 | A | --- |

| | | | | | | | | | |
|---|---|---|---|---|---|---|---|---|---|
| 252527_at | 824.1 | P | 1036.1 | P | 610.4 | P | 966.6 | P | UXS5 |
| 252528_at | 779.4 | P | 766.8 | P | 660.8 | P | 800.9 | P | UBC13 |
| 252529_at | 46.7 | A | 178.7 | P | 70.9 | A | 207 | P | --- |
| 252530_at | 2.7 | A | 6.4 | A | 24.4 | A | 3.5 | A | --- |
| 252531_at | 5.9 | A | 2.1 | A | 3.6 | A | 15.6 | A | ACT12 |
| 252532_at | 13.6 | A | 22.2 | A | 45.2 | A | 36.2 | P | --- |
| 252533_at | 234.3 | P | 192.5 | P | 169.9 | P | 180.7 | P | --- |
| 252534_at | 18.8 | A | 72.8 | P | 7.1 | A | 50.1 | P | MYB111 |
| 252535_at | 40.8 | A | 63 | A | 35.5 | A | 53 | A | --- |
| 252536_at | 7.8 | A | 5.1 | A | 19.4 | A | 5.4 | A | --- |
| 252537_at | 17.4 | A | 23.2 | A | 6.2 | A | 5.6 | A | --- |
| 252538_at | 47.4 | P | 63.7 | P | 52 | M | 16.8 | A | --- |
| 252539_at | 331 | P | 99 | P | 193.2 | P | 87.4 | P | --- |
| 252540_at | 229.2 | P | 256.4 | P | 276.4 | P | 255.3 | P | --- |
| 252541_at | 84.9 | P | 70.8 | P | 141 | P | 85.7 | P | --- |
| 252542_at | 956.7 | P | 934.5 | P | 968.4 | P | 894.7 | P | --- |
| 252543_at | 1492.8 | P | 1710.2 | P | 1464.8 | P | 1652.2 | P | PHOT1 |
| 252544_at | 9.5 | A | 3.2 | A | 5 | A | 8.9 | A | --- |
| 252545_at | 17.9 | A | 1.6 | A | 3.9 | A | 20 | A | --- |
| 252546_at | 262.1 | P | 261.5 | P | 224 | P | 200.4 | P | --- |
| 252547_at | 7.7 | A | 6.3 | A | 8.4 | A | 4.5 | A | --- |
| 252548_at | 238.7 | P | 369.6 | P | 277.3 | P | 348.5 | P | --- |
| 252549_at | 285.9 | P | 583.2 | P | 321.9 | P | 442.7 | P | --- |
| 252550_at | 37.3 | A | 46.9 | A | 73.3 | A | 66.2 | A | --- |
| 252551_at | 101 | P | 46.1 | M | 100.4 | P | 78.7 | A | --- |
| 252552_at | 295.5 | P | 291.3 | P | 188.1 | P | 269.3 | P | --- |
| 252553_at | 3.1 | A | 12.6 | A | 16.5 | A | 6.4 | A | --- |
| 252554_s_at | 16.1 | A | 1.8 | A | 14.1 | A | 0.6 | A | MEE39 |
| 252555_at | 6.6 | A | 3.5 | A | 1.7 | A | 1.6 | A | --- |
| 252556_at | 21.7 | A | 23.4 | A | 39.9 | A | 10.9 | A | --- |
| 252557_at | 2.2 | A | 22 | A | 6.8 | A | 15.2 | A | ATEXLA3 |
| 252558_at | 13.3 | A | 33 | A | 12.5 | A | 21 | P | --- |
| 252559_at | 68.7 | P | 80.3 | P | 107.9 | P | 57 | P | --- |
| 252560_at | 1110.6 | P | 1201.6 | P | 877.1 | P | 1128.1 | P | HTB11 |
| 252561_at | 1846.9 | P | 2471.5 | P | 1903.9 | P | 2424 | P | HTB9 |
| 252562_s_at | 583.4 | P | 876.5 | P | 621 | P | 857.5 | P | --- |
| 252563_at | 1128.5 | P | 1064.4 | P | 656.5 | P | 1012.6 | P | ATEXLA1 |
| 252564_at | 5252.5 | P | 4978.6 | P | 4894.5 | P | 5226 | P | ADF1 |
| 252565_at | 2732.7 | P | 2334.5 | P | 2433.3 | P | 2510.1 | P | ADF2 |
| 252566_at | 5564.7 | P | 4873.5 | P | 6510.1 | P | 5491.8 | P | RPS15AD |
| 252567_at | 19 | A | 20.2 | A | 34.4 | A | 3.6 | A | --- |
| 252568_at | 21.1 | A | 21 | A | 6.9 | A | 29.4 | A | --- |
| 252569_at | 42.5 | P | 32.9 | P | 62 | P | 51 | P | --- |
| 252570_at | 657.4 | P | 473.4 | P | 417.2 | P | 443.1 | P | IVD |
| 252571_at | 34.6 | A | 33.2 | A | 25.4 | A | 6.4 | A | SYP72 |
| 252572_at | 145.1 | P | 233.7 | P | 205.2 | P | 244.7 | P | MLO3 |
| 252573_at | 295.6 | P | 334.4 | P | 235.8 | P | 313.3 | P | --- |
| 252574_at | 22.7 | A | 35.4 | A | 46.5 | P | 34.6 | P | --- |
| 252575_at | 48.4 | A | 6.4 | A | 34.4 | A | 34.9 | A | --- |
| 252576_s_at | 483.7 | P | 700.5 | P | 735.3 | P | 667.7 | P | --- |
| 252577_at | 29 | A | 25.9 | A | 8.9 | A | 15.2 | A | --- |
| 252578_at | 11.9 | A | 16.3 | P | 14.2 | A | 16.2 | A | --- |
| 252579_at | 2.4 | A | 8.4 | A | 23.2 | A | 9.7 | A | --- |
| 252580_at | 15.5 | A | 7.6 | A | 5 | A | 2.1 | A | --- |
| 252581_at | 8.9 | A | 22.7 | A | 6.7 | A | 8 | A | --- |
| 252582_at | 9.2 | A | 1.2 | A | 49.3 | A | 1.3 | A | --- |
| 252583_at | 16.5 | A | 40.8 | A | 8.1 | A | 30.7 | A | --- |
| 252584_at | 5.8 | A | 15.8 | A | 4.4 | A | 6.2 | A | --- |
| 252585_s_at | 0.4 | A | 0.6 | A | 2.1 | A | 0.7 | A | --- |
| 252586_at | 9.4 | A | 11.2 | A | 8.5 | A | 6.7 | A | --- |
| 252587_at | 462 | P | 690.7 | P | 504.3 | P | 672.9 | P | --- |
| 252588_at | 270 | P | 271.5 | P | 266.4 | P | 328.8 | P | --- |
| 252589_s_at | 20.1 | A | 71.2 | P | 55.8 | A | 44.3 | A | NAXT1 |
| 252590_at | 4.2 | A | 27.8 | A | 24.5 | A | 26.7 | A | --- |
| 252591_at | 592.1 | P | 772.6 | P | 551.5 | P | 720.3 | P | TET3 |
| 252592_at | 1396.3 | P | 1915.7 | P | 1349.7 | P | 1690.1 | P | NAXT1 |
| 252593_at | 160.7 | P | 152.3 | P | 169.2 | P | 160.2 | P | ATSEN1 |
| 252594_at | 65.1 | P | 94.6 | P | 182.9 | P | 75.4 | P | --- |
| 252595_at | 15.8 | A | 21.8 | A | 39.1 | A | 32 | A | --- |
| 252596_at | 2.4 | A | 15.4 | A | 31.9 | A | 16.6 | A | --- |
| 252597_at | 3.5 | A | 2.1 | A | 0.1 | A | 0.1 | A | --- |
| 252598_at | 0.8 | A | 0.1 | A | 0.4 | A | 0.1 | A | --- |
| 252599_at | 4.8 | A | 35.8 | A | 38.3 | A | 33.7 | A | --- |
| 252600_at | 0.9 | A | 2.5 | A | 1.2 | A | 0.9 | A | VPS24.2 |
| 252601_s_at | 5932.4 | P | 5021.5 | P | 6465.4 | P | 5914.8 | P | --- |
| 252602_at | 508.3 | P | 474.5 | P | 362.9 | P | 398.8 | P | --- |
| 252603_at | 693.3 | P | 1103.4 | P | 692.2 | P | 1049.5 | P | --- |
| 252604_at | 9.8 | A | 30.4 | A | 12.9 | A | 47.3 | A | NRT2.6 |
| 252605_s_at | 4.2 | A | 43.5 | P | 21 | A | 24.9 | P | --- |
| 252606_at | 1189 | P | 893.1 | P | 1187.2 | P | 977.2 | P | scpl48 |
| 252607_at | 322.5 | P | 2342.8 | P | 570 | P | 2454.2 | P | XTR8 |
| 252608_at | 227.1 | P | 235 | P | 259.2 | P | 321.1 | P | --- |
| 252609_at | 49.6 | M | 20.9 | P | 32.1 | A | 26.2 | A | --- |
| 252610_x_at | 22.8 | A | 1 | A | 15.3 | A | 22 | A | --- |
| 252611_at | 424.7 | P | 56 | P | 169.5 | P | 79.5 | P | LAS1 |
| 252612_at | 44.3 | P | 576.9 | P | 375.4 | P | 664.1 | P | --- |
| 252613_at | 284 | P | 315.3 | P | 324.5 | P | 280.2 | P | --- |
| 252614_at | 31.9 | A | 1 | A | 17.3 | A | 2.8 | A | --- |
| 252615_at | 235.8 | P | 424.6 | P | 364.5 | P | 439.2 | P | --- |
| 252616_at | 444.8 | P | 352.2 | P | 506.1 | P | 523.8 | P | GRIK1 |
| 252617_at | 337.4 | P | 344.2 | P | 399.9 | P | 316.9 | P | SETH2 |
| 252618_at | 16727.7 | P | 9821.6 | P | 13436.1 | P | 13110.4 | P | LOX2 |
| 252619_at | 197.3 | P | 357.7 | P | 277.4 | P | 340.8 | P | --- |
| 252620_at | 1.2 | A | 1.3 | A | 2.8 | A | 3 | A | --- |
| 252621_at | 22 | A | 3.2 | A | 1.8 | A | 15.5 | A | --- |
| 252622_at | 1439.4 | P | 1350 | P | 1354.4 | P | 1349.9 | P | --- |
| 252623_at | 16.4 | A | 5 | A | 12.9 | A | 3.1 | A | --- |
| 252624_at | 273.9 | P | 400.2 | P | 284.2 | P | 394.4 | P | PSK1 |
| 252625_at | 1327.9 | P | 950.2 | P | 1804.1 | P | 1081.9 | P | HDA3 |
| 252626_at | 69.8 | P | 128.2 | P | 107 | P | 103.2 | P | --- |
| 252627_at | 57.8 | A | 29.1 | A | 79.3 | A | 46.7 | A | --- |
| 252628_at | 45.4 | P | 35.2 | P | 41.2 | A | 24 | P | --- |
| 252629_at | 1514.8 | P | 1090.5 | P | 1531.6 | P | 1088.5 | P | --- |
| 252630_at | 9.2 | A | 3.4 | A | 4.9 | A | 3.2 | A | --- |
| 252631_s_at | 15.6 | A | 0.7 | A | 5.5 | A | 1 | A | --- |
| 252632_s_at | 102.7 | M | 110.9 | M | 160.9 | P | 128.4 | P | RPP1 |
| 252633_x_at | 367.5 | P | 341.2 | P | 332.7 | P | 427.7 | P | hda17 |
| 252634_x_at | 12.1 | A | 14.8 | A | 19.2 | A | 31.8 | A | --- |
| 252635_at | 25.3 | A | 11.2 | A | 24.1 | A | 23 | P | --- |

| | | | | | | | | | |
|---|---|---|---|---|---|---|---|---|---|
| 252636_at | 1.3 | A | 22.2 | A | 1.5 | A | 1.3 | A | --- |
| 252637_at | 167.3 | P | 209 | P | 228.4 | P | 220.9 | P | HIRA |
| 252638_at | 15.3 | A | 28.9 | P | 56.2 | P | 14.3 | A | FAR4 |
| 252639_at | 229.3 | P | 84.2 | P | 100.8 | P | 73.4 | P | FAR5 |
| 252640_at | 9.3 | A | 3 | A | 5.2 | A | 1.4 | A | FAR8 |
| 252641_at | 1.1 | A | 8.1 | A | 10.5 | A | 4.7 | A | --- |
| 252642_at | 4.8 | A | 2.3 | A | 40.3 | A | 20.3 | A | --- |
| 252643_at | 3308.8 | P | 2986.5 | P | 3824.6 | P | 3577.2 | P | --- |
| 252644_at | 565.7 | P | 486.9 | P | 405.4 | P | 473 | P | CYP71 |
| 252645_at | 16.8 | A | 26.4 | A | 6.5 | A | 8.3 | A | DPBF2 |
| 252646_at | 153.2 | P | 139.4 | P | 153.6 | P | 185.7 | P | --- |
| 252647_at | 1716.4 | P | 1836.3 | P | 1506.5 | P | 1834.3 | P | --- |
| 252648_at | 644.9 | P | 787.8 | P | 537.9 | P | 660.3 | P | --- |
| 252649_at | 151.4 | P | 167.8 | P | 170.1 | P | 208.7 | P | HDA9 |
| 252650_at | 32.6 | A | 50 | A | 47.3 | A | 51.1 | P | --- |
| 252651_at | 22.2 | A | 26.7 | P | 30.8 | P | 29.9 | A | --- |
| 252652_at | 1537.4 | P | 807.9 | P | 1341.1 | P | 906.2 | P | ADT4 |
| 252653_at | 44.2 | P | 58.9 | P | 97.8 | P | 56.5 | P | --- |
| 252654_at | 116.6 | P | 116.4 | P | 137.1 | P | 126.9 | P | --- |
| 252655_at | 14.6 | A | 16.3 | A | 20.4 | A | 15.4 | A | --- |
| 252656_at | 32.5 | A | 41 | A | 33.2 | A | 33 | A | --- |
| 252657_at | 14.2 | A | 20.3 | A | 18.2 | A | 13 | A | --- |
| 252658_at | 1.4 | A | 3.8 | A | 1.2 | A | 13.2 | A | --- |
| 252659_at | 233.5 | P | 244.8 | P | 293 | P | 254.9 | P | --- |
| 252660_at | 29.6 | M | 15.8 | P | 37.8 | A | 18.5 | A | --- |
| 252661_at | 761.1 | P | 1092.8 | P | 730.7 | P | 1022.9 | P | --- |
| 252662_at | 1032.8 | P | 1125.8 | P | 1013 | P | 1045.4 | P | CEF |
| 252663_at | 61.5 | P | 71.2 | P | 101.7 | P | 55 | P | --- |
| 252664_at | 1.1 | A | 13.3 | A | 11.2 | A | 11.9 | A | --- |
| 252665_at | 99.9 | P | 115.3 | P | 149.5 | P | 151.9 | P | --- |
| 252666_at | 46.2 | A | 47 | A | 56.1 | A | 51.1 | A | --- |
| 252667_at | 155.8 | P | 228.5 | P | 228.3 | P | 288.8 | P | NEK6 |
| 252668_at | 1.4 | A | 9.6 | A | 17.8 | A | 18.9 | A | --- |
| 252669_at | 1813.3 | P | 1773.2 | P | 1468.9 | P | 1938.6 | P | --- |
| 252670_at | 4658.2 | P | 4600.7 | P | 5429.7 | P | 5029.5 | P | ATJ3 |
| 252671_at | 660.2 | P | 832.2 | P | 536.7 | P | 800.9 | P | --- |
| 252672_at | 25.1 | A | 23.3 | A | 47.7 | P | 24.7 | A | --- |
| 252673_at | 23.1 | P | 25.3 | P | 38.4 | P | 18.9 | P | --- |
| 252674_at | 23.4 | A | 19 | A | 35.7 | A | 4 | A | CYP71B38 |
| 252675_at | 2.4 | A | 1.3 | A | 1.5 | A | 1.5 | A | --- |
| 252676_at | 583.8 | P | 649.7 | P | 380.6 | P | 523 | P | --- |
| 252677_at | 625.6 | P | 246.4 | P | 657.6 | P | 283.7 | P | NIT3 |
| 252678_s_at | 18853.2 | P | 16630.2 | P | 18394.4 | P | 17888.1 | P | NIT2 |
| 252679_at | 233.4 | P | 196.6 | P | 272 | P | 207.3 | P | --- |
| 252680_at | 760.6 | P | 809.4 | P | 760.3 | P | 747.2 | P | --- |
| 252681_at | 1.1 | A | 33.4 | P | 4.7 | A | 33.7 | A | anac061 |
| 252682_at | 175 | P | 178.1 | P | 210.5 | P | 176 | P | --- |
| 252683_at | 186.3 | P | 230.1 | P | 224.3 | P | 256.3 | P | --- |
| 252684_at | 61.7 | P | 85.6 | P | 60.8 | P | 51.4 | P | --- |
| 252685_at | 33.2 | A | 55.5 | P | 60.8 | P | 30.3 | A | --- |
| 252686_at | 42.1 | A | 21.1 | A | 41.7 | A | 13.6 | A | --- |
| 252687_at | 11.2 | A | 12.8 | A | 53.2 | A | 43.3 | A | --- |
| 252688_at | 326.4 | P | 340.7 | P | 337.5 | P | 251.6 | P | --- |
| 252689_at | 7.3 | A | 14.4 | A | 17.1 | A | 30.3 | A | --- |
| 252690_at | 1.7 | A | 4.6 | A | 6.8 | A | 6.6 | A | --- |
| 252691_at | 57.1 | P | 87 | P | 66 | P | 87 | P | --- |
| 252692_at | 107.1 | P | 185 | P | 74.2 | P | 144.9 | P | --- |
| 252693_s_at | 9359.3 | P | 9211.7 | P | 9270.3 | P | 10211.6 | P | --- |
| 252694_at | 5.3 | A | 13.1 | A | 62.5 | A | 32.2 | A | --- |
| 252695_s_at | 1.4 | A | 7.9 | A | 1.7 | A | 1.6 | A | --- |
| 252696_at | 0.5 | A | 0.9 | A | 1.7 | A | 2.5 | A | --- |
| 252697_at | 53.8 | P | 21.2 | P | 55.3 | P | 32 | A | --- |
| 252698_at | 475.6 | P | 237.4 | P | 215 | P | 299.1 | P | --- |
| 252699_at | 6.2 | A | 3.8 | A | 7.6 | A | 5.1 | A | --- |
| 252700_at | 89 | P | 60.7 | P | 105.1 | P | 92.6 | P | --- |
| 252701_at | 125.7 | P | 161.9 | P | 193.4 | P | 152.7 | P | ATBPM6 |
| 252702_at | 8.6 | A | 0.8 | A | 7.7 | A | 10.5 | A | --- |
| 252703_at | 696.8 | P | 676.3 | P | 435.4 | P | 549.1 | P | --- |
| 252704_at | 10.4 | A | 2.6 | A | 2.9 | A | 2 | A | --- |
| 252705_at | 2.9 | A | 1.3 | A | 5.8 | A | 2.2 | A | --- |
| 252706_at | 1.7 | A | 10.2 | A | 21.9 | A | 5 | A | --- |
| 252707_at | 153.2 | P | 171.2 | P | 159.8 | P | 174.3 | P | ZIFL2 |
| 252708_at | 2.1 | A | 2.2 | A | 1.7 | A | 2.3 | A | --- |
| 252709_at | 13.3 | A | 5.8 | A | 12.4 | A | 12.8 | A | --- |
| 252710_at | 6 | A | 7.5 | A | 28 | A | 8.7 | A | AtGH9A4 |
| 252711_at | 1705.5 | P | 2410.7 | P | 2322.3 | P | 2346 | P | --- |
| 252712_at | 498.5 | P | 825.4 | P | 526.3 | P | 715.3 | P | ATGSTU27 |
| 252713_at | 2310.7 | P | 2462.9 | P | 1757.2 | P | 2520.3 | P | CAM7 |
| 252714_x_at | 7.2 | A | 7.2 | A | 22.1 | A | 10 | A | --- |
| 252715_x_at | 44.9 | A | 37.4 | A | 64.2 | A | 32.9 | M | --- |
| 252716_at | 76.7 | P | 110.1 | P | 63.5 | P | 120.7 | P | DCL3 |
| 252717_at | 22.5 | A | 15.4 | A | 12.4 | A | 16.4 | A | --- |
| 252718_at | 3.3 | A | 16.2 | A | 17.7 | A | 4.1 | A | --- |
| 252719_at | 3.7 | A | 8.5 | A | 2.7 | A | 1.3 | A | --- |
| 252720_at | 1.3 | A | 3.6 | A | 1.3 | A | 1 | A | --- |
| 252721_at | 2.5 | A | 1.3 | A | 2.1 | A | 3.4 | A | --- |
| 252722_at | 3 | A | 12.5 | A | 24 | A | 3.4 | A | --- |
| 252723_at | 2462.6 | P | 2506.9 | P | 2733.8 | P | 2432.3 | P | --- |
| 252724_at | 759.1 | P | 783.9 | P | 795.4 | P | 679 | P | --- |
| 252725_at | 2.7 | A | 17.7 | A | 17.3 | A | 3 | A | --- |
| 252726_at | 2.5 | A | 2.9 | A | 2.5 | A | 6.5 | A | --- |
| 252727_s_at | 0.3 | A | 12.7 | A | 1.2 | A | 1.4 | A | --- |
| 252728_at | 16 | A | 3.1 | A | 3.9 | A | 3.2 | A | --- |
| 252729_at | 12.8 | A | 11.5 | A | 0.5 | A | 7 | A | --- |
| 252730_at | 36.9 | P | 24.4 | A | 45.6 | A | 32.1 | A | --- |
| 252731_at | 29.1 | A | 21.4 | A | 2.3 | A | 10.9 | A | --- |
| 252732_at | 1.2 | A | 1.4 | A | 4.9 | A | 1.9 | A | --- |
| 252733_at | 14.7 | A | 5.8 | A | 10.1 | A | 3.2 | A | --- |
| 252734_at | 3.9 | A | 17.8 | A | 12.6 | A | 20 | A | MEE38 |
| 252735_at | 21.7 | A | 7.5 | A | 26.8 | A | 2.7 | A | --- |
| 252736_at | 73.6 | P | 115.9 | P | 58.8 | A | 97.5 | P | TES |
| 252737_at | 369.5 | P | 413.5 | P | 333.5 | P | 377.6 | P | --- |
| 252738_at | 314.3 | P | 300.6 | P | 256 | P | 314.5 | P | --- |
| 252739_at | 43.6 | P | 2.3 | A | 6.4 | A | 11.7 | A | --- |
| 252740_at | 2469 | P | 915.6 | P | 677.3 | P | 1310.7 | P | --- |
| 252741_at | 31.8 | A | 11.7 | A | 21.6 | A | 21.7 | A | --- |
| 252742_at | 36.8 | A | 61.9 | A | 78.7 | A | 76.5 | A | --- |
| 252743_at | 731.5 | P | 617.5 | P | 480.2 | P | 555.7 | P | ATMIN7 |
| 252744_at | 0.9 | A | 3.1 | A | 1.1 | A | 0.6 | A | --- |

| | | | | | | | | | |
|---|---|---|---|---|---|---|---|---|---|
| 252745_at | 555.6 | P | 495.2 | P | 445.6 | P | 482.3 | P | --- |
| 252746_at | 28.1 | A | 50.4 | P | 51 | A | 40.5 | P | SUS4 |
| 252747_at | 20.3 | A | 38.8 | A | 41.9 | A | 38.9 | A | --- |
| 252748_at | 10.1 | A | 0.7 | A | 2.1 | A | 0.7 | A | --- |
| 252749_at | 1.7 | A | 1.8 | A | 3.7 | A | 1.5 | A | --- |
| 252750_at | 7.1 | A | 1.8 | A | 11.2 | A | 0.9 | A | --- |
| 252751_at | 53.8 | A | 95.3 | A | 53.3 | A | 72.6 | A | --- |
| 252752_s_at | 0.9 | A | 0.8 | A | 2.5 | A | 1.8 | A | --- |
| 252753_at | 1.7 | A | 1.7 | A | 5.2 | A | 4.4 | A | --- |
| 252754_at | 347.9 | P | 278.1 | P | 293.2 | P | 353.9 | P | --- |
| 252755_at | 0.3 | A | 5.4 | A | 0.6 | A | 0.4 | A | --- |
| 252756_s_at | 0.8 | A | 0.9 | A | 27.1 | A | 5.9 | A | --- |
| 252757_at | 7.7 | A | 19 | A | 30.2 | A | 12.7 | A | --- |
| 252758_at | 2.3 | A | 6.6 | A | 1.9 | A | 5.3 | A | --- |
| 252759_at | 298.4 | P | 378.1 | P | 306.6 | P | 296.7 | P | --- |
| 252760_x_at | 32.2 | A | 2.8 | A | 20 | A | 8 | A | --- |
| 252761_at | 4 | A | 2.7 | A | 4.7 | A | 2.7 | A | --- |
| 252762_at | 5.1 | A | 2.5 | A | 44.1 | A | 5.9 | A | --- |
| 252763_at | 38.3 | P | 61.9 | P | 53.6 | P | 57.4 | P | --- |
| 252764_at | 1141.8 | P | 1345.2 | P | 1158.3 | P | 1175.5 | P | AL3 |
| 252765_at | 23.8 | A | 47.5 | P | 18.4 | A | 36.9 | P | --- |
| 252766_at | 28.3 | P | 24.6 | P | 75.3 | P | 47.3 | P | --- |
| 252767_at | 40 | A | 40.7 | A | 21.1 | A | 26.1 | A | --- |
| 252768_at | 1 | A | 1.4 | A | 2.6 | A | 1.6 | A | --- |
| 252769_at | 16.7 | A | 22.2 | A | 42.8 | A | 12.5 | A | --- |
| 252770_at | 67.7 | P | 74.4 | P | 77.7 | P | 62.5 | P | --- |
| 252771_at | 1.3 | A | 1.3 | A | 3.3 | A | 6.4 | A | --- |
| 252772_at | 2.6 | A | 32.9 | A | 12.9 | A | 2.5 | A | --- |
| 252773_at | 22.8 | A | 13.8 | A | 31.2 | A | 8.6 | A | --- |
| 252774_at | 22.3 | A | 34.3 | A | 54.8 | A | 7.9 | A | --- |
| 252775_s_at | 1.4 | A | 0.1 | A | 2 | A | 0.2 | A | --- |
| 252776_at | 3.8 | A | 38.6 | A | 12.9 | A | 5.1 | A | --- |
| 252777_at | 1.2 | A | 0.8 | A | 2.9 | A | 0.7 | A | --- |
| 252778_at | 4.4 | A | 1.6 | A | 17.6 | A | 3.8 | A | --- |
| 252779_at | 0.8 | A | 2 | A | 3.3 | A | 1.1 | A | --- |
| 252780_at | 30.3 | A | 33.5 | A | 44.4 | A | 15.2 | A | ATA1 |
| 252781_at | 143.3 | P | 186.4 | P | 120.4 | P | 164.4 | P | --- |
| 252782_at | 5 | A | 1.7 | A | 1.5 | A | 0.6 | A | --- |
| 252783_at | 8.3 | A | 14.7 | A | 14.8 | A | 6.8 | A | --- |
| 252784_at | 12.6 | A | 0.7 | A | 14.3 | A | 1.7 | A | --- |
| 252785_at | 166.2 | P | 202.4 | P | 179.3 | P | 191.2 | P | --- |
| 252786_at | 97.6 | P | 122.6 | P | 140.7 | P | 102.1 | P | CHR38 |
| 252787_at | 2.9 | A | 2.3 | A | 2.8 | A | 10.6 | A | --- |
| 252788_at | 25 | A | 23.1 | A | 30 | A | 5.1 | A | --- |
| 252789_s_at | 1238 | P | 1135 | P | 1196.2 | P | 1109 | P | --- |
| 252790_at | 7.2 | A | 32.2 | A | 37.9 | M | 2.4 | A | --- |
| 252791_at | 14.7 | A | 15.3 | A | 20.9 | A | 15.2 | A | --- |
| 252792_at | 7.3 | A | 6.3 | A | 34.4 | A | 14.2 | A | --- |
| 252793_at | 0.3 | A | 8 | A | 6.9 | A | 1 | A | --- |
| 252794_at | 25.3 | A | 30.9 | P | 52 | A | 26.3 | A | --- |
| 252795_at | 32.9 | P | 9.2 | A | 23.5 | P | 27.6 | P | --- |
| 252796_at | 1.6 | A | 7.3 | A | 3.6 | A | 1.6 | A | --- |
| 252797_at | 20.5 | A | 27.6 | A | 10.2 | A | 7.8 | A | --- |
| 252798_at | 1.5 | A | 13.6 | A | 3.1 | A | 7.6 | A | --- |
| 252799_at | 2 | A | 1.4 | A | 2.1 | A | 1.7 | A | --- |
| 252800_at | 3.3 | A | 5.9 | A | 17.2 | A | 6 | A | --- |
| 252801_at | 0.8 | A | 1.4 | A | 2.1 | A | 1.5 | A | --- |
| 252802_at | 0.7 | A | 24.5 | A | 1.4 | A | 1.3 | A | --- |
| 252803_at | 26.5 | A | 22.7 | A | 29.8 | A | 21.4 | A | --- |
| 252804_at | 6.9 | A | 7.2 | A | 2.1 | A | 22.3 | A | --- |
| 252805_at | 49 | A | 27.4 | A | 24 | A | 10.6 | A | --- |
| 252806_at | 1.8 | A | 2.6 | A | 2.4 | A | 7.1 | A | --- |
| 252807_at | 3 | A | 4 | A | 0.8 | A | 2.9 | A | --- |
| 252808_at | 5.1 | A | 3 | A | 7.2 | A | 5.9 | A | --- |
| 252809_at | 0.5 | A | 0.8 | A | 0.6 | A | 0.4 | A | --- |
| 252810_at | 1.5 | A | 4.1 | A | 1.5 | A | 12 | A | --- |
| 252811_at | 2.7 | A | 18.7 | A | 45.5 | A | 4.4 | A | --- |
| 252812_at | 3 | A | 2.4 | A | 17.6 | A | 9.9 | A | --- |
| 252813_at | 20.5 | A | 0.3 | A | 0.5 | A | 16.5 | A | --- |
| 252814_at | 1.8 | A | 3.8 | A | 38.3 | A | 20.5 | A | --- |
| 252815_at | 4.7 | A | 5.8 | A | 6.3 | A | 3.5 | A | --- |
| 252816_at | 20.7 | A | 19.1 | A | 23.9 | A | 2.5 | A | --- |
| 252817_at | 15.4 | A | 4.2 | A | 33.8 | A | 24.8 | A | --- |
| 252818_at | 16.6 | A | 5.7 | A | 2.6 | A | 20.8 | A | --- |
| 252819_at | 15.5 | A | 24.6 | A | 14.9 | A | 25.4 | M | --- |
| 252820_at | 113.1 | P | 110.3 | P | 106.2 | P | 118.2 | P | AHA8 |
| 252821_at | 210.9 | P | 297.9 | P | 290.7 | P | 296.3 | P | --- |
| 252822_at | 727.7 | P | 486.4 | P | 689.5 | P | 469 | P | --- |
| 252823_at | 631.1 | P | 665.1 | P | 565.2 | P | 648.9 | P | --- |
| 252824_at | 5121.8 | P | 5775.8 | P | 6616 | P | 6325.7 | P | --- |
| 252825_at | 114.2 | P | 101.2 | P | 108 | A | 63.7 | A | AtRABH1c |
| 252826_at | 343.5 | P | 415.8 | P | 321.8 | P | 405.2 | P | --- |
| 252827_at | 6185.9 | P | 1989.7 | P | 5056.7 | P | 2259.9 | P | CYP79B2 |
| 252828_at | 929.7 | P | 1006.3 | P | 1076.4 | P | 1126.3 | P | --- |
| 252829_at | 553.8 | P | 637.7 | P | 544.5 | P | 580.3 | P | ATHB16 |
| 252830_at | 784.9 | P | 742.7 | P | 632.3 | P | 753.6 | P | PXA1 |
| 252831_at | 4194.9 | P | 2830.6 | P | 3415.3 | P | 3083.3 | P | DHS1 |
| 252832_at | 867.8 | P | 957.5 | P | 855.1 | P | 830.8 | P | ATUBP3 |
| 252833_at | 19.7 | A | 19.9 | A | 27 | A | 15.5 | A | AGP3 |
| 252834_at | 35 | P | 32.9 | M | 53 | A | 55.8 | P | --- |
| 252835_at | 5.7 | A | 3.4 | A | 3.6 | A | 3.8 | A | --- |
| 252836_at | 38.7 | A | 28.5 | A | 43.3 | A | 41 | A | --- |
| 252837_at | 0.1 | A | 8.2 | A | 9.7 | A | 2.7 | A | --- |
| 252838_at | 0.8 | A | 0.3 | A | 0.5 | A | 2.2 | A | --- |
| 252839_at | 10.2 | A | 2.4 | A | 5.2 | A | 18.1 | A | --- |
| 252840_at | 30.4 | A | 4.5 | A | 17.3 | A | 25.8 | A | --- |
| 252841_at | 3.2 | A | 6.7 | A | 3.5 | A | 1.1 | A | --- |
| 252842_at | 0.9 | A | 0.6 | A | 1 | A | 0.8 | A | --- |
| 252843_at | 2.9 | A | 5 | A | 17.4 | A | 15.4 | A | --- |
| 252844_at | 47.7 | M | 54.6 | P | 60.9 | P | 44 | P | --- |
| 252845_at | 3659.9 | P | 3699.1 | P | 4412.7 | P | 3734.9 | P | --- |
| 252846_at | 7.4 | A | 1.8 | A | 17.2 | A | 1.5 | A | --- |
| 252847_at | 376.6 | P | 468.9 | P | 368.1 | P | 498.1 | P | --- |
| 252848_at | 31 | A | 11.1 | A | 16.9 | A | 19.3 | A | --- |
| 252849_at | 4583.8 | P | 6030.5 | P | 5671.2 | P | 6070.4 | P | --- |
| 252850_at | 1352.2 | P | 1105.4 | P | 1183.4 | P | 1185.9 | P | --- |
| 252851_at | 136.8 | P | 77.1 | P | 106.6 | P | 109.5 | P | --- |
| 252852_at | 483.9 | P | 750.7 | P | 715.3 | P | 749.4 | P | --- |
| 252853_at | 308.1 | P | 649 | P | 342.3 | P | 583.3 | P | --- |

| | | | | | | | | | |
|---|---|---|---|---|---|---|---|---|---|
| 252854_at | 147.8 | P | 165.6 | P | 153.5 | P | 161 | P | UBP27 |
| 252855_at | 936.8 | P | 921.3 | P | 775 | P | 777.3 | P | AGT2 |
| 252856_at | 658.4 | P | 733.5 | P | 678.8 | P | 694.5 | P | --- |
| 252857_at | 2.3 | A | 15.4 | A | 20.3 | A | 12.9 | A | --- |
| 252858_at | 16.7 | A | 35.9 | A | 55.9 | A | 26.3 | A | --- |
| 252859_at | 98.1 | P | 143.7 | P | 153 | P | 117.4 | P | --- |
| 252860_at | 40.7 | P | 56.5 | P | 53.8 | P | 54.9 | P | --- |
| 252861_at | 319.6 | P | 506.3 | P | 355.2 | P | 481 | P | --- |
| 252862_at | 97.6 | P | 266.4 | P | 74.9 | P | 139.8 | P | --- |
| 252863_at | 2123.7 | P | 2864 | P | 2464.1 | P | 3884.6 | P | MIPS1 |
| 252864_at | 141.8 | P | 192.7 | P | 176.5 | P | 195.6 | P | --- |
| 252865_at | 9.4 | A | 11.1 | A | 40.5 | A | 3.4 | A | --- |
| 252866_at | 856.4 | P | 950.2 | P | 752.4 | P | 802.4 | P | --- |
| 252867_at | 238.1 | P | 292.3 | P | 256.3 | P | 271.9 | P | --- |
| 252868_at | 166.9 | P | 184.9 | P | 223.3 | P | 206.7 | P | POR |
| 252869_at | 44.4 | M | 43.7 | P | 49.2 | P | 38.3 | P | --- |
| 252870_at | 3920.4 | P | 2223.3 | P | 4077.3 | P | 2533.5 | P | AKN2 |
| 252871_at | 28.5 | A | 91.1 | P | 90.3 | P | 109.8 | P | --- |
| 252872_at | 106.2 | P | 93 | P | 121.5 | P | 67.7 | A | SNRK2.7 |
| 252873_at | 13.8 | A | 10 | A | 14.2 | A | 7.8 | A | --- |
| 252874_at | 29 | A | 37.9 | A | 29.4 | A | 55.2 | M | --- |
| 252875_at | 682.4 | P | 833.3 | P | 560.2 | P | 875.7 | P | RABA4B |
| 252876_at | 899.9 | P | 973.6 | P | 894.5 | P | 1128.9 | P | --- |
| 252877_at | 59 | P | 90.4 | P | 92.8 | P | 89.1 | P | --- |
| 252878_at | 1541.6 | P | 1504.2 | P | 1455.1 | P | 1481.1 | P | SAMC1 |
| 252879_at | 389.3 | P | 380.4 | P | 370.1 | P | 308.3 | P | NST-K1 |
| 252880_at | 5842 | P | 5507.9 | P | 5240.9 | P | 6573.3 | P | --- |
| 252881_at | 2.8 | A | 27.2 | A | 38 | A | 32.8 | A | --- |
| 252882_at | 38.7 | P | 4.4 | A | 39.5 | A | 15.4 | A | --- |
| 252883_at | 1780.5 | P | 1595.8 | P | 1891.4 | P | 1512.8 | P | --- |
| 252884_at | 666 | P | 812.4 | P | 663.5 | P | 800.3 | P | --- |
| 252885_at | 9665.6 | P | 8987.8 | P | 9083.7 | P | 9468.7 | P | GR-RBP8 |
| 252886_at | 1472 | P | 1620.1 | P | 1412.8 | P | 1455 | P | CESA2 |
| 252887_at | 478.2 | P | 472.1 | P | 488.6 | P | 416.5 | P | --- |
| 252888_at | 1080 | P | 1304.9 | P | 1010 | P | 1562.2 | P | APL3 |
| 252889_at | 58.9 | P | 51.5 | P | 51.3 | A | 56.9 | P | --- |
| 252890_at | 1400.2 | P | 1250.6 | P | 1314.7 | P | 1356.6 | P | BRI1 |
| 252891_at | 50.3 | P | 47.8 | A | 70.7 | P | 32.6 | A | WRKY13 |
| 252892_at | 17.1 | A | 25.7 | A | 43.7 | A | 13.5 | A | --- |
| 252893_at | 8.4 | A | 3.3 | A | 7.9 | A | 4.6 | A | --- |
| 252894_at | 56.9 | A | 43.1 | A | 45.3 | A | 24.4 | A | --- |
| 252895_at | 440.6 | P | 371.1 | P | 341.8 | P | 372.5 | P | --- |
| 252896_at | 26.1 | P | 50.2 | P | 44.8 | P | 28.9 | P | CYP96A9 |
| 252897_at | 10.1 | A | 3 | A | 4.2 | A | 4.5 | A | CYP96A9 |
| 252898_at | 32.1 | A | 3.1 | A | 28.5 | M | 12.4 | A | CYP96A11 |
| 252899_at | 106.6 | P | 93.6 | P | 123 | P | 142.8 | P | --- |
| 252900_at | 1230.2 | P | 819.1 | P | 1286.8 | P | 881.1 | P | --- |
| 252901_at | 126.2 | P | 99.6 | P | 135.9 | P | 118.9 | P | --- |
| 252902_at | 119.4 | P | 122.6 | A | 127.8 | P | 85 | P | --- |
| 252903_at | 200.5 | P | 250.9 | P | 277.4 | P | 271.7 | P | --- |
| 252904_at | 433 | P | 532.4 | P | 478.6 | P | 523.2 | P | EMB2453 |
| 252905_at | 55.4 | P | 73.9 | P | 100.1 | P | 67.2 | P | --- |
| 252906_at | 319.1 | P | 459.2 | P | 163.2 | P | 242.2 | P | GGT1 |
| 252907_at | 3 | A | 20.8 | A | 23.4 | A | 1.8 | A | GGT2 |
| 252908_at | 184.9 | P | 150 | P | 97.3 | P | 156.6 | P | --- |
| 252909_at | 770.5 | P | 985.6 | P | 967.9 | P | 964.7 | P | --- |
| 252910_at | 3.5 | A | 3.6 | A | 5.1 | A | 3.8 | A | --- |
| 252911_at | 210.6 | P | 521.4 | P | 301.2 | P | 443.8 | P | CYP96A12 |
| 252912_at | 6552.9 | P | 6073.9 | P | 5506.1 | P | 7674.8 | P | --- |
| 252913_at | 868.5 | P | 1004.8 | P | 822.2 | P | 865.4 | P | ATRER1A |
| 252914_at | 10.4 | A | 9.9 | A | 25 | A | 13.9 | A | --- |
| 252915_at | 1148.4 | P | 906.3 | P | 963.6 | P | 944.2 | P | --- |
| 252916_at | 59.8 | P | 194.8 | P | 128.6 | P | 242 | P | --- |
| 252917_at | 77.1 | A | 151.7 | P | 114.3 | A | 107.1 | P | --- |
| 252918_at | 192.9 | P | 197.4 | P | 208.8 | P | 222 | P | --- |
| 252919_at | 4.8 | A | 12.8 | A | 28.4 | A | 3 | A | AtGH9B16 |
| 252920_at | 20.4 | A | 24.9 | A | 20.5 | A | 14 | A | AtGH9B17 |
| 252921_at | 385.5 | P | 389.8 | P | 315.3 | P | 280.8 | P | EDS5 |
| 252922_at | 1967.6 | P | 2382.8 | P | 1992.9 | P | 2211.4 | P | --- |
| 252923_at | 264.1 | P | 385.8 | P | 364.6 | P | 324.4 | P | --- |
| 252924_at | 100.8 | P | 66.6 | P | 118.9 | P | 75.8 | P | --- |
| 252925_at | 687.8 | P | 713.2 | P | 591.8 | P | 595.4 | P | SHL1 |
| 252926_at | 6031.3 | P | 6565.4 | P | 5873.9 | P | 6465.2 | P | ATVHA-C3 |
| 252927_at | 8045.7 | P | 7844.2 | P | 7644.8 | P | 7496.6 | P | RD19 |
| 252928_at | 110.2 | P | 96 | P | 124.8 | P | 108.3 | P | --- |
| 252929_at | 18438.5 | P | 18469 | P | 19826.3 | P | 19929.7 | P | --- |
| 252930_at | 41.4 | A | 60.9 | P | 70.6 | P | 81.4 | P | AtGH9B18 |
| 252931_at | 454.2 | P | 476.8 | P | 452.1 | P | 475 | P | --- |
| 252932_at | 2930.9 | P | 3012.3 | P | 3235.7 | P | 2548.4 | P | VHA-A3 |
| 252933_at | 4.9 | A | 1.7 | A | 47.1 | A | 9.8 | A | --- |
| 252934_at | 354 | P | 454.6 | P | 467 | P | 439 | P | IMPL2 |
| 252935_at | 774.6 | P | 828.9 | P | 729.7 | P | 758.2 | P | --- |
| 252936_at | 104.5 | P | 172.2 | P | 105.9 | P | 135.6 | P | --- |
| 252937_at | 81.8 | P | 160.9 | P | 84.1 | P | 134.2 | P | SEC14 |
| 252938_at | 70.7 | P | 65.4 | P | 51.5 | P | 54.9 | P | --- |
| 252939_at | 127.8 | P | 174.2 | P | 193 | P | 172.6 | P | --- |
| 252940_at | 250.5 | P | 199.8 | P | 166.1 | P | 158.3 | P | --- |
| 252941_at | 1251.1 | P | 1389.7 | P | 1324.3 | P | 1379.7 | P | --- |
| 252942_at | 638.9 | P | 510.8 | P | 652.4 | P | 547 | P | --- |
| 252943_at | 2980.8 | P | 4733.1 | P | 3093.3 | P | 4155.2 | P | CAD9 |
| 252944_at | 369.8 | P | 101.5 | P | 123.3 | P | 90.6 | P | --- |
| 252945_at | 382.6 | P | 367.5 | P | 295.4 | P | 329.5 | P | --- |
| 252946_at | 120.2 | P | 155.2 | P | 180 | P | 152.9 | P | --- |
| 252947_at | 969.3 | P | 716 | P | 887.8 | P | 812.3 | P | KAK |
| 252948_at | 62.3 | P | 72.6 | P | 80.2 | A | 73.4 | P | KAK |
| 252949_at | 178.8 | P | 164.9 | P | 225.6 | P | 155.4 | P | --- |
| 252950_at | 227.5 | P | 234.8 | P | 172.7 | P | 259.1 | P | --- |
| 252951_at | 49.6 | P | 37.6 | P | 59.3 | P | 64.8 | P | --- |
| 252952_at | 1143.5 | P | 1004.3 | P | 1082.2 | P | 1022.5 | P | --- |
| 252953_at | 264.4 | P | 305 | P | 333.9 | P | 289 | P | PIS2 |
| 252954_at | 493.5 | P | 808 | P | 555.2 | P | 882.3 | P | --- |
| 252955_at | 2058.6 | P | 2030 | P | 2045.8 | P | 2221.1 | P | RPN10 |
| 252956_at | 1417.8 | P | 1517.7 | P | 1430.1 | P | 1684.8 | P | ATFP6 |
| 252957_at | 2361.9 | P | 1917.5 | P | 2802.4 | P | 2292.2 | P | GRP2 |
| 252958_at | 268.2 | P | 229.4 | P | 310.6 | P | 244.9 | P | MYB4 |
| 252959_at | 742.2 | P | 812.7 | P | 786.9 | P | 856.3 | P | --- |
| 252960_at | 415.6 | P | 406.9 | P | 429.2 | P | 402.1 | P | --- |
| 252961_at | 52.6 | A | 54.1 | A | 109.7 | A | 40.1 | A | --- |
| 252962_at | 56.9 | A | 58.3 | A | 71.4 | M | 72.3 | P | --- |

| | | | | | | | | | |
|---|---|---|---|---|---|---|---|---|---|
| 252963_at | 2.7 | A | 3.3 | A | 3.5 | A | 3.8 | A | --- |
| 252964_at | 23.6 | A | 23 | A | 30.8 | A | 25.3 | A | --- |
| 252965_at | 116.6 | P | 829.1 | P | 125.4 | P | 671.8 | P | --- |
| 252966_at | 38.9 | P | 53.6 | P | 40.2 | A | 37.5 | A | --- |
| 252967_at | 64.3 | P | 56.2 | P | 52.4 | A | 36.5 | A | ATASE3 |
| 252968_at | 961.7 | P | 972.9 | P | 1120.5 | P | 1064.7 | P | --- |
| 252969_at | 263.6 | P | 302.3 | P | 270.1 | P | 277.1 | P | --- |
| 252970_at | 17.7 | A | 112 | P | 7.8 | A | 59 | P | SAUR15 |
| 252971_at | 7957 | P | 9296.1 | P | 8590.4 | P | 9596 | P | PRP4 |
| 252972_at | 467.6 | P | 1129.6 | P | 866.9 | P | 1074.9 | P | --- |
| 252973_s_at | 4472.5 | P | 4853.4 | P | 5043.9 | P | 5713.6 | P | ROC1 |
| 252974_at | 6025.4 | P | 5837 | P | 6293.7 | P | 5420.6 | P | MTN1 |
| 252975_s_at | 571.1 | P | 925 | P | 735.4 | P | 947.3 | P | ROPGEF1 |
| 252976_s_at | 566.1 | P | 1086.9 | P | 561.8 | P | 853.1 | P | --- |
| 252977_at | 55.1 | P | 166.3 | P | 64 | P | 121.6 | P | --- |
| 252978_at | 62.3 | P | 66.4 | M | 64.1 | P | 65.5 | A | BGAL14 |
| 252979_at | 993.2 | P | 1293.5 | P | 1263.7 | P | 1085.6 | P | --- |
| 252980_at | 634 | P | 588.7 | P | 599.6 | P | 629.6 | P | --- |
| 252981_at | 199.3 | P | 224.4 | P | 202.3 | P | 178.3 | P | --- |
| 252982_at | 954.6 | P | 1102.9 | P | 1070 | P | 1085.9 | P | HD1 |
| 252983_at | 2567.8 | P | 1420.1 | P | 1624.6 | P | 2371.6 | P | ELI3-1 |
| 252984_at | 1970.8 | P | 175.2 | P | 424.6 | P | 374.8 | P | ELI3-2 |
| 252985_at | 471.4 | P | 504.1 | P | 480 | P | 542.9 | P | --- |
| 252986_at | 100.3 | P | 117.7 | P | 167.1 | P | 114.8 | P | --- |
| 252987_at | 10.2 | A | 6.5 | A | 8.5 | A | 4.9 | A | --- |
| 252988_at | 39.8 | A | 33.6 | A | 49.7 | A | 36.1 | P | --- |
| 252989_at | 307.7 | P | 267.7 | P | 279.4 | P | 256.3 | P | sks9 |
| 252990_at | 160.1 | P | 121.4 | P | 146.9 | P | 132.2 | P | --- |
| 252991_at | 257 | P | 185.6 | P | 319.5 | P | 194.2 | P | --- |
| 252992_at | 1268.8 | P | 1355.6 | P | 1314.7 | P | 1509.4 | P | --- |
| 252993_at | 2225.1 | P | 1127.3 | P | 1938.3 | P | 1191.1 | P | --- |
| 252994_at | 102.5 | A | 109.4 | A | 86 | A | 113.5 | A | --- |
| 252995_at | 258 | P | 283.4 | P | 348.8 | P | 293 | P | --- |
| 252996_s_at | 1431.8 | P | 1835.3 | P | 1532 | P | 1690.6 | P | GGR |
| 252997_at | 120.3 | P | 86.3 | P | 129.5 | P | 128.2 | P | ATEXLA2 |
| 252998_at | 2349.5 | P | 2419.9 | P | 2334.1 | P | 2546.6 | P | --- |
| 252999_at | 370.4 | P | 454.1 | P | 357.3 | P | 371.8 | P | --- |
| 253000_at | 595.9 | P | 651 | P | 703.3 | P | 676 | P | --- |
| 253001_at | 1131.5 | P | 1133 | P | 1220.7 | P | 1122.6 | P | --- |
| 253002_at | 91.5 | P | 87.4 | P | 145.6 | P | 85.8 | P | ATPLC1 |
| 253003_s_at | 48.1 | P | 44 | A | 50.8 | P | 52.7 | P | --- |
| 253004_at | 717.6 | P | 907.5 | P | 778 | P | 881.9 | P | --- |
| 253005_at | 1131.3 | P | 1066.4 | P | 1262.2 | P | 1179.9 | P | --- |
| 253006_at | 3049.7 | P | 2686.8 | P | 2900.6 | P | 2878.4 | P | --- |
| 253007_at | 579 | P | 811.8 | P | 730.5 | P | 808.8 | P | --- |
| 253008_at | 258.1 | P | 115.2 | P | 204 | P | 196.8 | P | ATEXPA20 |
| 253009_at | 11830.7 | P | 11719.7 | P | 14404.8 | P | 11862 | P | SHM1 |
| 253010_at | 119.8 | P | 214.2 | P | 207.8 | P | 240.7 | P | ANT |
| 253011_at | 73.1 | P | 79.5 | P | 88.4 | P | 108 | P | EDA40 |
| 253012_at | 29.8 | A | 22.8 | A | 1.7 | A | 6.7 | A | --- |
| 253013_at | 4544.2 | P | 2895.5 | P | 5089.1 | P | 3340.9 | P | mtHsc70-1 |
| 253014_at | 2.8 | A | 4.4 | A | 3.2 | A | 2 | A | AGL21 |
| 253015_at | 2.9 | A | 2.8 | A | 2.6 | A | 6.5 | A | --- |
| 253016_at | 10.9 | A | 20.4 | A | 5.1 | A | 8.4 | A | --- |
| 253017_at | 42.3 | A | 55.2 | M | 75.6 | P | 42 | A | CAD6 |
| 253018_at | 46.1 | A | 54.8 | A | 39.6 | A | 10.3 | A | --- |
| 253019_at | 138.2 | P | 112.6 | P | 122.1 | P | 109.6 | P | --- |
| 253020_at | 378.7 | P | 444.7 | P | 345.7 | P | 398.5 | P | --- |
| 253021_at | 167.1 | P | 220.6 | P | 169.8 | P | 222.8 | P | --- |
| 253022_at | 541.5 | P | 517.3 | P | 546.7 | P | 530 | P | --- |
| 253023_at | 21.1 | P | 47.2 | P | 38.9 | P | 9.2 | A | --- |
| 253024_at | 14.5 | A | 129.1 | P | 102.4 | P | 112.6 | P | --- |
| 253025_at | 213.4 | P | 224.8 | P | 175.4 | P | 196.4 | P | --- |
| 253026_at | 2 | A | 21 | A | 6.2 | A | 4.9 | A | --- |
| 253027_at | 439.9 | P | 348.5 | P | 586.5 | P | 305.6 | P | --- |
| 253028_at | 1100.6 | P | 1274 | P | 1193.8 | P | 1193 | P | pde191 |
| 253029_at | 161.6 | P | 126 | P | 144.6 | P | 104.4 | P | FRS9 |
| 253030_at | 87.7 | P | 114 | P | 133.6 | P | 102.2 | P | FRS5 |
| 253031_at | 85.1 | M | 99.2 | P | 128.4 | P | 86.4 | A | CSLD4 |
| 253032_at | 306.9 | P | 307.5 | P | 294.9 | P | 335.1 | P | --- |
| 253033_at | 1297.8 | P | 1472.9 | P | 1273.9 | P | 1227.1 | P | --- |
| 253034_at | 39.1 | P | 68.8 | P | 42.1 | P | 55.1 | P | CPK26 |
| 253035_at | 532.6 | P | 392.5 | P | 441.9 | P | 386.7 | P | CGL1 |
| 253036_at | 1.4 | A | 1.9 | A | 5.6 | A | 3 | A | --- |
| 253037_at | 323.5 | P | 324.3 | P | 322.8 | P | 249.7 | P | GAUT3 |
| 253038_at | 322 | P | 322.7 | P | 326.3 | P | 282.7 | P | HAT22 |
| 253039_at | 1312.1 | P | 1560.6 | P | 1971.1 | P | 1611.2 | P | SQE3 |
| 253040_at | 2364.2 | P | 4521.8 | P | 2747.8 | P | 4539.3 | P | --- |
| 253041_at | 11555.9 | P | 5191.7 | P | 12935.8 | P | 6086.5 | P | PCK1 |
| 253042_at | 601.8 | P | 837.8 | P | 1052.5 | P | 852.1 | P | --- |
| 253043_at | 101.3 | P | 143.5 | P | 152.5 | P | 147 | P | LBD39 |
| 253044_at | 79.8 | P | 34.3 | A | 61.8 | A | 52.8 | A | --- |
| 253045_at | 33.1 | P | 78.4 | P | 54.7 | P | 70.6 | P | --- |
| 253046_at | 730.1 | P | 306.1 | P | 295.9 | P | 326.4 | P | CYP81D8 |
| 253047_at | 57.4 | P | 35.7 | P | 29.7 | A | 48.2 | A | --- |
| 253048_at | 332.8 | P | 579.5 | P | 355.7 | P | 545.5 | P | --- |
| 253049_at | 1507.4 | P | 1528.9 | P | 1465.1 | P | 1475.3 | P | MEE59 |
| 253050_at | 450.2 | P | 935.7 | P | 528.3 | P | 869.2 | P | AGP18 |
| 253051_at | 82.4 | P | 122.4 | P | 134.9 | P | 141.4 | P | CYCB1;1 |
| 253052_at | 1065.8 | P | 754 | P | 810.7 | P | 816.4 | P | CYP81H1 |
| 253053_at | 1965.3 | P | 2097.1 | P | 1808.7 | P | 2214.6 | P | --- |
| 253054_at | 32.6 | A | 8.2 | A | 66.5 | P | 73.2 | P | HLS1 |
| 253055_at | 132.1 | P | 134.1 | P | 173.8 | P | 118.9 | P | CYCD5;1 |
| 253056_at | 105.1 | P | 127.1 | P | 127 | P | 142.2 | P | SHR |
| 253057_at | 290.6 | P | 224.9 | P | 154.1 | P | 246.3 | P | --- |
| 253058_at | 526.1 | P | 495.1 | P | 588.2 | P | 474.3 | P | --- |
| 253059_s_at | 972.3 | P | 934.7 | P | 922.3 | P | 981.4 | P | HHP5 |
| 253060_at | 25.2 | A | 22.8 | A | 25.1 | A | 0.9 | A | --- |
| 253061_at | 307 | P | 191.5 | P | 197.6 | P | 107.2 | P | BT5 |
| 253062_at | 460.5 | P | 374.2 | P | 483.6 | P | 397.7 | P | NPY5 |
| 253063_at | 594.9 | P | 832.9 | P | 692.4 | P | 646.2 | P | ACA2 |
| 253064_at | 36.7 | P | 63.6 | P | 47.2 | A | 40.1 | P | AtbZIP7 |
| 253065_at | 150.3 | P | 202 | P | 180.2 | P | 168.1 | P | AtGRF2 |
| 253066_at | 28.7 | A | 32.4 | A | 50.1 | A | 6.3 | A | ACS8 |
| 253067_at | 0.8 | A | 1.3 | A | 8.4 | A | 2.5 | A | MYB87 |
| 253068_at | 129.7 | P | 153 | P | 195 | P | 131.5 | P | --- |
| 253069_at | 3.7 | A | 2.4 | A | 1.8 | A | 3.8 | A | HKL3 |
| 253070_at | 2.3 | A | 0.6 | A | 3.3 | A | 0.4 | A | --- |
| 253071_at | 568.9 | P | 665.6 | P | 760.6 | P | 571.4 | P | --- |

| | | | | | | | | | |
|---|---|---|---|---|---|---|---|---|---|
| 253072_at | 10094 | P | 8797.9 | P | 8890.1 | P | 9099.4 | P | --- |
| 253073_at | 2787.1 | P | 401.4 | P | 2508.9 | P | 576.6 | P | CYP81F4 |
| 253074_at | 96.3 | P | 160.8 | P | 129 | P | 107.2 | P | --- |
| 253075_at | 99.4 | P | 121.2 | P | 106.1 | A | 99.8 | P | --- |
| 253076_at | 28.8 | P | 37 | P | 45.5 | A | 27.8 | P | ANAC076 |
| 253077_at | 2.1 | A | 0.6 | A | 1 | A | 2.1 | A | --- |
| 253078_at | 260.3 | P | 343.5 | P | 293.2 | P | 395.1 | P | --- |
| 253079_s_at | 2351.8 | P | 1797.4 | P | 1359.9 | P | 2046 | P | --- |
| 253080_at | 23 | A | 18.8 | A | 7.1 | A | 4.1 | A | --- |
| 253081_at | 1117.3 | P | 1364.1 | P | 863.1 | P | 1235.6 | P | --- |
| 253082_at | 35.7 | A | 22.6 | A | 24.7 | A | 18.7 | A | --- |
| 253083_at | 1356.6 | P | 1956.6 | P | 1396.3 | P | 1916 | P | ALDH3F1 |
| 253084_at | 35.4 | A | 41.7 | A | 51.2 | A | 41 | A | STY2 |
| 253085_s_at | 66.9 | A | 81.7 | P | 77.6 | A | 67 | P | --- |
| 253086_at | 1.9 | A | 2.3 | A | 2.6 | A | 3.4 | A | --- |
| 253087_at | 31.1 | A | 34.3 | A | 46.4 | A | 25.5 | A | PAP25 |
| 253088_at | 656.3 | P | 783.8 | P | 611.8 | P | 678.3 | P | FAH1 |
| 253089_at | 204.4 | P | 250.4 | P | 233.4 | P | 224.7 | P | --- |
| 253090_at | 1495.6 | P | 2509.9 | P | 1522.3 | P | 2504.4 | P | BGAL3 |
| 253091_at | 47.5 | A | 34.8 | A | 40 | A | 31.8 | P | CYP81D2 |
| 253092_at | 222.5 | P | 201.8 | P | 173.9 | P | 178.5 | P | --- |
| 253093_at | 564 | P | 621 | P | 446.8 | P | 500.8 | P | SRFR1 |
| 253094_at | 117.4 | P | 130.3 | P | 159.4 | P | 168.4 | P | --- |
| 253095_at | 1172 | P | 1629.3 | P | 1574.6 | P | 1589.1 | P | --- |
| 253096_at | 382 | P | 421.5 | P | 403.8 | P | 455.9 | P | CYP81D4 |
| 253097_at | 263.1 | P | 204.2 | P | 235.2 | P | 193.4 | P | CYP81D5 |
| 253098_at | 4.5 | A | 5.4 | A | 6.4 | A | 5.9 | A | CYP81D3 |
| 253099_s_at | 610.7 | P | 597.5 | P | 385.9 | P | 497.2 | P | --- |
| 253100_at | 31.8 | A | 27.5 | A | 13.6 | A | 39.9 | A | CYP81F3 |
| 253101_at | 161.5 | P | 181 | P | 190.4 | P | 217.3 | P | CYP91A2 |
| 253102_at | 4.3 | A | 49.8 | A | 15.2 | A | 20.1 | A | --- |
| 253103_at | 36.4 | A | 52.6 | P | 38.7 | A | 32.6 | M | --- |
| 253104_at | 765.2 | P | 479.7 | P | 492.8 | P | 541 | P | --- |
| 253105_at | 176.8 | P | 156.8 | P | 106.4 | M | 120.4 | P | --- |
| 253106_at | 130.4 | P | 153.3 | P | 143.5 | P | 161.6 | P | --- |
| 253107_at | 144.5 | P | 184.6 | P | 145.7 | P | 196.7 | P | --- |
| 253108_at | 60 | A | 32.3 | A | 84.1 | A | 52.2 | A | FD |
| 253109_at | 446.2 | P | 606.6 | P | 509.3 | P | 615.4 | P | MCA1 |
| 253110_at | 72.3 | A | 71.6 | P | 83.6 | A | 92.4 | P | --- |
| 253111_at | 169.1 | P | 135.5 | P | 163.8 | P | 180.6 | P | --- |
| 253112_at | 59.7 | P | 87.9 | P | 76.9 | P | 90.2 | P | APX5 |
| 253113_at | 88.9 | P | 204.3 | P | 73.2 | P | 139.2 | P | --- |
| 253114_at | 722.5 | P | 698.9 | P | 700.6 | P | 633.5 | P | ATGB2 |
| 253115_at | 279 | P | 309 | P | 275.6 | P | 254.4 | P | --- |
| 253116_at | 1228.7 | P | 1033 | P | 1319.1 | P | 1136.9 | P | --- |
| 253117_at | 187.8 | P | 296.7 | P | 247.1 | P | 278.1 | P | --- |
| 253118_at | 570.3 | P | 723.2 | P | 651.5 | P | 723 | P | ARAC6 |
| 253119_at | 609.8 | P | 639.3 | P | 669.3 | P | 751.1 | P | --- |
| 253120_at | 1376.3 | P | 1198.9 | P | 1314.9 | P | 1053.6 | P | PLDDELTA |
| 253121_at | 1021.2 | P | 1022.8 | P | 1106.3 | P | 925 | P | --- |
| 253122_at | 195.8 | P | 206.9 | P | 214.1 | P | 184.9 | P | --- |
| 253123_at | 21 | A | 9 | A | 11.5 | A | 8 | A | --- |
| 253124_at | 119.7 | P | 122.6 | P | 114.8 | P | 99.9 | P | ARO3 |
| 253125_at | 1281.4 | P | 1588.6 | P | 892.1 | P | 1015.9 | P | --- |
| 253126_at | 139.6 | P | 129.6 | P | 129.6 | P | 109.6 | P | --- |
| 253127_at | 40.8 | P | 39.3 | P | 83.1 | P | 56.8 | P | --- |
| 253128_at | 1.1 | A | 2.2 | A | 1.6 | A | 2.9 | A | CPK18 |
| 253129_at | 1130.1 | P | 772 | P | 1219.1 | P | 955.7 | P | CSDP1 |
| 253130_at | 217.1 | P | 160.9 | P | 201.6 | P | 208.7 | P | --- |
| 253131_at | 277.4 | P | 237.2 | P | 281.3 | P | 288 | P | WOX13 |
| 253132_at | 261.2 | P | 302.4 | P | 270.9 | P | 237.2 | P | NTL9 |
| 253133_at | 265.3 | P | 257.5 | P | 306.5 | P | 243.9 | P | NRPB1 |
| 253134_at | 1.9 | A | 2.7 | A | 4.4 | A | 1.1 | A | --- |
| 253135_at | 6782 | P | 5565.4 | P | 6159.3 | P | 5271.2 | P | --- |
| 253136_at | 324.8 | P | 419.3 | P | 362 | P | 382.4 | P | --- |
| 253137_at | 153.2 | P | 125.6 | P | 128.3 | P | 144.4 | P | --- |
| 253138_at | 2440.2 | P | 2161.5 | P | 2477.4 | P | 2139.6 | P | MRPL11 |
| 253139_at | 3474.8 | P | 4314.4 | P | 3354.9 | P | 4416.6 | P | AKR2 |
| 253140_at | 174.8 | P | 268.9 | P | 203.2 | P | 223.4 | P | RHA3B |
| 253141_at | 241.9 | P | 220.8 | P | 206.4 | P | 247.2 | P | --- |
| 253142_at | 2.4 | A | 2.9 | A | 10.7 | A | 29.5 | A | MLH3 |
| 253143_at | 278.5 | P | 268.7 | P | 280.6 | P | 289.4 | P | --- |
| 253144_at | 108.6 | P | 101.6 | P | 129.5 | P | 99.4 | P | --- |
| 253145_at | 58.8 | P | 57 | P | 45.3 | A | 53.6 | P | --- |
| 253146_at | 54 | A | 6.8 | A | 45.4 | A | 6.5 | A | --- |
| 253147_at | 471.6 | P | 276.6 | P | 376.5 | P | 301.1 | P | CONNEXIN 32 |
| 253148_at | 121.4 | P | 125.8 | P | 111.9 | P | 125.2 | P | CYCB2;2 |
| 253149_at | 2.8 | A | 1.9 | A | 4.9 | A | 5.6 | A | --- |
| 253150_at | 19.1 | A | 25.7 | A | 23.4 | A | 1.9 | A | --- |
| 253151_at | 1.3 | A | 0.5 | A | 0.8 | A | 0.7 | A | --- |
| 253152_at | 26.6 | A | 21.6 | A | 32.4 | A | 24.9 | A | --- |
| 253153_at | 4.8 | A | 5.6 | A | 3.1 | A | 6.7 | A | --- |
| 253154_at | 18.6 | A | 19 | A | 34.2 | P | 13.8 | A | --- |
| 253155_at | 219.4 | P | 137.5 | P | 157.4 | P | 150.1 | P | --- |
| 253156_at | 147.6 | P | 150.7 | P | 166.6 | P | 141.7 | P | --- |
| 253157_at | 121.6 | P | 147.1 | P | 147 | P | 130.7 | P | RecQl3 |
| 253158_at | 237.3 | P | 229.4 | P | 259.6 | P | 213.5 | P | --- |
| 253159_at | 578.2 | P | 496.3 | P | 556.1 | P | 513.9 | P | HMGB5 |
| 253160_at | 1339.7 | P | 1432.8 | P | 1451 | P | 1381.8 | P | --- |
| 253161_at | 513.6 | P | 138.3 | P | 390.5 | P | 68.3 | P | SEN1 |
| 253162_at | 10223 | P | 4235.2 | P | 8075.1 | P | 5314.5 | P | PSAT |
| 253163_at | 3379.8 | P | 3334 | P | 3011.4 | P | 2838.5 | P | --- |
| 253164_at | 10 | A | 26 | A | 11 | A | 3.9 | A | --- |
| 253165_at | 178.1 | P | 292.1 | P | 124.3 | P | 307.1 | P | --- |
| 253166_at | 4.2 | A | 43.4 | A | 60.4 | A | 87 | A | GLUR2 |
| 253167_at | 962.2 | P | 1275.6 | P | 1172.3 | P | 1289.9 | P | CPK5 |
| 253168_at | 51.3 | P | 25.9 | A | 37.7 | A | 56.8 | P | --- |
| 253169_at | 27.9 | P | 32 | M | 29.2 | A | 24.5 | A | --- |
| 253170_at | 68.9 | P | 80.5 | P | 116.1 | P | 98 | P | --- |
| 253171_at | 504 | P | 580.5 | P | 539.4 | P | 583.3 | P | --- |
| 253172_at | 823.4 | P | 359.5 | P | 512.8 | P | 482 | P | --- |
| 253173_at | 240.4 | P | 132 | P | 183.5 | P | 109.8 | P | --- |
| 253174_at | 7679.6 | P | 6957.3 | P | 8401.4 | P | 6924.7 | P | CAT2 |
| 253175_at | 372.2 | P | 316.6 | P | 289.1 | P | 268 | P | MSI3 |
| 253176_at | 128.5 | P | 105.6 | P | 124.4 | P | 111.1 | P | --- |
| 253177_s_at | 59.3 | P | 25.8 | A | 57.6 | P | 24.3 | A | --- |
| 253178_at | 30.4 | A | 29.4 | A | 11.2 | A | 33.8 | P | --- |
| 253179_at | 1.1 | A | 2.4 | A | 1.4 | A | 2.8 | A | --- |
| 253180_at | 2.9 | A | 1.3 | A | 2.2 | A | 1.7 | A | --- |

| | | | | | | | | | |
|---|---|---|---|---|---|---|---|---|---|
| 253181_at | 113.2 | P | 115.8 | P | 121 | P | 73.8 | P | LHT7 |
| 253182_at | 24.9 | A | 48.5 | P | 62.7 | P | 19 | A | --- |
| 253183_at | 1584.1 | P | 1382.6 | P | 1540.9 | P | 1389.9 | P | --- |
| 253184_at | 1476.1 | P | 1357.4 | P | 1168.4 | P | 1382.6 | P | BSK1 |
| 253185_at | 275.6 | P | 333.5 | P | 362.7 | P | 342 | P | --- |
| 253186_at | 113.3 | P | 170.1 | P | 142.1 | P | 164.1 | P | --- |
| 253187_at | 23.3 | A | 16.6 | A | 6.8 | A | 6.3 | A | --- |
| 253188_at | 595 | P | 813.2 | P | 583.7 | P | 803.5 | P | TMT2 |
| 253189_at | 359.7 | P | 487.6 | P | 565.4 | P | 554.2 | P | --- |
| 253190_at | 338.7 | P | 363.1 | P | 357.2 | P | 400.4 | P | --- |
| 253191_at | 191.5 | P | 313.2 | P | 269.8 | P | 377.5 | P | XCP1 |
| 253192_at | 33.3 | A | 10.9 | A | 8.1 | A | 23 | A | --- |
| 253193_at | 63.4 | A | 49.8 | A | 48.1 | A | 57.7 | A | --- |
| 253194_s_at | 7.3 | A | 2.8 | A | 23.7 | A | 34.1 | A | --- |
| 253195_at | 104.9 | A | 79.9 | A | 88.3 | A | 81.3 | A | --- |
| 253196_at | 1419 | P | 1375.8 | P | 1393.3 | P | 1363.5 | P | IDH1 |
| 253197_at | 2178.7 | P | 2786.7 | P | 2513.7 | P | 3227.3 | P | --- |
| 253198_at | 885.3 | P | 1057 | P | 794.1 | P | 1041.1 | P | --- |
| 253199_at | 235.5 | P | 259.8 | P | 173.5 | P | 307.4 | P | --- |
| 253200_at | 3509.6 | P | 3890.1 | P | 3824.8 | P | 4492.2 | P | AVA-P1 |
| 253201_at | 10313.6 | P | 10543.3 | P | 10219.2 | P | 10870.9 | P | SSR16 |
| 253202_at | 524.5 | P | 510.6 | P | 704.4 | P | 553.6 | P | --- |
| 253203_at | 18321.2 | P | 7828 | P | 22263.8 | P | 10521.1 | P | ADC2 |
| 253204_at | 1071.3 | P | 1109.3 | P | 977.6 | P | 1074.9 | P | AGB1 |
| 253205_at | 1679.9 | P | 1859.8 | P | 2119.4 | P | 2111.1 | P | ATCAP1 |
| 253206_at | 1156.3 | P | 1265.2 | P | 1139.7 | P | 1227.6 | P | SQS1 |
| 253207_at | 19.3 | A | 212.3 | P | 7.1 | A | 210 | P | --- |
| 253208_at | 1161.3 | P | 1244.5 | P | 1362.9 | P | 1472.2 | P | --- |
| 253209_at | 474.6 | P | 523.6 | P | 619 | P | 540.2 | P | --- |
| 253210_at | 187.6 | P | 201.3 | P | 215.3 | P | 159 | P | MTN2 |
| 253211_at | 20.3 | A | 20.6 | A | 29 | A | 6.7 | A | --- |
| 253212_s_at | 545.7 | P | 453.7 | P | 413 | P | 467 | P | XDH1 |
| 253213_at | 210.1 | P | 230.8 | P | 234.3 | P | 148.8 | P | --- |
| 253214_at | 10.2 | A | 11.9 | A | 38.6 | M | 20.9 | A | ARO1 |
| 253215_at | 338.9 | P | 3258.5 | P | 467.4 | P | 3008.6 | P | --- |
| 253216_at | 308.7 | P | 420.4 | P | 365.4 | P | 429.2 | P | --- |
| 253217_at | 1.1 | A | 57.3 | A | 26.6 | A | 32.7 | A | ADF9 |
| 253218_at | 1028.8 | P | 1195.7 | P | 785.9 | P | 1193.4 | P | SLP2 |
| 253219_at | 242.1 | P | 180.4 | P | 233 | P | 196.8 | P | AtMYB32 |
| 253220_s_at | 834.6 | P | 875.4 | P | 804.5 | P | 1219.7 | P | --- |
| 253221_at | 8869.9 | P | 8446.4 | P | 7725.6 | P | 8639 | P | ROC5 |
| 253222_at | 3.8 | A | 5.1 | A | 26.3 | A | 7.3 | A | --- |
| 253223_at | 4236.6 | P | 4758.5 | P | 4673.1 | P | 4912 | P | APX3 |
| 253224_at | 272.2 | P | 211.1 | P | 371.9 | P | 312.6 | P | --- |
| 253225_at | 464 | P | 510.1 | P | 522.7 | P | 505.8 | P | ARAC3 |
| 253226_at | 12 | A | 1.5 | A | 4.1 | A | 9.2 | A | BGAL11 |
| 253227_at | 53.7 | P | 67.5 | P | 65.2 | P | 59.4 | P | --- |
| 253228_at | 1223.4 | P | 1461.3 | P | 917.4 | P | 1361.7 | P | --- |
| 253229_at | 292.3 | P | 283.3 | P | 257.3 | P | 310.3 | P | --- |
| 253230_at | 5022.8 | P | 4682.5 | P | 5135.4 | P | 4856 | P | --- |
| 253231_at | 2516.2 | P | 3101.9 | P | 2732.2 | P | 3001.3 | P | --- |
| 253232_at | 115.7 | P | 178.7 | P | 140.7 | P | 155.5 | P | TAF8 |
| 253233_at | 1027.4 | P | 1859 | P | 1401 | P | 1725 | P | --- |
| 253234_at | 1042.6 | P | 965.7 | P | 1103.8 | P | 946.3 | P | --- |
| 253235_at | 2272.6 | P | 2617.2 | P | 2315.8 | P | 2489.8 | P | HDR |
| 253236_at | 380.3 | P | 355.7 | P | 344.1 | P | 378.3 | P | ARI1 |
| 253237_at | 1079.7 | P | 1143.7 | P | 923.2 | P | 1145.5 | P | ALDH3I1 |
| 253238_at | 935.3 | P | 1154.1 | P | 876.4 | P | 994.5 | P | --- |
| 253239_at | 322.2 | P | 288.4 | P | 213.9 | P | 365.7 | P | --- |
| 253240_at | 3.1 | A | 8.8 | A | 14.5 | A | 7.6 | A | KCS17 |
| 253241_at | 53.4 | M | 34.1 | A | 30.6 | A | 19.1 | A | KCS18 |
| 253242_at | 307.8 | P | 232.7 | P | 323.8 | P | 225.9 | P | CIB1 |
| 253243_at | 107.5 | P | 113.5 | P | 144.7 | P | 106.8 | P | --- |
| 253244_at | 110.2 | M | 45.5 | A | 91.6 | A | 66.3 | A | COW1 |
| 253245_at | 4760.4 | P | 2029.5 | P | 4281.8 | P | 2557.1 | P | GBF6 |
| 253246_at | 24.1 | A | 52.2 | P | 28 | A | 36.3 | A | --- |
| 253247_at | 355.8 | P | 365.6 | P | 473.3 | P | 394.5 | P | BLH6 |
| 253248_at | 8134.4 | P | 6176.2 | P | 8901.2 | P | 6417.4 | P | --- |
| 253249_at | 336 | P | 271.4 | P | 284.9 | P | 291.6 | P | --- |
| 253250_at | 17.2 | A | 41.6 | A | 52.5 | A | 25.9 | A | --- |
| 253251_at | 288.7 | P | 457.3 | P | 336.3 | P | 456.7 | P | --- |
| 253252_at | 468.7 | P | 540.7 | P | 455.4 | P | 411.8 | P | ATASE2 |
| 253253_at | 162.4 | M | 371.9 | P | 166.6 | M | 328.3 | P | --- |
| 253254_at | 54.3 | A | 52.3 | P | 14.3 | A | 47.9 | A | SQS2 |
| 253255_at | 153.4 | P | 759.3 | P | 348.9 | P | 769 | P | --- |
| 253256_at | 123.7 | P | 126.3 | P | 175.2 | P | 136.7 | P | --- |
| 253257_at | 338.3 | P | 462.7 | P | 466.9 | P | 309.2 | P | XLG2 |
| 253258_at | 28.1 | P | 27.4 | A | 32.8 | A | 32.2 | P | --- |
| 253259_at | 20.2 | A | 30.2 | P | 25.9 | P | 20.3 | A | RRTF1 |
| 253260_at | 37.8 | P | 51.9 | P | 32.5 | A | 78.4 | P | AT4G34420 /// CHB3 |
| 253261_at | 178.3 | P | 193.9 | P | 181.7 | P | 205.1 | P | CHB3 |
| 253262_at | 8.7 | A | 2.5 | A | 31.5 | A | 23.5 | A | --- |
| 253263_at | 267.8 | P | 312.4 | P | 245.4 | P | 307.5 | P | ABF3 |
| 253264_at | 506.1 | P | 491.9 | P | 628 | P | 582.5 | P | OST1 |
| 253265_at | 649 | P | 575.6 | P | 602.5 | P | 608.5 | P | --- |
| 253266_s_at | 273.7 | P | 365.5 | P | 303.6 | P | 324.7 | P | --- |
| 253267_at | 859.9 | P | 1199.3 | P | 959.2 | P | 1119.3 | P | --- |
| 253268_s_at | 3314.7 | P | 3248.8 | P | 3330.8 | P | 3994.6 | P | UGT73B2 |
| 253269_at | 157.7 | P | 212.9 | P | 99.8 | P | 154.5 | P | --- |
| 253270_at | 331.9 | P | 559.1 | P | 504.8 | P | 515.4 | P | CYCD3;1 |
| 253271_s_at | 103.4 | P | 55.9 | A | 103.7 | P | 47 | A | ASK12 |
| 253272_at | 5005.3 | P | 6132 | P | 6540.7 | P | 6472.5 | P | 41518 |
| 253273_at | 3471.9 | P | 3245.7 | P | 2178.1 | P | 3094.3 | P | --- |
| 253274_at | 6443.1 | P | 2846 | P | 5885.8 | P | 3947 | P | EDA9 |
| 253275_at | 302.3 | P | 422.6 | P | 275.9 | P | 386.8 | P | --- |
| 253276_at | 9791.3 | P | 6824.5 | P | 9016.9 | P | 8281.1 | P | --- |
| 253277_at | 2393.5 | P | 1405.3 | P | 1481.8 | P | 1500.9 | P | ATCAD5 |
| 253278_at | 110.2 | P | 264.3 | P | 210.9 | P | 244.9 | P | --- |
| 253279_at | 893.5 | P | 952.8 | P | 826.7 | P | 803.9 | P | MCCB |
| 253280_at | 4701.9 | P | 4908.6 | P | 5274.9 | P | 4987.7 | P | PAB2 |
| 253281_at | 525 | P | 466.9 | P | 781.9 | P | 642.5 | P | UGT73B1 |
| 253282_at | 3766.6 | P | 4466.2 | P | 3274 | P | 4469.1 | P | LEJ1 |
| 253283_at | 738.4 | P | 911.9 | P | 743.3 | P | 878.5 | P | --- |
| 253284_at | 561.1 | P | 832.3 | P | 569.3 | P | 607.6 | P | --- |
| 253285_at | 135.6 | P | 256.2 | P | 178.6 | P | 225.2 | P | KCS16 |
| 253286_at | 1645 | P | 1516.2 | P | 1688.8 | P | 1341.3 | P | FUC95A |
| 253287_at | 588.9 | P | 467.5 | P | 572.1 | P | 520.7 | P | --- |
| 253288_at | 117.2 | P | 114.3 | P | 155.3 | P | 111.5 | P | --- |
| 253289_at | 17.5 | A | 4.2 | A | 21.6 | A | 10.9 | A | --- |

| | | | | | | | | | |
|---|---|---|---|---|---|---|---|---|---|
| 253290_at | 1.4 | A | 2 | A | 3.7 | A | 0.8 | A | --- |
| 253291_at | 9329.3 | P | 8315.3 | P | 10004.5 | P | 9066.1 | P | --- |
| 253292_at | 252.1 | P | 239 | P | 236.2 | P | 274.4 | P | --- |
| 253293_at | 75.9 | P | 33.9 | A | 78.9 | P | 87.4 | P | --- |
| 253294_at | 773.6 | P | 346.9 | P | 578.6 | P | 416.9 | P | --- |
| 253295_at | 703 | P | 691.6 | P | 718.8 | P | 708.7 | P | --- |
| 253296_at | 148.4 | P | 154.3 | P | 133.1 | P | 135.4 | P | --- |
| 253297_at | 102.4 | P | 77.2 | P | 120.3 | P | 111.5 | P | --- |
| 253298_at | 6.6 | A | 37.2 | P | 33.1 | A | 26.6 | P | --- |
| 253299_at | 14 | A | 11.4 | A | 36.1 | A | 31.2 | A | --- |
| 253300_at | 1554.2 | P | 1620.3 | P | 1826.3 | P | 1615.4 | P | BCA5 |
| 253301_at | 5.3 | A | 6 | A | 10.1 | A | 4 | A | --- |
| 253302_at | 1066.4 | P | 1337.2 | P | 1211.3 | P | 1174.8 | P | --- |
| 253303_at | 1198.9 | P | 954.1 | P | 1174.5 | P | 1012.8 | P | --- |
| 253304_at | 2603.8 | P | 2537.8 | P | 2404.5 | P | 2068.9 | P | --- |
| 253305_at | 336.1 | P | 465.4 | P | 387.2 | P | 405.9 | P | --- |
| 253306_at | 542 | P | 441.9 | P | 457.8 | P | 507.6 | P | DRP3A |
| 253307_at | 2690.9 | P | 2285.7 | P | 2354.4 | P | 2464.9 | P | --- |
| 253308_at | 4610.3 | P | 4706.9 | P | 5328 | P | 4685.9 | P | AGD2 |
| 253309_at | 40.7 | P | 60.4 | P | 51.6 | A | 51.9 | M | CER4 |
| 253310_at | 651.2 | P | 607.9 | P | 552.3 | P | 595.5 | P | EX1 |
| 253311_s_at | 61.7 | P | 41.7 | P | 58.6 | P | 44.5 | M | --- |
| 253312_s_at | 1.2 | A | 3 | A | 1.6 | A | 2.6 | A | --- |
| 253313_at | 38 | A | 27.6 | A | 62 | M | 41.6 | P | --- |
| 253314_at | 449.1 | P | 404.6 | P | 469.5 | P | 474.5 | P | --- |
| 253315_at | 1.6 | A | 19.6 | A | 3 | A | 0.9 | A | --- |
| 253316_s_at | 28.9 | A | 30 | A | 40.1 | A | 18.4 | A | --- |
| 253317_at | 34.9 | A | 85.5 | P | 56.4 | A | 25.4 | A | --- |
| 253318_at | 20.8 | A | 25.2 | A | 6.1 | A | 3.3 | A | --- |
| 253319_at | 63.8 | A | 68 | A | 68 | A | 55.7 | P | EMB2758 |
| 253320_at | 197.7 | P | 209.9 | P | 198.6 | P | 161 | P | --- |
| 253321_at | 600.4 | P | 577.3 | P | 452.9 | P | 553.8 | P | --- |
| 253322_at | 612.7 | P | 336.4 | P | 314.9 | P | 363.5 | P | --- |
| 253323_at | 450.5 | P | 344.1 | P | 250.6 | P | 331.9 | P | --- |
| 253324_at | 480.7 | P | 482.4 | P | 460.8 | P | 368.9 | P | --- |
| 253325_at | 89 | A | 91.1 | A | 99.6 | A | 79.2 | A | --- |
| 253326_at | 197.4 | P | 203.8 | P | 156.3 | P | 219.3 | P | --- |
| 253327_at | 13.2 | A | 30.2 | A | 4.8 | A | 14.1 | A | MYB69 |
| 253328_at | 316.9 | P | 413.2 | P | 302.8 | P | 318.1 | P | ATNAP13 |
| 253329_at | 468.3 | P | 677.2 | P | 467.8 | P | 579.1 | P | --- |
| 253330_at | 897.2 | P | 700.7 | P | 923 | P | 734.6 | P | KUP5 |
| 253331_at | 829.1 | P | 656.5 | P | 628.8 | P | 704.6 | P | --- |
| 253332_at | 347.4 | P | 240 | P | 381.1 | P | 234.7 | P | --- |
| 253333_at | 3404.2 | P | 3882.1 | P | 3886.4 | P | 3895.9 | P | DHS2 |
| 253334_at | 1188.9 | P | 905.9 | P | 945.9 | P | 840.7 | P | --- |
| 253335_at | 1082.3 | P | 1084 | P | 978.2 | P | 1110 | P | --- |
| 253336_at | 3540.9 | P | 2868 | P | 3524.5 | P | 2947.8 | P | EIF3K |
| 253337_at | 580.8 | P | 988.6 | P | 718.3 | P | 1006.1 | P | hda14 |
| 253338_at | 848.4 | P | 750.9 | P | 778.7 | P | 826.2 | P | BAK1 |
| 253339_at | 345.8 | P | 215.3 | P | 347.3 | P | 249 | P | PAA1 |
| 253340_s_at | 371.5 | P | 636.8 | P | 395.6 | P | 736.8 | P | CDC20.2 |
| 253341_at | 485 | P | 505.1 | P | 465.5 | P | 531.6 | P | --- |
| 253342_at | 773.7 | P | 952.9 | P | 935.3 | P | 1042 | P | PAA1 |
| 253343_at | 3722.2 | P | 2081.2 | P | 2347.1 | P | 2674.3 | P | --- |
| 253344_at | 73.8 | P | 168.3 | P | 74.2 | P | 235.4 | P | --- |
| 253345_at | 37.1 | A | 18.5 | P | 15.9 | A | 17.4 | A | BCA5 |
| 253346_at | 1.5 | A | 3.4 | A | 2.8 | A | 1.8 | A | --- |
| 253347_at | 2.3 | A | 6.7 | A | 23.5 | A | 22.8 | A | --- |
| 253348_at | 24.1 | P | 40.8 | A | 22.3 | A | 26 | A | --- |
| 253349_at | 63 | P | 35.7 | A | 93.8 | A | 80.1 | M | --- |
| 253350_at | 102.1 | P | 114 | P | 107.1 | P | 101.2 | P | --- |
| 253351_at | 811.3 | P | 689.7 | P | 760.5 | P | 730.6 | P | --- |
| 253352_at | 4.6 | A | 29.5 | A | 44.9 | A | 31.7 | A | --- |
| 253353_at | 2.1 | A | 5 | A | 3.3 | A | 9.2 | A | --- |
| 253354_at | 4.9 | A | 3.5 | A | 37.1 | A | 19.4 | A | --- |
| 253355_at | 1231.7 | P | 1513.9 | P | 490 | P | 1962.5 | P | --- |
| 253356_at | 7.4 | A | 26.4 | A | 2.5 | A | 3.7 | A | --- |
| 253357_at | 191.1 | P | 290.4 | P | 244.6 | P | 256.2 | P | --- |
| 253358_at | 4114.6 | P | 8294 | P | 2975.1 | P | 7931 | P | GAMMA-VPE |
| 253359_at | 458 | P | 779.3 | P | 454.7 | P | 702.9 | P | --- |
| 253360_at | 2236.4 | P | 1813.5 | P | 1967.4 | P | 1907.9 | P | APM1 |
| 253361_at | 115.5 | P | 146.6 | P | 149.3 | P | 97.8 | P | --- |
| 253362_s_at | 1068.5 | P | 819.7 | P | 986.3 | P | 840.3 | P | --- |
| 253363_at | 61 | A | 85.5 | M | 87.8 | A | 96.4 | A | --- |
| 253364_at | 99.8 | P | 107 | P | 78 | A | 119.1 | P | --- |
| 253365_at | 61.8 | P | 62.4 | P | 66.9 | P | 58.3 | M | --- |
| 253366_at | 164 | P | 120.2 | P | 158.4 | P | 164.9 | P | --- |
| 253367_at | 655.5 | P | 549.6 | P | 716.3 | P | 533.7 | P | --- |
| 253368_at | 262.5 | P | 283.2 | P | 258.2 | P | 260.3 | P | XI-I |
| 253369_at | 605.1 | P | 585.5 | P | 597.6 | P | 579.7 | P | --- |
| 253370_at | 4 | A | 4.4 | A | 33.1 | A | 1.5 | A | --- |
| 253371_at | 197.6 | P | 262.1 | P | 180.7 | P | 216.9 | P | --- |
| 253372_at | 896.5 | P | 1856.1 | P | 784.6 | P | 1540.8 | P | PME44 |
| 253373_at | 1197.8 | P | 526 | P | 576.9 | P | 425.3 | P | --- |
| 253374_at | 374.7 | P | 381.7 | P | 438 | P | 428.9 | P | --- |
| 253375_at | 17.3 | A | 8.9 | A | 19.2 | A | 37 | A | --- |
| 253376_at | 11.8 | A | 1.7 | A | 3.6 | A | 3 | A | --- |
| 253377_at | 1082.5 | P | 1477.9 | P | 941.6 | P | 988.2 | P | ADR1-L1 |
| 253378_at | 40.2 | A | 25.7 | A | 13.2 | A | 20.2 | A | --- |
| 253379_at | 32.2 | A | 39 | A | 59.9 | A | 57.4 | A | PGSIP3 |
| 253380_at | 41.2 | A | 60.6 | A | 46.9 | A | 21.8 | A | PGSIP3 |
| 253381_at | 1573.4 | P | 1483.6 | P | 1498.5 | P | 1422.8 | P | --- |
| 253382_at | 1036.6 | P | 1014.4 | P | 850.6 | P | 1054.8 | P | --- |
| 253383_at | 377.7 | P | 392 | P | 300.3 | P | 356.5 | P | --- |
| 253384_at | 822.2 | P | 960.8 | P | 859.4 | P | 929.5 | P | --- |
| 253385_at | 912.4 | P | 884.1 | P | 909.5 | P | 886.6 | P | --- |
| 253386_at | 2143.4 | P | 2258.2 | P | 1994.6 | P | 2075.4 | P | SQD1 |
| 253387_at | 9787.9 | P | 9171.5 | P | 14468.6 | P | 10356.9 | P | AtGLDP1 |
| 253388_at | 819 | P | 707.3 | P | 885.3 | P | 730 | P | --- |
| 253389_at | 301.4 | P | 346.3 | P | 266.5 | P | 363.9 | P | --- |
| 253390_at | 154.4 | P | 124.4 | P | 274.4 | P | 146.5 | P | --- |
| 253391_at | 931.3 | P | 890.1 | P | 1000.9 | P | 1003 | P | --- |
| 253392_at | 93.8 | P | 78.7 | P | 54.2 | A | 77.9 | P | ATKC1 |
| 253393_at | 336.2 | P | 319.9 | P | 326 | P | 349.5 | P | GLB3 |
| 253394_at | 526.5 | P | 475.9 | P | 416.3 | P | 521.5 | P | VTE1 |
| 253395_at | 880.1 | P | 652.1 | P | 679.7 | P | 647.6 | P | AME3 |
| 253396_at | 988.8 | P | 803.1 | P | 1164.1 | P | 990.3 | P | AtLa1 |
| 253397_at | 148.4 | P | 158.7 | P | 166.8 | P | 171.6 | P | --- |
| 253398_at | 72 | P | 76.3 | P | 99.3 | P | 77.1 | P | CCD8 |

| | | | | | | | | | |
|---|---|---|---|---|---|---|---|---|---|
| 253399_at | 809.6 | P | 792.4 | P | 556.2 | P | 748 | P | nPAP |
| 253400_at | 3.8 | A | 3.5 | A | 6.9 | A | 3.9 | A | --- |
| 253401_at | 36.5 | A | 156.1 | P | 80.8 | A | 173.6 | P | --- |
| 253402_at | 137.1 | A | 151.8 | P | 181.8 | A | 190.8 | P | ATHB-8 |
| 253403_at | 259.9 | P | 380.9 | P | 276.5 | P | 384 | P | AtAUR1 |
| 253404_at | 238.6 | P | 253.3 | P | 318.6 | P | 250.8 | P | PFK6 |
| 253405_at | 113.7 | P | 211.5 | P | 86.3 | A | 208.2 | P | --- |
| 253406_at | 64.2 | P | 58.1 | P | 95.7 | P | 109.8 | P | --- |
| 253407_at | 375.3 | P | 318.6 | P | 480.3 | P | 272.6 | P | --- |
| 253408_at | 28.1 | A | 67.1 | M | 43.5 | A | 52 | P | --- |
| 253409_at | 749.2 | P | 808.4 | P | 801.8 | P | 738.1 | P | --- |
| 253410_at | 108.6 | P | 123 | P | 175.8 | P | 95.4 | P | --- |
| 253411_at | 809.2 | P | 856.6 | P | 959.5 | P | 887.2 | P | ATH1 |
| 253412_at | 358.6 | P | 592.7 | P | 495.5 | P | 551 | P | CBL10 |
| 253413_at | 7.6 | A | 46.3 | A | 69.2 | A | 19 | A | ZIP9 |
| 253414_at | 1782.7 | P | 1412.4 | P | 1405.3 | P | 1084.9 | P | EDA39 |
| 253415_at | 422.3 | P | 329.5 | P | 487 | P | 357 | P | --- |
| 253416_at | 76.1 | P | 43.8 | P | 42 | A | 31.6 | A | --- |
| 253417_at | 280.8 | P | 221.7 | P | 307.9 | P | 280.2 | P | PC-MYB1 |
| 253418_at | 617.7 | P | 804.7 | P | 800 | P | 784.9 | P | --- |
| 253419_at | 6.4 | A | 5.4 | A | 2.4 | A | 13.1 | A | --- |
| 253420_at | 8517.7 | P | 9801 | P | 10546.3 | P | 10313.8 | P | --- |
| 253421_at | 2482.6 | P | 2157.3 | P | 1545.5 | P | 1873.1 | P | --- |
| 253422_at | 1081.9 | P | 875.5 | P | 1080.6 | P | 934.1 | P | --- |
| 253423_at | 22.4 | A | 28.8 | A | 5.3 | A | 27.9 | A | IAA29 |
| 253424_at | 645.4 | P | 812.5 | P | 756.9 | P | 793.1 | P | --- |
| 253425_at | 228.8 | P | 334.7 | P | 303.6 | P | 301.4 | P | --- |
| 253426_at | 7.9 | A | 17.6 | A | 4.3 | A | 6.8 | A | --- |
| 253427_at | 1524.8 | P | 1553.1 | P | 1720 | P | 1671 | P | --- |
| 253428_at | 1415.2 | P | 2034.4 | P | 1464 | P | 1867.4 | P | CESA1 |
| 253429_at | 755.6 | P | 733.2 | P | 775.7 | P | 851.8 | P | --- |
| 253430_at | 49.2 | P | 49.5 | P | 81.6 | A | 72.7 | P | --- |
| 253431_at | 149.1 | P | 167.5 | P | 127.4 | P | 137 | P | --- |
| 253432_at | 121.2 | P | 112.1 | P | 146.1 | P | 118.5 | P | --- |
| 253433_s_at | 22.2 | A | 11.9 | A | 36.5 | A | 13.5 | A | --- |
| 253434_at | 35.2 | P | 27.4 | A | 51.7 | A | 38 | A | AKT5 |
| 253435_at | 546.3 | P | 376.6 | P | 598.5 | P | 489.8 | P | SHS1 |
| 253436_at | 6919.6 | P | 5841.9 | P | 5196.7 | P | 6067.9 | P | --- |
| 253437_at | 569.2 | P | 1571.1 | P | 458 | P | 941.5 | P | --- |
| 253438_at | 1453.8 | P | 1171.4 | P | 1882.3 | P | 1274.9 | P | SHM3 |
| 253439_at | 36.1 | A | 50 | A | 52.9 | A | 46.8 | A | YUC1 |
| 253440_at | 507.2 | P | 842.6 | P | 572.8 | P | 797.3 | P | TIFY8 |
| 253441_at | 190.8 | P | 188.4 | P | 160.7 | P | 152.5 | P | --- |
| 253442_at | 2302.5 | P | 2371.2 | P | 2265 | P | 2537.6 | P | --- |
| 253443_at | 696.7 | P | 834.8 | P | 824.8 | P | 806.8 | P | LUG |
| 253444_at | 573.5 | P | 512.1 | P | 356.3 | P | 545.9 | P | --- |
| 253445_at | 485.9 | P | 477.6 | P | 519 | P | 501 | P | --- |
| 253446_at | 364.6 | P | 350.6 | P | 291.5 | P | 389.9 | P | --- |
| 253447_at | 30.5 | A | 51.9 | P | 81 | A | 31.9 | A | --- |
| 253448_at | 783.2 | P | 794 | P | 837.9 | P | 845.3 | P | --- |
| 253449_at | 22.8 | A | 21.5 | A | 24.3 | A | 14.8 | A | --- |
| 253450_at | 32.2 | A | 49.6 | P | 61.7 | A | 28.5 | A | --- |
| 253451_at | 240.5 | P | 209.8 | P | 243.1 | P | 247.7 | P | PC-MYB1 |
| 253452_at | 274.9 | P | 241.7 | P | 255 | P | 205.3 | P | --- |
| 253453_at | 2347.4 | P | 1360.6 | P | 1328.6 | P | 1421.4 | P | --- |
| 253454_at | 4.9 | A | 5.9 | A | 4.9 | A | 3.9 | A | --- |
| 253455_at | 1358.1 | P | 2179.6 | P | 1475.5 | P | 1966.5 | P | --- |
| 253456_at | 206.8 | P | 160.8 | P | 191.6 | P | 129.3 | P | --- |
| 253457_at | 2876.8 | P | 3713.9 | P | 2819.7 | P | 3383.5 | P | --- |
| 253458_at | 304.5 | P | 243.7 | P | 248.1 | P | 354.2 | P | --- |
| 253459_at | 11.3 | A | 23.2 | A | 28.6 | A | 2.5 | A | --- |
| 253460_at | 317.3 | P | 470.5 | P | 382.6 | P | 327.5 | P | --- |
| 253461_at | 1.6 | A | 18.7 | A | 3.2 | A | 4.4 | A | CYP96A2 |
| 253462_at | 2096.4 | P | 1897.4 | P | 2057.6 | P | 1805.6 | P | ATVAMP711 |
| 253463_at | 33 | A | 29.7 | A | 52.8 | A | 20.3 | A | --- |
| 253464_at | 271 | P | 274.4 | P | 240.6 | P | 270.1 | P | --- |
| 253465_at | 607.3 | P | 516.1 | P | 484.7 | P | 677.1 | P | --- |
| 253466_at | 567.1 | P | 521.4 | P | 428.3 | P | 423.2 | P | KNAT5 |
| 253467_at | 333.6 | P | 422.3 | P | 342.3 | P | 432.4 | P | --- |
| 253468_at | 266.1 | P | 287.5 | P | 330.6 | P | 268.3 | P | --- |
| 253469_at | 590 | P | 604 | P | 627.1 | P | 520 | P | ATPANK2 |
| 253470_s_at | 3212.8 | P | 3090.3 | P | 3126.1 | P | 3226.1 | P | SDH3-2 |
| 253471_at | 23.3 | A | 27.5 | A | 37.1 | A | 3.1 | A | --- |
| 253472_at | 13.5 | A | 4.9 | A | 20.4 | A | 6.3 | A | --- |
| 253473_at | 583.4 | P | 547.5 | P | 576.9 | P | 514.8 | P | --- |
| 253474_at | 197.9 | P | 312.5 | P | 205.5 | P | 278.2 | P | --- |
| 253475_at | 210.6 | P | 283.4 | P | 257.8 | P | 283.4 | P | --- |
| 253476_at | 239.7 | P | 259.8 | P | 204.9 | P | 257.4 | P | SD2-5 |
| 253477_at | 457.3 | P | 527.7 | P | 372.6 | P | 508.6 | P | APX6 |
| 253478_at | 92.3 | P | 190.1 | P | 109.5 | P | 172.6 | P | --- |
| 253479_at | 176.3 | P | 168.3 | P | 176.8 | P | 149.3 | P | --- |
| 253480_at | 550.9 | P | 1026 | P | 784.8 | P | 1077.5 | P | --- |
| 253481_at | 4739.7 | P | 4145.8 | P | 4706.1 | P | 4523.5 | P | ASP5 |
| 253482_at | 4599.7 | P | 3859.6 | P | 4221.5 | P | 4020.1 | P | --- |
| 253483_at | 13 | A | 70.8 | A | 45.6 | A | 32.2 | A | --- |
| 253484_at | 974.7 | P | 972.2 | P | 804.4 | P | 1046.5 | P | TAFII15 |
| 253485_at | 765.2 | P | 418.8 | P | 655.4 | P | 336.1 | P | WRKY18 |
| 253486_at | 207.4 | P | 277.4 | P | 229.4 | P | 305 | P | --- |
| 253487_at | 6375.5 | P | 5420.3 | P | 6695.7 | P | 6120.8 | P | RPS6 |
| 253488_at | 7 | A | 1 | A | 1.7 | A | 13.9 | A | REM1 |
| 253489_at | 1941 | P | 1632.5 | P | 1714.6 | P | 1582.7 | P | MGD1 |
| 253490_at | 637.2 | P | 566.1 | P | 820 | P | 639.3 | P | --- |
| 253491_at | 156.8 | P | 165.1 | P | 225.4 | P | 177.2 | P | --- |
| 253492_at | 495.2 | P | 383.5 | P | 594.1 | P | 448 | P | --- |
| 253493_at | 210.8 | P | 226.1 | P | 198.8 | P | 224.4 | P | ENP |
| 253494_at | 12.8 | A | 3.5 | A | 17.2 | A | 2.8 | A | --- |
| 253495_at | 418.8 | P | 516.9 | P | 472.5 | P | 535.7 | P | PGR3 |
| 253496_at | 167.4 | P | 296.6 | P | 253.9 | P | 346.6 | P | ATGPX7 |
| 253497_at | 395.6 | P | 436.5 | P | 457.8 | P | 425.4 | P | --- |
| 253498_at | 115.7 | P | 117.8 | P | 102.2 | P | 99.1 | P | --- |
| 253499_at | 4.2 | A | 4.3 | A | 3.9 | A | 4.4 | A | --- |
| 253500_at | 266.2 | P | 374 | P | 415.2 | P | 372.1 | P | ARR10 |
| 253501_at | 234.6 | P | 253 | P | 288.3 | P | 307.9 | P | HSL1 |
| 253502_at | 1.3 | A | 6.8 | A | 10.9 | A | 0.6 | A | CYP82C4 |
| 253503_at | 2.6 | A | 3.4 | A | 1 | A | 0.8 | A | CYP82C3 |
| 253504_at | 19.4 | A | 26.2 | A | 25.4 | A | 12 | A | --- |
| 253505_at | 1.1 | A | 1.3 | A | 13.6 | A | 0.9 | A | CYP82C2 |
| 253506_at | 153.6 | P | 195.2 | P | 159.9 | P | 157.6 | P | --- |
| 253507_at | 28.7 | P | 75.1 | M | 65 | A | 35.1 | A | --- |

| | | | | | | | | | |
|---|---|---|---|---|---|---|---|---|---|
| 253508_at | 0.8 | A | 10.8 | A | 2.4 | A | 3.9 | A | --- |
| 253509_at | 37.5 | P | 32 | P | 61.4 | P | 40.7 | P | GLR2.4 |
| 253510_at | 30.7 | P | 93.8 | P | 74.1 | P | 110.5 | P | GDU1 |
| 253511_at | 5.7 | A | 1.7 | A | 2.2 | A | 3.7 | A | --- |
| 253512_at | 217.8 | P | 246.4 | P | 146.5 | P | 265.2 | P | WIN2 |
| 253513_at | 8.4 | A | 25.7 | A | 5.8 | A | 2.3 | A | --- |
| 253514_at | 73.5 | P | 94.9 | P | 147.1 | P | 108.3 | P | --- |
| 253515_at | 17.2 | A | 15.6 | A | 10.1 | A | 4.9 | A | --- |
| 253516_at | 84.2 | P | 124.3 | P | 154.7 | P | 112.9 | P | --- |
| 253517_at | 601.7 | P | 708.6 | P | 834.1 | P | 744.5 | P | --- |
| 253518_at | 118.3 | P | 141.3 | P | 116 | P | 129.3 | P | --- |
| 253519_at | 341.4 | P | 250 | P | 280.4 | P | 208.2 | P | --- |
| 253520_at | 379.5 | P | 280.4 | P | 345.4 | P | 296.8 | P | --- |
| 253521_at | 3434.8 | P | 2988.7 | P | 3427.3 | P | 3007.5 | P | PBA1 |
| 253522_at | 1566.7 | P | 1155.7 | P | 1711.3 | P | 1348.8 | P | --- |
| 253523_at | 1790.7 | P | 2078 | P | 1890.4 | P | 1991.2 | P | --- |
| 253524_at | 274.8 | P | 354.2 | P | 367.7 | P | 361.7 | P | --- |
| 253525_at | 59.1 | A | 122.6 | P | 121.6 | P | 121.9 | A | --- |
| 253526_at | 755.7 | P | 710.1 | P | 668.7 | P | 638.2 | P | --- |
| 253527_at | 1.8 | A | 2.3 | A | 4 | A | 8 | A | --- |
| 253528_s_at | 1548.5 | P | 2015.3 | P | 1678.5 | P | 1980 | P | --- |
| 253529_at | 37.9 | P | 17.7 | A | 28.5 | P | 20.7 | P | --- |
| 253530_at | 1181.1 | P | 1173.1 | P | 1310.4 | P | 1128.1 | P | --- |
| 253531_at | 254 | P | 263.5 | P | 298.6 | P | 350.8 | P | ATEXO70G1 |
| 253532_at | 61.3 | P | 51.1 | P | 41.6 | A | 61.7 | P | --- |
| 253533_at | 659.8 | P | 563.4 | P | 661.8 | P | 611.9 | P | ATCSLC5 |
| 253534_at | 10707 | P | 7373.7 | P | 7881.7 | P | 6876.3 | P | CYP83B1 |
| 253535_at | 268.1 | P | 403.8 | P | 395 | P | 415.4 | P | WRKY11 |
| 253536_at | 1387.4 | P | 1370.2 | P | 1691.6 | P | 1471 | P | SRZ-22 |
| 253537_at | 450 | P | 515.5 | P | 414.9 | P | 480.1 | P | HCF153 |
| 253538_at | 536.3 | P | 487.8 | P | 634.1 | P | 575.6 | P | --- |
| 253539_at | 267.7 | P | 321.6 | P | 375.5 | P | 324.8 | P | --- |
| 253540_at | 7 | A | 18.9 | A | 5.2 | A | 4.3 | A | --- |
| 253541_at | 26.5 | A | 15.7 | A | 24.4 | A | 3.6 | A | --- |
| 253542_at | 284 | P | 391.1 | P | 319.3 | P | 307.4 | P | UBP18 |
| 253543_at | 68.8 | P | 73.4 | P | 74.5 | P | 81.6 | P | --- |
| 253544_at | 1.8 | A | 1 | A | 2.1 | A | 3.1 | A | --- |
| 253545_at | 621.2 | P | 713.9 | P | 689.3 | P | 788.2 | P | --- |
| 253546_at | 915 | P | 1080.9 | P | 1170.9 | P | 1041.9 | P | --- |
| 253547_at | 6438.2 | P | 6228.9 | P | 6620.1 | P | 6640.6 | P | FAD6 |
| 253548_at | 1129.9 | P | 1171.8 | P | 945.9 | P | 1186.5 | P | --- |
| 253549_at | 777.3 | P | 643.9 | P | 826.9 | P | 701.6 | P | NFD1 |
| 253550_at | 3727.8 | P | 2509.6 | P | 3421.4 | P | 2798.8 | P | SIP3 |
| 253551_at | 1047.8 | P | 1116.6 | P | 1050.6 | P | 1102.5 | P | --- |
| 253552_at | 915.6 | P | 977.2 | P | 807.7 | P | 969.7 | P | UBP24 |
| 253553_at | 281.1 | P | 312.4 | P | 373 | P | 313.3 | P | --- |
| 253554_at | 84.8 | M | 45.5 | P | 83.3 | M | 61.3 | P | --- |
| 253555_at | 750.8 | P | 865.1 | P | 783.7 | P | 819.8 | P | --- |
| 253556_at | 9.7 | A | 13.1 | A | 2 | A | 1.1 | A | --- |
| 253557_at | 107.8 | P | 83.4 | P | 92.6 | P | 110.9 | P | --- |
| 253558_at | 809.2 | P | 779.7 | P | 789.5 | P | 748.8 | P | SKB1 |
| 253559_at | 759.3 | P | 305.7 | P | 767.8 | P | 453.1 | P | --- |
| 253560_at | 410.6 | P | 375.5 | P | 446.1 | P | 421.1 | P | DCAF1 |
| 253561_at | 2291.1 | P | 1725.3 | P | 2117.8 | P | 1804 | P | --- |
| 253562_at | 808.3 | P | 1271.4 | P | 1077.8 | P | 1351.6 | P | --- |
| 253563_at | 162.2 | P | 225.7 | P | 235.4 | P | 188.7 | P | --- |
| 253564_at | 853.5 | P | 865.5 | P | 771 | P | 932.9 | P | --- |
| 253565_at | 224 | P | 246.8 | P | 322.2 | P | 266 | P | --- |
| 253566_at | 717.7 | P | 707.1 | P | 737 | P | 692.9 | P | --- |
| 253567_at | 2.8 | A | 4.5 | A | 7 | A | 2.3 | A | --- |
| 253568_at | 1.1 | A | 0.9 | A | 2 | A | 2.1 | A | --- |
| 253569_at | 100 | M | 84 | P | 104 | A | 99.5 | P | --- |
| 253570_at | 11.6 | A | 1.3 | A | 16.1 | A | 1.9 | A | --- |
| 253571_at | 153.4 | P | 198.7 | P | 152.9 | P | 152.8 | P | --- |
| 253572_at | 152 | P | 218.9 | P | 183.6 | P | 155.6 | P | --- |
| 253573_at | 44 | P | 59.9 | P | 56.5 | P | 54.8 | P | --- |
| 253574_at | 182 | P | 178.7 | P | 124.4 | P | 115.9 | P | --- |
| 253575_at | 44.5 | P | 91.1 | P | 69.5 | P | 80.6 | P | --- |
| 253576_at | 45.8 | P | 37.8 | P | 36 | A | 43.9 | P | --- |
| 253577_at | 418.9 | P | 492.1 | P | 411.1 | P | 426.4 | P | --- |
| 253578_at | 257.3 | P | 268.2 | P | 208.7 | P | 267.7 | P | ATDGK7 |
| 253579_at | 129.9 | P | 227.1 | P | 207.6 | P | 311.9 | P | BRS1 |
| 253580_at | 97.1 | P | 114.9 | P | 75.4 | M | 140 | P | --- |
| 253581_at | 3282.7 | P | 2716.1 | P | 2519.7 | P | 2947.8 | P | --- |
| 253582_at | 336.3 | P | 127.9 | P | 264.1 | P | 153.7 | P | --- |
| 253583_at | 101.9 | P | 92.4 | M | 59 | A | 64.9 | A | --- |
| 253584_at | 74.9 | P | 99.9 | P | 122.8 | P | 97.9 | P | --- |
| 253585_at | 274.4 | P | 378.4 | P | 292.4 | P | 369.4 | P | --- |
| 253586_s_at | 40.9 | P | 67.6 | P | 35.3 | P | 67.3 | P | --- |
| 253587_at | 5.5 | A | 5.1 | A | 2.6 | A | 17.1 | A | --- |
| 253588_at | 253.2 | P | 289.6 | P | 318.5 | P | 267.4 | P | --- |
| 253589_at | 454.7 | P | 450.6 | P | 359.2 | P | 437.3 | P | --- |
| 253590_at | 105.4 | P | 132.2 | P | 135 | P | 109.7 | P | HHP2 |
| 253591_at | 25.2 | A | 33.9 | A | 16.6 | A | 41.6 | A | MUS81 |
| 253592_at | 207.3 | P | 253 | P | 253.5 | P | 298.5 | P | --- |
| 253593_at | 1912.1 | P | 1543.1 | P | 2036.3 | P | 1508 | P | --- |
| 253594_at | 129.1 | P | 111.6 | P | 160.4 | P | 105.9 | P | --- |
| 253595_at | 1.7 | A | 12.3 | A | 1.4 | A | 3.4 | A | --- |
| 253596_s_at | 378.8 | P | 301.3 | P | 383.9 | P | 273.6 | P | --- |
| 253597_at | 710.8 | P | 748.8 | P | 535 | P | 691 | P | --- |
| 253598_at | 362.3 | P | 455 | P | 592.3 | P | 429.7 | P | --- |
| 253599_at | 129.7 | P | 152.3 | P | 158.8 | P | 165.5 | P | SDG4 |
| 253600_at | 845.9 | P | 1494.4 | P | 958.6 | P | 1420.6 | P | scpl29 |
| 253601_at | 328 | P | 411.6 | P | 292.3 | P | 299.5 | P | --- |
| 253602_s_at | 1839.1 | P | 2396.5 | P | 2161.6 | P | 2380.1 | P | --- |
| 253603_at | 275.9 | P | 258.6 | P | 342.2 | P | 253.2 | P | WRKY32 |
| 253604_at | 7.1 | A | 0.5 | A | 0.9 | A | 1.4 | A | --- |
| 253605_at | 199.1 | P | 131.3 | P | 197.4 | P | 150 | P | --- |
| 253606_at | 7611.4 | P | 4076 | P | 8034 | P | 4624 | P | --- |
| 253607_at | 580.1 | P | 669.8 | P | 417.2 | P | 723.8 | P | --- |
| 253608_at | 459.8 | P | 434.5 | P | 355.8 | P | 312.6 | P | XTH19 |
| 253609_at | 3952.1 | P | 4316.1 | P | 4415.3 | P | 4249.2 | P | AHA2 |
| 253610_at | 1336.2 | P | 1305.3 | P | 1608.6 | P | 1298.9 | P | VEL1 |
| 253611_at | 4.5 | A | 2 | A | 1.3 | A | 2.3 | A | NAP15 |
| 253612_at | 1284.1 | P | 1010.3 | P | 1131.5 | P | 1045.9 | P | --- |
| 253613_at | 9.7 | A | 36.9 | A | 15.9 | A | 7 | A | --- |
| 253614_at | 643.4 | P | 540.6 | P | 438.5 | P | 485.7 | P | --- |
| 253615_at | 113.8 | P | 189.3 | P | 151.7 | P | 157.4 | P | CNGC17 |
| 253616_at | 4 | A | 2.1 | A | 1.7 | A | 1.5 | A | EXLB2 |

| | | | | | | | | | |
|---|---|---|---|---|---|---|---|---|---|
| 253617_at | 88.9 | A | 109.9 | P | 72.8 | P | 141.5 | P | --- |
| 253618_at | 29 | P | 50.4 | P | 50.4 | P | 41 | P | --- |
| 253619_at | 193.4 | P | 173.4 | P | 287.2 | P | 173.8 | P | --- |
| 253620_at | 190.4 | P | 242.2 | P | 270.6 | P | 252.8 | P | --- |
| 253621_at | 12.9 | A | 3 | A | 3.5 | A | 2.5 | A | --- |
| 253622_at | 126.6 | P | 146.2 | P | 144.1 | P | 116.7 | P | ATCNGC9 |
| 253623_at | 73.4 | P | 58.9 | P | 72 | P | 77.3 | P | --- |
| 253624_at | 1032.5 | P | 1051.8 | P | 1053.1 | P | 847.7 | P | ATS2 |
| 253625_at | 3416.6 | P | 2154.3 | P | 2321.2 | P | 2309.6 | P | --- |
| 253626_at | 7 | A | 23.2 | A | 3.6 | A | 2 | A | --- |
| 253627_at | 2882.2 | P | 4327.5 | P | 2763.6 | P | 5692.8 | P | --- |
| 253628_at | 579.9 | P | 238.7 | P | 330.8 | P | 171.3 | P | XTH18 |
| 253629_at | 327.6 | P | 259.5 | P | 494.4 | P | 327.6 | P | --- |
| 253630_at | 769.8 | P | 863.7 | P | 790.3 | P | 768.4 | P | --- |
| 253631_at | 1582 | P | 2035.4 | P | 1470.1 | P | 1920.5 | P | GAE1 |
| 253632_at | 64.1 | P | 72.6 | P | 74.6 | A | 67.3 | P | TET9 |
| 253633_at | 493.9 | P | 511.9 | P | 447.7 | P | 517.1 | P | --- |
| 253634_at | 16.9 | A | 42.1 | M | 56.2 | A | 53.7 | A | --- |
| 253635_at | 449.1 | P | 708.8 | P | 659.5 | P | 674.3 | P | --- |
| 253636_at | 473.8 | P | 463.3 | P | 482.9 | P | 475.9 | P | --- |
| 253637_at | 407.5 | P | 373.9 | P | 380.2 | P | 326 | P | --- |
| 253638_at | 1090.6 | P | 737.6 | P | 1164.1 | P | 1207.3 | P | --- |
| 253639_at | 770.9 | P | 1191.1 | P | 672.8 | P | 1063.5 | P | --- |
| 253640_at | 368.1 | P | 306.2 | P | 248.3 | P | 326.7 | P | --- |
| 253641_at | 15.7 | A | 2.4 | A | 31.7 | A | 23 | A | --- |
| 253642_at | 414 | P | 492.3 | P | 543.5 | P | 486.6 | P | --- |
| 253643_at | 172.9 | P | 188.2 | P | 213.7 | P | 169.4 | P | --- |
| 253644_at | 596 | P | 797.7 | P | 691.3 | P | 736 | P | --- |
| 253645_at | 451 | P | 523.7 | P | 472 | P | 593.3 | P | VIP3 |
| 253646_at | 924.9 | P | 1171.9 | P | 945.5 | P | 1145.2 | P | ATMKK2 |
| 253647_at | 666 | P | 496.9 | P | 493 | P | 494.1 | P | --- |
| 253648_at | 122.9 | P | 72.7 | P | 59.5 | P | 75.6 | P | PRHA |
| 253649_at | 160.5 | P | 210.8 | P | 105.3 | P | 166.7 | P | --- |
| 253650_at | 519.8 | P | 853.3 | P | 505.4 | P | 1039.7 | P | --- |
| 253651_at | 15.9 | A | 2 | A | 17.9 | A | 3.5 | A | --- |
| 253652_at | 31.7 | A | 9.9 | A | 31.9 | A | 34 | A | --- |
| 253653_at | 14 | A | 12.6 | A | 3.4 | A | 23.6 | P | --- |
| 253654_at | 81.6 | A | 127.9 | P | 90.6 | A | 133.6 | P | --- |
| 253655_at | 5.9 | A | 2.2 | A | 16 | A | 2.9 | A | LCR59 |
| 253656_at | 30.7 | A | 7.3 | A | 32.1 | A | 26 | A | emb1353 |
| 253657_at | 5.2 | A | 52.8 | A | 18.1 | A | 46.2 | A | HMA2 |
| 253658_at | 11.4 | A | 9.8 | A | 6.8 | A | 5.6 | A | HMA3 |
| 253659_at | 172.3 | P | 139.8 | P | 209.6 | P | 125.5 | P | --- |
| 253660_at | 744.5 | P | 479.5 | P | 708.4 | P | 499 | P | --- |
| 253661_at | 4.8 | A | 22.5 | A | 3.4 | A | 4.3 | A | --- |
| 253662_at | 315.7 | P | 316.4 | P | 260.8 | P | 290.8 | P | ARF16 |
| 253663_at | 276.7 | P | 294.2 | P | 276.3 | P | 262.3 | P | VLN4 |
| 253664_at | 1751.2 | P | 1154.4 | P | 1316.4 | P | 1287.9 | P | ATR2 |
| 253665_at | 41.7 | A | 42.6 | A | 11.4 | A | 33.6 | A | --- |
| 253666_at | 9370.5 | P | 7480.1 | P | 8055.3 | P | 7753.1 | P | MERI5B |
| 253667_at | 3.1 | A | 69.2 | P | 32.6 | A | 55.2 | P | --- |
| 253668_at | 1634.4 | P | 1625.3 | P | 1714.8 | P | 1623.7 | P | RUXF |
| 253669_at | 94.2 | P | 85.6 | P | 145.4 | P | 118.6 | P | --- |
| 253670_at | 4673.9 | P | 4762.3 | P | 4674.9 | P | 4315.1 | P | --- |
| 253671_at | 27.1 | P | 33.8 | A | 33.4 | A | 37.3 | A | --- |
| 253672_at | 346.6 | P | 277.8 | P | 280.7 | P | 240.1 | P | CFIM-25 |
| 253673_at | 132.7 | P | 145.4 | P | 144.3 | P | 118 | P | --- |
| 253674_at | 56.7 | A | 66.5 | P | 79.9 | P | 54.7 | A | --- |
| 253675_at | 118.7 | P | 127 | P | 135 | P | 100.9 | P | --- |
| 253676_at | 30.3 | A | 36.3 | A | 12.8 | A | 36.6 | A | --- |
| 253677_at | 58.8 | A | 21.5 | A | 99.7 | M | 26.1 | A | --- |
| 253678_at | 40.9 | P | 39.7 | P | 32.2 | M | 11.8 | A | --- |
| 253679_at | 273.8 | P | 237.2 | P | 169.7 | P | 230.3 | P | --- |
| 253680_at | 5 | A | 3.5 | A | 5.8 | A | 3.8 | A | --- |
| 253681_at | 0.8 | A | 6.7 | A | 1.1 | A | 2.4 | A | --- |
| 253682_at | 17.5 | P | 21.8 | M | 6.4 | A | 26.1 | A | --- |
| 253683_at | 42.8 | A | 34.9 | A | 66.2 | A | 17.5 | A | --- |
| 253684_at | 56 | P | 8.7 | A | 69.9 | P | 25.3 | A | --- |
| 253685_at | 1.6 | A | 14.3 | A | 16.5 | A | 0.6 | A | --- |
| 253686_at | 478 | P | 571.2 | P | 396.9 | P | 521.4 | P | --- |
| 253687_at | 671.2 | P | 651.7 | P | 459.2 | P | 732.6 | P | --- |
| 253688_at | 642.8 | P | 818.3 | P | 651 | P | 651.5 | P | --- |
| 253689_at | 29.5 | A | 19.5 | A | 45.8 | A | 24.8 | A | --- |
| 253690_at | 25.6 | A | 20.9 | A | 18.8 | A | 31.3 | A | --- |
| 253691_at | 450.8 | P | 416.1 | P | 391 | P | 357.1 | P | EMB2752 |
| 253692_at | 287.5 | P | 458.4 | P | 403.5 | P | 405.4 | P | ATPAO5 |
| 253693_at | 1785.3 | P | 1927 | P | 1714.4 | P | 2016.2 | P | ACHT2 |
| 253694_at | 1650.4 | P | 1875.1 | P | 1594.5 | P | 1924 | P | --- |
| 253695_at | 1139.1 | P | 851.4 | P | 982 | P | 920 | P | PRMT11 |
| 253696_at | 123 | P | 113.3 | P | 152 | P | 105.9 | P | CKX4 |
| 253697_at | 397.9 | P | 215.9 | P | 541 | P | 169 | P | --- |
| 253698_at | 907.7 | P | 861.5 | P | 976.4 | P | 908.5 | P | --- |
| 253699_at | 1.1 | A | 22.4 | A | 49.6 | A | 6.9 | A | PLP8 |
| 253700_at | 3004.5 | P | 2276.1 | P | 2505.7 | P | 2650.7 | P | MTO2 |
| 253701_at | 405.6 | P | 423.3 | P | 414.5 | P | 376.2 | P | --- |
| 253702_at | 1338.3 | P | 1283.6 | P | 1032.1 | P | 1015.5 | P | ACA10 |
| 253703_at | 150.9 | P | 213.4 | P | 163.4 | P | 155.8 | P | ORC5 |
| 253704_at | 1023.9 | P | 984.2 | P | 1029.1 | P | 1045.8 | P | --- |
| 253705_at | 1940.4 | P | 2086.5 | P | 2263.3 | P | 2050.8 | P | HXK1 |
| 253706_at | 795.3 | P | 937.6 | P | 999.1 | P | 936.1 | P | --- |
| 253707_at | 24.1 | A | 30.3 | A | 28.7 | A | 21.2 | A | --- |
| 253708_at | 806.3 | P | 508.5 | P | 763.2 | P | 543.2 | P | GGT4 |
| 253709_at | 1349.8 | P | 1343.7 | P | 1453.2 | P | 1335.1 | P | PFK1 |
| 253710_at | 18.8 | A | 8.7 | A | 27.7 | A | 9.1 | A | anac075 |
| 253711_at | 3.6 | A | 13.5 | A | 14.5 | A | 9.6 | A | DER1 |
| 253712_at | 463.4 | P | 565.8 | P | 386.5 | P | 540.6 | P | DER1 |
| 253713_at | 56.3 | A | 32.3 | P | 39.6 | M | 45.5 | A | --- |
| 253714_at | 170.4 | P | 205.2 | P | 222.8 | P | 184.6 | P | --- |
| 253715_at | 4694.7 | P | 4350 | P | 5164.6 | P | 4648.1 | P | --- |
| 253716_at | 119.5 | P | 105 | P | 159.8 | P | 91.9 | P | --- |
| 253717_at | 104.4 | P | 144 | P | 80.4 | P | 109.8 | P | --- |
| 253718_at | 3.7 | A | 3.8 | A | 27.5 | A | 7.8 | A | --- |
| 253719_at | 524.5 | P | 539.4 | P | 372.6 | P | 604.6 | P | --- |
| 253720_at | 60.4 | P | 203.5 | P | 146.7 | P | 257.5 | P | --- |
| 253721_at | 11.6 | A | 15.4 | A | 11.3 | A | 20.2 | A | --- |
| 253722_at | 390.2 | P | 540.1 | P | 431.2 | P | 491.5 | P | --- |
| 253723_at | 73.1 | P | 100 | P | 85.4 | P | 105.3 | P | --- |
| 253724_at | 12.2 | A | 7.2 | A | 22.8 | A | 19.3 | A | LCR24 |
| 253725_at | 47.8 | P | 5.4 | A | 26.7 | A | 17.8 | A | PRF4 |

| | | | | | | | | | |
|---|---|---|---|---|---|---|---|---|---|
| 253726_at | 240.4 | P | 217.8 | P | 206.1 | P | 207.4 | P | rps15ae |
| 253727_at | 8509.6 | P | 8072.5 | P | 7125.9 | P | 8515 | P | PFN2 |
| 253728_at | 4039 | P | 2683.4 | P | 3727.4 | P | 2988.3 | P | --- |
| 253729_at | 72.1 | P | 182 | P | 166.6 | P | 180.7 | P | --- |
| 253730_at | 3643.7 | P | 3361.2 | P | 2996.5 | P | 3899.7 | P | --- |
| 253731_at | 455.7 | P | 500.4 | P | 477.9 | P | 503.5 | P | --- |
| 253732_at | 129.3 | P | 140 | P | 127.2 | P | 142.7 | P | --- |
| 253733_at | 108.7 | P | 74.2 | P | 89.9 | P | 92 | P | ATMND1 |
| 253734_at | 31.6 | A | 40.8 | A | 14.7 | A | 27.8 | A | --- |
| 253735_at | 895.2 | P | 769.4 | P | 706.8 | P | 658.2 | P | SNF7.1 |
| 253736_at | 283.7 | P | 335.3 | P | 361.2 | P | 389.6 | P | --- |
| 253737_at | 47 | A | 20.5 | A | 22 | A | 48.1 | A | --- |
| 253738_at | 15403.7 | P | 14120 | P | 14859.9 | P | 14064 | P | PSAE-1 |
| 253739_at | 556.9 | P | 476.2 | P | 613.7 | P | 504.9 | P | --- |
| 253740_at | 733.3 | P | 890.7 | P | 649.5 | P | 892.6 | P | --- |
| 253741_at | 2.6 | A | 16.5 | A | 19.9 | A | 22.8 | A | --- |
| 253742_at | 45.4 | A | 44.2 | A | 11.2 | A | 37.3 | A | --- |
| 253743_at | 1.6 | A | 52.9 | A | 88.6 | A | 75.9 | A | --- |
| 253744_s_at | 5.2 | A | 0.4 | A | 6.7 | A | 5.8 | A | --- |
| 253745_at | 1.7 | A | 3.5 | A | 3.5 | A | 8.7 | A | --- |
| 253746_at | 354.5 | P | 396.2 | P | 468 | P | 409 | P | --- |
| 253747_at | 71.9 | P | 76 | P | 74.2 | A | 55.9 | P | --- |
| 253748_at | 185.6 | P | 203.1 | P | 279.8 | P | 198.5 | P | --- |
| 253749_at | 452.9 | P | 483.7 | P | 787.8 | P | 644 | P | PAP2 |
| 253750_at | 103.6 | P | 110 | P | 143.5 | P | 138 | P | --- |
| 253751_at | 1157.1 | P | 1280.8 | P | 1116.4 | P | 1156.8 | P | --- |
| 253752_at | 1046.4 | P | 887.2 | P | 1134.9 | P | 882.1 | P | --- |
| 253753_at | 1119.5 | P | 1275 | P | 788.7 | P | 1253.3 | P | --- |
| 253754_at | 3339.9 | P | 5795.6 | P | 5396.9 | P | 6452.7 | P | --- |
| 253755_at | 2270.3 | P | 2233.8 | P | 1857.4 | P | 2103.9 | P | RPT2a |
| 253756_at | 315.3 | P | 367.2 | P | 271.8 | P | 333 | P | CAK1AT |
| 253757_at | 90.6 | P | 99.1 | P | 117.4 | P | 124.3 | P | ROP9 |
| 253758_at | 4079.1 | P | 4042 | P | 4520.9 | P | 4290.3 | P | emb2726 |
| 253759_at | 4155.1 | P | 3405.3 | P | 3655 | P | 3504.7 | P | AIM1 |
| 253760_at | 15.8 | A | 23.6 | A | 8.2 | A | 3.4 | A | AT4G28815 |
| 253761_at | 101 | P | 76.1 | P | 135.6 | P | 83.3 | P | --- |
| 253762_at | 276 | P | 313.2 | P | 334.4 | P | 289.5 | P | --- |
| 253763_at | 2.4 | A | 2.6 | A | 4.8 | A | 4 | A | --- |
| 253764_s_at | 379.3 | P | 353.9 | P | 293.7 | P | 376.5 | P | ckl4 |
| 253765_at | 1345.6 | P | 1436.7 | P | 1402.5 | P | 1439.1 | P | --- |
| 253766_at | 1033.5 | P | 1154.3 | P | 933.9 | P | 1058.1 | P | --- |
| 253767_at | 26.5 | M | 38.3 | P | 59 | A | 34.5 | A | CRU3 |
| 253768_at | 78.5 | A | 81 | A | 45.7 | A | 61.3 | A | --- |
| 253769_at | 46.2 | A | 27.1 | A | 29.5 | A | 35.4 | A | RIC7 |
| 253770_at | 256.5 | P | 314.6 | P | 266.9 | P | 274.9 | P | CKL6 |
| 253771_at | 16 | A | 79 | P | 42.9 | A | 61.1 | P | --- |
| 253772_at | 23.8 | A | 1.5 | A | 2.4 | A | 1.9 | A | A7 |
| 253773_s_at | 1683 | P | 1366.4 | P | 1592.2 | P | 1649.7 | P | --- |
| 253774_at | 41.1 | A | 48.2 | P | 89.5 | P | 82.3 | P | anac074 |
| 253775_at | 1539.4 | P | 1518.6 | P | 1825.7 | P | 1816.9 | P | --- |
| 253776_at | 775.3 | P | 713.4 | P | 637.3 | P | 569.9 | P | AAC3 |
| 253777_at | 653.7 | P | 449.8 | P | 810.5 | P | 569 | P | --- |
| 253778_at | 726.4 | P | 684.9 | P | 540 | P | 688.6 | P | --- |
| 253779_at | 261 | P | 242.4 | P | 218.3 | P | 248.5 | P | HAE |
| 253780_at | 538.6 | P | 670.8 | P | 464.9 | P | 658.2 | P | --- |
| 253781_at | 3.1 | A | 3.6 | A | 28 | A | 31.8 | A | --- |
| 253782_at | 238.1 | P | 258.5 | P | 248.7 | P | 253.7 | P | --- |
| 253783_at | 808.8 | P | 652.4 | P | 793.4 | P | 712.5 | P | NPGR2 |
| 253784_at | 247.2 | P | 259.7 | P | 257.4 | P | 236.5 | P | PHR1 |
| 253785_s_at | 69.9 | P | 60.3 | P | 94 | M | 67.3 | P | ATM2 |
| 253786_at | 39.1 | A | 46.9 | A | 47.5 | A | 46.3 | A | --- |
| 253787_at | 18.2 | A | 2.7 | A | 5.7 | A | 2.5 | A | --- |
| 253788_at | 39.9 | A | 127 | P | 41.2 | A | 83.9 | P | --- |
| 253789_at | 506 | P | 517.8 | P | 467.2 | P | 468.3 | P | --- |
| 253790_at | 2603.6 | P | 3076.8 | P | 3230.2 | P | 3108.8 | P | PSB28 |
| 253791_at | 281.7 | P | 258.5 | P | 313.3 | P | 259.3 | P | IAA11 |
| 253792_at | 9.2 | A | 4.2 | A | 8.9 | A | 5.2 | A | AMT1;4 |
| 253793_at | 69.1 | P | 112.1 | P | 93.6 | P | 86.2 | P | XIH |
| 253794_at | 61.6 | P | 11.6 | A | 60.6 | A | 36.4 | A | --- |
| 253795_at | 17.9 | A | 10.8 | A | 8.7 | A | 2.8 | A | --- |
| 253796_at | 34.2 | P | 28.4 | A | 45.1 | P | 35.4 | A | --- |
| 253797_at | 285.7 | P | 285.7 | P | 350.4 | P | 277.4 | P | RPN1B |
| 253798_at | 13.3 | A | 21.5 | A | 14 | A | 40.8 | A | ANAC073 |
| 253799_at | 39.1 | M | 52.7 | P | 72.2 | P | 62.1 | P | --- |
| 253800_at | 104.3 | P | 154.9 | P | 67.4 | A | 165.5 | P | --- |
| 253801_at | 26.2 | A | 64.3 | A | 59.1 | A | 18.7 | A | --- |
| 253802_at | 55.1 | A | 46.4 | A | 69 | A | 53.3 | A | --- |
| 253803_at | 756.8 | P | 626.1 | P | 764.6 | P | 620.2 | P | --- |
| 253804_at | 96.1 | P | 119 | P | 77.5 | P | 123.2 | P | --- |
| 253805_at | 379.4 | P | 429.2 | P | 336.3 | P | 475.1 | P | --- |
| 253806_at | 373 | P | 454 | P | 376.9 | P | 385.8 | P | --- |
| 253807_at | 6.6 | A | 26.2 | A | 7.5 | A | 16.2 | A | --- |
| 253808_at | 473.7 | P | 535 | P | 403.2 | P | 411.5 | P | --- |
| 253809_at | 52.4 | P | 108.3 | P | 37 | A | 102.1 | P | --- |
| 253810_at | 407 | P | 379 | P | 486.4 | P | 390 | P | NDB1 |
| 253811_at | 129.8 | P | 202.8 | P | 228 | P | 165.9 | P | ULT1 |
| 253812_at | 528.2 | P | 741.6 | P | 450.5 | P | 505.8 | P | --- |
| 253813_at | 89.5 | P | 87.9 | P | 113.8 | P | 83.8 | P | --- |
| 253814_at | 151 | P | 202.1 | P | 198.8 | P | 230.3 | P | --- |
| 253815_at | 239.2 | P | 598.2 | P | 269.3 | P | 647.8 | P | ATEXPB3 |
| 253816_at | 578.4 | P | 772.4 | P | 756.9 | P | 788.9 | P | emb1923 |
| 253817_at | 250.4 | P | 453.2 | P | 371.5 | P | 472.1 | P | --- |
| 253818_at | 131.3 | P | 87 | P | 120.8 | P | 103 | P | --- |
| 253819_at | 28.8 | P | 35.2 | P | 41.2 | A | 50.8 | P | --- |
| 253820_at | 214.8 | P | 267.1 | P | 176.4 | P | 267.7 | P | --- |
| 253821_at | 4.3 | A | 2.5 | A | 2.8 | A | 1.5 | A | --- |
| 253822_at | 16.9 | A | 2.9 | A | 3.6 | A | 4.1 | A | --- |
| 253823_at | 1295.5 | P | 1282.3 | P | 1290.3 | P | 1232.3 | P | --- |
| 253824_at | 351.3 | P | 277.6 | P | 302 | P | 312.8 | P | MTM1 |
| 253825_at | 1642.5 | P | 1696.9 | P | 1790.2 | P | 1597 | P | --- |
| 253826_s_at | 11653.8 | P | 11147.9 | P | 11087.6 | P | 12076.8 | P | UBC9 |
| 253827_at | 96.1 | P | 55.1 | P | 82.3 | P | 31.3 | P | --- |
| 253828_at | 102.9 | P | 94.5 | P | 122.7 | P | 58.6 | P | SLAH2 |
| 253829_at | 639.7 | P | 119.7 | P | 398.9 | P | 152.3 | P | --- |
| 253830_at | 289.6 | P | 232.1 | P | 256.8 | P | 230.1 | P | --- |
| 253831_at | 13.7 | A | 5.8 | A | 7.9 | A | 7.6 | A | --- |
| 253832_at | 52 | P | 18.8 | P | 60 | P | 32.2 | A | --- |
| 253833_at | 32.9 | A | 34.5 | A | 56.5 | A | 27.5 | A | --- |
| 253834_at | 2778.4 | P | 3163.6 | P | 2680 | P | 3096.1 | P | --- |

| | | | | | | | | | |
|---|---|---|---|---|---|---|---|---|---|
| 253835_at | 579.6 | P | 466.1 | P | 610.3 | P | 581.4 | P | BGLU9 |
| 253836_at | 259.2 | P | 205.7 | P | 191.7 | P | 253.8 | P | --- |
| 253837_at | 43.7 | A | 40.8 | P | 83.6 | A | 23.6 | A | --- |
| 253838_at | 385.5 | P | 481 | P | 389.6 | P | 345.2 | P | --- |
| 253839_at | 39.2 | A | 29.8 | A | 66.1 | P | 17.2 | A | --- |
| 253840_at | 178.4 | P | 241.4 | P | 229.1 | P | 198.3 | P | ACBP2 |
| 253841_at | 2936.1 | P | 1915.6 | P | 2217.3 | P | 2081.1 | P | BGLU10 |
| 253842_at | 1059.8 | P | 323.9 | P | 950.1 | P | 430.1 | P | --- |
| 253843_at | 162.8 | P | 146.9 | P | 120.3 | P | 144.6 | P | SDG16 |
| 253844_at | 3.8 | A | 10 | A | 32.1 | A | 7.7 | A | --- |
| 253845_at | 48.5 | A | 43.8 | A | 76.1 | A | 30.5 | A | --- |
| 253846_at | 28.9 | A | 7 | A | 45.5 | A | 7.8 | A | --- |
| 253847_at | 48.2 | P | 67.4 | P | 64.8 | P | 59.9 | P | --- |
| 253848_at | 163.8 | P | 122.5 | P | 147.1 | P | 153 | P | --- |
| 253849_at | 1349.6 | P | 1723.1 | P | 1619.2 | P | 1732.6 | P | --- |
| 253850_at | 2.2 | A | 46.5 | A | 23 | A | 44.9 | A | sks10 |
| 253851_at | 29.3 | A | 47 | M | 58.2 | A | 29.9 | A | AtMYB41 |
| 253852_at | 20.7 | A | 4 | A | 27.3 | A | 9.6 | A | --- |
| 253853_at | 20 | P | 1.5 | A | 2.1 | A | 7.4 | A | --- |
| 253854_at | 475.8 | P | 777.4 | P | 547.4 | P | 767 | P | --- |
| 253855_at | 543.5 | P | 750.1 | P | 639.2 | P | 837.1 | P | TET7 |
| 253856_at | 398.7 | P | 526 | P | 451.3 | P | 442.7 | P | --- |
| 253857_at | 1640.3 | P | 1624.7 | P | 1467.9 | P | 1703.3 | P | --- |
| 253858_at | 1529.9 | P | 1690.7 | P | 1564.2 | P | 1717.9 | P | --- |
| 253859_at | 51.8 | P | 56.9 | P | 87.9 | P | 66.2 | P | --- |
| 253860_at | 1188.1 | P | 1906.9 | P | 1405.7 | P | 2047.1 | P | --- |
| 253861_at | 678 | P | 659.4 | P | 658.9 | P | 744.5 | P | --- |
| 253862_at | 3.6 | A | 5.1 | A | 7.7 | A | 17.5 | A | SPL |
| 253863_at | 38.4 | A | 42.5 | A | 21 | A | 30.6 | A | --- |
| 253864_at | 83.5 | P | 17.4 | A | 34.3 | M | 49.9 | P | --- |
| 253865_at | 263.1 | P | 397.3 | P | 247.5 | P | 297.2 | P | --- |
| 253866_at | 100.3 | P | 101.8 | P | 90 | P | 85.4 | P | --- |
| 253867_at | 194.3 | P | 172.5 | P | 242.7 | P | 224.4 | P | --- |
| 253868_at | 3643.4 | P | 3058.7 | P | 2721.8 | P | 2864.1 | P | PPI1 |
| 253869_at | 57.3 | P | 72.3 | P | 71.4 | P | 57.7 | P | --- |
| 253870_at | 5 | A | 14.2 | A | 35.7 | A | 5.4 | A | --- |
| 253871_at | 3510.7 | P | 5826.6 | P | 4128.9 | P | 5054.6 | P | PORB |
| 253872_at | 378.8 | P | 210.2 | P | 229.8 | P | 209.5 | P | RD26 |
| 253873_at | 344.9 | P | 253.9 | P | 276.9 | P | 263.3 | P | PRA1.H |
| 253874_at | 96.2 | P | 283.4 | P | 131.3 | P | 163.4 | P | --- |
| 253875_at | 1975.2 | P | 2234.7 | P | 2186.8 | P | 2701.8 | P | --- |
| 253876_at | 448.3 | P | 487.1 | P | 453.2 | P | 536.4 | P | CIP7 |
| 253877_at | 130.2 | P | 175.5 | P | 177.7 | P | 159.7 | P | --- |
| 253878_at | 12.6 | A | 16.7 | A | 13.2 | A | 24.3 | A | TPS4 |
| 253879_s_at | 2381.3 | P | 1647.5 | P | 2281.6 | P | 1968.9 | P | --- |
| 253880_at | 33.2 | A | 42.3 | P | 22.4 | A | 25.7 | A | --- |
| 253881_at | 585.8 | P | 517.2 | P | 956.8 | P | 621.5 | P | --- |
| 253882_at | 469.2 | P | 512.1 | P | 489.2 | P | 491 | P | PEL1 |
| 253883_at | 75.5 | P | 73 | P | 77.3 | P | 94.2 | P | --- |
| 253884_at | 17.1 | A | 2.2 | A | 3 | A | 6.3 | A | HSP21 |
| 253885_at | 437.3 | P | 470.4 | P | 374.4 | P | 407.5 | P | VPS26B |
| 253886_at | 380.3 | P | 523.5 | P | 340.6 | P | 384.1 | P | CYP709B3 |
| 253887_at | 92.8 | A | 152.6 | P | 55.9 | A | 85.5 | P | OPT6 |
| 253888_at | 641.5 | P | 617.2 | P | 552.5 | P | 563.3 | P | ISI1 |
| 253889_at | 410.4 | P | 653.5 | P | 493.5 | P | 602.3 | P | FEY |
| 253890_s_at | 911.7 | P | 753.4 | P | 783.2 | P | 753.1 | P | --- |
| 253891_at | 568.6 | P | 1348 | P | 683.3 | P | 1140.7 | P | --- |
| 253892_at | 243.3 | P | 222.7 | P | 249.8 | P | 212.5 | P | --- |
| 253893_at | 380.2 | P | 453.4 | P | 306.4 | P | 374.4 | P | --- |
| 253894_at | 5.3 | A | 14 | P | 25 | P | 5.8 | A | --- |
| 253895_at | 40.8 | A | 3.4 | A | 3.5 | A | 10.1 | A | AT2S3 |
| 253896_at | 1.5 | A | 1.1 | A | 2.7 | A | 1.3 | A | --- |
| 253897_at | 408.5 | P | 461.7 | P | 417.4 | P | 396.8 | P | --- |
| 253898_s_at | 4848.5 | P | 2614.4 | P | 3818.2 | P | 2574.1 | P | TSB2 |
| 253899_at | 678 | P | 852.5 | P | 682 | P | 885.9 | P | ATPDIL5-4 |
| 253900_at | 2283.7 | P | 2035.4 | P | 1759.1 | P | 2061.3 | P | --- |
| 253901_at | 9055.5 | P | 8696.1 | P | 9833.3 | P | 8963.8 | P | --- |
| 253902_at | 4.9 | A | 10.9 | A | 19.2 | A | 4.3 | A | --- |
| 253903_at | 471.9 | P | 582.8 | P | 487.3 | P | 573.6 | P | ATK2 |
| 253904_at | 79.2 | P | 46.8 | A | 63.8 | P | 33.7 | A | --- |
| 253905_s_at | 3.6 | A | 14.4 | A | 23 | A | 4.2 | A | --- |
| 253906_at | 27.2 | A | 2.8 | A | 20.6 | A | 15.7 | A | --- |
| 253907_at | 35.3 | A | 42 | A | 70.3 | P | 27.7 | A | --- |
| 253908_at | 134.8 | P | 300.1 | P | 180.6 | P | 296.1 | P | WES1 |
| 253909_at | 207.9 | P | 317.2 | P | 235.4 | P | 266.4 | P | --- |
| 253910_at | 2.1 | A | 11.7 | A | 4 | A | 3.1 | A | --- |
| 253911_at | 761.5 | P | 865.8 | P | 652.8 | P | 742.8 | P | --- |
| 253912_at | 389.2 | P | 373.6 | P | 575.8 | P | 400.3 | P | --- |
| 253913_at | 22.7 | A | 42.6 | P | 49.1 | P | 50.3 | P | VIIIB |
| 253914_at | 31.4 | A | 47.3 | M | 36.3 | A | 21.7 | A | --- |
| 253915_at | 893.9 | P | 736 | P | 607.3 | P | 634.3 | P | --- |
| 253916_at | 52.4 | P | 99.7 | P | 64.4 | P | 70.3 | P | --- |
| 253917_at | 469.1 | P | 474.4 | P | 467.4 | P | 499.1 | P | --- |
| 253918_at | 867.7 | P | 872.7 | P | 624.5 | P | 691.2 | P | PHOS34 |
| 253919_at | 154.1 | P | 169.1 | P | 155.8 | P | 179.9 | P | --- |
| 253920_at | 653.8 | P | 1130.9 | P | 742.5 | P | 1098.6 | P | HTA2 |
| 253921_at | 786.8 | P | 735.1 | P | 996.4 | P | 717.8 | P | AT-HF |
| 253922_at | 1856.6 | P | 3380.5 | P | 2520.1 | P | 3502.8 | P | VTC2 |
| 253923_at | 413.3 | P | 440.7 | P | 381.6 | P | 380.1 | P | TOR1 |
| 253924_at | 7.4 | A | 12.9 | A | 25.6 | A | 30.9 | A | COBL11 |
| 253925_at | 639.5 | P | 698 | P | 722.2 | P | 705.7 | P | SHV3 |
| 253926_at | 484.3 | P | 630.6 | P | 609.9 | P | 593.5 | P | --- |
| 253927_at | 6884.9 | P | 5656.8 | P | 6129 | P | 5192.9 | P | --- |
| 253928_at | 550.1 | P | 529 | P | 511.6 | P | 477.2 | P | PPX1 |
| 253929_s_at | 12.4 | A | 20.3 | A | 43 | A | 38.4 | A | --- |
| 253930_at | 4.2 | A | 15.1 | A | 5.6 | A | 5.2 | A | ATS1 |
| 253931_at | 3.3 | A | 16.4 | A | 1.8 | A | 3.2 | A | --- |
| 253932_at | 1.5 | A | 2.1 | A | 1.1 | A | 2 | A | --- |
| 253933_at | 4.9 | A | 37.8 | A | 37.9 | A | 7.3 | A | --- |
| 253934_at | 26.1 | A | 23.9 | P | 39.5 | P | 50.6 | P | --- |
| 253935_at | 1171.3 | P | 1181.9 | P | 1340.6 | P | 1193.3 | P | --- |
| 253936_at | 13.1 | A | 23.5 | A | 39.6 | A | 3.5 | A | --- |
| 253937_at | 24.4 | A | 29.4 | A | 17.3 | A | 27.4 | A | MAPKKK16 |
| 253938_at | 19.6 | A | 44.3 | A | 34.4 | A | 37.8 | A | --- |
| 253939_at | 24.5 | A | 2 | A | 22.5 | A | 13 | A | MYB97 |
| 253940_at | 181.7 | P | 174.2 | P | 202.2 | P | 143.4 | P | --- |
| 253941_at | 205.6 | P | 243.1 | P | 199.9 | P | 224.3 | P | --- |
| 253942_at | 371.5 | P | 190.8 | P | 345.3 | P | 234.9 | P | --- |
| 253943_at | 323.3 | P | 457.4 | P | 561.8 | P | 557.2 | P | --- |

| | | | | | | | | | |
|---|---|---|---|---|---|---|---|---|---|
| 253944_at | 851.6 | P | 832.2 | P | 680.1 | P | 781 | P | VPS22 |
| 253945_at | 207.1 | P | 173.1 | P | 243.7 | P | 230.3 | P | --- |
| 253946_at | 158.7 | P | 172.7 | P | 182.4 | P | 129 | P | --- |
| 253947_at | 164 | P | 346.4 | P | 201.2 | P | 354.2 | P | MAP65-2 |
| 253948_at | 1196.6 | P | 1255.9 | P | 1031.9 | P | 1279.2 | P | --- |
| 253949_at | 446.9 | P | 401.4 | P | 532.7 | P | 451.2 | P | AR192 |
| 253950_at | 2053.1 | P | 2035.1 | P | 1910.9 | P | 1941.7 | P | --- |
| 253951_at | 476.8 | P | 711.1 | P | 537.7 | P | 694.7 | P | --- |
| 253952_at | 1872 | P | 2075.7 | P | 1996.5 | P | 2329.4 | P | SUMO1 |
| 253953_at | 89.2 | P | 120.7 | P | 148.7 | P | 137 | P | --- |
| 253954_at | 3503.5 | P | 2340.1 | P | 3022.2 | P | 2601.6 | P | --- |
| 253955_at | 3496.4 | P | 2697.1 | P | 3868.6 | P | 2653.1 | P | ATRBP45C |
| 253956_at | 732 | P | 977.7 | P | 829.7 | P | 891.3 | P | FIM1 |
| 253957_at | 2.2 | A | 12.3 | A | 34.6 | A | 21 | A | AGP13 |
| 253958_at | 713.3 | P | 744.3 | P | 754 | P | 623.9 | P | --- |
| 253959_at | 1682.7 | P | 1776.2 | P | 1805.2 | P | 1756.5 | P | --- |
| 253960_at | 38 | A | 29.6 | A | 33.1 | A | 49.8 | A | GAMT1 |
| 253961_at | 1.8 | A | 1.3 | A | 1.5 | A | 2.5 | A | WRKY34 |
| 253962_at | 24.3 | P | 26.8 | P | 26.1 | A | 22.6 | P | --- |
| 253963_at | 77.5 | P | 88.8 | P | 55.5 | A | 47.5 | A | --- |
| 253964_at | 899.5 | P | 852.8 | P | 1164 | P | 716.4 | P | --- |
| 253965_at | 85.3 | P | 76.5 | P | 130.9 | P | 96.5 | M | --- |
| 253966_at | 215.6 | P | 387.1 | P | 315.4 | P | 344.1 | P | --- |
| 253967_at | 262 | P | 276.9 | P | 215.2 | P | 250.7 | P | --- |
| 253968_at | 35.2 | A | 22.1 | A | 39.2 | A | 17 | A | CBL7 |
| 253969_at | 344.9 | P | 365.6 | P | 304.6 | P | 345.4 | P | CSN6B |
| 253970_at | 862 | P | 1171.5 | P | 708.6 | P | 1007.8 | P | --- |
| 253971_at | 1572.8 | P | 2873.7 | P | 2104.3 | P | 2318.9 | P | --- |
| 253972_at | 1294.4 | P | 1030.6 | P | 1508.7 | P | 1070.4 | P | CPSUFE |
| 253973_at | 382 | P | 565.7 | P | 372.5 | P | 471.8 | P | --- |
| 253974_at | 7.3 | A | 33.1 | A | 71.2 | A | 41.8 | A | --- |
| 253975_at | 268.4 | P | 249.6 | P | 324.9 | P | 216.3 | P | --- |
| 253976_at | 346 | P | 410.2 | P | 393.4 | P | 314 | P | D6PKL1 |
| 253977_at | 1239.6 | P | 1242.5 | P | 1387.7 | P | 1184.9 | P | --- |
| 253978_at | 61.4 | M | 83.6 | P | 44 | A | 96.3 | P | --- |
| 253979_at | 21.8 | A | 63.4 | A | 70.1 | A | 47 | A | --- |
| 253980_at | 470.9 | P | 683.8 | P | 511 | P | 674.3 | P | --- |
| 253981_at | 1427.3 | P | 1528.1 | P | 1428.2 | P | 1452.9 | P | --- |
| 253982_at | 625.3 | P | 725 | P | 535.9 | P | 710.7 | P | ATCBL3 |
| 253983_at | 375.4 | P | 436.5 | P | 518.9 | P | 440.2 | P | WRKY20 |
| 253984_at | 22.3 | A | 50.5 | A | 56 | A | 41.6 | A | OPT5 |
| 253985_at | 4.7 | A | 5.8 | A | 15.5 | A | 4.9 | A | --- |
| 253986_at | 1017.6 | P | 913.6 | P | 1096 | P | 1117.4 | P | --- |
| 253987_at | 380.8 | P | 465.2 | P | 248.4 | P | 471 | P | PFK3 |
| 253988_at | 496.6 | P | 505.3 | P | 501.8 | P | 458.7 | P | --- |
| 253989_at | 669.6 | P | 795.3 | P | 668.2 | P | 770.4 | P | --- |
| 253990_at | 179.8 | P | 184.1 | P | 147.7 | P | 190.6 | P | ACHT1 |
| 253991_at | 424 | P | 371.7 | P | 399.3 | P | 378.1 | P | PEP |
| 253992_at | 371.5 | P | 308.1 | P | 204 | P | 288.4 | P | --- |
| 253993_at | 359.6 | P | 570.6 | P | 310.9 | P | 397.6 | P | MEK1 |
| 253994_at | 1155.4 | P | 1173.2 | P | 718.4 | P | 1356.4 | P | ABI1 |
| 253995_at | 533.8 | P | 544.7 | P | 641.5 | P | 505.3 | P | CK1 |
| 253996_at | 1418.4 | P | 1042 | P | 1723.8 | P | 928 | P | NAP1;1 |
| 253997_at | 80.8 | A | 145.6 | P | 73.3 | P | 98 | P | RPS2 |
| 253998_at | 8.1 | A | 31 | A | 1.6 | A | 1.6 | A | --- |
| 253999_at | 54.3 | A | 54.7 | P | 62.1 | P | 86.5 | P | ACS7 |
| 254000_at | 11.1 | A | 2.3 | A | 2 | A | 2.2 | A | AtGolS6 |
| 254001_at | 22.3 | A | 22.5 | A | 15.4 | A | 10.8 | A | MIOX4 |
| 254002_at | 32.1 | A | 42.8 | A | 29.2 | A | 23.5 | A | --- |
| 254003_at | 5.9 | A | 6 | A | 6.3 | A | 0.4 | A | --- |
| 254004_at | 1080.9 | P | 1074.5 | P | 1210.3 | P | 1135.4 | P | emb1027 |
| 254005_at | 26.9 | A | 34.6 | A | 24.9 | A | 43.7 | A | UNE17 |
| 254006_at | 2.9 | A | 4.8 | A | 34.7 | A | 20.4 | A | --- |
| 254007_s_at | 11.1 | A | 14.3 | A | 37.1 | A | 13 | A | --- |
| 254008_at | 21.1 | A | 1.7 | A | 19.1 | A | 4.4 | A | --- |
| 254009_at | 9.4 | A | 9.8 | A | 22.1 | A | 9.3 | A | --- |
| 254010_at | 345.2 | P | 384.1 | P | 306.2 | P | 316.5 | P | --- |
| 254011_at | 937.5 | P | 1147.4 | P | 1089 | P | 1169.7 | P | --- |
| 254012_at | 6224.3 | P | 5795.7 | P | 6550.7 | P | 6190.3 | P | --- |
| 254013_at | 6 | A | 9.8 | A | 13.4 | A | 8 | A | --- |
| 254014_at | 80.2 | P | 19.8 | A | 65 | P | 40.7 | P | --- |
| 254015_at | 47.7 | P | 53.9 | P | 97.8 | P | 82.3 | P | BGAL12 |
| 254016_at | 282.2 | P | 288.1 | P | 365 | P | 251.4 | P | CGA1 |
| 254017_at | 1.3 | A | 12.6 | A | 1.4 | A | 10 | A | --- |
| 254018_at | 110.4 | P | 131 | P | 153.3 | A | 92.1 | A | --- |
| 254019_at | 144.5 | P | 178.9 | P | 292.9 | P | 210 | P | --- |
| 254020_at | 1300.7 | P | 1520.4 | P | 2693.9 | P | 1905.6 | P | BETA-OHASE 1 |
| 254021_at | 2456.2 | P | 2701 | P | 2298.1 | P | 2637.4 | P | ACD1-LIKE |
| 254022_at | 31 | P | 60.3 | P | 48.7 | A | 62.4 | P | --- |
| 254023_at | 224.5 | P | 223.6 | P | 205 | A | 243.2 | P | --- |
| 254024_at | 138.7 | P | 122.1 | P | 179.8 | P | 99.5 | P | --- |
| 254025_at | 6.5 | A | 3.1 | A | 29.1 | A | 19.3 | A | --- |
| 254026_at | 17.6 | A | 30.5 | A | 50 | A | 45 | A | --- |
| 254027_at | 74 | P | 61.2 | P | 66.1 | P | 67.3 | P | --- |
| 254028_s_at | 25.2 | A | 17.6 | A | 23.3 | A | 15.3 | A | ORP4B |
| 254029_at | 344.6 | P | 424.8 | P | 332.3 | P | 443.5 | P | --- |
| 254030_at | 1221.8 | P | 910 | P | 1911.3 | P | 1150.6 | P | --- |
| 254031_at | 38.3 | M | 29 | P | 43.1 | P | 5.5 | A | --- |
| 254032_at | 70.1 | P | 68.7 | P | 128.8 | P | 84.2 | P | --- |
| 254033_at | 3.9 | A | 38.4 | A | 1 | A | 11.1 | A | VATG3 |
| 254034_at | 590.8 | P | 718.1 | P | 528.1 | P | 760.5 | P | PGP2 |
| 254035_at | 1128.1 | P | 1234.6 | P | 902.6 | P | 1248.2 | P | PSD3 |
| 254036_at | 19 | A | 50.4 | P | 49.5 | A | 28.3 | A | --- |
| 254037_at | 54.9 | A | 83.5 | A | 49.8 | A | 70.5 | A | ATGDU2 |
| 254038_at | 927.3 | P | 1019.4 | P | 1117 | P | 1161.9 | P | NFU3 |
| 254039_at | 376.2 | P | 471.6 | P | 542.9 | P | 470.5 | P | GPP1 |
| 254040_at | 1163.3 | P | 1294.4 | P | 1026.3 | P | 1167.1 | P | --- |
| 254041_at | 175.4 | P | 292.7 | P | 170.6 | P | 341.9 | P | --- |
| 254042_at | 428.9 | P | 150.3 | P | 275.6 | P | 106.2 | P | XTR6 |
| 254043_at | 66.2 | A | 71.6 | P | 80.6 | P | 65.8 | A | CIL |
| 254044_at | 20.4 | A | 23.6 | A | 19.3 | A | 1.3 | A | XTR9 |
| 254045_at | 733.1 | P | 915 | P | 802.3 | P | 1034.1 | P | APUM6 |
| 254046_at | 5.3 | A | 0.2 | A | 0.4 | A | 0.7 | A | --- |
| 254047_at | 1.7 | A | 6.9 | A | 4.8 | A | 12 | A | --- |
| 254048_at | 306.4 | P | 296.6 | P | 270.4 | P | 260.7 | P | --- |
| 254049_at | 2870.5 | P | 2821.7 | P | 2883.2 | P | 2989.2 | P | --- |
| 254050_s_at | 1180 | P | 1316.8 | P | 1122.8 | P | 1211 | P | --- |
| 254051_at | 97.2 | P | 101.1 | P | 123.5 | P | 150.8 | P | --- |
| 254052_at | 226.8 | P | 311.6 | P | 188.7 | P | 340.8 | P | --- |

| | | | | | | | | | |
|---|---|---|---|---|---|---|---|---|---|
| 254053_s_at | 125.5 | P | 137.6 | P | 118.6 | P | 135.5 | P | --- |
| 254054_at | 341.2 | P | 319.3 | P | 362.8 | P | 292.1 | P | --- |
| 254055_at | 3.1 | A | 4.5 | A | 23.4 | A | 29.8 | A | --- |
| 254056_at | 10.9 | A | 5.8 | A | 42.3 | A | 6.8 | A | --- |
| 254057_at | 426 | P | 755.2 | P | 601.3 | P | 642.3 | P | --- |
| 254058_at | 1336.5 | P | 840.1 | P | 1006.6 | P | 1036 | P | --- |
| 254059_at | 19.6 | A | 58.2 | A | 58.4 | A | 37.8 | A | ATHSP23.6-MITO |
| 254060_at | 13.9 | A | 48.2 | A | 11.1 | A | 32.9 | A | SHB1 |
| 254061_at | 274.7 | P | 373.3 | P | 297.8 | P | 374.5 | P | --- |
| 254062_at | 36.9 | M | 20 | A | 53.9 | P | 38.3 | A | --- |
| 254063_at | 209.1 | P | 209.4 | M | 132 | A | 169.3 | P | --- |
| 254064_at | 1.2 | A | 17.7 | A | 28 | A | 11.8 | A | --- |
| 254065_at | 60.3 | P | 50.8 | M | 66.7 | A | 42.2 | A | GA20OX1 |
| 254066_at | 46.7 | P | 56.7 | P | 24.3 | A | 61.3 | P | DREB1A |
| 254067_at | 470.6 | P | 331.5 | P | 425.5 | P | 315.2 | P | NAP8 |
| 254068_at | 1212.2 | P | 683.8 | P | 1561.3 | P | 978.8 | P | NAP8 |
| 254069_at | 5.4 | A | 20.7 | A | 19.7 | A | 15 | A | ATNUDT10 |
| 254070_at | 4.7 | A | 10.8 | A | 2.9 | A | 7.7 | A | --- |
| 254071_at | 1 | A | 8.5 | A | 6 | A | 7.4 | A | --- |
| 254072_at | 2300.9 | P | 3165.2 | P | 2956.5 | P | 3039.5 | P | --- |
| 254073_at | 480 | P | 520.4 | P | 420.5 | P | 569.7 | P | RSP35 |
| 254074_at | 10.9 | A | 8.6 | A | 3.1 | A | 14.6 | A | CBF1 |
| 254075_at | 33.2 | A | 56.6 | P | 68.6 | A | 44.2 | A | CBF2 |
| 254076_at | 469.4 | P | 465.3 | P | 420.3 | P | 430 | P | --- |
| 254077_at | 2339.2 | P | 2179.5 | P | 2309.1 | P | 2438.6 | P | --- |
| 254078_at | 401.3 | P | 353.6 | P | 540.8 | P | 376.1 | P | --- |
| 254079_at | 557.4 | P | 405 | P | 637.9 | P | 493.4 | P | --- |
| 254080_at | 1744.2 | P | 1281 | P | 2246.5 | P | 1624.9 | P | FIB2 |
| 254081_at | 606.2 | P | 784.5 | P | 426.6 | P | 506.9 | P | --- |
| 254082_at | 772.9 | P | 691.8 | P | 777.3 | P | 755 | P | QC |
| 254083_at | 3203.1 | P | 3960.4 | P | 2394.9 | P | 3726.8 | P | --- |
| 254084_at | 771.9 | P | 869.3 | P | 971.6 | P | 849.2 | P | SAE1A |
| 254085_at | 710.5 | P | 1234 | P | 759 | P | 1056.7 | P | HVA22D |
| 254086_at | 2236.8 | P | 2526.3 | P | 2211.9 | P | 2560.3 | P | --- |
| 254087_at | 592 | P | 751.9 | P | 527.8 | P | 697.5 | P | --- |
| 254088_at | 216.9 | P | 205.9 | P | 243.2 | P | 181 | P | --- |
| 254089_at | 394.1 | P | 506.1 | P | 512.8 | P | 425.4 | P | --- |
| 254090_at | 17.7 | A | 42.5 | P | 53.8 | A | 42.1 | A | --- |
| 254091_at | 28.6 | A | 59.3 | A | 8.7 | A | 34 | A | --- |
| 254092_at | 2 | A | 1.3 | A | 4 | A | 3.8 | A | --- |
| 254093_at | 41.4 | A | 55.3 | A | 13.2 | A | 50.1 | A | AtMC2 |
| 254094_at | 97.1 | P | 116.3 | P | 133.6 | P | 113.2 | P | --- |
| 254095_at | 20.5 | P | 24.7 | A | 35.7 | A | 22.1 | A | OLEO1 |
| 254096_at | 45.3 | P | 106.3 | P | 59.3 | P | 70.3 | P | --- |
| 254097_at | 1.4 | A | 4.5 | A | 3.1 | A | 14.8 | P | --- |
| 254098_at | 17007 | P | 15533.9 | P | 18159.4 | P | 15790.4 | P | FSD1 |
| 254099_at | 3840.8 | P | 4714.9 | P | 4411.4 | P | 5002.3 | P | --- |
| 254100_at | 160.4 | A | 62.9 | A | 122 | A | 57.6 | A | --- |
| 254101_at | 336.9 | P | 191.9 | P | 85.6 | P | 168.2 | P | AMY1 |
| 254102_at | 5407 | P | 6633.9 | P | 4764.2 | P | 5792.5 | P | ACP4 |
| 254103_at | 1576.4 | P | 1068.9 | P | 1428.3 | P | 1217.7 | P | --- |
| 254104_at | 48.2 | A | 20.4 | A | 53.4 | A | 38.5 | A | --- |
| 254105_at | 3053 | P | 3698.8 | P | 3341.4 | P | 3614.6 | P | CHLM |
| 254106_at | 932.6 | P | 831.4 | P | 688.5 | P | 840.5 | P | ATGP4 |
| 254107_at | 31.7 | A | 29.1 | A | 44.8 | A | 22.8 | A | --- |
| 254108_at | 1369.5 | P | 995.4 | P | 1152.8 | P | 1147.1 | P | RIN2 |
| 254109_at | 52.1 | P | 126.9 | P | 42.5 | P | 105.6 | P | SKS1 |
| 254110_at | 221.4 | P | 424.3 | P | 355 | P | 341.3 | P | --- |
| 254111_at | 44.5 | A | 31.1 | A | 51.7 | A | 32.8 | P | PAP24 |
| 254112_at | 76.3 | P | 83.2 | P | 65.3 | P | 107.2 | P | --- |
| 254113_at | 124.7 | P | 165.1 | P | 123.5 | A | 191.3 | P | --- |
| 254114_at | 25.6 | A | 23.5 | A | 21.4 | A | 24.6 | A | --- |
| 254115_at | 40.2 | A | 59.8 | A | 51.9 | A | 80.9 | A | --- |
| 254116_at | 395.9 | P | 453.2 | P | 414 | P | 509.8 | P | --- |
| 254117_at | 1290.9 | P | 1574.1 | P | 1298.6 | P | 1331.2 | P | --- |
| 254118_at | 65.9 | P | 118.5 | P | 125 | P | 138.7 | P | --- |
| 254119_at | 756.4 | P | 1972.5 | P | 1174.8 | P | 1949.5 | P | --- |
| 254120_at | 296.2 | P | 306 | P | 264.1 | P | 239.3 | P | DIC2 |
| 254121_at | 464 | P | 310.9 | P | 595.4 | P | 407.7 | P | --- |
| 254122_at | 496.3 | P | 728.2 | P | 563.8 | P | 607.2 | P | CER2 |
| 254123_at | 41.6 | A | 37.9 | A | 37.3 | M | 25.8 | A | APPB1 |
| 254124_at | 190.6 | P | 215.7 | P | 175 | P | 200.3 | P | --- |
| 254125_at | 49.1 | P | 136 | P | 65.6 | P | 115.1 | P | TAR2 |
| 254126_at | 5924.9 | P | 5936.6 | P | 5498.6 | P | 5914.1 | P | RBP31 |
| 254127_at | 1428 | P | 1476.4 | P | 1354.2 | P | 1400.1 | P | ATR1 |
| 254128_at | 354 | P | 410.7 | P | 368.6 | P | 432.3 | P | UBP16 |
| 254129_at | 18.5 | A | 8.3 | A | 9.3 | A | 7.9 | A | IPT4 |
| 254130_at | 55.5 | P | 168.1 | P | 94.4 | P | 166.7 | P | AGL24 |
| 254131_at | 711.8 | P | 721.9 | P | 656.2 | P | 738.5 | P | AFC2 |
| 254132_at | 252.5 | P | 208.3 | P | 283.7 | P | 255.8 | P | ATHB22 |
| 254133_at | 323.7 | P | 344.2 | P | 287.3 | P | 295.6 | P | --- |
| 254134_at | 1985.4 | P | 1507.7 | P | 2306.3 | P | 1469.8 | P | --- |
| 254135_at | 38.2 | P | 36.4 | A | 51.3 | A | 29.3 | A | --- |
| 254136_at | 4.6 | A | 22.1 | A | 19.1 | A | 26.6 | A | --- |
| 254137_at | 1472.1 | P | 1416.9 | P | 1695.4 | P | 1518.8 | P | --- |
| 254138_at | 1 | A | 2.1 | A | 1.7 | A | 2.4 | A | --- |
| 254139_at | 14 | A | 7.4 | A | 18.2 | A | 1.8 | A | --- |
| 254140_at | 134.8 | P | 255.7 | P | 197.7 | P | 288.8 | P | --- |
| 254141_at | 1668.9 | P | 1852.1 | P | 1756.6 | P | 2084.5 | P | PGI1 |
| 254142_at | 103.8 | P | 114.3 | P | 175.2 | M | 117.6 | P | --- |
| 254143_at | 216.3 | P | 218.9 | P | 190.2 | P | 162.3 | P | --- |
| 254144_at | 4232.4 | P | 3681 | P | 2525.4 | P | 3376.5 | P | --- |
| 254145_at | 203.4 | P | 236 | P | 312.7 | P | 306.5 | P | --- |
| 254146_at | 160.6 | P | 124.3 | P | 181.8 | P | 148.7 | P | ATGH9A3 |
| 254147_at | 213.8 | P | 201.7 | P | 300.4 | P | 208 | P | --- |
| 254148_at | 3333.5 | P | 2998 | P | 3630 | P | 2860 | P | cpHsc70-1 |
| 254149_at | 3.8 | A | 15.8 | A | 3.5 | A | 27.7 | A | --- |
| 254150_at | 1157.2 | P | 600 | P | 1004.4 | P | 756.2 | P | --- |
| 254151_at | 241.6 | P | 219.2 | P | 192.3 | P | 288.6 | P | --- |
| 254152_at | 1 | A | 1.6 | A | 1.1 | A | 0.6 | A | --- |
| 254153_at | 316.6 | P | 459.5 | P | 308.5 | P | 436.3 | P | PWD |
| 254154_at | 437.8 | P | 446.7 | P | 393.2 | P | 469.2 | P | --- |
| 254155_at | 102.5 | P | 95.7 | P | 75.8 | A | 74.6 | A | --- |
| 254156_at | 357.6 | P | 351.4 | P | 397.1 | P | 358.8 | P | --- |
| 254157_at | 5689.8 | P | 4652.2 | P | 4279.4 | P | 4812 | P | VEP1 |
| 254158_at | 1278.6 | P | 654.3 | P | 846.2 | P | 663.4 | P | --- |
| 254159_at | 156.2 | P | 309.9 | P | 189.8 | P | 283.3 | P | WRKY7 |
| 254160_at | 446.5 | P | 460.1 | P | 418 | P | 409.8 | P | SLY1 |
| 254161_at | 240 | P | 280.7 | P | 197.3 | P | 243.2 | P | --- |

| | | | | | | | | | |
|---|---|---|---|---|---|---|---|---|---|
| 254162_at | 828.6 | P | 739.5 | P | 671.5 | P | 738.6 | P | --- |
| 254163_s_at | 2249.7 | P | 867.2 | P | 2038.7 | P | 1244.4 | P | --- |
| 254164_at | 608.5 | P | 659.1 | P | 639.9 | P | 692.6 | P | TIFY1 |
| 254165_at | 1288.3 | P | 1270.2 | P | 1214.2 | P | 1262.2 | P | --- |
| 254166_at | 6252.8 | P | 7486.3 | P | 5675.9 | P | 8052.4 | P | SHD |
| 254167_at | 606.8 | P | 655.5 | P | 551.9 | P | 612.7 | P | CIPK8 |
| 254168_at | 39.2 | A | 24 | A | 32.5 | A | 28.5 | A | MLO13 |
| 254169_at | 316.5 | P | 345.5 | P | 328.6 | P | 415.7 | P | --- |
| 254170_at | 21.7 | A | 35.7 | P | 75.4 | P | 26.2 | A | --- |
| 254171_at | 188.7 | P | 230 | P | 188.3 | P | 215.5 | P | --- |
| 254172_at | 1549.8 | P | 1363.6 | P | 1457.8 | P | 1406.9 | P | --- |
| 254173_at | 5 | A | 3.6 | A | 2.5 | A | 2.2 | A | REN1 |
| 254174_at | 903.4 | P | 934 | P | 853.7 | P | 1062.4 | P | YSL1 |
| 254175_at | 41.8 | P | 73.9 | A | 90.2 | A | 79.8 | P | --- |
| 254176_at | 213.7 | P | 225.1 | P | 220.7 | P | 188.6 | P | --- |
| 254177_at | 132.1 | P | 150.9 | P | 178.5 | P | 142.3 | P | --- |
| 254178_at | 81.6 | P | 107 | P | 72.3 | P | 111.3 | P | --- |
| 254179_at | 426.1 | P | 349.4 | P | 385.1 | P | 364 | P | --- |
| 254180_at | 93.1 | P | 76.2 | P | 105.4 | P | 113 | P | --- |
| 254181_at | 436.3 | P | 407.3 | P | 423.3 | P | 458.3 | P | --- |
| 254182_at | 150.3 | P | 154.2 | P | 128.7 | P | 181.7 | P | --- |
| 254183_at | 2 | A | 15.7 | A | 34 | A | 2.3 | A | --- |
| 254184_at | 26.8 | A | 3.4 | A | 52.4 | A | 33.1 | A | --- |
| 254185_at | 285.5 | P | 311.2 | P | 415.5 | P | 290.6 | P | CSLG3 |
| 254186_at | 40.2 | A | 37.3 | A | 46.5 | A | 66.4 | P | CSLG1 |
| 254187_at | 2164.9 | P | 2247.4 | P | 2634.1 | P | 1882 | P | --- |
| 254188_at | 392.5 | P | 313.6 | P | 409.9 | P | 268.5 | P | UGE2 |
| 254189_at | 966.4 | P | 291.6 | P | 560.1 | P | 355 | P | CSLG2 |
| 254190_at | 333.5 | P | 366.4 | P | 208.1 | P | 299.2 | P | --- |
| 254191_at | 265.4 | P | 415.6 | P | 253 | P | 472.4 | P | --- |
| 254192_at | 3895.1 | P | 4320.8 | P | 3096.2 | P | 3878.3 | P | --- |
| 254193_at | 88.5 | P | 107.2 | P | 63.1 | P | 73.5 | P | --- |
| 254194_at | 229.4 | P | 247.8 | P | 181 | P | 227.2 | P | ARF9 |
| 254195_at | 441.7 | P | 373.6 | P | 321.6 | P | 339.2 | P | NLP7 |
| 254196_at | 11.3 | A | 6.2 | A | 49 | A | 26.6 | A | --- |
| 254197_at | 232.9 | P | 154.7 | P | 113.8 | P | 145.6 | P | TRE1 |
| 254198_at | 428.1 | P | 578.9 | P | 415.5 | P | 532.7 | P | --- |
| 254199_at | 80 | P | 49.3 | P | 69 | A | 72.9 | A | --- |
| 254200_at | 156.7 | P | 48.7 | P | 94.2 | P | 52.5 | P | --- |
| 254201_at | 644.4 | P | 281.4 | P | 488.1 | P | 298.6 | P | --- |
| 254202_at | 32.6 | A | 94.4 | P | 44.9 | A | 50.2 | P | --- |
| 254203_at | 15.8 | A | 2.5 | A | 47.1 | A | 24.4 | A | AtGRF8 |
| 254204_at | 718.4 | P | 785.6 | P | 606.5 | P | 856.9 | P | --- |
| 254205_at | 16.3 | A | 27.5 | A | 3.3 | A | 18.4 | A | --- |
| 254206_at | 4.3 | A | 39.4 | A | 38.5 | A | 6.6 | A | --- |
| 254207_at | 163.2 | P | 213.2 | P | 166 | P | 202.4 | P | --- |
| 254208_at | 265.2 | P | 336.4 | P | 278.7 | P | 337.8 | P | --- |
| 254209_at | 178.6 | P | 134.3 | P | 159.6 | P | 146.2 | P | --- |
| 254210_at | 50.2 | P | 61.7 | P | 45.5 | A | 29.4 | A | --- |
| 254211_at | 698 | P | 1149.2 | P | 706.1 | P | 902.5 | P | SGT1A |
| 254212_at | 10.8 | A | 14.6 | A | 39.3 | A | 28.8 | A | --- |
| 254213_at | 48.7 | A | 5.3 | A | 47.6 | A | 36.1 | A | --- |
| 254214_at | 754.5 | P | 826.7 | P | 698.5 | P | 818.9 | P | TRH1 |
| 254215_at | 5.7 | A | 27.1 | A | 29.2 | A | 42.8 | A | ATCHX17 |
| 254216_at | 792.9 | P | 721.3 | P | 667.3 | P | 819.2 | P | VAG2 |
| 254217_at | 61.1 | A | 49.9 | P | 69.2 | P | 75.9 | P | --- |
| 254218_at | 301.5 | P | 386.6 | P | 337.1 | P | 448.3 | P | --- |
| 254219_at | 49.2 | P | 4.9 | A | 24.4 | A | 8.6 | A | --- |
| 254220_at | 13.2 | A | 4.2 | A | 5.7 | A | 7.9 | A | --- |
| 254221_at | 232 | P | 1287.9 | P | 301.4 | P | 996.1 | P | --- |
| 254222_at | 38.4 | A | 52.6 | M | 54.6 | A | 52.6 | A | --- |
| 254223_at | 351.6 | P | 451 | P | 391.6 | P | 465.1 | P | --- |
| 254224_at | 1428.5 | P | 1795.3 | P | 1405.8 | P | 1580.3 | P | CDPK6 |
| 254225_at | 9894.4 | P | 8027.4 | P | 9614.4 | P | 8606.5 | P | --- |
| 254226_at | 64 | P | 139.1 | P | 137.3 | P | 204.3 | P | --- |
| 254227_at | 4666.5 | P | 5477.3 | P | 3931.6 | P | 5772.7 | P | BTI1 |
| 254228_at | 377 | P | 475.7 | P | 563.5 | P | 508.2 | P | --- |
| 254229_at | 37.3 | P | 105.6 | P | 50.6 | P | 62.4 | P | --- |
| 254230_at | 171.5 | P | 197.4 | P | 206.1 | P | 208.2 | P | AtPPT1 |
| 254231_at | 34 | P | 364.1 | P | 68.2 | P | 222.3 | P | WRKY53 |
| 254232_at | 17960.8 | P | 5461.8 | P | 13899.1 | P | 8684.8 | P | CORI3 |
| 254233_at | 375.3 | P | 732.5 | P | 442.5 | P | 767.7 | P | --- |
| 254234_at | 8200.1 | P | 4136.3 | P | 6956.8 | P | 4608.2 | P | --- |
| 254235_at | 87.2 | P | 86 | P | 100.5 | P | 73.7 | A | CRF2 |
| 254236_at | 93.9 | P | 74.4 | P | 95.3 | P | 107.7 | P | --- |
| 254237_at | 24.3 | A | 3.5 | A | 24.9 | A | 6 | A | --- |
| 254238_at | 150.1 | P | 114.9 | P | 148.4 | P | 115.6 | P | --- |
| 254239_at | 808.3 | P | 1820.7 | P | 888.7 | P | 1705.5 | P | PIP1;5 |
| 254240_at | 12.9 | A | 29 | A | 10.5 | A | 58.8 | A | SP1L5 |
| 254241_at | 157.9 | P | 170.6 | P | 93.4 | P | 129 | P | CRK11 |
| 254242_at | 351.4 | P | 334.6 | P | 157.1 | P | 237 | P | --- |
| 254243_at | 195.4 | P | 237.2 | P | 115.8 | P | 112 | P | CRK13 |
| 254244_at | 1 | A | 49.3 | P | 9.1 | A | 12.3 | A | --- |
| 254245_at | 72.2 | P | 57.4 | P | 44.2 | P | 57.2 | P | --- |
| 254246_at | 44.4 | A | 52.7 | P | 59 | M | 46.7 | P | EMB1290 |
| 254247_at | 665 | P | 606.4 | P | 578.1 | P | 593.7 | P | --- |
| 254248_at | 457 | P | 421.4 | P | 358.8 | P | 395 | P | --- |
| 254249_at | 25.3 | P | 56.8 | P | 18.1 | A | 27.7 | P | --- |
| 254250_at | 292.7 | P | 397.1 | P | 362.8 | P | 419.1 | P | --- |
| 254251_at | 233.5 | P | 310.2 | P | 301.3 | P | 307.1 | P | --- |
| 254252_at | 2.9 | A | 46.6 | A | 3.8 | A | 21.6 | A | --- |
| 254253_at | 23.6 | A | 60.2 | A | 39 | A | 45.7 | A | --- |
| 254254_at | 65.9 | P | 104.4 | P | 85.7 | A | 76.1 | P | --- |
| 254255_at | 79.1 | P | 296.2 | P | 78.2 | P | 190.3 | P | --- |
| 254256_at | 1067.3 | P | 1240.6 | P | 759.6 | P | 998.8 | P | CRK10 |
| 254257_s_at | 16.3 | A | 19.4 | A | 10.1 | A | 24.1 | A | --- |
| 254258_at | 77 | P | 88.4 | P | 72.9 | P | 86.8 | P | TET5 |
| 254259_s_at | 603.3 | P | 794.4 | P | 717.6 | P | 889.1 | P | --- |
| 254260_at | 145.8 | P | 169.5 | P | 158.8 | P | 171.2 | P | --- |
| 254261_at | 468.4 | P | 569.5 | P | 376.7 | P | 461.8 | P | --- |
| 254262_at | 1413.4 | P | 1334.6 | P | 1149.6 | P | 1087.7 | P | --- |
| 254263_at | 44.2 | A | 58.5 | A | 14.7 | A | 66 | A | --- |
| 254264_at | 17 | A | 23 | A | 22.2 | A | 20.9 | M | --- |
| 254265_s_at | 55.2 | P | 898 | P | 48.6 | P | 689 | P | CRK6 |
| 254266_at | 65.1 | A | 189.1 | P | 66.3 | P | 113.5 | P | CRK5 |
| 254267_at | 634.3 | P | 699.4 | P | 663.6 | P | 648.1 | P | PYR4 |
| 254268_at | 168.3 | P | 218 | P | 215 | P | 224.4 | P | --- |
| 254269_at | 628.5 | P | 512.1 | P | 765.7 | P | 632.7 | P | --- |
| 254270_at | 11661.6 | P | 7078 | P | 12524.5 | P | 7635.3 | P | GSH1 |

| | | | | | | | | | |
|---|---|---|---|---|---|---|---|---|---|
| 254271_at | 35.4 | A | 182.5 | P | 46 | A | 111.1 | P | --- |
| 254272_at | 130 | P | 183.5 | P | 162.5 | P | 154.8 | P | --- |
| 254273_at | 594.4 | P | 510.5 | P | 598.2 | P | 579.4 | P | --- |
| 254274_at | 189.1 | P | 190.3 | P | 188.6 | P | 158.6 | P | --- |
| 254275_at | 2262.3 | P | 2604.4 | P | 2089.1 | P | 2536.5 | P | AtHip1 |
| 254276_at | 1113.6 | P | 897.4 | P | 846.4 | P | 989.5 | P | --- |
| 254277_at | 8.8 | A | 7.7 | A | 4.8 | A | 11.3 | A | MYB85 |
| 254278_at | 949 | P | 1097.6 | P | 795.5 | P | 1085.1 | P | --- |
| 254279_at | 729.6 | P | 857.9 | P | 872.2 | P | 810.8 | P | --- |
| 254280_at | 325.8 | P | 396.7 | P | 396.7 | P | 448 | P | SMO1-2 |
| 254281_at | 190.2 | P | 284.8 | P | 151.1 | P | 232.9 | P | --- |
| 254282_at | 47.8 | P | 80.3 | P | 99.4 | P | 85 | P | --- |
| 254283_s_at | 2182 | P | 380 | P | 3081.4 | P | 1046 | P | --- |
| 254284_at | 197.9 | P | 273 | P | 263.8 | P | 243.9 | P | FZR2 |
| 254285_at | 23.6 | P | 32.1 | A | 13.8 | A | 33 | P | --- |
| 254286_at | 40.9 | P | 22.1 | A | 42.4 | A | 33.5 | A | AGL19 |
| 254287_at | 45.8 | P | 40.3 | P | 72 | P | 53.7 | P | --- |
| 254288_at | 73.5 | P | 65.7 | P | 64.4 | A | 56.2 | P | AESP |
| 254289_at | 126.7 | P | 100.3 | P | 92.3 | P | 110.1 | P | --- |
| 254290_at | 68.9 | P | 94.4 | P | 69.5 | P | 91.3 | P | --- |
| 254291_at | 1860.3 | P | 1977 | P | 1222 | P | 1642.3 | P | UTR2 |
| 254292_at | 43.9 | A | 29.4 | A | 20.6 | A | 36.1 | A | --- |
| 254293_at | 419.2 | P | 314.2 | P | 411.8 | P | 472.3 | P | IQD22 |
| 254294_at | 44.9 | A | 36.7 | A | 45.8 | A | 45.3 | A | ATRBL7 |
| 254295_at | 3.7 | A | 4.3 | A | 2.1 | A | 3.2 | A | --- |
| 254296_at | 33.1 | A | 30.3 | A | 55.9 | A | 21.7 | A | --- |
| 254297_at | 1.3 | A | 1 | A | 17.1 | A | 3 | A | --- |
| 254298_at | 4777.7 | P | 5931 | P | 5656.1 | P | 6140.3 | P | PGR5-LIKE A |
| 254299_at | 935.7 | P | 695.8 | P | 683.3 | P | 588.5 | P | NYE1 |
| 254300_at | 61.9 | P | 143.1 | P | 89 | P | 125 | P | ACR7 |
| 254301_at | 111.8 | P | 113.4 | P | 149.7 | P | 108.7 | P | --- |
| 254302_at | 1.3 | A | 2 | A | 18.6 | A | 12.3 | A | --- |
| 254303_at | 446.7 | P | 413.8 | P | 468.2 | P | 419.5 | P | --- |
| 254304_at | 87.6 | P | 85 | P | 58.2 | P | 72.9 | P | --- |
| 254305_at | 375.7 | P | 366.5 | P | 375.6 | P | 394.6 | P | AKT2/3 |
| 254306_at | 854.8 | P | 984.4 | P | 854.9 | P | 1068.4 | P | ATCES1 |
| 254307_at | 8.8 | A | 5.1 | A | 17.5 | A | 2.2 | A | --- |
| 254308_at | 97.6 | P | 90.4 | P | 123 | P | 108.2 | P | --- |
| 254309_s_at | 6 | A | 22.2 | A | 11.6 | A | 3.4 | A | --- |
| 254310_at | 3.1 | A | 36 | P | 45.8 | A | 32.5 | A | --- |
| 254311_at | 45.6 | A | 42.2 | A | 67.1 | A | 20.2 | A | --- |
| 254312_s_at | 48.7 | P | 27 | P | 29.8 | A | 34 | A | --- |
| 254313_at | 34.9 | A | 54.4 | A | 61.9 | A | 37 | A | --- |
| 254314_at | 651.7 | P | 446.6 | P | 201.5 | P | 260 | P | --- |
| 254315_at | 100.2 | P | 71.2 | A | 117.5 | P | 104.6 | P | --- |
| 254316_at | 32.9 | A | 45 | A | 41 | A | 29.1 | A | --- |
| 254317_at | 65.8 | P | 32.7 | P | 37 | P | 29.1 | P | --- |
| 254318_at | 600.5 | P | 449.2 | P | 320.1 | P | 389.4 | P | --- |
| 254319_at | 102.6 | P | 141.1 | P | 103.5 | P | 146.2 | P | --- |
| 254320_at | 151.9 | P | 197.4 | P | 171.9 | P | 197 | P | --- |
| 254321_at | 299.1 | P | 262.4 | P | 256.1 | P | 251.2 | P | CPuORF27 |
| 254322_at | 28.7 | A | 26.6 | A | 62.3 | A | 27.2 | A | --- |
| 254323_at | 23.5 | A | 12.6 | A | 7.5 | A | 16.6 | A | --- |
| 254324_at | 30.5 | A | 1.8 | A | 2.9 | A | 2.9 | A | --- |
| 254325_at | 2.3 | A | 31.3 | A | 26.8 | A | 25.7 | A | --- |
| 254326_at | 23 | A | 23.8 | A | 46 | A | 25.6 | A | --- |
| 254327_at | 2161.2 | P | 2955.8 | P | 2237.8 | P | 3033.9 | P | --- |
| 254328_at | 680.1 | P | 912.9 | P | 896.6 | P | 1072.9 | P | APT3 |
| 254329_at | 259.3 | P | 398.8 | P | 248.4 | P | 321.1 | P | ORP2A |
| 254330_at | 4.7 | A | 4 | A | 3.5 | A | 3.9 | A | LBD32 |
| 254331_s_at | 5498.6 | P | 5712.4 | P | 4626.5 | P | 4793.4 | P | CYP706A2 |
| 254332_at | 160.2 | P | 127.7 | P | 82 | P | 139.5 | P | --- |
| 254333_at | 1765.8 | P | 895.4 | P | 1601.3 | P | 854 | P | SMO1-3 |
| 254334_at | 47.3 | A | 38.6 | A | 62.9 | A | 51.5 | P | --- |
| 254335_at | 1171.8 | P | 940.9 | P | 941.6 | P | 924.3 | P | IM |
| 254336_at | 20.2 | A | 2.3 | A | 15.2 | A | 22.8 | A | --- |
| 254337_at | 20.5 | A | 4.3 | A | 6.1 | A | 6.3 | A | --- |
| 254338_s_at | 0.7 | A | 2.1 | A | 0.8 | A | 0.6 | A | --- |
| 254339_at | 37.5 | A | 47.5 | A | 38.4 | A | 37.5 | A | BGLU3 |
| 254340_at | 166.7 | P | 258.2 | P | 210.7 | P | 326.1 | P | --- |
| 254341_at | 197.5 | P | 434.2 | P | 244.3 | P | 395.6 | P | SRF8 |
| 254342_at | 485 | P | 536 | P | 488.3 | P | 545.2 | P | --- |
| 254343_at | 1747.8 | P | 1554.6 | P | 1543.3 | P | 1443.8 | P | 41367 |
| 254344_at | 21.6 | A | 30.8 | P | 41.3 | P | 10.5 | A | --- |
| 254345_at | 1291.2 | P | 927.3 | P | 1250.3 | P | 980 | P | --- |
| 254346_at | 1254.2 | P | 952.3 | P | 762.1 | P | 789 | P | APG8A |
| 254347_at | 11.8 | A | 26.9 | A | 20.9 | A | 22.2 | A | WRKY31 |
| 254348_at | 790.8 | P | 954.5 | P | 789 | P | 882.8 | P | PUX3 |
| 254349_at | 116.2 | P | 117.8 | P | 123.9 | P | 150.9 | P | --- |
| 254350_at | 77.3 | P | 73.7 | M | 115.3 | P | 83.5 | M | --- |
| 254351_at | 231.6 | P | 318.5 | P | 262 | P | 342.5 | P | SOBER1 |
| 254352_at | 181.1 | P | 146.6 | P | 158.4 | P | 174 | P | --- |
| 254353_s_at | 713.2 | P | 625.3 | P | 842.9 | P | 609.9 | P | --- |
| 254354_at | 20.1 | A | 20.7 | M | 35.1 | A | 8.6 | A | --- |
| 254355_at | 669.7 | P | 487.2 | P | 778.9 | P | 533 | P | --- |
| 254356_at | 454.9 | P | 734 | P | 676 | P | 708.7 | P | --- |
| 254357_at | 2461.6 | P | 2978.2 | P | 2635.4 | P | 2918 | P | ISU1 |
| 254358_at | 3022.5 | P | 3054.7 | P | 2995.7 | P | 3390.2 | P | --- |
| 254359_at | 295.6 | P | 308.6 | P | 306.4 | P | 324.5 | P | --- |
| 254360_at | 285.9 | P | 341 | P | 291.7 | P | 325.6 | P | --- |
| 254361_at | 36.4 | P | 95.6 | P | 64.9 | P | 70.4 | P | --- |
| 254362_at | 52.9 | A | 116.2 | P | 112.3 | A | 117.5 | P | --- |
| 254363_at | 427.4 | P | 950.1 | P | 531.8 | P | 718.3 | P | sks4 |
| 254364_at | 24.8 | A | 6.1 | A | 11.1 | A | 9.3 | A | --- |
| 254365_at | 4.1 | A | 2.9 | A | 7.3 | A | 1.9 | A | --- |
| 254366_s_at | 4.3 | A | 1.8 | A | 2.3 | A | 5.6 | A | --- |
| 254367_at | 140.6 | P | 100 | P | 98.2 | P | 101.7 | P | --- |
| 254368_at | 957.1 | P | 698.7 | P | 843.2 | P | 752.2 | P | NRPB2 |
| 254369_at | 114.1 | P | 102.6 | P | 98.5 | P | 151.7 | P | --- |
| 254370_at | 983.9 | P | 1181.3 | P | 1171.2 | P | 1783.8 | P | ATML1 |
| 254371_at | 30.3 | P | 113.3 | P | 75 | P | 111.1 | P | BGLU47 |
| 254372_at | 2258.1 | P | 2365.9 | P | 1571.8 | P | 2259 | P | --- |
| 254373_at | 20.8 | A | 3.6 | A | 8.4 | A | 35.7 | A | --- |
| 254374_at | 46.8 | A | 40.2 | A | 32.9 | A | 5.1 | A | --- |
| 254375_at | 525 | P | 467.6 | P | 450.5 | P | 449.9 | P | QQT2 |
| 254376_at | 1715.2 | P | 1400.1 | P | 1496.8 | P | 1430.6 | P | TOM1 |
| 254377_at | 72.2 | P | 121.2 | P | 153.3 | P | 141.3 | P | --- |
| 254378_at | 641 | P | 676.9 | P | 575.5 | P | 573.9 | P | DER2.1 |
| 254379_at | 29.2 | A | 37.6 | P | 17.6 | A | 41.1 | P | --- |

| | | | | | | | | | |
|---|---|---|---|---|---|---|---|---|---|
| 254380_at | 150.9 | P | 63.1 | P | 144.7 | P | 130.2 | P | --- |
| 254381_at | 34.6 | A | 42.4 | P | 41.7 | P | 47.7 | P | --- |
| 254382_at | 67.5 | P | 104.5 | P | 85.7 | P | 142.9 | P | --- |
| 254383_at | 0.9 | A | 4.7 | A | 1.8 | A | 1.3 | A | --- |
| 254384_at | 502.6 | P | 1201 | P | 841.7 | P | 1004.5 | P | --- |
| 254385_s_at | 1599.1 | P | 140.5 | P | 631.1 | P | 102.6 | P | ATMSRB7 |
| 254386_at | 13105.6 | P | 17931.7 | P | 13792.9 | P | 19002.9 | P | PRXR1 |
| 254387_at | 3691.2 | P | 794.6 | P | 3732.4 | P | 731.1 | P | ATMSRB9 |
| 254388_at | 4381.3 | P | 4213.4 | P | 4785.9 | P | 4378.9 | P | MSRB2 |
| 254389_s_at | 1882.9 | P | 900.2 | P | 1551.8 | P | 1258.9 | P | --- |
| 254390_at | 101.5 | P | 111.3 | P | 142.9 | P | 135.9 | P | CPK15 |
| 254391_at | 69.7 | A | 55.4 | A | 83.7 | A | 61.2 | A | ENDO3 |
| 254392_at | 27.4 | A | 51.1 | P | 67.9 | P | 34.7 | P | ENDO5 |
| 254393_at | 1866.6 | P | 1237.3 | P | 1415.6 | P | 1348.5 | P | --- |
| 254394_at | 1.8 | A | 3.4 | A | 2.7 | A | 6.5 | A | --- |
| 254395_at | 1.4 | A | 2 | A | 1.7 | A | 0.9 | A | --- |
| 254396_at | 621 | P | 130.3 | P | 290.4 | P | 151.9 | P | --- |
| 254397_at | 33.2 | A | 15.3 | A | 9.1 | A | 2.2 | A | GA3OX3 |
| 254398_at | 6478.4 | P | 8024 | P | 6727.1 | P | 7647.5 | P | PSBQA |
| 254399_at | 2369.2 | P | 1726.1 | P | 1988.9 | P | 1993.7 | P | --- |
| 254400_at | 83.2 | A | 102.1 | P | 83.1 | A | 89.1 | A | ATK1 |
| 254401_at | 38 | A | 23.9 | A | 42.4 | A | 44 | P | --- |
| 254402_at | 62.7 | P | 131.4 | P | 112.5 | P | 90.7 | P | --- |
| 254403_at | 4.1 | A | 3 | A | 8.1 | A | 4.4 | A | --- |
| 254404_at | 3.3 | A | 2 | A | 3 | A | 5.4 | A | B70 |
| 254405_at | 284.5 | P | 256.1 | P | 303.3 | P | 247 | P | PUB8 |
| 254406_at | 3 | A | 0.7 | A | 2.3 | A | 0.7 | A | --- |
| 254407_at | 19.4 | A | 17.4 | A | 20.4 | A | 24.9 | P | --- |
| 254408_at | 63.9 | P | 47.9 | A | 31.3 | A | 34.8 | M | B120 |
| 254409_at | 68.7 | P | 102.3 | P | 57.7 | A | 52.3 | P | --- |
| 254410_at | 337.3 | P | 381.8 | P | 275.7 | P | 362 | P | --- |
| 254411_at | 2.2 | A | 10.9 | A | 3.6 | A | 5.9 | A | --- |
| 254412_at | 140.2 | P | 159.8 | P | 189.2 | P | 175.8 | P | B160 |
| 254413_at | 21.4 | A | 22.1 | A | 12.3 | A | 19 | A | ATMYB102 |
| 254414_at | 66.5 | P | 73.5 | P | 75 | P | 83.9 | P | HSA32 |
| 254415_at | 88 | P | 75.2 | A | 84.9 | A | 85.5 | P | ATSBT3.12 |
| 254416_at | 168.8 | P | 217.7 | P | 131.1 | P | 175.2 | P | ARK3 |
| 254417_at | 1467.8 | P | 1303.6 | P | 1473.2 | P | 1413 | P | ATFMN/FHY |
| 254418_at | 178.5 | P | 115.8 | P | 85.8 | P | 139.1 | P | --- |
| 254419_at | 88.2 | A | 54.9 | A | 83 | A | 81.4 | A | NDB3 |
| 254420_at | 16 | A | 12.2 | A | 45.9 | A | 39.4 | A | --- |
| 254421_at | 89.4 | A | 67.8 | A | 71.6 | A | 67 | A | VAL3 |
| 254422_at | 465.6 | P | 495.3 | P | 425 | P | 440.4 | P | VPS28-1 |
| 254423_at | 364.5 | P | 335.2 | P | 380.3 | P | 289.9 | P | LOL2 |
| 254424_at | 285.9 | P | 150.3 | P | 244.7 | P | 232.9 | P | --- |
| 254425_at | 1358.9 | P | 1320 | P | 1021.2 | P | 1230.6 | P | --- |
| 254426_at | 1.2 | A | 8.8 | A | 7.5 | A | 26.1 | A | --- |
| 254427_at | 214.9 | P | 197.8 | P | 229.9 | P | 226.2 | P | emb1417 |
| 254428_at | 1373.5 | P | 1048.4 | P | 1001.7 | P | 1137 | P | ZAC |
| 254429_at | 6137.3 | P | 6130.1 | P | 4390.3 | P | 5491.5 | P | --- |
| 254430_at | 2.8 | A | 20.4 | A | 12.7 | A | 12.9 | A | --- |
| 254431_at | 244.8 | P | 207.9 | P | 214 | P | 207.4 | P | --- |
| 254432_at | 1809.8 | P | 1260.2 | P | 691.1 | P | 1030.3 | P | --- |
| 254433_at | 2442 | P | 2226.7 | P | 2302.2 | P | 2440.9 | P | TPP2 |
| 254434_at | 578.6 | P | 345.2 | P | 455.7 | P | 380.7 | P | --- |
| 254435_at | 32.7 | A | 38.7 | A | 23.5 | A | 26.6 | A | MS5 |
| 254436_at | 3.8 | A | 2.1 | A | 1.3 | A | 2.6 | A | --- |
| 254437_s_at | 73.9 | A | 78.4 | A | 88.2 | A | 73.8 | A | --- |
| 254438_at | 1887 | P | 1553 | P | 1242.5 | P | 1448.1 | P | --- |
| 254439_s_at | 29.2 | A | 4.6 | A | 20.3 | A | 1.5 | A | ACA6 |
| 254440_at | 1.9 | A | 1.9 | A | 1 | A | 0.9 | A | --- |
| 254441_at | 1.7 | A | 1.6 | A | 1.8 | A | 7 | A | --- |
| 254442_at | 60.3 | P | 101.9 | P | 99.7 | P | 119.7 | P | --- |
| 254443_at | 92.1 | A | 96.4 | P | 93.8 | A | 114.3 | P | BRCA1 |
| 254444_at | 646.1 | P | 777.6 | P | 752.8 | P | 772.6 | P | --- |
| 254445_at | 506 | P | 497 | P | 519.4 | P | 504.9 | P | --- |
| 254446_at | 2331.6 | P | 1910.9 | P | 2701.2 | P | 2400.3 | P | TUB9 |
| 254447_at | 553.7 | P | 362.9 | P | 434.5 | P | 289.7 | P | --- |
| 254448_at | 589 | P | 687.9 | P | 615.7 | P | 776.1 | P | FAH2 |
| 254449_at | 79.7 | P | 153.3 | P | 137.4 | P | 172.4 | P | HEN1 |
| 254450_at | 9.4 | A | 1.9 | A | 35.5 | P | 21 | A | --- |
| 254451_at | 511.5 | P | 440.2 | P | 511 | P | 446 | P | --- |
| 254452_at | 570.2 | P | 469.8 | P | 533.7 | P | 560.2 | P | DDB1B |
| 254453_at | 132.4 | M | 183.2 | P | 102.3 | A | 129.5 | A | AAT1 |
| 254454_at | 3.3 | A | 2.4 | A | 2.3 | A | 3.8 | A | EMB2271 |
| 254455_at | 429.4 | P | 317.6 | P | 512.8 | P | 342 | P | --- |
| 254456_at | 1473.3 | P | 1834.9 | P | 1522.1 | P | 1811.7 | P | HAP6 |
| 254457_at | 78.8 | P | 116 | P | 86.4 | P | 125.5 | P | --- |
| 254458_at | 300.4 | P | 379.9 | P | 303.1 | P | 321.6 | P | ATERDJ2B |
| 254459_at | 2.9 | A | 48.4 | A | 48.8 | A | 28.4 | M | GA2OX8 |
| 254460_at | 2641.3 | P | 2620.6 | P | 2490.4 | P | 2622.8 | P | ATRP1 |
| 254461_at | 32.8 | A | 56.3 | A | 27.5 | A | 46.3 | A | --- |
| 254462_at | 7329.6 | P | 6625.5 | P | 6439.5 | P | 7144.8 | P | --- |
| 254463_at | 423.7 | P | 381.9 | P | 332.5 | P | 364.8 | P | TAF11 |
| 254464_at | 220.7 | P | 260.7 | P | 250.5 | P | 273.5 | P | --- |
| 254465_at | 29.2 | A | 28.3 | A | 22.3 | A | 34.1 | A | --- |
| 254466_at | 202.6 | P | 235.9 | P | 284.3 | P | 385.9 | P | --- |
| 254467_at | 38.9 | A | 29.6 | A | 39.4 | A | 8.2 | A | --- |
| 254468_at | 69.4 | P | 102.5 | P | 63.9 | P | 39.7 | P | --- |
| 254469_at | 2.5 | A | 5.5 | A | 6.7 | A | 5.8 | A | --- |
| 254470_at | 1.1 | A | 1 | A | 4.3 | A | 1.5 | A | --- |
| 254471_s_at | 4 | A | 1.8 | A | 8.3 | A | 13.1 | A | --- |
| 254472_at | 3.1 | A | 2.6 | A | 4.3 | A | 1.4 | A | --- |
| 254473_s_at | 1.9 | A | 4 | A | 4.2 | A | 6 | A | --- |
| 254474_at | 33.5 | A | 99.4 | P | 84 | P | 112.1 | P | --- |
| 254475_at | 749.4 | P | 795.8 | P | 894.5 | P | 731.2 | P | smB |
| 254476_at | 525.6 | P | 660.8 | P | 558.3 | P | 660.7 | P | GSNAP |
| 254477_at | 656.6 | P | 774 | P | 588.9 | P | 670.2 | P | --- |
| 254478_at | 1031.6 | P | 1107.2 | P | 872 | P | 948.2 | P | --- |
| 254479_at | 132 | P | 77 | P | 118.1 | P | 68.9 | P | --- |
| 254480_at | 13858.6 | P | 13791.5 | P | 15127.5 | P | 13929.7 | P | ATRABE1B |
| 254481_at | 1192.9 | P | 848.6 | P | 1130.2 | P | 818.8 | P | --- |
| 254482_at | 40 | A | 76.1 | A | 47.7 | A | 33.4 | A | TSF |
| 254483_at | 128.7 | P | 116.6 | P | 152.5 | P | 159.1 | P | --- |
| 254484_at | 275.7 | P | 429.1 | P | 374.8 | P | 447.4 | P | --- |
| 254485_at | 586.8 | P | 770.2 | P | 708.7 | P | 746.6 | P | --- |
| 254486_at | 34.1 | A | 14.2 | A | 38.3 | A | 16.2 | A | --- |
| 254487_at | 215.8 | P | 251.7 | P | 191 | P | 247.2 | P | --- |
| 254488_at | 16.6 | A | 0.3 | A | 6.3 | A | 1.7 | A | --- |

| | | | | | | | | | |
|---|---|---|---|---|---|---|---|---|---|
| 254489_at | 6.9 | A | 1.3 | A | 1.8 | A | 3.5 | A | --- |
| 254490_at | 56.1 | P | 84.1 | P | 73.4 | P | 91.2 | P | --- |
| 254491_at | 429.3 | P | 377.1 | P | 506 | P | 376 | P | --- |
| 254492_at | 2617.3 | P | 3335.3 | P | 2093.2 | P | 3137.8 | P | PCAP1 |
| 254493_at | 681.4 | P | 617.2 | P | 913.5 | P | 604 | P | --- |
| 254494_at | 60.4 | P | 50.2 | P | 56.4 | P | 56.2 | P | QRT3 |
| 254495_at | 62.8 | A | 79.7 | P | 105.8 | P | 85.8 | A | EMB1895 |
| 254496_at | 2573.8 | P | 1481.6 | P | 1944.7 | P | 1736.2 | P | ATAAH |
| 254497_at | 43.8 | A | 24.8 | A | 35.9 | A | 4.2 | A | --- |
| 254498_at | 52.3 | A | 70.8 | A | 18.5 | A | 60.2 | A | EMB1025 |
| 254499_at | 25.9 | M | 23.4 | A | 32.2 | M | 16.8 | A | --- |
| 254500_at | 319.6 | P | 417.1 | P | 156 | P | 265.9 | P | --- |
| 254501_at | 408.6 | P | 574.8 | P | 395.5 | P | 509.5 | P | PTAC14 |
| 254502_at | 699.8 | P | 1035.6 | P | 852.7 | P | 906.1 | P | PTAC14 |
| 254503_at | 711.6 | P | 842.8 | P | 930.1 | P | 733 | P | PTAC9 |
| 254504_at | 290 | P | 421.6 | P | 315.3 | P | 344.6 | P | --- |
| 254505_at | 459.1 | P | 594.2 | P | 577.3 | P | 612.8 | P | --- |
| 254506_at | 43.7 | P | 29.1 | P | 82.9 | P | 58.7 | P | GSO1 |
| 254507_at | 2.9 | A | 4.6 | A | 28.9 | A | 3.4 | A | --- |
| 254508_at | 629.7 | P | 463.1 | P | 596.3 | P | 548.7 | P | --- |
| 254509_at | 2.2 | A | 2.7 | A | 1.3 | A | 5.5 | A | --- |
| 254510_at | 2.1 | A | 2.3 | A | 3.3 | A | 10 | A | --- |
| 254511_at | 18.4 | A | 14.7 | A | 12.6 | A | 13 | A | --- |
| 254512_at | 2.2 | A | 4.7 | A | 7.6 | A | 36.9 | A | --- |
| 254513_at | 1.1 | A | 19.7 | P | 17.5 | P | 16.4 | A | CYP71A27 |
| 254514_at | 10.3 | A | 1.2 | A | 16.2 | A | 6.8 | A | --- |
| 254515_at | 516.9 | P | 480.4 | P | 700.3 | P | 643.8 | P | BAM3 |
| 254516_at | 1 | A | 3.7 | A | 1.6 | A | 2.6 | A | --- |
| 254517_at | 114.3 | P | 129.6 | P | 187.7 | P | 116.1 | P | --- |
| 254518_at | 11.1 | A | 5.3 | A | 2.2 | A | 17.7 | A | --- |
| 254519_at | 16.5 | A | 41.1 | A | 67.4 | A | 8.1 | A | --- |
| 254520_at | 289.3 | P | 368.4 | P | 215.5 | P | 277.1 | P | --- |
| 254521_at | 57.9 | P | 54.7 | P | 41 | A | 77.4 | P | --- |
| 254522_at | 25.1 | A | 24 | A | 4.8 | A | 8.2 | A | --- |
| 254523_at | 98.7 | A | 119.6 | A | 83.8 | A | 89.4 | A | FRS1 |
| 254524_at | 2.1 | A | 7.2 | A | 8 | A | 2.3 | A | --- |
| 254525_at | 4.7 | A | 2.8 | A | 18.3 | A | 16.4 | A | --- |
| 254526_at | 388.9 | P | 357.1 | P | 273.7 | P | 321.7 | P | CYCT1;4 |
| 254527_at | 319.5 | P | 263.7 | P | 239.2 | P | 261.5 | P | --- |
| 254528_at | 1.5 | A | 1.2 | A | 1.4 | A | 2.6 | A | --- |
| 254529_at | 319.2 | P | 283 | P | 369.5 | P | 292.7 | P | INDL |
| 254530_at | 2770.9 | P | 2680.9 | P | 2902.4 | P | 2504.8 | P | ARA7 |
| 254531_at | 82.3 | A | 76.6 | A | 70.2 | A | 84.7 | A | --- |
| 254532_at | 219.6 | P | 353.3 | P | 240.4 | P | 376.6 | P | NPR4 |
| 254533_at | 492.4 | P | 556.1 | P | 542.6 | P | 566 | P | --- |
| 254534_at | 12.8 | A | 2.6 | A | 2.4 | A | 3.5 | A | IRT2 |
| 254535_at | 585.1 | P | 670.8 | P | 666.5 | P | 763.2 | P | AK-HSDH II |
| 254536_at | 8.5 | A | 2 | A | 13.4 | A | 24.1 | A | --- |
| 254537_at | 21.1 | A | 5.5 | A | 26.2 | A | 25.9 | P | --- |
| 254538_at | 19.2 | A | 3.9 | A | 5.9 | A | 6.7 | A | --- |
| 254539_s_at | 1.1 | A | 1.8 | A | 3.1 | A | 1 | A | --- |
| 254540_s_at | 23.6 | P | 4.2 | A | 5 | A | 16.7 | A | --- |
| 254541_at | 7.6 | A | 4.9 | A | 21.7 | M | 1.2 | A | --- |
| 254542_s_at | 7.3 | A | 46.8 | P | 37.9 | A | 30.9 | A | --- |
| 254543_at | 233.8 | P | 222.6 | P | 153.5 | P | 192.5 | P | --- |
| 254544_at | 21.4 | A | 20.5 | A | 44.2 | A | 41.4 | P | --- |
| 254545_at | 210.8 | P | 237.6 | P | 136.9 | P | 228 | P | --- |
| 254546_at | 4.1 | A | 23.3 | A | 12.6 | A | 22.3 | A | PP2-A2 |
| 254547_at | 1085.9 | P | 1467.5 | P | 1008.7 | P | 1227.5 | P | --- |
| 254548_at | 71 | P | 50.7 | P | 62.2 | A | 58.2 | P | --- |
| 254549_at | 3381.5 | P | 2980.3 | P | 3010.8 | P | 2872.4 | P | --- |
| 254550_at | 1.9 | A | 1.1 | A | 12.4 | A | 1.1 | A | IRT1 |
| 254551_at | 1995.2 | P | 1584.4 | P | 1589.7 | P | 1598.7 | P | ATPP2-A1 |
| 254552_at | 81.5 | P | 93.1 | P | 128.3 | P | 118.6 | A | --- |
| 254553_at | 505.5 | P | 454.6 | P | 591.2 | P | 463.4 | P | --- |
| 254554_at | 36.8 | P | 44.7 | P | 34.8 | P | 36 | P | --- |
| 254555_at | 19.1 | A | 1.2 | A | 2.9 | A | 6.8 | A | --- |
| 254556_at | 41.8 | P | 54 | P | 55 | P | 47.8 | P | --- |
| 254557_at | 894.6 | P | 682 | P | 922.5 | P | 713.3 | P | --- |
| 254558_at | 273.3 | P | 343.7 | P | 183.9 | P | 351.7 | P | --- |
| 254559_at | 2008.3 | P | 3015.2 | P | 1855.8 | P | 2814.7 | P | --- |
| 254560_at | 1054.3 | P | 1211.5 | P | 995.7 | P | 1130.9 | P | --- |
| 254561_at | 612.7 | P | 797.5 | P | 455.5 | P | 685.2 | P | --- |
| 254562_at | 85.8 | P | 66.3 | A | 39 | A | 64.4 | A | CYP707A1 |
| 254563_at | 1760.1 | P | 2080.9 | P | 1334.4 | P | 2300.7 | P | ERD3 |
| 254564_at | 257.5 | P | 640.4 | P | 232.3 | P | 345 | P | NCED4 |
| 254565_at | 61.5 | A | 53.9 | A | 33.4 | A | 49.2 | A | --- |
| 254566_at | 3.3 | A | 10.1 | A | 12.6 | A | 20.2 | A | --- |
| 254567_s_at | 0.9 | A | 1 | A | 18.8 | A | 11.4 | A | --- |
| 254568_x_at | 16 | A | 27.3 | A | 18.7 | A | 26.4 | A | --- |
| 254569_s_at | 19.1 | A | 20.9 | A | 5.4 | A | 15.3 | A | --- |
| 254570_s_at | 9.4 | A | 3.2 | A | 3.9 | A | 6.8 | A | --- |
| 254571_at | 70.3 | A | 75 | M | 55.8 | A | 49 | A | --- |
| 254572_at | 50 | P | 61.7 | P | 78.7 | P | 112.6 | P | --- |
| 254573_at | 862.2 | P | 784.9 | P | 594.7 | P | 667 | P | --- |
| 254574_at | 305.9 | P | 1556.3 | P | 569.6 | P | 967.6 | P | --- |
| 254575_at | 73.4 | P | 54.9 | P | 67.2 | P | 59.7 | P | --- |
| 254576_at | 394.5 | P | 427.9 | P | 551.4 | P | 430 | P | EMB3006 |
| 254577_at | 796.2 | P | 801.4 | P | 1079.7 | P | 935.9 | P | --- |
| 254578_at | 9221.3 | P | 7133.8 | P | 7622.9 | P | 7497.4 | P | --- |
| 254579_at | 337.2 | P | 295.7 | P | 377.8 | P | 261.6 | P | --- |
| 254580_at | 603.7 | P | 590.4 | P | 478.3 | P | 606.2 | P | --- |
| 254581_at | 90 | P | 66.8 | P | 91.5 | P | 101.2 | P | --- |
| 254582_at | 4.5 | A | 11.7 | A | 7.1 | A | 23.2 | P | --- |
| 254583_at | 5.7 | A | 6.1 | A | 8.7 | A | 6.9 | A | --- |
| 254584_at | 248.3 | P | 296.8 | P | 278.8 | P | 263.1 | P | VPS54 |
| 254585_at | 473.3 | P | 487.3 | P | 493.1 | P | 470.3 | P | --- |
| 254586_at | 204 | P | 188.9 | P | 133.5 | P | 182.3 | P | --- |
| 254587_at | 80 | A | 45.2 | A | 60.9 | A | 61.2 | A | --- |
| 254588_at | 115 | P | 107.7 | P | 109.2 | P | 117.6 | P | --- |
| 254589_at | 0.8 | A | 2.4 | A | 1.2 | A | 0.9 | A | --- |
| 254590_at | 19.7 | A | 16.1 | A | 36.8 | A | 19.4 | A | --- |
| 254591_at | 4.2 | A | 1.7 | A | 2.7 | A | 4.6 | A | --- |
| 254592_at | 309.7 | P | 441.1 | P | 260.9 | P | 328.2 | P | HSF A4A |
| 254593_s_at | 390.9 | P | 258.5 | P | 349.7 | P | 298.3 | P | --- |
| 254594_at | 1780.5 | P | 1664 | P | 1165.9 | P | 1579.8 | P | --- |
| 254595_at | 18 | A | 31.5 | P | 16.8 | A | 25.2 | M | AG |
| 254596_at | 363.4 | P | 459.9 | P | 301.4 | P | 398.9 | P | --- |
| 254597_at | 11.9 | A | 6.6 | A | 4.1 | A | 19 | A | --- |

| | | | | | | | | | |
|---|---|---|---|---|---|---|---|---|---|
| 254598_at | 50.8 | P | 16.3 | A | 27.3 | A | 36.9 | A | --- |
| 254599_at | 4.1 | A | 19.7 | A | 19.7 | A | 16.4 | A | --- |
| 254600_at | 144.4 | P | 187.7 | P | 201.8 | P | 137.9 | P | --- |
| 254601_at | 148.4 | P | 142.5 | P | 160.3 | P | 161.8 | P | CMT2 |
| 254602_at | 552.9 | P | 620.6 | P | 601.7 | P | 680.6 | P | EDR2 |
| 254603_at | 121.9 | P | 145.1 | P | 199.9 | P | 142.3 | P | --- |
| 254604_at | 195.1 | P | 222.2 | P | 239.8 | P | 202.8 | P | --- |
| 254605_at | 428.8 | P | 304.2 | P | 324.4 | P | 302.6 | P | --- |
| 254606_at | 215.6 | P | 134 | P | 161.8 | P | 205.8 | P | NLM1 |
| 254607_at | 1.7 | A | 12.7 | A | 9.1 | A | 12.1 | A | --- |
| 254608_at | 265.3 | P | 300.6 | P | 213.8 | P | 244.1 | P | NIP1;2 |
| 254609_at | 303.6 | P | 1017.2 | P | 484.9 | P | 655.3 | P | --- |
| 254610_at | 187.1 | P | 270.6 | P | 183.7 | P | 257.4 | P | --- |
| 254611_at | 34.5 | A | 62.9 | P | 63.5 | A | 27.2 | A | --- |
| 254612_at | 505.4 | P | 656 | P | 568.3 | P | 641.6 | P | --- |
| 254613_at | 340.2 | P | 255.1 | P | 366.1 | P | 233.6 | P | --- |
| 254614_at | 45.5 | P | 53.2 | P | 73.8 | P | 23.7 | P | --- |
| 254615_at | 2819.2 | P | 1898.9 | P | 2996.4 | P | 2070.9 | P | RLI2 |
| 254616_at | 1154.9 | P | 1164.6 | P | 1285.1 | P | 1315.6 | P | BIN2 |
| 254617_s_at | 8555.9 | P | 7145.6 | P | 7983.9 | P | 7731.1 | P | RPL16B |
| 254618_at | 90.3 | P | 124.9 | P | 107.8 | P | 170.6 | P | IRX1 |
| 254619_at | 34.9 | A | 38.2 | A | 68.9 | A | 44.9 | A | MYB98 |
| 254620_at | 586.2 | P | 361.8 | P | 534.5 | P | 322 | P | --- |
| 254621_at | 157.7 | P | 140.5 | P | 172.3 | P | 187.8 | P | WAVE5 |
| 254622_at | 50 | A | 49 | A | 41.3 | A | 49.9 | A | --- |
| 254623_at | 3923.4 | P | 5227.7 | P | 4600.1 | P | 5055.1 | P | CHLI1 |
| 254624_at | 285.9 | P | 231.3 | P | 311.1 | P | 273.4 | P | --- |
| 254625_at | 53.7 | P | 40.3 | P | 57.4 | P | 33.9 | P | SNI1 |
| 254626_at | 405.6 | P | 459 | P | 358.8 | P | 413.2 | P | --- |
| 254627_at | 16.1 | A | 13 | A | 8.6 | A | 13.7 | A | --- |
| 254628_at | 471.7 | P | 456.5 | P | 556.9 | P | 466 | P | --- |
| 254629_at | 30.7 | A | 65.3 | P | 77.7 | P | 76.8 | P | --- |
| 254630_at | 841.5 | P | 261.3 | P | 526.8 | P | 228.8 | P | --- |
| 254631_at | 39.4 | A | 57.1 | P | 60.1 | A | 69.2 | A | LSH9 |
| 254632_at | 86 | P | 54.5 | A | 46.9 | P | 40.4 | M | --- |
| 254633_at | 52 | P | 100.1 | P | 110 | P | 121.5 | P | MRH1 |
| 254634_at | 28.1 | A | 8.3 | A | 46.9 | A | 34.5 | A | --- |
| 254635_at | 45.5 | P | 57 | P | 62.4 | P | 65 | P | --- |
| 254636_at | 911.6 | P | 728.5 | P | 805.3 | P | 811.9 | P | CIPK12 |
| 254637_at | 15.3 | A | 4.3 | A | 37.2 | A | 15.9 | A | --- |
| 254638_at | 291 | P | 335.3 | P | 258.6 | P | 278.5 | P | --- |
| 254639_at | 98.1 | P | 103.7 | P | 126.5 | P | 108.4 | P | DOT4 |
| 254640_at | 4.5 | A | 16.1 | A | 68.5 | M | 33.1 | P | NRAMP5 |
| 254641_at | 284.1 | P | 299 | P | 261.5 | P | 282.2 | P | ATRABA1D |
| 254642_at | 842.1 | P | 1027.4 | P | 909.1 | P | 1030.7 | P | --- |
| 254643_at | 246.4 | P | 300.7 | P | 295.2 | P | 334.4 | P | --- |
| 254644_at | 10.8 | A | 12.1 | A | 6.2 | A | 4.8 | A | CLE2 |
| 254645_at | 293 | P | 244.1 | P | 256.6 | P | 216.3 | P | --- |
| 254646_at | 169 | P | 115.2 | P | 130.9 | P | 144.1 | P | --- |
| 254647_at | 3.5 | A | 3.2 | A | 10.7 | A | 8.7 | A | --- |
| 254648_at | 13.2 | A | 6.5 | A | 6.3 | A | 30.7 | A | --- |
| 254649_at | 994.6 | P | 571.6 | P | 902.8 | P | 721.5 | P | --- |
| 254650_at | 41.6 | A | 31.2 | A | 70.5 | A | 8.6 | A | --- |
| 254651_at | 324.9 | P | 234 | P | 295.7 | P | 284 | P | KCO6 |
| 254652_at | 174.8 | P | 32.1 | A | 75.1 | P | 45.5 | A | WRKY28 |
| 254653_at | 6.1 | A | 5.1 | A | 45.5 | A | 2.7 | A | --- |
| 254654_at | 2184.3 | P | 2101.4 | P | 2751.8 | P | 2120.5 | P | EIF4E |
| 254655_s_at | 9675.5 | P | 7387 | P | 6711.7 | P | 7936.3 | P | --- |
| 254656_at | 785.6 | P | 1192.6 | P | 984.5 | P | 1082.2 | P | --- |
| 254657_s_at | 321.2 | P | 240.5 | P | 360.4 | P | 216.6 | P | --- |
| 254658_at | 509.6 | P | 544.3 | P | 366.5 | P | 393.1 | P | --- |
| 254659_at | 806.4 | P | 1294.5 | P | 727.8 | P | 1160.2 | P | SS4 |
| 254660_at | 81.7 | P | 66.5 | P | 65.6 | P | 48.4 | P | --- |
| 254661_at | 108.9 | P | 109.3 | P | 78 | P | 137.1 | P | --- |
| 254662_at | 131.6 | A | 157.5 | P | 197.2 | A | 178.9 | P | TRANS11 |
| 254663_at | 256.8 | P | 126.8 | P | 594 | P | 267.7 | P | KAT2 |
| 254664_at | 136.3 | P | 160.3 | P | 184.9 | P | 179.1 | P | --- |
| 254665_at | 30.1 | A | 59.7 | P | 33.5 | A | 44.1 | A | --- |
| 254666_at | 37.3 | A | 42.2 | A | 80.3 | A | 22.5 | A | --- |
| 254667_at | 219.5 | P | 703 | P | 195 | P | 638.4 | P | --- |
| 254668_at | 48 | A | 3.4 | A | 32 | A | 7.7 | A | NCED2 |
| 254669_at | 1117.2 | P | 1381.8 | P | 1534 | P | 1453 | P | DEG5 |
| 254670_at | 758 | P | 1070.2 | P | 928.9 | P | 1019.6 | P | --- |
| 254671_at | 11.1 | A | 25 | A | 14.6 | A | 1.3 | A | --- |
| 254672_at | 5.5 | A | 5.3 | A | 7.4 | A | 2.3 | A | --- |
| 254673_at | 48.7 | P | 64.8 | P | 61.3 | P | 64.4 | M | AtRABA1e |
| 254674_at | 5.8 | A | 3 | A | 5.7 | A | 31.7 | A | --- |
| 254675_at | 131.9 | P | 170.6 | P | 205.8 | P | 179.2 | P | --- |
| 254676_at | 139.4 | P | 248.6 | P | 245.2 | P | 264.4 | P | --- |
| 254677_at | 38.6 | P | 51.7 | P | 19.7 | A | 38.3 | M | --- |
| 254678_at | 2.3 | A | 0.9 | A | 1.4 | A | 1.5 | A | --- |
| 254679_at | 612.1 | P | 504.5 | P | 560.6 | P | 689.6 | P | AML3 |
| 254680_at | 552.9 | P | 525 | P | 543.6 | P | 565.4 | P | PHYE |
| 254681_at | 237.1 | P | 186.6 | P | 187.9 | P | 131.9 | P | --- |
| 254682_at | 518.2 | P | 470.1 | P | 338.1 | P | 437.7 | P | UNE16 |
| 254683_at | 69.4 | P | 76.1 | P | 68.9 | P | 56.8 | P | --- |
| 254684_at | 9293.9 | P | 6454.5 | P | 10178.6 | P | 7213.7 | P | GR-RBP2 |
| 254685_at | 31.4 | P | 2.3 | A | 3.1 | A | 3.6 | A | --- |
| 254686_at | 1103.5 | P | 1091.9 | P | 1123.7 | P | 1264.9 | P | --- |
| 254687_at | 5775.4 | P | 4065.8 | P | 7420.1 | P | 4236.9 | P | CYP83A1 |
| 254688_at | 435.8 | P | 625 | P | 435.6 | P | 530.3 | P | J20 |
| 254689_at | 66.8 | P | 71.2 | P | 90.5 | P | 103.1 | P | --- |
| 254690_at | 1265 | P | 1317.3 | P | 851.5 | P | 1153.4 | P | --- |
| 254691_at | 2053.6 | P | 1872.6 | P | 1912.2 | P | 1994 | P | --- |
| 254692_at | 6.2 | A | 56.6 | A | 6.5 | A | 23 | A | --- |
| 254693_at | 201.6 | P | 246.5 | P | 236.8 | P | 267.7 | P | --- |
| 254694_at | 866.5 | P | 761.2 | P | 687.2 | P | 656 | P | --- |
| 254695_at | 61.1 | P | 75 | P | 70.2 | P | 53.4 | P | ATL4H |
| 254696_at | 1.7 | A | 0.5 | A | 9.3 | A | 2.5 | A | --- |
| 254697_at | 370.7 | P | 307.7 | P | 456.6 | P | 322.5 | P | --- |
| 254698_at | 5.3 | A | 24.1 | A | 22.2 | A | 29.7 | A | anac071 |
| 254699_at | 17.3 | A | 4.2 | A | 6.1 | A | 12.8 | A | --- |
| 254700_at | 24.2 | A | 7.8 | A | 12.3 | A | 4.9 | A | --- |
| 254701_at | 1623.2 | P | 2381.2 | P | 1362.3 | P | 2430.3 | P | --- |
| 254702_at | 708.4 | P | 681.5 | P | 633.4 | P | 650.3 | P | --- |
| 254703_at | 177.9 | P | 189.3 | P | 218 | P | 177.7 | P | --- |
| 254704_at | 338.2 | P | 488.9 | P | 527.6 | P | 488.6 | P | APRR2 |
| 254705_at | 920.3 | P | 1381.1 | P | 1284.2 | P | 1273.6 | P | --- |
| 254706_at | 238.2 | P | 231.5 | P | 239.6 | P | 200.6 | P | --- |

| | | | | | | | | | |
|---|---|---|---|---|---|---|---|---|---|
| 254707_at | 820.1 | P | 674 | P | 600.9 | P | 602 | P | AT5PTASE2 |
| 254708_at | 152.1 | P | 163.8 | P | 135 | P | 121.4 | P | UBP20 |
| 254709_at | 719.2 | P | 1065.8 | P | 926.9 | P | 1090 | P | AGD8 |
| 254710_at | 174.6 | P | 121.1 | P | 123.1 | P | 124.2 | P | PGP9 |
| 254711_at | 461.4 | P | 520.3 | P | 489.1 | P | 536 | P | --- |
| 254712_at | 4.4 | A | 3.4 | A | 2.6 | A | 5.8 | A | --- |
| 254713_at | 16.3 | P | 15 | A | 44.2 | A | 14.9 | A | --- |
| 254714_at | 12 | A | 13.4 | A | 25.9 | A | 1.7 | A | AtMYB79 |
| 254715_at | 486.3 | P | 459.1 | P | 411.8 | P | 361.1 | P | --- |
| 254716_at | 146.2 | P | 75.7 | P | 87.6 | P | 91.2 | P | UNE15 |
| 254717_at | 30.1 | A | 15.6 | A | 44.7 | P | 31.5 | P | HTA4 |
| 254718_at | 3.3 | A | 3.9 | A | 6 | A | 4.1 | A | --- |
| 254719_at | 38.4 | P | 60.6 | P | 84.1 | P | 65.4 | P | --- |
| 254720_at | 29.3 | A | 46 | P | 26.4 | A | 34.2 | A | MEE57 |
| 254721_at | 2440 | P | 3010.1 | P | 2432.2 | P | 3186.9 | P | SMAP1 |
| 254722_at | 1296.3 | P | 1381.5 | P | 1098.4 | P | 1287.2 | P | --- |
| 254723_at | 1322.8 | P | 2995.3 | P | 1505.6 | P | 2275.4 | P | AMT1;1 |
| 254724_at | 249.3 | P | 231.1 | P | 190.6 | P | 193.5 | P | --- |
| 254725_at | 70.9 | P | 61.8 | P | 103 | P | 66.1 | P | --- |
| 254726_at | 65.7 | P | 146.5 | P | 46.3 | A | 136.4 | P | PRR2 |
| 254727_at | 702.4 | P | 1052.8 | P | 719.8 | P | 913.3 | P | PTAC5 |
| 254728_at | 4 | A | 4.7 | A | 6.8 | A | 30.6 | A | --- |
| 254729_at | 2.9 | A | 12.5 | A | 7.9 | A | 4.9 | A | PAP23 |
| 254730_at | 572.5 | P | 546.4 | P | 636.9 | P | 514.1 | P | --- |
| 254731_at | 3 | A | 31 | M | 5.6 | A | 36.4 | P | --- |
| 254732_at | 202.3 | P | 151 | P | 236.2 | P | 164.2 | P | --- |
| 254733_at | 24.1 | A | 18 | A | 34.3 | A | 0.9 | A | --- |
| 254734_at | 1214 | P | 1419.8 | P | 1229.2 | P | 1011.5 | P | --- |
| 254735_at | 86.9 | P | 97.3 | P | 82.9 | P | 89.2 | P | AtRLP47 |
| 254736_at | 55.7 | A | 37.4 | A | 58.5 | A | 63.4 | A | --- |
| 254737_at | 2182.8 | P | 1935.6 | P | 2180.3 | P | 2089 | P | --- |
| 254738_at | 206.3 | P | 47.7 | P | 94.9 | P | 39.6 | P | AT4G13860 |
| 254739_at | 1.5 | A | 10.5 | A | 27 | A | 10.1 | A | AtRLP48 |
| 254740_s_at | 2664.5 | P | 2781.9 | P | 2527.2 | P | 3143.8 | P | --- |
| 254741_s_at | 158.5 | P | 252.9 | P | 135.6 | P | 189.6 | P | AtRLP49 |
| 254742_at | 8982.4 | P | 6327.5 | P | 9127.7 | P | 6691 | P | IIL1 |
| 254743_at | 158.7 | P | 37.9 | A | 158.8 | P | 43.6 | A | HAK5 |
| 254744_at | 154.9 | P | 239.7 | P | 208.7 | P | 219.9 | P | MEE55 |
| 254745_at | 528.3 | P | 521.5 | P | 754.3 | P | 564.7 | P | SUVH9 |
| 254746_at | 95.1 | P | 187.7 | P | 185.6 | P | 213.4 | P | --- |
| 254747_at | 451.3 | P | 593.8 | P | 432 | P | 570.9 | P | MHK |
| 254748_at | 137.8 | P | 102.3 | A | 141.3 | A | 113.7 | A | --- |
| 254749_at | 6.9 | A | 45.3 | A | 7.7 | A | 33.3 | M | --- |
| 254750_at | 545.2 | P | 699.9 | P | 326.2 | P | 642.8 | P | --- |
| 254751_at | 58.5 | P | 52.8 | P | 69.1 | A | 86.9 | P | --- |
| 254752_at | 74.2 | P | 84.9 | P | 62.6 | P | 80.5 | P | --- |
| 254753_at | 6.2 | A | 3.8 | A | 15.7 | A | 17.8 | A | --- |
| 254754_at | 50.5 | P | 80.7 | P | 57.7 | P | 95.7 | P | --- |
| 254755_at | 825.9 | P | 1074 | P | 753 | P | 964.5 | P | --- |
| 254756_at | 27.5 | A | 10.5 | A | 14.2 | A | 35.2 | A | EDA21 |
| 254757_at | 4.3 | A | 29.3 | A | 33.1 | A | 28 | A | ROPGEF9 |
| 254758_at | 137.9 | P | 121.5 | P | 174.1 | P | 144.7 | P | YUC2 |
| 254759_at | 2351.2 | P | 1697.5 | P | 1952.8 | P | 1902.6 | P | --- |
| 254760_at | 1089.6 | P | 1563.3 | P | 1074 | P | 1495.8 | P | --- |
| 254761_at | 295.2 | P | 213.9 | P | 447.9 | P | 260.8 | P | CLE44 |
| 254762_at | 42.1 | A | 33.5 | A | 35.2 | A | 8 | A | --- |
| 254763_at | 2810.2 | P | 2535.3 | P | 3471.8 | P | 2870.4 | P | --- |
| 254764_at | 4805.4 | P | 2522.8 | P | 2729.5 | P | 2691.9 | P | --- |
| 254765_at | 598.7 | P | 701.4 | P | 434.3 | P | 591.4 | P | --- |
| 254766_at | 2.2 | A | 24.7 | A | 7.2 | A | 12.1 | A | TPS12 |
| 254767_s_at | 0.9 | A | 19.5 | P | 10.4 | A | 19.1 | A | CYP71A19 |
| 254768_at | 1.4 | A | 5.5 | A | 35.2 | A | 1.9 | A | --- |
| 254769_at | 106.2 | P | 97.3 | P | 143.7 | P | 80.5 | P | --- |
| 254770_at | 212.4 | P | 129 | P | 219.1 | P | 146.5 | P | --- |
| 254771_at | 6.4 | A | 0.6 | A | 2.3 | A | 1.3 | A | MEE56 |
| 254772_at | 0.8 | A | 1.3 | A | 15.1 | A | 14.3 | A | --- |
| 254773_at | 177.4 | P | 146.8 | P | 225.1 | P | 214.3 | P | ATCSLA15 |
| 254774_at | 27.7 | A | 11 | A | 32.9 | A | 25.7 | A | --- |
| 254775_at | 30.1 | P | 19.4 | A | 45.8 | A | 12.3 | A | --- |
| 254776_at | 1607.2 | P | 1346.1 | P | 1516.7 | P | 1425.1 | P | --- |
| 254777_at | 18.3 | A | 5.1 | A | 1.6 | A | 11.7 | A | --- |
| 254778_at | 395.6 | P | 418.5 | P | 481.4 | P | 415.3 | P | --- |
| 254779_at | 294.5 | P | 298.4 | P | 285.9 | P | 229.1 | P | --- |
| 254780_s_at | 141.3 | P | 153.1 | P | 149.3 | P | 131.2 | P | --- |
| 254781_at | 49.8 | P | 103.1 | P | 75.8 | P | 110.4 | P | --- |
| 254782_at | 29.6 | P | 7 | A | 22.3 | A | 6.4 | A | UNE14 |
| 254783_at | 304.2 | P | 385.7 | P | 313.7 | P | 298.8 | P | --- |
| 254784_at | 3079.6 | P | 3470 | P | 2537.9 | P | 2669.2 | P | NUDT7 |
| 254785_at | 2139.5 | P | 1502.7 | P | 1882.2 | P | 1862.3 | P | FLA2 |
| 254786_at | 33.1 | A | 63.4 | P | 48.6 | P | 68.6 | P | --- |
| 254787_at | 85.7 | P | 69.9 | P | 54.3 | P | 77.8 | P | --- |
| 254788_at | 1090.4 | P | 875.6 | P | 1066.3 | P | 853.2 | P | --- |
| 254789_at | 839.2 | P | 2182.7 | P | 890.8 | P | 2018.6 | P | --- |
| 254790_at | 9625.7 | P | 11132.1 | P | 9903.4 | P | 10745 | P | PSAL |
| 254791_at | 282.2 | P | 369 | P | 330.2 | P | 468.9 | P | scpl20 |
| 254792_at | 2.3 | A | 1.6 | A | 6 | A | 4 | A | --- |
| 254793_at | 1.6 | A | 14.6 | A | 1.6 | A | 12.3 | A | --- |
| 254794_at | 698.5 | P | 680.4 | P | 875.1 | P | 803.4 | P | --- |
| 254795_at | 93.4 | P | 98.8 | P | 108.9 | P | 109.2 | P | --- |
| 254796_at | 57 | P | 75 | P | 86.8 | P | 50.3 | M | --- |
| 254797_at | 82.8 | M | 92.6 | P | 93.2 | P | 100.4 | P | --- |
| 254798_at | 511 | P | 543.6 | P | 549.3 | P | 483.1 | P | --- |
| 254799_at | 0.8 | A | 17.3 | A | 20.2 | A | 1.6 | A | --- |
| 254800_at | 71.6 | P | 68.5 | P | 77.8 | P | 89.8 | P | --- |
| 254801_at | 23.5 | A | 31.9 | A | 36.3 | A | 15.7 | A | --- |
| 254802_at | 0.9 | A | 4 | A | 2.9 | A | 13.2 | A | --- |
| 254803_at | 306.2 | P | 418.6 | P | 293.2 | P | 510.4 | P | --- |
| 254804_at | 2011.6 | P | 2460.5 | P | 2049.7 | P | 2185.4 | P | --- |
| 254805_at | 3867 | P | 5648.1 | P | 2184 | P | 3762.1 | P | pEARLI 1 |
| 254806_at | 132 | P | 182.9 | P | 169.3 | P | 169.4 | P | --- |
| 254807_at | 202 | P | 191.6 | P | 235.7 | P | 203.2 | P | --- |
| 254808_at | 23.1 | A | 41.7 | P | 34.2 | A | 36.9 | A | --- |
| 254809_at | 45.5 | A | 35.8 | A | 20.5 | A | 26.4 | A | --- |
| 254810_at | 1006.1 | P | 851.9 | P | 1323 | P | 921.9 | P | PME1 |
| 254811_at | 622 | P | 765.3 | P | 493.7 | P | 690.7 | P | --- |
| 254812_at | 259.6 | P | 203 | P | 182 | P | 201.3 | P | GAE5 |
| 254813_at | 4.7 | A | 2.2 | A | 2.5 | A | 2.3 | A | MYB42 |
| 254814_at | 835.1 | P | 696.4 | P | 736.1 | P | 637.2 | P | --- |
| 254815_at | 1412 | P | 2018.9 | P | 1767.6 | P | 2059 | P | SKU5 |

| | | | | | | | | | |
|---|---|---|---|---|---|---|---|---|---|
| 254816_at | 9.8 | A | 30.3 | A | 9.3 | A | 29.2 | A | APT4 |
| 254817_at | 225.8 | P | 285 | P | 187.9 | P | 257.9 | P | ORP2B |
| 254818_at | 3404.4 | P | 5960.6 | P | 2425.9 | P | 4280.4 | P | --- |
| 254819_at | 150 | P | 308.5 | P | 141.5 | P | 279.4 | P | --- |
| 254820_s_at | 63.3 | A | 105 | P | 102.8 | P | 69.4 | P | --- |
| 254821_at | 26.6 | A | 30.2 | A | 21.4 | M | 24.6 | A | --- |
| 254822_at | 272.8 | P | 382.1 | P | 325.1 | P | 277.5 | P | UPL5 |
| 254823_at | 15.2 | A | 13.6 | M | 24.4 | A | 35.3 | P | --- |
| 254824_s_at | 47.5 | P | 104.6 | P | 110.6 | P | 83 | P | ORC1B |
| 254825_at | 85.3 | P | 94.2 | P | 94.8 | P | 98.4 | P | --- |
| 254826_at | 233.1 | P | 234.1 | P | 205.5 | P | 210.7 | P | --- |
| 254827_at | 1569.5 | P | 1840 | P | 1651.1 | P | 1828.7 | P | --- |
| 254828_at | 50.1 | P | 45.4 | P | 69.4 | P | 45.6 | P | AIR1 |
| 254829_at | 25.5 | A | 14 | A | 21.6 | A | 1.2 | A | --- |
| 254830_at | 1740.8 | P | 2222.4 | P | 1639.7 | P | 2218.5 | P | --- |
| 254831_at | 2984.8 | P | 2494.2 | P | 3619.7 | P | 2973.8 | P | --- |
| 254832_at | 1741.9 | P | 1790.7 | P | 762.9 | P | 1259.6 | P | --- |
| 254833_s_at | 2670.7 | P | 2786.1 | P | 2195.8 | P | 2747.1 | P | --- |
| 254834_at | 394 | P | 456.9 | P | 461.5 | P | 428.7 | P | CYP706A4 |
| 254835_s_at | 1329.3 | P | 1729.3 | P | 1904 | P | 1847.1 | P | CYP706A5 |
| 254836_at | 11.1 | A | 26.5 | A | 3 | A | 12.8 | A | CYP706A7 |
| 254837_at | 3.2 | A | 16.8 | A | 30.5 | A | 15.6 | A | --- |
| 254838_at | 3.5 | A | 1.9 | A | 13.7 | A | 17.6 | A | --- |
| 254839_at | 64.8 | P | 88.1 | P | 62.8 | P | 72.2 | P | --- |
| 254840_at | 2 | A | 22.2 | A | 12.1 | A | 1.7 | A | --- |
| 254841_at | 26.6 | A | 32.2 | A | 54.1 | A | 44 | A | --- |
| 254842_at | 50.1 | A | 54.4 | A | 99.5 | A | 63.7 | M | --- |
| 254843_at | 357.8 | P | 312.5 | P | 303.1 | P | 271.2 | P | --- |
| 254844_at | 367.9 | P | 357 | P | 467.8 | P | 367.4 | P | --- |
| 254845_at | 919.3 | P | 779.6 | P | 975.8 | P | 799.4 | P | MVA1 |
| 254846_at | 230.2 | P | 255.5 | P | 174.2 | P | 270.7 | P | PLDGAMMA2 |
| 254847_at | 98.6 | P | 92.1 | P | 120.2 | P | 96.5 | P | PLDGAMMA1 |
| 254848_at | 800.7 | P | 1121.7 | P | 770.2 | P | 1037.4 | P | PGRL1B |
| 254849_at | 142.8 | P | 96.2 | P | 145.4 | P | 174.1 | P | --- |
| 254850_at | 242.7 | P | 220.7 | P | 240.5 | P | 312 | P | --- |
| 254851_at | 225.5 | P | 243.1 | P | 177.8 | P | 185 | P | --- |
| 254852_at | 149.2 | P | 125.6 | P | 147.3 | P | 128.1 | P | WRKY19 |
| 254853_at | 293.1 | P | 166.6 | P | 178.4 | P | 170.5 | P | --- |
| 254854_at | 303.9 | P | 221.1 | P | 357.9 | P | 201.9 | P | --- |
| 254855_s_at | 13.3 | A | 20.3 | A | 28.8 | A | 33.5 | A | --- |
| 254856_at | 0.9 | A | 8.9 | A | 3.3 | A | 18.9 | A | --- |
| 254857_at | 90.7 | P | 89.3 | P | 63.3 | A | 89.9 | P | SEC1B |
| 254858_at | 275.3 | P | 344.3 | P | 271.1 | P | 324.8 | P | --- |
| 254859_at | 1939.4 | P | 2150.9 | P | 1950.3 | P | 1969.3 | P | --- |
| 254860_at | 732.6 | P | 686.6 | P | 733.9 | P | 759.9 | P | SMO1-1 |
| 254861_at | 1216.3 | P | 856.3 | P | 927.8 | P | 903.9 | P | --- |
| 254862_at | 1864.3 | P | 1240.3 | P | 2688.2 | P | 1343.5 | P | --- |
| 254863_at | 380.7 | P | 456.4 | P | 318.1 | P | 467.7 | P | ATNUDX14 |
| 254864_at | 10.3 | A | 4.3 | A | 16.9 | A | 11.1 | A | --- |
| 254865_s_at | 23.2 | P | 16.2 | A | 48.5 | A | 6.5 | A | --- |
| 254866_at | 2.5 | A | 4.3 | A | 26.4 | A | 25.9 | A | --- |
| 254867_at | 134.1 | P | 104 | P | 93.7 | P | 91.2 | P | --- |
| 254868_at | 1.1 | A | 2.5 | A | 2.6 | A | 2.2 | A | --- |
| 254869_at | 288.7 | P | 417.4 | P | 76.4 | P | 232.7 | P | --- |
| 254870_at | 48.5 | P | 95.2 | P | 53.3 | P | 113 | P | --- |
| 254871_at | 45.5 | A | 56.6 | A | 75.9 | A | 59.6 | M | CCS52A2 |
| 254872_at | 509.8 | P | 570.8 | P | 505.7 | P | 540.1 | P | --- |
| 254873_at | 1400.8 | P | 1144.3 | P | 1519.4 | P | 1263 | P | EIF3A |
| 254874_at | 1990.6 | P | 2221.2 | P | 1680.2 | P | 1800.8 | P | --- |
| 254875_at | 0.8 | A | 23.3 | A | 21.3 | A | 1.8 | A | --- |
| 254876_at | 42.5 | P | 57.9 | A | 77.4 | P | 37.5 | A | --- |
| 254877_at | 176.2 | P | 164.2 | P | 243.6 | P | 167.8 | P | ATSR |
| 254878_at | 75.8 | P | 132.2 | P | 124.5 | P | 112.9 | P | AT-HSFB2B |
| 254879_at | 262.4 | P | 288.2 | P | 325.7 | P | 274.4 | P | --- |
| 254880_at | 48.3 | A | 48.6 | A | 47.8 | A | 33.6 | A | --- |
| 254881_at | 2.6 | A | 9.1 | A | 9.7 | A | 5.3 | A | --- |
| 254882_s_at | 6.4 | A | 1.2 | A | 3.3 | A | 1.3 | A | --- |
| 254883_at | 1.7 | A | 2.7 | A | 20.4 | A | 19.1 | A | HAP2 |
| 254884_at | 10.2 | A | 5 | A | 4.7 | A | 3.8 | A | --- |
| 254885_at | 2 | A | 3.3 | A | 3.1 | A | 2.4 | A | --- |
| 254886_at | 5.8 | A | 3.4 | A | 7.7 | A | 24.5 | A | LCR17 |
| 254887_at | 26.2 | A | 22.9 | A | 30.7 | A | 32.2 | A | --- |
| 254888_at | 11.8 | A | 18.3 | A | 28.2 | P | 17.1 | A | --- |
| 254889_at | 4.2 | A | 129.3 | P | 121.3 | P | 186 | P | ATOSM34 |
| 254890_at | 8752.7 | P | 8064.3 | P | 7356.7 | P | 9685.5 | P | ATGPX6 |
| 254891_at | 401.1 | P | 470.7 | P | 348.7 | P | 358 | P | SAY1 |
| 254892_at | 434.7 | P | 468 | P | 465.6 | P | 474.4 | P | --- |
| 254893_at | 107 | P | 107.6 | P | 110.9 | P | 71.2 | P | PLDGAMMA2 |
| 254894_at | 41.1 | P | 48.1 | P | 32.3 | P | 46.3 | P | PLDGAMMA3 |
| 254895_at | 19.5 | A | 23.7 | A | 29.7 | A | 27.1 | A | --- |
| 254896_at | 0.8 | A | 2.6 | A | 2.2 | A | 2 | A | AGL14 |
| 254897_at | 17.5 | A | 23 | A | 18 | A | 23.9 | A | --- |
| 254898_at | 0.5 | A | 2.2 | A | 2.7 | A | 7 | A | --- |
| 254899_at | 20.5 | A | 18.3 | A | 28.4 | A | 8.7 | A | --- |
| 254900_at | 14.2 | A | 8.7 | A | 34.6 | A | 0.8 | A | RALFL28 |
| 254901_at | 47.3 | P | 28.3 | P | 45.3 | A | 56.5 | M | --- |
| 254902_at | 7 | A | 11.7 | A | 23.8 | A | 4.1 | A | --- |
| 254903_at | 8639.8 | P | 7884.8 | P | 8808.2 | P | 8388.1 | P | TUF |
| 254904_at | 673.8 | P | 556 | P | 659.2 | P | 544.8 | P | --- |
| 254905_at | 33.5 | P | 69 | P | 64.1 | P | 34.4 | A | --- |
| 254906_at | 25.4 | A | 13.3 | A | 10.6 | A | 14.2 | A | --- |
| 254907_at | 28.1 | P | 30.5 | P | 112.9 | P | 43.7 | P | --- |
| 254908_at | 24.7 | A | 1.4 | A | 26.2 | A | 14.9 | A | --- |
| 254909_at | 8.3 | A | 44 | P | 13.7 | A | 17.6 | A | --- |
| 254910_at | 1342.2 | P | 1419.4 | P | 921.3 | P | 1451.5 | P | --- |
| 254911_at | 491.1 | P | 568.3 | P | 363.6 | P | 529.3 | P | --- |
| 254912_at | 40.9 | P | 42.5 | P | 43.9 | A | 32.7 | A | --- |
| 254913_at | 373.1 | P | 433.3 | P | 367.9 | P | 347.4 | P | --- |
| 254914_at | 75.4 | A | 156.2 | P | 138.6 | P | 126.3 | P | --- |
| 254915_s_at | 753.8 | P | 804.7 | P | 742.6 | P | 811.5 | P | --- |
| 254916_at | 43.3 | P | 12.7 | A | 31.5 | A | 13.9 | A | --- |
| 254917_at | 211.6 | P | 126.5 | P | 139.7 | P | 97.4 | P | --- |
| 254918_at | 1652.7 | P | 1613.6 | P | 1289.5 | P | 1526.6 | P | SGT1B |
| 254919_at | 439.2 | P | 700.6 | P | 460.2 | P | 724.3 | P | RHA1B |
| 254920_at | 1094.9 | P | 1251.9 | P | 1228 | P | 1216.9 | P | BTI2 |
| 254921_at | 128.1 | P | 151.6 | P | 119.1 | P | 97.4 | P | --- |
| 254922_at | 61.2 | P | 67.4 | P | 41.7 | A | 75.9 | P | RHA1A |
| 254923_at | 752.5 | P | 694.7 | P | 1019.8 | P | 699.5 | P | TOPP7 |
| 254924_at | 255 | P | 304.5 | P | 241.7 | P | 281.1 | P | ATMPK5 |

| | | | | | | | | | |
|---|---|---|---|---|---|---|---|---|---|
| 254925_at | 1183.8 | P | 1459.6 | P | 1309.9 | P | 1503.9 | P | --- |
| 254926_at | 235.5 | P | 258.3 | P | 246.2 | P | 267.3 | P | ACS6 |
| 254927_at | 6.4 | A | 19.7 | A | 17.1 | A | 7 | A | --- |
| 254928_at | 699.6 | P | 646.8 | P | 731 | P | 625.8 | P | --- |
| 254929_at | 71.8 | P | 69.2 | P | 79.2 | A | 37.8 | P | --- |
| 254930_at | 86.4 | P | 159.7 | P | 160.1 | P | 159.3 | P | --- |
| 254931_at | 89.2 | P | 164.9 | P | 110 | P | 143.3 | P | --- |
| 254932_at | 561.9 | P | 565.6 | P | 637.6 | P | 596.7 | P | --- |
| 254933_at | 71.3 | P | 103.2 | P | 93.6 | P | 79.3 | P | RDR2 |
| 254934_at | 62.8 | P | 40.8 | P | 54.6 | P | 64 | P | CRF1 |
| 254935_at | 3.5 | A | 11 | A | 9.5 | A | 3.9 | A | --- |
| 254936_at | 359.5 | P | 472.9 | P | 470.7 | P | 424.4 | P | --- |
| 254937_at | 1020.7 | P | 719.7 | P | 856.3 | P | 796.8 | P | --- |
| 254938_at | 365.6 | P | 334.9 | P | 416.6 | P | 340.5 | P | OPT7 |
| 254939_at | 31.7 | P | 63.2 | P | 33.6 | A | 61.9 | P | --- |
| 254940_at | 12.4 | A | 17.5 | A | 44.8 | A | 27.9 | A | --- |
| 254941_s_at | 5.4 | A | 0.2 | A | 3.7 | A | 0.1 | A | --- |
| 254942_at | 2.2 | A | 18.4 | A | 3.8 | A | 15.8 | A | --- |
| 254943_at | 130.9 | P | 156.9 | P | 215.3 | P | 159.3 | P | --- |
| 254944_at | 96.7 | P | 103.4 | P | 117.4 | P | 109.5 | P | --- |
| 254945_at | 143.7 | A | 118.4 | A | 139.2 | P | 129.9 | A | --- |
| 254946_at | 30.3 | A | 21.7 | A | 33.3 | A | 20.2 | A | --- |
| 254947_at | 11.3 | A | 4.4 | A | 4.9 | A | 23.1 | A | --- |
| 254948_at | 38.6 | A | 71.7 | A | 34.8 | A | 75.1 | A | --- |
| 254949_at | 1.7 | A | 0.9 | A | 1.2 | A | 2.1 | A | --- |
| 254950_at | 683.5 | P | 775.1 | P | 562.5 | P | 685.9 | P | --- |
| 254951_at | 1057 | P | 1285.1 | P | 1069.5 | P | 1372.6 | P | --- |
| 254952_at | 566.4 | P | 408.7 | P | 248.3 | P | 405.2 | P | UGE5 |
| 254953_at | 274.7 | P | 351.5 | P | 372.7 | P | 336.9 | P | --- |
| 254954_at | 1.2 | A | 31.6 | A | 36.3 | A | 37 | A | --- |
| 254955_at | 722.9 | P | 741.5 | P | 629.6 | P | 762.5 | P | KELP |
| 254956_at | 2.3 | A | 3.4 | A | 24.5 | A | 14.5 | A | --- |
| 254957_at | 929.4 | P | 1073.6 | P | 980.1 | P | 976.9 | P | --- |
| 254958_at | 7320.4 | P | 6192.1 | P | 7771.2 | P | 7055.3 | P | NDPK3 |
| 254959_at | 193 | P | 281.4 | P | 204.7 | P | 227.9 | P | UGE5 |
| 254960_at | 20.8 | A | 17.7 | A | 20.6 | A | 12.6 | A | --- |
| 254961_at | 14.1 | A | 12.7 | A | 42.1 | A | 13.3 | A | --- |
| 254962_at | 23.5 | A | 29.9 | A | 6.8 | A | 15.9 | A | AtGH9C3 |
| 254963_at | 336.8 | P | 358.4 | P | 399.3 | P | 309.8 | P | MTSSB |
| 254964_at | 133.1 | P | 130.7 | P | 121.9 | P | 189.7 | P | --- |
| 254965_at | 281.7 | P | 300.3 | P | 210 | P | 306.6 | P | --- |
| 254966_at | 329.4 | P | 407.5 | P | 404.9 | P | 360.2 | P | SPA2 |
| 254967_at | 10.5 | A | 2.8 | A | 2.8 | A | 4.3 | A | --- |
| 254968_at | 44.8 | P | 41.3 | P | 52.5 | A | 58.5 | P | --- |
| 254969_at | 663.8 | P | 631.9 | P | 686 | P | 542.6 | P | --- |
| 254970_at | 19585 | P | 18959.5 | P | 16702.5 | P | 18968.1 | P | LHCB5 |
| 254971_at | 282.8 | P | 238.1 | P | 409.4 | P | 254.7 | P | NIP5;1 |
| 254972_at | 7.2 | A | 3.1 | A | 5.1 | A | 7.4 | A | --- |
| 254973_at | 17.7 | A | 16.7 | A | 4.3 | A | 8.5 | A | --- |
| 254974_at | 27.4 | P | 36.8 | A | 41.4 | A | 8.4 | A | --- |
| 254975_at | 19.3 | A | 159.9 | P | 16.7 | A | 119.9 | P | --- |
| 254976_at | 4.2 | A | 3.8 | A | 5.7 | A | 3.9 | A | --- |
| 254977_s_at | 3 | A | 3.3 | A | 11.9 | A | 7 | A | --- |
| 254978_at | 1.5 | A | 16.7 | A | 3.1 | A | 17.4 | A | --- |
| 254979_at | 108.6 | P | 113.1 | P | 45 | A | 86 | P | --- |
| 254980_at | 2177.6 | P | 1881.7 | P | 2567.1 | P | 2054.6 | P | --- |
| 254981_at | 3261.5 | P | 3025.5 | P | 3938.2 | P | 3555.7 | P | --- |
| 254982_at | 185.7 | P | 217.4 | P | 171.4 | P | 184.9 | P | --- |
| 254983_at | 14.6 | A | 13.5 | A | 21.4 | A | 20.3 | A | MEE53 |
| 254984_s_at | 843.6 | P | 854.8 | P | 527.2 | P | 733 | P | UBP10 |
| 254985_x_at | 53.9 | P | 43.6 | P | 37.6 | P | 30.8 | A | --- |
| 254986_at | 39.8 | A | 6.7 | A | 34.7 | A | 6.4 | A | IQD16 |
| 254987_s_at | 2.3 | A | 26.8 | P | 13.7 | A | 2.2 | A | --- |
| 254988_at | 20.3 | A | 35 | P | 32.6 | A | 19.2 | A | GTC2 |
| 254989_at | 18.1 | A | 56.5 | A | 30.6 | A | 36 | A | --- |
| 254990_at | 1599 | P | 1497.4 | P | 1100.3 | P | 1286.3 | P | CID12 |
| 254991_at | 277.5 | P | 264.3 | P | 310.5 | P | 262.6 | P | --- |
| 254992_at | 640.7 | P | 632.8 | P | 745.5 | P | 666.1 | P | SPT16 |
| 254993_at | 134.4 | P | 156.6 | P | 193 | P | 113.9 | P | --- |
| 254994_at | 330.9 | P | 309.8 | P | 332 | P | 372.3 | P | MTA |
| 254995_at | 7.5 | A | 17.5 | A | 39.5 | A | 3.3 | A | --- |
| 254996_at | 212.3 | P | 82.3 | P | 174.9 | P | 106.2 | P | --- |
| 254997_s_at | 91.1 | P | 71.4 | P | 121.7 | P | 90.3 | P | --- |
| 254998_at | 1214.5 | P | 1220 | P | 839.2 | P | 990.2 | P | --- |
| 254999_at | 1492.5 | P | 1286 | P | 1692 | P | 1223.4 | P | --- |
| 255000_at | 6192.2 | P | 5192.5 | P | 6928.8 | P | 5508.2 | P | RPS18C |
| 255001_at | 2.2 | A | 8.2 | A | 20.2 | A | 1.1 | A | --- |
| 255002_at | 2.4 | A | 2.6 | A | 6.1 | A | 3.6 | A | --- |
| 255003_at | 62.1 | A | 50.9 | P | 111 | A | 44.7 | A | --- |
| 255004_at | 181.6 | P | 150.2 | P | 207.3 | P | 210.5 | P | --- |
| 255005_at | 21.1 | A | 38.1 | P | 43.8 | A | 50.1 | A | --- |
| 255006_at | 46.9 | A | 34.1 | A | 33.5 | A | 32.5 | A | --- |
| 255007_at | 6.3 | A | 3.6 | A | 12.9 | A | 2 | A | AtHSD5 |
| 255008_at | 1361.6 | P | 1422.4 | P | 1289.4 | P | 1180.3 | P | --- |
| 255009_at | 206.5 | P | 195.7 | P | 245 | P | 231.5 | P | --- |
| 255010_at | 499 | P | 514.3 | P | 468.6 | P | 558.8 | P | --- |
| 255011_at | 2181.7 | P | 1986.5 | P | 2445.8 | P | 2407.8 | P | CYTC-2 |
| 255012_at | 487 | P | 284.3 | P | 471.8 | P | 335.7 | P | --- |
| 255013_at | 365 | P | 463.2 | P | 351.9 | P | 429.6 | P | --- |
| 255014_at | 3 | A | 24.6 | A | 9.8 | A | 14.2 | A | STK |
| 255015_at | 148.7 | P | 182.5 | P | 139.6 | P | 119.2 | P | EMB1691 |
| 255016_at | 628.8 | P | 1335.7 | P | 1008.7 | P | 1263.8 | P | ATSPS4F |
| 255017_at | 2009.6 | P | 1556.3 | P | 1945.6 | P | 1648.8 | P | CNX7 |
| 255018_at | 271.4 | P | 244.5 | P | 210.4 | P | 261.1 | P | --- |
| 255019_at | 12 | A | 24.4 | A | 29.2 | A | 33.7 | A | --- |
| 255020_at | 2196.1 | P | 1921.9 | P | 2568.8 | P | 2151.1 | P | --- |
| 255021_at | 316.5 | P | 374.6 | P | 296.1 | P | 378 | P | --- |
| 255022_at | 34.9 | A | 24.3 | A | 32.6 | A | 47.3 | A | ANAC070 |
| 255023_at | 1.8 | A | 9.1 | A | 27.2 | A | 15.7 | A | --- |
| 255024_at | 0.6 | A | 1.3 | A | 9.1 | A | 0.9 | A | --- |
| 255025_at | 643.1 | P | 557.1 | P | 532.2 | P | 490.2 | P | MES12 |
| 255026_at | 305.4 | P | 251.7 | P | 218.2 | P | 304.2 | P | MES12 |
| 255027_at | 1.4 | A | 4.3 | A | 39 | A | 13 | A | --- |
| 255028_at | 245.1 | P | 250.1 | P | 197 | P | 245.4 | P | PMP |
| 255029_x_at | 25.3 | A | 29.1 | A | 32.3 | A | 11.9 | A | --- |
| 255030_at | 18.6 | A | 9.5 | A | 5.8 | A | 16.3 | A | --- |
| 255031_at | 121.1 | P | 111.6 | P | 105.1 | P | 123.4 | P | --- |
| 255032_at | 162.1 | P | 186.5 | P | 108.8 | P | 194.2 | P | --- |
| 255033_at | 210.6 | P | 216.9 | P | 219.8 | P | 240.9 | P | --- |

| | | | | | | | | | |
|---|---|---|---|---|---|---|---|---|---|
| 255034_at | 20.8 | A | 11.2 | A | 2.2 | A | 0.7 | A | --- |
| 255035_at | 742.5 | P | 787.3 | P | 804 | P | 851.8 | P | --- |
| 255036_at | 107.7 | P | 103.5 | P | 108.8 | M | 105.8 | P | --- |
| 255037_at | 299 | P | 459.9 | P | 294.5 | P | 398.4 | P | ATMYB6 |
| 255038_at | 573.4 | P | 473.6 | P | 666.7 | P | 546.9 | P | CINV2 |
| 255039_at | 739.6 | P | 785.1 | P | 576.8 | P | 557.1 | P | CPK4 |
| 255040_at | 242 | P | 302.2 | P | 249.2 | P | 294.6 | P | --- |
| 255041_at | 878 | P | 860.4 | P | 790.6 | P | 817.7 | P | --- |
| 255042_at | 234 | P | 257.1 | P | 267.4 | P | 322.3 | P | --- |
| 255043_at | 283.3 | P | 280.4 | P | 275.2 | P | 278.2 | P | --- |
| 255044_at | 199 | P | 147.7 | P | 169.8 | P | 151.1 | P | --- |
| 255045_at | 2.1 | A | 3.3 | A | 3 | A | 4.6 | A | --- |
| 255046_at | 3968.8 | P | 6176.1 | P | 5402.4 | P | 5806.7 | P | ATPD |
| 255047_at | 1065.4 | P | 1257.5 | P | 1189.9 | P | 1313 | P | --- |
| 255048_at | 18 | A | 10.8 | A | 13 | A | 8.7 | A | GASA3 |
| 255049_at | 4.6 | A | 2.3 | A | 36.5 | A | 2.5 | A | GASA2 |
| 255050_at | 7.6 | A | 8 | A | 4.7 | A | 15.6 | A | --- |
| 255051_at | 57.5 | P | 62.7 | P | 137.7 | P | 44.1 | P | --- |
| 255052_at | 1244.6 | P | 1736.4 | P | 1411.4 | P | 1713.2 | P | ATRABG3A |
| 255053_at | 679.5 | P | 753.6 | P | 829.3 | P | 833.6 | P | --- |
| 255054_s_at | 2.8 | A | 23.8 | A | 47.2 | P | 28.1 | A | AtGH9B15 |
| 255055_at | 182.1 | P | 204.7 | P | 163.8 | P | 196.1 | P | --- |
| 255056_at | 112.4 | P | 2.1 | A | 128.7 | P | 59.8 | P | TT8 |
| 255057_at | 58.6 | P | 54.1 | A | 54 | P | 56.9 | P | --- |
| 255058_s_at | 20.4 | A | 5.1 | A | 28.1 | A | 6.6 | A | --- |
| 255059_at | 83.6 | P | 100.2 | P | 127.3 | P | 77.4 | P | --- |
| 255060_at | 1.5 | A | 1.7 | A | 2.2 | A | 1 | A | --- |
| 255061_at | 323 | P | 380.8 | P | 375.1 | P | 326.1 | P | ATAPRL6 |
| 255062_at | 118.6 | P | 121.5 | P | 178.4 | P | 159.7 | P | --- |
| 255063_at | 2.7 | A | 5.3 | A | 5.9 | A | 1.4 | A | --- |
| 255064_at | 2108.3 | P | 794.9 | P | 1861.5 | P | 998.5 | P | EXO |
| 255065_s_at | 5813.6 | P | 2035.7 | P | 4319.9 | P | 2590.5 | P | --- |
| 255066_at | 1369 | P | 1364.5 | P | 1431.3 | P | 1368.6 | P | --- |
| 255067_at | 603.9 | P | 551.4 | P | 626.1 | P | 558.3 | P | --- |
| 255068_at | 616.7 | P | 704.8 | P | 615.2 | P | 626.8 | P | CRY1 |
| 255069_s_at | 47.3 | P | 7.9 | A | 59.9 | P | 35 | A | --- |
| 255070_at | 1781.3 | P | 2158.7 | P | 1715.1 | P | 2566.3 | P | ISA3 |
| 255071_at | 134.7 | P | 259.9 | P | 141.1 | P | 238 | P | --- |
| 255072_at | 46.7 | A | 18.2 | A | 8.4 | A | 28.2 | A | TOC75-IV |
| 255073_at | 1.8 | A | 1.2 | A | 25 | A | 1.8 | A | --- |
| 255074_at | 12.4 | A | 5.3 | A | 19.5 | A | 12.8 | A | --- |
| 255075_at | 6.7 | A | 24.6 | A | 36.6 | A | 10.8 | A | --- |
| 255076_at | 2.4 | A | 15.9 | A | 22.3 | A | 2.7 | A | --- |
| 255077_at | 504.7 | P | 558.9 | P | 447.1 | P | 508.2 | P | --- |
| 255078_at | 3175.8 | P | 4800.8 | P | 3908 | P | 4404.4 | P | APX4 |
| 255079_s_at | 7643.8 | P | 9804.8 | P | 8232 | P | 10777.9 | P | GF14 PHI |
| 255080_at | 1709.8 | P | 910.2 | P | 1411.1 | P | 977.2 | P | AGP10 |
| 255081_at | 178.9 | P | 179.8 | P | 142.7 | A | 157.1 | P | MLH1 |
| 255082_at | 111.5 | P | 180.8 | P | 99.8 | P | 130.7 | P | --- |
| 255083_s_at | 0.3 | A | 1.8 | A | 1.8 | A | 0.6 | A | --- |
| 255084_s_at | 20.2 | A | 6.2 | A | 7.5 | A | 3.6 | A | --- |
| 255085_at | 0.7 | A | 7.3 | A | 17.4 | A | 10 | A | --- |
| 255086_at | 21 | A | 22 | A | 4.9 | A | 11.1 | A | --- |
| 255087_at | 226.4 | P | 309.4 | P | 220.7 | P | 256.9 | P | --- |
| 255088_at | 363.4 | P | 491 | P | 462.7 | P | 555.2 | P | --- |
| 255089_at | 12854.4 | P | 11217.8 | P | 13143 | P | 12225.9 | P | NDPK1 |
| 255090_at | 2.1 | A | 2.2 | A | 16.5 | A | 1.7 | A | --- |
| 255091_at | 18.9 | A | 1.6 | A | 13.7 | A | 25.4 | A | --- |
| 255092_s_at | 1.1 | A | 1.5 | A | 1.6 | A | 12.9 | A | --- |
| 255093_s_at | 514.6 | P | 474.4 | P | 424.8 | P | 449.7 | P | --- |
| 255094_at | 26.2 | A | 70.7 | M | 70.6 | A | 62.9 | M | ORTHL |
| 255095_at | 373.8 | P | 354.8 | P | 260.8 | P | 373.6 | P | MEKK1 |
| 255096_at | 4 | A | 28.3 | A | 14.8 | A | 3 | A | --- |
| 255097_at | 16.6 | A | 12.6 | A | 14.1 | A | 11.3 | A | --- |
| 255098_at | 24.4 | A | 15.5 | A | 7.7 | A | 23.1 | A | --- |
| 255099_at | 2.6 | A | 11 | A | 7.6 | A | 4.8 | A | --- |
| 255100_at | 2.7 | A | 2.5 | A | 1.9 | A | 1.5 | A | --- |
| 255101_at | 8.9 | A | 16.2 | A | 17.2 | A | 1.1 | A | --- |
| 255102_s_at | 17.9 | A | 6.6 | A | 20.1 | A | 1.2 | A | --- |
| 255103_at | 296.3 | P | 299.4 | P | 326.6 | P | 272.1 | P | --- |
| 255104_at | 1445.1 | P | 2141.6 | P | 1675.8 | P | 2443.1 | P | SAH7 |
| 255105_at | 3.7 | A | 9.3 | A | 9.4 | A | 6.3 | A | SULTR1;1 |
| 255106_at | 10 | A | 9.4 | A | 28.1 | A | 0.2 | A | --- |
| 255107_at | 35 | P | 34.6 | M | 59.5 | P | 37.3 | P | --- |
| 255108_at | 8.1 | A | 5 | A | 8.4 | A | 12 | A | --- |
| 255109_s_at | 0.6 | A | 3.5 | A | 17.3 | A | 0.7 | A | --- |
| 255110_at | 79.9 | A | 133 | P | 101.2 | A | 144.4 | P | --- |
| 255111_at | 44.8 | A | 63.2 | P | 49.4 | A | 68.4 | P | --- |
| 255112_at | 9.6 | A | 10.7 | A | 5.7 | A | 4.9 | A | --- |
| 255113_at | 3.6 | A | 2.3 | A | 3.3 | A | 1.6 | A | --- |
| 255114_at | 38.6 | A | 42.4 | A | 59.6 | P | 46.5 | A | --- |
| 255115_at | 43.3 | A | 31.3 | P | 36.1 | A | 15.3 | A | APUM11 |
| 255116_at | 620.3 | P | 1260.7 | P | 542.1 | P | 964.8 | P | --- |
| 255117_s_at | 86.1 | P | 88.5 | P | 91.8 | P | 114.4 | P | ATPUP13 |
| 255118_at | 271.5 | P | 320.7 | P | 358.7 | P | 327.9 | P | --- |
| 255119_at | 3.6 | A | 9.2 | A | 15.6 | P | 2.3 | A | --- |
| 255120_x_at | 3.7 | A | 11.2 | A | 11.2 | A | 38 | A | --- |
| 255121_at | 88.2 | P | 99 | P | 70.6 | P | 88.9 | P | MAPKKK9 |
| 255122_at | 11.6 | M | 8 | A | 4.2 | A | 11.2 | A | --- |
| 255123_at | 17.3 | A | 25.3 | A | 17.2 | A | 38.1 | P | --- |
| 255124_at | 1.7 | A | 1.8 | A | 2.1 | A | 2.3 | A | APUM15 |
| 255125_at | 31 | A | 30.5 | A | 41.8 | A | 33.9 | A | --- |
| 255126_at | 2.4 | A | 22.4 | A | 19.3 | A | 3.8 | A | --- |
| 255127_at | 64.5 | P | 99.9 | P | 48.3 | A | 68.5 | P | --- |
| 255128_at | 399.4 | P | 423.6 | P | 436.3 | P | 374.4 | P | --- |
| 255129_at | 309 | P | 244 | P | 157 | P | 209 | P | --- |
| 255130_at | 240.6 | P | 380.5 | P | 326.9 | P | 360.2 | P | --- |
| 255131_at | 1216.8 | P | 1392.2 | P | 1168.7 | P | 1281.6 | P | --- |
| 255132_at | 427.7 | P | 412.9 | P | 446.3 | P | 396.2 | P | --- |
| 255133_at | 296.1 | P | 303.4 | P | 267.5 | P | 318.1 | P | --- |
| 255134_at | 192.6 | P | 200 | P | 235.6 | P | 230.2 | P | --- |
| 255135_at | 22.3 | A | 32.2 | A | 13 | A | 21 | A | --- |
| 255136_s_at | 404.8 | P | 386 | P | 404.7 | P | 438.3 | P | --- |
| 255137_at | 23.8 | A | 1.6 | A | 13.5 | A | 1 | A | --- |
| 255138_at | 3.4 | A | 5.7 | A | 6.1 | A | 10.4 | A | --- |
| 255139_s_at | 41.3 | A | 32.5 | A | 55.7 | A | 12.8 | A | --- |
| 255140_x_at | 70.9 | A | 89.3 | A | 104.6 | A | 72 | A | --- |
| 255141_at | 14 | A | 5.9 | A | 18.7 | A | 10.9 | A | --- |
| 255142_at | 4527.2 | P | 2262.1 | P | 4774.2 | P | 3339.1 | P | SAPX |

| | | | | | | | | | |
|---|---|---|---|---|---|---|---|---|---|
| 255143_at | 8.6 | A | 13.6 | A | 3.8 | A | 1.5 | A | --- |
| 255144_at | 0.3 | A | 3.8 | A | 1 | A | 1.5 | A | --- |
| 255145_at | 11.6 | A | 2.5 | A | 4.8 | A | 3.4 | A | --- |
| 255146_at | 1.7 | A | 0.4 | A | 5 | A | 2 | A | --- |
| 255147_at | 566.4 | P | 489.9 | P | 633.8 | P | 509.8 | P | --- |
| 255148_at | 486.7 | P | 829.7 | P | 384.5 | P | 582.7 | P | MAPKKK10 |
| 255149_at | 92.1 | P | 308.8 | P | 275.7 | P | 345 | P | KNAT1 |
| 255150_at | 41.3 | P | 63 | P | 78 | P | 88.2 | P | --- |
| 255151_at | 312.2 | P | 321.9 | P | 249.9 | P | 295.6 | P | ORP1C |
| 255152_at | 8.8 | A | 2.1 | A | 0.5 | A | 2.3 | A | --- |
| 255153_at | 78.8 | P | 53.1 | P | 79.9 | M | 69.7 | P | --- |
| 255154_at | 383 | P | 387.7 | P | 324.2 | P | 351 | P | --- |
| 255155_at | 87.5 | P | 124.5 | P | 153.5 | P | 129.1 | P | --- |
| 255156_at | 0.8 | A | 0.3 | A | 8.7 | A | 0.3 | A | --- |
| 255157_at | 21.7 | A | 9.9 | A | 15 | A | 26.4 | A | --- |
| 255158_s_at | 1.3 | A | 0.3 | A | 8.8 | A | 1.1 | A | --- |
| 255159_at | 27.6 | A | 22.4 | A | 39 | A | 25.7 | A | --- |
| 255160_at | 43 | A | 69.9 | A | 91 | A | 55.9 | A | --- |
| 255161_at | 7.3 | A | 13.5 | A | 13.7 | A | 20.5 | P | --- |
| 255162_at | 0.4 | A | 15.6 | A | 24.3 | A | 1.9 | A | --- |
| 255163_at | 6.4 | A | 1.1 | A | 11.1 | A | 10.7 | A | --- |
| 255164_at | 13 | A | 6.4 | A | 25.3 | A | 16.3 | A | --- |
| 255165_at | 2.4 | A | 11.7 | A | 1.5 | A | 1.9 | A | --- |
| 255166_at | 0.9 | A | 1.3 | A | 3.9 | A | 1 | A | --- |
| 255167_at | 22.1 | A | 36.2 | P | 31.9 | A | 22.9 | A | --- |
| 255168_at | 2.1 | A | 3 | A | 2.3 | A | 3.9 | A | --- |
| 255169_x_at | 28.2 | A | 24.6 | A | 56.4 | M | 16.9 | A | --- |
| 255170_at | 0.4 | A | 2.9 | A | 1.6 | A | 8.3 | A | --- |
| 255171_at | 304.4 | P | 296.5 | P | 269.5 | P | 311.5 | P | --- |
| 255172_x_at | 14.5 | A | 5.7 | A | 6 | A | 1.3 | A | --- |
| 255173_at | 6.1 | A | 5.5 | A | 40.5 | A | 12.4 | A | --- |
| 255174_at | 8.3 | A | 1.7 | A | 21.9 | A | 1.6 | A | --- |
| 255175_at | 49.8 | P | 41.6 | P | 54.3 | P | 50.8 | P | ATCSLC12 |
| 255176_s_at | 1580.5 | P | 1331.7 | P | 1356.2 | P | 1468.7 | P | --- |
| 255177_at | 7.2 | A | 45 | A | 6.4 | A | 30.3 | A | ACS11 |
| 255178_at | 0.5 | A | 0.6 | A | 1.1 | A | 0.6 | A | --- |
| 255179_at | 3 | A | 2.9 | A | 34.8 | A | 14.4 | A | --- |
| 255180_at | 0.9 | A | 0.2 | A | 0.3 | A | 0.8 | A | --- |
| 255181_at | 136.5 | P | 123.9 | P | 93.7 | P | 158.2 | P | --- |
| 255182_at | 2.3 | A | 1.5 | A | 4.3 | A | 1.9 | A | --- |
| 255183_at | 4.5 | A | 35.7 | A | 29.4 | A | 17.2 | A | --- |
| 255184_at | 18 | P | 6.5 | A | 22.3 | A | 9.5 | A | --- |
| 255185_at | 18.3 | A | 28.3 | A | 32 | A | 32.7 | A | --- |
| 255186_at | 14.3 | A | 23.6 | A | 6.2 | A | 4.5 | A | --- |
| 255187_s_at | 0.6 | A | 1.4 | A | 2.3 | A | 1 | A | --- |
| 255188_at | 3.6 | A | 6.4 | A | 1.5 | A | 2.1 | A | --- |
| 255189_at | 2.6 | A | 1.7 | A | 0.6 | A | 2.4 | A | --- |
| 255190_x_at | 27.6 | M | 15.4 | A | 26.8 | A | 14.1 | A | --- |
| 255191_at | 6.2 | A | 14.1 | A | 22.6 | A | 4.2 | A | --- |
| 255192_at | 22 | A | 19.9 | A | 25.7 | A | 10 | A | --- |
| 255193_at | 76.7 | P | 67.4 | P | 99.2 | P | 77.2 | P | VFB3 |
| 255194_at | 407 | P | 326.2 | P | 467.7 | P | 365.2 | P | --- |
| 255195_at | 18.1 | A | 2.1 | A | 2.7 | A | 3 | A | --- |
| 255196_x_at | 23.5 | A | 18.4 | A | 8.5 | A | 20 | A | --- |
| 255197_x_at | 37.4 | P | 43.4 | A | 70.6 | P | 12.3 | A | --- |
| 255198_at | 5.8 | A | 1.1 | A | 1.2 | A | 9.9 | A | --- |
| 255199_at | 874.9 | P | 760 | P | 831.9 | P | 833 | P | --- |
| 255200_at | 17 | A | 27.9 | A | 32.8 | A | 11.5 | A | --- |
| 255201_at | 0.7 | A | 13.6 | A | 0.6 | A | 0.9 | A | --- |
| 255202_at | 19.5 | A | 19.7 | A | 26.6 | A | 38.7 | A | --- |
| 255203_at | 9.9 | A | 20.6 | A | 36.5 | A | 24.9 | A | --- |
| 255204_at | 3.6 | A | 2.4 | A | 3.1 | A | 3 | A | --- |
| 255205_at | 1.5 | A | 6.3 | A | 12.7 | A | 8.3 | A | --- |
| 255206_at | 0.4 | A | 0.8 | A | 1.9 | A | 7.5 | A | --- |
| 255207_at | 2.2 | A | 2.5 | A | 1.6 | A | 1.1 | A | --- |
| 255208_at | 2 | A | 1 | A | 8.3 | A | 4.4 | A | --- |
| 255209_at | 12.3 | A | 14.8 | A | 22.8 | A | 7.3 | A | --- |
| 255210_at | 7.4 | A | 4.6 | A | 5.4 | A | 23.8 | A | --- |
| 255211_at | 0.8 | A | 1 | A | 0.8 | A | 7.9 | A | --- |
| 255212_at | 0.9 | A | 1.8 | A | 2 | A | 20.1 | A | --- |
| 255213_at | 5.7 | A | 0.4 | A | 2.2 | A | 0.7 | A | --- |
| 255214_at | 19.3 | A | 16.6 | A | 18.6 | A | 1.1 | A | --- |
| 255215_at | 0.9 | A | 1.8 | A | 1.2 | A | 13.6 | A | --- |
| 255216_s_at | 67 | A | 80.1 | P | 23.8 | A | 92.4 | P | --- |
| 255217_s_at | 20.4 | A | 28.7 | P | 32.3 | A | 30.4 | P | --- |
| 255218_at | 0.8 | A | 0.4 | A | 0.3 | A | 1.7 | A | --- |
| 255219_at | 1.7 | A | 8.2 | A | 45.9 | A | 26.1 | A | --- |
| 255220_at | 9317 | P | 10786.7 | P | 11961.5 | P | 10980 | P | UBQ10 |
| 255221_at | 1533.5 | P | 1980.9 | P | 1609.2 | P | 1833.4 | P | --- |
| 255222_at | 5.4 | A | 6.2 | A | 4.2 | A | 0.6 | A | --- |
| 255223_at | 107.5 | P | 1.5 | A | 95.3 | P | 20.7 | A | --- |
| 255224_at | 340.9 | P | 363.8 | P | 427 | P | 350.4 | P | --- |
| 255225_at | 505.2 | P | 399.9 | P | 535.1 | P | 439.5 | P | --- |
| 255226_at | 797.4 | P | 761.9 | P | 687 | P | 802 | P | DDB1A |
| 255227_at | 354.3 | P | 329.8 | P | 385.3 | P | 304.6 | P | EDA35 |
| 255228_at | 15.9 | A | 16.2 | A | 35 | A | 21.4 | A | --- |
| 255229_at | 17.2 | A | 13.2 | A | 3.8 | A | 12.8 | A | --- |
| 255230_at | 1108.3 | P | 1070.8 | P | 1087.6 | P | 902.2 | P | ATRFNR1 |
| 255231_at | 369.4 | P | 443.9 | P | 368.5 | P | 449.4 | P | --- |
| 255232_at | 212.4 | P | 247.6 | P | 274.4 | P | 225.1 | P | AGD13 |
| 255233_at | 358.5 | P | 391.5 | P | 440.9 | P | 438.1 | P | MFDX1 |
| 255234_at | 1.7 | A | 4.3 | A | 6.6 | A | 2.8 | A | --- |
| 255235_at | 14.9 | A | 6.7 | A | 34.8 | A | 2.3 | A | --- |
| 255236_at | 216.4 | P | 432.8 | P | 180 | P | 410.3 | P | ATEHD2 |
| 255237_at | 1.9 | A | 10.8 | A | 16.2 | A | 0.9 | A | --- |
| 255238_at | 11.8 | A | 1.5 | A | 2.8 | A | 0.7 | A | --- |
| 255239_at | 7.8 | A | 5.2 | A | 16.8 | A | 0.4 | A | --- |
| 255240_at | 988.8 | P | 1259 | P | 1162.8 | P | 1180 | P | IBR1 |
| 255241_at | 20 | A | 18.7 | A | 2.6 | A | 0.4 | A | --- |
| 255242_at | 6.4 | A | 12.5 | A | 6.9 | A | 0.7 | A | --- |
| 255243_at | 309.2 | P | 406.1 | P | 308.3 | P | 329 | P | --- |
| 255244_at | 48.8 | P | 16.6 | A | 45.2 | A | 33.3 | A | --- |
| 255245_at | 21 | A | 10.1 | A | 6 | A | 3 | A | --- |
| 255246_at | 2.1 | A | 16 | A | 51.7 | A | 13.7 | A | --- |
| 255247_at | 0.6 | A | 3.5 | A | 1.7 | A | 14 | A | --- |
| 255248_at | 10181.3 | P | 10243.2 | P | 8980.8 | P | 9687.7 | P | PSBQ-2 |
| 255249_at | 231.8 | P | 286 | P | 250.6 | P | 263.9 | P | --- |
| 255250_at | 32.2 | P | 19.5 | A | 34 | A | 30 | A | AtMYB74 |
| 255251_at | 2.8 | A | 2.8 | A | 10.8 | A | 4.8 | A | --- |

| | | | | | | | | | |
|---|---|---|---|---|---|---|---|---|---|
| 255252_at | 2.5 | A | 2.3 | A | 13 | A | 1.5 | A | --- |
| 255253_at | 375 | P | 438.4 | P | 330.6 | P | 390.3 | P | VPS28-2 |
| 255254_at | 1.4 | A | 31.6 | A | 14.5 | A | 5.2 | A | --- |
| 255255_at | 419.9 | P | 567.9 | P | 373.7 | P | 379.6 | P | --- |
| 255256_at | 6.2 | A | 1.2 | A | 18 | A | 1.9 | A | --- |
| 255257_at | 6727.2 | P | 7579.3 | P | 6531.2 | P | 8064.7 | P | UBQ11 |
| 255258_at | 260.8 | P | 353.6 | P | 306.1 | P | 274.5 | P | --- |
| 255259_at | 1385.5 | P | 755.7 | P | 1009.3 | P | 769.5 | P | NDB2 |
| 255260_at | 107.5 | P | 146.1 | P | 184.4 | P | 138.4 | P | --- |
| 255261_s_at | 93.1 | P | 137 | P | 103.4 | P | 134.3 | P | FUR1 |
| 255262_at | 2.5 | A | 6.2 | A | 21.6 | A | 2.7 | A | --- |
| 255263_at | 1862 | P | 1559.9 | P | 1597.5 | P | 1521.2 | P | --- |
| 255264_at | 21.7 | A | 31.7 | P | 52.9 | P | 28.3 | A | --- |
| 255265_at | 106.6 | P | 263.7 | P | 170 | P | 223.9 | P | ATK5 |
| 255266_at | 4.8 | A | 34.3 | A | 14.2 | A | 14.8 | A | --- |
| 255267_at | 75.2 | P | 57.9 | P | 57 | P | 64.5 | P | --- |
| 255268_at | 16.9 | A | 27.3 | M | 6 | A | 36.2 | A | --- |
| 255269_at | 1.1 | A | 0.8 | A | 5 | A | 1.7 | A | --- |
| 255270_at | 5.3 | A | 5.3 | A | 3.7 | A | 32.2 | A | --- |
| 255271_at | 45.8 | P | 35.4 | P | 58.3 | A | 43.4 | A | --- |
| 255272_at | 20.8 | A | 12.9 | A | 17.5 | A | 8 | A | --- |
| 255273_at | 2.4 | A | 12.4 | A | 21.3 | A | 1.4 | A | --- |
| 255274_at | 5.2 | A | 15.4 | A | 22.5 | A | 5.4 | A | --- |
| 255275_at | 16.6 | A | 10.5 | A | 21.1 | A | 1.9 | A | --- |
| 255276_at | 28.8 | P | 15.1 | A | 2.5 | A | 24 | A | DES-1-LIKE |
| 255277_at | 809.6 | P | 1034.7 | P | 803.8 | P | 1062.3 | P | PDF2 |
| 255278_at | 267.9 | P | 218.5 | P | 359.8 | P | 214.4 | P | --- |
| 255279_at | 505 | P | 557.8 | P | 620.2 | P | 519.7 | P | --- |
| 255280_at | 171.1 | P | 142.9 | P | 137.7 | P | 99.7 | P | --- |
| 255281_at | 161.8 | P | 247.6 | P | 157.3 | P | 180.6 | P | GSL1 |
| 255282_at | 23.7 | A | 16.2 | M | 18.8 | A | 39 | P | --- |
| 255283_at | 691.5 | P | 490.8 | P | 646.3 | P | 439.1 | P | ATG8B |
| 255284_at | 2511.4 | P | 1741.9 | P | 2697.6 | P | 1608.1 | P | 41365 |
| 255285_at | 176.2 | P | 126 | P | 189.9 | P | 117.5 | P | --- |
| 255286_at | 25.3 | A | 9.6 | A | 7.1 | A | 4.5 | A | --- |
| 255287_at | 12.8 | A | 32.2 | A | 40.6 | A | 23.6 | A | --- |
| 255288_at | 144 | P | 172.1 | P | 155 | P | 114.2 | P | --- |
| 255289_at | 36.1 | A | 33.9 | A | 16.5 | A | 17.7 | A | --- |
| 255290_at | 12293.4 | P | 12395.4 | P | 12358.2 | P | 12373 | P | ATPC1 |
| 255291_at | 29.4 | P | 65.4 | P | 83.2 | P | 62.6 | P | CPK27 |
| 255292_s_at | 470.1 | P | 593.5 | P | 534.5 | P | 530 | P | CPK22 |
| 255293_at | 17.1 | A | 16 | A | 25.4 | A | 9.5 | A | --- |
| 255294_at | 457.5 | P | 320.5 | P | 497.9 | P | 377.1 | P | --- |
| 255295_at | 48.9 | A | 63.3 | A | 86.5 | A | 86.2 | A | --- |
| 255296_at | 128.7 | P | 192.5 | P | 105.3 | P | 111 | P | --- |
| 255297_x_at | 5.2 | A | 3.7 | A | 7.5 | A | 4.4 | A | --- |
| 255298_at | 670.8 | P | 443.7 | P | 1309.7 | P | 688 | P | ATMSRB6 |
| 255299_at | 451 | P | 466.7 | P | 542.9 | P | 455 | P | --- |
| 255300_at | 615.6 | P | 448.6 | P | 766 | P | 602.6 | P | CLS |
| 255301_at | 367.5 | P | 477.5 | P | 349.8 | P | 404.3 | P | ATMSRB3 |
| 255302_at | 2228.9 | P | 1649.2 | P | 3801.2 | P | 1550.4 | P | ATMSRB5 |
| 255303_at | 502.5 | P | 533 | P | 583.9 | P | 531.3 | P | DER2.2 |
| 255304_at | 436.4 | P | 675.1 | P | 583.3 | P | 690.4 | P | KEA3 |
| 255305_at | 2625.6 | P | 2636.5 | P | 2273.7 | P | 2737.9 | P | ATABC1 |
| 255306_at | 52.6 | P | 64.9 | P | 51.5 | P | 88.7 | P | CPK23 |
| 255307_at | 18.5 | A | 35 | P | 42.7 | P | 16.7 | M | RIC10 |
| 255308_at | 785.9 | P | 1141.8 | P | 955 | P | 1189.9 | P | NSF |
| 255309_at | 218.1 | P | 256.9 | P | 176.5 | P | 155.8 | P | SFR6 |
| 255310_at | 171 | P | 514.6 | P | 187.4 | P | 476.4 | P | ATALN |
| 255311_at | 3.7 | A | 27.4 | A | 37.1 | A | 23.5 | A | --- |
| 255312_at | 1.3 | A | 0.6 | A | 0.4 | A | 0.9 | A | --- |
| 255313_at | 3.2 | A | 32.2 | P | 32.5 | A | 3.5 | A | --- |
| 255314_at | 0.9 | A | 2.1 | A | 17.3 | A | 14.2 | A | --- |
| 255315_at | 17.8 | A | 2.5 | A | 6.5 | A | 9.9 | A | --- |
| 255316_at | 32.4 | A | 26.2 | A | 19.8 | A | 14.1 | A | --- |
| 255317_at | 108.2 | A | 89.6 | A | 134.9 | A | 111.7 | M | --- |
| 255318_at | 177.4 | P | 227.6 | P | 192.7 | P | 190.6 | P | --- |
| 255319_at | 67.6 | P | 158.5 | P | 76.1 | P | 104.1 | P | AtRLP46 |
| 255320_at | 16 | A | 1.5 | A | 27.5 | A | 13.8 | A | --- |
| 255321_at | 13.9 | A | 33.7 | A | 4.3 | A | 46.2 | A | --- |
| 255322_at | 0.7 | A | 1.8 | A | 1.6 | A | 0.9 | A | --- |
| 255323_at | 19.6 | A | 18 | P | 13.8 | A | 17.2 | A | --- |
| 255324_s_at | 21.2 | P | 26.9 | P | 30.1 | P | 26.8 | A | --- |
| 255325_at | 1162.2 | P | 1345.1 | P | 1010.8 | P | 1204.4 | P | PUX4 |
| 255326_at | 0.3 | A | 4.2 | A | 0.6 | A | 7.8 | A | --- |
| 255327_at | 885.4 | P | 981.1 | P | 897.4 | P | 984.9 | P | --- |
| 255328_at | 527.2 | P | 600.6 | P | 548.7 | P | 631.9 | P | EMB2369 |
| 255329_at | 69.3 | P | 84.1 | P | 59.1 | P | 102.5 | P | --- |
| 255330_s_at | 37.3 | A | 2.8 | A | 3.2 | A | 1.4 | A | --- |
| 255331_at | 899.6 | P | 1121.9 | P | 1200.2 | P | 1059.7 | P | --- |
| 255332_at | 1014.4 | P | 1301.7 | P | 989.5 | P | 1309.1 | P | --- |
| 255333_at | 10.3 | A | 8.3 | A | 2.1 | A | 5.4 | A | --- |
| 255334_at | 0.3 | A | 0.3 | A | 0.5 | A | 3.3 | A | --- |
| 255335_at | 0.3 | A | 1 | A | 0.1 | A | 0.3 | A | --- |
| 255336_at | 1.6 | A | 8.4 | A | 5.5 | A | 1.2 | A | --- |
| 255337_at | 30.7 | A | 16.5 | A | 44.5 | A | 4.8 | A | WRKY42 |
| 255338_at | 1076.3 | P | 1148.5 | P | 1352.3 | P | 1124.6 | P | PMP22 |
| 255339_at | 14.8 | A | 39.3 | A | 65.1 | A | 20.3 | A | --- |
| 255340_at | 83.9 | P | 147.5 | P | 38.1 | P | 93.3 | P | --- |
| 255341_at | 69.7 | P | 209.2 | P | 47.4 | P | 114.2 | P | --- |
| 255342_at | 9.8 | A | 2.6 | A | 20.2 | A | 6.8 | A | --- |
| 255343_at | 6.5 | A | 1 | A | 4.5 | A | 1.8 | A | --- |
| 255344_s_at | 807.2 | P | 661.4 | P | 698.1 | P | 549.1 | P | --- |
| 255345_at | 76.5 | P | 102.5 | P | 93.5 | P | 119.2 | P | --- |
| 255346_at | 11.8 | A | 12.9 | A | 1.5 | A | 4.6 | A | --- |
| 255347_at | 0.9 | A | 0.2 | A | 0.2 | A | 0.6 | A | --- |
| 255348_at | 287.9 | P | 269.7 | P | 225.2 | P | 302.8 | P | --- |
| 255349_at | 5 | A | 4.9 | A | 13.2 | A | 27.1 | A | --- |
| 255350_at | 6.3 | A | 3.7 | A | 2.8 | A | 1.7 | A | --- |
| 255351_x_at | 14.1 | A | 10.8 | A | 38.9 | A | 4.4 | A | --- |
| 255352_at | 2.2 | A | 4.7 | A | 31.9 | A | 3.2 | A | --- |
| 255353_at | 25.5 | A | 31.8 | A | 34.2 | A | 39.4 | A | --- |
| 255354_at | 11.4 | A | 0.9 | A | 12.2 | A | 15.9 | A | --- |
| 255355_at | 1.7 | A | 11.2 | A | 15.8 | A | 9.3 | A | --- |
| 255356_x_at | 15.5 | A | 30.6 | A | 1.9 | A | 29.2 | A | --- |
| 255357_at | 2.8 | A | 1.4 | A | 2.9 | A | 1.5 | A | --- |
| 255358_at | 23 | A | 32.1 | A | 4.6 | A | 3.7 | A | --- |
| 255359_at | 37.2 | P | 40.9 | P | 38.7 | P | 30 | P | --- |
| 255360_at | 356.4 | P | 424.4 | P | 365.5 | P | 472 | P | --- |

| | | | | | | | | | |
|---|---|---|---|---|---|---|---|---|---|
| 255361_at | 0.5 | A | 0.7 | A | 1.2 | A | 5.2 | A | --- |
| 255362_at | 18.3 | A | 15.2 | A | 7.6 | A | 22.7 | A | --- |
| 255363_x_at | 43.9 | A | 52.5 | A | 69.8 | A | 56 | A | --- |
| 255364_s_at | 4058.1 | P | 4087.6 | P | 3637.7 | P | 4823.6 | P | --- |
| 255365_at | 1074.6 | P | 1670.8 | P | 1229.5 | P | 1393.4 | P | MEE51 |
| 255366_x_at | 29.5 | A | 43.5 | P | 52.6 | P | 46.7 | P | --- |
| 255367_at | 2.1 | A | 6.2 | A | 3 | A | 1.8 | A | --- |
| 255368_at | 0.8 | A | 6.2 | A | 1.8 | A | 2.2 | A | --- |
| 255369_at | 3 | A | 22.5 | A | 5.1 | A | 2.6 | A | ISU3 |
| 255370_at | 3.4 | A | 2.9 | A | 3.1 | A | 3.1 | A | --- |
| 255371_at | 13.9 | A | 17.1 | A | 30.3 | A | 15.8 | M | --- |
| 255372_at | 13 | A | 9.2 | A | 64.2 | A | 69.8 | A | --- |
| 255373_s_at | 24.8 | A | 19.8 | A | 23.1 | A | 52.6 | P | --- |
| 255374_at | 0.6 | A | 1.5 | A | 1.6 | A | 5.6 | A | --- |
| 255375_at | 0.3 | A | 0.3 | A | 1.3 | A | 0.8 | A | --- |
| 255376_x_at | 25.1 | A | 22.8 | P | 14.5 | A | 23.6 | P | --- |
| 255377_at | 38.8 | A | 13.1 | A | 50.2 | A | 41.7 | P | --- |
| 255378_at | 550.8 | P | 563.9 | P | 473.2 | P | 549.9 | P | ATGSL05 |
| 255379_at | 8060.5 | P | 8370.3 | P | 8355.6 | P | 8447.9 | P | ATHM2 |
| 255380_at | 1558.7 | P | 1550.5 | P | 1709.8 | P | 1426.3 | P | ATTPC1 |
| 255381_at | 321.9 | P | 317.1 | P | 269.8 | P | 244 | P | RMA1 |
| 255382_at | 640.5 | P | 519.9 | P | 434.1 | P | 445.5 | P | EMB2770 |
| 255383_at | 0.7 | A | 8.8 | A | 5.3 | A | 2.9 | A | --- |
| 255384_at | 10.2 | A | 15.7 | A | 1.3 | A | 18.7 | A | --- |
| 255385_at | 45.2 | P | 84.2 | P | 84.2 | P | 53.2 | P | --- |
| 255386_at | 0.8 | A | 2.1 | A | 1.1 | A | 2.3 | A | --- |
| 255387_at | 2.3 | A | 6 | A | 5.2 | A | 18 | A | --- |
| 255388_at | 10.9 | A | 11.2 | A | 13.9 | A | 30.7 | A | --- |
| 255389_at | 2.5 | A | 2.8 | A | 23.6 | A | 0.8 | A | --- |
| 255390_at | 1.3 | A | 14.2 | A | 15.9 | A | 12.3 | A | --- |
| 255391_at | 19.1 | A | 2.4 | A | 1 | A | 2.4 | A | --- |
| 255392_at | 0.7 | A | 1.2 | A | 3.7 | A | 0.5 | A | --- |
| 255393_at | 2.1 | A | 1.4 | A | 2.4 | A | 0.9 | A | --- |
| 255394_at | 15.5 | A | 10.4 | A | 60.8 | P | 2.4 | A | --- |
| 255395_at | 5.8 | A | 3.8 | A | 1.1 | A | 5 | A | --- |
| 255396_at | 16.8 | P | 20.9 | P | 18.8 | A | 18.9 | A | --- |
| 255397_at | 18.7 | A | 19.9 | A | 9.4 | A | 16.1 | A | --- |
| 255398_at | 4.9 | A | 2 | A | 18.4 | A | 2.3 | A | --- |
| 255399_at | 5 | A | 28.3 | A | 19.4 | A | 18.2 | A | --- |
| 255400_at | 22.8 | A | 7.1 | A | 2.7 | A | 20 | A | --- |
| 255401_at | 37.5 | A | 57.1 | A | 42.2 | A | 59.5 | A | --- |
| 255402_at | 164.6 | A | 185 | M | 144.9 | A | 182.5 | A | hemf2 |
| 255403_at | 386 | P | 570.3 | P | 425.3 | P | 504.1 | P | DFL2 |
| 255404_at | 17.7 | A | 3.3 | A | 11.7 | A | 3.4 | A | --- |
| 255405_s_at | 4.9 | A | 19.7 | P | 25.9 | A | 3.5 | A | --- |
| 255406_at | 47.4 | A | 163.3 | P | 67.1 | A | 105.1 | P | --- |
| 255407_at | 5.1 | A | 1.4 | A | 11.5 | A | 6.9 | A | --- |
| 255408_at | 18.5 | A | 2 | A | 6 | A | 2.5 | A | --- |
| 255409_at | 71.9 | P | 57.3 | P | 38.4 | P | 65.3 | P | --- |
| 255410_at | 99.9 | A | 160.6 | P | 82 | A | 137.3 | P | --- |
| 255411_at | 184.6 | P | 333.3 | P | 273.1 | P | 279.2 | P | --- |
| 255412_at | 470.4 | P | 607.9 | P | 612.9 | P | 671 | P | ABP1 |
| 255413_at | 98.6 | P | 129.9 | P | 115.6 | P | 95.4 | P | --- |
| 255414_at | 8.3 | A | 1.2 | A | 0.7 | A | 0.6 | A | --- |
| 255415_at | 9.2 | A | 0.9 | A | 1.1 | A | 12.2 | A | --- |
| 255416_at | 1.8 | A | 1.1 | A | 1.1 | A | 1.7 | A | --- |
| 255417_at | 113.8 | P | 330 | P | 168 | P | 237.6 | P | GRH1 |
| 255418_at | 676.1 | P | 878.8 | P | 635.4 | P | 829.9 | P | --- |
| 255419_at | 37.3 | A | 68.4 | A | 35.5 | A | 34.6 | A | --- |
| 255420_at | 248.6 | P | 297.9 | P | 326.7 | P | 349.9 | P | FH |
| 255421_at | 1471.7 | P | 1372.6 | P | 1793.4 | P | 1399.2 | P | --- |
| 255422_at | 13.1 | A | 30.1 | A | 30.6 | A | 71.1 | P | CYCD6;1 |
| 255423_at | 9.2 | A | 4.3 | A | 11.5 | A | 0.2 | A | --- |
| 255424_at | 24.5 | A | 31.6 | A | 51.8 | A | 51.6 | P | --- |
| 255425_s_at | 0.5 | A | 2.8 | A | 2.3 | A | 2 | A | --- |
| 255426_at | 19.1 | A | 5.1 | A | 31.8 | A | 36.4 | A | --- |
| 255427_at | 20 | A | 10.1 | A | 2.8 | A | 17 | A | --- |
| 255428_at | 438.9 | P | 586.4 | P | 423.4 | P | 511.5 | P | SRF3 |
| 255429_at | 540.5 | P | 745.4 | P | 627.1 | P | 630.9 | P | --- |
| 255430_at | 617.3 | P | 236.5 | P | 442.6 | P | 185.6 | P | tic20-IV |
| 255431_at | 452.4 | P | 537 | P | 619.2 | P | 549.3 | P | --- |
| 255432_at | 21.8 | A | 13.8 | A | 24.8 | A | 14.9 | A | SYP123 |
| 255433_at | 2067 | P | 3186.3 | P | 3600.3 | P | 3405.6 | P | XTH9 |
| 255434_at | 435.2 | P | 358.5 | P | 425.5 | P | 368.5 | P | --- |
| 255435_at | 15180.5 | P | 15746.7 | P | 15039.2 | P | 15521.2 | P | PETC |
| 255436_at | 615.5 | P | 713.1 | P | 662.6 | P | 557.3 | P | --- |
| 255437_at | 1019 | P | 642.3 | P | 1501.3 | P | 599.7 | P | AOP2 |
| 255438_at | 217.3 | P | 251.9 | P | 228.5 | P | 339 | P | AOP1 |
| 255439_at | 442.8 | P | 545.3 | P | 402.8 | P | 490.1 | P | BSL1 |
| 255440_at | 4033 | P | 3837.8 | P | 2895.2 | P | 4068.1 | P | --- |
| 255441_at | 1541 | P | 1717.9 | P | 1727.2 | P | 1605.5 | P | ATCUL1 |
| 255442_at | 3302.7 | P | 4041.9 | P | 4398.8 | P | 4353.7 | P | --- |
| 255443_at | 167.3 | P | 104.6 | P | 259.6 | P | 141.3 | P | SULTR3;2 |
| 255444_at | 429.8 | P | 416.5 | P | 447.2 | P | 355.1 | P | LD |
| 255445_at | 3.2 | A | 16 | A | 3.3 | A | 1.6 | A | --- |
| 255446_at | 96.5 | P | 65.6 | P | 111.6 | P | 99.9 | P | --- |
| 255447_at | 1215.2 | P | 1510.3 | P | 1576.4 | P | 1407.4 | P | --- |
| 255448_at | 33.7 | P | 29.6 | P | 77.5 | P | 36.9 | P | --- |
| 255449_at | 205 | P | 156.7 | P | 191.4 | P | 142.5 | P | --- |
| 255450_at | 186.9 | P | 163.5 | P | 239.8 | P | 258.1 | P | --- |
| 255451_at | 1799.4 | P | 1616.9 | P | 1777.8 | P | 1617.3 | P | --- |
| 255452_at | 273.4 | P | 301.9 | P | 219.9 | P | 245.9 | P | --- |
| 255453_at | 49.9 | A | 46.2 | P | 79.6 | A | 54.5 | M | --- |
| 255454_at | 3.4 | A | 2.9 | A | 15.1 | A | 4.2 | A | --- |
| 255455_at | 1913 | P | 1707 | P | 2560 | P | 1947.1 | P | --- |
| 255456_at | 2057.8 | P | 2611.3 | P | 1831.8 | P | 2482.3 | P | --- |
| 255457_at | 7368.6 | P | 8396 | P | 7394.9 | P | 7844.8 | P | PSAD-1 |
| 255458_at | 860.3 | P | 1017.2 | P | 777.9 | P | 972.1 | P | --- |
| 255459_at | 7833.5 | P | 7439.6 | P | 7138.5 | P | 6049.1 | P | UBQ14 |
| 255460_at | 119.2 | P | 288.4 | P | 167.2 | P | 269.3 | P | --- |
| 255461_at | 19.3 | A | 43.9 | A | 51.3 | A | 10.1 | A | GA1 |
| 255462_at | 1513 | P | 921 | P | 1295.4 | P | 1197.9 | P | --- |
| 255463_at | 3.3 | A | 1.2 | A | 6.1 | A | 2.2 | A | --- |
| 255464_s_at | 11.7 | A | 33.9 | A | 33 | A | 7.2 | A | AT7SL-1 |
| 255465_at | 686.6 | P | 809.8 | P | 855.4 | P | 823.5 | P | --- |
| 255466_at | 104.1 | P | 112.5 | P | 130 | P | 154.4 | P | --- |
| 255467_at | 49.6 | A | 33.9 | A | 89.1 | A | 39.2 | A | --- |
| 255468_at | 1756 | P | 1748.9 | P | 1833.3 | P | 1585.5 | P | --- |
| 255469_at | 883.2 | P | 639.8 | P | 621.9 | P | 601 | P | --- |

| | | | | | | | | | |
|---|---|---|---|---|---|---|---|---|---|
| 255470_at | 0.9 | A | 24.9 | A | 5.9 | A | 3.6 | A | --- |
| 255471_at | 2.6 | A | 16.9 | A | 41.6 | A | 23.7 | A | AOP3 |
| 255472_at | 217.5 | P | 208.1 | P | 260 | P | 239.2 | P | --- |
| 255473_at | 1584.1 | P | 1488.2 | P | 1489 | P | 1647.9 | P | --- |
| 255474_at | 1787.8 | P | 1626.7 | P | 1526.3 | P | 1479.5 | P | --- |
| 255475_at | 214.5 | P | 337.6 | P | 563.6 | P | 352.5 | P | --- |
| 255476_at | 18.7 | A | 32.7 | A | 60.3 | A | 11.8 | A | --- |
| 255477_at | 1891.9 | P | 1768.9 | P | 1597.9 | P | 1835.7 | P | --- |
| 255478_at | 404.2 | P | 577.9 | P | 486.7 | P | 623.8 | P | EID1 |
| 255479_at | 4385.1 | P | 4049.9 | P | 3057.5 | P | 3372 | P | SAG21 |
| 255480_at | 89.9 | P | 95.6 | P | 92.3 | P | 106.5 | P | --- |
| 255481_at | 139.6 | P | 117.1 | P | 209.7 | P | 142.5 | P | PMS1 |
| 255482_at | 2031.6 | P | 1708.4 | P | 2295.6 | P | 1656 | P | TOC159 |
| 255483_at | 1498.3 | P | 1901.7 | P | 1495.5 | P | 1954.2 | P | XT2 |
| 255484_at | 269.7 | P | 377.5 | P | 280 | P | 316 | P | --- |
| 255485_at | 211.6 | P | 227.3 | P | 181.9 | P | 232.6 | P | --- |
| 255486_at | 628.9 | P | 782.6 | P | 710.2 | P | 780.9 | P | MLO1 |
| 255487_at | 489.1 | P | 439.7 | P | 471.2 | P | 540 | P | --- |
| 255488_at | 348.7 | P | 330.1 | P | 255.3 | P | 302.4 | P | --- |
| 255489_at | 10.9 | A | 50.3 | A | 49.5 | A | 13.4 | A | AT4G02650 /// AT4G02655 |
| 255490_at | 19.7 | A | 21 | A | 8.7 | A | 18.9 | A | --- |
| 255491_at | 61.4 | A | 44.3 | A | 39.8 | A | 18.8 | A | AtIDD12 |
| 255492_at | 720.6 | P | 812.4 | P | 739.7 | P | 1004.4 | P | EOL1 |
| 255493_at | 33 | P | 36.4 | A | 32.9 | A | 31.9 | A | --- |
| 255494_at | 73 | P | 68.8 | P | 84 | P | 60.6 | P | --- |
| 255495_at | 692.7 | P | 702 | P | 652.9 | P | 655.1 | P | --- |
| 255496_at | 504.3 | P | 691.9 | P | 524.9 | P | 644.5 | P | BZO2H1 |
| 255497_at | 513.5 | P | 512.7 | P | 436 | P | 493.3 | P | UNE12 |
| 255498_at | 3122.6 | P | 3539.9 | P | 3343.9 | P | 3514.9 | P | --- |
| 255499_at | 241.1 | P | 231.2 | P | 167.2 | P | 228.4 | P | --- |
| 255500_at | 83.2 | P | 165.6 | P | 93.9 | P | 129.1 | P | APP |
| 255501_at | 436.8 | P | 355.6 | P | 380.4 | P | 315.9 | P | --- |
| 255502_at | 696.1 | P | 615.6 | P | 525.1 | P | 566.6 | P | --- |
| 255503_at | 390.1 | P | 251.2 | P | 346.2 | P | 333.2 | P | --- |
| 255504_at | 576 | P | 907.1 | P | 967.1 | P | 645.9 | P | --- |
| 255505_at | 2540.4 | P | 2830.2 | P | 3072.4 | P | 2525 | P | ATSAR2 |
| 255506_at | 801.1 | P | 884.9 | P | 729.8 | P | 897.7 | P | GATL6 |
| 255507_at | 761.4 | P | 795.4 | P | 704.8 | P | 768.9 | P | MOS6 |
| 255508_at | 244.1 | P | 164.7 | P | 255.4 | P | 206.3 | P | --- |
| 255509_at | 166.4 | P | 226.5 | P | 196.6 | P | 192.6 | P | --- |
| 255510_at | 302.5 | P | 316.8 | P | 353.2 | P | 329.4 | P | SWN |
| 255511_at | 66 | A | 84.8 | P | 71.4 | A | 97.5 | A | PIT1 |
| 255512_at | 188.4 | P | 228.5 | P | 140.6 | P | 233.5 | P | SYP42 |
| 255513_at | 254.9 | P | 425.1 | P | 347.7 | P | 388.6 | P | PRL |
| 255514_s_at | 14.5 | A | 0.5 | A | 15.3 | A | 0.5 | A | AGL78 |
| 255515_at | 1.3 | A | 1.5 | A | 35.4 | A | 5.1 | A | --- |
| 255516_at | 32.3 | A | 39.8 | P | 33.1 | A | 10 | A | --- |
| 255517_at | 109.9 | P | 508.6 | P | 245.6 | P | 475 | P | AtGH9B13 |
| 255518_at | 25.7 | A | 8.2 | A | 6.1 | A | 5.6 | A | --- |
| 255519_at | 2.3 | A | 3.9 | A | 7.3 | A | 5.5 | A | --- |
| 255520_at | 4653 | P | 3700.5 | P | 4812.8 | P | 3895.5 | P | --- |
| 255521_at | 251.5 | P | 107.2 | P | 139.3 | P | 110.8 | P | SUS3 |
| 255522_at | 1010 | P | 1047.6 | P | 838.4 | P | 1068.8 | P | RSH1 |
| 255523_at | 5 | A | 5.4 | A | 6.2 | A | 4.5 | A | --- |
| 255524_at | 1557.5 | P | 1312.9 | P | 867.4 | P | 1045.3 | P | ATPMEPCRB |
| 255525_at | 830.9 | P | 925.8 | P | 688.2 | P | 755.4 | P | --- |
| 255526_at | 611.8 | P | 580 | P | 629 | P | 611.6 | P | --- |
| 255527_at | 359.8 | P | 90.9 | A | 169.7 | P | 77.1 | P | --- |
| 255528_at | 3.8 | A | 2.5 | A | 4.1 | A | 1.7 | A | --- |
| 255529_at | 586.1 | P | 592.7 | P | 452.2 | P | 504.8 | P | --- |
| 255530_at | 27.3 | A | 11.9 | A | 21.6 | A | 3.3 | A | --- |
| 255531_at | 55.2 | A | 48.6 | A | 32 | A | 32.3 | A | --- |
| 255532_at | 22.2 | A | 33.2 | A | 12.8 | A | 26.9 | A | --- |
| 255533_at | 10.7 | A | 10.4 | A | 48 | A | 42.7 | A | --- |
| 255534_at | 10.7 | A | 1.9 | A | 33.9 | A | 3.6 | A | --- |
| 255535_at | 87.1 | P | 107.9 | P | 128 | P | 75.5 | P | --- |
| 255536_at | 137.6 | P | 134.8 | P | 104.2 | P | 174 | P | --- |
| 255537_at | 2273.7 | P | 2622.4 | P | 2850 | P | 2441.3 | P | PPOX |
| 255538_at | 48.9 | A | 121.9 | P | 58.2 | A | 111 | P | MYB55 |
| 255539_at | 253.3 | P | 239.9 | P | 206.9 | P | 272.3 | P | CRK |
| 255540_at | 1216.3 | P | 1324.3 | P | 1056.3 | P | 1243.1 | P | --- |
| 255541_s_at | 20.1 | A | 1.9 | A | 12.2 | A | 2.1 | A | PGP5 |
| 255542_at | 211.1 | P | 204.5 | P | 210.3 | P | 230.5 | P | --- |
| 255543_at | 2974.9 | P | 2105.7 | P | 1504.1 | P | 1828.6 | P | --- |
| 255544_at | 241 | P | 283 | P | 248.3 | P | 232.3 | P | --- |
| 255545_at | 28.2 | A | 54.8 | A | 63.9 | A | 48.8 | A | --- |
| 255546_at | 76.2 | P | 57 | A | 72.2 | M | 84.4 | P | --- |
| 255547_at | 37.7 | P | 21.5 | A | 20.4 | A | 23.4 | P | --- |
| 255548_at | 6.8 | A | 4.1 | A | 41.7 | A | 18.9 | A | --- |
| 255549_at | 302.7 | P | 264.8 | P | 168.4 | P | 337.9 | P | GPAT3 |
| 255550_at | 30.4 | P | 27.8 | A | 25.1 | A | 16.6 | A | AtSTS |
| 255551_at | 73.4 | P | 68.9 | P | 95.6 | P | 68.9 | P | KCO5 |
| 255552_at | 7732.2 | P | 5703.8 | P | 5624.1 | P | 5512.9 | P | SAM-2 |
| 255553_at | 479.6 | P | 484.8 | P | 393.3 | P | 453.7 | P | --- |
| 255554_at | 423 | P | 492.5 | P | 439.1 | P | 502.3 | P | --- |
| 255555_at | 614.2 | P | 632.2 | P | 507.8 | P | 534.6 | P | --- |
| 255556_at | 1616.1 | P | 1994.1 | P | 1860.3 | P | 1841.5 | P | NFU1 |
| 255557_at | 164 | P | 132 | P | 155.4 | P | 129.5 | P | --- |
| 255558_at | 2242.6 | P | 1874.2 | P | 2199.9 | P | 1999.6 | P | GLB1 |
| 255559_at | 276.8 | P | 333.1 | P | 235.2 | P | 342.8 | P | --- |
| 255560_at | 349 | P | 375.2 | P | 305.1 | P | 320 | P | --- |
| 255561_at | 352 | P | 328.6 | P | 313.8 | P | 358.4 | P | --- |
| 255562_at | 80.2 | P | 87.1 | P | 105.8 | P | 82.3 | P | MSH6 |
| 255563_at | 4 | A | 45.2 | A | 39.5 | A | 53.5 | A | --- |
| 255564_s_at | 64.5 | P | 56.2 | P | 33 | P | 41.2 | P | RGXT2 |
| 255565_at | 50.4 | P | 33.9 | P | 44 | A | 28.8 | A | --- |
| 255566_s_at | 218.8 | P | 242.3 | P | 189.9 | P | 267.1 | P | --- |
| 255567_at | 12203.4 | P | 12456 | P | 13036.8 | P | 12864.5 | P | --- |
| 255568_at | 77.2 | P | 78 | P | 103 | P | 77.3 | P | WRKY22 |
| 255569_at | 1327.9 | P | 1437 | P | 1328.2 | P | 1348.4 | P | ATSTE24 |
| 255570_at | 1566.1 | P | 1645.2 | P | 1813.7 | P | 1743.1 | P | ADNT1 |
| 255571_at | 293.7 | P | 372.1 | P | 319.1 | P | 302.9 | P | --- |
| 255572_at | 2781.6 | P | 3007.6 | P | 2449.5 | P | 2840.7 | P | --- |
| 255573_at | 483.9 | P | 568.3 | P | 400.4 | P | 465.3 | P | --- |
| 255574_at | 26.6 | A | 13 | A | 6.1 | A | 9.1 | A | CBL5 |
| 255575_at | 1227.1 | P | 320.7 | P | 352 | P | 363.8 | P | --- |
| 255576_at | 52.3 | P | 121.2 | P | 87.9 | P | 103.2 | P | --- |
| 255577_at | 210.6 | P | 302.2 | P | 212.7 | P | 221.1 | P | --- |
| 255578_at | 210.5 | P | 284.9 | P | 272.2 | P | 265.3 | P | --- |

| | | | | | | | | | |
|---|---|---|---|---|---|---|---|---|---|
| 255579_at | 106.3 | P | 141.1 | P | 109.9 | P | 157.2 | P | --- |
| 255580_at | 58 | A | 54.4 | A | 57.6 | A | 74.3 | P | TIP1;3 |
| 255581_at | 3.8 | A | 39 | A | 31.1 | A | 6.9 | A | --- |
| 255582_at | 70.5 | A | 57.1 | A | 102.6 | M | 59.3 | A | NGA4 |
| 255583_at | 211.7 | P | 191.9 | P | 159.8 | P | 171.6 | P | ARV2 |
| 255584_at | 1.6 | A | 7.4 | A | 12.4 | A | 1.1 | A | --- |
| 255585_at | 231.6 | P | 269.9 | P | 160.7 | P | 207.5 | P | anac069 |
| 255586_at | 1004.1 | P | 955.5 | P | 1130 | P | 1052.6 | P | MEE49 |
| 255587_at | 1420.5 | P | 1183.5 | P | 1538.5 | P | 1256.3 | P | AtPPa5 |
| 255588_at | 182.1 | P | 200.2 | P | 218.5 | P | 218.1 | P | --- |
| 255589_s_at | 542 | P | 526.8 | P | 526.1 | P | 524.3 | P | --- |
| 255590_at | 4282.6 | P | 4594.6 | P | 3202.6 | P | 4235.4 | P | --- |
| 255591_at | 8.9 | A | 45.7 | A | 60.4 | A | 49.1 | A | ATEXPA17 |
| 255592_s_at | 14.4 | A | 24.5 | M | 36.7 | M | 3 | A | --- |
| 255593_at | 258.8 | P | 401 | P | 244.8 | P | 326.7 | P | --- |
| 255594_at | 527.4 | P | 491.2 | P | 584.3 | P | 509.8 | P | ATABC1 |
| 255595_at | 237.9 | P | 250.3 | P | 220.5 | P | 222.4 | P | --- |
| 255596_at | 236.4 | P | 298 | P | 282 | P | 224.3 | P | WRKY47 |
| 255597_at | 91.6 | P | 146.2 | P | 152.4 | P | 131.5 | P | --- |
| 255598_at | 713.1 | P | 553 | P | 564.8 | P | 619.8 | P | --- |
| 255599_at | 181.4 | P | 160.2 | P | 164.5 | P | 138.9 | P | ATCNGC13 |
| 255600_s_at | 289.7 | P | 258.8 | P | 393 | P | 301.7 | P | --- |
| 255601_at | 17.8 | A | 37.6 | P | 43.2 | A | 54.4 | A | --- |
| 255602_at | 405.1 | P | 629.9 | P | 322.9 | P | 431.6 | P | --- |
| 255603_at | 502.8 | P | 561.3 | P | 506.6 | P | 660.1 | P | --- |
| 255604_at | 999.1 | P | 575.2 | P | 806.9 | P | 1606.5 | P | --- |
| 255605_at | 147 | P | 165.6 | P | 102.4 | P | 155.9 | P | --- |
| 255606_at | 3.7 | A | 31.8 | A | 1.2 | A | 14.7 | A | --- |
| 255607_at | 343.3 | P | 409.1 | P | 247 | P | 400.5 | P | --- |
| 255608_at | 26.7 | A | 25.2 | A | 55.9 | A | 32 | A | --- |
| 255609_s_at | 38.7 | P | 10.4 | A | 28.7 | P | 1.4 | A | --- |
| 255610_at | 35.2 | A | 31.9 | A | 40.5 | A | 36.9 | P | ATPIPK10 |
| 255611_at | 709.3 | P | 629.1 | P | 529.8 | P | 658.1 | P | --- |
| 255612_at | 3.2 | A | 30.2 | A | 47.7 | A | 2.6 | A | --- |
| 255613_at | 51.4 | A | 68.9 | P | 63 | A | 60 | P | --- |
| 255614_at | 246 | P | 300.6 | P | 276.2 | P | 310.2 | P | --- |
| 255615_at | 1227.1 | P | 1074.8 | P | 807.5 | P | 1168.4 | P | --- |
| 255616_at | 181.9 | P | 183.6 | P | 169.4 | P | 230.3 | P | --- |
| 255617_at | 215.3 | P | 233.7 | P | 177.6 | P | 193.2 | P | --- |
| 255618_at | 2.1 | A | 16.9 | A | 3 | A | 13.2 | A | --- |
| 255619_s_at | 393.2 | P | 394.3 | P | 384.9 | P | 392.4 | P | --- |
| 255620_at | 30.8 | A | 13.9 | A | 31 | A | 17.4 | A | --- |
| 255621_at | 456.7 | P | 605.8 | P | 775.2 | P | 691.5 | P | --- |
| 255622_at | 2310.4 | P | 1950.5 | P | 1995.4 | P | 2201.8 | P | GT72B1 |
| 255623_at | 5269.2 | P | 6880.6 | P | 6505.7 | P | 6585.8 | P | --- |
| 255624_at | 1222.9 | P | 1432.9 | P | 1057.7 | P | 1312.4 | P | ATMPK4 |
| 255625_at | 231.3 | P | 237.7 | P | 250.5 | P | 250.3 | P | GBF2 |
| 255626_at | 1896.2 | P | 1893.8 | P | 2896.3 | P | 1912.2 | P | --- |
| 255627_at | 76.7 | P | 95.7 | P | 57.5 | M | 104.3 | P | --- |
| 255628_at | 158.7 | P | 194.3 | P | 227.5 | P | 166.8 | P | GIF3 |
| 255629_at | 5099 | P | 4764.8 | P | 5362.2 | P | 4786.6 | P | ATOZI1 |
| 255630_at | 37.8 | A | 210.3 | P | 46.8 | A | 116.3 | P | --- |
| 255631_at | 644.8 | P | 811.4 | P | 626.4 | P | 839.1 | P | BSK3 |
| 255632_at | 16.1 | A | 0.9 | A | 12.2 | A | 3 | A | ADF8 |
| 255633_at | 1721.5 | P | 1602.8 | P | 1361.8 | P | 1672.8 | P | --- |
| 255634_at | 77.7 | P | 51.1 | P | 63.8 | P | 59.7 | A | FRI |
| 255635_at | 873.7 | P | 1055.1 | P | 852.6 | P | 848.7 | P | ATSK32 |
| 255636_at | 622.2 | P | 756.7 | P | 740.7 | P | 703.7 | P | ANL2 |
| 255637_at | 630.9 | P | 376.6 | P | 461.2 | P | 404.8 | P | --- |
| 255638_at | 654.9 | P | 696.6 | P | 718.2 | P | 795 | P | --- |
| 255639_at | 176.1 | P | 182.8 | P | 253.5 | P | 140.6 | P | APRR8 |
| 255640_at | 124.9 | P | 146.3 | P | 175.9 | P | 142.8 | P | --- |
| 255641_at | 66.2 | M | 77.6 | A | 118.3 | M | 50.2 | A | --- |
| 255642_at | 66.9 | P | 122.2 | P | 65.8 | P | 139.7 | P | iqd17 |
| 255643_at | 80.8 | P | 86.6 | P | 74 | P | 75.5 | P | --- |
| 255644_at | 2.2 | A | 2.6 | A | 4.8 | A | 2.9 | A | --- |
| 255645_at | 355.8 | P | 698.9 | P | 475.7 | P | 731.8 | P | --- |
| 255646_at | 1.8 | A | 3.8 | A | 5.8 | A | 4 | A | --- |
| 255647_at | 238.8 | P | 221.5 | P | 227.8 | P | 176.2 | P | ECA2 |
| 255648_at | 3.3 | A | 2.9 | A | 34.6 | A | 17.8 | A | --- |
| 255649_at | 0.4 | A | 0.5 | A | 25.3 | A | 10.6 | A | --- |
| 255650_s_at | 198.6 | P | 231.5 | P | 140.2 | P | 169.1 | P | CIP4.1 |
| 255651_at | 44.2 | P | 53.7 | P | 102 | P | 44.4 | P | --- |
| 255652_at | 135.3 | P | 155.1 | P | 131.8 | P | 144.6 | P | MEE47 |
| 255653_at | 4.5 | A | 17.6 | A | 16.9 | A | 1.7 | A | --- |
| 255654_at | 28.8 | P | 26.6 | A | 24.5 | A | 8.9 | A | --- |
| 255655_at | 137.5 | P | 123.1 | P | 144.2 | P | 110.9 | P | --- |
| 255656_at | 191.8 | P | 222.7 | P | 172.7 | P | 193.5 | P | --- |
| 255657_at | 3715.4 | P | 3613.1 | P | 3325.2 | P | 4220.2 | P | --- |
| 255658_at | 3.3 | A | 39.3 | P | 5.6 | A | 26.2 | A | --- |
| 255659_at | 1610.4 | P | 2103.7 | P | 1814.6 | P | 1981.6 | P | --- |
| 255660_at | 307.5 | P | 400 | P | 313 | P | 354.3 | P | --- |
| 255661_at | 14.2 | A | 2.5 | A | 21.4 | A | 22.7 | A | ULP1B |
| 255662_at | 222.5 | P | 516.6 | P | 350 | P | 376.8 | P | GPAT8 |
| 255663_at | 161 | P | 183.1 | P | 186.8 | P | 241.9 | P | --- |
| 255664_at | 289.1 | P | 335.7 | P | 314.4 | P | 335.1 | P | --- |
| 255665_at | 257.3 | P | 190.3 | P | 202.4 | P | 165.3 | P | CRP |
| 255666_at | 190.1 | P | 115.8 | P | 128.3 | P | 109.4 | P | --- |
| 255667_at | 46.2 | P | 77 | P | 48.5 | A | 67.8 | P | PLDBETA2 |
| 255668_s_at | 417 | P | 346.1 | P | 447.2 | P | 341 | P | --- |
| 255669_at | 0.9 | A | 7.8 | A | 14.6 | A | 0.6 | A | MBD3 |
| 255670_at | 288 | P | 234.7 | P | 166.9 | P | 243.4 | P | RHB1A |
| 255671_at | 640.2 | P | 760 | P | 642.1 | P | 655.8 | P | --- |
| 255672_at | 30.4 | P | 43.5 | P | 48.7 | A | 57 | A | EDA8 |
| 255673_at | 167.6 | P | 236 | P | 194.3 | P | 273.7 | P | RLK4 |
| 255674_at | 4342.5 | P | 4654.6 | P | 4306.4 | P | 5478.3 | P | PIP1;4 |
| 255675_at | 190.7 | P | 161.7 | P | 312.8 | P | 183.9 | P | ATMYC1 |
| 255676_at | 937.7 | P | 1109.1 | P | 879.2 | P | 1249.3 | P | BAM2 |
| 255677_at | 220.6 | P | 289.8 | P | 256.2 | P | 295.2 | P | --- |
| 255678_at | 167.1 | P | 219.4 | P | 157.3 | P | 194.8 | P | AT4G00520 |
| 255679_at | 197 | P | 237.9 | P | 266.9 | P | 225.5 | P | --- |
| 255680_at | 82.7 | P | 122.2 | P | 103 | P | 61.2 | P | --- |
| 255681_at | 206.6 | P | 229.7 | P | 180.6 | P | 178.8 | P | DGD2 |
| 255682_at | 381.5 | P | 460.6 | P | 407.9 | P | 420.6 | P | --- |
| 255683_at | 899.5 | P | 1079.2 | P | 935.2 | P | 1075.4 | P | --- |
| 255684_at | 169.8 | P | 142.1 | P | 140.9 | P | 135.8 | P | --- |
| 255685_s_at | 678.8 | P | 447.6 | P | 900.6 | P | 473.6 | P | --- |
| 255686_at | 864.2 | P | 978.3 | P | 810.7 | P | 1142.3 | P | KEA2 |
| 255687_at | 77.1 | P | 82 | P | 76.8 | P | 99.4 | P | KEA2 |

| | | | | | | | | | |
|---|---|---|---|---|---|---|---|---|---|
| 255688_at | 665.2 | P | 703.6 | P | 838.7 | P | 717.4 | P | --- |
| 255689_at | 59.4 | P | 37.1 | P | 18.4 | A | 28.7 | P | --- |
| 255690_at | 1066.6 | P | 1313 | P | 1188.3 | P | 1624.4 | P | CYP86A2 |
| 255691_at | 1439.4 | P | 2185.8 | P | 2466.3 | P | 2378.8 | P | ANTR2 |
| 255692_at | 391.4 | P | 629.4 | P | 573 | P | 527.8 | P | GPAT8 |
| 255693_s_at | 62.2 | P | 78.7 | P | 105.2 | P | 97.9 | P | BRCA2(IV) |
| 255694_at | 1003.8 | P | 792.9 | P | 1615.3 | P | 1132.1 | P | UNE10 |
| 255695_at | 28.3 | A | 4.1 | A | 8 | A | 19.5 | A | UNE11 |
| 255696_at | 308.2 | P | 246.1 | P | 279.1 | P | 263 | P | GAE3 |
| 255697_at | 2.5 | A | 0.3 | A | 16.3 | A | 1.9 | A | --- |
| 255698_at | 398.7 | P | 326.7 | P | 372.8 | P | 293.4 | P | --- |
| 255699_at | 31.2 | A | 20.5 | A | 65.8 | A | 13.6 | A | PME38 |
| 255700_at | 47.7 | P | 73.4 | P | 47.4 | P | 57.2 | P | --- |
| 255701_at | 4.3 | A | 5.4 | A | 7.3 | A | 3.8 | A | JLO |
| 255702_at | 148 | P | 166.9 | P | 163.2 | P | 160.3 | P | XSP1 |
| 255703_at | 375.4 | P | 257.6 | P | 207.3 | P | 306.8 | P | --- |
| 255704_at | 500.1 | P | 540.1 | P | 589.9 | P | 531.7 | P | --- |
| 255705_at | 9.6 | A | 0.5 | A | 21.1 | P | 1 | A | --- |
| 255706_at | 6447.3 | P | 5601.1 | P | 7150.1 | P | 5864.2 | P | ATRPS13A |
| 255707_at | 155.3 | P | 180 | P | 244.4 | P | 175.8 | P | MEE50 |
| 255708_at | 2.7 | A | 27.3 | A | 35.2 | A | 15.9 | A | --- |
| 255709_at | 510.3 | P | 494.1 | P | 562.4 | P | 513.7 | P | YAB3 |
| 255710_at | 488 | P | 441.7 | P | 405.1 | P | 464.2 | P | --- |
| 255711_at | 694.2 | P | 615.1 | P | 663.8 | P | 576.2 | P | --- |
| 255712_at | 27.2 | P | 13.7 | A | 31.4 | A | 4.7 | A | --- |
| 255713_s_at | 162.4 | P | 231 | P | 184.9 | P | 194.7 | P | --- |
| 255714_at | 817.6 | P | 1021.9 | P | 723.8 | P | 938.3 | P | --- |
| 255715_s_at | 13.4 | A | 28.7 | A | 11.6 | A | 16.2 | A | --- |
| 255716_at | 752.4 | P | 889.3 | P | 614.8 | P | 832.1 | P | CRCK2 |
| 255717_at | 2.7 | A | 2.2 | A | 42.8 | A | 23.2 | A | --- |
| 255718_at | 1249.4 | P | 1648.2 | P | 1182.7 | P | 1180.8 | P | ATNSI |
| 255719_at | 3231.6 | P | 3583.4 | P | 4179.8 | P | 3778.7 | P | --- |
| 255720_at | 9284.2 | P | 11236.6 | P | 11768.6 | P | 10977 | P | PRK |
| 255721_at | 717.8 | P | 792.4 | P | 799.4 | P | 831.4 | P | --- |
| 255722_at | 24.9 | A | 34.8 | A | 35.3 | A | 23.2 | A | --- |
| 255723_at | 469.3 | P | 317.2 | P | 312 | P | 326.1 | P | AFP3 |
| 255724_at | 37.2 | P | 44.5 | P | 51.3 | A | 40.2 | A | BRCA2(IV) |
| 255725_at | 304.6 | P | 306.1 | P | 292.7 | P | 286 | P | PFT1 |
| 255726_at | 36.2 | A | 139.7 | P | 82.2 | A | 151.4 | P | --- |
| 255727_at | 27.8 | A | 61.2 | A | 46.8 | A | 79.9 | M | --- |
| 255728_at | 250.9 | P | 176.2 | P | 222.1 | P | 173.3 | P | --- |
| 255729_at | 447.6 | P | 387.2 | P | 397.6 | P | 371.9 | P | --- |
| 255730_at | 43.8 | P | 37.6 | P | 52.8 | P | 26.5 | P | --- |
| 255731_at | 1329 | P | 1716.2 | P | 1622 | P | 1676 | P | RCN1 |
| 255732_at | 803.3 | P | 895.5 | P | 918.6 | P | 1230.3 | P | KCS5 |
| 255733_at | 135.7 | P | 106.2 | P | 99.9 | P | 90.3 | P | --- |
| 255734_at | 552.9 | P | 350.5 | P | 536.7 | P | 284 | P | --- |
| 255735_at | 893.4 | P | 1059 | P | 967.1 | P | 924.6 | P | --- |
| 255736_at | 707.2 | P | 765.7 | P | 758.7 | P | 749.2 | P | --- |
| 255737_at | 578.9 | P | 601.6 | P | 589.9 | P | 452.7 | P | --- |
| 255738_at | 371 | P | 400.2 | P | 404.7 | P | 417.6 | P | --- |
| 255739_at | 153.3 | P | 196.3 | P | 145 | P | 195.9 | P | ANAC008 |
| 255740_at | 428.3 | P | 450.4 | P | 414.1 | P | 424.7 | P | --- |
| 255741_at | 1.2 | A | 4.1 | A | 1.4 | A | 1.6 | A | ATIPT6 |
| 255742_at | 569.2 | P | 728.3 | P | 461.1 | P | 633.8 | P | TEM1 |
| 255743_at | 257.9 | P | 367.5 | P | 286.7 | P | 326.6 | P | --- |
| 255744_at | 4 | A | 21.3 | A | 3 | A | 1.5 | A | --- |
| 255745_at | 4.1 | A | 2.3 | A | 1.8 | A | 10.4 | A | --- |
| 255746_at | 2 | A | 6.8 | A | 3.2 | A | 1.1 | A | --- |
| 255747_s_at | 20.3 | A | 2.1 | A | 4 | A | 1.6 | A | --- |
| 255748_at | 0.9 | A | 1.1 | A | 0.9 | A | 2.5 | A | --- |
| 255749_at | 1590.3 | P | 1244.2 | P | 1887.2 | P | 1452.5 | P | STRS1 |
| 255750_at | 13.1 | A | 6.5 | A | 22.2 | A | 11.9 | A | --- |
| 255751_at | 12.2 | A | 22.3 | A | 9.8 | A | 4.7 | A | --- |
| 255752_at | 3999.5 | P | 4346.9 | P | 3919 | P | 4030.6 | P | --- |
| 255753_at | 308.4 | P | 271.5 | P | 234 | P | 248.2 | P | MYB51 |
| 255754_at | 16.1 | A | 3.9 | A | 6 | A | 30 | A | --- |
| 255755_s_at | 0.8 | A | 1.5 | A | 0.9 | A | 1.6 | A | --- |
| 255756_at | 29.8 | A | 41.1 | P | 6.4 | A | 33.5 | A | AtGH9B5 |
| 255757_at | 10.8 | A | 39.7 | P | 33.9 | P | 33 | P | ROPGEF3 |
| 255758_at | 14.5 | A | 28.1 | A | 24.5 | A | 34.8 | A | --- |
| 255759_at | 201.2 | P | 220.9 | P | 253.7 | P | 244.3 | P | --- |
| 255760_at | 301.8 | P | 358.1 | P | 279.7 | P | 438.4 | P | --- |
| 255761_at | 44.1 | A | 37.6 | A | 54.4 | P | 54.3 | A | --- |
| 255762_at | 312.2 | P | 320.8 | P | 299.7 | P | 271.5 | P | HAC12 |
| 255763_at | 181 | P | 222.4 | P | 153.5 | A | 261.2 | P | --- |
| 255764_at | 1112 | P | 1738 | P | 1666.3 | P | 1332.5 | P | HCF173 |
| 255765_at | 3.1 | A | 17.9 | A | 2.9 | A | 2.1 | A | --- |
| 255766_at | 289.5 | P | 271.9 | P | 381.9 | P | 320.7 | P | --- |
| 255767_at | 1463.5 | P | 1168.7 | P | 1474.8 | P | 1287.8 | P | --- |
| 255768_at | 13.8 | A | 32.6 | A | 39.9 | A | 39.5 | A | --- |
| 255769_at | 118.3 | P | 122.8 | P | 107.1 | P | 99.9 | P | --- |
| 255770_at | 83.4 | P | 80 | P | 75.4 | P | 75.7 | P | --- |
| 255771_at | 5.6 | A | 50.3 | A | 50.1 | A | 13.8 | A | --- |
| 255772_at | 0.7 | A | 1.7 | A | 1.1 | A | 1.6 | A | --- |
| 255773_at | 2803.3 | P | 2077 | P | 3454.9 | P | 1925.4 | P | SOT17 |
| 255774_at | 466.8 | P | 464.2 | P | 438.7 | P | 434.4 | P | --- |
| 255775_at | 391.9 | P | 456.3 | P | 430.8 | P | 413.2 | P | RBL12 |
| 255776_at | 6422.2 | P | 5386.5 | P | 6805.8 | P | 5815 | P | --- |
| 255777_at | 433.6 | P | 552.1 | P | 529.9 | P | 465.2 | P | GR-RBP6 |
| 255778_at | 562 | P | 656.6 | P | 614.8 | P | 645.2 | P | PSP |
| 255779_at | 456.2 | P | 537.7 | P | 551.6 | P | 624.9 | P | PDCB3 |
| 255780_at | 237.5 | P | 296.6 | P | 232.4 | P | 277.6 | P | GAUT11 |
| 255781_at | 293.3 | P | 350.9 | P | 334.3 | P | 351 | P | --- |
| 255782_at | 241.3 | P | 230.4 | P | 272.9 | P | 318.5 | P | MP |
| 255783_at | 270.2 | P | 287.4 | P | 322.6 | P | 326.8 | P | iqd32 |
| 255784_at | 187.2 | P | 227.7 | P | 173.6 | P | 205.1 | P | --- |
| 255785_at | 1727.8 | P | 2136.7 | P | 1888.1 | P | 2219.7 | P | APS2 |
| 255786_at | 7227.9 | P | 2728.6 | P | 7328.4 | P | 3877.1 | P | ATCLH1 |
| 255787_at | 915.1 | P | 807.9 | P | 1049 | P | 707.4 | P | --- |
| 255788_at | 742.7 | P | 764.6 | P | 708.6 | P | 823 | P | IAA13 |
| 255789_at | 5851.6 | P | 5295.9 | P | 7742.7 | P | 5322.2 | P | --- |
| 255790_at | 36.1 | A | 58.3 | A | 51 | A | 80.2 | P | --- |
| 255791_at | 1351.7 | P | 1780.4 | P | 1659.1 | P | 1636.9 | P | DAL1 |
| 255792_at | 515.6 | P | 710.2 | P | 683 | P | 731.2 | P | --- |
| 255793_at | 1100.1 | P | 1138.9 | P | 1214.5 | P | 1184.6 | P | --- |
| 255794_at | 504.7 | P | 348.3 | P | 445.4 | P | 333.4 | P | ANAC041 |
| 255795_at | 1917.1 | P | 968.9 | P | 932.2 | P | 1953.8 | P | RD20 |
| 255796_at | 96.9 | P | 135.2 | P | 183.6 | P | 131.7 | P | SUVH2 |

| | | | | | | | | | |
|---|---|---|---|---|---|---|---|---|---|
| 255797_at | 1451.5 | P | 1331.7 | P | 1415.3 | P | 1240.9 | P | --- |
| 255798_at | 745.4 | P | 940.9 | P | 837.9 | P | 1027.6 | P | --- |
| 255799_at | 676 | P | 743.8 | P | 820.1 | P | 664.8 | P | DET1 |
| 255800_at | 0.5 | A | 0.7 | A | 2.2 | A | 2.4 | A | --- |
| 255801_at | 198 | P | 171.6 | P | 151.1 | P | 198.7 | P | --- |
| 255802_s_at | 24.6 | A | 24.2 | M | 24.6 | A | 34.1 | P | --- |
| 255803_at | 223.9 | P | 238.2 | P | 235.5 | P | 218.3 | P | --- |
| 255804_at | 12.8 | A | 18.1 | A | 11.9 | A | 24.8 | A | --- |
| 255805_at | 16.7 | A | 2.4 | A | 3.3 | A | 13.3 | A | --- |
| 255806_at | 32.1 | A | 26.4 | A | 7.1 | A | 1.7 | A | --- |
| 255807_at | 52.6 | P | 139.2 | P | 72.8 | P | 91.9 | P | --- |
| 255808_at | 512.2 | P | 384 | P | 482.2 | P | 398.2 | P | --- |
| 255809_at | 3329.5 | P | 3805.8 | P | 3613.3 | P | 4408.1 | P | --- |
| 255810_at | 696.7 | P | 622 | P | 677 | P | 592.6 | P | --- |
| 255811_at | 59.1 | A | 82.1 | A | 58.4 | A | 46.6 | A | ATHSP22.0 |
| 255812_at | 1.3 | A | 34.1 | A | 1.6 | A | 2.7 | A | HKT1 |
| 255813_at | 47.5 | A | 41.9 | A | 46.9 | A | 10.2 | A | --- |
| 255814_at | 2.7 | A | 12.7 | A | 17.5 | A | 5.8 | A | --- |
| 255815_at | 2.6 | A | 1.7 | A | 1.8 | A | 6.9 | A | MGH3 |
| 255816_at | 2197.7 | P | 2184.9 | P | 2290.7 | P | 2185.3 | P | GLTP1 |
| 255817_at | 411.5 | P | 506.8 | P | 614.1 | P | 583.2 | P | PDLP3 |
| 255818_at | 345.1 | P | 420.8 | P | 269.4 | P | 426.8 | P | --- |
| 255819_s_at | 3953 | P | 3099 | P | 3552.3 | P | 3262 | P | --- |
| 255820_at | 6.5 | A | 18.1 | A | 23.9 | A | 8.2 | A | --- |
| 255821_at | 331.9 | P | 273.9 | P | 288 | P | 245.9 | P | --- |
| 255822_at | 58.3 | P | 200.3 | P | 74.3 | A | 168.3 | P | ATEXPA8 |
| 255823_at | 3.1 | A | 3.3 | A | 9.3 | A | 3.5 | A | LBD15 |
| 255824_at | 2.9 | A | 24.8 | A | 3.4 | A | 4.8 | A | --- |
| 255825_at | 292 | P | 394.3 | P | 334.5 | P | 368.6 | P | --- |
| 255826_at | 1770.4 | P | 2738.9 | P | 1994.8 | P | 2691.9 | P | HEME2 |
| 255827_at | 709.5 | P | 956.9 | P | 651 | P | 947.2 | P | --- |
| 255828_at | 150.1 | P | 158.9 | P | 197.1 | P | 130.9 | P | --- |
| 255829_at | 391.5 | P | 546.6 | P | 433.3 | P | 575.7 | P | KT2 |
| 255830_at | 2015.7 | P | 1862.5 | P | 2644.1 | P | 1926.1 | P | --- |
| 255831_at | 30.1 | A | 32.2 | A | 31.9 | A | 37.3 | A | --- |
| 255832_at | 93 | P | 112.5 | P | 118.4 | P | 119.8 | P | --- |
| 255833_at | 203.3 | P | 205.3 | P | 167.9 | P | 226.5 | P | --- |
| 255834_at | 1008 | P | 1080.4 | P | 1051.6 | P | 1057 | P | --- |
| 255835_at | 5 | A | 29.1 | A | 40.8 | A | 34.1 | A | --- |
| 255836_at | 205.2 | P | 241.3 | P | 203.2 | P | 283.5 | P | --- |
| 255837_at | 40.1 | A | 48.5 | A | 53.9 | A | 40.8 | A | RIC1 |
| 255838_at | 163.2 | M | 182.6 | A | 157.1 | A | 258.3 | M | --- |
| 255839_at | 143.4 | P | 172.1 | P | 176.7 | P | 169.5 | P | --- |
| 255840_at | 27.2 | A | 29 | A | 41.3 | A | 30.9 | A | --- |
| 255841_at | 7.8 | A | 36.7 | A | 6.9 | A | 23 | A | --- |
| 255842_at | 410.9 | P | 490.1 | P | 460.7 | P | 513.8 | P | scpl46 |
| 255843_at | 78.8 | P | 91.8 | P | 77.7 | A | 54.2 | P | CPL3 |
| 255844_at | 279.6 | P | 353.5 | P | 167.5 | P | 259.1 | P | --- |
| 255845_at | 414.4 | P | 553.5 | P | 385.7 | P | 483.2 | P | --- |
| 255846_at | 284.4 | P | 283.4 | P | 322.5 | P | 269.5 | P | ATSWI3B |
| 255847_at | 6.9 | A | 38.5 | A | 63.1 | A | 6.1 | A | ACHT3 |
| 255848_at | 23.4 | A | 24 | A | 7.5 | A | 19.3 | A | --- |
| 255849_at | 7.3 | A | 9 | A | 5 | A | 24.9 | A | --- |
| 255850_at | 6403.7 | P | 7129 | P | 4924.6 | P | 7048.2 | P | --- |
| 255851_at | 113.4 | P | 116.4 | P | 154.3 | P | 150.1 | P | --- |
| 255852_at | 578.4 | P | 1818.2 | P | 728.4 | P | 1352.9 | P | SVL2 |
| 255853_at | 62.8 | A | 57.2 | A | 50.4 | A | 50.4 | A | --- |
| 255854_at | 96.9 | P | 204.2 | P | 158.8 | P | 194.9 | P | --- |
| 255855_at | 43.2 | P | 36.6 | A | 3.7 | A | 18.5 | A | PDR11 |
| 255856_at | 459.9 | P | 1223.8 | P | 714.3 | P | 987.1 | P | --- |
| 255857_at | 824.7 | P | 1070 | P | 937.7 | P | 1021.2 | P | ABA4 |
| 255858_at | 31.1 | A | 51.2 | P | 40.1 | A | 59.3 | P | ZFP6 |
| 255859_at | 344.3 | P | 337.1 | P | 412.9 | P | 398.9 | P | --- |
| 255860_at | 352.4 | P | 330.6 | P | 325.2 | P | 424.1 | P | AtGUS3 |
| 255861_at | 41.1 | A | 10.8 | A | 53 | A | 8.5 | A | --- |
| 255862_at | 30.7 | A | 27.9 | M | 36.5 | A | 48.1 | P | --- |
| 255863_s_at | 0.4 | A | 0.7 | A | 1.5 | A | 6.1 | A | --- |
| 255864_at | 255.6 | P | 340.4 | P | 315.1 | P | 372.1 | P | --- |
| 255865_at | 361.9 | P | 272.1 | P | 289.8 | P | 314.6 | P | --- |
| 255866_at | 204 | P | 176.1 | P | 182.2 | P | 210.9 | P | --- |
| 255867_at | 9.9 | A | 5.3 | A | 7.1 | A | 2.8 | A | --- |
| 255868_at | 1.6 | A | 1.7 | A | 2.5 | A | 2 | A | --- |
| 255869_at | 536.5 | P | 569.8 | P | 526.5 | P | 557.5 | P | --- |
| 255870_at | 251.9 | P | 176 | P | 155.6 | P | 184.3 | P | --- |
| 255871_at | 643.4 | P | 697.4 | P | 849.5 | P | 653.7 | P | U2B" |
| 255872_at | 197.1 | P | 114.1 | P | 110.1 | P | 156.5 | P | SIP4 |
| 255873_at | 21.5 | A | 4.9 | A | 3.1 | A | 16.8 | A | LBD13 |
| 255874_at | 378.9 | P | 406.7 | P | 267.7 | P | 390.2 | P | ETG1 |
| 255875_s_at | 27.6 | A | 21.6 | A | 28.7 | P | 23.6 | A | --- |
| 255876_at | 88.6 | P | 146.8 | P | 168.3 | P | 171.6 | P | --- |
| 255877_at | 405.6 | P | 401.5 | P | 570.2 | P | 449.6 | P | --- |
| 255878_at | 107.1 | P | 123.7 | P | 115.8 | P | 128.5 | P | --- |
| 255879_at | 2.3 | A | 2.1 | A | 11.1 | A | 18.6 | A | --- |
| 255880_at | 242.5 | P | 269.9 | P | 220.7 | P | 307.9 | P | --- |
| 255881_at | 116.2 | P | 157.1 | P | 174 | P | 150.2 | P | DIN9 |
| 255882_x_at | 27.2 | A | 33 | A | 45.2 | A | 10.2 | A | --- |
| 255883_at | 184.7 | P | 170.2 | P | 204.3 | P | 249.9 | P | --- |
| 255884_at | 88.1 | P | 72.4 | P | 85.3 | M | 109.9 | P | --- |
| 255885_at | 2415.8 | P | 2186.8 | P | 3028.7 | P | 2444.7 | P | SMT2 |
| 255886_at | 16258.1 | P | 17123.1 | P | 15756.5 | P | 16764.5 | P | DRT112 |
| 255887_at | 166.8 | P | 212.5 | P | 179.8 | P | 243.6 | P | --- |
| 255888_at | 297 | P | 252.6 | P | 339.7 | P | 266.3 | P | --- |
| 255889_at | 968.8 | P | 1346.7 | P | 952.8 | P | 1329.3 | P | WBC11 |
| 255890_at | 197.7 | P | 274 | P | 276.9 | P | 340.3 | P | --- |
| 255891_at | 55.3 | A | 93.8 | P | 51.4 | A | 57.7 | P | EGY3 |
| 255892_at | 3.9 | A | 4.6 | A | 27 | A | 10 | A | --- |
| 255893_at | 110.3 | P | 18.9 | A | 174 | P | 13.7 | A | --- |
| 255894_at | 389.9 | P | 402.4 | P | 304.4 | P | 403.6 | P | --- |
| 255895_at | 988.8 | P | 753 | P | 823.7 | P | 601.4 | P | --- |
| 255896_at | 24 | A | 26.1 | A | 44.5 | P | 35 | M | --- |
| 255897_at | 0.8 | A | 0.1 | A | 0.3 | A | 0.1 | A | --- |
| 255898_at | 288.9 | P | 269.5 | P | 254.4 | P | 317.8 | P | --- |
| 255899_at | 265.5 | P | 284.1 | P | 307.7 | P | 278.8 | P | --- |
| 255900_at | 319.7 | P | 224.9 | P | 292.1 | P | 234.4 | P | --- |
| 255901_at | 469.7 | P | 499.7 | P | 411 | P | 559.8 | P | GER2 |
| 255902_at | 5935.9 | P | 4916.3 | P | 6429.7 | P | 5098 | P | --- |
| 255903_at | 4.2 | A | 4.2 | A | 1.9 | A | 1.5 | A | MYB52 |
| 255904_at | 613.9 | P | 693.5 | P | 669.5 | P | 592.8 | P | --- |
| 255905_at | 14.5 | A | 4 | A | 2.7 | A | 1.6 | A | BETA-TIP |

| | | | | | | | | | |
|---|---|---|---|---|---|---|---|---|---|
| 255906_at | 99.2 | P | 105.7 | P | 118.4 | P | 107.6 | P | --- |
| 255907_at | 52.5 | P | 50 | P | 47 | P | 44.4 | P | HDG12 |
| 255908_s_at | 365.5 | P | 353.3 | P | 342.7 | P | 314.1 | P | --- |
| 255909_at | 111.1 | P | 138.4 | P | 153.4 | P | 120 | P | CDKD1;3 |
| 255910_at | 146.8 | P | 193.2 | P | 164.7 | P | 171.5 | P | --- |
| 255911_at | 1.7 | A | 8.2 | A | 8.9 | A | 13 | A | --- |
| 255912_at | 4.5 | A | 20.1 | A | 40 | A | 26.7 | A | --- |
| 255913_at | 400.8 | P | 522.8 | P | 417.7 | P | 521.8 | P | --- |
| 255914_at | 0.6 | A | 14.9 | P | 16.7 | A | 15 | A | --- |
| 255915_at | 2.8 | A | 2.4 | A | 23.1 | A | 11.7 | A | --- |
| 255916_at | 25.5 | A | 30.2 | A | 4.7 | A | 25.3 | A | --- |
| 255917_at | 47.1 | A | 33 | A | 26.6 | A | 34.4 | A | --- |
| 255918_at | 5.9 | A | 10.8 | A | 5.3 | A | 3.3 | A | --- |
| 255919_at | 6.8 | A | 40.9 | A | 15.6 | A | 36.8 | A | --- |
| 255920_at | 67.2 | A | 69.2 | A | 24.5 | A | 74.8 | A | --- |
| 255921_at | 15.4 | A | 22.8 | A | 42.1 | A | 41.4 | A | APUM8 |
| 255922_at | 27.9 | P | 15 | A | 5.4 | A | 11.7 | A | --- |
| 255923_at | 174.5 | P | 236.2 | P | 211.1 | P | 151.7 | P | --- |
| 255924_at | 152.6 | P | 238.5 | P | 246.6 | P | 211.5 | P | --- |
| 255925_at | 952.7 | P | 1007.7 | P | 840.3 | P | 1110 | P | --- |
| 255926_at | 346.7 | P | 453.6 | P | 359.2 | P | 428.5 | P | --- |
| 255927_at | 46.6 | A | 98.9 | A | 57.8 | A | 13.3 | A | --- |
| 255928_at | 648.7 | P | 545.6 | P | 597.4 | P | 532.5 | P | --- |
| 255929_at | 483.1 | P | 287.6 | P | 418.1 | P | 332.4 | P | --- |
| 255930_at | 21 | A | 10.5 | A | 47.6 | A | 10.4 | A | --- |
| 255931_at | 157.3 | P | 316.4 | P | 164.7 | P | 264.9 | P | AtPP2-A12 |
| 255932_at | 26.8 | A | 48 | A | 60.1 | A | 28 | A | --- |
| 255933_at | 314.3 | P | 346 | P | 338.3 | P | 344 | P | ATRBL6 |
| 255934_at | 1 | A | 1.3 | A | 4.2 | A | 2.4 | A | CYP87A2 |
| 255935_at | 60.7 | P | 50 | A | 59.4 | A | 45.6 | P | --- |
| 255936_at | 265.8 | P | 305.6 | P | 227.5 | P | 297.1 | P | --- |
| 255937_at | 1.2 | A | 3.5 | A | 3.7 | A | 2.3 | A | DDF1 |
| 255938_at | 67.1 | P | 50.8 | P | 55.3 | A | 39.5 | A | --- |
| 255939_at | 86.8 | P | 94.3 | P | 83.5 | P | 104.8 | P | --- |
| 255940_at | 231.2 | P | 233.7 | P | 222.9 | P | 231.9 | P | --- |
| 255941_at | 98.7 | P | 143.3 | P | 114.6 | P | 115.3 | P | ATTIM17-1 |
| 255942_at | 1826.8 | P | 960.3 | P | 1538.9 | P | 1091 | P | AtUGT85A2 |
| 255943_at | 421.5 | P | 297.6 | P | 352.2 | P | 299.5 | P | AtUGT85A5 |
| 255944_at | 36.1 | A | 31.8 | A | 8.4 | A | 39.9 | A | --- |
| 255945_at | 17 | A | 15.4 | A | 2.5 | A | 32 | A | --- |
| 255946_at | 375.8 | P | 401.9 | P | 448.7 | P | 368.5 | P | SHM6 |
| 255947_at | 284.8 | P | 316.4 | P | 270.1 | P | 264 | P | --- |
| 255948_at | 238.3 | P | 332.7 | P | 273.2 | P | 329.6 | P | --- |
| 255949_at | 2.1 | A | 20.9 | A | 2.1 | A | 19.8 | A | --- |
| 255950_at | 11.4 | A | 8.4 | A | 34.4 | A | 19.2 | A | --- |
| 255951_at | 250.3 | P | 272.7 | P | 223 | P | 258.9 | P | MUB6 |
| 255952_at | 24.1 | A | 0.8 | A | 2 | A | 9.7 | A | AGL104 |
| 255953_at | 622 | P | 1071.5 | P | 616.7 | P | 1007.9 | P | TGA3 |
| 255954_at | 2.4 | A | 1.1 | A | 13.8 | A | 2 | A | emb2204 |
| 255955_at | 81.5 | A | 140 | A | 77.2 | A | 83.2 | A | --- |
| 255956_at | 3.3 | A | 5.3 | A | 4.9 | A | 1.9 | A | DD46 |
| 255957_at | 393.6 | P | 835.7 | P | 518.3 | P | 888.7 | P | --- |
| 255958_at | 4.2 | A | 18.5 | A | 12.8 | A | 6.3 | A | SULTR1;3 |
| 255959_at | 480.9 | P | 319.8 | P | 401.9 | P | 294 | P | ATPIP5K1 |
| 255960_at | 1169.1 | P | 1126.4 | P | 1099.5 | P | 1218.4 | P | --- |
| 255961_at | 117.3 | P | 87.3 | P | 170.3 | P | 76 | A | AtUGT85A7 |
| 255962_at | 33.1 | P | 83.4 | P | 66.9 | P | 97.4 | P | --- |
| 255963_at | 118.1 | P | 102.1 | P | 132.1 | P | 106.6 | P | MBD8 |
| 255964_at | 3.1 | A | 25.7 | A | 42.1 | A | 9.6 | A | ZYP1b |
| 255965_at | 43.8 | P | 23.5 | A | 48.2 | P | 0.8 | A | --- |
| 255966_at | 4672.8 | P | 5834.2 | P | 4742.2 | P | 5858.6 | P | GRF10 |
| 255967_at | 1009.8 | P | 1004.5 | P | 794.4 | P | 988.8 | P | --- |
| 255968_at | 1019.3 | P | 730.7 | P | 863.2 | P | 974.7 | P | --- |
| 255969_at | 37.4 | P | 60.6 | P | 70.1 | P | 101.5 | P | --- |
| 255970_s_at | 17.2 | A | 16.9 | A | 16.1 | A | 21.5 | A | --- |
| 255971_at | 12.3 | A | 0.3 | A | 3.8 | A | 0.2 | A | --- |
| 255972_at | 10.5 | A | 11.7 | A | 2.1 | A | 1.2 | A | --- |
| 255973_at | 70 | A | 80.9 | P | 108.9 | M | 80.8 | A | --- |
| 255974_at | 16.4 | A | 20.9 | A | 6.1 | A | 22.5 | P | --- |
| 255975_at | 16.4 | A | 40.6 | A | 47.4 | A | 29.9 | A | --- |
| 255976_at | 8.8 | A | 5.5 | A | 4 | A | 25.4 | A | --- |
| 255977_at | 3300.9 | P | 2856.3 | P | 3578.2 | P | 2984.3 | P | --- |
| 255978_at | 208.5 | P | 221.9 | P | 238.6 | P | 296.7 | P | --- |
| 255979_at | 560.7 | P | 486 | P | 528.8 | P | 569.2 | P | UPF3 |
| 255980_at | 1236.1 | P | 1046.7 | P | 649.8 | P | 957.6 | P | --- |
| 255981_at | 561.8 | P | 587.1 | P | 672.5 | P | 627.6 | P | --- |
| 255982_at | 3103.9 | P | 3949.1 | P | 2617.2 | P | 3864.9 | P | OHP2 |
| 255983_at | 2321.7 | P | 1938.1 | P | 1886.2 | P | 2009.3 | P | MES14 |
| 255984_at | 102.3 | P | 111 | P | 126.9 | P | 114.9 | P | IP5PI |
| 255985_at | 260.7 | P | 207 | P | 163.1 | P | 169.3 | P | --- |
| 255986_x_at | 1.2 | A | 2.4 | A | 4.2 | A | 6.9 | A | --- |
| 255987_s_at | 18.8 | A | 24.9 | A | 17.5 | A | 14 | A | --- |
| 255988_at | 40.7 | A | 27.6 | A | 19.9 | A | 26.1 | A | --- |
| 255989_at | 19.6 | A | 21.3 | A | 5.7 | A | 7.9 | A | --- |
| 255990_at | 12 | A | 18.9 | A | 39.4 | P | 5.7 | A | --- |
| 255991_at | 94.5 | P | 113.5 | P | 143.4 | P | 126.3 | P | --- |
| 255992_at | 542.9 | P | 668.3 | P | 709.5 | P | 753.3 | P | --- |
| 255993_at | 1.5 | A | 2.1 | A | 9.8 | A | 4.8 | A | --- |
| 255994_at | 377.8 | P | 345.3 | P | 370.7 | P | 339.4 | P | --- |
| 255995_at | 207.6 | P | 196.8 | P | 220.6 | P | 183.5 | P | RKF1 |
| 255996_s_at | 2.4 | A | 9.6 | A | 7 | A | 4.3 | A | --- |
| 255997_s_at | 18254.1 | P | 18677.8 | P | 18304.1 | P | 18839.2 | P | CAB3 |
| 255998_at | 18.2 | A | 8.7 | A | 41.2 | A | 18.5 | A | --- |
| 255999_at | 25.3 | A | 51.2 | A | 47 | M | 31.1 | A | WRKY71 |
| 256000_at | 1658.4 | P | 1698.2 | P | 1936.8 | P | 1554.6 | P | --- |
| 256001_at | 1391.8 | P | 1478.4 | P | 1337.4 | P | 1548.9 | P | --- |
| 256002_at | 2230.9 | P | 1704.4 | P | 2597.2 | P | 1859.4 | P | CARB |
| 256003_at | 27.5 | A | 22.6 | A | 31.1 | A | 37.8 | A | --- |
| 256004_at | 1.6 | A | 8.6 | A | 3.4 | A | 3.9 | A | --- |
| 256005_at | 2.8 | A | 24.9 | A | 3.8 | A | 4.5 | A | --- |
| 256006_at | 23.6 | A | 2.7 | A | 45.5 | A | 3.6 | A | --- |
| 256007_at | 195.2 | P | 199.8 | P | 206.8 | P | 237.6 | P | SAMC2 |
| 256008_s_at | 217.9 | P | 506.1 | P | 193.5 | P | 609.3 | P | --- |
| 256009_at | 14.4 | A | 26.8 | A | 67.4 | A | 27.5 | A | --- |
| 256010_at | 89.8 | P | 125.6 | P | 58.5 | A | 98.3 | P | ARF19 |
| 256011_at | 5.3 | A | 28.9 | P | 46.3 | A | 21.7 | A | --- |
| 256012_at | 10.5 | A | 92.5 | P | 6.5 | A | 34 | A | FMO1 |
| 256013_at | 151.6 | P | 255.4 | P | 182.1 | P | 173.9 | P | DA1 |
| 256014_at | 110.3 | P | 47.9 | M | 111.2 | P | 41.4 | A | --- |

| | | | | | | | | | |
|---|---|---|---|---|---|---|---|---|---|
| 256015_at | 2102.7 | P | 2737.2 | P | 2163.3 | P | 2781.5 | P | LHCA6 |
| 256016_at | 428.5 | P | 371.5 | P | 438.7 | P | 412.1 | P | --- |
| 256017_at | 1755 | P | 1011.2 | P | 1412.9 | P | 812.5 | P | JAZ1 |
| 256018_at | 53.3 | P | 14.2 | A | 33.1 | A | 2.6 | A | ho4 |
| 256019_at | 14.5 | A | 3.6 | A | 23.9 | A | 2.4 | A | --- |
| 256020_at | 1011.9 | P | 692.1 | P | 1014.5 | P | 753.7 | P | HEMA1 |
| 256021_at | 1350.6 | P | 786.9 | P | 1111.6 | P | 879.2 | P | ZW9 |
| 256022_at | 1774.3 | P | 1466 | P | 1640.8 | P | 1850.8 | P | AAP1 |
| 256023_at | 7.9 | A | 10.8 | A | 19.9 | A | 4.7 | A | ZW2 |
| 256024_at | 140.8 | P | 74 | P | 150.2 | P | 105.2 | P | ZF14 |
| 256025_at | 104 | P | 148 | P | 138.9 | P | 110 | P | RXF12 |
| 256026_at | 8.3 | A | 14 | A | 26.8 | A | 4.3 | A | --- |
| 256027_at | 100.6 | P | 128.5 | P | 120 | P | 108.2 | P | --- |
| 256028_at | 1.2 | A | 1.8 | A | 1.7 | A | 5 | A | PAB1 |
| 256029_at | 796.8 | P | 1107.4 | P | 859.3 | P | 1036.4 | P | STT3B |
| 256030_at | 82.8 | P | 95.6 | P | 89.4 | P | 129.1 | P | --- |
| 256031_at | 8.4 | A | 17.2 | A | 9.9 | A | 6.3 | A | --- |
| 256032_at | 0.7 | A | 1.7 | A | 2.8 | A | 3.2 | A | --- |
| 256033_at | 735.8 | P | 670 | P | 604.4 | P | 633.2 | P | UGT71C4 |
| 256034_at | 3164.8 | P | 2596.5 | P | 2423.1 | P | 2871.1 | P | --- |
| 256035_at | 4815.8 | P | 4753.1 | P | 4589.6 | P | 4972.6 | P | SIRANBP |
| 256036_at | 1441.8 | P | 1511.3 | P | 1654.2 | P | 1467.9 | P | F2KP |
| 256037_at | 58.5 | A | 16.8 | A | 34 | A | 11.8 | A | --- |
| 256038_at | 267.7 | P | 287.8 | P | 219.9 | P | 313.8 | P | --- |
| 256039_at | 24.8 | A | 4.7 | A | 22 | A | 21.1 | A | --- |
| 256040_at | 37.4 | A | 57.6 | P | 62 | M | 64.2 | P | --- |
| 256041_at | 422.1 | P | 426.4 | P | 300.4 | P | 379.3 | P | --- |
| 256042_at | 254.8 | P | 227.3 | P | 189.3 | P | 205.5 | P | --- |
| 256043_at | 619 | P | 723.1 | P | 712.4 | P | 696 | P | --- |
| 256044_at | 2.1 | A | 6.4 | A | 6.7 | A | 4.7 | A | --- |
| 256045_at | 41.7 | P | 48.3 | P | 39.3 | M | 29.4 | P | MAPKKK13 |
| 256046_at | 198.7 | P | 208.9 | P | 188.3 | P | 172.4 | P | --- |
| 256047_at | 22 | A | 2.3 | A | 4.5 | A | 18.2 | A | --- |
| 256048_at | 4.5 | A | 18.7 | A | 26.1 | A | 15.8 | A | --- |
| 256049_at | 1084.3 | P | 1248.4 | P | 1202.3 | P | 1363.4 | P | --- |
| 256050_at | 493.6 | P | 438.5 | P | 360.1 | P | 356.8 | P | ATEXO70B2 |
| 256051_at | 18 | A | 11.3 | A | 30.9 | A | 2.5 | A | CHX14 |
| 256052_at | 179.9 | P | 149.9 | P | 196.4 | P | 164 | P | --- |
| 256053_at | 77.3 | P | 46.1 | A | 85 | P | 76.6 | P | UGT71C3 |
| 256054_at | 3.8 | A | 4.1 | A | 5.4 | A | 13.7 | A | --- |
| 256055_at | 512.3 | P | 426.4 | P | 422.8 | P | 488.8 | P | --- |
| 256056_at | 1 | A | 1.7 | A | 2.7 | A | 7.5 | A | --- |
| 256057_at | 382.9 | P | 324.3 | P | 433 | P | 411.5 | P | NDA1 |
| 256058_at | 433.2 | P | 425.5 | P | 442.8 | P | 448.1 | P | UGT71C5 |
| 256059_at | 4.3 | A | 1.9 | A | 7 | A | 4 | A | --- |
| 256060_at | 803.7 | P | 603.3 | P | 637.9 | P | 720.9 | P | --- |
| 256061_at | 1825.6 | P | 1563.4 | P | 1545.4 | P | 1670.6 | P | --- |
| 256062_at | 934.1 | P | 858.5 | P | 1032.7 | P | 598.5 | P | LSH6 |
| 256063_at | 68.6 | A | 49.3 | P | 61.8 | P | 77.3 | P | STN1 |
| 256064_at | 288.5 | P | 385.1 | P | 327.3 | P | 287.7 | P | --- |
| 256065_at | 524.7 | P | 638 | P | 710.4 | P | 673.1 | P | --- |
| 256066_at | 64.7 | A | 125.8 | P | 56.1 | A | 126.2 | P | --- |
| 256067_at | 975.1 | P | 694.1 | P | 854.1 | P | 755.9 | P | --- |
| 256068_at | 532.8 | P | 621.5 | P | 699.9 | P | 628.1 | P | ATE1 |
| 256069_at | 441.2 | P | 478.8 | P | 545.5 | P | 460.4 | P | AFP2 |
| 256070_at | 287.4 | P | 475.4 | P | 363.9 | P | 534 | P | --- |
| 256071_at | 413.4 | P | 343.1 | P | 394.6 | P | 353.3 | P | --- |
| 256072_at | 9333.5 | P | 8437.3 | P | 10824.5 | P | 8852.3 | P | ATARCA |
| 256073_at | 30.7 | A | 26.4 | A | 3.3 | A | 23.1 | A | E12A11 |
| 256074_at | 875.3 | P | 1037.1 | P | 868.8 | P | 817.1 | P | --- |
| 256075_at | 611.3 | P | 731.2 | P | 445.1 | P | 811.2 | P | ATMPK8 |
| 256076_at | 1578.2 | P | 1994.6 | P | 1872.7 | P | 2096.9 | P | --- |
| 256077_at | 133.2 | P | 155.7 | P | 145.3 | P | 204.9 | P | --- |
| 256078_at | 189 | P | 201.7 | P | 191.6 | P | 180.3 | P | --- |
| 256079_at | 1.2 | A | 0.9 | A | 20.3 | A | 11.2 | A | --- |
| 256080_at | 11.7 | A | 14 | A | 15.3 | A | 7.7 | A | --- |
| 256081_at | 24.4 | P | 32.7 | A | 57.3 | P | 31.8 | P | WOX14 |
| 256082_at | 18 | A | 42.5 | A | 35.2 | A | 43.2 | P | --- |
| 256083_at | 24.8 | A | 35 | A | 10.5 | A | 8.5 | A | --- |
| 256084_at | 12.8 | A | 1.7 | A | 22.7 | A | 4.4 | A | --- |
| 256085_at | 449.3 | P | 522.4 | P | 405.3 | P | 541.5 | P | --- |
| 256086_at | 604.1 | P | 743.2 | P | 662.1 | P | 781.6 | P | --- |
| 256087_at | 41.4 | A | 33.3 | A | 39.3 | M | 31.9 | P | --- |
| 256088_at | 603.3 | P | 771.3 | P | 655 | P | 807.1 | P | --- |
| 256089_at | 196.5 | P | 261.3 | P | 246.3 | P | 232.5 | P | --- |
| 256090_at | 218 | P | 190.5 | P | 238.2 | P | 220.7 | P | --- |
| 256091_at | 3273.2 | P | 2587.4 | P | 2081.8 | P | 2250.7 | P | HMGB2 |
| 256092_at | 6499.4 | P | 4890.4 | P | 5009.6 | P | 5500.3 | P | HMGB3 |
| 256093_at | 87.5 | P | 139.6 | P | 122.5 | P | 100.6 | P | --- |
| 256094_at | 86.2 | P | 100.8 | P | 142.9 | P | 123 | P | SAUL1 |
| 256095_at | 5.1 | A | 29.2 | A | 34.5 | A | 28 | A | --- |
| 256096_at | 1.5 | A | 2.6 | A | 5 | A | 24.6 | A | --- |
| 256097_at | 39.4 | A | 75.2 | P | 49.8 | A | 79.1 | A | --- |
| 256098_at | 51.2 | A | 72.2 | A | 15 | A | 24.9 | A | --- |
| 256099_at | 129.6 | A | 265.6 | P | 192.6 | P | 254.4 | P | CYP78A5 |
| 256100_at | 66.5 | P | 253.9 | P | 77.7 | P | 194 | P | --- |
| 256101_at | 4.4 | A | 0.9 | A | 7.9 | A | 8.2 | A | --- |
| 256102_at | 0.9 | A | 1 | A | 2.6 | A | 3.3 | A | --- |
| 256103_at | 0.5 | A | 2.7 | A | 0.8 | A | 9.5 | A | --- |
| 256104_at | 2.6 | A | 13.9 | A | 8.4 | A | 12.9 | A | ATTPS2 |
| 256105_at | 43.1 | A | 25.8 | A | 30.7 | A | 39.9 | A | LSH8 |
| 256106_at | 1595.6 | P | 1398.9 | P | 1986.6 | P | 1450.8 | P | --- |
| 256107_at | 178.7 | P | 122.2 | P | 200.3 | P | 76.6 | P | --- |
| 256108_at | 37.4 | A | 13.2 | A | 31.4 | A | 32.5 | A | --- |
| 256109_at | 1.1 | A | 1.3 | A | 28.1 | A | 1.4 | A | --- |
| 256110_at | 302.7 | P | 454.3 | P | 418.7 | P | 479.3 | P | --- |
| 256111_at | 36.5 | P | 36.6 | A | 37.3 | A | 42.3 | P | --- |
| 256112_at | 721 | P | 1377.6 | P | 765.2 | P | 1221 | P | ATRABA1B |
| 256113_at | 1510.8 | P | 1594.7 | P | 1482.4 | P | 1726.1 | P | --- |
| 256114_at | 147.3 | P | 196.6 | P | 46.2 | A | 286.4 | P | --- |
| 256115_at | 7206.1 | P | 6992.7 | P | 7774.1 | P | 7068.2 | P | --- |
| 256116_at | 378.8 | P | 487.7 | P | 250.5 | P | 542.6 | P | CPuORF55 |
| 256117_at | 733.7 | P | 870.3 | P | 688.5 | P | 842.8 | P | --- |
| 256118_at | 276.9 | P | 317.4 | P | 337.5 | P | 335.9 | P | KU70 |
| 256119_at | 1687.5 | P | 1678.5 | P | 1527.3 | P | 1837 | P | --- |
| 256120_at | 21.5 | A | 12.5 | A | 35 | A | 3.7 | A | --- |
| 256121_at | 501.6 | P | 441.1 | P | 439.2 | P | 462.2 | P | --- |
| 256122_at | 130.6 | P | 131.4 | P | 138.6 | P | 114.6 | P | --- |
| 256123_at | 150.8 | P | 220.7 | P | 211 | P | 210.9 | P | GC2 |

| | | | | | | | | | |
|---|---|---|---|---|---|---|---|---|---|
| 256124_at | 4.7 | A | 12.5 | A | 2 | A | 2.2 | A | ATPUP9 |
| 256125_at | 474.6 | P | 956.1 | P | 724.8 | P | 946.4 | P | ATLP-1 |
| 256126_at | 1.8 | A | 1.5 | A | 2.7 | A | 3.1 | A | --- |
| 256127_at | 28.9 | A | 45 | P | 65.3 | P | 41.2 | P | AtRABA6b |
| 256128_at | 35.4 | P | 34.4 | P | 53.8 | P | 32 | P | LAC1 |
| 256129_at | 2259 | P | 3642 | P | 2443.5 | P | 3307.4 | P | --- |
| 256130_at | 646.8 | P | 597.8 | P | 513.2 | P | 677.9 | P | --- |
| 256131_at | 21.2 | A | 51.6 | P | 46.9 | M | 41.6 | A | AtbZIP58 |
| 256132_at | 2.7 | A | 51.2 | A | 81 | A | 29.3 | P | --- |
| 256133_at | 250.1 | P | 316.8 | P | 334.2 | P | 281 | P | --- |
| 256134_at | 1906.5 | P | 1987.6 | P | 2216.7 | P | 2141 | P | AAPT1 |
| 256135_at | 6.8 | A | 17.4 | A | 9.5 | A | 4.5 | A | --- |
| 256136_s_at | 0.1 | A | 0.1 | A | 0.6 | A | 0.1 | A | --- |
| 256137_at | 1.5 | A | 1.8 | A | 2.7 | A | 2.8 | A | --- |
| 256138_at | 13.5 | A | 4.8 | A | 2.5 | A | 1 | A | --- |
| 256139_at | 17.5 | A | 12.6 | A | 24.4 | P | 38.2 | P | --- |
| 256140_at | 759.8 | P | 684.3 | P | 823.9 | P | 714.3 | P | --- |
| 256141_at | 31.5 | A | 14.7 | A | 42.8 | A | 19.4 | A | --- |
| 256142_at | 1045.9 | P | 1129.3 | P | 972.5 | P | 1016.9 | P | --- |
| 256143_at | 4786 | P | 3949.2 | P | 5108.2 | P | 4030.2 | P | --- |
| 256144_at | 4408.6 | P | 3832.2 | P | 4824 | P | 4325.3 | P | RACK1B_AT |
| 256145_at | 993.8 | P | 546 | P | 851 | P | 611.8 | P | --- |
| 256146_at | 186.2 | P | 177.1 | P | 166.4 | P | 166.7 | P | delta-ADR |
| 256147_at | 252.5 | P | 176.6 | P | 232.2 | P | 150.1 | P | --- |
| 256148_at | 864.9 | P | 730.2 | P | 727.6 | P | 735 | P | --- |
| 256149_at | 667.9 | P | 349.5 | P | 663.7 | P | 363.2 | P | AtIDD7 |
| 256150_at | 179.9 | P | 181.3 | P | 222.2 | P | 179.2 | P | ATFRUCT5 |
| 256151_at | 354.6 | P | 537 | P | 459.8 | P | 463.9 | P | --- |
| 256152_at | 752.9 | P | 694.7 | P | 692.9 | P | 712 | P | --- |
| 256153_at | 7.8 | A | 7.1 | A | 17.4 | A | 12 | A | UBQ12 |
| 256154_at | 5.1 | A | 5.2 | A | 24.3 | A | 18.3 | A | --- |
| 256155_at | 2.6 | A | 35.2 | A | 5.3 | A | 3.8 | A | MYB83 |
| 256156_at | 1820.7 | P | 2167.7 | P | 1504.5 | P | 2092.8 | P | ATPLC2 |
| 256157_at | 158 | P | 215.8 | P | 180.4 | P | 183.1 | P | LAG13 |
| 256158_at | 17.7 | A | 25.3 | A | 4.3 | A | 17.4 | A | ATPSK1 |
| 256159_at | 77.3 | P | 27 | A | 109.4 | P | 36.4 | P | JAZ8 |
| 256160_at | 2674.5 | P | 2400.5 | P | 2707.7 | P | 2593.7 | P | PDH-E1 BETA |
| 256161_at | 546 | P | 468.5 | P | 474 | P | 410.3 | P | --- |
| 256162_at | 34.4 | P | 17 | A | 21.1 | A | 39 | A | --- |
| 256163_at | 14.7 | A | 10.6 | A | 24.6 | A | 16.9 | A | --- |
| 256164_at | 2.4 | A | 1.7 | A | 9.6 | A | 1.4 | A | --- |
| 256165_at | 35.9 | P | 55.3 | P | 60.5 | A | 52.7 | P | --- |
| 256166_at | 1.8 | A | 7 | A | 4.3 | A | 3.1 | A | --- |
| 256167_at | 0.3 | A | 0.6 | A | 0.9 | A | 5.6 | A | --- |
| 256168_at | 827.4 | P | 1171.3 | P | 733.5 | P | 1093.7 | P | --- |
| 256169_at | 64.5 | P | 227.7 | P | 22.3 | A | 142 | P | --- |
| 256170_at | 53.1 | P | 176.3 | P | 26.7 | A | 78.1 | P | --- |
| 256171_s_at | 2.2 | A | 4.9 | A | 15.8 | A | 4.4 | A | --- |
| 256172_at | 217.1 | P | 226.2 | P | 310.6 | P | 234.1 | P | --- |
| 256173_at | 1330.7 | P | 1056.3 | P | 1253.1 | P | 1226 | P | --- |
| 256174_at | 522.4 | P | 583.6 | P | 441.4 | P | 646.7 | P | --- |
| 256175_at | 31.3 | A | 30.6 | A | 17.8 | A | 28.1 | A | --- |
| 256176_at | 26.4 | A | 29.3 | A | 5.2 | A | 41.3 | A | ATEXO70G2 |
| 256177_at | 76.7 | P | 121.1 | P | 65.7 | P | 65.9 | P | --- |
| 256178_s_at | 1834.1 | P | 531.4 | P | 1394.1 | P | 650.4 | P | ILL5 |
| 256179_at | 1822.1 | P | 1678.1 | P | 1836.5 | P | 1727.9 | P | UBP6 |
| 256180_at | 0.9 | A | 30.1 | A | 39.8 | A | 16.2 | A | --- |
| 256181_at | 37.6 | P | 50.4 | P | 37.5 | P | 47.2 | P | --- |
| 256182_at | 213.1 | P | 245.7 | P | 218.3 | P | 202.1 | P | SYP81 |
| 256183_at | 424.4 | P | 461.5 | P | 331.8 | P | 387.3 | P | MKK4 |
| 256184_at | 4927.6 | P | 3800.5 | P | 3338.3 | P | 3692.9 | P | --- |
| 256185_at | 276 | P | 562.1 | P | 351.5 | P | 514.2 | P | ADOF1 |
| 256186_at | 3037.7 | P | 2540.9 | P | 3428.7 | P | 2690.2 | P | 4CL1 |
| 256187_at | 599.6 | P | 699.1 | P | 670.6 | P | 732.2 | P | ATB ALPHA |
| 256188_at | 56.2 | A | 60.8 | P | 98.9 | P | 75 | P | --- |
| 256189_at | 2.4 | A | 9.8 | A | 18.1 | A | 12.9 | A | --- |
| 256190_at | 4.6 | A | 14.8 | A | 23.1 | A | 2.4 | A | NCED5 |
| 256191_at | 525.7 | P | 479.8 | P | 774.6 | P | 535.3 | P | --- |
| 256192_at | 99 | P | 181.4 | P | 136.7 | P | 181.8 | P | ATNUDX25 |
| 256193_at | 325.6 | P | 406.8 | P | 335.5 | P | 423 | P | --- |
| 256194_at | 9.6 | A | 1.3 | A | 5.5 | A | 1 | A | --- |
| 256195_at | 3.4 | A | 1.4 | A | 3.3 | A | 18.1 | A | --- |
| 256196_at | 44.7 | A | 17.4 | A | 66.6 | A | 39.8 | A | --- |
| 256197_at | 1.9 | A | 4.3 | A | 8.4 | A | 2.5 | A | --- |
| 256198_at | 138.7 | P | 120.9 | P | 71.8 | P | 138.3 | P | --- |
| 256199_at | 402.5 | P | 384.4 | P | 316.7 | P | 455.7 | P | SAB |
| 256200_at | 3 | A | 1.6 | A | 4.1 | A | 2.8 | A | EMB1674 |
| 256201_at | 201.5 | P | 277.9 | P | 210 | P | 227.4 | P | --- |
| 256202_s_at | 37.9 | A | 15.2 | A | 25.4 | A | 2.2 | A | --- |
| 256203_at | 0.5 | A | 1.1 | A | 1.1 | A | 1.5 | A | --- |
| 256204_at | 705.4 | P | 598.2 | P | 666.8 | P | 623.9 | P | POLGAMMA2 |
| 256205_at | 76 | P | 86.1 | P | 94.5 | P | 119.4 | P | --- |
| 256206_at | 45.1 | P | 60.7 | P | 71 | P | 46.7 | P | --- |
| 256207_at | 2673.4 | P | 1815.5 | P | 2567.1 | P | 2134.3 | P | --- |
| 256208_at | 0.8 | A | 0.7 | A | 30.1 | A | 0.8 | A | --- |
| 256209_at | 555.6 | P | 565.1 | P | 501.6 | P | 534.3 | P | ETFALPHA |
| 256210_at | 64.4 | P | 141.8 | P | 119.2 | P | 99.4 | P | --- |
| 256211_at | 4.6 | A | 1.9 | A | 1.5 | A | 10.8 | A | GA2OX7 |
| 256212_at | 74.5 | P | 75 | P | 70.8 | P | 96.2 | P | --- |
| 256213_at | 19.2 | A | 30.1 | A | 67.5 | P | 45.5 | A | --- |
| 256214_x_at | 1.8 | A | 3.4 | A | 1.1 | A | 6.7 | A | --- |
| 256215_at | 2463.3 | P | 3779.5 | P | 3215.3 | P | 3390.8 | P | --- |
| 256216_at | 3653.2 | P | 3884.3 | P | 3206.5 | P | 4119 | P | CRT1 |
| 256217_at | 51.5 | A | 74.2 | P | 46.9 | A | 85.6 | P | --- |
| 256218_at | 29.8 | P | 35.6 | A | 41.6 | A | 46.1 | P | --- |
| 256219_at | 65.3 | P | 126.4 | P | 136.9 | P | 105.7 | P | --- |
| 256220_at | 241.8 | P | 257.5 | P | 278.6 | P | 263.7 | P | --- |
| 256221_at | 2749.9 | P | 3182.1 | P | 2099.7 | P | 2984 | P | --- |
| 256222_at | 95.6 | P | 152.4 | P | 213.6 | P | 167.7 | P | --- |
| 256223_at | 5522 | P | 4811.9 | P | 4895.2 | P | 4800.8 | P | emb1303 |
| 256224_at | 3264.9 | P | 3174.1 | P | 3276.1 | P | 3395.2 | P | ATSAR1B |
| 256225_at | 2343.7 | P | 3502.5 | P | 2769.6 | P | 2384.8 | P | --- |
| 256226_at | 6696.1 | P | 7231.8 | P | 10532.6 | P | 6948.9 | P | ATDI19 |
| 256227_at | 195.8 | P | 218.9 | P | 309.3 | P | 197.5 | P | --- |
| 256228_at | 3305.3 | P | 3659.6 | P | 3350.7 | P | 3623.4 | P | --- |
| 256229_at | 834.6 | P | 867.1 | P | 650.4 | P | 830.1 | P | MSL3 |
| 256230_at | 151.5 | P | 130.4 | P | 137.7 | P | 130.7 | P | --- |
| 256231_at | 2289.6 | P | 2363.8 | P | 2561.1 | P | 2961.6 | P | --- |
| 256232_at | 666.3 | P | 583.5 | P | 628.2 | P | 629.7 | P | FYD |

| | | | | | | | | | |
|---|---|---|---|---|---|---|---|---|---|
| 256233_at | 720.8 | P | 794.6 | P | 584.8 | P | 709.4 | P | ITN1 |
| 256234_at | 673.2 | P | 678 | P | 931.2 | P | 687.7 | P | --- |
| 256235_at | 3830 | P | 3318 | P | 3305 | P | 3338.7 | P | ATCYSB |
| 256236_at | 353.8 | P | 384.7 | P | 396.7 | P | 360.9 | P | --- |
| 256237_at | 532.1 | P | 1875.9 | P | 593.3 | P | 1626.3 | P | DRT100 |
| 256238_at | 620 | P | 621.4 | P | 616.8 | P | 569.4 | P | ELC |
| 256239_at | 5.2 | A | 5.3 | A | 37.7 | A | 18.9 | A | --- |
| 256240_at | 753.9 | P | 817.4 | P | 739.7 | P | 765.6 | P | atnudt16 |
| 256241_at | 3491.8 | P | 3331.9 | P | 4610.3 | P | 3646.2 | P | --- |
| 256242_at | 227.9 | P | 307.8 | P | 302.8 | P | 292.5 | P | ATARP5 |
| 256243_at | 69 | A | 106.3 | P | 18.7 | A | 96.5 | P | ATHCHIB |
| 256244_at | 158.3 | P | 159.3 | P | 282.5 | P | 171.3 | P | SULTR4;2 |
| 256245_at | 1010.5 | P | 710 | P | 932.9 | P | 657.4 | P | HSP70 |
| 256246_at | 479.6 | P | 604.6 | P | 634.3 | P | 537.9 | P | ALDH22a1 |
| 256247_at | 2.3 | A | 4.3 | A | 20.1 | A | 1.6 | A | AGL91 |
| 256248_at | 98.6 | P | 62.6 | P | 117.9 | P | 94.1 | P | --- |
| 256249_at | 618.6 | P | 798.7 | P | 580.3 | P | 762.8 | P | MEE34 |
| 256250_at | 231.9 | P | 256.5 | P | 172 | P | 218.6 | P | --- |
| 256251_at | 465.4 | P | 416.4 | P | 455.7 | P | 403.3 | P | --- |
| 256252_at | 413.8 | P | 79.6 | P | 175.7 | P | 261.5 | P | --- |
| 256253_at | 799.4 | P | 755.9 | P | 899 | P | 809.2 | P | --- |
| 256254_at | 228.2 | P | 281.1 | P | 293.2 | P | 252.5 | P | --- |
| 256255_at | 789.5 | P | 911.9 | P | 1035.6 | P | 844 | P | --- |
| 256256_at | 720.6 | P | 707.4 | P | 634 | P | 610.8 | P | --- |
| 256257_at | 80.9 | P | 106 | P | 119.5 | P | 113 | P | ATE2 |
| 256258_at | 384.9 | P | 337.5 | P | 362.5 | P | 331.7 | P | NF-YC11 |
| 256259_at | 19.2 | A | 29.1 | A | 54.9 | A | 36 | A | --- |
| 256260_at | 21.4 | A | 2.7 | A | 35.3 | A | 24.1 | A | --- |
| 256261_at | 1.5 | A | 3.7 | A | 9.6 | A | 1.2 | A | RABA4D |
| 256262_at | 90.9 | P | 113.5 | P | 90.8 | P | 80.7 | P | --- |
| 256263_at | 3171.8 | P | 3450.6 | P | 3308.6 | P | 3345.9 | P | --- |
| 256264_at | 343.3 | P | 330.9 | P | 348.9 | P | 292.7 | P | --- |
| 256265_at | 8.7 | A | 38.6 | A | 3.7 | A | 2.1 | A | scpl16 |
| 256266_at | 66.6 | P | 258.2 | P | 42.3 | P | 142.8 | P | --- |
| 256267_at | 4127.7 | P | 5409.8 | P | 4901 | P | 5328.1 | P | --- |
| 256268_at | 428.2 | P | 522.6 | P | 457.5 | P | 511.1 | P | RBR1 |
| 256269_at | 111.4 | P | 132.2 | P | 130.3 | P | 135 | P | TGA6 |
| 256270_at | 255.2 | P | 366.1 | P | 260.3 | P | 319.1 | P | --- |
| 256271_at | 234.7 | P | 229.8 | P | 360.2 | P | 247.9 | P | --- |
| 256272_at | 334.3 | P | 313.2 | P | 367.9 | P | 379.1 | P | --- |
| 256273_at | 64.5 | P | 55.1 | A | 54.3 | A | 50.4 | A | TET6 |
| 256274_at | 1416.8 | P | 1629.6 | P | 1541.3 | P | 1535.8 | P | emb2738 |
| 256275_at | 146.2 | P | 210.1 | P | 186.8 | P | 246.8 | P | ACT11 |
| 256276_at | 116.1 | P | 86.7 | P | 108.8 | P | 89.3 | P | --- |
| 256277_at | 11620.2 | P | 11084.2 | P | 15088.5 | P | 11092.7 | P | FAD2 |
| 256278_at | 661.2 | P | 532.9 | P | 691.8 | P | 663.3 | P | --- |
| 256279_at | 837.1 | P | 826.7 | P | 649.8 | P | 801 | P | --- |
| 256280_at | 225.5 | P | 308.2 | P | 227.2 | P | 237.2 | P | AT3G12590 |
| 256281_at | 284.9 | P | 339.5 | P | 278.7 | P | 339.5 | P | TRFL9 |
| 256282_at | 101.1 | P | 155.5 | P | 162.5 | M | 157.9 | P | --- |
| 256283_at | 2 | A | 1.2 | A | 2 | A | 1.3 | A | --- |
| 256284_at | 180.6 | P | 233.1 | P | 271.7 | P | 200.6 | P | PSF2 |
| 256285_at | 60.3 | A | 40.3 | A | 57.5 | A | 32.8 | A | --- |
| 256286_at | 331.3 | P | 367.6 | P | 349 | P | 419.9 | P | --- |
| 256287_at | 2.6 | A | 3 | A | 5.3 | A | 0.3 | A | --- |
| 256288_at | 360.5 | P | 258.8 | P | 362.2 | P | 233.9 | P | PRMT3 |
| 256289_s_at | 0.4 | A | 0.4 | A | 10.7 | A | 0.5 | A | scpl14 |
| 256290_at | 1.1 | A | 2 | A | 4 | A | 16.6 | A | scpl17 |
| 256291_at | 414.1 | P | 448.3 | P | 378.9 | P | 482.7 | P | AtNek7 |
| 256292_at | 60.1 | A | 51.5 | A | 43.7 | A | 9.9 | A | --- |
| 256293_at | 64.3 | P | 60.2 | M | 45.4 | A | 85.7 | P | AGO7 |
| 256294_at | 379.6 | P | 537.8 | P | 308.3 | P | 444.6 | P | --- |
| 256295_at | 3.2 | A | 2 | A | 14.5 | A | 4.3 | A | --- |
| 256296_at | 168.2 | P | 92.4 | P | 108.2 | P | 78.7 | A | --- |
| 256297_at | 1.7 | A | 2.6 | A | 29.7 | A | 6.6 | A | --- |
| 256298_at | 25 | M | 14.6 | A | 14.9 | A | 26.2 | A | AGL94 |
| 256299_at | 1014.2 | P | 2770.2 | P | 1269.8 | P | 2593 | P | ATEXPA1 |
| 256300_at | 42.3 | A | 67.4 | P | 44.7 | A | 19.5 | A | NAP |
| 256301_at | 2317.1 | P | 2189.3 | P | 2294.2 | P | 2393.8 | P | --- |
| 256302_at | 53.5 | P | 57.7 | P | 56.5 | P | 105.2 | P | --- |
| 256303_at | 230.6 | P | 229.4 | P | 181.8 | P | 194.1 | P | --- |
| 256304_at | 129.2 | P | 165.8 | P | 203.5 | P | 232.9 | P | --- |
| 256305_at | 1165.1 | P | 1110.8 | P | 966.7 | P | 1118 | P | MRP1 |
| 256306_at | 10.7 | A | 29.7 | A | 51.1 | A | 17.3 | A | --- |
| 256307_at | 17.7 | A | 41.7 | A | 26 | A | 24 | A | --- |
| 256308_s_at | 584.5 | P | 438.3 | P | 537.6 | P | 543.6 | P | ATMRP13 |
| 256309_at | 19254.1 | P | 19509.8 | P | 19626.5 | P | 18987.2 | P | PSAK |
| 256310_at | 1452.4 | P | 2597.3 | P | 1691.7 | P | 2645.4 | P | ERD4 |
| 256311_at | 364.2 | P | 420.8 | P | 379.3 | P | 452.2 | P | ARF6 |
| 256312_x_at | 2.2 | A | 21.4 | A | 20.1 | A | 5.4 | A | --- |
| 256313_s_at | 1.3 | A | 0.8 | A | 1.5 | A | 1 | A | APUM17 |
| 256314_at | 20.7 | A | 45.1 | A | 23.3 | A | 27.2 | A | TOC75-I |
| 256315_at | 22.9 | A | 11.1 | A | 26.8 | A | 1.9 | A | --- |
| 256316_at | 29.4 | A | 4.6 | A | 49.9 | A | 14.8 | A | --- |
| 256317_at | 1.9 | A | 1.8 | A | 9.7 | A | 3.9 | A | --- |
| 256318_s_at | 0.8 | A | 1.8 | A | 8.2 | A | 5.8 | A | --- |
| 256319_at | 155.2 | P | 16.1 | A | 59.2 | P | 3.1 | A | --- |
| 256320_at | 134 | P | 130.5 | P | 128.5 | P | 168 | P | --- |
| 256321_at | 297.3 | P | 253.5 | P | 359 | P | 291.5 | P | LOX1 |
| 256322_at | 981.9 | P | 903.1 | P | 872.3 | P | 908.3 | P | AXR4 |
| 256323_at | 251.1 | P | 319.6 | P | 264.2 | P | 296.9 | P | --- |
| 256324_at | 1733.2 | P | 167.7 | P | 1760.4 | P | 274.8 | P | --- |
| 256325_at | 53.3 | M | 53.9 | A | 39.9 | A | 69.5 | M | --- |
| 256326_at | 270.8 | P | 289.5 | P | 186.9 | P | 183.1 | P | --- |
| 256327_at | 884.4 | P | 1067.9 | P | 825 | P | 1099.9 | P | GAUT9 |
| 256328_at | 4817.1 | P | 4321.5 | P | 4797.4 | P | 4353.3 | P | --- |
| 256329_at | 603.9 | P | 770.1 | P | 505.1 | P | 765.7 | P | SEC5A |
| 256330_at | 54.3 | P | 71.4 | P | 84.8 | P | 91.1 | P | --- |
| 256331_at | 326.7 | P | 423.6 | P | 404.5 | P | 455.3 | P | --- |
| 256332_at | 181.3 | P | 226.6 | P | 131.4 | P | 225.9 | P | GT2 |
| 256333_at | 710.9 | P | 743.7 | P | 603.5 | P | 825.1 | P | --- |
| 256334_at | 22 | A | 21.9 | A | 33.3 | A | 33.9 | P | --- |
| 256335_at | 3.6 | A | 2.5 | A | 28.4 | A | 4.4 | A | --- |
| 256336_at | 1167.1 | P | 1515.5 | P | 1240.4 | P | 1564.6 | P | --- |
| 256337_at | 63.9 | P | 169.2 | P | 91.1 | P | 115.6 | P | --- |
| 256338_at | 5.2 | A | 1.1 | A | 2.1 | A | 3.7 | A | --- |
| 256339_at | 4.1 | A | 17.2 | A | 3.2 | A | 4.3 | A | --- |
| 256340_at | 82.3 | P | 58.5 | A | 118.7 | P | 51.6 | A | --- |
| 256341_at | 699.2 | P | 687.7 | P | 887.9 | P | 786.7 | P | --- |

| | | | | | | | | | |
|---|---|---|---|---|---|---|---|---|---|
| 256342_at | 4567.4 | P | 4612.4 | P | 5312.3 | P | 4679.3 | P | --- |
| 256343_at | 355 | P | 408.3 | P | 369.6 | P | 374 | P | --- |
| 256344_at | 677.9 | P | 558.9 | P | 627 | P | 550 | P | TFIIIA |
| 256345_at | 19.9 | A | 18.5 | M | 54.7 | P | 6.7 | A | --- |
| 256346_s_at | 8.4 | A | 23.4 | P | 25.9 | P | 22 | M | --- |
| 256347_at | 1.9 | A | 11.3 | A | 18.7 | A | 0.9 | A | --- |
| 256348_at | 0.7 | A | 0.9 | A | 2 | A | 1.1 | A | --- |
| 256349_at | 29.1 | P | 21.1 | A | 22.2 | A | 6.1 | A | --- |
| 256350_at | 15.9 | A | 13.7 | P | 33 | A | 19 | A | PGSIP4 |
| 256351_at | 68.2 | P | 124.1 | P | 168.5 | P | 129.3 | P | ANP2 |
| 256352_at | 24.8 | A | 19.3 | A | 53.1 | P | 40 | A | ATPRP1 |
| 256353_at | 1070.7 | P | 1320.8 | P | 1040 | P | 1216 | P | --- |
| 256354_at | 11.6 | A | 5.2 | A | 7.2 | A | 31.5 | A | --- |
| 256355_at | 122.9 | A | 49.4 | A | 71.6 | A | 60.3 | A | --- |
| 256356_s_at | 43.4 | P | 102.1 | P | 42.1 | P | 74.6 | P | --- |
| 256357_at | 1.6 | A | 11.1 | A | 23.7 | A | 2.7 | A | --- |
| 256358_at | 11 | A | 5.2 | A | 5.9 | A | 24.7 | A | --- |
| 256359_at | 2.9 | A | 23.7 | A | 16 | A | 26.5 | P | --- |
| 256360_at | 28.8 | A | 24.4 | A | 7.5 | A | 27.6 | A | --- |
| 256361_at | 200.1 | P | 224 | P | 247.9 | P | 233.1 | P | pde194 |
| 256362_at | 23.4 | A | 47.5 | P | 34.3 | A | 27 | P | --- |
| 256363_at | 103.8 | P | 127.4 | P | 85.3 | A | 144.7 | P | --- |
| 256364_at | 10.9 | A | 10.3 | A | 30.5 | A | 5.1 | A | WRKY67 |
| 256365_at | 517.8 | P | 493.6 | P | 511.9 | P | 467.4 | P | --- |
| 256366_at | 458.4 | P | 706 | P | 331.8 | P | 523.1 | P | --- |
| 256367_at | 86.1 | P | 94.1 | P | 88.4 | P | 95.3 | P | --- |
| 256368_at | 1.4 | A | 12.9 | A | 18.8 | A | 5.4 | A | --- |
| 256369_at | 0.7 | A | 7 | A | 3.6 | A | 4.5 | A | --- |
| 256370_at | 62 | P | 36.7 | P | 45.8 | P | 55.4 | P | --- |
| 256371_at | 23 | A | 30.6 | A | 32.9 | P | 21.8 | A | --- |
| 256372_at | 287.1 | P | 280.2 | P | 375.1 | P | 200.8 | P | CAK4 |
| 256373_at | 364.8 | P | 339 | P | 347 | P | 355.3 | P | SGA2 |
| 256374_at | 151.7 | P | 174 | P | 214.9 | P | 130.2 | P | --- |
| 256375_at | 7.2 | A | 2.4 | A | 3.3 | A | 30 | A | --- |
| 256376_s_at | 58.9 | P | 27.8 | P | 51.1 | P | 40.5 | P | PXMT1 |
| 256377_at | 79.3 | P | 40.9 | P | 132.3 | P | 43.5 | P | --- |
| 256378_at | 44.5 | P | 44 | P | 18.3 | A | 50.4 | P | --- |
| 256379_at | 141.5 | P | 170 | P | 195 | P | 149.9 | P | --- |
| 256380_at | 881.8 | P | 681.5 | P | 756.1 | P | 792.7 | P | AR401 |
| 256381_at | 22.1 | A | 10.4 | A | 30.1 | A | 37.4 | A | --- |
| 256382_at | 15.3 | A | 12.6 | A | 51.2 | A | 21.5 | A | --- |
| 256383_at | 923.5 | P | 929.6 | P | 926.8 | P | 963.2 | P | --- |
| 256384_at | 309.1 | P | 303.7 | P | 337.1 | P | 264.3 | P | --- |
| 256385_at | 9379.7 | P | 6754.4 | P | 8949.5 | P | 7424.2 | P | --- |
| 256386_at | 141.7 | P | 125.8 | P | 143.1 | P | 140.2 | P | --- |
| 256387_at | 884.8 | P | 906.5 | P | 1009.1 | P | 782.4 | P | --- |
| 256388_at | 88.6 | P | 90.4 | P | 91.6 | P | 76 | P | --- |
| 256389_at | 3.4 | A | 11.5 | A | 8.9 | A | 31 | A | --- |
| 256390_at | 2.9 | A | 21.4 | A | 4.8 | A | 3.1 | A | ATMKK8 |
| 256391_at | 14 | A | 21.5 | A | 21.8 | A | 44 | P | --- |
| 256392_at | 35.8 | A | 31.7 | A | 19.1 | A | 25.2 | A | GATL4 |
| 256393_at | 31.9 | P | 37.9 | P | 37.7 | P | 39.7 | P | --- |
| 256394_at | 132.4 | P | 117.9 | P | 94.7 | P | 103 | P | --- |
| 256395_at | 51.1 | P | 36.4 | P | 73.1 | P | 50.1 | P | MUTE |
| 256396_at | 379 | P | 529.8 | P | 375.7 | P | 518.3 | P | --- |
| 256397_at | 797.3 | P | 691.3 | P | 875.3 | P | 854.1 | P | MKP2 |
| 256398_at | 267.4 | P | 113.1 | A | 183.4 | A | 124.9 | A | NIP7;1 |
| 256399_at | 389 | P | 371.5 | P | 269.3 | P | 351.7 | P | --- |
| 256400_at | 68.3 | P | 104.2 | P | 106.2 | P | 134.2 | P | --- |
| 256401_at | 381 | P | 344.5 | P | 351.5 | P | 329.6 | P | --- |
| 256402_at | 135 | P | 208.1 | P | 305.2 | P | 280.2 | P | --- |
| 256403_at | 462.4 | P | 557.1 | P | 463.5 | P | 567.5 | P | BPM2 |
| 256404_at | 198.5 | P | 177.6 | P | 193.7 | P | 218.9 | P | FRS7 |
| 256405_at | 92.7 | P | 58.1 | P | 119.1 | P | 67.3 | P | CPK33 |
| 256406_at | 140.5 | P | 159.3 | P | 228.9 | P | 157.6 | P | --- |
| 256407_at | 97.5 | P | 21 | A | 19.7 | A | 25.5 | A | ATSUC7 |
| 256408_at | 5.8 | A | 6.3 | A | 5.9 | A | 2.3 | A | --- |
| 256409_at | 214.2 | P | 152.4 | P | 200.2 | P | 169.3 | P | --- |
| 256410_at | 7.3 | A | 5.3 | A | 4.7 | A | 15.6 | A | --- |
| 256411_at | 3344.5 | P | 4452.2 | P | 4021.6 | P | 4081.5 | P | CLPP3 |
| 256412_at | 159.1 | P | 220.7 | P | 333.7 | P | 155.2 | P | ELO1 |
| 256413_at | 182.7 | P | 231.8 | P | 141 | P | 201.2 | P | --- |
| 256414_at | 1260.4 | P | 1393.8 | P | 1787.4 | P | 1423.1 | P | AL2 |
| 256415_at | 235.3 | P | 300.5 | P | 348.7 | P | 238.4 | P | --- |
| 256416_at | 139.5 | P | 90.4 | A | 183.2 | P | 97.1 | P | ATFER2 |
| 256417_s_at | 5979.8 | P | 4919.3 | P | 5126.3 | P | 5080.8 | P | FAD7 |
| 256418_at | 1.7 | A | 36.1 | A | 11.5 | A | 11.4 | A | --- |
| 256419_at | 155.1 | P | 220.5 | P | 125.2 | P | 235.7 | P | --- |
| 256420_at | 48.1 | P | 59.5 | A | 48.3 | A | 15.4 | A | --- |
| 256421_at | 2.1 | A | 11.8 | A | 1.8 | A | 13.8 | A | --- |
| 256422_at | 211.2 | P | 265.6 | P | 218.7 | P | 242.6 | P | MOS2 |
| 256423_at | 31.6 | P | 25.4 | P | 24.2 | A | 11.8 | A | scpl18 |
| 256424_at | 1239.8 | P | 1073.9 | P | 1136 | P | 1045.3 | P | --- |
| 256425_at | 954.7 | P | 833.2 | P | 787.3 | P | 744.3 | P | ADR1 |
| 256426_at | 205.3 | P | 145.8 | P | 167.3 | P | 161.5 | P | --- |
| 256427_at | 427.9 | P | 525.6 | P | 408.9 | P | 481 | P | LBD21 |
| 256428_at | 74.4 | P | 46.8 | P | 5.3 | A | 19.9 | A | AtRLP35 |
| 256429_at | 49.8 | P | 101.7 | P | 83.7 | P | 86.7 | P | --- |
| 256430_at | 109.3 | P | 125.6 | P | 138.3 | P | 147.7 | P | DREB2B |
| 256431_s_at | 390.3 | P | 367.6 | P | 116.8 | P | 226.5 | P | AtRLP34 |
| 256432_at | 20.7 | A | 16.8 | A | 12.8 | A | 28.8 | A | --- |
| 256433_at | 1061.2 | P | 1080.7 | P | 802.9 | P | 686.8 | P | SAG20 |
| 256434_at | 1.5 | A | 30.1 | A | 4.5 | A | 4.9 | A | --- |
| 256435_at | 3.4 | A | 10.6 | A | 13.1 | A | 2.6 | A | --- |
| 256436_at | 354.8 | P | 8.8 | A | 192.3 | M | 32.9 | A | --- |
| 256437_s_at | 2742.9 | P | 3030.4 | P | 3525.7 | P | 2565.9 | P | --- |
| 256438_s_at | 3765 | P | 2977.5 | P | 3934 | P | 3016.5 | P | --- |
| 256439_at | 5241.3 | P | 4300.4 | P | 5949.3 | P | 5205.3 | P | MSD1 |
| 256440_at | 2258.2 | P | 2506 | P | 2266.3 | P | 1941.9 | P | --- |
| 256441_at | 987 | P | 980.8 | P | 1138.6 | P | 1128.6 | P | --- |
| 256442_at | 106.8 | P | 135.3 | P | 48.3 | P | 41 | P | --- |
| 256443_at | 134.4 | P | 115.4 | P | 155 | P | 138.7 | P | --- |
| 256444_at | 14 | A | 22.7 | P | 37.4 | A | 27.8 | A | --- |
| 256445_at | 2.6 | A | 2.5 | A | 4.9 | A | 2.8 | A | --- |
| 256446_at | 68.4 | A | 92.6 | A | 115.7 | P | 81.8 | M | --- |
| 256447_at | 23.5 | A | 63.5 | P | 59.2 | A | 39.2 | P | --- |
| 256448_s_at | 1.3 | A | 1.6 | A | 5.8 | A | 2.6 | A | MED21 |
| 256449_at | 16.1 | A | 4.3 | A | 1.8 | A | 1.9 | A | --- |
| 256450_at | 27.8 | A | 27.4 | A | 50.8 | A | 31.2 | A | --- |

| | | | | | | | | | |
|---|---|---|---|---|---|---|---|---|---|
| 256451_s_at | 316.8 | P | 341.9 | P | 253.7 | P | 335.6 | P | --- |
| 256452_at | 287.4 | P | 407.1 | P | 570 | P | 384.3 | P | AtHB33 |
| 256453_at | 2267.7 | P | 1831.4 | P | 2422.9 | P | 1980 | P | DHAR2 |
| 256454_at | 1950.5 | P | 2972.9 | P | 2104.7 | P | 3475.6 | P | --- |
| 256455_at | 210 | P | 184.6 | P | 246.5 | P | 203.7 | P | --- |
| 256456_at | 145.8 | P | 349 | P | 190.2 | P | 285.4 | P | --- |
| 256457_at | 196.1 | P | 242.2 | P | 210.9 | A | 224.9 | P | --- |
| 256458_at | 1114.6 | P | 895.5 | P | 1040 | P | 788.4 | P | --- |
| 256459_at | 113.7 | P | 79.4 | A | 173.8 | P | 49.4 | P | ACC2 |
| 256460_at | 46.4 | P | 56.2 | P | 48.3 | P | 28.7 | A | --- |
| 256461_s_at | 440.9 | P | 354.3 | P | 412.7 | P | 433.5 | P | --- |
| 256462_at | 36.7 | A | 21.4 | A | 28.8 | A | 24.1 | A | --- |
| 256463_s_at | 7 | A | 1.2 | A | 2.7 | A | 1.8 | A | --- |
| 256464_at | 15.9 | A | 3.1 | A | 28.5 | A | 2.6 | A | --- |
| 256465_at | 28.6 | A | 37 | P | 46.7 | A | 16.2 | A | --- |
| 256466_at | 842.1 | P | 672.3 | P | 887.9 | P | 668.3 | P | --- |
| 256467_at | 11.5 | A | 3.1 | A | 1.8 | A | 2.2 | A | --- |
| 256468_at | 1795.3 | P | 2064.4 | P | 1877.7 | P | 1872.8 | P | --- |
| 256469_at | 226.7 | P | 169.9 | P | 213.9 | P | 165.9 | P | LOL1 |
| 256470_at | 7 | A | 0.6 | A | 5.4 | A | 11.3 | A | --- |
| 256471_at | 8.3 | A | 8.6 | P | 2.4 | A | 4.8 | A | --- |
| 256472_at | 1.5 | A | 2.5 | A | 2.3 | A | 2.1 | A | --- |
| 256473_at | 2 | A | 1.4 | A | 2.7 | A | 2.5 | A | --- |
| 256474_at | 1 | A | 1.1 | A | 0.9 | A | 0.5 | A | --- |
| 256475_s_at | 1.5 | A | 9.9 | A | 11.8 | A | 0.9 | A | --- |
| 256476_at | 2.1 | A | 4.8 | A | 2.4 | A | 2.6 | A | --- |
| 256477_at | 3.3 | A | 3.5 | A | 23.5 | A | 1.3 | A | --- |
| 256478_at | 25.1 | A | 16.1 | A | 47 | A | 32.6 | A | --- |
| 256479_at | 252.9 | P | 246 | P | 278 | P | 220.1 | P | --- |
| 256480_at | 601.2 | P | 565.4 | P | 711.4 | P | 662.8 | P | SAR1 |
| 256481_at | 32.2 | A | 4.9 | A | 38.3 | A | 20.3 | A | --- |
| 256482_at | 1165.3 | P | 1742.1 | P | 1260.9 | P | 1964.9 | P | FEI1 |
| 256483_at | 356 | P | 349.7 | P | 318.2 | P | 342.2 | P | --- |
| 256484_at | 92.1 | P | 61.3 | P | 50.8 | A | 70.8 | P | --- |
| 256485_at | 488.7 | P | 631 | P | 499 | P | 518.2 | P | --- |
| 256486_at | 6.6 | A | 11.1 | A | 6.5 | A | 3.4 | A | --- |
| 256487_at | 31 | P | 35.2 | P | 25.2 | A | 42.2 | P | --- |
| 256488_at | 197.1 | P | 163.6 | P | 224.1 | P | 181.8 | P | NFD4 |
| 256489_at | 1312.9 | P | 550.6 | P | 971.4 | P | 677.9 | P | --- |
| 256490_at | 104.4 | P | 143.2 | P | 141.6 | P | 134.1 | P | --- |
| 256491_at | 251.2 | P | 305.1 | P | 318.8 | P | 269.2 | P | --- |
| 256492_at | 19.6 | A | 16.7 | A | 3.3 | A | 19.3 | A | --- |
| 256493_at | 318.8 | P | 404 | P | 425 | P | 373.5 | P | --- |
| 256494_at | 25.2 | A | 49.7 | A | 15.7 | A | 21.6 | A | --- |
| 256495_at | 2.1 | A | 2 | A | 3 | A | 1.8 | A | --- |
| 256496_at | 13.4 | A | 19 | A | 26 | A | 23.5 | A | --- |
| 256497_at | 8004.9 | P | 15897.2 | P | 9562.5 | P | 14951.5 | P | ECS1 |
| 256498_at | 8.4 | A | 15.4 | A | 10.4 | A | 1.1 | A | --- |
| 256499_at | 9.5 | A | 1.8 | A | 11 | A | 0.4 | A | --- |
| 256500_at | 34 | P | 31.6 | P | 60 | P | 28.7 | P | --- |
| 256501_at | 31.7 | A | 12.7 | A | 6.1 | A | 9.1 | A | --- |
| 256502_at | 1979.4 | P | 1805.7 | P | 1980.2 | P | 2142.1 | P | --- |
| 256503_at | 1.7 | A | 1 | A | 2.6 | A | 2.3 | A | ATRL6 |
| 256504_at | 716 | P | 756.5 | P | 635.9 | P | 712 | P | --- |
| 256505_at | 302.7 | P | 342.6 | P | 363.1 | P | 347.9 | P | --- |
| 256506_at | 2.2 | A | 18.4 | A | 6.5 | A | 21.6 | A | --- |
| 256507_at | 58.3 | P | 69.1 | P | 68.9 | P | 53.3 | P | --- |
| 256508_at | 229.5 | P | 226.5 | P | 250 | P | 190.8 | P | --- |
| 256509_at | 83.7 | A | 73.6 | A | 72.4 | A | 66 | A | --- |
| 256510_at | 267 | P | 184.6 | P | 199.1 | P | 201.8 | P | --- |
| 256511_at | 255 | P | 176.5 | P | 213.6 | P | 175.3 | P | --- |
| 256512_at | 612.7 | P | 627.1 | P | 536.6 | P | 661.9 | P | --- |
| 256513_at | 193.2 | P | 178.9 | P | 177.4 | P | 145.2 | P | SGR2 |
| 256514_at | 531.1 | P | 550.1 | P | 602.3 | P | 557.7 | P | --- |
| 256515_at | 10.7 | A | 26.7 | A | 42.2 | A | 2.5 | A | --- |
| 256516_at | 1183.8 | P | 1196.3 | P | 1191.5 | P | 1352.8 | P | TMK1 |
| 256517_at | 11.3 | A | 5.5 | A | 27.3 | P | 12.5 | A | MMD1 |
| 256518_at | 129.4 | P | 185.1 | P | 189.4 | P | 213.2 | P | --- |
| 256519_at | 23.1 | A | 4.9 | A | 5 | A | 7.6 | A | --- |
| 256520_at | 1.3 | A | 3.7 | A | 2 | A | 4 | A | CYP96A14P |
| 256521_at | 19.2 | A | 3.7 | A | 2.8 | A | 22 | A | --- |
| 256522_at | 271 | P | 158.7 | P | 174.5 | P | 109.1 | P | CMPG1 |
| 256523_at | 520.4 | P | 520 | P | 585.9 | P | 672.8 | P | --- |
| 256524_at | 7842.2 | P | 8462 | P | 8154.9 | P | 8488.9 | P | ATGSR2 |
| 256525_at | 5841.8 | P | 2240.3 | P | 5736.5 | P | 2953.5 | P | --- |
| 256526_at | 292.8 | P | 340.2 | P | 216.7 | P | 198 | P | --- |
| 256527_at | 6158.5 | P | 406.9 | P | 5299.5 | P | 937 | P | --- |
| 256528_at | 81.6 | P | 91.6 | P | 121.1 | P | 131.6 | P | ZFP4 |
| 256529_at | 206 | P | 128 | P | 151.5 | P | 147.2 | P | --- |
| 256530_at | 943.3 | P | 1004.5 | P | 806.3 | P | 876.6 | P | --- |
| 256531_at | 15.9 | A | 22.9 | A | 64.4 | A | 17.9 | A | --- |
| 256532_at | 59.9 | P | 85 | P | 91.9 | P | 53 | P | --- |
| 256533_at | 275.5 | P | 301.7 | P | 238.7 | P | 268.8 | P | --- |
| 256534_at | 447 | P | 560.2 | P | 545.9 | P | 427.8 | P | --- |
| 256535_at | 8.9 | A | 1 | A | 4.2 | A | 18.9 | A | ANAC015 |
| 256536_at | 482.4 | P | 463.9 | P | 439.5 | P | 477.5 | P | --- |
| 256537_at | 76.9 | A | 77.7 | A | 135.9 | A | 76 | A | --- |
| 256538_x_at | 33.5 | A | 18.2 | A | 56.8 | A | 45.2 | P | --- |
| 256539_at | 12.3 | A | 8.2 | A | 28.4 | A | 0.3 | A | --- |
| 256540_at | 1.4 | A | 0.4 | A | 1.7 | A | 2.7 | A | --- |
| 256541_at | 385.7 | P | 358.5 | P | 409.4 | P | 358.5 | P | GLR3.3 |
| 256542_at | 931.1 | P | 865.9 | P | 964.7 | P | 976.2 | P | PMI1 |
| 256543_at | 561.2 | P | 777.5 | P | 565.3 | P | 573.7 | P | --- |
| 256544_at | 4 | A | 17.2 | A | 4.9 | A | 3.3 | A | MLO9 |
| 256545_at | 132 | P | 163.6 | P | 132.9 | P | 143.9 | P | --- |
| 256546_at | 2.3 | A | 1 | A | 1.9 | A | 11.5 | A | --- |
| 256547_at | 827.4 | P | 1317.4 | P | 810.1 | P | 1123.7 | P | --- |
| 256548_at | 95.9 | P | 149.7 | P | 138.2 | P | 145.5 | P | --- |
| 256549_at | 3.1 | A | 1.3 | A | 3.5 | A | 6.7 | A | --- |
| 256550_at | 1.7 | A | 1.7 | A | 2.4 | A | 3.8 | A | --- |
| 256551_at | 19.1 | A | 26.8 | A | 42.6 | A | 1.9 | A | --- |
| 256552_at | 6.3 | A | 3 | A | 31.6 | A | 18.7 | A | --- |
| 256553_at | 31.4 | P | 19.8 | A | 34.1 | A | 12.3 | A | --- |
| 256554_at | 1.2 | A | 0.6 | A | 9.6 | A | 0.6 | A | --- |
| 256555_at | 17.3 | A | 0.5 | A | 14.5 | A | 11 | A | --- |
| 256556_at | 4.3 | A | 1 | A | 6 | A | 1.8 | A | --- |
| 256557_at | 5.1 | A | 0.6 | A | 2 | A | 16.7 | A | --- |
| 256558_at | 0.8 | A | 0.5 | A | 1.7 | A | 1.2 | A | --- |
| 256559_at | 0.3 | A | 0.5 | A | 17.7 | A | 5.3 | A | --- |

| | | | | | | | | | |
|---|---|---|---|---|---|---|---|---|---|
| 256560_s_at | 2 | A | 1.4 | A | 4.5 | A | 2 | A | --- |
| 256561_s_at | 0.2 | A | 5.2 | A | 2.9 | A | 2.1 | A | --- |
| 256562_at | 1.2 | A | 3.5 | A | 6.3 | A | 0.7 | A | --- |
| 256563_at | 1.3 | A | 2.6 | A | 1.7 | A | 1.4 | A | RALFL27 |
| 256564_at | 166.2 | P | 211.1 | P | 170 | P | 165.5 | P | MES11 |
| 256565_at | 36 | A | 44.3 | P | 54.8 | P | 52.4 | P | AT3G19515 |
| 256566_at | 4.9 | A | 1.9 | A | 4.7 | A | 5.6 | A | --- |
| 256567_at | 475.7 | P | 510.4 | P | 389.3 | P | 405.4 | P | --- |
| 256568_at | 856.5 | P | 619.7 | P | 594.9 | P | 619.3 | P | --- |
| 256569_at | 26.1 | P | 31 | A | 32 | A | 13.8 | A | --- |
| 256570_at | 432.8 | P | 486.5 | P | 526 | P | 468.3 | P | --- |
| 256571_at | 15.1 | P | 30 | P | 36.5 | P | 18.6 | P | --- |
| 256572_at | 14.1 | A | 9.1 | A | 38.2 | A | 9.9 | A | --- |
| 256573_at | 441.8 | P | 440.2 | P | 459.5 | P | 460.5 | P | --- |
| 256574_at | 46 | A | 44.2 | M | 46 | A | 14.4 | A | --- |
| 256575_at | 508.6 | P | 744.7 | P | 686.4 | P | 672.9 | P | RHM3 |
| 256576_at | 144.5 | P | 76 | P | 69.9 | A | 73.8 | P | PMZ |
| 256577_at | 5307.5 | P | 1981.7 | P | 3990.4 | P | 3893.3 | P | --- |
| 256578_at | 586.9 | P | 619.3 | P | 737.5 | P | 672.9 | P | --- |
| 256579_at | 779.6 | P | 777.5 | P | 830.1 | P | 794.3 | P | --- |
| 256580_s_at | 5.7 | A | 4.6 | A | 29.6 | A | 6.7 | A | --- |
| 256581_at | 1.5 | A | 2 | A | 3.1 | A | 1.4 | A | --- |
| 256582_at | 8.2 | A | 21.6 | A | 56.4 | A | 16.4 | A | --- |
| 256583_at | 192.6 | P | 120.4 | P | 99.5 | P | 126.3 | P | --- |
| 256584_at | 24.2 | A | 24.8 | A | 36.9 | A | 31.4 | A | --- |
| 256585_at | 147.5 | P | 180.4 | P | 154.3 | P | 207.3 | P | --- |
| 256586_at | 26.6 | A | 2.4 | A | 12 | A | 18.8 | A | --- |
| 256587_at | 2.6 | A | 3.2 | A | 5.4 | A | 1.4 | A | --- |
| 256588_at | 5 | A | 3.9 | A | 11.8 | A | 5.2 | A | --- |
| 256589_at | 3225 | P | 1699.5 | P | 3352.6 | P | 1755.4 | P | CYP81D1 |
| 256590_at | 489.8 | P | 562.6 | P | 547 | P | 589.1 | P | ATHMG |
| 256591_at | 0.9 | A | 2.6 | A | 1.2 | A | 0.6 | A | --- |
| 256592_at | 1 | A | 3.2 | A | 7.1 | A | 1.8 | A | --- |
| 256593_at | 18.2 | A | 20.7 | A | 1.5 | A | 15 | A | --- |
| 256594_at | 1.9 | A | 3.3 | A | 2.4 | A | 3.1 | A | --- |
| 256595_x_at | 149 | P | 111.4 | P | 108.7 | P | 86 | P | --- |
| 256596_at | 165.4 | A | 520.4 | P | 121.6 | P | 310.6 | P | --- |
| 256597_at | 182.3 | M | 149.3 | A | 372.9 | P | 149.8 | P | --- |
| 256598_at | 537.1 | P | 530.2 | P | 583.9 | P | 491.2 | P | BR6OX2 |
| 256599_at | 88.6 | P | 117.4 | P | 49.2 | A | 138.9 | P | --- |
| 256600_at | 74.3 | P | 177.2 | P | 71.3 | A | 143.4 | P | --- |
| 256601_s_at | 3041.2 | P | 320.2 | P | 2204.2 | P | 980.3 | P | AT14A |
| 256602_at | 8.1 | A | 29.2 | A | 10.6 | A | 25.5 | A | --- |
| 256603_at | 1832.2 | P | 321.1 | P | 2956.5 | P | 1176 | P | --- |
| 256604_at | 0.1 | A | 11.3 | A | 4 | A | 1.5 | A | --- |
| 256605_at | 289.4 | P | 276.5 | P | 243.3 | P | 244 | P | --- |
| 256606_at | 5.7 | A | 5.9 | A | 32.9 | A | 7.4 | A | --- |
| 256607_at | 2404.9 | P | 2710.3 | P | 2312.9 | P | 2616.7 | P | --- |
| 256608_at | 2.3 | A | 0.7 | A | 5.8 | A | 1.1 | A | --- |
| 256609_at | 10.7 | A | 14.2 | A | 4.7 | A | 4.3 | A | --- |
| 256610_at | 0.4 | A | 0.7 | A | 15.2 | A | 0.2 | A | --- |
| 256611_at | 381.7 | P | 373.2 | P | 364 | P | 365.5 | P | --- |
| 256612_at | 158.7 | P | 203.1 | P | 145.9 | P | 224 | P | --- |
| 256613_at | 340.6 | P | 257.6 | P | 364.7 | P | 272.2 | P | emb2076 |
| 256614_at | 44 | P | 25.5 | P | 42.2 | P | 41.8 | P | --- |
| 256615_at | 37.7 | A | 33 | A | 35.3 | A | 47.3 | P | --- |
| 256616_at | 133.2 | P | 125.3 | P | 110 | P | 106.1 | P | --- |
| 256617_at | 5438.7 | P | 1855.6 | P | 7040.8 | P | 3006.5 | P | --- |
| 256618_at | 306.7 | P | 223.7 | P | 300.7 | P | 276 | P | VRN5 |
| 256619_at | 42.4 | A | 59.6 | P | 47.3 | A | 63.1 | A | --- |
| 256620_at | 147.8 | P | 131.4 | P | 212.2 | P | 129 | P | --- |
| 256621_at | 14.9 | A | 53.3 | P | 64 | P | 61.2 | P | --- |
| 256622_at | 590.5 | P | 597.9 | P | 597 | P | 564.2 | P | AtHB34 |
| 256623_at | 344 | P | 417.5 | P | 376.7 | P | 432.1 | P | ATM1 |
| 256624_at | 618 | P | 526.5 | P | 468 | P | 467.2 | P | --- |
| 256625_at | 123.1 | P | 149.8 | P | 139.4 | P | 131.2 | P | --- |
| 256626_at | 122.6 | P | 166.8 | P | 153.4 | P | 188.8 | P | --- |
| 256627_at | 238.2 | P | 178.7 | P | 170.2 | P | 201.4 | P | --- |
| 256628_at | 1643.4 | P | 1282.3 | P | 1882.2 | P | 1506.4 | P | TOM40 |
| 256629_at | 1062.5 | P | 1158.2 | P | 1059.8 | P | 1149.9 | P | ATFYPP3 |
| 256630_at | 34.1 | P | 54.1 | P | 63.1 | P | 45.8 | A | --- |
| 256631_at | 40.4 | A | 20.5 | A | 50.5 | A | 32 | A | --- |
| 256632_at | 39.1 | A | 37.1 | M | 72.2 | P | 65.8 | P | --- |
| 256633_at | 212.1 | P | 192.8 | P | 138.9 | P | 170.7 | P | GATL10 |
| 256634_s_at | 2.8 | A | 0.5 | A | 1.1 | A | 0.9 | A | --- |
| 256635_at | 0.3 | A | 0.6 | A | 0.7 | A | 5 | A | --- |
| 256636_at | 11 | A | 3.8 | A | 29.5 | A | 2.8 | A | --- |
| 256637_at | 636 | P | 704.2 | P | 712.6 | P | 655.2 | P | --- |
| 256638_at | 19.9 | A | 64.6 | A | 45.6 | A | 64 | A | --- |
| 256639_at | 10.9 | A | 3.2 | A | 1 | A | 1.1 | A | --- |
| 256640_at | 33.4 | P | 23.3 | A | 42.1 | P | 16.2 | A | --- |
| 256641_at | 71.9 | A | 69.1 | A | 100.5 | A | 48.5 | A | --- |
| 256642_at | 0.5 | A | 0.7 | A | 1.1 | A | 12.8 | A | --- |
| 256643_at | 0.8 | A | 0.5 | A | 0.7 | A | 0.1 | A | --- |
| 256644_at | 4.3 | A | 3.9 | A | 15.4 | A | 3.5 | A | --- |
| 256645_at | 12.8 | A | 75.1 | P | 60.9 | A | 100.3 | P | --- |
| 256646_at | 44.3 | A | 69.7 | A | 89.5 | P | 87.1 | P | --- |
| 256647_at | 54 | A | 186 | P | 67.7 | P | 77.4 | P | --- |
| 256648_at | 2938.7 | P | 2069.9 | P | 3968.7 | P | 2450.3 | P | --- |
| 256649_at | 483.6 | P | 578.5 | P | 554.2 | P | 543.3 | P | SCL30A |
| 256650_at | 270.8 | P | 361 | P | 348.3 | P | 368.3 | P | --- |
| 256651_at | 51.7 | A | 65.6 | P | 65.9 | A | 65.7 | P | --- |
| 256652_at | 147.7 | P | 379.9 | P | 228 | P | 349.3 | P | LPAT5 |
| 256653_at | 207.3 | P | 229.1 | P | 172.7 | P | 171.2 | P | --- |
| 256654_at | 27.7 | A | 57.7 | P | 88.4 | P | 56.3 | P | --- |
| 256655_at | 837.5 | P | 1032.1 | P | 847 | P | 923 | P | --- |
| 256656_at | 38.2 | A | 43.6 | A | 14.4 | A | 4.9 | A | --- |
| 256657_at | 919.5 | P | 950.5 | P | 873 | P | 927.9 | P | --- |
| 256658_at | 495 | P | 700.4 | P | 639.6 | P | 582.7 | P | --- |
| 256659_at | 134.9 | P | 158.4 | P | 129.5 | P | 126.1 | P | --- |
| 256660_at | 15.6 | A | 21.1 | A | 19.6 | A | 3.1 | A | --- |
| 256661_at | 478.1 | P | 402 | P | 490.2 | P | 395.2 | P | --- |
| 256662_at | 1.4 | A | 2 | A | 2.3 | A | 2.8 | A | MS2 |
| 256663_at | 1451.5 | P | 1458.9 | P | 1748.8 | P | 1574.1 | P | --- |
| 256664_at | 105.7 | P | 83.6 | P | 80.7 | P | 98.5 | P | --- |
| 256665_at | 41.4 | M | 40.4 | A | 52.2 | P | 35.3 | A | --- |
| 256666_at | 1377.7 | P | 1889.3 | P | 1401.5 | P | 1929.5 | P | HTA13 |
| 256667_at | 4.2 | A | 3.3 | A | 3.7 | A | 1.4 | A | --- |
| 256668_at | 30.9 | A | 40.2 | A | 48.6 | A | 34.8 | A | --- |

| | | | | | | | | | |
|---|---|---|---|---|---|---|---|---|---|
| 256669_s_at | 4.7 | A | 2.1 | A | 1.3 | A | 9.5 | A | --- |
| 256670_at | 4.2 | A | 6.1 | A | 5 | A | 1.5 | A | --- |
| 256671_at | 171 | P | 242.8 | P | 221.6 | P | 290.5 | P | IQD3 |
| 256672_at | 28.8 | A | 21.1 | A | 38 | A | 23 | A | --- |
| 256673_at | 279.9 | P | 319.3 | P | 305.8 | P | 346.8 | P | FLA15 |
| 256674_at | 508.8 | P | 393.2 | P | 436.8 | P | 483 | P | --- |
| 256675_at | 294 | P | 336.5 | P | 418.2 | P | 320.3 | P | --- |
| 256676_at | 3542.2 | P | 2928.9 | P | 3858.9 | P | 3656.5 | P | SEX4 |
| 256677_at | 477.8 | P | 515.3 | P | 458.5 | P | 515.7 | P | PHF1 |
| 256678_at | 3037.6 | P | 3934.3 | P | 2382.4 | P | 3584 | P | CP33 |
| 256679_at | 6744.1 | P | 6315.7 | P | 4220.6 | P | 5381.5 | P | ATPQ |
| 256680_at | 2032.2 | P | 2087 | P | 2141 | P | 2184.2 | P | --- |
| 256681_at | 856.9 | P | 921.1 | P | 869.7 | P | 969.4 | P | SPP2 |
| 256682_at | 825.9 | P | 746 | P | 820.3 | P | 864.6 | P | LTA3 |
| 256683_at | 898.4 | P | 804.8 | P | 962.1 | P | 835.5 | P | --- |
| 256684_at | 2.1 | A | 1.6 | A | 3.1 | A | 1.6 | A | --- |
| 256685_at | 1.9 | A | 0.5 | A | 1.9 | A | 1.3 | A | --- |
| 256686_at | 9.9 | A | 4.3 | A | 20.7 | A | 26.4 | A | --- |
| 256687_x_at | 3.2 | A | 0.8 | A | 1.8 | A | 0.9 | A | --- |
| 256688_at | 10.3 | A | 0.6 | A | 16.6 | A | 14.1 | A | --- |
| 256689_at | 2.4 | A | 31.4 | A | 6.3 | A | 3.6 | A | --- |
| 256690_at | 3.8 | A | 31.2 | A | 15.9 | A | 13.5 | A | --- |
| 256691_at | 23.4 | A | 23 | A | 27.4 | A | 9.2 | A | --- |
| 256692_at | 5.3 | A | 2.4 | A | 6.2 | A | 5 | A | --- |
| 256693_at | 0.7 | A | 1.4 | A | 1.8 | A | 3 | A | --- |
| 256694_at | 10.9 | A | 16.5 | A | 8.7 | A | 18.9 | A | --- |
| 256695_at | 18.2 | A | 17.6 | A | 39 | A | 4.1 | A | --- |
| 256696_at | 523.8 | P | 457.7 | P | 482.7 | P | 540.6 | P | --- |
| 256697_at | 316.4 | P | 141.8 | P | 301.1 | P | 151.8 | P | AtOCT4 |
| 256698_at | 340.6 | P | 387.5 | P | 276.2 | P | 325.9 | P | --- |
| 256699_at | 1.8 | A | 1.6 | A | 5.8 | A | 3.2 | A | --- |
| 256700_at | 131.1 | P | 141 | P | 188.9 | P | 119.6 | P | --- |
| 256701_at | 15.4 | A | 16.2 | A | 14.1 | A | 3 | A | --- |
| 256702_at | 78.1 | P | 70.2 | P | 74.6 | P | 60.1 | P | --- |
| 256703_at | 8.7 | A | 2.5 | A | 3.1 | A | 1.6 | A | AGL79 |
| 256704_at | 17.9 | A | 39 | A | 10.8 | A | 34.4 | A | --- |
| 256705_at | 38.6 | A | 12.8 | A | 5.1 | A | 1.3 | A | CYP702A8 |
| 256706_at | 534.8 | P | 523.2 | P | 538.8 | P | 626.9 | P | --- |
| 256707_at | 6 | A | 10.7 | A | 1.3 | A | 3.2 | A | --- |
| 256708_at | 18 | A | 6.5 | A | 17.7 | A | 4.5 | A | --- |
| 256709_at | 4 | A | 2.6 | A | 9 | A | 18.8 | A | --- |
| 256710_at | 17.5 | A | 2.7 | A | 7.1 | A | 17.5 | A | --- |
| 256711_at | 4624.5 | P | 4809.4 | P | 4009.6 | P | 4734.6 | P | --- |
| 256712_at | 10.4 | A | 18.8 | A | 16.4 | A | 3.2 | A | --- |
| 256713_at | 27.6 | A | 95.3 | A | 75.7 | A | 76.7 | A | --- |
| 256714_at | 115.5 | P | 37.1 | A | 67.8 | A | 47.5 | A | --- |
| 256715_at | 345.1 | P | 324.5 | P | 354.3 | P | 401.5 | P | MEE18 |
| 256716_at | 1.6 | A | 1 | A | 1.9 | A | 2.1 | A | --- |
| 256717_at | 1.4 | A | 2.2 | A | 4.7 | A | 1.8 | A | --- |
| 256718_at | 0.4 | A | 0.1 | A | 1 | A | 0.1 | A | --- |
| 256719_at | 16.6 | A | 16.6 | A | 2 | A | 12.2 | A | --- |
| 256720_at | 39.4 | A | 40.4 | A | 11.1 | A | 65 | P | --- |
| 256721_at | 89 | P | 84.4 | P | 111.8 | P | 88.4 | P | WAVE1 |
| 256722_at | 266.9 | P | 284.5 | P | 306.8 | P | 310.4 | P | --- |
| 256723_at | 657.6 | P | 701 | P | 653.9 | P | 632 | P | --- |
| 256724_at | 774.5 | P | 769.6 | P | 698.5 | P | 720.9 | P | --- |
| 256725_at | 2406.6 | P | 864.1 | P | 1875.1 | P | 1102.4 | P | --- |
| 256726_at | 17.4 | A | 85.1 | A | 107.1 | A | 85.3 | A | --- |
| 256727_at | 314.2 | P | 384.9 | P | 345.2 | P | 396.2 | P | --- |
| 256728_at | 1439.2 | P | 1843.8 | P | 1200.4 | P | 1819.6 | P | --- |
| 256729_at | 334.1 | P | 327.1 | P | 267.1 | P | 275.6 | P | --- |
| 256730_at | 1.9 | A | 11.2 | A | 1 | A | 5.2 | A | AGL79 |
| 256731_at | 78.2 | A | 106.2 | P | 165 | P | 144.5 | P | --- |
| 256732_at | 4.2 | A | 1.3 | A | 4.8 | A | 13.4 | A | --- |
| 256733_at | 1 | A | 0.8 | A | 2.6 | A | 17.7 | A | --- |
| 256734_at | 248 | P | 302.5 | P | 263 | P | 309.8 | P | RIK |
| 256735_at | 211.7 | P | 311.2 | P | 283.5 | P | 350.4 | P | ATEXO70E1 |
| 256736_at | 29.1 | A | 44.9 | A | 29.6 | A | 37.3 | A | --- |
| 256737_at | 1.1 | A | 0.9 | A | 3.1 | A | 2 | A | --- |
| 256738_at | 3.5 | A | 13.2 | A | 6 | A | 6.3 | A | --- |
| 256739_s_at | 11.6 | A | 7.2 | A | 18.9 | P | 17.8 | P | --- |
| 256740_at | 83.8 | P | 67.1 | P | 83 | P | 72.2 | P | --- |
| 256741_at | 141.3 | P | 211.7 | P | 218.7 | P | 190.2 | P | --- |
| 256742_at | 12 | A | 40.5 | M | 28.5 | A | 18.2 | A | --- |
| 256743_at | 2 | A | 22.2 | A | 11.2 | A | 2.8 | A | --- |
| 256744_at | 916 | P | 1123.8 | P | 938.6 | P | 1019.2 | P | AHP2 |
| 256745_at | 2685.9 | P | 2848.1 | P | 3072.7 | P | 3094.1 | P | --- |
| 256746_at | 2249 | P | 2993.9 | P | 2005 | P | 2837.3 | P | --- |
| 256747_at | 600.5 | P | 775.1 | P | 609 | P | 715.2 | P | --- |
| 256748_x_at | 20.3 | A | 18.3 | A | 48.2 | A | 6.6 | A | --- |
| 256749_at | 2.1 | A | 17.4 | A | 25.8 | A | 3.1 | A | --- |
| 256750_at | 6.6 | A | 29.8 | A | 76.5 | A | 29.8 | A | --- |
| 256751_at | 74.9 | P | 69.7 | P | 105.6 | P | 48 | P | CLC-B |
| 256752_at | 513.3 | P | 566.9 | P | 484 | P | 598 | P | --- |
| 256753_at | 3052.8 | P | 3475.3 | P | 2780.3 | P | 3137.1 | P | GHS1 |
| 256754_at | 2419.9 | P | 2538 | P | 2622.9 | P | 2540.1 | P | CHUP1 |
| 256755_at | 456.6 | P | 615.8 | P | 322.7 | P | 545.4 | P | --- |
| 256756_at | 194 | P | 231.3 | P | 211.4 | P | 172.3 | P | --- |
| 256757_at | 88.7 | P | 56.3 | P | 80.5 | P | 93.6 | P | --- |
| 256758_at | 0.8 | A | 9.7 | A | 1.9 | A | 1 | A | --- |
| 256759_at | 70.9 | P | 97.5 | P | 69.8 | P | 89.6 | P | --- |
| 256760_at | 1.8 | A | 3.6 | A | 4.3 | A | 2.3 | A | ASK15 |
| 256761_at | 29.5 | P | 25.4 | M | 52.9 | A | 44.1 | P | --- |
| 256762_at | 25.6 | P | 26.7 | A | 46.5 | P | 17.8 | A | IDL1 |
| 256763_at | 220.1 | P | 147.5 | P | 161.4 | P | 137 | P | COBL8 |
| 256764_at | 154.4 | P | 136 | P | 115.7 | P | 149.7 | P | --- |
| 256765_at | 11970.3 | P | 5442.2 | P | 12094.5 | P | 6883.3 | P | POP2 |
| 256766_at | 212.5 | P | 790 | P | 399.5 | P | 1751.8 | P | PCC1 |
| 256767_at | 29.4 | A | 14.2 | A | 52.3 | M | 24.7 | A | --- |
| 256768_at | 38.9 | P | 66.4 | P | 81.3 | P | 59.4 | P | LDL2 |
| 256769_at | 408.5 | P | 527.3 | P | 376.9 | P | 557.8 | P | --- |
| 256770_at | 242.1 | P | 198 | P | 261.8 | P | 301.9 | P | --- |
| 256771_at | 81.8 | P | 86.2 | P | 61.3 | M | 61.6 | P | --- |
| 256772_at | 1254.8 | P | 2528.4 | P | 1572.6 | P | 2489.5 | P | BGAL1 |
| 256773_at | 19.1 | A | 13.8 | A | 60.7 | A | 41.3 | A | --- |
| 256774_at | 2.9 | A | 14 | A | 2.2 | A | 1.6 | A | --- |
| 256775_at | 56.7 | P | 58.7 | P | 63 | P | 51.9 | P | --- |
| 256776_at | 1402.9 | P | 1663.7 | P | 1432.2 | P | 1699.2 | P | --- |
| 256777_at | 161.4 | P | 94.1 | P | 126.8 | P | 111.1 | P | --- |

| | | | | | | | | | |
|---|---|---|---|---|---|---|---|---|---|
| 256778_at | 51.8 | A | 33.8 | A | 39.1 | A | 6.6 | A | NAP1;4 |
| 256779_at | 41.8 | P | 31.8 | P | 52.7 | P | 20.4 | A | AtcwINV5 |
| 256780_at | 12.4 | A | 36.9 | A | 36.1 | A | 14.2 | A | ATRLI1 |
| 256781_at | 32.1 | A | 84.1 | P | 17.9 | A | 110.8 | A | --- |
| 256782_at | 36.1 | A | 21.9 | A | 38.7 | A | 35.4 | A | --- |
| 256783_at | 1668.9 | P | 1604.3 | P | 1624.1 | P | 1628.9 | P | --- |
| 256784_at | 113.2 | P | 152 | P | 106.6 | P | 134.4 | P | --- |
| 256785_at | 832.7 | P | 773.2 | P | 586.6 | P | 723.3 | P | PRA8 |
| 256786_at | 469.1 | P | 652.5 | P | 495 | P | 584 | P | --- |
| 256787_at | 836.1 | P | 441.2 | P | 463 | P | 356.1 | P | ATBFRUCT1 |
| 256788_at | 104 | P | 114.7 | P | 123.6 | P | 113.3 | P | CYP90D1 |
| 256789_at | 158.8 | P | 215.8 | P | 149 | P | 210 | P | --- |
| 256790_at | 268.6 | P | 276.7 | P | 308.2 | P | 236.6 | P | ARR1 |
| 256791_at | 15.6 | A | 23.5 | A | 46.2 | A | 11.6 | A | --- |
| 256792_at | 227.8 | P | 407.1 | P | 294.6 | P | 397.9 | P | --- |
| 256793_at | 465.9 | P | 479 | P | 481.8 | A | 432.1 | P | --- |
| 256794_at | 3706 | P | 3194.8 | P | 4155.2 | P | 3512.2 | P | --- |
| 256795_at | 2148.9 | P | 2359.5 | P | 2145.9 | P | 2411.8 | P | PAC1 |
| 256796_at | 483.7 | P | 3001.9 | P | 542.6 | P | 2265.6 | P | --- |
| 256797_at | 1160.5 | P | 720.2 | P | 1086.9 | P | 897.9 | P | --- |
| 256798_at | 45.9 | A | 59.6 | P | 49.6 | A | 79.4 | P | --- |
| 256799_at | 77 | P | 59 | P | 59.5 | A | 99.6 | P | --- |
| 256800_at | 40.5 | A | 52.3 | A | 29.3 | A | 5.2 | A | --- |
| 256801_at | 1.2 | A | 1.4 | A | 2.3 | A | 4.5 | A | CYP705A30 |
| 256802_at | 1.6 | A | 0.8 | A | 27.8 | A | 5.5 | A | CYP705A32 |
| 256803_at | 46.2 | P | 33.8 | P | 41.5 | A | 26.1 | A | CYP705A33 |
| 256804_at | 758.8 | P | 756.2 | P | 805.6 | P | 821.2 | P | --- |
| 256805_at | 430.7 | P | 607.3 | P | 436.3 | P | 554.1 | P | --- |
| 256806_at | 172.4 | P | 209.6 | P | 213.7 | P | 191.5 | P | NF-YA9 |
| 256807_at | 776.8 | P | 605.8 | P | 760 | P | 643.2 | P | NFU4 |
| 256808_at | 321.9 | P | 230.1 | P | 280.5 | P | 224.4 | P | ALY3 |
| 256809_at | 47.4 | A | 74.8 | A | 57.7 | A | 62.9 | A | ALY3 |
| 256810_at | 142.3 | P | 162 | P | 149 | P | 215.8 | P | --- |
| 256811_at | 23.9 | A | 19.9 | A | 1.5 | A | 5.7 | A | --- |
| 256812_at | 513.9 | P | 569.1 | P | 565.9 | P | 557.6 | P | --- |
| 256813_at | 262.2 | P | 180.3 | P | 223 | P | 219.4 | P | --- |
| 256814_at | 22.8 | A | 1.9 | A | 14.6 | A | 5.1 | A | BGLU19 |
| 256815_at | 2.6 | A | 3.5 | A | 8.3 | A | 5.1 | A | --- |
| 256816_at | 213.1 | P | 212.9 | P | 244.3 | P | 181.7 | P | --- |
| 256817_at | 6 | A | 4.3 | A | 11.4 | A | 5.1 | A | --- |
| 256818_at | 914.4 | P | 389.7 | P | 692.8 | P | 429.3 | P | --- |
| 256819_at | 300 | P | 439.7 | P | 317.1 | P | 418.1 | P | --- |
| 256820_at | 435.5 | P | 454.8 | P | 423.4 | P | 482 | P | FHY3 |
| 256821_at | 65.6 | P | 114.2 | P | 54.3 | P | 110.5 | P | --- |
| 256822_at | 608.4 | P | 456.9 | P | 522.9 | P | 507.2 | P | IQD5 |
| 256823_at | 256.2 | P | 240.8 | P | 262.5 | P | 237.6 | P | --- |
| 256824_at | 39.9 | A | 65.6 | P | 88.6 | P | 55 | A | --- |
| 256825_at | 6258 | P | 7178.4 | P | 7682.9 | P | 6346.6 | P | CWLP |
| 256826_at | 43.5 | A | 44 | A | 33 | A | 31 | A | --- |
| 256827_at | 14.8 | A | 25 | A | 19.9 | A | 8.3 | A | --- |
| 256828_at | 766.2 | P | 1004.3 | P | 737.3 | P | 862.4 | P | --- |
| 256829_at | 595.5 | P | 605.3 | P | 690.5 | P | 532.8 | P | --- |
| 256830_at | 16.7 | P | 27.8 | A | 66.9 | P | 28.5 | A | TIF3C2 |
| 256831_at | 18.6 | A | 35.4 | A | 32 | A | 41.7 | A | --- |
| 256832_at | 205.4 | P | 523.8 | P | 215.4 | P | 483.8 | P | DMC1 |
| 256833_at | 111.8 | P | 229.9 | P | 29.4 | A | 103.7 | P | --- |
| 256834_at | 4.7 | A | 4.3 | A | 28.9 | A | 6 | A | --- |
| 256835_at | 9146.2 | P | 6920.6 | P | 7363.3 | P | 6929.6 | P | APS1 |
| 256836_at | 2666.3 | P | 2619.2 | P | 3374.6 | P | 2843.2 | P | PKP-ALPHA |
| 256837_at | 90.3 | P | 127.8 | P | 126.8 | P | 101.7 | P | NRPD7 |
| 256838_at | 1373.7 | P | 1441.6 | P | 1456.1 | P | 1439.4 | P | ATARFC1 |
| 256839_at | 80.6 | P | 75.4 | P | 76.2 | P | 66.7 | P | --- |
| 256840_x_at | 2.8 | A | 1.5 | A | 2.7 | A | 6.4 | A | --- |
| 256841_x_at | 7.7 | A | 10.2 | A | 16.4 | A | 2.9 | A | --- |
| 256842_at | 4.1 | A | 7.8 | A | 16.6 | A | 10.2 | A | --- |
| 256843_at | 1.7 | A | 2.2 | A | 5.7 | A | 1.8 | A | --- |
| 256844_s_at | 0.5 | A | 1.9 | A | 2.3 | A | 1.9 | A | --- |
| 256845_x_at | 8.4 | A | 33.3 | A | 36.8 | A | 34.8 | A | --- |
| 256846_at | 3.9 | A | 0.4 | A | 1 | A | 0 | A | --- |
| 256847_at | 24.3 | A | 20.4 | A | 17.9 | A | 16.2 | A | --- |
| 256848_at | 516.9 | P | 280.2 | P | 417.8 | P | 351.5 | P | --- |
| 256849_at | 6.6 | A | 1.8 | A | 14.5 | A | 12 | A | --- |
| 256850_at | 17.3 | A | 5.9 | A | 5 | A | 2.3 | A | --- |
| 256851_at | 236.8 | P | 265.6 | P | 236.1 | P | 252.2 | P | --- |
| 256852_at | 97.2 | A | 29.2 | A | 141.7 | A | 37.7 | A | ATRANGAP1 |
| 256853_at | 271.8 | P | 288.4 | P | 302.4 | P | 248.8 | P | --- |
| 256854_at | 366.9 | P | 241 | P | 318.9 | P | 287.6 | P | --- |
| 256855_at | 6347.1 | P | 9045.3 | P | 5835 | P | 8871.6 | P | --- |
| 256856_at | 890.5 | P | 1339.3 | P | 988.8 | P | 1157.3 | P | --- |
| 256857_at | 3.8 | A | 4.3 | A | 4.9 | A | 19.7 | A | CUC1 |
| 256858_at | 200.4 | P | 178.7 | P | 213.5 | P | 219.5 | P | --- |
| 256859_at | 14.4 | A | 2.3 | A | 10.3 | A | 2.4 | A | --- |
| 256860_at | 158.1 | P | 142.5 | P | 203.7 | P | 155.1 | P | --- |
| 256861_at | 1371.5 | P | 1514.7 | P | 1427.1 | P | 1415.8 | P | BAM1 |
| 256862_at | 2174.2 | P | 1541.9 | P | 2637.1 | P | 1954.4 | P | --- |
| 256863_at | 140.4 | P | 149.3 | P | 187.6 | P | 209.6 | P | --- |
| 256864_at | 131.2 | P | 269.4 | P | 261.3 | P | 315.2 | P | TOPII |
| 256865_at | 1918.6 | P | 3697 | P | 1789.8 | P | 4096.5 | P | GAE6 |
| 256866_at | 728.4 | P | 805.1 | P | 653.4 | P | 636.5 | P | --- |
| 256867_at | 15.4 | A | 2.8 | A | 3 | A | 7.9 | A | --- |
| 256868_at | 333.5 | P | 286.9 | P | 311.1 | P | 303.6 | P | EIF4B1 |
| 256869_at | 310.4 | P | 196.2 | P | 212.5 | P | 239.2 | P | --- |
| 256870_at | 553.4 | P | 487.5 | P | 692.7 | P | 532.6 | P | CYP71B34 |
| 256871_at | 36.5 | A | 52.2 | P | 62.3 | P | 48.7 | P | --- |
| 256872_at | 60.6 | P | 37.1 | P | 67.4 | M | 37.2 | P | --- |
| 256873_at | 139.3 | P | 189.4 | P | 191.6 | P | 150.8 | P | CYP71B34 |
| 256874_at | 61.5 | P | 85.7 | P | 66.5 | P | 76.7 | P | CYP71B36 |
| 256875_at | 117.1 | P | 57.2 | P | 63.5 | P | 39.9 | P | CYP71B37 |
| 256876_at | 261.5 | P | 259.1 | P | 332.3 | P | 303.4 | P | --- |
| 256877_at | 464.2 | P | 397.8 | P | 352.2 | P | 230.7 | P | --- |
| 256878_at | 7.8 | A | 11 | A | 28.1 | A | 30.4 | P | --- |
| 256879_at | 566.3 | P | 603.7 | P | 452.6 | P | 589.4 | P | --- |
| 256880_at | 3831.9 | P | 3392.4 | P | 2641.8 | P | 3580.9 | P | --- |
| 256881_at | 284.8 | P | 236.9 | P | 396.3 | P | 246.7 | P | --- |
| 256882_at | 1561.4 | P | 1566.3 | P | 1849 | P | 2060.5 | P | --- |
| 256883_at | 349.6 | P | 160.4 | P | 343.1 | P | 169.5 | P | --- |
| 256884_at | 21.1 | A | 20.5 | A | 11.7 | A | 6.5 | A | --- |
| 256885_at | 403.2 | P | 323.6 | P | 342.6 | P | 336.9 | P | --- |
| 256886_at | 28.9 | P | 31.5 | P | 44.7 | P | 14.3 | A | --- |

| | | | | | | | | | |
|---|---|---|---|---|---|---|---|---|---|
| 256887_at | 124 | P | 131 | P | 163 | P | 133.2 | P | --- |
| 256888_at | 438 | P | 496 | P | 486.9 | P | 471 | P | --- |
| 256889_at | 603.1 | P | 466.1 | P | 469.9 | P | 591.8 | P | ING1 |
| 256890_at | 1158.6 | P | 1024.7 | P | 1853.7 | P | 1307.7 | P | GRP4 |
| 256891_at | 187 | P | 63.6 | A | 121.3 | P | 59.5 | P | --- |
| 256892_at | 927.5 | P | 1012.7 | P | 1109.3 | P | 943.7 | P | --- |
| 256893_x_at | 15.8 | A | 2.1 | A | 9.1 | A | 2.9 | A | --- |
| 256894_at | 146 | P | 220.4 | P | 178.2 | P | 215.5 | P | CYCP2;1 |
| 256895_at | 45.8 | A | 36.1 | A | 54.8 | A | 16.4 | A | --- |
| 256896_at | 29.7 | A | 64.7 | P | 27.2 | A | 25 | P | --- |
| 256897_at | 14.6 | A | 8.3 | A | 18.3 | A | 9.3 | A | --- |
| 256898_at | 4.6 | A | 24 | A | 22.8 | A | 36.8 | A | ABI3 |
| 256899_at | 59 | P | 109.6 | P | 98 | P | 111 | P | TMKL1 |
| 256900_at | 115.6 | P | 230.5 | P | 182.5 | P | 169.9 | P | --- |
| 256901_at | 3 | A | 2.8 | A | 5.7 | A | 0.6 | A | --- |
| 256902_s_at | 269.5 | P | 426.9 | P | 314 | P | 514.5 | P | --- |
| 256903_at | 17.5 | A | 23.1 | A | 52.5 | P | 15.4 | A | --- |
| 256904_at | 399.4 | P | 373.7 | P | 438.3 | P | 329.5 | P | --- |
| 256905_at | 3213.5 | P | 2232.3 | P | 3613.9 | P | 2755.8 | P | HSP60 |
| 256906_at | 272.7 | P | 153.5 | P | 395.6 | P | 209 | P | --- |
| 256907_at | 386.6 | P | 426.3 | P | 460.4 | P | 356.1 | P | --- |
| 256908_at | 425 | P | 548 | P | 471 | P | 473.8 | P | --- |
| 256909_at | 2.5 | A | 4.6 | A | 23.9 | A | 6.1 | A | --- |
| 256910_at | 1016.2 | P | 486.7 | P | 1118.7 | P | 654.4 | P | --- |
| 256911_at | 73.1 | P | 109.7 | P | 84.6 | P | 104.3 | P | --- |
| 256912_at | 2.9 | A | 8.4 | A | 31.8 | A | 10.7 | A | --- |
| 256913_at | 62.6 | P | 50.7 | P | 64.5 | P | 28.7 | P | --- |
| 256914_at | 101.6 | P | 102.2 | P | 115.3 | P | 82 | P | --- |
| 256915_at | 423.6 | P | 443.8 | P | 488.5 | P | 486.6 | P | --- |
| 256916_at | 991.6 | P | 1072.8 | P | 1064.9 | P | 1015.3 | P | --- |
| 256917_at | 134.1 | P | 179.1 | P | 147.1 | A | 153.1 | P | --- |
| 256918_s_at | 109.5 | P | 221 | P | 173.2 | P | 278.6 | P | --- |
| 256919_at | 65.7 | P | 78 | P | 70.2 | P | 84.2 | P | --- |
| 256920_at | 214.1 | P | 274 | P | 196.8 | P | 276.1 | P | --- |
| 256921_at | 413.5 | P | 497.9 | P | 599.6 | P | 527.6 | P | --- |
| 256922_at | 920.8 | P | 1077.7 | P | 553.7 | P | 944.8 | P | --- |
| 256923_at | 25.1 | A | 29 | A | 52.2 | A | 31.7 | A | --- |
| 256924_at | 363 | P | 79.5 | P | 486.5 | P | 223.9 | P | AT5MAT |
| 256925_at | 0.6 | A | 18.1 | A | 4.8 | A | 13.4 | A | --- |
| 256926_at | 32.9 | P | 57.8 | P | 53 | P | 46.2 | P | --- |
| 256927_at | 841.1 | P | 593.6 | P | 945.7 | P | 648.9 | P | --- |
| 256928_at | 572.2 | P | 407 | P | 631.1 | P | 528.5 | P | --- |
| 256929_at | 36.1 | A | 40.4 | P | 40.6 | A | 25.6 | A | --- |
| 256930_at | 124.5 | P | 109.5 | P | 79.3 | P | 90.7 | P | OASA2 |
| 256931_at | 4 | A | 2.6 | A | 2.2 | A | 3.7 | A | --- |
| 256932_at | 326.4 | P | 410.6 | P | 306.9 | P | 432.4 | P | --- |
| 256933_at | 137.7 | P | 244.8 | P | 119.5 | P | 175 | P | --- |
| 256934_at | 854.9 | P | 730.9 | P | 849 | P | 781.7 | P | --- |
| 256935_at | 7.9 | A | 7.7 | A | 5.2 | A | 23.1 | A | --- |
| 256936_at | 626.8 | P | 661.2 | P | 759.9 | P | 643.8 | P | PDF2 |
| 256937_at | 38.1 | A | 125.1 | P | 105.9 | A | 120.6 | P | --- |
| 256938_at | 1.1 | A | 20.3 | A | 23.3 | A | 3.5 | A | ATECP31 |
| 256939_at | 2038 | P | 2208.4 | P | 2541.7 | P | 2337.9 | P | PBD1 |
| 256940_at | 420 | P | 223.5 | P | 487.2 | P | 276.9 | P | QQS |
| 256941_at | 28.6 | A | 4.4 | A | 25.9 | A | 4.1 | A | --- |
| 256942_at | 222.8 | P | 344.8 | P | 285.8 | P | 380.9 | P | LSH4 |
| 256943_at | 182.4 | P | 143.6 | P | 162.1 | P | 166.8 | P | --- |
| 256944_at | 401.8 | P | 439.7 | P | 463.9 | P | 451.5 | P | VRN1 |
| 256945_at | 5.3 | A | 21.7 | A | 2.8 | A | 14.2 | A | --- |
| 256946_at | 19.2 | A | 33.4 | A | 71.9 | A | 32.2 | A | HAF2 |
| 256947_at | 2.8 | A | 4.7 | A | 1.8 | A | 4.5 | A | POK2 |
| 256948_at | 233.9 | P | 198.1 | P | 190.5 | P | 275.8 | P | --- |
| 256949_at | 1 | A | 0.8 | A | 8.8 | A | 0.9 | A | --- |
| 256950_at | 32.2 | A | 56.5 | A | 86.3 | A | 49.1 | P | --- |
| 256951_at | 30.3 | A | 15.2 | A | 39.8 | A | 4.9 | A | --- |
| 256952_at | 211.7 | P | 274.1 | P | 308 | P | 218.9 | P | --- |
| 256953_at | 25.6 | A | 32.8 | A | 56.5 | A | 37 | A | --- |
| 256954_at | 396.1 | P | 405.8 | P | 293 | P | 317.4 | P | FUS9 |
| 256955_at | 30.3 | A | 23.4 | A | 28.8 | A | 24.7 | A | sks11 |
| 256956_at | 4462.9 | P | 4853.2 | P | 4538.3 | P | 5297 | P | --- |
| 256957_at | 24 | P | 42.9 | A | 47.6 | P | 20.5 | A | --- |
| 256958_at | 196.4 | P | 192.7 | P | 224.1 | P | 185.5 | P | --- |
| 256959_at | 512.7 | P | 380.5 | P | 501.8 | P | 401.8 | P | --- |
| 256960_at | 277.3 | P | 384.8 | P | 404.1 | P | 404.7 | P | --- |
| 256961_at | 333.9 | P | 346.4 | P | 250.7 | P | 272.4 | P | TBP1 |
| 256962_at | 320 | P | 549 | P | 444.3 | P | 613.1 | P | --- |
| 256963_at | 571.7 | P | 576.8 | P | 713.6 | P | 619.8 | P | OVA5 |
| 256964_at | 3672.8 | P | 4316.5 | P | 3897.2 | P | 4532.8 | P | AGP12 |
| 256965_at | 660.4 | P | 496.2 | P | 578 | P | 466.9 | P | DIN4 |
| 256966_at | 19.1 | A | 40 | A | 38.9 | A | 25.6 | A | sks13 |
| 256967_at | 257.1 | P | 332.2 | P | 258.5 | P | 320.6 | P | --- |
| 256968_at | 125 | P | 118.3 | P | 119.1 | P | 104.1 | P | NADK1 |
| 256969_at | 47.7 | A | 72.9 | A | 17.1 | A | 71.1 | A | --- |
| 256970_at | 63.4 | A | 60.6 | A | 80.7 | A | 52.5 | A | --- |
| 256971_at | 2.9 | A | 37.7 | A | 67.4 | A | 44.9 | A | --- |
| 256972_at | 458.7 | P | 509.4 | P | 589.6 | P | 494.6 | P | --- |
| 256973_at | 21.8 | A | 10.5 | A | 30.5 | A | 32.7 | A | --- |
| 256974_at | 11.5 | A | 1.5 | A | 2.5 | A | 1.8 | A | --- |
| 256975_at | 21.1 | A | 19.9 | A | 3.8 | A | 0.8 | A | --- |
| 256976_at | 100.6 | P | 70.7 | P | 137.8 | P | 67.6 | P | --- |
| 256977_at | 14.6 | A | 9.6 | A | 8.3 | A | 7.1 | A | --- |
| 256978_at | 410.1 | P | 336.4 | P | 432.4 | P | 354.5 | P | PUR7 |
| 256979_at | 8031.5 | P | 10612.2 | P | 8003.4 | P | 8540.8 | P | PSBTN |
| 256980_at | 94.8 | P | 111.6 | P | 91.4 | P | 112.9 | P | DRB3 |
| 256981_at | 293.3 | P | 269.3 | P | 351.6 | P | 254.9 | P | BRL3 |
| 256982_at | 2865.4 | P | 2477.3 | P | 3351.3 | P | 2567.8 | P | ECT2 |
| 256983_at | 2879 | P | 4750.6 | P | 3287.9 | P | 4789.1 | P | --- |
| 256984_at | 470.6 | P | 620.7 | P | 525.2 | P | 619 | P | --- |
| 256985_at | 46.4 | A | 43.2 | A | 60.4 | A | 55.2 | A | ATMYB5 |
| 256986_at | 361 | P | 444.6 | P | 384.3 | P | 489 | P | --- |
| 256987_at | 1.4 | A | 3.4 | A | 2.7 | A | 1.5 | A | --- |
| 256988_s_at | 2717.2 | P | 2700 | P | 2156.4 | P | 2454.1 | P | --- |
| 256989_at | 64.6 | P | 60.2 | P | 29 | A | 41 | P | --- |
| 256990_at | 172.2 | A | 135.9 | A | 114.6 | A | 170.9 | M | --- |
| 256991_at | 53.3 | P | 3.3 | A | 44.5 | A | 23.5 | A | --- |
| 256992_at | 18.4 | A | 27 | P | 37.1 | A | 12.3 | A | --- |
| 256993_at | 174.7 | P | 234.9 | P | 232.6 | P | 214.6 | P | --- |
| 256994_s_at | 48.6 | A | 59.7 | P | 94 | A | 58.9 | P | ATTPS-CIN |
| 256995_at | 415.7 | P | 445 | P | 391.7 | P | 400.7 | P | --- |

| | | | | | | | | | |
|---|---|---|---|---|---|---|---|---|---|
| 256996_at | 21.8 | A | 28.6 | P | 9.3 | A | 25.8 | A | --- |
| 256997_at | 1197.7 | P | 1714.4 | P | 922.7 | P | 1665.5 | P | --- |
| 256998_at | 465 | P | 351.9 | P | 473.9 | P | 350.2 | P | --- |
| 256999_at | 331.6 | P | 543 | P | 320.7 | P | 430.8 | P | --- |
| 257000_at | 305.5 | P | 318.8 | P | 310.9 | P | 278.8 | P | --- |
| 257001_at | 403.3 | P | 547.6 | P | 405.2 | P | 577.7 | P | --- |
| 257002_at | 3468.8 | P | 2891.2 | P | 3021.9 | P | 3101.2 | P | --- |
| 257003_at | 2238.5 | P | 2375.1 | P | 2601.5 | P | 2101.2 | P | FLU |
| 257004_s_at | 343.9 | P | 312.4 | P | 320.4 | P | 298.9 | P | --- |
| 257005_at | 211.4 | P | 306.2 | P | 205.1 | P | 403 | P | --- |
| 257006_at | 234.8 | P | 243.5 | P | 158.8 | P | 281.8 | P | --- |
| 257007_at | 590.8 | P | 519 | P | 522.1 | P | 563.3 | P | --- |
| 257008_at | 15894.4 | P | 16463.6 | P | 17796.6 | P | 17487.2 | P | ESM1 |
| 257009_at | 135.5 | P | 99.6 | P | 174.2 | P | 132.2 | P | --- |
| 257010_at | 363 | P | 308.5 | P | 297.1 | P | 274.3 | P | ATEXO70D3 |
| 257011_at | 108 | A | 50.7 | A | 79.7 | A | 74.4 | A | CAX9 |
| 257012_at | 4.6 | A | 3.6 | A | 31.9 | A | 5.8 | A | TEL1 |
| 257013_at | 85.4 | A | 56.5 | A | 111.6 | A | 72.6 | A | --- |
| 257014_at | 26.8 | A | 32.8 | A | 70.2 | M | 24.2 | A | --- |
| 257015_at | 17.2 | A | 38.7 | A | 36.2 | A | 28.1 | A | --- |
| 257016_at | 379.4 | P | 366.3 | P | 277.9 | P | 395.2 | P | --- |
| 257017_at | 168.2 | P | 151.3 | P | 252.2 | P | 186.3 | P | --- |
| 257018_at | 432.9 | P | 367.1 | P | 564.1 | P | 380.5 | P | --- |
| 257019_at | 942.6 | P | 735.1 | P | 825 | P | 664.3 | P | --- |
| 257020_at | 278.7 | P | 450.7 | P | 274.8 | P | 364 | P | --- |
| 257021_at | 9868.1 | P | 7109.4 | P | 13499 | P | 6067.6 | P | BCAT4 |
| 257022_at | 214.6 | P | 126.3 | P | 169 | P | 103.7 | P | AZF2 |
| 257023_at | 2157.1 | P | 2189.4 | P | 2554.1 | P | 2359.7 | P | --- |
| 257024_at | 763.5 | P | 530.6 | P | 944 | P | 700.2 | P | --- |
| 257025_at | 158.3 | P | 155.3 | P | 125.2 | P | 174.5 | P | ATG2 |
| 257026_at | 32.9 | A | 43.2 | A | 59.9 | M | 56.5 | P | --- |
| 257027_at | 40.1 | A | 54.4 | P | 57.5 | A | 46.2 | A | ATRAD54 |
| 257028_at | 32 | A | 32 | A | 52.4 | P | 30.3 | P | --- |
| 257029_at | 405.5 | P | 475 | P | 300.5 | P | 407.6 | P | --- |
| 257030_at | 82.2 | P | 41.6 | A | 67.6 | A | 73.8 | A | --- |
| 257031_at | 74.2 | P | 95.5 | P | 92.5 | P | 89 | P | FUT11 |
| 257032_at | 33.3 | P | 17.5 | A | 26.4 | A | 22.5 | P | --- |
| 257033_at | 4474.3 | P | 3972.2 | P | 3798.7 | P | 3927.4 | P | ATPREP1 |
| 257034_at | 2.1 | A | 4 | A | 37.2 | A | 1.8 | A | --- |
| 257035_at | 86.9 | P | 115.4 | P | 141 | P | 110.4 | P | CYP707A4 |
| 257036_at | 123.4 | P | 145.5 | P | 132.2 | P | 203.9 | P | --- |
| 257037_at | 1103.6 | P | 1135.6 | P | 1314 | P | 1039.3 | P | ATRBP47B |
| 257038_at | 556.7 | P | 539.6 | P | 472.2 | P | 498.5 | P | LOH2 |
| 257039_at | 51.3 | P | 17.9 | A | 30.1 | A | 51 | P | ATIPT8 |
| 257040_at | 261.2 | P | 276.3 | P | 316.6 | P | 295.8 | P | SCO2 |
| 257041_at | 1.5 | A | 0.7 | A | 4.1 | A | 1 | A | --- |
| 257042_s_at | 137.9 | P | 83.4 | P | 120.1 | P | 103.9 | P | --- |
| 257043_at | 29.5 | A | 8.1 | A | 21.3 | A | 26.1 | A | IKU2 |
| 257044_at | 817.4 | P | 1194.7 | P | 803.1 | P | 1118.2 | P | ARC5 |
| 257045_at | 72.1 | P | 146.7 | P | 145.8 | P | 153.1 | P | ARC5 |
| 257046_at | 736.8 | P | 674.3 | P | 683.1 | P | 708.2 | P | --- |
| 257047_at | 109.6 | P | 202.7 | P | 161.7 | P | 199.5 | P | --- |
| 257048_at | 54.4 | A | 85.3 | P | 101.3 | A | 72.8 | M | --- |
| 257049_at | 23.1 | A | 3.5 | A | 33.7 | A | 11.8 | A | --- |
| 257050_at | 441.9 | P | 403.7 | P | 304.8 | P | 376.4 | P | --- |
| 257051_at | 17.3 | A | 32.7 | A | 9.5 | A | 16.9 | A | SPL5 |
| 257052_at | 545.5 | P | 465.5 | P | 580.1 | P | 495 | P | --- |
| 257053_at | 528.1 | P | 504.2 | P | 459.6 | P | 496.5 | P | ERF4 |
| 257054_at | 13039.5 | P | 15846.7 | P | 16512.5 | P | 16613.3 | P | MT3 |
| 257055_at | 301.2 | P | 252.9 | P | 248.5 | P | 241.2 | P | --- |
| 257056_at | 595.9 | P | 727 | P | 587.4 | P | 772.5 | P | --- |
| 257057_at | 71.4 | P | 56.3 | P | 212.9 | P | 91.8 | P | --- |
| 257058_at | 547.2 | P | 330.3 | P | 461.4 | P | 469.1 | P | COX17 |
| 257059_at | 17.5 | A | 19.9 | A | 37.4 | P | 15 | A | --- |
| 257060_at | 332.2 | P | 368 | P | 269.4 | P | 285.6 | P | --- |
| 257061_at | 36.5 | A | 205.8 | P | 14.9 | A | 62.3 | A | --- |
| 257062_at | 793.5 | P | 1847.5 | P | 2108.2 | P | 1935.6 | P | EMB2454 |
| 257063_s_at | 306.2 | P | 312.4 | P | 324.2 | P | 470.4 | P | --- |
| 257064_at | 34 | P | 42.2 | P | 53.2 | P | 42.8 | P | --- |
| 257065_at | 1.8 | A | 2.1 | A | 5.1 | A | 1.9 | A | --- |
| 257066_at | 654.4 | P | 1583.6 | P | 928.6 | P | 1441.8 | P | --- |
| 257067_at | 371.1 | P | 442.8 | P | 374.1 | P | 459 | P | CYP77A5P |
| 257068_at | 1 | A | 15.6 | A | 4.8 | A | 15.7 | A | --- |
| 257069_at | 24.4 | A | 3.7 | A | 32.8 | A | 18.8 | A | --- |
| 257070_at | 6 | A | 21.5 | A | 23 | A | 3.1 | A | --- |
| 257071_at | 1852.3 | P | 1255.3 | P | 1246.9 | P | 1219.2 | P | ATCSLC04 |
| 257072_at | 1487.4 | P | 1712.9 | P | 1224.5 | P | 1561.5 | P | --- |
| 257073_at | 206.2 | P | 160.8 | P | 218.1 | P | 221.7 | P | --- |
| 257074_at | 335.3 | P | 244.9 | P | 203.3 | P | 275.2 | P | --- |
| 257075_at | 142.7 | P | 169.5 | P | 165.4 | P | 150.8 | P | --- |
| 257076_at | 423.6 | P | 461.3 | P | 435.6 | P | 378.2 | P | --- |
| 257077_at | 10.5 | A | 38.4 | A | 26 | A | 1.4 | A | --- |
| 257078_at | 176.2 | P | 141.1 | P | 192.8 | P | 155.6 | P | --- |
| 257079_at | 1.9 | A | 17.6 | A | 1.9 | A | 1.7 | A | --- |
| 257080_at | 2.4 | A | 4.7 | A | 4.7 | A | 21 | A | --- |
| 257081_at | 92.9 | P | 97.4 | P | 131 | M | 94.5 | P | --- |
| 257082_at | 11 | A | 3.7 | A | 28.7 | A | 4.5 | A | COBL10 |
| 257083_s_at | 119.8 | P | 146.6 | P | 121.5 | P | 132.3 | P | --- |
| 257084_at | 293.5 | P | 317.8 | P | 247.7 | P | 296.8 | P | --- |
| 257085_at | 459.6 | P | 503.5 | P | 462.2 | P | 511.8 | P | UBP14 |
| 257086_at | 100.6 | P | 145 | P | 123.4 | P | 125.7 | P | --- |
| 257087_at | 1403.8 | P | 1958.6 | P | 1359.7 | P | 1750.1 | P | PAP18 |
| 257088_at | 1225.2 | P | 989.6 | P | 883.8 | P | 1058.8 | P | --- |
| 257089_at | 7.4 | A | 13.8 | A | 7.8 | A | 10.9 | A | SVL3 |
| 257090_at | 6.1 | A | 1.8 | A | 1.9 | A | 4.7 | A | --- |
| 257091_at | 6.8 | A | 52.4 | A | 63.8 | A | 38.2 | M | POLGAMMA1 |
| 257092_at | 311.8 | P | 346.2 | P | 281.9 | P | 309 | P | DDL |
| 257093_at | 155.6 | P | 333.9 | P | 306.8 | P | 343.7 | P | --- |
| 257094_at | 67.7 | A | 58.6 | M | 55.4 | A | 49.6 | A | --- |
| 257095_at | 177.4 | P | 280.9 | P | 236.4 | P | 229.3 | P | ATPDIL5-3 |
| 257096_at | 23.5 | A | 37.6 | A | 9.6 | A | 36.7 | A | --- |
| 257097_at | 20 | A | 10 | A | 25.5 | A | 24.8 | A | --- |
| 257098_at | 3.8 | A | 12.1 | A | 7 | A | 4.2 | A | --- |
| 257099_s_at | 555 | P | 78.2 | P | 206 | P | 123.3 | P | --- |
| 257100_at | 188.9 | P | 365.4 | P | 119 | P | 263.2 | P | AtRLP41 |
| 257101_at | 69.9 | P | 71.7 | P | 44.2 | A | 62.4 | P | AtRLP42 |
| 257102_at | 11.9 | A | 3.3 | A | 3.8 | A | 1.9 | A | XTH3 |
| 257103_at | 14.3 | A | 21.5 | A | 4.8 | A | 6.8 | A | --- |
| 257104_at | 1242.9 | P | 1582.2 | P | 1342.5 | P | 1606.4 | P | --- |

| | | | | | | | | | |
|---|---|---|---|---|---|---|---|---|---|
| 257105_at | 98.7 | P | 81.5 | P | 89 | P | 79.9 | P | --- |
| 257106_at | 23.2 | A | 7.9 | A | 3.4 | A | 15.6 | A | --- |
| 257107_at | 3.9 | A | 30.4 | A | 21.3 | A | 10.2 | A | --- |
| 257108_x_at | 14.8 | A | 14.6 | A | 3.8 | A | 14.1 | A | --- |
| 257109_s_at | 31.5 | A | 30.4 | A | 6.4 | A | 2.2 | A | --- |
| 257110_x_at | 23 | A | 10.6 | A | 31.3 | A | 8.2 | A | --- |
| 257111_x_at | 1.7 | A | 10.3 | A | 32.4 | A | 3.4 | A | --- |
| 257112_at | 222.8 | P | 232 | P | 220.4 | P | 200.2 | P | CYP705A21 |
| 257113_at | 15.2 | A | 30.3 | A | 29.7 | A | 38.6 | A | CYP705A22 |
| 257114_at | 17.6 | A | 17.4 | A | 10.5 | A | 25.2 | A | CYP705A23 |
| 257115_at | 91.8 | P | 113 | P | 87.2 | P | 137.7 | P | --- |
| 257116_at | 116.7 | P | 129.2 | P | 188.1 | P | 179.6 | P | PRMT6 |
| 257117_at | 4.6 | A | 1.7 | A | 4.7 | A | 3.5 | A | --- |
| 257118_at | 12.8 | A | 1.6 | A | 3 | A | 1.5 | A | --- |
| 257119_at | 4.1 | A | 1.1 | A | 22.8 | A | 5.7 | A | --- |
| 257120_at | 12.6 | A | 26.9 | A | 41 | A | 50.5 | A | --- |
| 257121_at | 31.8 | A | 19.9 | A | 3.5 | A | 18.6 | A | --- |
| 257122_at | 460.1 | P | 517 | P | 572 | P | 573.2 | P | APUM5 |
| 257123_at | 20 | A | 6.6 | A | 2 | A | 3 | A | --- |
| 257124_at | 58.5 | A | 92.7 | P | 105.4 | P | 73.4 | P | ATHXK4 |
| 257125_at | 1523.5 | P | 1426.3 | P | 1724.3 | P | 1586.6 | P | ATTCP-1 |
| 257126_at | 1528.3 | P | 1399.9 | P | 1395.4 | P | 1406.2 | P | UBC19 |
| 257127_at | 72 | P | 100.2 | P | 90.9 | P | 107.1 | P | TTN9 |
| 257128_at | 61.3 | P | 67.6 | M | 45.9 | A | 71.9 | P | CYP705A15 |
| 257129_at | 385.1 | P | 567 | P | 341.5 | P | 756.2 | P | CYP705A19 |
| 257130_at | 76.2 | P | 40.3 | P | 100.1 | P | 49.4 | P | DELTA-VPE |
| 257131_at | 249.2 | P | 393.4 | P | 457.1 | P | 459.5 | P | --- |
| 257132_at | 1659.7 | P | 2192.4 | P | 1900.3 | P | 1868 | P | --- |
| 257133_at | 1 | A | 2.6 | A | 1.3 | A | 0.6 | A | --- |
| 257134_at | 167.5 | P | 281.6 | P | 184.5 | P | 259.6 | P | --- |
| 257135_at | 23.4 | A | 25.8 | A | 12.5 | A | 6.5 | A | --- |
| 257136_at | 35.5 | A | 48.4 | P | 9.5 | A | 59.2 | P | ASML2 |
| 257137_at | 577.7 | P | 767.2 | P | 709.1 | P | 690.7 | P | ABCB19 |
| 257138_at | 11.2 | A | 0.6 | A | 16.6 | A | 1.5 | A | --- |
| 257139_at | 52.7 | P | 27 | A | 30.3 | A | 26.8 | A | AtRLP43 |
| 257140_at | 128.1 | P | 193.7 | P | 169.2 | P | 191.3 | P | MYB30 |
| 257141_at | 2965 | P | 2881.9 | P | 3152.7 | P | 3282.9 | P | --- |
| 257142_at | 33.1 | P | 31.8 | P | 53.7 | P | 33.8 | P | CYP705A18 |
| 257143_at | 21.9 | P | 13 | A | 24.6 | A | 25.9 | A | CYP705A20 |
| 257144_at | 1515.9 | P | 1402.7 | P | 1280.9 | P | 1439.9 | P | G6PD5 |
| 257145_at | 240.6 | P | 292.9 | P | 290.7 | P | 314 | P | --- |
| 257146_at | 501.1 | P | 503.5 | P | 475.5 | P | 506.5 | P | GTE8 |
| 257147_at | 112.5 | P | 150 | P | 173.8 | P | 97.9 | P | --- |
| 257148_at | 4118.3 | P | 2883.5 | P | 3584.4 | P | 3125.8 | P | --- |
| 257149_at | 731 | P | 671.6 | P | 918.6 | P | 679.2 | P | ATPHB4 |
| 257150_at | 724.8 | P | 934 | P | 827.9 | P | 1011.4 | P | --- |
| 257151_at | 211 | P | 286.7 | P | 302.4 | P | 341.8 | P | --- |
| 257152_at | 529 | P | 476 | P | 355.7 | P | 472.9 | P | PUX1 |
| 257153_at | 117.1 | P | 107.4 | P | 171.6 | P | 108 | P | --- |
| 257154_at | 412.9 | P | 341.8 | P | 173.9 | P | 321.4 | P | --- |
| 257155_at | 3.5 | A | 10.5 | A | 52.4 | A | 40.2 | A | --- |
| 257156_at | 2.9 | A | 1.2 | A | 3.8 | A | 26.9 | A | --- |
| 257157_at | 571 | P | 621.3 | P | 564.8 | P | 611 | P | SYP32 |
| 257158_at | 55.3 | A | 47 | A | 62.8 | P | 35.4 | M | --- |
| 257159_at | 24.8 | A | 4.4 | A | 6.7 | A | 3.4 | A | --- |
| 257160_at | 3.1 | A | 3.9 | A | 4 | A | 1.8 | A | --- |
| 257161_at | 16.7 | A | 1.9 | A | 6.6 | A | 3.2 | A | SMAP2 |
| 257162_s_at | 2.4 | A | 9.8 | A | 1.1 | A | 3 | A | AMT1;3 |
| 257163_at | 0.5 | A | 0.9 | A | 1.6 | A | 0.5 | A | MYB305 |
| 257164_at | 95.9 | P | 109.4 | P | 96.3 | P | 96 | P | MSH1 |
| 257165_at | 7.7 | A | 5.1 | A | 9.5 | A | 7.3 | A | --- |
| 257166_at | 16.1 | A | 18.7 | A | 2.8 | A | 1.4 | A | --- |
| 257167_at | 21.7 | A | 7.6 | A | 34.4 | A | 23.2 | A | chr40 |
| 257168_at | 1353.7 | P | 1645.4 | P | 1446.9 | P | 1564.1 | P | HCF101 |
| 257169_at | 518.4 | P | 650.9 | P | 603.8 | P | 569.6 | P | AtSec20 |
| 257170_at | 3.4 | A | 16.9 | A | 36.8 | A | 24.1 | A | --- |
| 257171_at | 730.2 | P | 820.1 | P | 698.1 | P | 864.8 | P | --- |
| 257172_at | 1844.9 | P | 2273.7 | P | 1738.1 | P | 2069.1 | P | --- |
| 257173_at | 4866.6 | P | 10468.8 | P | 5356.7 | P | 9984 | P | SAHH2 |
| 257174_at | 180.3 | P | 269 | P | 216.8 | P | 213 | P | --- |
| 257175_s_at | 26.6 | P | 43.4 | P | 38.1 | P | 29.5 | P | --- |
| 257176_s_at | 271.4 | P | 510.8 | P | 375.8 | P | 608.6 | P | --- |
| 257177_at | 1982.9 | P | 2321.2 | P | 2012.7 | P | 2284.9 | P | CYN |
| 257178_at | 1333.1 | P | 1279.2 | P | 987.4 | P | 1315.7 | P | --- |
| 257179_at | 23.9 | A | 40.4 | A | 27 | A | 3.3 | A | --- |
| 257180_at | 395.8 | P | 579.8 | P | 386.4 | P | 588 | P | --- |
| 257181_at | 126 | P | 99.1 | P | 101.4 | P | 107.5 | P | --- |
| 257182_s_at | 122.6 | M | 86.2 | A | 103.5 | P | 66.1 | P | --- |
| 257183_at | 14.7 | A | 18 | A | 13.1 | A | 15.9 | A | --- |
| 257184_at | 74.7 | P | 56.3 | P | 58 | P | 71.4 | P | ATMRP8 |
| 257185_at | 40.1 | A | 98.3 | P | 34.6 | A | 82.5 | P | ATMRP7 |
| 257186_at | 7.8 | A | 76.2 | A | 39.2 | A | 45.8 | A | --- |
| 257187_at | 4.9 | A | 17.4 | A | 34.1 | A | 16.9 | A | --- |
| 257188_at | 172 | P | 163.1 | P | 173.2 | P | 144.8 | P | --- |
| 257189_at | 244.8 | P | 269.6 | P | 363.1 | P | 268.8 | P | GIP1 |
| 257190_at | 4458 | P | 7011.5 | P | 5544.3 | P | 6724.7 | P | --- |
| 257191_at | 147.1 | P | 203.5 | P | 150.3 | P | 179.9 | P | --- |
| 257192_at | 1472.4 | P | 1306.9 | P | 1321.7 | P | 1373 | P | EMB2769 |
| 257193_at | 664.6 | P | 673.6 | P | 518.6 | P | 664 | P | --- |
| 257194_at | 4801.4 | P | 2310.9 | P | 4465.2 | P | 2677.7 | P | ATSERAT2;2 |
| 257195_at | 1.5 | A | 25.2 | A | 4.6 | A | 2.7 | A | --- |
| 257196_at | 385.8 | P | 433.5 | P | 383.6 | P | 397.2 | P | --- |
| 257197_at | 13.1 | A | 33.4 | A | 9.1 | A | 30.4 | A | SBP3 |
| 257198_at | 415.3 | P | 547.9 | P | 435 | P | 510.2 | P | --- |
| 257199_at | 347.8 | P | 411.4 | P | 467.3 | P | 362.4 | P | --- |
| 257200_at | 0.1 | A | 1.3 | A | 0.5 | A | 0.9 | A | --- |
| 257201_at | 16.9 | A | 94 | P | 55.5 | A | 79.2 | P | --- |
| 257202_at | 613 | P | 548.9 | P | 531.7 | P | 585.5 | P | --- |
| 257203_at | 474.5 | P | 732.9 | P | 594.2 | P | 615.7 | P | --- |
| 257204_at | 181.4 | P | 333.7 | P | 293.3 | P | 324 | P | RALFL24 |
| 257205_at | 1924.1 | P | 1652.7 | P | 1892.9 | P | 1512.3 | P | UGT88A1 |
| 257206_at | 4667.8 | P | 1633.1 | P | 5032.8 | P | 1829.4 | P | --- |
| 257207_at | 228 | P | 167.7 | P | 250.8 | P | 199.3 | P | --- |
| 257208_at | 156 | P | 154.5 | P | 224.9 | P | 157.4 | P | --- |
| 257209_at | 293.8 | P | 359.3 | P | 329.9 | P | 345.4 | P | --- |
| 257210_at | 10 | A | 24.6 | A | 15.8 | A | 15.1 | A | TTL2 |
| 257211_at | 357 | P | 277 | P | 372 | P | 281.8 | P | --- |
| 257212_at | 483.1 | P | 598.7 | P | 685.3 | P | 629.2 | P | --- |
| 257213_at | 962.8 | P | 769.8 | P | 1148.4 | P | 880.2 | P | --- |

| | | | | | | | | | |
|---|---|---|---|---|---|---|---|---|---|
| 257214_at | 590.8 | P | 600.1 | P | 418.1 | P | 476.4 | P | --- |
| 257215_at | 237.4 | P | 251.7 | P | 231.8 | P | 248.4 | P | --- |
| 257216_at | 8443 | P | 4154 | P | 7773.5 | P | 4466.1 | P | --- |
| 257217_at | 39.4 | P | 102.1 | P | 209.8 | P | 130.8 | P | ATPPC3 |
| 257218_at | 1711.9 | P | 1523 | P | 2536.3 | P | 1729.6 | P | --- |
| 257219_at | 1032.2 | P | 1754.9 | P | 1263 | P | 1386.3 | P | HEME1 |
| 257220_at | 1 | A | 0.8 | A | 3.8 | A | 1.5 | A | ATMYB21 |
| 257221_at | 47.3 | P | 48.1 | P | 49.2 | P | 59.9 | P | ATMYB0 |
| 257222_at | 1656.7 | P | 2044.2 | P | 1712.1 | P | 1907 | P | DEGP1 |
| 257223_at | 45.8 | A | 57.6 | P | 30.8 | A | 54.9 | A | RPL12-B |
| 257224_at | 278.9 | P | 172.6 | P | 225.9 | P | 180 | P | --- |
| 257225_s_at | 12195.2 | P | 12446.6 | P | 9783.1 | P | 12957.5 | P | RPL12-C |
| 257226_at | 175.1 | P | 190.2 | P | 220.3 | P | 181.7 | P | --- |
| 257227_at | 2511 | P | 2530.2 | P | 2201 | P | 2662.6 | P | MDAR4 |
| 257228_at | 1344.8 | P | 1462 | P | 1288.4 | P | 1323.7 | P | NQR |
| 257229_at | 44.7 | P | 48.7 | P | 100 | P | 50.2 | P | IQD26 |
| 257230_at | 46.3 | P | 18.4 | A | 28.4 | A | 18 | A | DegP11 |
| 257231_at | 12.3 | A | 2.1 | A | 3.7 | A | 32.6 | P | DEGP12 |
| 257232_at | 540.2 | P | 412 | P | 749.2 | P | 443.2 | P | PAP1 |
| 257233_at | 31.8 | A | 8.9 | A | 40.7 | A | 32.3 | A | IQD10 |
| 257234_at | 39.6 | A | 25.1 | A | 35.2 | A | 53.1 | A | --- |
| 257235_at | 307.1 | P | 267.6 | P | 253.5 | P | 272.6 | P | AtRABA1g |
| 257236_at | 510.1 | P | 462.8 | P | 577.3 | P | 502.5 | P | --- |
| 257237_at | 105.6 | P | 193.1 | P | 140.7 | P | 171.4 | P | --- |
| 257238_at | 332.3 | P | 432.8 | P | 379.1 | P | 475 | P | --- |
| 257239_at | 1255.2 | P | 922.1 | P | 846.1 | P | 858 | P | --- |
| 257240_at | 790.4 | P | 808.9 | P | 773.6 | P | 795.3 | P | --- |
| 257241_at | 4.7 | A | 3.1 | A | 2.9 | A | 1.8 | A | --- |
| 257242_at | 1.4 | A | 2.2 | A | 3.5 | A | 3.8 | A | NCED6 |
| 257243_at | 24.8 | P | 21.7 | A | 14.8 | A | 11.8 | A | --- |
| 257244_at | 37.1 | A | 43.4 | A | 64.9 | A | 38.2 | A | --- |
| 257245_at | 20.6 | A | 28.5 | P | 57 | P | 51.8 | P | --- |
| 257246_at | 5.7 | A | 26.2 | A | 31.8 | A | 29.1 | A | --- |
| 257247_at | 294.9 | P | 267.8 | P | 218.9 | P | 295.3 | P | FMA |
| 257248_at | 160.6 | P | 194.1 | P | 185.5 | P | 202.3 | P | --- |
| 257249_at | 794.2 | P | 698.7 | P | 764.8 | P | 658.7 | P | --- |
| 257250_at | 528.1 | P | 446.9 | P | 590.6 | P | 387 | P | --- |
| 257251_at | 3769.8 | P | 4122.9 | P | 3566.7 | P | 3994.5 | P | PMP |
| 257252_at | 3981.4 | P | 3003.1 | P | 3675.4 | P | 3183 | P | ATGR1 |
| 257253_at | 1405 | P | 1436.1 | P | 1547.4 | P | 1892.5 | P | --- |
| 257254_at | 205.6 | P | 250 | P | 182.9 | P | 288.5 | P | --- |
| 257255_at | 28 | A | 30.2 | A | 5.7 | A | 20.9 | A | --- |
| 257256_at | 11.6 | A | 38.9 | A | 39.3 | A | 45.7 | A | --- |
| 257257_at | 13.6 | A | 19.7 | A | 42.4 | A | 2.9 | A | --- |
| 257258_at | 13.3 | A | 5.8 | A | 8 | A | 1.6 | A | --- |
| 257259_at | 17.8 | P | 4 | A | 3.3 | A | 25.3 | P | --- |
| 257260_at | 361.2 | P | 435.9 | P | 372 | P | 426.2 | P | --- |
| 257261_s_at | 27.1 | A | 12.2 | A | 23.9 | A | 15.4 | A | --- |
| 257262_at | 51.8 | A | 115.1 | P | 59.1 | A | 204.9 | P | --- |
| 257263_at | 60.8 | A | 84.9 | A | 86.4 | A | 88.5 | A | --- |
| 257264_at | 73.3 | P | 279 | P | 36 | A | 228.8 | P | --- |
| 257265_at | 99.1 | P | 173.1 | P | 110.1 | P | 143.5 | P | --- |
| 257266_at | 733.1 | P | 736 | P | 937.1 | P | 853.1 | P | --- |
| 257267_at | 434.2 | P | 459 | P | 449.9 | P | 528.4 | P | TCP4 |
| 257268_at | 326.5 | P | 330.3 | P | 348.8 | P | 328.2 | P | --- |
| 257269_at | 463.4 | P | 555.1 | P | 577.4 | P | 571.2 | P | NPSN13 |
| 257270_at | 163.6 | P | 192.6 | P | 183.7 | P | 152.3 | P | --- |
| 257271_at | 352.5 | P | 70.3 | P | 186.9 | P | 105.9 | P | --- |
| 257272_at | 149.6 | P | 340 | P | 144.6 | P | 327.8 | P | --- |
| 257273_at | 95.6 | A | 91.5 | P | 112.1 | A | 130.4 | M | UVH3 |
| 257274_at | 7.4 | A | 5.2 | A | 23.9 | A | 2.8 | A | --- |
| 257275_at | 307 | P | 120.9 | P | 240.4 | P | 105.7 | P | CID9 |
| 257276_at | 5.1 | A | 1.5 | A | 1.1 | A | 1.2 | A | --- |
| 257277_at | 41.2 | P | 59.1 | P | 27 | A | 49.7 | P | --- |
| 257278_at | 4.9 | A | 6.2 | A | 2.7 | A | 8.8 | A | --- |
| 257279_at | 536.5 | P | 706.5 | P | 354.2 | P | 581.8 | P | --- |
| 257280_at | 88.3 | P | 3.3 | A | 32.2 | A | 35 | A | NCED3 |
| 257281_s_at | 39.6 | A | 26 | A | 49.3 | A | 17.6 | A | --- |
| 257282_at | 0.2 | A | 0.3 | A | 12.3 | A | 4.4 | A | --- |
| 257283_at | 19.3 | A | 31.5 | A | 2.9 | A | 13.1 | A | --- |
| 257284_at | 6.1 | A | 6.2 | A | 20.4 | A | 13.4 | A | --- |
| 257285_at | 345.9 | P | 435.9 | P | 335.1 | P | 466.1 | P | --- |
| 257286_s_at | 31.1 | A | 0.6 | A | 2.6 | A | 19.9 | A | --- |
| 257287_at | 17.4 | A | 22.2 | P | 11.6 | A | 2.6 | A | --- |
| 257288_at | 13.6 | A | 40.9 | A | 74.3 | A | 48.3 | A | --- |
| 257289_at | 1.7 | A | 1.6 | A | 10.8 | A | 1.8 | A | --- |
| 257290_at | 31.9 | P | 97.5 | P | 70.1 | P | 91 | P | --- |
| 257291_at | 1178.4 | P | 822.1 | P | 1206.4 | P | 890.7 | P | --- |
| 257292_s_at | 2.7 | A | 0.7 | A | 1.2 | A | 3.1 | A | --- |
| 257293_at | 2391.4 | P | 2385.3 | P | 1408.7 | P | 2198.2 | P | APG8H |
| 257294_at | 289.9 | P | 483.4 | P | 290.3 | P | 567.9 | P | --- |
| 257295_at | 263.8 | P | 288.3 | P | 231.1 | P | 295.3 | P | GPK1 |
| 257296_at | 48.7 | A | 52.4 | A | 27.1 | A | 37.9 | A | --- |
| 257297_at | 891 | P | 1250.9 | P | 1046.7 | P | 1171.1 | P | --- |
| 257298_at | 19.7 | A | 21.9 | P | 22 | P | 11 | A | --- |
| 257299_at | 3165.4 | P | 2281.3 | P | 2404.6 | P | 2048.9 | P | --- |
| 257300_at | 504.6 | P | 505.6 | P | 511.4 | P | 565.3 | P | --- |
| 257301_at | 0.5 | A | 0.9 | A | 2.1 | A | 1.5 | A | --- |
| 257302_at | 17.8 | A | 20.4 | A | 20.2 | A | 25.3 | A | --- |
| 257303_at | 14.6 | A | 28.1 | A | 38.1 | A | 28.1 | A | --- |
| 257304_at | 16.8 | A | 5.9 | A | 17.9 | A | 13.7 | A | --- |
| 257305_at | 3.9 | A | 3 | A | 5.3 | A | 3.8 | A | --- |
| 257306_at | 1.4 | A | 1.5 | A | 3.6 | A | 1.1 | A | --- |
| 257307_at | 25 | A | 11.6 | A | 30.1 | A | 33.6 | A | MYB121 |
| 257308_at | 11.8 | A | 13.3 | A | 1.3 | A | 0.8 | A | --- |
| 257309_at | 38.2 | A | 29.2 | P | 32.9 | A | 27.5 | P | --- |
| 257310_at | 695.7 | P | 640.5 | P | 611.9 | P | 614 | P | --- |
| 257311_at | 2991.1 | P | 3850.8 | P | 3815.7 | P | 3799.8 | P | PHT2;1 |
| 257312_at | 29.3 | A | 28.8 | A | 49 | A | 45.4 | A | AT3G26590 |
| 257313_at | 14699.2 | P | 13616.9 | P | 14648.7 | P | 14676.8 | P | TIP2 |
| 257314_at | 361.6 | P | 368.2 | P | 456.9 | P | 361.8 | P | --- |
| 257315_at | 3354.4 | P | 575 | P | 3205.9 | P | 829.8 | P | ERD5 |
| 257316_at | 1 | A | 0.8 | A | 1.2 | A | 1.4 | A | --- |
| 257317_at | 4.3 | A | 11.4 | A | 19.7 | A | 2.9 | A | ArthMp093 |
| 257318_at | 71.9 | P | 50.8 | A | 75.1 | P | 89.6 | A | --- |
| 257319_at | 68.5 | P | 17.9 | A | 41.4 | A | 30.5 | P | ArthMp096 |
| 257320_at | 8.2 | A | 1.9 | A | 1.9 | A | 3.4 | A | --- |
| 257321_at | 124.5 | P | 76.4 | A | 148.4 | P | 102.9 | A | nad1 |
| 257322_at | 66.1 | P | 66.5 | P | 75.3 | P | 92.8 | P | nad1 |

| | | | | | | | | | |
|---|---|---|---|---|---|---|---|---|---|
| 257323_at | 80 | P | 48.3 | P | 60.2 | P | 42.9 | P | --- |
| 257324_at | 95.6 | A | 59.5 | A | 118.9 | A | 102.5 | A | nad1 |
| 257325_at | 88.6 | A | 64 | M | 86.9 | A | 72.2 | A | --- |
| 257326_s_at | 15.1 | A | 4.5 | A | 13.3 | A | 11.1 | A | ArthMp101 |
| 257327_at | 34.2 | A | 7.3 | A | 41.4 | A | 44.4 | A | nad1 |
| 257328_s_at | 40.6 | A | 26.8 | A | 37.6 | A | 54.7 | A | ArthMp102 |
| 257329_at | 10.4 | A | 7.6 | A | 46.8 | A | 47.9 | A | nad1 |
| 257330_at | 12.1 | A | 22.3 | A | 69.8 | P | 9.5 | A | ArthMp105 |
| 257331_at | 6.5 | A | 27.4 | A | 67.4 | P | 7 | A | nad2 |
| 257332_at | 35.6 | A | 38.6 | A | 28.5 | A | 23 | A | ArthMp108 |
| 257333_at | 123 | P | 132.4 | P | 140.3 | P | 106.3 | P | cox1 |
| 257334_at | 210.4 | P | 229.9 | P | 168.1 | P | 255.1 | P | ArthMp110 |
| 257335_at | 17.9 | A | 10.4 | A | 10.6 | A | 0.9 | A | ArthMp111 |
| 257336_at | 59.2 | P | 35.6 | P | 69.9 | P | 39.6 | P | ArthMp112 |
| 257337_at | 59 | A | 27.1 | A | 53.3 | A | 18.2 | A | nad5 |
| 257338_s_at | 57.5 | P | 58.1 | P | 63.2 | P | 42.2 | P | --- |
| 257339_s_at | 463.1 | P | 409.5 | P | 527.8 | P | 368.2 | P | --- |
| 257340_at | 24.4 | A | 14.1 | A | 38.6 | A | 5.4 | A | --- |
| 257341_at | 0.5 | A | 0.9 | A | 0.5 | A | 1.1 | A | --- |
| 257342_at | 53.5 | A | 70.2 | A | 57.8 | A | 62.5 | A | --- |
| 257343_s_at | 21.2 | A | 16.6 | A | 23.9 | A | 19.8 | A | --- |
| 257344_s_at | 11.7 | A | 3 | A | 4 | A | 1.2 | A | --- |
| 257345_s_at | 28.1 | P | 23 | A | 11.1 | A | 39 | A | --- |
| 257346_at | 7.8 | A | 13.5 | A | 15.1 | A | 5.3 | A | --- |
| 257347_at | 2.2 | A | 4.8 | A | 1.6 | A | 2.8 | A | --- |
| 257348_at | 2.2 | A | 6.9 | A | 2.2 | A | 1.8 | A | --- |
| 257349_at | 23.6 | A | 4.5 | A | 22.8 | A | 3.2 | A | --- |
| 257350_x_at | 26.4 | A | 19.3 | A | 30.2 | A | 32.1 | P | RALFL12 |
| 257351_at | 256.9 | P | 314.1 | P | 338.9 | P | 321.7 | P | MBAC1 |
| 257352_at | 340.2 | P | 266 | P | 307 | P | 230.8 | P | IMB1 |
| 257353_at | 2.5 | A | 2.4 | A | 2.7 | A | 1.5 | A | NPY4 |
| 257354_x_at | 3.4 | A | 4.4 | A | 9.2 | A | 3.6 | A | --- |
| 257355_at | 12.4 | A | 28.8 | A | 61.6 | P | 28.1 | A | --- |
| 257356_s_at | 23.4 | A | 27.2 | A | 27.1 | A | 29.7 | A | --- |
| 257357_at | 27.9 | A | 10.3 | A | 56.1 | A | 57.1 | A | --- |
| 257358_at | 11 | A | 10.1 | A | 3.2 | A | 3.2 | A | --- |
| 257359_x_at | 35.1 | A | 60.7 | A | 38.9 | A | 28.3 | A | --- |
| 257360_at | 24.5 | A | 4.3 | A | 8.3 | A | 30.6 | A | --- |
| 257361_at | 8.7 | A | 2.3 | A | 2.2 | A | 2.8 | A | --- |
| 257362_at | 5.9 | A | 20.2 | A | 7.5 | A | 9 | A | --- |
| 257363_at | 30.5 | P | 37.2 | A | 7 | A | 13.6 | A | BAP2 |
| 257364_at | 22.6 | A | 26.6 | A | 32.7 | A | 5.5 | A | --- |
| 257365_x_at | 1031.9 | P | 1643.5 | P | 651.7 | P | 1271.5 | P | PDF1.2b |
| 257366_s_at | 1.9 | A | 2 | A | 20.6 | A | 1.5 | A | --- |
| 257367_at | 19.6 | A | 14.7 | A | 9.3 | A | 6.7 | A | --- |
| 257368_at | 44.1 | A | 46.7 | A | 46.3 | A | 23 | A | --- |
| 257369_at | 69.6 | P | 80.8 | P | 121.1 | A | 77.7 | M | BPC7 |
| 257370_at | 7.8 | A | 38.7 | A | 4.2 | A | 15 | A | --- |
| 257371_at | 26.1 | A | 3.8 | A | 36.6 | P | 13.2 | A | NF-YB5 |
| 257372_at | 11.6 | A | 5.3 | A | 42.3 | A | 15.7 | A | --- |
| 257373_at | 4.8 | A | 7.5 | A | 4 | A | 24.5 | A | --- |
| 257374_at | 227.3 | P | 274.4 | P | 206.2 | P | 202.1 | P | --- |
| 257375_at | 136.4 | P | 183 | P | 104.7 | P | 154.7 | P | --- |
| 257376_at | 7.6 | A | 3.1 | A | 29.2 | A | 13.6 | A | --- |
| 257377_at | 267.7 | P | 463.2 | P | 330.9 | P | 434.8 | P | PLL4 |
| 257378_s_at | 1 | A | 6.3 | A | 2.5 | A | 0.9 | A | --- |
| 257379_x_at | 8.5 | A | 5.8 | A | 35.7 | A | 5.8 | A | --- |
| 257380_at | 4.6 | A | 2.2 | A | 5.6 | A | 7.2 | A | --- |
| 257381_at | 60.4 | P | 56.7 | P | 80.7 | P | 52.8 | P | --- |
| 257382_at | 14.2 | A | 23.7 | A | 14 | A | 61.6 | A | WRKY54 |
| 257383_at | 1.9 | A | 7.2 | A | 1.8 | A | 0.7 | A | --- |
| 257384_at | 5.7 | A | 4.2 | A | 34.2 | A | 5 | A | --- |
| 257385_at | 17.1 | A | 0.8 | A | 2.7 | A | 8.5 | A | --- |
| 257386_at | 0.7 | A | 2.2 | A | 2.8 | A | 1.1 | A | --- |
| 257387_s_at | 0.6 | A | 1.8 | A | 14.6 | A | 0.9 | A | --- |
| 257388_at | 1.1 | A | 18.5 | A | 8.9 | A | 19.1 | A | --- |
| 257389_at | 171.1 | P | 166.4 | P | 315 | P | 179.3 | P | --- |
| 257390_x_at | 3.3 | A | 18.6 | A | 10.3 | A | 9.2 | A | --- |
| 257391_at | 19.8 | A | 1.3 | A | 2.4 | A | 2 | A | --- |
| 257392_at | 4.4 | A | 3 | A | 6 | A | 3 | A | FLA3 |
| 257393_at | 5.9 | A | 9.2 | A | 2.8 | A | 2.7 | A | --- |
| 257394_at | 1.5 | A | 1.6 | A | 6.2 | A | 2.5 | A | --- |
| 257395_at | 124 | P | 134.2 | P | 116.5 | P | 113.2 | P | --- |
| 257396_at | 174.4 | P | 197.8 | P | 252 | P | 226.1 | P | EPF1 |
| 257397_at | 38.4 | A | 15.6 | A | 5.9 | A | 42.2 | A | RIC6 |
| 257398_at | 119.3 | P | 178.2 | P | 205.7 | P | 135.2 | P | --- |
| 257399_at | 15.2 | A | 3.1 | A | 20.8 | A | 21.7 | A | --- |
| 257400_s_at | 15.5 | A | 13.6 | A | 24.1 | A | 20.6 | A | --- |
| 257401_at | 223.7 | P | 162.1 | P | 208.7 | P | 140.2 | P | SRO2 |
| 257402_at | 0.7 | A | 3 | A | 2.5 | A | 12.2 | A | --- |
| 257403_at | 36.2 | A | 120.5 | A | 9.7 | A | 73.5 | A | --- |
| 257404_at | 45.6 | P | 88.2 | P | 60.1 | P | 42.8 | P | SUVR1 |
| 257405_at | 14.3 | A | 3.4 | A | 3 | A | 2.7 | A | --- |
| 257406_at | 104.4 | P | 130.4 | P | 129.7 | P | 128.4 | P | --- |
| 257407_at | 550.4 | P | 788.8 | P | 617.2 | P | 536.4 | P | --- |
| 257408_at | 4.3 | A | 18.9 | A | 24.9 | A | 26.8 | A | SON1 |
| 257409_at | 56.3 | A | 32.1 | A | 100.1 | A | 52.7 | A | --- |
| 257410_at | 384.6 | P | 349.3 | P | 360.5 | P | 299.7 | P | emb2421 |
| 257411_s_at | 214.8 | P | 8.9 | A | 78.5 | P | 31 | A | --- |
| 257412_at | 17.1 | A | 45.2 | A | 30.7 | A | 24.6 | A | --- |
| 257413_at | 335.3 | P | 409.3 | P | 424.3 | P | 403.8 | P | --- |
| 257414_at | 27 | A | 39.7 | A | 34.2 | A | 49.6 | A | --- |
| 257415_at | 2.6 | A | 2.1 | A | 8.7 | A | 8.2 | A | --- |
| 257416_at | 11.3 | A | 31.3 | A | 45 | A | 31.3 | A | --- |
| 257417_at | 33.3 | A | 38.4 | A | 58 | A | 50.1 | A | --- |
| 257418_at | 32.6 | A | 3.4 | A | 38.7 | A | 3.5 | A | --- |
| 257419_at | 12.3 | A | 22.3 | A | 23.1 | A | 19.8 | A | --- |
| 257420_at | 109.1 | P | 132.9 | P | 127.4 | P | 148.3 | P | --- |
| 257421_at | 35.6 | A | 62.9 | P | 306.9 | P | 170.7 | P | --- |
| 257422_at | 106.2 | P | 102.4 | P | 100.4 | P | 129.6 | P | --- |
| 257423_at | 117.2 | P | 117.4 | P | 137.5 | P | 135.3 | P | --- |
| 257424_at | 26.3 | A | 36.4 | P | 6.3 | A | 22 | P | --- |
| 257425_at | 28.3 | A | 15 | A | 78 | A | 12 | A | --- |
| 257426_at | 292.2 | P | 362.4 | P | 336.1 | P | 324.5 | P | --- |
| 257427_at | 44.5 | A | 24.4 | A | 50.6 | A | 52.4 | A | --- |
| 257428_at | 1.2 | A | 4.9 | A | 2.1 | A | 9.8 | A | --- |
| 257429_at | 17.2 | A | 6.6 | A | 20.7 | A | 1.3 | A | --- |
| 257430_at | 28.8 | A | 65.3 | A | 50 | A | 51 | A | --- |
| 257431_at | 404.5 | P | 474.5 | P | 351.8 | P | 457.7 | P | --- |

| | | | | | | | | | |
|---|---|---|---|---|---|---|---|---|---|
| 257432_at | 42.9 | A | 51.9 | P | 76.7 | M | 59.2 | M | --- |
| 257433_at | 22.2 | A | 7.1 | A | 19.4 | A | 25 | A | --- |
| 257434_at | 20.8 | A | 18.5 | A | 4.6 | A | 14.9 | A | --- |
| 257435_at | 608.6 | P | 662.2 | P | 710.6 | P | 562.3 | P | --- |
| 257436_s_at | 37.3 | P | 89 | P | 44.3 | A | 78.8 | P | --- |
| 257437_s_at | 6.3 | A | 1.5 | A | 27.7 | A | 26.5 | A | --- |
| 257438_at | 4.8 | A | 2.3 | A | 8.5 | A | 13.8 | A | --- |
| 257439_at | 19.9 | A | 2.8 | A | 20.9 | A | 2.6 | A | --- |
| 257440_at | 10.2 | A | 9 | A | 12.9 | A | 9 | A | --- |
| 257441_at | 9.1 | A | 6.3 | A | 7.5 | A | 32.9 | A | --- |
| 257442_at | 13.3 | A | 6.1 | A | 10.5 | A | 3.1 | A | --- |
| 257443_at | 1.9 | A | 1.6 | A | 2.7 | A | 2.7 | A | --- |
| 257444_at | 310.7 | P | 240 | P | 272.4 | P | 223.3 | P | --- |
| 257445_x_at | 22.8 | A | 38.9 | A | 36.6 | A | 14.4 | A | --- |
| 257446_at | 0.7 | A | 4 | A | 20 | A | 0.5 | A | --- |
| 257447_at | 326.4 | P | 284.8 | P | 337 | P | 247 | P | --- |
| 257448_s_at | 12.3 | A | 16.6 | A | 19.7 | A | 30.8 | A | --- |
| 257449_at | 36.5 | P | 6.8 | A | 66.3 | A | 18 | M | --- |
| 257450_at | 63.2 | A | 13.1 | A | 26.2 | A | 2.9 | A | --- |
| 257451_at | 103.6 | P | 127.5 | P | 122.4 | P | 128.5 | P | BT3 |
| 257452_at | 44.2 | A | 38.4 | P | 38.3 | P | 34.7 | P | --- |
| 257453_at | 26.7 | A | 24.9 | A | 34.3 | A | 26.3 | A | --- |
| 257454_at | 35.7 | P | 2.7 | A | 17.7 | A | 16.9 | A | --- |
| 257455_s_at | 19.9 | A | 7.7 | A | 37.9 | A | 21.2 | A | --- |
| 257456_at | 31.3 | P | 38.6 | P | 86.8 | P | 97.4 | P | SRS4 |
| 257457_at | 15.3 | A | 2.6 | A | 4.7 | A | 7.3 | A | --- |
| 257458_at | 1.5 | A | 1.6 | A | 1.9 | A | 1.4 | A | --- |
| 257459_at | 581.8 | P | 594.2 | P | 642.9 | P | 562.8 | P | --- |
| 257460_at | 45.4 | P | 252.1 | P | 90.2 | P | 252.7 | P | --- |
| 257461_at | 46 | P | 48 | P | 83.2 | P | 41.2 | P | ATPUP15 |
| 257462_at | 3.1 | A | 16.9 | A | 23.7 | A | 9.3 | A | --- |
| 257463_at | 565.9 | P | 759.7 | P | 646.3 | P | 864.5 | P | --- |
| 257464_at | 29.1 | A | 27.2 | P | 38.3 | P | 27.8 | P | --- |
| 257465_at | 35.4 | A | 37.1 | A | 61 | A | 41.7 | A | --- |
| 257466_at | 69.2 | P | 78.9 | P | 48.2 | A | 72.3 | P | --- |
| 257467_at | 41.4 | A | 63.7 | A | 62 | A | 62.4 | A | LBD4 |
| 257468_at | 4.5 | A | 1.3 | A | 2.1 | A | 2.5 | A | --- |
| 257469_at | 18 | A | 19.3 | A | 16.2 | A | 17.6 | A | --- |
| 257470_at | 158.8 | P | 179.9 | P | 131.1 | P | 167.2 | P | --- |
| 257471_at | 3.2 | A | 16.9 | A | 7.3 | A | 4.4 | A | ATBZIP4 |
| 257472_at | 14.7 | A | 22.5 | A | 5.5 | A | 5.4 | A | --- |
| 257473_at | 15.5 | A | 49.8 | M | 37 | A | 40.5 | A | --- |
| 257474_at | 174.1 | P | 221.9 | P | 289.9 | P | 182 | P | --- |
| 257475_at | 93.9 | P | 105.2 | A | 75.1 | A | 90.7 | P | --- |
| 257476_at | 70.1 | P | 70.4 | P | 67.7 | P | 61.4 | P | --- |
| 257477_at | 860.1 | P | 617.4 | P | 721.4 | P | 627.1 | P | --- |
| 257478_at | 73.5 | P | 56.6 | A | 25.1 | A | 38.4 | P | WAKL2 |
| 257479_at | 30.9 | A | 7.3 | A | 4.7 | A | 3.2 | A | WAKL4 |
| 257480_at | 4.6 | A | 8.8 | A | 77.8 | A | 28.2 | A | --- |
| 257481_at | 3380.6 | P | 127.2 | P | 4565.2 | P | 640.3 | P | ALMT1 |
| 257482_x_at | 0.9 | A | 29.2 | A | 32.9 | A | 27.2 | P | --- |
| 257483_at | 135.3 | P | 149 | P | 151.4 | P | 149.4 | P | ICK5 |
| 257484_at | 306.8 | P | 771.8 | P | 338.9 | P | 731.8 | P | --- |
| 257485_at | 42.7 | P | 41.2 | P | 59.8 | P | 32.1 | A | --- |
| 257486_at | 2.2 | A | 10.9 | A | 27.7 | A | 3.3 | A | --- |
| 257487_at | 144.1 | P | 170.8 | P | 174.3 | P | 196 | P | --- |
| 257488_s_at | 19.7 | P | 37.3 | A | 14 | A | 27.5 | A | --- |
| 257489_at | 37 | A | 25.8 | A | 6.3 | A | 33.6 | A | --- |
| 257490_x_at | 43.1 | P | 16.8 | P | 11.6 | A | 26.4 | M | ETC1 |
| 257491_at | 7.7 | A | 28.5 | A | 16.2 | A | 1.5 | A | --- |
| 257492_at | 43.1 | A | 28.4 | A | 36.4 | A | 2.9 | A | ARR19 |
| 257493_at | 18.4 | A | 4.9 | A | 4.7 | A | 4 | A | --- |
| 257494_at | 40.3 | A | 22.7 | A | 56.6 | A | 37.3 | A | --- |
| 257495_at | 585.8 | P | 785.6 | P | 633.2 | P | 730.9 | P | ATPDIL5-1 |
| 257496_s_at | 32.1 | P | 38.3 | A | 68.3 | A | 56 | M | --- |
| 257497_at | 121.4 | P | 125.1 | P | 177.7 | P | 122.2 | P | --- |
| 257498_at | 22.5 | A | 29.4 | A | 54.3 | A | 39.9 | A | --- |
| 257499_at | 319.9 | P | 376.9 | P | 349.9 | P | 372.7 | P | --- |
| 257500_s_at | 13.8 | A | 13.4 | A | 18.7 | A | 2.2 | A | scpl2 |
| 257501_at | 2.2 | A | 2 | A | 3.7 | A | 23.5 | A | --- |
| 257502_at | 91.2 | P | 80.1 | P | 79.9 | P | 62.8 | P | --- |
| 257503_at | 74.1 | P | 49.1 | P | 100.8 | P | 70.7 | P | --- |
| 257504_at | 61.4 | A | 90 | P | 130.8 | P | 80.6 | P | ROPGEF11 |
| 257505_at | 22.3 | A | 15.5 | A | 20.3 | A | 29.6 | A | --- |
| 257506_at | 26.7 | A | 69.9 | P | 48.3 | M | 70.6 | P | --- |
| 257507_at | 3.6 | A | 4.8 | A | 24.4 | A | 19.7 | A | --- |
| 257508_at | 11.8 | A | 8.1 | A | 1.8 | A | 0.7 | A | --- |
| 257509_at | 1 | A | 16.1 | A | 24.5 | A | 14.6 | A | --- |
| 257510_at | 595.9 | P | 1475.4 | P | 1348.4 | P | 1493.1 | P | --- |
| 257511_at | 27.6 | P | 2.7 | A | 31.7 | A | 38 | A | --- |
| 257512_at | 20.6 | A | 46.1 | A | 28.6 | A | 53.1 | P | --- |
| 257513_s_at | 445.4 | P | 469 | P | 530.4 | P | 491.9 | P | --- |
| 257514_at | 17.1 | A | 11.5 | A | 2.6 | A | 25.5 | A | --- |
| 257515_at | 15.5 | A | 4.7 | A | 37 | A | 27.1 | A | DOT5 |
| 257516_at | 196.1 | P | 239.2 | P | 271.9 | P | 209.6 | P | ACR4 |
| 257517_at | 1252.9 | P | 212.3 | P | 747.1 | P | 289.8 | P | --- |
| 257518_at | 146.7 | P | 133.8 | P | 111.3 | P | 111.7 | P | --- |
| 257519_at | 415.8 | P | 326.2 | P | 267.5 | P | 313.2 | P | --- |
| 257520_at | 1.2 | A | 4.7 | A | 1.3 | A | 1.9 | A | --- |
| 257521_at | 4.4 | A | 3 | A | 7.3 | A | 3.1 | A | --- |
| 257522_at | 266.4 | P | 144.3 | P | 303.9 | P | 176.7 | P | --- |
| 257523_at | 6 | A | 55 | A | 85.1 | A | 48.2 | A | --- |
| 257524_at | 46.3 | P | 69.5 | P | 70.3 | P | 76.2 | P | DEL3 |
| 257525_at | 11.4 | A | 5.4 | A | 34.7 | A | 4.5 | A | --- |
| 257526_s_at | 25.9 | A | 29.9 | A | 23.1 | A | 8.5 | A | --- |
| 257527_at | 30.2 | A | 19.7 | A | 39.4 | A | 37.1 | A | --- |
| 257528_at | 2.4 | A | 2.6 | A | 12.2 | A | 7.7 | A | --- |
| 257529_at | 10.9 | A | 1.4 | A | 7.7 | A | 27.4 | A | --- |
| 257530_at | 322.6 | P | 330.4 | P | 307.6 | P | 315.7 | P | --- |
| 257531_at | 12.4 | A | 10.3 | A | 6.6 | A | 26.2 | A | --- |
| 257532_at | 26.1 | A | 27.5 | P | 44.1 | A | 29.8 | P | --- |
| 257533_at | 588.9 | P | 828.5 | P | 684.5 | P | 835.5 | P | --- |
| 257534_at | 145.3 | P | 125.8 | P | 165.8 | P | 152.4 | P | --- |
| 257535_at | 95.9 | P | 112.7 | P | 111.8 | P | 68.1 | P | --- |
| 257536_at | 52.6 | P | 93 | P | 93.1 | P | 65.5 | P | --- |
| 257537_at | 8.4 | A | 27.3 | A | 17.7 | A | 18.5 | A | --- |
| 257538_at | 25.8 | A | 3 | A | 9.4 | A | 1.7 | A | --- |
| 257539_at | 2.9 | A | 2.2 | A | 5.9 | A | 4.8 | A | --- |
| 257540_at | 3.1 | A | 4.8 | A | 8.8 | A | 4.4 | A | --- |

| | | | | | | | | | |
|---|---|---|---|---|---|---|---|---|---|
| 257541_at | 56.1 | A | 48.9 | P | 83.5 | A | 24.7 | A | --- |
| 257542_at | 19.3 | A | 40.2 | P | 43.3 | A | 35.7 | A | --- |
| 257543_at | 86.2 | P | 54.2 | P | 78 | P | 62.3 | P | --- |
| 257544_at | 3.9 | A | 31 | A | 6.3 | A | 30.3 | A | --- |
| 257545_at | 192.8 | P | 161.8 | P | 165.5 | P | 169.1 | P | --- |
| 257546_s_at | 21.9 | A | 1.7 | A | 32.1 | A | 15.1 | A | --- |
| 257547_at | 95.5 | P | 145.1 | P | 119.2 | P | 157.3 | P | --- |
| 257548_s_at | 9.6 | A | 10.4 | A | 9.8 | A | 8.8 | A | LBD23 |
| 257549_at | 1.2 | A | 12.2 | A | 17.6 | A | 10.3 | A | agl103 |
| 257550_at | 23.3 | A | 20 | A | 40.3 | P | 29 | A | --- |
| 257551_at | 19.2 | A | 6.9 | A | 29.6 | A | 26.3 | A | --- |
| 257552_at | 0.5 | A | 1.3 | A | 2.5 | A | 0.5 | A | --- |
| 257553_at | 402.4 | P | 384.2 | P | 436.2 | P | 405.8 | P | TPR2 |
| 257554_at | 109.8 | P | 130.4 | P | 113 | P | 157.8 | P | VAMP728 |
| 257555_at | 70.3 | P | 24.4 | A | 27 | P | 21.7 | A | --- |
| 257556_at | 250.8 | P | 271.2 | P | 248.2 | P | 229.9 | P | --- |
| 257557_at | 7.1 | A | 7.9 | A | 34.8 | A | 21.3 | A | --- |
| 257558_s_at | 1.7 | A | 1.4 | A | 1.5 | A | 2.1 | A | --- |
| 257559_at | 2.7 | A | 9.7 | A | 4.3 | A | 10.7 | A | --- |
| 257560_at | 383.2 | P | 355.9 | P | 368.3 | P | 329.6 | P | --- |
| 257561_at | 27.9 | A | 16.5 | A | 24.7 | A | 13.4 | A | APUM25 |
| 257562_at | 24.3 | M | 59 | P | 53.5 | M | 34.3 | A | --- |
| 257563_at | 19.8 | A | 29.1 | P | 20 | A | 16.1 | P | --- |
| 257564_at | 19.4 | A | 21.3 | A | 31.1 | A | 38.7 | A | --- |
| 257565_at | 13.2 | A | 5.9 | A | 4.7 | A | 1.3 | A | --- |
| 257566_x_at | 0.4 | A | 2.4 | A | 3.7 | A | 0.3 | A | --- |
| 257567_at | 66.8 | P | 35.2 | A | 101.6 | P | 58.7 | P | --- |
| 257568_s_at | 23.5 | A | 14.4 | A | 3.7 | A | 30 | A | --- |
| 257569_at | 45.6 | P | 52.2 | P | 26 | A | 55.5 | P | --- |
| 257570_at | 7.9 | A | 18.5 | A | 16.9 | A | 2.9 | A | --- |
| 257571_at | 264.7 | P | 257.9 | P | 298.5 | P | 250.7 | P | --- |
| 257572_at | 24.6 | A | 3.7 | A | 16.4 | A | 29.6 | A | --- |
| 257573_at | 103 | P | 133.4 | P | 60.1 | P | 144.3 | P | iqd9 |
| 257574_at | 34.3 | A | 6 | A | 4.3 | A | 4.8 | A | --- |
| 257575_at | 21.7 | A | 13.4 | A | 13.5 | A | 27.4 | A | --- |
| 257576_at | 1.3 | A | 3.7 | A | 4.1 | A | 2.5 | A | --- |
| 257577_at | 2 | A | 3 | A | 11 | A | 2.6 | A | --- |
| 257578_x_at | 98.7 | P | 122.3 | P | 125.6 | A | 136.7 | P | --- |
| 257579_at | 5.8 | A | 29.1 | A | 37.2 | A | 63.2 | A | --- |
| 257580_at | 138.5 | P | 141.4 | P | 114.6 | A | 128.6 | P | --- |
| 257581_s_at | 2.2 | A | 2.3 | A | 1.4 | A | 6 | A | --- |
| 257582_at | 1.3 | A | 10.1 | A | 16.8 | A | 0.8 | A | --- |
| 257583_at | 405.8 | P | 222.5 | P | 359.8 | P | 197.6 | P | PMI2 |
| 257584_at | 3.8 | A | 7.4 | A | 12.2 | A | 6.1 | A | --- |
| 257585_at | 15.5 | A | 3.6 | A | 4.3 | A | 2.8 | A | --- |
| 257586_at | 30.9 | A | 15.7 | A | 3.7 | A | 9.6 | A | --- |
| 257587_at | 137.9 | P | 162.3 | P | 146.8 | P | 135.3 | P | --- |
| 257588_x_at | 50.2 | A | 8.8 | A | 7.8 | A | 26.9 | A | --- |
| 257589_at | 27.9 | A | 30.3 | A | 50.6 | A | 25.5 | A | --- |
| 257590_s_at | 166.8 | P | 138.3 | P | 136.4 | P | 119.3 | P | --- |
| 257591_at | 41.7 | P | 72.8 | P | 58.3 | A | 35.2 | P | AtRLP39 |
| 257592_at | 2471.8 | P | 127.4 | P | 528.8 | P | 378.6 | P | --- |
| 257593_at | 50.6 | A | 82.3 | P | 60.6 | M | 58.6 | P | --- |
| 257594_at | 298.7 | P | 378.1 | P | 341.8 | P | 369 | P | --- |
| 257595_at | 3.5 | A | 1.3 | A | 6.2 | A | 7.2 | A | --- |
| 257596_at | 762.5 | P | 631.5 | P | 727.6 | P | 678.9 | P | --- |
| 257597_at | 6.5 | A | 4 | A | 20.4 | A | 5.8 | A | --- |
| 257598_at | 304.7 | P | 386 | P | 317 | P | 336.2 | P | PRT1 |
| 257599_at | 7196.5 | P | 7579.8 | P | 7922 | P | 7569.3 | P | --- |
| 257600_at | 1126.9 | P | 955.9 | P | 1111.2 | P | 1113.2 | P | CLE41 |
| 257601_at | 227.7 | P | 174.7 | P | 204.5 | P | 175.7 | P | --- |
| 257602_at | 166.2 | P | 281 | P | 267.8 | P | 234.2 | P | --- |
| 257603_at | 23.1 | A | 8.6 | A | 3.9 | A | 19.8 | A | --- |
| 257604_s_at | 2.9 | A | 3.9 | A | 1.2 | A | 9.9 | A | --- |
| 257605_at | 17.7 | A | 15.4 | A | 6.8 | A | 23.6 | A | --- |
| 257606_at | 1127.5 | P | 1313.8 | P | 1009.7 | P | 1253 | P | RHD3 |
| 257607_at | 82.5 | P | 76 | P | 89.5 | P | 59.2 | P | --- |
| 257608_at | 1226.3 | P | 1093.7 | P | 1218.6 | P | 1172.1 | P | HSP60-3A |
| 257609_at | 443.9 | P | 479.7 | P | 368.9 | P | 519.2 | P | --- |
| 257610_at | 220.5 | P | 121.4 | P | 108.9 | P | 96.2 | P | AtIDD11 |
| 257611_at | 3489.6 | P | 3052.5 | P | 3333.7 | P | 3145.8 | P | --- |
| 257612_at | 334.4 | P | 361.9 | P | 291.5 | P | 311.5 | P | ARO4 |
| 257613_at | 41.6 | A | 31.9 | A | 52 | A | 2.1 | A | --- |
| 257614_at | 19.8 | A | 28.2 | A | 72.1 | A | 51.7 | A | --- |
| 257615_at | 1761.7 | P | 1220.1 | P | 1577.9 | P | 1130.6 | P | --- |
| 257616_at | 91.4 | P | 77.2 | P | 94.8 | P | 91.5 | P | --- |
| 257617_at | 13.3 | A | 29.5 | P | 35.7 | A | 29.1 | A | --- |
| 257618_at | 51.3 | P | 49.9 | P | 7.9 | A | 39.1 | P | --- |
| 257619_at | 83.9 | P | 76.6 | P | 82.7 | P | 102.1 | P | ICK3 |
| 257620_at | 6.6 | A | 4.9 | A | 6.7 | A | 6.5 | A | --- |
| 257621_at | 462.5 | P | 398.9 | P | 507.2 | P | 403.7 | P | CPK9 |
| 257622_at | 124.2 | P | 150.3 | P | 129.7 | P | 121.2 | P | --- |
| 257623_at | 194.1 | P | 551.2 | P | 86.8 | P | 362.9 | P | CYP71B23 |
| 257624_at | 212.7 | P | 245 | P | 150.8 | P | 173.4 | P | CYP71B3 |
| 257625_at | 73.4 | P | 237.4 | P | 50.8 | P | 100 | P | CYP71B24 |
| 257626_s_at | 44.3 | A | 32.1 | P | 33.2 | A | 52.4 | P | --- |
| 257627_at | 1.3 | A | 14.3 | A | 5.1 | A | 4.8 | A | CYP71B25 |
| 257628_at | 3437.4 | P | 2697.3 | P | 2749.1 | P | 2553.4 | P | CYP71B26 |
| 257629_at | 28.6 | A | 17.1 | A | 55.5 | A | 32.8 | P | --- |
| 257630_at | 5.9 | A | 5.3 | A | 1.3 | A | 3.1 | A | CYP71B16 |
| 257631_at | 40.4 | A | 47.6 | A | 7.4 | A | 31.7 | A | CYP71B17 |
| 257632_at | 0.5 | A | 20.7 | A | 4.5 | A | 1.3 | A | CYP71B21 |
| 257633_at | 38.3 | P | 40.7 | P | 50.1 | P | 50.1 | P | CYP86C2 |
| 257634_s_at | 1312.6 | P | 1332 | P | 1140.8 | P | 1129.4 | P | CYP71B19 |
| 257635_at | 1040.9 | P | 769.5 | P | 649.3 | P | 670.4 | P | CYP71B4 |
| 257636_at | 47.9 | M | 267.1 | P | 52.3 | A | 177.1 | P | CYP71B22 |
| 257637_at | 4.6 | A | 16.3 | A | 10.1 | A | 7.7 | A | --- |
| 257638_at | 17.7 | A | 18.5 | A | 41.4 | A | 3.5 | A | ATTPS-CIN |
| 257639_at | 22.9 | A | 46.4 | A | 65 | A | 44.7 | A | --- |
| 257640_at | 1 | A | 2.7 | A | 3.8 | A | 1.1 | A | --- |
| 257641_s_at | 13195.4 | P | 8373.6 | P | 10293.3 | P | 8537 | P | AOC1 |
| 257642_at | 34 | P | 36.1 | P | 79.4 | P | 57.4 | P | BHLH32 |
| 257643_at | 55.2 | P | 69.3 | P | 87.7 | P | 61.2 | P | --- |
| 257644_at | 774 | P | 333.1 | P | 646.7 | P | 274.4 | P | AOC3 |
| 257645_at | 3.1 | A | 40.1 | A | 4.2 | A | 3.3 | A | --- |
| 257646_at | 398.3 | P | 356.9 | P | 358.7 | P | 307.7 | P | MAP1B |
| 257647_at | 645.3 | P | 810.5 | P | 643.1 | P | 829.4 | P | --- |
| 257648_at | 298.1 | P | 231.1 | P | 341 | P | 275.2 | P | --- |
| 257649_at | 55.8 | P | 170 | P | 151.3 | P | 120.7 | P | ARR1 |

| | | | | | | | | | |
|---|---|---|---|---|---|---|---|---|---|
| 257650_at | 134.2 | P | 118.5 | P | 125.1 | A | 78.2 | A | --- |
| 257651_at | 565.7 | P | 695.1 | P | 514.9 | P | 673.8 | P | --- |
| 257652_at | 484.9 | P | 392.5 | P | 630.8 | P | 376.8 | P | APUM24 |
| 257653_at | 120.8 | P | 137.8 | P | 138.1 | M | 140.5 | P | --- |
| 257654_at | 434.8 | P | 474.8 | P | 458 | P | 511 | P | --- |
| 257655_at | 155.1 | P | 214.3 | P | 208.7 | P | 195.2 | P | --- |
| 257656_at | 18.2 | A | 23.8 | M | 35.1 | A | 43.8 | P | --- |
| 257657_at | 1070.2 | P | 1303.9 | P | 1044.6 | P | 1184 | P | --- |
| 257658_at | 1223.1 | P | 1030.5 | P | 1507.4 | P | 1150.5 | P | --- |
| 257659_at | 502.5 | P | 441.4 | P | 542.6 | P | 492.4 | P | --- |
| 257660_at | 337.3 | P | 469.6 | P | 406.9 | P | 553.3 | P | WIP3 |
| 257661_at | 453.9 | P | 474.1 | P | 517.8 | P | 554.7 | P | --- |
| 257662_at | 371.8 | P | 439 | P | 383.3 | P | 432.2 | P | CAX2 |
| 257663_at | 107.1 | P | 160.5 | P | 155.4 | P | 195.7 | P | --- |
| 257664_at | 2.1 | A | 9.5 | A | 15.3 | A | 23.7 | A | EMB2743 |
| 257665_at | 417.6 | P | 476.6 | P | 509.1 | P | 452.7 | P | --- |
| 257666_at | 702.1 | P | 571.2 | P | 883.5 | P | 584.4 | P | --- |
| 257667_at | 140.7 | P | 179 | P | 210 | P | 180.4 | P | EMB2729 |
| 257668_at | 34.3 | P | 26.3 | A | 44.7 | A | 13.9 | A | --- |
| 257669_at | 42.6 | P | 45.4 | P | 34.2 | P | 45.4 | A | MSH5 |
| 257670_at | 56.1 | P | 117.2 | P | 22.7 | A | 91.4 | P | --- |
| 257671_at | 7.3 | A | 36.7 | A | 14.9 | A | 7.3 | A | --- |
| 257672_at | 700.9 | P | 407.6 | P | 652.4 | P | 510.3 | P | --- |
| 257673_at | 1153.8 | P | 328.2 | P | 1198.8 | P | 468.7 | P | --- |
| 257674_at | 6842.2 | P | 6024.9 | P | 6430.9 | P | 6117.8 | P | --- |
| 257675_at | 524.8 | P | 450.9 | P | 383.1 | P | 422.6 | P | ERF7 |
| 257676_at | 828.1 | P | 842.6 | P | 906.4 | P | 857.3 | P | TGD2 |
| 257677_at | 1119.3 | P | 933 | P | 939.1 | P | 962 | P | ATEHD1 |
| 257678_at | 77.1 | P | 44.1 | P | 70.3 | P | 58.4 | P | RTL2 |
| 257679_at | 391.9 | P | 1026.5 | P | 490.7 | P | 784.4 | P | GRP5 |
| 257680_at | 1895.7 | P | 1426.1 | P | 2358.2 | P | 1455.4 | P | --- |
| 257681_at | 2.8 | A | 0.6 | A | 8.3 | A | 2.3 | A | --- |
| 257682_at | 11.4 | M | 19.4 | P | 17.7 | P | 19.5 | P | --- |
| 257683_at | 2.8 | A | 34.3 | A | 47.4 | A | 41.6 | A | --- |
| 257684_s_at | 2031.6 | P | 1998.2 | P | 2023.4 | P | 2168.1 | P | VCS |
| 257685_at | 87.1 | P | 96.4 | P | 144.6 | P | 158.2 | P | --- |
| 257686_at | 3201.3 | P | 2369.5 | P | 2254.8 | P | 2336.2 | P | SDRB |
| 257687_at | 3366.3 | P | 2928.6 | P | 3208.4 | P | 3132.5 | P | SDRB |
| 257688_at | 37.5 | P | 46.1 | P | 52.4 | P | 34.9 | P | PIE1 |
| 257689_at | 68.9 | A | 64.3 | M | 78.9 | A | 42.9 | P | AtMYB10 |
| 257690_at | 30.1 | A | 250.7 | P | 30.8 | A | 170 | P | --- |
| 257691_at | 19.1 | A | 6.4 | A | 32 | A | 35.3 | A | FLA14 |
| 257692_at | 3.8 | A | 2.1 | A | 2.4 | A | 9.6 | A | --- |
| 257693_at | 28.8 | A | 32.6 | A | 72 | A | 62.2 | A | --- |
| 257694_at | 185 | P | 121.3 | P | 155.6 | P | 148.1 | P | --- |
| 257695_at | 378.4 | P | 348.4 | P | 395.6 | P | 318.6 | P | HUA1 |
| 257696_at | 113.3 | P | 126 | P | 153.7 | P | 130.2 | P | AGC1.5 |
| 257697_at | 197.6 | P | 237.1 | P | 264.8 | P | 287.7 | P | --- |
| 257698_at | 154.1 | P | 157.3 | P | 156.8 | P | 137.7 | P | --- |
| 257699_at | 12175.6 | P | 11057.9 | P | 9766 | P | 10586.5 | P | PGK1 |
| 257700_at | 2455.2 | P | 1785.9 | P | 1494.9 | P | 1803 | P | ALIS1 |
| 257701_at | 62 | P | 351.6 | P | 117.1 | P | 290.9 | P | --- |
| 257702_at | 1696.2 | P | 723 | P | 1399 | P | 945.1 | P | emb2742 |
| 257703_at | 587 | P | 509 | P | 613.8 | P | 538.3 | P | --- |
| 257704_at | 2.4 | A | 2.3 | A | 2.4 | A | 2.1 | A | ATMYB67 |
| 257705_at | 1765.9 | P | 1929.6 | P | 1972.1 | P | 1924.7 | P | --- |
| 257706_at | 428.2 | P | 501.7 | P | 382.6 | P | 460.2 | P | --- |
| 257707_at | 737.7 | P | 521.7 | P | 704.2 | P | 495.4 | P | --- |
| 257708_at | 580 | P | 688.9 | P | 476.3 | P | 670.4 | P | --- |
| 257709_at | 102.1 | P | 145.2 | P | 158.6 | P | 180.2 | P | --- |
| 257710_at | 136.3 | P | 190.7 | P | 221.7 | P | 201.5 | P | --- |
| 257711_at | 1811.5 | P | 1746.3 | P | 1913.9 | P | 1849.2 | P | PBB1 |
| 257712_at | 480.3 | P | 298.9 | P | 394 | P | 321.8 | P | --- |
| 257713_at | 2893.8 | P | 2316.9 | P | 2739.1 | P | 2502.8 | P | SDH2-1 |
| 257714_at | 364.3 | P | 462.7 | P | 416.5 | P | 501.3 | P | --- |
| 257715_at | 122.1 | P | 163.4 | P | 251.2 | P | 190.9 | P | ZIP1 |
| 257716_at | 10.6 | A | 3.8 | A | 38.5 | A | 10.1 | A | --- |
| 257717_at | 982.5 | P | 1383.9 | P | 960.2 | P | 1179.5 | P | EMB1865 |
| 257718_at | 19.3 | A | 39.6 | P | 51.7 | A | 33.3 | A | anac058 |
| 257719_at | 249.1 | P | 274.1 | P | 130.6 | P | 301.6 | P | AtALMT9 |
| 257720_at | 23.8 | A | 3.4 | A | 16 | A | 26.1 | A | --- |
| 257721_at | 159.5 | P | 104.7 | P | 110.7 | P | 115.1 | P | --- |
| 257722_at | 3381 | P | 5752.8 | P | 3263.1 | P | 4757.6 | P | --- |
| 257723_at | 241.9 | P | 397.3 | P | 230.7 | P | 414.1 | P | --- |
| 257724_at | 453.3 | P | 330.5 | P | 312.6 | P | 411.2 | P | --- |
| 257725_at | 72.4 | P | 127.2 | P | 150.7 | P | 124.4 | P | MSH2 |
| 257726_at | 6.3 | A | 33.5 | A | 2.9 | A | 17 | A | --- |
| 257727_at | 253.8 | P | 310.9 | P | 339.2 | P | 281.5 | P | --- |
| 257728_at | 190.5 | P | 173.5 | P | 141.7 | P | 168 | P | --- |
| 257729_at | 414 | P | 352.5 | P | 418 | P | 368.9 | P | HDA15 |
| 257730_at | 1351.5 | P | 1614.7 | P | 1718.7 | P | 1707.4 | P | --- |
| 257731_at | 1063.8 | P | 895.1 | P | 840.8 | P | 890.8 | P | --- |
| 257732_at | 532 | P | 546.8 | P | 428 | P | 554.9 | P | AtCASP |
| 257733_at | 235.8 | P | 269.7 | P | 228.6 | P | 236.2 | P | --- |
| 257734_at | 229.2 | P | 441.3 | P | 153.3 | P | 353.1 | P | ATSYTF |
| 257735_at | 387.9 | P | 126.6 | P | 253 | P | 178.1 | P | --- |
| 257736_at | 21.2 | A | 25.1 | A | 24 | A | 1.7 | A | --- |
| 257737_at | 4.9 | A | 28.4 | A | 18.2 | A | 3.4 | A | --- |
| 257738_at | 17.3 | A | 23.9 | A | 26.4 | A | 27.2 | A | --- |
| 257739_at | 163.7 | P | 163.4 | P | 172.8 | P | 194.5 | P | --- |
| 257740_at | 39.8 | P | 84.1 | P | 50.3 | P | 56.3 | P | --- |
| 257741_at | 506.5 | P | 498.5 | P | 417.5 | P | 434.5 | P | --- |
| 257742_at | 3.8 | A | 3.7 | A | 1.5 | A | 4.8 | A | SDH2-1 |
| 257743_at | 604.3 | P | 602.3 | P | 480.8 | P | 545.4 | P | --- |
| 257744_at | 465.9 | P | 490.1 | P | 436.3 | P | 609.2 | P | --- |
| 257745_at | 1689.1 | P | 2011 | P | 1721.6 | P | 1624.5 | P | --- |
| 257746_at | 1017.5 | P | 858.5 | P | 1047.4 | P | 934.8 | P | CM1 |
| 257747_at | 16.9 | A | 2.6 | A | 4 | A | 27.4 | A | --- |
| 257748_at | 150.7 | P | 76.6 | P | 138.6 | P | 123.9 | P | PUB29 |
| 257749_at | 5347 | P | 5759.6 | P | 5951.1 | P | 6538.6 | P | ACT2 |
| 257750_at | 146.6 | P | 207.5 | P | 180.8 | P | 148.1 | P | --- |
| 257751_at | 208.5 | P | 151.6 | P | 213.6 | P | 196.6 | P | MKS1 |
| 257752_at | 1.4 | A | 4 | A | 3.3 | A | 4.7 | A | --- |
| 257753_at | 10751.7 | P | 9484.1 | P | 13432 | P | 10706.5 | P | --- |
| 257754_at | 2114.5 | P | 1973.1 | P | 1565.1 | P | 1957.7 | P | ATRAB7B |
| 257755_at | 689.7 | P | 644.4 | P | 820.8 | P | 705.4 | P | --- |
| 257756_at | 2067.3 | P | 2575.4 | P | 2605.4 | P | 2521.9 | P | --- |
| 257757_at | 61.1 | A | 57.2 | P | 57 | P | 50.3 | P | PGSIP1 |
| 257758_at | 392.6 | P | 342.4 | P | 327.5 | P | 289.4 | P | --- |

| | | | | | | | | | |
|---|---|---|---|---|---|---|---|---|---|
| 257759_at | 500.7 | P | 471.3 | P | 480.6 | P | 513.7 | P | --- |
| 257760_s_at | 1.6 | A | 0.6 | A | 7.9 | A | 1 | A | --- |
| 257761_at | 43.1 | A | 55.6 | P | 61.4 | A | 38.8 | P | --- |
| 257762_at | 79.5 | P | 113.7 | P | 108.3 | P | 117.5 | P | XRCC4 |
| 257763_s_at | 241.1 | P | 792.9 | P | 149.5 | P | 491.6 | P | AtRLP37 |
| 257764_at | 10.5 | A | 14.3 | A | 1.8 | A | 12.2 | A | AtRLP36 |
| 257765_at | 117.7 | P | 127.1 | P | 116.9 | P | 94.2 | P | --- |
| 257766_at | 356.6 | P | 396.7 | P | 258.7 | P | 394 | P | IAA2 |
| 257767_at | 3.1 | A | 6 | A | 3.4 | A | 4.3 | A | --- |
| 257768_at | 19.1 | A | 21.1 | A | 12.2 | A | 23.2 | A | --- |
| 257769_at | 959 | P | 2008.2 | P | 932.8 | P | 1706.7 | P | IAA7 |
| 257770_at | 573.2 | P | 663.3 | P | 491.6 | P | 594.7 | P | LFR |
| 257771_at | 563.6 | P | 1211.2 | P | 638.5 | P | 1102.7 | P | CIPK7 |
| 257772_at | 305.5 | P | 358.6 | P | 237.7 | P | 343.5 | P | --- |
| 257773_at | 1302.5 | P | 1417.8 | P | 1141.8 | P | 1351.6 | P | --- |
| 257774_at | 66.4 | A | 5.7 | A | 51.5 | A | 23.7 | A | --- |
| 257775_at | 19.6 | A | 21.4 | A | 4.1 | A | 8.5 | A | --- |
| 257776_at | 5.7 | A | 26 | A | 5 | A | 2.5 | A | --- |
| 257777_x_at | 44.3 | P | 44.6 | P | 49.3 | A | 42.4 | A | --- |
| 257778_at | 1.8 | A | 1.1 | A | 3.6 | A | 5 | A | --- |
| 257779_at | 24.5 | A | 11.3 | A | 21.5 | A | 29.2 | A | CDG1 |
| 257780_at | 418.7 | P | 359.2 | P | 341.8 | P | 349.5 | P | --- |
| 257781_at | 24.5 | A | 22.3 | A | 60.6 | A | 54.1 | A | --- |
| 257782_at | 31.3 | A | 6 | A | 4.3 | A | 37.1 | A | --- |
| 257783_at | 345.3 | P | 369.6 | P | 349.9 | P | 332.5 | P | --- |
| 257784_at | 448.5 | P | 462.9 | P | 359.5 | P | 362.1 | P | MUB4 |
| 257785_at | 309.6 | P | 319 | P | 275.3 | P | 282.9 | P | MUB4 |
| 257786_at | 156.8 | P | 168.6 | P | 141.2 | P | 145.3 | P | --- |
| 257787_at | 201.9 | A | 214.3 | A | 147.6 | A | 195.7 | A | ARP2 |
| 257788_at | 141.9 | P | 124 | P | 102.2 | P | 104.4 | P | TCP20 |
| 257789_at | 499.5 | P | 855 | P | 660.6 | P | 757 | P | YSL6 |
| 257790_at | 2251.2 | P | 2658.9 | P | 3049.1 | P | 2696.5 | P | --- |
| 257791_at | 668.9 | P | 725.8 | P | 522.2 | P | 706.5 | P | --- |
| 257792_at | 752 | P | 1113.8 | P | 843.4 | P | 1027.6 | P | TOM20-3 |
| 257793_at | 262.8 | P | 377.3 | P | 552.7 | P | 422.5 | P | --- |
| 257794_at | 565.4 | P | 838.3 | P | 648.2 | P | 712 | P | --- |
| 257795_at | 50.8 | P | 39.5 | P | 81.7 | P | 30.9 | P | --- |
| 257796_at | 4.6 | A | 13.6 | A | 36.9 | P | 12.3 | A | --- |
| 257797_at | 430.8 | P | 559.7 | P | 467.2 | P | 508.1 | P | --- |
| 257798_at | 2639.7 | P | 1325.2 | P | 2444.8 | P | 1537.4 | P | NAI2 |
| 257799_at | 70.4 | P | 86.6 | P | 94.6 | P | 93.6 | P | --- |
| 257800_at | 801.4 | P | 658.3 | P | 799.5 | P | 745.4 | P | --- |
| 257801_at | 97.1 | P | 184.9 | P | 239 | P | 232.3 | P | WNK6 |
| 257802_at | 66.1 | P | 87.7 | P | 84.5 | P | 54.1 | P | --- |
| 257803_at | 612.3 | P | 440.3 | P | 413.7 | P | 431.6 | P | --- |
| 257804_at | 31.6 | P | 18.2 | P | 21.7 | M | 19.9 | A | --- |
| 257805_at | 1572.6 | P | 1573.3 | P | 905.7 | P | 1415.3 | P | ATPLT5 |
| 257806_at | 10.9 | A | 24.9 | A | 48.8 | A | 23.9 | A | --- |
| 257807_at | 19320.2 | P | 19860.5 | P | 25236.9 | P | 21176.1 | P | GAPA |
| 257808_at | 368.1 | P | 336.6 | P | 413 | P | 363.3 | P | --- |
| 257809_at | 2311.3 | P | 3438.1 | P | 3022.2 | P | 3636.1 | P | TSO2 |
| 257810_at | 14.7 | A | 31.1 | A | 19.2 | A | 9.1 | A | TOM20-1 |
| 257811_at | 42 | A | 33.3 | A | 39.6 | A | 20.7 | A | --- |
| 257812_at | 1.7 | A | 9.3 | A | 31.6 | A | 9.8 | A | --- |
| 257813_at | 73.4 | P | 123.4 | P | 139.2 | P | 129.9 | P | CDC45 |
| 257814_at | 146.9 | P | 162.2 | P | 153.8 | P | 230.3 | P | AtFaTA |
| 257815_at | 54 | P | 82.6 | P | 49.7 | P | 114.6 | P | --- |
| 257816_at | 1898.1 | P | 2373 | P | 2180.4 | P | 2186.3 | P | QUA1 |
| 257817_at | 1093.5 | P | 1174.3 | P | 1444.2 | P | 1225.7 | P | --- |
| 257818_at | 930 | P | 909.2 | P | 1073.2 | P | 964.4 | P | --- |
| 257819_at | 9.7 | A | 3.4 | A | 5 | A | 1.9 | A | RALFL25 |
| 257820_at | 1433.8 | P | 2127 | P | 1537.8 | P | 1691 | P | FKBP15-1 |
| 257821_at | 6.3 | A | 5.1 | A | 3.4 | A | 3.7 | A | RALFL26 |
| 257822_at | 1084.3 | P | 1235.8 | P | 1365.8 | P | 1258 | P | ROF1 |
| 257823_at | 6.5 | A | 95.3 | A | 69.1 | A | 122.8 | A | --- |
| 257824_at | 433.1 | P | 210.4 | P | 303.1 | P | 256.6 | P | --- |
| 257825_at | 153.1 | P | 219.1 | P | 184.9 | P | 246.7 | P | --- |
| 257826_at | 697 | P | 706.6 | P | 657.5 | P | 655.8 | P | --- |
| 257827_at | 307.3 | P | 297.7 | P | 315.7 | P | 316.6 | P | --- |
| 257828_at | 449.1 | P | 369 | P | 333.9 | P | 352.7 | P | --- |
| 257829_at | 114.8 | P | 118.3 | P | 122.3 | P | 125.7 | P | SNM1 |
| 257830_at | 1248.8 | P | 545.7 | P | 1167.8 | P | 656.7 | P | ATNUDX13 |
| 257831_at | 3234.5 | P | 3032.8 | P | 3528.2 | P | 2719.8 | P | CCB1 |
| 257832_at | 9998.1 | P | 10642.1 | P | 10677.2 | P | 10721.1 | P | CCL |
| 257833_at | 369.1 | P | 393 | P | 334.3 | P | 304.6 | P | LWD2 |
| 257834_at | 768.6 | P | 849.4 | P | 929.7 | P | 866.9 | P | --- |
| 257835_at | 355.4 | P | 50.5 | A | 129.6 | P | 68.4 | P | CYP82G1 |
| 257836_at | 3.6 | A | 1.7 | A | 14.6 | A | 2.2 | A | --- |
| 257837_at | 2.3 | A | 27.6 | A | 59.1 | A | 25 | A | --- |
| 257838_at | 140.2 | P | 118.2 | P | 159.1 | P | 129.6 | P | --- |
| 257839_at | 46.1 | A | 15.1 | A | 2.9 | A | 34.1 | A | --- |
| 257840_at | 73.5 | P | 33.6 | A | 46.7 | A | 45.9 | P | AGC2-1 |
| 257841_at | 26.2 | A | 40.2 | A | 32.5 | A | 37.5 | A | --- |
| 257842_at | 2.2 | A | 4.2 | A | 1.6 | A | 2.2 | A | PGP18 |
| 257843_at | 5.6 | A | 9.7 | A | 40 | A | 11.8 | A | --- |
| 257844_at | 744.5 | P | 849.9 | P | 662 | P | 888.4 | P | --- |
| 257845_at | 147.4 | P | 157.6 | P | 174.3 | P | 135.2 | P | --- |
| 257846_at | 4.6 | A | 4.1 | A | 2.3 | A | 4 | A | --- |
| 257847_at | 2.1 | A | 32.4 | A | 28.9 | A | 33.5 | A | --- |
| 257848_at | 119.7 | P | 140.1 | P | 141 | P | 154.5 | P | --- |
| 257849_at | 583 | P | 719.1 | P | 510.6 | P | 642.3 | P | ECT5 |
| 257850_at | 10.8 | A | 9.5 | A | 9.6 | A | 4.1 | A | SRF4 |
| 257851_at | 104.7 | P | 130.2 | P | 107.9 | P | 84.8 | P | --- |
| 257852_at | 201.9 | P | 254.7 | P | 216.8 | P | 254.2 | P | --- |
| 257853_at | 2.5 | A | 1.9 | A | 1.3 | A | 6.6 | A | --- |
| 257854_at | 308.7 | P | 347.2 | P | 310 | P | 367 | P | HAC5 |
| 257855_at | 1031.1 | P | 822.1 | P | 856.8 | P | 963.5 | P | --- |
| 257856_at | 956.7 | P | 1626 | P | 1292.4 | P | 1563.8 | P | --- |
| 257857_s_at | 451.7 | P | 412.6 | P | 541.6 | P | 410.1 | P | RRP45a |
| 257858_at | 390.7 | P | 704.6 | P | 351.4 | P | 510.4 | P | --- |
| 257859_at | 13.5 | A | 28.1 | P | 29.5 | A | 26.7 | A | --- |
| 257860_at | 352 | P | 472.2 | P | 412.3 | P | 360.3 | P | --- |
| 257861_at | 262.1 | P | 267.9 | P | 275.3 | P | 265.8 | P | --- |
| 257862_s_at | 12.1 | A | 0.7 | A | 1.6 | A | 8.9 | A | --- |
| 257863_at | 28.7 | A | 52.7 | A | 9 | A | 25.6 | A | anac057 |
| 257864_at | 248.4 | P | 183.8 | P | 230 | P | 177.6 | P | --- |
| 257865_at | 290.5 | P | 394.3 | P | 385.2 | P | 405.1 | P | --- |
| 257866_at | 323.9 | P | 516.3 | P | 278.4 | P | 415.1 | P | --- |
| 257867_at | 643.5 | P | 680.9 | P | 618.1 | P | 701.7 | P | --- |

| | | | | | | | | | |
|---|---|---|---|---|---|---|---|---|---|
| 257868_at | 664.2 | P | 788.8 | P | 514.3 | P | 728.7 | P | RIN4 |
| 257869_at | 36.1 | P | 35.5 | P | 30.1 | A | 18.8 | A | --- |
| 257870_at | 21.9 | A | 9.9 | A | 4 | A | 1.1 | A | --- |
| 257871_at | 15.7 | A | 13.5 | A | 10.5 | A | 12.2 | A | --- |
| 257872_at | 0.7 | A | 3.9 | A | 9.8 | A | 2.3 | A | PGP16 |
| 257873_at | 1 | A | 1.4 | A | 2.1 | A | 0.8 | A | PGP17 |
| 257874_at | 536.1 | P | 150.5 | P | 285.7 | P | 192.6 | P | --- |
| 257875_at | 202.9 | P | 228.6 | P | 244.7 | P | 217.2 | P | --- |
| 257876_at | 65.8 | P | 167.5 | P | 115.4 | P | 176.4 | P | --- |
| 257877_at | 3.7 | A | 2.7 | A | 5 | A | 6.8 | A | --- |
| 257878_at | 35 | A | 5.2 | A | 47.3 | A | 22.7 | A | --- |
| 257879_at | 250.7 | P | 289.5 | P | 234.4 | P | 345.5 | P | --- |
| 257880_at | 696.5 | P | 691.4 | P | 734.6 | P | 667.6 | P | AAE7 |
| 257881_at | 22.9 | A | 24.3 | P | 37.2 | A | 25.5 | A | scpl33 |
| 257882_at | 1.7 | A | 1.1 | A | 1.6 | A | 2.1 | A | --- |
| 257883_at | 387.8 | P | 368.9 | P | 312.3 | P | 351.7 | P | --- |
| 257884_at | 7.2 | A | 17.4 | A | 2.7 | A | 22.4 | A | --- |
| 257885_at | 13.6 | A | 3.1 | A | 37.3 | A | 22.8 | A | --- |
| 257886_at | 2.1 | A | 2.3 | A | 2.3 | A | 2.1 | A | --- |
| 257887_at | 863.7 | P | 924.7 | P | 867.4 | P | 891.6 | P | --- |
| 257888_at | 270.6 | P | 312.2 | P | 184.1 | P | 397.7 | P | --- |
| 257889_at | 1.7 | A | 2.9 | A | 2.8 | A | 1.7 | A | --- |
| 257890_s_at | 295 | P | 300.6 | P | 209.7 | P | 308 | P | --- |
| 257891_at | 1040 | P | 1675.8 | P | 1266.4 | P | 1619.6 | P | RFC3 |
| 257892_at | 5801.7 | P | 5852.7 | P | 5889.5 | P | 6161.3 | P | --- |
| 257893_at | 704.4 | P | 766.7 | P | 579.1 | P | 737.5 | P | UBC32 |
| 257894_at | 495.7 | P | 500.5 | P | 351.2 | P | 449.7 | P | --- |
| 257895_at | 808.9 | P | 624.6 | P | 999.4 | P | 750.7 | P | LPD1 |
| 257896_at | 74.6 | P | 137.6 | P | 117.8 | P | 161.1 | P | --- |
| 257897_at | 337.5 | P | 275.6 | P | 356.5 | P | 242.8 | P | HAL3A |
| 257898_s_at | 1 | A | 1.1 | A | 3.1 | A | 2.6 | A | --- |
| 257899_at | 9.6 | A | 31.9 | M | 4.3 | A | 25.9 | A | --- |
| 257900_at | 27.3 | A | 40.8 | A | 45.1 | A | 50.7 | A | --- |
| 257901_at | 32.7 | A | 19.2 | A | 51.1 | A | 26.7 | A | --- |
| 257902_at | 440.5 | P | 461 | P | 385 | P | 393.6 | P | --- |
| 257903_at | 608.8 | P | 774.6 | P | 664.6 | P | 866.1 | P | --- |
| 257904_at | 3.7 | A | 18.9 | A | 23.9 | A | 15 | A | TDF1 |
| 257905_at | 98.2 | M | 82.4 | A | 29.1 | A | 93.9 | A | --- |
| 257906_at | 5597.9 | P | 4909.2 | P | 5760.9 | P | 5449 | P | ATL5 |
| 257907_at | 2701.2 | P | 2948.1 | P | 2766.5 | P | 2862.3 | P | --- |
| 257908_at | 714.7 | P | 818.9 | P | 809.6 | P | 930.5 | P | --- |
| 257909_at | 712.6 | P | 865.6 | P | 727.6 | P | 813 | P | --- |
| 257910_at | 257.7 | P | 391.9 | P | 267.6 | P | 239.9 | P | --- |
| 257911_at | 1976.6 | P | 2654 | P | 1883.5 | P | 2490.8 | P | GLYR1 |
| 257912_at | 467.6 | P | 521.4 | P | 502.3 | P | 582.8 | P | AFH1 |
| 257913_at | 483.3 | P | 589.4 | P | 587.1 | P | 663.1 | P | LAG1 |
| 257914_at | 357 | P | 454.7 | P | 405.9 | P | 431.6 | P | --- |
| 257915_at | 21.2 | A | 15.1 | A | 12.6 | A | 15.5 | A | SUP |
| 257916_at | 146.5 | P | 280.3 | P | 207.9 | P | 238.2 | P | --- |
| 257917_at | 4.3 | A | 2.3 | A | 4 | A | 0.9 | A | --- |
| 257918_at | 27.9 | A | 25 | A | 23.3 | A | 17.3 | A | --- |
| 257919_at | 105.6 | P | 57.9 | P | 89.2 | P | 48.3 | P | MYB15 |
| 257920_at | 33.4 | A | 40.9 | P | 25.7 | A | 29.1 | M | --- |
| 257921_at | 0.7 | A | 3.7 | A | 15 | A | 9.2 | A | --- |
| 257922_at | 63.8 | P | 195 | P | 55 | P | 136.1 | P | ETR2 |
| 257923_at | 71.8 | P | 157 | P | 113.6 | P | 179.6 | P | --- |
| 257924_at | 44.2 | A | 40.8 | A | 41.4 | A | 31 | A | --- |
| 257925_at | 260.1 | P | 299.5 | P | 186.9 | P | 310 | P | --- |
| 257926_at | 1805.5 | P | 2283.2 | P | 1678.9 | P | 2279.5 | P | --- |
| 257927_at | 34.1 | A | 38.2 | A | 45.1 | A | 43.7 | A | ERF1 |
| 257928_at | 24.6 | A | 46.7 | P | 58.6 | A | 67.8 | P | PPR40 |
| 257929_at | 37 | A | 40.1 | P | 43.8 | A | 31.1 | A | NRPB9A |
| 257930_at | 27.6 | A | 48.4 | A | 7.3 | A | 6.2 | A | --- |
| 257931_at | 106.9 | P | 127.8 | P | 144.6 | P | 118.8 | P | --- |
| 257932_at | 398.5 | P | 515 | P | 469.7 | P | 420.6 | P | HCF107 |
| 257933_at | 1026.1 | P | 894.6 | P | 1157.2 | P | 919.7 | P | AAPT2 |
| 257934_at | 33 | A | 27.1 | A | 18.6 | A | 44.8 | M | scpl21 |
| 257935_at | 41.9 | P | 43.6 | P | 52.5 | P | 50.1 | P | --- |
| 257936_at | 1.2 | A | 12.4 | A | 2.3 | A | 1.2 | A | --- |
| 257937_at | 1318.9 | P | 1359.3 | P | 1466 | P | 1276.1 | P | --- |
| 257938_at | 3337.3 | P | 5392.4 | P | 5981.4 | P | 5721.5 | P | DWF1 |
| 257939_at | 384.7 | P | 684.6 | P | 320.5 | P | 609.6 | P | STP4 |
| 257940_at | 757.3 | P | 489.4 | P | 628.8 | P | 583 | P | --- |
| 257941_at | 21.8 | A | 8.2 | A | 46.6 | A | 24.9 | A | UGT71B8 |
| 257942_at | 0.8 | A | 1.7 | A | 1.2 | A | 1.7 | A | ASK8 |
| 257943_at | 19.6 | P | 9.7 | P | 13.3 | A | 17.2 | A | ASK7 |
| 257944_at | 2.7 | A | 0.5 | A | 15.1 | A | 2.4 | A | ASK9 |
| 257945_at | 6.1 | A | 3.7 | A | 9.4 | A | 1.5 | A | ASK10 |
| 257946_at | 36.8 | A | 54.7 | A | 75.9 | P | 69.1 | P | --- |
| 257947_at | 19.1 | A | 4.9 | A | 6.3 | A | 4.3 | A | ICL |
| 257948_at | 222.5 | P | 195.2 | P | 230.9 | P | 175.9 | P | APO4 |
| 257949_at | 1864.5 | P | 1112.7 | P | 2131.7 | P | 1531.8 | P | UGT71B1 |
| 257950_at | 47.1 | P | 32.7 | A | 36.6 | A | 21.9 | A | UGT71B6 |
| 257951_at | 420.5 | P | 289.1 | P | 222 | P | 349.5 | P | SGP2 |
| 257952_at | 136.7 | P | 310.6 | P | 334.7 | P | 344 | P | --- |
| 257953_at | 840.8 | P | 879.4 | P | 919.9 | P | 880.8 | P | PEX22 |
| 257954_at | 683.3 | P | 587.5 | P | 866.6 | P | 708.1 | P | --- |
| 257955_at | 138.3 | P | 171.3 | P | 131.2 | P | 129 | P | --- |
| 257956_at | 114.8 | P | 160.2 | P | 103.6 | P | 119.5 | P | --- |
| 257957_at | 11.3 | A | 2.1 | A | 34.3 | A | 33.7 | A | --- |
| 257958_at | 16 | A | 2.5 | A | 14.1 | A | 0.9 | A | --- |
| 257959_at | 177.3 | P | 172.6 | P | 150.4 | P | 147.5 | P | NIK2 |
| 257960_at | 218.3 | P | 248.3 | P | 285 | P | 306.2 | P | --- |
| 257961_at | 2.6 | A | 0.7 | A | 0.2 | A | 3.6 | A | --- |
| 257962_at | 396.5 | P | 492.5 | P | 545.2 | P | 576.6 | P | --- |
| 257963_at | 64.3 | P | 188.6 | P | 97.9 | P | 164.4 | P | --- |
| 257964_at | 86.7 | P | 112.9 | P | 88.8 | P | 137.3 | P | --- |
| 257965_at | 582.5 | P | 720 | P | 512.3 | P | 569.1 | P | --- |
| 257966_at | 1289 | P | 1420.9 | P | 1411.8 | P | 1322.2 | P | --- |
| 257967_at | 1068.6 | P | 972.1 | P | 947 | P | 928.7 | P | --- |
| 257968_at | 53.2 | P | 49.4 | P | 100.7 | P | 60.7 | P | --- |
| 257969_at | 76.4 | P | 98.1 | P | 87.8 | M | 82.4 | P | --- |
| 257970_at | 1033.8 | P | 1188.6 | P | 1232.8 | P | 1591.7 | P | --- |
| 257971_at | 280.5 | P | 323.6 | P | 274.5 | P | 305.7 | P | GC6 |
| 257972_at | 375 | P | 372.7 | P | 282.4 | P | 321.4 | P | ATN1 |
| 257973_at | 332.9 | P | 310 | P | 311.6 | P | 314.7 | P | --- |
| 257974_at | 1132.1 | P | 1598.8 | P | 1440.6 | P | 1637.4 | P | --- |
| 257975_at | 38 | A | 41.3 | A | 58.6 | A | 57.2 | A | --- |
| 257976_at | 23.5 | A | 19.6 | A | 22.3 | A | 10.8 | A | PLT1 |

| | | | | | | | | | |
|---|---|---|---|---|---|---|---|---|---|
| 257977_at | 11.6 | A | 20.5 | A | 16 | A | 20.4 | A | --- |
| 257978_at | 142.5 | P | 137.4 | P | 118.2 | P | 128.2 | P | ATNEK5 |
| 257979_at | 62.7 | P | 82.5 | P | 102.7 | P | 94.6 | P | --- |
| 257980_at | 5.3 | A | 3.8 | A | 5.4 | A | 3.4 | A | --- |
| 257981_at | 1345.3 | P | 1549.4 | P | 1280.3 | P | 1328.3 | P | EIN3 |
| 257982_at | 119.1 | P | 149.3 | P | 157 | P | 168.9 | P | ATTOP6B |
| 257983_at | 1013.9 | P | 1369.8 | P | 988.3 | P | 1425 | P | --- |
| 257984_at | 1121.1 | P | 1062.4 | P | 1062.3 | P | 1104.6 | P | --- |
| 257985_at | 1698.3 | P | 1036.9 | P | 2044.4 | P | 1257.6 | P | --- |
| 257986_at | 76 | A | 66.4 | A | 33.9 | A | 67.1 | A | AGP40 |
| 257987_at | 252.8 | P | 335.3 | P | 247.1 | P | 428 | P | FIE |
| 257988_at | 944.7 | P | 938.4 | P | 998.6 | P | 1031.5 | P | --- |
| 257989_at | 354.5 | P | 448.2 | P | 327.3 | P | 426.4 | P | VPS9A |
| 257990_at | 440.3 | P | 501.6 | P | 515 | P | 438.1 | P | --- |
| 257991_at | 184.4 | P | 260.9 | P | 250.3 | P | 215.1 | P | --- |
| 257992_at | 1 | A | 1.8 | A | 6.3 | A | 12.9 | A | --- |
| 257993_at | 1 | A | 13.9 | A | 21.3 | A | 8.7 | A | --- |
| 257994_at | 3 | A | 2.4 | A | 2.9 | A | 3.4 | A | --- |
| 257995_at | 2.3 | A | 29.5 | A | 10.7 | A | 12.8 | A | --- |
| 257996_at | 620.6 | P | 565.8 | P | 560.6 | P | 573.2 | P | --- |
| 257997_at | 8.6 | A | 23.1 | A | 3.4 | A | 5.6 | A | --- |
| 257998_at | 3.8 | A | 4 | A | 8.3 | A | 1.4 | A | --- |
| 257999_at | 194.6 | P | 281 | P | 233.6 | P | 249.1 | P | --- |
| 258000_at | 6715.4 | P | 6291.2 | P | 5993.4 | P | 5806.5 | P | --- |
| 258001_at | 534.6 | P | 621.3 | P | 497.7 | P | 597.3 | P | --- |
| 258002_at | 1587.3 | P | 902.3 | P | 1217.8 | P | 610.3 | P | AIG2 |
| 258003_at | 314.1 | P | 1305 | P | 350.7 | P | 1175.6 | P | EXPA5 |
| 258004_at | 24.8 | A | 13 | A | 17.4 | A | 20.9 | A | MYB110 |
| 258005_at | 100.7 | P | 50.5 | A | 80.8 | P | 57.4 | A | --- |
| 258006_at | 313.4 | P | 387.1 | P | 237.8 | P | 292.1 | P | --- |
| 258007_at | 751 | P | 833.6 | P | 749.7 | P | 765.5 | P | ATPEN2 |
| 258008_at | 32.4 | P | 15.5 | A | 48.1 | P | 34.8 | P | --- |
| 258009_at | 127.6 | P | 129.9 | P | 148.9 | P | 124.8 | P | --- |
| 258010_at | 18.5 | A | 63.8 | M | 31.3 | A | 40.4 | A | --- |
| 258011_at | 157.8 | P | 130 | P | 209.7 | P | 141.1 | P | HAT3.1 |
| 258012_at | 30.7 | A | 20.6 | P | 63.7 | M | 32.5 | P | --- |
| 258013_at | 5.1 | A | 43.7 | A | 10.8 | A | 25.5 | A | --- |
| 258014_at | 69.1 | P | 88.2 | P | 122.2 | P | 74.1 | P | --- |
| 258015_at | 489.9 | P | 529.9 | P | 487.5 | P | 498.8 | P | --- |
| 258016_at | 137.6 | P | 79.1 | P | 103.4 | P | 93.8 | P | MPC |
| 258017_at | 142.3 | P | 177.5 | P | 168.1 | P | 191.5 | P | --- |
| 258018_at | 110.4 | M | 83.3 | A | 109.5 | P | 86.7 | A | --- |
| 258019_at | 165.8 | P | 117.2 | P | 109 | P | 94.5 | P | --- |
| 258020_at | 1867.5 | P | 1761.5 | P | 1819.5 | P | 1969.1 | P | NHD1 |
| 258021_at | 143.5 | P | 103.7 | P | 142.5 | P | 115.5 | P | PUB25 |
| 258022_at | 367.4 | P | 417.5 | P | 290.1 | P | 356.6 | P | --- |
| 258023_at | 961.9 | P | 1091.2 | P | 1462.9 | P | 1309.8 | P | ATCAD4 |
| 258024_at | 117.9 | P | 98.5 | P | 226 | P | 117.9 | P | --- |
| 258025_at | 671.8 | P | 527 | P | 570.8 | P | 627.3 | P | --- |
| 258026_at | 345.6 | P | 250.8 | P | 283.5 | P | 220 | P | ABF4 |
| 258027_at | 331.8 | P | 527.6 | P | 473.5 | P | 473.7 | P | --- |
| 258028_at | 5.6 | A | 4.5 | A | 1 | A | 2.6 | A | --- |
| 258029_at | 139.3 | P | 213.1 | P | 210.4 | P | 172.3 | P | ATPK7 |
| 258030_at | 4.2 | A | 5.8 | A | 7.2 | A | 3.2 | A | --- |
| 258031_s_at | 14.5 | A | 26.4 | A | 14.7 | A | 2 | A | --- |
| 258032_at | 5.2 | A | 18.5 | A | 34.4 | A | 6.9 | A | --- |
| 258033_at | 2045.3 | P | 1453.9 | P | 2046.8 | P | 1787.9 | P | MRP6 |
| 258034_at | 258.9 | P | 292 | P | 234.6 | P | 330.2 | P | --- |
| 258035_at | 32.7 | A | 29.1 | A | 34.2 | A | 7.8 | A | ACA9 |
| 258036_at | 36.4 | A | 31.9 | A | 20.6 | A | 21.7 | A | --- |
| 258037_at | 389.9 | P | 201 | P | 287 | P | 195.9 | P | 4CL5 |
| 258038_at | 512.6 | P | 561.2 | P | 519.2 | P | 502.2 | P | --- |
| 258039_at | 1918.8 | P | 2641.9 | P | 2247.6 | P | 2652.8 | P | --- |
| 258040_at | 628.3 | P | 1030.6 | P | 683.6 | P | 1010.9 | P | --- |
| 258041_at | 168 | P | 214.8 | P | 172 | P | 199.8 | P | ZML1 |
| 258042_at | 72.6 | P | 37.7 | P | 70.9 | P | 57.7 | P | --- |
| 258043_at | 137.6 | P | 136.3 | P | 141.6 | P | 144.7 | P | --- |
| 258044_at | 253.5 | P | 252.3 | P | 409.1 | P | 360.7 | P | ADOF2 |
| 258045_at | 515.7 | P | 438.8 | P | 466.1 | P | 486.4 | P | UBP7 |
| 258046_at | 462.9 | P | 533.6 | P | 532.1 | P | 425.2 | P | MKK5 |
| 258047_at | 728.8 | P | 444.2 | P | 674.1 | P | 562.1 | P | 4CL2 |
| 258048_at | 889.2 | P | 1034.9 | P | 1036.6 | P | 1079.6 | P | EMB2083 |
| 258049_at | 696.5 | P | 590.6 | P | 648 | P | 605.8 | P | --- |
| 258050_at | 227.9 | P | 284.6 | P | 284.8 | P | 316.1 | P | --- |
| 258051_at | 321.1 | P | 409.2 | P | 364.4 | P | 448.1 | P | TRZ4 |
| 258052_at | 1117.3 | P | 894.8 | P | 1380.8 | P | 954.5 | P | --- |
| 258053_at | 533.7 | P | 537.6 | P | 496.8 | P | 476.9 | P | --- |
| 258054_at | 2981.1 | P | 6543.1 | P | 4984.7 | P | 7069.1 | P | DELTA-TIP |
| 258055_at | 993.9 | P | 1895.3 | P | 1408.5 | P | 1914.7 | P | NDF4 |
| 258056_at | 98.2 | P | 114.9 | P | 101 | P | 126.4 | P | --- |
| 258057_at | 758.3 | P | 706.4 | P | 738 | P | 824.5 | P | AAR3 |
| 258058_at | 19.1 | A | 23.4 | A | 6.7 | A | 55.7 | A | --- |
| 258059_at | 144.6 | P | 221.7 | P | 107 | P | 177.4 | P | ATNAC3 |
| 258060_at | 715.5 | P | 873.7 | P | 660.3 | P | 647.4 | P | ATB' DELTA |
| 258061_at | 1528.6 | P | 1210.7 | P | 1598.6 | P | 1245.2 | P | --- |
| 258062_at | 600.6 | P | 581.8 | P | 474.8 | P | 518.8 | P | --- |
| 258063_at | 3999.3 | P | 2259.7 | P | 3848.2 | P | 2351.7 | P | CYP72A8 |
| 258064_at | 44 | M | 30.6 | A | 3.5 | A | 30.5 | A | CYP72A14 |
| 258065_at | 19.9 | A | 16.2 | A | 28.1 | A | 14.5 | A | --- |
| 258066_at | 66.6 | P | 76.1 | P | 128.7 | P | 78.9 | P | --- |
| 258067_at | 267 | P | 434.4 | P | 281.9 | P | 330.1 | P | --- |
| 258068_at | 86.7 | P | 76.9 | P | 85.7 | P | 60.2 | P | --- |
| 258069_at | 19.5 | A | 5.6 | A | 8.5 | A | 5.4 | A | --- |
| 258070_at | 8.8 | A | 2.7 | A | 4.4 | A | 14.3 | A | --- |
| 258071_s_at | 4223.7 | P | 4297.3 | P | 4213.6 | P | 4436.3 | P | --- |
| 258072_at | 493.7 | P | 529.9 | P | 346.7 | P | 468.4 | P | RGS1 |
| 258073_at | 25.6 | P | 14.8 | A | 20 | A | 17.8 | A | --- |
| 258074_at | 467.7 | P | 511.8 | P | 726 | P | 579.3 | P | --- |
| 258075_at | 178.1 | P | 222.7 | P | 208.2 | P | 209 | P | HMT-1 |
| 258076_at | 5054 | P | 6076 | P | 5583.1 | P | 6349.7 | P | RPL15 |
| 258077_at | 5.4 | A | 9.1 | A | 4.5 | A | 4.7 | A | --- |
| 258078_at | 14.8 | A | 92.6 | P | 99.1 | P | 78.9 | P | --- |
| 258079_at | 532.3 | P | 454.7 | P | 543.3 | P | 448.3 | P | --- |
| 258080_at | 0.8 | A | 19.7 | A | 18 | A | 16.2 | A | --- |
| 258081_at | 1133.6 | P | 1173.1 | P | 1100.4 | P | 1125.1 | P | --- |
| 258082_at | 59 | A | 58.4 | P | 79.3 | A | 74.2 | M | CLE27 |
| 258083_at | 16.2 | A | 26.4 | P | 36.7 | A | 11 | A | --- |
| 258084_at | 418.8 | P | 335.4 | P | 335 | P | 323.2 | P | --- |
| 258085_at | 748.8 | P | 812.2 | P | 831.7 | P | 651.3 | P | --- |

| | | | | | | | | | |
|---|---|---|---|---|---|---|---|---|---|
| 258086_at | 2508.9 | P | 2829.7 | P | 2910.8 | P | 2823.1 | P | LTA2 |
| 258087_at | 5889.1 | P | 7666.5 | P | 6551 | P | 8110.1 | P | ATPRX Q |
| 258088_at | 54.9 | A | 81.3 | P | 108.8 | P | 72.2 | P | --- |
| 258089_at | 247 | P | 240.8 | P | 206.4 | P | 210.9 | P | --- |
| 258090_at | 4408.2 | P | 3551.4 | P | 4166.3 | P | 4101.1 | P | --- |
| 258091_at | 54.8 | P | 124.9 | P | 102.2 | P | 118.2 | P | --- |
| 258092_at | 854 | P | 1263.1 | P | 585.8 | P | 1145.9 | P | --- |
| 258093_at | 154.8 | P | 208.6 | P | 114.1 | P | 145.8 | P | --- |
| 258094_at | 2310.7 | P | 1733.2 | P | 1500.2 | P | 1848.9 | P | CYP72A15 |
| 258095_at | 132.6 | P | 144.3 | P | 183.7 | P | 156.5 | P | --- |
| 258096_at | 3.7 | A | 3.3 | A | 4.8 | A | 7.7 | A | --- |
| 258097_at | 488.8 | P | 528.3 | P | 518.4 | P | 499.8 | P | --- |
| 258098_at | 49.2 | P | 97.3 | P | 113.9 | P | 84.1 | P | KINESIN-12B |
| 258099_at | 286.8 | P | 379.2 | P | 308 | P | 289.6 | P | --- |
| 258100_at | 925.8 | P | 321.8 | P | 697.4 | P | 419.9 | P | --- |
| 258101_at | 393.7 | P | 398.4 | P | 382.6 | P | 390.4 | P | RFR1 |
| 258102_at | 3939.3 | P | 4536.6 | P | 3272.3 | P | 4293.1 | P | --- |
| 258103_at | 25.1 | A | 34.5 | A | 41.4 | A | 40.9 | A | IPT7 |
| 258104_at | 1316.9 | P | 933.8 | P | 1346.3 | P | 985.7 | P | --- |
| 258105_at | 438.8 | P | 562 | P | 284.2 | P | 401.8 | P | --- |
| 258106_at | 1056.5 | P | 968.5 | P | 1498.9 | P | 948.1 | P | RNR2A |
| 258107_at | 1149.5 | P | 707.2 | P | 965.1 | P | 654.5 | P | ALF5 |
| 258108_at | 3660.5 | P | 1711.8 | P | 3306.2 | P | 1724 | P | --- |
| 258109_at | 742.2 | P | 820 | P | 766.7 | P | 795.9 | P | HGL1 |
| 258110_at | 151.6 | P | 149.2 | P | 141.2 | P | 157.9 | P | CYP72A7 |
| 258111_at | 17.9 | A | 20.8 | A | 46.5 | M | 26 | A | CYP72A9 |
| 258112_at | 72.2 | A | 97.4 | P | 95.1 | P | 75.3 | P | CYP72A10 |
| 258113_at | 1264.9 | P | 1153.5 | P | 1485.6 | P | 1134.9 | P | CYP72A11 |
| 258114_at | 3402.8 | P | 2762 | P | 2744.6 | P | 2940.8 | P | CYP72A13 |
| 258115_at | 7.2 | A | 18.3 | A | 9.3 | A | 5.2 | A | --- |
| 258116_at | 1 | A | 18.8 | M | 11.5 | A | 11 | A | --- |
| 258117_at | 103.6 | P | 55.8 | P | 116.1 | P | 52.9 | P | --- |
| 258118_at | 26.7 | A | 27.3 | A | 48.9 | A | 36.4 | P | --- |
| 258119_at | 400.2 | P | 563.7 | P | 400.1 | P | 578.7 | P | MPK19 |
| 258120_at | 69.5 | P | 72.5 | P | 100 | P | 73.9 | P | --- |
| 258121_s_at | 30 | A | 34.5 | P | 39.3 | A | 25.6 | A | GGPS3 |
| 258122_at | 62.1 | P | 96 | P | 75.7 | P | 124.8 | P | ATGSL04 |
| 258123_at | 490.4 | P | 588.3 | P | 557.8 | P | 566.9 | P | MPK9 |
| 258124_at | 908.1 | P | 797.8 | P | 924 | P | 737.8 | P | --- |
| 258125_s_at | 113.9 | P | 246.4 | P | 115.1 | P | 224.8 | P | --- |
| 258126_at | 512.9 | P | 482.4 | P | 508.4 | P | 529.1 | P | --- |
| 258127_at | 810 | P | 782.3 | P | 919.3 | P | 813.2 | P | --- |
| 258128_at | 906.1 | P | 1396.9 | P | 1328.5 | P | 1362.3 | P | PLSP1 |
| 258129_at | 2.9 | A | 8.2 | A | 4.3 | A | 11.3 | A | ROPGEF8 |
| 258130_at | 14.8 | A | 9.5 | A | 1.6 | A | 1.6 | A | --- |
| 258131_at | 4.3 | A | 5.5 | A | 5.1 | A | 6 | A | --- |
| 258132_at | 1656.3 | P | 1339.9 | P | 1750.9 | P | 1547.9 | P | ATPERK1 |
| 258133_at | 411.3 | P | 707.8 | P | 409.3 | P | 579.7 | P | MBF1C |
| 258134_at | 187.4 | P | 213.8 | P | 168.8 | P | 203.3 | P | --- |
| 258135_at | 402.9 | P | 458.4 | P | 451.9 | P | 409.3 | P | --- |
| 258136_at | 179.6 | P | 140.5 | P | 202.3 | P | 148.1 | P | RSY3 |
| 258137_at | 129.8 | P | 153.8 | P | 150.9 | P | 146.2 | P | UBC37 |
| 258138_at | 146.5 | P | 152.1 | P | 154.9 | P | 170.3 | P | MSH7 |
| 258139_at | 76.6 | A | 74.4 | A | 59.7 | A | 77.4 | P | HSFC1 |
| 258140_at | 1933.8 | P | 1997 | P | 2527.7 | P | 2060.6 | P | ALDH2C4 |
| 258141_at | 723 | P | 555.2 | P | 539.9 | P | 634.4 | P | HON4 |
| 258142_at | 3.6 | A | 2.2 | A | 6.3 | A | 1.9 | A | --- |
| 258143_at | 17.8 | A | 53.6 | P | 5.4 | A | 48.2 | P | --- |
| 258144_at | 8.7 | A | 5 | A | 16.7 | A | 1.2 | A | --- |
| 258145_at | 6.5 | A | 9.5 | A | 9.5 | A | 8.8 | A | --- |
| 258146_at | 1203.7 | P | 1666.4 | P | 1151.8 | P | 1425.1 | P | --- |
| 258147_at | 3.1 | A | 14.3 | A | 20.1 | A | 17.7 | M | BGLU43 |
| 258148_s_at | 414.5 | P | 455 | P | 437.7 | P | 451.1 | P | NRPD2B |
| 258149_at | 290.5 | P | 409.5 | P | 430.8 | P | 370.2 | P | EMB1270 |
| 258150_at | 187.2 | P | 243.9 | P | 278.3 | P | 248.8 | P | --- |
| 258151_at | 6041.6 | P | 6700.7 | P | 4956.8 | P | 5774.9 | P | BGLU44 |
| 258152_at | 1989.1 | P | 1734.1 | P | 2589.7 | P | 2061.2 | P | --- |
| 258153_at | 439.8 | P | 634.8 | P | 628.3 | P | 574.6 | P | MOS4 |
| 258154_at | 291.1 | P | 337.9 | P | 368 | P | 396.8 | P | --- |
| 258155_at | 2262.9 | P | 1688.9 | P | 2511.2 | P | 2181.3 | P | RACK1C_AT |
| 258156_at | 820.8 | P | 1116.9 | P | 856 | P | 1267.5 | P | --- |
| 258157_at | 77.4 | P | 98.1 | P | 109.8 | P | 85.8 | P | MYB4R1 |
| 258158_at | 1197.7 | P | 1403 | P | 905.1 | P | 1315.1 | P | PAP17 |
| 258159_at | 326.5 | P | 761.2 | P | 766.4 | P | 935.9 | P | RLK902 |
| 258160_at | 3559.6 | P | 3193.2 | P | 3988.9 | P | 2964 | P | GLN1.3 |
| 258161_at | 4025.5 | P | 3591.4 | P | 3317.7 | P | 3590.1 | P | --- |
| 258162_at | 1673.1 | P | 1351.4 | P | 1056.2 | P | 1373.7 | P | --- |
| 258163_at | 1452.2 | P | 1412.7 | P | 1155.8 | P | 1480.9 | P | --- |
| 258164_at | 279.5 | P | 315.8 | P | 389.2 | P | 307.8 | P | SURF1 |
| 258165_at | 408.1 | P | 446.7 | P | 330.6 | P | 428.5 | P | --- |
| 258166_at | 237.6 | P | 147.3 | P | 255.5 | P | 166.1 | P | --- |
| 258167_at | 1290.6 | P | 1123.2 | P | 1753.8 | P | 1481.3 | P | UGT84A2 |
| 258168_at | 1.4 | A | 10.2 | A | 36 | P | 3.9 | A | --- |
| 258169_at | 0.9 | A | 9.7 | A | 7.9 | A | 16 | A | --- |
| 258170_at | 146.3 | P | 191.3 | P | 307.7 | P | 184.9 | P | --- |
| 258171_at | 461.9 | P | 510.5 | P | 463.1 | P | 440.7 | P | --- |
| 258172_at | 3.1 | A | 10.3 | A | 2.3 | A | 4.6 | A | --- |
| 258173_at | 542.3 | P | 535.4 | P | 483.5 | P | 504.2 | P | CERK1 |
| 258174_at | 67.7 | P | 55.9 | P | 18.6 | A | 57.6 | P | --- |
| 258175_at | 176.6 | P | 168.3 | P | 150.2 | P | 174.8 | P | TWD1 |
| 258176_at | 233.8 | P | 407.1 | P | 274 | P | 377.6 | P | --- |
| 258177_at | 27.4 | A | 25.4 | A | 16.1 | A | 38.2 | A | --- |
| 258178_at | 19.5 | A | 3.7 | A | 25.2 | A | 0.9 | A | --- |
| 258179_at | 3825.9 | P | 2426.2 | P | 4432.3 | P | 2764.5 | P | --- |
| 258180_at | 82 | P | 68.4 | P | 41.5 | P | 88.3 | P | --- |
| 258181_at | 2611.5 | P | 1348.7 | P | 1790.8 | P | 1538.5 | P | --- |
| 258182_at | 132.6 | P | 10.6 | A | 11.1 | A | 56.1 | A | DXPS1 |
| 258183_at | 148.7 | P | 344.9 | P | 325.7 | P | 397.2 | P | --- |
| 258184_at | 88.1 | P | 124.5 | P | 118.7 | P | 83.4 | P | AHP1 |
| 258185_at | 201.2 | P | 263.2 | P | 295.5 | P | 331.9 | P | --- |
| 258186_s_at | 1126.9 | P | 959.2 | P | 867.1 | P | 973.2 | P | --- |
| 258187_at | 314.5 | P | 368.5 | P | 369.3 | P | 330.4 | P | ATTDX |
| 258188_at | 2225.2 | P | 3236.9 | P | 1879.6 | P | 2854.5 | P | --- |
| 258189_at | 547.8 | P | 242.4 | P | 493 | P | 327.4 | P | JAZ3 |
| 258190_at | 2.1 | A | 1.2 | A | 3.7 | A | 1.1 | A | --- |
| 258191_at | 85.5 | P | 79.5 | P | 100.9 | P | 73.6 | P | VTI13 |
| 258192_at | 27.8 | A | 17.3 | A | 18.9 | A | 4.4 | A | --- |
| 258193_at | 601.6 | P | 493.8 | P | 654.7 | P | 463.6 | P | PME31 |
| 258194_at | 413 | P | 408.3 | P | 320.3 | P | 336.2 | P | --- |

| | | | | | | | | | |
|---|---|---|---|---|---|---|---|---|---|
| 258195_at | 36.1 | A | 28.1 | A | 5.6 | A | 28.2 | A | ATMYB26 |
| 258196_at | 66.2 | A | 149.5 | P | 114 | P | 160.5 | P | --- |
| 258197_at | 84.1 | P | 67.5 | P | 58.3 | P | 66.7 | P | ATBRXL2 |
| 258198_at | 283.5 | P | 431.9 | P | 347.6 | P | 384.3 | P | NF-YA6 |
| 258199_at | 32.2 | A | 6.7 | A | 23.8 | A | 38.9 | P | --- |
| 258200_at | 37.6 | A | 43.8 | A | 65.9 | P | 52.5 | A | --- |
| 258201_at | 743.1 | P | 461.8 | P | 578.2 | P | 425.3 | P | --- |
| 258202_at | 607.7 | P | 473.7 | P | 686.1 | P | 569.9 | P | --- |
| 258203_at | 79.4 | A | 168.8 | P | 56.2 | A | 81.5 | A | --- |
| 258204_at | 73 | P | 86.3 | P | 108.3 | P | 72.9 | P | AtGRF5 |
| 258205_at | 23.3 | M | 60 | A | 53 | A | 44.4 | A | --- |
| 258206_at | 1142.5 | P | 1017.3 | P | 1175.5 | P | 1098.4 | P | CID4 |
| 258207_at | 497.1 | P | 685.9 | P | 840.6 | P | 921.2 | P | RSH2 |
| 258208_at | 1551.8 | P | 1583.3 | P | 1720.3 | P | 1411.7 | P | --- |
| 258209_at | 245.4 | P | 197.9 | P | 220 | P | 192.1 | P | GAPC1 |
| 258210_at | 6280.9 | P | 5593.9 | P | 7423.8 | P | 5842 | P | EIF4A1 |
| 258211_at | 226.8 | P | 136.9 | P | 191.8 | P | 125.8 | P | --- |
| 258212_at | 253.4 | P | 252.6 | P | 255.3 | P | 228.6 | P | --- |
| 258213_at | 117.1 | P | 147.8 | P | 139.1 | P | 153.5 | P | --- |
| 258214_at | 320.3 | P | 345.8 | P | 400.2 | P | 361.1 | P | atToc64-III |
| 258215_at | 497.6 | P | 509 | P | 477.6 | P | 561 | P | atToc64-III |
| 258216_at | 30.7 | A | 36.5 | A | 64.7 | A | 53.2 | A | --- |
| 258217_at | 1202.8 | P | 2633.4 | P | 1764.1 | P | 2779.8 | P | XPL1 |
| 258218_at | 2587.7 | P | 3618.5 | P | 2894.8 | P | 4223.6 | P | XPL1 |
| 258219_at | 652.5 | P | 577.4 | P | 633.6 | P | 554.5 | P | ATTDX |
| 258220_at | 91.9 | A | 99.7 | A | 97.6 | M | 94 | A | --- |
| 258221_at | 591.1 | P | 699.8 | P | 700.9 | P | 735.5 | P | AKIN11 |
| 258222_at | 269.5 | P | 537.8 | P | 497.3 | P | 641.4 | P | --- |
| 258223_at | 8697.9 | P | 8770 | P | 6581.9 | P | 7637.5 | P | PIFI |
| 258224_at | 2.1 | A | 1.3 | A | 2.5 | A | 11.7 | A | --- |
| 258225_at | 232.3 | P | 320.8 | P | 151.6 | P | 283.5 | P | --- |
| 258226_at | 2242.7 | P | 2588.8 | P | 1842.3 | P | 2078.8 | P | PLDALPHA1 |
| 258227_at | 93.9 | P | 101.9 | P | 113.4 | P | 81.9 | P | UVR3 |
| 258228_at | 186.5 | P | 200.4 | P | 191.8 | P | 182.7 | P | --- |
| 258229_at | 172.9 | P | 257.2 | P | 271.7 | P | 221 | P | --- |
| 258230_s_at | 19.8 | P | 9.1 | A | 39 | P | 16.7 | A | ARI13 |
| 258231_at | 7.6 | A | 31.7 | A | 3.3 | A | 21.8 | A | RCK |
| 258232_at | 168 | P | 179.2 | P | 170.3 | P | 211.9 | P | --- |
| 258233_at | 54.5 | A | 40.8 | A | 27.9 | A | 35.3 | A | MYB118 |
| 258234_at | 7.7 | A | 2.3 | A | 41.4 | A | 1.3 | A | MYB118 |
| 258235_at | 42.8 | P | 9 | A | 45.2 | A | 4 | A | AOX1C |
| 258236_at | 18.2 | A | 12.2 | A | 25.8 | A | 1.7 | A | --- |
| 258237_at | 2.2 | A | 22 | A | 3.1 | A | 18.3 | A | ATMYB21 |
| 258238_at | 367.6 | P | 435.7 | P | 383.8 | P | 399.3 | P | RST1 |
| 258239_at | 383.9 | P | 2100.5 | P | 1821.3 | P | 1816.4 | P | LHCB2.3 |
| 258240_at | 9.2 | A | 82.2 | P | 10.2 | A | 49.5 | P | OLEO4 |
| 258241_at | 8 | A | 112.2 | A | 15 | A | 55.4 | A | LBD25 |
| 258242_at | 68.7 | P | 109.1 | P | 114.1 | P | 91.9 | P | --- |
| 258243_at | 2120.6 | P | 1685.1 | P | 2123.4 | P | 1661.2 | P | CARA |
| 258244_at | 4736.3 | P | 3430.7 | P | 4146.8 | P | 3477.2 | P | --- |
| 258245_at | 625.3 | P | 640.6 | P | 435.3 | P | 562.8 | P | --- |
| 258246_s_at | 88.3 | P | 84.3 | P | 83.9 | P | 44.6 | P | --- |
| 258247_at | 137.9 | P | 128.8 | P | 144.3 | P | 143.9 | P | --- |
| 258248_at | 34.7 | A | 27.7 | A | 15.1 | A | 15.7 | A | --- |
| 258249_s_at | 690 | P | 451 | P | 493.1 | P | 488.1 | P | --- |
| 258250_at | 2532.2 | P | 3147.9 | P | 3703.8 | P | 2883.8 | P | FAD5 |
| 258251_at | 1303.4 | P | 1344.7 | P | 1153.5 | P | 1193.1 | P | --- |
| 258252_at | 472.6 | P | 335.8 | P | 514.1 | P | 323.7 | P | --- |
| 258253_at | 53.8 | P | 76.1 | P | 74.6 | P | 76.7 | P | --- |
| 258254_at | 227.4 | M | 218 | P | 165.4 | P | 204.9 | P | --- |
| 258255_at | 88.1 | P | 29.1 | A | 86.1 | P | 40.8 | A | --- |
| 258256_at | 683.5 | P | 730.4 | P | 743.4 | P | 674.5 | P | --- |
| 258257_at | 46 | A | 65.1 | P | 72.3 | P | 81.7 | P | --- |
| 258258_at | 36.9 | A | 39.4 | A | 47.5 | A | 40.2 | A | FUS3 |
| 258259_s_at | 866.4 | P | 362.9 | P | 482.3 | P | 306.1 | P | --- |
| 258260_at | 152.3 | P | 119.5 | P | 96 | P | 123.9 | P | --- |
| 258261_at | 29 | A | 42 | P | 54.4 | P | 30.2 | A | --- |
| 258262_at | 242.8 | P | 176.3 | P | 113.5 | P | 129.5 | P | --- |
| 258263_at | 320.2 | P | 287.4 | P | 303 | P | 282.4 | P | --- |
| 258264_at | 325.9 | P | 303.7 | P | 367.5 | P | 333.1 | P | MBD11 |
| 258265_at | 64.5 | A | 18.1 | A | 49 | A | 60.3 | A | --- |
| 258266_at | 13.9 | A | 4 | A | 24.5 | A | 11.9 | A | --- |
| 258267_at | 2.3 | A | 2.9 | A | 4.3 | A | 1.4 | A | --- |
| 258268_at | 634.4 | P | 642.9 | P | 686.9 | P | 676.2 | P | WSIP2 |
| 258269_at | 794.2 | P | 763.2 | P | 840.4 | P | 713.9 | P | --- |
| 258270_at | 375.5 | P | 207.1 | P | 288.3 | P | 192.9 | P | --- |
| 258271_at | 29.5 | A | 14.8 | A | 15.2 | A | 28 | A | --- |
| 258272_at | 820 | P | 942.2 | P | 761.2 | P | 904 | P | --- |
| 258273_at | 2464.2 | P | 1915.9 | P | 2518.6 | P | 1962.8 | P | GRX4 |
| 258274_at | 2794.4 | P | 1974.7 | P | 3056.4 | P | 2407.2 | P | --- |
| 258275_at | 164.9 | P | 175.9 | P | 123.4 | P | 165.4 | P | --- |
| 258276_at | 1571.9 | P | 1681.9 | P | 1668.8 | P | 1707.9 | P | --- |
| 258277_at | 887.8 | P | 297 | P | 395.9 | P | 209.2 | P | PAD3 |
| 258278_at | 1.1 | A | 1.5 | A | 1.7 | A | 0.7 | A | --- |
| 258279_at | 1.2 | A | 4.7 | A | 13.7 | A | 1.7 | A | --- |
| 258280_at | 17.2 | A | 8.1 | A | 19.8 | A | 1.8 | A | --- |
| 258281_at | 1673.7 | P | 1875.8 | P | 2018.5 | P | 1648.6 | P | --- |
| 258282_at | 147.5 | P | 123.5 | P | 129.7 | P | 125.2 | P | --- |
| 258283_at | 85.3 | P | 99 | P | 103.2 | P | 109.9 | P | --- |
| 258284_at | 5958.4 | P | 4859.5 | P | 6185.2 | P | 5427 | P | --- |
| 258285_at | 7789.9 | P | 8969 | P | 7278.1 | P | 8801.4 | P | PSAH-1 |
| 258286_at | 530.4 | P | 636.5 | P | 605.3 | P | 681.9 | P | --- |
| 258287_at | 118.9 | P | 96 | P | 139.7 | P | 107.2 | P | SULTR3;4 |
| 258288_at | 91.9 | P | 169.4 | P | 186.4 | P | 169.4 | P | LSH4 |
| 258289_at | 172.9 | P | 207.6 | P | 177.4 | P | 244.8 | P | --- |
| 258290_at | 0.9 | A | 2.1 | A | 1.7 | A | 3.3 | A | --- |
| 258291_at | 285.6 | P | 369.4 | P | 318.7 | P | 351.1 | P | --- |
| 258292_at | 124.6 | P | 91.5 | P | 115.6 | P | 75 | P | --- |
| 258293_at | 99.7 | A | 158.5 | P | 209.2 | P | 150.9 | P | PHO1 |
| 258294_at | 0.7 | A | 19.3 | A | 15.9 | A | 1.7 | A | --- |
| 258295_at | 3834.3 | P | 4799.3 | P | 3571.5 | P | 4627.5 | P | --- |
| 258296_at | 6322.7 | P | 5048.8 | P | 6694.1 | P | 5417.4 | P | --- |
| 258297_at | 1475.8 | P | 1265 | P | 1514.1 | P | 1284.3 | P | --- |
| 258298_at | 814.6 | P | 1151.4 | P | 902.4 | P | 991 | P | --- |
| 258299_at | 779.1 | P | 598.3 | P | 834.7 | P | 590.7 | P | --- |
| 258300_at | 202.9 | P | 199.8 | P | 215.3 | P | 278 | P | ckl10 |
| 258301_at | 10.7 | A | 12.5 | A | 2.7 | A | 1.2 | A | --- |
| 258302_at | 10.7 | A | 2.5 | A | 8.8 | A | 0.4 | A | --- |
| 258303_at | 1.6 | A | 1.9 | A | 2.6 | A | 5.6 | A | --- |

| | | | | | | | | | |
|---|---|---|---|---|---|---|---|---|---|
| 258304_at | 1.1 | A | 2.4 | A | 1.6 | A | 11.9 | A | --- |
| 258305_at | 21.6 | A | 2.9 | A | 3.4 | A | 1.1 | A | --- |
| 258306_at | 29.7 | A | 13.9 | A | 17.8 | A | 38.1 | P | ATBZIP42 |
| 258307_x_at | 12.1 | A | 2 | A | 3.9 | A | 0.8 | A | --- |
| 258308_at | 16.7 | A | 9.8 | A | 11.2 | A | 7.2 | A | --- |
| 258309_at | 2.1 | A | 12.7 | A | 28.2 | A | 13.9 | A | --- |
| 258310_at | 796.2 | P | 852.5 | P | 850.4 | P | 979.1 | P | ICE1 |
| 258311_at | 374.6 | P | 448.8 | P | 549.5 | P | 421.8 | P | AFB2 |
| 258312_at | 362.7 | P | 245.8 | P | 184.2 | P | 264.3 | P | --- |
| 258313_at | 393.9 | P | 312.9 | P | 399.9 | P | 366.2 | P | --- |
| 258314_at | 993 | P | 899.3 | P | 973.2 | P | 1041.4 | P | ATRABG3C |
| 258315_at | 68.2 | P | 140.8 | P | 90.9 | P | 104.5 | P | --- |
| 258316_at | 633.4 | P | 382.4 | P | 691.5 | P | 442.3 | P | --- |
| 258317_at | 78.7 | P | 104.3 | P | 101.8 | P | 101.6 | P | --- |
| 258318_at | 666.6 | P | 488.1 | P | 718.8 | P | 566.7 | P | --- |
| 258319_at | 1.1 | A | 2.9 | A | 5.4 | A | 3.5 | A | --- |
| 258320_at | 26.5 | A | 10.3 | A | 21.4 | A | 22.6 | A | --- |
| 258321_at | 875.2 | P | 1208.1 | P | 1666.1 | P | 1654.4 | P | ELIP1 |
| 258322_at | 1486.8 | P | 315.8 | P | 1512.5 | P | 345.2 | P | HMT3 |
| 258323_at | 186.1 | P | 242.5 | P | 178.2 | P | 226.9 | P | --- |
| 258324_at | 345.8 | P | 360.7 | P | 396.5 | P | 418.6 | P | TSO1 |
| 258325_at | 22 | A | 21.7 | A | 4 | A | 22.8 | P | HSFA6B |
| 258326_at | 188.5 | P | 218.6 | P | 269.8 | P | 265.8 | P | SOL1 |
| 258327_at | 37.7 | M | 52.1 | P | 24.1 | A | 12 | A | PAP85 |
| 258328_at | 168 | P | 251.3 | P | 251.6 | P | 238.8 | P | --- |
| 258329_at | 522.1 | P | 632.1 | P | 596.3 | P | 620.1 | P | ATPDIL1-6 |
| 258330_at | 11.8 | A | 26.6 | A | 50.6 | P | 28 | A | ROPGEF13 |
| 258331_at | 789.5 | P | 799.1 | P | 568.9 | P | 742.7 | P | --- |
| 258332_at | 193.6 | P | 589.9 | P | 229.8 | P | 539.9 | P | --- |
| 258333_at | 914.8 | P | 1108.5 | P | 1075.1 | P | 1212.6 | P | MFP1 |
| 258334_at | 230.1 | P | 246.7 | P | 173.5 | P | 186 | P | --- |
| 258335_at | 1.8 | A | 13.7 | A | 3.7 | A | 13.3 | A | --- |
| 258336_at | 441.4 | P | 411.6 | P | 325.9 | P | 497 | P | PDX1.2 |
| 258337_at | 1.2 | A | 12 | A | 10.7 | A | 6.2 | A | --- |
| 258338_at | 162.3 | P | 136.3 | P | 141.8 | P | 209.1 | P | --- |
| 258339_at | 48 | A | 60.2 | M | 56.2 | A | 57.5 | M | --- |
| 258340_at | 30 | P | 28.8 | A | 43 | P | 36.7 | P | --- |
| 258341_at | 86.4 | P | 206.8 | P | 117.1 | P | 179.9 | P | --- |
| 258342_at | 8.8 | A | 17.6 | A | 8.1 | A | 39.6 | A | --- |
| 258343_at | 66.2 | P | 83.2 | P | 58.3 | P | 81.2 | P | --- |
| 258344_at | 2 | A | 1.8 | A | 4.4 | A | 4.4 | A | CEG |
| 258345_at | 1607.7 | P | 1513.8 | P | 1384 | P | 1333.8 | P | --- |
| 258346_at | 58.3 | P | 46.5 | P | 60 | P | 55.6 | P | --- |
| 258347_at | 3.2 | A | 13.2 | A | 1.2 | A | 1 | A | --- |
| 258348_at | 124.9 | P | 94.5 | P | 117.7 | P | 80.4 | P | --- |
| 258349_at | 452.3 | P | 356.1 | P | 533.2 | P | 389.3 | P | HYH |
| 258350_at | 683.9 | P | 475 | P | 846.7 | P | 526.8 | P | CIPK1 |
| 258351_at | 162.7 | P | 188.7 | P | 128.7 | P | 179.3 | P | CNBT1 |
| 258352_at | 33.1 | A | 17.2 | A | 24 | A | 26.7 | A | IAA31 |
| 258353_s_at | 418.2 | P | 517.4 | P | 381.1 | P | 426.9 | P | YSL5 |
| 258354_at | 49.9 | A | 25.9 | A | 52.9 | A | 37.2 | A | --- |
| 258355_at | 179.8 | P | 232.3 | P | 234.1 | P | 258.9 | P | --- |
| 258356_at | 180.9 | P | 118.2 | P | 169.1 | P | 98 | P | --- |
| 258357_at | 717.5 | P | 892.8 | P | 704.8 | P | 820.2 | P | SRF7 |
| 258358_at | 49.6 | P | 40.4 | P | 73.1 | P | 72.7 | P | --- |
| 258359_s_at | 9703.8 | P | 10599.4 | P | 10983.6 | P | 10902.5 | P | --- |
| 258360_at | 3.6 | A | 1.9 | A | 6.5 | A | 1.8 | A | --- |
| 258361_at | 504.8 | P | 464 | P | 340.2 | P | 418.1 | P | --- |
| 258362_at | 35.7 | P | 91.1 | P | 29 | A | 76.6 | P | --- |
| 258363_at | 14 | A | 25.2 | A | 28.2 | A | 25.3 | A | ATPMEPCRC |
| 258364_at | 29.2 | P | 29.2 | M | 47.1 | A | 22.3 | A | GLIP4 |
| 258365_s_at | 2866.8 | P | 2252.4 | P | 3453.5 | P | 2409.9 | P | --- |
| 258366_at | 1910.7 | P | 2271 | P | 1907.7 | P | 2090.1 | P | RAP2.2 |
| 258367_at | 18.8 | A | 51.3 | P | 29.7 | A | 39.6 | P | WAG2 |
| 258368_at | 1077.9 | P | 1350.6 | P | 1421.6 | P | 1383.2 | P | --- |
| 258369_at | 4489.1 | P | 4481.9 | P | 4658.7 | P | 4613 | P | ATPME3 |
| 258370_at | 2.9 | A | 2.7 | A | 14.7 | A | 9.1 | A | --- |
| 258371_at | 342.9 | P | 431.2 | P | 533.8 | P | 372.9 | P | --- |
| 258372_at | 359.5 | P | 385.4 | P | 384.7 | P | 471.5 | P | UBP25 |
| 258373_at | 1610.7 | P | 1459.3 | P | 1448 | P | 1539.4 | P | PAE2 |
| 258374_at | 212.1 | P | 160 | P | 204.8 | P | 227.3 | P | --- |
| 258375_at | 208.6 | P | 322.7 | P | 179.9 | P | 263 | P | --- |
| 258376_at | 149.7 | P | 246.9 | P | 187.7 | P | 208.9 | P | --- |
| 258377_at | 39.6 | A | 57.3 | M | 46.3 | A | 33.7 | A | CNGC19 |
| 258378_at | 9.7 | A | 18.4 | A | 29 | A | 16.6 | A | --- |
| 258379_at | 80.4 | P | 124.5 | P | 91 | P | 141.1 | P | --- |
| 258380_at | 183.8 | P | 185.8 | P | 209.5 | P | 189.1 | P | --- |
| 258381_at | 643.4 | P | 762.7 | P | 602.8 | P | 755.2 | P | KINESIN-13A |
| 258382_at | 119.9 | P | 160 | P | 82 | P | 146.7 | P | UBC25 |
| 258383_at | 5.6 | A | 6.1 | A | 13.8 | A | 5.7 | A | --- |
| 258384_at | 0.9 | A | 1.8 | A | 2.4 | A | 3.2 | A | --- |
| 258385_at | 263 | P | 171 | P | 167.7 | P | 134.6 | P | ATNAC2 |
| 258386_at | 697.3 | P | 796.2 | P | 691.4 | P | 738.7 | P | --- |
| 258387_at | 30.1 | A | 55.8 | A | 38.4 | A | 60 | A | --- |
| 258388_at | 34.5 | A | 4 | A | 6.3 | A | 34.8 | A | ATEXPA12 |
| 258389_at | 44.5 | P | 78.1 | P | 70 | P | 74.3 | P | SDE5 |
| 258390_at | 382.3 | P | 494.4 | P | 432.2 | P | 507.5 | P | --- |
| 258391_at | 114.7 | P | 125.7 | P | 83.7 | P | 113.3 | P | --- |
| 258392_at | 80.1 | A | 62.6 | A | 48.3 | A | 79.8 | A | ATA20 |
| 258393_at | 792.2 | P | 1383.4 | P | 943.8 | P | 1403.5 | P | --- |
| 258394_at | 850.4 | P | 1051.6 | P | 709.3 | P | 1018 | P | --- |
| 258395_at | 263.4 | P | 63.5 | A | 193.9 | P | 79.5 | A | ANAC055 |
| 258396_at | 376.9 | P | 454 | P | 437.6 | P | 449.8 | P | --- |
| 258397_at | 320.2 | P | 117.7 | P | 391.2 | P | 118 | P | --- |
| 258398_at | 7850 | P | 8770.6 | P | 7145.4 | P | 8195.6 | P | TRX-M4 |
| 258399_at | 82.7 | P | 125.6 | P | 52.2 | A | 125 | P | IAA19 |
| 258400_at | 180.7 | P | 220.2 | P | 205.3 | P | 136.9 | P | --- |
| 258401_at | 384.1 | P | 521.4 | P | 357.8 | P | 482.2 | P | --- |
| 258402_at | 1204.6 | P | 510.1 | P | 906.6 | P | 467.7 | P | --- |
| 258403_at | 363.3 | P | 364.7 | P | 356.8 | P | 364.2 | P | --- |
| 258404_at | 770.2 | P | 662.5 | P | 864.9 | P | 775 | P | RPL3P |
| 258405_at | 400.4 | P | 373.4 | P | 463.5 | P | 399 | P | BSH |
| 258406_at | 451.7 | P | 346.6 | P | 510 | P | 383.5 | P | ATRBL14 |
| 258407_at | 4.8 | A | 4.2 | A | 8.5 | A | 1.9 | A | --- |
| 258408_at | 59.4 | P | 40.9 | P | 83.6 | P | 68.8 | P | ATCHX19 |
| 258409_at | 80.5 | A | 65.9 | P | 35.9 | P | 72.9 | P | --- |
| 258410_at | 1712.7 | P | 1627.9 | P | 2152.1 | P | 1762.6 | P | --- |
| 258411_at | 25.5 | A | 41.5 | P | 46.5 | A | 50.7 | A | --- |
| 258412_at | 4328.4 | P | 3597.6 | P | 5227.1 | P | 4127.7 | P | HS1 |

| | | | | | | | | | |
|---|---|---|---|---|---|---|---|---|---|
| 258413_at | 747.5 | P | 538.2 | P | 808.8 | P | 534.4 | P | --- |
| 258414_at | 75.9 | P | 132.2 | P | 105.6 | P | 127 | P | --- |
| 258415_at | 10601.6 | P | 12339.4 | P | 12937.6 | P | 12562.2 | P | MTO3 |
| 258416_at | 210.2 | P | 243.3 | P | 241.1 | P | 199 | P | --- |
| 258417_at | 457.4 | P | 392.5 | P | 360.1 | P | 349.4 | P | --- |
| 258418_at | 269.7 | P | 483.5 | P | 544.5 | P | 651.1 | P | --- |
| 258419_at | 740.8 | P | 1556.5 | P | 1711.4 | P | 1989.1 | P | --- |
| 258420_at | 10.3 | A | 4.2 | A | 27.1 | A | 10.8 | P | --- |
| 258421_at | 267.1 | P | 126.6 | P | 330.8 | P | 112.6 | P | --- |
| 258422_at | 45.5 | P | 47.2 | P | 56.1 | P | 25.8 | A | --- |
| 258423_at | 36.2 | P | 62.6 | P | 52.8 | P | 65.4 | P | --- |
| 258424_at | 74 | P | 93.9 | P | 59.9 | P | 92.9 | P | --- |
| 258425_at | 334 | P | 334.5 | P | 326.7 | P | 353.7 | P | --- |
| 258426_at | 41.6 | A | 70.1 | P | 52.5 | A | 51.5 | M | PLDP1 |
| 258427_at | 2.7 | A | 7 | A | 21.7 | A | 1.8 | A | --- |
| 258428_at | 76.8 | A | 60.7 | M | 82.6 | A | 109 | P | --- |
| 258429_at | 124.9 | P | 154.2 | P | 182.4 | P | 176.9 | P | TOC120 |
| 258430_at | 315.8 | P | 339.3 | P | 196.5 | P | 280.9 | P | PLDP1 |
| 258431_at | 6.2 | A | 2.4 | A | 6.4 | A | 7 | A | --- |
| 258432_at | 1236.9 | P | 1421.8 | P | 1258 | P | 1538.3 | P | RALFL23 |
| 258433_at | 12913.9 | P | 12716.5 | P | 15019.2 | P | 13122.5 | P | TCTP |
| 258434_at | 4773.4 | P | 4541 | P | 4800.5 | P | 4394.5 | P | ATEBP |
| 258435_at | 389.8 | P | 375 | P | 373.6 | P | 384 | P | --- |
| 258436_at | 327.9 | P | 275.9 | P | 228.8 | P | 249.9 | P | ATL2 |
| 258437_at | 405.6 | P | 450 | P | 498.1 | P | 408.2 | P | --- |
| 258438_at | 16.2 | A | 30.4 | A | 54.5 | M | 15.6 | A | --- |
| 258439_at | 2892.7 | P | 2264.7 | P | 3177.6 | P | 2576.3 | P | mtLPD2 |
| 258440_at | 237.2 | P | 245.6 | P | 192.8 | P | 162 | P | --- |
| 258441_at | 2.1 | A | 1.2 | A | 13.8 | A | 2.2 | A | AT3G17265 |
| 258442_at | 1.3 | A | 17.9 | A | 59.1 | A | 19.9 | A | --- |
| 258443_at | 115.3 | P | 109.1 | P | 119.1 | P | 115.3 | P | --- |
| 258444_at | 1099.5 | P | 1069.7 | P | 1109.8 | P | 1063.1 | P | TIC |
| 258445_at | 37 | P | 72.6 | P | 75.4 | P | 97.2 | P | LOX5 |
| 258446_at | 85.7 | P | 103.3 | P | 100.4 | P | 92.5 | P | --- |
| 258447_at | 112.1 | P | 145.5 | P | 149.5 | P | 101.2 | P | --- |
| 258448_at | 905.5 | P | 759.4 | P | 831.8 | P | 737.6 | P | --- |
| 258449_s_at | 1551.7 | P | 1047.5 | P | 1780.8 | P | 1169.1 | P | PMH1 |
| 258450_at | 1.6 | A | 2.1 | A | 2.6 | A | 2.9 | A | --- |
| 258451_at | 11.6 | A | 7.2 | A | 8.1 | A | 6.7 | A | AOX1B |
| 258452_at | 4061.9 | P | 2237.7 | P | 3123.2 | P | 2217.3 | P | AOX1A |
| 258453_at | 2010.4 | P | 1330.5 | P | 1481.3 | P | 1367.1 | P | NRPB5 |
| 258454_at | 837.4 | P | 700.8 | P | 1060.1 | P | 804.1 | P | RPS10 |
| 258455_at | 916.5 | P | 1094.6 | P | 1069.5 | P | 1103.5 | P | --- |
| 258456_at | 240.6 | P | 428.9 | P | 312.3 | P | 395.6 | P | WNK2 |
| 258457_at | 343.6 | P | 450.7 | P | 493.5 | P | 465.6 | P | IGPD |
| 258458_at | 114.8 | P | 104.2 | P | 91.6 | P | 106.6 | P | --- |
| 258459_at | 1.5 | A | 1.9 | A | 2.6 | A | 2.4 | A | --- |
| 258460_at | 83.4 | P | 175.5 | P | 130.4 | P | 199.7 | P | ECT6 |
| 258461_at | 67.4 | P | 110 | P | 36.1 | P | 109.4 | P | --- |
| 258462_at | 108.7 | P | 204.8 | P | 135 | P | 155.3 | P | --- |
| 258463_at | 1148.4 | P | 1091.2 | P | 1022.5 | P | 1002.1 | P | --- |
| 258464_at | 38 | A | 60.7 | M | 31.3 | A | 63.7 | P | POK1 |
| 258465_at | 20 | A | 6.5 | A | 30.6 | A | 11 | A | PMEI2 |
| 258466_at | 1643.6 | P | 1445.3 | P | 1231.7 | P | 1406.7 | P | --- |
| 258467_at | 489.8 | P | 532.2 | P | 484.3 | P | 581.7 | P | --- |
| 258468_at | 38.9 | P | 223.2 | P | 58 | P | 104.6 | P | --- |
| 258469_at | 3334.7 | P | 2954.8 | P | 3489.6 | P | 2871.3 | P | PRXIIF |
| 258470_at | 745.6 | P | 946.6 | P | 862 | P | 1089.5 | P | --- |
| 258471_at | 104.9 | P | 160.1 | P | 147.9 | P | 197.2 | P | ANP3 |
| 258472_at | 484.6 | P | 437.7 | P | 295.1 | P | 385.8 | P | --- |
| 258473_s_at | 133.9 | P | 51.5 | A | 140.2 | P | 79 | A | --- |
| 258474_at | 512.2 | P | 455.7 | P | 515.6 | P | 429.6 | P | --- |
| 258475_at | 603.8 | P | 660.4 | P | 621.8 | P | 664.2 | P | emb2768 |
| 258476_at | 42.3 | P | 48.6 | P | 44 | P | 32.9 | P | --- |
| 258477_at | 48.9 | P | 49.8 | P | 72 | P | 63.8 | P | NBS1 |
| 258478_at | 449.1 | P | 420.1 | P | 442.2 | P | 377.3 | P | --- |
| 258479_at | 15.7 | A | 20 | A | 12.7 | A | 21.9 | A | --- |
| 258480_at | 342.4 | P | 656.1 | P | 417.3 | P | 707.2 | P | --- |
| 258481_at | 628.3 | P | 787 | P | 827.4 | P | 832.2 | P | LPP3 |
| 258482_at | 932 | P | 1381.5 | P | 1277.4 | P | 1438.2 | P | --- |
| 258483_at | 1083.6 | P | 1286.9 | P | 1417.4 | P | 1533 | P | MEE31 |
| 258484_at | 540.9 | P | 931.8 | P | 792.6 | P | 856.5 | P | STE1 |
| 258485_at | 1178.5 | P | 1283.1 | P | 1407.5 | P | 1559.6 | P | --- |
| 258486_at | 6167.6 | P | 4830.8 | P | 6656.4 | P | 5352.3 | P | --- |
| 258487_at | 70.8 | P | 326.5 | P | 153.5 | P | 181.9 | P | LBD41 |
| 258488_at | 726.1 | P | 746.7 | P | 668 | P | 825.1 | P | --- |
| 258489_at | 2167.3 | P | 2865.8 | P | 2517.5 | P | 2689.4 | P | GRF7 |
| 258490_at | 6.8 | A | 4.6 | A | 22.1 | A | 5.7 | A | --- |
| 258491_at | 718.6 | P | 990 | P | 734.3 | P | 716.2 | P | --- |
| 258492_at | 33.1 | P | 44.5 | P | 36.8 | P | 33 | P | --- |
| 258493_at | 157.7 | P | 128 | P | 171.6 | P | 142.9 | P | --- |
| 258494_at | 728.5 | P | 935.4 | P | 697.4 | P | 969.1 | P | --- |
| 258495_at | 875.8 | P | 940.5 | P | 1085.5 | P | 979.9 | P | --- |
| 258496_at | 360.6 | P | 306.8 | P | 244.7 | P | 316.3 | P | --- |
| 258497_at | 50.2 | P | 192.5 | P | 75.3 | P | 165.9 | P | COL2 |
| 258498_at | 544.1 | P | 122.1 | P | 646.4 | P | 171.5 | P | --- |
| 258499_at | 534.9 | P | 513 | P | 556.3 | P | 588.8 | P | RAD23-3 |
| 258500_at | 16087.8 | P | 13262.8 | P | 14250.1 | P | 13409.5 | P | CPuORF9 |
| 258501_at | 861.1 | P | 606 | P | 634.2 | P | 546.2 | P | --- |
| 258502_at | 248.4 | P | 204.8 | P | 210.6 | P | 195.6 | P | --- |
| 258503_at | 44.3 | A | 99.7 | P | 139.3 | P | 88.5 | M | --- |
| 258504_at | 61.6 | P | 71.6 | P | 44.9 | P | 63.8 | A | --- |
| 258505_at | 500.5 | P | 360.7 | P | 498.5 | P | 384.7 | P | --- |
| 258506_at | 54.7 | P | 82.9 | A | 61.4 | P | 57.5 | P | --- |
| 258507_at | 1338.5 | P | 774 | P | 1113.9 | P | 752.6 | P | --- |
| 258508_at | 3.9 | A | 4 | A | 5.3 | A | 2.5 | A | --- |
| 258509_at | 134.2 | P | 119.9 | P | 118.5 | P | 147.1 | P | --- |
| 258510_at | 53.1 | A | 50.2 | P | 88 | P | 62.5 | P | --- |
| 258511_at | 278 | P | 368.4 | P | 303.3 | P | 368.8 | P | --- |
| 258512_at | 2751.2 | P | 2563.5 | P | 2732.5 | P | 2832.5 | P | --- |
| 258513_at | 18.5 | A | 10.5 | A | 1.3 | A | 11.9 | A | --- |
| 258514_at | 865.7 | P | 910.5 | P | 993.3 | P | 953.9 | P | --- |
| 258515_at | 1849 | P | 2094.5 | P | 1803.2 | P | 2028 | P | ACLB-1 |
| 258516_at | 60.6 | P | 32.7 | P | 39.9 | A | 25 | A | MYB108 |
| 258517_at | 1239.1 | P | 1350.4 | P | 967.5 | P | 1362.6 | P | GALK |
| 258518_at | 245.2 | P | 150.6 | P | 235.9 | P | 176.5 | P | --- |
| 258519_at | 885.7 | P | 725.8 | P | 911.9 | P | 804.1 | P | --- |
| 258520_at | 37.1 | A | 36.8 | A | 53 | A | 30 | A | --- |
| 258521_at | 3160.2 | P | 2616.6 | P | 3358.7 | P | 3085.2 | P | --- |

| | | | | | | | | | |
|---|---|---|---|---|---|---|---|---|---|
| 258522_at | 145.4 | P | 204.7 | P | 181.2 | P | 228 | P | --- |
| 258523_at | 13 | A | 34.5 | P | 32.9 | A | 39.6 | P | --- |
| 258524_at | 787.5 | P | 628.9 | P | 765.4 | P | 630.1 | P | IBR3 |
| 258525_at | 310.9 | P | 241.2 | P | 212.9 | P | 235.6 | P | IBR3 |
| 258526_at | 974.2 | P | 802.1 | P | 1052.2 | P | 819.7 | P | --- |
| 258527_at | 621.3 | P | 687.5 | P | 536.9 | P | 602.7 | P | BCE2 |
| 258528_at | 301.1 | P | 372.3 | P | 374.7 | P | 392.4 | P | --- |
| 258529_at | 129.1 | P | 195.2 | P | 174.6 | P | 203.8 | P | --- |
| 258530_at | 92.6 | M | 114.5 | P | 113.5 | P | 131.8 | P | --- |
| 258531_at | 1333 | P | 1056.1 | P | 1376.9 | P | 1138.9 | P | IMPA-1 |
| 258532_at | 4457.8 | P | 4172.9 | P | 3793.2 | P | 4436.3 | P | --- |
| 258533_at | 335 | P | 356.3 | P | 384.5 | P | 366 | P | --- |
| 258534_at | 833.9 | P | 1044.4 | P | 832.1 | P | 1036.5 | P | --- |
| 258535_at | 861 | P | 1180.1 | P | 980.6 | P | 1091.2 | P | --- |
| 258536_at | 343.3 | P | 353.1 | P | 354.2 | P | 306.2 | P | --- |
| 258537_at | 358.9 | P | 493.4 | P | 462.3 | P | 435.4 | P | --- |
| 258538_at | 884.9 | P | 1122 | P | 1008.5 | P | 1044.9 | P | --- |
| 258539_at | 15.6 | A | 3.9 | A | 2.7 | A | 6.3 | A | --- |
| 258540_at | 11.1 | A | 23.6 | A | 5.1 | A | 18.5 | A | --- |
| 258541_at | 8 | A | 40.9 | A | 21.9 | A | 32.1 | M | --- |
| 258542_at | 513 | P | 417.1 | P | 435.3 | P | 467.6 | P | --- |
| 258543_at | 34.6 | P | 59 | P | 33 | P | 48.9 | P | --- |
| 258544_at | 150.5 | P | 151.3 | P | 184.9 | P | 127.4 | P | RPM1 |
| 258545_at | 851.3 | P | 646.9 | P | 902.9 | P | 641.9 | P | --- |
| 258546_at | 80.6 | P | 100.9 | P | 91.4 | P | 111.7 | P | emb1974 |
| 258547_at | 1 | A | 0.6 | A | 1.3 | A | 1.3 | A | --- |
| 258548_at | 154.6 | P | 172.6 | P | 185.6 | P | 168.3 | P | ULP1A |
| 258549_at | 446.4 | P | 392 | P | 561.9 | P | 405.9 | P | PRMT4B |
| 258550_at | 225.8 | P | 237.4 | P | 281.5 | P | 224.6 | P | --- |
| 258551_at | 134.8 | P | 95.3 | P | 187.7 | P | 139.1 | P | --- |
| 258552_at | 109.7 | P | 356.3 | P | 95.7 | A | 319.8 | P | --- |
| 258553_at | 473.2 | P | 396.7 | P | 317.7 | P | 356.9 | P | PDE320 |
| 258554_at | 744.2 | P | 1102.7 | P | 1105.5 | P | 1154.7 | P | --- |
| 258555_at | 1938.5 | P | 1833.5 | P | 1908.7 | P | 1827.8 | P | MFP2 |
| 258556_at | 471.2 | P | 530.2 | P | 449.9 | P | 529.7 | P | --- |
| 258557_at | 175.6 | P | 190.3 | P | 146.5 | P | 188.3 | P | --- |
| 258558_at | 6.9 | A | 5 | A | 20.3 | A | 3.5 | A | --- |
| 258559_at | 185.5 | P | 212.5 | P | 180.2 | P | 221.1 | P | ATCHR12 |
| 258560_at | 28.7 | A | 20.5 | A | 53.2 | A | 49.4 | A | --- |
| 258561_at | 52.1 | A | 62.3 | A | 83.2 | A | 48.1 | A | STP6 |
| 258562_at | 32.6 | A | 44.2 | P | 32.7 | A | 35.1 | P | --- |
| 258563_at | 2408.2 | P | 2023.9 | P | 1559.9 | P | 2049.3 | P | LACS6 |
| 258564_at | 23.2 | A | 14.9 | A | 30.2 | A | 13.3 | A | --- |
| 258565_at | 397.4 | P | 553.8 | P | 617.6 | P | 487.5 | P | --- |
| 258566_at | 189.8 | P | 238.6 | P | 148.5 | P | 203 | P | GLR1.1 |
| 258567_at | 340.7 | P | 348.8 | P | 338.7 | P | 378.7 | P | ATAPY1 |
| 258568_at | 474.6 | P | 563.8 | P | 562.7 | P | 589 | P | --- |
| 258569_at | 16984.7 | P | 16139.7 | P | 18913.2 | P | 17458.4 | P | emb2171 |
| 258570_at | 59.3 | A | 64.3 | A | 48.4 | A | 23.7 | A | PPCK2 |
| 258571_at | 21.7 | A | 28 | A | 4.6 | A | 3.3 | A | anac048 |
| 258572_at | 32.3 | P | 31.9 | P | 8.1 | A | 36.5 | P | --- |
| 258573_at | 704.9 | P | 852.8 | P | 711.1 | P | 774.4 | P | PTAC3 |
| 258574_at | 1 | A | 5.8 | A | 1.8 | A | 2.9 | A | --- |
| 258575_at | 910.5 | P | 739.8 | P | 658.4 | P | 764.6 | P | SEC |
| 258576_at | 784.5 | P | 513.4 | P | 560 | P | 576.1 | P | --- |
| 258577_at | 146.4 | P | 124.8 | P | 110.9 | P | 97.4 | P | --- |
| 258578_at | 4.5 | A | 32.3 | A | 21 | A | 6.7 | A | --- |
| 258579_s_at | 0.7 | A | 0.8 | A | 8 | A | 6.4 | A | --- |
| 258580_at | 23.5 | A | 38.2 | A | 37.1 | A | 11.8 | A | --- |
| 258581_at | 52.3 | P | 44.1 | P | 42.5 | A | 52 | P | --- |
| 258582_at | 53.6 | A | 31.8 | A | 74.3 | P | 48.3 | P | --- |
| 258583_at | 4.7 | A | 3 | A | 31.2 | A | 2.9 | A | AGL57 |
| 258584_at | 1812.5 | P | 2131.1 | P | 1693 | P | 1766.1 | P | SIP1A |
| 258585_at | 688.7 | P | 789.8 | P | 914.5 | P | 833.9 | P | emb2458 |
| 258586_s_at | 0.6 | A | 0.9 | A | 1.2 | A | 1 | A | --- |
| 258587_at | 148.4 | P | 216 | P | 78.9 | P | 155.1 | P | --- |
| 258588_s_at | 15529.8 | P | 14871 | P | 13360.7 | P | 16428.2 | P | GAPC1 |
| 258589_at | 3586.5 | P | 4859.1 | P | 3247 | P | 3964.4 | P | LTL1 |
| 258590_at | 0.8 | A | 3.2 | A | 1.7 | A | 3.8 | A | ARR22 |
| 258591_at | 31.1 | A | 21.5 | A | 31.6 | A | 6.8 | A | --- |
| 258592_at | 131.1 | P | 135.2 | P | 193.1 | P | 107.4 | P | --- |
| 258593_at | 533.6 | P | 817.8 | P | 592.1 | P | 884.4 | P | --- |
| 258594_at | 40.9 | A | 73 | P | 127.5 | A | 78.5 | P | --- |
| 258595_at | 358.7 | P | 453.2 | P | 414.1 | P | 477.8 | P | --- |
| 258596_at | 57.7 | A | 54.4 | P | 71.9 | A | 56.8 | P | LSH2 |
| 258597_x_at | 3 | A | 11.6 | A | 4.3 | A | 3.3 | A | --- |
| 258598_at | 0.9 | A | 14.9 | A | 12.2 | A | 14 | A | --- |
| 258599_at | 2496.6 | P | 1388 | P | 2900.2 | P | 1402.6 | P | THA2 |
| 258600_at | 0.9 | A | 0.9 | A | 2.1 | A | 2.2 | A | --- |
| 258601_at | 646.2 | P | 555.4 | P | 760.6 | P | 643.7 | P | --- |
| 258602_at | 762.6 | P | 1021.3 | P | 772.4 | P | 1159.5 | P | --- |
| 258603_at | 43.5 | P | 67.2 | P | 69.8 | P | 79.6 | P | ATHSFA1E |
| 258604_at | 75.8 | P | 88.2 | P | 102.5 | P | 83.6 | P | --- |
| 258605_at | 21.3 | A | 10.9 | A | 18.1 | A | 13.2 | A | EXL6 |
| 258606_at | 59.3 | P | 64.7 | P | 35 | A | 48.3 | P | --- |
| 258607_at | 3491.4 | P | 4779.3 | P | 4051.9 | P | 5050.3 | P | TRXF1 |
| 258608_at | 242.8 | P | 251.4 | P | 156.2 | P | 254.2 | P | --- |
| 258609_at | 1445.7 | P | 2699.4 | P | 1806.7 | P | 2667.2 | P | --- |
| 258610_at | 415.3 | P | 217.7 | P | 336.9 | P | 218.5 | P | ILR1 |
| 258611_at | 155.3 | P | 99.8 | P | 200 | P | 105.4 | P | --- |
| 258612_at | 247.8 | P | 255.7 | P | 254.9 | P | 272.9 | P | --- |
| 258613_at | 642.6 | P | 925.1 | P | 651.3 | P | 1003.7 | P | VTC4 |
| 258614_at | 729.8 | P | 1036.7 | P | 674.9 | P | 898.1 | P | --- |
| 258615_at | 1344.3 | P | 1137 | P | 1171.9 | P | 1032.8 | P | --- |
| 258616_at | 813.9 | P | 830.9 | P | 909.5 | P | 820.8 | P | --- |
| 258617_at | 86.2 | P | 93.9 | P | 72.7 | P | 102.9 | P | --- |
| 258618_at | 204.5 | P | 102 | P | 178.2 | P | 158.5 | P | GASA5 |
| 258619_at | 1826.1 | P | 2194.6 | P | 1882.5 | P | 2041.2 | P | IPP2 |
| 258620_at | 1.2 | A | 16.4 | A | 6.3 | A | 2.9 | A | MYB107 |
| 258621_at | 915.7 | P | 1275.1 | P | 1177.5 | P | 1375.7 | P | ZFN1 |
| 258622_at | 1138.8 | P | 1178.1 | P | 1418 | P | 1074.8 | P | --- |
| 258623_at | 931.4 | P | 692.4 | P | 1422.4 | P | 858.2 | P | --- |
| 258624_at | 169.6 | P | 192.6 | P | 153.7 | P | 182.3 | P | --- |
| 258625_at | 20 | A | 28.3 | A | 63.7 | A | 32 | A | PDLP4 |
| 258626_at | 67.6 | P | 52.4 | P | 79.1 | A | 46 | P | --- |
| 258627_at | 312.9 | P | 319.7 | P | 267.7 | P | 284.8 | P | PEX12 |
| 258628_at | 129.9 | P | 137.7 | P | 111.2 | P | 143.2 | P | --- |
| 258629_at | 8 | A | 17 | A | 5.6 | A | 24.2 | A | SKOR |
| 258630_at | 190.8 | P | 292.8 | P | 261.8 | P | 225.3 | P | --- |

| | | | | | | | | | |
|---|---|---|---|---|---|---|---|---|---|
| 258631_at | 14.4 | A | 3.9 | A | 19.4 | A | 12.4 | A | QRT2 |
| 258632_s_at | 303.2 | P | 309.2 | P | 220.7 | P | 429.2 | P | MAPKKK7 |
| 258633_at | 73.7 | P | 276.7 | P | 88.4 | P | 233.1 | P | SCPL27 |
| 258634_at | 198.9 | P | 146.2 | P | 180.4 | P | 135.5 | P | --- |
| 258635_at | 175.6 | P | 209.2 | P | 195.8 | P | 230.8 | P | --- |
| 258636_at | 853.7 | P | 864.7 | P | 805.5 | P | 860.2 | P | --- |
| 258637_at | 954 | P | 975 | P | 967.7 | P | 961.6 | P | --- |
| 258638_at | 683.3 | P | 780.3 | P | 781.2 | P | 928.7 | P | --- |
| 258639_at | 2.2 | A | 3.5 | A | 2 | A | 1 | A | --- |
| 258640_at | 197.4 | P | 226 | P | 181.9 | P | 208.7 | P | --- |
| 258641_at | 2251.9 | P | 3248.6 | P | 2301.3 | P | 2947.5 | P | --- |
| 258642_at | 1092.1 | P | 997.9 | P | 1224.2 | P | 1025.4 | P | --- |
| 258643_at | 1640.9 | P | 1756.1 | P | 1933.7 | P | 1790.5 | P | ATAB2 |
| 258644_at | 494.9 | P | 512.1 | P | 525 | P | 490 | P | --- |
| 258645_s_at | 27.5 | A | 4.9 | A | 9.9 | A | 5.5 | A | --- |
| 258646_at | 153.2 | P | 230.3 | P | 231.5 | P | 254.3 | P | FRD3 |
| 258647_at | 514.6 | P | 391.8 | P | 409.4 | P | 389.6 | P | --- |
| 258648_at | 8.6 | A | 16 | A | 31.6 | A | 11 | A | --- |
| 258649_at | 4547.1 | P | 5467.4 | P | 4486 | P | 5603.4 | P | CDC48 |
| 258650_at | 408.9 | P | 435 | P | 395.8 | P | 318.3 | P | --- |
| 258651_at | 192 | P | 211.1 | P | 193.3 | P | 190.3 | P | PIP5K9 |
| 258652_at | 5.2 | A | 25 | A | 5 | A | 24.8 | A | ATRABC2B |
| 258653_at | 38.8 | A | 54.2 | P | 21.8 | A | 43.4 | M | --- |
| 258654_at | 149.8 | P | 114.6 | P | 191 | P | 121.9 | P | --- |
| 258655_at | 495.4 | P | 421.7 | P | 474.2 | P | 460.5 | P | --- |
| 258656_at | 802 | P | 882.1 | P | 938.5 | P | 785.5 | P | ATRABE1E |
| 258657_at | 339.8 | P | 296 | P | 425.8 | P | 295 | P | --- |
| 258658_at | 4636.3 | P | 5085.6 | P | 5137.1 | P | 6025.4 | P | ADK1 |
| 258659_at | 367.1 | P | 381.6 | P | 408.5 | P | 374 | P | ATB' BETA |
| 258660_at | 667.5 | P | 695.2 | P | 782.1 | P | 705.7 | P | --- |
| 258661_at | 61.1 | P | 34.4 | P | 80.1 | P | 55.9 | P | --- |
| 258662_at | 943.3 | P | 1087.6 | P | 1124.4 | P | 1230.9 | P | --- |
| 258663_at | 286.6 | P | 299.2 | P | 280.5 | P | 357.3 | P | --- |
| 258664_at | 33.4 | A | 36.1 | A | 28.5 | A | 41 | P | UBC12 |
| 258665_at | 873.2 | P | 754.9 | P | 581 | P | 695.9 | P | ATH9 |
| 258666_at | 789.3 | P | 1007.2 | P | 730 | P | 1079.5 | P | KOB1 |
| 258667_at | 3.9 | A | 3.4 | A | 6 | A | 3.1 | A | --- |
| 258668_at | 236.8 | P | 344.7 | P | 302.6 | P | 286.1 | P | --- |
| 258669_at | 47.6 | A | 98.8 | P | 50.1 | P | 62.6 | P | --- |
| 258670_at | 49.7 | P | 34.6 | A | 35.4 | P | 43.1 | P | --- |
| 258671_at | 3.7 | A | 2.5 | A | 11.8 | A | 4 | A | VHA-E2 |
| 258672_at | 46.4 | P | 58.1 | M | 25.4 | A | 35.2 | A | --- |
| 258673_at | 164.1 | P | 205.3 | P | 168.4 | P | 200.3 | P | --- |
| 258674_at | 6677.9 | P | 7301.8 | P | 7487.1 | P | 6907.6 | P | --- |
| 258675_at | 2533.1 | P | 3450.4 | P | 2990.7 | P | 3420.3 | P | LTP6 |
| 258676_at | 181.1 | P | 371.1 | P | 255.5 | P | 308.7 | P | --- |
| 258677_at | 874 | P | 868.3 | P | 847 | P | 831.8 | P | PK1 |
| 258678_at | 1619.8 | P | 1264 | P | 1478.5 | P | 1142 | P | UBC11 |
| 258679_at | 4096.9 | P | 3174.4 | P | 5107.6 | P | 3905.4 | P | --- |
| 258680_at | 14686.7 | P | 15120.9 | P | 14467.8 | P | 15185.3 | P | AAC1 |
| 258681_at | 8074.1 | P | 9572.9 | P | 8720.5 | P | 8870.4 | P | --- |
| 258682_at | 439.3 | P | 240.6 | P | 303.2 | P | 202.7 | P | S6K2 |
| 258683_at | 575.3 | P | 475.5 | P | 536.7 | P | 386.6 | P | ATSIK |
| 258684_at | 646.1 | P | 781.9 | P | 752 | P | 848.5 | P | --- |
| 258685_at | 2.6 | A | 33.6 | A | 6.4 | A | 6.7 | A | --- |
| 258686_at | 3.7 | A | 33.7 | A | 9.6 | A | 3.2 | A | --- |
| 258687_at | 251.6 | P | 214.7 | P | 291.5 | P | 235.8 | P | --- |
| 258688_at | 9.3 | A | 3.4 | A | 33.9 | A | 17.1 | A | --- |
| 258689_at | 314.8 | P | 206.7 | P | 321.2 | P | 249.1 | P | --- |
| 258690_at | 54.6 | P | 59.8 | A | 60.8 | A | 57.7 | M | --- |
| 258691_at | 289.9 | P | 240.7 | P | 309.2 | P | 235.3 | P | --- |
| 258692_at | 2540.4 | P | 1616.1 | P | 2680.5 | P | 1819.1 | P | --- |
| 258693_at | 1033.9 | P | 1107.5 | P | 1018.7 | P | 1091.7 | P | --- |
| 258694_at | 22.1 | A | 40.3 | P | 48.9 | A | 18.2 | A | --- |
| 258695_at | 2.5 | A | 42.2 | P | 42.1 | A | 9.1 | A | APX2 |
| 258696_at | 432 | P | 441.5 | P | 653.5 | P | 587.5 | P | HCF152 |
| 258697_at | 117.8 | A | 120.1 | M | 93.5 | A | 99.2 | A | MCM8 |
| 258698_at | 271.4 | P | 289.8 | P | 312.9 | P | 287.7 | P | --- |
| 258699_at | 851.6 | P | 877 | P | 825 | P | 824.4 | P | --- |
| 258700_at | 41.2 | P | 45.9 | P | 69.7 | P | 45.2 | A | IQD1 |
| 258701_at | 205.7 | P | 184.3 | P | 239.8 | P | 233.7 | P | --- |
| 258702_at | 87.8 | P | 77.2 | P | 84.7 | P | 77.4 | P | --- |
| 258703_at | 27.7 | A | 28.4 | A | 40.6 | A | 29.7 | A | --- |
| 258704_at | 90.7 | P | 99.5 | P | 115.5 | P | 96.2 | P | CCR1 |
| 258705_at | 457.1 | P | 537.8 | P | 444.4 | P | 487.3 | P | --- |
| 258706_at | 1277.4 | P | 1185.4 | P | 1475.8 | P | 1253.3 | P | --- |
| 258707_at | 111 | P | 83.3 | P | 63.5 | P | 67.9 | P | --- |
| 258708_at | 242.5 | P | 405.2 | P | 236.9 | P | 339.8 | P | --- |
| 258709_at | 5803.5 | P | 5346.3 | P | 6143.5 | P | 5385.7 | P | --- |
| 258710_s_at | 618.9 | P | 909.6 | P | 651.2 | P | 907.9 | P | --- |
| 258711_at | 1.7 | A | 2.7 | A | 2.5 | A | 2.2 | A | UBQ8 |
| 258712_s_at | 11567.7 | P | 11665.9 | P | 14521.5 | P | 12633 | P | --- |
| 258713_at | 869.3 | P | 1133.7 | P | 872.1 | P | 952.3 | P | --- |
| 258714_at | 386.7 | P | 310.3 | P | 351.8 | P | 302.4 | P | --- |
| 258715_at | 8827.3 | P | 6942.5 | P | 7166.2 | P | 6941.8 | P | --- |
| 258716_at | 235.4 | P | 181 | P | 194.3 | P | 217.5 | P | --- |
| 258717_at | 835.8 | P | 1012.9 | P | 886.6 | P | 1020.1 | P | SYP71 |
| 258718_at | 203.8 | P | 187.6 | P | 172.9 | P | 169.8 | P | --- |
| 258719_at | 250.3 | P | 206.7 | P | 237.3 | P | 277.7 | P | --- |
| 258720_at | 49.2 | P | 57.2 | P | 90.9 | M | 59.1 | A | --- |
| 258721_at | 274.2 | P | 245 | P | 273.8 | P | 243.5 | P | --- |
| 258722_at | 2.4 | A | 5.4 | A | 34.1 | A | 23.3 | A | --- |
| 258723_at | 18.3 | A | 65.7 | P | 51.9 | A | 65.5 | P | --- |
| 258724_at | 5.1 | A | 29.8 | P | 19.4 | A | 15.2 | P | --- |
| 258725_at | 34.5 | A | 13.5 | A | 4.8 | A | 13.7 | A | --- |
| 258726_at | 149.6 | P | 133 | P | 134.1 | P | 140.4 | P | --- |
| 258727_at | 3818.3 | P | 3636.1 | P | 4416 | P | 3415.7 | P | --- |
| 258728_at | 854.5 | P | 1072.4 | P | 837.7 | P | 965.7 | P | --- |
| 258729_at | 806.6 | P | 554.8 | P | 758.7 | P | 578.4 | P | ANT1 |
| 258730_at | 932 | P | 1018.7 | P | 1130.1 | P | 972.6 | P | ATFP8 |
| 258731_at | 566.9 | P | 409.1 | P | 399.1 | P | 430.4 | P | --- |
| 258732_at | 11.3 | A | 31.9 | A | 9.4 | A | 17.7 | A | --- |
| 258733_at | 188.2 | P | 223 | P | 234.1 | P | 261.4 | P | --- |
| 258734_at | 1.5 | A | 5.1 | A | 1.8 | A | 0.9 | A | --- |
| 258735_at | 8353.8 | P | 6819.5 | P | 6699.4 | P | 7712.6 | P | RCI2A |
| 258736_at | 165.4 | P | 261.5 | P | 145.4 | P | 224.7 | P | --- |
| 258737_at | 244.8 | P | 314.7 | P | 211 | P | 276.9 | P | --- |
| 258738_at | 210.4 | P | 231.8 | P | 247.8 | P | 226.8 | P | --- |
| 258739_s_at | 0.9 | A | 1.6 | A | 0.9 | A | 1.1 | A | --- |

| | | | | | | | | | |
|---|---|---|---|---|---|---|---|---|---|
| 258740_at | 10.4 | A | 21.6 | A | 31.8 | A | 19 | A | LON3 |
| 258741_at | 10.4 | A | 13.7 | A | 32.8 | A | 15.8 | A | LON4 |
| 258742_at | 193.7 | P | 191.6 | P | 201.1 | P | 249 | P | --- |
| 258743_s_at | 2049.6 | P | 2334.8 | P | 1768.2 | P | 2481.2 | P | ATSK12 |
| 258744_at | 110.3 | P | 139.2 | P | 92.6 | P | 106.9 | P | --- |
| 258745_at | 2 | A | 1.7 | A | 3.3 | A | 1.5 | A | --- |
| 258746_at | 23.6 | A | 14.5 | A | 47.1 | A | 4 | A | --- |
| 258747_at | 436.2 | P | 420.7 | P | 517 | P | 467.5 | P | --- |
| 258748_at | 12.4 | A | 23.5 | A | 39.7 | A | 31.1 | A | GLP8 |
| 258749_at | 744.6 | P | 862 | P | 757.2 | P | 773.5 | P | --- |
| 258750_at | 1451.4 | P | 1514.3 | P | 1255.1 | P | 1629.5 | P | --- |
| 258751_at | 83.9 | P | 152.9 | P | 110.6 | P | 183.3 | P | RCI2B |
| 258752_at | 64.9 | M | 120.1 | P | 86.9 | A | 114.2 | A | ATEXO70H4 |
| 258753_at | 5.9 | A | 4.8 | A | 4.1 | A | 2.9 | A | ATEXO70H3 |
| 258754_at | 21.5 | A | 2.6 | A | 5.6 | A | 1.5 | A | --- |
| 258755_at | 1587.1 | P | 1804.4 | P | 1729 | P | 1709.5 | P | --- |
| 258756_at | 248.5 | P | 187.8 | P | 179.4 | P | 166.9 | P | --- |
| 258757_at | 205.7 | P | 47.9 | A | 113.3 | P | 71.8 | P | --- |
| 258758_at | 452.2 | P | 405.6 | P | 446.3 | P | 394.9 | P | --- |
| 258759_at | 663.4 | P | 725.6 | P | 625.9 | P | 638.1 | P | BZIP28 |
| 258760_at | 22.7 | A | 31.4 | A | 43.2 | A | 6.7 | A | --- |
| 258761_at | 369.5 | P | 495.6 | P | 368.8 | P | 501 | P | --- |
| 258762_at | 20 | A | 8.9 | A | 34.2 | A | 24.7 | A | --- |
| 258763_s_at | 17.3 | A | 1.8 | A | 1.6 | A | 0.7 | A | --- |
| 258764_at | 656.9 | P | 963.1 | P | 811.2 | P | 751 | P | --- |
| 258765_at | 1.2 | A | 1.4 | A | 2.7 | A | 1.6 | A | --- |
| 258766_at | 195.6 | P | 185.6 | P | 189.2 | P | 287.4 | P | --- |
| 258767_at | 18 | A | 5.9 | A | 2.6 | A | 3.5 | A | --- |
| 258768_at | 18.6 | P | 27.2 | P | 6.8 | A | 40.1 | P | --- |
| 258769_at | 129.1 | P | 200.7 | P | 150.2 | P | 187.6 | P | MES17 |
| 258770_at | 27.9 | A | 18.5 | A | 31.9 | P | 28.6 | A | --- |
| 258771_at | 1592.9 | P | 1959.5 | P | 1888.2 | P | 1822.1 | P | --- |
| 258772_at | 6513.6 | P | 6016.5 | P | 5484.4 | P | 5287.5 | P | --- |
| 258773_at | 218.7 | P | 255.9 | P | 290.7 | P | 233.4 | P | --- |
| 258774_at | 538.5 | P | 855.5 | P | 560.5 | P | 671.9 | P | ASD1 |
| 258775_at | 3003.2 | P | 3057.5 | P | 2866.8 | P | 2894.2 | P | GLY2 |
| 258776_at | 2099.8 | P | 1561.3 | P | 1745.2 | P | 1519.6 | P | --- |
| 258777_at | 135.7 | P | 178 | P | 112.9 | P | 176.8 | P | --- |
| 258778_s_at | 105.5 | P | 73 | P | 92 | P | 109.4 | P | --- |
| 258779_at | 4.5 | A | 4.5 | A | 6.1 | A | 3.4 | A | --- |
| 258780_at | 1084.3 | P | 1021 | P | 1247.3 | P | 1071.3 | P | UBP13 |
| 258781_at | 12.7 | A | 10.2 | A | 48.2 | A | 20.9 | A | --- |
| 258782_at | 92.1 | P | 117.3 | P | 104.5 | P | 95.4 | P | --- |
| 258783_at | 196.4 | P | 228.1 | P | 225.9 | P | 228.1 | P | --- |
| 258784_at | 772.5 | P | 734.8 | P | 936.8 | P | 703.6 | P | --- |
| 258785_at | 17.1 | A | 23.5 | A | 2.8 | A | 2.5 | A | --- |
| 258786_at | 651.5 | P | 1005.2 | P | 562.7 | P | 780.1 | P | SYP121 |
| 258787_at | 138.9 | P | 129.9 | P | 90 | P | 123.5 | P | PUB24 |
| 258788_at | 1312.2 | P | 1456.8 | P | 1097.3 | P | 1158.5 | P | --- |
| 258789_at | 367.9 | P | 384.6 | P | 319.9 | P | 390.9 | P | --- |
| 258790_at | 1063.1 | P | 1004.5 | P | 1166.6 | P | 1047.9 | P | FLK |
| 258791_at | 3913.8 | P | 3780.8 | P | 3504.5 | P | 2822.6 | P | PR4 |
| 258792_at | 513.6 | P | 347.9 | P | 318.1 | P | 320.3 | P | --- |
| 258793_at | 697 | P | 775.5 | P | 624.8 | P | 817.4 | P | --- |
| 258794_at | 355.2 | P | 330.4 | P | 364.9 | P | 295 | P | --- |
| 258795_at | 18.8 | A | 10.2 | A | 29.8 | A | 29.9 | A | --- |
| 258796_at | 269.4 | P | 346.4 | P | 329.4 | P | 295.1 | P | WDL1 |
| 258797_at | 1673.6 | P | 1605.1 | P | 2134 | P | 1783.8 | P | IAA16 |
| 258798_at | 9.1 | A | 10.1 | A | 13.8 | A | 22.6 | A | --- |
| 258799_at | 479.6 | P | 398 | P | 653 | P | 414.7 | P | RPSAb |
| 258800_at | 3429 | P | 2957.5 | P | 3514.9 | P | 3377.4 | P | --- |
| 258801_at | 440 | P | 374.3 | P | 391.6 | P | 366.3 | P | --- |
| 258802_at | 218.4 | P | 360.8 | P | 222.8 | P | 303.1 | P | --- |
| 258803_at | 731.1 | P | 496.5 | P | 539.8 | P | 548.2 | P | WRKY39 |
| 258804_at | 1075 | P | 1951.1 | P | 1472.6 | P | 1558.5 | P | --- |
| 258805_at | 189.6 | P | 227.4 | P | 131.8 | P | 252.5 | P | --- |
| 258806_at | 75.9 | P | 96.1 | P | 101.5 | P | 109.5 | P | --- |
| 258807_at | 91.1 | P | 101 | P | 124.5 | P | 115.5 | P | --- |
| 258808_at | 12.6 | A | 30.4 | A | 29.9 | A | 37.7 | A | --- |
| 258809_at | 697.5 | P | 364.1 | P | 235 | P | 316 | P | anac047 |
| 258810_at | 85.5 | P | 125.7 | P | 142.2 | P | 106 | P | --- |
| 258811_at | 2374.5 | P | 2262.7 | P | 2342.6 | P | 2269.6 | P | --- |
| 258812_at | 816.9 | P | 743.6 | P | 804.4 | P | 770.2 | P | ECT1 |
| 258813_at | 115.4 | P | 75 | P | 97.1 | P | 70.9 | P | anac046 |
| 258814_at | 530.3 | P | 570.7 | P | 668 | P | 601.9 | P | --- |
| 258815_at | 3733.8 | P | 1039.8 | P | 2907.8 | P | 1614.9 | P | --- |
| 258816_at | 1299.7 | P | 1314.8 | P | 1628.8 | P | 1463.7 | P | --- |
| 258817_at | 108.8 | P | 117.5 | P | 107.2 | P | 88.2 | P | --- |
| 258818_at | 242.9 | P | 248.9 | P | 253.2 | P | 287.3 | P | EIN4 |
| 258819_at | 473.7 | P | 474 | P | 483.5 | P | 420.1 | P | --- |
| 258820_at | 1224.6 | P | 959.1 | P | 962.2 | P | 1009.7 | P | --- |
| 258821_at | 3173.2 | P | 2799.6 | P | 3047 | P | 2788.1 | P | --- |
| 258822_s_at | 405.2 | P | 480.4 | P | 422.8 | P | 430.2 | P | --- |
| 258823_at | 394 | P | 454 | P | 516 | P | 516.9 | P | --- |
| 258824_at | 234.5 | P | 194.2 | P | 254 | P | 136.4 | P | --- |
| 258825_at | 336.1 | P | 480.6 | P | 380 | P | 438.1 | P | --- |
| 258826_at | 1018.2 | P | 1151.6 | P | 961.5 | P | 1165 | P | ATGSL10 |
| 258827_at | 22.5 | A | 14.1 | A | 46.6 | A | 34.3 | A | --- |
| 258828_at | 109.4 | P | 78.2 | P | 126.3 | P | 74.4 | P | PAP15 |
| 258829_at | 905.6 | P | 946 | P | 825.9 | P | 881.5 | P | --- |
| 258830_at | 1261.3 | P | 2032.8 | P | 1161.8 | P | 1459.9 | P | --- |
| 258831_at | 263 | P | 331.4 | P | 325.8 | P | 357.2 | P | --- |
| 258832_at | 1.7 | A | 2.5 | A | 2.3 | A | 1.5 | A | --- |
| 258833_at | 247.3 | P | 244.9 | P | 268.8 | P | 246 | P | --- |
| 258834_at | 807.9 | P | 951.5 | P | 684.4 | P | 981.2 | P | --- |
| 258835_at | 22.3 | P | 31.1 | M | 42.7 | P | 31.7 | A | --- |
| 258836_at | 98 | P | 104.2 | P | 77.3 | A | 124.2 | P | --- |
| 258837_at | 7702.3 | P | 6829.6 | P | 9324 | P | 7702.6 | P | --- |
| 258838_at | 641.8 | P | 765.8 | P | 833.5 | P | 889.6 | P | --- |
| 258839_at | 488.6 | P | 563.5 | P | 575.7 | P | 575.3 | P | --- |
| 258840_at | 81.4 | P | 38.2 | P | 77.3 | P | 53.7 | P | --- |
| 258841_at | 7.6 | A | 10.8 | A | 12.5 | A | 8.3 | A | --- |
| 258842_at | 435.5 | P | 434.5 | P | 424 | P | 429.2 | P | CLPS3 |
| 258843_at | 1.1 | A | 3.4 | A | 5.1 | A | 4.9 | A | --- |
| 258844_at | 312 | P | 286.2 | P | 264 | P | 292.4 | P | SWP |
| 258845_at | 2991 | P | 2672.6 | P | 2782.3 | P | 2217.9 | P | --- |
| 258846_at | 4140.6 | P | 3682.1 | P | 3785.3 | P | 3845.7 | P | --- |
| 258847_at | 5217.3 | P | 3893.6 | P | 3751.8 | P | 4163.7 | P | --- |
| 258848_at | 414.1 | P | 549.8 | P | 415.4 | P | 517.9 | P | DCL2 |

| | | | | | | | | | |
|---|---|---|---|---|---|---|---|---|---|
| 258849_at | 3087.5 | P | 3759.5 | P | 2801.6 | P | 4554 | P | UGP |
| 258850_at | 670 | P | 561.6 | P | 592.4 | P | 566.7 | P | CSLD3 |
| 258851_at | 712.2 | P | 533.9 | P | 1031.3 | P | 500.9 | P | ATGSTF11 |
| 258852_at | 451.2 | P | 555.4 | P | 375.9 | P | 396.8 | P | AT-P4H-2 |
| 258853_at | 211.5 | P | 243.9 | P | 250 | P | 218.4 | P | --- |
| 258854_at | 24.4 | A | 12.3 | A | 32 | A | 5.8 | A | --- |
| 258855_at | 448.6 | P | 437.3 | P | 461 | P | 374.7 | P | --- |
| 258856_at | 269.1 | P | 202.6 | P | 96.3 | P | 159.5 | P | SRG3 |
| 258857_at | 666.8 | P | 808.6 | P | 759.5 | P | 895.5 | P | scpl25 |
| 258858_at | 7169.8 | P | 5988.5 | P | 8415.2 | P | 6702.6 | P | --- |
| 258859_at | 171.1 | P | 404.6 | P | 215.8 | P | 341.2 | P | --- |
| 258860_at | 764.6 | P | 851.1 | P | 621 | P | 845.6 | P | KUP3 |
| 258861_at | 499.4 | P | 609.1 | P | 613.2 | P | 538.3 | P | --- |
| 258862_at | 7362.8 | P | 6061 | P | 6785.4 | P | 6161.6 | P | MPPBETA |
| 258863_at | 232.5 | P | 266.1 | P | 194.8 | P | 210.9 | P | DCL2 |
| 258864_at | 285.2 | P | 313.3 | P | 339.9 | P | 331.3 | P | --- |
| 258865_at | 4.9 | A | 27.9 | A | 28.5 | A | 19.1 | A | anac045 |
| 258866_at | 103.4 | P | 101.4 | P | 81.1 | P | 109.4 | P | --- |
| 258867_at | 85.9 | P | 96.5 | P | 94.5 | P | 94.2 | P | --- |
| 258868_at | 250.8 | P | 362.9 | P | 284.1 | P | 374.2 | P | XPO1B |
| 258869_at | 319.3 | P | 433.1 | P | 408 | P | 389.3 | P | AtVGT1 |
| 258870_at | 35.7 | P | 27.7 | A | 35.8 | P | 27.5 | A | --- |
| 258871_at | 196.5 | P | 150.7 | P | 288.3 | A | 242.9 | P | --- |
| 258872_at | 3.8 | A | 4.3 | A | 7.8 | A | 4.6 | A | HDG8 |
| 258873_at | 21.2 | A | 47.4 | P | 6.6 | A | 35.1 | P | --- |
| 258874_at | 34.9 | A | 56.7 | A | 47.3 | A | 34.1 | A | --- |
| 258875_at | 936.4 | P | 1133 | P | 1030.9 | P | 1165.4 | P | --- |
| 258876_at | 227.5 | P | 249.8 | P | 209.3 | P | 193.7 | P | ATARFB1C |
| 258877_at | 292.4 | P | 543 | P | 436.7 | P | 516.8 | P | ATEXPA13 |
| 258878_at | 50.3 | M | 47.5 | A | 47 | A | 19.8 | A | --- |
| 258879_at | 688.2 | P | 915.5 | P | 755.4 | P | 981.6 | P | --- |
| 258880_at | 500.1 | P | 403.3 | P | 271 | P | 381.7 | P | ATG8H |
| 258881_at | 884.4 | P | 756.9 | P | 923.9 | P | 744.8 | P | --- |
| 258882_at | 393 | P | 583.9 | P | 304 | P | 436.8 | P | --- |
| 258883_at | 882.8 | P | 891.7 | P | 732.3 | P | 886.5 | P | --- |
| 258884_at | 1313.4 | P | 875.7 | P | 1652.8 | P | 1013.4 | P | OMR1 |
| 258885_at | 298.9 | P | 336.6 | P | 292.5 | P | 350.3 | P | --- |
| 258886_at | 13.8 | A | 4.5 | A | 2.8 | A | 15.2 | A | IMPA-7 |
| 258887_at | 85.1 | P | 83.2 | P | 83.3 | A | 77.8 | P | PLDP2 |
| 258888_at | 35.3 | A | 74.7 | P | 43.4 | A | 80.1 | P | --- |
| 258889_at | 1.3 | A | 3 | A | 5.4 | A | 1.7 | A | --- |
| 258890_at | 886.7 | P | 1194.5 | P | 991.8 | P | 1183.5 | P | NF-YA2 |
| 258891_at | 71.5 | A | 63.7 | P | 84.2 | P | 67.6 | P | EMB2016 |
| 258892_at | 376.8 | P | 316.7 | P | 398.3 | P | 357.8 | P | --- |
| 258893_at | 47 | P | 113.5 | P | 79.2 | P | 122.3 | P | AtRLP33 |
| 258894_at | 274.1 | P | 320.8 | P | 243.2 | P | 343.8 | P | AtRLP32 |
| 258895_at | 93.6 | P | 381.9 | P | 153.5 | P | 342.2 | P | --- |
| 258896_at | 235 | P | 311.8 | P | 252.6 | P | 323.6 | P | SYP43 |
| 258897_at | 3089 | P | 4912 | P | 5140.6 | P | 5798.5 | P | --- |
| 258898_at | 52.6 | P | 51.1 | P | 52.9 | A | 58.7 | P | RECQI1 |
| 258899_at | 100 | P | 140.6 | P | 83 | P | 101.8 | P | --- |
| 258900_at | 7969.6 | P | 7084.4 | P | 7302.4 | P | 7074.9 | P | RPL18 |
| 258901_at | 235 | P | 158.9 | P | 127.7 | P | 217.3 | P | --- |
| 258902_at | 1840 | P | 2104.6 | P | 1729 | P | 1683.6 | P | PDK |
| 258903_at | 689.5 | P | 642.6 | P | 428.8 | P | 586.7 | P | --- |
| 258904_at | 1205.8 | P | 1238 | P | 1254.1 | P | 1217.7 | P | CHR11 |
| 258905_at | 3.8 | A | 2.9 | A | 29.5 | A | 13.3 | A | --- |
| 258906_at | 396.5 | P | 377.6 | P | 355.1 | P | 271.8 | P | ATTLP9 |
| 258907_at | 45.9 | A | 42.1 | A | 54.9 | A | 29.9 | A | NHX4 |
| 258908_at | 1823.1 | P | 1677.4 | P | 1818.5 | P | 1646.3 | P | MEE32 |
| 258909_at | 263.5 | P | 279.2 | P | 247.3 | P | 221.4 | P | --- |
| 258910_at | 311.3 | P | 351.6 | P | 358.2 | P | 367 | P | --- |
| 258911_at | 533.4 | P | 537.5 | P | 495.4 | P | 478 | P | --- |
| 258912_at | 48.6 | A | 81.5 | A | 81.9 | A | 57.6 | A | --- |
| 258913_at | 872 | P | 939.3 | P | 834.6 | P | 935.8 | P | --- |
| 258914_at | 7.3 | A | 25.3 | A | 10.5 | A | 29.7 | A | AGP27 |
| 258915_at | 513.9 | P | 473.5 | P | 307.2 | P | 430.3 | P | VPS60.1 |
| 258916_at | 1704.2 | P | 1686.3 | P | 2224.9 | P | 1666.2 | P | ATNAP7 |
| 258917_at | 307.1 | P | 341 | P | 246.3 | P | 276.1 | P | --- |
| 258918_at | 24.7 | A | 28.1 | A | 30.6 | A | 49.1 | A | UNE9 |
| 258919_at | 1326 | P | 988.2 | P | 1201.8 | P | 1177.1 | P | --- |
| 258920_at | 1747.2 | P | 1780.4 | P | 2352.4 | P | 1875.6 | P | --- |
| 258921_at | 998 | P | 1012.7 | P | 788.9 | P | 826.6 | P | anac053 |
| 258922_at | 1681.5 | P | 1393.2 | P | 2035.7 | P | 1630.3 | P | --- |
| 258923_at | 738 | P | 328.3 | P | 577.8 | P | 281 | P | SCPL7 |
| 258924_at | 617.9 | P | 669.5 | P | 686.9 | P | 648.9 | P | ATNUDX26 |
| 258925_at | 785.9 | P | 911.2 | P | 687.7 | P | 645 | P | --- |
| 258926_s_at | 553.8 | P | 467.2 | P | 344.6 | P | 455.9 | P | anac052 |
| 258927_at | 437.6 | P | 523.4 | P | 459.5 | P | 637.8 | P | DFC |
| 258928_at | 151.1 | P | 187.9 | P | 210.2 | P | 177.7 | P | TAF12 |
| 258929_at | 1046.5 | P | 1411.6 | P | 1058.7 | P | 1391.4 | P | --- |
| 258930_at | 19 | A | 65 | P | 54 | P | 71.6 | P | --- |
| 258931_at | 120.5 | P | 175.9 | P | 148 | P | 189.5 | P | DML2 |
| 258932_at | 41.7 | A | 46.3 | A | 3.8 | A | 45.6 | M | PAP16 |
| 258933_at | 315 | P | 561.7 | P | 405.9 | P | 602.3 | P | --- |
| 258934_at | 271.8 | P | 326.5 | P | 290.6 | P | 305.4 | P | RECA3 |
| 258935_at | 64.2 | P | 155.4 | P | 216.8 | P | 170.2 | P | --- |
| 258936_s_at | 1.2 | A | 1.8 | A | 13.2 | A | 1.8 | A | --- |
| 258937_at | 1563.6 | P | 1512.8 | P | 2251.2 | P | 1322.9 | P | --- |
| 258938_at | 390.1 | P | 532.1 | P | 617.6 | P | 595.5 | P | --- |
| 258939_at | 983.9 | P | 589.8 | P | 665.4 | P | 630 | P | --- |
| 258940_at | 14.6 | A | 22.4 | A | 46.7 | A | 20 | A | --- |
| 258941_at | 230.2 | P | 39.2 | A | 153.5 | P | 25 | A | MDHAR |
| 258942_at | 16.1 | A | 23.3 | A | 34 | A | 27.7 | P | --- |
| 258943_at | 103.9 | P | 93.8 | P | 90.2 | P | 86.2 | P | --- |
| 258944_at | 166.7 | P | 133.6 | P | 182.4 | P | 156.4 | P | FLD |
| 258945_at | 50.3 | P | 103.9 | P | 49.8 | A | 64.9 | P | CPK2 |
| 258946_at | 369.8 | P | 403.6 | P | 330.9 | P | 346.9 | P | --- |
| 258947_at | 160.6 | P | 356 | P | 76.8 | P | 173.8 | P | --- |
| 258948_at | 3039.4 | P | 2839.5 | P | 2755.8 | P | 2899.3 | P | SOX |
| 258949_at | 1057.7 | P | 947.2 | P | 851.7 | P | 994.5 | P | ATCFM2 |
| 258950_at | 129.8 | P | 187.9 | P | 108.4 | P | 169.7 | P | MBD9 |
| 258951_at | 259.9 | P | 302.1 | P | 311.9 | P | 303.6 | P | --- |
| 258952_at | 106.5 | A | 207.8 | M | 183.2 | A | 194.3 | P | --- |
| 258953_at | 206.1 | P | 146.5 | P | 106.4 | A | 128.5 | A | --- |
| 258954_at | 256.9 | P | 244.4 | P | 226.3 | P | 246.1 | P | --- |
| 258955_s_at | 38.5 | P | 48.1 | P | 43.9 | P | 69 | P | --- |
| 258956_at | 446.4 | P | 805 | P | 653 | P | 831.9 | P | --- |
| 258957_at | 21.5 | A | 74.2 | M | 76.4 | P | 49.1 | A | DOX1 |

| | | | | | | | | | |
|---|---|---|---|---|---|---|---|---|---|
| 258958_at | 2582.3 | P | 2830.2 | P | 2343.7 | P | 2543.7 | P | VMA10 |
| 258959_at | 35.2 | A | 5.3 | A | 46.6 | A | 33 | A | CAT7 |
| 258960_at | 15.1 | A | 40.9 | A | 46.3 | A | 21 | A | --- |
| 258961_at | 32.7 | A | 41.3 | P | 65 | A | 28.5 | A | --- |
| 258962_at | 102.5 | P | 114.2 | P | 147.3 | P | 202.7 | P | CYP77A6 |
| 258963_at | 208.8 | P | 212.1 | P | 242.3 | P | 187 | P | --- |
| 258964_at | 150 | P | 214.8 | P | 167.1 | P | 228.4 | P | --- |
| 258965_at | 454.9 | P | 353.3 | P | 526.5 | P | 447.7 | P | --- |
| 258966_at | 790.4 | P | 933.3 | P | 1047.5 | P | 1022.8 | P | --- |
| 258967_at | 46.6 | P | 49.3 | P | 42.9 | A | 29.8 | A | --- |
| 258968_at | 27.1 | A | 22 | A | 51.8 | A | 13.8 | A | --- |
| 258969_at | 29.3 | A | 30.7 | A | 58.3 | P | 34.9 | A | --- |
| 258970_at | 3900.4 | P | 4342.7 | P | 3925 | P | 4439 | P | scpl49 |
| 258971_at | 47.3 | A | 38 | A | 29.5 | A | 47.1 | A | ACR6 |
| 258972_at | 548.9 | P | 700 | P | 458.6 | P | 523.3 | P | --- |
| 258973_at | 6.8 | A | 3.7 | A | 8.7 | A | 8.1 | A | CYP94B2 |
| 258974_at | 144.7 | P | 88.5 | P | 160.6 | P | 39.9 | M | --- |
| 258975_at | 894.9 | P | 345.7 | P | 736 | P | 328.6 | P | WRKY45 |
| 258976_at | 352.1 | P | 325 | P | 362.7 | P | 345.1 | P | --- |
| 258977_s_at | 1751.8 | P | 1509.2 | P | 2151.3 | P | 1391.1 | P | AK3 |
| 258978_at | 303.5 | P | 328.1 | P | 272.3 | P | 350.2 | P | --- |
| 258979_at | 3961.3 | P | 5052 | P | 4234.3 | P | 4517.4 | P | --- |
| 258980_at | 24.1 | A | 34.9 | A | 29.7 | A | 31.5 | A | RGP3 |
| 258981_at | 33.9 | P | 55.4 | P | 50.8 | P | 49.7 | P | --- |
| 258982_at | 40.7 | M | 63.6 | P | 3 | A | 64.5 | P | --- |
| 258983_at | 87.6 | P | 130.7 | P | 101.6 | P | 118.9 | P | --- |
| 258984_at | 185.6 | P | 175 | P | 108 | P | 139.2 | P | ATERDJ3A |
| 258985_at | 212.6 | P | 207.3 | P | 172.3 | P | 225.8 | P | --- |
| 258986_at | 142.1 | P | 116.3 | P | 165.6 | P | 138.2 | P | --- |
| 258987_at | 440.3 | P | 390.1 | P | 512.1 | P | 436.9 | P | --- |
| 258988_at | 284.3 | P | 293.3 | P | 357.8 | P | 300.4 | P | --- |
| 258989_at | 891.8 | P | 1226.8 | P | 691.9 | P | 1095.8 | P | --- |
| 258990_at | 206.8 | P | 211.1 | P | 214.8 | P | 200.3 | P | --- |
| 258991_at | 43 | P | 29.1 | A | 65.4 | P | 72.7 | P | --- |
| 258992_at | 1100.8 | P | 1505 | P | 858.6 | P | 1441.6 | P | --- |
| 258993_at | 3120.5 | P | 5599.5 | P | 2681 | P | 4380.5 | P | LHCB4.2 |
| 258994_at | 311.4 | P | 334.7 | P | 311.9 | P | 338.1 | P | RAPTOR1 |
| 258995_at | 1640.6 | P | 1439.1 | P | 1714.1 | P | 1680.5 | P | --- |
| 258996_at | 404.1 | P | 329.2 | P | 369.8 | P | 315.3 | P | --- |
| 258997_at | 401.5 | P | 521.6 | P | 433.8 | P | 527.6 | P | --- |
| 258998_at | 601.3 | P | 493.3 | P | 693.9 | P | 577.3 | P | --- |
| 258999_at | 846.7 | P | 601.4 | P | 633.2 | P | 577.6 | P | --- |
| 259000_at | 292.4 | P | 392.2 | P | 393.1 | P | 442.4 | P | --- |
| 259001_at | 42.1 | P | 33.7 | M | 61.8 | P | 62.3 | P | --- |
| 259002_at | 7.2 | A | 38.9 | A | 10.5 | A | 33.2 | A | ROXY1 |
| 259003_at | 75.5 | P | 82.7 | P | 83.9 | P | 61.5 | P | --- |
| 259004_at | 18.8 | A | 26.4 | A | 3.3 | A | 8 | A | --- |
| 259005_at | 319.1 | P | 313.5 | P | 329 | P | 296 | P | --- |
| 259006_at | 13285.9 | P | 11476.7 | P | 14931.9 | P | 13304.7 | P | --- |
| 259007_at | 153.9 | P | 186.8 | P | 198.9 | P | 173.2 | P | MYB3R-3 |
| 259008_at | 20519.1 | P | 18598.1 | P | 21974.6 | P | 19696.5 | P | MT2A |
| 259009_at | 10963.6 | P | 7729.1 | P | 10698.8 | P | 8910.9 | P | PYK10 |
| 259010_at | 81.7 | P | 74.4 | P | 93.1 | P | 60.9 | P | --- |
| 259011_s_at | 4.2 | A | 1.3 | A | 1.7 | A | 1.6 | A | --- |
| 259012_at | 502.8 | P | 357.5 | P | 437 | P | 423.5 | P | PUB9 |
| 259013_at | 3852.6 | P | 3779.7 | P | 4782.4 | P | 3516.3 | P | emb1990 |
| 259014_at | 388.9 | P | 617.6 | P | 381.9 | P | 591.7 | P | --- |
| 259015_at | 49.7 | A | 64.5 | A | 80.8 | A | 28.3 | A | --- |
| 259016_at | 4744.7 | P | 4542.3 | P | 4944.1 | P | 4616.6 | P | --- |
| 259017_at | 508.9 | P | 690.1 | P | 650.8 | P | 593.5 | P | --- |
| 259018_at | 1101.1 | P | 1118.2 | P | 669.5 | P | 1272.8 | P | AIR12 |
| 259019_at | 225.4 | P | 206.5 | P | 220.6 | P | 185.3 | P | CHIP |
| 259020_at | 1814 | P | 2084.9 | P | 2171.3 | P | 2283.8 | P | --- |
| 259021_at | 205.6 | P | 234.1 | P | 201.5 | P | 250.6 | P | --- |
| 259022_at | 81.1 | P | 125.3 | P | 118.5 | P | 92.1 | P | NS2 |
| 259023_at | 285.1 | P | 315.4 | P | 280 | P | 351.7 | P | --- |
| 259024_at | 5.2 | A | 28.1 | A | 29.9 | A | 26.7 | A | --- |
| 259025_at | 61.6 | A | 25 | A | 46.2 | A | 20.1 | A | --- |
| 259026_at | 10.7 | A | 6.9 | A | 3.3 | A | 5.5 | A | --- |
| 259027_at | 32.1 | A | 20.5 | A | 53.8 | A | 19.9 | A | --- |
| 259028_at | 2.2 | A | 1.3 | A | 2.1 | A | 1.1 | A | TAC1 |
| 259029_at | 962.2 | P | 922.2 | P | 1024.6 | P | 983.1 | P | ORP3B |
| 259030_at | 131.4 | P | 160.8 | P | 159.1 | P | 158.7 | P | --- |
| 259031_at | 203.5 | P | 122.7 | P | 174.4 | P | 95.9 | P | --- |
| 259032_at | 2.7 | A | 4.8 | A | 30 | A | 28.9 | A | --- |
| 259033_at | 348.7 | P | 147.5 | P | 224.3 | P | 107.8 | P | --- |
| 259034_at | 261.7 | P | 360.1 | P | 306.5 | P | 303.9 | P | --- |
| 259035_at | 26 | A | 18.4 | A | 2.9 | A | 6.7 | A | --- |
| 259036_at | 28.6 | A | 41.2 | P | 59.1 | P | 36.6 | P | LAC7 |
| 259037_at | 354.4 | P | 469.6 | P | 381.3 | P | 392.6 | P | --- |
| 259038_at | 1125.1 | P | 1342.8 | P | 1395.3 | P | 1229.9 | P | PTAC13 |
| 259039_at | 999.4 | P | 1131.2 | P | 904.7 | P | 995.1 | P | --- |
| 259040_at | 732.7 | P | 480.7 | P | 632.6 | P | 476.9 | P | ATGSTU8 |
| 259041_at | 254.6 | P | 372 | P | 337.2 | P | 304.8 | P | ATCSLC6 |
| 259042_at | 57.8 | P | 71.9 | P | 91.9 | P | 88.5 | P | RGL2 |
| 259043_at | 98.1 | P | 127.8 | P | 101.4 | P | 137.7 | P | --- |
| 259044_at | 1.5 | A | 2.9 | A | 17.5 | A | 23.3 | A | --- |
| 259045_at | 0.6 | A | 1 | A | 1.2 | A | 0.8 | A | --- |
| 259046_at | 2.7 | A | 1.9 | A | 2.1 | A | 2 | A | --- |
| 259047_at | 1754.3 | P | 1493.2 | P | 1405.5 | P | 1386 | P | DegP7 |
| 259048_at | 426 | P | 501.4 | P | 483.3 | P | 466.5 | P | DegP7 |
| 259049_at | 59.1 | P | 41.2 | A | 97.4 | P | 45.7 | A | --- |
| 259050_at | 79.1 | M | 157.3 | P | 166.7 | P | 191.4 | P | --- |
| 259051_at | 627.7 | P | 817.8 | P | 709.1 | P | 697.4 | P | --- |
| 259052_at | 549.4 | P | 481.4 | P | 477.4 | P | 432.2 | P | PEX19-1 |
| 259053_at | 184.1 | P | 222.2 | P | 222.1 | P | 227.1 | P | --- |
| 259054_at | 262.7 | P | 54 | A | 157.3 | P | 87.2 | P | CHAT |
| 259055_at | 663.8 | P | 648.2 | P | 500.4 | P | 714.9 | P | UNE6 |
| 259056_at | 559.7 | P | 553.5 | P | 514 | P | 564.6 | P | --- |
| 259057_at | 188.4 | P | 157.4 | P | 117.9 | P | 202.8 | P | --- |
| 259058_at | 1400.8 | P | 1068.8 | P | 769 | P | 843.8 | P | CYP89A9 |
| 259059_at | 688.9 | P | 573.9 | P | 696.1 | P | 633.3 | P | --- |
| 259060_at | 384.3 | P | 357.5 | P | 316.8 | P | 320.8 | P | --- |
| 259061_at | 285 | P | 319.7 | P | 202.6 | P | 312.5 | P | AtRABA5b |
| 259062_at | 234.7 | P | 241.9 | P | 312 | P | 291.4 | P | --- |
| 259063_at | 33.1 | A | 17.3 | A | 29.3 | A | 17.6 | A | --- |
| 259064_at | 19.2 | A | 14.5 | A | 20.5 | A | 2.3 | A | AGD11 |
| 259065_at | 138.2 | P | 214.6 | P | 151.3 | P | 169 | P | GLR1.4 |
| 259066_at | 106.7 | P | 86.2 | P | 77.4 | P | 73.8 | P | --- |

| | | | | | | | | | |
|---|---|---|---|---|---|---|---|---|---|
| 259067_at | 195.1 | P | 239.7 | P | 165.6 | P | 234.7 | P | --- |
| 259068_at | 1771.5 | P | 1325.7 | P | 1700.3 | P | 1353.3 | P | PEX13 |
| 259069_at | 2920.5 | P | 2691.7 | P | 4394.5 | P | 2739.6 | P | ATKRS-1 |
| 259070_at | 676.5 | P | 827.8 | P | 769.8 | P | 782.9 | P | DGD1 |
| 259071_at | 230.3 | P | 269.9 | P | 216.3 | P | 241.8 | P | NHL2 |
| 259072_at | 976.9 | P | 1360.6 | P | 961.3 | P | 1428.2 | P | FLA18 |
| 259073_at | 195.4 | P | 257.2 | P | 254.1 | P | 268.3 | P | --- |
| 259074_at | 271 | P | 273.6 | P | 300.9 | P | 265.1 | P | RPK2 |
| 259075_at | 253.7 | P | 264.3 | P | 291.3 | P | 261.2 | P | --- |
| 259076_at | 29 | A | 30.6 | A | 68.6 | A | 26.5 | A | TMAC2 |
| 259077_s_at | 4856.1 | P | 6043.4 | P | 5809.7 | P | 6602.4 | P | RGP1 |
| 259078_at | 213.4 | P | 179.9 | P | 172.2 | P | 211.9 | P | HST |
| 259079_at | 90.1 | P | 106.4 | P | 106.9 | P | 97.3 | P | --- |
| 259080_at | 1825.8 | P | 1788.3 | P | 1749 | P | 1608 | P | WNK1 |
| 259081_at | 197.8 | P | 266.5 | P | 168 | P | 321.7 | P | NHX2 |
| 259082_at | 224.1 | P | 219.7 | P | 297.5 | P | 259.5 | P | --- |
| 259083_at | 437.9 | P | 412.2 | P | 391.1 | P | 438.5 | P | ATNEK2 |
| 259084_at | 16.2 | A | 14.9 | A | 5.3 | A | 19.8 | A | ATTIM23-3 |
| 259085_at | 754.5 | P | 896.3 | P | 831.7 | P | 956.9 | P | --- |
| 259086_at | 0.6 | A | 0.4 | A | 1 | A | 1.1 | A | --- |
| 259087_at | 47.5 | P | 19.1 | P | 43.4 | A | 64.2 | M | --- |
| 259088_at | 196.6 | P | 194.9 | P | 142.4 | P | 193 | P | --- |
| 259089_at | 20.1 | A | 37.6 | A | 15.1 | A | 20.9 | A | --- |
| 259090_at | 2854.2 | P | 3026.1 | P | 3718.3 | P | 3140.9 | P | --- |
| 259091_at | 417.2 | P | 371.3 | P | 411.8 | P | 376.9 | P | --- |
| 259092_at | 2057.4 | P | 2151.5 | P | 2313.5 | P | 2294.5 | P | ZDS |
| 259093_at | 295 | P | 208.2 | P | 328.7 | P | 224.6 | P | --- |
| 259094_at | 542.8 | P | 674.3 | P | 709.7 | P | 727.8 | P | CYSD1 |
| 259095_at | 1226.8 | P | 830.4 | P | 1370.7 | P | 979.9 | P | ACP1 |
| 259096_at | 7072.3 | P | 6580 | P | 8092.2 | P | 6868.2 | P | --- |
| 259097_at | 909.1 | P | 938.7 | P | 643.8 | P | 869.2 | P | --- |
| 259098_at | 5992.2 | P | 6810.9 | P | 6778.2 | P | 7282.3 | P | --- |
| 259099_at | 413.9 | P | 426.6 | P | 402.9 | P | 382.2 | P | --- |
| 259100_at | 550.2 | P | 529 | P | 567.3 | P | 518 | P | DRT102 |
| 259101_at | 3.4 | A | 9.1 | A | 41.4 | A | 4.3 | A | --- |
| 259102_at | 1428.3 | P | 1294.4 | P | 893.9 | P | 1196.6 | P | NHL1 |
| 259103_at | 129.5 | P | 147.3 | P | 128.4 | P | 149 | P | --- |
| 259104_at | 144.9 | P | 282.3 | P | 131.1 | P | 180.8 | P | LNG2 |
| 259105_at | 1923.8 | P | 1714.7 | P | 1908.4 | P | 1671.9 | P | --- |
| 259106_at | 701.6 | P | 598.5 | P | 816.1 | P | 703.7 | P | RALFL22 |
| 259107_at | 11.4 | A | 19.4 | A | 2.9 | A | 3.6 | A | --- |
| 259108_at | 20.9 | A | 12.6 | A | 18.8 | A | 29.1 | A | --- |
| 259109_at | 152 | P | 250.8 | P | 166.4 | A | 165.6 | P | --- |
| 259110_at | 327.6 | P | 247.2 | P | 291.1 | P | 251.6 | P | --- |
| 259111_at | 496.1 | P | 726.5 | P | 560.7 | P | 776.3 | P | --- |
| 259112_at | 7370.6 | P | 6884.5 | P | 8016.7 | P | 8201 | P | --- |
| 259113_at | 185.6 | P | 158.4 | P | 135.7 | P | 187.7 | P | --- |
| 259114_at | 3415.9 | P | 3244.3 | P | 2729.5 | P | 3239.4 | P | RPT5A |
| 259115_at | 694.4 | P | 712.5 | P | 562.3 | P | 622 | P | --- |
| 259116_at | 160.4 | P | 172.6 | P | 227.9 | P | 144.3 | P | --- |
| 259117_at | 386.3 | P | 334.3 | P | 285.9 | P | 344.5 | P | SNL1 |
| 259118_at | 1815.3 | P | 1512.9 | P | 1607.3 | P | 1602.6 | P | --- |
| 259119_at | 874 | P | 861.4 | P | 945.2 | P | 840.9 | P | --- |
| 259120_at | 49.1 | P | 9.6 | P | 23.7 | A | 21.1 | A | --- |
| 259121_at | 1707.8 | P | 1280.6 | P | 1680.7 | P | 1231.6 | P | --- |
| 259122_at | 111.6 | P | 128.7 | P | 140.6 | P | 150.8 | P | COBL1 |
| 259123_at | 2706.3 | P | 2149.5 | P | 3342.7 | P | 1879.3 | P | --- |
| 259124_at | 4 | A | 2.8 | A | 23.7 | A | 3.9 | A | 41519 |
| 259125_at | 518.7 | P | 621.8 | P | 445.2 | P | 495.2 | P | --- |
| 259126_at | 194.2 | P | 205.2 | P | 205.6 | P | 188 | P | --- |
| 259127_at | 3.4 | A | 2.3 | A | 4.9 | A | 12.7 | A | --- |
| 259128_at | 695.6 | P | 528.7 | P | 520.5 | P | 510.8 | P | BIG |
| 259129_at | 507.1 | P | 266.9 | P | 404.3 | P | 230.5 | P | PTF1 |
| 259130_at | 1094.6 | P | 789.1 | P | 1043.1 | P | 815.2 | P | --- |
| 259131_at | 1753.5 | P | 1993.4 | P | 1672.8 | P | 2208.8 | P | SP1L3 |
| 259132_at | 417.4 | P | 567.9 | P | 416.9 | P | 554.2 | P | --- |
| 259133_at | 58 | P | 56.5 | P | 125.1 | P | 75.3 | P | --- |
| 259134_at | 40.6 | P | 42.8 | P | 35.2 | A | 27.6 | P | --- |
| 259135_at | 22.7 | A | 53.7 | A | 7.5 | A | 15.1 | A | RAD9 |
| 259136_at | 94.4 | P | 108.2 | P | 97.2 | P | 117.1 | P | --- |
| 259137_at | 961.6 | P | 975.1 | P | 653.2 | P | 883.6 | P | --- |
| 259138_s_at | 613.3 | P | 819.6 | P | 633.6 | P | 849.6 | P | --- |
| 259139_at | 2.9 | A | 19 | A | 17.1 | A | 18.1 | A | --- |
| 259140_at | 986.5 | P | 1062.6 | P | 934.4 | P | 975.8 | P | LYC |
| 259141_at | 591.6 | P | 549.1 | P | 667.5 | P | 559.5 | P | --- |
| 259142_at | 33.7 | A | 63.8 | P | 10.1 | A | 74.6 | P | --- |
| 259143_at | 391.2 | P | 290.2 | P | 328.8 | P | 274.2 | P | --- |
| 259144_at | 2.2 | A | 5.5 | A | 26.8 | A | 7.6 | A | --- |
| 259145_at | 25.6 | A | 44.9 | P | 29.6 | A | 39.9 | M | --- |
| 259146_at | 732.2 | P | 663.9 | P | 690.4 | P | 784.7 | P | SDP6 |
| 259147_at | 169.1 | P | 251.3 | P | 186.2 | P | 212.1 | P | APUM4 |
| 259148_at | 1239 | P | 1158.8 | P | 1389.4 | P | 1248.6 | P | --- |
| 259149_at | 63 | P | 60 | P | 31.9 | A | 86.2 | P | PAL4 |
| 259150_at | 48.5 | A | 23.3 | A | 5.7 | A | 3.2 | A | --- |
| 259151_at | 47.5 | P | 80.8 | P | 75 | P | 114.8 | P | --- |
| 259152_at | 381.8 | P | 409.6 | P | 350.3 | P | 323.2 | P | --- |
| 259153_at | 253.9 | P | 355.9 | P | 300.8 | P | 318 | P | --- |
| 259154_at | 1342.2 | P | 1080 | P | 1070.2 | P | 1086.8 | P | --- |
| 259155_at | 308.5 | P | 278.3 | P | 241.8 | P | 266.3 | P | --- |
| 259156_at | 1157.9 | P | 1258.8 | P | 1037.9 | P | 1084.7 | P | SEC8 |
| 259157_at | 29.1 | A | 24.9 | A | 51.4 | A | 7.7 | A | --- |
| 259158_at | 71.9 | A | 104.8 | P | 109.1 | A | 62.2 | A | --- |
| 259159_at | 686.4 | P | 620.6 | P | 591.3 | P | 742.2 | P | ACBP4 |
| 259160_at | 527.7 | P | 539.6 | P | 576.3 | P | 576 | P | --- |
| 259161_at | 14629.9 | P | 16194.2 | P | 16535.2 | P | 16593.2 | P | CA1 |
| 259162_at | 467.4 | P | 432.3 | P | 446.3 | P | 410.2 | P | --- |
| 259163_at | 119.7 | P | 170 | P | 179.6 | P | 229.5 | P | --- |
| 259164_at | 435 | P | 370 | P | 462 | P | 380.5 | P | ATBET10 |
| 259165_at | 680.6 | P | 955 | P | 690.5 | P | 782.9 | P | ATHB-1 |
| 259166_at | 276.1 | P | 354.6 | P | 327.9 | P | 315.6 | P | --- |
| 259167_at | 36.4 | A | 63.6 | A | 71.1 | A | 41.8 | A | --- |
| 259168_at | 354 | P | 388.1 | P | 465.8 | P | 406.6 | P | --- |
| 259169_at | 175.8 | P | 178.3 | P | 205.2 | P | 179 | P | --- |
| 259170_at | 21.2 | A | 33.5 | A | 49.6 | A | 39.4 | A | --- |
| 259171_at | 65.7 | P | 105.9 | P | 132.5 | P | 104.4 | P | --- |
| 259172_at | 300.2 | P | 346.4 | P | 354.5 | P | 344.1 | P | CS26 |
| 259173_at | 902.4 | P | 1883.3 | P | 428.8 | P | 1468.9 | P | BGLU25 |
| 259174_at | 173.3 | P | 143.2 | P | 197.5 | P | 195.6 | P | UNE7 |
| 259175_at | 383.3 | P | 389.6 | P | 379.8 | P | 315 | P | --- |

| | | | | | | | | | |
|---|---|---|---|---|---|---|---|---|---|
| 259176_at | 187.4 | P | 147.8 | P | 180.3 | P | 164.1 | P | CDC48C |
| 259177_at | 23 | A | 9.2 | A | 5.7 | A | 32.3 | A | --- |
| 259178_at | 146.5 | P | 217.5 | P | 147.6 | P | 199.5 | P | RGLG1 |
| 259179_at | 398.7 | P | 481.7 | P | 444.9 | P | 484.7 | P | --- |
| 259180_at | 314.2 | P | 447.2 | P | 503.9 | P | 398.9 | P | --- |
| 259181_at | 944.5 | P | 1078.2 | P | 830.1 | P | 1071.6 | P | --- |
| 259182_at | 86.1 | A | 73.9 | M | 75.2 | A | 85 | P | --- |
| 259183_at | 1.2 | A | 22.4 | A | 2 | A | 1.4 | A | --- |
| 259184_at | 1431.8 | P | 1588.2 | P | 1335.4 | P | 1404.9 | P | --- |
| 259185_at | 33.5 | P | 267 | P | 86.1 | A | 164.1 | P | PPT2 |
| 259186_at | 432.6 | P | 439.7 | P | 459 | P | 474.9 | P | --- |
| 259187_at | 6.1 | A | 6.4 | A | 4.1 | A | 1.9 | A | AtMYB57 |
| 259188_at | 717.3 | P | 672.1 | P | 813.6 | P | 650.5 | P | --- |
| 259189_at | 1 | A | 1.7 | A | 1.9 | A | 2.6 | A | AGP11 |
| 259190_at | 338.5 | P | 337.9 | P | 416.2 | P | 349.4 | P | TPLATE |
| 259191_at | 400.8 | P | 548.7 | P | 460 | P | 520.3 | P | --- |
| 259192_at | 846.6 | P | 703.2 | P | 821.4 | P | 775.3 | P | --- |
| 259193_at | 1684.4 | P | 2440.3 | P | 1722.2 | P | 2259.8 | P | CYP38 |
| 259194_at | 3524.7 | P | 3004.3 | P | 3466.1 | P | 3074.7 | P | DRH1 |
| 259195_at | 25.5 | A | 4.4 | A | 23 | A | 38.5 | A | --- |
| 259196_at | 1283.1 | P | 1142.6 | P | 1294.2 | P | 1109.8 | P | RPS2 |
| 259197_at | 22.3 | A | 22.3 | A | 39.6 | A | 19.8 | A | --- |
| 259198_at | 352.5 | P | 273.7 | P | 307.1 | P | 226.5 | P | --- |
| 259199_at | 226.8 | P | 329.1 | P | 272.8 | P | 353.2 | P | --- |
| 259200_at | 50 | P | 79.4 | P | 132.6 | P | 80.5 | P | --- |
| 259201_at | 42.8 | A | 43.6 | A | 51 | A | 40.1 | A | --- |
| 259202_at | 475.2 | P | 482.4 | P | 418.5 | P | 546.5 | P | --- |
| 259203_at | 19.4 | A | 10.3 | A | 27.1 | A | 2.9 | A | --- |
| 259204_s_at | 24.4 | A | 3.1 | A | 5.9 | A | 22.3 | A | --- |
| 259205_at | 190.2 | P | 200 | P | 151.7 | P | 194.5 | P | --- |
| 259206_at | 35.8 | A | 46.5 | A | 59.7 | A | 32.6 | A | --- |
| 259207_at | 225.4 | P | 349.5 | P | 253.9 | P | 337.6 | P | --- |
| 259208_at | 37.3 | A | 42 | P | 53.8 | P | 65.8 | A | --- |
| 259209_at | 66 | P | 72.1 | P | 80.8 | P | 86 | P | --- |
| 259210_at | 625 | P | 482.7 | P | 587.7 | P | 478 | P | HY2 |
| 259211_at | 31.6 | A | 52.7 | P | 42.8 | A | 37.1 | P | --- |
| 259212_at | 353.5 | P | 350.4 | P | 434.5 | P | 349.8 | P | --- |
| 259213_at | 172.4 | P | 176.6 | P | 160.1 | P | 164.8 | P | --- |
| 259214_at | 474.9 | P | 416.5 | P | 362.7 | P | 435.4 | P | --- |
| 259215_at | 733.6 | P | 661.6 | P | 706.7 | P | 673.1 | P | DEX1 |
| 259216_at | 129.1 | P | 128 | P | 114.5 | P | 147 | P | --- |
| 259217_at | 17.9 | A | 26.5 | A | 36.8 | A | 27.7 | A | --- |
| 259218_at | 87.1 | P | 111.7 | P | 91.1 | M | 150.4 | P | --- |
| 259219_at | 225.2 | P | 322.1 | P | 275 | P | 336.7 | P | --- |
| 259220_at | 259.2 | P | 170.1 | P | 148.7 | P | 225.6 | P | --- |
| 259221_s_at | 8.7 | A | 39.1 | A | 36.1 | A | 44.8 | A | NPC5 |
| 259222_at | 151.7 | P | 159.4 | P | 167.4 | P | 166.9 | P | --- |
| 259223_at | 2.1 | A | 11 | A | 26.4 | A | 30.2 | A | WOX11 |
| 259224_at | 617.5 | P | 759.2 | P | 655 | P | 879.9 | P | --- |
| 259225_at | 245.9 | P | 298.4 | P | 244.9 | P | 296.7 | P | --- |
| 259226_at | 1176.8 | P | 990.2 | P | 1019 | P | 1096.3 | P | --- |
| 259227_at | 342.9 | P | 236.7 | P | 477.7 | P | 267.1 | P | --- |
| 259228_at | 2779 | P | 1816.6 | P | 2707.7 | P | 1645.8 | P | --- |
| 259229_at | 151.3 | P | 198.6 | P | 158 | P | 148.9 | P | ADA2A |
| 259230_at | 324.2 | P | 235.9 | P | 229.4 | P | 228.2 | P | OBE1 |
| 259231_at | 591.6 | P | 482.9 | P | 365.2 | P | 616 | P | PP2CA |
| 259232_at | 328.9 | P | 253.9 | P | 244.7 | P | 253.9 | P | --- |
| 259233_at | 2.1 | A | 0.8 | A | 3.6 | A | 2.3 | A | --- |
| 259234_at | 267.5 | P | 268.5 | P | 313.8 | P | 275 | P | --- |
| 259235_at | 258.9 | P | 276.5 | P | 236 | P | 257.2 | P | --- |
| 259236_at | 618.1 | P | 631.5 | P | 598.4 | P | 584.4 | P | --- |
| 259237_at | 11051.1 | P | 12094.2 | P | 12925.6 | P | 12162.6 | P | --- |
| 259238_at | 1844.7 | P | 1758 | P | 2051.9 | P | 1851.6 | P | EIF3G1 |
| 259239_at | 7265.5 | P | 6556.2 | P | 7171.2 | P | 6496.8 | P | --- |
| 259240_at | 426.9 | P | 366.4 | P | 422.4 | P | 444.7 | P | --- |
| 259241_at | 443.3 | P | 374.1 | P | 393.8 | P | 426.3 | P | --- |
| 259242_at | 601.6 | P | 586.9 | P | 593.2 | P | 587 | P | ATARP6 |
| 259243_at | 459.7 | P | 513.3 | P | 553.9 | P | 519.6 | P | --- |
| 259244_at | 528.7 | P | 379.1 | P | 535.1 | P | 423.4 | P | COL9 |
| 259245_at | 740.6 | P | 740.5 | P | 791.1 | P | 736.5 | P | --- |
| 259246_s_at | 93.4 | P | 62.8 | P | 63.6 | P | 87 | P | --- |
| 259247_at | 508.1 | P | 627.7 | P | 392.7 | P | 527.5 | P | --- |
| 259248_at | 1234 | P | 1253.6 | P | 1486.2 | P | 1378.4 | P | --- |
| 259249_at | 992 | P | 799.5 | P | 1016.3 | P | 768.7 | P | --- |
| 259250_at | 82.8 | A | 112.7 | P | 86.5 | A | 113.3 | P | --- |
| 259251_at | 36.8 | A | 35.6 | A | 6.4 | A | 27.3 | A | --- |
| 259252_at | 180.1 | P | 142.9 | P | 165.6 | P | 120.2 | P | IBM1 |
| 259253_at | 815 | P | 730.9 | P | 754.1 | P | 823.7 | P | --- |
| 259254_at | 988.4 | P | 842.2 | P | 794.3 | P | 670.5 | P | ADT2 |
| 259255_at | 187.7 | P | 174.1 | P | 146.3 | P | 128.6 | P | --- |
| 259256_at | 1741.2 | P | 1638.7 | P | 1846.3 | P | 2126.1 | P | --- |
| 259257_at | 815.7 | P | 801.1 | P | 822.4 | P | 884.5 | P | --- |
| 259258_at | 646.9 | P | 710.3 | P | 753.9 | P | 799.5 | P | --- |
| 259259_at | 487.6 | P | 610.7 | P | 665.1 | P | 539.8 | P | SPY |
| 259260_at | 2.7 | A | 29.6 | A | 8.5 | A | 18 | A | --- |
| 259261_at | 9.9 | A | 8.1 | A | 6.6 | A | 4.1 | A | --- |
| 259262_at | 3.1 | A | 7.3 | A | 24.8 | A | 10 | A | ISU2 |
| 259263_at | 15.1 | A | 10.4 | A | 51 | A | 16.7 | A | --- |
| 259264_at | 2.3 | A | 11.1 | A | 27.1 | A | 2.6 | A | --- |
| 259265_at | 1.5 | A | 2.7 | A | 2.5 | A | 0.8 | A | --- |
| 259266_at | 1 | A | 0.4 | A | 1.6 | A | 9.2 | A | --- |
| 259267_at | 398.1 | P | 333.9 | P | 443.5 | P | 460.2 | P | --- |
| 259268_at | 36.6 | A | 36.2 | A | 49.8 | M | 25.4 | A | --- |
| 259269_at | 20.3 | P | 33.1 | P | 44.8 | P | 22.7 | A | --- |
| 259270_at | 4811.9 | P | 4192.9 | P | 4799.3 | P | 4276.9 | P | VDAC1 |
| 259271_at | 391.2 | P | 309.7 | P | 297.6 | P | 347.5 | P | --- |
| 259272_at | 865.5 | P | 1453.1 | P | 725.7 | P | 888.4 | P | --- |
| 259273_s_at | 240.7 | P | 345.6 | P | 339.1 | P | 369.2 | P | --- |
| 259274_at | 28.8 | A | 50.7 | P | 45.6 | P | 63 | A | ATHB20 |
| 259275_at | 425.3 | P | 645.4 | P | 532.5 | P | 532.7 | P | --- |
| 259276_at | 6.3 | A | 2.3 | A | 15.5 | A | 5.4 | A | --- |
| 259277_at | 568.5 | P | 529.7 | P | 537.2 | P | 567.5 | P | AtSS2 |
| 259278_at | 272.5 | P | 233.3 | P | 384.5 | P | 253.5 | P | --- |
| 259279_at | 9948.2 | P | 7687.5 | P | 12625.3 | P | 8561.9 | P | MTO1 |
| 259280_at | 518 | P | 539.7 | P | 566 | P | 498.3 | P | PTB1 |
| 259281_at | 208.6 | P | 176.5 | P | 254.6 | P | 224.9 | P | MYB106 |
| 259282_at | 18.7 | A | 35.8 | A | 12.9 | A | 28 | A | GPAT5 |
| 259283_at | 88 | P | 130.8 | P | 62.2 | P | 101.2 | P | MYB65 |
| 259284_at | 596.9 | P | 570.2 | P | 625.2 | P | 567.2 | P | --- |

| | | | | | | | | | |
|---|---|---|---|---|---|---|---|---|---|
| 259285_at | 121.1 | P | 83.1 | P | 123.2 | P | 96.6 | P | --- |
| 259286_at | 761.3 | P | 2.7 | A | 246.5 | P | 71.2 | P | BSMT1 |
| 259287_at | 210.3 | P | 232.9 | P | 243.1 | P | 275.3 | P | --- |
| 259288_at | 797.2 | P | 814.3 | P | 825.6 | P | 830.9 | P | --- |
| 259289_at | 2 | A | 1.8 | A | 6.4 | A | 3 | A | --- |
| 259290_at | 83.3 | P | 175.8 | P | 131.3 | P | 147.7 | P | CYCB1;3 |
| 259291_at | 34.8 | A | 5.4 | A | 3.8 | A | 1.4 | A | --- |
| 259292_at | 770.4 | P | 887.1 | P | 893.4 | P | 783.2 | P | --- |
| 259293_at | 451.1 | P | 218.3 | P | 489.3 | P | 240.5 | P | --- |
| 259294_at | 85.3 | P | 96.9 | P | 83.7 | P | 102.7 | P | ATN |
| 259295_at | 247.2 | P | 367.8 | P | 307.3 | P | 256.2 | P | --- |
| 259296_at | 1037.5 | P | 972.6 | P | 1067.7 | P | 976 | P | --- |
| 259297_at | 285.1 | P | 130.5 | P | 179.9 | P | 105.3 | P | AtRLP30 |
| 259298_at | 104.7 | P | 79.2 | P | 93.8 | P | 88.6 | P | AtRLP31 |
| 259299_at | 45.9 | P | 90.5 | P | 130.4 | P | 86 | P | --- |
| 259300_at | 365.1 | P | 411.6 | P | 383.2 | P | 475.4 | P | --- |
| 259301_at | 21.3 | A | 19.7 | A | 12.3 | A | 4.2 | A | --- |
| 259302_at | 1002.3 | P | 814 | P | 1125.7 | P | 973.3 | P | GID1A |
| 259303_at | 200.4 | P | 135.1 | P | 136 | P | 142.1 | P | --- |
| 259304_at | 257.4 | P | 306.3 | P | 228.6 | P | 341.2 | P | ERCC1 |
| 259305_at | 566.9 | P | 615.7 | P | 624.7 | P | 586.4 | P | --- |
| 259306_at | 378.5 | P | 360.5 | P | 475.2 | P | 345.6 | P | PNC1 |
| 259307_at | 477 | P | 574.2 | P | 534.9 | P | 526.1 | P | --- |
| 259308_at | 588.1 | P | 730.8 | P | 665.8 | P | 823.3 | P | --- |
| 259309_at | 223.8 | P | 216.8 | P | 225.7 | P | 179.7 | P | --- |
| 259310_s_at | 850.6 | P | 901.4 | P | 816.3 | P | 727.3 | P | --- |
| 259311_at | 1208.3 | P | 999.7 | P | 1550.9 | P | 1096.1 | P | --- |
| 259312_at | 740.7 | P | 503.2 | P | 481 | P | 475.6 | P | ATL6 |
| 259313_at | 675.3 | P | 699.8 | P | 794.9 | P | 704 | P | --- |
| 259314_at | 37.2 | A | 37.1 | A | 72.6 | A | 14.6 | A | --- |
| 259315_at | 632.1 | P | 694.2 | P | 589.4 | P | 569.6 | P | AtRP2 |
| 259316_at | 12.7 | A | 17.4 | A | 4.2 | A | 18.7 | P | --- |
| 259317_at | 33.3 | A | 91.9 | A | 124.2 | A | 64.4 | A | HYP1 |
| 259318_at | 380.5 | P | 344.2 | P | 332.2 | P | 284.9 | P | HYP1 |
| 259319_at | 1006.3 | P | 1012.9 | P | 1074 | P | 1038.8 | P | AKIN10 |
| 259320_at | 13.9 | A | 2.1 | A | 5.2 | A | 3.5 | A | WRKY58 |
| 259321_at | 356.4 | P | 408.1 | P | 342.5 | P | 387.7 | P | GAUT13 |
| 259322_at | 157 | P | 200 | P | 123.1 | P | 201.2 | P | --- |
| 259323_at | 477.4 | P | 597.5 | P | 529.6 | P | 557.3 | P | --- |
| 259324_at | 11 | A | 4.9 | A | 6.5 | A | 5.4 | A | MIRO3 |
| 259325_at | 81.8 | P | 108.8 | P | 69.8 | P | 75.7 | P | --- |
| 259326_at | 717.4 | P | 799.4 | P | 702.3 | P | 671.5 | P | MPPalpha |
| 259327_at | 2723.5 | P | 2015.6 | P | 3760.9 | P | 2530.2 | P | --- |
| 259328_at | 7.5 | A | 2.9 | A | 19.1 | A | 6.1 | A | ATMLP-300B |
| 259329_at | 135.5 | P | 107.3 | P | 154.4 | P | 190.7 | P | AHP4 |
| 259330_at | 506.4 | P | 534.4 | P | 528.6 | P | 464.7 | P | --- |
| 259331_at | 0.9 | A | 39.5 | P | 4.3 | A | 23.1 | A | --- |
| 259332_at | 13.3 | A | 41.9 | P | 11.8 | A | 19.9 | A | --- |
| 259333_at | 187.1 | P | 205.6 | P | 154.6 | P | 190.9 | P | EDA30 |
| 259334_at | 376.3 | P | 414.4 | P | 306.1 | P | 425 | P | --- |
| 259335_s_at | 294 | P | 321.2 | P | 296.2 | P | 311.3 | P | --- |
| 259336_at | 72.3 | P | 83 | P | 89.7 | A | 98.3 | P | SDG20 |
| 259337_at | 985.2 | P | 839.7 | P | 729.9 | P | 867.7 | P | CAT4 |
| 259338_at | 5.5 | A | 32.5 | A | 7.5 | A | 4.4 | A | SYP131 |
| 259339_at | 169.1 | P | 155.4 | P | 188.6 | P | 205.7 | P | --- |
| 259340_at | 328.4 | P | 379.3 | P | 212.6 | P | 337.7 | P | PYK10 |
| 259341_at | 399.6 | P | 419.5 | P | 357.3 | P | 432.7 | P | ATBPM4 |
| 259342_at | 1426.8 | P | 1294.6 | P | 1268.9 | P | 1273.5 | P | --- |
| 259343_s_at | 9525.6 | P | 9321.5 | P | 10288.2 | P | 9955.2 | P | ATMS2 |
| 259344_at | 455.2 | P | 626.3 | P | 595.7 | P | 638.1 | P | RIF10 |
| 259345_s_at | 8.4 | A | 45.4 | A | 47.4 | A | 10.2 | A | --- |
| 259346_at | 37.3 | A | 75 | P | 101.5 | P | 80.2 | P | GDH3 |
| 259347_at | 1366.4 | P | 1168.3 | P | 1745.9 | P | 1232.6 | P | --- |
| 259348_at | 447.7 | P | 475.5 | P | 466.9 | P | 529.7 | P | --- |
| 259349_at | 788.4 | P | 632.1 | P | 531.5 | P | 483.8 | P | ATAPRL5 |
| 259350_at | 29.7 | A | 32 | A | 30.1 | A | 27.3 | A | RBK2 |
| 259351_at | 4.5 | A | 4.9 | A | 3.5 | A | 2.6 | A | --- |
| 259352_at | 94.6 | P | 92 | P | 58.6 | P | 99.8 | P | --- |
| 259353_at | 175.5 | P | 98.3 | P | 235.9 | P | 167.1 | P | --- |
| 259354_at | 2839.8 | P | 1890.1 | P | 2107.4 | P | 2027.8 | P | --- |
| 259355_at | 60.2 | P | 33.3 | A | 55.8 | A | 40.1 | A | --- |
| 259356_at | 174.4 | P | 173.3 | P | 151.2 | P | 131.9 | P | --- |
| 259357_at | 513.3 | P | 717.3 | P | 550.9 | P | 700.4 | P | --- |
| 259358_at | 212.3 | P | 407.1 | P | 271.3 | P | 409.7 | P | GATL3 |
| 259359_at | 211.9 | P | 242 | P | 192.3 | P | 249.7 | P | --- |
| 259360_at | 9 | A | 1.5 | A | 23.5 | A | 5.1 | A | --- |
| 259361_at | 14907.7 | P | 14665.8 | P | 17810.3 | P | 14695.7 | P | GAPC2 |
| 259362_s_at | 485.3 | P | 373.6 | P | 393 | P | 335.6 | P | --- |
| 259363_at | 2447 | P | 2804.7 | P | 2350 | P | 2764.3 | P | MAP1C |
| 259364_at | 464.7 | P | 556.3 | P | 406.9 | P | 402.4 | P | RAV1 |
| 259365_at | 1.2 | A | 3.4 | A | 4 | A | 5.2 | A | --- |
| 259366_at | 1292.5 | P | 1118 | P | 1140.4 | P | 1038.7 | P | AOC4 |
| 259367_at | 382.7 | P | 318.6 | P | 372.6 | P | 295.2 | P | --- |
| 259368_at | 28.1 | A | 26 | A | 39.5 | A | 4.7 | A | --- |
| 259369_s_at | 1.9 | A | 5.8 | A | 22.9 | A | 14.5 | A | --- |
| 259370_at | 7.5 | A | 12.6 | A | 49.7 | A | 1.3 | A | --- |
| 259371_at | 82.1 | P | 87.1 | P | 53.4 | A | 120 | P | --- |
| 259372_at | 31.5 | A | 7.5 | A | 18.5 | A | 21.1 | A | AP1 |
| 259373_at | 241.8 | P | 212.9 | P | 258.3 | P | 222.8 | P | --- |
| 259374_at | 79.4 | P | 92.7 | P | 90 | P | 77.7 | P | --- |
| 259375_at | 3900.9 | P | 5914.7 | P | 4897.7 | P | 5890.4 | P | --- |
| 259376_at | 66.6 | P | 55.1 | P | 58.5 | P | 46.7 | P | CDC27a |
| 259377_at | 65.8 | P | 64.8 | P | 94 | P | 75.6 | P | PAB6 |
| 259378_at | 794.1 | P | 547.3 | P | 934.2 | P | 547.8 | P | --- |
| 259379_at | 98.8 | P | 117.7 | P | 88.8 | P | 94.1 | P | --- |
| 259380_at | 113.2 | P | 119 | P | 157.7 | P | 146.9 | P | PDR1 |
| 259381_s_at | 8418.4 | P | 3285 | P | 7468.3 | P | 4152.1 | P | NSP1 |
| 259382_s_at | 10976.5 | P | 7505.5 | P | 11073.1 | P | 8842.5 | P | JAL31 |
| 259383_at | 10977.2 | P | 3462.1 | P | 9247.9 | P | 5638.3 | P | JR1 |
| 259384_at | 2243.9 | P | 1427 | P | 3010.8 | P | 1801.7 | P | --- |
| 259385_at | 45.6 | P | 219.9 | P | 20.5 | A | 101.9 | P | --- |
| 259386_at | 69 | A | 22.4 | A | 47 | A | 33.7 | A | NUB |
| 259387_at | 11.8 | A | 40.1 | A | 17.3 | A | 48.5 | A | --- |
| 259388_at | 3.2 | A | 27.1 | A | 4.3 | A | 3.1 | A | ST4B |
| 259389_at | 5.7 | A | 16.7 | A | 3.7 | A | 5.7 | A | --- |
| 259390_at | 37.6 | A | 14.6 | A | 4.5 | A | 33.3 | A | --- |
| 259391_s_at | 276.6 | P | 796.8 | P | 290.7 | P | 685.2 | P | --- |
| 259392_at | 447.5 | P | 326.8 | P | 369.9 | P | 326.2 | P | --- |
| 259393_at | 1378.1 | P | 1400.4 | P | 1676.1 | P | 1318.9 | P | ATTPS7 |

| | | | | | | | | | |
|---|---|---|---|---|---|---|---|---|---|
| 259394_at | 2.4 | A | 45.6 | A | 34.3 | A | 37.8 | A | --- |
| 259395_at | 1059.3 | P | 1122.6 | P | 1019 | P | 1228.8 | P | ARA-2 |
| 259396_at | 715.8 | P | 843 | P | 655.8 | P | 838 | P | GSK1 |
| 259397_at | 233.7 | P | 246 | P | 271 | P | 212.2 | P | --- |
| 259398_at | 54.8 | P | 165.7 | P | 116.8 | P | 175.5 | P | PRA1.F1 |
| 259399_at | 26.4 | A | 62.3 | P | 77.6 | A | 41.4 | P | --- |
| 259400_at | 187.9 | P | 62 | P | 149.3 | P | 45.4 | P | --- |
| 259401_at | 183.5 | P | 191.3 | P | 278.5 | P | 212.2 | P | CSTF77 |
| 259402_at | 735.6 | P | 746.1 | P | 638.5 | P | 729.4 | P | VPS46.1 |
| 259403_at | 10317.6 | P | 4756.5 | P | 11560.3 | P | 6525.1 | P | PGDH |
| 259404_at | 617.1 | P | 625.8 | P | 630.4 | P | 642.6 | P | ATB BETA |
| 259405_at | 140.8 | P | 81.6 | P | 166.8 | P | 97.8 | P | --- |
| 259406_at | 539.7 | P | 522.9 | P | 591.9 | P | 433.7 | P | --- |
| 259407_at | 1260.3 | P | 1419.8 | P | 1328.6 | P | 1386.7 | P | PP2AA3 |
| 259408_at | 1150 | P | 1349.9 | P | 1321.9 | P | 1296.9 | P | PP2AA2 |
| 259409_at | 29.1 | P | 37.4 | P | 62.3 | P | 53.5 | P | --- |
| 259410_at | 205.3 | P | 236.1 | P | 103.4 | P | 166.2 | P | --- |
| 259411_at | 143.9 | P | 100.8 | P | 155.8 | P | 120.4 | P | --- |
| 259412_at | 307.6 | P | 280.3 | P | 338.6 | P | 265.8 | P | --- |
| 259413_at | 4.6 | A | 3.3 | A | 1.3 | A | 2.7 | A | --- |
| 259414_at | 99.7 | P | 90.1 | P | 77.8 | P | 81.7 | P | --- |
| 259415_at | 499.9 | P | 443.2 | P | 474.9 | P | 446.4 | P | --- |
| 259416_at | 2852.3 | P | 3056.9 | P | 2291.4 | P | 2873 | P | --- |
| 259417_at | 2.1 | A | 41.3 | P | 23.7 | A | 27 | A | HFR1 |
| 259418_at | 63.8 | P | 72.8 | P | 48.8 | A | 96.5 | P | GPAT2 |
| 259419_at | 19.3 | A | 42.1 | A | 28.6 | A | 27.5 | A | ACX6 |
| 259420_at | 543 | P | 526.8 | P | 537.2 | P | 525 | P | --- |
| 259421_at | 90.5 | M | 143.3 | P | 137.9 | P | 132.3 | P | --- |
| 259422_at | 279.1 | P | 200.9 | P | 231.2 | P | 192.9 | P | --- |
| 259423_at | 286.4 | P | 264.3 | P | 187.8 | P | 277.5 | P | --- |
| 259424_at | 43.6 | A | 48.1 | P | 39 | A | 39.5 | A | --- |
| 259425_at | 7.8 | A | 1.7 | A | 5.2 | A | 3.1 | A | PIPK11 |
| 259426_at | 4090 | P | 5771.1 | P | 2542.4 | P | 5135.4 | P | LEA14 |
| 259427_at | 14.6 | A | 23.4 | A | 21.9 | A | 18.6 | A | AGL28 |
| 259428_at | 150.1 | P | 244.2 | P | 128.3 | P | 141.4 | P | ATMPK11 |
| 259429_at | 141.7 | P | 205.2 | P | 113 | P | 222 | P | CYP86A4 |
| 259430_at | 1082.1 | P | 1409.8 | P | 1150.1 | P | 1230.4 | P | GPAT4 |
| 259431_at | 1362.7 | P | 2648 | P | 1246.8 | P | 2512.7 | P | PIP1C |
| 259432_at | 32.3 | P | 33.8 | P | 43.7 | P | 49.6 | P | --- |
| 259433_at | 79.3 | P | 97.9 | P | 141.3 | P | 98.9 | P | --- |
| 259434_at | 3809.3 | P | 3075.7 | P | 3210.6 | P | 2607.1 | P | --- |
| 259435_at | 2.1 | A | 33.2 | A | 29.2 | A | 31.2 | A | --- |
| 259436_at | 728.9 | P | 437.8 | P | 489.9 | P | 437.1 | P | --- |
| 259437_at | 648.2 | P | 664.8 | P | 742.6 | P | 692.3 | P | --- |
| 259438_at | 577 | P | 770.2 | P | 652.5 | P | 664.2 | P | AN |
| 259439_at | 49.2 | A | 14.1 | A | 54.5 | A | 4.3 | A | ACS2 |
| 259440_at | 1611.1 | P | 2202.7 | P | 1620.8 | P | 2025.2 | P | BPS1 |
| 259441_at | 428.5 | P | 536.9 | P | 361 | P | 509.6 | P | --- |
| 259442_at | 53.1 | A | 34 | A | 33.8 | A | 38.2 | A | --- |
| 259443_at | 2.5 | A | 19.4 | A | 24.7 | A | 4.2 | A | --- |
| 259444_at | 156.2 | P | 152.1 | P | 209.3 | P | 134.9 | P | --- |
| 259445_at | 91.4 | P | 117.6 | P | 57.1 | P | 82.7 | P | GA2OX6 |
| 259446_at | 851.7 | P | 766.3 | P | 731.3 | P | 726.1 | P | --- |
| 259447_s_at | 13.2 | A | 24.4 | A | 30.3 | A | 2.4 | A | ATARFD1A |
| 259448_at | 39.7 | P | 72.7 | P | 51.6 | P | 68.8 | P | --- |
| 259449_at | 30.3 | A | 26.7 | A | 57.9 | A | 31.6 | A | --- |
| 259450_at | 273.7 | P | 304.3 | P | 267.9 | P | 298.4 | P | DRL1 |
| 259451_at | 1.5 | A | 0.9 | A | 3.9 | A | 0.6 | A | SNAP30 |
| 259452_at | 7.5 | A | 3.4 | A | 11.1 | A | 7.7 | A | --- |
| 259453_at | 11.4 | A | 22.1 | A | 3.9 | A | 2.9 | A | ATGA20OX5 |
| 259454_at | 0.7 | A | 5.5 | A | 40.5 | A | 1.1 | A | --- |
| 259455_x_at | 43.6 | A | 29.8 | A | 36.7 | A | 10.2 | A | --- |
| 259456_s_at | 31.1 | A | 1.9 | A | 13.4 | A | 4.3 | A | --- |
| 259457_at | 7.6 | A | 2.4 | A | 9.2 | A | 1.4 | A | ARF23 |
| 259458_at | 22.4 | P | 18.8 | A | 47.7 | P | 19.8 | A | --- |
| 259459_at | 1.5 | A | 15.9 | A | 4.7 | A | 1.4 | A | --- |
| 259460_at | 873.4 | P | 1373.7 | P | 952.5 | P | 1477.9 | P | --- |
| 259461_at | 353.4 | P | 317 | P | 451.5 | P | 320.1 | P | --- |
| 259462_at | 45.2 | A | 31.9 | A | 51.8 | A | 66.2 | P | --- |
| 259463_at | 187.7 | P | 183.3 | P | 171.6 | P | 205.2 | P | --- |
| 259464_at | 2.6 | A | 1.4 | A | 0.7 | A | 1.4 | A | --- |
| 259465_at | 3.5 | A | 3.1 | A | 2.9 | A | 13.1 | A | --- |
| 259466_at | 873.7 | P | 1022.7 | P | 874.6 | P | 1060.1 | P | ARR7 |
| 259467_at | 10.6 | A | 22.7 | A | 31.4 | A | 4.5 | A | --- |
| 259468_at | 359.4 | P | 366 | P | 464 | P | 301.6 | P | TTN10 |
| 259469_at | 101.8 | P | 105 | P | 136.9 | P | 100.4 | P | --- |
| 259470_at | 756 | P | 996.8 | P | 1084.1 | P | 1112 | P | --- |
| 259471_at | 6.4 | A | 4.3 | A | 21.2 | A | 3.4 | A | --- |
| 259472_at | 139.2 | P | 181.7 | P | 184.9 | P | 199.7 | P | --- |
| 259473_at | 71.8 | P | 51.5 | P | 50.2 | M | 63.7 | P | --- |
| 259474_at | 1270.2 | P | 1095 | P | 1071.5 | P | 949.4 | P | --- |
| 259475_at | 841.9 | P | 744.8 | P | 1039.8 | P | 847.1 | P | --- |
| 259476_at | 1230 | P | 1679.6 | P | 1866.5 | P | 1455.8 | P | --- |
| 259477_at | 1071.1 | P | 1079.6 | P | 1133.8 | P | 917.3 | P | --- |
| 259478_at | 14.2 | A | 20.8 | A | 7.4 | A | 5.2 | A | --- |
| 259479_at | 1120.9 | P | 647.6 | P | 676.5 | P | 458.9 | P | --- |
| 259480_at | 144.5 | P | 158.5 | P | 123.2 | P | 118.8 | P | --- |
| 259481_at | 3.8 | A | 25.9 | A | 45.8 | P | 16.1 | A | GLP4 |
| 259482_at | 14.8 | A | 4.6 | A | 6 | A | 1.6 | A | RKF2 |
| 259483_at | 4.5 | A | 1.9 | A | 3.2 | A | 27.8 | A | --- |
| 259484_at | 217.3 | P | 300.5 | P | 231.3 | P | 317.5 | P | PEPKR1 |
| 259485_at | 161.7 | P | 136 | P | 199.7 | P | 136.4 | P | ATPC2 |
| 259486_at | 555.1 | P | 554.1 | P | 566.6 | P | 521.7 | P | --- |
| 259487_at | 67.1 | P | 86.7 | A | 85.7 | A | 57.9 | M | TRFL5 |
| 259488_at | 395.3 | P | 350.2 | P | 354.3 | P | 291.2 | P | --- |
| 259489_at | 160.3 | P | 145.6 | P | 129.8 | P | 125.9 | P | --- |
| 259490_at | 154.4 | P | 109 | P | 172.5 | P | 109.2 | P | --- |
| 259491_at | 9330.8 | P | 11168.3 | P | 7456 | P | 9712.4 | P | LHCB6 |
| 259492_at | 28.1 | A | 53.9 | A | 39 | A | 12.5 | A | --- |
| 259493_at | 75.8 | A | 34.4 | A | 79.3 | A | 40.5 | A | --- |
| 259494_at | 22.3 | A | 23.3 | A | 57.4 | P | 24.9 | A | --- |
| 259495_at | 65.3 | P | 49.6 | P | 36.1 | A | 41.9 | A | --- |
| 259496_at | 50.6 | P | 26.7 | A | 70.1 | A | 16.6 | A | --- |
| 259497_at | 468 | P | 431.7 | P | 441.7 | P | 382.8 | P | --- |
| 259498_at | 669.1 | P | 658.2 | P | 577.1 | P | 637.7 | P | GOS11 |
| 259499_at | 1085.5 | P | 987.3 | P | 969.6 | P | 888.5 | P | --- |
| 259500_at | 800.9 | P | 925.5 | P | 631.2 | P | 740.8 | P | --- |
| 259501_at | 1448 | P | 1327.6 | P | 1450.7 | P | 1445.4 | P | TPL |
| 259502_at | 1121.6 | P | 699.3 | P | 483.5 | P | 618.3 | P | --- |

| | | | | | | | | | |
|---|---|---|---|---|---|---|---|---|---|
| 259503_at | 82.7 | P | 66.3 | P | 86.1 | P | 92.7 | P | --- |
| 259504_at | 9584.1 | P | 11179.6 | P | 11696.5 | P | 11608.6 | P | AVP1 |
| 259505_at | 599.9 | P | 655.7 | P | 645.5 | P | 723 | P | --- |
| 259506_at | 679.3 | P | 640.6 | P | 601.3 | P | 740.4 | P | --- |
| 259507_at | 2880.3 | P | 2004.8 | P | 3256.3 | P | 1705 | P | --- |
| 259508_at | 6.2 | A | 37 | A | 25.9 | A | 33.4 | A | --- |
| 259509_at | 9.2 | A | 1.4 | A | 2.3 | A | 11.8 | A | --- |
| 259510_at | 15.4 | A | 3 | A | 11.2 | A | 3 | A | --- |
| 259511_at | 1162.3 | P | 356.1 | P | 541.4 | P | 791.8 | P | ATCCS |
| 259512_at | 794.1 | P | 737.8 | P | 615.4 | P | 693.2 | P | KEU |
| 259513_at | 291 | P | 430.5 | P | 405.8 | P | 413.9 | P | ARK3 |
| 259514_at | 138.5 | P | 129.5 | P | 133.9 | P | 132.7 | P | OZS1 |
| 259515_at | 925.6 | P | 826.4 | P | 827.5 | P | 777.4 | P | --- |
| 259516_at | 4407.4 | P | 4385.7 | P | 2845.9 | P | 5210.4 | P | ERD10 |
| 259517_at | 2080.4 | P | 1289.7 | P | 1614.7 | P | 1373.7 | P | CAT1 |
| 259518_at | 1522.2 | P | 1105.1 | P | 1562.4 | P | 1117.3 | P | OPCL1 |
| 259519_at | 597 | P | 627.6 | P | 629.1 | P | 633 | P | --- |
| 259520_at | 4.2 | A | 70.4 | M | 7.8 | A | 56.4 | A | --- |
| 259521_at | 2759.2 | P | 2795.6 | P | 2728.5 | P | 2718.7 | P | CLP2 |
| 259522_at | 14.2 | A | 2.3 | A | 29.4 | A | 2.9 | A | --- |
| 259523_at | 278.2 | P | 373.7 | P | 346.4 | P | 379.3 | P | --- |
| 259524_at | 137 | P | 174.2 | P | 226.8 | P | 174.8 | P | --- |
| 259525_at | 32.4 | A | 8.1 | A | 38.6 | A | 26.7 | A | ATEXPA7 |
| 259526_at | 60.3 | M | 70.4 | P | 55.7 | A | 119.3 | P | --- |
| 259527_at | 12.8 | A | 5.6 | A | 51.2 | A | 4.6 | A | --- |
| 259528_at | 295.8 | P | 257.2 | P | 304.8 | P | 214.6 | P | --- |
| 259529_at | 707.2 | P | 653.7 | P | 669.2 | P | 663.5 | P | --- |
| 259530_at | 165.8 | P | 190.1 | P | 174.6 | P | 173.6 | P | --- |
| 259531_at | 117.1 | P | 122.2 | A | 99.5 | A | 162.5 | P | --- |
| 259532_at | 467.1 | P | 479.1 | P | 442.7 | P | 501 | P | --- |
| 259533_at | 345.2 | P | 270.8 | P | 170.4 | P | 280.9 | P | --- |
| 259534_at | 63 | P | 62.1 | M | 39.3 | A | 38.2 | A | --- |
| 259535_at | 83.8 | P | 126.7 | P | 106.6 | M | 111.9 | P | --- |
| 259536_s_at | 329.7 | P | 215.9 | P | 277.5 | P | 229.1 | P | ATCOAB |
| 259537_at | 72.6 | P | 145.2 | P | 149.7 | P | 130.7 | P | PHR1 |
| 259538_at | 1442.9 | P | 1608.9 | P | 1522.1 | P | 1690.9 | P | --- |
| 259539_at | 56.4 | P | 66.7 | P | 47.4 | P | 60.7 | P | --- |
| 259540_at | 184.1 | P | 238.1 | P | 255.9 | P | 248.3 | P | --- |
| 259541_at | 497.3 | P | 562.6 | P | 518.5 | P | 526.2 | P | --- |
| 259542_at | 511.6 | P | 532 | P | 491.2 | P | 487.9 | P | --- |
| 259543_at | 741.8 | P | 906.6 | P | 895.6 | P | 811.2 | P | --- |
| 259544_at | 14796.3 | P | 15866.7 | P | 15730.1 | P | 16091.4 | P | CAT3 |
| 259545_at | 938.8 | P | 751.2 | P | 844.2 | P | 747 | P | AAE1 |
| 259546_at | 439.9 | P | 461.1 | P | 328.2 | P | 444.4 | P | --- |
| 259547_at | 39.5 | P | 24.1 | M | 33.5 | P | 25.6 | P | --- |
| 259548_at | 86.5 | P | 104.5 | P | 173.4 | P | 111.8 | P | MLP165 |
| 259549_at | 59.5 | P | 57.7 | P | 111.4 | P | 78.9 | P | --- |
| 259550_at | 110.2 | A | 280 | P | 79.5 | A | 226.3 | P | AGP5 |
| 259551_at | 311 | P | 308.1 | P | 305.8 | P | 405.1 | P | --- |
| 259552_at | 17.4 | A | 34.9 | A | 11.9 | A | 17.8 | A | --- |
| 259553_x_at | 200.5 | P | 132.7 | P | 151.7 | A | 143.9 | P | ATEXT3 |
| 259554_at | 49.1 | P | 54.8 | M | 60.2 | M | 42.5 | P | --- |
| 259555_at | 3.1 | A | 4 | A | 7.9 | A | 5.1 | A | --- |
| 259556_at | 19.2 | A | 15 | A | 4.3 | A | 24.7 | A | --- |
| 259557_at | 1.7 | A | 4.1 | A | 24 | A | 2.3 | A | WAK4 |
| 259558_at | 19.2 | P | 15.5 | A | 1.7 | A | 2.3 | A | WAK5 |
| 259559_at | 11.6 | A | 70.9 | P | 10.2 | A | 23.4 | A | WAK3 |
| 259560_at | 337.9 | P | 532.6 | P | 242 | P | 392.4 | P | WAK2 |
| 259561_at | 1177.7 | P | 2366.5 | P | 1168.4 | P | 1938.9 | P | WAK1 |
| 259562_at | 101.2 | P | 113.4 | P | 87.1 | P | 118.8 | P | --- |
| 259563_s_at | 94.1 | P | 120.5 | P | 83.9 | P | 108.3 | P | --- |
| 259564_at | 565.4 | P | 661 | P | 671 | P | 654.2 | P | --- |
| 259565_at | 28.7 | A | 23.8 | M | 46.4 | A | 16.6 | P | --- |
| 259566_at | 5.3 | A | 23.3 | A | 45.8 | A | 20.2 | A | --- |
| 259567_at | 16.4 | A | 19.7 | P | 15.1 | A | 10.3 | A | --- |
| 259568_at | 245.5 | P | 39.7 | A | 219.2 | P | 64.9 | P | --- |
| 259569_at | 74.5 | A | 8.7 | A | 47.8 | A | 7.4 | A | --- |
| 259570_at | 4366.2 | P | 4307.7 | P | 2654.9 | P | 4398.4 | P | COR47 |
| 259571_at | 278.3 | P | 272.1 | P | 310.2 | P | 283.5 | P | --- |
| 259572_at | 2 | A | 3.1 | A | 7.5 | A | 3.9 | A | --- |
| 259573_at | 116.3 | P | 93.4 | P | 162.6 | P | 80.9 | P | --- |
| 259574_at | 25.4 | A | 49.5 | P | 45.4 | P | 48.8 | P | MLP168 |
| 259575_at | 100.6 | P | 58.9 | P | 117.1 | P | 72.1 | P | --- |
| 259576_at | 25.8 | A | 12.2 | A | 14.3 | A | 18.4 | A | --- |
| 259577_at | 739.6 | P | 819 | P | 787 | P | 797.6 | P | --- |
| 259578_at | 25.3 | A | 18.9 | A | 6.3 | A | 4.4 | A | --- |
| 259579_at | 93.1 | P | 75.9 | P | 116.2 | P | 59.2 | P | PGP14 |
| 259580_at | 38 | P | 37 | A | 57.6 | P | 29.5 | P | --- |
| 259581_at | 32.6 | A | 23.5 | A | 61.1 | A | 65.4 | A | --- |
| 259582_at | 466.1 | P | 492.1 | P | 626.3 | P | 490.3 | P | --- |
| 259583_at | 168.7 | P | 140 | P | 138.2 | P | 175 | P | --- |
| 259584_at | 39.3 | A | 41.3 | A | 62.4 | P | 59.3 | A | --- |
| 259585_at | 219.4 | P | 240.4 | P | 278.5 | P | 254.4 | P | --- |
| 259586_at | 223.6 | P | 298.1 | P | 275.7 | P | 305.3 | P | --- |
| 259587_at | 1062.1 | P | 1057.3 | P | 929.7 | P | 1020.9 | P | --- |
| 259588_at | 1142.8 | P | 1351.4 | P | 921.8 | P | 1159.4 | P | --- |
| 259589_at | 9.6 | A | 1.5 | A | 1.5 | A | 3.2 | A | --- |
| 259590_at | 25.4 | A | 37.5 | A | 35.8 | M | 31.3 | A | --- |
| 259591_at | 1410.8 | P | 1886.1 | P | 1898.9 | P | 1425.1 | P | --- |
| 259592_at | 2534.6 | P | 3121.6 | P | 2480.9 | P | 3023.1 | P | LTPG1 |
| 259593_at | 2745.1 | P | 3505.4 | P | 2615.8 | P | 3373.4 | P | NTF2B |
| 259594_at | 2421.5 | P | 2740.9 | P | 2582 | P | 2493.3 | P | --- |
| 259595_at | 408.8 | P | 325.5 | P | 343.5 | P | 377.2 | P | --- |
| 259596_at | 342.2 | P | 288.6 | P | 257.2 | P | 267.3 | P | GH3.17 |
| 259597_at | 401.5 | P | 422.8 | P | 403.4 | P | 524.8 | P | --- |
| 259598_at | 994.4 | P | 747.4 | P | 950.8 | P | 712.9 | P | DPL1 |
| 259599_at | 281.7 | P | 498.8 | P | 500.1 | P | 503.2 | P | SCPL45 |
| 259600_at | 139.6 | P | 168.1 | P | 111.7 | P | 168.6 | P | --- |
| 259601_s_at | 2.2 | A | 3.7 | A | 1.9 | A | 2.4 | A | --- |
| 259602_at | 67.4 | A | 68.1 | P | 32.7 | A | 75.5 | A | --- |
| 259603_at | 1994.2 | P | 2378.9 | P | 2347.4 | P | 2573 | P | --- |
| 259604_at | 2657.8 | P | 3168.6 | P | 2506 | P | 3203.1 | P | PBG1 |
| 259605_at | 169.4 | P | 178.2 | P | 179.5 | A | 192.4 | P | PUB45 |
| 259606_at | 32.2 | P | 53.3 | P | 39.1 | P | 92.7 | P | MAP65-8 |
| 259607_at | 49 | P | 70.4 | A | 46.8 | A | 113.8 | P | PGP13 |
| 259608_at | 106.9 | P | 151.4 | P | 178.4 | P | 123.3 | P | ECT9 |
| 259609_at | 3420.2 | P | 1113.4 | P | 2510.4 | P | 1426.9 | P | TSA1 |
| 259610_at | 394.2 | P | 400.1 | P | 361.9 | P | 384.9 | P | --- |
| 259611_at | 478.4 | P | 551.9 | P | 456.6 | P | 559 | P | AtRABG3d |

| | | | | | | | | | |
|---|---|---|---|---|---|---|---|---|---|
| 259612_at | 12627.4 | P | 10418.6 | P | 10786.1 | P | 10717.5 | P | --- |
| 259613_at | 7.8 | A | 1.9 | A | 0.5 | A | 9.5 | A | --- |
| 259614_at | 3.3 | A | 3.9 | A | 12.5 | A | 1.8 | A | ATGA2OX4 |
| 259615_at | 5.3 | A | 40.9 | A | 47.4 | A | 36.1 | A | --- |
| 259616_at | 410.4 | P | 397.4 | P | 322.8 | P | 403.8 | P | C/VIF1 |
| 259617_at | 676.5 | P | 690.9 | P | 1045.2 | P | 609.5 | P | --- |
| 259618_at | 58.1 | P | 54.3 | A | 61.7 | P | 91 | P | MYB112 |
| 259619_at | 11.9 | A | 22 | A | 22.8 | A | 14.2 | A | --- |
| 259620_s_at | 31 | A | 14.9 | A | 29.2 | A | 37.4 | A | --- |
| 259621_at | 2.5 | A | 2.4 | A | 1 | A | 1.1 | A | --- |
| 259622_at | 19.4 | A | 10.2 | A | 3.1 | A | 8.5 | A | --- |
| 259623_at | 8.8 | A | 26.7 | A | 10 | A | 44.4 | A | --- |
| 259624_at | 16.7 | A | 12.3 | A | 35.8 | A | 3.1 | A | --- |
| 259625_at | 15833.8 | P | 15421.7 | P | 15418.3 | P | 16242.8 | P | GAPB |
| 259626_at | 1297.8 | P | 1180.3 | P | 959.2 | P | 1056.4 | P | ATBZIP60 |
| 259627_at | 5404.1 | P | 4944.4 | P | 5038.2 | P | 5273.5 | P | --- |
| 259628_at | 108.5 | P | 168 | P | 152.3 | P | 148.3 | P | --- |
| 259629_at | 400.6 | P | 508.5 | P | 386.6 | P | 349.5 | P | WRR4 |
| 259630_at | 208.3 | P | 207.3 | P | 202.7 | P | 225.1 | P | --- |
| 259631_at | 17.2 | A | 6.7 | A | 24.6 | A | 8 | A | ERD2 |
| 259632_at | 112.4 | P | 729.7 | P | 587.8 | P | 745.6 | P | NAS4 |
| 259633_at | 661.2 | P | 919 | P | 807.8 | P | 876.7 | P | --- |
| 259634_at | 3.6 | A | 4.6 | A | 30.1 | A | 3.7 | A | --- |
| 259635_at | 2.7 | A | 10.2 | A | 14.3 | A | 9.3 | A | PAP6 |
| 259636_at | 91.5 | P | 69.5 | P | 127.3 | P | 88.1 | P | --- |
| 259637_at | 148.1 | P | 178.4 | P | 166.1 | P | 162.4 | P | ATPDIL1-5 |
| 259638_at | 657.7 | P | 735 | P | 693.7 | P | 687.1 | P | --- |
| 259639_at | 664.5 | P | 932.2 | P | 679.8 | P | 729.9 | P | --- |
| 259640_at | 15337.3 | P | 9216 | P | 15637.2 | P | 11027.6 | P | BGLU18 |
| 259641_at | 73 | P | 65.6 | P | 117.2 | P | 58.6 | P | --- |
| 259642_at | 449.9 | P | 590.5 | P | 456.1 | P | 513.4 | P | --- |
| 259643_at | 53.3 | P | 79.1 | P | 55.7 | A | 42.2 | M | --- |
| 259644_at | 108.8 | P | 121.6 | P | 68.7 | P | 69.2 | P | --- |
| 259645_at | 287.5 | P | 453.8 | P | 280.8 | P | 475.6 | P | BIM2 |
| 259646_at | 193.7 | P | 197.7 | P | 186.3 | P | 190.5 | P | DEK1 |
| 259647_at | 167 | P | 175.7 | P | 185.4 | P | 151.4 | P | --- |
| 259648_at | 73.1 | P | 86.6 | P | 96.4 | P | 62.5 | M | --- |
| 259649_at | 199.1 | P | 257.5 | P | 267 | P | 243.7 | P | TAF7 |
| 259650_at | 28.9 | A | 22.3 | A | 5.5 | A | 45.2 | A | --- |
| 259651_at | 107.4 | P | 135.6 | P | 111.5 | P | 103.1 | P | --- |
| 259652_at | 65.4 | P | 92.7 | P | 99.2 | P | 74.3 | P | HUB2 |
| 259653_at | 22 | A | 24.3 | A | 30.4 | A | 27.2 | P | --- |
| 259654_at | 2.2 | A | 1 | A | 2.3 | A | 1.9 | A | --- |
| 259655_at | 2350.1 | P | 1793.8 | P | 1671.9 | P | 2133.2 | P | --- |
| 259656_at | 57.6 | P | 53.2 | P | 47.9 | P | 39.7 | P | --- |
| 259657_at | 35.7 | A | 21 | A | 37.5 | A | 37.8 | A | PLDEPSILON |
| 259658_at | 278.1 | P | 394.8 | P | 354 | P | 367 | P | NDF5 |
| 259659_at | 332.4 | P | 289.1 | P | 277.1 | P | 321.2 | P | --- |
| 259660_at | 2185.8 | P | 2494.6 | P | 2612.2 | P | 2570.9 | P | --- |
| 259661_at | 170.2 | P | 321.9 | P | 156.9 | P | 220.8 | P | --- |
| 259662_at | 227 | P | 221.4 | P | 203.4 | P | 221.5 | P | HUB2 |
| 259663_at | 554.6 | P | 524.2 | P | 574.3 | P | 468.4 | P | PRA7 |
| 259664_at | 5623 | P | 5833.2 | P | 7016.5 | P | 7310.5 | P | AGP21 |
| 259665_at | 710.4 | P | 732.5 | P | 760.4 | P | 774.8 | P | --- |
| 259666_at | 580.2 | P | 560.8 | P | 516.6 | P | 583 | P | SR33 |
| 259667_at | 332.2 | P | 291.9 | P | 352.7 | P | 251.6 | P | --- |
| 259668_at | 4 | A | 10.6 | A | 6.7 | A | 7.7 | A | --- |
| 259669_at | 307.4 | P | 359.8 | P | 282.4 | P | 355 | P | ABA2 |
| 259670_at | 459.3 | P | 522.4 | P | 494 | P | 525.1 | P | --- |
| 259671_at | 237.3 | P | 291.7 | P | 307.2 | P | 327.9 | P | --- |
| 259672_at | 474.2 | P | 292.9 | P | 398.2 | P | 355 | P | --- |
| 259673_at | 358.6 | P | 363.4 | P | 298.5 | P | 416.4 | P | --- |
| 259674_at | 63 | A | 76.8 | P | 79.5 | A | 91.1 | P | --- |
| 259675_at | 4334.4 | P | 3729.3 | P | 4119.4 | P | 3652.9 | P | --- |
| 259676_at | 26.5 | A | 20 | A | 10.2 | A | 11.7 | A | --- |
| 259677_at | 435.3 | P | 419.5 | P | 409.8 | P | 483.4 | P | --- |
| 259678_at | 640.8 | P | 938.4 | P | 655.5 | P | 937.5 | P | --- |
| 259679_at | 65.3 | P | 79.1 | P | 74.2 | A | 101.5 | P | --- |
| 259680_at | 122.6 | P | 249.1 | P | 170.9 | P | 275.8 | P | LAX3 |
| 259681_at | 7042.5 | P | 714.4 | P | 7309.1 | P | 1345.9 | P | NIA1 |
| 259682_at | 24.5 | A | 30.5 | A | 65.7 | A | 36.6 | A | --- |
| 259683_at | 288.1 | P | 299.3 | P | 274.8 | P | 210.4 | P | --- |
| 259684_at | 7.5 | A | 11 | A | 0.5 | A | 0.7 | A | --- |
| 259685_at | 966.4 | P | 945.9 | P | 966.4 | P | 852.9 | P | AtPP2-A11 |
| 259686_at | 60.9 | P | 138.7 | P | 78 | P | 116.1 | P | --- |
| 259687_at | 527.9 | P | 598.9 | P | 535 | P | 569.3 | P | --- |
| 259688_at | 537.7 | P | 644.8 | P | 519.1 | P | 696.4 | P | ATRBL2 |
| 259689_x_at | 399.6 | P | 515.5 | P | 500.7 | P | 569.5 | P | --- |
| 259690_at | 220.8 | P | 133.6 | P | 180.2 | P | 235.7 | P | --- |
| 259691_at | 21.2 | A | 54.8 | P | 57.7 | P | 52.1 | A | --- |
| 259692_at | 82.6 | P | 93.8 | P | 118.7 | P | 101.9 | P | --- |
| 259693_at | 0.9 | A | 2 | A | 2.8 | A | 1.8 | A | --- |
| 259694_at | 97.9 | P | 91.8 | P | 125.2 | P | 69.6 | P | UGE3 |
| 259695_at | 425.1 | P | 497.6 | P | 515.9 | P | 467.9 | P | --- |
| 259696_at | 49 | P | 54.7 | P | 65.7 | A | 56.1 | P | --- |
| 259697_at | 119 | P | 160.8 | P | 143.2 | P | 170.3 | P | --- |
| 259698_at | 74.4 | P | 81.9 | A | 99.4 | P | 81.7 | P | --- |
| 259699_at | 6.2 | A | 10.4 | A | 5.3 | A | 3.9 | A | --- |
| 259700_at | 73.3 | P | 120.1 | P | 97.4 | P | 140.8 | P | --- |
| 259701_at | 480.8 | P | 436.8 | P | 318.5 | P | 433.3 | P | --- |
| 259702_at | 7.9 | A | 14.3 | A | 41.5 | P | 24 | A | --- |
| 259703_at | 21 | A | 7.5 | A | 37.2 | A | 22.5 | P | --- |
| 259704_at | 339.4 | P | 379.7 | P | 436.7 | P | 425.1 | P | --- |
| 259705_at | 1491.7 | P | 822.2 | P | 1223.7 | P | 1024.7 | P | anac032 |
| 259706_at | 656 | P | 424 | P | 517.7 | P | 448.4 | P | --- |
| 259707_at | 922 | P | 1087.4 | P | 1090.6 | P | 901.3 | P | TAPX |
| 259708_at | 515.9 | P | 403.6 | P | 517.9 | P | 341.5 | P | --- |
| 259709_at | 6.2 | A | 6.2 | A | 38.9 | A | 46.4 | A | --- |
| 259710_at | 696.7 | P | 855.1 | P | 1056.1 | P | 783.2 | P | --- |
| 259711_at | 135.1 | P | 107.2 | P | 120 | P | 101.6 | P | --- |
| 259712_at | 1367.7 | P | 1429.7 | P | 1218.1 | P | 1403.6 | P | PBC2 |
| 259713_at | 612 | P | 781.8 | P | 782.1 | P | 756 | P | --- |
| 259714_at | 25.3 | A | 7.5 | A | 30.6 | A | 24 | A | ATGA20OX4 |
| 259715_at | 434.6 | P | 364.4 | P | 387.6 | P | 469.1 | P | --- |
| 259716_at | 245.7 | P | 229.2 | P | 250 | P | 208.3 | P | --- |
| 259717_at | 447.3 | P | 558.1 | P | 609.8 | P | 501.4 | P | CPSF73-I |
| 259718_at | 252.1 | P | 245 | P | 296.3 | P | 255.8 | P | VIP5 |
| 259719_at | 7.8 | A | 9.5 | A | 35.8 | A | 17.2 | A | LCR66 |
| 259720_at | 1.9 | A | 3.1 | A | 4.3 | A | 1.2 | A | --- |

| | | | | | | | | | |
|---|---|---|---|---|---|---|---|---|---|
| 259721_at | 170.9 | P | 324.3 | P | 234.5 | P | 406.3 | P | --- |
| 259722_at | 34.6 | P | 68.2 | P | 64.4 | P | 76.9 | P | ATRECQ4B |
| 259723_at | 653.3 | P | 364.6 | P | 699.2 | P | 550.1 | P | IRT3 |
| 259724_at | 1393.7 | P | 1401.2 | P | 1179.7 | P | 1414.5 | P | SNRK2.10 |
| 259725_at | 176.1 | P | 146.5 | P | 187.9 | P | 142.6 | P | --- |
| 259726_at | 29.2 | A | 33.7 | A | 24.8 | A | 11.1 | A | SCRL6 |
| 259727_at | 14935.3 | P | 13873.7 | P | 17091.2 | P | 13597.2 | P | FED A |
| 259728_at | 17.5 | A | 12.6 | A | 12.2 | A | 20 | A | --- |
| 259729_at | 59.8 | P | 45 | A | 58.2 | P | 31 | A | --- |
| 259730_at | 91.5 | P | 92.2 | P | 86.2 | A | 82.4 | P | --- |
| 259731_at | 175.4 | P | 258.7 | P | 206.3 | P | 284.2 | P | --- |
| 259732_at | 284.4 | P | 300.8 | P | 427.6 | P | 332.4 | P | --- |
| 259733_at | 429.5 | P | 347.6 | P | 404.8 | P | 332.3 | P | --- |
| 259734_at | 136.1 | P | 112.8 | P | 91.2 | P | 81.3 | P | --- |
| 259735_at | 23.4 | A | 70 | P | 26.5 | A | 71.6 | A | --- |
| 259736_at | 705.6 | P | 1815 | P | 836.7 | P | 1747.3 | P | AtGH9C2 |
| 259737_at | 123.4 | P | 283.1 | P | 148.9 | P | 278.5 | P | --- |
| 259738_at | 477.5 | P | 662.7 | P | 548.4 | P | 674.8 | P | --- |
| 259739_at | 379.7 | P | 367.4 | P | 354.6 | P | 382.2 | P | SEH1H |
| 259740_at | 46.5 | A | 79.6 | P | 23.5 | A | 76 | P | --- |
| 259741_at | 164.7 | P | 287 | P | 233.5 | P | 254.7 | P | --- |
| 259742_at | 17 | A | 9.9 | A | 16.7 | A | 40.3 | A | GLIP6 |
| 259743_at | 1615.1 | P | 757.6 | P | 815.3 | P | 608.1 | P | --- |
| 259744_at | 5.4 | A | 22.5 | A | 22.6 | A | 29 | A | KCS7 |
| 259745_at | 476.8 | P | 552.5 | P | 556.9 | P | 501 | P | SAG18 |
| 259746_at | 105.9 | P | 94.9 | P | 99.3 | P | 96.5 | P | --- |
| 259747_at | 319 | P | 249.8 | P | 279.4 | P | 250.7 | P | --- |
| 259748_at | 234 | P | 200.6 | P | 293.8 | P | 266.5 | P | --- |
| 259749_at | 420.7 | P | 331.9 | P | 398.2 | P | 390 | P | RSW10 |
| 259750_at | 223 | P | 247.7 | P | 184.5 | P | 251.4 | P | --- |
| 259751_at | 521.8 | P | 549.1 | P | 439.9 | P | 306.7 | P | MYBL2 |
| 259752_at | 1465.5 | P | 1516 | P | 1623.1 | P | 1247.7 | P | LPR2 |
| 259753_at | 152.7 | P | 239.4 | P | 186.6 | P | 239.3 | P | --- |
| 259754_at | 699.5 | P | 955.2 | P | 824.2 | P | 927.1 | P | --- |
| 259755_at | 427.3 | P | 469.8 | P | 414.5 | P | 508 | P | --- |
| 259756_at | 880.4 | P | 614.9 | P | 842.8 | P | 585.9 | P | --- |
| 259757_at | 1922.7 | P | 1888.7 | P | 1163.2 | P | 2214.2 | P | ATPDIL1-2 |
| 259758_s_at | 42.3 | A | 19.2 | A | 66.6 | P | 25.4 | A | --- |
| 259759_at | 408.8 | P | 308.6 | P | 421.4 | P | 338.4 | P | --- |
| 259760_at | 83.1 | P | 91.6 | P | 110.6 | P | 97.8 | P | --- |
| 259761_at | 533 | P | 591 | P | 487.8 | P | 553.5 | P | LACS9 |
| 259762_at | 156.5 | P | 197.4 | P | 167.5 | P | 127.9 | P | --- |
| 259763_at | 138.5 | P | 219.4 | P | 242.2 | P | 278.3 | P | --- |
| 259764_at | 572 | P | 765.2 | P | 383.4 | P | 668.8 | P | NPR1 |
| 259765_at | 2423.6 | P | 1690.9 | P | 2363.8 | P | 2035.9 | P | --- |
| 259766_at | 505.1 | P | 493 | P | 797.1 | P | 510 | P | --- |
| 259767_s_at | 986.8 | P | 1150.5 | P | 1103.1 | P | 1146.3 | P | --- |
| 259768_at | 1939 | P | 2800.1 | P | 1585.4 | P | 2783.6 | P | COR314-TM2 |
| 259769_at | 1222.9 | P | 1326.2 | P | 1367.9 | P | 1309.7 | P | AML5 |
| 259770_s_at | 478.2 | P | 394 | P | 452.3 | P | 452.7 | P | PAI3 |
| 259771_at | 3686.6 | P | 3998.3 | P | 3974.5 | P | 3905.6 | P | --- |
| 259772_at | 19.4 | A | 1.2 | A | 2.1 | A | 2 | A | --- |
| 259773_at | 15.3 | A | 75.3 | P | 30.8 | A | 70.1 | P | --- |
| 259774_at | 85.5 | P | 126.6 | P | 126.6 | P | 107.7 | P | --- |
| 259775_at | 324.8 | P | 456.2 | P | 498.3 | P | 558.6 | P | --- |
| 259776_s_at | 145.3 | P | 237.7 | P | 190.8 | P | 285.7 | P | --- |
| 259777_at | 2 | A | 1 | A | 2 | A | 0.7 | A | --- |
| 259778_at | 0.9 | A | 2.8 | A | 17 | A | 8.2 | A | --- |
| 259779_s_at | 13.1 | A | 1.7 | A | 2.2 | A | 7.8 | A | --- |
| 259780_at | 26.4 | A | 35.2 | A | 25.5 | A | 33.9 | A | --- |
| 259781_at | 7.2 | A | 6.1 | A | 10.1 | A | 2.3 | A | --- |
| 259782_at | 51 | A | 33.1 | A | 73.9 | A | 50.7 | A | --- |
| 259783_at | 23.3 | M | 93.5 | P | 34.1 | P | 75 | P | SAUR68 |
| 259784_at | 2.1 | A | 56 | A | 4.7 | A | 25.8 | A | --- |
| 259785_at | 1.4 | A | 3.1 | A | 8 | A | 3.1 | A | --- |
| 259786_at | 3109 | P | 2731 | P | 3466.4 | P | 3255.3 | P | --- |
| 259787_at | 1.3 | A | 1.7 | A | 1 | A | 8.5 | A | --- |
| 259788_at | 2028.4 | P | 5183.5 | P | 2146.6 | P | 4298.9 | P | --- |
| 259789_at | 416.2 | P | 1093.4 | P | 293.9 | P | 1376.4 | P | COR414-TM1 |
| 259790_s_at | 18.2 | P | 167.8 | P | 45.2 | P | 117.7 | P | --- |
| 259791_at | 562.3 | P | 765.4 | P | 592.7 | P | 791.4 | P | --- |
| 259792_at | 224.5 | P | 221.5 | P | 136.7 | P | 183.1 | P | CAD1 |
| 259793_at | 33.4 | P | 28.6 | P | 35.1 | A | 25.8 | A | --- |
| 259794_at | 136.6 | P | 190.6 | P | 118 | P | 239.1 | P | --- |
| 259795_at | 21.5 | A | 25.7 | A | 37.9 | A | 26.1 | A | --- |
| 259796_at | 140.4 | P | 132.5 | P | 129.7 | P | 118.4 | P | --- |
| 259797_at | 9.5 | A | 2.1 | A | 2.8 | A | 14.3 | A | --- |
| 259798_at | 17.5 | P | 42.6 | P | 27.6 | P | 29.9 | A | --- |
| 259799_at | 2.7 | A | 2.4 | A | 5.4 | A | 1.6 | A | --- |
| 259800_at | 434 | P | 349.4 | P | 404.4 | P | 336.6 | P | --- |
| 259801_at | 1.7 | A | 89.2 | P | 37.2 | A | 114.3 | P | --- |
| 259802_at | 436 | P | 42.1 | P | 251.3 | P | 98.6 | P | THI2.1 |
| 259803_at | 2058.2 | P | 2177.3 | P | 1856.9 | P | 2163.9 | P | PATL1 |
| 259804_at | 1409.2 | P | 1512.8 | P | 1175.4 | P | 1338 | P | --- |
| 259805_at | 29.7 | A | 38.4 | A | 17.5 | A | 15.4 | A | AtRLP7 |
| 259806_at | 249.9 | P | 223.7 | P | 232.8 | P | 298.2 | P | --- |
| 259807_at | 25.7 | A | 12.5 | A | 3.4 | A | 9.7 | A | --- |
| 259808_at | 37.1 | M | 31.8 | A | 39.3 | P | 37.1 | P | --- |
| 259809_at | 3.1 | A | 10.4 | A | 23.3 | A | 20.2 | A | --- |
| 259810_at | 2.2 | A | 3 | A | 6.8 | A | 8.1 | A | ATNHD2 |
| 259811_at | 32.9 | P | 58.8 | P | 72.7 | P | 49.8 | P | --- |
| 259812_at | 344.4 | P | 214.7 | P | 282.8 | P | 234.7 | P | --- |
| 259813_at | 1.3 | A | 15.2 | A | 20.4 | A | 18.4 | A | ATGSTF14 |
| 259814_at | 25.4 | A | 52.5 | A | 46.7 | A | 52 | A | --- |
| 259815_at | 54.5 | A | 63.1 | P | 66.5 | A | 83.8 | P | --- |
| 259816_at | 399.6 | P | 377.3 | P | 340 | P | 332.2 | P | --- |
| 259817_at | 142.7 | P | 121.1 | P | 117.6 | P | 149.3 | P | --- |
| 259818_at | 192 | P | 221.8 | P | 246 | P | 253.5 | P | --- |
| 259819_at | 1062.9 | P | 1125.6 | P | 946 | P | 1160.8 | P | ATMTK |
| 259820_at | 18.8 | A | 18.3 | A | 32.9 | A | 21.4 | A | --- |
| 259821_at | 34.2 | A | 39.3 | A | 56.9 | A | 39.9 | A | --- |
| 259822_at | 17.5 | A | 72.6 | P | 39 | A | 66 | P | MYB20 |
| 259823_at | 189.9 | P | 269.1 | P | 158.5 | P | 325.4 | P | --- |
| 259824_at | 6869.5 | P | 6104.5 | P | 6196.6 | P | 6513.6 | P | ATX1 |
| 259825_at | 1274.2 | P | 1173.5 | P | 1456.9 | P | 1028.1 | P | --- |
| 259826_at | 1154 | P | 1064.8 | P | 777.6 | P | 1032.4 | P | PUB17 |
| 259827_at | 15.8 | A | 1.9 | A | 19.4 | A | 26.9 | A | --- |
| 259828_at | 10.7 | A | 3 | A | 9.2 | A | 5.9 | A | --- |
| 259829_at | 18.5 | A | 6.6 | A | 5.2 | A | 24.4 | A | --- |

| | | | | | | | | | |
|---|---|---|---|---|---|---|---|---|---|
| 259830_at | 100.1 | P | 172.4 | P | 140.2 | P | 180.9 | P | --- |
| 259831_at | 1.7 | A | 2.1 | A | 5.4 | A | 2.1 | A | ZFHD1 |
| 259832_at | 103.4 | P | 103.4 | P | 179.7 | P | 119.5 | P | --- |
| 259833_at | 10.5 | A | 36.4 | A | 31.1 | A | 23.8 | A | MYB105 |
| 259834_at | 91.8 | A | 133.5 | P | 38.8 | A | 89.5 | P | --- |
| 259835_at | 419 | P | 452.9 | P | 383.5 | P | 391.9 | P | TRZ3 |
| 259836_at | 3.8 | A | 4.6 | A | 3.2 | A | 3.3 | A | ROPGEF11 |
| 259837_at | 76.2 | P | 65.4 | P | 65.3 | P | 62.6 | P | --- |
| 259838_at | 3315.4 | P | 4252.4 | P | 3118 | P | 3857 | P | --- |
| 259839_at | 163 | P | 1241.4 | P | 286.6 | P | 1031.8 | P | --- |
| 259840_at | 9857.1 | P | 12078.5 | P | 9251.9 | P | 10705.7 | P | PSAH2 |
| 259841_at | 483.5 | P | 1106.5 | P | 282.6 | P | 765.5 | P | --- |
| 259842_at | 488.9 | P | 981.7 | P | 848.4 | P | 1231.7 | P | CPuORF32 |
| 259843_at | 55.1 | A | 58.4 | P | 69.7 | P | 68.6 | P | --- |
| 259844_at | 3.3 | A | 2.9 | A | 27 | A | 1.4 | A | --- |
| 259845_at | 76.5 | P | 128.9 | P | 172.1 | P | 160.1 | P | PIN1 |
| 259846_at | 116.5 | P | 101.5 | P | 142.5 | P | 100.2 | P | --- |
| 259847_at | 847.2 | P | 1182.5 | P | 799.3 | P | 1161 | P | --- |
| 259848_at | 1367.3 | P | 1208 | P | 1275.8 | P | 1261.3 | P | --- |
| 259849_at | 300.4 | P | 315.1 | P | 350.5 | P | 337 | P | --- |
| 259850_at | 30.6 | P | 105 | P | 33.9 | A | 68.9 | P | --- |
| 259851_at | 61.7 | A | 103.7 | P | 98.3 | P | 108.6 | P | --- |
| 259852_at | 206.5 | P | 128.5 | P | 145 | P | 170.3 | P | AERO1 |
| 259853_at | 147.4 | P | 148.8 | P | 153.8 | P | 165.2 | P | --- |
| 259854_at | 13.7 | A | 43.4 | P | 66.3 | P | 71.8 | P | --- |
| 259855_at | 6.5 | A | 3.6 | A | 2.9 | A | 0.6 | A | --- |
| 259856_at | 474.6 | P | 472.1 | P | 623.9 | P | 480.2 | P | --- |
| 259857_at | 55.8 | P | 124.5 | P | 86.4 | P | 110.5 | P | --- |
| 259858_at | 213.1 | P | 249.2 | P | 326.1 | P | 253.5 | P | --- |
| 259859_at | 1002 | P | 600.9 | P | 539.2 | P | 695.3 | P | --- |
| 259860_at | 812.6 | P | 696.3 | P | 982 | P | 764.8 | P | --- |
| 259861_at | 579.8 | P | 710.1 | P | 789.7 | P | 579.2 | P | --- |
| 259862_at | 209.6 | P | 243.4 | P | 235.8 | P | 300.7 | P | TRFL6 |
| 259863_at | 415.8 | P | 496.5 | P | 286.8 | P | 404.3 | P | ELF4-L2 |
| 259864_at | 467.7 | P | 380 | P | 425.3 | P | 334.6 | P | --- |
| 259865_at | 1466.7 | P | 1696.9 | P | 1398.8 | P | 1566.1 | P | CKL2 |
| 259866_at | 266.8 | P | 1 | A | 135.2 | P | 2 | A | --- |
| 259867_at | 29.1 | A | 57.6 | P | 32.7 | A | 57 | P | --- |
| 259868_at | 76 | P | 164.2 | P | 168.7 | P | 136.3 | P | ATY1 |
| 259869_at | 7.6 | A | 11.7 | A | 27.2 | A | 7.7 | A | --- |
| 259870_at | 18.2 | A | 9 | A | 16.6 | A | 13.6 | P | --- |
| 259871_at | 77.3 | A | 133.4 | P | 104.9 | A | 74.6 | M | --- |
| 259872_at | 1079.8 | P | 1091.6 | P | 1100.3 | P | 1137.3 | P | --- |
| 259873_s_at | 85.7 | P | 51.4 | P | 63 | P | 40 | P | --- |
| 259874_at | 227.2 | P | 283.1 | P | 282.2 | P | 301.4 | P | --- |
| 259875_s_at | 4010.1 | P | 6092.8 | P | 3727.2 | P | 5749.4 | P | OPR1 |
| 259876_at | 232.8 | P | 269.1 | P | 200.2 | P | 244.9 | P | --- |
| 259877_at | 132.7 | P | 113.6 | P | 121.8 | P | 113.8 | P | SDG26 |
| 259878_at | 6637.6 | P | 1399.5 | P | 4964.9 | P | 2783.1 | P | --- |
| 259879_at | 300.1 | P | 194.3 | P | 201.2 | P | 157.2 | P | CML38 |
| 259880_at | 1342.4 | P | 1522.8 | P | 1734.5 | P | 1599.2 | P | --- |
| 259881_at | 113.4 | P | 83.1 | P | 74.8 | P | 82.9 | P | --- |
| 259882_at | 1412.8 | P | 1805.2 | P | 1298.3 | P | 1763.5 | P | --- |
| 259883_at | 919 | P | 1015.5 | P | 1182.9 | P | 1009.3 | P | --- |
| 259884_at | 459.8 | P | 417.1 | P | 311.4 | P | 369.5 | P | --- |
| 259885_at | 150.8 | P | 141.7 | P | 216 | P | 153.1 | P | --- |
| 259886_at | 12.2 | A | 3.1 | A | 2.6 | A | 2.1 | A | --- |
| 259887_at | 349.4 | P | 353.9 | P | 254.1 | P | 300.1 | P | --- |
| 259888_at | 166 | P | 225.7 | P | 164.7 | P | 197.2 | P | --- |
| 259889_at | 324.8 | P | 504.5 | P | 365.2 | P | 555.4 | P | --- |
| 259890_at | 1653.1 | P | 1158.4 | P | 1546.3 | P | 1263.9 | P | ATTIM23-2 |
| 259891_at | 224.6 | P | 206.2 | P | 241.7 | P | 234.5 | P | --- |
| 259892_at | 9082.4 | P | 12674.9 | P | 13111.6 | P | 14711.5 | P | GER1 |
| 259893_at | 5.5 | A | 21.2 | A | 38 | A | 4 | A | AtRLP11 |
| 259894_at | 343.1 | P | 252.1 | P | 365.9 | P | 259.4 | P | --- |
| 259895_at | 237.8 | P | 257.1 | P | 233.1 | P | 220.1 | P | PFI |
| 259896_at | 2642.3 | P | 3098.9 | P | 2913.3 | P | 3018.2 | P | --- |
| 259897_at | 25.1 | A | 37.7 | A | 26.1 | A | 38.4 | A | ATCEL3 |
| 259898_at | 22.9 | A | 50.3 | A | 7.8 | A | 14.4 | A | --- |
| 259899_at | 252.6 | P | 199.4 | P | 271 | P | 238.9 | P | --- |
| 259900_at | 129.4 | P | 110.9 | P | 102.2 | P | 131.9 | P | CSN5B |
| 259901_at | 75.7 | P | 64.2 | P | 57.5 | A | 61.4 | P | AtRLP14 |
| 259902_at | 76.7 | P | 86.1 | P | 118.3 | P | 113.1 | P | AtRLP13 |
| 259903_at | 167.4 | P | 230.6 | P | 239.9 | P | 181.1 | P | --- |
| 259904_at | 36.5 | P | 46.7 | A | 70.9 | M | 42.9 | A | --- |
| 259905_s_at | 22.5 | A | 23.4 | A | 58.8 | A | 5.4 | A | --- |
| 259906_at | 47.7 | P | 82.4 | A | 77.2 | A | 84.2 | P | --- |
| 259907_at | 118.6 | P | 168.7 | P | 205.5 | P | 180.6 | P | AGD2 |
| 259908_at | 510.1 | P | 348.1 | P | 602.6 | P | 349.2 | P | ATRPAC42 |
| 259909_at | 1297.3 | P | 1284.1 | P | 1446.6 | P | 1357.1 | P | MEE9 |
| 259910_at | 521.8 | P | 406.7 | P | 461.4 | P | 419.7 | P | --- |
| 259911_at | 5038.3 | P | 2062.3 | P | 3435.8 | P | 2938.2 | P | CAD1 |
| 259912_at | 58.8 | A | 171.7 | P | 73.5 | A | 129.6 | P | iqd8 |
| 259913_at | 17.5 | A | 39.6 | A | 19.6 | A | 21 | A | --- |
| 259914_at | 190.2 | P | 284.3 | P | 251.1 | P | 194.4 | P | --- |
| 259915_at | 156.5 | P | 122.2 | P | 117.9 | P | 159.6 | P | --- |
| 259916_at | 13 | A | 33.6 | A | 4.2 | A | 30.6 | A | --- |
| 259917_at | 0.4 | A | 1.1 | A | 0.7 | A | 7.3 | A | --- |
| 259918_at | 1.3 | A | 1.4 | A | 2 | A | 1.5 | A | --- |
| 259919_at | 194 | P | 220.6 | P | 209.5 | P | 191.2 | P | PSD |
| 259920_at | 1366.9 | P | 1340.8 | P | 1674 | P | 1278 | P | --- |
| 259921_at | 35.6 | A | 27.6 | A | 4.1 | A | 27.5 | A | --- |
| 259922_at | 1081.9 | P | 853.1 | P | 860.4 | P | 907.9 | P | HAB1 |
| 259923_at | 4.7 | A | 18.3 | A | 3.5 | A | 19.3 | A | --- |
| 259924_at | 554.5 | P | 419.8 | P | 562.2 | P | 441.2 | P | --- |
| 259925_at | 1238.6 | P | 740.9 | P | 676.3 | P | 723.3 | P | PR5 |
| 259926_at | 96.5 | P | 93.4 | P | 142.7 | P | 95.2 | P | --- |
| 259927_at | 198.3 | P | 327.7 | P | 228 | P | 239.8 | P | JAC1 |
| 259928_at | 247.3 | P | 327.5 | P | 329 | P | 311.2 | P | --- |
| 259929_at | 60.8 | P | 104.7 | P | 116.5 | P | 83.2 | P | --- |
| 259930_at | 73.2 | P | 80.6 | P | 54.5 | P | 62.5 | P | --- |
| 259931_at | 11 | A | 2.3 | A | 11.8 | A | 4.8 | A | --- |
| 259932_at | 1157.2 | P | 963.6 | P | 1076.3 | P | 1078.4 | P | STOP1 |
| 259933_at | 245.8 | P | 322.9 | P | 243.9 | P | 346.4 | P | --- |
| 259934_at | 209.6 | P | 188.6 | P | 173.9 | P | 178.8 | P | --- |
| 259935_at | 2 | A | 3.4 | A | 5.7 | A | 1.3 | A | --- |
| 259936_at | 766.4 | P | 910.6 | P | 793.2 | P | 765.9 | P | EBS1 |
| 259937_s_at | 754.9 | P | 506.8 | P | 539.3 | P | 468.5 | P | ATNAP5 |
| 259938_at | 242.8 | P | 307 | P | 344.9 | P | 324.5 | P | --- |

| | | | | | | | | | |
|---|---|---|---|---|---|---|---|---|---|
| 259939_s_at | 294.5 | P | 386 | P | 281 | P | 346.5 | P | --- |
| 259940_at | 10 | P | 17.1 | M | 17.7 | P | 8.9 | A | --- |
| 259941_s_at | 95.4 | P | 35.5 | A | 68.8 | P | 43.7 | A | --- |
| 259942_at | 1473 | P | 1395.5 | P | 1603.7 | P | 1536 | P | ATWHY2 |
| 259943_at | 561.4 | P | 757.5 | P | 704.6 | P | 805.2 | P | --- |
| 259944_at | 8.3 | A | 24.8 | P | 28.6 | A | 17.1 | A | --- |
| 259945_at | 288.3 | P | 256.7 | P | 287.6 | P | 265.6 | P | --- |
| 259946_at | 27.8 | A | 25.6 | A | 30.6 | A | 36.3 | A | --- |
| 259947_at | 4.9 | A | 1.2 | A | 9.8 | A | 1.3 | A | --- |
| 259948_at | 403.9 | P | 317.6 | P | 367.3 | P | 348.6 | P | --- |
| 259949_at | 47.4 | P | 42.4 | P | 45 | P | 36.4 | A | --- |
| 259950_at | 318.3 | P | 316 | P | 340.9 | P | 330.6 | P | --- |
| 259951_at | 42.7 | P | 35.1 | P | 30.2 | A | 34.2 | A | --- |
| 259952_at | 160.7 | P | 193 | P | 182.6 | P | 176 | P | AtRLP12 |
| 259953_at | 444.5 | P | 358.7 | P | 337.7 | P | 469.8 | P | BOR5 |
| 259954_at | 402 | P | 440 | P | 364.7 | P | 374.8 | P | CYP721A1 |
| 259955_s_at | 1200.7 | P | 1823.3 | P | 1242.8 | P | 1580.7 | P | BZR1 |
| 259956_at | 22.8 | A | 60 | A | 19 | A | 44.4 | A | RRA2 |
| 259957_at | 287.7 | P | 372.9 | P | 320.4 | P | 352.7 | P | --- |
| 259958_at | 483.6 | P | 679.3 | P | 560.6 | P | 666.8 | P | SRF6 |
| 259959_at | 169.2 | P | 224.1 | P | 227.6 | P | 289.5 | P | CYP59 |
| 259960_at | 132.5 | P | 211.2 | P | 149.4 | P | 194.5 | P | --- |
| 259961_at | 10.5 | A | 7.4 | A | 3.7 | A | 2.9 | A | WAG1 |
| 259962_at | 3.6 | A | 1.1 | A | 2.5 | A | 7.9 | A | --- |
| 259963_at | 38 | P | 61.8 | A | 73.2 | A | 38 | A | --- |
| 259964_at | 37.5 | A | 52 | A | 34.8 | A | 31.1 | P | ATGSTU28 |
| 259965_at | 2208.5 | P | 1766.8 | P | 2155.1 | P | 1733.8 | P | MSRB1 |
| 259966_at | 40.9 | A | 43.9 | A | 59.7 | A | 74.6 | P | SOB3 |
| 259967_at | 192 | P | 205.7 | P | 150.5 | P | 230.5 | P | --- |
| 259968_at | 241.8 | P | 114.5 | P | 271.5 | P | 176.5 | P | --- |
| 259969_at | 716.2 | P | 615.3 | P | 819.7 | P | 710.7 | P | --- |
| 259970_at | 354.5 | P | 451.2 | P | 505.5 | P | 513.7 | P | --- |
| 259971_at | 249.3 | P | 378.4 | P | 245.9 | P | 279 | P | --- |
| 259972_at | 2.3 | A | 37.3 | P | 19.8 | P | 39.4 | P | CUC3 |
| 259973_at | 145.2 | P | 194.3 | P | 164.3 | P | 155.4 | P | --- |
| 259974_at | 8.5 | A | 3 | A | 13.1 | A | 7.2 | A | --- |
| 259975_at | 175.4 | P | 51.4 | M | 111.5 | P | 36.6 | A | --- |
| 259976_at | 117.2 | P | 168.5 | P | 132.7 | P | 157.4 | P | CP12-3 |
| 259977_at | 457.3 | P | 323.1 | P | 250.8 | P | 322.3 | P | --- |
| 259978_at | 213 | P | 343.1 | P | 249.5 | P | 355.1 | P | CDKB2;1 |
| 259979_at | 594.7 | P | 612.5 | P | 586.6 | P | 525.7 | P | --- |
| 259980_at | 6710.6 | P | 3755 | P | 5722.8 | P | 4267.2 | P | --- |
| 259981_at | 1454.7 | P | 1824.9 | P | 1287.2 | P | 1721.4 | P | --- |
| 259982_at | 42.8 | M | 69 | P | 60 | A | 73.2 | P | ATL8 |
| 259983_at | 3402.8 | P | 2461.8 | P | 2815.4 | P | 2644.3 | P | HMG1 |
| 259984_at | 549.5 | P | 425.1 | P | 438.8 | P | 468.7 | P | --- |
| 259985_at | 148.3 | P | 234.2 | P | 103.2 | P | 242.1 | P | --- |
| 259986_at | 36.4 | A | 41.9 | A | 31.6 | A | 10.5 | A | --- |
| 259987_at | 34.5 | A | 63.7 | P | 62.4 | A | 63.4 | P | ATLP-3 |
| 259988_at | 0.6 | A | 0.4 | A | 8.6 | A | 4 | A | --- |
| 259989_at | 1.4 | A | 5.3 | A | 16.9 | A | 3.3 | A | --- |
| 259990_s_at | 835.3 | P | 692.2 | P | 614.6 | P | 774.9 | P | FKF1 |
| 259991_at | 28.8 | A | 28.1 | A | 51.1 | A | 35.3 | A | --- |
| 259992_at | 452.2 | P | 533.2 | P | 342.3 | P | 486.9 | P | HSFA8 |
| 259993_at | 285.6 | P | 235.1 | P | 266.7 | P | 244.7 | P | --- |
| 259994_at | 377.9 | P | 275.4 | P | 359.3 | P | 293.1 | P | AtIDD14 |
| 259995_at | 552.5 | P | 768.7 | P | 499.4 | P | 555.9 | P | --- |
| 259996_at | 161 | P | 113.6 | P | 185.5 | P | 124.4 | P | --- |
| 259997_at | 244 | P | 220.8 | P | 243.6 | P | 230.9 | P | --- |
| 259998_at | 57.6 | A | 49.6 | P | 55.6 | P | 42.9 | A | BPC3 |
| 259999_at | 120.8 | P | 162.7 | P | 183.9 | P | 180.6 | P | --- |
| 260000_at | 218.3 | P | 355.4 | P | 215.1 | P | 302.4 | P | ATMAP70-1 |
| 260001_at | 11 | A | 16.5 | A | 18.5 | A | 11.4 | A | TSM1 |
| 260002_at | 644.4 | P | 735.9 | P | 545 | P | 613.4 | P | ATNAP3 |
| 260003_at | 435.4 | P | 502.2 | P | 379.8 | P | 423.1 | P | IAR1 |
| 260004_at | 584.2 | P | 673 | P | 737.1 | P | 614.3 | P | --- |
| 260005_at | 29.5 | A | 48.4 | P | 40 | A | 24.6 | A | --- |
| 260006_at | 537.5 | P | 655.5 | P | 467.9 | P | 498 | P | ATPIS1 |
| 260007_at | 1102 | P | 1386.9 | P | 1484.1 | P | 1609.6 | P | --- |
| 260008_at | 453.3 | P | 429.2 | P | 489.5 | P | 427.6 | P | --- |
| 260009_at | 394.5 | P | 452.4 | P | 327.9 | P | 443.5 | P | --- |
| 260010_at | 259.1 | P | 183.2 | P | 208.2 | P | 152.6 | P | ATTPS6 |
| 260011_at | 60.7 | P | 72.9 | P | 76.3 | P | 54.9 | P | --- |
| 260012_at | 886.7 | P | 981.3 | P | 804.5 | P | 1283.1 | P | --- |
| 260013_at | 1.8 | A | 12 | A | 3.6 | A | 1.2 | A | ANN5 |
| 260014_at | 7933.5 | P | 8662.1 | P | 8570.9 | P | 8227.9 | P | HPR |
| 260015_at | 49.2 | P | 42.6 | M | 64.4 | P | 41.5 | P | CCOAMT |
| 260016_at | 0.9 | A | 0.7 | A | 1.5 | A | 4.7 | A | --- |
| 260017_at | 19.4 | A | 27.1 | A | 35.7 | A | 25.2 | P | --- |
| 260018_at | 8 | A | 10.1 | A | 4.4 | A | 1.8 | A | --- |
| 260019_at | 1.7 | A | 16.6 | A | 3.8 | A | 13.2 | A | --- |
| 260020_at | 904.1 | P | 796.6 | P | 697.2 | P | 774.4 | P | PDF6 |
| 260021_at | 116.5 | P | 113.6 | P | 111 | P | 135.3 | P | --- |
| 260022_at | 23.8 | P | 30.1 | P | 37 | P | 15.1 | A | --- |
| 260023_at | 128.1 | P | 108.4 | A | 106.6 | A | 80.4 | A | ATGA2OX2 |
| 260024_at | 17 | A | 19.8 | A | 44 | A | 26.9 | A | --- |
| 260025_at | 408.8 | P | 469.3 | P | 368.9 | P | 454.5 | P | --- |
| 260026_at | 757.7 | P | 958.4 | P | 843.7 | P | 836.3 | P | RPL18AA |
| 260027_at | 472.2 | P | 473.9 | P | 460.2 | P | 466.5 | P | CPuORF34 |
| 260028_at | 850 | P | 864 | P | 952.4 | P | 885.2 | P | --- |
| 260029_at | 306 | P | 323.3 | P | 283.6 | P | 312.4 | P | --- |
| 260030_at | 17.3 | A | 15.2 | P | 32.4 | P | 22.7 | P | AtbZIP |
| 260031_at | 362.6 | P | 435.4 | P | 483 | P | 454.9 | P | LINC3 |
| 260032_at | 15.4 | A | 28.7 | A | 56.9 | P | 44.8 | M | ATPPC4 |
| 260033_at | 204.8 | P | 279.2 | P | 259.3 | P | 268.4 | P | ATNUDX1 |
| 260034_at | 79.2 | P | 144.6 | P | 176 | P | 200.3 | P | --- |
| 260035_at | 24.8 | A | 27.9 | P | 45.1 | P | 45.1 | P | --- |
| 260036_at | 1097.5 | P | 1450.6 | P | 982.3 | P | 1255.2 | P | STN7 |
| 260037_at | 253.6 | P | 652.7 | P | 321 | P | 514.4 | P | RAV2 |
| 260038_at | 1.2 | A | 14.7 | A | 13.9 | A | 4.1 | A | --- |
| 260039_at | 52.5 | P | 33.4 | P | 48.9 | A | 58.7 | P | CLE12 |
| 260040_at | 27.7 | A | 7.2 | A | 30.4 | A | 28.4 | A | IDA |
| 260041_at | 141.1 | P | 144.8 | P | 207.6 | P | 171 | P | --- |
| 260042_at | 406.1 | P | 403.7 | P | 290.9 | P | 360.2 | P | --- |
| 260043_at | 1.9 | A | 1.9 | A | 5.8 | A | 2.1 | A | --- |
| 260044_at | 1549.9 | P | 2375.6 | P | 1224.2 | P | 2162.6 | P | --- |
| 260045_at | 229.8 | P | 316.6 | P | 246.4 | P | 286.2 | P | ATMPK15 |
| 260046_at | 159.9 | P | 197.2 | P | 84.4 | P | 142.8 | P | --- |
| 260047_at | 179.2 | P | 208.4 | P | 205.5 | P | 187.8 | P | --- |

| | | | | | | | | | |
|---|---|---|---|---|---|---|---|---|---|
| 260048_at | 405.4 | P | 552.9 | P | 405.9 | P | 427.8 | P | --- |
| 260049_at | 868.7 | P | 551.6 | P | 1075 | P | 594.9 | P | NRPA2 |
| 260050_at | 25 | A | 0.9 | A | 8.8 | A | 0.8 | A | AGL64 /// AT1G29960 |
| 260051_at | 776 | P | 716.2 | P | 600.3 | P | 781 | P | --- |
| 260052_at | 16.6 | A | 22.9 | A | 3.6 | A | 12.8 | A | GRF13 |
| 260053_at | 7.5 | A | 46.8 | A | 13.1 | A | 43.5 | A | --- |
| 260054_at | 101.1 | P | 153.2 | P | 84.9 | P | 124.9 | P | UNE2 |
| 260055_at | 745.5 | P | 943.7 | P | 929.9 | P | 832.5 | P | --- |
| 260056_at | 1005.5 | P | 945.8 | P | 951.5 | P | 842.9 | P | --- |
| 260057_at | 627.8 | P | 811.9 | P | 634 | P | 795.1 | P | --- |
| 260058_at | 239.4 | P | 424.7 | P | 347.8 | P | 437.2 | P | --- |
| 260059_at | 15.8 | A | 27.1 | A | 3.5 | A | 8.7 | A | ATTPPB |
| 260060_at | 403.1 | P | 515.3 | P | 387.4 | P | 402.1 | P | --- |
| 260061_at | 45.5 | A | 28.1 | A | 5.8 | A | 29.6 | A | CDKD1;1 |
| 260062_at | 174.2 | P | 192.4 | P | 226.8 | P | 168.8 | P | --- |
| 260063_at | 277 | P | 385.1 | P | 327.9 | P | 313.7 | P | --- |
| 260064_at | 299.6 | P | 214.7 | P | 304.9 | P | 235.4 | P | EIL3 |
| 260065_at | 587.9 | P | 541.5 | P | 511 | P | 581.8 | P | --- |
| 260066_at | 19.1 | A | 23.9 | A | 52.7 | A | 7.7 | A | --- |
| 260067_at | 21.3 | A | 14.2 | A | 1.3 | A | 12.1 | A | --- |
| 260068_at | 65.7 | P | 98.7 | P | 61.6 | P | 93.3 | P | --- |
| 260069_at | 361.6 | P | 379.6 | P | 370.7 | P | 363.1 | P | --- |
| 260070_at | 68 | P | 251.6 | P | 121.4 | P | 177.6 | P | BEE3 |
| 260071_at | 465.9 | P | 505.4 | P | 502.3 | P | 551.8 | P | ESP1 |
| 260072_at | 1091.5 | P | 1404 | P | 1020.6 | P | 1239.4 | P | --- |
| 260073_at | 233.2 | P | 250.9 | P | 265.5 | P | 260.1 | P | --- |
| 260074_at | 61.9 | P | 74.2 | P | 92.2 | P | 88.3 | P | AtRABA6a |
| 260075_at | 86.2 | P | 46.8 | P | 93.9 | P | 39.8 | P | --- |
| 260076_at | 180.5 | P | 238.1 | P | 246.1 | P | 229.4 | P | --- |
| 260077_at | 143.1 | P | 318.6 | P | 177.7 | P | 308.9 | P | --- |
| 260078_at | 411.3 | P | 345.3 | P | 292.4 | P | 376.4 | P | --- |
| 260079_s_at | 567.8 | P | 636.6 | P | 536.5 | P | 622.9 | P | --- |
| 260080_at | 34.9 | A | 45.5 | A | 46.9 | A | 38.6 | P | APUM7 |
| 260081_at | 185.6 | P | 375.5 | P | 239.6 | P | 256.6 | P | --- |
| 260082_at | 372.7 | P | 439.1 | P | 433.8 | P | 459.3 | P | --- |
| 260083_at | 213.7 | P | 287.8 | P | 186.1 | P | 219.9 | P | --- |
| 260084_at | 3.5 | A | 3.4 | A | 6.7 | A | 2.6 | A | --- |
| 260085_at | 944.7 | P | 876.2 | P | 1041.8 | P | 975.9 | P | --- |
| 260086_at | 114.9 | P | 196.3 | P | 139 | P | 157.6 | P | --- |
| 260087_at | 381.5 | P | 402.9 | P | 347.9 | P | 343.9 | P | --- |
| 260088_at | 8.9 | A | 15.7 | A | 58.5 | A | 4.1 | A | TIP3;1 |
| 260089_at | 1220.2 | P | 1478.5 | P | 1299.3 | P | 1396.2 | P | --- |
| 260090_at | 57.8 | P | 49.2 | P | 39.7 | P | 37.1 | P | scpl4 |
| 260091_at | 13.3 | A | 5.2 | A | 3 | A | 1.8 | A | scpl5 |
| 260092_at | 12.2 | A | 2.8 | A | 14.7 | A | 1.2 | A | scpl3 |
| 260093_at | 1.2 | A | 9 | A | 1.8 | A | 0.8 | A | scpl6 |
| 260094_at | 234.7 | P | 316.1 | P | 297 | P | 303.5 | P | GER1 |
| 260095_at | 3873.9 | P | 3916.6 | P | 4728.2 | P | 4512.7 | P | --- |
| 260096_at | 240.1 | P | 235 | P | 301.4 | P | 300.1 | P | --- |
| 260097_at | 1.8 | A | 1.4 | A | 3.4 | A | 1.1 | A | AtOCT1 |
| 260098_at | 108.1 | P | 121.8 | P | 132.8 | P | 130.3 | P | --- |
| 260099_at | 690.6 | P | 565.4 | P | 772 | P | 597.3 | P | --- |
| 260100_at | 332.1 | P | 257.2 | P | 301.6 | P | 258.6 | P | BNS |
| 260101_at | 138 | P | 176.1 | P | 118.2 | A | 146.4 | P | --- |
| 260102_at | 344.5 | P | 422.5 | P | 325.1 | P | 362.4 | P | --- |
| 260103_at | 146.4 | P | 187.3 | P | 117.5 | P | 133.3 | P | --- |
| 260104_x_at | 38 | A | 14.1 | A | 7.2 | A | 15.5 | A | --- |
| 260105_at | 7.8 | M | 2.4 | A | 10.9 | A | 6.5 | A | --- |
| 260106_at | 1687.7 | P | 2198.8 | P | 1906.9 | P | 2134.3 | P | --- |
| 260107_at | 1365.1 | P | 1348.3 | P | 1481 | P | 1602.5 | P | --- |
| 260108_at | 12.7 | A | 16.4 | A | 13.5 | A | 23.5 | A | --- |
| 260109_at | 139.9 | P | 298.2 | P | 171.2 | P | 270.3 | P | TET10 |
| 260110_at | 130.5 | P | 83.6 | P | 130.9 | P | 78.4 | P | --- |
| 260111_at | 18.3 | A | 2.7 | A | 1.1 | A | 4.8 | A | --- |
| 260112_at | 86.7 | P | 169.8 | P | 175.2 | P | 185.2 | P | --- |
| 260113_at | 23.4 | P | 37.6 | P | 38.2 | A | 44.2 | A | --- |
| 260114_at | 1.8 | A | 18 | A | 17.8 | A | 9.3 | A | --- |
| 260115_at | 21.6 | A | 19 | A | 11.2 | A | 8.5 | A | --- |
| 260116_at | 1301.7 | P | 736.6 | P | 646.4 | P | 361 | P | AIG1 |
| 260117_at | 52.4 | P | 40.9 | P | 34.6 | A | 12.2 | A | --- |
| 260118_s_at | 187.6 | P | 280.7 | P | 150.2 | P | 276.8 | P | --- |
| 260119_at | 28.8 | A | 33.9 | P | 44.1 | A | 26.2 | M | --- |
| 260120_at | 6.3 | A | 5.9 | A | 37.4 | A | 22.9 | A | ATPP2-A4 |
| 260121_at | 3.3 | A | 23.9 | A | 23.4 | A | 0.9 | A | --- |
| 260122_at | 2 | A | 4.3 | A | 1.5 | A | 1.7 | A | --- |
| 260123_at | 33.1 | P | 21.9 | P | 35.9 | M | 30.4 | P | --- |
| 260124_at | 4.1 | A | 2.1 | A | 6.1 | A | 3 | A | UBC31 |
| 260125_at | 515.5 | P | 707.7 | P | 520.3 | P | 639.6 | P | --- |
| 260126_at | 531.9 | P | 470.9 | P | 491.2 | P | 443.7 | P | SHM7 |
| 260127_at | 2113.1 | P | 1998.5 | P | 2070.6 | P | 2197 | P | --- |
| 260128_at | 999.8 | P | 1208.5 | P | 1178.9 | P | 1223.7 | P | --- |
| 260129_at | 624.1 | P | 664.9 | P | 496.9 | P | 464 | P | --- |
| 260130_s_at | 13.4 | A | 47.2 | P | 52.2 | A | 29.3 | P | BGLU21 |
| 260131_at | 13.2 | A | 3.8 | A | 10.7 | A | 24.3 | A | --- |
| 260132_s_at | 2.2 | A | 11.5 | A | 5.3 | A | 17.2 | A | --- |
| 260133_at | 315.1 | P | 365.8 | P | 293 | P | 288.5 | P | ETR1 |
| 260134_at | 207.2 | P | 36.9 | A | 258.6 | P | 74.1 | P | MYB113 |
| 260135_at | 242.2 | P | 235.1 | P | 183.1 | P | 179.1 | P | --- |
| 260136_at | 25.1 | A | 4.2 | A | 3.1 | A | 6.1 | A | --- |
| 260137_at | 603.1 | P | 1136.9 | P | 984.8 | P | 856.9 | P | --- |
| 260138_at | 3081.7 | P | 3272 | P | 3403.6 | P | 3326.9 | P | CAM4 |
| 260139_at | 20.8 | A | 9.2 | A | 4.2 | A | 2.5 | A | MYB114 |
| 260140_at | 652.1 | P | 417 | P | 655.3 | P | 997 | P | MYB90 |
| 260141_at | 271.5 | P | 250.2 | P | 240.7 | P | 254.6 | P | RGL1 |
| 260142_at | 289.8 | P | 337.2 | P | 459 | P | 375.2 | P | --- |
| 260143_at | 1031.8 | P | 1456.5 | P | 1371.1 | P | 1599.2 | P | SUC1 |
| 260144_at | 161.8 | P | 177.3 | P | 137.2 | P | 157.4 | P | --- |
| 260145_at | 23 | A | 13.2 | A | 35.9 | A | 57.2 | A | GPCR |
| 260146_at | 1063 | P | 591.7 | P | 796.5 | P | 635.4 | P | --- |
| 260147_at | 4.1 | A | 1.2 | A | 6.5 | A | 11.1 | A | --- |
| 260148_at | 23.1 | A | 52.6 | A | 6 | A | 48.2 | A | --- |
| 260149_at | 2.1 | A | 36.9 | P | 60.7 | P | 15.6 | A | --- |
| 260150_at | 17.5 | A | 13.5 | A | 8.7 | A | 2.1 | A | --- |
| 260151_at | 190.2 | P | 352.4 | P | 281.4 | P | 344.9 | P | --- |
| 260152_at | 28.4 | A | 41.9 | A | 56.1 | A | 36 | A | IAA6 |
| 260153_at | 3338.3 | P | 2498.9 | P | 2757.1 | P | 2540.5 | P | --- |
| 260154_at | 2937 | P | 3063 | P | 3282.2 | P | 3334.1 | P | HTA9 |
| 260155_at | 1763.4 | P | 3156.2 | P | 1854.2 | P | 2970.9 | P | --- |
| 260156_at | 523.6 | P | 561.9 | P | 379 | P | 615.1 | P | NAM |

| | | | | | | | | | |
|---|---|---|---|---|---|---|---|---|---|
| 260157_at | 1299.4 | P | 1100.4 | P | 1340.8 | P | 1253.2 | P | --- |
| 260158_at | 42.8 | A | 44.3 | A | 35.6 | A | 45.5 | A | --- |
| 260159_at | 15.7 | A | 22.9 | A | 37.2 | A | 19.9 | A | --- |
| 260160_at | 121.3 | P | 135.4 | P | 128.3 | P | 112.4 | P | --- |
| 260161_at | 21.4 | A | 8.2 | A | 25.5 | A | 14.4 | A | ROPGEF12 |
| 260162_at | 1100.3 | P | 1285.9 | P | 1040.8 | P | 1221.7 | P | GC5 |
| 260163_at | 85.5 | P | 29.5 | A | 7.1 | A | 40 | A | BAC2 |
| 260164_at | 1257.1 | P | 1393.6 | P | 1272.2 | P | 1360.3 | P | --- |
| 260165_at | 5742.8 | P | 6678.4 | P | 6656.1 | P | 6464.1 | P | RPS17 |
| 260166_at | 151.7 | P | 88.5 | M | 94.7 | A | 128.9 | P | GL2 |
| 260167_at | 276.2 | P | 297.5 | P | 189.1 | P | 202.7 | P | --- |
| 260168_at | 863.5 | P | 741.6 | P | 839.5 | P | 796 | P | --- |
| 260169_at | 223.2 | P | 185.9 | P | 214.7 | P | 225.5 | P | FUT13 |
| 260170_at | 68.4 | P | 69 | A | 54.7 | P | 36.4 | A | SUC5 |
| 260171_at | 70 | P | 120.8 | P | 102.8 | P | 72.5 | P | --- |
| 260172_s_at | 465.6 | P | 507.5 | P | 431.8 | P | 539.3 | P | --- |
| 260173_at | 45.7 | P | 53.5 | P | 44.1 | A | 41.4 | A | VND7 |
| 260174_at | 226.2 | P | 312.7 | P | 256.6 | P | 280 | P | --- |
| 260175_at | 28.1 | A | 6.3 | A | 31.3 | A | 30.1 | P | --- |
| 260176_at | 1101 | P | 1130.8 | P | 1075.1 | P | 1085 | P | --- |
| 260177_at | 66.9 | P | 68.5 | M | 25.3 | A | 36 | P | --- |
| 260178_at | 3.9 | A | 3.2 | A | 9.7 | A | 2.8 | A | --- |
| 260179_at | 76.5 | A | 103.6 | A | 24.1 | A | 53.9 | A | HWI1 |
| 260180_at | 1477.3 | P | 1606.8 | P | 1473.6 | P | 1571.6 | P | --- |
| 260181_at | 199.1 | P | 362.1 | P | 251.8 | P | 296.9 | P | ATGH9B1 |
| 260182_at | 194.3 | P | 175.6 | P | 241.9 | P | 190.9 | P | --- |
| 260183_at | 9.3 | A | 1.5 | A | 3.7 | A | 1.4 | A | --- |
| 260184_s_at | 1.3 | A | 0.9 | A | 4.5 | A | 4.7 | A | --- |
| 260185_at | 5.1 | A | 4.1 | A | 0.7 | A | 0.6 | A | --- |
| 260186_at | 4 | A | 19.1 | A | 18.7 | A | 17.5 | A | --- |
| 260187_at | 2.4 | A | 2.3 | A | 3.3 | A | 1.5 | A | LBD5 |
| 260188_at | 4.4 | A | 3.2 | A | 11.5 | A | 4.3 | A | --- |
| 260189_at | 256.1 | P | 184.1 | P | 258.1 | P | 178 | P | URE |
| 260190_at | 533.2 | P | 401.8 | P | 410.4 | P | 438.1 | P | --- |
| 260191_at | 550.4 | P | 502.5 | P | 685.2 | P | 613.6 | P | --- |
| 260192_at | 121 | P | 223.5 | P | 225.2 | P | 205.8 | P | POLA2 |
| 260193_at | 6.8 | A | 24.1 | A | 72.3 | A | 54.1 | A | --- |
| 260194_at | 198.1 | P | 191.5 | P | 193.1 | P | 170.4 | P | --- |
| 260195_at | 21.1 | A | 2.7 | A | 50.3 | A | 17.1 | A | --- |
| 260196_at | 269.2 | P | 222.9 | P | 338.4 | P | 188.1 | P | --- |
| 260197_at | 11.3 | A | 6.7 | A | 31.7 | P | 12.7 | A | --- |
| 260198_at | 2.7 | A | 8.2 | A | 11 | A | 22.7 | A | --- |
| 260199_at | 72.4 | P | 95.7 | P | 66.7 | P | 64.9 | P | --- |
| 260200_at | 199.9 | P | 124.7 | P | 182.7 | P | 189.6 | P | --- |
| 260201_at | 42.5 | A | 51.4 | A | 49.2 | A | 60.1 | A | --- |
| 260202_at | 6.7 | A | 7.1 | A | 2.5 | A | 9.7 | A | --- |
| 260203_at | 763.3 | P | 418 | P | 264.9 | P | 345.6 | P | ANAC019 |
| 260204_at | 21.1 | P | 7 | A | 19 | A | 9.2 | A | --- |
| 260205_at | 1606.1 | P | 703 | P | 1777 | P | 1029.9 | P | TIFY7 |
| 260206_at | 180.6 | P | 168 | P | 161.1 | P | 157.3 | P | --- |
| 260207_at | 1466.4 | P | 1684.7 | P | 1784.1 | P | 1871.8 | P | --- |
| 260208_s_at | 495.7 | P | 382.6 | P | 444.1 | P | 342.8 | P | --- |
| 260209_at | 687.1 | P | 522.1 | P | 666.5 | P | 644.2 | P | CPuORF53 |
| 260210_at | 79.6 | P | 68.8 | P | 103 | P | 61.9 | P | FUT3 |
| 260211_at | 225.5 | P | 289.5 | P | 167.5 | P | 220.9 | P | --- |
| 260212_at | 35.3 | A | 23.8 | A | 33.6 | A | 26.7 | P | --- |
| 260213_at | 35.2 | A | 22.1 | A | 15.2 | A | 24.3 | A | --- |
| 260214_at | 254 | P | 283.3 | P | 290.5 | P | 303.8 | P | --- |
| 260215_at | 412.2 | P | 471 | P | 601.1 | P | 518.3 | P | --- |
| 260216_at | 32 | A | 35.2 | A | 33.5 | A | 26.7 | A | --- |
| 260217_at | 89.7 | P | 86.5 | P | 126.9 | P | 140 | P | --- |
| 260218_at | 30.1 | A | 28.1 | A | 35.6 | A | 37.7 | A | --- |
| 260219_at | 35.5 | M | 17.3 | A | 71.7 | P | 37.8 | P | --- |
| 260220_at | 32.5 | P | 38.2 | P | 32.6 | A | 70.1 | A | MYB31 |
| 260221_at | 198.5 | P | 903.6 | P | 441.4 | P | 694.4 | P | --- |
| 260222_at | 839.9 | P | 834.2 | P | 524.1 | P | 1192.3 | P | XXT5 |
| 260223_at | 140.5 | P | 116.3 | P | 159.4 | P | 144.7 | P | --- |
| 260224_at | 55.8 | P | 83.9 | P | 102.2 | P | 78.5 | P | --- |
| 260225_at | 8.4 | A | 39.5 | A | 2.5 | A | 4.2 | A | GSTU10 |
| 260226_at | 27.8 | A | 42.9 | A | 38.8 | A | 7.2 | A | MIF1 |
| 260227_at | 514.2 | P | 569.4 | P | 566.9 | P | 764.9 | P | --- |
| 260228_at | 6.3 | A | 5.8 | A | 3.3 | A | 1.5 | A | CYP98A8 |
| 260229_at | 189.7 | P | 301.6 | P | 331.6 | P | 225.5 | P | --- |
| 260230_at | 7.5 | A | 0.5 | A | 0.8 | A | 2.7 | A | --- |
| 260231_at | 380.3 | P | 409.1 | P | 347.6 | P | 420.5 | P | --- |
| 260232_at | 898.8 | P | 1111.2 | P | 796.9 | P | 1047.7 | P | --- |
| 260233_at | 40.7 | P | 55.9 | P | 37 | A | 37.2 | M | CYP98A9 |
| 260234_at | 27.6 | A | 101.2 | P | 97.8 | M | 120.3 | P | --- |
| 260235_at | 1668.8 | P | 1491.9 | P | 2205.4 | P | 1655.7 | P | NRP1 |
| 260236_at | 5088 | P | 7156.9 | P | 6737.9 | P | 6890.1 | P | --- |
| 260237_at | 326 | P | 216.9 | P | 256.5 | P | 297.2 | P | MYB95 |
| 260238_at | 1267.6 | P | 1483.3 | P | 1378.7 | P | 1495.2 | P | ATHVA22A |
| 260239_at | 100.9 | P | 78.7 | P | 85.9 | P | 66.1 | P | --- |
| 260240_at | 17.6 | A | 12.5 | A | 31.3 | A | 17.9 | A | ATIPT1 |
| 260241_at | 34.9 | A | 53.3 | P | 17 | A | 66.5 | P | CYP86A7 |
| 260242_at | 65.4 | P | 41 | P | 28.3 | A | 59.1 | P | EGL3 |
| 260243_at | 400 | P | 298.3 | P | 209.9 | P | 203.1 | P | --- |
| 260244_at | 353.9 | P | 416.7 | P | 405 | P | 373.3 | P | --- |
| 260245_at | 1278.2 | P | 1460 | P | 1318.5 | P | 1364.8 | P | GR-RBP5 |
| 260246_at | 14.9 | A | 5.3 | A | 28.3 | A | 2 | A | --- |
| 260247_at | 47.8 | P | 36.8 | P | 67.2 | A | 32.4 | P | AtRLP16 |
| 260248_at | 118.4 | P | 171 | P | 121.2 | P | 126.1 | P | ATHSP101 |
| 260249_s_at | 146.9 | P | 188 | P | 110.6 | P | 157.7 | P | --- |
| 260250_at | 1312.4 | P | 1191.4 | P | 1487.5 | P | 1204.6 | P | PUR4 |
| 260251_at | 500.5 | P | 302.2 | P | 467.4 | P | 350.2 | P | --- |
| 260252_at | 258.5 | P | 210.3 | P | 333.2 | P | 289.5 | P | --- |
| 260253_at | 6.4 | A | 2.5 | A | 5.5 | A | 0.9 | A | AtRLP15 |
| 260254_at | 543.1 | P | 976.4 | P | 605.2 | P | 837.9 | P | --- |
| 260255_at | 118.7 | P | 87.7 | P | 158.3 | P | 77 | P | --- |
| 260256_at | 48.4 | A | 69.5 | P | 75.8 | M | 72.1 | P | --- |
| 260257_at | 1263.6 | P | 1220.4 | P | 1226.2 | P | 1056.8 | P | --- |
| 260258_at | 2048.7 | P | 1608.9 | P | 2063.1 | P | 1658.2 | P | --- |
| 260259_at | 74.9 | A | 149.4 | P | 113.9 | M | 102.2 | P | --- |
| 260260_at | 291.1 | P | 273 | P | 370.6 | P | 333.3 | P | --- |
| 260261_at | 73 | P | 68.8 | P | 114.4 | P | 76.7 | P | --- |
| 260262_at | 152.9 | P | 135.7 | P | 167 | P | 146.2 | P | --- |
| 260263_at | 24.3 | P | 24 | A | 42.1 | P | 33.1 | A | JAG |
| 260264_at | 120.8 | P | 195.1 | P | 201.1 | P | 240.3 | P | --- |
| 260265_at | 18.1 | A | 3 | A | 17.9 | A | 2.3 | A | LBD42 |

| | | | | | | | | | |
|---|---|---|---|---|---|---|---|---|---|
| 260266_at | 574.3 | P | 533.8 | P | 575.6 | P | 399.4 | P | --- |
| 260267_at | 3501.9 | P | 3039.2 | P | 2815.4 | P | 3452.7 | P | KCS6 |
| 260268_at | 1028.3 | P | 1106.4 | P | 993.1 | P | 922.3 | P | --- |
| 260269_at | 57.3 | P | 64.2 | P | 67 | P | 85.7 | P | --- |
| 260270_at | 39.9 | P | 61.7 | P | 57.9 | P | 55 | P | --- |
| 260271_at | 859.5 | P | 850.2 | P | 714.8 | P | 757.6 | P | --- |
| 260272_at | 95.3 | P | 100.8 | P | 102.4 | P | 102.1 | P | --- |
| 260273_at | 163.7 | P | 193.3 | P | 117 | P | 163.5 | P | --- |
| 260274_at | 633.4 | P | 1004.8 | P | 645.8 | P | 757.7 | P | NHO1 |
| 260275_at | 19.8 | A | 38 | P | 35.6 | A | 30.9 | P | --- |
| 260276_at | 95.3 | M | 102.9 | P | 73.1 | P | 105.7 | P | --- |
| 260277_at | 78.2 | P | 126 | P | 133.8 | P | 117.6 | P | --- |
| 260278_at | 25.8 | A | 30.9 | P | 55.3 | A | 14.2 | A | WRKY66 |
| 260279_at | 206 | P | 245.8 | P | 255 | P | 176 | P | --- |
| 260280_at | 9.2 | A | 3.4 | A | 4.1 | A | 2 | A | --- |
| 260281_at | 1084.3 | P | 1256.4 | P | 1060.4 | P | 1348.2 | P | --- |
| 260282_at | 1071.4 | P | 997.6 | P | 1245.5 | P | 978.5 | P | EMB2753 |
| 260283_at | 2172 | P | 2282.4 | P | 2868.2 | P | 2600.1 | P | PTAC17 |
| 260284_at | 2807.2 | P | 2355.7 | P | 2629.4 | P | 2584.5 | P | --- |
| 260285_at | 1163.8 | P | 948.4 | P | 1269.2 | P | 988.5 | P | --- |
| 260286_at | 3838.5 | P | 3269.1 | P | 3536.8 | P | 3492.2 | P | WIN1 |
| 260287_at | 222.4 | P | 238.5 | P | 127.1 | P | 156.3 | P | --- |
| 260288_at | 954.9 | P | 1362.8 | P | 1091.2 | P | 1243.5 | P | --- |
| 260289_at | 328.1 | P | 420.6 | P | 350.6 | P | 429.5 | P | ERH3 |
| 260290_at | 212.9 | P | 283.4 | P | 217.4 | P | 202.8 | P | --- |
| 260291_at | 245.3 | P | 358.4 | P | 256.1 | P | 360.5 | P | YDA |
| 260292_at | 485.1 | P | 529 | P | 528.4 | P | 448.7 | P | MURE |
| 260293_at | 214.4 | P | 260.9 | P | 227.9 | P | 244.6 | P | --- |
| 260294_at | 1077.8 | P | 931.8 | P | 1483.9 | P | 1045.5 | P | --- |
| 260295_at | 4291.6 | P | 3767.9 | P | 4614.9 | P | 3925.6 | P | --- |
| 260296_at | 266.7 | P | 217.8 | P | 149.1 | P | 183.4 | P | --- |
| 260297_at | 134.4 | P | 284.1 | P | 147.2 | P | 266.9 | P | --- |
| 260298_at | 2.9 | A | 6 | A | 7.4 | A | 6.4 | A | --- |
| 260299_at | 40.8 | A | 7.3 | A | 7.8 | A | 8.6 | A | GA3OX4 |
| 260300_at | 4 | A | 10.2 | A | 12 | A | 5.7 | A | GA3OX2 |
| 260301_at | 128.4 | P | 174.1 | P | 135.8 | P | 144.6 | P | --- |
| 260302_at | 586.3 | P | 450.6 | P | 546 | P | 511.5 | P | --- |
| 260303_at | 215.9 | P | 339 | P | 195.6 | P | 281.1 | P | --- |
| 260304_at | 904.3 | P | 565.8 | P | 630 | P | 591.2 | P | --- |
| 260305_at | 3473.3 | P | 3967.5 | P | 3599.2 | P | 4304 | P | ARFA1D |
| 260306_at | 6.2 | A | 3.9 | A | 2.2 | A | 8.6 | A | EDA24 |
| 260307_at | 422.1 | P | 486.6 | P | 411.7 | P | 463.3 | P | --- |
| 260308_at | 440.5 | P | 590.1 | P | 543.6 | P | 568.8 | P | ATTAP1 |
| 260309_at | 546.2 | P | 641.7 | P | 432 | P | 764.8 | P | AOAT2 |
| 260310_at | 226 | P | 244.6 | P | 227.6 | P | 254.2 | P | --- |
| 260311_at | 520.2 | P | 504.8 | P | 467.1 | P | 486.7 | P | --- |
| 260312_at | 774.5 | P | 913.6 | P | 520.8 | P | 707.9 | P | --- |
| 260313_at | 20.2 | A | 8.9 | A | 3.7 | A | 5.4 | A | --- |
| 260314_at | 2127.7 | P | 1946.9 | P | 1817.4 | P | 1855.5 | P | --- |
| 260315_at | 71.7 | P | 52.3 | M | 95 | P | 42.3 | A | --- |
| 260316_at | 476.4 | P | 369.3 | P | 535.1 | P | 281.5 | P | --- |
| 260317_at | 3775.8 | P | 1991.9 | P | 5001.3 | P | 1811.1 | P | UBC5 |
| 260318_at | 17.2 | A | 4.9 | A | 29.1 | A | 42.3 | A | --- |
| 260319_at | 2 | A | 5.1 | A | 1.7 | A | 1.6 | A | --- |
| 260320_at | 2.8 | A | 3.7 | A | 11.7 | A | 4.1 | A | --- |
| 260321_at | 817.1 | P | 718.5 | P | 708 | P | 686.4 | P | --- |
| 260322_at | 424.4 | P | 397.5 | P | 399.6 | M | 385.9 | P | --- |
| 260323_at | 924.3 | P | 669 | P | 1121.8 | P | 753.9 | P | IMP4 |
| 260324_at | 2856.3 | P | 3207.3 | P | 2577.1 | P | 3171.9 | P | ISPF |
| 260325_at | 1383.5 | P | 1464.6 | P | 1295.4 | P | 1558.2 | P | MDAR6 |
| 260326_at | 28.1 | A | 25.3 | A | 30.7 | P | 37.8 | A | AtMYB103 |
| 260327_at | 2153.8 | P | 1112.6 | P | 2137.1 | P | 1143.2 | P | --- |
| 260328_at | 1161.8 | P | 957.2 | P | 1058 | P | 912.4 | P | --- |
| 260329_at | 51.1 | A | 79 | P | 102.1 | A | 93.2 | P | CYCA2;4 |
| 260330_at | 507.1 | P | 441.7 | P | 355 | P | 456.1 | P | --- |
| 260331_at | 558.8 | P | 437 | P | 730 | P | 536.5 | P | --- |
| 260332_at | 157.3 | P | 161.7 | P | 143.3 | P | 161.6 | P | --- |
| 260333_at | 29.1 | A | 33.3 | A | 56.4 | A | 55 | M | --- |
| 260334_at | 34.2 | A | 33.1 | P | 107.2 | P | 53.7 | P | KNAT2 |
| 260335_at | 40.1 | A | 65.6 | A | 29.1 | A | 53.6 | P | SS3 |
| 260336_at | 903.5 | P | 879.5 | P | 829.8 | P | 686.1 | P | SPPA |
| 260337_at | 419.6 | P | 228.5 | P | 299 | P | 226 | P | WRKY57 |
| 260338_at | 495.1 | P | 568.2 | P | 484.7 | P | 580.4 | P | --- |
| 260339_at | 383.9 | P | 426.3 | P | 370 | P | 384.2 | P | SIK1 |
| 260340_at | 3.1 | A | 3 | A | 6.6 | A | 15.4 | A | --- |
| 260341_at | 62.2 | A | 120 | P | 104.2 | M | 62 | M | --- |
| 260342_at | 53.1 | P | 39 | P | 56.4 | A | 44.9 | P | --- |
| 260343_at | 793.6 | P | 953.2 | P | 946.2 | P | 898.2 | P | --- |
| 260344_at | 5 | A | 5.6 | A | 6.3 | A | 4.8 | A | MES15 |
| 260345_at | 185.3 | P | 217.9 | P | 171.9 | P | 196.1 | P | RPK1 |
| 260346_at | 8.8 | A | 42.3 | A | 13.3 | A | 7.5 | A | CLE10 |
| 260347_at | 408.1 | P | 519.5 | P | 488.3 | P | 603.2 | P | --- |
| 260348_at | 67.9 | P | 71 | P | 59.4 | P | 44 | A | --- |
| 260349_at | 45.5 | P | 44.3 | P | 58.6 | P | 59.8 | P | --- |
| 260350_at | 4861.2 | P | 4776.5 | P | 3578.5 | P | 4664.7 | P | ELF5A-3 |
| 260351_at | 204.1 | P | 221.7 | P | 218.7 | P | 215.9 | P | --- |
| 260352_at | 853.2 | P | 735.6 | P | 953.7 | P | 805.7 | P | PDCB4 |
| 260353_at | 88 | P | 91.5 | P | 94.5 | P | 89.1 | P | SP1L2 |
| 260354_at | 174.8 | P | 227 | P | 166.6 | P | 260.7 | P | --- |
| 260355_at | 1.3 | A | 4.3 | A | 2.6 | A | 1.6 | A | CRC |
| 260356_at | 1002.1 | P | 1201.4 | P | 1043.3 | P | 1174.3 | P | ATMINE1 |
| 260357_at | 177.1 | P | 70.9 | M | 85.9 | M | 64.8 | P | AFP1 |
| 260358_at | 744.8 | P | 768.8 | P | 635 | P | 788 | P | --- |
| 260359_at | 212.7 | P | 196.9 | P | 228.5 | P | 244.2 | P | --- |
| 260360_at | 378.9 | P | 298.8 | P | 380.2 | P | 301.2 | P | CM3 |
| 260361_at | 218.6 | P | 167.9 | P | 150.8 | P | 180.9 | P | --- |
| 260362_at | 245.8 | P | 366.1 | P | 253.6 | P | 344.8 | P | --- |
| 260363_at | 111.4 | P | 120.4 | P | 114 | P | 123.3 | P | --- |
| 260364_at | 554.7 | P | 489.8 | P | 560 | P | 590.2 | P | TAA1 |
| 260365_at | 66.7 | P | 72.7 | P | 60.5 | M | 63.8 | P | --- |
| 260366_at | 34 | A | 7.8 | A | 9.1 | A | 6.1 | A | --- |
| 260367_at | 497.6 | P | 465.8 | P | 674.3 | P | 374 | P | --- |
| 260368_at | 446.4 | P | 705.9 | P | 506 | P | 623.5 | P | ATHVA22C |
| 260369_at | 7607.1 | P | 7219 | P | 8106.6 | P | 7510.9 | P | RPL34 |
| 260370_at | 3623.6 | P | 4072.6 | P | 3646.5 | P | 3755.5 | P | HEMB1 |
| 260371_at | 366.3 | P | 554.6 | P | 401.7 | P | 463.9 | P | --- |
| 260372_at | 517.4 | P | 613.5 | P | 393.1 | P | 498.9 | P | --- |
| 260373_at | 222.7 | P | 269.3 | P | 220.1 | P | 236 | P | --- |
| 260374_at | 330.7 | P | 298.2 | P | 370 | P | 271.7 | P | TAF2 |

| | | | | | | | | | |
|---|---|---|---|---|---|---|---|---|---|
| 260375_at | 113 | P | 199.4 | P | 110.8 | P | 195.6 | P | --- |
| 260376_at | 2.2 | A | 1.6 | A | 2 | A | 1.7 | A | CYP78A10 |
| 260377_at | 311.2 | P | 294.5 | P | 480.2 | P | 326.3 | P | --- |
| 260378_at | 3.6 | A | 21.6 | A | 45.3 | A | 17.6 | A | ATARP4A |
| 260379_at | 982.4 | P | 726.7 | P | 867.1 | P | 840.1 | P | UGT89B1 |
| 260380_at | 62.5 | P | 72.5 | P | 22.4 | A | 45.9 | P | --- |
| 260381_at | 17.7 | A | 32.1 | A | 11.7 | A | 15 | A | --- |
| 260382_at | 53.1 | P | 25.9 | A | 77.9 | P | 33.3 | A | --- |
| 260383_s_at | 5866 | P | 5575.7 | P | 5320.3 | P | 5833.7 | P | --- |
| 260384_at | 432.1 | P | 346.9 | P | 421.5 | P | 370 | P | IMS1 |
| 260385_at | 4261.5 | P | 2795.1 | P | 5517 | P | 3016.2 | P | SOT18 |
| 260386_at | 103.9 | P | 44.3 | A | 79 | P | 31.5 | A | --- |
| 260387_at | 5857.4 | P | 3505.3 | P | 2952.3 | P | 2457.7 | P | SOT16 |
| 260388_at | 211 | P | 279.7 | P | 314.3 | P | 333.8 | P | --- |
| 260389_at | 23.6 | A | 54.1 | P | 15.2 | A | 56.9 | P | --- |
| 260390_at | 692.4 | P | 617.4 | P | 763.7 | P | 759.5 | P | --- |
| 260391_at | 869.3 | P | 640.5 | P | 685.4 | P | 618.4 | P | SS2 |
| 260392_at | 528.7 | P | 430.6 | P | 546.5 | P | 491.7 | P | --- |
| 260393_at | 668.4 | P | 800 | P | 546.1 | P | 528.3 | P | --- |
| 260394_at | 0.8 | A | 1.5 | A | 1.9 | A | 3.8 | A | MYB122 |
| 260395_at | 306.6 | P | 339 | P | 275.4 | P | 341.5 | P | ATHB13 |
| 260396_at | 47 | M | 89.3 | P | 49.3 | A | 90.4 | P | ho3 |
| 260397_at | 261.2 | P | 236.4 | P | 277.2 | P | 340.7 | P | --- |
| 260398_at | 359.1 | P | 344.1 | P | 384.1 | P | 399.2 | P | APUM23 |
| 260399_at | 764.7 | P | 110.6 | P | 266.1 | P | 156 | P | --- |
| 260400_at | 9.5 | A | 38.1 | A | 4.6 | A | 16.7 | A | --- |
| 260401_at | 2222.9 | P | 2534.7 | P | 1771.3 | P | 1770.4 | P | --- |
| 260402_at | 19.9 | A | 29.9 | P | 2 | A | 16.8 | A | GGT3 |
| 260403_at | 4.4 | A | 21.1 | A | 32.6 | A | 19.5 | A | WRKY36 |
| 260404_at | 121.6 | P | 130.2 | P | 156.5 | P | 166.6 | P | --- |
| 260405_at | 182.7 | P | 59.8 | P | 143 | P | 48.5 | A | ATGSTU11 |
| 260406_at | 17.7 | A | 1.9 | A | 12.7 | A | 17.8 | A | ATGSTU12 |
| 260407_at | 183.6 | P | 174.6 | P | 186.3 | P | 158.3 | P | --- |
| 260408_at | 857.7 | P | 152.1 | P | 507.5 | P | 187.4 | P | ATH8 |
| 260409_at | 603.1 | P | 719.5 | P | 733.5 | P | 810.7 | P | SHW1 |
| 260410_at | 4086.3 | P | 2491.9 | P | 3645.9 | P | 2784.9 | P | --- |
| 260411_at | 374 | P | 177.1 | P | 323.3 | P | 220.9 | P | --- |
| 260412_at | 2998.1 | P | 3289.3 | P | 3210.8 | P | 3488 | P | AMY3 |
| 260413_at | 476.9 | P | 426.8 | P | 526.5 | P | 432.1 | P | --- |
| 260414_at | 301.7 | P | 615.1 | P | 279.5 | P | 573.6 | P | ATNRT1:2 |
| 260415_at | 66 | P | 32.7 | P | 63.6 | P | 49.9 | P | --- |
| 260416_at | 196.8 | P | 196.4 | P | 143.7 | P | 222.1 | P | CUL3B |
| 260417_at | 334.4 | P | 381.4 | P | 256.3 | P | 395.5 | P | CMT3 |
| 260418_s_at | 894.5 | P | 687.1 | P | 794.6 | P | 606.9 | P | COX19-1 |
| 260419_at | 642.8 | P | 666 | P | 670 | P | 617.1 | P | --- |
| 260420_at | 212 | P | 147.7 | P | 176.2 | P | 150.6 | P | --- |
| 260421_at | 1181.6 | P | 1213.9 | P | 1151.6 | P | 1199.7 | P | SBH1 |
| 260422_at | 1.7 | A | 13.8 | P | 15.9 | A | 4 | A | --- |
| 260423_at | 321.8 | P | 191.1 | P | 246.1 | P | 252.1 | P | ATEXO70D1 |
| 260424_at | 47.2 | P | 59.1 | P | 54.6 | A | 60.3 | M | --- |
| 260425_at | 468.1 | P | 317.2 | P | 573.3 | P | 406.3 | P | EDA25 |
| 260426_at | 8944 | P | 8236.5 | P | 9459.7 | P | 8951.5 | P | P40 |
| 260427_at | 1442 | P | 1507.1 | P | 1757.1 | P | 1091.9 | P | --- |
| 260428_at | 426 | P | 490.4 | P | 437.7 | P | 484.2 | P | --- |
| 260429_at | 1559 | P | 901.6 | P | 1810.9 | P | 993.8 | P | JAZ6 |
| 260430_at | 37.3 | A | 63.8 | A | 29.7 | A | 10.1 | A | --- |
| 260431_at | 97.7 | A | 103.7 | A | 57.3 | A | 52.4 | A | --- |
| 260432_at | 9 | A | 6.1 | A | 2.3 | A | 27.5 | A | WRKY9 |
| 260433_at | 2.7 | A | 1.8 | A | 5.1 | A | 2.4 | A | --- |
| 260434_at | 36.2 | P | 47.1 | P | 69.1 | P | 49.8 | P | --- |
| 260435_at | 21.2 | A | 34.5 | A | 37.7 | A | 36.6 | A | MYB62 |
| 260436_at | 562.9 | P | 540.4 | P | 554.8 | P | 587.6 | P | --- |
| 260437_at | 21.9 | A | 15.9 | A | 29.7 | A | 21.2 | A | --- |
| 260438_at | 257.9 | P | 125.5 | P | 145.4 | P | 123.3 | P | ENDO 2 |
| 260439_at | 68.6 | P | 34.5 | P | 51.5 | M | 36.7 | A | --- |
| 260440_at | 313.4 | P | 346.6 | P | 394 | P | 367.9 | P | --- |
| 260441_at | 489.8 | P | 581 | P | 573.1 | P | 506.9 | P | --- |
| 260442_at | 675.2 | P | 867.8 | P | 808.7 | P | 1007.5 | P | --- |
| 260443_at | 206.7 | P | 239.4 | P | 252.1 | P | 282.1 | P | --- |
| 260444_at | 1731.3 | P | 1447.7 | P | 1230 | P | 1379 | P | --- |
| 260445_at | 612.2 | P | 642.7 | P | 744.6 | P | 771.4 | P | ARG1 |
| 260446_at | 416.2 | P | 402 | P | 421.4 | P | 420.3 | P | --- |
| 260447_at | 30 | A | 24.1 | M | 41 | A | 11.9 | A | --- |
| 260448_at | 183.8 | P | 286.8 | P | 310.9 | P | 382 | P | --- |
| 260449_at | 217.3 | P | 205.4 | P | 238.6 | P | 228.5 | P | --- |
| 260450_at | 64.9 | P | 39.4 | P | 33.2 | A | 65.6 | P | --- |
| 260451_at | 64.9 | A | 93.9 | P | 77.9 | A | 100.8 | M | --- |
| 260452_at | 4.4 | A | 25.5 | A | 6 | A | 21.8 | A | --- |
| 260453_s_at | 637 | P | 519.4 | P | 422.2 | P | 465.6 | P | --- |
| 260454_at | 368.1 | P | 419.8 | P | 346.7 | P | 385.8 | P | ATL3 |
| 260455_at | 537.2 | P | 631.3 | P | 557.1 | P | 575.1 | P | --- |
| 260456_at | 44.4 | P | 19.8 | A | 73.8 | A | 27.8 | A | --- |
| 260457_at | 273.6 | P | 373.5 | P | 352.9 | P | 351.2 | P | --- |
| 260458_at | 37.2 | P | 20.9 | A | 66.8 | P | 37.2 | P | --- |
| 260459_at | 22.8 | A | 27 | A | 6.9 | A | 15.5 | A | --- |
| 260460_at | 48 | P | 77.9 | P | 68.4 | P | 43.8 | P | --- |
| 260461_at | 52.1 | A | 34.6 | A | 45.5 | A | 40.1 | A | --- |
| 260462_at | 103.9 | P | 147.6 | P | 170.4 | P | 175.7 | P | ZIP4 |
| 260463_at | 91.7 | P | 175.2 | P | 167.9 | P | 136.6 | P | RECQ4A |
| 260464_at | 78.2 | P | 99.1 | P | 93.1 | P | 98.9 | P | LOV1 |
| 260465_at | 420.5 | P | 569.5 | P | 553.5 | P | 493.4 | P | --- |
| 260466_at | 228 | P | 294.4 | P | 269 | P | 271 | P | --- |
| 260467_at | 35 | P | 22 | A | 65.2 | A | 44.1 | A | --- |
| 260468_at | 36.4 | P | 42.1 | P | 25.3 | P | 65.8 | P | --- |
| 260469_at | 286.6 | P | 303.2 | P | 231.5 | P | 293.1 | P | AGD4 |
| 260470_at | 77.3 | A | 58.6 | A | 81.3 | A | 81.1 | A | --- |
| 260471_at | 22.8 | A | 1 | A | 24.7 | A | 2.5 | A | --- |
| 260472_at | 69 | A | 121.8 | P | 112.2 | A | 139.3 | P | --- |
| 260473_at | 8 | A | 8.5 | A | 8.1 | A | 8.7 | A | --- |
| 260474_at | 290.2 | P | 255.7 | P | 232.2 | P | 306.5 | P | --- |
| 260475_at | 77.7 | P | 259.9 | P | 373.7 | P | 229.1 | P | scpl31 |
| 260476_at | 209.6 | P | 172.9 | P | 223.4 | P | 202.6 | P | --- |
| 260477_at | 341.8 | P | 292.8 | P | 337.7 | P | 283.9 | P | --- |
| 260478_at | 33 | A | 17.9 | A | 23.9 | A | 37.3 | A | --- |
| 260479_at | 337.7 | P | 349.1 | P | 332.7 | P | 280.3 | P | --- |
| 260480_at | 800.3 | P | 772.5 | P | 687.7 | P | 747.7 | P | SNRK2.4 |
| 260481_at | 9482.7 | P | 13647.8 | P | 10473.8 | P | 13272.3 | P | ATFD1 |
| 260482_at | 2529.6 | P | 2931.3 | P | 2183.8 | P | 2765.8 | P | --- |
| 260483_at | 11.3 | A | 69.4 | P | 13.5 | A | 64.2 | P | MLO4 |

| | | | | | | | | | |
|---|---|---|---|---|---|---|---|---|---|
| 260484_at | 112.5 | A | 40.1 | A | 77.3 | A | 13.4 | A | --- |
| 260485_at | 0.8 | A | 10.7 | A | 1.9 | A | 4 | A | --- |
| 260486_at | 176.5 | P | 147.7 | P | 142.2 | P | 120.1 | P | --- |
| 260487_at | 1820 | P | 2261.9 | P | 2110 | P | 2228.5 | P | Y14 |
| 260488_at | 202.4 | P | 232.6 | P | 241.9 | P | 246.3 | P | TIFY2A |
| 260489_at | 664.7 | P | 1039.1 | P | 664.5 | P | 1022 | P | --- |
| 260490_at | 1378.1 | P | 1904 | P | 1637.5 | P | 1927.5 | P | CER5 |
| 260491_at | 35 | A | 56.6 | A | 60.8 | A | 97.2 | P | --- |
| 260492_at | 27.6 | A | 31.8 | A | 25.9 | A | 4.7 | A | PGAZAT |
| 260493_at | 175.6 | P | 122.2 | P | 155.3 | P | 181.7 | P | --- |
| 260494_at | 170.8 | P | 268.1 | P | 199 | P | 288.9 | P | --- |
| 260495_at | 28.8 | A | 28.8 | A | 2.1 | A | 7.5 | A | --- |
| 260496_at | 379.7 | P | 505.2 | P | 337.3 | P | 433.5 | P | --- |
| 260497_at | 8931.1 | P | 7550.1 | P | 9278.8 | P | 8420.3 | P | --- |
| 260498_at | 191.2 | P | 192.4 | P | 184.9 | P | 237.4 | P | --- |
| 260499_at | 322.7 | P | 382.2 | P | 300.8 | P | 334.5 | P | --- |
| 260500_at | 471.2 | P | 461.3 | P | 488.7 | P | 525.9 | P | --- |
| 260501_at | 346.3 | P | 429.4 | P | 300.5 | P | 425.4 | P | --- |
| 260502_at | 186.6 | P | 237.2 | P | 283.9 | P | 279.8 | P | AtTLP6 |
| 260503_at | 899.5 | P | 940.8 | P | 952.4 | P | 1073.1 | P | PAF2 |
| 260504_at | 26.2 | A | 3.8 | A | 19 | A | 32 | A | CYCA3;3 |
| 260505_at | 132 | P | 174.5 | P | 107.8 | P | 192.6 | P | CYCA3;4 |
| 260506_at | 507.6 | P | 418.5 | P | 518.5 | P | 514.8 | P | CYCA3;2 |
| 260507_at | 579.7 | P | 846 | P | 706 | P | 743.8 | P | WPP2 |
| 260508_at | 3454.5 | P | 3237.4 | P | 3648.1 | P | 3258.5 | P | GAMMA CA2 |
| 260509_at | 346.7 | P | 427.4 | P | 341 | P | 453.2 | P | NRAMP2 |
| 260510_at | 458.9 | P | 367.9 | P | 494 | P | 432.5 | P | --- |
| 260511_at | 1041.3 | P | 1139.4 | P | 1175.5 | P | 1096.9 | P | --- |
| 260512_at | 17.1 | A | 21.3 | P | 33.9 | P | 29.6 | M | --- |
| 260513_at | 23.8 | A | 21.2 | P | 32.9 | A | 2.1 | A | BGLU36 |
| 260514_at | 13.4 | A | 9.6 | A | 12.5 | A | 12.1 | A | --- |
| 260515_at | 179.3 | P | 189.4 | P | 196.2 | P | 165.9 | P | --- |
| 260516_at | 143.9 | P | 135.9 | P | 159.7 | P | 145.4 | P | --- |
| 260517_at | 41.1 | A | 45.3 | M | 47.9 | A | 45.5 | P | SPP1 |
| 260518_at | 28.5 | A | 11 | A | 26.2 | A | 6.1 | A | --- |
| 260519_at | 457.4 | P | 402.5 | P | 430.6 | P | 416.6 | P | --- |
| 260520_at | 243.2 | P | 283.3 | P | 256.4 | P | 256.3 | P | --- |
| 260521_at | 420.3 | P | 505.1 | P | 385.2 | P | 487.1 | P | VLN2 |
| 260522_x_at | 707.5 | P | 256.6 | P | 626.8 | P | 265.6 | P | --- |
| 260523_at | 608 | P | 736.9 | P | 616.1 | P | 686.4 | P | EMB2654 |
| 260524_at | 38.7 | P | 91 | P | 82.4 | P | 90.6 | P | --- |
| 260525_at | 634 | P | 672.2 | P | 807.4 | P | 714 | P | --- |
| 260526_at | 293.2 | P | 377.2 | P | 389 | P | 403.7 | P | --- |
| 260527_at | 31 | P | 32.3 | M | 28.8 | P | 11.2 | A | --- |
| 260528_at | 128.6 | P | 130.8 | P | 87.1 | M | 97.6 | P | WRKY23 |
| 260529_at | 8476.6 | P | 9875.4 | P | 8285 | P | 10168.9 | P | CP12-1 |
| 260530_at | 1808.8 | P | 2194.6 | P | 1765.6 | P | 2346.1 | P | --- |
| 260531_at | 373.7 | P | 665.9 | P | 413 | P | 670.8 | P | CER8 |
| 260532_at | 406.8 | P | 399.6 | P | 470.9 | P | 290.2 | P | --- |
| 260533_at | 213.3 | P | 135.6 | P | 280.4 | P | 166.9 | P | --- |
| 260534_at | 3.3 | A | 10.1 | A | 23.9 | A | 11.8 | A | --- |
| 260535_at | 0.5 | A | 0.4 | A | 2.2 | A | 2 | A | --- |
| 260536_at | 337.7 | P | 315.3 | P | 276.9 | P | 275.7 | P | ETFQO |
| 260537_at | 4.9 | A | 19.9 | P | 27.3 | P | 14.9 | A | --- |
| 260538_at | 4259.9 | P | 3879.3 | P | 4434.7 | P | 4230.5 | P | --- |
| 260539_at | 16.1 | A | 18.2 | A | 33.8 | A | 29.8 | P | --- |
| 260540_at | 238.8 | P | 357.4 | P | 196.3 | P | 346.4 | P | --- |
| 260541_at | 7589.3 | P | 3226.4 | P | 4667.9 | P | 3816.9 | P | --- |
| 260542_at | 2023.7 | P | 2243.3 | P | 2100.7 | P | 2023.4 | P | --- |
| 260543_at | 295.8 | P | 300.1 | P | 278 | P | 291.7 | P | INT1 |
| 260544_at | 326.6 | P | 295 | P | 269 | P | 235.7 | P | --- |
| 260545_at | 774.5 | P | 853.6 | P | 541.8 | P | 505.8 | P | ATGPX3 |
| 260546_at | 867.1 | P | 448.8 | P | 833.5 | P | 382.5 | P | ATTI2 |
| 260547_at | 5087.9 | P | 2896.5 | P | 5003.2 | P | 3923.8 | P | --- |
| 260548_at | 998.8 | P | 1072.6 | P | 1213.6 | P | 1131.1 | P | BIO2 |
| 260549_at | 1512 | P | 656.1 | P | 1270.1 | P | 643.2 | P | --- |
| 260550_at | 553.3 | P | 523.9 | P | 502.4 | P | 481 | P | --- |
| 260551_at | 7336.7 | P | 1744.9 | P | 3572 | P | 1655.9 | P | ATTI1 |
| 260552_at | 512.1 | P | 611.4 | P | 518.5 | P | 541.9 | P | GLX2-1 |
| 260553_at | 316.7 | P | 5.9 | A | 76.6 | A | 53.6 | A | --- |
| 260554_at | 857.1 | P | 937.4 | P | 772.5 | P | 787.9 | P | --- |
| 260555_at | 45.9 | P | 49 | P | 44.7 | P | 46.6 | P | --- |
| 260556_at | 456.1 | P | 1263.8 | P | 213.7 | P | 772.1 | P | --- |
| 260557_at | 37.7 | P | 79.5 | P | 92.3 | P | 88.2 | P | --- |
| 260558_at | 4.1 | A | 3 | A | 6.4 | A | 5.7 | A | --- |
| 260559_at | 24.2 | A | 2.9 | A | 3.6 | A | 1.6 | A | --- |
| 260560_at | 331 | P | 642.6 | P | 583.6 | P | 445.4 | P | --- |
| 260561_at | 8.4 | A | 19.5 | A | 27.8 | A | 38.4 | A | --- |
| 260562_at | 204.3 | P | 285.8 | P | 170.5 | P | 206.3 | P | --- |
| 260563_at | 61.4 | A | 57.3 | A | 63.7 | A | 65 | A | UGT74F1 |
| 260564_at | 454.3 | P | 504 | P | 479.4 | P | 494.4 | P | --- |
| 260565_at | 295.3 | P | 392.4 | P | 426.1 | P | 362.6 | P | --- |
| 260566_at | 2522.1 | P | 2321.2 | P | 2302.3 | P | 2354.4 | P | OASB |
| 260567_at | 4054 | P | 1513.9 | P | 3127.7 | P | 2038.9 | P | UGT74F2 |
| 260568_at | 452.9 | P | 1344.3 | P | 236 | P | 1200 | P | --- |
| 260569_at | 1009.8 | P | 926.1 | P | 962.5 | P | 989.7 | P | --- |
| 260570_at | 2231.8 | P | 2590.2 | P | 2889.4 | P | 2734.7 | P | SSI2 |
| 260571_at | 582.3 | P | 765.7 | P | 726.5 | P | 711.6 | P | ATMPK6 |
| 260572_at | 1250.4 | P | 1199.2 | P | 1469.4 | P | 1236.6 | P | --- |
| 260573_at | 16.7 | A | 36.1 | A | 53.5 | A | 20.9 | A | --- |
| 260574_at | 26.2 | A | 17.9 | A | 22.9 | A | 32.7 | A | --- |
| 260575_at | 92.9 | P | 81.5 | P | 108 | A | 80.9 | P | --- |
| 260576_at | 91.7 | P | 117.8 | P | 153.2 | P | 93.3 | P | --- |
| 260577_at | 3 | A | 3.2 | A | 17 | A | 3.9 | A | --- |
| 260578_at | 176.9 | M | 179.9 | P | 183.6 | P | 191.3 | P | --- |
| 260579_at | 5940.3 | P | 5309.4 | P | 4204.9 | P | 5273.8 | P | --- |
| 260580_at | 2044.7 | P | 2198.7 | P | 2080.5 | P | 2105.1 | P | --- |
| 260581_at | 120.3 | P | 49.4 | P | 84 | A | 52.8 | A | MYB2 |
| 260582_at | 10.4 | A | 23.5 | P | 34.5 | P | 36.2 | P | --- |
| 260583_x_at | 49.5 | A | 100.2 | A | 55.8 | A | 65.7 | A | --- |
| 260584_at | 4 | A | 32.9 | A | 64.7 | P | 32.8 | A | --- |
| 260585_at | 299.2 | P | 232.7 | P | 240.9 | P | 283.6 | P | EMB2777 |
| 260586_at | 989.2 | P | 1294 | P | 1068.8 | P | 1342.5 | P | --- |
| 260587_at | 3285.4 | P | 4661.4 | P | 2809.6 | P | 4095.7 | P | --- |
| 260588_at | 1827.5 | P | 1347.5 | P | 1474.6 | P | 1311.7 | P | AtTLP7 |
| 260589_at | 855.1 | P | 786.9 | P | 685.8 | P | 737 | P | --- |
| 260590_at | 1278.9 | P | 2265.3 | P | 1172.4 | P | 2235 | P | ATPPC1 |
| 260591_at | 0.5 | A | 0.9 | A | 1.2 | A | 0.4 | A | --- |
| 260592_at | 2770 | P | 2761.8 | P | 2336.4 | P | 2733.1 | P | CSLE1 |

| | | | | | | | | | |
|---|---|---|---|---|---|---|---|---|---|
| 260593_at | 113.4 | P | 122.6 | P | 157.8 | P | 127.4 | P | AHG2 |
| 260594_at | 185.3 | P | 210.5 | P | 211.6 | P | 160.9 | M | --- |
| 260595_at | 821.2 | P | 882.7 | P | 1232.5 | P | 818.7 | P | --- |
| 260596_at | 715.5 | P | 701.3 | P | 808.3 | P | 750.1 | P | TIM50 |
| 260597_at | 102.2 | P | 97.7 | P | 84.6 | A | 62.1 | P | --- |
| 260598_at | 347.7 | P | 244.1 | P | 321.7 | P | 254.2 | P | --- |
| 260599_at | 54.2 | P | 65.8 | P | 72.3 | A | 58.2 | P | CYP708A1 |
| 260600_at | 1.3 | A | 1.5 | A | 18.1 | A | 1.5 | A | --- |
| 260601_at | 483.1 | P | 861.4 | P | 696.2 | P | 914.5 | P | ZIP11 |
| 260602_at | 1448.3 | P | 1316.8 | P | 1046.1 | P | 1151.2 | P | ATSERAT2;1 |
| 260603_at | 314.2 | P | 450.9 | P | 380.7 | P | 411.6 | P | --- |
| 260604_at | 1937.9 | P | 2229.2 | P | 2728.9 | P | 2602.7 | P | --- |
| 260605_at | 565.9 | P | 472.1 | P | 660.3 | P | 473 | P | --- |
| 260606_at | 727.6 | P | 656.3 | P | 871.1 | P | 684.9 | P | --- |
| 260607_at | 66.3 | P | 40.3 | P | 44.6 | A | 25.4 | A | --- |
| 260608_at | 3.6 | A | 75.8 | P | 7 | A | 44.8 | A | --- |
| 260609_at | 73.8 | P | 35.4 | P | 80.4 | P | 60.3 | P | --- |
| 260610_at | 1000.8 | P | 967.9 | P | 1039.5 | P | 964.3 | P | IQD14 |
| 260611_at | 3.1 | A | 19.8 | A | 50.8 | A | 20.3 | A | --- |
| 260612_at | 13.3 | A | 32.7 | A | 13.9 | A | 20.1 | A | --- |
| 260613_at | 310.2 | P | 385 | P | 337.6 | P | 337.7 | P | --- |
| 260614_at | 586.3 | P | 540.2 | P | 436.9 | P | 637 | P | --- |
| 260615_at | 7899.6 | P | 7232 | P | 8541.2 | P | 7368.9 | P | --- |
| 260616_at | 2158 | P | 1790.3 | P | 2388.7 | P | 2002 | P | --- |
| 260617_at | 50.3 | P | 45.7 | P | 68 | A | 57.5 | P | --- |
| 260618_at | 959.1 | P | 915.5 | P | 927.9 | P | 880 | P | TCP3 |
| 260619_at | 2154.9 | P | 2340.4 | P | 2295.8 | P | 2542.5 | P | --- |
| 260620_at | 76.2 | P | 72.2 | P | 84.2 | P | 71.5 | P | --- |
| 260621_at | 25.5 | A | 19.2 | A | 15.7 | A | 13 | A | ACA5 |
| 260622_at | 261.9 | P | 217.6 | P | 180.5 | P | 249.9 | P | NF-YC10 |
| 260623_at | 7.9 | A | 40 | A | 31.4 | A | 59.4 | A | ATNRT2:1 |
| 260624_at | 20.6 | A | 5.4 | A | 5.2 | A | 3.2 | A | NRT2.2 |
| 260625_at | 110 | A | 130.7 | A | 157.1 | A | 98.6 | A | MOM |
| 260626_at | 663 | P | 621.3 | P | 596 | P | 649 | P | --- |
| 260627_at | 133.5 | P | 103.5 | P | 85.4 | P | 88.7 | P | --- |
| 260628_at | 3.9 | A | 2.8 | A | 2.7 | A | 2.5 | A | --- |
| 260629_at | 251.1 | P | 363.7 | P | 370.8 | P | 408.9 | P | --- |
| 260630_at | 15.6 | A | 37 | A | 31.2 | A | 36.8 | A | ALE1 |
| 260631_at | 91.4 | P | 74.9 | P | 79.5 | A | 40.6 | P | --- |
| 260632_at | 67.1 | A | 151.6 | P | 153.8 | P | 174.1 | P | STM |
| 260633_at | 30.1 | A | 62.5 | A | 11.2 | A | 41.5 | A | HT1 |
| 260634_at | 1.2 | A | 3.5 | A | 3.4 | A | 1.2 | A | --- |
| 260635_at | 678 | P | 586.7 | P | 996.5 | P | 547.1 | P | --- |
| 260636_at | 738.1 | P | 946.3 | P | 750.3 | P | 934.1 | P | ATCDS1 |
| 260637_at | 6928.1 | P | 10232.4 | P | 7236.5 | P | 9127.1 | P | ACO2 |
| 260638_at | 1564.5 | P | 1634.5 | P | 1338.7 | P | 1664.4 | P | --- |
| 260639_at | 39.4 | P | 79.7 | P | 81.9 | P | 33.5 | A | --- |
| 260640_at | 73.8 | P | 99.6 | P | 105.8 | P | 71.2 | P | --- |
| 260641_at | 180.9 | P | 134.2 | P | 163.8 | P | 146 | P | --- |
| 260642_at | 41.9 | A | 29.9 | P | 85.8 | M | 45.5 | P | --- |
| 260643_at | 13.7 | A | 5.1 | A | 5.7 | A | 5.1 | A | --- |
| 260644_at | 292.5 | P | 442.7 | P | 442.4 | P | 407.7 | P | --- |
| 260645_at | 307.3 | P | 443.9 | P | 337.9 | P | 408.3 | P | TTL1 |
| 260646_at | 24.9 | A | 21.4 | P | 53.4 | A | 28.7 | A | --- |
| 260647_at | 260.6 | P | 318.6 | P | 280.6 | P | 290.9 | P | --- |
| 260648_at | 133.3 | P | 195.8 | P | 160.4 | P | 160.2 | P | --- |
| 260649_at | 5.3 | A | 36.2 | A | 40 | A | 23.2 | A | ACA7 |
| 260650_at | 266.8 | P | 342.5 | P | 308.4 | P | 297.6 | P | TOM2B |
| 260651_at | 1498.9 | P | 1709.1 | P | 1291.8 | P | 1446.6 | P | --- |
| 260652_at | 512.9 | P | 575.5 | P | 564.7 | P | 540.8 | P | --- |
| 260653_at | 556.5 | P | 568.9 | P | 761 | P | 678.6 | P | PKp3 |
| 260654_at | 86.7 | P | 80.8 | P | 119.3 | P | 75.2 | P | --- |
| 260655_at | 4.5 | A | 35.5 | A | 41.7 | A | 57 | A | --- |
| 260656_at | 980.4 | P | 1508.1 | P | 869.9 | P | 1122 | P | --- |
| 260657_at | 4.4 | A | 11.8 | A | 3.6 | A | 3.4 | A | --- |
| 260658_at | 17.9 | A | 11.3 | A | 13.8 | A | 27 | A | --- |
| 260659_s_at | 1.5 | A | 2.1 | A | 2.1 | A | 1.7 | A | --- |
| 260660_at | 220.8 | P | 244.7 | P | 270.5 | P | 257.9 | P | --- |
| 260661_at | 1.4 | A | 5.2 | A | 4.8 | A | 3.2 | A | --- |
| 260662_at | 64.6 | P | 82.3 | P | 98.7 | P | 150.7 | P | --- |
| 260663_at | 529.4 | P | 446.5 | P | 605 | P | 457.8 | P | NFD5 |
| 260664_at | 46.6 | A | 39.3 | A | 62.8 | P | 56.5 | P | ATRL5 |
| 260665_at | 846 | P | 1026.7 | P | 814.4 | P | 1050.3 | P | --- |
| 260666_at | 308.8 | P | 414.8 | P | 279 | P | 413.6 | P | PARVUS |
| 260667_at | 259.2 | P | 373 | P | 295.7 | P | 385.1 | P | KCS4 |
| 260668_at | 2.7 | A | 21.2 | A | 1.2 | A | 14 | A | --- |
| 260669_at | 128 | P | 116.3 | P | 117 | A | 148 | P | --- |
| 260670_at | 357 | P | 305.3 | P | 303.8 | P | 320.1 | P | NFD5 |
| 260671_at | 427.7 | P | 527.5 | P | 311.2 | P | 469.5 | P | --- |
| 260672_at | 162.5 | P | 119 | P | 136.7 | P | 188.5 | P | --- |
| 260673_at | 168.8 | P | 351.6 | P | 241.6 | P | 365.1 | P | --- |
| 260674_at | 221.7 | P | 259.4 | P | 257.4 | P | 300.2 | P | --- |
| 260675_at | 404.4 | P | 464.3 | P | 365.1 | P | 460.7 | P | --- |
| 260676_at | 1027.2 | P | 1444.8 | P | 1380.3 | P | 1351 | P | --- |
| 260677_at | 180.3 | P | 218.2 | P | 207.7 | P | 204 | P | RNL |
| 260678_at | 320.9 | P | 321 | P | 393.6 | P | 318 | P | --- |
| 260679_at | 333.8 | P | 388.2 | P | 326.4 | P | 353.3 | P | --- |
| 260680_s_at | 115.6 | P | 152.3 | P | 76.5 | P | 153 | P | --- |
| 260681_at | 51.8 | P | 83 | P | 90.1 | P | 65.9 | P | TPST |
| 260682_at | 682.3 | P | 645.1 | P | 651.9 | P | 677.1 | P | --- |
| 260683_at | 437.5 | P | 308.2 | P | 399.5 | P | 382.1 | P | HLL |
| 260684_at | 161 | A | 257.6 | P | 158.9 | A | 219.8 | P | NF-YA8 |
| 260685_at | 1054.6 | P | 1044 | P | 1169.2 | P | 1152.1 | P | GLYR2 |
| 260686_at | 575.3 | P | 471.2 | P | 403.9 | P | 492.2 | P | --- |
| 260687_at | 791.8 | P | 696.8 | P | 822 | P | 831.6 | P | ATTIM23-1 |
| 260688_at | 245.4 | P | 173.2 | P | 201.1 | P | 186.8 | P | --- |
| 260689_at | 114.2 | A | 146.5 | A | 204.4 | A | 138.8 | A | --- |
| 260690_at | 506.2 | P | 432.9 | P | 481.8 | P | 515 | P | NHL8 |
| 260691_s_at | 1 | A | 16 | M | 27.7 | A | 5.9 | A | --- |
| 260692_at | 53.7 | P | 52.1 | P | 61.9 | P | 53.7 | P | --- |
| 260693_at | 83.4 | P | 308.8 | P | 79 | A | 227.2 | P | NRT1.5 |
| 260694_at | 1.6 | A | 2.4 | A | 5.7 | A | 21.5 | A | --- |
| 260695_at | 493.5 | P | 483.1 | P | 553.8 | P | 431 | P | ESP3 |
| 260696_at | 352.6 | P | 429.7 | P | 357.4 | P | 439.5 | P | --- |
| 260697_at | 317.8 | P | 221 | P | 256.9 | P | 223.8 | P | --- |
| 260698_at | 1.8 | A | 16.5 | A | 16.7 | A | 5 | A | --- |
| 260699_at | 8.1 | A | 33.1 | A | 29.6 | P | 34.8 | A | ATMKK10 |
| 260700_at | 243.4 | P | 326.8 | P | 281.9 | P | 285.6 | P | --- |
| 260701_at | 242.7 | P | 384.2 | P | 330.5 | P | 247.9 | P | ATHSFA1D |

| | | | | | | | | | |
|---|---|---|---|---|---|---|---|---|---|
| 260702_at | 3.1 | A | 1.7 | A | 2.8 | A | 3.1 | A | --- |
| 260703_at | 30.6 | P | 22.8 | A | 24.6 | A | 23.4 | A | ATSYP24 |
| 260704_at | 6127.2 | P | 6314 | P | 6543.4 | P | 7070.3 | P | --- |
| 260705_at | 1680.1 | P | 1715.3 | P | 1757.7 | P | 1585.2 | P | TOM2A |
| 260706_at | 225.5 | P | 89 | M | 125.4 | P | 38.9 | A | AOX1D |
| 260707_s_at | 743.6 | P | 807.3 | P | 753.4 | P | 787.5 | P | --- |
| 260708_at | 418.2 | P | 355.3 | P | 411.9 | P | 371.8 | P | --- |
| 260709_at | 1256.2 | P | 1098.7 | P | 1169.7 | P | 1082 | P | ATNAP6 |
| 260710_at | 3.5 | A | 22.6 | A | 36.8 | A | 45.2 | A | --- |
| 260711_at | 360.4 | P | 361 | P | 375.1 | P | 344.6 | P | MYA1 |
| 260712_at | 401.3 | P | 423.3 | P | 313.3 | P | 410 | P | HAB2 |
| 260713_at | 1.8 | A | 1.1 | A | 2.1 | A | 1.5 | A | --- |
| 260714_at | 4825.6 | P | 3957.6 | P | 4695.1 | P | 4558 | P | CPN10 |
| 260715_at | 416.3 | P | 324.6 | P | 411.9 | P | 239.8 | P | --- |
| 260716_at | 2 | A | 3.6 | A | 7.2 | A | 4.9 | A | ATPER1 |
| 260717_at | 87.6 | P | 78 | P | 79.8 | P | 86.3 | P | --- |
| 260718_at | 391.5 | P | 545.8 | P | 368.6 | P | 486.2 | P | ECT7 |
| 260719_at | 398.5 | P | 427.3 | P | 393.1 | P | 402.5 | P | --- |
| 260720_at | 5.2 | A | 33.9 | A | 23.5 | A | 34.5 | A | --- |
| 260721_at | 2.7 | A | 4.7 | A | 7.5 | A | 3.8 | A | --- |
| 260722_at | 113.7 | P | 128.2 | P | 114.6 | P | 92.7 | P | --- |
| 260723_at | 8 | A | 8.2 | A | 3.7 | A | 1.8 | A | --- |
| 260724_at | 617.2 | P | 698.9 | P | 597.7 | P | 651.6 | P | --- |
| 260725_at | 600.3 | P | 571 | P | 623.6 | P | 636.1 | P | --- |
| 260726_at | 565 | P | 576.5 | P | 529.3 | P | 549.2 | P | --- |
| 260727_at | 91.6 | P | 123.3 | P | 130.2 | P | 221.6 | P | --- |
| 260728_at | 671.2 | P | 834.6 | P | 609.6 | P | 761.2 | P | --- |
| 260729_at | 199.9 | P | 229.9 | P | 244.5 | P | 200.4 | P | KU80 |
| 260730_at | 9794.3 | P | 8063 | P | 11054.5 | P | 9300.5 | P | mtLPD1 |
| 260731_at | 294.5 | P | 325.3 | P | 254.8 | P | 272.6 | P | --- |
| 260732_at | 144.3 | P | 168.3 | P | 217.1 | P | 179 | P | --- |
| 260733_at | 80.6 | P | 92.4 | P | 105.8 | P | 88.4 | P | --- |
| 260734_at | 23.7 | A | 65.9 | P | 35.7 | A | 64.3 | A | --- |
| 260735_at | 51.6 | A | 99.1 | P | 40.5 | A | 61.4 | A | --- |
| 260736_at | 130.8 | P | 65.2 | P | 112.9 | P | 111.8 | P | --- |
| 260737_at | 4.3 | A | 40.7 | A | 44.3 | A | 25.3 | A | --- |
| 260738_at | 515.2 | P | 471.8 | P | 510.3 | P | 484.7 | P | --- |
| 260739_at | 336.2 | P | 212.3 | P | 386.4 | P | 264.2 | P | scpl50 |
| 260740_at | 215.5 | P | 192.2 | P | 151.5 | P | 185.4 | P | ATQSOX1 |
| 260741_at | 7.2 | A | 3.8 | A | 6.7 | A | 4.1 | A | --- |
| 260742_at | 14.5 | A | 4.7 | A | 23.4 | A | 34.3 | P | IAA34 |
| 260743_at | 4 | A | 3.3 | A | 6.8 | A | 4.1 | A | --- |
| 260744_at | 220.7 | P | 527.5 | P | 190.4 | P | 173.3 | P | --- |
| 260745_at | 10183.8 | P | 10531.7 | P | 14976 | P | 9781.3 | P | ATGSTU20 |
| 260746_at | 15558.4 | P | 12036.6 | P | 17049.6 | P | 14644 | P | ATGSTU19 |
| 260747_at | 501 | P | 461 | P | 525.2 | P | 443.7 | P | --- |
| 260748_at | 1.3 | A | 24.1 | A | 2.1 | A | 3.6 | A | --- |
| 260749_at | 24.9 | A | 60.7 | A | 81.6 | A | 39.8 | A | --- |
| 260750_at | 21 | A | 10.5 | A | 22.8 | A | 1.8 | A | --- |
| 260751_at | 509.4 | P | 463.7 | P | 475 | P | 418.5 | P | SCD1 |
| 260752_at | 4.5 | A | 8.4 | A | 28.5 | A | 34.2 | A | --- |
| 260753_at | 60.8 | P | 223.7 | P | 117.3 | P | 236.1 | P | --- |
| 260754_at | 42.3 | A | 30.3 | A | 8 | A | 4.7 | A | --- |
| 260755_at | 17 | A | 21 | P | 29.7 | A | 20.6 | P | --- |
| 260756_at | 257.6 | P | 238.8 | P | 324.4 | P | 222.5 | P | --- |
| 260757_at | 14.3 | A | 12.6 | A | 37.7 | A | 18.3 | A | --- |
| 260758_at | 25.7 | A | 37.7 | A | 79.5 | P | 51 | A | AtGH9C1 |
| 260759_at | 85 | P | 128.1 | P | 111.3 | P | 102.4 | P | --- |
| 260760_at | 100.7 | P | 84.7 | P | 120.8 | P | 108.3 | P | --- |
| 260761_at | 38.9 | P | 33.7 | P | 57.4 | M | 24 | A | --- |
| 260762_at | 24.8 | A | 5.8 | A | 49.1 | A | 12.1 | A | --- |
| 260763_at | 4.9 | A | 4.8 | A | 5.8 | A | 19.4 | A | --- |
| 260764_at | 256.4 | P | 273.6 | P | 287.2 | P | 193.6 | P | --- |
| 260765_at | 2372.7 | P | 2402.4 | P | 2558.1 | P | 2425.4 | P | ACT8 |
| 260766_at | 196.1 | P | 168.8 | P | 171.8 | P | 173 | P | --- |
| 260767_s_at | 9813.6 | P | 8843.1 | P | 8630.2 | P | 8689.5 | P | --- |
| 260768_at | 229.2 | P | 161.9 | P | 183.5 | P | 188 | P | --- |
| 260769_at | 503.8 | P | 440.8 | P | 553.3 | P | 482.6 | P | --- |
| 260770_at | 36.3 | P | 79.4 | P | 64.1 | P | 69.9 | P | --- |
| 260771_at | 79.7 | P | 36.8 | A | 76.3 | P | 31 | A | WNK7 |
| 260772_at | 528 | P | 746.4 | P | 368.4 | P | 570.5 | P | --- |
| 260773_at | 2.9 | A | 22.3 | A | 27 | A | 22.1 | A | ATGA2OX1 |
| 260774_at | 160.2 | P | 363.3 | P | 231.1 | P | 257.3 | P | --- |
| 260775_at | 1243.6 | P | 1761 | P | 1211.5 | P | 1628 | P | GRF2 |
| 260776_at | 280.6 | P | 183.4 | P | 199.8 | P | 151.4 | P | --- |
| 260777_at | 163.5 | P | 235.5 | P | 235 | P | 203.9 | P | --- |
| 260778_at | 22.8 | A | 67.5 | A | 39.2 | A | 50.4 | P | --- |
| 260779_at | 306.1 | P | 237.6 | P | 319.6 | P | 257.2 | P | --- |
| 260780_at | 1353 | P | 1176.3 | P | 1421.2 | P | 1147 | P | TWN2 |
| 260781_at | 1021.8 | P | 747.3 | P | 1204.3 | P | 743 | P | DECOY |
| 260782_at | 1343.7 | P | 1477.4 | P | 1558.1 | P | 1342.1 | P | --- |
| 260783_at | 117.6 | P | 187 | P | 88.5 | P | 181.6 | P | ORA59 |
| 260784_at | 540.7 | P | 321 | P | 467.4 | P | 375.3 | P | ATMYB13 |
| 260785_at | 2381.7 | P | 2070.7 | P | 1768.7 | P | 1910.9 | P | --- |
| 260786_s_at | 1137.7 | P | 1060.4 | P | 1395.6 | P | 1135.9 | P | MEE5 |
| 260787_at | 314.5 | P | 241.6 | P | 370.8 | P | 253.6 | P | GTE4 |
| 260788_at | 44.2 | A | 35.3 | A | 42.7 | A | 29.9 | A | --- |
| 260789_s_at | 1908.3 | P | 2012.9 | P | 1393 | P | 1815.6 | P | ACX3 |
| 260790_at | 256.5 | P | 297 | P | 291.7 | P | 323.9 | P | --- |
| 260791_at | 26.1 | A | 16.7 | A | 45.2 | P | 25.4 | A | --- |
| 260792_at | 131 | A | 134.5 | A | 145.6 | A | 110 | M | --- |
| 260793_at | 1398 | P | 1663 | P | 1732.3 | P | 1947.8 | P | --- |
| 260794_at | 392 | P | 523.3 | P | 414.5 | P | 508.1 | P | --- |
| 260795_at | 12 | A | 3.7 | A | 3.2 | A | 20 | A | CLE3 |
| 260796_at | 49 | A | 35 | A | 78.4 | A | 15.5 | A | ATGSTU21 |
| 260797_at | 1.9 | A | 1.7 | A | 2.2 | A | 1.3 | A | NCED9 |
| 260798_at | 389.4 | P | 444.1 | P | 434 | P | 412.6 | P | --- |
| 260799_at | 203.1 | P | 175.7 | P | 148.8 | P | 187.2 | P | AtUGT85A4 |
| 260800_at | 898.6 | P | 1010.6 | P | 942.9 | P | 943.8 | P | TSD2 |
| 260801_at | 31.5 | A | 102.1 | P | 66 | P | 137.1 | P | --- |
| 260802_at | 31 | A | 4.1 | A | 38 | A | 4.3 | A | --- |
| 260803_at | 35.6 | P | 48 | M | 80.7 | P | 23.4 | A | ATGSTU22 |
| 260804_at | 128.9 | P | 126.2 | P | 35.8 | A | 189.6 | P | --- |
| 260805_at | 10.7 | A | 46.1 | P | 25.5 | A | 16.1 | A | ATGSTU23 |
| 260806_at | 102.3 | P | 107.4 | P | 127.4 | P | 117.8 | P | --- |
| 260807_at | 188.9 | P | 234.5 | P | 207.2 | P | 168.3 | P | --- |
| 260808_at | 34.5 | A | 57.3 | A | 21.2 | A | 21.1 | A | --- |
| 260809_at | 3.8 | A | 23.4 | A | 29.3 | A | 12 | A | --- |
| 260810_s_at | 4.1 | A | 1.3 | A | 4.6 | A | 11.3 | A | --- |

| | | | | | | | | | |
|---|---|---|---|---|---|---|---|---|---|
| 260811_s_at | 18.1 | A | 22.8 | A | 20.9 | A | 9.6 | A | --- |
| 260812_at | 102 | P | 85.4 | P | 108.3 | P | 93.9 | P | --- |
| 260813_at | 861 | P | 724 | P | 727 | P | 636.3 | P | VIP1 |
| 260814_at | 2011.1 | P | 1920.2 | P | 1847.5 | P | 1980.9 | P | emb1075 |
| 260815_at | 2731.1 | P | 2368.3 | P | 2560.2 | P | 2345.3 | P | TIC110 |
| 260816_at | 42.5 | P | 52.2 | P | 66.7 | A | 37.8 | P | --- |
| 260817_at | 731.7 | P | 639.3 | P | 844.5 | P | 660 | P | --- |
| 260818_at | 1631.9 | P | 1922.2 | P | 1668.7 | P | 1985 | P | --- |
| 260819_at | 114 | P | 191.5 | P | 122.6 | P | 184.7 | P | AtbZIP52 |
| 260820_at | 202.2 | P | 233 | P | 249.2 | P | 245.1 | P | --- |
| 260821_at | 894.8 | P | 797.8 | P | 991.6 | P | 849.6 | P | CRTISO |
| 260822_at | 314.9 | P | 296.8 | P | 395.1 | P | 346.4 | P | --- |
| 260823_at | 117.2 | P | 124.3 | P | 120.1 | P | 115.1 | P | DRIP1 |
| 260824_at | 530.8 | P | 348.9 | P | 522.4 | P | 352.4 | P | --- |
| 260825_at | 26.2 | A | 14.5 | A | 40.1 | A | 17.2 | A | --- |
| 260826_at | 77.9 | P | 56.5 | P | 120.1 | P | 77.1 | P | --- |
| 260827_at | 134.5 | P | 117.2 | P | 167.7 | P | 136.1 | P | AT1G06750 |
| 260828_at | 4.7 | A | 5.1 | A | 10.4 | A | 3.7 | A | --- |
| 260829_s_at | 2010.9 | P | 1628.3 | P | 1870.8 | P | 1492 | P | --- |
| 260830_at | 1620.3 | P | 2124.3 | P | 1505.7 | P | 2111.4 | P | --- |
| 260831_at | 110.7 | P | 209.7 | P | 290.2 | P | 247.6 | P | --- |
| 260832_at | 283.4 | P | 233.8 | P | 221.3 | P | 214.6 | P | GAUT6 |
| 260833_at | 134.9 | P | 108.1 | P | 139.3 | P | 94.7 | P | --- |
| 260834_at | 235.8 | P | 208.2 | P | 189.9 | P | 221.4 | P | --- |
| 260835_at | 1131.5 | P | 1224.8 | P | 1319.2 | P | 1257.9 | P | --- |
| 260836_at | 0.1 | A | 7 | A | 1.8 | A | 3 | A | --- |
| 260837_at | 3059.2 | P | 3345.4 | P | 3020.5 | P | 3119.4 | P | --- |
| 260838_at | 9.3 | A | 30.8 | P | 26.8 | P | 29.6 | A | --- |
| 260839_at | 564.9 | P | 662.1 | P | 523.4 | P | 726.8 | P | --- |
| 260840_at | 256.1 | P | 186.9 | P | 240.2 | P | 190.1 | P | --- |
| 260841_at | 450.3 | P | 172 | P | 170.2 | P | 115.1 | P | --- |
| 260842_at | 2524.6 | P | 3031.8 | P | 2774.3 | P | 2689.4 | P | ATS9 |
| 260843_at | 130 | P | 135.9 | P | 87.8 | P | 152.8 | P | --- |
| 260844_at | 895.4 | P | 939.5 | P | 1016.9 | P | 948.9 | P | PEX7 |
| 260845_at | 15 | A | 23 | A | 4.6 | A | 23.1 | A | --- |
| 260846_at | 11.1 | A | 16.8 | A | 5 | A | 2.7 | A | --- |
| 260847_s_at | 6382.9 | P | 5420.4 | P | 4824.5 | P | 4239.8 | P | AlaAT1 |
| 260848_at | 6.8 | A | 35.3 | A | 30.7 | A | 17.6 | A | sks8 |
| 260849_at | 2 | A | 3.5 | A | 5.2 | A | 3.5 | A | sks7 |
| 260850_at | 32.1 | P | 35.2 | A | 39.5 | A | 38.8 | A | --- |
| 260851_at | 2 | A | 151.7 | P | 53.8 | A | 113.7 | A | --- |
| 260852_at | 882.6 | P | 903.6 | P | 854.7 | P | 868.7 | P | --- |
| 260853_at | 1.2 | A | 1.9 | A | 23.9 | A | 1.7 | A | --- |
| 260854_at | 3.7 | A | 41.6 | A | 30.9 | A | 16.8 | A | LEC1 |
| 260855_at | 427.7 | P | 479.6 | P | 202.5 | P | 434.2 | P | --- |
| 260856_at | 274.5 | P | 296.3 | P | 229.3 | P | 242.8 | P | --- |
| 260857_at | 462.2 | P | 634.4 | P | 485.4 | P | 604 | P | LYM1 |
| 260858_at | 117 | P | 97.8 | P | 94.3 | P | 126.8 | P | --- |
| 260859_at | 35 | A | 26.5 | A | 61.8 | A | 29.9 | A | scpl44 |
| 260860_at | 22.1 | A | 31.6 | A | 36.8 | P | 53.5 | P | --- |
| 260861_at | 1 | A | 9 | A | 22.6 | A | 8.5 | A | --- |
| 260862_at | 479.1 | P | 522.4 | P | 501.5 | P | 507.3 | P | SEU |
| 260863_at | 4.6 | A | 1.3 | A | 2.3 | A | 3 | A | --- |
| 260864_at | 6 | A | 21.8 | A | 1.7 | A | 3 | A | --- |
| 260865_at | 47.9 | P | 11.7 | A | 13 | A | 57.6 | A | --- |
| 260866_at | 3.5 | A | 2.8 | A | 30.9 | A | 3 | A | --- |
| 260867_at | 302.1 | P | 525.8 | P | 514.8 | P | 513.7 | P | TED6 |
| 260868_at | 751.2 | P | 557.5 | P | 624.2 | P | 569.6 | P | --- |
| 260869_at | 1.3 | A | 37.1 | A | 2.6 | A | 14 | A | --- |
| 260870_at | 1376.7 | P | 1382.1 | P | 1113.5 | P | 1333 | P | ATRAB18 |
| 260871_at | 621.2 | P | 731.2 | P | 630.2 | P | 687.9 | P | --- |
| 260872_at | 690.7 | P | 951.3 | P | 787.5 | P | 929.3 | P | --- |
| 260873_at | 86 | P | 114.2 | P | 84.5 | P | 100.8 | P | --- |
| 260874_at | 54.2 | P | 40.3 | P | 58.2 | P | 41.3 | P | emb2170 |
| 260875_at | 957.6 | P | 631.1 | P | 925.2 | P | 686.4 | P | SKP2A |
| 260876_at | 1149.6 | P | 1072.5 | P | 1265.3 | P | 987 | P | --- |
| 260877_at | 834.9 | P | 1320.3 | P | 666.6 | P | 1018.4 | P | --- |
| 260878_at | 336 | P | 367.4 | P | 312.9 | P | 325.6 | P | SCL1 |
| 260879_at | 233.3 | P | 238.8 | P | 218 | P | 283.9 | P | --- |
| 260880_at | 490.2 | P | 583.3 | P | 579.6 | P | 600.8 | P | --- |
| 260881_at | 58 | P | 223.6 | P | 35.5 | A | 116.1 | P | --- |
| 260882_at | 126 | P | 130.3 | P | 153.5 | P | 116.3 | P | WRKY65 |
| 260883_at | 21.4 | A | 86.2 | P | 13.9 | A | 55.1 | P | --- |
| 260884_at | 138.3 | P | 289.5 | P | 194.7 | P | 187.3 | P | --- |
| 260885_at | 41.5 | P | 29.3 | P | 60.4 | P | 52.7 | P | CIPK18 |
| 260886_at | 25.7 | A | 42.8 | A | 26.1 | A | 37.7 | A | --- |
| 260887_at | 95.7 | P | 106.5 | P | 119.3 | P | 129.8 | P | --- |
| 260888_at | 15.1 | A | 3.6 | A | 8 | A | 16.1 | A | --- |
| 260889_at | 457.3 | P | 570.7 | P | 609.9 | P | 576.6 | P | --- |
| 260890_at | 4.3 | A | 21.4 | A | 23.6 | A | 2.6 | A | --- |
| 260891_at | 4.6 | A | 4 | A | 2.7 | A | 2.8 | A | --- |
| 260892_at | 13.5 | A | 16.8 | A | 7 | A | 34.8 | A | --- |
| 260893_at | 25.1 | A | 22.8 | A | 50 | A | 11.5 | A | --- |
| 260894_at | 260.1 | P | 357.3 | P | 262.1 | P | 279.8 | P | --- |
| 260895_at | 3363.6 | P | 2596.2 | P | 3259.1 | P | 3088.2 | P | --- |
| 260896_at | 596 | P | 644.8 | P | 558.8 | P | 539.8 | P | --- |
| 260897_at | 420.8 | P | 197.3 | P | 353.2 | P | 258.6 | P | ERD2 |
| 260898_at | 6092.8 | P | 7221.1 | P | 6779.4 | P | 6462.4 | P | --- |
| 260899_at | 486.6 | P | 488.2 | P | 440.9 | P | 540.5 | P | --- |
| 260900_s_at | 784.2 | P | 358 | P | 361.5 | P | 371.9 | P | --- |
| 260901_at | 30.7 | A | 9.9 | A | 11.2 | A | 17.3 | A | YUC11 |
| 260902_at | 3932.4 | P | 3470.5 | P | 4707.9 | P | 3291.9 | P | --- |
| 260903_at | 84.9 | P | 41 | A | 56.6 | A | 63.2 | P | --- |
| 260904_at | 39 | P | 72.2 | P | 35.1 | A | 93.5 | P | NIMIN1 |
| 260905_at | 27.7 | A | 30.5 | A | 16.4 | A | 21.1 | A | --- |
| 260906_at | 1.7 | A | 1.3 | A | 5.1 | A | 1.5 | A | --- |
| 260907_at | 49.8 | A | 51.4 | A | 53.5 | A | 28.3 | A | --- |
| 260908_at | 2.8 | A | 4.2 | A | 3.1 | A | 3.2 | A | MEA |
| 260909_at | 38.9 | A | 1.4 | A | 28.7 | A | 21.4 | A | --- |
| 260910_at | 248.4 | P | 339.1 | P | 322.5 | P | 280 | P | IMPA-6 |
| 260911_at | 2.5 | A | 2.1 | A | 2.4 | A | 11.4 | A | --- |
| 260912_at | 6545.2 | P | 6082.3 | P | 6696.4 | P | 6136 | P | CLPP5 |
| 260913_at | 8118.8 | P | 6687.3 | P | 7985.2 | P | 7056.5 | P | SAM1 |
| 260914_at | 57.4 | A | 169.2 | P | 100.9 | P | 152 | P | BXL2 |
| 260915_at | 108.1 | P | 145.1 | P | 129.7 | P | 104.5 | P | --- |
| 260916_at | 2413.4 | P | 2530.4 | P | 1701.4 | P | 2487.1 | P | --- |
| 260917_at | 17.2 | A | 58.2 | P | 27 | A | 38 | A | --- |
| 260918_at | 37 | A | 38.7 | A | 29.9 | A | 25.1 | A | --- |
| 260919_at | 957.3 | P | 656.7 | P | 406.8 | P | 479.6 | P | --- |

| | | | | | | | | | |
|---|---|---|---|---|---|---|---|---|---|
| 260920_at | 16.7 | A | 6.4 | A | 24 | A | 5.2 | A | --- |
| 260921_at | 26.9 | A | 53.4 | A | 41.5 | A | 69.5 | A | --- |
| 260922_at | 314 | P | 445.4 | P | 347.7 | P | 479.3 | P | --- |
| 260923_at | 842.2 | P | 815 | P | 966.1 | P | 819.4 | P | --- |
| 260924_at | 349.5 | P | 655.5 | P | 385.9 | P | 611.3 | P | --- |
| 260925_at | 37.1 | A | 40.8 | A | 6.3 | A | 32.6 | A | --- |
| 260926_at | 2.7 | A | 18.6 | A | 4.5 | A | 2.6 | A | GLTP2 |
| 260927_at | 1073.1 | P | 1130.8 | P | 1051.7 | P | 1001.5 | P | CAT9 |
| 260928_at | 110.2 | P | 104.4 | P | 108.7 | P | 81.8 | P | GATL5 |
| 260929_at | 238 | P | 227.1 | P | 208.5 | P | 185 | P | TAF13 |
| 260930_at | 83.7 | P | 73.7 | P | 79.6 | M | 42.2 | A | --- |
| 260931_at | 14.2 | A | 6.7 | A | 45.3 | A | 6.8 | A | --- |
| 260932_s_at | 26.3 | A | 27 | A | 5.8 | A | 13.5 | A | PGP12 |
| 260933_at | 508.4 | P | 212.1 | P | 313.8 | P | 197.4 | P | --- |
| 260934_at | 20.6 | A | 2.1 | A | 7.6 | A | 4.2 | A | --- |
| 260935_at | 155.5 | P | 109.3 | P | 225.8 | P | 142.4 | P | --- |
| 260936_at | 217.4 | P | 117.5 | P | 116.9 | P | 127.8 | P | --- |
| 260937_at | 459.9 | P | 449.9 | P | 499.2 | P | 405 | P | --- |
| 260938_at | 271 | P | 363.6 | P | 381 | P | 342.1 | P | --- |
| 260939_at | 70.6 | P | 96.7 | P | 116.5 | P | 98.4 | P | --- |
| 260940_at | 2094.5 | P | 1907.9 | P | 1838.4 | P | 2073.5 | P | --- |
| 260941_at | 20.1 | P | 56.3 | P | 44.6 | P | 33.1 | P | --- |
| 260942_s_at | 2.1 | A | 2.2 | A | 1.5 | A | 1.2 | A | --- |
| 260943_at | 1703.5 | P | 1296.2 | P | 954.3 | P | 1149.1 | P | ATTRX5 |
| 260944_at | 250.4 | P | 548.9 | P | 178.5 | P | 363.5 | P | BGAL5 |
| 260945_at | 49.6 | A | 49.6 | A | 45.7 | A | 47.8 | A | --- |
| 260946_at | 464.4 | P | 413.6 | P | 525.8 | P | 484.1 | P | --- |
| 260947_at | 24.5 | A | 7.1 | A | 9.3 | A | 25.6 | A | --- |
| 260948_at | 44.7 | P | 65.9 | P | 74.9 | P | 101.9 | P | --- |
| 260949_at | 788.1 | P | 833 | P | 654.1 | P | 715.9 | P | SKIP16 |
| 260950_s_at | 61.1 | P | 41.2 | A | 23.2 | A | 57 | P | --- |
| 260951_at | 100 | P | 72.7 | P | 74.4 | P | 82.8 | P | --- |
| 260952_at | 36.1 | A | 18.2 | A | 45.8 | A | 54.8 | P | --- |
| 260953_at | 277.4 | P | 307.7 | P | 265.6 | P | 272.5 | P | --- |
| 260954_at | 651.2 | P | 498.9 | P | 529.9 | P | 470.7 | P | GLX2-4 |
| 260955_at | 1491 | P | 1333.8 | P | 1788.7 | P | 1964.4 | P | --- |
| 260956_at | 2393.8 | P | 3639.7 | P | 2174.2 | P | 2928.2 | P | STO |
| 260957_at | 47.4 | A | 111.2 | P | 56.8 | A | 98 | P | ADS1 |
| 260958_at | 392.9 | P | 439.6 | P | 426.2 | P | 429.7 | P | --- |
| 260959_at | 126.2 | P | 125.7 | P | 115.6 | P | 131.1 | P | --- |
| 260960_at | 2.3 | A | 2.4 | A | 23.3 | A | 29.5 | A | --- |
| 260961_at | 385.9 | P | 328.9 | P | 408.5 | P | 357.2 | P | --- |
| 260962_at | 4.5 | A | 17.4 | A | 23.4 | A | 14.5 | A | PME7 |
| 260963_at | 39.5 | A | 38.1 | A | 41.6 | A | 11.4 | A | --- |
| 260964_at | 487.7 | P | 399.7 | P | 397.8 | P | 384.4 | P | ATUBC2-1 |
| 260965_s_at | 1.9 | A | 0.9 | A | 2.4 | A | 1.3 | A | --- |
| 260966_at | 81.3 | P | 97 | P | 146.6 | P | 109.4 | P | RPS5 |
| 260967_at | 1953.4 | P | 1673 | P | 1804.3 | P | 1644.7 | P | --- |
| 260968_at | 955.5 | P | 1375.3 | P | 1173.4 | P | 1147.6 | P | --- |
| 260969_at | 1562.4 | P | 1407.7 | P | 1926.5 | P | 1552.5 | P | ATBETAFRUCT4 |
| 260970_at | 102.7 | P | 78.7 | P | 118.8 | P | 104.7 | P | --- |
| 260971_at | 7.9 | A | 29.1 | A | 52.5 | A | 34 | A | --- |
| 260972_at | 213.6 | P | 318.7 | P | 307.6 | P | 309.2 | P | --- |
| 260973_at | 51.7 | P | 77.7 | P | 47.8 | A | 65.9 | P | --- |
| 260974_at | 376.2 | P | 442.1 | P | 455.2 | P | 547.7 | P | --- |
| 260975_at | 336 | P | 451.2 | P | 344 | P | 398 | P | --- |
| 260976_at | 274.3 | P | 260.4 | P | 271.4 | P | 281 | P | CID8 |
| 260977_at | 7.3 | A | 0.6 | A | 0.7 | A | 2.3 | A | --- |
| 260978_at | 31 | A | 101.4 | P | 31.5 | A | 89.1 | P | --- |
| 260979_at | 221.9 | P | 327.2 | P | 255.3 | P | 293.9 | P | ATMPK18 |
| 260980_at | 34.4 | A | 11.1 | A | 62 | A | 28.3 | A | --- |
| 260981_at | 694.7 | P | 555.3 | P | 739 | P | 678.4 | P | --- |
| 260982_at | 489.7 | P | 989.9 | P | 600.4 | P | 936 | P | --- |
| 260983_at | 1302 | P | 668.2 | P | 752 | P | 588.9 | P | --- |
| 260984_at | 1008.2 | P | 895.2 | P | 994 | P | 962 | P | --- |
| 260985_at | 758.8 | P | 1005.6 | P | 769 | P | 961.3 | P | MUM4 |
| 260986_at | 2028.7 | P | 1030.6 | P | 1245 | P | 937.2 | P | GLY3 |
| 260987_at | 195.5 | P | 257.8 | P | 148 | P | 247.7 | P | NTMC2T6.1 |
| 260988_at | 548.2 | P | 476.5 | P | 391.4 | P | 472.6 | P | MAP3KA |
| 260989_at | 187.8 | P | 180.1 | P | 271.5 | P | 220.2 | P | --- |
| 260990_at | 6.5 | A | 9.8 | A | 36.8 | A | 4.2 | A | --- |
| 260991_at | 56.5 | P | 46.8 | A | 70.6 | P | 42.1 | P | --- |
| 260992_at | 12 | A | 1.2 | A | 1.5 | A | 8.8 | A | --- |
| 260993_at | 647 | P | 631.7 | P | 525.6 | P | 641.4 | P | FMO GS-OX5 |
| 260994_at | 9.4 | A | 4.7 | A | 1.5 | A | 4.3 | A | --- |
| 260995_at | 638.2 | P | 598.1 | P | 764.7 | P | 753 | P | --- |
| 260996_at | 59.7 | P | 66.2 | P | 62.8 | P | 63.8 | P | RFL1 |
| 260997_at | 38.8 | P | 40.7 | P | 53.9 | P | 32.5 | P | --- |
| 260998_at | 46.3 | P | 34.4 | P | 62.8 | P | 19 | P | --- |
| 260999_at | 547.6 | P | 589.5 | P | 458.1 | P | 572.1 | P | --- |
| 261000_at | 58.7 | P | 122.5 | P | 80 | A | 94.4 | P | --- |
| 261001_at | 24.1 | A | 35.4 | A | 24.7 | A | 18.9 | A | --- |
| 261002_at | 195.1 | P | 133.2 | P | 217.8 | P | 129.5 | P | --- |
| 261003_at | 89.2 | P | 78.1 | M | 68.8 | A | 84.1 | P | --- |
| 261004_at | 45.1 | P | 64 | P | 93 | P | 56.1 | P | --- |
| 261005_at | 71.8 | P | 52.8 | A | 44.7 | A | 16.8 | A | --- |
| 261006_at | 6.7 | A | 3.3 | A | 8.5 | A | 11.1 | A | --- |
| 261007_at | 4.1 | A | 1.5 | A | 19.4 | A | 11.8 | A | --- |
| 261008_at | 225.6 | P | 235.6 | P | 297.6 | P | 271.3 | P | --- |
| 261009_at | 25.7 | A | 20.7 | A | 50.7 | A | 25.6 | A | MES13 |
| 261010_at | 11.7 | A | 21.2 | P | 45.2 | A | 27.5 | A | --- |
| 261011_at | 211.5 | P | 288.6 | P | 298.4 | P | 293.5 | P | CB5-A |
| 261012_at | 97.9 | P | 89.7 | P | 101.9 | P | 126.6 | P | CLE9 |
| 261013_at | 289.8 | P | 276.4 | P | 202.6 | P | 219.2 | P | ATUPS5 |
| 261014_at | 678 | P | 608.6 | P | 529.1 | P | 729.7 | P | --- |
| 261015_at | 1 | A | 5.2 | A | 18.8 | A | 1.8 | A | GRF12 |
| 261016_at | 747.9 | P | 860.6 | P | 599.2 | P | 869.3 | P | BGLU40 |
| 261017_at | 330.6 | P | 401.7 | P | 353.7 | P | 357.4 | P | UGD1 |
| 261018_at | 4380.5 | P | 4538.9 | P | 3781.1 | P | 4574.7 | P | --- |
| 261019_at | 1321.9 | P | 957.1 | P | 1452.9 | P | 913.8 | P | --- |
| 261020_at | 24.5 | A | 25.3 | A | 27.9 | A | 16.1 | A | --- |
| 261021_at | 90 | P | 29 | A | 50.9 | P | 31.9 | A | --- |
| 261022_at | 25.1 | A | 58.1 | M | 56.4 | A | 33.4 | P | ANAC007 |
| 261023_at | 754.2 | P | 877.8 | P | 802.3 | P | 931.1 | P | --- |
| 261024_at | 6 | A | 1.6 | A | 25.8 | A | 3.8 | A | --- |
| 261025_at | 41.5 | A | 71.6 | P | 36.1 | A | 78 | P | --- |
| 261026_at | 136.9 | P | 244.8 | P | 72.1 | A | 172 | P | --- |
| 261027_at | 97.9 | P | 82.6 | P | 129.7 | P | 44.4 | P | ATCNGC10 |
| 261028_at | 152.1 | P | 155.9 | P | 138 | P | 134.4 | P | --- |

| | | | | | | | | | |
|---|---|---|---|---|---|---|---|---|---|
| 261029_at | 95.6 | P | 108.5 | P | 121.6 | P | 120.9 | P | --- |
| 261030_at | 137.6 | P | 114.1 | P | 162.6 | P | 125 | P | --- |
| 261031_at | 352.4 | P | 547.5 | P | 483.2 | P | 570.4 | P | --- |
| 261032_at | 146.1 | P | 163.9 | P | 131.1 | P | 166.6 | P | --- |
| 261033_at | 872.3 | P | 279.6 | P | 728.2 | P | 356.4 | P | JAZ5 |
| 261034_s_at | 77.5 | P | 95.9 | P | 70 | P | 79.5 | P | --- |
| 261035_at | 1.6 | A | 0.5 | A | 1.2 | A | 0.5 | A | --- |
| 261036_at | 1551.4 | P | 1295.6 | P | 1557.8 | P | 1296.7 | P | ATDRG1 |
| 261037_at | 924.5 | P | 167.9 | P | 373.5 | P | 243.4 | P | LOX3 |
| 261038_at | 935 | P | 685.9 | P | 1558.8 | P | 772.2 | P | --- |
| 261039_at | 104.2 | P | 96.7 | P | 89.8 | P | 78.2 | P | ELF4-L4 |
| 261040_at | 1107.7 | P | 1151.2 | P | 1140.5 | P | 1233.6 | P | UBP1B |
| 261041_at | 435.2 | P | 462.4 | P | 365.4 | P | 479.6 | P | EER4 |
| 261042_at | 46.2 | A | 47.9 | A | 14.5 | A | 25.9 | A | ATRABA3 |
| 261043_at | 339.3 | P | 454.3 | P | 381.3 | P | 613.7 | P | --- |
| 261044_at | 512.3 | P | 432.2 | P | 439.3 | P | 424.5 | P | CNX3 |
| 261045_at | 3.4 | A | 2.2 | A | 3.5 | A | 1.7 | A | --- |
| 261046_at | 67.2 | P | 86.4 | P | 115.6 | P | 115.9 | P | --- |
| 261047_at | 24.4 | A | 1.1 | A | 1.6 | A | 4.7 | A | APUM22 |
| 261048_at | 187.4 | P | 245.7 | P | 239.5 | P | 274.7 | P | UGT72B3 |
| 261049_at | 1151.8 | P | 1173.7 | P | 1193.8 | P | 1248.3 | P | --- |
| 261050_at | 260.7 | P | 192.8 | P | 195.7 | P | 209.4 | P | --- |
| 261051_at | 3.7 | A | 1.8 | A | 4.2 | A | 13.5 | A | CYP703A2 |
| 261052_at | 138.2 | P | 150.2 | P | 145.5 | P | 157.1 | P | --- |
| 261053_at | 2181.9 | P | 2085.8 | P | 2759.1 | P | 2087.7 | P | --- |
| 261054_at | 232.7 | P | 187.4 | P | 259.1 | P | 241.2 | P | --- |
| 261055_at | 971 | P | 1700.6 | P | 1061.6 | P | 1606.8 | P | --- |
| 261056_at | 334.3 | P | 372.7 | P | 242.3 | P | 332.7 | P | --- |
| 261057_at | 819.7 | P | 882.6 | P | 699.5 | P | 728.7 | P | --- |
| 261058_at | 31.7 | P | 88.2 | P | 64.4 | P | 101.3 | P | HTR12 |
| 261059_at | 58 | A | 45.2 | A | 55.9 | A | 56.8 | A | --- |
| 261060_at | 312.1 | P | 238.6 | P | 211.6 | P | 211.2 | P | --- |
| 261061_at | 8.6 | A | 2.3 | A | 3.5 | A | 1.7 | A | TRFL2 |
| 261062_at | 316.9 | P | 364.2 | P | 370 | P | 412.4 | P | SCL14 |
| 261063_at | 112 | P | 90.8 | P | 126.4 | P | 87.6 | P | --- |
| 261064_at | 748.8 | P | 657.9 | P | 825.6 | P | 675.5 | P | ftsh10 |
| 261065_at | 21.7 | A | 37.4 | A | 102.6 | A | 58.1 | A | --- |
| 261066_at | 8.5 | A | 26.3 | P | 1.9 | A | 15.4 | A | --- |
| 261067_at | 1.3 | A | 1.1 | A | 2.8 | A | 18.4 | A | --- |
| 261068_at | 40 | P | 89.8 | P | 81.8 | M | 146.7 | P | --- |
| 261069_at | 19.3 | A | 62.6 | P | 41.1 | A | 44.8 | P | ATRABA2B |
| 261070_at | 327.1 | P | 295.6 | P | 359.8 | P | 287.3 | P | AtRLP1 |
| 261071_at | 189.7 | P | 202.8 | P | 169.8 | P | 229.4 | P | --- |
| 261072_at | 16.1 | A | 4.8 | A | 5.3 | A | 7.8 | A | ATSTP2 |
| 261073_at | 66.9 | P | 124.4 | P | 121.1 | P | 122 | P | --- |
| 261074_at | 38 | A | 57.7 | A | 60.5 | A | 21.4 | A | --- |
| 261075_at | 1015.6 | P | 980.3 | P | 1043.5 | P | 1210.6 | P | --- |
| 261076_at | 482.1 | P | 595 | P | 594.5 | P | 703.7 | P | SMO2-1 |
| 261077_at | 82.9 | P | 50.9 | P | 39.2 | A | 69.7 | P | --- |
| 261078_at | 3946.6 | P | 6550.4 | P | 5484.2 | P | 6004.9 | P | RPL4 |
| 261079_s_at | 881.2 | P | 863.2 | P | 691.3 | P | 781.3 | P | --- |
| 261080_at | 401.2 | P | 756.4 | P | 444.4 | P | 725.1 | P | PCNA1 |
| 261081_at | 929.7 | P | 1383.2 | P | 1622.2 | P | 1237.1 | P | --- |
| 261082_at | 717.5 | P | 796.7 | P | 855.2 | P | 729.4 | P | --- |
| 261083_at | 529.9 | P | 627.1 | P | 392.3 | P | 550.9 | P | --- |
| 261084_at | 1027.2 | P | 1438.8 | P | 938 | P | 1430.9 | P | --- |
| 261085_at | 83.2 | P | 135.4 | P | 110.5 | P | 93.6 | P | IQD7 |
| 261086_at | 108 | P | 241 | P | 124.7 | P | 243.7 | P | TRFL3 |
| 261087_at | 444.9 | P | 428.8 | P | 388.3 | P | 395 | P | --- |
| 261088_at | 130.9 | P | 84.8 | P | 61.4 | P | 100.5 | P | --- |
| 261089_at | 194.1 | P | 222.3 | P | 209.6 | P | 242.9 | P | APK1A |
| 261090_at | 15.1 | A | 23.4 | A | 39.5 | A | 39.8 | M | --- |
| 261091_at | 8.3 | A | 11.8 | A | 55.8 | P | 37.3 | A | --- |
| 261092_at | 93.9 | P | 113 | P | 130.2 | P | 106.5 | P | --- |
| 261093_at | 8.4 | A | 5 | A | 14.4 | A | 8.8 | A | AT1G62880 |
| 261094_at | 41.7 | A | 49 | A | 56.6 | A | 37.7 | A | --- |
| 261095_at | 92.1 | P | 77.4 | P | 102.3 | P | 79.8 | P | --- |
| 261096_at | 1.3 | A | 5.3 | A | 41 | A | 8.8 | A | ACOS5 |
| 261097_at | 86.9 | P | 93.5 | P | 104.9 | P | 62 | P | --- |
| 261098_at | 72.3 | P | 93.4 | P | 140.2 | P | 104.5 | P | --- |
| 261099_at | 7.5 | A | 42.5 | A | 49 | A | 24.4 | A | ATEXPA18 |
| 261100_at | 131.4 | P | 119.2 | P | 101 | A | 122.7 | P | NRPD1A |
| 261101_at | 1.5 | A | 5.7 | A | 4.4 | A | 3.5 | A | ddf2 |
| 261102_at | 204.9 | P | 189.1 | P | 188.1 | P | 178.3 | P | --- |
| 261103_s_at | 7.2 | A | 9.9 | A | 7.9 | A | 41.1 | A | --- |
| 261104_at | 17.9 | A | 25 | P | 35.4 | P | 23.5 | A | --- |
| 261105_at | 2351.6 | P | 2749.6 | P | 2646.1 | P | 2865.8 | P | NRS/ER |
| 261106_at | 57.9 | P | 53.9 | P | 72.8 | P | 71.9 | P | KNAT7 |
| 261107_at | 2369.9 | P | 1399.9 | P | 2337.6 | P | 1662.9 | P | --- |
| 261108_at | 486.6 | P | 712.1 | P | 528.7 | P | 644.3 | P | ACS10 |
| 261109_at | 225.4 | P | 86.6 | P | 175.1 | P | 98.9 | P | CKX5 |
| 261110_at | 1005.5 | P | 467.4 | P | 852 | P | 553.7 | P | UBC16 |
| 261111_at | 37.2 | A | 35.9 | A | 52.1 | A | 64.4 | A | BLH11 |
| 261112_at | 309.6 | P | 323.5 | P | 247.8 | P | 303.3 | P | --- |
| 261113_at | 244.3 | P | 248.3 | P | 191.5 | P | 249.1 | P | --- |
| 261114_at | 177.6 | P | 320.1 | P | 224.7 | P | 279.3 | P | AtbZIP44 |
| 261115_at | 1.2 | A | 6.3 | A | 13.9 | A | 3.2 | A | --- |
| 261116_at | 758.6 | P | 696.3 | P | 659.3 | P | 652.8 | P | --- |
| 261117_at | 149.7 | P | 185.3 | P | 173.9 | P | 142.6 | P | AUL1 |
| 261118_at | 805.5 | P | 1175.6 | P | 944.9 | P | 946.6 | P | --- |
| 261119_at | 6441.7 | P | 8059.2 | P | 6099.3 | P | 7874 | P | emb2184 |
| 261120_at | 639.9 | P | 609 | P | 537 | P | 511.3 | P | BLH3 |
| 261121_at | 167.6 | P | 191.6 | P | 142.2 | P | 191.3 | P | --- |
| 261122_at | 3101.7 | P | 2393.7 | P | 3085.7 | P | 2523.7 | P | OTC |
| 261123_at | 116.8 | P | 103.6 | P | 93.9 | A | 94.1 | P | --- |
| 261124_at | 283.5 | P | 249.2 | P | 307.6 | P | 320.8 | P | --- |
| 261125_at | 1182.1 | P | 969.6 | P | 1130.9 | P | 1092.9 | P | --- |
| 261126_at | 231.9 | P | 341.3 | P | 275.8 | P | 298.4 | P | TAF6 |
| 261127_at | 230.7 | P | 228 | P | 195.6 | P | 212.2 | P | --- |
| 261128_at | 383.8 | P | 372.7 | P | 386.9 | P | 378.1 | P | UBP2 |
| 261129_at | 1910 | P | 3542.9 | P | 2520.3 | P | 3695 | P | TUA4 |
| 261130_at | 716.6 | P | 636.3 | P | 858.2 | P | 655.3 | P | PRMT10 |
| 261131_at | 667 | P | 719.6 | P | 737.6 | P | 647.3 | P | --- |
| 261132_at | 867.1 | P | 1098.4 | P | 782.3 | P | 1207 | P | TGD1 |
| 261133_at | 332.7 | P | 506.2 | P | 497.4 | P | 509 | P | --- |
| 261134_at | 18.1 | A | 55 | A | 72.7 | A | 56.2 | A | CYP722A1 |
| 261135_at | 575.1 | P | 109.6 | P | 149.2 | P | 52.3 | P | PDF1.4 |
| 261136_at | 1209.7 | P | 780.4 | P | 1224.8 | P | 783.2 | P | --- |
| 261137_at | 30.7 | A | 39.1 | P | 28.4 | A | 35.9 | P | --- |

| | | | | | | | | | |
|---|---|---|---|---|---|---|---|---|---|
| 261138_at | 221.4 | P | 228.8 | P | 162.2 | P | 255.5 | P | --- |
| 261139_at | 180.6 | P | 229.6 | P | 212.2 | P | 181.4 | P | BEL10 |
| 261140_at | 651 | P | 495.8 | P | 811.5 | P | 536.7 | P | --- |
| 261141_at | 1326.6 | P | 1840.7 | P | 934.3 | P | 1658.2 | P | --- |
| 261142_at | 0.7 | A | 0.7 | A | 1.1 | A | 1.3 | A | ATCNGC8 |
| 261143_at | 706.4 | P | 947.6 | P | 668 | P | 879.5 | P | ATPUP14 |
| 261144_s_at | 5585.1 | P | 6834.7 | P | 4625.6 | P | 5891.5 | P | --- |
| 261145_at | 833.5 | P | 605.6 | P | 766.4 | P | 618.5 | P | ATTRX4 |
| 261146_at | 29 | M | 45.2 | P | 63.7 | P | 51.7 | P | --- |
| 261147_at | 201.9 | P | 243 | P | 210.2 | P | 255.7 | P | --- |
| 261148_at | 544.8 | P | 534.2 | P | 434.6 | P | 557 | P | GAMMA CA1 |
| 261149_s_at | 17738 | P | 10069.5 | P | 14238.9 | P | 10978.8 | P | DHAR1 |
| 261150_at | 151.8 | P | 5.4 | A | 81.8 | A | 35.2 | A | JMT |
| 261151_at | 165.2 | P | 191.7 | P | 219.4 | P | 168.5 | P | --- |
| 261152_at | 42.4 | A | 65.7 | P | 113.4 | A | 81.3 | P | --- |
| 261153_at | 1428.8 | P | 1354.1 | P | 1059.5 | P | 1382.2 | P | --- |
| 261154_at | 1268.1 | P | 1096.5 | P | 1043.3 | P | 1027.2 | P | --- |
| 261155_at | 778.8 | P | 685.4 | P | 638.2 | P | 680.5 | P | --- |
| 261156_s_at | 22.2 | A | 21.7 | A | 8.6 | A | 16.7 | A | --- |
| 261157_at | 3.3 | A | 1.8 | A | 2.1 | A | 2.9 | A | --- |
| 261158_at | 20.3 | A | 28.6 | A | 15.3 | A | 25.5 | A | --- |
| 261159_s_at | 132.2 | P | 204.1 | P | 117.5 | P | 169.4 | P | CYCB1;5 |
| 261160_at | 3.7 | A | 15.7 | A | 21.2 | A | 24 | P | --- |
| 261161_at | 141.7 | P | 111.4 | P | 103.6 | P | 97.2 | P | --- |
| 261162_at | 54.6 | P | 57.4 | P | 57.5 | P | 51.6 | P | --- |
| 261163_x_at | 2.6 | A | 4.7 | A | 4.2 | A | 4.7 | A | ARF21 |
| 261164_at | 237.8 | P | 329.9 | P | 342 | P | 287.1 | P | --- |
| 261165_at | 1692.7 | P | 1616.3 | P | 1665.9 | P | 1573.1 | P | EMB3003 |
| 261166_s_at | 173.4 | P | 255.3 | P | 200.3 | P | 249.2 | P | --- |
| 261167_at | 669.9 | P | 596.6 | P | 501.6 | P | 639 | P | ATPDIL2-2 |
| 261168_at | 255 | P | 170.8 | P | 261.6 | P | 251.2 | P | --- |
| 261169_at | 63.3 | A | 34.9 | A | 51.8 | A | 62 | M | ATSPS3F |
| 261170_at | 415.2 | P | 643.8 | P | 481.8 | P | 524.9 | P | --- |
| 261171_at | 43.1 | P | 36.6 | P | 47.4 | P | 37.9 | A | --- |
| 261172_at | 18.4 | A | 32.6 | A | 45 | A | 22.3 | A | --- |
| 261173_at | 579.2 | P | 508.4 | P | 602 | P | 499.4 | P | --- |
| 261174_at | 698.6 | P | 673.2 | P | 731.2 | P | 691.9 | P | --- |
| 261175_at | 975.9 | P | 1751.1 | P | 1270.4 | P | 1509.6 | P | --- |
| 261176_at | 551.7 | P | 530.5 | P | 500.4 | P | 459.1 | P | --- |
| 261177_at | 259.7 | P | 416.7 | P | 254.8 | P | 273.6 | P | --- |
| 261178_at | 79.3 | P | 71.3 | P | 79.8 | P | 93.8 | P | ATVAMP726 |
| 261179_at | 753.2 | P | 711.1 | P | 838.7 | P | 658.4 | P | --- |
| 261180_at | 0.9 | A | 1.9 | A | 16 | A | 1.9 | A | --- |
| 261181_at | 107.9 | P | 68.4 | P | 120.8 | P | 70.5 | P | --- |
| 261182_at | 3.9 | A | 4.5 | A | 8.7 | A | 12.7 | A | --- |
| 261183_at | 519 | P | 585.7 | P | 605.3 | P | 563.6 | P | EMB2756 |
| 261184_at | 14.1 | A | 8.3 | A | 2.7 | A | 7.3 | A | --- |
| 261185_at | 7.1 | A | 21.6 | A | 10.5 | A | 23.5 | A | CYP94D1 |
| 261186_at | 1.8 | A | 0.9 | A | 1.7 | A | 2.2 | A | --- |
| 261187_at | 195.1 | P | 175.4 | P | 254.8 | P | 158.9 | P | --- |
| 261188_at | 7.1 | A | 15.6 | A | 24.5 | A | 7.3 | A | --- |
| 261189_at | 1826.6 | P | 2296.4 | P | 1833.7 | P | 2185.6 | P | NACA5 |
| 261190_at | 3623.4 | P | 4543.8 | P | 4052.9 | P | 4831.5 | P | PRPL11 |
| 261191_at | 492 | P | 1238.6 | P | 661.6 | P | 1319.1 | P | --- |
| 261192_at | 334.7 | P | 199.3 | P | 341.2 | P | 181 | P | ANAC13 |
| 261193_at | 2190.4 | P | 1046.3 | P | 1080 | P | 999.8 | P | --- |
| 261194_at | 52.6 | P | 48.8 | P | 49.4 | P | 29.3 | P | --- |
| 261195_at | 452.3 | P | 459.1 | P | 500 | P | 424.1 | P | AFB3 |
| 261196_at | 519.6 | P | 521.8 | P | 475.2 | P | 628.9 | P | SCRM2 |
| 261197_at | 8658.8 | P | 11712.9 | P | 10730.3 | P | 11391 | P | GAPA-2 |
| 261198_at | 34 | P | 92.5 | M | 35.5 | A | 71.1 | P | ATNRT2.5 |
| 261199_at | 36.5 | A | 38.8 | A | 73 | A | 50.9 | A | --- |
| 261200_at | 33.5 | P | 47.6 | A | 79 | A | 40.5 | A | --- |
| 261201_at | 550.4 | P | 842.9 | P | 515.5 | P | 795.4 | P | --- |
| 261202_at | 820.7 | P | 812.5 | P | 899.9 | P | 730.1 | P | ATAN11 |
| 261203_at | 1031.5 | P | 942.2 | P | 950.3 | P | 1016.9 | P | --- |
| 261204_s_at | 2783.7 | P | 2256.6 | P | 2686.3 | P | 2455.4 | P | ERF1-2 |
| 261205_at | 85.6 | P | 93.8 | P | 108.6 | P | 90.7 | P | --- |
| 261206_at | 1825.2 | P | 2524.7 | P | 1745.3 | P | 2408.5 | P | --- |
| 261207_at | 292.4 | P | 282.7 | P | 344.9 | P | 289.5 | P | --- |
| 261208_at | 237.9 | P | 308.2 | P | 326.4 | P | 294.3 | P | --- |
| 261209_at | 308.5 | P | 334.5 | P | 337.5 | P | 288.6 | P | --- |
| 261210_at | 4345.5 | P | 4375 | P | 4015 | P | 4703.5 | P | DET3 |
| 261211_at | 2157.3 | P | 1690.1 | P | 1894.3 | P | 1565.4 | P | UGE1 |
| 261212_at | 269.8 | P | 236.5 | P | 140.2 | P | 158.4 | P | atnudt12 |
| 261213_at | 84.3 | P | 131.4 | P | 81.1 | P | 101 | P | --- |
| 261214_at | 1.6 | A | 8.9 | A | 4 | A | 1 | A | --- |
| 261215_at | 2.1 | A | 10.2 | A | 6.1 | A | 15.8 | A | --- |
| 261216_at | 487.4 | P | 200.8 | P | 216.3 | P | 185.2 | P | --- |
| 261217_at | 9.6 | A | 16.5 | A | 13.4 | A | 18.4 | A | UBP11 |
| 261218_at | 11065.2 | P | 11378.3 | P | 10764.6 | P | 10880.8 | P | FNR2 |
| 261219_at | 89.5 | P | 101.2 | P | 95.1 | P | 78.5 | P | --- |
| 261220_at | 227.2 | P | 181.4 | P | 200.3 | P | 220.8 | P | --- |
| 261221_at | 381.4 | P | 269.9 | P | 333.5 | P | 236.2 | P | --- |
| 261222_at | 33.8 | A | 48.1 | A | 38.5 | A | 30.6 | A | --- |
| 261223_at | 105.6 | P | 123.5 | P | 108.2 | P | 105.6 | P | HVA22H |
| 261224_at | 386.3 | P | 369.7 | P | 456.6 | P | 490.1 | P | ATSBT5.2 |
| 261225_at | 1325.1 | P | 1055.9 | P | 1270.8 | P | 1068.1 | P | --- |
| 261226_at | 314.5 | P | 325 | P | 209.8 | P | 264.5 | P | ATEXPA11 |
| 261227_at | 1234.9 | P | 1508.3 | P | 1261 | P | 1441.6 | P | EMB2719 |
| 261228_at | 2117.4 | P | 2124.6 | P | 2039.7 | P | 2141.4 | P | HYD1 |
| 261229_at | 622.1 | P | 1044.9 | P | 851.1 | P | 1095.8 | P | ROP2 |
| 261230_at | 1176.9 | P | 2155.6 | P | 1444.9 | P | 2184.9 | P | TUB5 |
| 261231_at | 2051.6 | P | 1873.6 | P | 1570 | P | 1727 | P | --- |
| 261232_at | 1446.6 | P | 1236.9 | P | 1744.7 | P | 1381 | P | --- |
| 261233_at | 61.6 | A | 62.2 | A | 63.2 | A | 62 | A | --- |
| 261234_x_at | 7.8 | A | 2.5 | A | 2.4 | A | 6.6 | A | --- |
| 261235_x_at | 33.7 | A | 66.9 | A | 41.4 | A | 15.8 | A | --- |
| 261236_at | 2.9 | A | 1.6 | A | 4.7 | A | 4.7 | A | --- |
| 261237_at | 13.9 | A | 1.4 | A | 23.3 | A | 1.4 | A | --- |
| 261238_at | 251.7 | P | 221.9 | P | 265.9 | P | 268.6 | P | --- |
| 261239_at | 183.9 | P | 206.3 | P | 196.6 | P | 183.2 | P | --- |
| 261240_at | 485.1 | P | 192 | P | 187.8 | P | 183.3 | P | SBT3.5 |
| 261241_at | 36.7 | A | 6.3 | A | 53.7 | A | 8.3 | A | --- |
| 261242_at | 305.7 | P | 93.6 | P | 192.4 | P | 81.7 | A | SBT3.3 |
| 261243_at | 10.2 | A | 6.9 | A | 21.7 | A | 1 | A | --- |
| 261244_at | 11.7 | A | 26.5 | A | 45.5 | A | 26.2 | A | --- |
| 261245_at | 11.3 | A | 20.4 | A | 18.7 | A | 14.2 | A | --- |
| 261246_at | 5.7 | A | 3.8 | A | 19.6 | A | 3.2 | A | SYTB |

| | | | | | | | | | |
|---|---|---|---|---|---|---|---|---|---|
| 261247_at | 243.3 | P | 345.4 | P | 137.1 | P | 389.7 | P | --- |
| 261248_at | 209.1 | P | 349.2 | P | 301.3 | P | 441.1 | P | --- |
| 261249_at | 8.9 | A | 32.5 | P | 27.1 | A | 24.5 | A | --- |
| 261250_at | 459.6 | P | 432 | P | 433.5 | P | 414.1 | P | --- |
| 261251_at | 3.5 | A | 2.2 | A | 3.8 | A | 3.6 | A | --- |
| 261252_at | 361 | P | 369 | P | 339.2 | P | 316.9 | P | RABA5E |
| 261253_at | 368.1 | P | 389.9 | P | 308.5 | P | 310.6 | P | --- |
| 261254_at | 434.8 | P | 467 | P | 448.5 | P | 442.5 | P | --- |
| 261255_at | 200.9 | P | 255 | P | 299.6 | P | 216.9 | P | ATNTH2 |
| 261256_at | 70.7 | P | 97.4 | P | 121.7 | P | 117 | P | RTM1 |
| 261257_at | 271.5 | P | 297.3 | P | 325.1 | P | 369 | P | --- |
| 261258_at | 507.9 | P | 664.6 | P | 571 | P | 587.5 | P | --- |
| 261259_at | 204 | P | 110.8 | P | 150.5 | P | 152.7 | P | --- |
| 261260_at | 3.2 | A | 1.6 | A | 4.4 | A | 3.8 | A | --- |
| 261261_at | 162.5 | P | 142 | P | 131 | P | 76.3 | P | --- |
| 261262_at | 66.9 | P | 53.6 | P | 41.9 | P | 74.4 | P | SDG35 |
| 261263_at | 1.9 | A | 3.6 | A | 26.6 | A | 2.6 | A | --- |
| 261264_at | 15.9 | P | 28.9 | P | 38.2 | P | 20.6 | P | --- |
| 261265_at | 155.6 | P | 180.5 | P | 133.8 | P | 137.7 | P | --- |
| 261266_at | 7067.5 | P | 4706.4 | P | 6975.5 | P | 6435.2 | P | ATEXPA10 |
| 261267_at | 977.7 | P | 982 | P | 1172.9 | P | 985.4 | P | --- |
| 261268_at | 450 | P | 563.2 | P | 634 | P | 601.6 | P | --- |
| 261269_at | 586.1 | P | 477.4 | P | 395.1 | P | 446.3 | P | --- |
| 261270_at | 13216.6 | P | 12485.2 | P | 13498.4 | P | 13501 | P | FBR12 |
| 261271_at | 23.1 | A | 23.4 | A | 24.6 | A | 8.6 | A | --- |
| 261272_at | 1134.3 | P | 901 | P | 979 | P | 966.1 | P | --- |
| 261273_at | 433 | P | 555.7 | P | 338 | P | 492.7 | P | --- |
| 261274_at | 22.4 | A | 35.8 | A | 65.6 | A | 36.3 | A | MYB117 |
| 261275_at | 21 | A | 31.4 | A | 50.5 | A | 26.6 | A | MLO14 |
| 261276_at | 1654.6 | P | 1613.4 | P | 1506.5 | P | 1705.2 | P | VTI1B |
| 261277_at | 140.7 | P | 150.7 | P | 158.1 | P | 124.4 | P | --- |
| 261278_at | 47.3 | A | 4.5 | A | 28 | A | 5.1 | A | DGL |
| 261279_at | 5308.2 | P | 6723.2 | P | 5268.3 | P | 6855.8 | P | POM1 |
| 261280_at | 70.1 | P | 70.5 | P | 83.3 | P | 123.6 | P | --- |
| 261281_at | 3.9 | A | 1.6 | A | 0.7 | A | 13.1 | A | --- |
| 261282_at | 3.5 | A | 4.2 | A | 21.3 | A | 3.1 | A | APUM10 |
| 261283_s_at | 0.9 | A | 3.8 | A | 1.2 | A | 5.8 | A | --- |
| 261284_at | 1.7 | A | 3 | A | 6.4 | A | 15.1 | A | --- |
| 261285_at | 9335 | P | 8381.9 | P | 9167 | P | 8732.6 | P | ANNAT1 |
| 261286_at | 423.4 | P | 673.6 | P | 650.5 | P | 724.4 | P | --- |
| 261287_at | 23.7 | P | 24.3 | P | 9.2 | A | 33.5 | A | --- |
| 261288_at | 24.6 | A | 6.1 | A | 2.7 | A | 4.4 | A | --- |
| 261289_at | 5.3 | A | 14 | A | 52.2 | A | 3.8 | A | --- |
| 261290_at | 212.8 | P | 241.2 | P | 206.2 | P | 230.4 | P | --- |
| 261291_at | 28.4 | A | 13.3 | A | 15.9 | A | 29 | A | --- |
| 261292_at | 15.9 | A | 77.1 | P | 23 | A | 87 | P | --- |
| 261293_at | 2338.5 | P | 2540.4 | P | 2239.3 | P | 2309.3 | P | --- |
| 261294_at | 237.1 | P | 289.1 | P | 209.5 | P | 217.8 | P | --- |
| 261295_at | 2553.8 | P | 2386.6 | P | 2195.5 | P | 2355.6 | P | --- |
| 261296_at | 374.6 | P | 423.5 | P | 471.6 | P | 414.3 | P | --- |
| 261297_at | 4.2 | A | 8.4 | A | 4.2 | A | 14.6 | A | JAZ4 |
| 261298_at | 2.9 | A | 21.3 | A | 5.9 | A | 3.2 | A | --- |
| 261299_at | 191.3 | P | 249.9 | P | 202.3 | P | 202.6 | P | --- |
| 261300_at | 79.4 | P | 86.4 | P | 31.7 | A | 68.3 | P | --- |
| 261301_at | 776 | P | 850.6 | P | 846.1 | P | 944.4 | P | --- |
| 261302_at | 13.5 | A | 21 | P | 51.4 | A | 33.8 | P | --- |
| 261303_at | 418.1 | P | 448.8 | P | 468.8 | P | 499.1 | P | HON5 |
| 261304_at | 4090 | P | 3608.5 | P | 3100 | P | 3210.1 | P | --- |
| 261305_at | 23.4 | A | 23.4 | A | 65.9 | A | 36.9 | A | GLN1;5 |
| 261306_at | 156.9 | P | 251.5 | P | 186.7 | P | 293.6 | P | --- |
| 261307_at | 732.9 | P | 918.9 | P | 727.8 | P | 883.6 | P | GATB |
| 261308_at | 227.3 | P | 275.3 | P | 274 | P | 299.2 | P | RKL1 |
| 261309_at | 892.7 | P | 1061.7 | P | 1223.3 | P | 1192.4 | P | CPuORF31 |
| 261310_at | 148.4 | P | 146.9 | P | 169 | P | 127.9 | P | PDE247 |
| 261311_at | 3.6 | A | 4.3 | A | 3 | A | 2.8 | A | --- |
| 261312_at | 450 | P | 389.2 | P | 296.2 | P | 372.7 | P | --- |
| 261313_at | 1.9 | A | 4.6 | A | 13 | A | 6.1 | A | --- |
| 261314_at | 1391.2 | P | 971.6 | P | 1532.9 | P | 1093.6 | P | --- |
| 261315_at | 346.4 | P | 329.9 | P | 319.5 | P | 300.5 | P | ERF8 |
| 261316_at | 280.4 | P | 391.1 | P | 263.2 | P | 304.3 | P | ATMAP4K ALPHA1 |
| 261317_at | 492.9 | P | 418.5 | P | 436.5 | P | 445 | P | --- |
| 261318_at | 1028.7 | P | 922.7 | P | 878.4 | P | 996.5 | P | --- |
| 261319_at | 325 | P | 433.5 | P | 395.1 | P | 406 | P | SPA4 |
| 261320_at | 476.6 | P | 464.5 | P | 406 | P | 482.5 | P | --- |
| 261321_at | 34.9 | A | 24.2 | A | 9.4 | A | 27.4 | A | --- |
| 261322_at | 451 | P | 519.6 | P | 519.7 | P | 484.4 | P | ATPUP11 |
| 261323_at | 340 | P | 332.8 | P | 361.7 | P | 407.2 | P | --- |
| 261324_at | 578.3 | P | 781 | P | 539.9 | P | 666.6 | P | --- |
| 261325_at | 129 | P | 134.2 | P | 127.9 | P | 142.2 | P | --- |
| 261326_s_at | 408.7 | P | 597.4 | P | 534.4 | P | 561.2 | P | --- |
| 261327_at | 54.5 | P | 424.8 | P | 103.2 | P | 164.8 | P | --- |
| 261328_at | 369.6 | P | 383.1 | P | 509.7 | P | 438.2 | P | --- |
| 261329_at | 124 | A | 67.2 | A | 107.4 | A | 104.3 | A | --- |
| 261330_at | 134.6 | P | 197.4 | P | 193 | P | 203.8 | P | MCM2 |
| 261331_at | 30.5 | A | 52.6 | M | 67.1 | M | 38.8 | A | --- |
| 261332_s_at | 0.9 | A | 0.7 | A | 0.9 | A | 2 | A | --- |
| 261333_at | 145.4 | P | 106.7 | P | 106.7 | P | 142.6 | P | --- |
| 261334_at | 246.4 | P | 234.1 | P | 249.3 | P | 245.5 | P | --- |
| 261335_at | 516.7 | P | 892.5 | P | 745.9 | P | 922.9 | P | --- |
| 261336_at | 1138.4 | P | 729.9 | P | 794.6 | P | 720 | P | --- |
| 261337_at | 416.2 | P | 457.3 | P | 399.6 | P | 409.4 | P | --- |
| 261338_at | 1485 | P | 1639.4 | P | 1587.7 | P | 1847.1 | P | --- |
| 261339_at | 345 | P | 1106 | P | 368.7 | P | 569.9 | P | --- |
| 261340_at | 15.5 | A | 26.4 | A | 5 | A | 13.5 | A | APUM9 |
| 261341_s_at | 4.2 | A | 11.4 | A | 7.9 | A | 5.8 | A | PAP5 |
| 261342_at | 5.3 | A | 3.4 | A | 3 | A | 4.5 | A | --- |
| 261343_s_at | 2.6 | A | 1 | A | 5.3 | A | 1.7 | A | --- |
| 261344_at | 213.1 | P | 211.4 | P | 264.3 | P | 258.5 | P | --- |
| 261345_at | 14 | A | 11 | A | 47.4 | A | 28.4 | P | DTA4 |
| 261346_at | 667.1 | P | 719.2 | P | 681.3 | P | 672.9 | P | --- |
| 261347_at | 567.3 | P | 560.3 | P | 559.1 | P | 517.8 | P | ELF7 |
| 261348_at | 773.4 | P | 688.3 | P | 792 | P | 644.6 | P | TED3 |
| 261349_at | 11.9 | A | 8 | A | 36.2 | A | 21.6 | A | NHX6 |
| 261350_at | 101.3 | P | 80.7 | P | 66.5 | P | 86.2 | P | --- |
| 261351_at | 833.5 | P | 659.8 | P | 693 | P | 646.6 | P | --- |
| 261352_at | 422.7 | P | 467.4 | P | 559 | P | 510.6 | P | RAD23 |
| 261353_at | 1717.3 | P | 1772.5 | P | 1642.6 | P | 1702.2 | P | --- |
| 261354_at | 2073.4 | P | 1530.3 | P | 1403 | P | 1720.9 | P | atnudt3 |
| 261355_at | 1755.7 | P | 1504.3 | P | 1668.3 | P | 1520.7 | P | ATNADP-ME4 |

| | | | | | | | | | |
|---|---|---|---|---|---|---|---|---|---|
| 261356_at | 396 | P | 447.1 | P | 434.6 | P | 462.6 | P | --- |
| 261357_at | 10.5 | A | 1.4 | A | 1.6 | A | 2.6 | A | --- |
| 261358_s_at | 1.6 | A | 8.7 | A | 1.2 | A | 17.7 | A | --- |
| 261359_at | 7.7 | A | 1.7 | A | 7.7 | A | 8.4 | A | --- |
| 261360_at | 27.1 | A | 30.5 | A | 17.1 | A | 34.7 | A | --- |
| 261361_at | 8.6 | A | 5.9 | A | 13.6 | A | 17.8 | A | --- |
| 261362_s_at | 7689.2 | P | 8211.3 | P | 8032.4 | P | 7989.3 | P | --- |
| 261363_at | 858.6 | P | 1529.1 | P | 905.7 | P | 1534.5 | P | SKS6 |
| 261364_at | 79.6 | M | 95 | P | 133.9 | A | 129 | P | --- |
| 261365_at | 115.1 | P | 92.1 | P | 148.4 | P | 97.3 | P | --- |
| 261366_at | 209.6 | P | 129.4 | P | 102.5 | P | 100.1 | P | --- |
| 261367_at | 3.8 | A | 7.4 | A | 8.2 | A | 21 | A | --- |
| 261368_at | 260.1 | P | 300.5 | P | 236.8 | P | 331.8 | P | --- |
| 261369_at | 132 | P | 162.8 | P | 161.8 | P | 138.8 | P | --- |
| 261370_at | 630 | P | 588.6 | P | 650.2 | P | 527.8 | P | --- |
| 261371_at | 21.5 | A | 32.3 | A | 47.6 | A | 27.7 | A | UBC26 |
| 261372_at | 29.8 | P | 27.7 | A | 44.4 | M | 15.8 | A | --- |
| 261373_at | 814.9 | P | 790.3 | P | 741.6 | P | 923.1 | P | --- |
| 261374_at | 15.9 | P | 48.8 | P | 41.9 | A | 40.1 | P | --- |
| 261375_at | 119.7 | P | 123.9 | P | 148.2 | P | 101.1 | P | SPL4 |
| 261376_at | 734.5 | P | 873.2 | P | 817 | P | 883.8 | P | --- |
| 261377_at | 400.4 | P | 358.5 | P | 516.4 | P | 386.7 | P | --- |
| 261378_at | 462.2 | P | 576.9 | P | 407.4 | P | 450 | P | ATCDPK1 |
| 261379_at | 2689.6 | P | 3114.3 | P | 2864.6 | P | 3101.5 | P | --- |
| 261380_at | 3.5 | A | 16.8 | A | 2.5 | A | 5.2 | A | --- |
| 261381_at | 75.7 | P | 89.3 | P | 101.7 | P | 127 | P | SDE3 |
| 261382_at | 20.1 | A | 97.8 | P | 71.8 | A | 91.3 | P | CVP2 |
| 261383_at | 540.2 | P | 461.3 | P | 557.2 | P | 468 | P | --- |
| 261384_at | 35.9 | P | 69.7 | P | 65.9 | P | 65.4 | P | --- |
| 261385_at | 14.4 | A | 14.2 | A | 25.8 | A | 2.1 | A | --- |
| 261386_at | 719.4 | P | 525.8 | P | 633 | P | 594.4 | P | --- |
| 261387_at | 286.5 | P | 241.7 | P | 263.6 | P | 313.3 | P | --- |
| 261388_at | 527.7 | P | 610.8 | P | 730.7 | P | 489.3 | P | --- |
| 261389_s_at | 0.9 | A | 0.4 | A | 1.7 | A | 13.1 | A | --- |
| 261390_at | 1.9 | A | 2.7 | A | 4.2 | A | 3.1 | A | --- |
| 261391_at | 4.3 | A | 2.1 | A | 18 | A | 14 | A | --- |
| 261392_at | 32.3 | A | 39.4 | A | 58.4 | A | 47.6 | A | --- |
| 261393_at | 24.2 | A | 20.7 | A | 31.9 | A | 30.5 | M | SMB |
| 261394_at | 121.9 | P | 61.7 | P | 94.6 | P | 45.7 | M | --- |
| 261395_at | 82.8 | P | 93.2 | P | 81.6 | A | 68.9 | P | --- |
| 261396_at | 32.1 | A | 27.8 | A | 51.1 | A | 36.9 | A | --- |
| 261397_at | 252.5 | P | 359 | P | 366 | P | 298 | P | SGB1 |
| 261398_at | 279.1 | P | 326.7 | P | 313.6 | P | 303.1 | P | NHX6 |
| 261399_at | 90.2 | A | 80.6 | A | 43.8 | A | 93.8 | A | --- |
| 261400_at | 144.7 | P | 132.6 | P | 189.8 | P | 139.8 | P | --- |
| 261401_at | 142.3 | P | 188.8 | P | 196.2 | P | 164.8 | P | --- |
| 261402_at | 173.2 | P | 200.3 | P | 107.1 | P | 199.3 | P | RFO1 |
| 261403_at | 722.5 | P | 729 | P | 558.2 | P | 613.3 | P | SYP52 |
| 261404_at | 82.5 | P | 92.6 | P | 118.1 | P | 102.6 | P | --- |
| 261405_at | 344.6 | P | 627.2 | P | 225.9 | P | 493.5 | P | --- |
| 261406_at | 599.9 | P | 550.9 | P | 830.3 | P | 580.4 | P | NRP2 |
| 261407_at | 78.2 | P | 80.5 | P | 148.1 | P | 100 | P | --- |
| 261408_s_at | 1091.2 | P | 1503.9 | P | 1002 | P | 1534.1 | P | --- |
| 261409_at | 468.2 | P | 663.5 | P | 659.9 | P | 637.3 | P | OBP2 |
| 261410_at | 1976.8 | P | 1778.4 | P | 1314.4 | P | 2124.6 | P | MT1C |
| 261411_at | 414.5 | P | 635.6 | P | 497.2 | P | 598.5 | P | HTB1 |
| 261412_at | 15478.8 | P | 12765.6 | P | 16382.9 | P | 14870.9 | P | APX1 |
| 261413_at | 253.9 | P | 316.2 | P | 220.9 | P | 244.8 | P | PLL5 |
| 261414_at | 56.1 | A | 39 | A | 51.1 | A | 49.8 | A | --- |
| 261415_at | 1072.8 | P | 805.9 | P | 821.4 | P | 787.2 | P | --- |
| 261416_at | 10039.4 | P | 8421.5 | P | 8940.9 | P | 9401.6 | P | RPS15A |
| 261417_at | 1535.1 | P | 1893.1 | P | 1649.8 | P | 1761.7 | P | --- |
| 261418_at | 1166.2 | P | 830.4 | P | 1256.9 | P | 942.2 | P | --- |
| 261419_at | 456.7 | P | 305.7 | P | 571.8 | P | 349.7 | P | --- |
| 261420_at | 978.4 | P | 872 | P | 774.4 | P | 986.3 | P | KCS3 |
| 261421_at | 260.6 | P | 401.4 | P | 243.5 | P | 364.2 | P | IQD30 |
| 261422_at | 2027.8 | P | 2313.3 | P | 2542.4 | P | 2330.8 | P | NDF6 |
| 261423_at | 122.4 | P | 120.1 | P | 168.7 | P | 108.5 | P | AGL65 |
| 261424_at | 840.8 | P | 803.3 | P | 950.6 | P | 857.5 | P | --- |
| 261425_at | 92.8 | P | 186 | P | 139.3 | P | 194 | P | --- |
| 261426_at | 410.3 | P | 361.3 | P | 251 | P | 311.6 | P | --- |
| 261427_at | 216.1 | P | 196.5 | P | 134.6 | P | 173.5 | P | IBS1 |
| 261428_at | 85.5 | P | 61.3 | P | 85.7 | P | 80 | P | ICS2 |
| 261429_at | 22 | A | 6.9 | A | 5.7 | A | 23.7 | A | WRKY61 |
| 261430_at | 40.8 | M | 18.8 | A | 52.8 | A | 51.6 | A | --- |
| 261431_at | 123.9 | P | 44.7 | A | 36.9 | A | 98.3 | P | AtMYB47 |
| 261432_at | 18.5 | A | 7.5 | A | 24.7 | A | 2.5 | A | --- |
| 261433_s_at | 811.9 | P | 927 | P | 839.1 | P | 859 | P | --- |
| 261434_at | 365.3 | P | 510.4 | P | 375.8 | P | 434.5 | P | --- |
| 261435_at | 219.6 | P | 137.3 | P | 149.7 | P | 107.5 | P | --- |
| 261436_at | 206.6 | P | 247.9 | P | 141.8 | P | 207.7 | P | --- |
| 261437_at | 89.4 | P | 49.3 | A | 29.3 | A | 33.4 | A | RAD51D |
| 261438_at | 19963.6 | P | 18208 | P | 15520.5 | P | 18862.7 | P | --- |
| 261439_at | 824.4 | P | 893.8 | P | 1094.6 | P | 873.5 | P | --- |
| 261440_at | 743.4 | P | 844.1 | P | 1046.4 | P | 894.2 | P | --- |
| 261441_at | 37.9 | P | 74.2 | P | 57.4 | A | 39.3 | P | ANAC010 |
| 261442_at | 6.3 | A | 4 | A | 7.5 | A | 3.7 | A | --- |
| 261443_at | 265.7 | P | 101.2 | P | 179.8 | P | 121.9 | P | GRX480 |
| 261444_at | 606.8 | P | 522.7 | P | 661.2 | P | 530.1 | P | SYP61 |
| 261445_at | 630.8 | P | 709.3 | P | 689.8 | P | 646.4 | P | NSL1 |
| 261446_at | 81.1 | P | 92.4 | P | 86.3 | A | 62.1 | P | SEC5B |
| 261447_at | 346.2 | P | 286.7 | P | 308.9 | P | 297.9 | P | --- |
| 261448_at | 37.4 | A | 31.3 | A | 84.4 | P | 46 | P | --- |
| 261449_at | 24.9 | P | 20.2 | P | 3.5 | A | 31.2 | P | --- |
| 261450_s_at | 35.4 | P | 39.7 | P | 22.1 | P | 48.1 | P | --- |
| 261451_at | 311.7 | P | 142 | P | 322.8 | P | 143.3 | P | --- |
| 261452_s_at | 12.2 | A | 20.4 | A | 3 | A | 13.9 | P | --- |
| 261453_at | 724.1 | P | 1020.7 | P | 425.8 | P | 899.1 | P | --- |
| 261454_at | 128.3 | P | 156.5 | P | 170.1 | P | 140.6 | P | --- |
| 261455_at | 216.6 | P | 246.9 | P | 260.1 | P | 246.7 | P | --- |
| 261456_at | 683.5 | P | 600.4 | P | 895.2 | P | 472.2 | P | --- |
| 261457_at | 2050 | P | 2457.2 | P | 2566.6 | P | 2315 | P | --- |
| 261458_at | 866.2 | P | 981.5 | P | 746.2 | P | 910.6 | P | --- |
| 261459_at | 154.2 | P | 59.5 | P | 86 | P | 61.7 | P | --- |
| 261460_at | 53.4 | P | 78.3 | P | 110.2 | P | 91.3 | P | ATMPK13 |
| 261461_at | 19.4 | A | 7.8 | A | 41.7 | A | 20.2 | A | --- |
| 261462_at | 2.5 | A | 1.2 | A | 22.8 | A | 4.5 | A | --- |
| 261463_at | 48.9 | P | 79 | P | 55.6 | A | 59.3 | P | --- |
| 261464_at | 72.8 | P | 72.8 | P | 69.2 | P | 68.8 | P | --- |

| | | | | | | | | | |
|---|---|---|---|---|---|---|---|---|---|
| 261465_at | 370.5 | P | 435.2 | P | 433.8 | P | 398.2 | P | --- |
| 261466_at | 2 | A | 17.4 | A | 28.9 | A | 22.5 | A | --- |
| 261467_at | 226.9 | P | 274.8 | P | 233.2 | P | 304.9 | P | VOZ1 |
| 261468_s_at | 27 | A | 30.6 | A | 51.6 | A | 24.9 | A | --- |
| 261469_at | 707.4 | P | 821.1 | P | 732.3 | P | 931.5 | P | AtRLP4 |
| 261470_at | 407.1 | P | 314.2 | P | 357.6 | P | 306.5 | P | ERF11 |
| 261471_at | 211.7 | P | 253.9 | P | 279.7 | P | 299.2 | P | --- |
| 261472_at | 15.6 | A | 18 | A | 36 | A | 17.4 | A | --- |
| 261473_at | 6.5 | A | 19 | A | 46.2 | A | 3.3 | A | --- |
| 261474_at | 4.7 | A | 2.1 | A | 7.7 | A | 1.9 | A | --- |
| 261475_at | 6.3 | A | 9.7 | A | 18 | A | 35.5 | A | --- |
| 261476_at | 129.2 | P | 108.1 | P | 122.3 | P | 119.6 | P | --- |
| 261477_at | 380.6 | P | 463.5 | P | 438.8 | P | 418.1 | P | --- |
| 261478_at | 16.5 | A | 38.4 | A | 22.4 | A | 36.6 | P | MIOX1 |
| 261479_at | 181.9 | P | 286.5 | P | 182.6 | P | 263.1 | P | IQD28 |
| 261480_at | 148 | P | 385.7 | P | 221.8 | P | 273.5 | P | PKS2 |
| 261481_at | 35.4 | A | 45.2 | P | 64.9 | A | 60.4 | P | --- |
| 261482_at | 167 | P | 172.9 | P | 121.6 | P | 162.1 | P | THH1 |
| 261483_at | 792.1 | P | 1126.7 | P | 899.4 | P | 1051.2 | P | --- |
| 261484_at | 2427.6 | P | 2770.9 | P | 2258.7 | P | 2705.8 | P | UBC1 |
| 261485_at | 992.6 | P | 999 | P | 811 | P | 937.1 | P | UTR3 |
| 261486_at | 1200.8 | P | 1248.9 | P | 1337.2 | P | 1217.3 | P | AL7 |
| 261487_at | 184 | P | 212.5 | P | 140.5 | P | 193.1 | P | --- |
| 261488_at | 1454.8 | P | 1724.6 | P | 2145.7 | P | 1702.9 | P | --- |
| 261489_at | 2253.2 | P | 1970.4 | P | 1748.2 | P | 1994.3 | P | --- |
| 261490_at | 10355.2 | P | 8996.7 | P | 11086.6 | P | 9839.5 | P | SAC52 |
| 261491_at | 83 | P | 136.6 | P | 56 | P | 135 | P | FLP |
| 261492_at | 291.1 | P | 428 | P | 347 | P | 414.2 | P | SBH2 |
| 261493_at | 422.2 | P | 789.2 | P | 536.1 | P | 708.2 | P | WHY1 |
| 261494_at | 207 | P | 180.9 | P | 190.9 | P | 192.3 | P | HB-1 |
| 261495_at | 317.7 | P | 302.9 | P | 341.5 | P | 358 | P | --- |
| 261496_at | 82 | A | 44.3 | A | 108.5 | A | 38.5 | A | ERF12 |
| 261497_x_at | 37.3 | A | 14.9 | A | 40.6 | A | 22.2 | A | AGL59 |
| 261498_at | 683.8 | P | 829 | P | 750.2 | P | 736.4 | P | HSL1 |
| 261499_at | 11.7 | A | 1.4 | A | 7 | A | 1.8 | A | CYP705A24 |
| 261500_at | 1803.2 | P | 1164.6 | P | 1239.4 | P | 1325.7 | P | --- |
| 261501_at | 182.5 | P | 114 | P | 151.7 | P | 148.5 | P | --- |
| 261502_at | 225.8 | P | 305.7 | P | 391.8 | P | 321.4 | P | AtHB31 |
| 261503_at | 32.7 | A | 50.9 | A | 50.2 | M | 52.4 | P | --- |
| 261504_at | 8.1 | A | 12.1 | A | 29 | A | 17.2 | A | AGL12 |
| 261505_at | 123.8 | P | 108.5 | P | 129.9 | P | 148.5 | P | SOL1 |
| 261506_at | 167.8 | P | 265.6 | P | 129.1 | P | 150 | P | ATCK1 |
| 261507_at | 784.7 | P | 860.2 | P | 713 | P | 814.6 | P | --- |
| 261508_at | 792 | P | 789.6 | P | 1027.6 | P | 792.1 | P | --- |
| 261509_at | 40.8 | A | 40.1 | A | 38.3 | A | 24.4 | A | --- |
| 261510_at | 18.8 | A | 23.6 | A | 2.9 | A | 18.6 | A | --- |
| 261511_at | 39.4 | P | 50.7 | P | 27.2 | A | 36.8 | P | PAB5 |
| 261512_at | 258.4 | P | 365.8 | P | 422.7 | P | 282.2 | P | --- |
| 261513_at | 1425.3 | P | 948 | P | 1741.1 | P | 1242 | P | --- |
| 261514_at | 65.5 | M | 67.7 | P | 111.9 | P | 70.3 | P | --- |
| 261515_at | 220.1 | P | 285 | P | 247.3 | P | 270.3 | P | --- |
| 261516_at | 130.6 | P | 137.4 | P | 96.9 | P | 135.8 | P | --- |
| 261517_at | 743 | P | 1044.7 | P | 950.8 | P | 863.2 | P | --- |
| 261518_at | 2924.7 | P | 2755.7 | P | 2851.3 | P | 2824.5 | P | --- |
| 261519_at | 331.6 | P | 410.5 | P | 376 | P | 382.2 | P | --- |
| 261520_at | 773.9 | P | 953.2 | P | 652.1 | P | 970 | P | SEC6 |
| 261521_at | 90.1 | P | 111.2 | P | 150.9 | P | 130.2 | P | SERK1 |
| 261522_at | 610.8 | P | 733.1 | P | 618.4 | P | 704.2 | P | --- |
| 261523_at | 1385.4 | P | 1553 | P | 1717.4 | P | 1491.8 | P | PTP1 |
| 261524_at | 578 | P | 457.3 | P | 690.2 | P | 512.4 | P | --- |
| 261525_at | 322 | P | 232.3 | P | 300.8 | P | 251.1 | P | --- |
| 261526_at | 298.4 | P | 343.9 | P | 177.7 | P | 211.2 | P | APK2A |
| 261527_at | 21.8 | A | 35.3 | A | 33.5 | A | 45 | A | --- |
| 261528_at | 2.2 | A | 3.3 | A | 4.1 | A | 5.5 | A | AT59 |
| 261529_at | 1.4 | A | 1.3 | A | 15.1 | A | 12.1 | A | --- |
| 261530_at | 633.1 | P | 720.9 | P | 464.4 | P | 654.4 | P | --- |
| 261531_at | 114.3 | P | 127.3 | P | 139.6 | P | 111.6 | P | --- |
| 261532_at | 3.6 | A | 4.1 | A | 8.1 | A | 5.5 | A | --- |
| 261533_at | 7.2 | A | 37.7 | A | 26.3 | A | 30.7 | A | --- |
| 261534_at | 1174.9 | P | 1237.5 | P | 1310.1 | P | 1186.3 | P | PEX11C |
| 261535_at | 275.9 | P | 206.7 | P | 163 | P | 183.8 | P | --- |
| 261536_at | 526.1 | P | 632.1 | P | 640.8 | P | 661.9 | P | KEA1 |
| 261537_at | 2122.5 | P | 2357.3 | P | 2345.5 | P | 2163.6 | P | --- |
| 261538_at | 111.3 | P | 167.9 | P | 110.9 | P | 124.5 | P | --- |
| 261539_at | 70.9 | P | 43 | P | 68.3 | P | 84.5 | P | --- |
| 261540_at | 626 | P | 880.6 | P | 685.1 | P | 837.6 | P | --- |
| 261541_at | 7.3 | A | 2.4 | A | 29.1 | A | 17.5 | A | --- |
| 261542_at | 5.3 | A | 4 | A | 1.9 | A | 5 | A | --- |
| 261543_at | 32.3 | A | 46.3 | P | 27.7 | A | 26.7 | M | --- |
| 261544_at | 1.6 | A | 2.9 | A | 2.8 | A | 2.7 | A | --- |
| 261545_at | 91.3 | P | 169.6 | P | 116 | P | 110.2 | P | --- |
| 261546_at | 16.6 | A | 43.8 | A | 34.6 | A | 31.9 | A | --- |
| 261547_at | 366.8 | P | 429.4 | P | 395.1 | P | 450.9 | P | --- |
| 261548_at | 107.8 | P | 147.7 | P | 109.7 | P | 130.3 | P | --- |
| 261549_at | 141.6 | P | 219.4 | P | 274.9 | P | 229.3 | P | --- |
| 261550_at | 3.6 | A | 4.1 | A | 14.1 | A | 28.6 | A | --- |
| 261551_at | 105.1 | P | 109.8 | P | 150.7 | M | 118.8 | P | HMA5 |
| 261552_at | 245.4 | P | 229.8 | P | 190.1 | P | 221.8 | P | --- |
| 261553_at | 518.2 | P | 444.7 | P | 421.4 | P | 553.3 | P | --- |
| 261554_at | 15.8 | A | 20.1 | P | 15.4 | M | 13.9 | A | --- |
| 261555_s_at | 35.4 | A | 49.7 | A | 13 | A | 51 | M | --- |
| 261556_s_at | 237.4 | P | 209.4 | A | 196.4 | A | 251.3 | A | --- |
| 261557_at | 177 | P | 166.9 | P | 161.6 | P | 190.3 | P | --- |
| 261558_at | 413.7 | P | 351.6 | P | 349.5 | P | 365.3 | P | --- |
| 261559_at | 313.5 | P | 168.1 | P | 384.4 | P | 240.6 | P | --- |
| 261560_at | 753.4 | P | 454.7 | P | 629.7 | P | 467.7 | P | --- |
| 261561_at | 613.5 | P | 615.6 | P | 638.7 | P | 645.8 | P | --- |
| 261562_at | 4.6 | A | 5.3 | A | 1.2 | A | 3.3 | A | ADF11 |
| 261563_at | 1476.7 | P | 1263.2 | P | 1132.3 | P | 1324.1 | P | --- |
| 261564_at | 1529.7 | P | 1183.9 | P | 1177.5 | P | 1368.5 | P | ATAF1 |
| 261565_at | 7.4 | A | 5.9 | A | 28.9 | A | 5.2 | A | --- |
| 261566_at | 502.9 | P | 494.8 | P | 542.8 | P | 491.4 | P | --- |
| 261567_at | 6.9 | A | 34.3 | A | 14.7 | A | 15.2 | A | --- |
| 261568_at | 114.5 | P | 93.9 | P | 106.4 | P | 57 | P | NGA3 |
| 261569_at | 13.2 | A | 296.1 | P | 45.1 | A | 179.1 | P | LHY |
| 261570_at | 351.4 | P | 415.4 | P | 387.3 | P | 444 | P | KCS1 |
| 261571_at | 10.1 | A | 22.5 | A | 20.1 | A | 12.7 | A | --- |
| 261572_at | 3250.5 | P | 3502.9 | P | 3178.5 | P | 3329.9 | P | --- |
| 261573_at | 36.6 | A | 22.5 | A | 14.9 | A | 12.1 | A | --- |

| | | | | | | | | | |
|---|---|---|---|---|---|---|---|---|---|
| 261574_at | 6 | A | 35.2 | A | 24.3 | A | 64.8 | A | CYP78A8 |
| 261575_at | 9 | A | 6.1 | A | 24.3 | A | 5.6 | A | --- |
| 261576_at | 120.9 | P | 266.1 | P | 162.3 | P | 205.8 | P | --- |
| 261577_at | 1256.7 | P | 1384.3 | P | 1130.4 | P | 1156.6 | P | --- |
| 261578_at | 10350.7 | P | 8443 | P | 9895 | P | 9589.9 | P | --- |
| 261579_at | 3052.4 | P | 3579.4 | P | 3783.7 | P | 3668 | P | AtPPa1 |
| 261580_at | 6.6 | A | 28.7 | A | 3.7 | A | 25.3 | A | IQD18 |
| 261581_at | 1456.5 | P | 2246.4 | P | 1155.7 | P | 1952.7 | P | CIPK9 |
| 261582_at | 437.2 | P | 488.9 | P | 451.4 | P | 533.5 | P | GIF2 |
| 261583_at | 6055.6 | P | 4908.2 | P | 6379.4 | P | 5363.8 | P | PDH-E1 ALPHA |
| 261584_at | 380.4 | P | 315.9 | P | 328.5 | P | 291 | P | DCL1 |
| 261585_at | 375.1 | P | 136.4 | P | 292.4 | P | 146 | P | ANAC001 |
| 261586_at | 103.3 | P | 131.7 | P | 74.1 | P | 132.2 | P | --- |
| 261587_at | 45.5 | A | 1.3 | A | 18.2 | A | 32.6 | A | --- |
| 261588_at | 62.2 | P | 69.4 | P | 56.2 | P | 86.4 | P | --- |
| 261589_at | 23.9 | A | 42.1 | A | 48.4 | A | 47.1 | A | --- |
| 261590_at | 15.3 | A | 22 | A | 6.4 | A | 21.2 | A | ROPGEF2 |
| 261591_at | 175.4 | P | 168.6 | P | 197.1 | P | 179.9 | P | --- |
| 261592_at | 3.7 | A | 3 | A | 4.5 | A | 4.6 | A | --- |
| 261593_at | 177.5 | P | 308 | P | 347.2 | P | 349.4 | P | --- |
| 261594_at | 436.7 | P | 537.1 | P | 413.4 | P | 476 | P | AT-GTL1 |
| 261595_at | 0.8 | A | 2 | A | 17.4 | A | 27.6 | A | --- |
| 261596_at | 146.5 | P | 122.1 | P | 176.7 | P | 146.9 | P | --- |
| 261597_at | 122.4 | P | 95.1 | P | 139.3 | P | 84.6 | P | PUB26 |
| 261598_at | 1200.7 | P | 1805.8 | P | 1259.8 | P | 1693.3 | P | --- |
| 261599_at | 59 | P | 72.6 | P | 112.1 | P | 52 | P | --- |
| 261600_x_at | 4 | A | 1.3 | A | 17.9 | A | 1.3 | A | --- |
| 261601_at | 2195.1 | P | 2002.5 | P | 1688.8 | P | 1852.2 | P | NQR |
| 261602_at | 344.9 | P | 434.4 | P | 358.3 | P | 350 | P | ATPREP2 |
| 261603_at | 1594.4 | P | 1637.7 | P | 1710.9 | P | 1703.5 | P | ATRBP47A |
| 261604_at | 159.7 | P | 170.6 | P | 231 | P | 173.2 | P | --- |
| 261605_at | 187.8 | P | 277.8 | P | 263.6 | P | 307.3 | P | --- |
| 261606_at | 29.8 | A | 61.6 | P | 57.8 | A | 60 | A | --- |
| 261607_at | 1317.6 | P | 1351.5 | P | 1936.6 | P | 1483.2 | P | AtCXE5 |
| 261608_at | 360.4 | P | 458.3 | P | 385.7 | P | 483.5 | P | --- |
| 261609_at | 516.2 | P | 335.5 | P | 497.2 | P | 350 | P | --- |
| 261610_at | 196.9 | P | 271.2 | P | 232.8 | P | 229.7 | P | --- |
| 261611_at | 134.7 | P | 128 | P | 123.4 | P | 137.4 | P | --- |
| 261612_at | 301.9 | P | 363.7 | P | 289.2 | P | 337.3 | P | FUT12 |
| 261613_at | 282 | P | 332.4 | P | 234.4 | P | 376.5 | P | ABF1 |
| 261614_at | 3323.4 | P | 2459.6 | P | 3651.6 | P | 2698.9 | P | PAB8 |
| 261615_at | 360.4 | P | 453.1 | P | 364.7 | P | 346.7 | P | --- |
| 261616_at | 144.9 | P | 214.1 | P | 201.6 | P | 225.6 | P | ANAC014 |
| 261617_s_at | 26.4 | M | 17 | A | 26.9 | A | 17.9 | A | --- |
| 261618_at | 1776.2 | P | 1156.1 | P | 1778.5 | P | 1316.8 | P | --- |
| 261619_x_at | 30 | P | 22.1 | P | 42.3 | P | 27.3 | A | --- |
| 261620_s_at | 7416.1 | P | 7313.5 | P | 9443.8 | P | 6499.1 | P | PGY2 |
| 261621_at | 855.4 | P | 866.9 | P | 859.1 | P | 889.5 | P | EDA10 |
| 261622_at | 665.6 | P | 922.6 | P | 732.2 | P | 927.5 | P | --- |
| 261623_at | 24.4 | A | 2.3 | A | 3.3 | A | 3.6 | A | ATSEC1A |
| 261624_at | 155.7 | P | 147 | P | 161.6 | P | 133.9 | P | GAE2 |
| 261625_at | 300.2 | P | 281.5 | P | 241.8 | P | 329.2 | P | --- |
| 261626_at | 312.1 | P | 231 | P | 246.8 | P | 248.2 | P | --- |
| 261627_at | 779 | P | 1002 | P | 910.5 | P | 960.4 | P | --- |
| 261628_at | 112 | P | 125.8 | P | 139 | P | 137 | P | --- |
| 261629_at | 213.4 | P | 192 | P | 230.3 | P | 219.7 | P | --- |
| 261630_at | 17.9 | A | 6.8 | A | 23.4 | A | 30.6 | A | --- |
| 261631_at | 1.6 | A | 23 | A | 19.5 | A | 12.7 | A | --- |
| 261632_at | 498.2 | P | 668.2 | P | 485.8 | P | 653.1 | P | --- |
| 261633_at | 175.6 | P | 124.5 | P | 163.4 | P | 127 | P | --- |
| 261634_at | 3703.3 | P | 3255.9 | P | 2954.3 | P | 3407.7 | P | CLPR1 |
| 261635_at | 2360.2 | P | 2526.8 | P | 2138 | P | 2468.2 | P | --- |
| 261636_at | 83.6 | P | 152.8 | P | 109 | P | 193.7 | P | --- |
| 261637_at | 507.5 | P | 627.3 | P | 639.6 | P | 600.6 | P | TRB1 |
| 261638_at | 445 | P | 724.6 | P | 324.1 | P | 651 | P | --- |
| 261639_at | 1539 | P | 2772.5 | P | 1693.3 | P | 2796.2 | P | TUA2 |
| 261640_at | 42.6 | M | 55.4 | P | 61.4 | P | 35.7 | P | --- |
| 261641_at | 14.5 | A | 39.1 | A | 26.1 | A | 29.7 | A | --- |
| 261642_at | 271.5 | P | 321.9 | P | 269.9 | P | 348.9 | P | APL2 |
| 261643_at | 24.2 | A | 4.5 | A | 45.1 | A | 3.9 | A | TAF4B |
| 261644_s_at | 454.8 | P | 515.8 | P | 415.2 | P | 540.6 | P | ATCSA-1 |
| 261645_at | 0.1 | A | 4.4 | A | 0.4 | A | 1.1 | A | --- |
| 261646_at | 250.9 | P | 286.8 | P | 296.6 | P | 279.5 | P | --- |
| 261647_at | 17.9 | A | 3.6 | A | 30.9 | A | 27.8 | A | --- |
| 261648_at | 349.4 | P | 479.7 | P | 362.2 | P | 371.7 | P | STZ |
| 261649_at | 365 | P | 389.5 | P | 451.3 | P | 367.9 | P | --- |
| 261650_at | 1557.9 | P | 1754 | P | 1443.2 | P | 1390.5 | P | ACA1 |
| 261651_at | 1446.7 | P | 980.1 | P | 1068.4 | P | 1176.6 | P | SAT32 |
| 261652_at | 229.9 | P | 264.1 | P | 246.1 | P | 274.8 | P | PFC1 |
| 261653_at | 181.3 | P | 222.3 | P | 176.6 | P | 276.8 | P | SBTI1.1 |
| 261654_at | 144.1 | P | 91.1 | P | 133.4 | P | 88.6 | P | --- |
| 261655_at | 263.9 | P | 337.1 | P | 327.8 | P | 416 | P | --- |
| 261656_at | 302 | P | 218.6 | P | 328.2 | P | 204.6 | P | ARK2 |
| 261657_at | 5.9 | A | 2 | A | 5.5 | A | 3.9 | A | --- |
| 261658_at | 40.6 | A | 25.9 | A | 44.8 | A | 20.3 | A | --- |
| 261659_at | 311.6 | P | 292.6 | P | 333.2 | P | 284.1 | P | TOR |
| 261660_at | 165.9 | P | 330.8 | P | 210.8 | P | 300.7 | P | HIK |
| 261661_at | 429.7 | P | 476 | P | 477.5 | P | 489.8 | P | --- |
| 261662_at | 38.6 | A | 46.6 | A | 27.3 | A | 52.4 | A | ATMKK7 |
| 261663_at | 77.4 | P | 392 | P | 184.8 | P | 285.8 | P | EPR1 |
| 261664_s_at | 222.7 | P | 213.4 | P | 318.2 | P | 207.3 | P | --- |
| 261665_at | 0.9 | A | 6.5 | A | 20.8 | A | 0.7 | A | --- |
| 261666_at | 320.1 | P | 322.4 | P | 459.4 | P | 277.5 | P | --- |
| 261667_at | 527.5 | P | 378.6 | P | 427.3 | P | 405.9 | P | --- |
| 261668_at | 1445.1 | P | 910.9 | P | 1767.6 | P | 1050.7 | P | MAML-4 |
| 261669_at | 252.2 | P | 289.3 | P | 309.1 | P | 328.8 | P | --- |
| 261670_at | 17.9 | A | 25.8 | A | 7.7 | A | 10.4 | A | TET11 |
| 261671_at | 200.5 | P | 286.8 | P | 222.9 | P | 240.9 | P | --- |
| 261672_at | 841.6 | P | 893.9 | P | 1021.1 | P | 915.7 | P | ATARP4 |
| 261673_at | 34 | A | 35.5 | A | 10.1 | A | 36.3 | A | --- |
| 261674_at | 375.7 | P | 317.3 | P | 284.8 | P | 292.2 | P | --- |
| 261675_at | 28.6 | A | 7.9 | A | 3.1 | A | 23.2 | A | --- |
| 261676_at | 540.6 | P | 418.8 | P | 488.9 | P | 397 | P | --- |
| 261677_at | 695.6 | P | 859.1 | P | 780.1 | P | 741.6 | P | --- |
| 261678_at | 17.1 | A | 32.6 | A | 3.6 | A | 7.5 | A | --- |
| 261679_at | 22.7 | A | 1.3 | A | 1.4 | A | 5.2 | A | --- |
| 261680_at | 17.7 | A | 2.7 | A | 2.3 | A | 2.9 | A | --- |
| 261681_at | 147 | P | 156.2 | P | 194.8 | P | 180.2 | P | --- |
| 261682_at | 408.9 | P | 482.3 | P | 446.5 | P | 438.5 | P | --- |

| | | | | | | | | | |
|---|---|---|---|---|---|---|---|---|---|
| 261683_at | 9.9 | A | 2.5 | A | 1.9 | A | 0.6 | A | --- |
| 261684_at | 212 | P | 1434.1 | P | 2279.1 | P | 2152.4 | P | --- |
| 261685_at | 226.4 | P | 412.6 | P | 290 | P | 326.2 | P | AT3BETAHSD/D1 |
| 261686_at | 371.2 | P | 485.5 | P | 381.8 | P | 426.9 | P | --- |
| 261687_at | 9.1 | A | 1.6 | A | 21.9 | A | 3.1 | A | --- |
| 261688_at | 286.4 | P | 387.4 | P | 259.8 | P | 409.1 | P | --- |
| 261689_at | 469.4 | P | 502 | P | 442.7 | P | 511.6 | P | --- |
| 261690_at | 27 | A | 4.2 | A | 4.5 | A | 16.8 | A | --- |
| 261691_at | 1.9 | A | 3.1 | A | 13.5 | A | 1.3 | A | --- |
| 261692_at | 953.8 | P | 1865.2 | P | 758.7 | P | 1639.1 | P | CRT3 |
| 261693_at | 612.4 | P | 678.9 | P | 550.3 | P | 667.3 | P | DCP1 |
| 261694_at | 6214.5 | P | 5044.2 | P | 6524.8 | P | 5094.2 | P | --- |
| 261695_at | 1882.7 | P | 2280.1 | P | 1834.1 | P | 1926.8 | P | CHLD |
| 261696_at | 569.5 | P | 718.3 | P | 515.6 | P | 644.2 | P | --- |
| 261697_at | 94.3 | P | 77.4 | P | 111.6 | P | 87.1 | P | --- |
| 261698_at | 176.1 | P | 127.4 | P | 219.1 | P | 96.3 | P | --- |
| 261699_at | 21.2 | A | 6 | A | 9.4 | A | 6.4 | A | --- |
| 261700_at | 158.3 | P | 146.8 | P | 176 | P | 141.3 | P | --- |
| 261701_at | 418.6 | P | 420.6 | P | 342.1 | P | 363.7 | P | HAF01 |
| 261702_at | 26.4 | P | 34 | P | 33.9 | A | 7.3 | A | --- |
| 261703_at | 37.7 | A | 27 | A | 1.3 | A | 14.4 | A | ANAC012 |
| 261704_at | 7.2 | A | 1 | A | 3.7 | A | 2.5 | A | --- |
| 261705_at | 0.8 | A | 6.4 | A | 15.2 | A | 0.2 | A | --- |
| 261706_at | 6.7 | A | 0.8 | A | 5.6 | A | 2 | A | --- |
| 261707_at | 0.5 | A | 1.3 | A | 1 | A | 0.9 | A | --- |
| 261708_at | 190.4 | P | 71.4 | P | 151.7 | P | 82.6 | P | --- |
| 261709_at | 2854.2 | P | 2311.9 | P | 2201.1 | P | 2375.5 | P | CID11 |
| 261710_at | 15.9 | A | 47 | P | 55.1 | M | 51 | P | --- |
| 261711_at | 493.9 | P | 882.4 | P | 366.8 | P | 573.2 | P | --- |
| 261712_at | 79.3 | P | 100.6 | P | 92.7 | P | 109.3 | P | --- |
| 261713_at | 1281.5 | P | 517.9 | P | 1268.2 | P | 636 | P | MYC2 |
| 261714_at | 3 | A | 3.1 | A | 7.8 | A | 2.2 | A | TET16 |
| 261715_at | 172.6 | P | 163.4 | P | 183.4 | P | 182.1 | P | --- |
| 261716_at | 1.5 | A | 2.1 | A | 2 | A | 1.3 | A | --- |
| 261717_at | 104.6 | P | 190.6 | P | 150 | P | 110 | P | BEE1 |
| 261718_at | 291.2 | P | 241.4 | P | 191.1 | P | 248.5 | P | --- |
| 261719_at | 103.8 | P | 105.9 | P | 81.2 | P | 84 | P | --- |
| 261720_at | 650.5 | P | 639.8 | P | 673.6 | P | 510.8 | P | HDA08 |
| 261721_at | 2639.4 | P | 2495.5 | P | 2322.1 | P | 2419.3 | P | --- |
| 261722_at | 1281 | P | 1470.3 | P | 1256.6 | P | 1387.8 | P | FATB |
| 261723_at | 4786.8 | P | 4905.1 | P | 4339.8 | P | 5159.5 | P | --- |
| 261724_at | 121.1 | P | 137.5 | P | 117 | P | 146.4 | P | --- |
| 261725_at | 991.4 | P | 738.7 | P | 1066.4 | P | 1034.1 | P | SmD3 |
| 261726_at | 127.5 | P | 165.5 | P | 146.1 | P | 147.3 | P | --- |
| 261727_at | 895.7 | P | 654.8 | P | 961.3 | P | 666.7 | P | SMT3 |
| 261728_at | 2519.2 | P | 2652.4 | P | 3154.5 | P | 2437.8 | P | sks5 |
| 261729_s_at | 39.7 | A | 24.6 | P | 45.6 | P | 76.7 | P | HXK3 |
| 261730_s_at | 0.9 | A | 0.3 | A | 0.5 | A | 0.7 | A | --- |
| 261731_s_at | 1 | A | 7.1 | A | 10.8 | A | 19.5 | A | --- |
| 261732_at | 36.4 | P | 33.6 | M | 49.5 | A | 32.5 | P | --- |
| 261733_at | 690.7 | P | 908.2 | P | 893.8 | P | 855.9 | P | --- |
| 261734_at | 18.5 | A | 9.4 | A | 13 | A | 18.7 | A | --- |
| 261735_at | 2.2 | A | 15.4 | A | 5.2 | A | 1.9 | A | --- |
| 261736_at | 26.4 | A | 18.6 | A | 21.8 | A | 12.9 | A | --- |
| 261737_at | 2.9 | A | 2.5 | A | 5.7 | A | 1.2 | A | --- |
| 261738_s_at | 442.1 | P | 405 | P | 310.2 | P | 562.5 | P | --- |
| 261739_at | 296.9 | P | 392.3 | P | 271.1 | P | 440.6 | P | PEX11A |
| 261740_at | 578.4 | P | 916.4 | P | 669.6 | P | 910.8 | P | --- |
| 261741_at | 261.8 | P | 211.4 | P | 266.5 | P | 218.2 | P | ATE2F2 |
| 261742_at | 106.3 | P | 175.1 | P | 128.8 | P | 168.6 | P | --- |
| 261743_s_at | 319.1 | P | 313.4 | P | 279 | P | 303.8 | P | BSL2 |
| 261744_at | 729.9 | P | 836.2 | P | 708.7 | P | 839.8 | P | CPNIFS |
| 261745_at | 35 | P | 190.3 | P | 80.9 | A | 118.6 | P | --- |
| 261746_at | 14346.9 | P | 16474 | P | 11978.7 | P | 15889.8 | P | PSAO |
| 261747_at | 259.5 | P | 241.8 | P | 130.7 | P | 270.8 | P | --- |
| 261748_at | 242.2 | P | 129.5 | P | 181.2 | A | 113.1 | P | --- |
| 261749_at | 3151.2 | P | 3415.5 | P | 2660.5 | P | 3434.6 | P | ERD14 |
| 261750_at | 194.5 | P | 192.6 | P | 202.3 | P | 141.9 | P | --- |
| 261751_at | 6622.9 | P | 7448.7 | P | 6917 | P | 7139.6 | P | CDSP32 |
| 261752_at | 32.9 | A | 1.9 | A | 6.9 | A | 5.6 | A | --- |
| 261753_at | 233.5 | P | 262.9 | P | 186.6 | P | 301.2 | P | --- |
| 261754_at | 385.7 | P | 437.4 | P | 488.4 | P | 509.9 | P | AMY2 |
| 261755_at | 35.3 | A | 30.1 | A | 9.1 | A | 5.5 | A | --- |
| 261756_at | 333 | P | 271.4 | P | 232.9 | P | 250.7 | P | --- |
| 261757_at | 351.2 | P | 424.1 | P | 443 | P | 475 | P | --- |
| 261758_at | 579.8 | P | 352.3 | P | 372.8 | P | 356.2 | P | ADT6 |
| 261759_at | 1.3 | A | 2.3 | A | 23.4 | A | 1.9 | A | --- |
| 261760_at | 20.7 | A | 29 | P | 47.6 | A | 10.1 | A | --- |
| 261761_at | 213.6 | P | 309.9 | P | 256.1 | P | 270.9 | P | CENP-C |
| 261762_at | 205.2 | P | 177.9 | P | 257.8 | P | 187 | P | --- |
| 261763_at | 107.4 | P | 78.8 | P | 101.5 | P | 59.7 | P | PDR12 |
| 261764_at | 58 | A | 46.2 | A | 60.4 | A | 75.2 | A | --- |
| 261765_at | 82.5 | P | 168.2 | P | 137.7 | P | 136.9 | P | CYCA2;3 |
| 261766_at | 3.4 | A | 31.6 | A | 4.4 | A | 4.8 | A | IAA5 |
| 261767_s_at | 3635.2 | P | 2950.5 | P | 3282.8 | P | 3080.8 | P | NTT1 |
| 261768_at | 244.2 | P | 161.2 | P | 343.6 | P | 125 | P | GA3OX1 |
| 261769_at | 1434 | P | 2758.8 | P | 1951.6 | P | 2519.3 | P | PETE1 |
| 261770_at | 1335.2 | P | 1378 | P | 1402 | P | 1437.9 | P | --- |
| 261771_at | 1974.9 | P | 1845.2 | P | 1865.8 | P | 1765.9 | P | --- |
| 261772_at | 49.3 | M | 114.3 | P | 47.9 | A | 115.9 | P | --- |
| 261773_at | 2 | A | 10 | A | 23.5 | A | 7.6 | A | --- |
| 261774_at | 306.7 | P | 345.9 | P | 386.1 | P | 291.2 | P | --- |
| 261775_at | 55 | P | 79.9 | P | 116.8 | P | 136.9 | P | --- |
| 261776_at | 84.2 | P | 101.1 | P | 79 | P | 102.7 | P | --- |
| 261777_at | 9.1 | A | 4.4 | A | 1.5 | A | 0.7 | A | --- |
| 261778_at | 2.6 | A | 3.1 | A | 17.1 | A | 11.1 | A | --- |
| 261779_at | 10.5 | A | 2.7 | A | 16.4 | A | 1.1 | A | --- |
| 261780_at | 85.6 | P | 174.6 | P | 100.1 | P | 147.5 | P | CYCB2;4 |
| 261781_at | 257.6 | P | 284.6 | P | 237.4 | P | 251.3 | P | FRS4 |
| 261782_at | 11.8 | A | 129.3 | P | 83.2 | P | 100.7 | P | --- |
| 261783_at | 773.4 | P | 1203.8 | P | 884.6 | P | 1057.2 | P | --- |
| 261784_at | 403.8 | P | 314.4 | P | 501.9 | P | 305.3 | P | --- |
| 261785_at | 339.6 | P | 256.2 | P | 217 | P | 228.9 | P | --- |
| 261786_at | 23.8 | A | 4.6 | A | 5 | A | 1.1 | A | ATCNGC7 |
| 261787_at | 343.3 | P | 356.3 | P | 444.3 | P | 350.6 | P | --- |
| 261788_at | 2600.3 | P | 3073.6 | P | 2429.2 | P | 2887.4 | P | NDF1 |
| 261789_at | 8965 | P | 8135.8 | P | 10371.5 | P | 8758.9 | P | --- |
| 261790_at | 731.6 | P | 791.1 | P | 635.9 | P | 770.7 | P | --- |
| 261791_at | 294.1 | P | 262.7 | P | 248.7 | P | 288.6 | P | --- |

| | | | | | | | | | |
|---|---|---|---|---|---|---|---|---|---|
| 261792_at | 1613 | P | 1872.4 | P | 1568.8 | P | 1859.7 | P | CCR1 |
| 261793_at | 864.4 | P | 1730.8 | P | 1005.1 | P | 1588.1 | P | --- |
| 261794_at | 431.3 | P | 289.3 | P | 658.5 | P | 297.2 | P | --- |
| 261795_at | 884.3 | P | 963.5 | P | 999.1 | P | 943 | P | --- |
| 261796_at | 294.4 | P | 381.3 | P | 359.3 | P | 390.3 | P | --- |
| 261797_at | 20.9 | A | 1.8 | A | 15.2 | A | 20 | A | --- |
| 261798_at | 96.5 | P | 108.1 | P | 140.2 | P | 103.9 | P | CPSF30 |
| 261799_at | 22.7 | A | 2.7 | A | 22 | A | 26.2 | A | --- |
| 261800_at | 702.4 | P | 646.2 | P | 671.5 | P | 616 | P | PHV |
| 261801_at | 338.1 | P | 481.3 | P | 392.6 | P | 465.6 | P | AAE14 |
| 261802_at | 124.6 | P | 140.9 | P | 128 | P | 119.9 | P | --- |
| 261803_at | 330.5 | P | 226.8 | P | 238.3 | P | 224.9 | P | NF-YA7 |
| 261804_at | 1494.9 | P | 1293.5 | P | 1416 | P | 1845.9 | P | UGT78D1 |
| 261805_at | 286.2 | P | 345 | P | 331 | P | 287.4 | P | --- |
| 261806_at | 295.1 | P | 367.6 | P | 258 | P | 360.4 | P | ATRFNR2 |
| 261807_at | 42.7 | P | 69.7 | P | 79.7 | P | 64.4 | P | --- |
| 261808_at | 308.5 | P | 286.7 | P | 304.5 | P | 286 | P | DRT111 |
| 261809_at | 30.4 | P | 45.5 | P | 50 | A | 50.2 | P | --- |
| 261810_at | 234.8 | P | 284.6 | P | 259.4 | P | 310.3 | P | ATLIG1 |
| 261811_at | 80.2 | P | 111.9 | P | 62.3 | P | 71.6 | P | TIL1 |
| 261812_at | 10.4 | A | 23.4 | A | 38.9 | A | 54.9 | P | --- |
| 261813_at | 396.8 | P | 319.7 | P | 495.5 | P | 361.8 | P | --- |
| 261814_at | 29.9 | M | 23.1 | A | 32.9 | A | 10.7 | A | --- |
| 261815_at | 5.3 | A | 7 | A | 7.5 | A | 5.6 | A | --- |
| 261816_at | 21.7 | A | 18.6 | A | 57.7 | P | 27.5 | A | CHX6B |
| 261817_at | 98.4 | P | 120.3 | P | 88.6 | P | 97.5 | P | --- |
| 261818_at | 202.3 | P | 204.2 | P | 257.1 | P | 221.6 | P | --- |
| 261819_at | 319.6 | P | 255 | P | 319 | P | 305.5 | P | --- |
| 261820_at | 75.6 | P | 77.8 | P | 63.8 | P | 71.9 | P | --- |
| 261821_at | 912.3 | P | 1269.6 | P | 1007 | P | 1315.5 | P | ATCXXS1 |
| 261822_at | 204.6 | P | 184.6 | P | 131.6 | P | 236.7 | P | --- |
| 261823_at | 244.7 | P | 264.1 | P | 314.3 | P | 299.8 | P | PYM |
| 261824_at | 2663.3 | P | 4178.2 | P | 3365.6 | P | 4109.6 | P | --- |
| 261825_at | 139.7 | P | 337.1 | P | 146.1 | P | 335.1 | P | --- |
| 261826_at | 1465.3 | P | 509.1 | P | 2092.3 | P | 639.7 | P | PMEPCRA |
| 261827_at | 504.5 | P | 556.2 | P | 487 | P | 521.9 | P | --- |
| 261828_at | 528.6 | P | 552 | P | 442.5 | P | 540.9 | P | --- |
| 261829_at | 1.3 | A | 2 | A | 3.5 | A | 2 | A | PGP10 |
| 261830_at | 3.7 | A | 2.6 | A | 3.9 | A | 2.8 | A | --- |
| 261831_at | 1968.8 | P | 2286.2 | P | 1773.9 | P | 2166.7 | P | ATARFA1F |
| 261832_at | 409.9 | P | 553.2 | P | 373.4 | P | 515.1 | P | --- |
| 261833_at | 2591.2 | P | 2949.5 | P | 3089.5 | P | 3343.5 | P | ACLA-1 |
| 261834_at | 81.5 | P | 226.8 | P | 155.5 | P | 264.7 | P | --- |
| 261835_at | 90.6 | P | 90.8 | P | 99.7 | P | 60.7 | P | --- |
| 261836_at | 59.1 | P | 71.1 | P | 7.2 | A | 68.8 | P | WAKL7 |
| 261837_s_at | 126.1 | P | 198.2 | P | 173.4 | P | 181.7 | P | --- |
| 261838_at | 183.5 | P | 128.6 | P | 198 | P | 189 | P | Hsp70b |
| 261839_at | 274.2 | P | 270.3 | P | 279.2 | P | 279 | P | --- |
| 261840_at | 97.3 | P | 114.8 | P | 55.8 | P | 98.8 | P | AtTLP8 |
| 261841_at | 522 | P | 440.6 | P | 420.6 | P | 380.8 | P | --- |
| 261842_at | 15.1 | A | 33.6 | M | 38.4 | A | 1.4 | A | WAKL5 |
| 261843_at | 4166 | P | 3852.1 | P | 3431.6 | P | 3466.6 | P | --- |
| 261844_at | 169.6 | P | 176.5 | P | 189.8 | P | 196.3 | P | --- |
| 261845_at | 158.1 | M | 155.9 | P | 224 | P | 153.5 | P | NRAMP6 |
| 261846_at | 86.9 | A | 72.1 | A | 114.7 | P | 58.1 | A | --- |
| 261847_at | 12.9 | P | 24.4 | P | 25.1 | A | 40.4 | A | --- |
| 261848_at | 2.7 | A | 6.9 | A | 5.1 | A | 1.5 | A | --- |
| 261849_at | 2.7 | A | 16.7 | A | 3.3 | A | 2.6 | A | --- |
| 261850_at | 8.8 | A | 16.9 | A | 9.1 | A | 28.7 | A | --- |
| 261851_at | 264.2 | P | 235.3 | P | 410.1 | P | 215.2 | P | HKL1 |
| 261852_at | 517.5 | P | 668.1 | P | 413.4 | P | 590 | P | --- |
| 261853_at | 394.7 | P | 502.3 | P | 472.5 | P | 556.9 | P | --- |
| 261854_at | 670.8 | P | 824.5 | P | 639.5 | P | 778.3 | P | --- |
| 261855_at | 365.3 | P | 505.9 | P | 346.3 | P | 394.6 | P | --- |
| 261856_at | 10.6 | A | 19 | P | 20.9 | P | 1.6 | A | --- |
| 261857_at | 109.8 | P | 156.9 | P | 157.8 | P | 156.9 | P | --- |
| 261858_at | 669.2 | P | 791.8 | P | 534.8 | P | 687.3 | P | --- |
| 261859_at | 482.8 | P | 707.1 | P | 520.7 | P | 661.6 | P | UBC20 |
| 261860_at | 485.7 | P | 402.2 | P | 343 | P | 447.9 | P | SCL5 |
| 261861_at | 849.5 | P | 1155.9 | P | 1160.1 | P | 1228.9 | P | --- |
| 261862_at | 335.8 | P | 305.9 | P | 238.6 | P | 264.2 | P | --- |
| 261863_at | 219.6 | P | 166.3 | P | 176.4 | P | 235.9 | P | --- |
| 261864_s_at | 3874.5 | P | 3671.8 | P | 3646.8 | P | 3257.1 | P | THFS |
| 261865_at | 1897.4 | P | 1664.8 | P | 1958 | P | 1694 | P | DWF5 |
| 261866_at | 916.2 | P | 412.6 | P | 1094.9 | P | 550.9 | P | SCL3 |
| 261867_at | 651.7 | P | 811.2 | P | 1077 | P | 916 | P | --- |
| 261868_s_at | 55.7 | P | 62.4 | P | 67.6 | P | 51.3 | P | --- |
| 261869_at | 2.2 | A | 3.5 | A | 20.3 | A | 3.4 | A | --- |
| 261870_at | 37.3 | A | 45.5 | A | 16 | A | 57.2 | A | NTL |
| 261871_at | 353.7 | P | 293.5 | P | 331.2 | P | 280.1 | P | --- |
| 261872_s_at | 510.1 | P | 563.3 | P | 722.1 | P | 490.1 | P | --- |
| 261873_at | 439.2 | P | 402.4 | P | 457.7 | P | 392.7 | P | SD1-13 |
| 261874_at | 757.8 | P | 677.7 | P | 577.4 | P | 733.8 | P | ERF3 |
| 261875_at | 7.4 | A | 24.1 | A | 5.8 | A | 6 | A | --- |
| 261876_at | 47.8 | P | 44.6 | P | 64.3 | P | 47 | P | --- |
| 261877_at | 1.9 | A | 1.9 | A | 6.8 | A | 1.8 | A | --- |
| 261878_at | 20.5 | A | 1.1 | A | 26.3 | A | 25.3 | A | CYP705A25 |
| 261879_at | 53.6 | A | 37.4 | A | 51.3 | A | 44.4 | A | CYP705A27 |
| 261880_at | 411.4 | P | 467.6 | P | 403.5 | P | 390.1 | P | HIT1 |
| 261881_at | 38.6 | A | 123.8 | P | 57 | A | 104.1 | P | NIP6;1 |
| 261882_at | 535.4 | P | 777.2 | P | 700.8 | P | 736.2 | P | PDE318 |
| 261883_at | 152.3 | P | 186.2 | P | 226.2 | P | 222.4 | P | --- |
| 261884_at | 344.8 | P | 458.7 | P | 550.1 | P | 515.1 | P | --- |
| 261885_at | 796.4 | P | 897.2 | P | 750.2 | P | 809 | P | --- |
| 261886_s_at | 566.9 | P | 482.2 | P | 598.7 | P | 518.5 | P | --- |
| 261887_at | 1623 | P | 1722.3 | P | 1730.6 | P | 1603.7 | P | --- |
| 261888_at | 153.2 | P | 317.2 | P | 195.4 | P | 332.6 | P | --- |
| 261889_at | 76.4 | P | 89.9 | P | 52.6 | M | 32.6 | P | --- |
| 261890_at | 28 | A | 32.3 | A | 3 | A | 14.1 | A | --- |
| 261891_at | 469.1 | P | 704.9 | P | 553.6 | P | 716.9 | P | --- |
| 261892_at | 280.7 | P | 439.4 | P | 253.4 | P | 323 | P | WRKY40 |
| 261893_at | 105.9 | P | 161.6 | P | 131.5 | P | 130.9 | P | --- |
| 261894_at | 108.3 | P | 125.2 | P | 147.2 | P | 92.1 | P | MGT1 |
| 261895_at | 623 | P | 763.3 | P | 734.1 | P | 786.5 | P | NRAMP1 |
| 261896_at | 733.9 | P | 975.2 | P | 955 | P | 998.1 | P | --- |
| 261897_at | 769.9 | P | 802.8 | P | 651.5 | P | 682.3 | P | --- |
| 261898_at | 153.4 | P | 145.1 | P | 187.1 | P | 162.7 | P | --- |
| 261899_at | 73.1 | P | 6.7 | A | 96 | A | 39.2 | P | CCR2 |
| 261900_at | 346.5 | P | 324 | P | 316 | P | 406.7 | P | --- |

| | | | | | | | | | |
|---|---|---|---|---|---|---|---|---|---|
| 261901_at | 2181.4 | P | 2338.2 | P | 1076.9 | P | 1469 | P | J8 |
| 261902_at | 632.7 | P | 669.1 | P | 780 | P | 763.8 | P | PLMT |
| 261903_at | 22.2 | A | 40.6 | A | 7.5 | A | 26.9 | A | CMT1 |
| 261904_at | 416.6 | P | 369 | P | 366.6 | P | 317.4 | P | AT1G65040 |
| 261905_at | 292.6 | P | 354.3 | P | 347.9 | P | 327 | P | --- |
| 261906_at | 122 | P | 92 | P | 142 | P | 101.5 | P | --- |
| 261907_at | 737.7 | P | 486.2 | P | 1021.1 | P | 869.7 | P | 4CL3 |
| 261908_at | 33 | A | 8.6 | A | 12.3 | A | 26.6 | A | --- |
| 261909_at | 654.4 | P | 637.4 | P | 768.2 | P | 636.9 | P | SAR3 |
| 261910_at | 1.6 | A | 9.7 | A | 20.2 | A | 4.3 | A | ZFP1 |
| 261911_at | 1478.3 | P | 1136.1 | P | 2043.2 | P | 1196.6 | P | --- |
| 261912_s_at | 12.8 | A | 81.9 | P | 24.8 | A | 63.3 | P | --- |
| 261913_at | 1234 | P | 642.9 | P | 2050.4 | P | 633 | P | FMO GS-OX1 |
| 261914_at | 40.5 | A | 54.8 | P | 90.2 | P | 62.2 | P | --- |
| 261915_at | 1.7 | A | 2.5 | A | 3.4 | A | 1.7 | A | BZO1 |
| 261916_at | 40.9 | A | 67 | P | 45.2 | P | 65.4 | P | anac028 |
| 261917_at | 104.1 | P | 112.9 | P | 127.8 | A | 81.7 | P | --- |
| 261918_at | 4.3 | A | 0.3 | A | 0.3 | A | 2.7 | A | --- |
| 261919_at | 7057.1 | P | 5932.9 | P | 7019.8 | P | 6073.6 | P | TPX1 |
| 261920_at | 6748.8 | P | 6754.2 | P | 7011.4 | P | 6257 | P | --- |
| 261921_at | 128.7 | P | 244.1 | P | 217.8 | P | 291.5 | P | --- |
| 261922_at | 71.1 | P | 2.8 | A | 93.6 | A | 16.4 | A | AAE12 |
| 261923_at | 1842.3 | P | 219.1 | P | 1282.1 | P | 241.5 | P | AtUGT85A3 |
| 261924_at | 165.8 | P | 91.7 | P | 187.1 | P | 137.7 | P | --- |
| 261925_at | 251 | P | 358.3 | P | 268.5 | P | 286 | P | --- |
| 261926_at | 1509.1 | P | 1072.1 | P | 1405.7 | P | 1255.6 | P | PATL2 |
| 261927_at | 2174.6 | P | 590.6 | P | 1966.2 | P | 871.2 | P | --- |
| 261928_at | 57.3 | A | 14.8 | A | 23.1 | A | 3.4 | A | --- |
| 261929_at | 5.1 | A | 23.3 | A | 20 | A | 9.6 | A | --- |
| 261930_at | 30.6 | A | 73.9 | P | 81.3 | P | 75.8 | P | --- |
| 261931_at | 852.7 | P | 864.5 | P | 955.6 | P | 851 | P | --- |
| 261932_at | 3 | A | 20.3 | A | 25.4 | A | 38.2 | A | --- |
| 261933_at | 2953.7 | P | 3273.7 | P | 3076.5 | P | 3249.4 | P | --- |
| 261934_at | 1810.4 | P | 1262.5 | P | 1311.8 | P | 1042.2 | P | UGT85A1 |
| 261935_at | 1102.2 | P | 955.4 | P | 1180.3 | P | 855 | P | --- |
| 261936_at | 38.3 | A | 53.4 | A | 64.7 | M | 33.7 | A | --- |
| 261937_at | 198.2 | P | 471.5 | P | 225.5 | P | 390.3 | P | --- |
| 261938_at | 379.4 | P | 443.7 | P | 276.6 | P | 355.4 | P | --- |
| 261939_at | 2488.5 | P | 2566.1 | P | 2902.9 | P | 2754.4 | P | COX6B |
| 261940_at | 871.7 | P | 981.8 | P | 1270.2 | P | 1119.6 | P | --- |
| 261941_at | 147.7 | P | 50.2 | P | 66.3 | P | 95.8 | P | --- |
| 261942_at | 344.9 | P | 360.4 | P | 504.4 | P | 423.2 | P | AGL87 |
| 261943_at | 10 | A | 22 | A | 3.3 | A | 2.7 | A | AHA9 |
| 261944_at | 1228 | P | 1504.1 | P | 775.1 | P | 1372.5 | P | --- |
| 261945_at | 79.7 | A | 114.1 | P | 106.3 | P | 134.5 | P | --- |
| 261946_at | 8.1 | A | 7.7 | A | 2.4 | A | 2.8 | A | --- |
| 261947_at | 63 | P | 102.8 | P | 89.5 | P | 121.9 | P | --- |
| 261948_at | 1674.7 | P | 1705.8 | P | 2013.9 | P | 1644.1 | P | --- |
| 261949_at | 221.5 | P | 287.5 | P | 296.3 | P | 370.5 | P | BDG1 |
| 261950_at | 101.3 | P | 124.8 | P | 159.2 | P | 130.1 | P | --- |
| 261951_at | 177 | P | 129.1 | P | 215.1 | P | 150.3 | P | --- |
| 261952_at | 555.4 | P | 517.7 | M | 572.5 | M | 526.5 | P | --- |
| 261953_at | 449.6 | P | 493.5 | P | 428.9 | P | 519 | P | RHD1 |
| 261954_at | 4703.8 | P | 6047.5 | P | 4109.7 | P | 5227.7 | P | --- |
| 261955_at | 1635.4 | P | 1871.1 | P | 1722.4 | P | 1731.8 | P | RPN12a |
| 261956_at | 23 | A | 72.2 | P | 72.9 | A | 71.2 | P | --- |
| 261957_at | 921.2 | P | 265.1 | P | 498.2 | P | 520.4 | P | ATMGL |
| 261958_at | 200.1 | P | 347.5 | P | 361.1 | P | 346.8 | P | --- |
| 261959_s_at | 5 | A | 4.2 | A | 5.8 | A | 4 | A | --- |
| 261960_at | 13.3 | A | 13 | A | 9.3 | A | 19.1 | P | --- |
| 261961_at | 3 | A | 4.2 | A | 6.2 | A | 2.3 | A | --- |
| 261962_s_at | 1.1 | A | 2.8 | A | 2.5 | A | 1.4 | A | --- |
| 261963_at | 2.3 | A | 6.1 | A | 2.2 | A | 16.2 | A | --- |
| 261964_at | 3.5 | A | 5.3 | A | 41.8 | A | 19.7 | A | --- |
| 261965_at | 0.4 | A | 1.8 | A | 1.4 | A | 4.2 | A | --- |
| 261966_x_at | 28.5 | P | 1.7 | A | 31.3 | A | 12.5 | A | --- |
| 261967_at | 4.7 | A | 5.4 | A | 26.1 | A | 2.9 | A | --- |
| 261968_at | 23.7 | A | 23 | A | 30.6 | A | 30 | A | --- |
| 261969_at | 378.5 | P | 435.4 | P | 272.6 | P | 320.4 | P | --- |
| 261970_at | 7964.6 | P | 8643 | P | 6945.7 | P | 9167.3 | P | GAD2 |
| 261971_at | 2 | A | 2.2 | A | 6.6 | A | 2.7 | A | --- |
| 261972_at | 221.9 | P | 185.3 | P | 258.5 | P | 193.8 | P | --- |
| 261973_at | 193.2 | P | 167.8 | P | 157 | P | 145.4 | P | --- |
| 261974_at | 325.4 | P | 345.1 | P | 454.6 | P | 363.7 | P | CPuORF57 |
| 261975_at | 181.4 | P | 429 | P | 341.3 | P | 447.3 | P | --- |
| 261976_at | 0.7 | A | 2 | A | 3.4 | A | 4.4 | A | --- |
| 261977_at | 35.8 | A | 24.5 | A | 43.6 | A | 6.9 | A | --- |
| 261978_at | 0.6 | A | 1.1 | A | 0.7 | A | 0.3 | A | --- |
| 261979_at | 4974.2 | P | 2945.4 | P | 4979.3 | P | 3435.5 | P | NIA2 |
| 261980_at | 9 | A | 15.5 | A | 6.5 | A | 14.5 | A | --- |
| 261981_at | 960.7 | P | 1211.3 | P | 1283.7 | P | 1206 | P | --- |
| 261982_at | 1204.8 | P | 1546.9 | P | 1077.7 | P | 1248.5 | P | --- |
| 261983_at | 31.7 | A | 21.2 | A | 3.8 | A | 23.4 | A | --- |
| 261984_at | 2.8 | A | 2.2 | A | 2.2 | A | 1.6 | A | --- |
| 261985_at | 1.5 | A | 2.7 | A | 2.2 | A | 2.2 | A | --- |
| 261986_s_at | 52.6 | P | 69.9 | P | 44 | A | 72.6 | P | CYP76C5 |
| 261987_at | 4.5 | A | 1.7 | A | 3 | A | 1.5 | A | --- |
| 261988_at | 313.5 | P | 269.1 | P | 281.2 | P | 263.1 | P | --- |
| 261989_at | 1.6 | A | 2.7 | A | 5.8 | A | 4.2 | A | --- |
| 261990_at | 148.6 | P | 99.9 | P | 139.5 | M | 126.7 | P | --- |
| 261991_at | 3 | A | 12.7 | A | 19.7 | A | 33.7 | A | --- |
| 261992_at | 373.2 | P | 410.4 | P | 459.7 | P | 457.1 | P | --- |
| 261993_at | 4.9 | A | 4.5 | A | 7.8 | A | 13.1 | A | --- |
| 261994_at | 21.6 | A | 4.1 | A | 40.4 | A | 2.9 | A | --- |
| 261995_at | 0.5 | A | 0.9 | A | 0.3 | A | 0.6 | A | --- |
| 261996_at | 5 | A | 4 | A | 3.2 | A | 11.3 | A | --- |
| 261997_at | 1.9 | A | 2.3 | A | 30.3 | A | 11 | A | --- |
| 261998_at | 2.4 | A | 2.1 | A | 9.5 | A | 11.8 | A | --- |
| 261999_at | 52.8 | P | 106.8 | P | 96.6 | P | 83.4 | P | --- |
| 262000_at | 1992.1 | P | 1860.7 | P | 1904.2 | P | 1941.5 | P | --- |
| 262001_at | 34.7 | A | 181.8 | P | 80.7 | A | 97.8 | P | --- |
| 262002_at | 117.3 | A | 231.5 | A | 208 | A | 197.7 | A | --- |
| 262003_at | 79.9 | P | 121.6 | P | 100.2 | P | 117.6 | P | --- |
| 262004_at | 49.5 | A | 15.6 | A | 53.7 | A | 37.3 | A | CBL8 |
| 262005_at | 1217.3 | P | 998.2 | P | 1226.8 | P | 1059.7 | P | ATGCN3 |
| 262006_at | 111.7 | P | 105.1 | P | 79.8 | P | 93.3 | P | --- |
| 262007_at | 142.5 | P | 139 | P | 170.1 | P | 175.5 | P | --- |
| 262008_s_at | 0.4 | A | 0.5 | A | 1 | A | 0.8 | A | --- |
| 262009_at | 6.9 | A | 23.4 | A | 39.5 | A | 4.6 | A | --- |

| | | | | | | | | | |
|---|---|---|---|---|---|---|---|---|---|
| 262010_at | 27.9 | P | 120.9 | P | 50.2 | P | 97.3 | P | --- |
| 262011_at | 5 | A | 1.7 | A | 2 | A | 3.6 | A | --- |
| 262012_s_at | 34.3 | A | 19.6 | A | 5.7 | A | 19.9 | A | --- |
| 262013_s_at | 10.3 | A | 32.1 | A | 18.6 | A | 9 | A | --- |
| 262014_at | 66.1 | P | 88.9 | P | 94.9 | P | 87 | P | --- |
| 262015_at | 2.8 | A | 1.8 | A | 3.9 | A | 5.3 | A | --- |
| 262016_at | 35.7 | P | 48.1 | P | 51.2 | P | 31.7 | P | --- |
| 262017_at | 268.8 | P | 186.2 | P | 234.8 | P | 247.5 | P | --- |
| 262018_at | 44.6 | P | 31.2 | P | 61 | A | 34.6 | A | --- |
| 262019_s_at | 9.8 | A | 3.3 | A | 19.6 | A | 2 | A | --- |
| 262020_at | 1.3 | A | 18.9 | A | 12.1 | A | 1 | A | --- |
| 262021_at | 32.8 | A | 2.6 | A | 35.9 | A | 36.5 | A | --- |
| 262022_at | 9.7 | A | 12.3 | A | 9.8 | A | 10.7 | A | --- |
| 262023_at | 3.2 | A | 4.1 | A | 4.2 | A | 17.4 | A | --- |
| 262024_at | 921.8 | P | 1096.2 | P | 720.4 | P | 985 | P | ATPDIL5-2 |
| 262025_at | 233.5 | P | 259.2 | P | 254.7 | P | 252.2 | P | --- |
| 262026_at | 495.8 | P | 404.1 | P | 341.3 | P | 344.9 | P | ATCDPK2 |
| 262027_at | 24.3 | A | 13.2 | A | 34.5 | A | 15.6 | A | HOS10 |
| 262028_at | 112.6 | P | 152.8 | P | 119.3 | P | 124.3 | P | --- |
| 262029_at | 5631.8 | P | 7785 | P | 8114 | P | 7182.3 | P | --- |
| 262030_s_at | 15.9 | A | 9.9 | A | 20.5 | A | 4 | A | --- |
| 262031_x_at | 39 | A | 35 | A | 8.6 | A | 19.5 | A | --- |
| 262032_at | 1.8 | A | 5.5 | A | 26.9 | A | 6.1 | A | HCS2 |
| 262033_at | 5.1 | A | 6.5 | A | 1.9 | A | 5.7 | A | MCT1 |
| 262034_at | 3 | A | 1.3 | A | 14.3 | A | 16.5 | A | --- |
| 262035_at | 0.7 | A | 2.8 | A | 5 | A | 7.3 | A | --- |
| 262036_at | 29.9 | P | 46.7 | P | 55.2 | A | 28.8 | A | --- |
| 262037_at | 4.7 | A | 1.5 | A | 2.6 | A | 1.5 | A | --- |
| 262038_at | 772.3 | P | 862.9 | P | 742 | P | 748.2 | P | CINV1 |
| 262039_at | 713.3 | P | 619.5 | P | 975.6 | P | 792.9 | P | APT2 |
| 262040_at | 207.7 | P | 254.9 | P | 331.1 | P | 264.9 | P | TMM |
| 262041_at | 2.6 | A | 2.2 | A | 4.4 | A | 4.5 | A | AHP6 |
| 262042_at | 7.4 | A | 24.6 | A | 31.9 | A | 25.8 | A | --- |
| 262043_at | 129.2 | P | 161.6 | P | 144.1 | P | 170.5 | P | PSF1 |
| 262044_s_at | 418.9 | P | 381.1 | P | 292.2 | P | 373.2 | P | --- |
| 262045_at | 44.9 | A | 17 | A | 22.5 | A | 3 | A | --- |
| 262046_at | 39.7 | A | 19.4 | A | 5.6 | A | 37.2 | A | OFP14 |
| 262047_at | 520.4 | P | 133.5 | P | 131 | P | 141.4 | P | --- |
| 262048_at | 1651.4 | P | 1430.7 | P | 1530.6 | P | 1563.8 | P | --- |
| 262049_at | 763.7 | P | 554.1 | P | 663.3 | P | 515.8 | P | --- |
| 262050_at | 828.5 | P | 1550.7 | P | 742.2 | P | 2084.8 | P | --- |
| 262051_at | 607.8 | P | 727.5 | P | 561.5 | P | 698.9 | P | --- |
| 262052_at | 186.8 | P | 203.6 | P | 176.9 | P | 221.4 | P | --- |
| 262053_at | 1005.4 | P | 999.1 | P | 1072 | P | 1004.3 | P | ATERDJ2A |
| 262054_s_at | 2564.4 | P | 2734.6 | P | 3038.5 | P | 3019.2 | P | --- |
| 262055_at | 856.6 | P | 1170.8 | P | 870.5 | P | 1235.7 | P | --- |
| 262056_at | 215.9 | P | 181.6 | P | 231.4 | P | 197.3 | P | --- |
| 262057_at | 1258 | P | 1192.2 | P | 1130.7 | P | 1249.8 | P | --- |
| 262058_at | 418.3 | P | 524.9 | P | 448.3 | P | 443.5 | P | --- |
| 262059_at | 809 | P | 753.6 | P | 931.4 | P | 861.9 | P | --- |
| 262060_at | 116.5 | P | 94 | P | 82.2 | P | 102.2 | P | --- |
| 262061_at | 154.9 | P | 73.6 | P | 101 | P | 86.8 | P | ATPP2-B11 |
| 262062_s_at | 151.4 | P | 155 | P | 146.8 | P | 170.9 | P | emb1427 |
| 262063_at | 357.8 | P | 538.4 | P | 362.2 | P | 466.7 | P | --- |
| 262064_at | 13147.5 | P | 12242.1 | P | 15570.7 | P | 13408.1 | P | LOS1 |
| 262065_at | 184 | P | 236.8 | P | 211.8 | P | 198.5 | P | --- |
| 262066_at | 460.6 | P | 305.8 | P | 405.5 | P | 323.7 | P | --- |
| 262067_at | 49.5 | A | 81.8 | P | 51.7 | P | 61.2 | P | --- |
| 262068_at | 1425.6 | P | 1081.3 | P | 1572.5 | P | 1115.4 | P | SUS2 |
| 262069_at | 36.5 | A | 15.2 | A | 18.9 | A | 31.6 | A | --- |
| 262070_at | 293.6 | P | 270.9 | P | 247.3 | P | 228.7 | P | FRS8 |
| 262071_at | 92.6 | P | 84.8 | P | 119.1 | P | 137.1 | P | CF9 |
| 262072_at | 101.2 | A | 51.6 | A | 10.5 | A | 34.1 | A | ZCF37 |
| 262073_at | 151.2 | P | 113 | P | 139.2 | P | 125 | P | ZCW32 |
| 262074_at | 38.9 | P | 49.6 | P | 26.8 | A | 68.4 | P | ATGSTU15 |
| 262075_at | 205.5 | P | 184.3 | P | 193.1 | P | 214.6 | P | ZCF61 |
| 262076_at | 422.3 | P | 536.3 | P | 443.3 | P | 446.5 | P | ATMPK2 |
| 262077_at | 1002.6 | P | 953.6 | P | 823.6 | P | 971 | P | ADL3 |
| 262078_at | 507.1 | P | 605 | P | 585.6 | P | 623.3 | P | CW7 |
| 262079_at | 278.6 | P | 253.5 | P | 271.5 | P | 252.7 | P | ZCW7 |
| 262080_at | 389.6 | P | 305.4 | P | 295.7 | P | 309.9 | P | CW14 |
| 262081_at | 88.3 | P | 143.3 | P | 76 | P | 106.5 | P | ZCF125 |
| 262082_s_at | 346.5 | P | 330.5 | P | 297.6 | P | 305.4 | P | --- |
| 262083_at | 2.5 | A | 9.2 | A | 16.5 | A | 2.8 | A | --- |
| 262084_at | 63.1 | P | 70 | P | 152.1 | P | 82.1 | P | --- |
| 262085_at | 114.6 | P | 152 | P | 111.5 | P | 102.6 | P | --- |
| 262086_at | 1003 | P | 1142.3 | P | 1151.3 | P | 1178.8 | P | --- |
| 262087_at | 5.5 | A | 1.2 | A | 27.2 | A | 18.1 | A | --- |
| 262088_at | 129.8 | A | 170.7 | M | 115 | A | 170.2 | M | --- |
| 262089_s_at | 421.6 | P | 415.3 | P | 461.8 | P | 399.4 | P | --- |
| 262090_at | 93.8 | P | 61.4 | P | 129.2 | P | 92.9 | P | HAC4 |
| 262091_at | 3.3 | A | 3.9 | A | 36.4 | A | 2.6 | A | MYB72 |
| 262092_at | 112.4 | P | 650.1 | P | 164.4 | P | 470.3 | P | --- |
| 262093_at | 229.6 | P | 201.8 | P | 264.6 | P | 189.6 | P | --- |
| 262094_at | 1679.1 | P | 1447.4 | P | 3306.3 | P | 1631.3 | P | NOP56 |
| 262095_at | 189.7 | P | 248.4 | P | 223 | P | 234.2 | P | --- |
| 262096_at | 108.3 | P | 164.5 | P | 108.7 | A | 127.9 | P | NAC1 |
| 262097_at | 1 | A | 15.7 | A | 0.7 | A | 1.7 | A | --- |
| 262098_at | 380.7 | P | 337.3 | P | 365.6 | P | 451.2 | P | NF-YC2 |
| 262099_s_at | 38.4 | A | 4 | A | 14.1 | A | 5.2 | A | GH3.4 |
| 262100_s_at | 5 | A | 51.1 | A | 29.9 | A | 8.4 | A | --- |
| 262101_at | 1.6 | A | 0.7 | A | 15.9 | A | 1.2 | A | --- |
| 262102_at | 26 | A | 2.7 | A | 16.7 | A | 27.2 | A | CUL2 |
| 262103_at | 163 | P | 68.6 | P | 131.2 | P | 78.8 | P | GSTF5 |
| 262104_at | 665.2 | P | 742 | P | 547.8 | P | 683 | P | LPA1 |
| 262105_at | 57.2 | P | 37.7 | P | 61.2 | P | 39.3 | P | --- |
| 262106_at | 78.8 | P | 71 | P | 89.4 | P | 86 | P | WEE1 |
| 262107_at | 418.8 | P | 333.5 | P | 443 | P | 318.1 | P | --- |
| 262108_at | 137.2 | P | 98.1 | P | 83.4 | P | 82.1 | P | --- |
| 262109_at | 338.9 | P | 552.3 | P | 412.3 | P | 671 | P | ATCSLD5 |
| 262110_at | 1090.6 | P | 1087.2 | P | 1237.4 | P | 1085.1 | P | SR1 |
| 262111_at | 232.7 | P | 216.9 | P | 255 | P | 245.1 | P | --- |
| 262112_at | 1250 | P | 901.8 | P | 1152.1 | P | 945.2 | P | --- |
| 262113_at | 9.2 | A | 35.5 | A | 6 | A | 40.3 | A | --- |
| 262114_at | 372.9 | P | 287.8 | P | 315.8 | P | 300.4 | P | NLA |
| 262115_at | 23.8 | M | 13 | A | 30.8 | P | 35.1 | M | --- |
| 262116_at | 708.2 | P | 588.2 | P | 465.2 | P | 695.6 | P | --- |
| 262117_at | 11236.1 | P | 10388.7 | P | 12203.3 | P | 11266.4 | P | emb2386 |
| 262118_at | 8508.7 | P | 2380.8 | P | 9029.6 | P | 3646.3 | P | BGLU11 |

| | | | | | | | | | |
|---|---|---|---|---|---|---|---|---|---|
| 262119_s_at | 11514.1 | P | 8500 | P | 11933 | P | 7050 | P | ATGSTF6 |
| 262120_at | 43.9 | P | 56.3 | P | 65.2 | P | 70.9 | P | ATGSTF4 |
| 262121_at | 27.3 | A | 93.6 | A | 46.2 | A | 66.6 | A | ATCEL2 |
| 262122_at | 2.2 | A | 32.8 | A | 36.4 | A | 20.5 | A | PGA4 |
| 262123_at | 2.6 | A | 4 | A | 27.6 | A | 0.8 | A | --- |
| 262124_at | 67.8 | P | 111.8 | P | 79.7 | P | 104.3 | P | --- |
| 262125_at | 9.7 | A | 13.9 | A | 6.6 | A | 15.8 | A | EDA1 |
| 262126_at | 107.1 | P | 102 | P | 88.7 | P | 77.9 | P | CW9 |
| 262127_at | 205.9 | P | 161 | P | 148.9 | P | 87.9 | P | --- |
| 262128_at | 55.7 | P | 27 | A | 45.4 | A | 26.8 | A | --- |
| 262129_at | 148 | P | 134.2 | P | 110.5 | P | 97.9 | P | ATMMH-1 |
| 262130_at | 645.2 | P | 604 | P | 579.9 | P | 580.9 | P | --- |
| 262131_at | 114.3 | P | 107.7 | P | 81.5 | P | 133.7 | P | RALF1 |
| 262132_at | 28.1 | A | 23.6 | A | 38 | A | 37.5 | A | --- |
| 262133_at | 67 | P | 122.9 | P | 96.6 | P | 128.2 | P | SULTR1;2 |
| 262134_at | 165.9 | P | 133.4 | P | 218.2 | P | 176.7 | P | AST56 |
| 262135_at | 2009.5 | P | 1646.4 | P | 1384.1 | P | 1547.4 | P | RAP2.4 |
| 262136_at | 173.9 | P | 175.8 | P | 135.9 | P | 180.7 | P | ARF17 |
| 262137_at | 1970.4 | P | 1260.6 | P | 1326.7 | P | 1337 | P | --- |
| 262138_at | 17.1 | A | 2.5 | A | 37.3 | A | 3.9 | A | --- |
| 262139_at | 17.1 | A | 18.6 | A | 40.8 | A | 3.8 | A | --- |
| 262140_at | 13.9 | A | 12.5 | A | 33.4 | A | 28.4 | A | --- |
| 262141_s_at | 1.5 | A | 1.7 | A | 3.4 | A | 5.5 | A | --- |
| 262142_at | 46.2 | P | 35.7 | P | 38.4 | P | 55.7 | P | --- |
| 262143_at | 97.5 | P | 144.1 | P | 99.6 | P | 137 | P | --- |
| 262144_at | 50.4 | P | 59.4 | P | 56.4 | P | 72.9 | P | --- |
| 262145_at | 802.7 | P | 739.8 | P | 928.7 | P | 745.7 | P | --- |
| 262146_at | 7.6 | A | 20.9 | A | 59.6 | A | 24.3 | A | ATRBL5 |
| 262147_at | 116.5 | P | 92.2 | P | 86.2 | P | 69.2 | P | PLDALPHA2 |
| 262148_at | 2.5 | A | 0.7 | A | 4.2 | A | 0.8 | A | --- |
| 262149_at | 71.5 | A | 86.5 | A | 99.4 | A | 73.5 | A | --- |
| 262150_at | 80.2 | P | 74.5 | P | 87.2 | P | 73.6 | P | FRS6 |
| 262151_at | 1103.5 | P | 1489.9 | P | 1152.6 | P | 1407.4 | P | --- |
| 262152_s_at | 40 | P | 43.9 | P | 34 | P | 69 | P | --- |
| 262153_at | 197.1 | P | 231.1 | P | 281.9 | P | 177.7 | P | --- |
| 262154_at | 78.2 | A | 75.7 | P | 131.2 | A | 84.9 | A | --- |
| 262155_at | 1045.6 | P | 1133.7 | P | 1071.3 | P | 1200.8 | P | --- |
| 262156_at | 7 | A | 11 | A | 6.3 | A | 12 | A | --- |
| 262157_at | 896.5 | P | 943.9 | P | 977.4 | P | 866.2 | P | --- |
| 262158_at | 227.6 | P | 211 | P | 172.7 | P | 228.2 | P | --- |
| 262159_at | 359.6 | P | 775.4 | P | 353.3 | P | 714.6 | P | --- |
| 262160_at | 1108.1 | P | 1205.1 | P | 1102.6 | P | 1164.2 | P | --- |
| 262161_at | 1754.8 | P | 1952.7 | P | 1817.5 | P | 1934.5 | P | --- |
| 262162_at | 1755.9 | P | 1800.4 | P | 1948.4 | P | 1923.1 | P | --- |
| 262163_at | 6932.3 | P | 6172.4 | P | 8158.9 | P | 6894.4 | P | --- |
| 262164_at | 127.2 | P | 273.7 | P | 104.3 | P | 270.8 | P | --- |
| 262165_at | 350.2 | P | 401.6 | P | 313.3 | P | 314.8 | P | LPAT4 |
| 262166_at | 414.4 | P | 544.3 | P | 494.1 | P | 481.2 | P | --- |
| 262167_at | 28.4 | P | 28.1 | M | 22.6 | A | 34.3 | M | --- |
| 262168_at | 4791.2 | P | 4812.9 | P | 4229.3 | P | 4947.7 | P | --- |
| 262169_at | 154.5 | P | 196.7 | P | 200.1 | P | 144 | P | --- |
| 262170_at | 284 | P | 418.5 | P | 247.7 | P | 437.8 | P | --- |
| 262171_at | 1577.4 | P | 909.3 | P | 1536.6 | P | 969.8 | P | TIFY10B |
| 262172_at | 4321.6 | P | 5118.1 | P | 3694.5 | P | 4856.6 | P | RPS9 |
| 262173_at | 3934.3 | P | 3301.6 | P | 3033.6 | P | 3289.9 | P | ALDH10A8 |
| 262174_at | 1564.9 | P | 1505.3 | P | 1681.6 | P | 1529.1 | P | --- |
| 262175_at | 1131.7 | P | 1092.5 | P | 1151.1 | P | 1166.1 | P | NDH-O |
| 262176_at | 1502.4 | P | 1643 | P | 1544.6 | P | 1511 | P | FAB1 |
| 262177_at | 206.5 | P | 594.3 | P | 186.2 | P | 369.6 | P | EDS16 |
| 262178_at | 12.5 | A | 52.7 | A | 58.7 | A | 29.8 | A | KOM |
| 262179_at | 19.6 | A | 20.2 | A | 35.4 | A | 32.2 | A | AGL66 |
| 262180_at | 68.2 | P | 69.9 | M | 76.9 | A | 67.4 | M | PGM |
| 262181_at | 528.9 | P | 802.2 | P | 570.6 | P | 849.9 | P | --- |
| 262182_at | 49.2 | P | 54.3 | P | 58.2 | P | 51 | P | --- |
| 262183_at | 297.7 | P | 173.3 | P | 320.6 | P | 176.4 | P | --- |
| 262184_at | 0.6 | A | 4.1 | A | 0.3 | A | 1.8 | A | --- |
| 262185_at | 1.6 | A | 3.1 | A | 14.5 | A | 0.7 | A | AGL67 |
| 262186_at | 4.7 | A | 18.7 | A | 9.1 | A | 26.6 | A | --- |
| 262187_at | 27.4 | A | 45.7 | A | 6.3 | A | 46.4 | A | --- |
| 262188_at | 314.8 | P | 288.9 | P | 251.2 | P | 270.6 | P | --- |
| 262189_at | 305.3 | P | 293.6 | P | 409.5 | P | 295.1 | P | --- |
| 262190_at | 9.4 | A | 23.9 | A | 47.9 | A | 22.5 | A | --- |
| 262191_at | 2.9 | A | 2.8 | A | 3.6 | A | 2.4 | A | --- |
| 262192_at | 18.8 | A | 23.1 | A | 6.6 | A | 21.3 | A | --- |
| 262193_at | 1058 | P | 845.3 | P | 1080.9 | P | 912.2 | P | --- |
| 262194_at | 962.2 | P | 1205.5 | P | 859.1 | P | 1141.8 | P | --- |
| 262195_at | 9337.3 | P | 11604.1 | P | 10403.3 | P | 11033 | P | --- |
| 262196_at | 44.7 | A | 97.9 | P | 92.7 | P | 53.7 | P | MUB5 |
| 262197_at | 1882.6 | P | 1500.7 | P | 1568.4 | P | 1614.6 | P | RAP2.12 |
| 262198_at | 76.9 | P | 21.2 | A | 13.1 | A | 23.3 | A | ATPME2 |
| 262199_at | 240.8 | P | 323.7 | P | 371.3 | P | 273.5 | P | --- |
| 262200_at | 302.4 | P | 358 | P | 347.6 | P | 329.6 | P | --- |
| 262201_at | 125.3 | P | 149.4 | P | 158.7 | P | 125 | P | ORC4 |
| 262202_at | 1549 | P | 2304.4 | P | 1225.3 | P | 1819 | P | APG2 |
| 262203_at | 755.4 | P | 896.4 | P | 866.5 | P | 862 | P | --- |
| 262204_at | 950.6 | P | 801.8 | P | 986.7 | P | 810.5 | P | --- |
| 262205_at | 245.8 | P | 183.1 | P | 249.1 | P | 180.4 | P | --- |
| 262206_at | 718.4 | P | 642.6 | P | 575.7 | P | 634.3 | P | --- |
| 262207_at | 358.6 | P | 537.7 | P | 456.4 | P | 531 | P | PTAC2 |
| 262208_at | 165.3 | P | 254.4 | P | 270.3 | P | 252.2 | P | --- |
| 262209_at | 576.3 | P | 552.2 | P | 537.9 | P | 518.2 | P | --- |
| 262210_at | 528.4 | P | 819.6 | P | 695.4 | P | 831.1 | P | IQD31 |
| 262211_at | 64.6 | P | 40.1 | P | 80.7 | P | 46.3 | P | ORA47 |
| 262212_at | 58.2 | A | 57.3 | P | 61.3 | M | 94.3 | P | ARR15 |
| 262213_at | 43.8 | A | 7.8 | A | 61 | A | 7.6 | A | --- |
| 262214_at | 32.3 | A | 42.8 | A | 67.5 | A | 48.7 | A | --- |
| 262215_at | 517.7 | P | 443.9 | P | 452.3 | P | 569.2 | P | --- |
| 262216_at | 201.3 | P | 182.1 | P | 202 | P | 195.7 | P | --- |
| 262217_at | 1 | A | 1 | A | 1.3 | A | 1.1 | A | --- |
| 262218_at | 15.3 | A | 50.8 | A | 72 | A | 58.8 | A | --- |
| 262219_at | 224.2 | P | 190.7 | P | 187.6 | P | 153.8 | P | --- |
| 262220_at | 113.1 | P | 113 | P | 84 | P | 113.5 | P | CPK30 |
| 262221_at | 293.9 | P | 423.4 | P | 263.7 | P | 315 | P | ARC3 |
| 262222_at | 151 | P | 179.7 | P | 128.3 | P | 174 | P | TRZ1 |
| 262223_at | 443.9 | P | 519.4 | P | 365.6 | P | 489.5 | P | --- |
| 262224_at | 90.2 | P | 75.3 | P | 56.6 | P | 85.4 | P | OTP43 |
| 262225_at | 779.6 | P | 865.7 | P | 664.2 | P | 795.3 | P | ATPME1 |
| 262226_at | 511.8 | P | 89.4 | P | 318.3 | P | 111.8 | P | --- |
| 262227_s_at | 3069.3 | P | 2578.2 | P | 2577 | P | 2445.1 | P | --- |

| | | | | | | | | | |
|---|---|---|---|---|---|---|---|---|---|
| 262228_at | 21.4 | A | 27 | A | 21.3 | A | 3.2 | A | --- |
| 262229_at | 125.1 | P | 33.2 | A | 30.6 | A | 5.7 | A | --- |
| 262230_at | 3455.7 | P | 2242.1 | P | 2677.8 | P | 2632.6 | P | XYL1 |
| 262231_at | 84.4 | A | 77.8 | P | 117.1 | P | 94.7 | P | --- |
| 262232_at | 501.7 | P | 322.3 | P | 263.1 | P | 383.3 | P | --- |
| 262233_at | 72.5 | P | 77.5 | P | 95 | P | 61 | P | CHR18 |
| 262234_at | 113.1 | P | 145.4 | P | 97.6 | P | 132 | P | GCR1 |
| 262235_at | 5596.6 | P | 6796 | P | 5570.4 | P | 6650.9 | P | --- |
| 262236_at | 65.4 | P | 228.3 | P | 97.3 | P | 208.1 | P | --- |
| 262237_at | 624.9 | P | 1081.8 | P | 656 | P | 846.8 | P | --- |
| 262238_at | 1910.8 | P | 2897.2 | P | 2204.3 | P | 2657.8 | P | --- |
| 262239_at | 396.5 | P | 444.7 | P | 330.9 | P | 529.8 | P | --- |
| 262240_at | 32.2 | A | 38.4 | A | 19.1 | A | 47.7 | A | --- |
| 262241_at | 43.2 | P | 40.2 | A | 28 | A | 10.5 | A | --- |
| 262242_at | 203.6 | P | 174.9 | P | 216.6 | P | 151.1 | P | --- |
| 262243_at | 19.7 | A | 6.9 | A | 25.1 | A | 19.4 | A | --- |
| 262244_at | 70.9 | A | 78.5 | M | 153.8 | P | 72.8 | P | CIPK17 |
| 262245_at | 326.8 | P | 378.3 | P | 259.6 | P | 402.9 | P | NPSN12 |
| 262246_at | 2269.9 | P | 2425.2 | P | 2727 | P | 2463.2 | P | AGO1 |
| 262247_at | 1994 | P | 1895.7 | P | 2102.9 | P | 1975.8 | P | D-CDES |
| 262248_at | 316.6 | P | 341.5 | P | 301.9 | P | 329.4 | P | YSL8 |
| 262249_at | 98.7 | P | 121.2 | P | 104.4 | P | 132 | P | RHL1 |
| 262250_at | 134.9 | P | 166.2 | P | 195.7 | P | 141.1 | P | --- |
| 262251_at | 610.7 | P | 450 | P | 535.2 | P | 447.2 | P | --- |
| 262252_at | 21.5 | A | 4.8 | A | 29.1 | A | 1.2 | A | --- |
| 262253_s_at | 877.7 | P | 752.3 | P | 851.6 | P | 770.8 | P | --- |
| 262254_at | 54 | P | 156 | P | 71.9 | P | 130.6 | P | GLIP5 |
| 262255_at | 134 | P | 79.3 | P | 82.7 | P | 97.3 | M | --- |
| 262256_at | 1.7 | A | 1.7 | A | 12.4 | A | 3 | A | --- |
| 262257_at | 36.7 | A | 39 | A | 26.1 | A | 24.7 | A | --- |
| 262258_at | 1517.9 | P | 1502.9 | P | 1531.1 | P | 1649.1 | P | PAE1 |
| 262259_s_at | 179.7 | P | 368.6 | P | 193.3 | P | 312.8 | P | --- |
| 262260_at | 32.1 | A | 191.2 | P | 94.7 | P | 242.3 | P | MLP34 |
| 262261_at | 96.5 | P | 87.4 | P | 104.2 | P | 95.6 | P | CLE17 |
| 262262_at | 8102.4 | P | 4315.7 | P | 10074.7 | P | 5633.4 | P | --- |
| 262263_at | 1079.8 | P | 1105.6 | P | 977.1 | P | 1028.1 | P | PIN3 |
| 262264_at | 529.3 | P | 467.6 | P | 634.5 | P | 427.9 | P | --- |
| 262265_at | 28.3 | A | 41.4 | A | 41.7 | A | 33 | A | --- |
| 262266_at | 26.1 | A | 22.8 | A | 43.7 | A | 28.4 | A | --- |
| 262267_at | 0.4 | A | 3.6 | A | 10.9 | A | 3.6 | A | --- |
| 262268_at | 7.8 | A | 2.1 | A | 29.6 | A | 16.1 | A | --- |
| 262269_at | 3 | A | 3.3 | A | 5.6 | A | 3.1 | A | --- |
| 262270_x_at | 1.4 | A | 3.8 | A | 3.3 | A | 2.2 | A | --- |
| 262271_x_at | 52.6 | A | 34.5 | A | 49.9 | A | 31.7 | A | --- |
| 262272_at | 827.8 | P | 605.5 | P | 797.8 | P | 679.6 | P | --- |
| 262273_at | 78.7 | A | 73.8 | A | 74.7 | A | 79.9 | A | AT1G68730 |
| 262274_at | 521.4 | P | 493.5 | P | 479.5 | P | 482.4 | P | TADA |
| 262275_at | 193.6 | P | 179.5 | P | 161.6 | P | 233.4 | P | --- |
| 262276_at | 19 | A | 4.1 | A | 24.7 | A | 7 | A | --- |
| 262277_at | 402.8 | P | 225.2 | P | 405.7 | P | 322 | P | --- |
| 262278_at | 3 | A | 15.4 | A | 3.9 | A | 2.1 | A | PAN |
| 262279_at | 28.9 | A | 28.2 | A | 31.5 | A | 35.5 | A | --- |
| 262280_at | 756.2 | P | 849.1 | P | 770.6 | P | 842.6 | P | --- |
| 262281_at | 3306.2 | P | 1905.1 | P | 2686.5 | P | 2768.8 | P | --- |
| 262282_at | 19.2 | A | 32.2 | P | 18.1 | A | 9.9 | A | --- |
| 262283_at | 5182.7 | P | 5447.4 | P | 4929.1 | P | 5085.1 | P | --- |
| 262284_at | 255 | P | 230.1 | P | 348.6 | P | 237.2 | P | --- |
| 262285_at | 667 | P | 567.5 | P | 571.2 | P | 597.6 | P | --- |
| 262286_at | 692.5 | P | 793.6 | P | 651.1 | P | 713.9 | P | --- |
| 262287_at | 2234.1 | P | 2424.2 | P | 2087.2 | P | 2585.2 | P | --- |
| 262288_at | 2011.2 | P | 2081.6 | P | 2517.1 | P | 2191.1 | P | CRR23 |
| 262289_at | 1537 | P | 1545.2 | P | 1569.2 | P | 1676.3 | P | --- |
| 262290_at | 4 | A | 68.5 | P | 14 | A | 62.5 | P | --- |
| 262291_at | 338.4 | P | 212.1 | P | 234.3 | P | 263.8 | P | --- |
| 262292_at | 222.4 | P | 208.8 | P | 270.3 | P | 251.7 | P | --- |
| 262293_at | 17.5 | A | 10 | A | 5.2 | A | 14.6 | A | --- |
| 262294_at | 4 | A | 20.3 | A | 26.2 | A | 17.7 | A | --- |
| 262295_at | 650.2 | P | 583.9 | P | 646 | P | 580.5 | P | ATU2AF35A |
| 262296_at | 686.3 | P | 605.3 | P | 667.4 | P | 621.7 | P | CYCT1;3 |
| 262297_at | 424.1 | P | 451 | P | 405.7 | P | 464.7 | P | --- |
| 262298_at | 2972.5 | P | 2830.7 | P | 2731.8 | P | 2939.2 | P | --- |
| 262299_at | 444.2 | P | 459.9 | P | 618.3 | P | 438.7 | P | --- |
| 262300_at | 10.1 | A | 10.9 | A | 0.9 | A | 2.5 | A | --- |
| 262301_at | 3.4 | A | 21.1 | A | 6.3 | A | 8.1 | A | --- |
| 262302_at | 22.4 | A | 7.7 | A | 18.5 | A | 16.1 | A | --- |
| 262303_at | 15.6 | A | 31.7 | A | 4.5 | A | 31.2 | A | ATHB18 |
| 262304_at | 3929.7 | P | 4754 | P | 3892 | P | 5457.9 | P | MLP43 |
| 262305_at | 19.3 | A | 4.2 | A | 11.2 | A | 13.8 | A | --- |
| 262306_s_at | 1258.3 | P | 830 | P | 1201.2 | P | 939.2 | P | SYNC3 |
| 262307_at | 19.2 | A | 24.4 | P | 1.9 | A | 39.5 | A | --- |
| 262308_at | 564.3 | P | 646.5 | P | 627.3 | P | 613.1 | P | --- |
| 262309_at | 1756.4 | P | 1358.8 | P | 1762.5 | P | 1451.9 | P | --- |
| 262310_at | 21.3 | A | 9.9 | A | 20.5 | A | 31.9 | A | MLP31 |
| 262311_at | 21.9 | A | 2.5 | A | 7 | A | 2.9 | A | ATHB18 |
| 262312_at | 1128.6 | P | 2070 | P | 2083.7 | P | 2059.4 | P | MLP28 |
| 262313_at | 561.3 | P | 633.9 | P | 504.8 | P | 562.9 | P | --- |
| 262314_at | 1753.9 | P | 1396 | P | 1243.7 | P | 1379.3 | P | --- |
| 262315_at | 36.6 | P | 35.8 | A | 43.1 | P | 25.3 | A | --- |
| 262316_at | 538.2 | P | 549.7 | P | 648.4 | P | 535.8 | P | PAC |
| 262317_at | 38.3 | M | 70.9 | P | 68.6 | P | 91.6 | P | EDA4 |
| 262318_at | 8.1 | A | 9.1 | A | 8.8 | A | 15.9 | A | --- |
| 262319_s_at | 699.2 | P | 664.2 | P | 537.3 | P | 579.2 | P | --- |
| 262320_at | 2.6 | A | 19.5 | A | 21.5 | A | 24.1 | A | --- |
| 262321_at | 17.4 | A | 11.2 | A | 37.8 | A | 38.7 | A | --- |
| 262322_at | 61.8 | A | 106.5 | A | 98.7 | A | 69.3 | P | --- |
| 262323_at | 1699 | P | 1461.7 | P | 1874.6 | P | 1685.1 | P | --- |
| 262324_at | 46.1 | P | 156.6 | P | 53 | A | 150.8 | P | ATCHX16 |
| 262325_at | 7.7 | A | 11.9 | A | 17.4 | A | 5 | A | --- |
| 262326_at | 1636.3 | P | 1455.5 | P | 1632.6 | P | 1562.9 | P | --- |
| 262327_at | 2.8 | A | 3.9 | A | 7.2 | A | 3.1 | A | --- |
| 262328_at | 10.9 | A | 25.1 | A | 10.8 | A | 19.2 | A | --- |
| 262329_at | 2120.5 | P | 2535.1 | P | 2245.5 | P | 2703.9 | P | --- |
| 262330_at | 29.8 | A | 33.3 | A | 48 | A | 37.6 | A | RLM1 |
| 262331_at | 216.7 | P | 218.3 | P | 256.8 | P | 214.4 | P | --- |
| 262332_at | 9.6 | A | 16.8 | A | 36.6 | A | 1.4 | A | --- |
| 262333_at | 1.6 | A | 2.4 | A | 1.8 | A | 2.4 | A | --- |
| 262334_at | 15.2 | A | 14.6 | A | 21.4 | A | 21.1 | A | --- |
| 262335_at | 28.4 | A | 13.4 | A | 8.2 | A | 29.2 | A | SPO11-2 |
| 262336_at | 114.1 | P | 51.1 | P | 181.9 | P | 68 | P | TOM7-2 |

| | | | | | | | | | |
|---|---|---|---|---|---|---|---|---|---|
| 262337_at | 2.1 | A | 27.3 | A | 11 | A | 12.3 | A | --- |
| 262338_at | 351.2 | P | 307.8 | P | 494.9 | P | 340.5 | P | --- |
| 262339_at | 18.6 | A | 14.2 | A | 41.6 | A | 11.4 | A | WRKY56 |
| 262340_at | 425.2 | P | 439 | P | 338 | P | 446.5 | P | ANAC027 |
| 262341_at | 9552.1 | P | 8217.1 | P | 7712.7 | P | 7299.7 | P | UBC28 |
| 262342_at | 553.8 | P | 604.9 | P | 604.2 | P | 650.7 | P | --- |
| 262343_at | 1641.5 | P | 1475.7 | P | 1647.7 | P | 1496.2 | P | TOPP3 |
| 262344_at | 673.3 | P | 374.6 | P | 743.7 | P | 456.2 | P | ATRBOH F |
| 262345_at | 104.8 | P | 111.3 | P | 101.7 | P | 107.7 | P | --- |
| 262346_at | 2345.9 | P | 1517.8 | P | 2108 | P | 1577.1 | P | --- |
| 262347_at | 44.5 | M | 17.6 | A | 63.6 | A | 3.2 | A | --- |
| 262348_at | 407.2 | P | 353.9 | P | 475.6 | P | 442.4 | P | --- |
| 262349_at | 15.9 | A | 47 | P | 56.1 | A | 91.4 | P | --- |
| 262350_at | 3.8 | A | 3.3 | A | 4.4 | A | 3.8 | A | ATGPX4 |
| 262351_at | 137.7 | P | 151.5 | P | 138.8 | P | 155.4 | P | BGAL17 |
| 262352_at | 27.4 | A | 30.2 | A | 24.7 | A | 45.8 | A | --- |
| 262353_at | 35.3 | A | 4.7 | A | 18.9 | A | 4.6 | A | --- |
| 262354_at | 961.8 | P | 403.6 | P | 809.1 | P | 545.2 | P | VHA-E3 |
| 262355_at | 384.5 | P | 341.9 | P | 286.3 | P | 305.8 | P | --- |
| 262356_at | 29.7 | A | 8.6 | A | 53.1 | A | 23.6 | A | --- |
| 262357_at | 498.4 | P | 375.2 | P | 368.6 | P | 363.8 | P | --- |
| 262358_at | 16.4 | A | 2.4 | A | 44 | A | 2.2 | A | --- |
| 262359_at | 5.1 | A | 3.4 | A | 5.7 | A | 13.4 | A | --- |
| 262360_at | 441.5 | P | 352.3 | P | 458.4 | P | 327.2 | P | --- |
| 262361_at | 84.4 | P | 72.1 | P | 87.6 | P | 67.3 | P | GTE3 |
| 262362_at | 66.3 | A | 70.9 | A | 23.8 | A | 33.3 | A | --- |
| 262363_at | 37.2 | M | 27.6 | A | 40.4 | A | 37.5 | P | --- |
| 262364_at | 50.2 | A | 38.4 | A | 61.4 | P | 20.9 | A | --- |
| 262365_at | 31.9 | A | 52.7 | P | 46.8 | A | 56.1 | P | --- |
| 262366_at | 190.8 | P | 222.3 | P | 195.1 | P | 202.7 | P | --- |
| 262367_at | 1237.3 | P | 1367.3 | P | 1344.6 | P | 1189.1 | P | VPS46.2 |
| 262368_at | 1699.8 | P | 2080.9 | P | 1739.2 | P | 2045.7 | P | --- |
| 262369_at | 150 | P | 102.9 | P | 146.6 | P | 105.2 | P | --- |
| 262370_at | 254.8 | P | 349.8 | P | 420.1 | P | 365.8 | P | --- |
| 262371_at | 6 | A | 6.5 | A | 3.5 | A | 10.4 | A | --- |
| 262372_at | 76.7 | P | 124.1 | P | 127.7 | P | 68.3 | P | --- |
| 262373_at | 1983.1 | P | 1406.5 | P | 3497.2 | P | 1707.5 | P | --- |
| 262374_s_at | 2296.9 | P | 3280.6 | P | 2310.9 | P | 3406.1 | P | TIR |
| 262375_at | 245.5 | P | 373.8 | P | 314.8 | P | 376.8 | P | SUVH3 |
| 262376_at | 688.7 | P | 1162.9 | P | 934 | P | 1226.5 | P | HTH |
| 262377_at | 781.1 | P | 802.5 | P | 811.1 | P | 803.3 | P | --- |
| 262378_at | 272.3 | P | 498.5 | P | 281.7 | P | 331.5 | P | NF-YA3 |
| 262379_at | 119.5 | P | 188.3 | P | 209.4 | P | 219.4 | P | --- |
| 262380_at | 904.4 | P | 876 | P | 1454 | P | 1001.3 | P | --- |
| 262381_at | 2625 | P | 1444.7 | P | 2225.3 | P | 1515.6 | P | --- |
| 262382_at | 480.2 | P | 266.5 | P | 358.4 | P | 294.9 | P | --- |
| 262383_at | 1012.8 | P | 534.3 | P | 728.3 | P | 528.9 | P | --- |
| 262384_at | 37.6 | A | 29.5 | A | 49.6 | A | 32.4 | A | --- |
| 262385_at | 2.8 | A | 13.3 | A | 3.7 | A | 11.3 | A | --- |
| 262386_at | 77.8 | P | 60.5 | P | 75.5 | P | 63.4 | P | --- |
| 262387_s_at | 237.1 | P | 259.4 | P | 324.1 | P | 287.1 | P | --- |
| 262388_at | 33.5 | A | 108.4 | P | 83.7 | A | 85.5 | P | --- |
| 262389_at | 4.6 | A | 5.7 | A | 7.6 | A | 14.7 | A | --- |
| 262390_at | 55.1 | P | 69.5 | P | 87.1 | P | 56.8 | P | --- |
| 262391_at | 57.1 | A | 31.9 | A | 60.9 | A | 10.8 | A | GGPS6 |
| 262392_at | 58.7 | P | 39.7 | P | 53.5 | A | 49.8 | A | --- |
| 262393_at | 10.4 | A | 5.1 | A | 28 | A | 15.5 | A | --- |
| 262394_at | 462.9 | P | 661.2 | P | 577.6 | P | 776.9 | P | emb1273 |
| 262395_at | 187.5 | P | 141 | A | 191.2 | A | 154.5 | A | --- |
| 262396_at | 330.6 | P | 458.8 | P | 314.1 | P | 474.7 | P | --- |
| 262397_at | 603.4 | P | 812.1 | P | 771.4 | P | 738.8 | P | --- |
| 262398_at | 244.8 | P | 282 | P | 254.2 | P | 281.6 | P | --- |
| 262399_at | 21.9 | P | 26.8 | A | 29.7 | A | 38.6 | P | --- |
| 262400_at | 826.3 | P | 873 | P | 940.7 | P | 847.5 | P | RTV1 |
| 262401_at | 1068.1 | P | 1230.8 | P | 995.8 | P | 1032.7 | P | RABG3E |
| 262402_at | 1130.5 | P | 996.4 | P | 1247.6 | P | 1070.1 | P | TOM6 |
| 262403_s_at | 9.9 | A | 6.1 | A | 2.5 | A | 0.9 | A | --- |
| 262404_at | 9.7 | A | 31.5 | A | 6.6 | A | 1.9 | A | --- |
| 262405_at | 10.3 | A | 24.2 | A | 33.8 | A | 32.5 | A | HDG10 |
| 262406_at | 28.6 | A | 20.2 | P | 34.1 | A | 46.2 | A | AtMYB93 |
| 262407_at | 178 | P | 242.8 | P | 180.7 | P | 252 | P | --- |
| 262408_at | 215.2 | P | 332.7 | P | 179.2 | P | 287.2 | P | --- |
| 262409_at | 2.9 | A | 13.5 | A | 4.7 | A | 2.8 | A | --- |
| 262410_at | 106 | P | 124.9 | P | 174.6 | P | 109.1 | P | --- |
| 262411_at | 269.6 | P | 266.1 | P | 243.7 | P | 329.1 | P | --- |
| 262412_at | 159.2 | P | 158.3 | P | 164.8 | P | 144.1 | P | GRF11 |
| 262413_at | 247.7 | P | 360 | P | 298 | P | 398.3 | P | ATAPRL4 |
| 262414_at | 374 | P | 415.6 | P | 411 | P | 468.6 | P | LACS2 |
| 262415_at | 826.7 | P | 807 | P | 915.3 | P | 773.4 | P | emb1129 |
| 262416_at | 32.9 | A | 20 | A | 41.4 | A | 11 | A | --- |
| 262417_at | 1142 | P | 1210.4 | P | 1315.7 | P | 1159 | P | ATSIRB |
| 262418_at | 3991.1 | P | 4249.1 | P | 3934.6 | P | 3700.7 | P | ATHX |
| 262419_at | 695.4 | P | 615.4 | P | 978.1 | P | 726.9 | P | --- |
| 262420_at | 400.7 | P | 384.9 | P | 358.2 | P | 374.3 | P | TAF15 |
| 262421_at | 100.7 | P | 117.7 | P | 101 | P | 99 | P | --- |
| 262422_at | 689.6 | P | 633.7 | P | 518.6 | P | 498.4 | P | --- |
| 262423_at | 22.4 | A | 13.1 | A | 40.5 | A | 28.1 | A | --- |
| 262424_at | 1.5 | A | 0.6 | A | 2.6 | A | 2.2 | A | --- |
| 262425_at | 1.3 | A | 6.4 | A | 4.6 | A | 5.2 | A | --- |
| 262426_s_at | 987 | P | 1059.6 | P | 997.5 | P | 1025.4 | P | --- |
| 262427_s_at | 33.7 | A | 4.3 | A | 72.8 | A | 34.1 | A | BGLU34 |
| 262428_at | 423.7 | P | 470.2 | P | 323.2 | P | 411.7 | P | --- |
| 262429_at | 5.8 | A | 19 | A | 4.9 | A | 28.8 | A | --- |
| 262430_s_at | 718.1 | P | 707.1 | P | 429.1 | P | 665.3 | P | --- |
| 262431_at | 20.4 | A | 0.8 | A | 25.4 | A | 25.7 | A | --- |
| 262432_at | 1293.5 | P | 1438.1 | P | 1442.7 | P | 1512.3 | P | --- |
| 262433_s_at | 2086.8 | P | 1764.6 | P | 2500.8 | P | 1775.7 | P | ATRBP47C' |
| 262434_at | 94.7 | P | 211.6 | P | 219.6 | P | 274.2 | P | --- |
| 262435_at | 3.9 | A | 15.4 | A | 15 | A | 16.9 | A | CYP96A8 |
| 262436_at | 27.7 | A | 43.4 | A | 41.2 | A | 45.3 | P | --- |
| 262437_at | 36.5 | A | 33.1 | A | 53.1 | A | 38.1 | A | --- |
| 262438_at | 112.1 | P | 161.8 | P | 151.6 | P | 162 | P | --- |
| 262439_at | 1 | A | 0.7 | A | 12.2 | A | 1.3 | A | --- |
| 262440_at | 1508 | P | 1712.2 | P | 1633.6 | P | 1627.4 | P | --- |
| 262441_at | 142.6 | P | 206.4 | P | 131.9 | P | 144.7 | P | OSB1 |
| 262442_at | 5263.8 | P | 5117.6 | P | 6586 | P | 5104.2 | P | --- |
| 262443_at | 51.1 | A | 54.1 | M | 48.5 | P | 64.9 | P | --- |
| 262444_at | 5.7 | A | 23.2 | A | 9.1 | A | 19.7 | A | --- |
| 262445_at | 16.2 | A | 14.7 | A | 16.4 | A | 20.3 | A | --- |

| | | | | | | | | | |
|---|---|---|---|---|---|---|---|---|---|
| 262446_at | 31.6 | A | 40.4 | P | 5.5 | A | 28.6 | A | --- |
| 262447_at | 1 | A | 1.5 | A | 2.3 | A | 11.4 | A | --- |
| 262448_at | 2.3 | A | 14.9 | A | 46.2 | A | 43 | A | --- |
| 262449_at | 25.3 | P | 39.2 | P | 30.3 | P | 37 | P | --- |
| 262450_at | 142.6 | P | 142.6 | P | 103.9 | P | 147.5 | P | --- |
| 262451_at | 251.5 | P | 373.2 | P | 354 | P | 488.4 | P | SUB |
| 262452_at | 309.3 | P | 405.8 | P | 198.4 | P | 542.3 | P | --- |
| 262453_at | 637.7 | P | 565.3 | P | 875.7 | P | 722.9 | P | --- |
| 262454_at | 60 | P | 84.8 | P | 82.1 | P | 75.7 | P | BFN1 |
| 262455_at | 1323.2 | P | 1477.9 | P | 1062.8 | P | 1255.7 | P | MLO2 |
| 262456_at | 434.4 | P | 664.6 | P | 326.9 | P | 499.6 | P | STP1 |
| 262457_at | 539.6 | P | 618.5 | P | 506.9 | P | 583.1 | P | --- |
| 262458_at | 445.8 | P | 327.3 | P | 500.5 | P | 365.9 | P | --- |
| 262459_at | 98.9 | P | 76.3 | M | 82.2 | M | 97.4 | M | --- |
| 262460_s_at | 5.7 | A | 12.9 | A | 19 | A | 26.8 | A | --- |
| 262461_at | 149.7 | P | 158.9 | P | 122.6 | P | 196.6 | P | VIIIA |
| 262462_at | 43.8 | P | 48.9 | P | 53 | P | 21.6 | P | --- |
| 262463_at | 25.3 | P | 52.4 | P | 79.4 | P | 63.4 | P | STP9 |
| 262464_at | 81.4 | P | 57.7 | P | 76.3 | P | 50.7 | P | --- |
| 262465_at | 78.1 | P | 78.5 | P | 97.9 | P | 85.7 | P | --- |
| 262466_at | 99.7 | P | 155.4 | P | 149.5 | M | 90.9 | M | NTMC2T5.1 |
| 262467_at | 143.5 | P | 219.8 | P | 134.8 | P | 225.4 | P | FU |
| 262468_at | 3178.8 | P | 2862.5 | P | 3281.3 | P | 3091.1 | P | ALATS |
| 262469_s_at | 17.8 | A | 2.9 | A | 11.1 | A | 1.4 | A | --- |
| 262470_at | 39.2 | P | 44.2 | P | 28.7 | P | 14.2 | A | --- |
| 262471_at | 103.9 | P | 121.2 | P | 83.2 | P | 90.2 | P | --- |
| 262472_at | 8.5 | A | 8.3 | A | 4.8 | A | 15.2 | A | --- |
| 262473_at | 3865.8 | P | 3623.1 | P | 3152.3 | P | 3904.1 | P | FTSH1 |
| 262474_at | 59.2 | A | 83.9 | P | 36.1 | P | 72.9 | P | FU |
| 262475_at | 1.1 | A | 19.8 | A | 5.1 | A | 3.5 | A | --- |
| 262476_at | 1569.7 | P | 1539.7 | P | 1524 | P | 1631.1 | P | --- |
| 262477_at | 38.2 | A | 85.9 | P | 37.1 | A | 74.7 | P | --- |
| 262478_at | 90 | P | 144.1 | P | 78.6 | P | 149.5 | P | --- |
| 262479_at | 89.3 | P | 240.1 | P | 194.2 | P | 208.6 | P | SUB |
| 262480_at | 58 | P | 62.2 | P | 58 | P | 44.8 | P | --- |
| 262481_at | 1625.3 | P | 1889.2 | P | 1304 | P | 1485.3 | P | --- |
| 262482_at | 1977.1 | P | 883.6 | P | 1103.3 | P | 739.5 | P | SRG1 |
| 262483_at | 3030.8 | P | 3437 | P | 3205.2 | P | 3617.3 | P | FUG1 |
| 262484_at | 291.1 | P | 375.2 | P | 303.7 | P | 299.7 | P | ATSWI3C |
| 262485_at | 56.1 | A | 95.8 | P | 95.8 | P | 92 | M | --- |
| 262486_at | 74.9 | M | 60 | P | 112.1 | A | 57.8 | P | --- |
| 262487_at | 175 | P | 171.1 | P | 178.7 | P | 179.8 | P | --- |
| 262488_at | 589.3 | P | 503.1 | P | 382.3 | P | 365 | P | --- |
| 262489_at | 260.3 | P | 279.2 | P | 345.7 | P | 188 | P | --- |
| 262490_at | 122.1 | P | 136.9 | P | 148.8 | P | 144.6 | P | UREF |
| 262491_at | 244.1 | P | 260.3 | P | 219.3 | P | 236.3 | P | --- |
| 262492_at | 561.8 | P | 694.4 | P | 561.2 | P | 714.5 | P | --- |
| 262493_at | 500.1 | P | 429.5 | P | 468.5 | P | 439.2 | P | --- |
| 262494_at | 50.5 | A | 78.2 | P | 64.6 | A | 85.2 | P | --- |
| 262495_at | 269.1 | P | 338.7 | P | 351.8 | P | 292.7 | P | --- |
| 262496_at | 237.9 | P | 247.6 | P | 204.7 | P | 210.1 | P | --- |
| 262497_at | 2318.8 | P | 2573.7 | P | 2667 | P | 3085.8 | P | PBC1 |
| 262498_at | 379.5 | P | 243.9 | P | 344.8 | P | 335 | P | OGG1 |
| 262499_at | 3146.9 | P | 2329.2 | P | 2844.3 | P | 2280.1 | P | --- |
| 262500_at | 1033.6 | P | 955.1 | P | 950.4 | P | 932.9 | P | FBP7 |
| 262501_at | 333.5 | P | 476.8 | P | 485.9 | P | 515.3 | P | emb1968 |
| 262502_at | 1052.5 | P | 1452.1 | P | 1494.8 | P | 1523.3 | P | PTAC6 |
| 262503_at | 1543.8 | P | 1584.9 | P | 974.7 | P | 1235.2 | P | --- |
| 262504_at | 3804.6 | P | 4436.1 | P | 3629.3 | P | 4810.9 | P | ATPDIL1-1 |
| 262505_at | 5041.7 | P | 4111.6 | P | 3498.4 | P | 4473.4 | P | --- |
| 262506_at | 630.6 | P | 758.7 | P | 602.4 | P | 753.7 | P | NADK2 |
| 262507_at | 401.3 | P | 378.8 | P | 269.9 | P | 360.7 | P | --- |
| 262508_at | 133.9 | P | 143.3 | P | 108.2 | P | 128.5 | P | --- |
| 262509_at | 152.2 | P | 135.1 | P | 90.3 | A | 113.1 | M | CRR22 |
| 262510_at | 194 | P | 159.6 | P | 209.7 | P | 148.6 | P | --- |
| 262511_at | 38.6 | A | 25.1 | A | 59 | A | 20.4 | A | SYP125 |
| 262512_at | 304.3 | P | 314.9 | P | 296.7 | P | 312.1 | P | --- |
| 262513_at | 190.3 | P | 121.2 | P | 136.4 | P | 165.8 | P | --- |
| 262514_at | 875.1 | P | 963 | P | 787.8 | P | 845.6 | P | anac017 |
| 262515_at | 317.2 | P | 368.7 | P | 366.4 | P | 327.1 | P | --- |
| 262516_at | 93.8 | P | 123.8 | P | 101.6 | P | 130.9 | P | ATGSTU26 |
| 262517_at | 215.1 | P | 73.2 | P | 110.9 | P | 53.9 | P | ATGSTU25 |
| 262518_at | 3839.7 | P | 2046.4 | P | 2544.5 | P | 2858.9 | P | ATGSTU24 |
| 262519_at | 682.5 | P | 587.8 | P | 539.6 | P | 575 | P | --- |
| 262520_at | 29.2 | A | 27.8 | A | 27.6 | A | 30.4 | A | --- |
| 262521_at | 315.2 | P | 346.4 | P | 370.7 | P | 360.4 | P | --- |
| 262522_at | 331.2 | P | 417.7 | P | 397.8 | P | 418.3 | P | CAT8 |
| 262523_at | 292.8 | P | 380.1 | P | 325.9 | P | 374.9 | P | UBP15 |
| 262524_at | 231.9 | P | 274 | P | 250.8 | P | 218.8 | P | --- |
| 262525_at | 36 | A | 49.5 | A | 38.7 | A | 40.3 | A | CYP72C1 |
| 262526_at | 591.8 | P | 705.6 | P | 592.5 | P | 721.5 | P | SPS2 |
| 262527_at | 1.3 | A | 1.1 | A | 1 | A | 1.6 | A | --- |
| 262528_at | 24.2 | P | 45.4 | P | 72 | P | 29.4 | P | AHA10 |
| 262529_at | 106.9 | P | 90.2 | P | 152.4 | P | 96 | A | AtRLP3 |
| 262530_at | 50.8 | A | 42 | P | 31.9 | A | 59.1 | P | AtRLP2 |
| 262531_at | 75.9 | P | 130 | P | 113.9 | P | 133.4 | P | --- |
| 262532_at | 568.2 | P | 591.2 | P | 582.9 | P | 657.1 | P | --- |
| 262533_at | 194.2 | P | 226.1 | P | 194.1 | P | 200.1 | P | --- |
| 262534_at | 109.2 | A | 125.4 | P | 107.4 | A | 115.7 | P | SHA |
| 262535_at | 0.7 | A | 4.5 | A | 18.2 | A | 8.6 | A | ATTPS3 |
| 262536_at | 6330 | P | 5564.8 | P | 6741.7 | P | 6573.5 | P | --- |
| 262537_s_at | 1155.3 | P | 1279.5 | P | 1114.2 | P | 1117.2 | P | UBC34 |
| 262538_at | 237.1 | P | 279.9 | P | 200.4 | P | 286.8 | P | --- |
| 262539_at | 942.9 | P | 1371.5 | P | 1282.8 | P | 1173.9 | P | --- |
| 262540_at | 257.9 | P | 319.2 | P | 273.8 | P | 240.5 | P | --- |
| 262541_at | 0.3 | A | 0.6 | A | 2 | A | 0.5 | A | --- |
| 262542_at | 147.9 | P | 142.8 | P | 123.9 | P | 100.5 | P | anac016 |
| 262543_at | 198 | P | 226.8 | P | 279.2 | P | 239.1 | P | --- |
| 262544_at | 739.9 | P | 572.1 | P | 734.5 | P | 529.4 | P | --- |
| 262545_at | 1.5 | A | 3.6 | A | 8.7 | A | 3.5 | A | --- |
| 262546_at | 1.4 | A | 2.1 | A | 5 | A | 1.8 | A | ZIP10 |
| 262547_at | 6.2 | A | 2.6 | A | 13.5 | A | 6.1 | A | --- |
| 262548_at | 175.8 | P | 215.9 | P | 235.7 | P | 213.8 | P | AGO2 |
| 262549_at | 98.1 | P | 55.5 | P | 50.3 | P | 60.1 | M | --- |
| 262550_at | 42.2 | P | 45.3 | P | 46.5 | P | 47.5 | P | --- |
| 262551_at | 1037.6 | P | 912.8 | P | 829.8 | P | 995.8 | P | RUB1 |
| 262552_at | 63.8 | P | 60.1 | M | 125.4 | P | 82.1 | P | --- |
| 262553_at | 90.1 | P | 61.4 | P | 75.8 | P | 35.9 | P | RECQL2 |
| 262554_at | 1.8 | A | 3.1 | A | 27.3 | A | 18.3 | A | --- |

| | | | | | | | | | |
|---|---|---|---|---|---|---|---|---|---|
| 262555_at | 12.1 | A | 1.9 | A | 20 | A | 2.9 | A | --- |
| 262556_at | 15.3 | A | 11.9 | A | 29.6 | A | 4.5 | A | --- |
| 262557_at | 15144.7 | P | 16241.8 | P | 14140.6 | P | 15806.4 | P | --- |
| 262558_at | 178.9 | P | 339.4 | P | 247 | P | 409.2 | P | --- |
| 262559_at | 607.6 | P | 604.9 | P | 508.1 | P | 585.2 | P | --- |
| 262560_at | 15.1 | A | 4.4 | A | 43.5 | A | 14.4 | A | --- |
| 262561_at | 299.1 | P | 307 | P | 315.2 | P | 249.2 | P | --- |
| 262562_at | 460.5 | P | 553.9 | P | 400.3 | P | 509.5 | P | --- |
| 262563_at | 604.3 | P | 595.6 | P | 618.5 | P | 666.9 | P | SERK2 |
| 262564_at | 1.6 | A | 2.7 | A | 2.1 | A | 3 | A | --- |
| 262565_at | 381.3 | P | 424.2 | P | 341.1 | P | 395.6 | P | --- |
| 262566_at | 553.5 | P | 850.8 | P | 1390.3 | P | 1049.5 | P | ARF12 |
| 262567_at | 301.4 | P | 333.4 | P | 325.9 | P | 291 | P | --- |
| 262568_at | 8.5 | A | 1.4 | A | 11.3 | A | 11.4 | A | AtRLP5 |
| 262569_at | 482.4 | P | 512.3 | P | 594.3 | P | 503.7 | P | --- |
| 262570_at | 646.1 | P | 369.1 | P | 667.6 | P | 513.9 | P | --- |
| 262571_at | 466.1 | P | 373.7 | P | 401.6 | P | 384.4 | P | --- |
| 262572_at | 1839.3 | P | 2242.5 | P | 1864.4 | P | 2361.7 | P | --- |
| 262573_at | 1758.2 | P | 1262.8 | P | 2065.6 | P | 1209.5 | P | PDF1A |
| 262574_at | 637.6 | P | 344.7 | P | 552.3 | P | 371 | P | --- |
| 262575_at | 40.8 | A | 38.5 | A | 28.8 | A | 91.1 | P | PDR7 |
| 262576_at | 185.9 | P | 261.7 | P | 201 | P | 225.4 | P | --- |
| 262577_at | 948.7 | P | 848.5 | P | 935.5 | P | 929.9 | P | --- |
| 262578_at | 69.2 | P | 74.3 | P | 67.4 | A | 84.8 | A | --- |
| 262579_at | 5.3 | A | 18.2 | A | 2.3 | A | 2.3 | A | --- |
| 262580_at | 50.7 | P | 52.2 | P | 58.6 | A | 34.1 | A | --- |
| 262581_at | 1158 | P | 1287.1 | P | 1186.6 | P | 1134.1 | P | --- |
| 262582_at | 583.8 | P | 721.7 | P | 673.9 | P | 720.1 | P | --- |
| 262583_at | 285.9 | P | 206.7 | P | 384.9 | P | 226.5 | P | --- |
| 262584_at | 594.8 | P | 436.9 | P | 662.5 | P | 433 | P | --- |
| 262585_at | 3.6 | A | 17.6 | A | 51.2 | A | 16.5 | A | BOR4 |
| 262586_at | 91.3 | A | 87 | A | 125.7 | A | 85.7 | M | --- |
| 262587_at | 97.7 | P | 115.7 | P | 109.5 | P | 88.6 | P | --- |
| 262588_at | 591.8 | P | 624.6 | P | 592.5 | P | 622.6 | P | --- |
| 262589_s_at | 60.2 | P | 33.7 | P | 101.1 | P | 58.8 | P | --- |
| 262590_at | 599.2 | P | 559.4 | P | 639 | P | 574.4 | P | RHA2A |
| 262591_at | 117.5 | P | 124.7 | P | 126.9 | P | 132.9 | P | CCMH |
| 262592_at | 1128.6 | P | 995.2 | P | 920.3 | P | 837.3 | P | --- |
| 262593_at | 3188.1 | P | 3104.8 | P | 3151.7 | P | 3526.8 | P | --- |
| 262594_at | 787.4 | P | 409.1 | P | 787.4 | P | 472.2 | P | --- |
| 262595_at | 3.7 | A | 4.2 | A | 8.5 | A | 11.1 | A | SHN1 |
| 262596_at | 93.1 | P | 93.7 | P | 90 | A | 92.5 | P | LPP2 |
| 262597_at | 408.7 | P | 362.7 | P | 404.4 | P | 428.6 | P | --- |
| 262598_at | 533.1 | P | 642.3 | P | 742.3 | P | 741.7 | P | --- |
| 262599_at | 191.5 | P | 246.3 | P | 223.1 | P | 233.4 | P | --- |
| 262600_at | 606.1 | P | 670.9 | P | 765.5 | P | 681.1 | P | MBD10 |
| 262601_at | 197.6 | P | 129.2 | P | 173.1 | P | 136.7 | P | ATHSRP54A |
| 262602_at | 2980.5 | P | 2683.5 | P | 2701.3 | P | 2772.1 | P | --- |
| 262603_at | 695.6 | P | 60.9 | A | 476 | P | 73.2 | A | --- |
| 262604_at | 237.9 | P | 307.4 | P | 317.7 | P | 356.6 | P | --- |
| 262605_at | 370.8 | P | 303.9 | P | 285.1 | P | 263.1 | P | --- |
| 262606_at | 7.3 | A | 5.9 | A | 7.3 | A | 3.8 | A | --- |
| 262607_at | 3265.1 | P | 2298.6 | P | 1850.7 | P | 2099 | P | --- |
| 262608_at | 573.8 | P | 130.6 | P | 448.3 | P | 177.2 | P | --- |
| 262609_at | 12333.4 | P | 11427.1 | P | 15160.9 | P | 12583.6 | P | --- |
| 262610_at | 332.2 | P | 370.3 | P | 395 | P | 348.7 | P | --- |
| 262611_at | 348.6 | P | 231.8 | P | 428.7 | P | 234.9 | P | --- |
| 262612_at | 1412.9 | P | 1778 | P | 1655.3 | P | 1905.9 | P | --- |
| 262613_at | 230.7 | P | 201.4 | P | 207.5 | P | 234.3 | P | WRKY4 |
| 262614_at | 667.1 | P | 687.9 | P | 730.1 | P | 736.3 | P | GN |
| 262615_at | 276.9 | P | 352.2 | P | 179.8 | P | 377.5 | P | ELF5A-1 |
| 262616_at | 236.3 | P | 112.8 | P | 301.9 | P | 73.1 | P | --- |
| 262617_at | 68.4 | P | 85.5 | P | 79.8 | P | 50.1 | P | POR C |
| 262618_at | 464.1 | P | 363 | P | 640.7 | P | 384.5 | P | --- |
| 262619_at | 492.6 | P | 597.1 | P | 700.5 | P | 795.9 | P | --- |
| 262620_at | 21.7 | A | 4.5 | A | 5.8 | A | 19.3 | A | --- |
| 262621_at | 1752.4 | P | 1205.8 | P | 1519.5 | P | 1298.8 | P | --- |
| 262622_at | 307.5 | P | 394.2 | P | 415.1 | P | 343.2 | P | --- |
| 262623_at | 443.2 | P | 447 | P | 434.4 | P | 503 | P | --- |
| 262624_at | 111.4 | P | 167.6 | P | 140.9 | P | 121.8 | P | --- |
| 262625_at | 124.9 | P | 61.8 | P | 58.2 | A | 55.1 | P | --- |
| 262626_at | 2655.8 | P | 2136.9 | P | 2602 | P | 2299.7 | P | FTSH8 |
| 262627_at | 40.3 | P | 25.5 | P | 47.9 | A | 25.1 | A | --- |
| 262628_at | 34.3 | A | 64.5 | P | 65.8 | A | 54.5 | P | ATGSL07 |
| 262629_at | 5894.3 | P | 5777.8 | P | 5749.3 | P | 5776.3 | P | ACD32.1 |
| 262630_at | 34.5 | A | 42.5 | P | 43.2 | A | 56.3 | A | GPAT1 |
| 262631_at | 178 | P | 242.1 | P | 205.3 | P | 234.8 | P | --- |
| 262632_at | 14457.2 | P | 15001.4 | P | 14186.1 | P | 14774.2 | P | PSBP-1 |
| 262633_at | 315.3 | P | 278.5 | P | 331.4 | P | 209.3 | P | --- |
| 262634_at | 650.7 | P | 646.5 | P | 669.4 | P | 693.3 | P | --- |
| 262635_at | 1293.5 | P | 910.2 | P | 1046.2 | P | 962.7 | P | PDS1 |
| 262636_at | 283.3 | P | 259.3 | P | 341.5 | P | 256 | P | NIH |
| 262637_at | 3129.3 | P | 2569.2 | P | 2485.5 | P | 2128.6 | P | --- |
| 262638_at | 1325 | P | 1403.9 | P | 1046 | P | 1125.8 | P | --- |
| 262639_at | 731.1 | P | 619.4 | P | 586.4 | P | 576.3 | P | --- |
| 262640_at | 70.9 | P | 32.9 | A | 84.4 | A | 13.9 | A | --- |
| 262641_at | 250.5 | P | 264.3 | P | 261.7 | P | 213.1 | P | --- |
| 262642_at | 1.3 | A | 3.1 | A | 4.9 | A | 7.9 | A | --- |
| 262643_at | 29.2 | P | 103.5 | P | 48.7 | M | 106 | P | --- |
| 262644_at | 553.9 | P | 486.2 | P | 383.9 | P | 509.5 | P | BETA-VPE |
| 262645_at | 4876 | P | 5037.9 | P | 6012.7 | P | 5584.9 | P | SCO1 |
| 262646_at | 690.7 | P | 468.3 | P | 749.6 | P | 434 | P | ASP4 |
| 262647_at | 140.7 | P | 233.4 | P | 208.3 | P | 248.8 | P | --- |
| 262648_at | 325.1 | P | 633.8 | P | 405.8 | P | 516.4 | P | --- |
| 262649_at | 163.3 | P | 150.8 | P | 140.5 | P | 131.6 | P | --- |
| 262650_at | 68.6 | P | 61.1 | P | 88.7 | P | 75 | P | --- |
| 262651_at | 28.3 | A | 22.7 | A | 19.9 | A | 33.7 | A | FUT9 |
| 262652_at | 3.9 | A | 28.9 | A | 9.6 | A | 2.3 | A | FUT9 |
| 262653_at | 760.5 | P | 577.9 | P | 488 | P | 565.5 | P | --- |
| 262654_at | 37.4 | A | 98.3 | P | 90.6 | A | 131.6 | P | --- |
| 262655_s_at | 21.3 | P | 21.5 | A | 26.5 | A | 23.5 | A | --- |
| 262656_at | 252.3 | P | 300.5 | P | 270 | P | 346 | P | --- |
| 262657_at | 465.6 | P | 558 | P | 458.9 | P | 554.1 | P | --- |
| 262658_at | 3.4 | A | 6.5 | A | 26.3 | A | 8.9 | A | --- |
| 262659_at | 22.9 | P | 44.4 | P | 40.9 | P | 41.7 | P | --- |
| 262660_at | 1295.9 | P | 1204.6 | P | 996.2 | P | 930.4 | P | VIK |
| 262661_s_at | 986.7 | P | 337 | P | 725.4 | P | 305.6 | P | --- |
| 262662_at | 42.9 | P | 62.6 | P | 71.5 | P | 85.2 | P | --- |
| 262663_at | 171.2 | P | 235 | P | 202.7 | P | 202.1 | P | --- |

| | | | | | | | | | |
|---|---|---|---|---|---|---|---|---|---|
| 262664_at | 42.6 | P | 28.8 | P | 5.9 | A | 36.4 | P | --- |
| 262665_at | 29.7 | A | 8.3 | A | 4.8 | A | 8.7 | A | FUT7 |
| 262666_at | 1 | A | 1.5 | A | 2.1 | A | 1.7 | A | FUT6 |
| 262667_at | 645.5 | P | 530.6 | P | 521.4 | P | 476.6 | P | --- |
| 262668_at | 188.7 | P | 206.2 | P | 166.3 | P | 139.3 | P | LDL1 |
| 262669_at | 468.8 | P | 399 | P | 384.8 | P | 350.9 | P | --- |
| 262670_s_at | 16.6 | A | 2.7 | A | 18.2 | A | 5 | A | AT2G30740 /// HAB1 |
| 262671_at | 210.7 | P | 299 | P | 177 | P | 162 | P | CPK29 |
| 262672_at | 313.8 | P | 361.1 | P | 374.5 | P | 327.8 | P | --- |
| 262673_at | 305.5 | P | 230.9 | P | 248.5 | P | 244.7 | P | EMB1793 |
| 262674_at | 24.7 | A | 21.7 | A | 49.4 | A | 26.8 | A | EXL4 |
| 262675_at | 1.6 | A | 12.6 | A | 7.6 | A | 2.3 | A | EXL6 |
| 262676_at | 6764.6 | P | 5420.9 | P | 6797.2 | P | 5189.2 | P | SKP1 |
| 262677_at | 631.6 | P | 544.8 | P | 511.5 | P | 470.9 | P | --- |
| 262678_at | 301.3 | P | 345.5 | P | 287.7 | P | 348 | P | --- |
| 262679_at | 1.9 | A | 5.2 | A | 3 | A | 0.8 | A | LCR67 |
| 262680_at | 123 | P | 114.1 | P | 90.6 | P | 110.7 | P | --- |
| 262681_at | 1 | A | 5 | A | 4.7 | A | 2.4 | A | --- |
| 262682_at | 423.9 | P | 532.6 | P | 588.8 | P | 698.4 | P | --- |
| 262683_at | 23.3 | A | 28.6 | A | 19.2 | A | 19.9 | A | --- |
| 262684_s_at | 4047.7 | P | 4559 | P | 4157.8 | P | 4568.3 | P | --- |
| 262685_at | 3339.2 | P | 2161 | P | 3774.3 | P | 2442.5 | P | --- |
| 262686_at | 411.2 | P | 521.8 | P | 442.8 | P | 496 | P | --- |
| 262687_at | 29 | A | 19.9 | A | 17.3 | A | 17.3 | A | --- |
| 262688_at | 37.4 | A | 10.4 | A | 25.8 | A | 26.7 | A | --- |
| 262689_at | 17.5 | A | 14.9 | A | 38.1 | A | 19.1 | A | ANAC026 |
| 262690_at | 55.3 | A | 23.8 | A | 108.5 | A | 71.4 | A | --- |
| 262691_at | 1260.7 | P | 1034 | P | 896.5 | P | 846.4 | P | --- |
| 262692_at | 8.2 | A | 2 | A | 19.7 | A | 1.7 | A | --- |
| 262693_at | 1405.8 | P | 1916.4 | P | 1253.5 | P | 1683.3 | P | --- |
| 262694_at | 1303.5 | P | 1255.6 | P | 1108.3 | P | 1148.7 | P | --- |
| 262695_at | 284.6 | P | 263.7 | P | 325.8 | P | 292.6 | P | VPS35B |
| 262696_at | 9.4 | A | 34.5 | A | 20.8 | A | 7.5 | A | --- |
| 262697_at | 23.8 | A | 2 | A | 1.1 | A | 2.6 | A | ATA27 |
| 262698_at | 81.2 | P | 86.4 | P | 170.1 | P | 101.7 | P | --- |
| 262699_at | 294.3 | P | 233.8 | P | 257.5 | P | 279.9 | P | --- |
| 262700_at | 120.7 | P | 288.6 | P | 165.4 | P | 290.5 | P | --- |
| 262701_at | 54.7 | A | 80.4 | A | 58 | A | 71.1 | M | --- |
| 262702_at | 69.5 | P | 76.9 | P | 73.6 | P | 69.8 | P | --- |
| 262703_at | 150.5 | P | 362.3 | P | 160.3 | P | 322.1 | P | --- |
| 262704_at | 43 | A | 85.7 | P | 54.4 | A | 99.2 | P | ASL9 |
| 262705_at | 233.8 | P | 318.9 | P | 262.6 | P | 296.2 | P | --- |
| 262706_at | 379.7 | P | 318.9 | P | 312.5 | P | 330.2 | P | --- |
| 262707_at | 55.2 | P | 54.8 | P | 76.7 | P | 53.8 | P | --- |
| 262708_at | 229.1 | P | 284.3 | P | 216.1 | P | 209.1 | P | --- |
| 262709_at | 1478.4 | P | 1481.7 | P | 1346.3 | P | 1395.1 | P | SYP51 |
| 262710_at | 1796.2 | P | 1690.3 | P | 1991.7 | P | 1779.4 | P | --- |
| 262711_at | 201.1 | P | 341.5 | P | 191 | P | 208 | P | --- |
| 262712_at | 1227.8 | P | 1158.9 | P | 786 | P | 1386.3 | P | ATRDH2 |
| 262713_at | 210 | P | 291.9 | P | 232.1 | P | 288 | P | --- |
| 262714_s_at | 590.6 | P | 561.9 | P | 658.6 | P | 623.9 | P | --- |
| 262715_at | 30.1 | A | 44.7 | A | 38 | A | 34.7 | A | MYB58 |
| 262716_at | 1921.3 | P | 1772.7 | P | 1693.4 | P | 1918.2 | P | PAB1 |
| 262717_s_at | 1906.1 | P | 1163.9 | P | 2238.1 | P | 1069.6 | P | CYP79F2 |
| 262718_at | 27.6 | A | 11.5 | A | 30.8 | A | 30.6 | A | --- |
| 262719_at | 95 | P | 58.8 | P | 101.1 | P | 44.4 | A | --- |
| 262720_s_at | 1.3 | A | 0.9 | A | 12.1 | A | 15.5 | A | --- |
| 262721_at | 3139.5 | P | 3289.9 | P | 2743.2 | P | 2946.4 | P | Aty2 |
| 262722_at | 559.3 | P | 691.9 | P | 427.4 | P | 649 | P | --- |
| 262723_at | 16.7 | A | 22.4 | A | 40.5 | P | 22.2 | A | --- |
| 262724_s_at | 21.4 | M | 15.2 | P | 30.5 | P | 21.3 | P | --- |
| 262725_at | 690.2 | P | 714 | P | 830.5 | P | 768.4 | P | --- |
| 262726_at | 25.4 | A | 42.3 | A | 45.8 | A | 58.2 | A | AtTLP5 |
| 262727_at | 1213.8 | P | 1433.9 | P | 898.9 | P | 1528.4 | P | --- |
| 262728_at | 381.1 | P | 694.2 | P | 543 | P | 730 | P | CLV1 |
| 262729_at | 549.7 | P | 675.2 | P | 683.7 | P | 884 | P | ARAC5 |
| 262730_at | 274.8 | P | 247 | P | 384.1 | P | 259.5 | P | ATOCT3 |
| 262731_at | 40.3 | P | 12.7 | A | 26.8 | A | 14.4 | A | MC8 |
| 262732_at | 20.1 | A | 9.7 | A | 4.5 | A | 3.7 | A | --- |
| 262733_s_at | 1900.7 | P | 2192.1 | P | 1680.6 | P | 2199.6 | P | --- |
| 262734_at | 18.2 | A | 25 | A | 22 | A | 28.9 | P | --- |
| 262735_at | 6.8 | A | 7.1 | A | 10.6 | A | 24.1 | A | --- |
| 262736_at | 450.7 | P | 419 | P | 521.3 | P | 431.5 | P | --- |
| 262737_at | 55.5 | P | 55.3 | P | 66 | P | 78.4 | P | SRD2 |
| 262738_at | 298.6 | P | 293.5 | P | 352.3 | P | 309.4 | P | --- |
| 262739_at | 16.2 | A | 20.1 | A | 29.9 | A | 15.9 | A | --- |
| 262740_at | 25.1 | A | 21.3 | A | 47.4 | P | 23.1 | A | --- |
| 262741_at | 147.8 | P | 135.5 | P | 173.7 | P | 135.6 | P | --- |
| 262742_at | 1.1 | A | 0.8 | A | 2.2 | A | 1.4 | A | AtRABA1i |
| 262743_at | 23.1 | A | 13.7 | A | 32.7 | A | 31 | A | --- |
| 262744_at | 394.3 | P | 421.6 | P | 352.7 | P | 446.7 | P | --- |
| 262745_at | 5020.3 | P | 3481.2 | P | 5122.2 | P | 3474.3 | P | --- |
| 262746_at | 411.6 | P | 346.6 | P | 331.1 | P | 313.5 | P | ATNUDX15 |
| 262747_at | 297.8 | P | 274.6 | P | 270.5 | P | 250.1 | P | --- |
| 262748_at | 620 | P | 455.5 | P | 604.1 | P | 497.3 | P | --- |
| 262749_at | 2347.6 | P | 2235 | P | 2135.9 | P | 2123.5 | P | --- |
| 262750_at | 150.3 | P | 160.3 | P | 164.1 | P | 155.8 | P | --- |
| 262751_at | 84.5 | P | 86.8 | P | 107.2 | P | 94.1 | P | --- |
| 262752_at | 23.3 | A | 92.9 | P | 79.5 | P | 105.8 | P | CYCB3;1 |
| 262753_at | 368.2 | P | 386.1 | P | 459.2 | P | 464.5 | P | ATKDSA2 |
| 262754_at | 1665.4 | P | 1638.9 | P | 2176.6 | P | 1764.9 | P | --- |
| 262755_at | 18.7 | A | 36.2 | A | 63.4 | A | 24.2 | A | --- |
| 262756_at | 62.2 | P | 77.9 | P | 111.9 | A | 88 | P | 41553 |
| 262757_at | 4.9 | A | 7.6 | A | 5.8 | A | 3.3 | A | ATCHX1 |
| 262758_at | 140.5 | P | 186.5 | P | 185.2 | P | 182.2 | P | --- |
| 262759_at | 51.8 | A | 43.6 | P | 5.6 | A | 33.7 | P | --- |
| 262760_at | 106.3 | P | 143.2 | P | 134.7 | P | 118.8 | P | --- |
| 262761_at | 181.3 | P | 213.9 | P | 168.1 | P | 233.4 | P | --- |
| 262762_at | 1351.1 | P | 486.4 | P | 768.6 | P | 546.4 | P | --- |
| 262763_at | 8.9 | A | 18.3 | A | 61.3 | P | 1.8 | A | --- |
| 262764_at | 40.8 | A | 42.8 | M | 52 | A | 32.2 | A | CYP86C3 |
| 262765_at | 38.6 | A | 56.1 | A | 38.1 | A | 13.9 | A | CYP86C4 |
| 262766_at | 368.7 | P | 269 | P | 552.8 | P | 276.4 | P | --- |
| 262767_at | 158 | P | 286.3 | P | 191.2 | P | 291.3 | P | ORP1D |
| 262768_at | 153.8 | P | 204.4 | P | 194.9 | P | 202.1 | P | --- |
| 262769_at | 254.3 | P | 289.4 | P | 215.2 | P | 248.1 | P | DIS1 |
| 262770_at | 494.4 | P | 448.1 | P | 423.7 | P | 526.9 | P | --- |
| 262771_at | 2.3 | A | 4.9 | A | 18.2 | A | 17.3 | A | --- |
| 262772_at | 280.6 | P | 352.7 | P | 215.9 | P | 286.1 | P | ACA.l |

| | | | | | | | | | |
|---|---|---|---|---|---|---|---|---|---|
| 262773_at | 135.6 | P | 175.7 | P | 152.9 | P | 135.6 | P | LINC2 |
| 262774_at | 16.3 | P | 23.5 | M | 29.8 | A | 17.2 | A | --- |
| 262775_at | 296.1 | P | 390.8 | P | 183.8 | P | 339.8 | P | --- |
| 262776_at | 456 | P | 647 | P | 561.2 | P | 660.4 | P | --- |
| 262777_at | 208.4 | P | 220.6 | P | 272.5 | P | 249.4 | P | --- |
| 262778_at | 100.6 | P | 90.6 | P | 84.1 | P | 71.4 | P | --- |
| 262779_at | 34 | A | 9.1 | A | 21.1 | A | 11.6 | A | --- |
| 262780_at | 594.9 | P | 628.2 | P | 494.9 | P | 590.2 | P | CYP71B28 |
| 262781_s_at | 3498.8 | P | 4289.2 | P | 3577.2 | P | 4234.5 | P | PBE1 |
| 262782_at | 342.2 | P | 285.3 | P | 249.9 | P | 260.9 | P | --- |
| 262783_at | 166.5 | P | 322.1 | P | 254.7 | P | 367.1 | P | --- |
| 262784_at | 5588.8 | P | 5817.7 | P | 5579.2 | P | 6154.8 | P | SEX1 |
| 262785_at | 66.4 | P | 73.6 | P | 106.6 | P | 83.4 | P | --- |
| 262786_at | 249.8 | P | 380.1 | P | 340.9 | P | 329.8 | P | --- |
| 262787_at | 316.6 | P | 273.9 | P | 324.4 | P | 254.2 | P | --- |
| 262788_at | 20.8 | A | 78.7 | P | 27 | A | 77.4 | M | --- |
| 262789_at | 19.9 | A | 26.1 | A | 19.8 | A | 5.7 | A | --- |
| 262790_at | 1637.9 | P | 1392.8 | P | 1764.3 | P | 1409.1 | P | TIF3H1 |
| 262791_at | 706.4 | P | 851.6 | P | 738.5 | P | 802 | P | --- |
| 262792_at | 16.4 | A | 22.7 | A | 1.7 | A | 22 | A | --- |
| 262793_at | 1134.2 | P | 1350.6 | P | 1148.2 | P | 1289.1 | P | CYP71B7 |
| 262794_at | 380.2 | P | 402.4 | P | 400.4 | P | 375.7 | P | emb1745 |
| 262795_at | 29 | A | 11 | A | 20.4 | A | 32.4 | A | --- |
| 262796_at | 436.4 | P | 940.2 | P | 990.4 | P | 960.4 | P | XCP2 |
| 262797_at | 1663.9 | P | 924.3 | P | 1524.3 | P | 961.4 | P | TMT1 |
| 262798_at | 1057.9 | P | 1363.1 | P | 1117.4 | P | 1042.6 | P | SPL14 |
| 262799_at | 27.5 | A | 31.9 | A | 70.2 | A | 37.8 | A | --- |
| 262800_at | 1079.1 | P | 953.4 | P | 1110 | P | 918.7 | P | emb1507 |
| 262801_at | 254.4 | P | 196 | P | 136.9 | P | 169.5 | P | --- |
| 262802_at | 199.8 | P | 259.7 | P | 263.6 | P | 262.6 | P | CDKB2;2 |
| 262803_at | 1106.8 | P | 1104 | P | 887.8 | P | 952.7 | P | --- |
| 262804_at | 167 | P | 211.7 | P | 152.8 | P | 208.3 | P | --- |
| 262805_at | 68.9 | P | 105.6 | P | 98.1 | P | 102.2 | P | ESC |
| 262806_at | 1379.6 | P | 1736.9 | P | 1604.4 | P | 2198.6 | P | --- |
| 262807_at | 46.2 | M | 12.4 | A | 22.9 | A | 22.2 | A | --- |
| 262808_at | 58.6 | P | 74.5 | P | 38.5 | A | 70.7 | P | --- |
| 262809_at | 449.4 | P | 497.3 | P | 492.8 | P | 507.2 | P | ATSS3 |
| 262810_at | 63.2 | P | 43.9 | A | 44.1 | A | 47.3 | P | --- |
| 262811_at | 358.4 | P | 513.5 | P | 420.1 | P | 488.1 | P | --- |
| 262812_at | 6.7 | A | 0.6 | A | 2.7 | A | 1.7 | A | --- |
| 262813_at | 240.2 | P | 173.8 | P | 263.4 | P | 213.8 | P | --- |
| 262814_at | 698.8 | P | 690.4 | P | 640.8 | P | 670.8 | P | --- |
| 262815_at | 0.4 | A | 0.7 | A | 1.7 | A | 0.4 | A | CYP71A18 |
| 262816_at | 285.9 | P | 379.6 | P | 378.7 | P | 368 | P | --- |
| 262817_at | 21.1 | M | 18.1 | M | 22.8 | M | 6 | A | --- |
| 262818_at | 271.3 | P | 294.5 | P | 292.7 | P | 267.6 | P | --- |
| 262819_at | 92.2 | P | 163.9 | P | 126.8 | P | 180.4 | P | CYP77B1 |
| 262820_at | 1279.8 | P | 1331.3 | P | 1447.5 | P | 1436.9 | P | CYP51G1 |
| 262821_at | 218.2 | P | 201.5 | P | 284.5 | P | 229.5 | P | --- |
| 262822_at | 264.9 | P | 211.4 | P | 257 | P | 220.7 | P | --- |
| 262823_at | 2299 | P | 2671.4 | P | 2466.2 | P | 2667.5 | P | CLPP6 |
| 262824_at | 4272.9 | P | 4018.5 | P | 4124.7 | P | 4121 | P | RBP45B |
| 262825_at | 731.3 | P | 841.7 | P | 887.7 | P | 889.2 | P | ADT1 |
| 262826_at | 283.4 | P | 457.8 | P | 235.1 | P | 326.1 | P | CYP71B2 |
| 262827_at | 89.2 | P | 104.6 | P | 85.5 | P | 107.5 | P | CYP71B29 |
| 262828_at | 25.4 | A | 44 | A | 32.3 | A | 19.5 | A | --- |
| 262829_at | 51.5 | P | 85.7 | P | 70.3 | M | 100.2 | P | --- |
| 262830_at | 242 | P | 483.6 | P | 188.9 | P | 429.9 | P | PAP3 |
| 262831_at | 16.4 | A | 62.1 | P | 61.6 | A | 50.5 | A | --- |
| 262832_s_at | 1931.7 | P | 1250.6 | P | 1403.7 | P | 1164.2 | P | --- |
| 262833_at | 29.5 | A | 16.4 | A | 31.9 | A | 3.1 | A | SDS |
| 262834_at | 54.9 | A | 52.1 | A | 70.4 | M | 64 | M | MAP65-7 |
| 262835_at | 69 | P | 87.5 | P | 83.3 | A | 77.3 | P | ATNHX8 |
| 262836_at | 62.4 | P | 29.5 | A | 47.6 | A | 44.7 | P | --- |
| 262837_at | 986.6 | P | 1041.3 | P | 915.2 | P | 1108.4 | P | ADL1C |
| 262838_at | 53.3 | A | 69 | A | 50.5 | A | 54.1 | A | --- |
| 262839_at | 31.9 | P | 74 | P | 55.2 | P | 57.9 | P | --- |
| 262840_at | 307.9 | P | 550.2 | P | 480.1 | P | 554.6 | P | HMGA |
| 262841_at | 4636 | P | 3197 | P | 5145.7 | P | 3430.4 | P | --- |
| 262842_at | 1651.5 | P | 1336.9 | P | 1332.3 | P | 1275.8 | P | XTR2 |
| 262843_at | 45.1 | P | 44.2 | P | 32 | A | 34.5 | A | AtHB32 |
| 262844_at | 350.9 | P | 853.6 | P | 480.3 | P | 659.4 | P | --- |
| 262845_at | 297.1 | P | 334.3 | P | 260.1 | P | 331.7 | P | --- |
| 262846_at | 1339.3 | P | 2062.4 | P | 1281.1 | P | 1786 | P | --- |
| 262847_at | 203.4 | P | 280.8 | P | 238.8 | P | 228.8 | P | ATMAP70-4 |
| 262848_at | 497.8 | P | 713.4 | P | 474.8 | P | 615.8 | P | BPC2 |
| 262849_at | 819.4 | P | 857.5 | P | 1022.2 | P | 894.4 | P | --- |
| 262850_at | 634.1 | P | 669.9 | P | 460.7 | P | 595.8 | P | GAI |
| 262851_at | 186.5 | P | 188.4 | P | 273.6 | P | 190 | P | --- |
| 262852_at | 188.9 | P | 140.8 | P | 185.3 | P | 224.9 | P | --- |
| 262853_at | 318.3 | P | 311.9 | P | 250.2 | P | 249.5 | P | --- |
| 262854_at | 52.8 | P | 82.9 | P | 41.7 | A | 70 | P | --- |
| 262855_at | 22.5 | A | 37.2 | A | 42.4 | A | 10.4 | A | --- |
| 262856_at | 345.1 | P | 388 | P | 293.5 | P | 367.4 | P | --- |
| 262857_at | 9.6 | A | 13.1 | A | 15.3 | A | 15.1 | A | --- |
| 262858_at | 3.1 | A | 29.7 | A | 3.8 | A | 16.5 | A | --- |
| 262859_at | 597.1 | P | 602.8 | P | 573.6 | P | 533.9 | P | --- |
| 262860_at | 589.3 | P | 595 | P | 527.7 | P | 659.2 | P | APO1 |
| 262861_at | 24.2 | A | 12.5 | A | 1.7 | A | 22.9 | A | --- |
| 262862_at | 2 | A | 6.9 | A | 1.7 | A | 1 | A | --- |
| 262863_at | 12.9 | A | 23 | P | 26.9 | A | 21.5 | P | --- |
| 262864_at | 13.5 | P | 8.7 | A | 18.1 | A | 1.5 | A | --- |
| 262865_at | 3.9 | A | 26.8 | A | 7.7 | A | 25.8 | A | CYP89A7 |
| 262866_at | 8.3 | A | 13.9 | P | 2.4 | A | 9.7 | A | CYP89A6 |
| 262867_at | 129.7 | P | 159.2 | P | 149.4 | P | 209.4 | P | --- |
| 262868_at | 759.2 | P | 1970.4 | P | 1760.3 | P | 2163.8 | P | --- |
| 262869_s_at | 430.1 | P | 482.4 | P | 431.4 | P | 474.7 | P | GTG1 |
| 262870_at | 1102.8 | P | 643.1 | P | 927.9 | P | 711.8 | P | --- |
| 262871_at | 286.4 | P | 383.1 | P | 264.7 | P | 394 | P | --- |
| 262872_at | 71.3 | P | 83.2 | P | 61.4 | P | 62 | P | --- |
| 262873_at | 143.5 | P | 176.2 | P | 164 | P | 116.2 | P | --- |
| 262874_at | 277.5 | P | 341.5 | P | 317.9 | P | 302.2 | P | --- |
| 262875_at | 509.6 | P | 737.5 | P | 564.5 | P | 672.2 | P | G-TMT |
| 262876_at | 887.4 | P | 834.8 | P | 954.8 | P | 924.6 | P | ATDSS1(I) |
| 262877_at | 2452.8 | P | 2684.4 | P | 2413.7 | P | 2375.1 | P | --- |
| 262878_at | 1166.8 | P | 1992.5 | P | 1244.1 | P | 2191.8 | P | NDF2 |
| 262879_at | 1103.9 | P | 1070.1 | P | 1377.9 | P | 1275.5 | P | SIGA |
| 262880_at | 1401 | P | 1272.2 | P | 1552.5 | P | 1331.4 | P | --- |
| 262881_at | 277.8 | P | 323.6 | P | 265.5 | P | 370.8 | P | --- |

| | | | | | | | | | |
|---|---|---|---|---|---|---|---|---|---|
| 262882_at | 1477 | P | 1321.8 | P | 1403.4 | P | 1378.5 | P | CYP89A2 |
| 262883_at | 1232.6 | P | 981.4 | P | 1741.3 | P | 1346.7 | P | ATAMT1;2 |
| 262884_at | 9463.8 | P | 8898 | P | 9604.8 | P | 7287.4 | P | CP5 |
| 262885_at | 558 | P | 529.4 | P | 458.3 | P | 456.9 | P | TUA1 |
| 262886_at | 62 | A | 77 | P | 75.4 | A | 55.3 | A | KNATM |
| 262887_at | 319.2 | P | 181 | P | 195.7 | P | 121.5 | P | --- |
| 262888_at | 152.7 | P | 195.7 | P | 95.9 | P | 151.3 | P | RDR1 |
| 262889_at | 7.8 | A | 21.9 | A | 29.1 | A | 3 | A | --- |
| 262890_at | 356 | P | 389.5 | P | 407 | P | 379.4 | P | NUP155 |
| 262891_at | 78.5 | P | 212.4 | P | 153.7 | P | 229.1 | P | GA2 |
| 262892_at | 1758.3 | P | 1287 | P | 1756.1 | P | 1406.8 | P | ALDH5F1 |
| 262893_at | 26.1 | A | 36.1 | P | 35.2 | P | 27.9 | A | --- |
| 262894_at | 16.8 | A | 17.5 | A | 21.5 | A | 28.1 | A | --- |
| 262895_at | 7 | A | 1.9 | A | 36.8 | A | 22.9 | A | --- |
| 262896_at | 317.6 | P | 416.5 | P | 278.6 | P | 354 | P | ALA3 |
| 262897_at | 338.2 | P | 588.8 | P | 318.5 | P | 526.4 | P | CCB4 |
| 262898_at | 12.3 | A | 77.5 | A | 40.6 | A | 50.3 | A | --- |
| 262899_at | 6050 | P | 5449.5 | P | 4451 | P | 4862.6 | P | PEN3 |
| 262900_at | 192.8 | P | 205.6 | P | 261.1 | P | 234 | P | SNL5 |
| 262901_at | 258.5 | P | 268 | P | 226.3 | P | 262.1 | P | --- |
| 262902_x_at | 26 | A | 10.3 | A | 50.3 | A | 12.5 | A | --- |
| 262903_at | 5.6 | A | 7.6 | A | 5.8 | A | 1.7 | A | --- |
| 262904_at | 12 | A | 32.4 | A | 43.2 | A | 35.5 | A | --- |
| 262905_at | 143.4 | P | 92.7 | P | 70.7 | P | 77.1 | P | ATH7 |
| 262906_at | 509.9 | P | 416.5 | P | 566.2 | P | 419.4 | P | --- |
| 262907_at | 69.7 | P | 67.1 | P | 104.7 | P | 121.4 | P | CRR28 |
| 262908_at | 3514.5 | P | 3451 | P | 3710.9 | P | 3671.2 | P | AT-E1 ALPHA |
| 262909_at | 562.3 | P | 631.2 | P | 547.5 | P | 635.9 | P | PP2A-1 |
| 262910_at | 469.7 | P | 521 | P | 578.5 | P | 551.8 | P | --- |
| 262911_s_at | 60.6 | P | 212.6 | P | 67.3 | A | 172.7 | P | --- |
| 262912_at | 146.9 | P | 214.4 | P | 177.6 | P | 198.2 | P | --- |
| 262913_at | 877 | P | 763 | P | 812.4 | P | 736.4 | P | --- |
| 262914_at | 361.2 | P | 335 | P | 380.8 | P | 335.5 | P | ARF1 |
| 262915_at | 7 | A | 2.2 | A | 5.4 | A | 5.3 | A | ARR3 |
| 262916_at | 1697.3 | P | 1163.7 | P | 1489.3 | P | 1278.3 | P | ATGSTU16 |
| 262917_at | 22.8 | A | 8.1 | A | 21.3 | A | 10.2 | A | --- |
| 262918_at | 114.8 | P | 109.7 | A | 115.2 | A | 110.2 | A | --- |
| 262919_at | 390 | P | 796.9 | P | 479.6 | P | 561.4 | P | --- |
| 262920_at | 264.6 | P | 396.8 | P | 390.5 | P | 410.4 | P | --- |
| 262921_at | 257.2 | P | 242.6 | P | 349.6 | P | 267.6 | P | APL |
| 262922_at | 68.1 | P | 88 | P | 51 | P | 88.6 | P | --- |
| 262923_at | 641.9 | P | 657.5 | P | 677.4 | P | 744.2 | P | XRN3 |
| 262924_s_at | 6202.2 | P | 6201.2 | P | 5923.5 | P | 5916.6 | P | AVA-P4 |
| 262925_at | 106.6 | P | 88.5 | P | 41.7 | A | 71 | P | YSL7 |
| 262926_s_at | 266.7 | P | 343.2 | P | 238.6 | P | 336.6 | P | ARK1 |
| 262927_at | 1.4 | A | 1.4 | A | 5.4 | A | 1.9 | A | --- |
| 262928_at | 14 | A | 3.8 | A | 12.6 | A | 2.9 | A | --- |
| 262929_at | 5083.4 | P | 5087.7 | P | 4756.1 | P | 5269.9 | P | --- |
| 262930_at | 201.3 | P | 138.9 | P | 43.3 | P | 64.9 | P | --- |
| 262931_at | 701.9 | P | 535.2 | P | 566.6 | P | 583.4 | P | --- |
| 262932_at | 6409.5 | P | 5041.7 | P | 6411.2 | P | 4951.3 | P | --- |
| 262933_at | 625.7 | P | 775 | P | 752.5 | P | 879.4 | P | ATPAO4 |
| 262934_s_at | 309 | P | 381 | P | 300.9 | P | 356.1 | P | --- |
| 262935_at | 1160.8 | P | 751.8 | P | 1075.4 | P | 1058.7 | P | AtOCT5 |
| 262936_at | 53.2 | P | 17.6 | M | 92.9 | P | 42.1 | P | ATCHX2 |
| 262937_at | 1328.3 | P | 1339.7 | P | 1367 | P | 1441.9 | P | FTSH12 |
| 262938_at | 161.3 | A | 152.4 | M | 221.7 | A | 177.3 | A | --- |
| 262939_s_at | 588 | P | 570.3 | P | 790 | P | 623.2 | P | GAPCP-2 |
| 262940_at | 382.8 | P | 253.1 | P | 317.6 | P | 341 | P | --- |
| 262941_at | 172 | P | 127.7 | P | 140.1 | P | 116.2 | P | EMB2217 |
| 262942_at | 22.1 | A | 39.5 | P | 2.1 | A | 24.9 | A | ALIS5 |
| 262943_at | 329.7 | P | 269.2 | P | 482.5 | P | 318.5 | P | --- |
| 262944_at | 6238.4 | P | 5702.2 | P | 7677.6 | P | 6358.5 | P | PGK |
| 262945_at | 878.6 | P | 988.4 | P | 1005.3 | P | 886.9 | P | --- |
| 262946_at | 858.9 | P | 745.7 | P | 999.1 | P | 779.2 | P | --- |
| 262947_at | 2842.4 | P | 1215 | P | 1181.7 | P | 1905.2 | P | GASA1 |
| 262948_at | 666.8 | P | 904.7 | P | 766.9 | P | 949.1 | P | --- |
| 262949_at | 26.2 | A | 27.8 | P | 53.2 | A | 13.2 | A | sks18 |
| 262950_at | 454.5 | P | 417.6 | P | 479.6 | P | 407.6 | P | --- |
| 262951_at | 953.3 | P | 2003.7 | P | 1207.6 | P | 1894.1 | P | --- |
| 262952_at | 99.3 | P | 110.3 | P | 117.6 | P | 113.7 | P | --- |
| 262953_at | 409 | P | 330.8 | P | 530.9 | P | 362.7 | P | --- |
| 262954_at | 2639.9 | P | 2717.7 | P | 2216.7 | P | 2616.6 | P | --- |
| 262955_at | 1547.1 | P | 1578 | P | 1268.5 | P | 1607.5 | P | --- |
| 262956_at | 4445.2 | P | 3340.9 | P | 4023.5 | P | 3869.4 | P | EIF4A-2 |
| 262957_s_at | 1611.1 | P | 1421.5 | P | 1259.1 | P | 1550 | P | NRPB8A |
| 262958_at | 10019.9 | P | 10144.2 | P | 10038.5 | P | 9854.5 | P | --- |
| 262959_at | 645.9 | P | 600.4 | P | 604.7 | P | 704.9 | P | --- |
| 262960_at | 1308 | P | 1588.4 | P | 1356.2 | P | 1290.7 | P | --- |
| 262961_at | 585.5 | P | 525.6 | P | 491 | P | 565.7 | P | XRN4 |
| 262962_at | 470.6 | P | 408.9 | P | 469.9 | P | 425.8 | P | ICDH |
| 262963_at | 669.4 | P | 534.1 | P | 550.1 | P | 564.3 | P | --- |
| 262964_at | 405.9 | P | 248.6 | P | 423.5 | P | 353.2 | P | --- |
| 262965_at | 290.2 | P | 330.7 | P | 487.5 | P | 264.3 | P | --- |
| 262966_at | 940.3 | P | 765.8 | P | 740.1 | P | 718.6 | P | --- |
| 262967_at | 158.1 | P | 214.4 | P | 189.7 | P | 150.5 | P | --- |
| 262968_at | 19.1 | A | 22.3 | P | 86.9 | A | 40.7 | P | --- |
| 262969_at | 468 | P | 527 | P | 353.3 | P | 507.2 | P | --- |
| 262970_at | 1947.9 | P | 2889.7 | P | 2200.2 | P | 2708.5 | P | --- |
| 262971_at | 54.5 | P | 79.4 | P | 69.1 | P | 82.8 | P | --- |
| 262972_at | 56.7 | A | 51.6 | A | 95.3 | A | 45.1 | A | --- |
| 262973_at | 0.3 | A | 0.7 | A | 1.5 | A | 0.3 | A | --- |
| 262974_at | 40.3 | A | 48.2 | P | 73.9 | P | 57.5 | P | --- |
| 262975_at | 291.7 | P | 435.2 | P | 424.8 | P | 304.8 | P | STH2 |
| 262976_at | 6.8 | A | 11.2 | A | 10.3 | A | 26.4 | A | SRS5 |
| 262977_at | 64.2 | A | 59.3 | A | 88.9 | A | 23.8 | A | --- |
| 262978_at | 161.4 | P | 513.9 | P | 201.9 | P | 481 | P | TUB1 |
| 262979_s_at | 0.4 | A | 0.5 | A | 0.1 | A | 2.6 | A | --- |
| 262980_at | 1318 | P | 1857.5 | P | 1279.4 | P | 1778.3 | P | AtGH9B7 |
| 262981_at | 6.1 | A | 35 | A | 31 | A | 30.2 | A | --- |
| 262982_at | 538.6 | P | 541.7 | P | 558.1 | P | 528.9 | P | ING2 |
| 262983_at | 10.3 | A | 20.9 | A | 31 | A | 3.4 | A | --- |
| 262984_at | 331.9 | P | 473.4 | P | 349.3 | P | 471.3 | P | --- |
| 262985_s_at | 13828.2 | P | 13280.3 | P | 16179.5 | P | 14430.7 | P | RPL27AB |
| 262986_at | 631.8 | P | 665.4 | P | 431.3 | P | 485.3 | P | --- |
| 262987_at | 4.3 | A | 28.3 | A | 23.2 | A | 14.2 | A | --- |
| 262988_at | 8738.6 | P | 9954.4 | P | 9640.7 | P | 9724.9 | P | GGT1 |
| 262989_at | 46.2 | A | 37.6 | A | 56.9 | A | 16.1 | A | INO |
| 262990_at | 919.1 | P | 958.5 | P | 860.8 | P | 882.3 | P | MMZ1 |

| | | | | | | | | | |
|---|---|---|---|---|---|---|---|---|---|
| 262991_at | 9 | A | 12.2 | A | 20.1 | A | 17.6 | A | BRX-LIKE3 |
| 262992_at | 222.7 | P | 297.7 | P | 156.4 | P | 217.1 | P | ATG12A |
| 262993_at | 28.8 | A | 37.7 | P | 48.8 | A | 18.3 | A | --- |
| 262994_at | 2.8 | A | 8.8 | A | 1.1 | A | 3.2 | A | --- |
| 262995_s_at | 3.2 | A | 8.3 | A | 26.7 | A | 14.6 | A | --- |
| 262996_at | 219.9 | P | 244.8 | P | 230.6 | P | 274.8 | P | --- |
| 262997_at | 38.7 | A | 20.2 | A | 48.6 | A | 31 | A | RPP27 |
| 262998_at | 56.2 | A | 4.7 | A | 22 | A | 22.8 | A | --- |
| 262999_at | 19.2 | A | 6.7 | A | 33.8 | A | 24.9 | A | ANAC020 |
| 263000_at | 898.2 | P | 1063.7 | P | 799.8 | P | 1040.4 | P | --- |
| 263001_at | 596.9 | P | 544 | P | 573.9 | P | 581.5 | P | TAF6B4 |
| 263002_at | 168.2 | P | 138.6 | P | 129.2 | P | 123.9 | P | --- |
| 263003_at | 71.3 | P | 64.7 | A | 49.5 | A | 67.4 | A | --- |
| 263004_at | 129.3 | P | 172.4 | P | 129.1 | P | 163.1 | P | ATNEK1 |
| 263005_at | 2 | A | 6.5 | A | 14 | A | 12.5 | A | --- |
| 263006_at | 2.1 | A | 1.3 | A | 9 | A | 2.6 | A | --- |
| 263007_at | 295.8 | P | 277.3 | P | 270.8 | P | 288 | P | --- |
| 263008_at | 3.9 | A | 1.2 | A | 2.4 | A | 2.7 | A | RPP27 |
| 263009_at | 563.1 | P | 550 | P | 577.9 | P | 463.5 | P | NHX5 |
| 263010_at | 319.3 | P | 269.2 | P | 307 | P | 234.7 | P | --- |
| 263011_at | 24.5 | A | 7 | A | 22.7 | A | 19.4 | A | --- |
| 263012_at | 25 | A | 51.7 | A | 46.7 | A | 58.1 | P | KNAT6 |
| 263013_at | 57.4 | A | 68.2 | P | 71.4 | A | 46.9 | A | KNAT6 |
| 263014_at | 678.4 | P | 964 | P | 854.9 | P | 892.4 | P | CAF2 |
| 263015_at | 1553.8 | P | 948.6 | P | 755.9 | P | 778.8 | P | --- |
| 263016_at | 339 | P | 196.7 | P | 301.8 | P | 250.6 | P | --- |
| 263017_at | 55.9 | P | 68.1 | P | 88.1 | P | 71.6 | P | CYCB2;1 |
| 263018_at | 18.2 | A | 23.1 | A | 53.1 | A | 28.6 | A | --- |
| 263019_at | 98.8 | P | 92.9 | P | 83.1 | P | 73.6 | P | ATTPS9 |
| 263020_at | 52.5 | P | 69.6 | P | 73.5 | P | 112.7 | P | --- |
| 263021_at | 1.2 | A | 6.1 | A | 14.7 | A | 0.2 | A | --- |
| 263022_s_at | 45.4 | P | 92.4 | P | 79.8 | P | 56.2 | P | --- |
| 263023_at | 1260.3 | P | 1143.1 | P | 1129.5 | P | 1175.9 | P | --- |
| 263024_at | 86.8 | P | 72.7 | P | 77 | P | 111.6 | P | --- |
| 263025_at | 2.5 | A | 12.4 | A | 1.5 | A | 15.2 | A | --- |
| 263026_at | 12.7 | A | 3.4 | A | 20.1 | A | 3.4 | A | --- |
| 263027_at | 6.3 | A | 0.8 | A | 1.7 | A | 1.4 | A | --- |
| 263028_at | 48.9 | A | 51 | A | 42.9 | A | 37.4 | P | --- |
| 263029_at | 1474.5 | P | 1403.8 | P | 1575.1 | P | 1408.5 | P | --- |
| 263030_at | 19 | A | 1.2 | A | 36.1 | A | 23.7 | A | --- |
| 263031_at | 668 | P | 225.6 | P | 588 | P | 350 | P | ATCSLA10 |
| 263032_at | 127.8 | P | 37.1 | P | 143.9 | P | 39.5 | M | --- |
| 263033_s_at | 1180.7 | P | 1165.5 | P | 1361.9 | P | 1192.3 | P | --- |
| 263034_at | 9582 | P | 9851.5 | P | 10862.1 | P | 11088.4 | P | MLP423 |
| 263035_at | 734.6 | P | 655.3 | P | 788.9 | P | 625 | P | RSZP21 |
| 263036_at | 140.6 | P | 225.7 | P | 206.2 | P | 200.6 | P | --- |
| 263037_at | 312.2 | P | 314.2 | P | 379.1 | P | 317.7 | P | --- |
| 263038_at | 2.9 | A | 4.4 | A | 8.8 | A | 3.7 | A | --- |
| 263039_at | 695.8 | P | 360.4 | P | 580.4 | P | 463.7 | P | --- |
| 263040_at | 25.6 | A | 19.1 | A | 50.4 | M | 8.8 | A | --- |
| 263041_at | 1.5 | A | 11.9 | A | 4.9 | A | 13.1 | A | TAR1 |
| 263042_at | 46.6 | P | 55.1 | A | 92.6 | A | 48.9 | P | --- |
| 263043_at | 3.2 | A | 8.6 | A | 4.3 | A | 4.2 | A | --- |
| 263044_at | 698.1 | P | 664.2 | P | 573.5 | P | 605.3 | P | --- |
| 263045_at | 6.2 | A | 6.7 | A | 68.1 | A | 9.8 | A | --- |
| 263046_at | 19119.7 | P | 14212.9 | P | 22388.4 | P | 15695.2 | P | GRP3S |
| 263047_at | 761.7 | P | 655.8 | P | 761.1 | P | 716.2 | P | --- |
| 263048_s_at | 2362 | P | 2554.1 | P | 2501.4 | P | 2522.2 | P | --- |
| 263049_at | 246.8 | P | 315.3 | P | 328.6 | A | 301.2 | P | --- |
| 263050_at | 627 | P | 590.5 | P | 548.5 | P | 633.4 | P | ATVPS11 |
| 263051_s_at | 1 | A | 1.8 | A | 17.5 | A | 1.7 | A | --- |
| 263052_at | 33 | M | 18.2 | A | 1.7 | A | 22.3 | A | --- |
| 263053_at | 312.6 | P | 299.1 | P | 346.4 | P | 301.6 | P | --- |
| 263054_at | 132.4 | P | 144.3 | P | 158.1 | P | 156.5 | P | --- |
| 263055_at | 13.2 | A | 1.9 | A | 3 | A | 13.6 | A | --- |
| 263056_at | 0.6 | A | 1.1 | A | 1.2 | A | 0.5 | A | --- |
| 263057_at | 29.3 | A | 15.6 | A | 10.2 | A | 14 | A | --- |
| 263058_at | 3.1 | A | 11.8 | A | 9.7 | A | 2.2 | A | --- |
| 263059_at | 17.7 | A | 28.7 | A | 43 | A | 31 | P | --- |
| 263060_s_at | 119.7 | P | 83.7 | A | 140.6 | A | 72.6 | A | --- |
| 263061_at | 554.7 | P | 44.3 | P | 373.2 | P | 30.5 | A | --- |
| 263062_at | 49.7 | P | 46.9 | P | 112.1 | P | 44.1 | P | --- |
| 263063_s_at | 42.4 | P | 150.6 | P | 131.7 | P | 259.5 | P | --- |
| 263064_at | 892.7 | P | 1194.2 | P | 1174.6 | P | 1101.1 | P | CPuORF1 |
| 263065_at | 822.3 | P | 723.1 | P | 656.2 | P | 787.8 | P | ATMPK7 |
| 263066_at | 287.8 | P | 253.5 | P | 305.2 | P | 250 | P | --- |
| 263067_at | 129.5 | P | 150.9 | P | 80 | P | 127.1 | P | --- |
| 263068_at | 105.3 | P | 124.9 | P | 119.6 | P | 169.4 | P | --- |
| 263069_at | 58.6 | A | 50.9 | A | 35.5 | A | 42.5 | A | --- |
| 263070_at | 12.3 | A | 22 | A | 24.8 | A | 26 | A | --- |
| 263071_at | 2.1 | A | 3.4 | A | 0.8 | A | 2 | A | --- |
| 263072_at | 349.9 | P | 361.7 | P | 287.3 | P | 355.9 | P | --- |
| 263073_at | 5140.4 | P | 1747.5 | P | 4267.6 | P | 2289.8 | P | --- |
| 263074_at | 539.6 | P | 831.9 | P | 695.8 | P | 892.3 | P | HMGB4 |
| 263075_at | 160.1 | P | 138.6 | P | 142.8 | P | 117.8 | P | --- |
| 263076_at | 676.9 | P | 602.4 | P | 644.1 | P | 534.9 | P | IRE1A |
| 263077_at | 509.4 | P | 462.1 | P | 599.9 | P | 429.3 | P | EMB2763 |
| 263078_at | 28.5 | M | 14.9 | A | 10.3 | A | 24.7 | A | --- |
| 263079_at | 32.5 | A | 27.1 | A | 47.4 | P | 17.6 | A | --- |
| 263080_at | 8.8 | A | 9.8 | A | 24.9 | A | 2.3 | A | --- |
| 263081_at | 18.6 | A | 16.1 | A | 38.5 | P | 26.6 | A | BLH5 |
| 263082_at | 297.4 | P | 273.4 | P | 282.6 | P | 300.2 | P | --- |
| 263083_at | 623.1 | P | 627.8 | P | 639.5 | P | 615.2 | P | PAP12 |
| 263084_at | 35.1 | A | 9 | A | 31.9 | A | 29 | A | --- |
| 263085_at | 169.2 | P | 203.6 | P | 178.1 | P | 228.3 | P | ARI12 |
| 263086_at | 60.3 | P | 40.4 | A | 48.6 | A | 48.5 | P | --- |
| 263087_s_at | 4.2 | A | 4.4 | A | 5 | A | 5.4 | A | --- |
| 263088_at | 7.7 | A | 1.7 | A | 1.2 | A | 4.7 | A | --- |
| 263089_at | 34.2 | A | 31.2 | P | 59.4 | P | 31 | A | --- |
| 263090_at | 16.3 | A | 1.5 | A | 11.8 | A | 16 | A | --- |
| 263091_at | 1.8 | A | 1.1 | A | 0.6 | A | 0.7 | A | --- |
| 263092_at | 20.7 | A | 2.7 | A | 15.9 | A | 17.4 | A | --- |
| 263093_at | 3.5 | A | 2.5 | A | 1.8 | A | 9.7 | A | --- |
| 263094_at | 19.4 | A | 2.4 | A | 33.5 | A | 22.6 | A | --- |
| 263095_at | 0.4 | A | 0.6 | A | 1.6 | A | 1.1 | A | --- |
| 263096_at | 36.9 | A | 89.3 | P | 10.5 | A | 56.7 | P | AHB1 |
| 263097_at | 647.7 | P | 644.4 | P | 600.4 | P | 730.3 | P | PDV2 |
| 263098_at | 4.5 | A | 36.7 | A | 24 | A | 78.5 | P | --- |
| 263099_at | 5.4 | A | 13.1 | A | 14.6 | A | 29.1 | A | --- |

| | | | | | | | | | |
|---|---|---|---|---|---|---|---|---|---|
| 263100_at | 196.6 | P | 191.9 | P | 188.7 | P | 202.2 | P | AtPOT1a |
| 263101_s_at | 532.7 | P | 528.7 | P | 436.2 | P | 538.7 | P | --- |
| 263102_at | 16.7 | A | 6.2 | A | 6.3 | A | 21.8 | A | --- |
| 263103_at | 2.3 | A | 2.2 | A | 13.5 | A | 3.1 | A | --- |
| 263104_at | 112.2 | P | 105.5 | P | 144 | P | 118.6 | P | --- |
| 263105_at | 231.4 | P | 218.1 | P | 217.1 | P | 185.9 | P | --- |
| 263106_at | 10.2 | A | 39.4 | A | 10.9 | A | 27.8 | A | --- |
| 263107_at | 25.2 | A | 14.8 | A | 19.1 | A | 0.4 | A | CYP705A6 |
| 263108_at | 128.5 | P | 132.8 | P | 110.9 | A | 80.6 | A | --- |
| 263109_at | 22.3 | A | 35.1 | A | 10.3 | A | 16.8 | A | --- |
| 263110_at | 896.5 | P | 955 | P | 1033.9 | P | 891.2 | P | --- |
| 263111_s_at | 1480.6 | P | 1206.4 | P | 2270.6 | P | 1579.8 | P | --- |
| 263112_at | 382.4 | P | 281.8 | P | 228.7 | P | 274.4 | P | --- |
| 263113_at | 408.3 | P | 446.7 | P | 467.3 | P | 464.2 | P | --- |
| 263114_at | 2504.5 | P | 5017.5 | P | 2140.8 | P | 3622.9 | P | PSAD-2 |
| 263115_at | 50.5 | A | 126.3 | P | 121.3 | P | 115.5 | P | --- |
| 263116_s_at | 899 | P | 884.2 | P | 841.2 | P | 715.8 | P | --- |
| 263117_at | 703.7 | P | 711.7 | P | 630.5 | P | 693.8 | P | --- |
| 263118_at | 533.1 | P | 375.6 | P | 470 | P | 384.7 | P | MCCA |
| 263119_at | 257.7 | P | 219.7 | P | 321.2 | P | 211.4 | P | --- |
| 263120_at | 546.3 | P | 924.6 | P | 804.5 | P | 940.3 | P | CYP708A3 |
| 263121_at | 53.6 | P | 33.8 | A | 49.9 | P | 62.4 | P | --- |
| 263122_at | 593.7 | P | 569.3 | P | 725.6 | P | 656.4 | P | SPS1 |
| 263123_at | 1.1 | A | 11.2 | A | 28 | A | 1.7 | A | --- |
| 263124_at | 45.5 | P | 65.4 | P | 67.5 | P | 45.7 | P | --- |
| 263125_at | 3.4 | A | 7.4 | A | 5 | A | 4.3 | A | --- |
| 263126_at | 1066.6 | P | 1545.7 | P | 840.2 | P | 1434.1 | P | --- |
| 263127_at | 372.2 | P | 250.5 | P | 333.4 | P | 289.3 | P | MSL6 |
| 263128_at | 510.2 | P | 913.8 | P | 591.2 | P | 835 | P | LZF1 |
| 263129_at | 859 | P | 1031.6 | P | 1003.7 | P | 1006.3 | P | --- |
| 263130_at | 162.3 | P | 205.3 | P | 179.5 | P | 161.1 | P | POLD3 |
| 263131_at | 5761.3 | P | 6365.3 | P | 5510.6 | P | 6579.6 | P | emb1473 |
| 263132_at | 824.4 | P | 808.9 | P | 743.2 | P | 725 | P | --- |
| 263133_at | 20.2 | A | 64.1 | A | 65.5 | A | 40.1 | A | --- |
| 263134_at | 1548.7 | P | 2263.1 | P | 2363.6 | P | 2926.9 | P | RHM1 |
| 263135_at | 155.8 | P | 190 | P | 118.1 | P | 164.3 | P | --- |
| 263136_at | 196.3 | P | 314.4 | P | 396.6 | P | 251.7 | P | ATTPS1 |
| 263137_at | 727.6 | P | 539.1 | P | 650.7 | P | 581.6 | P | ATGGH1 |
| 263138_at | 37.2 | A | 52.5 | A | 28.6 | A | 53.5 | A | --- |
| 263139_at | 11 | A | 5.4 | A | 5.9 | A | 2.7 | A | --- |
| 263140_at | 22.6 | A | 23.8 | P | 7.8 | A | 17.5 | P | --- |
| 263141_at | 20.7 | A | 28.9 | P | 29.8 | P | 25.1 | P | --- |
| 263142_at | 1702.4 | P | 2077.8 | P | 2018.6 | P | 1904 | P | --- |
| 263143_at | 122.9 | P | 80.8 | P | 116.6 | P | 127.8 | P | --- |
| 263144_at | 4.2 | A | 5.6 | A | 41.5 | A | 2.1 | A | --- |
| 263145_at | 363.5 | P | 391.2 | P | 419.3 | P | 369.2 | P | ATEXO70D2 |
| 263146_at | 5 | A | 1.8 | A | 9.7 | A | 1.5 | A | GLIP2 |
| 263147_at | 27.2 | P | 1.4 | A | 10 | A | 22 | A | --- |
| 263148_at | 4.2 | A | 4.2 | A | 1.3 | A | 1.1 | A | --- |
| 263149_at | 21.4 | P | 13.8 | P | 40.9 | A | 5.9 | A | --- |
| 263150_at | 345.9 | P | 246.8 | P | 298 | P | 326.5 | P | --- |
| 263151_at | 85.8 | P | 146.7 | P | 105.6 | A | 129.6 | P | --- |
| 263152_at | 295.8 | P | 373.1 | P | 281 | P | 330 | P | ASIL1 |
| 263153_s_at | 4932.2 | P | 1725 | P | 5483.1 | P | 2185.6 | P | --- |
| 263154_at | 206.1 | P | 183.3 | P | 240.3 | P | 192.2 | P | --- |
| 263155_at | 315.1 | P | 350.7 | P | 276.2 | P | 278.8 | P | TAFII21 |
| 263156_at | 2140.8 | P | 1153.8 | P | 1480.4 | P | 1307.6 | P | --- |
| 263157_at | 6219.9 | P | 3125.6 | P | 4395.8 | P | 3389.6 | P | ALDH7B4 |
| 263158_at | 243.1 | P | 307.2 | P | 258 | P | 292 | P | NF-YA5 |
| 263159_at | 1261.3 | P | 1767.3 | P | 1029.1 | P | 1661.3 | P | RSH3 |
| 263160_at | 3122.7 | P | 2582.4 | P | 2931.5 | P | 2858.1 | P | UBP1A |
| 263161_at | 1654.3 | P | 146.5 | P | 1017.4 | P | 343.5 | P | --- |
| 263162_at | 424.6 | P | 408.7 | P | 434 | P | 382.2 | P | --- |
| 263163_at | 532.6 | P | 608.9 | P | 536.5 | P | 607.1 | P | FZL |
| 263164_at | 69.1 | A | 84 | P | 70.2 | P | 64.1 | P | --- |
| 263165_at | 187 | P | 170.7 | P | 207.3 | P | 187.2 | P | --- |
| 263166_at | 3 | A | 3.7 | A | 4 | A | 1.9 | A | --- |
| 263167_at | 531.2 | P | 385.9 | P | 404.3 | P | 403.5 | P | --- |
| 263168_at | 5.5 | A | 1.6 | A | 40.6 | P | 19.5 | A | --- |
| 263169_at | 28.3 | M | 40.9 | P | 13.9 | A | 25.9 | P | --- |
| 263170_at | 449 | P | 473.5 | P | 497.9 | P | 569.4 | P | PEX6 |
| 263171_at | 826.8 | P | 760.4 | P | 668.6 | P | 687.5 | P | CCX4 |
| 263172_at | 311.9 | P | 382.1 | P | 391.2 | P | 385 | P | CID3 |
| 263173_at | 50 | P | 68 | P | 48 | A | 45.2 | A | BRX-LIKE3 |
| 263174_at | 462.6 | P | 550.9 | P | 349.2 | P | 719.8 | P | ESP |
| 263175_at | 31.3 | A | 3.6 | A | 7.4 | A | 6 | A | --- |
| 263176_at | 24.4 | A | 17.1 | A | 58.1 | A | 15.5 | A | UGT75B2 |
| 263177_at | 67.2 | A | 73.1 | P | 108.1 | A | 107.3 | P | --- |
| 263178_at | 43.4 | A | 30.1 | A | 72 | A | 5.2 | A | --- |
| 263179_at | 396 | P | 276.9 | P | 345.1 | P | 237.3 | P | --- |
| 263180_at | 840.4 | P | 499.8 | P | 799.6 | P | 626.9 | P | URH2 |
| 263181_at | 758.3 | P | 920.9 | P | 894.1 | P | 963.9 | P | --- |
| 263182_at | 245 | P | 116 | P | 200.1 | P | 115 | P | --- |
| 263183_at | 415.4 | P | 578 | P | 390.5 | P | 469.2 | P | CALS1 |
| 263184_at | 3642.3 | P | 2352.4 | P | 3653.1 | P | 3211.3 | P | UGT75B1 |
| 263185_at | 621.8 | P | 601.9 | P | 650.3 | P | 680.5 | P | --- |
| 263186_at | 17 | A | 15.9 | A | 10.5 | A | 31.5 | P | --- |
| 263187_at | 20.2 | A | 27.8 | A | 35.4 | A | 40.2 | A | --- |
| 263188_at | 21.6 | A | 1 | A | 1.6 | A | 17.8 | A | --- |
| 263189_at | 2.1 | A | 0.9 | A | 1.7 | A | 2.8 | A | --- |
| 263190_at | 11.8 | A | 11.3 | A | 1.1 | A | 12.1 | A | --- |
| 263191_at | 9.1 | A | 0.7 | A | 1.2 | A | 15.2 | A | --- |
| 263192_at | 667.8 | P | 525.2 | P | 695.9 | P | 609.3 | P | ACC1 |
| 263193_at | 621.9 | P | 664 | P | 597.1 | P | 569.6 | P | --- |
| 263194_at | 41.6 | A | 59.2 | A | 14.8 | A | 92.7 | A | --- |
| 263195_at | 5.8 | A | 5.7 | A | 17.1 | A | 9.1 | A | --- |
| 263196_at | 270.7 | P | 317.7 | P | 220.7 | P | 349.8 | P | --- |
| 263197_at | 1.5 | A | 1.7 | A | 2.8 | A | 1.2 | A | --- |
| 263198_at | 4.5 | A | 5.5 | A | 24.4 | A | 9.3 | A | GLIP3 |
| 263199_at | 116.6 | P | 155.3 | P | 135 | P | 144.9 | P | HEXO2 |
| 263200_at | 278.6 | P | 172.4 | P | 204 | P | 219.2 | P | --- |
| 263201_at | 22.2 | A | 40.2 | A | 22.1 | A | 26.1 | A | APS2 |
| 263202_at | 96.2 | P | 120.4 | P | 102.8 | P | 115.1 | P | 5PTASE13 |
| 263203_at | 2.2 | A | 1.1 | A | 2.7 | A | 14.2 | A | chr31 |
| 263204_at | 32.4 | A | 3.4 | A | 28.6 | A | 11.8 | A | chr31 |
| 263205_at | 482.1 | P | 506.3 | P | 427.1 | P | 495.7 | P | NTMC2T2.1 |
| 263206_at | 3521.5 | P | 2809.6 | P | 3013.3 | P | 2938.5 | P | --- |
| 263207_at | 46.2 | A | 44.5 | A | 60.8 | A | 52.1 | P | XTH33 |
| 263208_at | 38.9 | A | 31.3 | A | 18.3 | A | 29.6 | M | ZFP5 |

| | | | | | | | | | |
|---|---|---|---|---|---|---|---|---|---|
| 263209_at | 1007.8 | P | 1368.4 | P | 1028.5 | P | 1326.6 | P | --- |
| 263210_at | 1043.8 | P | 159.3 | P | 373.5 | P | 311.4 | P | --- |
| 263211_at | 20.4 | A | 11.2 | A | 7.1 | A | 7.3 | A | GLP7 |
| 263212_at | 747.1 | P | 803 | P | 733.5 | P | 856.1 | P | emb2004 |
| 263213_at | 28.7 | A | 16.7 | A | 26.4 | A | 20.7 | A | --- |
| 263214_at | 53.2 | P | 72.4 | P | 87.3 | P | 77.4 | P | --- |
| 263215_at | 9.3 | A | 2.4 | A | 5.2 | A | 9.2 | A | --- |
| 263216_s_at | 66.8 | A | 40.7 | M | 2.5 | A | 35.9 | A | --- |
| 263217_at | 1.3 | A | 0.8 | A | 3.6 | A | 2.2 | A | --- |
| 263218_at | 37.2 | A | 89.5 | P | 54.2 | A | 81.2 | P | --- |
| 263219_at | 126.4 | P | 232.7 | P | 158.5 | P | 267.6 | P | --- |
| 263220_at | 381.7 | P | 457.5 | P | 437.5 | P | 418.6 | P | emb2279 |
| 263221_at | 504.8 | P | 345.6 | P | 465.8 | P | 419.7 | P | MUR4 |
| 263222_at | 172.6 | P | 174.1 | P | 112.6 | P | 147.1 | P | --- |
| 263223_at | 1911 | P | 2296.9 | P | 1566.7 | P | 2238.2 | P | --- |
| 263224_at | 2832.8 | P | 2479 | P | 3435.7 | P | 2437.6 | P | --- |
| 263225_at | 46 | A | 52.7 | P | 61.6 | A | 64.7 | P | WRKY14 |
| 263226_at | 415.1 | P | 469.8 | P | 354.2 | P | 468.4 | P | --- |
| 263227_at | 32.4 | A | 36 | A | 39.8 | A | 21.5 | A | --- |
| 263228_at | 2.1 | A | 1.7 | A | 18 | A | 2.1 | A | --- |
| 263229_s_at | 27 | A | 0.4 | A | 37.3 | P | 0.6 | A | --- |
| 263230_at | 47.3 | P | 65.5 | P | 60.9 | M | 59 | P | --- |
| 263231_at | 3764.5 | P | 1124.5 | P | 2993.8 | P | 1953.4 | P | --- |
| 263232_at | 32.8 | A | 26.8 | A | 5.9 | A | 35.9 | A | --- |
| 263233_at | 3.4 | A | 11 | A | 2.1 | A | 2.1 | A | --- |
| 263234_at | 9.6 | A | 3.2 | A | 1.4 | A | 3.2 | A | --- |
| 263235_at | 402.4 | P | 445.7 | P | 341.3 | P | 492.7 | P | PP2A-2 |
| 263236_at | 2065.2 | P | 1684.9 | P | 1719.1 | P | 1625.5 | P | ARR4 |
| 263237_at | 182.5 | P | 94.2 | P | 138.8 | P | 122 | P | --- |
| 263238_at | 36.9 | P | 33.9 | P | 44.6 | P | 62.1 | A | --- |
| 263239_at | 104.7 | P | 63.2 | P | 92.4 | P | 63.9 | P | ATASE1 |
| 263240_s_at | 29.5 | A | 39.5 | A | 7.8 | A | 43.9 | A | --- |
| 263241_at | 1060.2 | P | 1484.9 | P | 1363.7 | P | 1506.5 | P | ADC1 |
| 263242_at | 2529 | P | 2896.9 | P | 2945.2 | P | 2978.2 | P | --- |
| 263243_at | 810 | P | 696 | P | 771.4 | P | 756.7 | P | GLX2-5 |
| 263244_at | 16.1 | A | 17 | A | 1.7 | A | 19.8 | A | --- |
| 263245_at | 10 | A | 2.6 | A | 3.7 | A | 9.2 | A | --- |
| 263246_at | 5.8 | A | 1.4 | A | 28.5 | A | 2.4 | A | --- |
| 263247_at | 465.7 | P | 579 | P | 620.9 | P | 480.1 | P | --- |
| 263248_at | 251.8 | P | 251.7 | P | 231.7 | P | 196.8 | P | ATNTH1 |
| 263249_at | 3270.9 | P | 3916.7 | P | 3323.9 | P | 4093.2 | P | ADS2 |
| 263250_at | 1889.4 | P | 1415.6 | P | 1694.4 | P | 1492.9 | P | --- |
| 263251_at | 1076.2 | P | 928.3 | P | 1018.8 | P | 1080.9 | P | --- |
| 263252_at | 192.5 | P | 528.2 | P | 313.3 | P | 445.1 | P | STH |
| 263253_at | 251.2 | P | 299.7 | P | 265.1 | P | 296.6 | P | --- |
| 263254_x_at | 64.4 | M | 86.4 | A | 80.1 | M | 84 | A | --- |
| 263255_at | 218.4 | P | 225.6 | P | 273.2 | P | 243.6 | P | --- |
| 263256_at | 1650.2 | P | 1917.1 | P | 1725.8 | P | 1928.3 | P | ATCPISCA |
| 263257_at | 5.2 | A | 6.9 | A | 8.1 | A | 7.2 | A | --- |
| 263258_at | 124.1 | P | 70.7 | P | 130 | P | 82.7 | P | --- |
| 263259_at | 19.6 | A | 9.1 | A | 19.5 | A | 28.3 | A | PUB18 |
| 263260_at | 403.1 | P | 277.5 | P | 457.2 | P | 356.7 | P | OTS2 |
| 263261_at | 606 | P | 496.1 | P | 549.4 | P | 577.7 | P | --- |
| 263262_at | 8.1 | A | 3.2 | A | 3.4 | A | 23.3 | A | --- |
| 263263_at | 123.4 | P | 170.5 | P | 120.5 | P | 199.5 | P | --- |
| 263264_at | 207.3 | P | 224.6 | P | 202.5 | P | 221.8 | P | HTA8 |
| 263265_at | 246.1 | P | 320.7 | P | 203.8 | P | 208.3 | P | --- |
| 263266_at | 0.7 | A | 1.6 | A | 3.5 | A | 0.4 | A | --- |
| 263267_at | 3507.5 | P | 3688.2 | P | 4561 | P | 4352.6 | P | --- |
| 263268_at | 59.3 | P | 79.1 | M | 95 | P | 60.7 | A | --- |
| 263269_at | 0.5 | A | 0.5 | A | 4.1 | A | 5.4 | A | --- |
| 263270_at | 1.2 | A | 0.5 | A | 1.7 | A | 1.4 | A | --- |
| 263271_s_at | 5.2 | A | 0.9 | A | 27.4 | A | 2.4 | A | --- |
| 263272_at | 1.1 | A | 3.6 | A | 4.9 | A | 0.9 | A | --- |
| 263273_x_at | 4.9 | A | 10.9 | A | 15.9 | P | 1.5 | A | --- |
| 263274_at | 737.9 | P | 696.7 | P | 471.4 | P | 559.8 | P | CRCK3 |
| 263275_at | 2110.9 | P | 2240.3 | P | 1789.3 | P | 2117.2 | P | ALDH6B2 |
| 263276_at | 17.1 | A | 26 | A | 44 | A | 8.8 | A | CYP705A13 |
| 263277_at | 200.6 | P | 214.9 | P | 183.7 | P | 221.5 | P | --- |
| 263278_at | 477.8 | P | 522.4 | P | 443 | P | 446 | P | DRP3B |
| 263279_x_at | 7.6 | A | 3.6 | A | 34.1 | A | 6.2 | A | --- |
| 263280_x_at | 2.2 | A | 1.1 | A | 10.1 | A | 0.6 | A | --- |
| 263281_at | 23.2 | A | 4.9 | A | 17.3 | A | 25.9 | A | --- |
| 263282_at | 3.5 | A | 12.7 | A | 49.3 | A | 5.8 | A | --- |
| 263283_at | 38.7 | A | 40.7 | A | 43.5 | A | 58.4 | A | --- |
| 263284_at | 28.2 | A | 62.7 | P | 66.2 | A | 47.7 | A | --- |
| 263285_at | 627 | P | 564.8 | P | 1014.1 | P | 591 | P | DOT1 |
| 263286_at | 5915.9 | P | 5570.6 | P | 5650.1 | P | 5276.5 | P | --- |
| 263287_at | 1817.2 | P | 1707.4 | P | 1641 | P | 1779 | P | --- |
| 263288_at | 541.3 | P | 624.7 | P | 601.4 | P | 614.5 | P | --- |
| 263289_at | 3956.9 | P | 3277.3 | P | 4501.6 | P | 3691.8 | P | --- |
| 263290_at | 46.1 | P | 39.2 | P | 107 | P | 57.1 | P | --- |
| 263291_at | 21.9 | A | 10.6 | A | 35.6 | A | 26.6 | A | --- |
| 263292_at | 6.9 | A | 11.6 | A | 11.6 | A | 14.2 | A | --- |
| 263293_x_at | 7 | A | 4.5 | A | 39.5 | A | 4.9 | A | --- |
| 263294_at | 11.6 | A | 17.3 | A | 56.9 | P | 22.7 | A | --- |
| 263295_at | 11.1 | A | 16.9 | A | 1.2 | A | 1.7 | A | AGL44 |
| 263296_at | 1211.6 | P | 954.5 | P | 955.6 | P | 931.5 | P | --- |
| 263297_at | 107.7 | P | 122.6 | P | 76.8 | P | 150.3 | P | ATARFB1A |
| 263298_at | 1615.6 | P | 2193.3 | P | 1944.5 | P | 2144.8 | P | TIC21 |
| 263299_s_at | 23.2 | A | 1.1 | A | 23.5 | A | 1.8 | A | --- |
| 263300_at | 13.5 | A | 14.3 | A | 7.5 | A | 22.5 | A | --- |
| 263301_x_at | 6.7 | A | 6.1 | A | 1.3 | A | 0.6 | A | --- |
| 263302_at | 9.6 | A | 0.1 | A | 20.1 | A | 4.3 | A | --- |
| 263303_at | 511.1 | P | 654.3 | P | 587.5 | P | 497.9 | P | --- |
| 263304_at | 0.5 | A | 0.4 | A | 3.5 | A | 5.5 | A | --- |
| 263305_at | 94.8 | P | 140.7 | P | 143.5 | P | 116.6 | P | BPC1 |
| 263306_at | 26.1 | A | 35 | P | 43.1 | P | 56 | P | SCPL43 |
| 263307_at | 1.1 | A | 0.6 | A | 3.5 | A | 1 | A | --- |
| 263308_at | 14.2 | A | 1.8 | A | 12.3 | A | 17.6 | A | --- |
| 263309_at | 7.6 | A | 10.3 | A | 8.4 | A | 10.4 | A | --- |
| 263310_s_at | 1.2 | A | 22.9 | A | 15.1 | A | 7.2 | A | --- |
| 263311_at | 47.8 | A | 4.7 | A | 30.1 | A | 12.6 | A | --- |
| 263312_at | 0.1 | A | 0.4 | A | 0.9 | A | 0.6 | A | --- |
| 263313_at | 4.5 | A | 7.4 | A | 19.8 | A | 19.1 | A | --- |
| 263314_at | 86.7 | P | 132.4 | P | 166.9 | P | 140.9 | P | --- |
| 263315_at | 135.7 | P | 197.2 | P | 139.5 | P | 174.9 | P | --- |
| 263316_s_at | 1.6 | A | 1.2 | A | 5.4 | A | 1.5 | A | ATGLR2.3 |
| 263317_s_at | 30.2 | A | 17.8 | A | 30.4 | A | 8.2 | A | SUVH7 |

| | | | | | | | | | |
|---|---|---|---|---|---|---|---|---|---|
| 263318_at | 161.5 | P | 306 | P | 362 | P | 285.3 | P | AtGDU4 |
| 263319_at | 142.6 | P | 271 | P | 203.9 | P | 277.8 | P | BOR1 |
| 263320_at | 381.5 | P | 152.4 | P | 342.9 | P | 128.1 | P | AtGolS1 |
| 263321_at | 4821.5 | P | 6016.2 | P | 5815.5 | P | 6086.1 | P | ARF1A1C |
| 263322_at | 190.9 | P | 224 | P | 280.8 | P | 263.8 | P | RNEE/G |
| 263323_at | 20.7 | A | 8.5 | A | 22 | A | 24.4 | A | --- |
| 263324_at | 5.6 | A | 5.1 | A | 8.9 | A | 4.2 | A | --- |
| 263325_at | 639.7 | P | 665.6 | P | 658.6 | P | 575.5 | P | XERICO |
| 263326_at | 1672.9 | P | 1882 | P | 1547.8 | P | 1831.3 | P | --- |
| 263327_at | 30.5 | A | 17.8 | A | 63.9 | A | 30.8 | A | --- |
| 263328_at | 179.4 | P | 274.4 | P | 246.8 | P | 325.8 | P | --- |
| 263329_at | 43.1 | A | 6.7 | A | 51.9 | A | 32.3 | A | --- |
| 263330_at | 331.4 | P | 321.2 | P | 275.2 | P | 319.3 | P | --- |
| 263331_at | 371 | P | 413.1 | P | 459.8 | P | 415.9 | P | --- |
| 263332_at | 1248.5 | P | 1211.3 | P | 1133.6 | P | 1278.6 | P | --- |
| 263333_at | 796 | P | 945.5 | P | 785.1 | P | 916.2 | P | --- |
| 263334_at | 1606.5 | P | 963 | P | 1508.1 | P | 1033.2 | P | --- |
| 263335_x_at | 14.4 | A | 12.2 | A | 7.2 | A | 0.9 | A | --- |
| 263336_x_at | 27.2 | A | 28.5 | M | 30 | A | 32.6 | A | --- |
| 263337_at | 2.5 | A | 1.4 | A | 24.2 | A | 14.9 | A | --- |
| 263338_at | 7 | A | 13 | A | 9 | A | 2.9 | A | --- |
| 263339_at | 13.5 | A | 3.9 | A | 4.2 | A | 0.8 | A | --- |
| 263340_at | 1.6 | A | 2.6 | A | 3.6 | A | 1.5 | A | --- |
| 263341_at | 12 | A | 26.6 | A | 26 | A | 26.1 | A | --- |
| 263342_at | 10.9 | A | 5.1 | A | 1.7 | A | 8.2 | A | --- |
| 263343_at | 0.7 | A | 0.4 | A | 1.5 | A | 0.7 | A | --- |
| 263344_at | 519.6 | P | 412.5 | P | 521 | P | 499.4 | P | --- |
| 263345_s_at | 7689.5 | P | 13556.3 | P | 8590.3 | P | 11846.7 | P | LHCB2.1 |
| 263346_at | 6.6 | A | 9.2 | A | 8.6 | A | 3.1 | A | --- |
| 263347_at | 55.2 | M | 58.3 | A | 86.6 | A | 49.1 | A | --- |
| 263348_at | 6615.3 | P | 5075.1 | P | 6279 | P | 5574.1 | P | --- |
| 263349_at | 254.9 | P | 286.6 | P | 243.2 | P | 273.9 | P | CHR5 |
| 263350_at | 11380.1 | P | 9788.4 | P | 12006.9 | P | 10417.4 | P | AGT |
| 263351_at | 255.2 | P | 271.8 | P | 193.2 | P | 281.1 | P | --- |
| 263352_at | 260.7 | P | 255.4 | P | 225.2 | P | 281.5 | P | --- |
| 263353_at | 6.4 | A | 20.9 | A | 33.4 | A | 46.8 | A | --- |
| 263354_at | 1 | A | 17.7 | A | 10.1 | A | 11.6 | A | --- |
| 263355_at | 220.8 | P | 236.9 | P | 257.1 | P | 207.9 | P | --- |
| 263356_at | 321.8 | P | 417.1 | P | 323.4 | P | 389.3 | P | UBA1A |
| 263357_s_at | 26.1 | P | 26 | P | 9.6 | A | 1.2 | A | --- |
| 263358_x_at | 35.2 | A | 16.3 | A | 13.5 | A | 5.1 | A | --- |
| 263359_at | 336.6 | P | 445.5 | P | 447.7 | P | 386.8 | P | ATLIP1 |
| 263360_at | 5.3 | A | 1.5 | A | 10 | A | 22.8 | A | --- |
| 263361_at | 14.1 | A | 14.6 | P | 26 | A | 17.8 | A | TET13 |
| 263362_at | 4.2 | A | 7.5 | A | 6.3 | A | 20.5 | A | --- |
| 263363_at | 14 | A | 2.5 | A | 4.9 | A | 1.1 | A | --- |
| 263364_at | 33.6 | A | 24.9 | A | 40.8 | A | 51.7 | A | --- |
| 263365_at | 56.1 | A | 104.4 | P | 88.7 | A | 65.4 | P | --- |
| 263366_at | 94.8 | A | 102.9 | P | 108.7 | M | 155.3 | P | --- |
| 263367_at | 27.2 | A | 30.9 | A | 56.1 | A | 25.1 | A | --- |
| 263368_at | 1.6 | A | 6.7 | A | 2.7 | A | 3.7 | A | --- |
| 263369_at | 45.2 | P | 52.4 | P | 80.9 | P | 54.1 | P | --- |
| 263370_at | 44.4 | P | 94.7 | P | 80.7 | P | 76.5 | P | --- |
| 263371_at | 1676.8 | P | 1313.7 | P | 2099.8 | P | 1474.3 | P | NOP10 |
| 263372_at | 1966.5 | P | 1695.9 | P | 2473.6 | P | 1815.5 | P | --- |
| 263373_at | 47.9 | M | 45.9 | P | 24.1 | A | 51.2 | P | --- |
| 263374_at | 166.7 | P | 189 | P | 151.4 | P | 162 | P | --- |
| 263375_s_at | 3447.7 | P | 2539.8 | P | 3342.2 | P | 2559.3 | P | ATPHB6 |
| 263376_at | 4.1 | A | 8.6 | A | 32.8 | A | 24.2 | A | FLA6 |
| 263377_at | 6.5 | A | 4.2 | A | 50.6 | A | 62.2 | A | ABI4 |
| 263378_at | 2 | A | 4.7 | A | 3 | A | 13.6 | A | ATHPP2C5 |
| 263379_at | 545.7 | P | 551.3 | P | 469.3 | P | 469.5 | P | CZF1 |
| 263380_at | 70.4 | P | 48 | P | 68.8 | P | 63.2 | P | --- |
| 263381_at | 11.7 | A | 7.1 | A | 9.7 | A | 13.9 | A | AGL48 |
| 263382_at | 110.4 | P | 126.9 | P | 161.5 | P | 111.3 | P | --- |
| 263383_at | 90.9 | P | 85.8 | P | 78.4 | P | 91.9 | P | --- |
| 263384_at | 7.4 | A | 5.6 | A | 9.9 | A | 6.9 | A | --- |
| 263385_at | 1.1 | A | 1.6 | A | 3 | A | 4.5 | A | GEA6 |
| 263386_at | 57.5 | P | 127.5 | P | 99.9 | P | 123.6 | P | --- |
| 263387_at | 27.8 | P | 53 | P | 63.4 | P | 61.5 | P | --- |
| 263388_at | 2.3 | A | 7.5 | A | 2.3 | A | 9.6 | A | --- |
| 263389_at | 2.5 | A | 1.6 | A | 19.8 | A | 1.6 | A | --- |
| 263390_at | 6 | A | 6.9 | A | 8.7 | A | 1.4 | A | --- |
| 263391_at | 152 | P | 178.6 | P | 95.7 | P | 176.6 | P | MGDC |
| 263392_at | 19.1 | A | 5 | A | 7.9 | P | 0.1 | A | --- |
| 263393_at | 4.2 | A | 23.2 | P | 1.2 | A | 13.9 | A | --- |
| 263394_at | 5.8 | A | 5.5 | A | 13.7 | A | 4.5 | A | --- |
| 263395_at | 9.6 | A | 3.4 | A | 3 | A | 7.3 | A | ATTIM44-1 |
| 263396_at | 390.1 | P | 442.5 | P | 374.2 | P | 529.4 | P | --- |
| 263397_s_at | 12.5 | A | 12.9 | A | 37.2 | A | 2.1 | A | --- |
| 263398_at | 272 | P | 302.2 | P | 277.1 | P | 300.4 | P | AtRABA5d |
| 263399_at | 2043.5 | P | 2026.1 | P | 1795.2 | P | 1927.6 | P | --- |
| 263400_s_at | 5932.4 | P | 5219.4 | P | 5096.3 | P | 5063.5 | P | --- |
| 263401_at | 120.5 | P | 40.6 | P | 79.5 | P | 41.8 | P | --- |
| 263402_at | 1478.8 | P | 88 | P | 946.6 | P | 163.8 | P | --- |
| 263403_at | 2149.2 | P | 673.7 | P | 1827.5 | P | 926.2 | P | TX1 |
| 263404_s_at | 188.6 | P | 128.6 | P | 219.8 | P | 117.1 | P | --- |
| 263405_s_at | 631 | P | 321.5 | P | 431.5 | P | 325.6 | P | --- |
| 263406_at | 3012.5 | P | 1314.3 | P | 2970.2 | P | 1844.1 | P | AIR3 |
| 263407_at | 2.6 | A | 2.2 | A | 1.6 | A | 15 | A | AtbZIP48 |
| 263408_at | 54.3 | A | 21.7 | A | 102.2 | A | 39.9 | A | --- |
| 263409_at | 8.4 | A | 16.5 | A | 4.5 | A | 1.7 | A | --- |
| 263410_at | 885.3 | P | 1531 | P | 1229.2 | P | 1357.6 | P | --- |
| 263411_at | 23.4 | A | 53.1 | A | 69.9 | A | 61.9 | P | --- |
| 263412_at | 332.5 | P | 353.1 | P | 268.9 | P | 379.5 | P | --- |
| 263413_at | 741 | P | 869.2 | P | 700.7 | P | 772.6 | P | BPC4 |
| 263414_at | 314.1 | P | 476.8 | P | 360.2 | P | 441.2 | P | --- |
| 263415_at | 586 | P | 542.5 | P | 768 | P | 611.1 | P | EMB2762 |
| 263416_at | 21.7 | A | 11.1 | A | 21.5 | A | 15.4 | A | --- |
| 263417_at | 2.7 | A | 1.7 | A | 3.6 | A | 2.8 | A | --- |
| 263418_at | 1.2 | A | 4.3 | A | 4.1 | A | 3.6 | A | --- |
| 263419_at | 468.7 | P | 723.3 | P | 485.8 | P | 638.2 | P | --- |
| 263420_at | 1941.7 | P | 2242.4 | P | 2066.1 | P | 2302.3 | P | --- |
| 263421_at | 814.9 | P | 1037.7 | P | 1012.8 | P | 1016.8 | P | EXL5 |
| 263422_s_at | 878.2 | P | 1129.2 | P | 978.7 | P | 892.8 | P | --- |
| 263423_at | 2.1 | A | 2.2 | A | 2 | A | 2.5 | A | --- |
| 263424_at | 1075.5 | P | 1080 | P | 856.3 | P | 1010.2 | P | --- |
| 263425_at | 386.9 | P | 476.9 | P | 485.7 | P | 431.3 | P | EMB2289 |
| 263426_at | 4653.9 | P | 4409.2 | P | 4780.1 | P | 4974.8 | P | ATGPX2 |

| | | | | | | | | | |
|---|---|---|---|---|---|---|---|---|---|
| 263427_at | 162.6 | P | 174.8 | P | 197.2 | P | 163 | P | --- |
| 263428_at | 170.7 | P | 233 | P | 199.7 | P | 223 | P | ATUBP4 |
| 263429_at | 334.6 | P | 353.2 | P | 348.4 | P | 330.8 | P | AAT |
| 263430_at | 134.4 | P | 120.9 | P | 155 | P | 134.8 | P | --- |
| 263431_at | 2274.5 | P | 3092 | P | 2574.1 | P | 3021.7 | P | --- |
| 263432_at | 1789.8 | P | 1677.9 | P | 2155 | P | 1733 | P | --- |
| 263433_at | 408 | P | 618.5 | P | 387.1 | P | 720.5 | P | MIPS2 |
| 263434_at | 10.5 | A | 1.9 | A | 30.8 | A | 1 | A | PRS |
| 263435_at | 600.8 | P | 508.7 | P | 781.2 | P | 491.7 | P | --- |
| 263436_at | 6.6 | A | 6.7 | A | 12.7 | A | 41.5 | A | --- |
| 263437_at | 69.6 | A | 94.9 | P | 100.6 | P | 79.3 | P | --- |
| 263438_at | 294.2 | P | 248.3 | P | 425.7 | P | 251.2 | P | --- |
| 263439_at | 53.4 | P | 35.5 | P | 77.5 | P | 60.2 | P | ATEXO70H8 |
| 263440_at | 26.8 | A | 34.8 | A | 26.1 | A | 17.4 | A | ATEXO70H5 |
| 263441_at | 96.5 | P | 128.2 | P | 56.4 | A | 127.7 | P | --- |
| 263442_at | 546.4 | P | 845.8 | P | 593.8 | P | 755.5 | P | --- |
| 263443_at | 65 | P | 97.6 | P | 67.5 | P | 90.4 | P | KCS12 |
| 263444_at | 3.3 | A | 3.3 | A | 3.1 | A | 0.9 | A | AGL46 |
| 263445_at | 5.3 | A | 3.2 | A | 6.2 | A | 4 | A | --- |
| 263446_at | 153.3 | P | 106.2 | P | 125 | P | 178.8 | P | ATX1 |
| 263447_s_at | 193 | P | 247.4 | P | 223.5 | P | 197.2 | P | ATX2 |
| 263448_at | 511.6 | P | 415.9 | P | 567.2 | P | 424.3 | P | SAD2 |
| 263449_at | 3207.4 | P | 3210.7 | P | 3368.7 | P | 3098.9 | P | AtRABA5d |
| 263450_at | 2.8 | A | 4.5 | A | 3.5 | A | 2.3 | A | CPK24 |
| 263451_at | 3.2 | A | 1.6 | A | 12.6 | A | 2.2 | A | --- |
| 263452_at | 939.1 | P | 403.8 | P | 816.3 | P | 468 | P | --- |
| 263453_at | 3 | A | 17.7 | A | 26.7 | A | 9.3 | A | --- |
| 263454_at | 3.2 | A | 1.1 | A | 1.2 | A | 1.3 | A | --- |
| 263455_at | 9.4 | A | 3.5 | A | 1.9 | A | 3 | A | --- |
| 263456_at | 731 | P | 940.5 | P | 802.7 | P | 1017.7 | P | --- |
| 263457_at | 363.9 | P | 447.9 | P | 393.3 | P | 340.1 | P | SR1 |
| 263458_at | 2.9 | A | 4.5 | A | 22.7 | A | 1.4 | A | ATRABH1D |
| 263459_at | 788.3 | P | 739.8 | P | 708.1 | P | 684.7 | P | --- |
| 263460_at | 1273.1 | P | 1162 | P | 1140.9 | P | 1037.9 | P | --- |
| 263461_at | 579.3 | P | 810.6 | P | 602.4 | P | 793.6 | P | --- |
| 263462_at | 19.9 | A | 22.4 | A | 10.5 | A | 12.6 | A | ARI11 |
| 263463_at | 1.7 | A | 5.7 | A | 0.9 | A | 17.3 | A | --- |
| 263464_at | 0.9 | A | 1.3 | A | 0.6 | A | 0.5 | A | --- |
| 263465_at | 42.2 | P | 56.8 | P | 52 | P | 52.2 | P | --- |
| 263466_at | 528 | P | 546 | P | 587.8 | P | 611 | P | --- |
| 263467_at | 331.3 | P | 470.2 | P | 405.2 | P | 364.2 | P | --- |
| 263468_at | 1.3 | A | 7.5 | A | 6.7 | A | 1.1 | A | --- |
| 263469_at | 0.8 | A | 3.6 | A | 1.2 | A | 9.6 | A | ATCHX21 |
| 263470_at | 33.4 | A | 15.5 | A | 38 | A | 36.6 | A | XIF |
| 263471_at | 999 | P | 1073.3 | P | 1240.7 | P | 1222.1 | P | RAP |
| 263472_at | 679 | P | 634.4 | P | 759.3 | P | 632.8 | P | CNX2 |
| 263473_at | 1850.2 | P | 1635.7 | P | 2206.7 | P | 1881.7 | P | UGT74D1 |
| 263474_at | 120 | P | 78.6 | P | 119 | P | 96.8 | P | --- |
| 263475_at | 77.6 | P | 46.7 | M | 66.8 | A | 40.7 | A | --- |
| 263476_at | 101.5 | P | 125.4 | P | 101.4 | P | 110.9 | P | TEJ |
| 263477_at | 2538.5 | P | 1815.1 | P | 3193.9 | P | 2046.9 | P | --- |
| 263478_at | 2137.2 | P | 2764.4 | P | 1144.3 | P | 1830.5 | P | --- |
| 263479_x_at | 3.4 | A | 3.6 | A | 16.9 | A | 2.5 | A | --- |
| 263480_at | 97.7 | P | 205.3 | P | 58.4 | A | 177.2 | P | ZIP7 |
| 263481_at | 1.2 | A | 3.2 | A | 16.9 | A | 6.3 | A | --- |
| 263482_at | 2260 | P | 1072.1 | P | 1905.8 | P | 1215.9 | P | --- |
| 263483_at | 5626.7 | P | 4974.8 | P | 6924.2 | P | 5297.2 | P | CR88 |
| 263484_at | 343.1 | P | 420.3 | P | 287.7 | P | 381.3 | P | --- |
| 263485_at | 81.4 | P | 130.5 | P | 105.3 | P | 120 | P | VLN1 |
| 263486_at | 20.8 | A | 9.3 | A | 33.4 | A | 4.4 | A | --- |
| 263487_at | 2.1 | A | 5.3 | A | 4.6 | A | 8.1 | A | --- |
| 263488_at | 689.4 | P | 873.3 | P | 666.9 | P | 833.8 | P | --- |
| 263489_at | 41.5 | A | 53 | A | 34.2 | A | 25.8 | A | --- |
| 263490_at | 201.9 | P | 253.7 | P | 198.5 | P | 196.4 | P | MAX2 |
| 263491_at | 814 | P | 1565 | P | 1116 | P | 1581.4 | P | ATPPC2 |
| 263492_at | 2 | A | 4.1 | A | 4.6 | A | 3.2 | A | --- |
| 263493_at | 915.2 | P | 851.5 | P | 932.1 | P | 878.8 | P | --- |
| 263494_at | 3079.9 | P | 4451 | P | 3295 | P | 3903.3 | P | GRF9 |
| 263495_at | 1876.5 | P | 2675.8 | P | 1595.2 | P | 4307 | P | COR15B |
| 263496_at | 282.8 | P | 220.2 | P | 151.5 | P | 241.8 | P | --- |
| 263497_at | 3682.4 | P | 4375 | P | 3150.1 | P | 8273.3 | P | COR15A |
| 263498_at | 127.4 | P | 835.8 | P | 215 | P | 889.9 | P | LSH10 |
| 263499_at | 534.2 | P | 600 | P | 547.8 | P | 641.3 | P | TTL3 |
| 263500_s_at | 4.2 | A | 3.1 | A | 2.7 | A | 2.1 | A | --- |
| 263501_s_at | 15.7 | A | 3.9 | A | 24 | A | 9.3 | A | --- |
| 263502_s_at | 61.7 | P | 47.5 | P | 21.1 | A | 43 | P | --- |
| 263503_s_at | 41.9 | A | 35.3 | A | 32 | A | 13.7 | A | --- |
| 263504_s_at | 2.9 | A | 6 | A | 2.8 | A | 2.9 | A | --- |
| 263505_s_at | 39.5 | A | 24.6 | A | 22.5 | A | 16.3 | A | --- |
| 263506_s_at | 3.2 | A | 4.4 | A | 6 | A | 6.7 | A | ArthMp070 |
| 263507_s_at | 16.2 | A | 22.8 | A | 39.3 | A | 16.8 | A | ArthMp069 |
| 263508_s_at | 28 | A | 4.7 | A | 42 | A | 23.7 | A | ArthMp068 |
| 263509_s_at | 147 | P | 95.7 | P | 106.5 | P | 94.9 | P | --- |
| 263510_s_at | 46.6 | A | 32.4 | A | 32.4 | A | 5.7 | A | --- |
| 263511_at | 0.6 | A | 10.9 | A | 25.2 | A | 14.1 | A | --- |
| 263512_at | 3.5 | A | 8.8 | A | 21.3 | A | 10.1 | A | --- |
| 263513_at | 153.6 | P | 365 | P | 186.9 | P | 320.8 | P | --- |
| 263514_at | 3.3 | A | 3.1 | A | 2.6 | A | 2.3 | A | --- |
| 263515_at | 1030.7 | P | 301.3 | P | 1039.2 | P | 321.3 | P | --- |
| 263516_at | 1105.5 | P | 1400 | P | 1064.7 | P | 1399.1 | P | ATRER1B |
| 263517_at | 4854.6 | P | 2817.9 | P | 3217.4 | P | 3368.2 | P | RD2 |
| 263518_at | 1.7 | A | 3.8 | A | 13.3 | A | 11.1 | A | --- |
| 263519_at | 2835.4 | P | 2150.6 | P | 2936.3 | P | 2757.1 | P | --- |
| 263520_at | 28.1 | A | 67.7 | P | 62.9 | P | 50 | P | --- |
| 263521_at | 154.5 | P | 158 | P | 153.7 | P | 122.5 | P | --- |
| 263522_at | 12.1 | A | 5.4 | A | 11.5 | A | 2.1 | A | --- |
| 263523_at | 17.3 | A | 15.7 | A | 7.3 | A | 7.3 | A | --- |
| 263524_x_at | 2.2 | A | 4.1 | A | 15.9 | A | 12.4 | A | --- |
| 263525_at | 12.6 | A | 64.5 | A | 86.8 | A | 32.2 | A | AGL61 |
| 263526_at | 252.9 | P | 304.3 | P | 333.1 | P | 254.9 | P | --- |
| 263527_at | 2.2 | A | 2.5 | A | 5.7 | A | 6.9 | A | --- |
| 263528_at | 41.2 | P | 41.4 | P | 64.2 | M | 76.5 | P | --- |
| 263529_at | 1440.9 | P | 1509.4 | P | 1486.4 | P | 1545.1 | P | ARF3 |
| 263530_at | 21.7 | A | 3.8 | A | 3.1 | A | 3 | A | --- |
| 263531_s_at | 5.8 | A | 21.6 | A | 2.3 | A | 1.5 | A | --- |
| 263532_s_at | 638.9 | P | 522.9 | P | 606.9 | P | 571.6 | P | --- |
| 263533_at | 2827.9 | P | 2513.9 | P | 2924.2 | P | 2491.6 | P | TIC55 |
| 263534_at | 3268.1 | P | 3956.6 | P | 3066 | P | 3508 | P | AtMAPR2 |
| 263535_at | 114.5 | P | 164.1 | P | 66 | P | 124.2 | P | --- |

| | | | | | | | | | |
|---|---|---|---|---|---|---|---|---|---|
| 263536_at | 417.1 | P | 326.7 | P | 240.8 | P | 214.6 | P | WRKY60 |
| 263537_at | 1558.5 | P | 1875.6 | P | 1670.1 | P | 1792.7 | P | COL3 |
| 263538_at | 305.5 | P | 258 | P | 314.7 | P | 269.4 | P | --- |
| 263539_at | 2384 | P | 910.3 | P | 1766.2 | P | 1373.9 | P | TAT3 |
| 263540_at | 11.8 | A | 1.8 | A | 3.9 | A | 10.9 | A | --- |
| 263541_at | 1645.5 | P | 1253.1 | P | 1279.6 | P | 1272.4 | P | --- |
| 263542_at | 116 | P | 163.8 | P | 140.1 | P | 127.3 | P | --- |
| 263543_at | 9.5 | A | 9.3 | A | 14.2 | A | 10.8 | A | PE11 |
| 263544_at | 116.8 | P | 260.4 | P | 185.8 | P | 276.8 | P | APL4 |
| 263545_at | 57 | P | 76.8 | P | 59.4 | P | 85.8 | P | --- |
| 263546_at | 46.3 | A | 58.3 | A | 59.3 | A | 49.8 | P | --- |
| 263547_at | 11.3 | A | 19.8 | A | 3.6 | A | 6.1 | A | --- |
| 263548_at | 20928.3 | P | 19576.1 | P | 21911.7 | P | 21280.1 | P | CCR2 |
| 263549_at | 403.3 | P | 680.2 | P | 738.2 | P | 832.8 | P | MEE3 |
| 263550_at | 716.2 | P | 890.3 | P | 906.9 | P | 773.5 | P | --- |
| 263551_at | 109 | P | 122.8 | P | 110.9 | P | 156.8 | P | --- |
| 263552_x_at | 45.9 | A | 60.1 | A | 75.2 | A | 49.4 | A | --- |
| 263553_at | 784.5 | P | 1566.2 | P | 1046.1 | P | 1417.1 | P | PAP10 |
| 263554_at | 15.5 | A | 3.2 | A | 27.6 | A | 20.8 | A | --- |
| 263555_at | 248.7 | P | 371.8 | P | 290.7 | P | 333.3 | P | --- |
| 263556_at | 673.8 | P | 706 | P | 571.2 | P | 762.7 | P | --- |
| 263557_at | 553.7 | P | 643.4 | P | 529.4 | P | 588.9 | P | BLH7 |
| 263558_at | 157.9 | P | 103.8 | P | 190.5 | P | 140.9 | P | --- |
| 263559_at | 19.8 | A | 23.1 | A | 50.5 | P | 24.8 | M | --- |
| 263560_s_at | 19.3 | A | 39.9 | A | 38.8 | P | 32.7 | P | FUT5 |
| 263561_at | 2 | A | 0.7 | A | 2.2 | A | 5 | A | --- |
| 263562_at | 3.1 | A | 2.9 | A | 7.1 | A | 1.7 | A | --- |
| 263563_at | 23.2 | A | 22.5 | A | 37.7 | A | 20.9 | A | --- |
| 263564_s_at | 681.1 | P | 656.7 | P | 617.2 | P | 731.3 | P | NRPB3 |
| 263565_at | 67.1 | P | 103.8 | P | 34.6 | P | 53.1 | P | FUT4 |
| 263566_at | 28.3 | A | 49.5 | A | 50.7 | A | 33.2 | A | --- |
| 263567_at | 107.8 | P | 80.3 | P | 130.9 | P | 107 | P | --- |
| 263568_at | 16.6 | A | 12.7 | A | 32.2 | A | 9.8 | A | --- |
| 263569_at | 138.8 | P | 172.6 | P | 151.8 | P | 143.1 | P | TTN7 |
| 263570_at | 403 | P | 419.3 | P | 277.2 | P | 368.7 | P | AAO3 |
| 263571_at | 0.7 | A | 1.1 | A | 1 | A | 1.1 | A | --- |
| 263572_at | 13.5 | A | 20.3 | A | 4.2 | A | 2 | A | --- |
| 263573_at | 109.6 | P | 136.2 | P | 122.9 | P | 101.8 | P | --- |
| 263574_at | 145.1 | P | 125.6 | P | 149.9 | P | 150.9 | P | --- |
| 263575_at | 9.7 | A | 6.8 | A | 15.3 | A | 11.5 | A | --- |
| 263576_at | 1.4 | A | 7 | A | 2 | A | 2.4 | A | --- |
| 263577_at | 3.2 | A | 4.9 | A | 5.1 | A | 4 | A | SSP |
| 263578_at | 213 | P | 228.9 | P | 260.6 | P | 255.8 | P | --- |
| 263579_at | 180.2 | P | 183.9 | P | 152.1 | P | 186.6 | P | --- |
| 263580_at | 33.1 | A | 36.7 | P | 62.1 | P | 39.8 | P | --- |
| 263581_at | 3.4 | A | 8.5 | A | 2.3 | A | 1.3 | A | --- |
| 263582_at | 991.1 | P | 1195.5 | P | 675.6 | P | 953.4 | P | LYM2 |
| 263583_at | 608.9 | P | 707.6 | P | 522.8 | P | 722.7 | P | IDH2 |
| 263584_at | 593.5 | P | 769.9 | P | 430.7 | P | 551 | P | anac036 |
| 263585_at | 2390 | P | 2034.2 | P | 2182.6 | P | 2261.4 | P | --- |
| 263586_at | 93.8 | P | 118.1 | P | 78 | P | 134.4 | P | --- |
| 263587_at | 21.6 | P | 36 | P | 63.5 | P | 46.9 | P | ATVAMP712 |
| 263588_s_at | 74.5 | P | 47.7 | P | 69.4 | P | 52.3 | P | --- |
| 263589_at | 800.9 | P | 744.5 | P | 711.3 | P | 742.6 | P | --- |
| 263590_at | 1198.9 | P | 1261.2 | P | 1154.4 | P | 1169.9 | P | --- |
| 263591_at | 211.8 | P | 366.2 | P | 340.9 | P | 323.9 | P | ATMAP65-6 |
| 263592_at | 11.7 | A | 32 | A | 57.6 | P | 34.2 | P | --- |
| 263593_at | 192.7 | P | 127.4 | P | 213.3 | P | 155.3 | P | EMB975 |
| 263594_at | 12.1 | A | 22.6 | A | 3.3 | A | 18.5 | A | PAP7 |
| 263595_at | 202.7 | P | 199.1 | P | 257.6 | P | 221.6 | P | PAP8 |
| 263596_at | 0.8 | A | 21.2 | A | 4 | A | 22.2 | A | --- |
| 263597_at | 504.8 | P | 661.9 | P | 519.5 | P | 565 | P | --- |
| 263598_at | 4285.1 | P | 2369 | P | 4451.4 | P | 2906.8 | P | EXGT-A3 |
| 263599_at | 220 | P | 192.5 | P | 248.1 | P | 265.5 | P | WOL |
| 263600_at | 95.4 | P | 101.7 | P | 108.3 | A | 91.6 | P | DRD1 |
| 263601_s_at | 369.3 | P | 437.4 | P | 431.3 | P | 431.3 | P | THY-1 |
| 263602_at | 52.3 | A | 39.7 | A | 42.1 | A | 61 | A | --- |
| 263603_at | 71.8 | P | 59.3 | P | 127.3 | P | 87.9 | M | --- |
| 263604_s_at | 1.5 | A | 2.1 | A | 21.4 | A | 6 | A | --- |
| 263605_at | 85.6 | P | 58.3 | P | 84.9 | P | 89.7 | P | --- |
| 263606_at | 1763.1 | P | 1509.3 | P | 1895 | P | 1561 | P | KCS9 |
| 263607_at | 75.7 | P | 96.6 | P | 73 | P | 148.3 | P | --- |
| 263608_at | 97.1 | A | 106.6 | P | 153.1 | P | 99.6 | P | --- |
| 263609_at | 86.1 | P | 126.2 | P | 82.8 | P | 133.9 | P | --- |
| 263610_at | 1 | A | 1.3 | A | 6.2 | A | 4.1 | A | --- |
| 263611_at | 5.4 | A | 1.6 | A | 4 | A | 7.8 | A | --- |
| 263612_at | 186.5 | P | 271.9 | P | 256.1 | P | 303.7 | P | MCM4 |
| 263613_at | 333.2 | P | 370.5 | P | 331 | P | 332.4 | P | --- |
| 263614_at | 24.4 | A | 1 | A | 52.6 | P | 25.9 | A | --- |
| 263615_at | 1.1 | A | 0.8 | A | 4.9 | A | 4.7 | A | MYB100 |
| 263616_at | 10.2 | A | 12.7 | A | 12.9 | A | 26.7 | A | --- |
| 263617_at | 6 | A | 6.2 | A | 1.3 | A | 1.4 | A | --- |
| 263618_at | 162 | P | 152.5 | P | 198.2 | P | 199.7 | P | APC2 |
| 263619_at | 73.8 | P | 141.5 | P | 99.9 | P | 129.4 | P | --- |
| 263620_at | 2.9 | A | 2.8 | A | 5.6 | A | 11.4 | A | --- |
| 263621_at | 1285.9 | P | 1068.7 | P | 862 | P | 933.9 | P | NRPB6B |
| 263622_at | 1.8 | A | 0.9 | A | 6.7 | A | 3 | A | --- |
| 263623_at | 20.2 | A | 6.8 | A | 50.2 | A | 18.4 | A | --- |
| 263624_at | 4453.6 | P | 5153.1 | P | 4457.7 | P | 5263.4 | P | --- |
| 263625_at | 24 | A | 30.8 | P | 55.8 | P | 26.1 | A | --- |
| 263626_at | 582.3 | P | 625.1 | P | 565.2 | P | 667.3 | P | ZAP1 |
| 263627_at | 0.9 | A | 14.4 | A | 31.2 | A | 1.9 | A | --- |
| 263628_at | 663.1 | P | 1260.2 | P | 812.3 | P | 1278.2 | P | FLA7 |
| 263629_at | 128.1 | P | 124.9 | P | 148.5 | P | 160.3 | P | --- |
| 263630_at | 327.5 | P | 377.5 | P | 301.6 | P | 375.6 | P | --- |
| 263631_at | 2340.5 | P | 2246.8 | P | 1505.4 | P | 1996.2 | P | --- |
| 263632_at | 405.2 | P | 480.6 | P | 673.7 | P | 344.8 | P | --- |
| 263633_at | 189.2 | P | 193.8 | P | 265.1 | P | 252.5 | P | SCL21 |
| 263634_at | 0.7 | A | 0.9 | A | 2.8 | A | 1 | A | --- |
| 263635_at | 12 | A | 10.5 | A | 4.1 | A | 2.7 | A | --- |
| 263636_at | 1926.5 | P | 1959.4 | P | 1665.8 | P | 1996 | P | --- |
| 263637_at | 526.1 | P | 529.6 | P | 608.6 | P | 595.8 | P | --- |
| 263638_at | 564.5 | P | 497.9 | P | 484.8 | P | 428.4 | P | --- |
| 263639_at | 138.1 | P | 215 | P | 221.2 | P | 156.2 | P | --- |
| 263640_at | 278.1 | P | 381.6 | P | 369.4 | P | 385 | P | --- |
| 263641_at | 1.2 | A | 1.1 | A | 4 | A | 2.1 | A | --- |
| 263642_at | 29.6 | P | 27 | P | 67.4 | P | 33.5 | P | --- |
| 263643_at | 194.1 | P | 215.1 | P | 264 | P | 247.8 | P | --- |
| 263644_at | 2.7 | A | 4.4 | A | 28.4 | A | 2.9 | A | --- |

| | | | | | | | | | |
|---|---|---|---|---|---|---|---|---|---|
| 263645_at | 0.6 | A | 6.3 | A | 2.8 | A | 15.1 | A | --- |
| 263646_at | 2.1 | A | 0.8 | A | 2.3 | A | 0.6 | A | --- |
| 263647_at | 430.6 | P | 538.2 | P | 401.5 | P | 419 | P | --- |
| 263648_at | 39.4 | A | 23.8 | M | 81.6 | P | 79.3 | M | --- |
| 263649_at | 33.5 | A | 28 | A | 26.9 | A | 37.3 | A | --- |
| 263650_at | 44.3 | A | 50.7 | A | 75.9 | A | 17.7 | A | --- |
| 263651_at | 549.4 | P | 632.5 | P | 669.5 | P | 615.9 | P | --- |
| 263652_at | 1.7 | A | 9.6 | A | 13.5 | A | 3.7 | A | --- |
| 263653_at | 132.7 | P | 206.2 | P | 112.4 | P | 141.7 | P | ERS2 |
| 263654_at | 674.5 | P | 569.1 | P | 513.3 | P | 497.5 | P | --- |
| 263655_at | 22.4 | A | 41 | A | 36.7 | A | 23 | A | --- |
| 263656_at | 78.2 | P | 179.7 | P | 158.5 | P | 241.4 | P | SHY2 |
| 263657_at | 590.8 | P | 574.5 | P | 498.1 | P | 477.9 | P | CKL13 |
| 263658_at | 18.6 | A | 17.5 | A | 14.2 | A | 15.2 | A | --- |
| 263659_at | 3.6 | A | 3.8 | A | 1.7 | A | 1.3 | A | AT1G04470 |
| 263660_at | 625.4 | P | 584.5 | P | 554 | P | 483.2 | P | MPI7 |
| 263661_at | 647.9 | P | 657.4 | P | 596.6 | P | 687.9 | P | --- |
| 263662_at | 2329.3 | P | 2322.1 | P | 1434.1 | P | 1940.2 | P | --- |
| 263663_at | 11403.5 | P | 10686.6 | P | 12678.3 | P | 11042.2 | P | --- |
| 263664_at | 459.9 | P | 515.2 | P | 577.4 | P | 517.7 | P | AXR3 |
| 263665_at | 5838.2 | P | 4883.3 | P | 6630.7 | P | 5768.9 | P | --- |
| 263666_at | 798 | P | 918.3 | P | 889.3 | P | 862.9 | P | --- |
| 263667_at | 8143 | P | 7673.8 | P | 9762 | P | 8110.1 | P | RPS15 |
| 263668_at | 1223.9 | P | 1311.5 | P | 1169.7 | P | 1293.2 | P | --- |
| 263669_at | 1990.7 | P | 2238.5 | P | 1674 | P | 2173.5 | P | CRY2 |
| 263670_at | 21.2 | A | 2.6 | A | 35.6 | A | 18.7 | A | --- |
| 263671_at | 5.1 | A | 0.2 | A | 1.2 | A | 1.1 | A | --- |
| 263672_at | 7 | A | 17 | A | 26.7 | A | 14.4 | A | --- |
| 263673_at | 2.4 | A | 14 | A | 4.4 | A | 39.2 | A | --- |
| 263674_at | 285 | P | 269.9 | P | 340.4 | P | 256.4 | P | --- |
| 263675_x_at | 40.3 | A | 27.3 | A | 32 | A | 33.7 | A | --- |
| 263676_at | 7404.2 | P | 8090.8 | P | 9397.2 | P | 7349.1 | P | CRB |
| 263677_at | 164.7 | P | 229.3 | P | 268.5 | P | 308.9 | P | PDLP2 |
| 263678_at | 1792.7 | P | 2063.8 | P | 1667.4 | P | 2125.6 | P | --- |
| 263679_at | 1105.6 | P | 1129.1 | P | 1076 | P | 1069.6 | P | --- |
| 263680_at | 581.5 | P | 131.2 | P | 178 | P | 142.7 | P | --- |
| 263681_at | 104 | P | 122.1 | P | 153.7 | P | 143.8 | P | ORC6 |
| 263682_at | 1 | A | 0.6 | A | 1.8 | A | 11.5 | A | AT5G49590 |
| 263683_at | 22.4 | A | 25.4 | A | 44.9 | A | 40.4 | M | FEZ |
| 263684_at | 135 | P | 118.8 | P | 112.7 | P | 92.5 | P | --- |
| 263685_at | 778.4 | P | 623.9 | P | 635.9 | P | 672.8 | P | ATCUL3 |
| 263686_at | 990.3 | P | 754.4 | P | 1188.7 | P | 805.6 | P | --- |
| 263687_at | 168.7 | P | 175.2 | P | 190.2 | P | 171.5 | P | --- |
| 263688_at | 519.6 | P | 542.9 | P | 472 | P | 433.8 | P | --- |
| 263689_at | 99.3 | P | 182 | P | 146.3 | P | 208.9 | P | RNS3 |
| 263690_at | 88.6 | P | 57.9 | P | 94.7 | P | 64.6 | P | AtHB23 |
| 263691_at | 5673.6 | P | 4734.2 | P | 5815.3 | P | 5319.6 | P | --- |
| 263692_at | 3655.7 | P | 3885.4 | P | 3650.2 | P | 4001.1 | P | --- |
| 263693_at | 52 | P | 54.7 | P | 80.2 | P | 71.1 | P | ATPP2-A9 |
| 263694_at | 1.2 | A | 18.9 | A | 2.9 | A | 17.5 | A | --- |
| 263695_at | 278.7 | P | 212.8 | P | 311.2 | P | 235.1 | P | --- |
| 263696_at | 975.8 | P | 928.6 | P | 1263.9 | P | 861.8 | P | AK-HSDH I |
| 263697_at | 3.1 | A | 4.3 | A | 22.6 | M | 3.6 | A | --- |
| 263698_at | 48.8 | A | 36.4 | A | 52.1 | A | 47.3 | A | --- |
| 263699_at | 234.6 | P | 134.3 | P | 151.6 | P | 161.3 | P | KUP10 |
| 263700_at | 64.7 | P | 84.8 | P | 118.9 | P | 84.9 | P | --- |
| 263701_at | 532.1 | P | 619.8 | P | 578.5 | P | 532.7 | P | --- |
| 263702_at | 65.4 | P | 40.5 | P | 112 | P | 70.7 | A | --- |
| 263703_at | 1377.9 | P | 1434.6 | P | 1351.5 | P | 1328.1 | P | SRX |
| 263704_at | 1438.6 | P | 1461.6 | P | 1331.6 | P | 1086.8 | P | --- |
| 263705_at | 1025.8 | P | 983.9 | P | 780.2 | P | 1068.9 | P | IMPL1 |
| 263706_s_at | 5075.7 | P | 3885 | P | 6619.1 | P | 3399.1 | P | --- |
| 263707_at | 282.1 | P | 255.2 | P | 373.6 | P | 299.6 | P | --- |
| 263708_at | 66.3 | P | 91.3 | P | 97 | P | 86.9 | P | --- |
| 263709_at | 13157.6 | P | 13856.4 | P | 13848.3 | P | 15052 | P | --- |
| 263710_at | 1149.5 | P | 1211.3 | P | 1452.5 | P | 1324.4 | P | --- |
| 263711_at | 1636.4 | P | 1947.9 | P | 1430.4 | P | 1770.1 | P | --- |
| 263712_at | 600.6 | P | 448.5 | P | 476.4 | P | 576.5 | P | NFD6 |
| 263713_at | 29.9 | A | 23 | A | 44.1 | A | 9 | A | --- |
| 263714_at | 3434.3 | P | 2193.8 | P | 3691.4 | P | 2336.2 | P | SUR1 |
| 263715_at | 428 | P | 938.6 | P | 551.6 | P | 879.3 | P | GPRI1 |
| 263716_at | 1064.7 | P | 1186.2 | P | 975.8 | P | 1047.6 | P | RPN1A |
| 263717_at | 2065.8 | P | 2532.6 | P | 1894.5 | P | 2377.4 | P | --- |
| 263718_at | 19.3 | A | 16.1 | A | 6.1 | A | 44.9 | A | NF-YB7 |
| 263719_at | 103.9 | P | 77.2 | P | 87.5 | P | 91.1 | P | --- |
| 263720_at | 3.3 | A | 5 | A | 2 | A | 2 | A | ATCHX15 |
| 263721_at | 7.5 | A | 2.7 | A | 13.9 | A | 1.4 | A | --- |
| 263722_at | 250.6 | P | 261.6 | P | 268.2 | P | 235.3 | P | GONST1 |
| 263723_at | 9.6 | A | 2 | A | 3.1 | A | 0.6 | A | --- |
| 263724_at | 92.7 | M | 77.5 | P | 102.4 | P | 56.5 | P | --- |
| 263725_at | 19 | A | 18.5 | A | 49.9 | A | 24.8 | A | --- |
| 263726_at | 236.2 | P | 435.8 | P | 327.6 | P | 435.5 | P | --- |
| 263727_at | 527.9 | P | 442.1 | P | 548.7 | P | 460.7 | P | ABH1 |
| 263728_at | 944.6 | P | 1148.3 | P | 1158.6 | P | 1202.7 | P | --- |
| 263729_at | 298.4 | P | 195.8 | P | 275.3 | P | 258.2 | P | --- |
| 263730_at | 50.7 | A | 26.4 | A | 69.3 | A | 32.2 | A | BGLU4 |
| 263731_at | 337.7 | P | 243.4 | P | 255 | P | 242.1 | P | --- |
| 263732_at | 11.7 | A | 100.7 | A | 38.4 | A | 72.2 | A | ARL2 |
| 263733_at | 1.5 | A | 1 | A | 2.7 | A | 22.4 | A | --- |
| 263734_at | 110.1 | P | 79 | P | 94.6 | P | 74.2 | P | NAT7 |
| 263735_s_at | 37.2 | A | 48.8 | P | 52.4 | A | 8.2 | A | AGL50 |
| 263736_at | 3037.7 | P | 3396.8 | P | 3297.9 | P | 3321.3 | P | --- |
| 263737_at | 2249.2 | P | 1884.6 | P | 1946.3 | P | 2006 | P | --- |
| 263738_at | 93.2 | A | 100.6 | P | 102.5 | A | 93.3 | A | --- |
| 263739_at | 210.7 | P | 691.5 | P | 249.2 | P | 575.8 | P | --- |
| 263740_at | 3 | A | 1.4 | A | 4 | A | 16.2 | A | RALFL14 |
| 263741_at | 28.9 | A | 18 | A | 7.2 | A | 34.1 | A | --- |
| 263742_at | 7.5 | A | 12.7 | A | 18.9 | A | 8.2 | A | --- |
| 263743_at | 1740.1 | P | 1823.2 | P | 1496.1 | P | 2426.9 | P | --- |
| 263744_at | 23.8 | A | 19.7 | A | 46.6 | A | 6 | A | --- |
| 263745_at | 3.9 | A | 35.5 | A | 45.8 | P | 20.6 | A | CHR34 |
| 263746_at | 2 | A | 7.7 | A | 3.7 | A | 17.9 | A | --- |
| 263747_at | 63.5 | P | 160.5 | P | 133.3 | P | 105 | P | SAE2 |
| 263748_at | 13.4 | A | 19 | A | 16.9 | A | 4.1 | A | --- |
| 263749_at | 993.7 | P | 1058.5 | P | 1007.2 | P | 1021.5 | P | --- |
| 263750_at | 1033.9 | P | 1100.8 | P | 784.8 | P | 1043 | P | --- |
| 263751_at | 129.2 | P | 292.1 | P | 215.2 | P | 287.1 | P | --- |
| 263752_at | 3.5 | A | 0.4 | A | 2 | A | 1.8 | A | --- |
| 263753_at | 8.1 | A | 2.3 | A | 3 | A | 1.1 | A | LEA |

| | | | | | | | | | |
|---|---|---|---|---|---|---|---|---|---|
| 263754_at | 67.4 | P | 80.5 | P | 46.7 | A | 54 | P | --- |
| 263755_at | 1096.5 | P | 1210.8 | P | 1150.6 | P | 1179.4 | P | --- |
| 263756_at | 798.5 | P | 753.6 | P | 622.4 | P | 662.4 | P | --- |
| 263757_at | 203.6 | P | 191.8 | P | 237.1 | P | 183.8 | P | --- |
| 263758_s_at | 3073.6 | P | 3070.8 | P | 2544.6 | P | 3088.7 | P | --- |
| 263759_at | 1554.5 | P | 1281.3 | P | 1460.7 | P | 1326.6 | P | --- |
| 263760_at | 1275.7 | P | 1577.2 | P | 1370 | P | 1655.8 | P | SULA |
| 263761_at | 6831.1 | P | 9403.5 | P | 8293 | P | 9116.4 | P | --- |
| 263762_at | 473.7 | P | 531 | P | 372 | P | 444.6 | P | --- |
| 263763_at | 922.2 | P | 889.5 | P | 977.4 | P | 894.8 | P | --- |
| 263764_at | 818.2 | P | 788.3 | P | 785.8 | P | 698.5 | P | VHA-A2 |
| 263765_at | 165.3 | P | 186.9 | P | 156.1 | P | 187.8 | P | SFH3 |
| 263766_at | 349.1 | P | 368.6 | P | 441 | P | 339.9 | P | --- |
| 263767_at | 0.3 | A | 9.3 | A | 1.8 | A | 1.1 | A | --- |
| 263768_x_at | 8.9 | A | 17.4 | A | 8.4 | A | 14.3 | A | --- |
| 263769_at | 4.2 | A | 1.7 | A | 2.7 | A | 1.4 | A | --- |
| 263770_at | 27.5 | A | 15.9 | A | 4.5 | A | 27.9 | A | --- |
| 263771_at | 16.9 | A | 33.1 | A | 26.2 | A | 28.4 | A | --- |
| 263772_at | 104.9 | P | 43.9 | P | 105.5 | P | 45.3 | P | ELF4-L3 |
| 263773_at | 403.4 | P | 370.2 | P | 301.8 | P | 368.7 | P | --- |
| 263774_at | 721.8 | P | 678.9 | P | 684.6 | P | 593.4 | P | --- |
| 263775_at | 53.8 | P | 43.7 | P | 43 | P | 47.4 | P | CPC |
| 263776_s_at | 347.4 | P | 577.1 | P | 284.8 | P | 432.9 | P | ATCNGC3 |
| 263777_at | 777.9 | P | 733 | P | 894.2 | P | 795.6 | P | ATCNGC12 |
| 263778_at | 400.5 | P | 353.5 | P | 377.7 | P | 325.2 | P | OXA1L |
| 263779_at | 988.6 | P | 1406.9 | P | 1182.8 | P | 1535.7 | P | SPA1 |
| 263780_at | 149.6 | P | 245.4 | P | 211.2 | P | 314.1 | P | SPA1 |
| 263781_at | 25.9 | A | 9 | A | 21.1 | A | 18.5 | M | --- |
| 263782_at | 97.4 | P | 109.3 | P | 80.6 | P | 108.6 | P | --- |
| 263783_at | 551.1 | P | 587.8 | P | 429.7 | P | 435 | P | WRKY46 |
| 263784_at | 1.9 | A | 1.7 | A | 3.5 | A | 2.7 | A | --- |
| 263785_at | 1286.7 | P | 1288.1 | P | 1422.2 | P | 1383.8 | P | --- |
| 263786_at | 1041.4 | P | 1008.2 | P | 858.6 | P | 917.8 | P | JAR1 |
| 263787_at | 600.6 | P | 860.2 | P | 835.4 | P | 960.9 | P | --- |
| 263788_at | 374.6 | P | 260.8 | P | 372 | P | 306.7 | P | --- |
| 263789_at | 24.1 | A | 28.1 | A | 69.2 | A | 7.1 | A | --- |
| 263790_at | 102.5 | P | 146.1 | P | 136.2 | P | 120.9 | P | --- |
| 263791_at | 242.6 | P | 209.2 | P | 246.2 | P | 195 | P | AHA5 |
| 263792_at | 11.1 | A | 44.8 | A | 77.5 | A | 49 | A | --- |
| 263793_at | 143.5 | M | 184.3 | P | 183 | P | 173.5 | P | ATCSLC08 |
| 263794_at | 1.4 | A | 1.6 | A | 14.5 | A | 1.2 | A | --- |
| 263795_at | 15.5 | A | 34.5 | P | 17.7 | A | 1.8 | A | ATCNGC14 |
| 263796_at | 266.5 | P | 266.7 | P | 287.5 | P | 301 | P | AFR |
| 263797_at | 122.2 | P | 168.4 | P | 143.8 | P | 202.5 | P | WRKY17 |
| 263798_at | 77 | P | 96.4 | P | 171.9 | P | 103.5 | P | UBP19 |
| 263799_at | 250.9 | P | 226.1 | P | 235.1 | P | 175.5 | P | --- |
| 263800_at | 914.4 | P | 669.1 | P | 704.9 | P | 633.2 | P | --- |
| 263801_at | 4.8 | A | 11.4 | A | 27.8 | A | 1.6 | A | --- |
| 263802_at | 829.5 | P | 692.6 | P | 825.6 | P | 777.3 | P | --- |
| 263803_at | 16 | A | 25.3 | A | 11.9 | A | 32.6 | A | --- |
| 263804_at | 430.2 | P | 611.2 | P | 323.4 | P | 456.9 | P | --- |
| 263805_at | 647.5 | P | 691.7 | P | 525.8 | P | 624.2 | P | --- |
| 263806_at | 291.9 | P | 281.7 | P | 261.6 | P | 249.6 | P | --- |
| 263807_at | 3897.4 | P | 2672.3 | P | 2966.2 | P | 2282.4 | P | --- |
| 263808_at | 309.3 | P | 331.2 | P | 457.5 | P | 324.7 | P | --- |
| 263809_at | 253.6 | P | 384.7 | P | 320.9 | P | 389.9 | P | --- |
| 263810_at | 3155.4 | P | 3001.5 | P | 2014.3 | P | 3105.8 | P | --- |
| 263811_at | 1052.9 | P | 1047.2 | P | 924.9 | P | 960.4 | P | --- |
| 263812_at | 5 | A | 5.3 | A | 11.8 | A | 11.7 | A | --- |
| 263813_s_at | 4.2 | A | 1.5 | A | 4.9 | A | 2.9 | A | --- |
| 263814_at | 0.3 | A | 1 | A | 22.2 | A | 6.4 | A | --- |
| 263815_at | 30.5 | A | 21.3 | A | 32.9 | A | 6.3 | A | --- |
| 263816_at | 0.9 | A | 3.9 | A | 5.1 | A | 12.4 | A | --- |
| 263817_at | 41.2 | P | 55.1 | P | 36.9 | A | 41.7 | M | --- |
| 263818_at | 2.5 | A | 3.1 | A | 5 | A | 1.6 | A | --- |
| 263819_x_at | 24.5 | A | 31.2 | P | 7.7 | A | 33.3 | M | --- |
| 263820_at | 3.7 | A | 6.3 | A | 11.4 | A | 4.7 | A | --- |
| 263821_s_at | 15964.6 | P | 13934.2 | P | 19262.2 | P | 16222.9 | P | --- |
| 263822_at | 134 | P | 53.6 | P | 122.9 | P | 134.8 | P | --- |
| 263823_s_at | 301 | P | 185.2 | P | 127.2 | P | 128 | P | --- |
| 263824_at | 1886.1 | P | 1527.3 | P | 1985.1 | P | 1551.7 | P | --- |
| 263825_at | 1.4 | A | 38.7 | A | 33.2 | A | 39.8 | A | LAC5 |
| 263826_at | 402.9 | P | 388.1 | P | 505.6 | P | 405.3 | P | --- |
| 263827_at | 256.5 | P | 309.7 | P | 259 | P | 290.5 | P | --- |
| 263828_at | 3.2 | A | 4.8 | A | 4 | A | 2.1 | A | --- |
| 263829_at | 1321.8 | P | 296 | P | 745.9 | P | 324.2 | P | --- |
| 263830_at | 55.2 | P | 89.1 | P | 93.9 | P | 85 | P | --- |
| 263831_at | 1271.1 | P | 814.8 | P | 1134.2 | P | 1068.7 | P | ATFER4 |
| 263832_at | 18.7 | P | 18 | A | 13.5 | A | 15.3 | A | --- |
| 263833_at | 121.3 | P | 124.3 | P | 111.8 | P | 130.5 | P | --- |
| 263834_at | 564 | P | 565.6 | P | 538 | P | 609.2 | P | --- |
| 263835_at | 1648.1 | P | 1622.7 | P | 1911.3 | P | 1663.6 | P | --- |
| 263836_at | 124.5 | P | 273.7 | P | 307.5 | P | 267.7 | P | --- |
| 263837_at | 27.2 | A | 5.8 | A | 10.8 | A | 6.2 | A | --- |
| 263838_at | 6180.1 | P | 6171.6 | P | 6770.2 | P | 6834.4 | P | MAT3 |
| 263839_at | 726.3 | P | 812.4 | P | 599.8 | P | 677.7 | P | MEMB11 |
| 263840_at | 1078.1 | P | 545.6 | P | 1153.1 | P | 993.3 | P | --- |
| 263841_at | 1337.9 | P | 2200.1 | P | 1534.8 | P | 2005.7 | P | --- |
| 263842_at | 2152.2 | P | 1998 | P | 2148.6 | P | 2212.7 | P | --- |
| 263843_at | 404.5 | P | 371.8 | P | 433.9 | P | 390.9 | P | --- |
| 263844_at | 3208.3 | P | 1791.1 | P | 3633.5 | P | 2081.7 | P | --- |
| 263845_at | 3731.9 | P | 2750.4 | P | 3861.3 | P | 4204.1 | P | pal1 |
| 263846_at | 1043.4 | P | 1013 | P | 1071.1 | P | 963.9 | P | SIGF |
| 263847_at | 1491.2 | P | 982.9 | P | 1177.8 | P | 1058.7 | P | --- |
| 263848_at | 19.2 | A | 2.9 | A | 6.7 | A | 4.4 | A | --- |
| 263849_at | 27.6 | A | 22 | A | 15.5 | A | 12.5 | A | --- |
| 263850_at | 15.8 | A | 46.5 | A | 21.1 | A | 19.3 | A | --- |
| 263851_at | 124.1 | P | 184.7 | P | 143.9 | P | 186.7 | P | --- |
| 263852_at | 83.6 | P | 326.9 | P | 27.7 | A | 175.5 | P | ATNUDT6 |
| 263853_at | 3.1 | A | 10.6 | A | 6 | A | 42.4 | A | --- |
| 263854_at | 40.6 | A | 87.4 | P | 68.7 | P | 68.1 | P | atnudt5 |
| 263855_at | 8.3 | A | 22.7 | P | 49.9 | A | 13.5 | A | --- |
| 263856_at | 1306.3 | P | 1349.4 | P | 1075.3 | P | 1364.3 | P | --- |
| 263857_at | 11.3 | A | 16.7 | A | 4.5 | A | 28.8 | A | --- |
| 263858_at | 0.6 | A | 1.6 | A | 2.4 | A | 4.6 | A | --- |
| 263859_at | 625.4 | P | 867 | P | 852 | P | 857 | P | --- |
| 263860_at | 6.1 | A | 5.8 | A | 11.3 | A | 9.9 | A | --- |
| 263861_at | 49.3 | A | 39.4 | A | 71.5 | A | 33.6 | A | --- |
| 263862_at | 382.3 | P | 366.4 | P | 356.4 | P | 282.9 | P | IBR5 |

| | | | | | | | | | |
|---|---|---|---|---|---|---|---|---|---|
| 263863_at | 243.5 | P | 216.6 | P | 229.1 | P | 203.8 | P | --- |
| 263864_at | 629 | P | 860.9 | P | 831.3 | P | 754.1 | P | CPZ |
| 263865_at | 1566 | P | 1820.5 | P | 1383.1 | P | 1796.1 | P | ABCB1 |
| 263866_at | 1402.2 | P | 811.5 | P | 1165.8 | P | 762.6 | P | --- |
| 263867_at | 3007.5 | P | 4931.8 | P | 3484.7 | P | 5096.6 | P | GAMMA-TIP |
| 263868_at | 66 | P | 80.6 | P | 81.3 | P | 90.9 | P | --- |
| 263869_at | 50.8 | P | 32.5 | M | 44 | M | 115.3 | P | PROPEP6 |
| 263870_at | 383.2 | P | 299.8 | P | 265.8 | P | 293.3 | P | RKP |
| 263871_at | 174 | P | 169.4 | P | 169 | P | 171.2 | P | RKP |
| 263872_at | 56.1 | P | 100.5 | P | 55.1 | P | 96.7 | P | CESA9 |
| 263873_at | 1603.9 | P | 1201.3 | P | 1799 | P | 1215.6 | P | --- |
| 263874_at | 5703 | P | 5890.8 | P | 5908.9 | P | 6444.6 | P | --- |
| 263875_at | 1639.2 | P | 2033.5 | P | 2038.7 | P | 2054.6 | P | 41519 |
| 263876_at | 91.8 | P | 108.8 | P | 92.9 | P | 100.1 | P | ATRAB7A |
| 263877_at | 29.5 | P | 8.3 | A | 12.5 | A | 11.8 | A | --- |
| 263878_s_at | 659.9 | P | 720.1 | P | 594.5 | P | 690 | P | --- |
| 263879_at | 2.4 | A | 1.7 | A | 8.5 | A | 24.2 | A | --- |
| 263880_at | 1238.7 | P | 1667.1 | P | 1439.6 | P | 1700.7 | P | --- |
| 263881_at | 31.9 | P | 20.9 | A | 41.3 | A | 14.1 | A | --- |
| 263882_at | 1034.9 | P | 1113.2 | P | 1199.2 | P | 1141.2 | P | RNR1 |
| 263883_at | 181.2 | P | 228.7 | P | 121.3 | P | 194.5 | P | --- |
| 263884_at | 11.6 | A | 21.4 | A | 5.9 | A | 4.5 | A | --- |
| 263885_at | 24.1 | P | 18.8 | P | 16.9 | A | 11.3 | A | --- |
| 263886_at | 222.9 | P | 276.3 | P | 240.7 | P | 243.5 | P | TKI1 |
| 263887_at | 58.4 | P | 57.2 | P | 55.5 | P | 74.8 | P | --- |
| 263888_at | 1.5 | A | 2.6 | A | 3.5 | A | 1.3 | A | --- |
| 263889_at | 36.2 | A | 12.1 | A | 41.7 | A | 23.8 | A | ATNAP12 |
| 263890_at | 3.2 | A | 2.8 | A | 18.5 | A | 8.7 | A | --- |
| 263891_at | 920.9 | P | 1109.3 | P | 880.5 | P | 1010.1 | P | GSL8 |
| 263892_at | 13.5 | A | 23.5 | A | 34.2 | A | 21.4 | A | RAX2 |
| 263893_at | 8.4 | A | 5.3 | A | 14.7 | A | 23 | A | WRKY59 |
| 263894_at | 9.1 | A | 2.9 | A | 6.1 | A | 1.8 | A | CYP96A5 |
| 263895_at | 33.3 | A | 24.4 | A | 32.7 | A | 35.8 | A | --- |
| 263896_at | 17.5 | A | 54.7 | P | 43.5 | P | 25.2 | A | --- |
| 263897_at | 235.5 | P | 225.9 | P | 311.5 | P | 278.5 | P | --- |
| 263898_at | 527.4 | P | 565.5 | P | 567.6 | P | 500.2 | P | SKIP6 |
| 263899_at | 100.9 | P | 119.6 | P | 129.6 | P | 109.3 | P | EMB2219 |
| 263900_at | 659.4 | P | 655.6 | P | 659.1 | P | 746 | P | --- |
| 263901_at | 455.4 | P | 725.5 | P | 180.1 | P | 490.1 | P | --- |
| 263902_at | 399.4 | P | 468.3 | P | 487.9 | P | 448.3 | P | APG10 |
| 263903_at | 9 | A | 2.1 | A | 19 | A | 1.8 | A | --- |
| 263904_at | 2187.4 | P | 1722.4 | P | 1698.2 | P | 1652.8 | P | PDR6 |
| 263905_at | 17 | A | 32.8 | A | 44.7 | A | 17.4 | A | AtcwINV4 |
| 263906_at | 1613.3 | P | 1551.7 | P | 1666 | P | 1822.5 | P | FTSZ2-1 |
| 263907_at | 185.2 | P | 60.4 | A | 136 | P | 40.3 | A | ABI5 |
| 263908_at | 270.3 | P | 285.5 | P | 271.9 | P | 282.8 | P | --- |
| 263909_at | 77.9 | A | 123 | P | 96.9 | A | 144.4 | P | DML1 |
| 263910_at | 1.6 | A | 2.8 | A | 7.3 | A | 2 | A | --- |
| 263911_at | 25.5 | A | 22.5 | A | 57.2 | P | 18.2 | A | --- |
| 263912_at | 1128.9 | P | 1267.3 | P | 1193.6 | P | 1410.9 | P | SBE2.1 |
| 263913_at | 186.3 | P | 348.6 | P | 253.5 | P | 354.2 | P | --- |
| 263914_at | 245.6 | P | 269.8 | P | 395 | P | 252.5 | P | AtGRF3 |
| 263915_at | 482.5 | P | 331.7 | P | 456.1 | P | 323.1 | P | --- |
| 263916_at | 4.1 | A | 0.5 | A | 2.2 | A | 14 | A | --- |
| 263917_at | 42.4 | A | 19.1 | A | 48.5 | A | 35.8 | A | HRD |
| 263918_at | 92 | A | 52.2 | A | 136.4 | P | 106.4 | M | ProT3 |
| 263919_at | 123.3 | P | 114.4 | P | 133.1 | P | 93.5 | P | --- |
| 263920_at | 471.2 | P | 531.5 | P | 469 | P | 564.9 | P | --- |
| 263921_at | 2959.2 | P | 3417.7 | P | 2939.9 | P | 3216.4 | P | --- |
| 263922_s_at | 2369.6 | P | 2060.3 | P | 2153.1 | P | 2212.3 | P | --- |
| 263923_at | 135 | P | 184.5 | P | 241.9 | P | 176.2 | P | --- |
| 263924_at | 9256.2 | P | 7561.3 | P | 9809.5 | P | 8173.2 | P | LOS2 |
| 263925_at | 12.1 | A | 2.4 | A | 6.9 | A | 3.7 | A | --- |
| 263926_at | 34 | P | 69.8 | P | 60.4 | P | 71.5 | P | ATEME1A |
| 263927_s_at | 43.3 | P | 64.6 | P | 60.6 | P | 33.3 | P | CAD2 |
| 263928_at | 278 | P | 365 | P | 213.4 | P | 306.8 | P | --- |
| 263929_at | 405 | P | 513.2 | P | 367.2 | P | 415.6 | P | URH1 |
| 263930_at | 526 | P | 598.2 | P | 541.1 | P | 642.5 | P | --- |
| 263931_at | 240.7 | P | 379.9 | P | 247.6 | P | 334 | P | --- |
| 263932_at | 20.3 | A | 35.9 | P | 46.4 | P | 24.3 | A | --- |
| 263933_s_at | 8.5 | A | 14.2 | A | 6.7 | A | 10.5 | A | --- |
| 263934_at | 26.9 | P | 46.7 | P | 52.4 | P | 36.9 | M | EDA12 |
| 263935_at | 85.1 | P | 87.3 | P | 115.9 | P | 70.6 | P | PUB23 |
| 263936_at | 270.2 | P | 272.7 | P | 293.7 | P | 361.7 | P | --- |
| 263937_at | 15.4 | A | 28.1 | A | 16.9 | A | 39.1 | A | --- |
| 263938_at | 488.3 | P | 499.9 | P | 499.1 | P | 487.3 | P | --- |
| 263939_at | 622.8 | P | 506.5 | P | 733.8 | P | 568.1 | P | ATTIM44-2 |
| 263940_at | 3.2 | A | 3.1 | A | 3 | A | 2 | A | CPK25 |
| 263941_at | 69.4 | P | 71.8 | P | 67.3 | P | 36.9 | P | --- |
| 263942_at | 619.3 | P | 644.5 | P | 531.8 | P | 796.7 | P | FLA16 |
| 263943_at | 515.3 | P | 560.3 | P | 597.9 | P | 606.7 | P | --- |
| 263944_at | 9.3 | A | 15.8 | A | 10.1 | A | 17.4 | A | --- |
| 263945_at | 17.1 | A | 23.7 | A | 23.6 | A | 13.6 | A | --- |
| 263946_at | 548.7 | P | 671.4 | P | 586 | P | 629.8 | P | --- |
| 263947_at | 639.2 | P | 462.8 | P | 604.1 | P | 422.3 | P | --- |
| 263948_at | 53.9 | A | 46.4 | A | 11.5 | A | 25.5 | A | YLS9 |
| 263949_at | 1603.5 | P | 1542.3 | P | 1399.7 | P | 1514.5 | P | MMZ3 |
| 263950_at | 29.1 | A | 3.1 | A | 6.6 | A | 14.2 | A | HVA22J |
| 263951_at | 482.7 | P | 432.5 | P | 683 | P | 555.5 | P | NHL12 |
| 263952_s_at | 2120.2 | P | 2565.5 | P | 2181.2 | P | 2593.2 | P | --- |
| 263953_at | 8.6 | A | 10.7 | A | 14.3 | A | 13.7 | A | OFP15 |
| 263954_at | 1995.5 | P | 1662.9 | P | 2228.2 | P | 1974.2 | P | --- |
| 263955_at | 81.5 | P | 98.8 | P | 141.1 | P | 115.8 | P | E2F3 |
| 263956_at | 1255.6 | P | 1487.6 | P | 1107.7 | P | 1408.5 | P | BLH1 |
| 263957_at | 583.9 | P | 554.7 | P | 488.9 | P | 500.6 | P | --- |
| 263958_at | 492.9 | P | 604 | P | 553.2 | P | 607.7 | P | --- |
| 263959_at | 1.7 | A | 6.2 | A | 3 | A | 2.1 | A | --- |
| 263960_at | 101.5 | A | 181.9 | A | 130.7 | A | 206.8 | A | --- |
| 263961_at | 79.5 | P | 84.2 | P | 62.8 | A | 97.1 | P | --- |
| 263962_at | 136.3 | P | 186.9 | P | 137.5 | P | 168.7 | P | --- |
| 263963_at | 267 | P | 340.1 | P | 315.7 | P | 194.7 | P | --- |
| 263964_at | 1.4 | A | 2.5 | A | 4.6 | A | 2.2 | A | --- |
| 263965_at | 2.3 | A | 2.4 | A | 28.7 | A | 3 | A | --- |
| 263966_s_at | 1.1 | A | 0.6 | A | 1.9 | A | 1.7 | A | --- |
| 263967_at | 3.7 | A | 0.4 | A | 1.4 | A | 3.9 | A | --- |
| 263968_s_at | 1.5 | A | 7.4 | A | 2.5 | A | 2 | A | --- |
| 263969_at | 1.9 | A | 2.1 | A | 4.4 | A | 1.2 | A | UNE4 |
| 263970_at | 2.6 | A | 2.4 | A | 3.6 | A | 2.7 | A | CYP718 |
| 263971_at | 142.4 | P | 171 | P | 205.2 | P | 163.7 | P | AtRLP29 |

| | | | | | | | | | |
|---|---|---|---|---|---|---|---|---|---|
| 263972_at | 390.5 | P | 129.9 | P | 226 | P | 107.2 | P | --- |
| 263973_at | 994 | P | 970.9 | P | 901.4 | P | 843.1 | P | RPL16A |
| 263974_at | 35.8 | A | 38.3 | A | 60.4 | A | 34.9 | A | --- |
| 263975_at | 541.9 | P | 464.1 | P | 616.1 | P | 557.8 | P | --- |
| 263976_at | 247.8 | P | 258.1 | P | 242.7 | P | 266.2 | P | --- |
| 263977_at | 4.3 | A | 35.7 | A | 20.6 | A | 4.5 | A | --- |
| 263978_at | 3874.2 | P | 3608.6 | P | 4086.2 | P | 4162.8 | P | MBF1A |
| 263979_at | 817 | P | 1479.2 | P | 985.5 | P | 1511.4 | P | PDF1 |
| 263980_at | 1090.6 | P | 1128.1 | P | 1207.7 | P | 1201.1 | P | --- |
| 263981_at | 69 | P | 109.1 | M | 52.1 | A | 75.8 | A | PAR1 |
| 263982_at | 23.6 | A | 8.5 | A | 46 | A | 14.1 | A | --- |
| 263983_at | 433.7 | P | 343.7 | P | 326.5 | P | 345.9 | P | --- |
| 263984_at | 207.1 | P | 292.8 | P | 229.2 | P | 231 | P | --- |
| 263985_at | 650.2 | P | 1446 | P | 1655.7 | P | 1529.1 | P | --- |
| 263986_at | 1542.9 | P | 1379.5 | P | 1038.1 | P | 1243.8 | P | CSY3 |
| 263987_at | 2141.1 | P | 2287.9 | P | 2519.2 | P | 2292.1 | P | --- |
| 263988_at | 31.4 | A | 6.3 | A | 46.1 | A | 7 | A | SHP2 |
| 263989_at | 607.8 | P | 790.9 | P | 553.8 | P | 707.8 | P | ATMPK20 |
| 263990_at | 804.1 | P | 824.6 | P | 1010.2 | P | 878.3 | P | PP5.2 |
| 263991_at | 41.9 | P | 18.3 | P | 17.2 | A | 18 | P | --- |
| 263992_at | 13.1 | A | 1.9 | A | 16.5 | A | 5.5 | A | --- |
| 263993_at | 13.7 | A | 9.2 | A | 52.7 | A | 5.4 | A | --- |
| 263994_at | 15 | A | 6.9 | A | 42 | A | 3 | A | --- |
| 263995_at | 262.1 | P | 309.8 | P | 302.7 | P | 264.9 | P | SVP |
| 263996_at | 141.7 | P | 208 | P | 221.8 | P | 155 | P | --- |
| 263997_at | 23.9 | A | 18.2 | A | 30.6 | A | 28 | A | --- |
| 263998_at | 17.4 | A | 56.7 | P | 48 | P | 53.8 | P | --- |
| 263999_at | 294.1 | P | 317.8 | P | 277.6 | P | 281.7 | P | --- |
| 264000_at | 1854.5 | P | 1776.9 | P | 1792.8 | P | 1707.6 | P | UCP5 |
| 264001_at | 209.6 | P | 132.9 | P | 163.4 | P | 150.9 | P | --- |
| 264002_at | 921.8 | P | 810.8 | P | 1015.6 | P | 812.5 | P | --- |
| 264003_at | 267.9 | P | 270.9 | P | 257.9 | P | 275.4 | P | GEM |
| 264004_at | 1927.9 | P | 1886.1 | P | 1575.9 | P | 1898.9 | P | --- |
| 264005_at | 679.8 | P | 179.1 | P | 114.1 | M | 185.5 | P | AGP2 |
| 264006_at | 1600.7 | P | 1749.8 | P | 1380.5 | P | 1594.9 | P | ATHB6 |
| 264007_at | 686.9 | P | 746.3 | P | 635.6 | P | 806.3 | P | ATPRP2 |
| 264008_at | 666.5 | P | 583.3 | P | 639.4 | P | 671.8 | P | --- |
| 264009_at | 18.6 | A | 10.7 | A | 21.8 | A | 24.3 | A | --- |
| 264010_at | 3.9 | A | 43.3 | A | 44.4 | A | 48 | A | --- |
| 264011_at | 82.9 | P | 73.7 | P | 118.1 | P | 126.8 | P | --- |
| 264012_at | 230.3 | P | 240.6 | P | 220.5 | P | 188.1 | P | --- |
| 264013_at | 94.4 | P | 137.7 | P | 121.1 | P | 114.2 | P | FIO1 |
| 264014_at | 28.4 | A | 112.7 | P | 14.9 | A | 113.3 | P | --- |
| 264015_at | 374.3 | P | 371.7 | P | 372.4 | P | 380.6 | P | XCT |
| 264016_at | 53.4 | A | 96.4 | P | 48.7 | A | 128 | P | --- |
| 264017_s_at | 873.5 | P | 937.9 | P | 763.8 | P | 851.4 | P | --- |
| 264018_at | 9541.5 | P | 8391 | P | 8181.4 | P | 9571.4 | P | TIM |
| 264019_at | 1916.2 | P | 1289.4 | P | 2134.1 | P | 1789.4 | P | --- |
| 264020_at | 2434.6 | P | 2920.6 | P | 2765.8 | P | 2668.8 | P | --- |
| 264021_at | 14.8 | A | 41.3 | A | 5.7 | A | 32.9 | A | --- |
| 264022_at | 474.6 | P | 531.4 | P | 387.2 | P | 406.9 | P | --- |
| 264023_at | 390.5 | P | 359.1 | P | 308 | P | 336.1 | P | --- |
| 264024_at | 469.1 | P | 370.6 | P | 454.5 | P | 393.7 | P | --- |
| 264025_at | 99.7 | P | 199.2 | P | 196 | P | 237.8 | P | --- |
| 264026_at | 1197.9 | P | 1139.2 | P | 1092.7 | P | 1226.9 | P | ATGRP2B |
| 264027_at | 199.4 | P | 154.8 | P | 265.3 | P | 159 | P | CDC48B |
| 264028_at | 2245.6 | P | 2791.9 | P | 2490.7 | P | 2611.4 | P | SPR1 |
| 264029_at | 2.5 | A | 16.2 | A | 4.5 | A | 16.5 | A | MRH6 |
| 264030_at | 3.6 | A | 2.6 | A | 2 | A | 3 | A | --- |
| 264031_at | 434.9 | P | 500.3 | P | 487 | P | 458.5 | P | --- |
| 264032_at | 418.4 | P | 329.7 | P | 335.7 | P | 352.4 | P | GEK1 |
| 264033_at | 21 | A | 13 | P | 30.1 | P | 6.3 | A | --- |
| 264034_s_at | 52.9 | P | 103 | A | 71.7 | A | 65.7 | A | ATUPS1 |
| 264035_at | 3.1 | A | 6.2 | A | 14 | A | 1.7 | A | --- |
| 264036_at | 82.3 | M | 121.3 | P | 96.7 | A | 102.3 | P | --- |
| 264037_at | 549.5 | P | 659 | P | 677.5 | P | 679.9 | P | --- |
| 264038_at | 773.7 | P | 724 | P | 523.8 | P | 696.9 | P | --- |
| 264039_at | 5.9 | A | 4.4 | A | 3.8 | A | 5 | A | --- |
| 264040_at | 214.4 | P | 180.7 | P | 206.9 | P | 194.5 | P | ACR5 |
| 264041_at | 124.2 | P | 105.5 | P | 153 | P | 91.5 | P | 41521 |
| 264042_at | 1949.1 | P | 651.7 | P | 1172.8 | P | 690.4 | P | ST |
| 264043_at | 1013.5 | P | 571.6 | P | 1053.5 | P | 665 | P | CYCD2;1 |
| 264044_at | 801.1 | P | 994.2 | P | 911.8 | P | 818.9 | P | PFK5 |
| 264045_at | 1385.4 | P | 943.4 | P | 1226.4 | P | 1114.2 | P | --- |
| 264046_at | 20.2 | A | 24.9 | A | 30.1 | A | 18.4 | A | --- |
| 264047_at | 96.2 | P | 85.4 | P | 68.2 | P | 106.5 | P | --- |
| 264048_at | 1007.9 | P | 1114.5 | P | 1356 | P | 967.5 | P | --- |
| 264049_at | 50.3 | P | 60.8 | P | 63.3 | P | 75.7 | P | --- |
| 264050_at | 7.6 | A | 2.9 | A | 7.8 | A | 9.1 | A | --- |
| 264051_at | 28.4 | A | 1.1 | A | 14.4 | A | 2.9 | A | --- |
| 264052_at | 4104.6 | P | 671 | P | 3194.8 | P | 938.4 | P | CYP79B3 |
| 264053_at | 96 | P | 106.3 | P | 81.8 | P | 72.2 | P | --- |
| 264054_at | 1771.1 | P | 2213.6 | P | 1686.8 | P | 2070.5 | P | SVP |
| 264055_at | 29.8 | P | 62.9 | P | 64.9 | P | 51.5 | P | --- |
| 264056_at | 564.3 | P | 520.3 | P | 410 | P | 460.2 | P | --- |
| 264057_at | 420.6 | P | 613.5 | P | 479.8 | P | 601.5 | P | RAP2.7 |
| 264058_at | 3.9 | A | 4.1 | A | 22.1 | A | 4.7 | A | --- |
| 264059_at | 140.7 | P | 156 | P | 133.4 | P | 125.1 | P | INH3 |
| 264060_at | 183.6 | P | 211.2 | P | 198.7 | P | 260.4 | P | --- |
| 264061_at | 434.3 | P | 640.8 | P | 504.8 | P | 649.8 | P | CKS2 |
| 264062_at | 134.5 | P | 175.5 | P | 71.9 | A | 194.9 | P | --- |
| 264063_at | 578.1 | P | 551.4 | P | 419.9 | P | 500.5 | P | --- |
| 264064_at | 295.1 | P | 329.6 | P | 287.7 | P | 292.9 | P | --- |
| 264065_at | 103.5 | P | 104.3 | P | 120.2 | A | 99.1 | P | --- |
| 264066_at | 8.6 | A | 2.6 | A | 16.7 | A | 9 | A | AGO5 |
| 264067_x_at | 2.9 | A | 1.5 | A | 2.5 | A | 1.4 | A | --- |
| 264068_at | 87.6 | A | 90.3 | P | 64.7 | A | 105 | A | BLH8 |
| 264069_at | 7348.6 | P | 8685.7 | P | 7439.3 | P | 8104.2 | P | CPN60A |
| 264070_at | 1125.3 | P | 1164.1 | P | 1186.2 | P | 1036.9 | P | CKS1 |
| 264071_at | 102.7 | P | 313.1 | P | 169.4 | P | 298.1 | P | SCPL51 |
| 264072_at | 0.9 | A | 2.9 | A | 18.2 | A | 11.7 | A | --- |
| 264073_at | 10.8 | A | 9.6 | A | 27.2 | A | 4 | A | --- |
| 264074_at | 2 | A | 0.6 | A | 1.6 | A | 1.1 | A | --- |
| 264075_at | 20.7 | A | 20.1 | A | 56.1 | A | 24.3 | A | --- |
| 264076_at | 561.4 | P | 479.1 | P | 622.8 | P | 479.3 | P | --- |
| 264077_at | 42.4 | A | 52.2 | A | 32.8 | A | 52.6 | A | RAD51B |
| 264078_at | 951.3 | P | 1690.3 | P | 962.8 | P | 1767 | P | BGAL8 |
| 264079_at | 3.5 | A | 4.6 | A | 3.6 | A | 2 | A | --- |
| 264080_at | 481.7 | P | 586.8 | P | 469.6 | P | 647.4 | P | VHA-A1 |

| | | | | | | | | | |
|---|---|---|---|---|---|---|---|---|---|
| 264081_at | 126 | P | 176.5 | P | 156.3 | P | 180.4 | P | --- |
| 264082_at | 82 | P | 200 | P | 61.1 | A | 152.8 | P | --- |
| 264083_at | 165.4 | P | 134.6 | P | 105.3 | P | 119.9 | P | ATERF15 |
| 264084_at | 365.8 | P | 352.6 | P | 382.8 | P | 385.8 | P | --- |
| 264085_at | 2.6 | A | 18.7 | A | 34.3 | A | 10.8 | A | HEMA3 |
| 264086_at | 225.6 | P | 219.3 | P | 297.6 | P | 214.6 | P | --- |
| 264087_at | 129.1 | P | 133.9 | P | 160.3 | P | 164.1 | P | emb1381 |
| 264088_at | 77.4 | P | 76.2 | P | 104.5 | A | 93.1 | P | emb1381 |
| 264089_at | 2447.5 | P | 2239.3 | P | 1967.6 | P | 2070.7 | P | ADF6 |
| 264090_at | 25.7 | A | 36 | P | 53.1 | A | 36.6 | M | CYP79C1 |
| 264091_at | 1534.3 | P | 848.1 | P | 1312.9 | P | 1080.4 | P | --- |
| 264092_at | 16052.1 | P | 15726.3 | P | 19225.5 | P | 17078.7 | P | PSBR |
| 264093_at | 64.2 | A | 84.5 | A | 115.7 | A | 76.1 | P | --- |
| 264094_at | 125.2 | P | 172.8 | P | 182.6 | P | 130.3 | P | --- |
| 264095_at | 2397.6 | P | 2364 | P | 2510 | P | 2569.6 | P | MST1 |
| 264096_at | 461.7 | P | 569.6 | P | 680.1 | P | 535.9 | P | --- |
| 264097_s_at | 2791.9 | P | 2925.3 | P | 2841.5 | P | 2851.2 | P | --- |
| 264098_at | 840.2 | P | 586 | P | 759.5 | P | 722.4 | P | --- |
| 264099_at | 881.3 | P | 921.8 | P | 794 | P | 829.6 | P | --- |
| 264100_at | 96.1 | P | 513.7 | P | 227.9 | P | 684.6 | P | LUP1 |
| 264101_at | 635.5 | P | 599.9 | P | 736.3 | P | 624.7 | P | HAC1 |
| 264102_at | 835 | P | 676.7 | P | 663 | P | 696.7 | P | ECT8 |
| 264103_at | 0.9 | A | 30.3 | P | 11.7 | A | 11.3 | A | --- |
| 264104_at | 1.3 | A | 4.6 | A | 2.8 | A | 4.7 | A | --- |
| 264105_x_at | 3.8 | A | 4.7 | A | 7.8 | A | 3.8 | A | --- |
| 264106_at | 67.9 | P | 51.4 | P | 42.3 | A | 68.4 | P | SERK4 |
| 264107_s_at | 850.7 | P | 1031.9 | P | 772.6 | P | 817.2 | P | SERK5 |
| 264108_at | 7.8 | A | 1.1 | A | 6.4 | A | 2 | A | CALS5 |
| 264109_at | 1.5 | A | 5.1 | A | 24.3 | A | 3.6 | A | --- |
| 264110_at | 0.8 | A | 0.6 | A | 0.8 | A | 13.4 | A | --- |
| 264111_at | 270.4 | P | 344.9 | P | 243 | P | 411.5 | P | --- |
| 264112_at | 38.6 | A | 59 | A | 50.9 | A | 47.5 | A | CALS5 |
| 264113_at | 225.3 | P | 278.9 | P | 254.3 | P | 247.2 | P | APG9 |
| 264114_at | 106.3 | P | 191.7 | P | 195.2 | P | 155.7 | P | CDT1A |
| 264115_at | 91.3 | P | 94.9 | P | 122.6 | P | 97 | P | --- |
| 264116_at | 120.7 | P | 165.2 | P | 188.2 | P | 192.5 | P | PARP2 |
| 264117_at | 0.6 | A | 10 | A | 1.2 | A | 14.5 | A | --- |
| 264118_at | 278.7 | P | 284.6 | P | 360.1 | P | 279.8 | P | --- |
| 264119_at | 41.9 | A | 19.8 | A | 32.5 | A | 14.1 | A | MYB63 |
| 264120_at | 1400.2 | P | 1661 | P | 1458.2 | P | 1579.5 | P | AtMC4 |
| 264121_at | 1576.3 | P | 2180.4 | P | 1430.5 | P | 1766.3 | P | TOC33 |
| 264122_at | 710.3 | P | 744.5 | P | 592.3 | P | 659.1 | P | ARA-5 |
| 264123_at | 533.2 | P | 821.5 | P | 517.3 | P | 815.6 | P | --- |
| 264124_at | 84.3 | P | 49.9 | P | 136.7 | P | 60.4 | P | ATOCT2 |
| 264125_at | 289.8 | P | 240.6 | P | 307 | P | 313.4 | P | EMB1135 |
| 264126_at | 395.8 | P | 422.2 | P | 468.2 | P | 415.6 | P | NUA |
| 264127_at | 5.2 | A | 21.7 | A | 30.3 | A | 5.8 | A | AGC1.7 |
| 264128_at | 134.3 | P | 139.1 | P | 108.2 | P | 126.7 | P | --- |
| 264129_at | 12.2 | A | 2.1 | A | 16.7 | A | 3.7 | A | --- |
| 264130_at | 390.3 | P | 270.4 | P | 349.1 | P | 323.6 | P | --- |
| 264131_at | 421 | P | 331 | P | 513 | P | 430.7 | P | --- |
| 264132_at | 2.3 | A | 1.3 | A | 4.7 | A | 14.7 | A | --- |
| 264133_at | 337.9 | P | 279.2 | P | 345.6 | P | 301.6 | P | --- |
| 264134_at | 111.1 | P | 59.2 | P | 109.4 | P | 111.6 | P | --- |
| 264135_at | 146.2 | P | 175.6 | P | 190 | P | 182.3 | P | --- |
| 264136_at | 22.8 | A | 28.3 | A | 39.7 | A | 21.2 | A | SRF5 |
| 264137_at | 157.2 | P | 124.1 | P | 193.6 | P | 102.9 | P | ATLUP2 |
| 264138_at | 34.1 | A | 29.1 | A | 47.3 | A | 21.9 | A | --- |
| 264139_at | 21.7 | A | 18.7 | P | 20.1 | A | 24.7 | M | --- |
| 264140_at | 976.4 | P | 1098.4 | P | 1126.1 | P | 1038.3 | P | --- |
| 264141_at | 785.8 | P | 790.9 | P | 747.9 | P | 752.8 | P | --- |
| 264142_at | 111.4 | P | 117.2 | P | 144 | P | 69.9 | P | --- |
| 264143_at | 9.5 | A | 2.9 | A | 11.1 | A | 19.4 | A | ATMC5 |
| 264144_at | 19.6 | A | 17.7 | A | 32.6 | A | 14.2 | A | AtMC6 |
| 264145_at | 17.4 | A | 12.6 | A | 17.4 | A | 27 | A | AtMC7 |
| 264146_at | 493.7 | P | 760.9 | P | 260.4 | P | 801.7 | P | CER1 |
| 264147_at | 227.2 | P | 385.2 | P | 168 | P | 449.9 | P | CER1 |
| 264148_at | 239.6 | P | 127.7 | P | 98.4 | P | 127.6 | P | ANAC003 |
| 264149_at | 3.1 | A | 25.6 | A | 25.6 | A | 8.2 | A | anac005 |
| 264150_at | 539.4 | P | 434 | P | 452.5 | P | 465.8 | P | FUS5 |
| 264151_at | 38.8 | A | 35.6 | A | 27.4 | A | 16.6 | A | --- |
| 264152_at | 3435 | P | 3233.6 | P | 3463 | P | 3496.6 | P | MAGO |
| 264153_at | 37.2 | P | 107.6 | P | 50.7 | A | 66.5 | P | ATPP2-A5 |
| 264154_at | 498 | P | 444.9 | P | 397.5 | P | 489.8 | P | --- |
| 264155_at | 300.4 | P | 208.1 | P | 239.2 | P | 244.9 | P | GTB1 |
| 264156_at | 503.4 | P | 424.9 | P | 452 | P | 478.8 | P | --- |
| 264157_at | 19.9 | A | 63.9 | P | 2.8 | A | 22 | A | XTH17 |
| 264158_at | 3640.4 | P | 4313.2 | P | 4153.3 | P | 4379.9 | P | PTAC4 |
| 264159_at | 809.7 | P | 917.1 | P | 829.9 | P | 843 | P | --- |
| 264160_at | 183.4 | P | 121 | P | 178.6 | P | 187.2 | P | --- |
| 264161_at | 349.4 | P | 485.9 | P | 420.7 | P | 461.4 | P | --- |
| 264162_at | 4324.1 | P | 4836.6 | P | 5994.2 | P | 5198.7 | P | mtACP2 |
| 264163_at | 57.2 | A | 59.9 | M | 59.3 | A | 72.9 | P | --- |
| 264164_at | 606.6 | P | 670.8 | P | 690.1 | P | 740.9 | P | --- |
| 264165_at | 751.5 | P | 758.1 | P | 553.9 | P | 838.7 | P | ATNAP11 |
| 264166_at | 561.4 | P | 407.3 | P | 559.1 | P | 436.8 | P | --- |
| 264167_at | 95.6 | P | 88.7 | P | 125.2 | P | 98.5 | P | --- |
| 264168_at | 1212.6 | P | 1073.6 | P | 1136.8 | P | 955.4 | P | --- |
| 264169_at | 196.2 | P | 198.9 | P | 240.6 | P | 197.5 | P | --- |
| 264170_at | 506.6 | P | 451.2 | P | 418.9 | P | 441.8 | P | --- |
| 264171_at | 564.8 | P | 476 | P | 540.6 | P | 481.8 | P | --- |
| 264172_at | 1166 | P | 1012.4 | P | 1426.8 | P | 1068.4 | P | VAD1 |
| 264173_at | 1141.5 | P | 941.8 | P | 1036.3 | P | 888.9 | P | --- |
| 264174_s_at | 9.4 | A | 22.3 | A | 23.2 | A | 12.6 | A | ANAC004 |
| 264175_at | 101.6 | P | 71.1 | P | 133.3 | P | 73.5 | P | --- |
| 264176_at | 292.6 | P | 227.5 | P | 283 | P | 242.2 | P | --- |
| 264177_at | 2144.2 | P | 2197.6 | P | 2542.4 | P | 2227.2 | P | --- |
| 264178_at | 655.7 | P | 688.4 | P | 634.2 | P | 657.7 | P | AMC1 |
| 264179_at | 129 | P | 157.9 | P | 148.2 | P | 144.7 | P | --- |
| 264180_at | 2.9 | A | 1.6 | A | 12.6 | A | 2.7 | A | --- |
| 264181_at | 63.4 | P | 46 | P | 49 | P | 53.5 | P | UBQ13 |
| 264182_at | 27 | P | 19.3 | A | 22.6 | A | 26.2 | P | AGL23 |
| 264183_at | 122 | P | 166.4 | P | 165.8 | P | 112 | P | CLV2 |
| 264184_at | 28.5 | A | 56.5 | P | 89 | P | 47 | P | --- |
| 264185_at | 6338.5 | P | 9100.2 | P | 5713.8 | P | 8153.9 | P | --- |
| 264186_at | 428.1 | P | 525.7 | P | 512.1 | P | 524.7 | P | --- |
| 264187_at | 2.6 | A | 2.3 | A | 2 | A | 1.4 | A | --- |
| 264188_at | 1185.3 | P | 1239 | P | 1255.4 | P | 1240.5 | P | GAMMA-H2AX |
| 264189_s_at | 4661.8 | P | 4810 | P | 4883.7 | P | 5553 | P | ACP3 |

| | | | | | | | | | |
|---|---|---|---|---|---|---|---|---|---|
| 264190_at | 388 | P | 434.8 | P | 422.9 | P | 460.4 | P | NF-YC3 |
| 264191_at | 184.4 | P | 247.9 | P | 223 | P | 211.6 | P | --- |
| 264192_at | 373.5 | P | 385.9 | P | 314.1 | P | 314.2 | P | ATATG18H |
| 264193_at | 745.9 | P | 914.6 | P | 1117.4 | P | 1025.4 | P | --- |
| 264194_at | 14.8 | P | 41.2 | A | 55.5 | A | 27.5 | P | --- |
| 264195_at | 1856.4 | P | 1177.6 | P | 2360.7 | P | 1324.9 | P | --- |
| 264196_at | 29.9 | A | 34.9 | A | 67.1 | A | 34.7 | A | --- |
| 264197_at | 271.4 | P | 331.9 | P | 231.3 | P | 294 | P | ATSAC1 |
| 264198_at | 348.5 | P | 298.2 | P | 313.8 | P | 330.9 | P | --- |
| 264199_at | 1161.4 | P | 1250.9 | P | 1284 | P | 1285.8 | P | --- |
| 264200_at | 47.1 | A | 132.8 | P | 51.6 | A | 94.5 | P | --- |
| 264201_at | 408.7 | P | 497.1 | P | 462.5 | P | 534.7 | P | --- |
| 264202_at | 25 | A | 17.9 | A | 4.5 | A | 2.5 | A | --- |
| 264203_at | 5905.6 | P | 5326.8 | P | 6866.4 | P | 5823.7 | P | PFL |
| 264204_at | 1281.6 | P | 1094.3 | P | 1157.2 | P | 1010.8 | P | SUC2 |
| 264205_at | 269.3 | P | 322.5 | P | 274.2 | P | 341.7 | P | --- |
| 264206_at | 426 | P | 331.6 | P | 360.6 | P | 359 | P | --- |
| 264207_at | 1314.7 | P | 1811.2 | P | 1531.7 | P | 1685.1 | P | --- |
| 264208_at | 47.9 | A | 40 | A | 28.4 | A | 47.1 | A | PAB3 |
| 264209_at | 129.4 | P | 207.3 | P | 116.6 | P | 181.9 | P | RABG3B |
| 264210_at | 220.2 | P | 226.1 | P | 135.6 | P | 168.1 | P | MYB3 |
| 264211_at | 961.8 | P | 879.1 | P | 496.5 | P | 797.2 | P | GI |
| 264212_at | 247.6 | P | 245.6 | P | 261.9 | P | 237.2 | P | --- |
| 264213_at | 85.2 | P | 142.9 | P | 126.8 | P | 145.9 | P | ATPP2-A5 |
| 264214_s_at | 2.5 | A | 3.8 | A | 4.4 | A | 1.3 | A | AGL38 |
| 264215_at | 5.3 | A | 3.4 | A | 21.9 | A | 11 | A | CYP96A3 |
| 264216_at | 13.1 | A | 3 | A | 14.3 | A | 0.9 | A | --- |
| 264217_at | 197.8 | P | 117.7 | A | 94.5 | A | 168.9 | M | --- |
| 264218_at | 398.4 | P | 390.7 | P | 506.7 | P | 375.3 | P | --- |
| 264219_at | 676.9 | P | 933.5 | P | 690 | P | 823 | P | --- |
| 264220_at | 269.7 | P | 246.9 | P | 333.8 | P | 280.7 | P | ARPC3 |
| 264221_s_at | 440.8 | P | 390.7 | P | 370.8 | P | 369.5 | P | emb1220 |
| 264222_at | 318.6 | P | 324 | P | 313.8 | P | 329.3 | P | --- |
| 264223_s_at | 151.9 | P | 156.2 | P | 95.3 | P | 116.6 | P | --- |
| 264224_at | 364.6 | P | 350.6 | P | 243.7 | P | 291.6 | P | emb1688 |
| 264225_at | 151.7 | P | 149.6 | P | 175.6 | P | 133.7 | P | --- |
| 264226_at | 130 | P | 120.9 | P | 169.1 | P | 181 | P | --- |
| 264227_at | 244.4 | P | 158.7 | P | 168.8 | P | 184.9 | P | ATREV3 |
| 264228_at | 303.5 | P | 295.6 | P | 293.2 | P | 285.6 | P | GCS1 |
| 264229_at | 675 | P | 984.4 | P | 959.5 | P | 770.7 | P | --- |
| 264230_at | 21.3 | M | 11.4 | A | 36.5 | P | 30.3 | A | --- |
| 264231_at | 1.9 | A | 30.9 | A | 17.9 | A | 13.2 | A | --- |
| 264232_at | 407.6 | P | 310.7 | P | 527.2 | P | 375.6 | P | --- |
| 264233_at | 10816.3 | P | 9385.8 | P | 11490.4 | P | 10136.5 | P | --- |
| 264234_at | 17.5 | A | 11.9 | A | 14.4 | A | 10.3 | A | --- |
| 264235_at | 16.1 | A | 29.3 | A | 23 | A | 23.1 | A | XIE |
| 264236_at | 182.3 | P | 193.2 | P | 140.6 | P | 144.9 | P | --- |
| 264237_at | 10.6 | A | 14.8 | A | 26 | A | 6.9 | A | --- |
| 264238_at | 208.8 | P | 179.8 | P | 286.6 | P | 193 | P | --- |
| 264239_at | 501.3 | P | 556.1 | P | 631 | P | 509.9 | P | --- |
| 264240_at | 195.3 | P | 362.8 | P | 267.9 | P | 325.3 | P | --- |
| 264241_at | 37.3 | A | 34.8 | P | 63.2 | P | 40.7 | P | --- |
| 264242_at | 25.7 | A | 32.3 | A | 14.2 | A | 18.5 | A | --- |
| 264243_at | 211.3 | P | 209.6 | P | 176 | P | 216.4 | P | --- |
| 264244_at | 1021.9 | P | 765.9 | P | 857.2 | P | 799.1 | P | ATPANK1 |
| 264245_at | 104.2 | P | 55.1 | A | 133 | A | 47.7 | A | AtGolS7 |
| 264246_at | 274.1 | P | 207.1 | P | 173.6 | P | 182.2 | P | ATTPS10 |
| 264247_at | 407.8 | P | 368.4 | P | 436.6 | P | 335.9 | P | --- |
| 264248_at | 227.3 | P | 185.5 | P | 273.2 | P | 157.1 | P | --- |
| 264249_at | 1221.9 | P | 1607.2 | P | 1199.4 | P | 1446.4 | P | AVP2 |
| 264250_at | 3334.9 | P | 2798 | P | 2792.8 | P | 2809.5 | P | ATGGH2 |
| 264251_at | 35.6 | P | 66.8 | P | 45.5 | P | 61.7 | A | --- |
| 264252_at | 132.3 | P | 85.7 | A | 96.2 | A | 114.7 | P | ATSARA1A |
| 264253_at | 8 | A | 17.5 | A | 8.2 | A | 17.2 | A | --- |
| 264254_at | 973.5 | P | 983.6 | P | 783.4 | P | 1115.7 | P | --- |
| 264255_at | 825.5 | P | 844.3 | P | 806.8 | P | 806.5 | P | --- |
| 264256_at | 518.6 | P | 716.6 | P | 558.9 | P | 633.4 | P | IMPA-4 |
| 264257_at | 180.4 | P | 163.8 | P | 225.2 | P | 237.8 | P | --- |
| 264258_at | 71.7 | P | 56.1 | P | 66.8 | P | 62.2 | P | --- |
| 264259_at | 159 | P | 212.4 | P | 171 | P | 176.6 | P | --- |
| 264260_at | 3113.9 | P | 3240.9 | P | 2655.5 | P | 3394.4 | P | --- |
| 264261_at | 503.8 | P | 885.6 | P | 408.3 | P | 668.9 | P | NAS3 |
| 264262_at | 1182.1 | P | 1888.7 | P | 1298.5 | P | 1780.6 | P | --- |
| 264263_at | 3.1 | A | 8.5 | A | 13 | A | 11 | A | AtPP2-B15 |
| 264264_at | 203.8 | P | 216.2 | P | 198.5 | P | 201.5 | P | --- |
| 264265_at | 261 | P | 318.6 | P | 296.4 | P | 279.8 | P | --- |
| 264266_at | 195.7 | P | 280.8 | P | 269.8 | P | 242.9 | P | --- |
| 264267_at | 102.8 | P | 132.5 | P | 161.3 | P | 141.1 | P | --- |
| 264268_at | 789.6 | P | 677.7 | P | 571.1 | P | 668.4 | P | ULP1D |
| 264269_at | 6.7 | A | 1.6 | A | 1.6 | A | 2.2 | A | --- |
| 264270_at | 358.7 | P | 279.3 | P | 304.6 | P | 262.3 | P | BGLU5 |
| 264271_at | 267.4 | P | 230 | P | 243 | P | 259 | P | BGLU6 |
| 264272_at | 1.6 | A | 1.2 | A | 1.4 | A | 2.9 | A | ANAC023 |
| 264273_s_at | 3.4 | A | 1.5 | A | 2.1 | A | 3.1 | A | --- |
| 264274_at | 14.1 | A | 7.2 | A | 15.4 | A | 34.8 | A | --- |
| 264275_at | 2.2 | A | 10.4 | A | 2.6 | A | 16.7 | A | anac024 |
| 264276_at | 9.7 | A | 4.2 | A | 14.7 | A | 3 | A | --- |
| 264277_at | 136.1 | P | 240.1 | P | 193.6 | P | 247.2 | P | --- |
| 264278_at | 3.4 | A | 3.5 | A | 2.2 | A | 2.4 | A | --- |
| 264279_s_at | 4307.5 | P | 3583.7 | P | 3449.3 | P | 3172.6 | P | --- |
| 264280_at | 47.5 | P | 38 | P | 50.5 | P | 35.3 | P | BGLU46 |
| 264281_at | 18.7 | A | 8.8 | A | 2.8 | A | 30.5 | P | --- |
| 264282_at | 1.6 | A | 1.9 | A | 7.6 | A | 9.7 | A | --- |
| 264283_at | 302.6 | P | 255.3 | P | 250 | P | 268.3 | P | --- |
| 264284_at | 22.4 | A | 25.7 | A | 12.7 | A | 17.7 | A | --- |
| 264285_at | 1014.9 | P | 865.8 | P | 851.8 | P | 803.9 | P | ATG8C |
| 264286_at | 885.4 | P | 861.1 | P | 1193.5 | P | 971.6 | P | PPR336 |
| 264287_at | 73.2 | P | 71.5 | P | 103.3 | P | 53 | P | --- |
| 264288_at | 367.1 | P | 159.5 | P | 268 | P | 182.2 | P | --- |
| 264289_at | 1741.7 | P | 698.5 | P | 1646.4 | P | 1024.1 | P | --- |
| 264290_at | 186.8 | P | 186.9 | P | 211.6 | P | 222 | P | --- |
| 264291_at | 257.7 | P | 234.1 | P | 188.6 | P | 228.8 | P | --- |
| 264292_at | 201.9 | P | 233.2 | P | 129.6 | P | 218.1 | P | --- |
| 264293_at | 115.1 | P | 121.5 | P | 120.5 | P | 151.5 | P | APC6 |
| 264294_at | 32.5 | A | 62.7 | P | 39.8 | P | 57.2 | A | --- |
| 264295_at | 23.4 | A | 8.7 | A | 9.5 | A | 38.8 | A | --- |
| 264296_at | 49.9 | A | 46 | A | 56.6 | A | 65.5 | A | --- |
| 264297_at | 4.4 | A | 30.6 | A | 2.1 | A | 18 | A | --- |
| 264298_at | 186.2 | P | 265.8 | P | 192.6 | P | 231.2 | P | --- |

| | | | | | | | | | |
|---|---|---|---|---|---|---|---|---|---|
| 264299_s_at | 4848.9 | P | 3166.3 | P | 6037.5 | P | 3532.1 | P | --- |
| 264300_at | 3223.2 | P | 2383.6 | P | 3002.2 | P | 2781.3 | P | ATGGH3 |
| 264301_at | 21.7 | A | 118.6 | P | 9.9 | A | 92.5 | P | --- |
| 264302_at | 6233.6 | P | 6471.7 | P | 6623.7 | P | 5805.3 | P | VHA-A |
| 264303_s_at | 3021.8 | P | 3052.7 | P | 2760.8 | P | 3127.7 | P | UBC35 |
| 264304_at | 309.2 | P | 321.6 | P | 306.2 | P | 273.6 | P | --- |
| 264305_at | 3.1 | A | 3.4 | A | 3.3 | A | 20.9 | A | LSH7 |
| 264306_at | 732.3 | P | 994.2 | P | 783.7 | P | 883.8 | P | --- |
| 264307_at | 381.1 | P | 460.4 | P | 366.8 | P | 470.6 | P | --- |
| 264308_at | 378.8 | P | 322.6 | P | 375.8 | P | 425.6 | P | --- |
| 264309_at | 1097.5 | P | 1309.6 | P | 1148.9 | P | 1250.9 | P | --- |
| 264310_at | 250.3 | P | 186.3 | P | 264.7 | P | 240.5 | P | --- |
| 264311_at | 1.8 | A | 1.6 | A | 2.5 | A | 2.1 | A | --- |
| 264312_at | 16.4 | A | 12.6 | A | 9.7 | A | 13.1 | A | --- |
| 264313_at | 1470.7 | P | 2589.7 | P | 1857.1 | P | 2464 | P | BCA4 |
| 264314_at | 74.5 | P | 104.2 | P | 70.6 | A | 89.7 | P | --- |
| 264315_at | 1873.4 | P | 2560.7 | P | 2437.3 | P | 2837.3 | P | --- |
| 264316_at | 933.2 | P | 1476 | P | 900.7 | P | 1215.1 | P | ENT1,AT |
| 264317_at | 2152.5 | P | 2424.4 | P | 2998.3 | P | 2524.4 | P | SPDS2 |
| 264318_at | 100.1 | P | 284.4 | P | 102.3 | A | 316.5 | P | KCS2 |
| 264319_at | 223.1 | P | 310.2 | P | 303.5 | P | 309.6 | P | SDD1 |
| 264320_at | 28.5 | A | 29.2 | A | 56 | A | 35.7 | A | --- |
| 264321_at | 254.9 | P | 317.8 | P | 205 | P | 341 | P | --- |
| 264322_at | 561 | P | 647.4 | P | 788.9 | P | 727.9 | P | --- |
| 264323_at | 4.9 | A | 1.2 | A | 5.1 | A | 2.9 | A | --- |
| 264324_at | 41.8 | P | 89 | P | 59.8 | P | 101.6 | P | XIB |
| 264325_at | 35.1 | P | 48.6 | P | 62.9 | P | 36.2 | P | --- |
| 264326_at | 337.5 | P | 337.6 | P | 392.7 | P | 299.2 | P | --- |
| 264327_at | 2240.9 | P | 2919.9 | P | 2894.4 | P | 2657.5 | P | EIF2 GAMMA |
| 264328_at | 147.2 | P | 111.7 | P | 95.6 | A | 113.7 | M | IAA10 |
| 264329_at | 827.5 | P | 921 | P | 966.1 | P | 845.7 | P | --- |
| 264330_at | 722.9 | P | 775.4 | P | 762.2 | P | 649 | P | ATMRP5 |
| 264331_at | 246.5 | P | 258.6 | P | 256.6 | P | 240.6 | P | --- |
| 264332_at | 75.4 | A | 81.2 | A | 180.2 | A | 85.5 | A | --- |
| 264333_at | 2.3 | A | 17.8 | A | 12 | A | 1.2 | A | AtTLP4 |
| 264334_at | 2.4 | A | 1.2 | A | 2 | A | 4.7 | A | CPK19 |
| 264335_s_at | 1658.7 | P | 1619.1 | P | 1183.9 | P | 1526.9 | P | UPL1 |
| 264336_at | 151.7 | P | 144.2 | P | 166.1 | P | 151.6 | P | --- |
| 264337_at | 270.2 | P | 289.2 | P | 359.3 | P | 241.9 | P | --- |
| 264338_at | 339.9 | P | 250.4 | P | 275.1 | P | 204.5 | P | KUP6 |
| 264339_at | 165.3 | P | 193.2 | P | 86.4 | P | 125.4 | P | TPS8 |
| 264340_at | 294 | P | 418 | P | 355.8 | P | 479.8 | P | --- |
| 264341_at | 29.8 | P | 40.6 | P | 48.4 | P | 37.8 | P | --- |
| 264342_at | 22.5 | P | 61.2 | P | 25 | A | 64.3 | P | --- |
| 264343_at | 282.2 | P | 553.9 | P | 355.3 | P | 468.3 | P | --- |
| 264344_at | 9609.2 | P | 9379.7 | P | 10317.3 | P | 9094.8 | P | --- |
| 264345_at | 4 | A | 8.2 | A | 6.2 | A | 7.6 | A | --- |
| 264346_at | 18.3 | A | 10.2 | A | 3.3 | A | 9.5 | A | --- |
| 264347_at | 6.8 | A | 1.2 | A | 9.5 | A | 2.7 | A | LRX1 |
| 264348_at | 5648.8 | P | 3128.2 | P | 6124.2 | P | 3866.3 | P | NRT1.1 |
| 264349_at | 1343 | P | 1605.3 | P | 1758 | P | 1419.3 | P | --- |
| 264350_at | 472.1 | P | 509.9 | P | 644.3 | P | 531.8 | P | SRS |
| 264351_at | 233.1 | P | 357.7 | P | 181.3 | P | 320.2 | P | --- |
| 264352_at | 129.4 | P | 90.6 | P | 120.8 | P | 107.4 | P | --- |
| 264353_at | 143.2 | P | 173.9 | P | 114.3 | P | 132.5 | P | --- |
| 264354_s_at | 60.8 | P | 69.1 | P | 57.6 | P | 46.3 | P | --- |
| 264355_at | 475.6 | P | 353.1 | P | 511 | P | 387 | P | --- |
| 264356_at | 181.1 | P | 189.6 | P | 201.3 | P | 200.2 | P | UVH6 |
| 264357_at | 319.2 | P | 242.4 | P | 376.7 | P | 273.6 | P | ATRRP4 |
| 264358_at | 55.5 | M | 86.4 | P | 83.2 | P | 44.7 | A | --- |
| 264359_at | 59.3 | A | 53.1 | A | 56.1 | A | 33.4 | A | --- |
| 264360_at | 1028.8 | P | 1387.5 | P | 1072.7 | P | 1495.3 | P | DBE1 |
| 264361_at | 65 | P | 64.5 | P | 85.3 | P | 72.9 | P | --- |
| 264362_at | 352.9 | P | 393 | P | 269.4 | P | 335.1 | P | --- |
| 264363_at | 28.7 | A | 45.7 | A | 8.7 | A | 37.8 | A | --- |
| 264364_at | 1097.9 | P | 1190.6 | P | 1243.9 | P | 1302 | P | --- |
| 264365_s_at | 1597.7 | P | 1141.6 | P | 851.2 | P | 982.7 | P | --- |
| 264366_at | 404.5 | P | 469.8 | P | 381.3 | P | 401.3 | P | --- |
| 264367_at | 452.6 | P | 605.7 | P | 625.6 | P | 643.6 | P | --- |
| 264368_at | 384.7 | P | 428.6 | P | 401.6 | P | 521.1 | P | --- |
| 264369_at | 27.4 | P | 51.7 | A | 53.8 | A | 76.8 | P | --- |
| 264370_at | 31.7 | A | 33.2 | A | 55.2 | A | 34.9 | A | --- |
| 264371_at | 8589.1 | P | 12100.3 | P | 8672.1 | P | 11814.7 | P | ELP |
| 264372_at | 7233.2 | P | 6882.4 | P | 5900.2 | P | 7799.8 | P | ATGLX1 |
| 264373_at | 1130.4 | P | 1295.3 | P | 968.3 | P | 1365.3 | P | SEC22 |
| 264374_at | 344.9 | P | 452.9 | P | 400.5 | P | 411.6 | P | ARR12 |
| 264375_at | 23.8 | A | 4.6 | A | 8.6 | A | 3.6 | A | CIPK16 |
| 264376_at | 1078.2 | P | 1153.8 | P | 932 | P | 1016.4 | P | --- |
| 264377_at | 517.9 | P | 781.2 | P | 724.3 | P | 1107.9 | P | --- |
| 264378_at | 78.4 | A | 120.4 | P | 78.4 | M | 128.2 | P | --- |
| 264379_at | 153.7 | P | 138.9 | P | 136.7 | P | 74 | P | --- |
| 264380_at | 277 | P | 307 | P | 207.9 | P | 297.5 | P | --- |
| 264381_at | 213.4 | P | 237.9 | P | 183.3 | P | 276 | P | --- |
| 264382_at | 2232.9 | P | 1998.7 | P | 1786.1 | P | 1957.5 | P | SDF2 |
| 264383_at | 4209.4 | P | 3776.5 | P | 4027.9 | P | 3416.3 | P | ATGPX1 |
| 264384_at | 387.6 | P | 457 | P | 421.5 | P | 491.1 | P | PKL |
| 264385_at | 271.8 | P | 185.8 | P | 300.9 | P | 216.7 | P | --- |
| 264386_at | 1226 | P | 1046.6 | P | 1138.8 | P | 1250.1 | P | --- |
| 264387_at | 9.2 | A | 11 | A | 30.4 | A | 29.5 | A | --- |
| 264388_at | 6.7 | A | 3.9 | A | 17.1 | A | 14.5 | A | --- |
| 264389_at | 171.5 | P | 372.9 | P | 160.9 | P | 269.9 | P | --- |
| 264390_at | 87.7 | P | 182.8 | P | 75.7 | P | 168.3 | P | --- |
| 264391_at | 1.4 | A | 1.2 | A | 1.9 | A | 2.4 | A | --- |
| 264392_at | 92.7 | P | 132.7 | P | 126.7 | P | 134.1 | P | --- |
| 264393_at | 42.5 | M | 76 | P | 74.4 | M | 61.8 | P | --- |
| 264394_at | 7969.9 | P | 8442.7 | P | 6380.1 | P | 7321.2 | P | --- |
| 264395_at | 12.2 | A | 20.2 | A | 27.7 | A | 46.2 | A | --- |
| 264396_at | 748.4 | P | 963.9 | P | 765.6 | P | 855.5 | P | --- |
| 264397_at | 631.1 | P | 868.9 | P | 906.1 | P | 910.3 | P | --- |
| 264398_at | 1635.2 | P | 1543.1 | P | 1372.4 | P | 1639.8 | P | --- |
| 264399_at | 1811.2 | P | 1727 | P | 1069.7 | P | 1686.8 | P | --- |
| 264400_at | 3294.5 | P | 2556.5 | P | 3530.5 | P | 4532.2 | P | GPT2 |
| 264401_at | 3.9 | A | 25 | A | 2.9 | A | 6.5 | A | BAN |
| 264402_at | 154.1 | P | 202.9 | P | 282.4 | P | 191 | P | CLPB4 |
| 264403_at | 12.3 | A | 15 | A | 28 | A | 2.6 | A | --- |
| 264404_at | 22.3 | A | 4.1 | A | 4.8 | A | 13.9 | A | CYP82F1 |
| 264405_at | 105.9 | P | 120 | P | 50.7 | P | 125.1 | P | ATMPK1 |
| 264406_at | 639.1 | P | 705.1 | P | 638.1 | P | 658 | P | ADL6 |
| 264407_at | 336.9 | P | 369 | P | 355.2 | P | 362 | P | --- |

| | | | | | | | | | |
|---|---|---|---|---|---|---|---|---|---|
| 264408_at | 178.3 | P | 141.7 | P | 140 | P | 116.3 | P | FRS11 |
| 264409_s_at | 42.3 | A | 1.2 | A | 1.6 | A | 1.9 | A | --- |
| 264410_s_at | 19.6 | A | 26 | A | 37.6 | A | 19.7 | A | --- |
| 264411_at | 3 | A | 3.2 | A | 24.7 | A | 3 | A | --- |
| 264412_at | 1.2 | A | 1.6 | A | 2.3 | A | 7.2 | A | --- |
| 264413_s_at | 11.2 | A | 3.1 | A | 10 | A | 1.4 | A | --- |
| 264414_s_at | 3.6 | A | 16.2 | A | 3 | A | 19 | A | --- |
| 264415_at | 146.7 | P | 5.3 | A | 91.1 | A | 3.7 | A | RAP2.6 |
| 264416_at | 3.1 | A | 7.9 | A | 12 | A | 17.4 | A | --- |
| 264417_at | 8.9 | A | 5.7 | A | 35.8 | A | 18.3 | A | --- |
| 264418_at | 812.4 | P | 1149.7 | P | 1041.2 | P | 1039.6 | P | --- |
| 264419_s_at | 3619.3 | P | 3471.9 | P | 4578.6 | P | 3897.9 | P | --- |
| 264420_at | 2.2 | A | 3.3 | A | 3.6 | A | 5.1 | A | --- |
| 264421_at | 10793.7 | P | 9190.4 | P | 11388.5 | P | 9800.7 | P | ARP1 |
| 264422_at | 746.7 | P | 870.8 | P | 675.2 | P | 862.1 | P | LCV2 |
| 264423_at | 168.8 | P | 223.2 | P | 188.5 | P | 219.8 | P | --- |
| 264424_at | 3199 | P | 2954.9 | P | 4499.6 | P | 3608.2 | P | --- |
| 264425_at | 24.4 | A | 10.2 | A | 15.4 | A | 1.3 | A | --- |
| 264426_at | 21.8 | A | 30.5 | P | 42.4 | A | 34.9 | P | --- |
| 264427_at | 1138.9 | P | 1162.2 | P | 1140.4 | P | 1107.3 | P | --- |
| 264428_at | 3736.4 | P | 4057.6 | P | 3824.2 | P | 3698 | P | --- |
| 264429_at | 356.8 | P | 270 | P | 206.1 | P | 341 | P | --- |
| 264430_at | 3 | A | 3 | A | 3.6 | A | 2.5 | A | TPS14 |
| 264431_at | 23.4 | A | 10.2 | A | 50.1 | A | 18.6 | A | --- |
| 264432_at | 17.3 | A | 14.2 | A | 40.5 | A | 32.5 | P | --- |
| 264433_at | 55.7 | P | 28 | P | 9.5 | A | 26 | P | BGLU45 |
| 264434_at | 127 | P | 492.4 | P | 62.8 | A | 207.4 | P | --- |
| 264435_at | 566.8 | P | 620.3 | P | 556.3 | P | 592.5 | P | ATGSTU18 |
| 264436_at | 1014.5 | P | 694.4 | P | 806.8 | P | 921.5 | P | ERD9 |
| 264437_at | 582.1 | P | 702.3 | P | 602.2 | P | 576.1 | P | --- |
| 264438_at | 4422.7 | P | 4144.8 | P | 5203.1 | P | 3892.5 | P | --- |
| 264439_at | 4821.5 | P | 5022 | P | 5545.3 | P | 5204.6 | P | APT1 |
| 264440_at | 702.8 | P | 777.9 | P | 782.3 | P | 756.8 | P | --- |
| 264441_at | 1269.2 | P | 1138.9 | P | 1288.1 | P | 1264.5 | P | --- |
| 264442_at | 768.7 | P | 1054.3 | P | 1143.8 | P | 1212.3 | P | --- |
| 264443_at | 908.1 | P | 937.1 | P | 869.7 | P | 1042.6 | P | --- |
| 264444_at | 102.6 | P | 93.9 | P | 89 | P | 94.1 | P | --- |
| 264445_at | 1053.5 | P | 1028.7 | P | 688 | P | 799.6 | P | --- |
| 264446_at | 1484.6 | P | 1894.8 | P | 1603.4 | P | 1889.2 | P | NTF2A |
| 264447_at | 822.8 | P | 501.4 | P | 686.6 | P | 483.2 | P | --- |
| 264448_at | 769.7 | P | 664.4 | P | 534 | P | 579.5 | P | AHK3 |
| 264449_at | 219.5 | P | 382.3 | P | 321.7 | P | 368.7 | P | NPGR1 |
| 264450_s_at | 160.2 | P | 122.4 | P | 166 | P | 118.7 | P | SNL6 |
| 264451_s_at | 17.1 | A | 2 | A | 3.6 | A | 22.4 | A | --- |
| 264452_at | 261.2 | P | 237.6 | P | 230.6 | P | 237.7 | P | GRP23 |
| 264453_at | 24.9 | A | 30.7 | A | 13.2 | A | 3.5 | A | --- |
| 264454_at | 127.5 | P | 94.9 | P | 92.6 | P | 110.9 | P | --- |
| 264455_at | 72.2 | P | 47.2 | A | 65.8 | A | 65.7 | A | --- |
| 264456_at | 443.4 | P | 355.7 | P | 309.2 | P | 297 | P | --- |
| 264457_at | 39 | P | 4.7 | A | 35.7 | A | 5.4 | A | --- |
| 264458_at | 1092.5 | P | 785.3 | P | 550.6 | P | 763.9 | P | --- |
| 264459_at | 46.2 | A | 76.6 | A | 72.5 | A | 69.9 | A | --- |
| 264460_at | 824.9 | P | 554 | P | 890.7 | P | 576.8 | P | ATNFXL1 |
| 264461_s_at | 6.2 | A | 1.5 | A | 11.7 | A | 8.2 | A | --- |
| 264462_at | 2098.5 | P | 2359.7 | P | 1791.2 | P | 2508.2 | P | WLIM1 |
| 264463_at | 253.1 | P | 379.7 | P | 248.4 | P | 343 | P | --- |
| 264464_at | 415.3 | P | 484.5 | P | 543.4 | P | 394.4 | P | --- |
| 264465_at | 345.5 | P | 370 | P | 382.4 | P | 350.9 | P | ASK18 |
| 264466_at | 119.3 | P | 150.3 | P | 212.4 | P | 132.2 | P | --- |
| 264467_at | 169.5 | P | 118.3 | P | 151.6 | P | 134.9 | P | --- |
| 264468_at | 673.4 | P | 736.5 | P | 753.8 | P | 651.9 | P | --- |
| 264469_at | 53.5 | A | 28.1 | A | 75.4 | P | 32.7 | A | LBD40 |
| 264470_at | 16 | A | 55.9 | P | 56.8 | A | 33.6 | A | CYP735A2 |
| 264471_at | 290.9 | P | 137.9 | P | 253.5 | P | 151.2 | P | --- |
| 264472_at | 254 | P | 274.7 | P | 198.3 | P | 237 | P | SWEETIE |
| 264473_at | 57.3 | A | 76.6 | P | 76.8 | A | 78.3 | A | --- |
| 264474_s_at | 27095.7 | P | 26957.6 | P | 29186.9 | P | 28913.7 | P | RBCS1A |
| 264475_s_at | 201.4 | P | 176.7 | P | 172.3 | P | 181.6 | P | --- |
| 264476_at | 820.7 | P | 635.4 | P | 694.7 | P | 767.8 | P | PGSIP2 |
| 264477_at | 1.5 | A | 13.4 | A | 3.5 | A | 11.3 | A | --- |
| 264478_at | 39.9 | P | 41.5 | P | 93.9 | P | 50.3 | P | --- |
| 264479_at | 305.3 | P | 164.2 | P | 170.5 | P | 217.4 | P | --- |
| 264480_at | 65.8 | P | 78.3 | P | 115.7 | P | 84.8 | P | EFS |
| 264481_at | 3.9 | A | 36.5 | A | 6.4 | A | 49.5 | A | --- |
| 264482_at | 236.7 | P | 262.4 | P | 233.7 | P | 212.8 | P | --- |
| 264483_at | 38.1 | A | 29 | A | 51.1 | A | 45.8 | M | --- |
| 264484_at | 617.6 | P | 490.4 | P | 555.3 | P | 543.6 | P | --- |
| 264485_at | 294 | P | 214.8 | P | 209.9 | P | 236.4 | P | --- |
| 264486_at | 1106.8 | P | 932.5 | P | 1210.1 | P | 988.2 | P | --- |
| 264487_at | 238.2 | P | 248.4 | P | 223.6 | P | 221.7 | P | VPS45 |
| 264488_s_at | 1786.9 | P | 1918.2 | P | 1480 | P | 1779.3 | P | --- |
| 264489_at | 125.2 | P | 106.7 | P | 95.1 | P | 105.6 | P | --- |
| 264490_at | 794.2 | P | 626.9 | P | 924.5 | P | 740.9 | P | TOM20-2 |
| 264491_at | 185 | P | 178.4 | P | 268.3 | P | 186.1 | P | --- |
| 264492_at | 241.9 | M | 184.2 | M | 127.7 | A | 154.4 | P | --- |
| 264493_at | 208 | P | 184.9 | P | 232.4 | P | 209.1 | P | GUT2 |
| 264494_at | 12.2 | A | 4.4 | A | 5.1 | A | 4.6 | A | --- |
| 264495_at | 2.8 | A | 2.3 | A | 3.2 | A | 3.4 | A | RIC2 |
| 264496_at | 23.3 | A | 23.5 | A | 20.6 | A | 59.6 | A | --- |
| 264497_at | 19.9 | A | 35.4 | P | 64.4 | P | 32.8 | P | ATPUP4 |
| 264498_at | 218 | P | 356.6 | P | 389.5 | P | 372.2 | P | DIS2 |
| 264499_at | 5.4 | A | 9.5 | A | 14.7 | A | 23.7 | A | --- |
| 264500_at | 8.7 | A | 0.4 | A | 0.9 | A | 0.7 | A | --- |
| 264501_at | 39.3 | P | 131.7 | P | 92.9 | P | 82.6 | P | --- |
| 264502_at | 13.2 | A | 2.3 | A | 5.8 | A | 3 | A | --- |
| 264503_at | 27.1 | P | 25.7 | A | 8.3 | A | 45.9 | P | --- |
| 264504_at | 1735.5 | P | 1544 | P | 1455.9 | P | 1371.8 | P | ACLA-3 |
| 264505_at | 43.8 | P | 43.8 | A | 32.1 | A | 33.9 | P | --- |
| 264506_at | 1382 | P | 1149.3 | P | 788.8 | P | 1081.8 | P | GLP5 |
| 264507_at | 460.1 | P | 404.9 | P | 328.3 | P | 369.7 | P | NIMIN-3 |
| 264508_at | 1014.3 | P | 895.2 | P | 963.7 | P | 843.9 | P | PHYA |
| 264509_at | 429 | P | 323.1 | P | 486.2 | P | 283.4 | P | --- |
| 264510_at | 126.8 | P | 229.5 | P | 145.3 | P | 158.6 | P | PIF3 |
| 264511_at | 15.1 | A | 65.1 | A | 11.7 | A | 55.1 | M | AtGolS3 |
| 264512_at | 55.8 | P | 50.6 | A | 40.9 | A | 77.8 | A | --- |
| 264513_at | 281.4 | P | 153.2 | P | 241.3 | P | 146.8 | P | G6PD4 |
| 264514_at | 1170.1 | P | 264.6 | P | 347.4 | P | 405.6 | P | --- |
| 264515_at | 372.5 | P | 588 | P | 339.5 | P | 487.7 | P | --- |
| 264516_at | 167.6 | P | 199.3 | P | 202.7 | P | 230.9 | P | --- |

| | | | | | | | | | |
|---|---|---|---|---|---|---|---|---|---|
| 264517_at | 102.5 | P | 161.1 | P | 119.4 | P | 155.9 | P | --- |
| 264518_at | 108.2 | P | 121.9 | P | 131.2 | P | 164.3 | P | --- |
| 264519_at | 22.1 | A | 25.3 | A | 73.1 | A | 23.3 | P | --- |
| 264520_at | 31.3 | A | 19.1 | A | 23.1 | A | 22.6 | A | AAP8 |
| 264521_at | 197.5 | P | 229.2 | P | 196.8 | P | 260.9 | P | --- |
| 264522_at | 83.6 | P | 111.8 | P | 69.6 | A | 86.9 | P | --- |
| 264523_at | 340 | P | 428.7 | P | 303.6 | P | 425.2 | P | ERG28 |
| 264524_at | 190.2 | P | 231.2 | P | 191.8 | P | 201.3 | P | ATBCAT-2 |
| 264525_at | 212.1 | P | 295.6 | P | 192.2 | P | 253.6 | P | ATBCAT-1 |
| 264526_at | 434.7 | P | 564.1 | P | 350.4 | P | 459.9 | P | ECA3 |
| 264527_at | 1.8 | A | 2.1 | A | 3.6 | A | 2.4 | A | --- |
| 264528_at | 103.7 | P | 125.1 | P | 135.3 | P | 112.6 | P | --- |
| 264529_at | 365.4 | P | 381.4 | P | 461.1 | P | 355.6 | P | --- |
| 264530_at | 27 | A | 3.8 | A | 24.4 | A | 7.7 | A | --- |
| 264531_at | 6.5 | A | 2.5 | A | 20.3 | A | 49.7 | A | --- |
| 264532_at | 56 | P | 61.5 | P | 37.2 | A | 61.7 | M | AtSIP1 |
| 264533_s_at | 295.3 | P | 368.6 | P | 271.2 | P | 355.4 | P | CAX5 |
| 264534_at | 18.6 | A | 26.4 | A | 42.5 | A | 7.2 | A | --- |
| 264535_at | 210.7 | P | 277 | P | 325.2 | P | 340.3 | P | --- |
| 264536_at | 7.2 | A | 24.8 | A | 38.5 | A | 2.8 | A | PRA1.G1 |
| 264537_at | 64.6 | P | 91.5 | P | 84.1 | P | 51.6 | P | BRL1 |
| 264538_at | 7.6 | A | 6.8 | A | 3.7 | A | 16.5 | A | WRKY10 |
| 264539_at | 128.8 | P | 139 | P | 118 | P | 171.9 | P | --- |
| 264540_at | 99.4 | P | 134.6 | P | 90.4 | P | 93.5 | P | --- |
| 264541_at | 4.5 | A | 2.9 | A | 2.4 | A | 2.5 | A | --- |
| 264542_at | 215.1 | P | 187.3 | P | 168.8 | P | 196.3 | P | --- |
| 264543_at | 34.7 | P | 21.6 | P | 13.4 | A | 34.9 | P | --- |
| 264544_s_at | 23.8 | A | 19.4 | A | 14.2 | A | 20 | A | --- |
| 264545_at | 13829.4 | P | 15989.8 | P | 16323.9 | P | 15636.4 | P | PSAG |
| 264546_at | 724.8 | P | 761 | P | 739.9 | P | 775.7 | P | --- |
| 264547_at | 190.7 | P | 219.3 | P | 202.1 | P | 251.8 | P | CLC-F |
| 264548_at | 581.7 | P | 846.4 | P | 604.7 | P | 787.4 | P | --- |
| 264549_at | 115.3 | P | 96.9 | P | 105.2 | P | 151.3 | P | --- |
| 264550_at | 53.9 | P | 92.7 | P | 80.1 | P | 72.8 | P | --- |
| 264551_at | 63.1 | P | 80.9 | P | 133.2 | P | 68.1 | P | --- |
| 264552_at | 7.3 | A | 47.1 | A | 89.3 | A | 63 | A | --- |
| 264553_s_at | 428 | P | 485.2 | P | 340.1 | P | 540.2 | P | --- |
| 264554_at | 40.5 | A | 24.2 | A | 62.3 | A | 40.8 | A | --- |
| 264555_at | 2.3 | A | 3.1 | A | 5.9 | A | 3.4 | A | --- |
| 264556_at | 34.7 | A | 21.9 | A | 44.6 | A | 37.1 | M | MYB61 |
| 264557_at | 30.4 | A | 36.7 | A | 13.3 | A | 33.1 | A | --- |
| 264558_at | 33.6 | P | 32.6 | A | 67.4 | P | 33.8 | A | --- |
| 264559_at | 26 | A | 27.4 | A | 10.6 | A | 22.6 | A | --- |
| 264560_at | 129.2 | P | 88.6 | A | 113 | A | 89 | P | --- |
| 264561_at | 608.7 | P | 348.8 | P | 540.7 | P | 379.2 | P | --- |
| 264562_at | 176.2 | P | 241.5 | P | 148.8 | P | 202.3 | P | --- |
| 264563_s_at | 127.1 | P | 137.2 | P | 159.8 | P | 140.9 | P | --- |
| 264564_at | 10.6 | A | 15.5 | A | 3 | A | 2.6 | A | --- |
| 264565_at | 8.4 | A | 23.5 | A | 1.7 | A | 3.3 | A | --- |
| 264566_at | 346.8 | P | 377.6 | P | 407.1 | P | 337.4 | P | --- |
| 264567_s_at | 47.8 | P | 39.9 | P | 30.3 | A | 26.3 | A | --- |
| 264568_at | 411.6 | P | 470.6 | P | 496.2 | P | 480.4 | P | --- |
| 264569_at | 31.3 | P | 32.8 | M | 59.9 | A | 57.4 | P | --- |
| 264570_at | 384.4 | P | 496.8 | P | 490.7 | P | 474.3 | P | --- |
| 264571_at | 2.3 | A | 27.7 | A | 1.5 | A | 2.8 | A | --- |
| 264572_at | 393.7 | P | 277.3 | P | 266 | P | 393.6 | P | --- |
| 264573_at | 72 | P | 54.1 | P | 70.7 | P | 72.5 | P | --- |
| 264574_at | 199.6 | P | 200.5 | P | 138.2 | P | 203.6 | P | ZIP5 |
| 264575_at | 4786.2 | P | 6469.5 | P | 5420.9 | P | 5964 | P | emb2394 |
| 264576_at | 308.3 | P | 267.7 | P | 332.6 | P | 267.4 | P | --- |
| 264577_at | 31.1 | A | 55.6 | P | 48.5 | M | 69.2 | P | RCI3 |
| 264578_at | 6.7 | A | 40 | A | 26.9 | A | 35.5 | A | --- |
| 264579_at | 536.3 | P | 514.9 | P | 562.1 | P | 589.2 | P | --- |
| 264580_at | 401.1 | P | 453.3 | P | 177.4 | P | 245.5 | P | --- |
| 264581_at | 202.8 | P | 267.8 | P | 197.4 | P | 275.6 | P | --- |
| 264582_at | 274.1 | P | 317.4 | P | 310.6 | P | 312.7 | P | HDG2 |
| 264583_at | 187.5 | P | 123.1 | P | 148.3 | P | 206.9 | P | --- |
| 264584_at | 1041.3 | P | 1164.4 | P | 834.3 | P | 1206.3 | P | --- |
| 264585_at | 894.2 | P | 921 | P | 921.7 | P | 1007.2 | P | AXR1 |
| 264586_at | 159.4 | P | 184.1 | P | 163.9 | P | 213.2 | P | CYP88A3 |
| 264587_at | 605.8 | P | 613.2 | P | 537.4 | P | 501 | P | ATGLR3.4 |
| 264588_at | 500.2 | P | 558.4 | P | 375.8 | P | 471 | P | --- |
| 264589_at | 201.9 | P | 78.7 | A | 191.1 | A | 122.9 | A | --- |
| 264590_at | 341.6 | P | 944.8 | P | 427.1 | P | 736.8 | P | --- |
| 264591_at | 655.4 | P | 439.3 | P | 776.8 | P | 491.4 | P | --- |
| 264592_at | 1079.9 | P | 1133.4 | P | 1106.1 | P | 1008.4 | P | --- |
| 264593_at | 492.5 | P | 577.8 | P | 534.5 | P | 656.1 | P | ARAC1 |
| 264594_at | 358.5 | P | 312.7 | P | 386.2 | P | 297.4 | P | ATSERAT3;1 |
| 264595_at | 2002.1 | P | 2313.9 | P | 2060.9 | P | 2075.9 | P | VAMP721 |
| 264596_at | 9.4 | A | 16.6 | A | 2 | A | 1.9 | A | --- |
| 264597_at | 449.6 | P | 540.3 | P | 520.8 | P | 634.5 | P | --- |
| 264598_at | 38.2 | A | 56.2 | A | 77.2 | A | 48 | A | --- |
| 264599_at | 17.5 | A | 20.3 | A | 39.8 | A | 22.7 | A | XIA |
| 264600_at | 84.3 | P | 92.7 | P | 133 | P | 113.8 | P | --- |
| 264601_at | 27.8 | A | 23.1 | A | 68.7 | A | 17.5 | A | --- |
| 264602_at | 33.6 | P | 32.7 | P | 45.1 | P | 44.4 | P | --- |
| 264603_at | 11.6 | A | 13.3 | A | 16.4 | A | 1.3 | A | --- |
| 264604_at | 79.6 | P | 83 | P | 44 | A | 62.5 | P | --- |
| 264605_at | 192.2 | P | 201.8 | P | 275.3 | P | 235.9 | P | IAA12 |
| 264606_at | 14.5 | A | 83 | A | 60.5 | A | 76.5 | A | --- |
| 264607_at | 2986.5 | P | 3560.5 | P | 3276.1 | P | 3221.4 | P | KAB1 |
| 264608_at | 754.4 | P | 740.4 | P | 939.3 | P | 798.1 | P | PKT4 |
| 264609_at | 372.1 | P | 656.7 | P | 365.4 | P | 529.7 | P | --- |
| 264610_at | 2.4 | A | 9 | A | 1.5 | A | 4.7 | A | --- |
| 264611_at | 794.4 | P | 1715.8 | P | 653.8 | P | 1690.5 | P | --- |
| 264612_at | 24 | A | 39.5 | A | 50.5 | A | 43.5 | A | --- |
| 264613_at | 1023.3 | P | 1183.9 | P | 977.9 | P | 1131.5 | P | LIP2 |
| 264614_s_at | 29 | A | 10.4 | A | 23.3 | A | 13.2 | A | --- |
| 264615_at | 552.2 | P | 544.9 | P | 739.3 | P | 642.5 | P | --- |
| 264616_at | 44.1 | A | 65.4 | A | 54.9 | M | 57.1 | P | --- |
| 264617_at | 16.6 | A | 52.8 | P | 15 | A | 54.8 | P | --- |
| 264618_at | 9.5 | A | 22.1 | A | 23.6 | P | 2.2 | A | --- |
| 264619_at | 303.7 | P | 356.2 | P | 267.1 | P | 312.2 | P | --- |
| 264620_at | 16 | A | 40.7 | A | 31.9 | A | 44.3 | A | ATBZIP27 |
| 264621_at | 313 | P | 248.8 | P | 288.3 | P | 284.9 | P | --- |
| 264622_at | 413.1 | P | 468 | P | 374.8 | P | 447.6 | P | VPS35A |
| 264623_at | 32 | A | 44.7 | A | 60.4 | A | 65.4 | A | ANP1 |
| 264624_at | 2490 | P | 2713.8 | P | 2557 | P | 2256.9 | P | ERD6 |
| 264625_at | 494.8 | P | 610.3 | P | 584.8 | P | 688.7 | P | SNF4 |

| | | | | | | | | | |
|---|---|---|---|---|---|---|---|---|---|
| 264626_at | 123.9 | P | 102.1 | P | 206.1 | P | 111 | P | AS2 |
| 264627_at | 266.4 | P | 216.2 | P | 218.2 | P | 165 | P | ECI1 |
| 264628_at | 329.8 | P | 456.9 | P | 406 | P | 387.1 | P | --- |
| 264629_at | 424.7 | P | 477.8 | P | 323.3 | P | 411.1 | P | --- |
| 264630_at | 53 | P | 143.1 | P | 114.3 | P | 123.9 | P | FAS1 |
| 264631_at | 11 | A | 20.4 | A | 14.7 | A | 12.7 | P | DegP3 |
| 264632_at | 1.6 | A | 18.8 | A | 4.9 | A | 3.8 | A | DegP4 |
| 264633_at | 816.9 | P | 691 | P | 1038.3 | P | 759.9 | P | SMP1 |
| 264634_at | 15.9 | A | 4.2 | A | 9.9 | A | 1.9 | A | CYP702A1 |
| 264635_at | 430.6 | P | 799.6 | P | 317.9 | P | 714.6 | P | --- |
| 264636_at | 1340.6 | P | 2541.1 | P | 1588 | P | 2206 | P | --- |
| 264637_at | 228.9 | P | 234.1 | P | 298.1 | P | 358.1 | P | --- |
| 264638_at | 23.1 | A | 38 | A | 63.4 | A | 29.5 | A | FT |
| 264639_at | 376 | P | 414.3 | P | 368.2 | P | 452.5 | P | UCH2 |
| 264640_at | 29.4 | A | 50.4 | A | 86.8 | A | 59.6 | A | ATEXPB2 |
| 264641_at | 3488.5 | P | 3344 | P | 3272.2 | P | 3341.1 | P | --- |
| 264642_at | 7.7 | A | 1.7 | A | 3.1 | A | 5 | A | NF-YB4 |
| 264643_at | 0.7 | A | 2.9 | A | 1.7 | A | 1.9 | A | PGSIP5 |
| 264644_at | 211.8 | P | 262 | P | 232 | P | 230.8 | P | CAX11 |
| 264645_at | 717.6 | P | 288.4 | P | 484.3 | P | 281.9 | P | --- |
| 264646_at | 10.1 | A | 3.7 | A | 2.7 | A | 19.7 | A | BON3 |
| 264647_at | 0.8 | A | 1.8 | A | 3.2 | A | 1.3 | A | ATRBOHB |
| 264648_at | 110.9 | P | 255.9 | P | 40 | P | 257.2 | P | BIP3 |
| 264649_at | 557.8 | P | 686.7 | P | 536.2 | P | 605.4 | P | --- |
| 264650_at | 746.8 | P | 834.2 | P | 998.5 | P | 861.2 | P | NF-YC9 |
| 264651_at | 1595 | P | 1644 | P | 1681.8 | P | 1605.4 | P | H2AXA |
| 264652_at | 391.3 | P | 196.5 | P | 394.9 | P | 210.2 | P | --- |
| 264653_at | 1973.4 | P | 1777.7 | P | 1728.9 | P | 1675.1 | P | AMI1 |
| 264654_s_at | 214.3 | P | 247.3 | P | 145.8 | P | 212.1 | P | --- |
| 264655_at | 4129.1 | P | 5224.2 | P | 3642.1 | P | 4460.8 | P | SRC2 |
| 264656_at | 654.7 | P | 733.8 | P | 462.1 | P | 587.7 | P | --- |
| 264657_at | 417.7 | P | 446.7 | P | 410.7 | P | 513.7 | P | RPT5B |
| 264658_at | 463.6 | P | 338.6 | P | 301.9 | P | 297.9 | P | --- |
| 264659_at | 3.1 | A | 2.8 | A | 7.3 | A | 0.8 | A | ATOPT2 |
| 264660_at | 240 | P | 206 | P | 244.9 | P | 233.5 | P | HEMA2 |
| 264661_at | 20.1 | A | 26.3 | A | 3.6 | A | 38.8 | A | --- |
| 264662_at | 250.6 | P | 238 | P | 369 | P | 295.7 | P | SUT4 |
| 264663_at | 4796.7 | P | 2962.6 | P | 3260.8 | P | 2717.8 | P | LRR XI-23 |
| 264664_at | 681 | P | 606.3 | P | 872.2 | P | 714.5 | P | --- |
| 264665_at | 593.2 | P | 576.2 | P | 658.5 | P | 639.9 | P | --- |
| 264666_at | 92.8 | P | 79.4 | P | 101.3 | P | 104.8 | P | --- |
| 264667_s_at | 580.7 | P | 631.6 | P | 565.8 | P | 670.5 | P | ZW18 |
| 264668_at | 2104.6 | P | 2492.4 | P | 2264.5 | P | 2560.4 | P | --- |
| 264669_at | 330.2 | P | 551.2 | P | 346.2 | P | 509.7 | P | ATRAB11C |
| 264670_s_at | 12219.5 | P | 10419 | P | 12031.9 | P | 10505.3 | P | --- |
| 264671_at | 561.4 | P | 634 | P | 492.2 | P | 574.7 | P | --- |
| 264672_at | 1329.8 | P | 3115.1 | P | 1441 | P | 3160.4 | P | --- |
| 264673_at | 1410.7 | P | 1445.4 | P | 1648.3 | P | 1430.5 | P | ATATP-PRT2 |
| 264674_at | 806.5 | P | 775.5 | P | 977.5 | P | 833 | P | POLD4 |
| 264675_at | 351.9 | P | 423.1 | P | 483.7 | P | 433 | P | --- |
| 264676_at | 472.7 | P | 509.6 | P | 403.1 | P | 442.8 | P | --- |
| 264677_at | 138.9 | P | 118.4 | P | 144.1 | P | 158.2 | P | HYL1 |
| 264678_at | 684.4 | P | 667.9 | P | 706.5 | P | 705.9 | P | U2A' |
| 264679_s_at | 6880.1 | P | 5902.4 | P | 9304.9 | P | 6633.7 | P | --- |
| 264680_at | 175.2 | P | 227.9 | P | 78.4 | P | 133.5 | P | --- |
| 264681_s_at | 1.3 | A | 6.4 | A | 0.9 | A | 0.9 | A | --- |
| 264682_at | 1.7 | A | 7.9 | A | 50.3 | A | 22.4 | A | --- |
| 264683_at | 1059.9 | P | 998 | P | 954.1 | P | 1091.3 | P | FRA3 |
| 264684_at | 40.5 | A | 70.6 | P | 66.2 | A | 91 | P | HEXO3 |
| 264685_at | 121.4 | P | 85.7 | P | 76.9 | P | 50.9 | M | KOR2 |
| 264686_at | 467.6 | P | 637.5 | P | 481.7 | P | 628 | P | ATSK41 |
| 264687_at | 1869.4 | P | 1540.2 | P | 1689.9 | P | 1422.6 | P | XBCP3 |
| 264688_at | 34.2 | A | 42.5 | A | 63.8 | A | 38.2 | P | --- |
| 264689_at | 353.1 | P | 453.6 | P | 457.1 | P | 511 | P | --- |
| 264690_at | 200 | P | 162.1 | P | 183.4 | P | 172.4 | P | SNL4 |
| 264691_s_at | 18 | A | 12.9 | A | 5.1 | A | 10.7 | A | --- |
| 264692_at | 734.9 | P | 433 | P | 1005.6 | P | 452.7 | P | --- |
| 264693_at | 30.3 | A | 31.3 | A | 49.5 | A | 8.4 | A | CLE26 |
| 264694_at | 114.1 | P | 152.5 | P | 139.4 | P | 160.8 | P | --- |
| 264695_at | 51.2 | P | 88.8 | P | 88.7 | P | 107.3 | P | --- |
| 264696_at | 298.7 | P | 271.4 | P | 364.1 | P | 298.5 | P | --- |
| 264697_at | 125.6 | P | 215.3 | P | 183.6 | P | 225.2 | P | CYCD1;1 |
| 264698_at | 440.1 | P | 741.5 | P | 589.2 | P | 604.2 | P | --- |
| 264699_at | 449.6 | P | 417.4 | P | 404 | P | 401.2 | P | --- |
| 264700_at | 309.2 | P | 377 | P | 326.1 | P | 343.4 | P | --- |
| 264701_at | 2662.4 | P | 2681.1 | P | 2084 | P | 2299.1 | P | --- |
| 264702_at | 503.3 | P | 463.2 | P | 541.4 | P | 467.7 | P | --- |
| 264703_at | 661.8 | P | 745.1 | P | 819.1 | P | 792 | P | PP2A |
| 264704_at | 547.1 | P | 407.1 | P | 667.5 | P | 457.3 | P | LGT8 |
| 264705_at | 1630.5 | P | 1153.2 | P | 1733.4 | P | 1388.4 | P | --- |
| 264706_at | 3.6 | A | 12.2 | A | 10 | A | 27.5 | M | --- |
| 264707_at | 250.1 | P | 252.7 | P | 236.8 | P | 214.2 | P | --- |
| 264708_at | 174.3 | P | 181.7 | P | 140 | P | 145.7 | P | --- |
| 264709_at | 1006.7 | P | 825.4 | P | 912.5 | P | 886.3 | P | ATCDC5 |
| 264710_at | 60.5 | P | 54.9 | P | 58 | P | 56.5 | P | COBL6 |
| 264711_at | 143.4 | P | 139 | P | 179.9 | P | 143.8 | P | --- |
| 264712_at | 172.2 | P | 160.2 | P | 274.7 | P | 155 | P | ECT11 |
| 264713_at | 113.9 | P | 151.9 | P | 66.4 | P | 136.8 | P | --- |
| 264714_at | 26.3 | P | 15.1 | A | 25.7 | A | 9.7 | A | --- |
| 264715_at | 200.1 | P | 208.4 | P | 139 | P | 208.5 | P | --- |
| 264716_at | 22.7 | A | 28 | A | 35.6 | A | 21.2 | A | MMP |
| 264717_at | 124.4 | P | 142.9 | P | 125.5 | P | 157 | P | ATFH8 |
| 264718_at | 28.7 | A | 4.3 | A | 2.2 | A | 3.2 | A | --- |
| 264719_at | 23.8 | A | 15.4 | A | 9.8 | A | 9 | A | --- |
| 264720_at | 173.3 | P | 28 | A | 97.8 | P | 54.8 | A | --- |
| 264721_at | 72.5 | P | 66.1 | M | 83.8 | P | 106.3 | P | --- |
| 264722_at | 333.4 | P | 302.6 | P | 359.2 | P | 287.7 | P | --- |
| 264723_at | 192.9 | P | 125.5 | P | 153.7 | P | 136.8 | P | --- |
| 264724_at | 734.1 | P | 838.5 | P | 842.8 | P | 849.9 | P | CSN5A |
| 264725_at | 443.5 | P | 395.5 | P | 367.2 | P | 352.7 | P | --- |
| 264726_at | 129 | P | 118.4 | P | 111.2 | P | 115.8 | P | --- |
| 264727_at | 11591.5 | P | 8203.1 | P | 11791.3 | P | 8382.7 | P | CYTC-1 |
| 264728_at | 1931.8 | P | 2632.6 | P | 2282.6 | P | 2383.5 | P | --- |
| 264729_at | 26.6 | A | 33.6 | A | 46.6 | A | 14.8 | A | --- |
| 264730_at | 101 | P | 113.2 | P | 159.6 | P | 100.1 | P | --- |
| 264731_at | 313.7 | P | 207.8 | P | 306.3 | P | 203.4 | P | --- |
| 264732_at | 0.6 | A | 1.3 | A | 4.9 | A | 10.9 | A | --- |
| 264733_at | 21.8 | A | 4.4 | A | 25.2 | A | 27 | A | --- |
| 264734_at | 28.3 | A | 30.3 | A | 59.9 | A | 26.9 | A | SLAH1 |

| | | | | | | | | | |
|---|---|---|---|---|---|---|---|---|---|
| 264735_s_at | 4.7 | A | 4.5 | A | 8.2 | A | 1.8 | A | --- |
| 264736_at | 882.2 | P | 654.9 | P | 884.5 | P | 534.8 | P | --- |
| 264737_at | 12.6 | A | 1.5 | A | 6.6 | A | 2.8 | A | --- |
| 264738_at | 344.9 | P | 645.4 | P | 439.9 | P | 638.6 | P | --- |
| 264739_at | 77.2 | P | 66.8 | M | 96.4 | A | 45.1 | A | --- |
| 264740_at | 2.7 | A | 1.8 | A | 18.2 | A | 14.1 | A | --- |
| 264741_at | 216 | P | 288.9 | P | 271.1 | P | 240.7 | P | --- |
| 264742_at | 85.4 | P | 114.2 | P | 82 | P | 83.2 | P | --- |
| 264743_at | 4.3 | A | 6.1 | A | 4.2 | A | 6.2 | A | --- |
| 264744_at | 252.8 | P | 194.1 | P | 272.7 | P | 215.2 | P | --- |
| 264745_at | 1294.1 | P | 1579.9 | P | 1520.5 | P | 1415.9 | P | 41366 |
| 264746_at | 161.4 | P | 148.6 | P | 76.4 | M | 111.8 | P | WRKY6 |
| 264747_at | 151.5 | P | 125.4 | P | 160 | P | 159.7 | P | --- |
| 264748_at | 407.2 | P | 505.2 | P | 433.6 | P | 484.5 | P | EMB25 |
| 264749_at | 81.3 | P | 88.6 | P | 74.2 | P | 56.2 | P | --- |
| 264750_at | 184.1 | P | 194.7 | P | 197.4 | P | 223.3 | P | --- |
| 264751_at | 354.7 | P | 1609.6 | P | 1457.4 | P | 1679.7 | P | FRO3 |
| 264752_at | 297.6 | P | 187.5 | P | 381.1 | P | 278.1 | P | LPR1 |
| 264753_at | 32.3 | P | 32.8 | A | 23.5 | A | 26.6 | P | --- |
| 264754_at | 106.9 | P | 59.2 | P | 113.6 | P | 63.8 | P | --- |
| 264755_at | 41.9 | M | 44.7 | A | 37.6 | A | 26.4 | A | --- |
| 264756_at | 110.4 | P | 121.7 | P | 133.2 | P | 152.7 | P | --- |
| 264757_at | 200.8 | P | 170 | P | 140.7 | P | 136.6 | P | --- |
| 264758_at | 214.3 | P | 174.7 | P | 186.3 | P | 222.6 | P | --- |
| 264759_at | 23.6 | A | 9.9 | A | 30.2 | A | 29.2 | A | --- |
| 264760_at | 22.7 | A | 13.2 | A | 17.6 | A | 21.1 | A | SYP124 |
| 264761_at | 92.2 | P | 96.7 | P | 127.9 | P | 74.6 | P | --- |
| 264762_at | 14.8 | A | 11 | A | 8 | A | 20.7 | A | --- |
| 264763_at | 52.2 | P | 94.3 | P | 50.4 | P | 85.1 | P | --- |
| 264764_at | 15.4 | A | 24.9 | A | 4.1 | A | 4.1 | A | --- |
| 264765_at | 20.5 | A | 16.3 | P | 8.5 | A | 21.4 | A | --- |
| 264766_at | 65.4 | P | 54.8 | P | 77.6 | P | 42.5 | P | --- |
| 264767_at | 205.9 | P | 417 | P | 221.7 | P | 303.1 | P | SD1-29 |
| 264768_at | 15.1 | A | 15.4 | A | 6.1 | A | 2.7 | A | --- |
| 264769_at | 281 | P | 320.5 | P | 297.6 | P | 269 | P | --- |
| 264770_at | 240 | P | 483.7 | P | 266.3 | P | 429.5 | P | --- |
| 264771_at | 725.9 | P | 707.9 | P | 741.6 | P | 731.6 | P | TH1 |
| 264772_at | 933.4 | P | 841.9 | P | 870.4 | P | 775.7 | P | --- |
| 264773_at | 105.8 | P | 92 | P | 113.2 | P | 118.4 | P | --- |
| 264774_at | 337.8 | P | 373.3 | P | 211.5 | P | 216.9 | P | --- |
| 264775_at | 53.2 | A | 56.1 | A | 63.4 | A | 44.2 | A | CEL5 |
| 264776_at | 239.9 | P | 214.8 | P | 296.8 | P | 188.6 | P | --- |
| 264777_at | 211.3 | P | 107.6 | P | 236.5 | P | 91.4 | P | THA1 |
| 264778_at | 378 | P | 428.8 | P | 422.9 | P | 400.1 | P | AIP3 |
| 264779_at | 570.8 | P | 1073.4 | P | 763.6 | P | 734.1 | P | ACHT4 |
| 264780_at | 200.5 | P | 368.3 | P | 237.5 | P | 269.9 | P | EDR1 |
| 264781_at | 1081.4 | P | 1415.2 | P | 1283.2 | P | 1432.9 | P | SIG2 |
| 264782_at | 149.9 | P | 189.4 | P | 212.9 | P | 188.6 | P | MYB60 |
| 264783_at | 61.3 | A | 394.6 | P | 54.9 | A | 278.1 | P | PPCK1 |
| 264784_s_at | 112.1 | P | 108.9 | P | 200.2 | P | 147.4 | P | --- |
| 264785_at | 32.5 | A | 64.1 | A | 56.9 | A | 23.3 | A | --- |
| 264786_at | 12.7 | A | 9.4 | A | 15.1 | A | 21.4 | A | --- |
| 264787_at | 582.4 | P | 1855.6 | P | 425 | P | 1427.2 | P | ERD7 |
| 264788_at | 41.5 | A | 68.4 | P | 2.2 | A | 25.5 | A | --- |
| 264789_at | 1564.9 | P | 1092.1 | P | 947.9 | P | 997.2 | P | --- |
| 264790_at | 280 | P | 454.6 | P | 300.6 | P | 312.7 | P | ATHK1 |
| 264791_at | 7.9 | A | 1.7 | A | 10.2 | A | 3.1 | A | --- |
| 264792_at | 404.5 | P | 500.5 | P | 488.9 | P | 483 | P | --- |
| 264793_at | 445 | P | 531.4 | P | 556 | P | 483 | P | --- |
| 264794_at | 8 | A | 20.9 | A | 37.8 | A | 11.2 | A | --- |
| 264795_at | 156.4 | P | 230.5 | P | 254.2 | P | 265.9 | P | ZIGA4 |
| 264796_at | 515.9 | P | 650.2 | P | 593.9 | P | 625 | P | ZIGA4 |
| 264797_at | 131.3 | P | 88.2 | P | 96 | P | 101.3 | P | --- |
| 264798_at | 52.9 | A | 53.3 | A | 22.1 | A | 23 | A | XIC |
| 264799_at | 1205.9 | P | 1435.5 | P | 1314.5 | P | 1360.1 | P | NPQ1 |
| 264800_at | 163.9 | P | 172.2 | P | 149.8 | P | 119 | P | --- |
| 264801_at | 97.7 | P | 66.1 | P | 95.9 | P | 63.6 | P | emb2411 |
| 264802_at | 426 | P | 860 | P | 528.8 | P | 849.2 | P | SYP111 |
| 264803_at | 614.9 | P | 572.6 | P | 584.1 | P | 629.7 | P | --- |
| 264804_at | 57.4 | P | 117.4 | P | 51.4 | P | 114.7 | A | --- |
| 264805_at | 187.8 | P | 201.6 | P | 238.7 | P | 198.6 | P | ATRX |
| 264806_at | 241.7 | P | 153.4 | P | 205.3 | P | 160.4 | P | --- |
| 264807_at | 398.8 | P | 537.7 | P | 435.2 | P | 494.2 | P | --- |
| 264808_at | 488.2 | P | 420.1 | P | 329 | P | 407.2 | P | PRA1.E |
| 264809_at | 6744 | P | 2064.7 | P | 2865.4 | P | 4366.8 | P | CSD1 |
| 264810_at | 357.2 | P | 472.2 | P | 435.5 | P | 391.6 | P | --- |
| 264811_at | 2456.6 | P | 2854.3 | P | 3167.2 | P | 2614.2 | P | --- |
| 264812_at | 16.8 | A | 2.4 | A | 34.6 | P | 23.4 | A | --- |
| 264813_at | 10.4 | A | 9.1 | A | 38.2 | A | 13.4 | A | CPK16 |
| 264814_at | 139.2 | P | 100.7 | P | 142.6 | P | 150.7 | P | SDG37 |
| 264815_at | 34.3 | P | 57 | P | 56.2 | M | 43.9 | P | --- |
| 264816_at | 113.3 | P | 107.5 | P | 123.6 | P | 133.1 | P | --- |
| 264817_at | 4.9 | A | 9.1 | A | 35.1 | A | 20.9 | A | --- |
| 264818_at | 213.8 | P | 180.8 | P | 261.6 | P | 239 | P | NAF1 |
| 264819_at | 78.4 | P | 80.2 | P | 89.5 | P | 76.7 | P | --- |
| 264820_at | 1949.4 | P | 2565.4 | P | 2359.5 | P | 2470.9 | P | LIN2 |
| 264821_at | 116.5 | P | 170.6 | P | 261.8 | P | 212.7 | P | --- |
| 264822_at | 66.7 | P | 66.9 | P | 88.7 | P | 53.8 | M | --- |
| 264823_at | 43.7 | A | 44.4 | A | 54.6 | A | 19.6 | A | BSU1 |
| 264824_at | 372.7 | P | 313.1 | P | 311.1 | P | 386.5 | P | 2A6 |
| 264825_at | 5.4 | A | 4.1 | A | 7.6 | A | 2.7 | A | --- |
| 264826_at | 48.2 | M | 85.7 | P | 89.5 | P | 82.3 | P | 2A6 |
| 264827_at | 2.2 | A | 2.4 | A | 19.8 | A | 13.5 | A | --- |
| 264828_at | 302.4 | P | 321.5 | P | 314.7 | P | 242.6 | P | ATATG18G |
| 264829_at | 1.6 | A | 12.8 | A | 19.7 | A | 18.2 | A | ANAC006 |
| 264830_at | 3.5 | A | 1.8 | A | 8.9 | A | 23.6 | A | --- |
| 264831_at | 198.1 | P | 175.4 | P | 301.5 | P | 199.7 | P | --- |
| 264832_at | 73.7 | P | 124.3 | P | 107.5 | P | 98.8 | P | --- |
| 264833_at | 363.6 | P | 348.7 | P | 345.3 | P | 341.6 | P | --- |
| 264834_at | 382.8 | P | 287.4 | P | 293.7 | P | 332 | P | --- |
| 264835_at | 114.4 | P | 157 | P | 104.2 | P | 145.3 | P | --- |
| 264836_at | 320 | P | 400.5 | P | 210.8 | P | 350.3 | P | --- |
| 264837_at | 6148.5 | P | 6583.6 | P | 6193.4 | P | 6409.8 | P | --- |
| 264838_at | 470.9 | P | 465.8 | P | 428 | P | 442 | P | AHP5 |
| 264839_at | 2976.5 | P | 3933.9 | P | 3924.9 | P | 3597.2 | P | POR C |
| 264840_at | 288.6 | P | 225.6 | P | 284.4 | P | 220.3 | P | --- |
| 264841_at | 205.9 | P | 266.4 | P | 133.6 | P | 208.4 | P | --- |
| 264842_at | 1.2 | A | 2.2 | A | 1.7 | A | 8.1 | A | --- |
| 264843_at | 279.8 | P | 343.6 | P | 284.4 | P | 284.5 | P | --- |

| | | | | | | | | | |
|---|---|---|---|---|---|---|---|---|---|
| 264844_at | 182.3 | P | 193.8 | P | 232.9 | P | 175.9 | P | --- |
| 264845_at | 4381 | P | 5124.5 | P | 4836.9 | P | 5178.5 | P | ATHM1 |
| 264846_at | 51.9 | A | 33.6 | M | 83 | M | 27.9 | A | --- |
| 264847_at | 1125.1 | P | 1296.9 | P | 984.4 | P | 1205.7 | P | AP19 |
| 264848_at | 473.2 | P | 432 | P | 358.4 | P | 458 | P | --- |
| 264849_at | 2973.6 | P | 2992.8 | P | 3827.7 | P | 3131.9 | P | --- |
| 264850_at | 1282 | P | 1670.3 | P | 1698.5 | P | 2055.6 | P | --- |
| 264851_at | 443.1 | P | 388 | P | 338.2 | P | 311.3 | P | CPK6 |
| 264852_at | 532.9 | P | 504.1 | P | 425.6 | P | 461.8 | P | MLO8 |
| 264853_at | 61.5 | P | 48.8 | A | 94.3 | P | 29.7 | A | GLR2 |
| 264854_at | 468.6 | P | 598.9 | P | 385.4 | P | 371.2 | P | RHA3A |
| 264855_at | 1277.2 | P | 1235.9 | P | 1613.3 | P | 1071.5 | P | HSK |
| 264856_at | 227 | P | 239.5 | P | 201.5 | P | 183.5 | P | HMG2 |
| 264857_at | 868.5 | P | 663.3 | P | 686.9 | P | 889.3 | P | LGT9 |
| 264858_at | 282 | P | 328.1 | P | 314.7 | P | 273.4 | P | SNL3 |
| 264859_at | 157.3 | P | 196.6 | P | 190.2 | P | 220.1 | P | G6PD3 |
| 264860_at | 186 | P | 199 | P | 170.5 | P | 181.2 | P | --- |
| 264861_at | 23.6 | A | 36.7 | A | 22.9 | A | 23.3 | A | --- |
| 264862_at | 130.1 | P | 105 | P | 110.2 | P | 93.1 | P | --- |
| 264863_at | 32 | A | 11.5 | A | 35.1 | A | 32.2 | A | --- |
| 264864_at | 214.8 | P | 171.5 | P | 213.4 | P | 176.4 | P | --- |
| 264865_at | 428.8 | P | 535.4 | P | 423.2 | P | 671.4 | P | ARL1 |
| 264866_at | 72 | M | 102.3 | P | 73.6 | A | 86 | P | --- |
| 264867_at | 181.9 | P | 142.5 | P | 148.7 | P | 147.6 | P | FH4 |
| 264868_at | 120.9 | P | 74.3 | P | 122.7 | P | 74.1 | P | --- |
| 264869_at | 499.3 | P | 511.7 | P | 454.9 | P | 450.1 | P | --- |
| 264870_at | 102 | P | 90.9 | P | 124.4 | P | 84.9 | P | --- |
| 264871_at | 1442.3 | P | 1401.8 | P | 1563.4 | P | 1366.8 | P | IAR4 |
| 264872_at | 83.5 | P | 83.5 | P | 102.1 | P | 72.9 | P | 41520 |
| 264873_at | 3834.6 | P | 2348.6 | P | 3921.4 | P | 2414.8 | P | UGT74B1 |
| 264874_at | 129.8 | P | 165.5 | P | 116.4 | P | 179.2 | P | --- |
| 264875_at | 2881.8 | P | 2644.8 | P | 2877.6 | P | 2456.5 | P | AKR2B |
| 264876_at | 119.4 | P | 160.4 | P | 101.2 | P | 90.6 | P | --- |
| 264877_at | 23.8 | A | 27.7 | A | 11.3 | A | 24.2 | A | CYP51G2 |
| 264878_at | 682.1 | P | 643.5 | P | 654 | P | 674.3 | P | --- |
| 264879_at | 72.2 | P | 94.5 | P | 90.1 | P | 62 | P | --- |
| 264880_at | 53.1 | P | 73.6 | P | 100.2 | P | 71.9 | M | --- |
| 264881_s_at | 21.9 | P | 1.6 | A | 5.9 | A | 0.5 | A | --- |
| 264882_at | 64.7 | P | 6.2 | A | 42.6 | A | 1.8 | A | anac025 |
| 264883_s_at | 2262.2 | P | 2320.6 | P | 1420.9 | P | 1848.9 | P | --- |
| 264884_at | 53.8 | P | 81.5 | P | 71.1 | P | 66.2 | P | --- |
| 264885_s_at | 1157.4 | P | 1140.4 | P | 1082.6 | P | 919.1 | P | --- |
| 264886_at | 1229.4 | P | 46.4 | A | 472.7 | P | 122.2 | P | TPS04 |
| 264887_at | 125 | P | 92.6 | P | 145.9 | P | 85.7 | P | --- |
| 264888_at | 33.1 | A | 33.6 | A | 46.4 | A | 9.3 | A | --- |
| 264889_at | 92.4 | A | 5.5 | A | 61.9 | A | 44.1 | A | --- |
| 264890_at | 486.8 | P | 620.4 | P | 546.3 | P | 609.5 | P | --- |
| 264891_at | 59.1 | M | 70.5 | A | 31.1 | A | 61.6 | A | --- |
| 264892_at | 9.8 | A | 21 | A | 3.7 | A | 2.5 | A | --- |
| 264893_at | 83.8 | P | 173.7 | P | 87.7 | P | 157.1 | P | --- |
| 264894_at | 751.4 | P | 638.9 | P | 529.2 | P | 677.6 | P | --- |
| 264895_at | 202.1 | P | 183.7 | P | 208.4 | P | 118.5 | P | --- |
| 264896_at | 7 | A | 1.8 | A | 3.9 | A | 1.6 | A | AtGH9B6 |
| 264897_at | 693.8 | P | 901.6 | P | 817.3 | P | 932.1 | P | --- |
| 264898_at | 101.2 | P | 321.4 | P | 111.8 | P | 422.6 | P | --- |
| 264899_at | 6982.9 | P | 7102.3 | P | 6558.4 | P | 7569.7 | P | --- |
| 264900_at | 364 | P | 619.5 | P | 252.6 | P | 694.1 | P | PIN7 |
| 264901_at | 569 | P | 1031 | P | 813.3 | P | 972.6 | P | AST91 |
| 264902_at | 109.8 | P | 113.7 | P | 60.3 | P | 96.6 | P | --- |
| 264903_at | 3317.6 | P | 2996.5 | P | 2788.3 | P | 3216.9 | P | --- |
| 264904_s_at | 1646.3 | P | 1406.1 | P | 1613.2 | P | 1473.9 | P | NTRA |
| 264905_at | 15.1 | A | 5.9 | A | 41.7 | P | 20.6 | A | MLO7 |
| 264906_at | 179.7 | P | 149.4 | P | 213.6 | P | 166.9 | P | --- |
| 264907_at | 817.2 | P | 630.2 | P | 991.4 | P | 859.6 | P | --- |
| 264908_at | 340.4 | P | 346.8 | P | 300.7 | P | 294.5 | P | --- |
| 264909_at | 101.6 | P | 103.2 | P | 125.1 | P | 95.2 | P | --- |
| 264910_at | 407.8 | P | 388.8 | P | 454.7 | P | 507 | P | --- |
| 264911_at | 244 | P | 302.9 | P | 229.7 | P | 253.6 | P | --- |
| 264912_at | 12.9 | A | 14.8 | A | 30.8 | A | 17.3 | A | --- |
| 264913_at | 614.5 | P | 611.5 | P | 860.9 | P | 618.9 | P | --- |
| 264914_at | 1691.6 | P | 2035.6 | P | 1852.4 | P | 2057 | P | HAP13 |
| 264915_at | 1.5 | A | 8.9 | A | 2.6 | A | 2.5 | A | --- |
| 264916_at | 420.5 | P | 607.8 | P | 502.1 | P | 520 | P | ACLA-2 |
| 264917_at | 6.8 | A | 5 | A | 13.7 | A | 2.8 | A | --- |
| 264918_at | 5 | A | 2.1 | A | 17.8 | A | 13 | A | --- |
| 264919_at | 6.9 | A | 9 | A | 41.3 | A | 1.8 | A | --- |
| 264920_at | 1537 | P | 1978.2 | P | 1876.7 | P | 2028.9 | P | ECHID |
| 264921_at | 353.3 | P | 361.7 | P | 358.1 | P | 388.8 | P | --- |
| 264922_s_at | 904.9 | P | 864.9 | P | 659.8 | P | 788.4 | P | --- |
| 264923_s_at | 430.7 | P | 72.2 | P | 214.4 | P | 80.1 | P | --- |
| 264924_at | 180.5 | P | 181.6 | P | 218.8 | P | 169.5 | P | ATRPAC43 |
| 264925_at | 157.8 | P | 164.2 | P | 163.9 | P | 204 | P | --- |
| 264926_at | 897.6 | P | 805.2 | P | 786.5 | P | 932.9 | P | CB5LP |
| 264927_at | 502.2 | P | 468.5 | P | 483.2 | P | 376.1 | P | VPS34 |
| 264928_at | 2237.7 | P | 1672 | P | 1706.9 | P | 1607.7 | P | ATB2 |
| 264929_at | 1814.9 | P | 720.2 | P | 1367.6 | P | 832.9 | P | --- |
| 264930_at | 390.4 | P | 557.5 | P | 507.8 | P | 551.7 | P | NIK3 |
| 264931_at | 96.2 | P | 516.8 | P | 156.8 | P | 530.9 | P | --- |
| 264932_at | 131.4 | P | 214.6 | P | 119.7 | P | 176 | P | --- |
| 264933_at | 7 | A | 3.9 | A | 24.1 | A | 3.2 | A | --- |
| 264934_at | 1.2 | A | 1.8 | A | 3.3 | A | 2.3 | A | --- |
| 264935_at | 728.4 | P | 642.2 | P | 510.9 | P | 672.4 | P | --- |
| 264936_at | 654.7 | P | 619.6 | P | 717.8 | P | 550.8 | P | EDA16 |
| 264937_at | 11.7 | A | 34.8 | A | 10 | A | 14.6 | A | SCPL32 |
| 264938_at | 412.3 | P | 235.4 | P | 438.3 | P | 191 | P | --- |
| 264939_at | 55.5 | P | 71.4 | P | 62.1 | P | 83.5 | P | --- |
| 264940_at | 27.4 | A | 64.1 | A | 54.7 | A | 8.3 | A | AtGolS4 |
| 264941_at | 55.3 | P | 72.4 | P | 52.7 | A | 89.6 | P | --- |
| 264942_at | 476.8 | P | 293.9 | P | 363.8 | P | 277.6 | P | --- |
| 264943_at | 2.9 | A | 18.2 | A | 7.8 | A | 2.3 | A | --- |
| 264944_at | 413.2 | P | 415.8 | P | 314.1 | P | 324.6 | P | --- |
| 264945_at | 86.2 | P | 145.8 | P | 105.8 | P | 78.4 | P | MAF1 |
| 264946_at | 130.3 | P | 87.1 | P | 97.6 | P | 72.5 | P | --- |
| 264947_at | 90.7 | P | 99.1 | P | 116.2 | P | 123.7 | P | --- |
| 264948_at | 227.9 | P | 209 | P | 309.9 | P | 226.6 | P | --- |
| 264949_at | 94.8 | P | 73.7 | P | 102.4 | P | 78 | P | MAF1 |
| 264950_at | 19.1 | A | 22.8 | A | 35.3 | A | 27.2 | A | --- |
| 264951_at | 125.8 | P | 165.3 | P | 156.3 | P | 145.5 | P | --- |
| 264952_at | 490.1 | P | 679.6 | P | 571.1 | P | 551.6 | P | AtTLP1 |

| | | | | | | | | | |
|---|---|---|---|---|---|---|---|---|---|
| 264953_at | 4877.5 | P | 2039 | P | 2417.2 | P | 2847.2 | P | ADH1 |
| 264954_at | 1135.4 | P | 1037.5 | P | 1218.7 | P | 1045 | P | --- |
| 264955_at | 693.4 | P | 725.4 | P | 723.3 | P | 611 | P | --- |
| 264956_at | 940.9 | P | 1532.6 | P | 1230.5 | P | 1539.2 | P | ACR3 |
| 264957_at | 436.3 | P | 408.3 | P | 388.8 | P | 292.3 | P | SKP2B |
| 264958_at | 362.8 | P | 1116 | P | 212.8 | P | 738.2 | P | --- |
| 264959_at | 1374.6 | P | 1600.9 | P | 1432.7 | P | 1431.3 | P | --- |
| 264960_at | 1742.4 | P | 926.4 | P | 1216.4 | P | 751.9 | P | ATEXT4 |
| 264961_at | 247 | P | 279.8 | P | 300.7 | P | 264.5 | P | PRAF1 |
| 264962_at | 62.4 | A | 90.5 | A | 92.8 | A | 105.4 | A | PIN6 |
| 264963_at | 539.2 | P | 767.2 | P | 520.7 | P | 781.4 | P | ABC4 |
| 264964_at | 44.2 | P | 50 | P | 52.9 | P | 50.1 | P | --- |
| 264965_at | 196.7 | P | 177.6 | P | 190.8 | P | 160.9 | P | --- |
| 264966_at | 43.7 | P | 23.7 | A | 58 | A | 34.3 | A | --- |
| 264967_at | 2083 | P | 2186.1 | P | 1869.5 | P | 2134.3 | P | --- |
| 264968_at | 1642 | P | 2108 | P | 1462.7 | P | 2459.2 | P | --- |
| 264969_at | 282.8 | P | 275.2 | P | 326 | P | 297.2 | P | --- |
| 264970_at | 4098 | P | 3837.7 | P | 3792.1 | P | 3718.9 | P | --- |
| 264971_at | 60.4 | A | 91.8 | P | 46.4 | A | 86.7 | M | --- |
| 264972_at | 22.1 | P | 33.9 | P | 42.2 | A | 52.3 | P | ASY1 |
| 264973_at | 31.8 | A | 19.4 | A | 67.1 | A | 3.4 | A | --- |
| 264974_at | 258.4 | P | 208.3 | P | 250.9 | P | 261.7 | P | ATHB54 |
| 264975_at | 128.4 | P | 152.2 | P | 157.7 | P | 123.4 | P | --- |
| 264976_at | 67.7 | A | 44.8 | P | 87.1 | P | 55.7 | M | NRT1.6 |
| 264977_at | 1859.9 | P | 1776.7 | P | 1744.2 | P | 1819.1 | P | --- |
| 264978_at | 97.6 | P | 206.6 | P | 113.5 | P | 249.2 | P | --- |
| 264979_s_at | 56 | P | 90.1 | P | 51.5 | P | 68.1 | P | --- |
| 264980_at | 540.7 | P | 853.4 | P | 886.6 | P | 838.2 | P | --- |
| 264981_at | 2.8 | A | 29.8 | A | 11.3 | A | 4.7 | A | --- |
| 264982_at | 9.9 | A | 6.2 | A | 10.1 | A | 2.6 | A | --- |
| 264983_at | 38.6 | P | 26.3 | P | 51 | P | 26.5 | P | --- |
| 264984_at | 2682.2 | P | 1875.9 | P | 2327.6 | P | 2121.1 | P | --- |
| 264985_at | 502.2 | P | 477.3 | P | 498.2 | P | 371.3 | P | --- |
| 264986_at | 1655 | P | 1183.2 | P | 1415.4 | P | 1009 | P | ATGSTU13 |
| 264987_at | 3673 | P | 1563.8 | P | 3899.4 | P | 1697.7 | P | --- |
| 264988_at | 6.3 | A | 13.1 | A | 5.8 | A | 34.6 | A | ATGSTU14 |
| 264989_at | 387.6 | P | 205.9 | P | 321.4 | P | 181.1 | P | --- |
| 264990_at | 203.4 | P | 274.5 | P | 215.8 | P | 233.5 | P | --- |
| 264991_s_at | 60.3 | P | 123 | P | 135.8 | P | 117.8 | P | --- |
| 264992_at | 629.1 | P | 512.5 | P | 527.4 | P | 493.9 | P | --- |
| 264993_at | 46.4 | P | 50.8 | P | 31.8 | A | 67.4 | A | --- |
| 264994_at | 2.6 | A | 5.7 | A | 7.5 | A | 1.2 | A | --- |
| 264995_at | 4.2 | A | 11.4 | A | 22.9 | A | 4.1 | A | --- |
| 264996_at | 373.9 | P | 416.8 | P | 456.5 | P | 409.5 | P | LINC1 |
| 264997_at | 35.4 | A | 26.7 | A | 76.9 | P | 73.8 | P | HAC2 |
| 264998_at | 108.5 | P | 227.3 | P | 103.6 | P | 167.8 | P | --- |
| 264999_at | 442.8 | P | 326.6 | P | 296.3 | P | 341.3 | P | --- |
| 265000_at | 158.8 | P | 126 | P | 142.1 | P | 108.9 | P | --- |
| 265001_at | 4528.5 | P | 4290.8 | P | 4173.1 | P | 4577.4 | P | --- |
| 265002_at | 123.6 | P | 136 | P | 135.8 | P | 196.5 | P | LHT2 |
| 265003_at | 48.8 | A | 32.2 | A | 74.4 | A | 22.6 | A | --- |
| 265004_at | 11.1 | A | 10.9 | A | 5.8 | A | 20.2 | A | --- |
| 265005_at | 251.5 | P | 216.2 | P | 247.6 | P | 188.5 | P | --- |
| 265006_at | 2301 | P | 1740 | P | 2355.4 | P | 1609.2 | P | TIM13 |
| 265007_s_at | 12.3 | A | 1 | A | 1.3 | A | 2.7 | A | RALFL8 |
| 265008_at | 56.5 | P | 34.3 | P | 54 | P | 25.3 | P | MLO6 |
| 265009_at | 9.4 | A | 11.4 | A | 24.1 | A | 12 | A | --- |
| 265010_at | 2387.5 | P | 2289 | P | 2236.1 | P | 2250.7 | P | --- |
| 265011_at | 840 | P | 710.2 | P | 980.5 | P | 803.9 | P | ALB4 |
| 265012_at | 31.2 | A | 46.8 | P | 59.6 | A | 68.7 | P | KCR2 |
| 265013_at | 417.3 | P | 369.8 | P | 328 | P | 397.7 | P | --- |
| 265014_at | 33.3 | A | 18.4 | A | 8.8 | A | 49.1 | A | --- |
| 265015_at | 26.2 | A | 37.9 | A | 66.2 | A | 31.2 | A | --- |
| 265016_at | 6.6 | A | 27.5 | A | 20.7 | A | 20.1 | A | --- |
| 265017_at | 10.2 | A | 2.1 | A | 20.1 | A | 1.2 | A | --- |
| 265018_s_at | 16.9 | A | 2.8 | A | 26.6 | A | 7.5 | A | --- |
| 265019_at | 295.2 | P | 290.9 | P | 252.2 | P | 284.4 | P | --- |
| 265020_at | 4.6 | A | 9.9 | A | 16.9 | A | 3.9 | A | CYP86C1 |
| 265021_at | 559.2 | P | 592.2 | P | 597.8 | P | 556 | P | --- |
| 265022_at | 89.1 | P | 54.9 | P | 105.4 | P | 46.6 | P | BCP1 |
| 265023_at | 456.8 | P | 410.8 | P | 410.7 | P | 412.2 | P | --- |
| 265024_at | 21.7 | A | 51.4 | A | 11.4 | A | 37.7 | A | --- |
| 265025_at | 273.5 | P | 245.2 | P | 278.4 | P | 289.2 | P | --- |
| 265026_at | 3224.1 | P | 3471.1 | P | 3559 | P | 3265.5 | P | --- |
| 265027_at | 298.3 | P | 418 | P | 341.7 | P | 343.3 | P | NFD2 |
| 265028_at | 142.8 | P | 477.6 | P | 220.2 | P | 289.5 | P | --- |
| 265029_at | 552.1 | P | 320.5 | P | 500 | P | 318.8 | P | ZFP7 |
| 265030_at | 129.4 | P | 25.4 | A | 97.3 | P | 44.2 | P | --- |
| 265031_at | 2.8 | A | 6.7 | A | 5.1 | A | 21.2 | A | --- |
| 265032_at | 422.8 | P | 568.3 | P | 451.3 | P | 652.1 | P | RPL3B |
| 265033_at | 17770.5 | P | 18249.6 | P | 18932.2 | P | 17173 | P | LHCA3 |
| 265034_at | 236.2 | P | 266.9 | P | 231.4 | P | 271.3 | P | --- |
| 265035_at | 805.1 | P | 724.1 | P | 799.1 | P | 682.2 | P | --- |
| 265036_at | 1188.2 | P | 1083.5 | P | 1145.8 | P | 1089.1 | P | PRP39 |
| 265037_at | 3007.6 | P | 2479.6 | P | 3634.6 | P | 2809.6 | P | ATPHB2 |
| 265038_at | 11 | A | 35.6 | P | 51.1 | A | 47.7 | P | --- |
| 265039_at | 133.6 | P | 169.4 | P | 159.3 | P | 186.2 | P | --- |
| 265040_at | 430 | P | 445.7 | P | 411.1 | P | 396 | P | GBF4 |
| 265041_at | 158.2 | P | 125.8 | P | 165.8 | P | 119.7 | P | TOM22-I |
| 265042_at | 1183.6 | P | 1724.2 | P | 1079.1 | P | 1515.8 | P | --- |
| 265043_at | 1310.3 | P | 1786.7 | P | 1131.8 | P | 1482.2 | P | ATNAP4 |
| 265044_at | 541.4 | P | 489.3 | P | 501.1 | P | 459.2 | P | VPS2.3 |
| 265045_s_at | 322.1 | P | 340.2 | P | 263.6 | P | 308 | P | ADK1 |
| 265046_s_at | 5.6 | A | 0.6 | A | 19.9 | A | 1.1 | A | --- |
| 265047_at | 21.7 | A | 6.9 | A | 10.5 | A | 24.5 | A | IQD27 |
| 265048_at | 8.1 | A | 18.5 | A | 64.8 | P | 17.6 | A | --- |
| 265049_at | 4.9 | A | 12.7 | A | 54 | A | 6.1 | A | --- |
| 265050_at | 12.7 | A | 15.2 | A | 37 | M | 7.4 | A | --- |
| 265051_at | 1395.5 | P | 239.6 | P | 1814.9 | P | 311.4 | P | --- |
| 265052_at | 0.7 | A | 11.4 | A | 1.8 | A | 0.9 | A | --- |
| 265053_at | 1915.1 | P | 578.2 | P | 1070 | P | 1103.4 | P | --- |
| 265054_x_at | 32.7 | A | 21.9 | A | 7 | A | 23.6 | A | --- |
| 265055_at | 4.1 | A | 0.8 | A | 2.4 | A | 1.2 | A | --- |
| 265056_at | 4873.8 | P | 4241.1 | P | 3776.6 | P | 4672.9 | P | --- |
| 265057_at | 228.5 | P | 203.1 | P | 254.3 | P | 160.6 | P | --- |
| 265058_s_at | 3920 | P | 751.3 | P | 2839.1 | P | 1639.9 | P | MBP2 |
| 265059_at | 319.3 | P | 239.7 | P | 299 | P | 260.1 | P | AR791 |
| 265060_at | 424.4 | P | 542.1 | P | 561.8 | P | 551.2 | P | ATHB-15 |
| 265061_at | 218.2 | P | 160.3 | P | 273 | P | 162.9 | P | --- |

| | | | | | | | | | |
|---|---|---|---|---|---|---|---|---|---|
| 265062_at | 28.6 | A | 1.9 | A | 27.8 | A | 14.7 | A | --- |
| 265063_at | 62.1 | M | 54.6 | P | 70.9 | P | 64.7 | P | --- |
| 265064_at | 26.1 | A | 18.7 | A | 46.3 | A | 42.6 | A | ENT7 |
| 265065_at | 14.2 | A | 32.6 | A | 43.5 | A | 40.1 | A | ATPCS2 |
| 265066_at | 835.2 | P | 1563.9 | P | 1527.4 | P | 1477.2 | P | FLA9 |
| 265067_at | 91.6 | P | 98.7 | P | 95.5 | P | 65.6 | P | --- |
| 265068_at | 58.3 | P | 91 | P | 98.9 | P | 64 | P | --- |
| 265069_at | 40.4 | A | 16.5 | A | 32.1 | A | 33.2 | A | --- |
| 265070_at | 498.2 | P | 438.2 | P | 432.2 | P | 454.7 | P | BCDH BETA1 |
| 265071_at | 379 | P | 469.4 | P | 425.4 | P | 448.3 | P | TBP2 |
| 265072_at | 39.8 | A | 47.2 | P | 51.6 | A | 46.4 | M | LAS |
| 265073_at | 1963.1 | P | 2493.7 | P | 1738.7 | P | 2396.5 | P | --- |
| 265074_at | 128.1 | P | 133.1 | P | 120.4 | P | 103.4 | P | emb1011 |
| 265075_at | 1485 | P | 1765.2 | P | 1499 | P | 1584.5 | P | --- |
| 265076_at | 7797.1 | P | 8019.9 | P | 6938.5 | P | 8267.3 | P | CPN60B |
| 265077_at | 683.6 | P | 820.2 | P | 857.6 | P | 729.8 | P | --- |
| 265078_at | 974.7 | P | 623 | P | 883.7 | P | 688.7 | P | ECT4 |
| 265079_at | 438.9 | P | 436.1 | P | 439.1 | P | 447.8 | P | --- |
| 265080_at | 2 | A | 4 | A | 2.2 | A | 3.9 | A | sks12 |
| 265081_at | 1.7 | A | 1.8 | A | 4.9 | A | 1.3 | A | MGP |
| 265082_at | 64.7 | P | 84.1 | A | 86.6 | M | 78.3 | P | --- |
| 265083_at | 610.5 | P | 237.5 | P | 397.6 | P | 318.9 | P | --- |
| 265084_at | 47 | A | 5.1 | A | 5.1 | A | 10.1 | A | SOM |
| 265085_at | 70.7 | A | 93.9 | P | 49.9 | A | 138.3 | P | TPX2 |
| 265086_at | 137 | P | 71.7 | P | 159.6 | P | 80.5 | P | RING1B |
| 265087_at | 221.6 | P | 142.7 | P | 191.4 | P | 157.9 | P | --- |
| 265088_at | 119.8 | P | 99.6 | P | 82.3 | P | 40.6 | P | SWI2 |
| 265089_at | 2 | A | 2.5 | A | 14.8 | A | 37.2 | A | SUVR1 |
| 265090_at | 137.5 | P | 107.3 | P | 114.7 | P | 123.5 | P | --- |
| 265091_s_at | 2028.2 | P | 337.5 | P | 2672.3 | P | 900.3 | P | --- |
| 265092_at | 908.7 | P | 995.5 | P | 922.5 | P | 946.7 | P | --- |
| 265093_at | 98.2 | P | 91.4 | P | 96.3 | M | 85 | M | --- |
| 265094_at | 4 | A | 3.5 | A | 5.6 | A | 2.5 | A | --- |
| 265095_at | 4.2 | A | 3.3 | A | 20 | A | 8.8 | A | CRU2 |
| 265096_at | 73.3 | P | 76.9 | P | 70.6 | P | 46.5 | P | --- |
| 265097_at | 259.4 | P | 227.4 | P | 235.9 | P | 241.6 | P | BARD1 |
| 265098_at | 166 | P | 186.9 | P | 161.1 | P | 208.9 | P | --- |
| 265099_at | 37.3 | A | 36.5 | A | 31.3 | A | 40 | A | --- |
| 265100_at | 136.5 | P | 106.7 | P | 147.7 | P | 118.4 | P | OSB4 |
| 265101_at | 1427.8 | P | 1239.8 | P | 1975 | P | 1299.6 | P | --- |
| 265102_at | 4.3 | A | 8.7 | A | 6.5 | A | 7.8 | A | --- |
| 265103_at | 184.7 | P | 171.3 | P | 214.6 | P | 190.6 | P | --- |
| 265104_at | 293.2 | P | 238.2 | P | 257.9 | P | 198.5 | P | ATO2 |
| 265105_s_at | 651.5 | P | 325.8 | P | 503.4 | P | 416.8 | P | --- |
| 265106_s_at | 64.4 | P | 93.8 | P | 95.1 | P | 66.3 | P | --- |
| 265107_s_at | 256.2 | P | 267.1 | P | 246.4 | P | 234.7 | P | --- |
| 265108_s_at | 22 | P | 27.2 | P | 32.8 | A | 50.4 | P | --- |
| 265109_s_at | 472.1 | P | 442.7 | P | 468.4 | P | 428.7 | P | --- |
| 265110_at | 3.1 | A | 2.6 | A | 3.7 | A | 7.9 | A | --- |
| 265111_at | 741.3 | P | 900.1 | P | 294.7 | P | 800.7 | P | --- |
| 265112_at | 4.8 | A | 3.7 | A | 21.5 | A | 2 | A | --- |
| 265113_at | 40.2 | A | 32.5 | M | 45.9 | P | 36 | P | --- |
| 265114_at | 1.1 | A | 6.3 | A | 29.9 | A | 14.5 | A | LRX2 |
| 265115_at | 37.9 | A | 34.4 | A | 46.9 | P | 28.4 | A | --- |
| 265116_at | 1198.5 | P | 2529.5 | P | 1812.8 | P | 2577.4 | P | --- |
| 265117_at | 337.5 | P | 623.7 | P | 555.7 | P | 657.5 | P | --- |
| 265118_at | 685 | P | 207.4 | P | 597.2 | P | 288.4 | P | --- |
| 265119_at | 1963.5 | P | 924.5 | P | 1190 | P | 1025.9 | P | FMO GS-OX4 |
| 265120_at | 575.1 | P | 780.3 | P | 646.4 | P | 759.1 | P | KAS III |
| 265121_at | 1983.5 | P | 1122.8 | P | 2393.7 | P | 1026.1 | P | FMO GS-OX3 |
| 265122_at | 583.3 | P | 197.9 | P | 448.2 | P | 333.2 | P | FMO GS-OX2 |
| 265123_at | 3.1 | A | 3.7 | A | 5.7 | A | 3.9 | A | --- |
| 265124_at | 3.5 | A | 7.7 | A | 4 | A | 2.7 | A | --- |
| 265125_at | 3.6 | A | 2.8 | A | 4.8 | A | 2.6 | A | --- |
| 265126_s_at | 27.2 | A | 29 | A | 57.2 | A | 21.7 | A | --- |
| 265127_at | 1.5 | A | 3.1 | A | 1.3 | A | 2.1 | A | sks14 |
| 265128_at | 9.1 | A | 48.9 | A | 28.9 | A | 33.8 | A | --- |
| 265129_at | 327.3 | P | 402.3 | P | 425.9 | P | 385.5 | P | SUF4 |
| 265130_at | 566.3 | P | 512.2 | P | 542.9 | P | 646.3 | P | --- |
| 265131_at | 6.8 | A | 8.3 | A | 23.4 | A | 36.3 | A | JP630 |
| 265132_at | 125.3 | P | 211.9 | P | 73 | P | 137.9 | P | --- |
| 265133_s_at | 23.4 | P | 1.6 | A | 3.2 | A | 43.8 | A | --- |
| 265134_at | 35.7 | A | 3 | A | 38.6 | A | 46.9 | A | LPAT3 |
| 265135_at | 64.5 | A | 123.8 | P | 133.1 | A | 102.5 | A | --- |
| 265136_at | 31.6 | A | 94.2 | P | 4.1 | A | 38.9 | P | --- |
| 265137_at | 2.4 | A | 6.5 | A | 19.4 | A | 14.1 | A | --- |
| 265138_at | 25.6 | A | 22.6 | A | 11.5 | A | 18.1 | A | --- |
| 265139_at | 507 | P | 447.1 | P | 388.2 | P | 395.4 | P | --- |
| 265140_at | 1.1 | A | 1.2 | A | 1.7 | A | 6.3 | A | --- |
| 265141_at | 450.6 | P | 375.7 | P | 397.4 | P | 349.9 | P | --- |
| 265142_at | 478.1 | P | 410.2 | P | 617.4 | P | 380.6 | P | DABB1 |
| 265143_at | 475 | P | 197.9 | P | 295.1 | P | 214.8 | P | NFU5 |
| 265144_at | 71.6 | P | 111.7 | P | 108.1 | P | 93.9 | P | --- |
| 265145_at | 3.9 | A | 13.5 | A | 7.6 | A | 8.8 | A | PLT2 |
| 265146_at | 41.9 | P | 22.5 | A | 14.8 | A | 23.1 | A | --- |
| 265147_at | 116.1 | P | 75.3 | P | 149.1 | P | 75 | P | --- |
| 265148_at | 1230.7 | P | 1367.3 | P | 1249.6 | P | 1340.9 | P | --- |
| 265149_at | 2429.9 | P | 4204.6 | P | 3146.8 | P | 3518.3 | P | --- |
| 265150_at | 4875.8 | P | 4734.5 | P | 5341.8 | P | 5109.1 | P | --- |
| 265151_at | 345.7 | P | 176.9 | P | 384.2 | P | 225.1 | P | --- |
| 265152_s_at | 47.1 | P | 27.6 | A | 42.7 | P | 26.7 | A | --- |
| 265153_at | 7.5 | A | 4.1 | A | 10.1 | A | 6.1 | A | UFO |
| 265154_at | 163.9 | P | 110.7 | P | 171.1 | P | 156.3 | P | --- |
| 265155_at | 4.4 | A | 4.8 | A | 2.8 | A | 3.4 | A | --- |
| 265156_at | 20.5 | A | 13.4 | A | 24.2 | A | 10.4 | A | --- |
| 265157_at | 21.8 | A | 36.6 | P | 32.2 | A | 32.4 | P | --- |
| 265158_at | 17.7 | A | 12.9 | A | 10.8 | A | 3.7 | A | --- |
| 265159_at | 7.5 | A | 1.4 | A | 35.3 | A | 5.1 | A | --- |
| 265160_at | 1.6 | A | 13.7 | A | 4.4 | A | 2.9 | A | --- |
| 265161_at | 82.5 | A | 403.5 | P | 109.4 | A | 346.7 | P | --- |
| 265162_at | 826.4 | P | 818 | P | 748.1 | P | 765.3 | P | --- |
| 265163_at | 28 | P | 37.1 | A | 83.5 | A | 7.1 | A | --- |
| 265164_at | 10.5 | A | 40.7 | A | 6.8 | A | 21.4 | A | --- |
| 265165_at | 2.3 | A | 0.4 | A | 1.1 | A | 2.1 | A | --- |
| 265166_at | 12.3 | A | 4.2 | A | 9.2 | A | 17.8 | A | --- |
| 265167_at | 8.5 | A | 0.7 | A | 5.5 | A | 23.4 | A | --- |
| 265168_at | 29.1 | A | 27.8 | M | 52.5 | P | 3.8 | A | --- |
| 265169_x_at | 30.2 | A | 76.5 | M | 49.9 | A | 68.3 | P | --- |
| 265170_at | 97.8 | P | 168.2 | P | 111.4 | P | 99.3 | P | BCA3 |

| | | | | | | | | | |
|---|---|---|---|---|---|---|---|---|---|
| 265171_at | 87 | A | 153.7 | P | 73.9 | A | 124.8 | P | --- |
| 265172_at | 3299 | P | 2503.3 | P | 2841.4 | P | 3207.4 | P | SPDS1 |
| 265173_at | 79.7 | P | 95.2 | P | 113.4 | A | 87.2 | P | --- |
| 265174_s_at | 162.7 | P | 231 | P | 189 | P | 211.3 | P | --- |
| 265175_at | 117.1 | P | 228.8 | P | 191.6 | P | 171.4 | P | ATCSLA03 |
| 265176_at | 26.3 | A | 5.9 | A | 28.2 | A | 19.5 | A | --- |
| 265177_at | 1.1 | A | 2.7 | A | 1 | A | 1.6 | A | --- |
| 265178_at | 2.4 | A | 3.8 | A | 7.4 | A | 1.8 | A | --- |
| 265179_at | 0.9 | A | 0.7 | A | 2.6 | A | 1.6 | A | --- |
| 265180_at | 3.9 | A | 10.6 | A | 41.6 | A | 24.6 | A | --- |
| 265181_at | 4 | A | 38.8 | A | 2.9 | A | 12.7 | A | --- |
| 265182_at | 1686.8 | P | 2812.8 | P | 1892.2 | P | 2787.5 | P | --- |
| 265183_at | 246.6 | P | 265.2 | P | 260.5 | P | 331 | P | --- |
| 265184_at | 208.5 | P | 140.4 | P | 143.4 | P | 135.9 | P | --- |
| 265185_at | 34.2 | P | 31.2 | A | 34.4 | A | 64.2 | P | --- |
| 265186_at | 1.4 | A | 12.6 | A | 15.3 | A | 1.4 | A | --- |
| 265187_at | 4840.3 | P | 5767 | P | 4383.1 | P | 5817.1 | P | --- |
| 265188_at | 1010.1 | P | 403.2 | P | 350.1 | P | 433.1 | P | ALDH2B7 |
| 265189_at | 103.5 | P | 121.8 | P | 67.2 | P | 83.3 | P | --- |
| 265190_at | 804.7 | P | 759.9 | P | 720.8 | P | 692.8 | P | --- |
| 265191_at | 195.5 | P | 200.5 | P | 276.7 | P | 180.5 | P | --- |
| 265192_at | 459.3 | P | 415.9 | P | 329.4 | P | 371.3 | P | --- |
| 265193_at | 364.5 | P | 440.3 | P | 353.5 | P | 513.9 | P | --- |
| 265194_at | 8992.2 | P | 7491.6 | P | 8087 | P | 7503.9 | P | EFE |
| 265195_at | 158.2 | P | 141.6 | P | 159.9 | P | 163.3 | P | --- |
| 265196_at | 140.4 | A | 159.7 | P | 161.9 | A | 193.5 | P | SWC2 |
| 265197_at | 817.4 | P | 309.4 | P | 470.9 | P | 462.1 | P | UGT73C1 |
| 265198_at | 27.7 | P | 33.4 | P | 55.6 | P | 24.3 | P | UGT73C2 |
| 265199_s_at | 326.9 | P | 52.4 | P | 102 | P | 58.5 | P | --- |
| 265200_s_at | 1467.3 | P | 999 | P | 1766.8 | P | 1106.4 | P | UGT73C6 |
| 265201_at | 312.9 | P | 364.5 | P | 236.5 | P | 374.7 | P | --- |
| 265202_at | 39.8 | A | 20.2 | A | 16 | A | 37.2 | A | AT2G36810 |
| 265203_at | 974.3 | P | 1033.3 | P | 728.9 | P | 1013.1 | P | --- |
| 265204_at | 33.2 | A | 22.7 | A | 28.5 | A | 15.4 | A | --- |
| 265205_at | 52.3 | P | 55.2 | P | 78.3 | M | 55.4 | P | PAB7 |
| 265206_at | 139.3 | P | 183.3 | P | 122.6 | P | 175.1 | P | --- |
| 265207_at | 321.7 | P | 400.7 | P | 370.2 | P | 388.4 | P | --- |
| 265208_at | 18.3 | A | 101.7 | P | 6.5 | A | 70.7 | P | --- |
| 265209_at | 16.7 | A | 1.7 | A | 4.5 | A | 2.7 | A | ATHB22 |
| 265210_at | 5884.9 | P | 5353.3 | P | 6033.1 | P | 6449.7 | P | RPL24A |
| 265211_at | 1.7 | A | 1.7 | A | 6.5 | A | 3.1 | A | ATECP63 |
| 265212_at | 236.2 | P | 289.3 | P | 306.2 | P | 345.1 | P | --- |
| 265213_at | 76.7 | A | 32 | A | 63.5 | M | 43.7 | A | --- |
| 265214_at | 842.5 | P | 299.6 | P | 653.9 | P | 343.8 | P | --- |
| 265215_at | 12.2 | A | 10.8 | A | 3 | A | 1.7 | A | --- |
| 265216_at | 23.9 | A | 6.4 | A | 16.6 | A | 6.5 | A | MAPKKK18 |
| 265217_s_at | 200.7 | P | 189.9 | P | 203.6 | P | 212.4 | P | --- |
| 265218_at | 363.7 | P | 404.3 | P | 406.4 | P | 383.9 | P | GTF2H2 |
| 265219_at | 1845.4 | P | 2553 | P | 2501.6 | P | 2571.7 | P | --- |
| 265220_at | 1164.5 | P | 1456.9 | P | 1071.5 | P | 1226.1 | P | PTR2 |
| 265221_s_at | 284.2 | P | 193.6 | P | 233.1 | P | 345 | P | GAD3 |
| 265222_at | 1343 | P | 1666.6 | P | 1303.3 | P | 1466.5 | P | --- |
| 265223_at | 10.3 | A | 22.6 | A | 3.9 | A | 3 | A | --- |
| 265224_at | 15.1 | A | 15.4 | A | 21.1 | A | 11.3 | A | --- |
| 265225_at | 396 | P | 444.5 | P | 453 | P | 437.4 | P | --- |
| 265226_at | 1032.5 | P | 1132.3 | P | 989.9 | P | 1095.2 | P | --- |
| 265227_s_at | 37.8 | P | 40.3 | P | 85.9 | P | 71 | P | --- |
| 265228_s_at | 615.7 | P | 469.4 | P | 486.7 | P | 369 | P | --- |
| 265229_s_at | 37 | P | 26.9 | P | 33.1 | M | 30.5 | A | --- |
| 265230_s_at | 214.4 | P | 203.2 | P | 156 | P | 217.1 | P | --- |
| 265231_s_at | 5.6 | A | 9.8 | A | 32.3 | A | 9 | A | --- |
| 265232_s_at | 149.4 | P | 112.4 | P | 207.2 | P | 141.8 | P | --- |
| 265233_s_at | 116.3 | P | 159.8 | P | 88.2 | A | 110.3 | P | --- |
| 265234_at | 4.7 | A | 12.5 | A | 16.9 | A | 31.2 | A | --- |
| 265235_s_at | 73.8 | P | 34.8 | P | 128.5 | P | 65.4 | P | --- |
| 265236_s_at | 2.2 | A | 14 | A | 1.5 | A | 3.4 | A | --- |
| 265237_s_at | 2.1 | A | 5.4 | A | 8 | A | 11.5 | A | --- |
| 265238_s_at | 72.6 | P | 38.1 | P | 60.4 | P | 38.5 | P | --- |
| 265239_s_at | 3 | A | 2.9 | A | 39.6 | A | 3.2 | A | --- |
| 265240_s_at | 26.5 | A | 29.1 | A | 27.1 | A | 8.7 | A | --- |
| 265241_at | 12.7 | A | 15.7 | A | 4.4 | A | 3.6 | A | --- |
| 265242_at | 47.7 | P | 45.6 | P | 56.5 | A | 30.8 | A | --- |
| 265243_at | 127.1 | P | 107.3 | P | 112.1 | P | 74.7 | P | NPG1 |
| 265244_at | 828.2 | P | 1028.8 | P | 484 | P | 884.5 | P | ATPAO2 |
| 265245_at | 143.3 | P | 101.5 | P | 61.5 | P | 171.4 | P | --- |
| 265246_at | 24.5 | A | 37.9 | A | 36.4 | A | 22 | A | ATPMEPCRD |
| 265247_at | 9699.3 | P | 10334.4 | P | 9875.7 | P | 10451.9 | P | --- |
| 265248_at | 362.2 | P | 395.2 | P | 222.1 | P | 294.8 | P | PIF4 |
| 265249_at | 68.5 | A | 49.1 | P | 61 | P | 40.8 | P | SGR5 |
| 265250_at | 129.2 | P | 215.4 | P | 185.8 | P | 199.9 | P | BRL2 |
| 265251_at | 22.1 | A | 13.9 | A | 20.6 | A | 26.4 | A | TET14 |
| 265252_at | 325.3 | P | 371.3 | P | 394 | P | 353 | P | SOS1 |
| 265253_at | 85.9 | M | 134.2 | P | 154.3 | P | 132 | P | --- |
| 265254_at | 305.6 | P | 397.9 | P | 338.9 | P | 408.9 | P | DRB2 |
| 265255_at | 37.2 | P | 39 | P | 54 | M | 38.3 | P | --- |
| 265256_at | 589.6 | P | 606.9 | P | 563.3 | P | 597.1 | P | --- |
| 265257_at | 3559.1 | P | 3792 | P | 4479.5 | P | 3985.3 | P | --- |
| 265258_at | 764 | P | 619.1 | P | 841.6 | P | 617.6 | P | --- |
| 265259_at | 674.4 | P | 502.2 | P | 985.5 | P | 547.3 | P | --- |
| 265260_at | 72.3 | P | 63.8 | M | 71.9 | P | 52.5 | P | anac042 |
| 265261_at | 42.9 | A | 34.5 | P | 41.3 | A | 47.2 | P | --- |
| 265262_at | 30.5 | P | 11.5 | A | 43.6 | P | 48.6 | P | --- |
| 265263_at | 15.8 | A | 4.7 | A | 24.4 | A | 36.5 | A | --- |
| 265264_at | 2.4 | A | 1 | A | 2.4 | A | 9.6 | A | --- |
| 265265_at | 45.5 | P | 113.9 | P | 92.8 | P | 107.1 | P | --- |
| 265266_at | 388.3 | P | 453.6 | P | 369.6 | P | 312 | P | AML2 |
| 265267_at | 78.6 | M | 58.1 | A | 95 | P | 44.1 | A | --- |
| 265268_at | 79.2 | P | 77.2 | P | 79.7 | A | 86.2 | P | --- |
| 265269_at | 268.4 | P | 258.5 | P | 274.6 | P | 224 | P | --- |
| 265270_at | 2856.2 | P | 2340.6 | P | 2673.8 | P | 2508.1 | P | --- |
| 265271_at | 188.8 | P | 195.8 | P | 178.1 | P | 167.3 | P | --- |
| 265272_at | 239.1 | P | 202.5 | P | 188.9 | P | 216.6 | P | ARF10 |
| 265273_at | 485.1 | P | 480.1 | P | 510.9 | P | 456.5 | P | --- |
| 265274_at | 528.5 | P | 353.3 | P | 610.2 | P | 440.9 | P | --- |
| 265275_at | 3.7 | A | 1.9 | A | 2.2 | A | 1.4 | A | --- |
| 265276_at | 732.3 | P | 249.1 | P | 340.5 | P | 264.6 | P | --- |
| 265277_at | 329.7 | P | 429.9 | P | 396.9 | P | 437.8 | P | --- |
| 265278_at | 118.8 | P | 111.5 | P | 105.3 | P | 107.4 | P | --- |
| 265279_at | 45.2 | A | 76.2 | P | 60.7 | P | 51.8 | P | --- |

| | | | | | | | | | |
|---|---|---|---|---|---|---|---|---|---|
| 265280_at | 12.3 | A | 8.3 | A | 31.6 | A | 4.2 | A | LCR5 |
| 265281_at | 877.9 | P | 981.5 | P | 745.9 | P | 996.5 | P | --- |
| 265282_at | 29.6 | A | 36 | A | 27.6 | A | 24 | A | --- |
| 265283_at | 389.1 | P | 442.5 | P | 309.8 | P | 427.2 | P | MUR3 |
| 265284_at | 1596.5 | P | 1854.6 | P | 1590.5 | P | 1674.5 | P | --- |
| 265285_at | 172.4 | P | 149.2 | P | 181.5 | P | 157.2 | P | --- |
| 265286_at | 2593.6 | P | 2681.1 | P | 2703.3 | P | 2888.2 | P | --- |
| 265287_at | 5167.4 | P | 5869.8 | P | 4569.8 | P | 5237.4 | P | PSAE-2 |
| 265288_at | 2381.8 | P | 2145.5 | P | 2130.1 | P | 2229 | P | --- |
| 265289_at | 301 | P | 288.7 | P | 292.5 | P | 275.4 | P | --- |
| 265290_at | 3.5 | A | 2.8 | A | 8.5 | A | 4.6 | A | --- |
| 265291_at | 527.7 | P | 535.6 | P | 518.4 | P | 528.6 | P | --- |
| 265292_s_at | 2.4 | A | 23.6 | A | 16.2 | A | 14.8 | A | --- |
| 265293_at | 27.9 | A | 29.9 | A | 43.9 | A | 24.6 | A | --- |
| 265294_at | 16.6 | A | 10.2 | A | 3.3 | A | 2.8 | A | --- |
| 265295_at | 79.1 | P | 102.8 | P | 115.4 | P | 114.9 | P | MCM9 |
| 265296_at | 38.4 | P | 36.3 | M | 61.7 | P | 30.3 | M | --- |
| 265297_at | 296.5 | P | 358.7 | P | 220.5 | P | 265.3 | P | --- |
| 265298_s_at | 2 | A | 8.8 | A | 1.3 | A | 1.2 | A | --- |
| 265299_at | 40.6 | A | 53.3 | A | 63.2 | A | 38 | A | --- |
| 265300_at | 3.6 | A | 2.1 | A | 34.2 | A | 8.8 | A | --- |
| 265301_s_at | 64.1 | P | 83.1 | P | 78.1 | P | 80.1 | P | --- |
| 265302_at | 81.9 | P | 149.2 | P | 104.2 | P | 150.3 | P | --- |
| 265303_at | 2.9 | A | 1.2 | A | 3.3 | A | 4.6 | A | --- |
| 265304_at | 19.7 | P | 1.3 | A | 6.1 | A | 1.7 | A | --- |
| 265305_at | 409.6 | P | 304.6 | P | 515.6 | P | 418.2 | P | --- |
| 265306_at | 158.3 | A | 173.9 | A | 113.7 | A | 150.5 | A | --- |
| 265307_at | 282.7 | P | 299.3 | P | 322.3 | P | 272.1 | P | RIN13 |
| 265308_at | 196 | P | 206.1 | P | 144.8 | P | 196 | P | ALE2 |
| 265309_at | 86.8 | P | 87.1 | P | 115.1 | P | 78.8 | P | XIG |
| 265310_at | 1204.3 | P | 1270 | P | 1221.2 | P | 1285 | P | --- |
| 265311_at | 56.5 | A | 45.9 | A | 65.4 | A | 33.3 | A | --- |
| 265312_at | 85.6 | P | 121.2 | P | 140.1 | P | 124.9 | P | --- |
| 265313_at | 17.1 | A | 45 | A | 60.6 | A | 58.7 | A | --- |
| 265314_at | 62.3 | A | 79.7 | A | 106 | A | 70.1 | A | --- |
| 265315_at | 79.8 | A | 41.8 | A | 23.6 | A | 8.3 | A | CLASP |
| 265316_at | 82.1 | P | 106.3 | P | 82.9 | P | 100.6 | P | --- |
| 265317_at | 196.3 | P | 226.2 | P | 270.4 | P | 180.3 | P | BRK1 |
| 265318_at | 243.9 | P | 203.9 | P | 185.8 | P | 207.7 | P | --- |
| 265319_at | 2193.6 | P | 3247.1 | P | 1836.7 | P | 2855.4 | P | IAA8 |
| 265320_at | 23.6 | A | 12.2 | A | 24.6 | A | 12.1 | A | --- |
| 265321_at | 1204 | P | 1244.5 | P | 1307.1 | P | 1138.5 | P | AtTLP2 |
| 265322_at | 27.1 | P | 39.1 | P | 38.4 | M | 23 | M | --- |
| 265323_at | 6.3 | A | 2 | A | 4 | A | 5.1 | A | SYP112 |
| 265324_at | 404 | P | 455.6 | P | 409 | P | 409.1 | P | ATCOAD |
| 265325_at | 243.2 | P | 304.9 | P | 175.2 | P | 329.7 | P | --- |
| 265326_at | 260.7 | P | 203.3 | P | 289 | P | 200.5 | P | --- |
| 265327_at | 112.9 | P | 68.8 | A | 65.1 | P | 45.4 | P | --- |
| 265328_at | 30.3 | M | 24.1 | A | 22.8 | A | 19.8 | A | --- |
| 265329_at | 3.7 | A | 3.3 | A | 1.6 | A | 2.5 | A | SDH1-2 |
| 265330_at | 1130.1 | P | 948.7 | P | 1307.5 | P | 877.8 | P | --- |
| 265331_at | 10 | A | 4.1 | A | 33.6 | A | 41.2 | A | --- |
| 265332_at | 233.2 | P | 134.9 | P | 169.8 | P | 162.5 | P | --- |
| 265333_at | 103.5 | P | 148.7 | P | 121.7 | P | 150.5 | P | AtHB24 |
| 265334_at | 57.5 | A | 224.3 | P | 112.6 | P | 179.4 | P | --- |
| 265335_at | 99.5 | P | 137.3 | P | 121 | P | 147.5 | P | --- |
| 265336_at | 326.2 | P | 678.8 | P | 308.1 | P | 519.9 | P | APC10 |
| 265337_at | 832.4 | P | 887.2 | P | 887.7 | P | 862.4 | P | TTN5 |
| 265338_at | 748.8 | P | 698.1 | P | 638.6 | P | 748.5 | P | --- |
| 265339_at | 674.9 | P | 1313.8 | P | 771.8 | P | 1538.3 | P | AtPPa2 |
| 265340_at | 543.2 | P | 440.8 | P | 656.4 | P | 441.8 | P | --- |
| 265341_at | 59.3 | P | 152.6 | P | 82.2 | A | 126.7 | P | --- |
| 265342_at | 58 | P | 287.7 | P | 167.2 | P | 245.5 | P | --- |
| 265343_at | 249.4 | P | 338.1 | P | 149.4 | P | 273.1 | P | NIC1 |
| 265344_at | 1735.8 | P | 1235.5 | P | 1472.6 | P | 1183.3 | P | --- |
| 265345_at | 77 | P | 54.2 | A | 90.6 | A | 68.3 | A | --- |
| 265346_at | 54.9 | A | 4.7 | A | 43.5 | A | 25 | A | --- |
| 265347_at | 313.2 | P | 305.6 | P | 322.7 | P | 345.6 | P | SUVH6 |
| 265348_at | 29 | A | 23.8 | A | 50.3 | A | 29.3 | A | --- |
| 265349_at | 65.8 | P | 91.1 | P | 74 | P | 95.7 | P | --- |
| 265350_at | 24.8 | A | 31.7 | A | 54.7 | A | 24.9 | A | --- |
| 265351_at | 16.9 | A | 6.3 | A | 11.5 | A | 22 | A | AGL17 |
| 265352_at | 8375.2 | P | 9661.9 | P | 8307.4 | P | 10707.7 | P | ROC3 |
| 265353_at | 889.4 | P | 1149.3 | P | 1165.3 | P | 1123.4 | P | --- |
| 265354_at | 841.6 | P | 1129.5 | P | 768.9 | P | 930.2 | P | ADF5 |
| 265355_at | 34.3 | A | 17.7 | A | 43.7 | A | 49.5 | A | --- |
| 265356_at | 174.7 | P | 143.9 | P | 163.7 | P | 155.4 | P | --- |
| 265357_at | 715.9 | P | 929 | P | 848.5 | P | 907.1 | P | UBC29 |
| 265358_at | 1119.6 | P | 1185.7 | P | 944.4 | P | 932.4 | P | --- |
| 265359_at | 98.8 | P | 107.2 | P | 97.3 | P | 111.5 | P | MYB7 |
| 265360_s_at | 3.6 | A | 9.8 | A | 28.2 | A | 3.5 | A | --- |
| 265361_at | 24 | A | 3 | A | 2.6 | A | 9.1 | A | --- |
| 265362_at | 1.8 | A | 2.9 | A | 2.1 | A | 3.6 | A | --- |
| 265363_at | 0.8 | A | 26.7 | A | 28.3 | A | 17.1 | A | --- |
| 265364_at | 3.1 | A | 8 | A | 23.6 | A | 4.2 | A | --- |
| 265365_at | 6.9 | A | 7.7 | A | 1.3 | A | 8.6 | A | --- |
| 265366_at | 395.2 | P | 306.1 | P | 287.8 | P | 308.6 | P | --- |
| 265367_at | 0.7 | A | 1.5 | A | 1.1 | A | 3.6 | A | --- |
| 265368_at | 16.9 | A | 37.1 | A | 51.4 | A | 21.4 | A | --- |
| 265369_s_at | 4.3 | A | 15.6 | A | 34.6 | A | 6 | A | --- |
| 265370_s_at | 15.2 | A | 3.3 | A | 7.9 | A | 0.8 | A | --- |
| 265371_at | 1.1 | A | 4.4 | A | 1.6 | A | 5.3 | A | --- |
| 265372_at | 19.9 | A | 11 | A | 16.8 | A | 16 | A | --- |
| 265373_at | 97.2 | P | 143.1 | P | 86.7 | P | 113.5 | P | RPA1A |
| 265374_at | 14217.3 | P | 14628.5 | P | 12366.9 | P | 13438.1 | P | PSBX |
| 265375_at | 1383.4 | P | 1395.7 | P | 1118 | P | 1179.7 | P | VPS2.1 |
| 265376_at | 143.1 | P | 220 | P | 188.7 | P | 231.6 | P | --- |
| 265377_at | 2223.1 | P | 2881.6 | P | 2913.9 | P | 3285.9 | P | --- |
| 265378_at | 36.5 | A | 56.6 | A | 53.8 | M | 37.2 | A | --- |
| 265379_at | 2.6 | A | 1.7 | A | 14.7 | A | 2.1 | A | --- |
| 265380_at | 5.9 | A | 1.8 | A | 3.5 | A | 6.1 | A | --- |
| 265381_at | 9.3 | A | 4.1 | A | 11.9 | A | 3 | A | --- |
| 265382_at | 179 | P | 226.2 | P | 133.7 | P | 177.9 | P | --- |
| 265383_at | 390.5 | P | 336.4 | P | 342.8 | P | 347.2 | P | MSI2 |
| 265384_at | 1808.2 | P | 1647.3 | P | 1420.2 | P | 1734.4 | P | --- |
| 265385_at | 528.8 | P | 515.4 | P | 450.9 | P | 547 | P | --- |
| 265386_at | 917.5 | P | 1152.4 | P | 774.7 | P | 1042 | P | --- |
| 265387_at | 245.6 | P | 81.7 | A | 120.5 | A | 76.4 | A | --- |
| 265388_s_at | 1985 | P | 1869.9 | P | 1453.5 | P | 1973.3 | P | --- |

| | | | | | | | | | |
|---|---|---|---|---|---|---|---|---|---|
| 265389_at | 878.4 | P | 881.5 | P | 955.5 | P | 977.5 | P | --- |
| 265390_at | 3877.7 | P | 3473 | P | 4031.4 | P | 3906.4 | P | --- |
| 265391_at | 495.7 | P | 410.4 | P | 423.9 | P | 471 | P | GAUT10 |
| 265392_at | 3092.4 | P | 2836.5 | P | 3060.7 | P | 2830.5 | P | LIP1 |
| 265393_at | 443.4 | P | 488.8 | P | 538.3 | P | 550.1 | P | --- |
| 265394_at | 402.7 | P | 514.3 | P | 473.2 | P | 508.7 | P | --- |
| 265395_at | 330.7 | P | 396.2 | P | 465 | P | 440.8 | P | SRF1 |
| 265396_at | 29.1 | A | 32.8 | A | 8.2 | A | 32.9 | A | --- |
| 265397_at | 5 | A | 2.1 | A | 2.4 | A | 0.6 | A | --- |
| 265398_at | 293.6 | P | 373.6 | P | 323.7 | P | 329.6 | P | ATMAK10 |
| 265399_at | 2.2 | A | 37.4 | A | 8.4 | A | 8.8 | A | --- |
| 265400_at | 3186.9 | P | 4999.6 | P | 4540.3 | P | 5351.3 | P | --- |
| 265401_at | 23.6 | A | 3.6 | A | 56.1 | A | 8.4 | A | --- |
| 265402_at | 493.3 | P | 443.3 | P | 379.2 | P | 391.1 | P | --- |
| 265403_at | 57.7 | P | 64.2 | A | 112.9 | P | 61.7 | A | --- |
| 265404_at | 13.4 | A | 32.4 | A | 26.4 | A | 3.7 | A | BGAL13 |
| 265405_at | 65.8 | P | 79 | P | 61.7 | P | 73.4 | P | --- |
| 265406_at | 15.8 | A | 9 | A | 1 | A | 4.9 | A | --- |
| 265407_at | 11.1 | A | 31.8 | A | 55.5 | A | 29.6 | A | --- |
| 265408_at | 0.6 | A | 1.1 | A | 1.9 | A | 0.4 | A | --- |
| 265409_at | 10.2 | A | 9 | A | 8.2 | A | 27.5 | A | PIP2;8 |
| 265410_at | 1.6 | A | 0.5 | A | 1.2 | A | 0.6 | A | --- |
| 265411_at | 488.4 | P | 705.4 | P | 543.9 | P | 597.7 | P | --- |
| 265412_at | 600.1 | P | 739.2 | P | 730.1 | P | 785 | P | TOC132 |
| 265413_s_at | 421 | P | 283.1 | P | 442.9 | P | 348.8 | P | --- |
| 265414_at | 864.6 | P | 2832.9 | P | 1201.3 | P | 2373.4 | P | --- |
| 265415_at | 5875.1 | P | 7085.6 | P | 6361.9 | P | 6746.7 | P | PSB29 |
| 265416_at | 962.7 | P | 1014.8 | P | 1019.7 | P | 1106.2 | P | --- |
| 265417_at | 1949.7 | P | 1955.2 | P | 2105.5 | P | 1934.8 | P | --- |
| 265418_at | 54.3 | A | 56.6 | A | 41 | A | 62 | A | --- |
| 265419_at | 169.6 | P | 186 | P | 161.3 | P | 224.2 | P | --- |
| 265420_s_at | 111.6 | P | 44.5 | P | 89.7 | P | 51.8 | P | BRX |
| 265421_s_at | 21.1 | A | 2.8 | A | 11.7 | A | 15.9 | A | --- |
| 265422_at | 109.3 | P | 52.3 | A | 101 | P | 22 | A | NDB4 |
| 265423_at | 429.7 | P | 432.2 | P | 406.1 | P | 416.2 | P | --- |
| 265424_at | 414.9 | P | 408 | P | 311.9 | P | 395.7 | P | --- |
| 265425_at | 71.4 | P | 42.2 | M | 87.9 | P | 64.6 | P | GCL2 |
| 265426_at | 811.5 | P | 770.7 | P | 734.8 | P | 728.8 | P | --- |
| 265427_at | 524.6 | P | 531.5 | P | 536.3 | P | 481.6 | P | --- |
| 265428_at | 77.9 | P | 22.1 | A | 68.4 | A | 16.9 | A | --- |
| 265429_at | 120.5 | P | 149 | P | 165.6 | P | 149.4 | P | --- |
| 265430_at | 47.5 | A | 52.2 | A | 52.5 | A | 22.1 | A | --- |
| 265431_at | 150.4 | P | 144.4 | P | 184.3 | P | 141.9 | P | --- |
| 265432_at | 114.9 | P | 161.7 | P | 130.4 | P | 176.1 | P | --- |
| 265433_at | 206.9 | P | 239.2 | P | 279 | P | 263 | P | --- |
| 265434_at | 14.2 | A | 9.7 | A | 23.9 | A | 2.7 | A | --- |
| 265435_s_at | 38.9 | A | 41.1 | A | 40.1 | A | 52.1 | A | --- |
| 265436_at | 1.3 | A | 0.3 | A | 2.3 | A | 1.5 | A | --- |
| 265437_at | 46.4 | A | 105.4 | P | 65.4 | A | 67.8 | P | MCM10 |
| 265438_at | 0.9 | A | 2.3 | A | 6.7 | A | 6.4 | A | --- |
| 265439_at | 25.7 | A | 37.8 | A | 54 | A | 25.6 | A | --- |
| 265440_at | 547.9 | P | 685.2 | P | 385.8 | P | 620.8 | P | pEARLI4 |
| 265441_at | 47.2 | P | 54.7 | P | 73.5 | P | 92.8 | P | --- |
| 265442_at | 467.5 | P | 437.8 | P | 619.4 | P | 508.2 | P | --- |
| 265443_at | 24.9 | A | 75.2 | P | 47.9 | A | 86.2 | M | ATEXPB1 |
| 265444_s_at | 1421.7 | P | 1367.5 | P | 1567.3 | P | 1398.1 | P | RD28 |
| 265445_at | 7941.7 | P | 5905.2 | P | 9419.1 | P | 6286.6 | P | --- |
| 265446_at | 2951.1 | P | 2730.2 | P | 2696.4 | P | 2634.4 | P | --- |
| 265447_at | 41.9 | P | 50.6 | A | 73.7 | A | 47.4 | P | LAC6 |
| 265448_at | 140.2 | P | 175.2 | P | 206.5 | P | 166.2 | P | DAG2 |
| 265449_at | 385.8 | P | 324.4 | P | 347.2 | P | 316.3 | P | --- |
| 265450_at | 363.2 | P | 311 | P | 328.1 | P | 292.3 | P | --- |
| 265451_at | 974.1 | P | 1386.7 | P | 758.9 | P | 1184.9 | P | --- |
| 265452_at | 854.7 | P | 446.2 | P | 599.8 | P | 307.5 | P | ATAIB |
| 265453_at | 1197.4 | P | 859.1 | P | 1465.5 | P | 934.6 | P | --- |
| 265454_at | 47 | P | 94.6 | P | 64.9 | P | 86 | P | ARF11 |
| 265455_at | 596.3 | P | 591.8 | P | 637.9 | P | 603.1 | P | --- |
| 265456_at | 2615.6 | P | 2809 | P | 2790.3 | P | 2790.2 | P | SDH4 |
| 265457_at | 625.1 | P | 655 | P | 544.6 | P | 489.8 | P | --- |
| 265458_at | 802.6 | P | 792.2 | P | 707.6 | P | 872.1 | P | --- |
| 265459_at | 2728.5 | P | 2359.9 | P | 2247.4 | P | 2663.5 | P | --- |
| 265460_at | 5570.4 | P | 4350.7 | P | 4134.7 | P | 3711.9 | P | --- |
| 265461_at | 578.5 | P | 700.3 | P | 486.9 | P | 582.8 | P | --- |
| 265462_at | 7.9 | A | 0.5 | A | 0.6 | A | 0.9 | A | --- |
| 265463_at | 71.3 | P | 79.1 | P | 75.7 | P | 99.3 | P | IRX9 |
| 265464_at | 227.2 | P | 195.2 | P | 226.6 | P | 355.9 | P | --- |
| 265465_at | 22.2 | A | 32 | A | 40.7 | A | 6.9 | A | --- |
| 265466_at | 254.5 | P | 115.6 | P | 172.6 | P | 202.6 | P | NF-YB8 |
| 265467_at | 475.1 | P | 729.1 | P | 715.7 | P | 728.9 | P | --- |
| 265468_at | 59.1 | A | 38 | A | 75 | A | 39.7 | A | --- |
| 265469_at | 174.9 | P | 236.8 | P | 199.3 | P | 209.4 | P | --- |
| 265470_at | 88.8 | P | 106.9 | P | 116 | P | 98.5 | P | --- |
| 265471_at | 77.7 | P | 435.5 | P | 137 | P | 291.1 | P | --- |
| 265472_at | 1094.9 | P | 1137.2 | P | 1111.6 | P | 1065.3 | P | --- |
| 265473_at | 1.3 | A | 2.1 | A | 15.2 | A | 13.2 | A | LCR10 |
| 265474_at | 503 | P | 510.7 | P | 592.6 | P | 477.6 | P | --- |
| 265475_at | 622.2 | P | 451.7 | P | 648.6 | P | 763.9 | P | NIR1 |
| 265476_at | 408.2 | P | 588.3 | P | 452.3 | P | 506.2 | P | ATHM3 |
| 265477_at | 25.4 | A | 4.4 | A | 54.7 | A | 36.3 | P | GAUT2 |
| 265478_at | 1790.2 | P | 1812.3 | P | 1619.1 | P | 1716.4 | P | MEE14 |
| 265479_at | 236.9 | P | 82.5 | P | 163 | P | 123.3 | P | --- |
| 265480_at | 3889.2 | P | 6514.8 | P | 3640.3 | P | 6041.2 | P | COR413-PM1 |
| 265481_at | 1331.1 | P | 1532 | P | 821.7 | P | 975.8 | P | --- |
| 265482_at | 21.8 | A | 21.6 | A | 24.1 | A | 6.5 | A | --- |
| 265483_at | 119.2 | P | 102.4 | P | 96 | P | 124.4 | P | SQN |
| 265484_at | 411.2 | P | 593.1 | P | 446.2 | P | 478.7 | P | OTP51 |
| 265485_at | 3.8 | A | 2.5 | A | 4.7 | A | 9 | A | --- |
| 265486_at | 27.7 | A | 43.5 | A | 36.1 | A | 46.6 | A | --- |
| 265487_at | 0.4 | A | 0.4 | A | 2.3 | A | 1.1 | A | --- |
| 265488_at | 2.7 | A | 4.6 | A | 6.1 | A | 3.5 | A | --- |
| 265489_at | 1.5 | A | 1.6 | A | 13.1 | A | 3.4 | A | --- |
| 265490_s_at | 8.3 | A | 0.3 | A | 0.8 | A | 1.4 | A | --- |
| 265491_s_at | 2.2 | A | 5.3 | A | 14 | A | 14 | A | --- |
| 265492_at | 23.7 | A | 15 | A | 33.4 | A | 13.2 | A | AGL95 |
| 265493_at | 0.6 | A | 16 | A | 7.9 | A | 4.7 | A | --- |
| 265494_at | 157.4 | P | 219.3 | P | 272 | P | 196 | P | --- |
| 265495_at | 728.7 | P | 821 | P | 796.3 | P | 765.8 | P | --- |
| 265496_at | 3.4 | A | 0.8 | A | 11.1 | A | 0.6 | A | --- |
| 265497_at | 23.5 | A | 3.3 | A | 17.3 | A | 30.5 | A | --- |

| | | | | | | | | | |
|---|---|---|---|---|---|---|---|---|---|
| 265498_at | 26 | M | 17.4 | M | 46.7 | P | 17.4 | A | --- |
| 265499_at | 402.5 | P | 340.1 | P | 275.8 | P | 306.6 | P | UGT73B5 |
| 265500_at | 30.6 | A | 32 | A | 37.2 | A | 38.8 | A | --- |
| 265501_at | 528.1 | P | 152.1 | P | 492.8 | P | 202.9 | P | UGT73B4 |
| 265502_at | 38.3 | A | 38.1 | A | 36.2 | A | 36.5 | A | --- |
| 265503_at | 17.7 | A | 17.5 | A | 8.4 | A | 12 | A | --- |
| 265504_at | 22.3 | A | 37.6 | P | 57.8 | P | 36.9 | P | --- |
| 265505_at | 389.9 | P | 348.2 | P | 314.3 | P | 387.8 | P | --- |
| 265506_at | 30 | A | 4.9 | A | 2.8 | A | 28.9 | A | --- |
| 265507_s_at | 0.7 | A | 1 | A | 3.4 | A | 1.3 | A | --- |
| 265508_at | 4.9 | A | 1.9 | A | 1.8 | A | 4.9 | A | --- |
| 265509_at | 30.1 | A | 3 | A | 14.1 | A | 13 | A | --- |
| 265510_at | 669.1 | P | 673.5 | P | 814.7 | P | 661.5 | P | ATG8D |
| 265511_at | 538 | P | 959.6 | P | 389.8 | P | 667.1 | P | --- |
| 265512_at | 357.1 | P | 444.2 | P | 296.6 | P | 369.2 | P | --- |
| 265513_at | 1.9 | A | 5.5 | A | 2.4 | A | 4 | A | --- |
| 265514_s_at | 1.2 | A | 1.9 | A | 8.7 | A | 4.6 | A | --- |
| 265515_at | 0.8 | A | 9.6 | A | 3 | A | 1.4 | A | --- |
| 265516_at | 1.1 | A | 0.7 | A | 0.9 | A | 1 | A | --- |
| 265517_at | 9.2 | A | 16.7 | A | 18.5 | A | 8.6 | A | --- |
| 265518_at | 76.2 | P | 61.8 | M | 76.8 | P | 92.1 | P | --- |
| 265519_at | 48.1 | P | 59.6 | P | 97.7 | P | 92.4 | P | --- |
| 265520_at | 12.7 | A | 13.2 | A | 28.1 | A | 4.8 | A | --- |
| 265521_at | 310.3 | P | 299.1 | P | 438.1 | P | 326.8 | P | ORG4 |
| 265522_at | 229.7 | P | 213 | P | 237.2 | P | 274.6 | P | ELF8 |
| 265523_at | 9.3 | A | 8.1 | A | 67.5 | A | 8.8 | A | --- |
| 265524_at | 3.2 | A | 1.4 | A | 10.4 | A | 6.6 | A | --- |
| 265525_at | 26.9 | A | 5.7 | A | 41.3 | A | 16.8 | A | --- |
| 265526_x_at | 17.9 | P | 4.6 | A | 54.2 | P | 17 | A | --- |
| 265527_at | 0.6 | A | 1.5 | A | 2.3 | A | 2.4 | A | --- |
| 265528_at | 1 | A | 12.5 | A | 52.9 | A | 16.5 | A | --- |
| 265529_at | 2.3 | A | 27.5 | A | 30.2 | A | 12.8 | A | --- |
| 265530_at | 5307.1 | P | 2876.9 | P | 6234 | P | 3205.8 | P | OPR3 |
| 265531_at | 3.4 | A | 22.2 | A | 3.5 | A | 21.4 | A | AtGRF6 |
| 265532_at | 3.8 | A | 5.9 | A | 3.5 | A | 3.3 | A | --- |
| 265533_at | 15.1 | P | 19.6 | A | 14.3 | A | 19.8 | A | --- |
| 265534_at | 18.4 | A | 14.2 | A | 7.3 | A | 8.5 | A | --- |
| 265535_at | 243.3 | P | 244 | P | 284.5 | P | 256 | P | --- |
| 265536_at | 37 | A | 71.7 | A | 70.6 | P | 44.7 | A | --- |
| 265537_at | 9.2 | A | 4.6 | A | 10.4 | A | 5.9 | A | --- |
| 265538_at | 286.5 | P | 349.7 | P | 313.6 | P | 305.5 | P | --- |
| 265539_at | 29.8 | P | 159 | P | 33.6 | A | 103.2 | P | --- |
| 265540_at | 3.8 | A | 5.8 | A | 5.8 | A | 4.6 | A | --- |
| 265541_at | 0.6 | A | 0.8 | A | 3.2 | A | 1 | A | SQN |
| 265542_at | 469.6 | P | 438.7 | P | 379.4 | P | 421.1 | P | SYD |
| 265543_at | 16 | A | 17.9 | A | 7.9 | A | 6.9 | A | --- |
| 265544_at | 71.6 | P | 112.4 | P | 82 | P | 109.5 | P | ATCNGC15 |
| 265545_at | 37.3 | A | 45.4 | P | 24.7 | A | 58.5 | A | NCRK |
| 265546_at | 70.4 | P | 70.1 | A | 98.2 | A | 53.1 | P | --- |
| 265547_at | 200.7 | P | 354.9 | P | 339.2 | P | 384.8 | P | --- |
| 265548_s_at | 20 | A | 0.9 | A | 1.7 | A | 3.9 | A | --- |
| 265549_at | 0.7 | A | 4.7 | A | 1.7 | A | 1.4 | A | --- |
| 265550_at | 2.7 | A | 11.4 | A | 4.9 | A | 2.8 | A | --- |
| 265551_at | 0.8 | A | 4.3 | A | 5.1 | A | 1.3 | A | --- |
| 265552_at | 5.4 | A | 2.4 | A | 25.8 | A | 1.6 | A | AHA6 |
| 265553_at | 0.8 | A | 3 | A | 1.2 | A | 2.8 | A | --- |
| 265554_at | 11.4 | A | 2.6 | A | 27.8 | A | 16.2 | A | --- |
| 265555_at | 14.8 | A | 30.6 | A | 5.4 | A | 6.2 | A | --- |
| 265556_at | 2.9 | A | 3.4 | A | 2 | A | 13.7 | A | --- |
| 265557_at | 21.9 | A | 24.9 | A | 13.4 | A | 18.6 | A | --- |
| 265558_at | 1.4 | A | 1 | A | 1.6 | A | 9.4 | A | --- |
| 265559_at | 18.7 | A | 20.4 | A | 65.8 | M | 25.8 | A | --- |
| 265560_at | 20006.2 | P | 19369.4 | P | 15602.6 | P | 21257.7 | P | GRP-3 |
| 265561_s_at | 18.7 | A | 63.3 | P | 85.7 | P | 70.7 | P | --- |
| 265562_at | 0.1 | A | 0.2 | A | 0.5 | A | 0.3 | A | --- |
| 265563_s_at | 5.8 | A | 19.9 | A | 15.6 | A | 14.3 | A | --- |
| 265564_s_at | 1.2 | A | 1 | A | 12.6 | A | 1.3 | A | --- |
| 265565_at | 3.4 | A | 26 | A | 49.9 | A | 37.1 | P | --- |
| 265566_at | 63.4 | A | 48.1 | A | 67.2 | A | 47.5 | A | --- |
| 265567_at | 1.7 | A | 3.4 | A | 5.1 | A | 5 | A | --- |
| 265568_s_at | 8.9 | A | 1.2 | A | 1.8 | A | 1.1 | A | --- |
| 265569_at | 1814.6 | P | 2465.7 | P | 2236.4 | P | 2348.2 | P | PGR5 |
| 265570_at | 364.3 | P | 476.6 | P | 486.8 | P | 495.5 | P | --- |
| 265571_s_at | 987.5 | P | 850.4 | P | 525.4 | P | 891 | P | --- |
| 265572_at | 91.8 | P | 389.9 | P | 69.4 | A | 259.9 | P | ATACA2 |
| 265573_at | 175.5 | P | 166 | P | 96.6 | P | 189.9 | P | --- |
| 265574_at | 1.7 | A | 1.4 | A | 0.9 | A | 0.8 | A | --- |
| 265575_at | 1573.4 | P | 1762.9 | P | 1694.4 | P | 1757.7 | P | PIP |
| 265576_at | 880.3 | P | 898.6 | P | 906.2 | P | 947.3 | P | CLASP |
| 265577_at | 67.4 | P | 77.1 | P | 52.2 | P | 47.1 | P | --- |
| 265578_at | 40.8 | A | 70.7 | P | 33 | A | 48.3 | P | --- |
| 265579_at | 15 | A | 2 | A | 3.7 | A | 1.9 | A | --- |
| 265580_at | 211.2 | P | 227.2 | P | 234.7 | P | 243.1 | P | --- |
| 265581_at | 569.6 | P | 525.1 | P | 666.3 | P | 532 | P | --- |
| 265582_at | 12.9 | A | 25.3 | P | 18.6 | A | 1.7 | A | --- |
| 265583_at | 363.9 | P | 387.9 | P | 424 | P | 348.3 | P | --- |
| 265584_at | 529.5 | P | 297.5 | P | 448.4 | P | 308.5 | P | PIL5 |
| 265585_at | 277.9 | P | 385.5 | P | 361.9 | P | 369.6 | P | HBT |
| 265586_at | 9.3 | A | 44.1 | A | 61.9 | A | 101 | P | PR-1-LIKE |
| 265587_at | 6.3 | A | 24 | A | 2.9 | A | 30 | A | --- |
| 265588_at | 83.2 | A | 121.6 | A | 114.8 | A | 93.7 | A | --- |
| 265589_at | 3.9 | A | 28.5 | A | 4.6 | A | 3.3 | A | --- |
| 265590_at | 17.8 | A | 11.9 | A | 37.8 | A | 23.6 | A | MEO |
| 265591_at | 31.5 | A | 25.6 | A | 22.5 | A | 23.5 | A | --- |
| 265592_at | 39.7 | P | 68.5 | P | 89.6 | P | 54.7 | P | --- |
| 265593_s_at | 803 | P | 945.8 | P | 991.2 | P | 774 | P | LCV1 |
| 265594_at | 1967.9 | P | 1523.8 | P | 2662.2 | P | 1791.8 | P | --- |
| 265595_at | 774.8 | P | 775.4 | P | 573.8 | P | 810.5 | P | --- |
| 265596_at | 595.9 | P | 564.2 | P | 552.3 | P | 607.9 | P | CAF1 |
| 265597_at | 266.3 | P | 337.7 | P | 218 | P | 186.1 | P | --- |
| 265598_at | 13.1 | A | 0.5 | A | 31.9 | A | 1.2 | A | --- |
| 265599_at | 181.2 | P | 254.4 | P | 229.8 | M | 201.4 | P | AT2G14255 |
| 265600_at | 0.8 | A | 4.5 | A | 6 | A | 15 | P | --- |
| 265601_at | 31.7 | A | 15.9 | A | 19.7 | A | 38.7 | A | --- |
| 265602_at | 3.2 | A | 6.1 | A | 2.5 | A | 5.7 | A | --- |
| 265603_at | 1.7 | A | 24.9 | A | 3.5 | A | 2.5 | A | --- |
| 265604_s_at | 0.6 | A | 1.2 | A | 1.2 | A | 1.4 | A | --- |
| 265605_at | 38.8 | A | 52 | P | 100.3 | A | 55.2 | A | CESA10 |
| 265606_s_at | 1.4 | A | 0.6 | A | 0.8 | A | 0.9 | A | --- |

| | | | | | | | | | |
|---|---|---|---|---|---|---|---|---|---|
| 265607_at | 1.9 | A | 1.6 | A | 3 | A | 1.7 | A | --- |
| 265608_at | 20.7 | A | 28.9 | A | 35.6 | A | 7.4 | A | --- |
| 265609_at | 56.3 | A | 70.8 | P | 81.9 | A | 99.6 | P | --- |
| 265610_at | 1116.6 | P | 1206.1 | P | 1197.8 | P | 1076.3 | P | --- |
| 265611_at | 5560 | P | 16223.5 | P | 4343.1 | P | 14642.7 | P | --- |
| 265612_at | 3.4 | A | 8.3 | A | 4.4 | A | 5.8 | A | --- |
| 265613_at | 244.9 | P | 327.3 | P | 302.3 | P | 228.5 | P | --- |
| 265614_at | 314.4 | P | 252.5 | P | 419.4 | P | 264.1 | P | --- |
| 265615_at | 12756 | P | 7422.6 | P | 12628 | P | 8117.9 | P | --- |
| 265616_at | 15.5 | P | 0.3 | A | 19.7 | A | 12.8 | A | --- |
| 265617_at | 1754.1 | P | 1811.2 | P | 1554.5 | P | 1756.3 | P | --- |
| 265618_at | 40.4 | A | 46.8 | A | 27.6 | A | 24.4 | A | --- |
| 265619_at | 28.5 | P | 45.8 | P | 51 | A | 36.7 | A | --- |
| 265620_at | 84.6 | P | 83.7 | P | 84.2 | P | 60 | P | --- |
| 265621_at | 3.5 | A | 17.3 | A | 4.2 | A | 2.4 | A | NTL8 |
| 265622_at | 12.1 | A | 2.8 | A | 10.9 | A | 4.3 | A | --- |
| 265623_at | 35.3 | A | 27.7 | A | 27.6 | P | 30.2 | A | --- |
| 265624_at | 4.9 | A | 2.9 | A | 9.4 | A | 3.8 | A | CLV3 |
| 265625_at | 9.4 | A | 4.1 | A | 5.8 | A | 6.6 | A | --- |
| 265626_at | 832.9 | P | 690.7 | P | 677.8 | P | 629.2 | P | --- |
| 265627_at | 520.6 | P | 440 | P | 590.1 | P | 497.3 | P | --- |
| 265628_at | 1918.8 | P | 1984.9 | P | 1530.3 | P | 1836.5 | P | --- |
| 265629_at | 670.8 | P | 422.8 | P | 998.9 | P | 772.6 | P | LHW |
| 265630_at | 493.6 | P | 521.2 | P | 527.6 | P | 538.7 | P | --- |
| 265631_at | 5.8 | A | 7.6 | A | 11.1 | A | 6.7 | A | --- |
| 265632_at | 109.9 | P | 10.9 | A | 102.1 | A | 18.9 | A | --- |
| 265633_at | 1164.3 | P | 1346.9 | P | 1320.5 | P | 1206.4 | P | EBF1 |
| 265634_at | 215.5 | P | 188.9 | P | 257.5 | P | 228.1 | P | --- |
| 265635_at | 13.3 | A | 2.4 | A | 13.4 | A | 1.1 | A | --- |
| 265636_at | 130.2 | P | 119.3 | P | 103.8 | A | 83.6 | M | --- |
| 265637_at | 341.9 | P | 304.9 | P | 214.2 | P | 282.6 | P | ATCOAE |
| 265638_at | 118.8 | P | 156.9 | P | 180.2 | P | 117 | P | --- |
| 265639_at | 19.7 | A | 21.6 | A | 25 | A | 27 | P | --- |
| 265640_at | 8.1 | A | 3.6 | A | 3.9 | A | 9 | A | --- |
| 265641_at | 292.8 | P | 328.6 | P | 263 | P | 394.5 | P | --- |
| 265642_at | 1023.9 | P | 1245.1 | P | 1043.8 | P | 1041.8 | P | NLP1 |
| 265643_at | 42.4 | P | 37.1 | A | 2.5 | A | 22.8 | A | --- |
| 265644_at | 26.9 | A | 77.3 | P | 71.9 | A | 73.8 | M | ATEPR1 |
| 265645_at | 18.2 | A | 18.3 | A | 10.9 | A | 5.4 | A | --- |
| 265646_at | 426.8 | P | 462.1 | P | 503.9 | P | 508.1 | P | --- |
| 265647_at | 26.5 | A | 32 | A | 22.5 | A | 38.7 | A | --- |
| 265648_at | 734.6 | P | 577.8 | P | 524.6 | P | 459.5 | P | --- |
| 265649_at | 1980.6 | P | 1358.7 | P | 2066.2 | P | 1374.7 | P | ATFD3 |
| 265650_at | 414.5 | P | 426.5 | P | 517.9 | P | 423 | P | --- |
| 265651_at | 1.1 | A | 1.3 | A | 1.4 | A | 1.2 | A | --- |
| 265652_x_at | 0.8 | A | 10.2 | A | 2.2 | A | 4.3 | A | --- |
| 265653_at | 6.1 | A | 3.9 | A | 4.1 | A | 1.9 | A | --- |
| 265654_s_at | 26.7 | A | 19.4 | A | 11.6 | A | 6.8 | A | --- |
| 265655_at | 1.8 | A | 28.9 | A | 18.9 | A | 19.1 | A | --- |
| 265656_at | 292.1 | P | 518.4 | P | 471.7 | P | 461.6 | P | --- |
| 265657_at | 135.7 | P | 171.1 | P | 221.3 | P | 178.3 | P | --- |
| 265658_at | 31.8 | A | 17.3 | A | 36 | A | 30.2 | A | ALD1 |
| 265659_at | 11.2 | A | 22.3 | P | 21.9 | A | 28.2 | A | AtRLP20 |
| 265660_at | 10.3 | A | 56.6 | P | 27.8 | A | 52.6 | P | AtRLP21 |
| 265661_at | 1368.4 | P | 1543.4 | P | 1217.8 | P | 1420.3 | P | --- |
| 265662_at | 1261.9 | P | 863.3 | P | 1311.3 | P | 1002.3 | P | FZF |
| 265663_at | 356.8 | P | 555.3 | P | 423.8 | P | 561.1 | P | --- |
| 265664_at | 1221.9 | P | 1223.1 | P | 1122.6 | P | 1017.2 | P | --- |
| 265665_at | 40.8 | P | 43 | P | 53.7 | P | 46.8 | P | --- |
| 265666_at | 1.2 | A | 1.5 | A | 11 | A | 7.4 | A | --- |
| 265667_at | 74.9 | P | 90.6 | P | 125.4 | P | 94.5 | P | NF-YB11 |
| 265668_at | 55.1 | P | 43.7 | A | 82.8 | P | 28.4 | A | --- |
| 265669_at | 519.6 | P | 590.4 | P | 631 | P | 507.5 | P | --- |
| 265670_s_at | 556.3 | P | 397.1 | P | 330.9 | P | 279.7 | P | --- |
| 265671_at | 3016 | P | 2902.4 | P | 3416.3 | P | 3442.7 | P | --- |
| 265672_at | 298.9 | P | 118.3 | P | 254.7 | P | 128.3 | P | --- |
| 265673_at | 1132.1 | P | 995.4 | P | 858.7 | P | 847.7 | P | --- |
| 265674_at | 333 | P | 169.7 | P | 119.3 | P | 90.6 | P | --- |
| 265675_at | 118.4 | P | 130.9 | P | 171.5 | P | 117.2 | P | HSP70T-2 |
| 265676_at | 977.8 | P | 924.2 | P | 928.7 | P | 919.2 | P | --- |
| 265677_at | 1664.8 | P | 1377 | P | 1819.8 | P | 1445.7 | P | PUR ALPHA-1 |
| 265678_at | 281.7 | P | 256.4 | P | 231.2 | P | 227.3 | P | RAD50 |
| 265679_at | 1223.2 | P | 1209.7 | P | 719.4 | P | 1017.8 | P | --- |
| 265680_at | 2244.7 | P | 1046.3 | P | 1852.2 | P | 1094.1 | P | --- |
| 265681_at | 3.8 | A | 21.3 | M | 14.1 | A | 1 | A | --- |
| 265682_at | 109 | P | 196.5 | P | 112 | P | 134.4 | P | --- |
| 265683_at | 2.8 | A | 4.9 | A | 3.7 | A | 11.8 | A | --- |
| 265684_at | 8.9 | A | 4.9 | A | 19.3 | A | 0.5 | A | --- |
| 265685_at | 34.8 | P | 40.3 | P | 67.6 | M | 42.7 | P | ANAC038 |
| 265686_at | 0.9 | A | 2.1 | A | 2.4 | A | 2.7 | A | --- |
| 265687_s_at | 19.3 | A | 19.9 | A | 16.5 | A | 28.5 | P | --- |
| 265688_at | 30 | A | 44.9 | P | 27.6 | A | 29 | A | --- |
| 265689_at | 29.9 | A | 40.9 | A | 57.5 | M | 67.5 | A | --- |
| 265690_at | 24.2 | A | 25 | A | 20 | A | 20.9 | A | --- |
| 265691_at | 326.2 | P | 363.5 | P | 365.4 | P | 325.7 | P | --- |
| 265692_at | 6.1 | A | 1.1 | A | 3.1 | A | 1.5 | A | --- |
| 265693_at | 26.6 | A | 9.1 | A | 4.1 | A | 34.1 | A | --- |
| 265694_at | 87.7 | P | 118.7 | P | 91.2 | P | 80 | P | --- |
| 265695_at | 227.8 | P | 280.7 | P | 289.6 | P | 313.7 | P | RPA2 |
| 265696_at | 37.1 | A | 36.3 | A | 62 | A | 15.3 | A | CNX2 |
| 265697_at | 971.5 | P | 1019.8 | P | 890.4 | P | 929.7 | P | --- |
| 265698_at | 33.1 | P | 159 | P | 31 | P | 128.3 | P | --- |
| 265699_at | 923.3 | P | 673.4 | P | 976.4 | P | 817.3 | P | --- |
| 265700_at | 592.8 | P | 610.9 | P | 491.3 | P | 560.3 | P | --- |
| 265701_at | 21.2 | A | 24.9 | A | 7.6 | A | 4 | A | --- |
| 265702_at | 130.9 | P | 154.4 | P | 153.5 | P | 116.9 | P | PAP9 |
| 265703_at | 305.3 | P | 298.1 | P | 290.4 | P | 349.1 | P | --- |
| 265704_at | 1869.9 | P | 1645.7 | P | 2198.5 | P | 1987.7 | P | --- |
| 265705_at | 11.6 | A | 3.8 | A | 44.2 | A | 18.7 | A | --- |
| 265706_at | 478.7 | P | 582.2 | P | 549.7 | P | 571.7 | P | --- |
| 265707_at | 1778.6 | P | 1604.3 | P | 1629.1 | P | 1525.1 | P | --- |
| 265708_at | 74.9 | P | 56.4 | A | 29.7 | A | 59.8 | P | --- |
| 265709_at | 3.2 | A | 79.2 | P | 3.6 | A | 27.1 | A | --- |
| 265710_at | 2.7 | A | 5.3 | A | 14.4 | A | 5.2 | A | --- |
| 265711_at | 3.9 | A | 3.2 | A | 4.1 | A | 19 | A | --- |
| 265712_s_at | 1133.3 | P | 1187.3 | P | 1263.1 | P | 1274.2 | P | --- |
| 265713_at | 292.3 | P | 487.4 | P | 266.4 | P | 315.7 | P | UPS2 |
| 265714_at | 4 | A | 4.9 | A | 5 | A | 3.1 | A | ATUPS4 |
| 265715_s_at | 411.5 | P | 477.6 | P | 414.5 | P | 511.9 | P | --- |

| | | | | | | | | | |
|---|---|---|---|---|---|---|---|---|---|
| 265716_at | 380.4 | P | 456.6 | P | 488.9 | P | 413.4 | P | --- |
| 265717_at | 1148.1 | P | 1232.4 | P | 1199.4 | P | 1142.5 | P | --- |
| 265718_at | 571.1 | P | 393 | P | 459.9 | P | 453.3 | P | WRKY3 |
| 265719_at | 50 | A | 70.6 | P | 91.5 | A | 67.2 | P | --- |
| 265720_at | 228 | P | 290.1 | P | 233.1 | P | 235.7 | P | --- |
| 265721_at | 345.3 | P | 362 | P | 344.6 | P | 306.4 | P | ATATH9 |
| 265722_at | 860.5 | P | 1517.7 | P | 739.1 | P | 1641.1 | P | LHCB4.3 |
| 265723_at | 168 | P | 100.8 | P | 98.2 | P | 60.9 | P | --- |
| 265724_at | 87.8 | P | 119 | P | 217.1 | P | 67.3 | P | OFP16 |
| 265725_at | 110.1 | P | 65.1 | P | 101.5 | P | 64.4 | P | --- |
| 265726_at | 102.7 | P | 95.6 | P | 140.6 | P | 78.9 | P | --- |
| 265727_at | 232.8 | P | 222.3 | P | 310.5 | P | 234.8 | P | --- |
| 265728_at | 17.2 | P | 59 | P | 56 | P | 47.5 | P | --- |
| 265729_at | 555.9 | P | 529.9 | P | 508 | P | 585.3 | P | ATGSL03 |
| 265730_at | 676.8 | P | 525.7 | P | 728.2 | P | 616.3 | P | --- |
| 265731_at | 92.2 | P | 97 | P | 110.1 | P | 88.8 | P | --- |
| 265732_at | 30.6 | A | 44 | A | 13.2 | A | 32.2 | A | --- |
| 265733_at | 385.5 | P | 315.7 | P | 372.5 | P | 297.6 | P | --- |
| 265734_at | 276.5 | P | 402.3 | P | 302.7 | P | 364.7 | P | --- |
| 265735_at | 2619 | P | 2189.8 | P | 2581.7 | P | 2326.9 | P | --- |
| 265736_at | 8907.5 | P | 8641.4 | P | 9012.6 | P | 9204.7 | P | --- |
| 265737_at | 375 | P | 235 | P | 295.7 | P | 239 | P | ATPAP1 |
| 265738_at | 2007.8 | P | 1693.7 | P | 1772 | P | 1758.5 | P | QPT |
| 265739_at | 929.7 | P | 982.9 | P | 927.8 | P | 881.2 | P | AtQSOX2 |
| 265740_at | 629.7 | P | 450 | P | 579 | P | 489.5 | P | RHA2B |
| 265741_at | 714.6 | P | 854.6 | P | 785.3 | P | 814.4 | P | --- |
| 265742_at | 790.6 | P | 921.4 | P | 1111.1 | P | 1059.2 | P | --- |
| 265743_s_at | 11.4 | A | 1 | A | 1.6 | A | 2.1 | A | --- |
| 265744_at | 10.1 | A | 7.4 | A | 2.6 | A | 16.1 | A | --- |
| 265745_at | 0.9 | A | 1.4 | A | 3.2 | A | 0.5 | A | --- |
| 265746_at | 1.3 | A | 9 | A | 0.3 | A | 0.1 | A | --- |
| 265747_s_at | 31.7 | A | 19.8 | A | 31.3 | A | 6.7 | A | --- |
| 265748_at | 2 | A | 0.9 | A | 11 | A | 8 | A | --- |
| 265749_at | 1.1 | A | 1.3 | A | 2.4 | A | 0.9 | A | --- |
| 265750_x_at | 43.4 | A | 37.1 | A | 34.7 | A | 47.4 | P | --- |
| 265751_at | 10.9 | A | 11.9 | A | 4.7 | A | 8.9 | A | --- |
| 265752_at | 0.9 | A | 0.8 | A | 12.8 | A | 1 | A | --- |
| 265753_at | 1.6 | A | 8.8 | A | 20.2 | A | 1.1 | A | --- |
| 265754_x_at | 37.7 | P | 47.5 | P | 76.2 | P | 41.5 | P | --- |
| 265755_at | 2.2 | A | 7 | A | 1.2 | A | 2.1 | A | --- |
| 265756_at | 22.4 | A | 25.8 | A | 2.8 | A | 15.5 | A | --- |
| 265757_x_at | 20.9 | A | 7 | A | 38.7 | A | 4.8 | A | --- |
| 265758_at | 8.9 | A | 2.3 | A | 17.3 | A | 0.9 | A | --- |
| 265759_at | 620 | P | 757.1 | P | 530.5 | P | 616.9 | P | --- |
| 265760_at | 829.9 | P | 654.2 | P | 890.3 | P | 705.7 | P | --- |
| 265761_at | 159.4 | P | 138.9 | P | 224.3 | P | 131.9 | P | --- |
| 265762_at | 4.3 | A | 5.7 | A | 7.1 | A | 1.5 | A | --- |
| 265763_at | 272.3 | P | 264.9 | P | 188.3 | P | 308.1 | P | --- |
| 265764_at | 70.2 | P | 125.9 | P | 101.7 | P | 131.5 | P | --- |
| 265765_at | 18.8 | A | 40.4 | A | 84.5 | A | 29.4 | A | --- |
| 265766_at | 49.4 | A | 58.3 | P | 69.5 | M | 48.4 | P | --- |
| 265767_at | 104.2 | P | 115.1 | P | 106.3 | P | 91.8 | P | REF4 |
| 265768_at | 1346.2 | P | 636.6 | P | 1022.5 | P | 692.4 | P | --- |
| 265769_at | 52.2 | P | 57.1 | P | 49.3 | A | 83.3 | P | --- |
| 265770_at | 769.3 | P | 686.9 | P | 721.3 | P | 723.3 | P | --- |
| 265771_at | 130.4 | P | 157 | P | 170.2 | P | 166.3 | P | --- |
| 265772_at | 290.9 | P | 149.8 | P | 167.8 | P | 159.9 | P | RKF3 |
| 265773_at | 1008.2 | P | 1390.4 | P | 970.8 | P | 1198.5 | P | RPH1 |
| 265774_at | 5.9 | A | 23 | A | 27.5 | A | 19.1 | A | --- |
| 265775_at | 26.6 | A | 22.4 | A | 17.3 | A | 23.2 | A | --- |
| 265776_at | 287.5 | P | 240.4 | P | 263.1 | P | 290.2 | P | --- |
| 265777_at | 837.1 | P | 719.1 | P | 1055.6 | P | 665.8 | P | --- |
| 265778_at | 1028.2 | P | 1364.2 | P | 1428 | P | 1262.4 | P | --- |
| 265779_at | 310.3 | P | 472.8 | P | 321.1 | P | 443.4 | P | --- |
| 265780_at | 2 | A | 10.3 | A | 4.1 | A | 10.9 | A | --- |
| 265781_at | 11.4 | A | 17.1 | A | 33.3 | A | 3.4 | A | --- |
| 265782_at | 13.4 | A | 27.1 | A | 4.5 | A | 8.6 | A | --- |
| 265783_at | 2.8 | A | 18.3 | A | 10.9 | A | 10.7 | A | --- |
| 265784_at | 0.9 | A | 2.6 | A | 1.9 | A | 1.3 | A | --- |
| 265785_at | 15.2 | A | 3 | A | 3 | A | 13.9 | A | --- |
| 265786_at | 9.9 | A | 12.8 | A | 24.9 | A | 2.8 | A | --- |
| 265787_at | 29 | A | 21.2 | M | 15 | A | 10.5 | A | --- |
| 265788_at | 15 | A | 12 | A | 2.1 | A | 11.7 | A | IAA32 |
| 265789_at | 59.2 | P | 82.5 | P | 73.3 | P | 77.8 | P | --- |
| 265790_at | 601.5 | P | 652.4 | P | 506.6 | P | 660.7 | P | BAT1 |
| 265791_at | 3.4 | A | 8.4 | A | 12.6 | A | 13.3 | A | MEE65 |
| 265792_at | 31.6 | A | 24.6 | A | 47.9 | A | 16.6 | A | --- |
| 265793_at | 595.3 | P | 612.2 | P | 666.5 | P | 631.2 | P | --- |
| 265794_at | 6 | A | 1.4 | A | 2.7 | A | 2.1 | A | --- |
| 265795_at | 1414.5 | P | 1614.8 | P | 1017.5 | P | 1541.9 | P | scpl26 |
| 265796_at | 67.1 | P | 3.1 | A | 23.6 | A | 3.8 | A | --- |
| 265797_at | 4.2 | A | 1.5 | A | 32.4 | A | 10.8 | A | --- |
| 265798_at | 861.1 | P | 630.1 | P | 904.3 | P | 721.7 | P | --- |
| 265799_at | 1495.4 | P | 1557.6 | P | 1541.3 | P | 1498.9 | P | --- |
| 265800_at | 520.3 | P | 491.2 | P | 402.2 | P | 501.5 | P | MOR1 |
| 265801_at | 25.4 | A | 3.9 | A | 3.8 | A | 13.6 | A | FIS2 |
| 265802_at | 33.2 | P | 70.8 | P | 79.4 | P | 28.3 | P | --- |
| 265803_at | 9.3 | A | 2.4 | A | 11.1 | A | 8.4 | A | --- |
| 265804_at | 520.2 | P | 517.6 | P | 541.7 | P | 503.3 | P | --- |
| 265805_s_at | 14920.1 | P | 12716.7 | P | 17523.3 | P | 14406.5 | P | EMB2296 |
| 265806_at | 8.9 | A | 11.4 | A | 37.4 | A | 14.3 | A | --- |
| 265807_at | 537.7 | P | 552.8 | P | 477.3 | P | 546.5 | P | --- |
| 265808_at | 4.3 | A | 4.9 | A | 3.5 | A | 4.3 | A | --- |
| 265809_at | 28.5 | A | 33.8 | A | 71.3 | A | 56.1 | A | --- |
| 265810_at | 574.5 | P | 541.7 | P | 672.2 | P | 608.5 | P | --- |
| 265811_at | 15.1 | A | 5.5 | A | 6.9 | A | 17.8 | A | EDA2 |
| 265812_at | 3.4 | A | 23.9 | A | 7.1 | A | 9.8 | A | --- |
| 265813_at | 44.5 | P | 28.2 | A | 42.3 | P | 23.9 | A | VND1 |
| 265814_at | 715 | P | 689.3 | P | 606 | P | 617.2 | P | --- |
| 265815_at | 202.4 | P | 233.8 | P | 216.9 | P | 220.5 | P | --- |
| 265816_s_at | 12887.6 | P | 9444 | P | 12766.4 | P | 10254.5 | P | --- |
| 265817_at | 166.5 | P | 86.2 | P | 130.5 | P | 61.2 | P | HIS1-3 |
| 265818_at | 2806.7 | P | 2788.6 | P | 2580 | P | 2696.5 | P | PIN1AT |
| 265819_at | 390.4 | P | 598.5 | P | 638.9 | P | 656.4 | P | --- |
| 265820_at | 33.4 | A | 15.6 | A | 3.6 | A | 9.7 | A | --- |
| 265821_at | 8.3 | A | 18.7 | A | 4.1 | A | 9.9 | A | WUS |
| 265822_at | 682.7 | P | 758.9 | P | 702.7 | P | 786.5 | P | ATSLY1 |
| 265823_at | 250.3 | P | 457.8 | P | 244 | P | 460.5 | P | --- |
| 265824_at | 197.6 | P | 288 | P | 278.1 | P | 293.3 | P | ATCSLA07 |

| | | | | | | | | | |
|---|---|---|---|---|---|---|---|---|---|
| 265825_at | 1189.7 | P | 1079.7 | P | 869.1 | P | 1082.3 | P | UBQ7 |
| 265826_at | 1653.8 | P | 1260.6 | P | 1410.6 | P | 1217.4 | P | --- |
| 265827_at | 204.9 | P | 224.9 | P | 199.3 | P | 216.9 | P | VOZ2 |
| 265828_at | 170.7 | P | 86 | P | 110.6 | P | 121.1 | P | --- |
| 265829_at | 1.7 | A | 21 | A | 4.8 | A | 4.7 | A | --- |
| 265830_at | 2.9 | A | 0.6 | A | 11.5 | A | 1.7 | A | --- |
| 265831_at | 417.3 | P | 419.3 | P | 408.5 | P | 500.9 | P | --- |
| 265832_at | 26.5 | A | 17.7 | A | 50 | A | 15.4 | A | --- |
| 265833_at | 21.5 | A | 3.1 | A | 32.1 | A | 6.1 | A | --- |
| 265834_at | 2.7 | A | 4.3 | A | 1.9 | A | 1.2 | A | --- |
| 265835_at | 21.2 | A | 25.8 | A | 8.2 | A | 1 | A | --- |
| 265836_at | 0.6 | A | 1.8 | A | 3.3 | A | 3.2 | A | --- |
| 265837_at | 365.5 | P | 855.4 | P | 278.7 | P | 812.4 | P | LURP1 |
| 265838_at | 21.4 | A | 4.6 | A | 2.1 | A | 1.4 | A | --- |
| 265839_at | 19.9 | A | 27.6 | A | 26.7 | A | 6.9 | A | --- |
| 265840_at | 132.8 | P | 189.7 | P | 149.3 | P | 225.1 | P | --- |
| 265841_at | 47.6 | P | 69.1 | M | 4.2 | A | 63 | A | --- |
| 265842_at | 145.4 | P | 165.7 | P | 163.7 | P | 185.6 | P | ERF38 |
| 265843_at | 223.2 | P | 228.3 | P | 284.8 | P | 258.1 | P | ACX5 |
| 265844_at | 641.7 | P | 720.4 | P | 634.9 | P | 803.6 | P | FEI2 |
| 265845_at | 406.2 | P | 493.7 | P | 497 | P | 453.4 | P | --- |
| 265846_at | 7.3 | A | 29 | A | 30.4 | A | 22.5 | A | scpl28 |
| 265847_at | 213.5 | P | 236.4 | P | 208.7 | P | 275.1 | P | --- |
| 265848_at | 29.7 | A | 52.7 | A | 47.2 | A | 41.5 | A | INT3 |
| 265849_at | 233.6 | P | 172.1 | P | 277.4 | P | 150.4 | P | --- |
| 265850_at | 311 | P | 262.8 | P | 275.4 | P | 257.1 | P | --- |
| 265851_at | 35.9 | A | 25.2 | A | 61.6 | M | 35.4 | A | --- |
| 265852_at | 40.5 | P | 21.3 | A | 51.7 | A | 18.8 | A | --- |
| 265853_at | 308.1 | P | 198.8 | P | 235.4 | P | 125.5 | P | --- |
| 265854_at | 23.1 | P | 36.6 | P | 27.4 | P | 31.3 | P | --- |
| 265855_at | 393.5 | P | 618.5 | P | 343.4 | P | 589.9 | P | --- |
| 265856_at | 38.9 | A | 44.6 | A | 61.6 | A | 37.4 | A | LBD16 |
| 265857_s_at | 1948.7 | P | 1907.7 | P | 2116.5 | P | 1854.7 | P | PP2A-4 |
| 265858_at | 2166.9 | P | 2359.9 | P | 2144 | P | 2170.3 | P | --- |
| 265859_at | 15.2 | A | 16.3 | A | 19.3 | A | 11.6 | A | --- |
| 265860_at | 28.3 | A | 27 | A | 30 | A | 31.5 | A | --- |
| 265861_at | 14.9 | A | 12.4 | A | 43.7 | A | 46 | A | --- |
| 265862_at | 3 | A | 2.1 | A | 3.1 | A | 4.7 | A | --- |
| 265863_at | 1.9 | A | 4.6 | A | 37.1 | A | 28.4 | A | VIT1 |
| 265864_at | 72.9 | P | 68.4 | P | 68.1 | P | 80.4 | P | ATMAP70-3 |
| 265865_at | 68.1 | P | 100.4 | P | 87 | P | 81.5 | P | --- |
| 265866_at | 543.3 | P | 510 | P | 448.1 | P | 536.8 | P | --- |
| 265867_at | 616.8 | P | 786.2 | P | 610.1 | P | 723.6 | P | MEE11 |
| 265868_at | 456.8 | P | 464.9 | P | 393.8 | P | 400.4 | P | PUX2 |
| 265869_at | 211.5 | P | 226.7 | P | 215.4 | P | 213.7 | P | ARR14 |
| 265870_at | 20.4 | A | 56.2 | M | 26.6 | A | 46.5 | M | PDLP6 |
| 265871_at | 321 | P | 394.5 | P | 348 | P | 387 | P | --- |
| 265872_at | 1290.8 | P | 2210.8 | P | 1175.7 | P | 1897 | P | atnudt17 |
| 265873_at | 158.2 | P | 185.3 | P | 146.1 | P | 176 | P | --- |
| 265874_at | 195.6 | P | 242 | P | 160.4 | P | 212.3 | P | --- |
| 265875_at | 379.8 | P | 467.5 | P | 358 | P | 412.5 | P | --- |
| 265876_at | 53.1 | P | 78.3 | P | 100.4 | P | 81 | P | --- |
| 265877_at | 3.8 | A | 43.2 | A | 3.2 | A | 33.7 | A | BZIP34 |
| 265878_at | 27.1 | M | 23 | A | 20.6 | A | 27.5 | A | ZFP11 |
| 265879_at | 306 | P | 313.3 | P | 370.8 | P | 311.7 | P | --- |
| 265880_at | 266.6 | P | 400.2 | P | 364.9 | P | 354.8 | P | --- |
| 265881_at | 1.8 | A | 1.5 | A | 3.4 | A | 5.2 | A | --- |
| 265882_at | 1507.9 | P | 1616 | P | 1653.2 | P | 1572.9 | P | --- |
| 265883_at | 2196.6 | P | 2481.5 | P | 2274.8 | P | 2613.2 | P | --- |
| 265884_at | 569.3 | P | 869.9 | P | 684.2 | P | 720.7 | P | --- |
| 265885_at | 135.1 | P | 114.3 | P | 115.2 | P | 98.4 | P | --- |
| 265886_at | 335.9 | P | 232.6 | P | 257.4 | P | 262.9 | P | --- |
| 265887_at | 307.8 | P | 427.1 | P | 455.9 | P | 489.7 | P | HCS1 |
| 265888_at | 183.5 | P | 198.7 | P | 179.5 | M | 198.9 | P | RIE1 |
| 265889_at | 7.9 | A | 1.6 | A | 30.1 | A | 5.4 | A | --- |
| 265890_s_at | 18.4 | A | 1.5 | A | 4.1 | A | 1.1 | A | --- |
| 265891_at | 5.4 | A | 2.8 | A | 4.6 | A | 11.7 | A | --- |
| 265892_at | 11.1 | A | 47.5 | P | 49.3 | A | 57.4 | P | --- |
| 265893_at | 59.1 | P | 71.3 | P | 28.3 | A | 62.6 | P | --- |
| 265894_at | 1734.7 | P | 2582.8 | P | 2118 | P | 2397.7 | P | LTP |
| 265895_at | 797.2 | P | 655.2 | P | 989.4 | P | 814.6 | P | --- |
| 265896_at | 241.5 | P | 261.6 | P | 244.8 | P | 259.3 | P | emb2410 |
| 265897_at | 142.4 | P | 148.3 | P | 91.5 | P | 172.8 | P | MOT1 |
| 265898_at | 145.8 | P | 145.4 | P | 238 | P | 173.6 | P | --- |
| 265899_s_at | 173.9 | P | 114.5 | P | 152.3 | P | 108.6 | P | ASK4 |
| 265900_at | 981.9 | P | 759 | P | 963.7 | P | 855.2 | P | --- |
| 265901_at | 148.2 | P | 101.1 | P | 142 | P | 126 | P | --- |
| 265902_at | 209.5 | P | 193.9 | P | 186.1 | P | 182.4 | P | --- |
| 265903_at | 5.1 | A | 9.6 | A | 7.4 | A | 3.5 | A | SPIK |
| 265904_at | 37.4 | A | 10.8 | A | 2.2 | A | 1.4 | A | BGLU14 |
| 265905_at | 127.7 | P | 87.7 | P | 103.5 | P | 143.5 | P | --- |
| 265906_at | 11.2 | A | 12.8 | A | 20.4 | A | 39 | A | --- |
| 265907_at | 103.1 | P | 156.3 | P | 108.3 | P | 158.9 | P | --- |
| 265908_at | 121.4 | P | 130.3 | P | 165.5 | P | 198.9 | P | --- |
| 265909_at | 319.4 | P | 231.9 | P | 278.4 | P | 272.8 | P | --- |
| 265910_at | 1290.9 | P | 1493.1 | P | 1357.3 | P | 1490.9 | P | --- |
| 265911_at | 911.8 | P | 607.6 | P | 767.7 | P | 687.5 | P | --- |
| 265912_at | 588.5 | P | 480.4 | P | 658.5 | P | 555.6 | P | --- |
| 265913_at | 475.2 | P | 415.3 | P | 182.4 | P | 379 | P | --- |
| 265914_at | 13.5 | A | 19.2 | A | 2.1 | A | 0.9 | A | --- |
| 265915_at | 2.1 | A | 1.6 | A | 2.5 | A | 1.6 | A | --- |
| 265916_at | 74.2 | P | 69 | P | 70.5 | P | 90.2 | A | --- |
| 265917_at | 86.3 | P | 90.8 | P | 58.7 | A | 56 | P | AtRLP19 |
| 265918_at | 1373.9 | P | 1328.2 | P | 1537.5 | P | 1227.6 | P | KCS8 |
| 265919_at | 1.4 | A | 2.2 | A | 6 | A | 2.2 | A | --- |
| 265920_s_at | 67.7 | P | 52.9 | P | 39.8 | P | 13.3 | A | --- |
| 265921_at | 8.8 | A | 0.6 | A | 37.1 | A | 1.1 | A | --- |
| 265922_at | 239.9 | P | 71.5 | A | 257.3 | P | 75.9 | P | --- |
| 265923_at | 20.4 | A | 35.4 | A | 45.6 | P | 30.6 | A | --- |
| 265924_at | 6.4 | A | 1.9 | A | 15 | A | 14.5 | A | --- |
| 265925_at | 23.9 | A | 2.1 | A | 4.5 | A | 25.1 | A | --- |
| 265926_at | 141 | P | 128.2 | P | 154.5 | P | 111 | P | --- |
| 265927_at | 51.9 | P | 53.4 | A | 89.9 | P | 88.9 | P | --- |
| 265928_at | 19.9 | A | 30.6 | A | 25.4 | A | 4 | A | --- |
| 265929_s_at | 64.9 | P | 102.7 | P | 137.2 | P | 136 | P | --- |
| 265930_at | 367.6 | P | 455.1 | P | 525.2 | P | 492.2 | P | emb2444 |
| 265931_at | 140.2 | P | 74 | P | 150.2 | P | 114.7 | P | --- |
| 265932_at | 0.8 | A | 0.6 | A | 1.7 | A | 2.3 | A | LBD8 |
| 265933_at | 27.5 | A | 4.4 | A | 3.4 | A | 13.5 | A | --- |

| | | | | | | | | | |
|---|---|---|---|---|---|---|---|---|---|
| 265934_at | 220.4 | P | 279.4 | P | 139.5 | P | 256.2 | P | EER5 |
| 265935_at | 113.4 | P | 224.7 | P | 128.1 | P | 198 | P | TET2 |
| 265936_at | 222.6 | P | 270.6 | P | 261.2 | P | 288.4 | P | ATKEA4 |
| 265937_at | 20.9 | A | 22.7 | A | 4.7 | A | 7.1 | A | --- |
| 265938_at | 224.2 | P | 307.3 | P | 240.7 | P | 390.4 | P | --- |
| 265939_at | 144.9 | P | 278.7 | P | 218.7 | P | 218.2 | P | --- |
| 265940_at | 3350 | P | 2361.3 | P | 3120.8 | P | 2716.8 | P | NAP1;2 |
| 265941_s_at | 115.8 | P | 103.7 | P | 142.7 | P | 127.5 | P | --- |
| 265942_at | 1020.4 | P | 975 | P | 1164.2 | P | 839.4 | P | ckl5 |
| 265943_at | 587.9 | P | 575.1 | P | 670.1 | P | 566 | P | CDA1 |
| 265944_at | 176.9 | P | 105 | P | 227.8 | P | 152.7 | P | ASHR2 |
| 265945_at | 52.5 | A | 53.7 | A | 47.2 | A | 38.1 | A | CKX2 |
| 265946_s_at | 738.2 | P | 894.5 | P | 849.2 | P | 936.3 | P | FVE |
| 265947_at | 2031.2 | P | 1294.6 | P | 2536.5 | P | 1310.6 | P | --- |
| 265948_at | 25.4 | A | 107.7 | P | 38.7 | A | 99.4 | P | ACO1 |
| 265949_at | 9.9 | A | 23.9 | A | 21.3 | A | 16.9 | A | --- |
| 265950_s_at | 17.3 | A | 18.9 | A | 50.1 | A | 12.8 | A | --- |
| 265951_at | 35 | A | 74.3 | A | 96.2 | A | 45.9 | A | OFP7 |
| 265952_at | 437.2 | P | 980.2 | P | 472.1 | P | 809.5 | P | --- |
| 265953_at | 685 | P | 1414.1 | P | 750 | P | 991.4 | P | --- |
| 265954_at | 109.9 | P | 50.1 | P | 112.5 | P | 64.9 | P | TTG2 |
| 265955_at | 39.4 | A | 61.5 | A | 90.5 | P | 80 | P | PDR5 |
| 265956_at | 2.8 | A | 11.2 | A | 24 | A | 1.8 | A | --- |
| 265957_at | 310.6 | P | 346.8 | P | 366 | P | 407.2 | P | --- |
| 265958_at | 1363.3 | P | 1034.6 | P | 1323.1 | P | 1214.5 | P | ADK |
| 265959_at | 1475.6 | P | 1253.2 | P | 1209.3 | P | 1726.5 | P | --- |
| 265960_at | 446.8 | P | 503 | P | 410.5 | P | 469 | P | --- |
| 265961_at | 531.1 | P | 491.9 | P | 644.6 | P | 519.6 | P | --- |
| 265962_at | 241.9 | P | 260.2 | P | 347 | P | 165.3 | P | --- |
| 265963_s_at | 11852.5 | P | 11009.7 | P | 14146.5 | P | 11381.3 | P | ATRPS5B |
| 265964_at | 159.9 | P | 108.4 | P | 136.2 | P | 119.9 | P | --- |
| 265965_at | 764 | P | 740.4 | P | 628.8 | P | 606.3 | P | --- |
| 265966_at | 7815.3 | P | 8121 | P | 8788.9 | P | 8493.3 | P | --- |
| 265967_at | 532 | P | 482.4 | P | 574.7 | P | 453.9 | P | --- |
| 265968_at | 2436.5 | P | 1891.3 | P | 3013.9 | P | 2089.4 | P | ATTIM17-2 |
| 265969_at | 0.8 | A | 2.6 | A | 1.9 | A | 1.5 | A | --- |
| 265970_at | 3.2 | A | 26.5 | A | 15.2 | A | 1.3 | A | --- |
| 265971_at | 1.7 | A | 1 | A | 1.3 | A | 2 | A | --- |
| 265972_at | 14.5 | A | 4 | A | 20.6 | A | 2.5 | A | --- |
| 265973_at | 21.5 | A | 37.5 | A | 15.9 | A | 13.9 | A | --- |
| 265974_at | 252.9 | P | 340.2 | P | 197.3 | P | 293.4 | P | --- |
| 265975_at | 33.6 | P | 22.7 | A | 24.2 | A | 6.9 | A | --- |
| 265976_at | 2.3 | A | 1.5 | A | 2.1 | A | 1.1 | A | --- |
| 265977_at | 2.3 | A | 7.7 | A | 5.5 | A | 3.2 | A | --- |
| 265978_at | 56.9 | P | 66.4 | P | 53.1 | P | 75.8 | P | --- |
| 265979_at | 2.4 | A | 5 | A | 24.5 | A | 5.2 | A | --- |
| 265980_at | 1.1 | A | 0.9 | A | 2 | A | 3 | A | --- |
| 265981_s_at | 27.4 | A | 6.5 | A | 4.3 | A | 2.9 | A | --- |
| 265982_at | 1.9 | A | 1.2 | A | 1.6 | A | 2.2 | A | --- |
| 265983_at | 33.9 | P | 32.5 | P | 49.4 | P | 31.4 | P | HB-2 |
| 265984_at | 5.1 | A | 23 | A | 28.9 | A | 5.8 | A | TPS10 |
| 265985_at | 111.1 | P | 123.6 | P | 168.5 | P | 144.1 | P | ATPUP5 |
| 265986_at | 105 | P | 186.5 | P | 150.3 | P | 168.9 | P | --- |
| 265987_at | 72.3 | P | 107.2 | P | 73.5 | A | 54.8 | P | --- |
| 265988_at | 29.6 | A | 44.4 | P | 45.6 | A | 39.4 | M | --- |
| 265989_at | 105.2 | P | 83.5 | P | 112.7 | P | 123.2 | P | --- |
| 265990_at | 1085.5 | P | 1018.4 | P | 1380.4 | P | 1164.1 | P | --- |
| 265991_at | 426.8 | P | 486.6 | P | 590.2 | P | 433.7 | P | SCA3 |
| 265992_at | 41.6 | P | 12.1 | A | 46.1 | M | 23.5 | M | --- |
| 265993_at | 41.7 | A | 87.3 | P | 71.7 | A | 78.2 | P | --- |
| 265994_at | 125.2 | P | 363.2 | P | 150 | P | 323.2 | P | --- |
| 265995_at | 0.5 | A | 14.3 | A | 7.9 | A | 13 | A | --- |
| 265996_at | 9779.1 | P | 6223.2 | P | 8953.7 | P | 7218.9 | P | --- |
| 265997_at | 46.2 | M | 23 | A | 45.1 | A | 66.8 | P | --- |
| 265998_at | 3866.3 | P | 4563.7 | P | 4421.8 | P | 4418.6 | P | ALDH11A3 |
| 265999_at | 582.3 | P | 331 | P | 447.6 | P | 352.8 | P | --- |
| 266000_at | 2081.9 | P | 1272 | P | 1507.4 | P | 1234.9 | P | CYP71B6 |
| 266001_at | 681.4 | P | 790.9 | P | 814.8 | P | 804 | P | HHP3 |
| 266002_at | 112.3 | P | 67.7 | P | 103 | P | 86.4 | P | --- |
| 266003_at | 40.6 | P | 31.1 | P | 44 | P | 51.4 | P | --- |
| 266004_at | 794.5 | P | 386.8 | P | 831.3 | P | 456.4 | P | ALS3 |
| 266005_at | 1903.3 | P | 1758.7 | P | 1629.2 | P | 1726.4 | P | RSZ33 |
| 266006_at | 40 | A | 30.5 | A | 54 | A | 47 | A | --- |
| 266007_at | 49.3 | P | 82.4 | P | 48.2 | A | 93.4 | P | --- |
| 266008_at | 89.5 | P | 124.1 | P | 74.2 | P | 123.1 | P | --- |
| 266009_at | 63.7 | P | 101.8 | P | 84.9 | P | 96.7 | P | --- |
| 266010_at | 2.6 | A | 4.6 | A | 16.7 | A | 28.2 | A | --- |
| 266011_at | 0.7 | A | 2.2 | A | 4.1 | A | 1.5 | A | --- |
| 266012_s_at | 192 | P | 146.7 | P | 234.8 | P | 222.2 | P | --- |
| 266013_s_at | 17.5 | A | 37.6 | A | 36.4 | A | 31.8 | A | --- |
| 266014_s_at | 40.1 | P | 44.7 | P | 62 | P | 80.1 | P | --- |
| 266015_at | 264.1 | P | 234.1 | P | 301.8 | P | 341.4 | P | --- |
| 266016_at | 156.1 | P | 254.6 | P | 192.4 | P | 243 | P | --- |
| 266017_at | 932.4 | P | 1325.6 | P | 475.8 | P | 607.6 | P | --- |
| 266018_at | 1747.9 | P | 1949.4 | P | 1817.8 | P | 1670.4 | P | SCY1 |
| 266019_at | 246.2 | P | 345 | P | 182.7 | P | 291.7 | P | --- |
| 266020_at | 5.6 | A | 6.7 | A | 9 | A | 3.1 | A | SDG11 |
| 266021_at | 36.2 | A | 58.8 | P | 77.7 | P | 54.6 | P | --- |
| 266022_at | 2003.4 | P | 2427.1 | P | 2592.3 | P | 2777.6 | P | --- |
| 266023_at | 11.6 | A | 19.9 | P | 19.9 | A | 12.4 | A | --- |
| 266024_at | 18.5 | A | 1.6 | A | 2 | A | 2.6 | A | --- |
| 266025_at | 0.3 | A | 0.8 | A | 1.6 | A | 0.7 | A | --- |
| 266026_at | 5.4 | A | 8.1 | A | 6.6 | A | 1.8 | A | --- |
| 266027_at | 2.5 | A | 0.8 | A | 1.5 | A | 0.9 | A | --- |
| 266028_at | 217 | P | 227.2 | P | 215.5 | P | 165.7 | P | --- |
| 266029_at | 2 | A | 3 | A | 4.4 | A | 3.6 | A | scpl38 |
| 266030_x_at | 12.7 | A | 6.4 | A | 9.9 | A | 17.8 | A | --- |
| 266031_at | 6.3 | A | 1.1 | A | 8 | A | 3.3 | A | --- |
| 266032_x_at | 4.2 | A | 10.8 | A | 26.1 | A | 39 | A | --- |
| 266033_at | 1144.4 | P | 1232.1 | P | 1237.2 | P | 1183 | P | --- |
| 266034_at | 41 | A | 53.6 | A | 19.6 | A | 74.4 | A | FIP1 |
| 266035_at | 1886.6 | P | 1824 | P | 2412.2 | P | 2004.9 | P | MOD1 |
| 266036_s_at | 2468.1 | P | 2377.8 | P | 2280.6 | P | 2827.5 | P | PAA2 |
| 266037_at | 571.6 | P | 515.8 | P | 606.7 | P | 630.5 | P | --- |
| 266038_at | 118.7 | P | 74.7 | P | 57.8 | P | 123.2 | P | ATMRP11 |
| 266039_s_at | 49.7 | P | 28.7 | A | 22 | A | 29.1 | A | --- |
| 266040_at | 2.8 | A | 3.9 | A | 11 | A | 2.4 | A | --- |
| 266041_s_at | 1.5 | A | 3 | A | 8.3 | A | 4.3 | A | ArthMp028 |
| 266042_s_at | 158.4 | P | 148.2 | P | 122.1 | P | 90.7 | P | --- |

| | | | | | | | | | |
|---|---|---|---|---|---|---|---|---|---|
| 266043_at | 77.8 | A | 62.7 | A | 78.9 | A | 47.4 | A | --- |
| 266044_s_at | 111 | P | 63.9 | A | 89.9 | P | 83.9 | P | --- |
| 266045_s_at | 89.9 | P | 46.3 | A | 56.6 | M | 27.6 | A | --- |
| 266046_at | 7.5 | A | 23.2 | A | 10.8 | A | 9.8 | A | --- |
| 266047_at | 246.7 | P | 198.3 | P | 224.8 | P | 216 | P | --- |
| 266048_at | 2 | A | 2.8 | A | 3.5 | A | 20.5 | A | ATCXXS2 |
| 266049_at | 140.4 | P | 103.9 | P | 124.1 | P | 106.8 | P | --- |
| 266050_at | 151.4 | P | 158 | P | 131.8 | P | 122.1 | P | --- |
| 266051_at | 338.7 | P | 326.4 | P | 302.4 | P | 398.6 | P | --- |
| 266052_at | 35.5 | A | 38.3 | A | 68.2 | M | 12.3 | A | WRKY55 |
| 266053_at | 247.5 | P | 353.6 | P | 258.4 | P | 344.6 | P | --- |
| 266054_at | 94.2 | P | 156.9 | P | 155.1 | P | 128.6 | P | --- |
| 266055_at | 359 | P | 250.8 | P | 258.6 | P | 241.7 | P | --- |
| 266056_at | 141.9 | P | 171.1 | P | 147.8 | P | 151 | P | ATATG18C |
| 266057_at | 1361.1 | P | 1414.6 | P | 1337.8 | P | 1330.7 | P | --- |
| 266058_at | 124.6 | P | 171.6 | P | 158.9 | P | 196.5 | P | GLY1 |
| 266059_at | 2113.2 | P | 1935 | P | 2229.3 | P | 2142.3 | P | --- |
| 266060_at | 219.2 | P | 257.6 | P | 293.9 | P | 271.3 | P | --- |
| 266061_at | 32 | A | 35.1 | A | 9.4 | A | 20.8 | A | --- |
| 266062_at | 624.8 | P | 599.3 | P | 528.8 | P | 448.5 | P | --- |
| 266063_at | 66.8 | P | 56.7 | P | 48.9 | A | 64.1 | P | CHR8 |
| 266064_at | 87.8 | P | 100.8 | P | 66.3 | M | 71.4 | P | --- |
| 266065_at | 996.6 | P | 943.1 | P | 975.7 | P | 1016.4 | P | PHYB |
| 266066_at | 3.2 | A | 10 | A | 14.3 | A | 1.1 | A | XTH21 |
| 266067_s_at | 1.2 | A | 17.1 | P | 17.6 | A | 16.9 | A | --- |
| 266068_at | 1.1 | A | 10.5 | A | 8.1 | A | 6.9 | A | GGPS4 |
| 266069_at | 6.7 | A | 35.5 | A | 9.7 | A | 28.5 | A | MEE16 |
| 266070_at | 684.3 | P | 2005.6 | P | 304.7 | P | 1521.6 | P | EXLB3 |
| 266071_at | 62.5 | P | 77.5 | P | 54.9 | A | 44.4 | P | --- |
| 266072_at | 402.8 | P | 302.4 | P | 167.8 | P | 209.3 | P | ATTPS11 |
| 266073_at | 379.3 | P | 385.2 | P | 290.6 | P | 361.3 | P | --- |
| 266074_at | 376.8 | P | 354 | P | 427.8 | P | 351.8 | P | --- |
| 266075_s_at | 2733.4 | P | 2210.4 | P | 2382.2 | P | 2494.7 | P | --- |
| 266076_at | 287.1 | P | 205.3 | P | 329.3 | P | 234.3 | P | --- |
| 266077_at | 27.1 | P | 20.7 | A | 28.7 | A | 7.4 | A | --- |
| 266078_at | 208.1 | P | 587.3 | P | 237.1 | P | 462.8 | P | ARR16 |
| 266079_at | 512.1 | P | 763.3 | P | 715.2 | P | 714.2 | P | LCD1 |
| 266080_at | 190.4 | P | 174.8 | P | 133.6 | P | 175.6 | P | --- |
| 266081_at | 81.4 | P | 135.6 | P | 139.9 | P | 136.1 | P | --- |
| 266082_at | 33.1 | A | 6.5 | A | 35 | A | 24.9 | A | --- |
| 266083_at | 1.7 | A | 4.4 | A | 4 | A | 2.8 | A | --- |
| 266084_at | 27.3 | A | 2.6 | A | 8.8 | A | 2.9 | A | --- |
| 266085_at | 2.3 | A | 11.2 | A | 19.4 | A | 25.4 | A | --- |
| 266086_at | 47.1 | A | 49.7 | A | 38.6 | A | 20.2 | A | PHT4;2 |
| 266087_at | 468.7 | P | 469.2 | P | 638.9 | P | 611.6 | P | --- |
| 266088_at | 39.3 | P | 61.4 | P | 76.8 | P | 49.1 | P | --- |
| 266089_at | 557.5 | P | 679 | P | 590 | P | 640.3 | P | --- |
| 266090_at | 377.9 | P | 351 | P | 277.7 | P | 369.2 | P | --- |
| 266091_at | 624 | P | 594.1 | P | 630.3 | P | 496.3 | P | emb1513 |
| 266092_at | 11.6 | A | 31.3 | A | 13.6 | A | 11.8 | A | --- |
| 266093_at | 687.5 | P | 775.8 | P | 980 | P | 793.8 | P | --- |
| 266094_at | 478.8 | P | 589.8 | P | 455.5 | P | 555.2 | P | --- |
| 266095_at | 609.8 | P | 434.8 | P | 541.5 | P | 371.4 | P | DET2 |
| 266096_at | 562.3 | P | 656 | P | 454.7 | P | 535.5 | P | VCL1 |
| 266097_at | 2584.3 | P | 2519.9 | P | 2177.2 | P | 1993.9 | P | SOUL-1 |
| 266098_at | 405.9 | P | 165.6 | P | 156.2 | P | 464.8 | P | --- |
| 266099_at | 1108.4 | P | 1276.7 | P | 1484.4 | P | 1268.1 | P | CAC3 |
| 266100_at | 28.5 | P | 56.4 | P | 39.3 | P | 39.2 | A | --- |
| 266101_at | 1011.1 | P | 1061.2 | P | 1010.8 | P | 1020.1 | P | --- |
| 266102_at | 79 | P | 107.3 | P | 106.1 | P | 125.9 | P | --- |
| 266103_at | 988.2 | P | 1068 | P | 788.8 | P | 1087.8 | P | PVA12 |
| 266104_at | 209.5 | P | 226.7 | P | 286.1 | P | 235.6 | P | --- |
| 266105_at | 2117.5 | P | 2536.3 | P | 2407.3 | P | 2512.4 | P | SEC61 BETA |
| 266106_at | 428.5 | P | 440.9 | P | 283.7 | P | 299.2 | P | ATATG8E |
| 266107_at | 15.9 | A | 26.6 | A | 46.5 | A | 3.3 | A | --- |
| 266108_at | 46.3 | A | 82.7 | P | 104.6 | P | 67.5 | P | --- |
| 266109_at | 171.7 | P | 153.7 | P | 156.3 | P | 175.7 | P | --- |
| 266110_at | 433.8 | P | 501.9 | P | 399.2 | P | 510.7 | P | AtIDD4 |
| 266111_at | 13.4 | A | 35.9 | A | 2.7 | A | 32.9 | A | --- |
| 266112_x_at | 12.7 | A | 3.2 | A | 30.7 | A | 28.7 | A | --- |
| 266113_x_at | 1.2 | A | 12.1 | A | 1.9 | A | 2.6 | A | --- |
| 266114_at | 139.2 | P | 70.4 | P | 88.4 | P | 56.4 | P | --- |
| 266115_at | 19.7 | A | 3.5 | A | 45.8 | A | 17.6 | A | LCR72 |
| 266116_at | 503.1 | P | 580.4 | P | 494.1 | P | 504.4 | P | TOM3 |
| 266117_at | 198.2 | P | 204.3 | P | 224.7 | P | 266.4 | P | --- |
| 266118_at | 1937.8 | P | 3208.7 | P | 2079.9 | P | 3460.2 | P | LCR68 |
| 266119_at | 7823.3 | P | 11019.2 | P | 9243 | P | 12265.1 | P | LCR69 |
| 266120_at | 713.6 | P | 907.4 | P | 874.8 | P | 992.1 | P | AtIDD5 |
| 266121_at | 673.7 | P | 487.7 | P | 704 | P | 527.5 | P | --- |
| 266122_at | 661.9 | P | 698.4 | P | 566.7 | P | 718.1 | P | ETL1 |
| 266123_at | 8245.9 | P | 12010.3 | P | 12187.2 | P | 10850.1 | P | --- |
| 266124_at | 83.9 | P | 323.4 | P | 139.7 | P | 122.8 | P | cycp3;1 |
| 266125_at | 80.6 | P | 142.4 | P | 93.9 | A | 130.6 | P | --- |
| 266126_at | 6.8 | A | 33.3 | A | 7.3 | A | 13.9 | A | --- |
| 266127_s_at | 1898.1 | P | 1686.8 | P | 2678.2 | P | 1698.1 | P | --- |
| 266128_at | 2.7 | A | 0.7 | A | 2.2 | A | 5.7 | A | EMB2766 |
| 266129_at | 57.4 | P | 78.3 | P | 65 | P | 69.5 | P | CCD7 |
| 266130_at | 157.3 | P | 152.4 | P | 133.7 | P | 122.2 | P | --- |
| 266131_at | 411.5 | P | 518.1 | P | 447.6 | P | 513.6 | P | --- |
| 266132_at | 22.4 | A | 8.2 | A | 31.9 | A | 6.2 | A | SPX3 |
| 266133_at | 67.8 | M | 50.9 | A | 89.5 | A | 71.5 | A | --- |
| 266134_s_at | 2.1 | A | 0.7 | A | 2.9 | A | 3 | A | --- |
| 266135_at | 176.8 | P | 220.2 | P | 169.6 | P | 140.5 | P | --- |
| 266136_at | 645.8 | P | 616.2 | P | 625.9 | P | 636.2 | P | --- |
| 266137_at | 428.5 | P | 607.8 | P | 446.4 | P | 478.5 | P | --- |
| 266138_at | 444.6 | P | 508.2 | P | 463.6 | P | 504.4 | P | --- |
| 266139_at | 145.5 | P | 65.9 | A | 172.4 | A | 97.1 | P | --- |
| 266140_at | 271.9 | P | 164.5 | P | 219.1 | P | 180.1 | P | --- |
| 266141_at | 52.2 | A | 28.6 | A | 14.3 | A | 42.6 | A | PDF2.1 |
| 266142_at | 10597.2 | P | 157.6 | P | 4623.9 | P | 626.5 | P | --- |
| 266143_at | 28.5 | P | 24.8 | A | 42.5 | A | 17.8 | A | --- |
| 266144_at | 1.8 | A | 7.7 | A | 17.5 | A | 11 | A | --- |
| 266145_at | 0.2 | A | 0.2 | A | 0.3 | A | 2.8 | A | --- |
| 266146_at | 17.1 | A | 21.4 | A | 17.1 | A | 28.3 | A | --- |
| 266147_at | 0.8 | A | 15.6 | A | 1.4 | A | 7.1 | A | --- |
| 266148_x_at | 20.2 | A | 30.4 | A | 76.2 | P | 34 | A | --- |
| 266149_x_at | 37.6 | A | 42.9 | A | 46.7 | A | 6 | A | --- |
| 266150_s_at | 466.1 | P | 353.7 | P | 320.4 | P | 304.9 | P | --- |
| 266151_x_at | 6.7 | A | 8.6 | A | 10.9 | A | 28 | A | --- |

| | | | | | | | | | |
|---|---|---|---|---|---|---|---|---|---|
| 266152_s_at | 29.7 | A | 30.1 | A | 46 | A | 10.5 | A | --- |
| 266153_at | 1.7 | A | 1.9 | A | 4.2 | A | 2.3 | A | --- |
| 266154_at | 88.4 | P | 41.8 | P | 125.4 | P | 82.7 | P | --- |
| 266155_at | 1371.8 | P | 1239.5 | P | 1444.5 | P | 1243.9 | P | CYP89A5 |
| 266156_at | 35.9 | A | 57.7 | P | 42 | A | 33.1 | P | FRA8 |
| 266157_at | 125 | P | 116.7 | P | 132.8 | P | 140.2 | P | ATFUC1 |
| 266158_at | 283.5 | P | 283.3 | P | 375.7 | P | 334.8 | P | --- |
| 266159_at | 1.8 | A | 11.9 | A | 5 | A | 10.9 | A | --- |
| 266160_at | 16.4 | A | 18.8 | A | 2.5 | A | 14.1 | A | ATCHX8 |
| 266161_at | 11.7 | A | 0.7 | A | 20.3 | A | 10 | A | ATCHX7 |
| 266162_at | 2.4 | A | 27.8 | A | 27.2 | A | 13.8 | A | FRU |
| 266163_at | 174.5 | P | 124.6 | P | 187 | P | 122.6 | P | --- |
| 266164_at | 47.7 | P | 65.9 | P | 99 | P | 83.7 | P | --- |
| 266165_at | 4056.6 | P | 1315.9 | P | 1471.3 | P | 2773.1 | P | CSD2 |
| 266166_at | 564.2 | P | 629.7 | P | 582.2 | P | 628.5 | P | --- |
| 266167_at | 3193.4 | P | 1946 | P | 2312.8 | P | 1531.1 | P | YLS5 |
| 266168_at | 2126.1 | P | 2443.6 | P | 950.9 | P | 1950.1 | P | --- |
| 266169_at | 2.2 | A | 15.1 | A | 3.3 | A | 3.8 | A | --- |
| 266170_at | 1346 | P | 561.2 | P | 874.2 | P | 677.7 | P | --- |
| 266171_at | 272.7 | P | 237.9 | P | 194 | P | 266.1 | P | NF-YB1 |
| 266172_at | 6789.4 | P | 5306.3 | P | 6034.3 | P | 5848.7 | P | PIP2E |
| 266173_at | 64.7 | P | 114.1 | P | 64.8 | P | 112.2 | P | --- |
| 266174_at | 12.9 | A | 18.6 | A | 60.8 | A | 28.1 | A | --- |
| 266175_at | 216.6 | P | 225.1 | P | 322.2 | P | 202 | P | ANAC035 |
| 266176_at | 9.8 | A | 3.2 | A | 6.1 | A | 19.1 | A | AtPP2-B2 |
| 266177_at | 25.3 | A | 23.3 | A | 30.5 | A | 15.6 | A | ATPP2-B3 |
| 266178_at | 0.9 | A | 10.4 | A | 2.6 | A | 1.2 | A | AtPP2-B4 |
| 266179_at | 0.8 | A | 2.8 | A | 4.1 | A | 1.8 | A | AtPP2-B5 |
| 266180_at | 439.8 | P | 415.6 | P | 464.4 | P | 470 | P | AL6 |
| 266181_at | 3467.9 | P | 1972.4 | P | 2368.1 | P | 1925.7 | P | ATGSTZ1 |
| 266182_at | 311.9 | P | 330.9 | P | 267.7 | P | 328.9 | P | --- |
| 266183_at | 46.2 | A | 29.5 | A | 42.2 | A | 37.6 | A | ATPT2 |
| 266184_s_at | 362.4 | P | 353 | P | 379.2 | P | 371 | P | ATPT2 |
| 266185_at | 413 | P | 453 | P | 477.9 | P | 510 | P | --- |
| 266186_at | 156.2 | P | 116.8 | P | 113.6 | P | 96.6 | P | AERO2 |
| 266187_at | 176 | P | 159 | P | 158.7 | P | 208.1 | P | --- |
| 266188_at | 653.8 | P | 722.7 | P | 511.3 | P | 601.9 | P | --- |
| 266189_at | 1206.6 | P | 640.5 | P | 921.9 | P | 628.6 | P | --- |
| 266190_at | 166.7 | P | 141.2 | P | 83.8 | P | 121 | P | --- |
| 266191_at | 15.5 | A | 6.7 | A | 19.8 | A | 6.4 | A | --- |
| 266192_at | 1981 | P | 1851.8 | P | 1712.1 | P | 1813 | P | --- |
| 266193_at | 1625.9 | P | 1493.9 | P | 1377.6 | P | 1449.8 | P | --- |
| 266194_at | 112.5 | P | 193.2 | P | 139.6 | P | 124.7 | P | --- |
| 266195_at | 157 | P | 117.3 | P | 170.5 | P | 131.2 | P | --- |
| 266196_at | 56.9 | P | 33.9 | A | 26.3 | A | 20.1 | A | --- |
| 266197_at | 86.3 | P | 62 | P | 72.3 | P | 82.9 | P | --- |
| 266198_at | 24.5 | A | 15.1 | A | 5 | A | 11.6 | A | --- |
| 266199_at | 16.7 | A | 20.2 | A | 70.7 | P | 22.4 | A | CPK20 |
| 266200_at | 17.9 | A | 4.5 | A | 3.2 | A | 5.8 | A | --- |
| 266201_at | 16.1 | A | 34.3 | A | 40.2 | A | 39.7 | P | --- |
| 266202_at | 509.4 | P | 534.2 | P | 480.5 | P | 496.3 | P | --- |
| 266203_at | 175.9 | P | 199.3 | P | 156.5 | P | 161.3 | P | AtPP2-B1 |
| 266204_at | 126.2 | P | 199.5 | P | 141.4 | P | 137 | P | --- |
| 266205_s_at | 5701.5 | P | 5518.4 | P | 5557.8 | P | 5809.8 | P | AXS2 |
| 266206_at | 3105.4 | P | 3284.4 | P | 3351.5 | P | 3694.2 | P | --- |
| 266207_at | 2625.2 | P | 3291 | P | 2968 | P | 3183.8 | P | --- |
| 266208_at | 128 | P | 161.9 | P | 135.9 | P | 161.8 | P | ATIPT2 |
| 266209_at | 21.6 | A | 1.9 | A | 5.6 | A | 2.9 | A | ATC |
| 266210_at | 5266.5 | P | 4598.1 | P | 5873.2 | P | 4720.8 | P | PGY1 |
| 266211_at | 15.4 | A | 3.7 | A | 47.1 | A | 27.9 | A | --- |
| 266212_at | 4 | A | 20.2 | A | 13.6 | A | 5.4 | A | --- |
| 266213_s_at | 3.9 | A | 6 | A | 31.4 | A | 10.2 | A | --- |
| 266214_at | 3.1 | A | 8.4 | A | 10.1 | A | 6.9 | A | --- |
| 266215_at | 2470.8 | P | 5381.4 | P | 3986.9 | P | 4765.5 | P | EXGT-A1 |
| 266216_at | 275.9 | P | 381.5 | P | 359.3 | P | 399.9 | P | --- |
| 266217_at | 22.2 | A | 7.7 | A | 18.9 | A | 22.9 | A | PUB12 |
| 266218_s_at | 22.6 | A | 25.6 | A | 10.1 | A | 17.7 | A | CYP710A4 |
| 266219_at | 89.6 | P | 125.3 | P | 150.4 | P | 152.7 | P | emb1997 |
| 266220_at | 1.1 | A | 8.8 | A | 1 | A | 13.9 | A | --- |
| 266221_at | 157.3 | P | 214.6 | P | 334.8 | P | 291.8 | P | UXS6 |
| 266222_at | 0.7 | A | 2.7 | A | 1.2 | A | 5.8 | A | --- |
| 266223_at | 1387.9 | P | 1294.1 | P | 1453.6 | P | 1284.1 | P | --- |
| 266224_at | 2951 | P | 3925.4 | P | 3222.8 | P | 3792.5 | P | ALB3 |
| 266225_at | 6233.3 | P | 7414.4 | P | 7279.2 | P | 9174.3 | P | ATOEP16-1 |
| 266226_at | 413.6 | P | 608.1 | P | 392.8 | P | 588.9 | P | HIS4 |
| 266227_at | 45.1 | P | 57 | P | 65.8 | P | 54.4 | P | --- |
| 266228_at | 4013.5 | P | 3220 | P | 3568.7 | P | 3145.8 | P | CXIP4 |
| 266229_at | 3074.8 | P | 3148.9 | P | 3534.9 | P | 3220.6 | P | --- |
| 266230_at | 108.4 | P | 108.4 | P | 87.8 | P | 96.9 | P | --- |
| 266231_at | 197.1 | P | 182.8 | P | 156.3 | P | 177.8 | P | PSKR1 |
| 266232_at | 37.3 | P | 28.7 | P | 32.3 | A | 0.6 | A | AtPP2-B6 |
| 266233_at | 19.5 | A | 12.9 | A | 21.5 | A | 11.9 | A | AtPP2-B8 |
| 266234_at | 122.3 | P | 132.3 | P | 120.4 | P | 79.5 | P | SKIP3 |
| 266235_at | 216.3 | P | 340.4 | P | 204.2 | P | 263.6 | P | AtPP2-B10 |
| 266236_at | 20 | A | 5.2 | A | 4.5 | A | 30.5 | A | ATGSTZ2 |
| 266237_at | 526.7 | P | 430.5 | P | 598.2 | P | 477.1 | P | ATRPC14 |
| 266238_at | 1752 | P | 1677.8 | P | 1691.3 | P | 1499.9 | P | TOPP1 |
| 266239_at | 1511.4 | P | 1219.4 | P | 1294.6 | P | 1277.7 | P | --- |
| 266240_at | 171.4 | P | 120.9 | P | 197 | P | 226.5 | P | --- |
| 266241_s_at | 89.6 | P | 116.2 | P | 140.5 | P | 106.3 | P | --- |
| 266242_at | 100.4 | P | 76.1 | P | 97.4 | A | 109 | P | --- |
| 266243_at | 2 | A | 1.8 | A | 3.8 | A | 1.5 | A | --- |
| 266244_at | 27.4 | A | 87.5 | P | 62.3 | A | 66.8 | P | --- |
| 266245_at | 5.1 | A | 5.5 | A | 3.4 | A | 1.1 | A | --- |
| 266246_at | 123.9 | P | 62.6 | P | 87.8 | P | 50.4 | M | CYP94C1 |
| 266247_at | 76.2 | P | 135.5 | P | 58.5 | A | 55.5 | P | --- |
| 266248_at | 17.4 | P | 26.4 | A | 47.1 | P | 20.4 | A | --- |
| 266249_at | 1.8 | A | 1.7 | A | 1.1 | A | 0.5 | A | --- |
| 266250_at | 2.3 | A | 3.3 | A | 5 | A | 2.3 | A | --- |
| 266251_s_at | 17.1 | P | 21.6 | M | 11.1 | A | 21.3 | A | --- |
| 266252_at | 29.7 | A | 29.1 | A | 33.5 | A | 14 | A | --- |
| 266253_at | 940.6 | P | 873.9 | P | 957.6 | P | 1013.1 | P | HDT4 |
| 266254_at | 595.5 | P | 621.9 | P | 650.9 | P | 663.4 | P | NAT12 |
| 266255_at | 36.7 | A | 41.9 | A | 21 | A | 57.3 | A | --- |
| 266256_at | 7500.4 | P | 7307.1 | P | 7896.4 | P | 7930.2 | P | --- |
| 266257_at | 994.1 | P | 544.5 | P | 922.8 | P | 640.4 | P | PD1 |
| 266258_at | 6511.9 | P | 6011.7 | P | 6324.9 | P | 6767.9 | P | --- |
| 266259_at | 715 | P | 281.4 | P | 483.2 | P | 284.8 | P | --- |
| 266260_at | 1790.3 | P | 2361.8 | P | 2441.7 | P | 1839.5 | P | SKD1 |

| | | | | | | | | | |
|---|---|---|---|---|---|---|---|---|---|
| 266261_at | 471.4 | P | 298.2 | P | 390.6 | P | 368.1 | P | --- |
| 266262_at | 197.1 | P | 180.3 | P | 244.6 | P | 159.7 | P | --- |
| 266263_at | 1.6 | A | 8.5 | A | 33 | A | 0.8 | A | --- |
| 266264_at | 388.8 | P | 295.4 | P | 310.3 | P | 324.4 | P | --- |
| 266265_at | 1962.9 | P | 1192.4 | P | 1783.3 | P | 1115.1 | P | --- |
| 266266_at | 596.3 | P | 719.9 | P | 812.3 | P | 783 | P | --- |
| 266267_at | 1673.9 | P | 476.3 | P | 1395.3 | P | 689 | P | ATGSTU4 |
| 266268_at | 350.2 | P | 404.6 | P | 347.3 | P | 467.9 | P | --- |
| 266269_at | 124.3 | P | 128.9 | M | 124.7 | A | 159.1 | P | ATGSTU2 |
| 266270_at | 46.8 | P | 28.6 | A | 46.3 | A | 21.5 | A | ATGSTU3 |
| 266271_at | 547.6 | P | 97.7 | P | 340.9 | P | 117.3 | P | ATGSTU6 |
| 266272_at | 40.7 | P | 15.9 | A | 52.1 | P | 39.6 | P | --- |
| 266273_at | 46.6 | P | 43.2 | A | 34.6 | A | 33.7 | A | MTPB1 |
| 266274_at | 48 | A | 26.4 | A | 11.3 | A | 30 | A | --- |
| 266275_at | 416.6 | P | 509.1 | P | 589.2 | P | 576.2 | P | --- |
| 266276_at | 1.9 | A | 35.7 | A | 47.4 | A | 21.2 | A | TRI |
| 266277_at | 552.5 | P | 985.4 | P | 599 | P | 925.8 | P | --- |
| 266278_at | 62.2 | P | 59.2 | P | 47.6 | M | 31.4 | P | --- |
| 266279_at | 580.5 | P | 864.3 | P | 787.6 | P | 777.8 | P | --- |
| 266280_at | 180.3 | P | 132.5 | P | 187.7 | P | 145.6 | P | --- |
| 266281_at | 5.8 | A | 2.6 | A | 4.2 | A | 5.1 | A | --- |
| 266282_at | 9.2 | A | 29.2 | A | 59.7 | A | 26.5 | A | --- |
| 266283_at | 591.4 | P | 484.7 | P | 505.8 | P | 452.2 | P | --- |
| 266284_at | 307.3 | P | 335.8 | P | 253.6 | P | 250.5 | P | APUM2 |
| 266285_at | 1019.8 | P | 1038 | P | 1050.4 | P | 1014.2 | P | --- |
| 266286_at | 36.9 | P | 40.3 | P | 34.3 | P | 32.3 | P | --- |
| 266287_at | 13.1 | A | 2.7 | A | 20.6 | A | 25.2 | A | --- |
| 266288_s_at | 700.3 | P | 631.9 | P | 597.8 | P | 634.3 | P | APUM1 |
| 266289_at | 916.3 | P | 941.8 | P | 900.2 | P | 1041.5 | P | SMO2-2 |
| 266290_at | 1677.9 | P | 921.3 | P | 1549.5 | P | 1352 | P | ATGSTU1 |
| 266291_at | 600.9 | P | 509.7 | P | 588 | P | 498.3 | P | --- |
| 266292_at | 1630 | P | 333 | P | 379.6 | P | 372.7 | P | SAG13 |
| 266293_at | 764.8 | P | 948.1 | P | 582.8 | P | 683.2 | P | --- |
| 266294_at | 856.1 | P | 293.1 | P | 692.8 | P | 368.5 | P | --- |
| 266295_at | 602.4 | P | 1173.1 | P | 926.5 | P | 1124.6 | P | TUB7 |
| 266296_at | 2482 | P | 2307.3 | P | 2020.8 | P | 2515.8 | P | ATGSTU7 |
| 266297_at | 466.3 | P | 702.3 | P | 648.4 | P | 648.6 | P | PCNA2 |
| 266298_at | 249.3 | P | 267.8 | P | 179 | P | 255.1 | P | --- |
| 266299_at | 3350.4 | P | 2223.8 | P | 3522.7 | P | 2791.3 | P | ATGSTU5 |
| 266300_at | 824.8 | P | 847.8 | P | 936.6 | P | 789.2 | P | PIN4 |
| 266301_at | 3.7 | A | 3.1 | A | 6 | A | 2.8 | A | AtMYB81 |
| 266302_at | 630.7 | P | 1058.3 | P | 586.4 | P | 902.1 | P | EIL1 |
| 266303_at | 248.7 | P | 330.2 | P | 412.2 | P | 284.9 | P | --- |
| 266304_at | 486.7 | P | 415.7 | P | 387.5 | P | 431.7 | P | --- |
| 266305_at | 1.8 | A | 14.9 | A | 1.1 | A | 3.1 | A | TIL2 |
| 266306_at | 295.2 | P | 403.6 | P | 232.7 | P | 342.9 | P | --- |
| 266307_at | 8.3 | A | 1.2 | A | 14.7 | A | 12 | A | CYP705A8 |
| 266308_at | 6.9 | A | 3 | A | 2.7 | A | 1.3 | A | CYP705A9 |
| 266309_at | 65.1 | P | 134.8 | P | 111.3 | P | 113.7 | P | --- |
| 266310_at | 597.1 | P | 599.4 | P | 625.5 | P | 658.9 | P | FUS12 |
| 266311_at | 216.9 | P | 276.1 | P | 273.9 | P | 279.9 | P | --- |
| 266312_at | 3375.4 | P | 3435.3 | P | 2692.2 | P | 3407.4 | P | PAG1 |
| 266313_at | 584.2 | P | 311.1 | P | 637 | P | 381.1 | P | CIPK3 |
| 266314_at | 519.9 | P | 629.2 | P | 629.6 | P | 630.9 | P | AGO4 |
| 266315_at | 423 | P | 350.5 | P | 419 | P | 338.9 | P | SE |
| 266316_at | 767.8 | P | 622.5 | P | 320 | P | 396.4 | P | --- |
| 266317_at | 1509.2 | P | 1740.6 | P | 1667.1 | P | 1802.9 | P | CAM5 |
| 266318_at | 177.1 | P | 206.6 | P | 244.9 | P | 201 | P | FRS3 |
| 266319_s_at | 270.2 | P | 243.1 | P | 290.2 | P | 279.5 | P | KCS13 |
| 266320_at | 190.6 | P | 151.4 | P | 197.9 | P | 172.6 | P | --- |
| 266321_at | 98.7 | P | 87.4 | P | 109.7 | P | 100.6 | P | CYP78A6 |
| 266322_at | 262.8 | P | 388.8 | P | 398.8 | P | 375.4 | P | --- |
| 266323_at | 247.9 | P | 190.5 | P | 264.4 | P | 225.6 | P | CRK3 |
| 266324_at | 201.7 | P | 290.8 | P | 270.1 | P | 249.9 | P | --- |
| 266325_at | 212.3 | P | 248.2 | P | 208.6 | P | 210.8 | P | --- |
| 266326_at | 4556 | P | 1778.2 | P | 5058.2 | P | 2067.8 | P | CB5-C |
| 266327_at | 750.4 | P | 528.1 | P | 453.5 | P | 442.3 | P | ATHB-7 |
| 266328_at | 497.5 | P | 536.7 | P | 602.6 | P | 531.8 | P | --- |
| 266329_at | 874.1 | P | 1020.5 | P | 1109.7 | P | 961.7 | P | CRR3 |
| 266330_at | 13.1 | A | 22.5 | P | 18.3 | A | 47.7 | P | MLP329 |
| 266331_at | 701 | P | 1068.5 | P | 580.6 | P | 1035.5 | P | RGA1 |
| 266332_at | 109.6 | M | 74.6 | A | 129.7 | A | 95.1 | P | --- |
| 266333_at | 95.7 | P | 119.4 | P | 62.8 | P | 131 | P | AXL |
| 266334_at | 250.8 | P | 395.1 | P | 292.3 | P | 368 | P | --- |
| 266335_at | 268.2 | P | 348.5 | P | 493.9 | P | 344.9 | P | KAO2 |
| 266336_at | 41.7 | A | 44.8 | A | 67.6 | A | 36.3 | A | ZIP3 |
| 266337_at | 151.7 | P | 258.8 | P | 168.6 | P | 185.3 | P | ATGLR3.5 |
| 266338_at | 299.4 | P | 320.1 | P | 268.4 | P | 347 | P | GLR5 |
| 266339_s_at | 23.3 | P | 11.9 | A | 11.2 | A | 21.5 | A | --- |
| 266340_at | 119.1 | P | 176 | P | 168.3 | P | 132.1 | P | --- |
| 266341_at | 79 | P | 83 | P | 68.2 | P | 86.5 | P | --- |
| 266342_at | 426 | P | 486.8 | P | 424.4 | P | 446.8 | P | --- |
| 266343_at | 25.3 | A | 32.5 | A | 47.3 | A | 43.1 | A | --- |
| 266344_at | 33.3 | A | 44.2 | P | 77.9 | A | 48.4 | A | --- |
| 266345_at | 560.9 | P | 356.2 | P | 507.9 | P | 383.6 | P | --- |
| 266346_at | 5.3 | A | 1.7 | A | 3.8 | A | 1.9 | A | ATHB17 |
| 266347_at | 138.5 | P | 178.7 | P | 203.2 | P | 209.9 | P | --- |
| 266348_at | 1211.9 | P | 1166.3 | P | 1163.1 | P | 881.3 | P | ATMPK17 |
| 266349_at | 89.6 | A | 143.8 | A | 61.6 | A | 128.9 | A | --- |
| 266350_at | 695.1 | P | 600.5 | P | 550.1 | P | 563.6 | P | STL2P |
| 266351_at | 3255.9 | P | 2511.3 | P | 2941 | P | 2545.9 | P | --- |
| 266352_at | 201.7 | P | 174 | P | 212.7 | P | 189.2 | P | --- |
| 266353_at | 0.8 | A | 148 | P | 104.9 | P | 175.2 | P | MLP328 |
| 266354_at | 11.1 | A | 4.1 | A | 2.9 | A | 6 | A | PFS2 |
| 266355_at | 166.3 | P | 140.1 | P | 137 | P | 144.8 | P | --- |
| 266356_at | 36.2 | P | 22.5 | P | 9.7 | A | 27.1 | P | UCC1 |
| 266357_at | 168.7 | P | 100.5 | P | 171.3 | P | 110.8 | P | BAM6 |
| 266358_at | 356.5 | P | 334.6 | P | 444.1 | P | 392.2 | P | --- |
| 266359_at | 567 | P | 516.8 | P | 500 | P | 499.2 | P | CCT1 |
| 266360_at | 173 | P | 183.1 | P | 191.1 | P | 224 | P | FRS2 |
| 266361_at | 158.8 | P | 168.7 | P | 150 | P | 153.6 | P | --- |
| 266362_at | 98.7 | P | 133.7 | P | 155.1 | P | 138.9 | P | --- |
| 266363_at | 170.3 | P | 369.3 | P | 129.9 | P | 263.8 | P | --- |
| 266364_at | 162.8 | P | 478.6 | P | 75.7 | P | 204.5 | P | --- |
| 266365_at | 563.9 | P | 538.7 | P | 600.3 | P | 565.4 | P | GLU2 |
| 266366_at | 2675.3 | P | 2622.6 | P | 2310.7 | P | 2376.5 | P | --- |
| 266367_s_at | 3 | A | 14.2 | A | 15.4 | A | 27.5 | A | --- |
| 266368_at | 3456.5 | P | 978.5 | P | 3188.4 | P | 1452.6 | P | --- |
| 266369_at | 158.9 | P | 174.3 | P | 221.1 | P | 156 | P | BOP2 |

| | | | | | | | | | |
|---|---|---|---|---|---|---|---|---|---|
| 266370_at | 290.6 | P | 295.5 | P | 238.4 | P | 307.6 | P | --- |
| 266371_at | 6031.2 | P | 5528.3 | P | 4910.6 | P | 4868.1 | P | --- |
| 266372_at | 114.1 | P | 87 | P | 200.6 | P | 113.7 | P | ATRR3 |
| 266373_x_at | 47.2 | A | 24.7 | A | 31.9 | A | 26.3 | A | --- |
| 266374_at | 14.1 | A | 17.1 | A | 8.2 | A | 15.9 | A | --- |
| 266375_at | 3.8 | A | 3.5 | A | 1.5 | A | 2.9 | A | --- |
| 266376_at | 33.1 | P | 87.8 | P | 59.1 | A | 65.4 | P | --- |
| 266377_at | 0.9 | A | 0.8 | A | 1.3 | A | 0.8 | A | --- |
| 266378_at | 9.5 | A | 21.5 | A | 25.1 | A | 34.5 | A | --- |
| 266379_at | 1.9 | A | 3 | A | 3.4 | A | 27.6 | A | --- |
| 266380_at | 230.7 | P | 251.2 | P | 223.5 | P | 288.6 | P | MEE13 |
| 266381_at | 1 | A | 1.9 | A | 30.9 | A | 3.9 | A | ATSUC8 |
| 266382_at | 38.8 | P | 49.2 | P | 40.9 | M | 34.7 | A | --- |
| 266383_at | 110.3 | P | 101.9 | P | 86 | P | 99.5 | P | ATPRB1 |
| 266384_at | 143.4 | P | 163.2 | P | 170.8 | P | 174.1 | P | --- |
| 266385_at | 76.6 | P | 754.3 | P | 60.3 | A | 363.2 | P | PR1 |
| 266386_at | 16.7 | A | 16.5 | A | 24.1 | A | 13.8 | A | HDG3 |
| 266387_at | 1.9 | A | 0.6 | A | 0.8 | A | 15.7 | A | --- |
| 266388_at | 55.9 | P | 63.1 | P | 122.5 | P | 49.5 | P | --- |
| 266389_s_at | 384.9 | P | 413.6 | P | 284.9 | P | 449 | P | --- |
| 266390_at | 5 | A | 33.5 | A | 41.8 | A | 32.8 | A | --- |
| 266391_at | 145.1 | P | 212.9 | P | 179.1 | P | 168.2 | P | --- |
| 266392_at | 2.9 | A | 7.6 | A | 8.7 | A | 2.8 | A | M10 |
| 266393_at | 5.2 | A | 0.8 | A | 1.9 | A | 3.8 | A | M17 |
| 266394_at | 334.1 | P | 532.4 | P | 393.7 | P | 467.7 | P | ARA4 |
| 266395_at | 3230.2 | P | 2844.5 | P | 3343.7 | P | 2293 | P | --- |
| 266396_at | 296 | P | 94.9 | P | 165.5 | P | 119 | P | --- |
| 266397_at | 23.6 | A | 29.5 | A | 45.8 | M | 14.7 | A | --- |
| 266398_at | 180.9 | P | 208.3 | P | 150.6 | P | 279.6 | P | --- |
| 266399_at | 1030.4 | P | 1274.6 | P | 1133.8 | P | 1433.1 | P | PECT1 |
| 266400_at | 223.7 | P | 242.7 | P | 194.3 | P | 370.5 | P | --- |
| 266401_s_at | 86 | P | 146 | P | 124.9 | P | 139.6 | P | CDKB1;2 |
| 266402_at | 329.4 | P | 434.1 | P | 291.3 | P | 392.9 | P | --- |
| 266403_at | 33.8 | A | 44.5 | P | 40.9 | P | 74.8 | P | --- |
| 266404_at | 16.5 | A | 21.3 | A | 4.9 | A | 2.9 | A | --- |
| 266405_at | 275.4 | P | 281 | P | 271 | P | 288.8 | P | --- |
| 266406_at | 75.9 | P | 68.5 | P | 104.1 | P | 80.2 | P | --- |
| 266407_at | 628.5 | P | 742.2 | P | 783.2 | P | 750.6 | P | TFIIS |
| 266408_at | 58.6 | A | 44.7 | A | 77.1 | A | 45.8 | A | --- |
| 266409_at | 1.5 | A | 1.6 | A | 2.6 | A | 14.9 | A | --- |
| 266410_at | 391.1 | P | 394.4 | P | 496.5 | P | 401.8 | P | EMB2765 |
| 266411_at | 236.4 | P | 235.6 | P | 293.7 | P | 266.2 | P | --- |
| 266412_at | 52.6 | M | 75.2 | P | 65.6 | A | 63.2 | P | MAP65-5 |
| 266413_at | 578.4 | P | 646.9 | P | 579.5 | P | 678.9 | P | --- |
| 266414_at | 801.8 | P | 742.7 | P | 776.6 | P | 761.5 | P | MVD1 |
| 266415_at | 12123.1 | P | 17276.8 | P | 8163.5 | P | 18408.6 | P | LTP2 |
| 266416_at | 1167.7 | P | 1394.4 | P | 1254.5 | P | 1395.9 | P | --- |
| 266417_at | 1780.1 | P | 1757.5 | P | 2031.4 | P | 1931.5 | P | --- |
| 266418_at | 982.1 | P | 335.1 | P | 977.4 | P | 454.4 | P | ANNAT4 |
| 266419_at | 177.4 | P | 73.4 | P | 103.7 | P | 56.1 | P | ANNAT3 |
| 266420_at | 1187.6 | P | 807.8 | P | 746.3 | P | 816.7 | P | --- |
| 266421_at | 15990.8 | P | 15298.1 | P | 13317.7 | P | 16412.3 | P | LP1 |
| 266422_at | 469.6 | P | 463.8 | P | 582.3 | P | 532.2 | P | GAUT7 |
| 266423_at | 122.7 | P | 139.4 | P | 191.3 | P | 184.1 | P | RPB5D |
| 266424_at | 184.4 | P | 95.4 | P | 145.9 | P | 143.1 | P | --- |
| 266425_at | 44.6 | A | 45.5 | A | 23.4 | A | 79.9 | A | --- |
| 266426_x_at | 83.2 | P | 67 | P | 158.9 | P | 94 | P | --- |
| 266427_at | 102.5 | P | 139.1 | M | 18.5 | A | 179.6 | P | --- |
| 266428_at | 630.6 | P | 765.5 | P | 624.8 | P | 581 | P | --- |
| 266429_at | 28.3 | P | 45.6 | P | 58.4 | P | 12.8 | A | --- |
| 266430_at | 34.2 | A | 14.8 | A | 39.8 | A | 21.9 | A | --- |
| 266431_at | 5.5 | A | 1.6 | A | 43.6 | A | 5.1 | A | --- |
| 266432_s_at | 0.6 | A | 1.2 | A | 1 | A | 0.4 | A | --- |
| 266433_at | 5.8 | A | 3.2 | A | 28 | A | 8.6 | A | --- |
| 266434_s_at | 1.9 | A | 1.7 | A | 2.4 | A | 1.8 | A | --- |
| 266435_s_at | 6 | A | 0.9 | A | 3.6 | A | 1.3 | A | --- |
| 266436_at | 140 | P | 240.7 | P | 211.2 | P | 224.9 | P | --- |
| 266437_at | 755.4 | P | 828.1 | P | 648.7 | P | 792.8 | P | --- |
| 266438_at | 155.7 | P | 88.5 | P | 134.4 | P | 79.1 | P | --- |
| 266439_s_at | 128.6 | P | 124.1 | P | 119 | P | 111.1 | P | --- |
| 266440_at | 245.4 | P | 254.2 | P | 257.3 | P | 330.6 | P | --- |
| 266441_at | 111.6 | P | 109.6 | P | 113.2 | P | 88 | P | --- |
| 266442_at | 217 | P | 187.8 | P | 225.9 | P | 228.1 | P | --- |
| 266443_at | 4 | A | 8 | A | 11.7 | A | 11.3 | A | --- |
| 266444_at | 31.3 | P | 32.1 | P | 50.1 | P | 16.3 | A | --- |
| 266445_at | 766 | P | 697 | P | 760.3 | P | 814.1 | P | SPPL3 |
| 266446_at | 66.9 | P | 68.6 | P | 82.1 | P | 54.6 | P | --- |
| 266447_at | 435.7 | P | 673.7 | P | 479.9 | P | 647 | P | MSS3 |
| 266448_s_at | 251.3 | P | 391.7 | P | 277.5 | P | 319.7 | P | --- |
| 266449_at | 482.4 | P | 433.1 | P | 393.2 | P | 463.3 | P | AT-P4H-1 |
| 266450_s_at | 302.2 | P | 284.4 | P | 349.4 | P | 276.2 | P | --- |
| 266451_at | 4140.7 | P | 3926.9 | P | 3610.5 | P | 3292.9 | P | --- |
| 266452_at | 129.4 | P | 110.5 | P | 141.9 | P | 132.6 | P | --- |
| 266453_at | 33.5 | P | 38.6 | A | 63.8 | P | 35.4 | A | --- |
| 266454_at | 1.3 | A | 11.4 | A | 4.9 | A | 18 | A | --- |
| 266455_at | 5.6 | A | 29 | A | 56.1 | A | 12.5 | A | --- |
| 266456_at | 182.9 | P | 133 | P | 213.2 | P | 78.7 | P | NAI1 |
| 266457_at | 2265.5 | P | 2005 | P | 2020.5 | P | 1994.1 | P | PMDH1 |
| 266458_at | 6031.8 | P | 4904.7 | P | 5937.5 | P | 4918.8 | P | --- |
| 266459_at | 1432.5 | P | 1079 | P | 1280.1 | P | 1093.3 | P | --- |
| 266460_at | 434.2 | P | 659.2 | P | 718.6 | P | 622.1 | P | AGP26 |
| 266461_at | 8646.5 | P | 7984.7 | P | 9730.6 | P | 8006.6 | P | ATGSTF8 |
| 266462_at | 201.3 | P | 49 | A | 97.6 | M | 37 | A | --- |
| 266463_at | 1407.9 | P | 1101 | P | 1395.8 | P | 1249.2 | P | --- |
| 266464_at | 920.5 | P | 626.6 | P | 821.9 | P | 736.4 | P | ATMRP4 |
| 266465_at | 50.3 | P | 140.7 | P | 95.8 | P | 99.2 | P | GH3.9 |
| 266466_at | 517 | P | 375.3 | P | 542.3 | P | 432.6 | P | --- |
| 266467_at | 571.3 | P | 498.4 | P | 607.7 | P | 460.8 | P | ALG3 |
| 266468_at | 445.1 | P | 607.3 | P | 448.3 | P | 577.1 | P | --- |
| 266469_at | 34.4 | P | 32.4 | P | 36.1 | P | 11.4 | A | MYB14 |
| 266470_at | 3.3 | A | 18.7 | A | 4 | A | 18.2 | A | --- |
| 266471_at | 949.2 | P | 726.2 | P | 1030.5 | P | 799.9 | P | --- |
| 266472_at | 113.3 | P | 179 | P | 206.5 | P | 126.8 | P | --- |
| 266473_at | 78.1 | A | 279 | P | 225.4 | A | 201 | P | --- |
| 266474_at | 184.1 | P | 409.9 | P | 263 | P | 422.4 | P | --- |
| 266475_at | 10.9 | A | 14.8 | A | 21.5 | A | 13.6 | A | --- |
| 266476_at | 936.5 | P | 1277 | P | 819.6 | P | 1132.1 | P | --- |
| 266477_at | 2.7 | A | 1.8 | A | 5.3 | A | 18.3 | A | CLE6 |
| 266478_at | 412.6 | P | 469.5 | P | 597.6 | P | 531.5 | P | SYCO ARATH |

| | | | | | | | | | |
|---|---|---|---|---|---|---|---|---|---|
| 266479_at | 125.7 | P | 174.1 | P | 146.2 | P | 144.5 | P | LSH3 |
| 266480_at | 155.7 | P | 124 | P | 87.8 | P | 146.9 | P | --- |
| 266481_at | 581.5 | P | 399.4 | P | 651.6 | P | 443.5 | P | TCP10 |
| 266482_at | 598.3 | P | 761.9 | P | 636.8 | P | 702.2 | P | --- |
| 266483_at | 1247.8 | P | 1342.3 | P | 1524.4 | P | 1351.3 | P | CRR6 |
| 266484_at | 461.6 | P | 506.6 | P | 458.1 | P | 521 | P | SCC3 |
| 266485_at | 321.5 | P | 457.6 | P | 402.6 | P | 416.1 | P | --- |
| 266486_at | 24.6 | A | 23.7 | A | 28.7 | A | 24.1 | A | --- |
| 266487_at | 1.5 | A | 6 | A | 15.6 | A | 0.6 | A | --- |
| 266488_at | 26.4 | A | 2.4 | A | 3.4 | A | 7.2 | A | --- |
| 266489_at | 239.4 | P | 378.3 | P | 196.8 | P | 394.1 | P | NPSN11 |
| 266490_at | 2.9 | A | 25 | A | 44.9 | A | 8.6 | A | --- |
| 266491_at | 5 | A | 17.9 | A | 1.7 | A | 6.5 | A | --- |
| 266492_at | 7.8 | A | 12.8 | A | 8.9 | A | 5.2 | A | --- |
| 266493_at | 15.8 | A | 16.1 | A | 11.9 | M | 26.1 | P | --- |
| 266494_at | 2.5 | A | 3.2 | A | 6.5 | A | 2.1 | A | PRK2A |
| 266495_at | 875.3 | P | 1396.2 | P | 1074.1 | P | 1375.6 | P | CAS1 |
| 266496_at | 22.8 | A | 45.1 | A | 37.1 | A | 30.4 | A | --- |
| 266497_at | 38.7 | A | 51 | A | 37.5 | A | 33 | A | --- |
| 266498_at | 37 | A | 46.1 | A | 52 | A | 37.2 | A | --- |
| 266499_at | 19 | A | 30.7 | A | 36 | A | 12.8 | A | --- |
| 266500_at | 966 | P | 1528.8 | P | 1012.1 | P | 1647.9 | P | PLA2-ALPHA |
| 266501_at | 356.2 | P | 284.7 | P | 319.5 | P | 269.4 | P | HEN2 |
| 266502_at | 36.3 | A | 44.7 | A | 53.7 | P | 23.9 | A | --- |
| 266503_at | 67.9 | A | 24.7 | A | 58.4 | A | 54.2 | P | --- |
| 266504_at | 66 | P | 55.8 | P | 73.3 | P | 65.2 | P | --- |
| 266505_at | 215.9 | P | 223.7 | P | 237.8 | P | 231.7 | P | --- |
| 266506_at | 414.1 | P | 516.2 | P | 398.2 | P | 445.1 | P | --- |
| 266507_at | 14.1 | A | 4.1 | A | 11.4 | A | 5.8 | A | --- |
| 266508_at | 35.9 | A | 34.8 | A | 3 | A | 52 | A | --- |
| 266509_at | 2434.7 | P | 2768.8 | P | 2629 | P | 2813.4 | P | DEGP2 |
| 266510_at | 471.9 | P | 357.9 | P | 468 | P | 363.4 | P | SWA1 |
| 266511_at | 111.6 | P | 187.4 | P | 163.7 | P | 135 | P | --- |
| 266512_at | 457 | P | 394.1 | P | 458.5 | P | 417.6 | P | --- |
| 266513_at | 361.9 | P | 366.8 | P | 360.4 | P | 438.3 | P | --- |
| 266514_at | 931.5 | P | 627.8 | P | 679.1 | P | 762.7 | P | --- |
| 266515_at | 431.3 | P | 516.7 | P | 369.8 | P | 454.6 | P | AtTLP3 |
| 266516_at | 150.4 | P | 388.6 | P | 154.4 | P | 479.4 | P | --- |
| 266517_at | 870.1 | P | 1141 | P | 1170.4 | P | 1191.1 | P | --- |
| 266518_at | 40.8 | A | 88.2 | P | 92.6 | A | 93.1 | A | --- |
| 266519_at | 313.6 | P | 439.7 | P | 381.4 | P | 429.6 | P | --- |
| 266520_at | 427.2 | P | 377.3 | P | 326.9 | P | 387.1 | P | ATCNGC6 |
| 266521_at | 1603.9 | P | 1788.5 | P | 1533.1 | P | 1666.7 | P | --- |
| 266522_at | 173.5 | P | 212.4 | P | 198 | P | 215.8 | P | UBC23 |
| 266523_at | 717.8 | P | 701.9 | P | 726.7 | P | 818.2 | P | TRN1 |
| 266524_at | 18.5 | A | 11.7 | A | 31.2 | A | 22.9 | A | --- |
| 266525_at | 27.6 | A | 68.7 | P | 90.8 | P | 46.6 | P | MEE15 |
| 266526_at | 19.1 | A | 8.9 | A | 7 | A | 16 | A | --- |
| 266527_at | 15.3 | A | 18.2 | A | 33.4 | A | 14.3 | A | --- |
| 266528_at | 53 | P | 14.3 | A | 40.9 | A | 38.4 | P | --- |
| 266529_at | 117.7 | P | 116.9 | P | 99 | P | 150.5 | P | --- |
| 266530_at | 27.1 | A | 19 | A | 33.3 | A | 13.6 | A | AMS |
| 266531_at | 2.8 | A | 12.4 | A | 36.5 | A | 3.6 | A | --- |
| 266532_at | 61.6 | P | 142.5 | P | 73.5 | P | 155.2 | P | --- |
| 266533_s_at | 6006.4 | P | 6176.3 | P | 6600 | P | 6783.2 | P | PIP2;8 |
| 266534_at | 700.6 | P | 597.3 | P | 624.9 | P | 740.9 | P | --- |
| 266535_s_at | 2231.6 | P | 1966.2 | P | 2157.9 | P | 2129.9 | P | --- |
| 266536_at | 463 | P | 507.6 | P | 533.5 | P | 504 | P | --- |
| 266537_at | 615.6 | P | 684.1 | P | 703.7 | P | 673.9 | P | --- |
| 266538_at | 135.1 | P | 207.3 | P | 246.1 | P | 233.8 | P | --- |
| 266539_at | 46.8 | A | 71.8 | P | 87.3 | A | 78.8 | P | SUVH5 |
| 266540_at | 12.7 | A | 49 | P | 61.8 | A | 51.4 | A | --- |
| 266541_at | 299.8 | P | 365.2 | P | 335.1 | P | 405.2 | P | GRL |
| 266542_at | 241.9 | P | 278.2 | P | 311.1 | P | 238.9 | P | ARAD1 |
| 266543_at | 26.6 | A | 22.5 | A | 15.6 | A | 36.3 | A | --- |
| 266544_at | 11 | A | 2.8 | A | 2.4 | A | 1.4 | A | --- |
| 266545_at | 199.9 | P | 246.8 | P | 187 | P | 175.4 | P | --- |
| 266546_at | 22.1 | A | 5.5 | A | 14.6 | A | 26.3 | A | --- |
| 266547_at | 571.1 | P | 566.8 | P | 489.5 | P | 527.4 | P | ATEYA |
| 266548_at | 7.6 | A | 5.8 | A | 7.8 | A | 16.9 | A | RPA |
| 266549_at | 9.8 | A | 27.1 | A | 36 | A | 42.7 | A | EXL1 |
| 266550_s_at | 26.9 | A | 10.6 | A | 8.8 | A | 21.2 | A | --- |
| 266551_at | 1103.1 | P | 1284.9 | P | 1326.7 | P | 1234 | P | --- |
| 266552_at | 2375 | P | 3922.7 | P | 2472.1 | P | 3603.2 | P | AGP16 |
| 266553_at | 1461.8 | P | 1534.9 | P | 1322.5 | P | 1597.2 | P | --- |
| 266554_s_at | 3679.4 | P | 2604.7 | P | 4362.8 | P | 2553 | P | --- |
| 266555_at | 171.2 | P | 148.4 | P | 147.7 | P | 184.3 | P | GBF3 |
| 266556_at | 2316.7 | P | 2007.5 | P | 2356.4 | P | 2209.5 | P | --- |
| 266557_at | 755.1 | P | 790.2 | P | 689.5 | P | 743.8 | P | --- |
| 266558_at | 15 | A | 3.2 | A | 2.9 | A | 1.8 | A | --- |
| 266559_at | 15.2 | A | 10.7 | A | 5 | A | 1.3 | A | --- |
| 266560_at | 136.5 | P | 180.3 | P | 204.6 | P | 194.8 | P | --- |
| 266561_at | 38.9 | P | 39.8 | P | 44.4 | A | 60.4 | P | --- |
| 266562_at | 24.9 | A | 15 | A | 38.8 | A | 26.2 | A | --- |
| 266563_at | 33.6 | A | 20.2 | A | 4.3 | A | 10.7 | A | --- |
| 266564_at | 1.5 | A | 2.7 | A | 23.3 | A | 5.3 | A | scpl22 |
| 266565_at | 22.8 | A | 7.6 | A | 8.1 | A | 46.8 | A | scpl23 |
| 266566_at | 541.7 | P | 570.8 | P | 343.5 | P | 445.3 | P | --- |
| 266567_at | 488.1 | P | 556.1 | P | 548 | P | 482.5 | P | --- |
| 266568_at | 50.4 | A | 40.8 | A | 36.4 | A | 62 | A | --- |
| 266569_at | 44.7 | A | 45.8 | A | 13.2 | A | 36.5 | A | --- |
| 266570_at | 2984.3 | P | 3277.7 | P | 3330.6 | P | 2714.8 | P | --- |
| 266571_at | 26 | A | 2.1 | A | 24.3 | A | 3.5 | A | --- |
| 266572_at | 468.5 | P | 631.3 | P | 312 | P | 521.6 | P | --- |
| 266573_at | 1.6 | A | 3.8 | A | 2.2 | A | 2.1 | A | --- |
| 266574_at | 193.2 | P | 216.1 | P | 184.4 | P | 219.6 | P | --- |
| 266575_at | 816.4 | P | 1151 | P | 949.9 | P | 1076.9 | P | --- |
| 266576_at | 1214.1 | P | 1388.5 | P | 1308.9 | P | 1391.6 | P | --- |
| 266577_at | 41.9 | P | 46.5 | P | 38 | P | 40.3 | P | --- |
| 266578_at | 1441.2 | P | 514.3 | P | 1486.2 | P | 1084.6 | P | --- |
| 266579_at | 2891.4 | P | 2717.3 | P | 2564.8 | P | 2133 | P | SNRNP-G |
| 266580_at | 424.1 | P | 402 | P | 350.5 | P | 448.8 | P | --- |
| 266581_at | 313 | P | 300.3 | P | 282.4 | P | 254.3 | P | --- |
| 266582_at | 944.8 | P | 1045.4 | P | 849.7 | P | 981.9 | P | --- |
| 266583_at | 1896.3 | P | 2226.6 | P | 1488.4 | P | 1335.1 | P | --- |
| 266584_s_at | 21.9 | M | 1.6 | A | 29.1 | A | 31.1 | A | ST4C |
| 266585_at | 2.5 | A | 4.6 | A | 3.6 | A | 1.7 | A | --- |
| 266586_s_at | 1.5 | A | 4.3 | A | 4.8 | A | 12.4 | A | --- |
| 266587_at | 1713.8 | P | 2187.5 | P | 2022.4 | P | 2172.4 | P | --- |

| | | | | | | | | | |
|---|---|---|---|---|---|---|---|---|---|
| 266588_at | 1948.5 | P | 2688.3 | P | 2209.3 | P | 2587.7 | P | AGP9 |
| 266589_at | 209.7 | P | 201.9 | P | 218 | P | 199.8 | P | --- |
| 266590_at | 106.4 | P | 108.7 | P | 95.3 | P | 104.3 | P | BAG6 |
| 266591_at | 377.7 | P | 674.2 | P | 462.6 | P | 634.6 | P | ABIL1 |
| 266592_at | 56.3 | P | 90.3 | P | 68.2 | P | 69.7 | P | --- |
| 266593_at | 73.8 | P | 48 | P | 71.9 | P | 89.1 | P | --- |
| 266594_at | 5.8 | A | 16.3 | A | 12 | A | 2.3 | A | --- |
| 266595_at | 339 | P | 368.2 | P | 217.9 | P | 367.7 | P | GC4 |
| 266596_at | 83.2 | P | 76.9 | P | 94.3 | P | 64 | M | --- |
| 266597_at | 15.5 | A | 2.3 | A | 9.6 | A | 12.3 | A | WRKY43 |
| 266598_at | 211.8 | P | 268.1 | P | 297.3 | P | 249.6 | P | KPHMT1 |
| 266599_at | 1692.7 | P | 1606.9 | P | 1081.8 | P | 1366.6 | P | --- |
| 266600_at | 194.5 | P | 204.1 | P | 208.9 | P | 186.1 | P | MPK12 |
| 266601_at | 334.7 | P | 377.5 | P | 290.8 | P | 390.3 | P | --- |
| 266602_at | 26.3 | A | 8.1 | A | 10.4 | A | 27.9 | A | --- |
| 266603_at | 27.2 | A | 42.8 | A | 52 | A | 48.7 | A | --- |
| 266604_at | 737.1 | P | 756.8 | P | 732.8 | P | 617.8 | P | UBC6 |
| 266605_at | 387.6 | P | 465.5 | P | 372.9 | P | 396.2 | P | BRM |
| 266606_at | 100.5 | P | 55.8 | A | 131.4 | P | 19.6 | A | CRF5 |
| 266607_at | 3 | A | 4.5 | A | 5.4 | A | 3.7 | A | --- |
| 266608_at | 1516.7 | P | 1564.8 | P | 1881.7 | P | 1546.5 | P | --- |
| 266609_at | 462.1 | P | 537.1 | P | 496.3 | P | 572.9 | P | SRO1 |
| 266610_at | 10.3 | A | 0.9 | A | 3.8 | A | 6.2 | A | --- |
| 266611_at | 3.9 | A | 3.8 | A | 14.8 | A | 8.6 | A | GH3.1 |
| 266612_at | 119.8 | P | 149.6 | P | 138.7 | P | 156.2 | P | --- |
| 266613_at | 1232 | P | 650.3 | P | 837.2 | P | 575.1 | P | --- |
| 266614_at | 4024.9 | P | 4362.9 | P | 4203.6 | P | 4253.2 | P | --- |
| 266615_s_at | 477.3 | P | 472.2 | P | 458.1 | P | 527.2 | P | CTF2A |
| 266616_at | 71.2 | P | 96 | P | 99.2 | P | 112.1 | P | CDC6 |
| 266617_at | 2008.3 | P | 2955.6 | P | 2022.5 | P | 3132.3 | P | --- |
| 266618_at | 253.7 | P | 187.8 | P | 193.2 | P | 184.7 | P | --- |
| 266619_at | 34.8 | P | 55.4 | P | 59.4 | P | 35.1 | P | --- |
| 266620_at | 429.9 | P | 554.3 | P | 423.8 | P | 565.1 | P | --- |
| 266621_at | 507.5 | P | 613.7 | P | 336.9 | P | 563.1 | P | --- |
| 266622_at | 62.5 | A | 44.8 | A | 100.3 | A | 76.2 | A | --- |
| 266623_at | 114.2 | P | 173.4 | P | 207.8 | P | 108.2 | P | --- |
| 266624_s_at | 778.5 | P | 642.3 | P | 754.9 | P | 715 | P | --- |
| 266625_at | 45.5 | A | 48.7 | A | 40.1 | A | 21.7 | A | --- |
| 266626_at | 200.5 | P | 165.9 | P | 171.3 | P | 159.6 | P | --- |
| 266627_at | 55.2 | P | 68.4 | P | 98.5 | P | 39 | A | MEE29 |
| 266628_at | 190.1 | P | 204.9 | P | 117.3 | P | 168.7 | P | --- |
| 266629_at | 26.6 | A | 27.5 | P | 55.6 | A | 30.4 | A | --- |
| 266630_at | 2.6 | A | 3.7 | A | 19.8 | A | 4.8 | A | --- |
| 266631_at | 42.8 | A | 3.3 | A | 58.3 | A | 1.6 | A | --- |
| 266632_at | 104.6 | P | 24.5 | A | 74.2 | P | 31.6 | A | --- |
| 266633_at | 140.1 | P | 127.6 | P | 198.3 | P | 135.7 | P | --- |
| 266634_at | 71.4 | P | 125.4 | P | 102.6 | P | 139 | P | --- |
| 266635_at | 383.2 | P | 481.7 | P | 426.3 | P | 412.5 | P | --- |
| 266636_at | 9903.8 | P | 9383.8 | P | 9972.6 | P | 9652 | P | GDCH |
| 266637_at | 2.5 | A | 11.3 | A | 55.2 | A | 33.5 | A | BRXL1 |
| 266638_at | 2670.9 | P | 2706.7 | P | 2471.2 | P | 2381.9 | P | --- |
| 266639_at | 768.1 | P | 866.4 | P | 668.8 | P | 836.1 | P | DAD2 |
| 266640_at | 12.2 | A | 53 | A | 30 | A | 37.1 | A | --- |
| 266641_at | 458.5 | P | 516.6 | P | 646.7 | P | 611.4 | P | --- |
| 266642_at | 2263.6 | P | 2361.5 | P | 2059.9 | P | 2052.8 | P | --- |
| 266643_s_at | 116.3 | P | 89.1 | P | 133.6 | P | 125.2 | P | --- |
| 266644_at | 269 | P | 212.5 | P | 183.8 | P | 258 | P | --- |
| 266645_at | 2.7 | A | 3.7 | A | 6.5 | A | 2.7 | A | --- |
| 266646_at | 1622.1 | P | 1578.8 | P | 1464 | P | 1974.3 | P | ATPH1 |
| 266647_at | 268.9 | P | 346.6 | P | 637.4 | P | 345.8 | P | --- |
| 266648_at | 780.4 | P | 735 | P | 797.8 | P | 840.7 | P | OVA4 |
| 266649_at | 111.6 | P | 86.5 | P | 72.8 | A | 104.5 | A | TIP4;1 |
| 266650_at | 503.2 | P | 522.2 | P | 455.6 | P | 554.6 | P | --- |
| 266651_at | 445.6 | P | 505.5 | P | 407.9 | P | 496.2 | P | --- |
| 266652_at | 730 | P | 765.1 | P | 690.2 | P | 755.6 | P | --- |
| 266653_at | 412.8 | P | 452.1 | P | 515.3 | P | 436.5 | P | --- |
| 266654_at | 19.4 | A | 3.3 | A | 5.2 | A | 3.1 | A | --- |
| 266655_at | 109.6 | P | 250.1 | P | 121 | P | 252.3 | P | AtAUR2 |
| 266656_at | 133.9 | P | 204.4 | P | 146.6 | P | 154.6 | P | ATCTH |
| 266657_at | 1596.4 | P | 1382.4 | P | 1578.4 | P | 1465.6 | P | --- |
| 266658_at | 197.4 | P | 97.9 | P | 115 | P | 127.8 | P | --- |
| 266659_at | 323 | P | 329.8 | P | 321.4 | P | 361 | P | --- |
| 266660_at | 216.2 | P | 252 | P | 245.3 | P | 242.7 | P | --- |
| 266661_at | 49.3 | A | 39.7 | A | 9 | A | 61.1 | A | --- |
| 266662_at | 1095.1 | P | 1285.6 | P | 875.8 | P | 1367.5 | P | --- |
| 266663_at | 148.9 | P | 153.8 | P | 202.7 | P | 174.9 | P | --- |
| 266664_at | 4.3 | A | 2.5 | A | 38.9 | A | 31.1 | A | --- |
| 266665_at | 8.7 | A | 9.6 | A | 2.7 | A | 2.8 | A | --- |
| 266666_at | 19.4 | A | 46.6 | P | 52.2 | M | 34.4 | P | --- |
| 266667_at | 2.4 | A | 1.2 | A | 4.7 | A | 14.6 | A | --- |
| 266668_at | 311.9 | P | 286.1 | P | 298.7 | P | 316.8 | P | --- |
| 266669_at | 4.9 | A | 5.1 | A | 5.4 | A | 3.5 | A | UGT71C1 |
| 266670_at | 13.1 | A | 42.3 | A | 71.7 | A | 48.7 | A | UGT71C2 |
| 266671_at | 378.9 | P | 259.3 | P | 325.5 | P | 296.8 | P | ASA2 |
| 266672_at | 1042.5 | P | 1607 | P | 850.4 | P | 1331.6 | P | PHT4;1 |
| 266673_at | 9786.3 | P | 9680.4 | P | 11393.3 | P | 10565.5 | P | THIC |
| 266674_at | 2.3 | A | 21.3 | A | 9.5 | A | 10.9 | A | --- |
| 266675_s_at | 1.4 | A | 1 | A | 8.5 | A | 1.3 | A | --- |
| 266676_s_at | 1.5 | A | 13.9 | A | 19.9 | A | 15.3 | A | --- |
| 266677_at | 12.3 | A | 25.4 | A | 43.1 | A | 20.7 | A | --- |
| 266678_at | 1486.3 | P | 1550.6 | P | 1100.9 | P | 1507.8 | P | --- |
| 266679_at | 247.6 | P | 335.7 | P | 266.1 | P | 261.8 | P | --- |
| 266680_s_at | 105.9 | P | 11.5 | A | 166 | P | 3.5 | A | --- |
| 266681_at | 1 | A | 0.9 | A | 13.8 | A | 7.4 | A | --- |
| 266682_at | 87.1 | P | 138.5 | P | 149 | P | 122.3 | P | --- |
| 266683_at | 173.7 | P | 208.2 | P | 208.3 | P | 164.2 | P | GC1 |
| 266684_at | 281.9 | P | 249.5 | P | 249.3 | P | 284.8 | P | rps15ab |
| 266685_at | 157.5 | P | 167.3 | P | 202.4 | P | 145.1 | P | --- |
| 266686_at | 1.6 | A | 3.1 | A | 2.1 | A | 1.7 | A | --- |
| 266687_at | 380 | P | 321 | P | 445.2 | P | 368.7 | P | PRMT1A |
| 266688_at | 1.1 | A | 9.7 | A | 21.7 | A | 3.5 | A | --- |
| 266689_at | 20.5 | A | 44.8 | A | 52.5 | A | 51.5 | P | --- |
| 266690_at | 93.3 | A | 59.2 | A | 81.2 | A | 40.9 | A | ATNADP-ME1 |
| 266691_at | 17.8 | A | 27.3 | A | 15.9 | A | 19.5 | A | --- |
| 266692_at | 14 | A | 10.9 | A | 22 | A | 2.1 | A | --- |
| 266693_at | 663.5 | P | 91.7 | P | 498.6 | P | 86.1 | P | MIOX2 |
| 266694_at | 1621.7 | P | 1656.2 | P | 1493.9 | P | 1774.9 | P | --- |
| 266695_at | 465.2 | P | 498.6 | P | 648.9 | P | 510.9 | P | --- |
| 266696_at | 1441.8 | P | 1184.2 | P | 1095.7 | P | 1220.6 | P | --- |

| | | | | | | | | | |
|---|---|---|---|---|---|---|---|---|---|
| 266697_at | 3.9 | A | 4.4 | A | 6.5 | A | 4.9 | A | PRF5 |
| 266698_at | 970 | P | 1002.3 | P | 992.4 | P | 920.7 | P | SNF7.2 |
| 266699_at | 8975.7 | P | 8522.9 | P | 10498.9 | P | 9443.8 | P | --- |
| 266700_at | 2203.1 | P | 1836.6 | P | 2479.7 | P | 2060.3 | P | --- |
| 266701_at | 1389.5 | P | 1449.6 | P | 1328.6 | P | 1561.8 | P | PRF1 |
| 266702_at | 792.1 | P | 827 | P | 904.7 | P | 862 | P | HXK2 |
| 266703_at | 265.1 | P | 347.2 | P | 316.8 | P | 381.1 | P | --- |
| 266704_at | 1436 | P | 1304.8 | P | 1576.3 | P | 1276 | P | --- |
| 266705_at | 2140.5 | P | 1872.6 | P | 2230.1 | P | 1945.4 | P | --- |
| 266706_at | 12.8 | A | 21.6 | A | 3.3 | A | 3.6 | A | --- |
| 266707_at | 425.6 | P | 549.1 | P | 364.5 | P | 481.1 | P | --- |
| 266708_at | 33.1 | A | 53.6 | A | 17.9 | A | 59 | A | --- |
| 266709_at | 1926.1 | P | 2381.8 | P | 2010.4 | P | 2343.2 | P | ATSPP |
| 266710_at | 4 | A | 26.9 | A | 33 | A | 10.4 | A | --- |
| 266711_at | 0.8 | A | 25.6 | A | 18.6 | A | 3.8 | A | --- |
| 266712_at | 3.7 | A | 40 | A | 35.9 | A | 33.7 | A | --- |
| 266713_at | 18.2 | A | 4.1 | A | 4.1 | A | 9.3 | A | --- |
| 266714_at | 5.9 | A | 6 | A | 3.3 | A | 4 | A | EMB2301 |
| 266715_at | 104.2 | P | 115.2 | P | 133.7 | P | 148.7 | P | --- |
| 266716_at | 7596.9 | P | 9949.2 | P | 7954.8 | P | 8267.2 | P | PSI-P |
| 266717_at | 323.3 | P | 290.4 | P | 310.1 | P | 296.3 | P | --- |
| 266718_at | 1147.3 | P | 937 | P | 931.3 | P | 893.5 | P | ZAT |
| 266719_at | 97.3 | P | 504 | P | 103.9 | P | 323.6 | P | CCA1 |
| 266720_s_at | 51.6 | P | 143.5 | P | 32 | P | 67.9 | P | APRR9 |
| 266721_at | 252.5 | P | 285.6 | P | 315.8 | P | 274.2 | P | FT1 |
| 266722_at | 16.7 | A | 18.2 | A | 57 | A | 13.1 | A | FUT2 |
| 266723_at | 0.8 | A | 1 | A | 1.2 | A | 1.4 | A | ASK16 |
| 266724_at | 6 | A | 13.3 | A | 15.5 | A | 6.8 | A | --- |
| 266725_at | 3.7 | A | 2.8 | A | 9.3 | A | 41.6 | A | ASK14 |
| 266726_at | 18.9 | A | 2.7 | A | 5.2 | A | 12 | A | ASK19 |
| 266727_at | 353.3 | P | 333.9 | P | 260.1 | P | 324.1 | P | emb1579 |
| 266728_at | 332.3 | P | 359.4 | P | 301.6 | P | 432.8 | P | --- |
| 266729_at | 95.8 | P | 21.4 | A | 107.3 | P | 32.4 | P | --- |
| 266730_at | 17.1 | A | 8.1 | A | 1.5 | A | 20.6 | A | --- |
| 266731_at | 65.8 | P | 72.7 | A | 80.4 | A | 78.5 | P | --- |
| 266732_at | 254.9 | P | 319.9 | P | 325.6 | P | 302.8 | P | --- |
| 266733_at | 148.6 | P | 135.3 | P | 155.4 | P | 159.7 | P | --- |
| 266734_at | 653.2 | P | 638 | P | 613.1 | P | 547.2 | P | --- |
| 266735_at | 748.9 | P | 939.2 | P | 730.1 | P | 859.4 | P | --- |
| 266736_at | 49.1 | M | 12.6 | A | 44.4 | A | 22.7 | A | CYP709B1 |
| 266737_at | 7 | A | 23.6 | P | 24.5 | A | 35.4 | A | --- |
| 266738_at | 145.2 | P | 185.7 | P | 207.5 | P | 151 | P | --- |
| 266739_at | 39.4 | A | 44.6 | A | 74.2 | A | 31.7 | A | --- |
| 266740_at | 40 | A | 23.2 | A | 32.4 | A | 37.2 | A | --- |
| 266741_s_at | 23.5 | A | 34.9 | A | 30.5 | A | 37.3 | A | --- |
| 266742_at | 207.6 | P | 265.2 | P | 334.2 | P | 266.1 | P | --- |
| 266743_at | 5646.4 | P | 1438 | P | 1738.7 | P | 1719 | P | RNS1 |
| 266744_at | 356.4 | P | 359.5 | P | 346.8 | P | 353.6 | P | --- |
| 266745_at | 317.1 | P | 302.8 | P | 372.5 | P | 300 | P | PKS1 |
| 266746_s_at | 8204.7 | P | 6650.6 | P | 6673.8 | P | 5159.9 | P | ATGSTF3 |
| 266747_at | 490.9 | P | 422.1 | P | 426.3 | P | 420.2 | P | --- |
| 266748_at | 388.3 | P | 581 | P | 587.4 | P | 463.7 | P | SPL1 |
| 266749_at | 345.2 | P | 511.3 | P | 439.7 | P | 520.8 | P | --- |
| 266750_s_at | 14.2 | A | 9.6 | A | 2.2 | A | 11.6 | A | VGDH1 |
| 266751_at | 129.9 | P | 145.9 | P | 208.7 | M | 137 | P | --- |
| 266752_at | 552 | P | 217.7 | P | 362.3 | P | 213.9 | P | ABCB4 |
| 266753_at | 4.6 | A | 4.8 | A | 33.1 | A | 4.3 | A | IAA20 |
| 266754_at | 24.7 | A | 44.6 | A | 2.7 | A | 26.9 | A | --- |
| 266755_at | 10.8 | A | 8.7 | A | 22.6 | A | 22.7 | A | --- |
| 266756_at | 16.4 | A | 23.2 | A | 54.2 | A | 29.7 | A | CYP709B2 |
| 266757_at | 8.5 | A | 17.7 | A | 7.1 | A | 26 | A | --- |
| 266758_at | 415.7 | P | 353.2 | P | 334.1 | P | 369.4 | P | POL |
| 266759_at | 48.6 | P | 32.6 | M | 89.3 | P | 34.9 | P | --- |
| 266760_at | 149.8 | P | 106.6 | P | 150.9 | P | 121 | P | NGA1 |
| 266761_at | 91 | P | 170.5 | P | 106.3 | P | 125.6 | P | --- |
| 266762_at | 3.1 | A | 2.8 | A | 2.5 | A | 1.7 | A | --- |
| 266763_at | 1554 | P | 1626.6 | P | 1654.6 | P | 1653 | P | SPL1 |
| 266764_at | 2.1 | A | 22.3 | A | 3.7 | A | 5.6 | A | --- |
| 266765_at | 20.8 | A | 4.9 | A | 1.8 | A | 19.8 | A | AtPPa3 |
| 266766_at | 12.9 | A | 15.5 | A | 44.1 | A | 17.8 | A | PAP14 |
| 266767_at | 1623 | P | 1585.2 | P | 1195.4 | P | 1223.3 | P | --- |
| 266768_s_at | 8518.8 | P | 8478.2 | P | 8715 | P | 8910 | P | UBQ6 |
| 266769_s_at | 9.7 | A | 11.4 | A | 18 | A | 10.4 | A | --- |
| 266770_at | 278.4 | P | 454 | P | 360 | P | 588.7 | P | ATEXPA15 |
| 266771_s_at | 118.2 | P | 105.5 | P | 116.2 | P | 84.3 | P | --- |
| 266772_s_at | 105.1 | P | 185 | P | 124.7 | P | 195.3 | P | --- |
| 266773_at | 14.6 | A | 4.1 | A | 24.3 | A | 12.7 | A | --- |
| 266774_at | 122.1 | P | 132.2 | P | 123.8 | P | 160.5 | P | ATRBL1 |
| 266775_at | 82.4 | P | 91.3 | P | 67.6 | A | 60.8 | P | --- |
| 266776_at | 197.1 | P | 180.2 | P | 216.1 | P | 175.5 | P | --- |
| 266777_at | 1257.1 | P | 1458.5 | P | 1268.3 | P | 1547.2 | P | ftsh3 |
| 266778_at | 41 | A | 302.6 | P | 82.6 | P | 218.1 | P | CYP707A2 |
| 266779_at | 1.5 | A | 2.7 | A | 1.8 | A | 2.5 | A | ATGLR2.9 |
| 266780_at | 42.8 | P | 45.8 | P | 39.7 | A | 39.5 | P | ATGLR2.8 |
| 266781_at | 98.6 | P | 146.5 | P | 102.5 | P | 110.9 | P | --- |
| 266782_at | 302.3 | P | 315.3 | P | 352.9 | P | 248 | P | ATGLR2.7 |
| 266783_at | 3.7 | A | 3.7 | A | 1.6 | A | 23.4 | A | LAC2 |
| 266784_at | 2.3 | A | 2.1 | A | 20.7 | A | 14.1 | A | --- |
| 266785_at | 12.1 | A | 16.9 | A | 10.8 | A | 2.6 | A | --- |
| 266786_at | 16.2 | A | 13.6 | A | 29.9 | A | 16.9 | A | --- |
| 266787_at | 4.9 | A | 10.1 | A | 1.5 | A | 5.4 | A | --- |
| 266788_at | 2.2 | A | 4.1 | A | 3.2 | A | 22.1 | A | --- |
| 266789_at | 1533.6 | P | 1928.3 | P | 1416.2 | P | 1802.5 | P | --- |
| 266790_at | 1202.1 | P | 2158.5 | P | 1445.6 | P | 1933.1 | P | ATEXPA6 |
| 266791_at | 384.7 | P | 460.9 | P | 513.5 | P | 379.4 | P | --- |
| 266792_at | 428.5 | P | 480.7 | P | 494.8 | P | 374.2 | P | SUT2 |
| 266793_at | 73.9 | P | 63.3 | A | 62.6 | A | 63.2 | P | AGL30 |
| 266794_at | 51.4 | P | 88.3 | P | 50.1 | P | 62.1 | P | --- |
| 266795_at | 220.7 | P | 259.7 | P | 284.8 | P | 256.3 | P | --- |
| 266796_at | 214.2 | P | 188.9 | P | 244.3 | P | 232.4 | P | --- |
| 266797_at | 136 | P | 170 | P | 175 | P | 180.8 | P | AtGRF1 |
| 266798_at | 46.6 | P | 38.9 | P | 26.3 | A | 47.6 | P | AtbZIP6 |
| 266799_at | 48.7 | P | 136.2 | P | 40.1 | P | 138.6 | P | ATPSK2 |
| 266800_at | 104.1 | P | 59.1 | A | 8.2 | A | 8.8 | A | --- |
| 266801_at | 616.4 | P | 811.7 | P | 834.5 | P | 839.1 | P | EMB2001 |
| 266802_at | 363.7 | P | 322.7 | P | 416.2 | P | 318.4 | P | --- |
| 266803_at | 349.4 | P | 459 | P | 387.6 | P | 477.5 | P | APK1B |
| 266804_at | 1317 | P | 1462.7 | P | 1200.7 | P | 1372.3 | P | CYP5 |
| 266805_at | 733.7 | P | 878 | P | 834.8 | P | 984 | P | --- |

| | | | | | | | | | |
|---|---|---|---|---|---|---|---|---|---|
| 266806_at | 1430.4 | P | 1432.5 | P | 1473.5 | P | 1537.2 | P | --- |
| 266807_at | 78.3 | P | 50.5 | P | 66 | P | 52.4 | A | --- |
| 266808_at | 209.8 | P | 172.6 | P | 223.5 | P | 198.1 | P | --- |
| 266809_at | 520.6 | P | 470.3 | P | 444.5 | P | 451.9 | P | --- |
| 266810_at | 13.4 | A | 18.3 | A | 8.5 | A | 19.3 | A | --- |
| 266811_at | 25.1 | A | 63.9 | A | 55.9 | A | 59.9 | A | --- |
| 266812_at | 119.9 | P | 131.8 | P | 163.4 | P | 130.4 | P | --- |
| 266813_at | 1309.6 | P | 1469.7 | P | 1292.1 | P | 1148.8 | P | --- |
| 266814_at | 34.1 | A | 19.5 | A | 13.3 | A | 9.6 | A | ATHB4 |
| 266815_at | 325.1 | P | 313.6 | P | 326.7 | P | 327.2 | P | --- |
| 266816_at | 844.8 | P | 794.9 | P | 707.2 | P | 825.1 | P | --- |
| 266817_at | 50.4 | P | 71.5 | P | 83.1 | P | 69.2 | P | HUB1 |
| 266818_at | 294.8 | P | 260.8 | P | 282.2 | P | 328.5 | P | HUB1 |
| 266819_at | 1853.5 | P | 2139.2 | P | 1985.3 | P | 2166.5 | P | --- |
| 266820_at | 189.9 | P | 271.5 | P | 211.9 | P | 256.8 | P | --- |
| 266821_at | 178.4 | P | 49.3 | P | 162.6 | P | 64.8 | P | ERF13 |
| 266822_at | 2059.4 | P | 1474.5 | P | 2377.8 | P | 1669.5 | P | --- |
| 266823_at | 22.1 | A | 62.4 | A | 74 | P | 64.9 | A | --- |
| 266824_at | 121.3 | P | 83.9 | P | 96.8 | P | 90.7 | P | HAT9 |
| 266825_at | 123.4 | P | 118.3 | P | 114.2 | P | 119.7 | P | --- |
| 266826_at | 449.4 | P | 297.2 | P | 262.1 | P | 294.5 | P | --- |
| 266827_at | 3.6 | A | 17.8 | P | 1.3 | A | 16.4 | A | SCPL12 |
| 266828_at | 210.4 | P | 43.6 | P | 247.7 | P | 44.2 | A | --- |
| 266829_at | 13.3 | A | 1.1 | A | 7.8 | A | 10.5 | A | --- |
| 266830_at | 34.6 | A | 48.7 | A | 57.2 | A | 32.1 | A | ACS4 |
| 266831_at | 103.8 | P | 141.2 | P | 99.4 | P | 107.6 | P | SQE2 |
| 266832_at | 164.1 | P | 72.3 | P | 90 | P | 60.8 | P | MAPKKK14 |
| 266833_at | 1.5 | A | 10.7 | A | 13.5 | A | 6.6 | A | --- |
| 266834_s_at | 168 | P | 151.1 | P | 120.5 | P | 165.1 | P | --- |
| 266835_at | 582.9 | P | 617.8 | P | 489.9 | P | 482.5 | P | NDA2 |
| 266836_at | 73.4 | P | 71 | P | 61.4 | P | 62.1 | P | --- |
| 266837_x_at | 32.5 | A | 2.2 | A | 15.3 | A | 18.8 | A | --- |
| 266838_at | 6.4 | A | 32 | A | 7.5 | A | 17.3 | A | --- |
| 266839_at | 607.2 | P | 545.1 | P | 676.9 | P | 602.7 | P | ELF3 |
| 266840_at | 1.4 | A | 1.6 | A | 1.3 | A | 0.7 | A | --- |
| 266841_at | 41.5 | P | 121.8 | P | 57 | P | 81.2 | P | ATHSFA2 |
| 266842_at | 785.8 | P | 861.6 | P | 707.3 | P | 799 | P | ftsh4 |
| 266843_at | 20.9 | P | 18.2 | A | 18.1 | A | 34.5 | P | --- |
| 266844_at | 3.4 | A | 14.7 | A | 6.5 | A | 17.1 | A | --- |
| 266845_at | 198.1 | P | 204.3 | P | 295.4 | P | 239.5 | P | --- |
| 266846_at | 1302 | P | 1094.8 | P | 1422.4 | P | 1115.2 | P | --- |
| 266847_at | 730.1 | P | 737.4 | P | 658 | P | 684.8 | P | emb1345 |
| 266848_at | 884.9 | P | 853.2 | P | 921.9 | P | 969.4 | P | --- |
| 266849_at | 2.2 | A | 15.8 | A | 4.5 | A | 4.1 | A | ALPHA-VPE |
| 266850_at | 16.1 | P | 38.5 | A | 66 | P | 55.7 | A | --- |
| 266851_at | 9.1 | A | 17.3 | A | 19.5 | A | 13.4 | A | ATPP2-A3 |
| 266852_at | 252.9 | P | 172.7 | P | 156.2 | P | 228.3 | P | --- |
| 266853_at | 35.4 | M | 16.1 | A | 30.7 | A | 15.3 | A | --- |
| 266854_at | 14 | A | 33 | A | 29 | A | 25.9 | A | --- |
| 266855_at | 476.7 | P | 402.4 | P | 396.4 | P | 428.3 | P | --- |
| 266856_at | 926.7 | P | 1086.3 | P | 1060 | P | 1071.4 | P | PDR4 |
| 266857_at | 4568.1 | P | 5422.2 | P | 5480.8 | P | 5621.9 | P | --- |
| 266858_at | 417.2 | P | 351.3 | P | 390.1 | P | 270.1 | P | GRV2 |
| 266859_at | 1.1 | A | 1.2 | A | 1.8 | A | 2.3 | A | AGL41 |
| 266860_at | 319.3 | P | 306.4 | P | 325.7 | P | 264.2 | P | --- |
| 266861_at | 421.5 | P | 468.7 | P | 332 | P | 449.7 | P | emb1187 |
| 266862_at | 8.5 | A | 15 | A | 27.7 | A | 10.6 | A | AtMYB104 |
| 266863_at | 640 | P | 865.9 | P | 723.3 | P | 787.2 | P | ATCDPMEK |
| 266864_at | 939.3 | P | 822.3 | P | 849.8 | P | 832.3 | P | --- |
| 266865_at | 3516.5 | P | 3194.8 | P | 4890.8 | P | 3524.8 | P | FAD3 |
| 266866_at | 65.1 | P | 60.6 | P | 88.3 | P | 51.3 | P | PDR3 |
| 266867_at | 420.8 | P | 609.7 | P | 538.9 | P | 502 | P | CPFTSY |
| 266868_at | 50.5 | P | 51 | P | 83.8 | P | 53.8 | P | --- |
| 266869_at | 227.6 | P | 323.3 | P | 323.8 | P | 326.2 | P | --- |
| 266870_at | 70 | A | 36.8 | A | 62.9 | A | 63.8 | A | --- |
| 266871_at | 349.8 | P | 321.1 | P | 303.3 | P | 357.3 | P | --- |
| 266872_at | 165.6 | P | 182.7 | P | 134.3 | P | 142.4 | P | --- |
| 266873_at | 42.9 | A | 129 | P | 85.4 | P | 83.8 | P | CYCP4;1 |
| 266874_at | 823 | P | 897.6 | P | 752.4 | P | 720.2 | P | --- |
| 266875_at | 0.8 | A | 16.6 | A | 6 | A | 13.3 | A | --- |
| 266876_at | 744.5 | P | 563.5 | P | 626.5 | P | 495.8 | P | --- |
| 266877_at | 41.1 | P | 13.5 | A | 57.9 | A | 22.2 | P | AtGH9B12 |
| 266878_at | 22.7 | A | 33.9 | A | 17.3 | A | 35.6 | A | --- |
| 266879_at | 3.1 | A | 4.4 | A | 19.8 | A | 14.6 | A | ADL1D |
| 266880_at | 428.5 | P | 661.4 | P | 543.2 | P | 623.9 | P | --- |
| 266881_at | 524.2 | P | 613.3 | P | 566 | P | 463.3 | P | CKB4 |
| 266882_at | 1558.6 | P | 2032.2 | P | 2073 | P | 1698.2 | P | --- |
| 266883_at | 31 | A | 3.9 | A | 29.2 | A | 10.4 | A | DAD1 |
| 266884_at | 85.7 | P | 196.2 | P | 163.2 | P | 206.3 | P | UCC2 |
| 266885_at | 1413.5 | P | 1663 | P | 1198 | P | 1652.7 | P | RAB6A |
| 266886_at | 3.5 | A | 28 | P | 66.9 | A | 8.1 | A | --- |
| 266887_at | 4958.4 | P | 5000.3 | P | 4444.8 | P | 3799.2 | P | CHL-CPN10 |
| 266888_s_at | 532.9 | P | 391.3 | P | 546.4 | P | 416.5 | P | TPK1 |
| 266889_at | 1217.3 | P | 1307.1 | P | 1485.7 | P | 1200.5 | P | --- |
| 266890_at | 2480.6 | P | 2509.8 | P | 1960.5 | P | 2347.4 | P | MTACP-1 |
| 266891_at | 64.3 | P | 89.9 | P | 99.3 | P | 83.4 | P | ARAC9 |
| 266892_at | 2803.6 | P | 3338.8 | P | 3553.1 | P | 3615.5 | P | AtGLDP2 |
| 266893_at | 284.1 | P | 426.6 | P | 411.5 | P | 383.8 | P | RTE1 |
| 266894_at | 24.4 | A | 24.3 | A | 47.9 | A | 31.7 | A | --- |
| 266895_at | 75.6 | A | 103.2 | A | 53.4 | A | 47.5 | A | --- |
| 266896_at | 873.5 | P | 1214.9 | P | 953.1 | P | 1010.2 | P | --- |
| 266897_at | 2114.4 | P | 2880.9 | P | 1996.1 | P | 2920.3 | P | --- |
| 266898_at | 1300.6 | P | 1371.5 | P | 1369.6 | P | 1567.4 | P | --- |
| 266899_at | 326.6 | P | 394.1 | P | 349.1 | P | 390.1 | P | --- |
| 266900_at | 5.8 | A | 2.8 | A | 4.6 | A | 15.1 | A | --- |
| 266901_at | 139 | P | 23.9 | A | 55.1 | P | 36.2 | P | JAZ7 |
| 266902_at | 2.2 | A | 2.2 | A | 1.6 | A | 2.2 | A | --- |
| 266903_at | 198.2 | P | 204.5 | P | 201.9 | P | 186.8 | P | MEE21 |
| 266904_at | 2089.4 | P | 1723.2 | P | 2044.7 | P | 1661 | P | --- |
| 266905_at | 1521.7 | P | 1117.8 | P | 1245.9 | P | 1228.8 | P | --- |
| 266906_at | 1597.1 | P | 1489.7 | P | 1572.2 | P | 1564.7 | P | --- |
| 266907_at | 42.3 | A | 81.9 | A | 74.2 | P | 66.7 | A | ATGA2OX3 |
| 266908_at | 114.6 | M | 128.6 | P | 90.6 | A | 112.7 | A | PID |
| 266909_at | 35.9 | A | 40.2 | P | 63.9 | A | 5.4 | A | BRM |
| 266910_at | 177.4 | P | 144.5 | P | 176.6 | P | 137.2 | P | --- |
| 266911_at | 170.9 | P | 190 | P | 134.2 | P | 198.2 | P | --- |
| 266912_at | 85.2 | P | 79.9 | P | 117.7 | P | 88.5 | P | --- |
| 266913_at | 28.6 | A | 41.1 | A | 7.4 | A | 52.5 | A | ROPGEF4 |
| 266914_at | 104.5 | P | 107.5 | P | 72.2 | P | 100.9 | P | BAM7 |

| | | | | | | | | | |
|---|---|---|---|---|---|---|---|---|---|
| 266915_at | 87.7 | P | 94.1 | P | 155.5 | P | 101.8 | P | --- |
| 266916_at | 438.6 | P | 379.1 | P | 503.6 | P | 376.5 | P | --- |
| 266917_at | 34.2 | M | 25.6 | M | 46.9 | P | 46 | P | DTA2 |
| 266918_at | 3.9 | A | 2.8 | A | 6.7 | A | 10.9 | A | --- |
| 266919_at | 20.1 | A | 29.8 | A | 12.9 | A | 2.2 | A | --- |
| 266920_at | 59.6 | P | 68.3 | A | 103.5 | M | 72.7 | P | --- |
| 266921_at | 261 | P | 383.4 | P | 369 | P | 411.4 | P | CYP86A8 |
| 266922_s_at | 476.4 | P | 564.6 | P | 498.6 | P | 566.2 | P | ASK20 |
| 266923_at | 1106 | P | 1116.5 | P | 905.5 | P | 1052.8 | P | --- |
| 266924_at | 512.6 | P | 508.4 | P | 648.6 | P | 568.3 | P | --- |
| 266925_at | 3817.5 | P | 3217.6 | P | 4215.3 | P | 3119.9 | P | PEX11D |
| 266926_at | 175.4 | P | 222.7 | P | 143.6 | P | 203.5 | P | --- |
| 266927_at | 5509.6 | P | 6784.4 | P | 7124 | P | 6853.4 | P | PIP1B |
| 266928_at | 3320.4 | P | 3342 | P | 2792.1 | P | 3246.6 | P | PMM |
| 266929_at | 308.3 | P | 293.7 | P | 292.8 | P | 322.9 | P | --- |
| 266930_at | 9.1 | A | 23.1 | A | 29 | A | 18.6 | A | --- |
| 266931_at | 3.8 | A | 2.1 | A | 3.5 | A | 3.1 | A | --- |
| 266932_s_at | 147.7 | P | 105.6 | P | 119.6 | P | 124.3 | P | --- |
| 266933_at | 28.3 | A | 22.9 | A | 9.6 | A | 10.4 | A | --- |
| 266934_at | 352.7 | P | 333.9 | P | 334.7 | P | 283.7 | P | --- |
| 266935_at | 289.1 | P | 289.2 | P | 231.7 | P | 278.2 | P | LKP2 |
| 266936_at | 2.3 | A | 26 | A | 28.7 | A | 29.3 | A | --- |
| 266937_at | 2.8 | A | 1.3 | A | 3.2 | A | 0.8 | A | --- |
| 266938_at | 713.8 | P | 893.8 | P | 857.8 | P | 839.2 | P | HPT1 |
| 266939_at | 12127.9 | P | 9656.8 | P | 13377.8 | P | 10571.8 | P | AHA1 |
| 266940_at | 3.5 | A | 16.7 | A | 6 | A | 4.7 | A | --- |
| 266941_at | 53.4 | M | 48.1 | P | 35.8 | A | 73.9 | P | --- |
| 266942_at | 386 | P | 478.2 | P | 483 | P | 490.9 | P | TXND9 |
| 266943_at | 46.6 | A | 40.3 | M | 59.3 | P | 52.6 | A | --- |
| 266944_at | 99 | P | 48.4 | P | 67.4 | P | 63.5 | P | --- |
| 266945_at | 1.8 | A | 1.1 | A | 4.3 | A | 1.5 | A | VEL2 |
| 266946_at | 98.2 | P | 135.7 | P | 117.6 | P | 132.1 | P | --- |
| 266947_at | 4.4 | A | 17.1 | A | 6.1 | A | 2.7 | A | --- |
| 266948_at | 221 | P | 144.7 | P | 244.6 | P | 162.5 | P | --- |
| 266949_s_at | 614.2 | P | 677.8 | P | 507.3 | P | 616.2 | P | --- |
| 266950_at | 459.3 | P | 774.6 | P | 557.8 | P | 792.1 | P | --- |
| 266951_at | 504.7 | P | 521.8 | P | 410.4 | P | 489.4 | P | --- |
| 266952_at | 14.2 | A | 41.4 | P | 57.5 | A | 25.9 | A | ATGA2OX3 |
| 266953_at | 29.6 | A | 39.6 | A | 37.8 | A | 41.1 | A | --- |
| 266954_at | 21.4 | A | 29.2 | A | 50.3 | A | 37.8 | M | --- |
| 266955_at | 1214.1 | P | 1005.4 | P | 1281.9 | P | 1060.4 | P | RPS14 |
| 266956_at | 568.6 | P | 581.8 | P | 625.5 | P | 632.5 | P | --- |
| 266957_at | 770.8 | P | 1013.8 | P | 992 | P | 935.3 | P | PTAC12 |
| 266958_at | 923.3 | P | 821.1 | P | 862.1 | P | 795.5 | P | GPS1 |
| 266959_at | 203.4 | P | 366.3 | P | 322.1 | P | 299.4 | P | MCM5 |
| 266960_at | 17.6 | A | 10.6 | A | 7.5 | A | 38.5 | A | --- |
| 266961_at | 40.7 | A | 14.6 | A | 46.6 | A | 6.3 | A | --- |
| 266962_at | 64.5 | P | 85.2 | P | 68.1 | P | 69.4 | P | --- |
| 266963_at | 2444.9 | P | 1193.9 | P | 1562.1 | P | 1359.1 | P | MTP11 |
| 266964_at | 297.3 | P | 308.9 | P | 256.9 | P | 336 | P | PGP6 |
| 266965_at | 82.7 | P | 227.7 | P | 273.5 | P | 309.9 | P | --- |
| 266966_at | 17.1 | A | 0.3 | A | 46.2 | A | 17.8 | A | --- |
| 266967_at | 46.3 | A | 36.7 | A | 18.1 | A | 16.5 | A | --- |
| 266968_at | 273.6 | P | 263.8 | P | 204.3 | P | 194.9 | P | --- |
| 266969_at | 34.2 | P | 39.2 | A | 21.8 | A | 62.9 | A | --- |
| 266970_at | 104.2 | P | 42.9 | M | 157.8 | P | 111.5 | P | --- |
| 266971_at | 309.9 | P | 297.9 | P | 244.3 | P | 286.1 | P | --- |
| 266972_at | 12.1 | A | 10.3 | A | 1.6 | A | 15.7 | A | --- |
| 266973_at | 85.7 | P | 62.6 | P | 66.5 | P | 67.6 | P | --- |
| 266974_at | 2.8 | A | 4.6 | A | 2.4 | A | 7.5 | A | --- |
| 266975_at | 55.5 | P | 56.1 | P | 78 | P | 41.2 | P | ATEXO70H2 |
| 266976_at | 4.9 | A | 20.3 | A | 55.8 | A | 5 | A | --- |
| 266977_at | 920.9 | P | 393.6 | P | 936.4 | P | 378 | P | --- |
| 266978_at | 17.5 | A | 30.6 | A | 64.6 | P | 34.3 | P | --- |
| 266979_at | 1790.3 | P | 2064.5 | P | 2183.7 | P | 2012.8 | P | PPL2 |
| 266980_at | 2246.9 | P | 1707 | P | 2298.8 | P | 1689.1 | P | --- |
| 266981_at | 7310.2 | P | 6875.3 | P | 9412.7 | P | 6503.7 | P | RPL23AA |
| 266982_at | 132.7 | P | 241.2 | P | 216.4 | P | 191.1 | P | --- |
| 266983_at | 273.8 | P | 319 | P | 151.5 | P | 271.9 | P | --- |
| 266984_at | 3053.1 | P | 1185.3 | P | 3689.4 | P | 1402.5 | P | --- |
| 266985_at | 340.3 | P | 369.4 | P | 467.1 | P | 341.3 | P | PGGT-I |
| 266986_at | 528.8 | P | 509.8 | P | 548.8 | P | 449.9 | P | --- |
| 266987_at | 147.3 | P | 137.6 | P | 115.6 | A | 187.6 | P | --- |
| 266988_at | 4275.4 | P | 1748.2 | P | 4029.5 | P | 2127.8 | P | JAL22 |
| 266989_at | 3346 | P | 454.1 | P | 2690.6 | P | 451.8 | P | JAL23 |
| 266990_at | 559 | P | 721.9 | P | 534.1 | P | 719.4 | P | ATATH8 |
| 266991_at | 542 | P | 784.7 | P | 661 | P | 729.6 | P | PGP1 |
| 266992_at | 31.4 | A | 37.2 | P | 30.9 | A | 28.1 | P | MLO12 |
| 266993_at | 262.4 | P | 795.8 | P | 212.8 | P | 450 | P | --- |
| 266994_at | 28.5 | A | 16.6 | A | 4.8 | A | 32.4 | A | AGL29 |
| 266995_at | 180.7 | P | 82.7 | M | 126.1 | P | 84.9 | A | CYP710A1 |
| 266996_at | 1170.9 | P | 839.9 | P | 1259.5 | P | 1068 | P | CYP710A2 |
| 266997_at | 129.4 | P | 183.3 | P | 144.5 | P | 191 | P | --- |
| 266998_at | 61.6 | A | 88.4 | P | 86.2 | A | 106.2 | A | --- |
| 266999_at | 116.4 | P | 73.4 | P | 155.8 | P | 89.1 | P | --- |
| 267000_at | 1114.7 | P | 913 | P | 1251 | P | 997.7 | P | --- |
| 267001_at | 1594.9 | P | 2126.3 | P | 1790.1 | P | 2106.3 | P | UREG |
| 267002_s_at | 17567.1 | P | 17025.6 | P | 16224 | P | 15969.2 | P | LHB1B1 |
| 267003_at | 109.4 | P | 80.7 | P | 109.1 | P | 83.1 | P | --- |
| 267004_at | 635.1 | P | 477.5 | P | 756.6 | P | 553.8 | P | --- |
| 267005_at | 1369.9 | P | 1507.9 | P | 1533.2 | P | 1531.7 | P | --- |
| 267006_at | 66.1 | P | 142.5 | P | 131.3 | P | 169.9 | P | --- |
| 267007_at | 10264.7 | P | 9288.8 | P | 10664.6 | P | 10056.8 | P | --- |
| 267008_at | 155.1 | P | 203.1 | P | 77.7 | P | 138.8 | P | --- |
| 267009_at | 223.4 | P | 241.6 | P | 255.1 | P | 229.7 | P | --- |
| 267010_at | 11.5 | A | 36.1 | A | 14.5 | A | 44.3 | A | SNZ |
| 267011_at | 48.5 | P | 54.7 | M | 80.8 | P | 79 | P | --- |
| 267012_at | 435 | P | 435.8 | P | 419.6 | P | 441.8 | P | PLP6 |
| 267013_at | 63.8 | P | 71.1 | P | 96 | P | 92.6 | P | CCR2 |
| 267014_at | 489.7 | P | 387.5 | P | 490.1 | P | 439 | P | --- |
| 267015_at | 384.6 | P | 402.8 | P | 515.2 | P | 449.4 | P | --- |
| 267016_at | 52.2 | A | 37.3 | A | 86.1 | M | 24.8 | M | --- |
| 267017_at | 340.2 | P | 253.2 | P | 282.4 | P | 252.2 | P | SVR1 |
| 267018_at | 552.6 | P | 770.4 | P | 788.3 | P | 756.8 | P | SVR1 |
| 267019_at | 491.8 | P | 500.4 | P | 549.6 | P | 593.4 | P | --- |
| 267020_at | 32.7 | P | 23.5 | P | 58.3 | P | 26.1 | P | --- |
| 267021_at | 32.9 | A | 33.4 | P | 63.7 | P | 45.3 | P | --- |
| 267022_at | 31.1 | A | 3 | A | 21.8 | A | 17 | A | --- |
| 267023_at | 3161.2 | P | 3156.7 | P | 3275.6 | P | 3393.8 | P | --- |

| | | | | | | | | | |
|---|---|---|---|---|---|---|---|---|---|
| 267024_s_at | 36.9 | P | 31.8 | A | 45.1 | A | 40.5 | A | NIP2;1 |
| 267025_at | 2.7 | A | 9.4 | A | 11.3 | A | 1.7 | A | --- |
| 267026_at | 64.9 | A | 23.7 | A | 32.9 | A | 15 | A | --- |
| 267027_at | 621.3 | P | 679.1 | P | 574.7 | P | 638.3 | P | --- |
| 267028_at | 560.8 | P | 535.7 | P | 356.8 | P | 418.8 | P | WRKY33 |
| 267029_at | 279 | P | 315.6 | P | 355.6 | P | 327.1 | P | ATIREG1 |
| 267030_at | 326.1 | P | 473.2 | P | 427.9 | P | 548 | P | SCAR2 |
| 267031_at | 115.7 | P | 111.6 | P | 111 | P | 94.4 | P | --- |
| 267032_at | 19.3 | A | 4.8 | A | 10.3 | A | 2.7 | A | CIPK22 |
| 267033_at | 1037.4 | P | 1234.1 | P | 1020.3 | P | 1168.5 | P | --- |
| 267034_at | 1825.7 | P | 3413.3 | P | 2577.6 | P | 2725.6 | P | --- |
| 267035_at | 1283 | P | 769.5 | P | 944.3 | P | 871 | P | AGT3 |
| 267036_at | 480.7 | P | 381.5 | P | 365.9 | P | 450.2 | P | --- |
| 267037_at | 47 | P | 74.5 | P | 94 | P | 79.2 | P | --- |
| 267038_at | 84.4 | P | 105.6 | P | 89.5 | P | 119.8 | P | --- |
| 267039_at | 21.4 | A | 15.6 | A | 44.9 | A | 17.1 | A | --- |
| 267040_at | 1224.4 | P | 1305.2 | P | 1104 | P | 1139.4 | P | --- |
| 267041_at | 15.2 | A | 19.7 | A | 4.1 | A | 8.8 | A | --- |
| 267042_at | 26.5 | A | 29 | A | 7.3 | A | 7.5 | A | --- |
| 267043_at | 20.7 | A | 25.9 | A | 24.1 | A | 12.4 | A | --- |
| 267044_at | 519 | P | 334.9 | P | 480.6 | P | 446 | P | --- |
| 267045_at | 2.9 | A | 8.2 | A | 4.6 | A | 15 | A | CIPK13 |
| 267046_at | 30.1 | A | 24.5 | A | 63.8 | A | 16.4 | A | --- |
| 267047_at | 4 | A | 6 | A | 15.6 | A | 3.9 | A | --- |
| 267048_at | 208.9 | P | 185.4 | P | 154.2 | P | 178.6 | P | --- |
| 267049_at | 3.1 | A | 5.2 | A | 5.1 | A | 2.4 | A | --- |
| 267050_at | 186.5 | P | 206.9 | P | 249.4 | P | 218.8 | P | --- |
| 267051_at | 66.8 | A | 6.2 | A | 34 | A | 8.6 | A | --- |
| 267052_at | 241.4 | P | 268 | P | 213 | P | 207.4 | P | --- |
| 267053_s_at | 64.2 | P | 403.5 | P | 290.9 | P | 428.2 | P | --- |
| 267054_at | 152.9 | P | 120.2 | P | 126.2 | P | 111 | P | --- |
| 267055_at | 420.3 | P | 469.9 | P | 357.5 | P | 577.3 | P | PRA1.B4 |
| 267056_at | 25.1 | A | 12.7 | A | 4 | A | 19.1 | A | --- |
| 267057_at | 368.4 | P | 584.4 | P | 451.1 | P | 602.5 | P | --- |
| 267058_at | 26.9 | A | 45.7 | A | 34.4 | A | 29.4 | A | MAPKKK17 |
| 267059_at | 1534.2 | P | 1902.2 | P | 1561.5 | P | 1944.7 | P | --- |
| 267060_at | 1024.4 | P | 1102.5 | P | 767 | P | 1134.2 | P | --- |
| 267061_at | 1507.7 | P | 2000.7 | P | 1823.5 | P | 1894.8 | P | --- |
| 267062_at | 579.5 | P | 460.8 | P | 510 | P | 523.7 | P | --- |
| 267063_at | 441.8 | P | 445.4 | P | 453.7 | P | 383.9 | P | --- |
| 267064_at | 1112.2 | P | 1387.9 | P | 869.4 | P | 1140 | P | CAM2 |
| 267065_at | 132.2 | P | 187.4 | P | 168.3 | P | 145.4 | P | --- |
| 267066_at | 377.3 | P | 312.4 | P | 330.1 | P | 375.1 | P | --- |
| 267067_at | 132.7 | P | 134.7 | P | 181.1 | A | 106.3 | P | --- |
| 267068_at | 141.3 | P | 161.2 | P | 145.9 | P | 128.4 | P | --- |
| 267069_at | 226.5 | P | 411.7 | P | 237.8 | P | 398.6 | P | ATCAMBP25 |
| 267070_at | 223.3 | P | 224 | P | 243.1 | P | 213.5 | P | --- |
| 267071_at | 265.3 | P | 280.6 | P | 303.1 | P | 296.4 | P | --- |
| 267072_at | 103 | P | 113.9 | P | 120.5 | P | 137.5 | P | --- |
| 267073_at | 325.7 | P | 491.4 | P | 330.2 | P | 419.7 | P | --- |
| 267074_s_at | 16.6 | A | 2.2 | A | 4.5 | A | 2.8 | A | --- |
| 267075_at | 138.8 | P | 82.8 | P | 105.3 | P | 80.7 | P | EEL |
| 267076_at | 3473.2 | P | 7398.2 | P | 3337.9 | P | 7401.6 | P | --- |
| 267077_at | 216.6 | P | 256 | P | 148 | P | 173.4 | P | --- |
| 267078_at | 177.5 | P | 161.4 | P | 165.8 | P | 150.8 | P | --- |
| 267079_at | 85 | P | 112.4 | P | 117.4 | P | 144.7 | P | --- |
| 267080_at | 130.6 | P | 122.1 | P | 132.3 | P | 124 | P | --- |
| 267081_at | 144.8 | P | 96.4 | A | 113.1 | P | 105.5 | P | PIP5K5 |
| 267082_at | 566.3 | P | 542.2 | P | 599.1 | P | 635.9 | P | CRK1 |
| 267083_at | 5560.2 | P | 6153 | P | 5038.3 | P | 4409.2 | P | TCH3 |
| 267084_at | 546.4 | P | 736.9 | P | 511 | P | 752 | P | --- |
| 267085_at | 21.5 | A | 19.6 | A | 39.2 | A | 19.7 | A | ATCSLB02 |
| 267086_at | 29.4 | A | 20.4 | A | 45.2 | A | 23.7 | A | --- |
| 267087_at | 22.6 | A | 39.1 | A | 49.8 | A | 9.2 | A | MYB101 |
| 267088_at | 3704.2 | P | 4330.3 | P | 3524.7 | P | 4452.7 | P | PSRP4 |
| 267089_at | 110.6 | P | 86.6 | P | 92.5 | P | 83 | P | --- |
| 267090_at | 1659.1 | P | 1966.4 | P | 1647 | P | 2086.9 | P | CXIP2 |
| 267091_at | 78 | P | 69.5 | P | 90.1 | A | 93.9 | P | --- |
| 267092_at | 379.7 | P | 648.9 | P | 580.6 | P | 722.8 | P | AUX1 |
| 267093_at | 5513.6 | P | 5128.6 | P | 5134.7 | P | 4481.7 | P | CAX1 |
| 267094_at | 150.8 | P | 186.2 | P | 247.8 | P | 308.7 | P | IRX12 |
| 267095_at | 1392.5 | P | 1326.2 | P | 1624.2 | P | 1317 | P | FAC1 |
| 267096_at | 261.5 | P | 302 | P | 276.4 | P | 269.1 | P | --- |
| 267097_at | 22.2 | A | 2 | A | 31.9 | A | 52 | A | --- |
| 267098_at | 2 | A | 1.2 | A | 1.9 | A | 1.4 | A | --- |
| 267099_at | 227.1 | P | 174 | P | 279.9 | P | 223.2 | P | ARP |
| 267100_at | 8.3 | A | 2.7 | A | 36.9 | A | 24.7 | A | --- |
| 267101_at | 239.3 | P | 101.1 | A | 137.8 | P | 94 | P | --- |
| 267102_at | 475.1 | P | 396.5 | P | 523 | P | 409.3 | P | LIS |
| 267103_at | 379.3 | P | 418.7 | P | 434.3 | P | 349.1 | P | GPT |
| 267104_at | 6429.4 | P | 8071.4 | P | 7313.6 | P | 7581.6 | P | ERD15 |
| 267105_at | 21.1 | A | 9.7 | A | 7.6 | A | 4.4 | A | --- |
| 267106_s_at | 1475.5 | P | 1947.7 | P | 1444.8 | P | 2234 | P | ATVSR3 |
| 267107_at | 2.5 | A | 2.8 | A | 2.1 | A | 1.7 | A | --- |
| 267108_at | 6.4 | A | 29.1 | A | 1.8 | A | 4.1 | A | --- |
| 267109_at | 1 | A | 2 | A | 0.9 | A | 2.7 | A | --- |
| 267110_at | 138.9 | P | 124.7 | P | 137.7 | P | 139.1 | P | --- |
| 267111_s_at | 34.5 | A | 29.5 | P | 52.8 | A | 38.9 | P | --- |
| 267112_at | 3200.6 | P | 2185.1 | P | 2867 | P | 2173.1 | P | APK |
| 267113_at | 147.9 | P | 127.9 | P | 122.3 | P | 129.3 | P | --- |
| 267114_at | 568 | P | 580.3 | P | 550 | P | 588.6 | P | --- |
| 267115_s_at | 498.1 | P | 720.7 | P | 485.8 | P | 483.2 | P | ATCSLB04 |
| 267116_at | 672.2 | P | 556.2 | P | 858.4 | P | 509.2 | P | --- |
| 267117_at | 23.7 | P | 10.2 | P | 30.7 | P | 16.9 | P | --- |
| 267118_at | 66.5 | P | 110.3 | P | 60.9 | A | 110.1 | P | --- |
| 267119_at | 17.2 | A | 32.3 | A | 9.6 | A | 2.7 | A | ATCSLB01 |
| 267120_at | 3.9 | A | 68.1 | P | 52.8 | A | 33.5 | A | --- |
| 267121_at | 4.7 | A | 125.9 | P | 105.4 | P | 114.1 | P | --- |
| 267122_at | 23.4 | A | 5.9 | A | 29.8 | M | 22.3 | A | MES6 |
| 267123_at | 134.6 | P | 55.3 | A | 168.1 | P | 101.9 | P | MES7 |
| 267124_at | 36 | A | 31.5 | A | 59.9 | A | 37.9 | A | MES19 |
| 267125_at | 13.3 | A | 22.1 | A | 6.4 | A | 20.7 | A | MES4 |
| 267126_s_at | 2578.7 | P | 4497.7 | P | 4336.1 | P | 3852.8 | P | ACL |
| 267127_at | 1112.1 | P | 1216.5 | P | 1193.8 | P | 1218.4 | P | MES3 |
| 267128_at | 99.6 | P | 51.4 | P | 104.6 | P | 75.3 | P | MES1 |
| 267129_at | 72.4 | P | 87.2 | P | 91.8 | A | 95.2 | P | CLF |
| 267130_at | 944.1 | P | 1162.3 | P | 909.4 | P | 1196.9 | P | --- |
| 267131_at | 0.7 | A | 0.4 | A | 0.9 | A | 1 | A | --- |
| 267132_at | 848.4 | P | 772.2 | P | 724.6 | P | 852 | P | NAPRT2 |

| | | | | | | | | | |
|---|---|---|---|---|---|---|---|---|---|
| 267133_at | 1 | A | 3.7 | A | 6.6 | A | 1.6 | A | --- |
| 267134_at | 465.7 | P | 511.4 | P | 395.4 | P | 384.4 | P | --- |
| 267135_at | 276.8 | P | 211.1 | P | 223.4 | P | 218.8 | P | ICK1 |
| 267136_at | 613.9 | P | 699.1 | P | 694.6 | P | 726.4 | P | XLG1 |
| 267137_at | 2.4 | A | 17.3 | A | 4.8 | A | 13.8 | A | ACPT |
| 267138_s_at | 3574.8 | P | 2772.5 | P | 4713.5 | P | 3316.3 | P | PDX1.1 |
| 267139_s_at | 10.6 | A | 25.4 | A | 18.3 | A | 16.1 | A | --- |
| 267140_at | 135.5 | P | 41.6 | A | 76.9 | P | 21.7 | A | --- |
| 267141_at | 75.5 | P | 105.1 | P | 79.9 | P | 112.5 | P | --- |
| 267142_at | 835.9 | P | 1199.5 | P | 728.2 | P | 906.7 | P | ATAMT2 |
| 267143_at | 22.4 | A | 43.4 | A | 42.6 | A | 15.5 | A | --- |
| 267144_at | 122.8 | P | 167.6 | P | 165 | P | 196.7 | P | GPAT6 |
| 267145_at | 545.1 | P | 556.6 | P | 436.9 | P | 502.9 | P | ATMAK3 |
| 267146_at | 35.7 | A | 53.9 | A | 33.8 | A | 35 | A | --- |
| 267147_at | 262.8 | P | 28.6 | A | 173.6 | M | 37.6 | A | --- |
| 267148_at | 96.7 | P | 131.7 | P | 168 | P | 100.3 | P | --- |
| 267149_s_at | 7.7 | A | 1.2 | A | 11.7 | A | 9 | A | --- |
| 267150_at | 1.5 | A | 2.8 | A | 4.3 | A | 0.7 | A | SDT |
| 267151_at | 3837.3 | P | 2895.1 | P | 3732.5 | P | 2992.6 | P | ASP1 |
| 267152_at | 3871 | P | 4082.7 | P | 3619.7 | P | 4275.4 | P | --- |
| 267153_at | 23050.9 | P | 18964.4 | P | 24322 | P | 20397.3 | P | ATGSTF9 |
| 267154_at | 14353.9 | P | 11835.5 | P | 13245.7 | P | 10975.4 | P | GSTF10 |
| 267155_at | 2.8 | A | 1 | A | 4.6 | A | 7 | A | --- |
| 267156_at | 16 | A | 4.7 | A | 20 | A | 15.8 | A | --- |
| 267157_at | 1049.9 | P | 1370.5 | P | 1444.6 | P | 1634.2 | P | AS1 |
| 267158_at | 372.2 | P | 1180.5 | P | 350.3 | P | 1109.2 | P | EXP3 |
| 267159_at | 128.2 | P | 118.6 | P | 109.6 | P | 143.6 | P | --- |
| 267160_at | 18.4 | A | 23.8 | A | 4.2 | A | 32.2 | A | --- |
| 267161_at | 317.3 | P | 340.2 | P | 255.7 | P | 363.5 | P | --- |
| 267162_s_at | 914.7 | P | 468.7 | P | 1249.9 | P | 508.7 | P | --- |
| 267163_at | 323.7 | P | 313.6 | P | 317.6 | P | 304.4 | P | --- |
| 267164_at | 3.3 | A | 1.5 | A | 9.6 | A | 2.9 | A | --- |
| 267165_at | 1000.7 | P | 1323.8 | P | 740.7 | P | 1170.3 | P | RLK |
| 267166_at | 39.3 | A | 41.8 | A | 37.8 | A | 33.3 | A | --- |
| 267167_at | 29.6 | M | 18.9 | A | 8.5 | A | 27.1 | A | ZFP10 |
| 267168_at | 6124.2 | P | 2348.5 | P | 4885.9 | P | 3578.9 | P | --- |
| 267169_at | 682.3 | P | 596.4 | P | 716.6 | P | 631.8 | P | --- |
| 267170_at | 158.1 | P | 162.3 | P | 151.2 | P | 145.9 | P | --- |
| 267171_at | 63.1 | P | 106.5 | P | 101.5 | P | 78.9 | P | DOF2.4 |
| 267172_at | 4704.9 | P | 6110.3 | P | 4837.4 | P | 5913 | P | --- |
| 267173_at | 68.1 | P | 121.5 | P | 92.3 | P | 137.9 | P | ORC2 |
| 267174_at | 881 | P | 706.5 | P | 1100 | P | 891.4 | P | --- |
| 267175_s_at | 830.3 | P | 852 | P | 783.6 | P | 899.8 | P | ACT1 |
| 267176_at | 26.6 | A | 4.3 | A | 15.8 | A | 6.5 | A | --- |
| 267177_at | 242.9 | P | 144 | P | 298.7 | P | 210.1 | P | --- |
| 267178_at | 183.5 | P | 115.1 | P | 91.4 | P | 89.8 | P | --- |
| 267179_at | 681.7 | P | 555.3 | P | 474.6 | P | 587.7 | P | AGD7 |
| 267180_at | 1060.2 | P | 684.7 | P | 969.3 | P | 801.1 | P | SLT1 |
| 267181_at | 5560.5 | P | 2521.8 | P | 5080.1 | P | 3128.9 | P | --- |
| 267182_at | 101.1 | P | 126.4 | P | 149 | P | 137.5 | P | --- |
| 267183_at | 241.9 | P | 203 | P | 241.5 | P | 219.6 | P | --- |
| 267184_at | 219.8 | P | 269.8 | P | 241.4 | P | 332.6 | P | ASHH3 |
| 267185_at | 849.4 | P | 990.6 | P | 790.3 | P | 911.7 | P | OEP37 |
| 267186_at | 249.6 | P | 336.9 | P | 267.6 | P | 345.2 | P | --- |
| 267187_s_at | 2228.7 | P | 2364.3 | P | 2510.5 | P | 2370 | P | MTHFR2 |
| 267188_at | 2314.9 | P | 2415.8 | P | 2530.3 | P | 2513.5 | P | COS1 |
| 267189_at | 519 | P | 699.1 | P | 494.3 | P | 605.6 | P | MAP2A |
| 267190_at | 9.6 | A | 24.5 | M | 31.3 | A | 28.8 | P | NMT2 |
| 267191_at | 20.6 | A | 3.2 | A | 25 | A | 5.9 | A | MLO15 |
| 267192_at | 109.6 | A | 110.5 | P | 137.5 | P | 163.9 | P | --- |
| 267193_at | 21.3 | A | 22.7 | A | 18.8 | A | 19 | A | --- |
| 267194_s_at | 268.2 | P | 337.9 | P | 210.1 | P | 264.8 | P | ARPC1A |
| 267195_at | 13.2 | A | 26.7 | A | 37.6 | A | 17.8 | A | --- |
| 267196_at | 8156.7 | P | 8785.7 | P | 9902.6 | P | 8484.7 | P | VAR2 |
| 267197_at | 107.9 | P | 80.7 | P | 114.5 | A | 115.4 | P | --- |
| 267198_at | 8.9 | A | 9.2 | A | 27 | A | 1.7 | A | --- |
| 267199_at | 560.1 | P | 600.6 | P | 518.3 | P | 509.8 | P | --- |
| 267200_at | 102.4 | M | 104 | A | 70.8 | A | 120.4 | M | --- |
| 267201_at | 38.5 | A | 127.5 | P | 66.8 | P | 92 | P | --- |
| 267202_s_at | 1.9 | A | 30 | A | 8.1 | A | 2.8 | A | ORP1A |
| 267203_at | 0.2 | A | 17.9 | A | 0.7 | A | 0.3 | A | --- |
| 267204_at | 4.9 | A | 32 | A | 23.4 | A | 37.7 | A | --- |
| 267205_at | 16.6 | A | 52.6 | A | 23.2 | A | 41.7 | P | --- |
| 267206_at | 335.2 | P | 34.3 | A | 137.3 | P | 25.6 | A | --- |
| 267207_at | 63.3 | P | 56.2 | P | 56.2 | M | 59.3 | P | --- |
| 267208_at | 1009.4 | P | 1119.9 | P | 1180.3 | P | 1215.3 | P | ASKdZeta |
| 267209_at | 553.7 | P | 503.7 | P | 432.5 | P | 568.9 | P | --- |
| 267210_at | 231.5 | P | 275.6 | P | 280.1 | P | 282.4 | P | ATCOQ3 |
| 267211_at | 1162.2 | P | 1075.7 | P | 1387.1 | P | 1107.8 | P | --- |
| 267212_at | 3145 | P | 3832.6 | P | 2788.5 | P | 3463.9 | P | --- |
| 267213_at | 6865.9 | P | 5810.1 | P | 7822 | P | 6806.3 | P | --- |
| 267214_at | 4031.9 | P | 3540.4 | P | 3849.5 | P | 3687.3 | P | --- |
| 267215_at | 3.6 | A | 3.3 | A | 2.8 | A | 2.8 | A | --- |
| 267216_at | 2.3 | A | 21.5 | A | 3.2 | A | 3.9 | A | --- |
| 267217_at | 18.4 | M | 3.1 | A | 14.9 | A | 10 | A | --- |
| 267218_at | 6.3 | A | 2.6 | A | 5.8 | A | 7 | A | --- |
| 267219_at | 107 | P | 183 | P | 105.6 | P | 179.2 | P | --- |
| 267220_at | 695.3 | P | 940.1 | P | 889.7 | P | 1016.9 | P | ISPD |
| 267221_at | 85.9 | P | 133.6 | P | 153.4 | P | 155.8 | P | STI |
| 267222_at | 32.8 | A | 30.1 | A | 42 | A | 44 | A | --- |
| 267223_at | 274.6 | P | 405.2 | P | 333.3 | P | 353.7 | P | AtITPK4 |
| 267224_at | 61.4 | P | 61.1 | P | 51.3 | A | 49.8 | A | --- |
| 267225_at | 26.5 | A | 10.5 | A | 3.3 | A | 4 | A | --- |
| 267226_at | 3.6 | A | 6.9 | A | 5.7 | A | 4.4 | A | --- |
| 267227_at | 49.7 | A | 15.1 | A | 65.1 | P | 8.1 | A | --- |
| 267228_at | 3.1 | A | 2.9 | A | 4.2 | A | 2.1 | A | --- |
| 267229_s_at | 477.6 | P | 286.7 | P | 330.5 | P | 280 | P | --- |
| 267230_at | 301.3 | P | 752 | P | 353.2 | P | 525.8 | P | ARL |
| 267231_at | 55.2 | A | 79.3 | P | 52.1 | A | 61.3 | A | --- |
| 267232_at | 42.8 | P | 89.1 | P | 43.2 | P | 80.5 | P | EDE1 |
| 267233_s_at | 4124 | P | 5040.2 | P | 4955.2 | P | 4719.9 | P | --- |
| 267234_x_at | 51.7 | A | 36.5 | A | 70.8 | A | 51.9 | A | --- |
| 267235_at | 396.2 | P | 453.8 | P | 494.7 | P | 486.2 | P | AT2G43945 |
| 267236_at | 3693.5 | P | 3678.1 | P | 3108.5 | P | 3500.8 | P | ATGDI1 |
| 267237_s_at | 601.5 | P | 633.3 | P | 641.3 | P | 595.3 | P | --- |
| 267238_at | 188.2 | P | 206.5 | P | 190.9 | P | 166.1 | P | --- |
| 267239_at | 2819.4 | P | 2788.7 | P | 2066.8 | P | 2669.6 | P | --- |
| 267240_at | 15 | A | 18.4 | A | 30.2 | A | 19.1 | A | --- |
| 267241_at | 13.2 | A | 20.4 | A | 40.2 | A | 16.8 | A | --- |

| | | | | | | | | | |
|---|---|---|---|---|---|---|---|---|---|
| 267242_at | 1.2 | A | 1.3 | A | 0.7 | A | 1.3 | A | --- |
| 267243_at | 5 | A | 14.4 | A | 1.6 | A | 5.6 | A | --- |
| 267244_at | 12.5 | A | 28.7 | A | 19.3 | A | 20.3 | A | --- |
| 267245_at | 0.3 | A | 1 | A | 5.5 | A | 0.3 | A | --- |
| 267246_at | 608.9 | P | 549.3 | P | 420.1 | P | 405.9 | P | WRKY25 |
| 267247_at | 1089 | P | 1213.1 | P | 1015.7 | P | 1173.7 | P | --- |
| 267248_at | 66.5 | P | 75.1 | P | 59.4 | P | 82.4 | P | --- |
| 267249_at | 21.6 | A | 43.8 | A | 44.4 | A | 39.1 | A | --- |
| 267250_at | 2.8 | A | 5 | A | 3.5 | A | 9.5 | A | --- |
| 267251_at | 2595.6 | P | 2307.3 | P | 2759.8 | P | 2462.8 | P | --- |
| 267252_at | 3.2 | A | 39 | P | 71.1 | A | 60.9 | P | --- |
| 267253_at | 2.7 | A | 7.2 | A | 17 | A | 3 | A | --- |
| 267254_at | 24.8 | A | 47.8 | A | 40.4 | A | 54.5 | A | SNRK2.9 |
| 267255_at | 22.5 | A | 13.7 | A | 39 | A | 36.6 | A | --- |
| 267256_s_at | 1585.6 | P | 714 | P | 1631.5 | P | 903.9 | P | scpl10 |
| 267257_at | 475.8 | P | 378.1 | P | 498.3 | P | 383.8 | P | --- |
| 267258_at | 223.2 | P | 158 | P | 199.4 | P | 169.4 | P | --- |
| 267259_at | 6725.1 | P | 5725.1 | P | 7397 | P | 6057.9 | P | --- |
| 267260_at | 367.6 | P | 443.7 | P | 412.4 | P | 465.7 | P | AGP17 |
| 267261_at | 3517.6 | P | 5694 | P | 3236.5 | P | 5192.1 | P | --- |
| 267262_at | 1129.6 | P | 1730.5 | P | 937.5 | P | 1229.4 | P | SNG1 |
| 267263_at | 7 | A | 4.5 | A | 3.4 | A | 3.2 | A | --- |
| 267264_at | 369 | P | 284.5 | P | 243.4 | P | 299.9 | P | SCPL11 |
| 267265_at | 390.4 | P | 462.4 | P | 413 | P | 399.8 | P | SCPL13 |
| 267266_at | 3157 | P | 1074.4 | P | 1687.3 | P | 1161.8 | P | NRAMP3 |
| 267267_at | 4.7 | A | 13.7 | A | 27.5 | P | 4.2 | A | CYP71B9 |
| 267268_at | 831.1 | P | 858.3 | P | 893.2 | P | 898.2 | P | --- |
| 267269_at | 601.3 | P | 686.4 | P | 512.3 | P | 644.1 | P | CAND1 |
| 267270_at | 1.3 | A | 35 | A | 13.6 | A | 2.2 | A | --- |
| 267271_at | 72 | A | 88.1 | P | 115.2 | P | 102.7 | M | ATHB21 |
| 267272_at | 3.5 | A | 22 | A | 29 | A | 4.1 | A | --- |
| 267273_at | 16.6 | A | 40.5 | P | 16.1 | A | 23.6 | P | --- |
| 267274_at | 459.8 | P | 392.1 | P | 473.2 | P | 383.2 | P | --- |
| 267275_at | 3.4 | A | 5.3 | A | 7.9 | A | 4.9 | A | ATCHX13 |
| 267276_at | 1.3 | A | 1.8 | A | 2.3 | A | 1.3 | A | ASL5 |
| 267277_at | 1031.5 | P | 1043.2 | P | 1166.6 | P | 1020.4 | P | --- |
| 267278_at | 313.7 | P | 384.6 | P | 290.2 | P | 304.6 | P | --- |
| 267279_at | 182.6 | P | 257.4 | P | 160.8 | P | 222.3 | P | --- |
| 267280_at | 1145 | P | 830.8 | P | 966.6 | P | 996.4 | P | TAG1 |
| 267281_at | 78.9 | P | 61.9 | P | 86.9 | P | 75.4 | P | --- |
| 267282_at | 86.3 | P | 86.4 | P | 51.9 | P | 81.6 | P | --- |
| 267283_at | 1.6 | A | 0.7 | A | 16.2 | A | 0.4 | A | --- |
| 267284_at | 123.7 | P | 151.8 | P | 130.5 | P | 139.1 | P | --- |
| 267285_at | 69.7 | A | 71.5 | A | 98 | A | 95.4 | A | --- |
| 267286_at | 6 | A | 18 | A | 67.1 | A | 4.8 | A | --- |
| 267287_at | 38.9 | P | 19.8 | A | 21.4 | A | 13.4 | A | sks16 |
| 267288_at | 330.7 | P | 385.4 | P | 310.9 | P | 362.1 | P | --- |
| 267289_at | 62.1 | P | 64.5 | P | 50.8 | A | 56.1 | P | --- |
| 267290_at | 335.3 | P | 357.2 | P | 331 | P | 330.6 | P | --- |
| 267291_at | 135.1 | P | 184.9 | P | 171.7 | P | 211.4 | P | --- |
| 267292_at | 29.1 | A | 19.6 | A | 54.1 | A | 25.9 | A | --- |
| 267293_at | 2000.7 | P | 2137.6 | P | 1304.8 | P | 1604.2 | P | TET8 |
| 267294_at | 2654.3 | P | 3689.8 | P | 3823.2 | P | 3815.1 | P | YCF37 |
| 267295_at | 3.4 | A | 37.1 | A | 8.1 | A | 8.2 | A | GGPS2 |
| 267296_at | 38.5 | A | 63.7 | A | 64.4 | A | 53.1 | A | LBD10 |
| 267297_at | 461.5 | P | 585.8 | P | 525.2 | P | 550.8 | P | --- |
| 267298_at | 714.7 | P | 442.4 | P | 587.8 | P | 448.2 | P | BLH4 |
| 267299_at | 651.5 | P | 552.6 | P | 564 | P | 556.7 | P | --- |
| 267300_at | 3052.9 | P | 1750.3 | P | 2274.3 | P | 1753.4 | P | --- |
| 267301_at | 2030.1 | P | 2026.8 | P | 1881.8 | P | 2070.5 | P | ATUBA1 |
| 267302_at | 483.5 | P | 572.4 | P | 417.2 | P | 500.2 | P | --- |
| 267303_at | 2.8 | A | 1.4 | A | 6.2 | A | 2.6 | A | --- |
| 267304_at | 159.4 | P | 140.9 | P | 140.8 | P | 143.8 | P | ZIP6 |
| 267305_at | 290.9 | P | 599.2 | P | 370 | P | 473.3 | P | ATKT1 |
| 267306_at | 1268.7 | P | 2151.7 | P | 1399.8 | P | 2035.6 | P | --- |
| 267307_at | 30.5 | A | 141.2 | P | 73.9 | P | 157.2 | P | LAC3 |
| 267308_at | 1263.8 | P | 1205.9 | P | 1175.9 | P | 1466 | P | --- |
| 267309_at | 500.5 | P | 340.5 | P | 415 | P | 354.1 | P | --- |
| 267310_at | 485.5 | P | 875.9 | P | 558.6 | P | 813.5 | P | AIR9 |
| 267311_at | 7 | A | 48.1 | A | 62.9 | A | 68.6 | A | --- |
| 267312_at | 1543.4 | P | 1051.3 | P | 1523.2 | P | 1127.7 | P | --- |
| 267313_at | 5.7 | A | 2.7 | A | 7.1 | A | 12 | A | --- |
| 267314_at | 309 | P | 223.3 | P | 292.6 | P | 242.8 | P | --- |
| 267315_at | 499.9 | P | 597.2 | P | 579.8 | P | 568.2 | P | NF-YA4 |
| 267316_at | 218.2 | P | 319.2 | P | 295.1 | P | 322.3 | P | PHB |
| 267317_at | 4.6 | A | 23.2 | A | 5.3 | A | 3.6 | A | --- |
| 267318_at | 669.8 | P | 721.4 | P | 637.2 | P | 703.7 | P | FAH1 |
| 267319_at | 1192.1 | P | 1197.7 | P | 1046 | P | 1246.1 | P | ATMRP2 |
| 267320_at | 1.3 | A | 6.4 | A | 3.7 | A | 9.2 | A | --- |
| 267321_at | 30 | A | 16.6 | A | 10.9 | A | 9.3 | A | --- |
| 267322_at | 31.2 | A | 34 | A | 59.1 | P | 15.9 | A | --- |
| 267323_at | 338.4 | P | 434.6 | P | 363.9 | P | 392.3 | P | --- |
| 267324_at | 12.8 | A | 1.2 | A | 2.2 | A | 4.1 | A | --- |
| 267325_at | 1.5 | A | 1.6 | A | 3.4 | A | 22.4 | A | --- |
| 267326_at | 68.2 | P | 68.5 | P | 71.3 | P | 60.7 | P | --- |
| 267327_at | 6.8 | A | 1.5 | A | 6.2 | A | 6.1 | A | --- |
| 267328_at | 4.7 | A | 2.5 | A | 6.2 | A | 2.4 | A | --- |
| 267329_at | 14.2 | A | 20 | A | 5.3 | A | 24.5 | A | --- |
| 267330_at | 470 | P | 583.9 | P | 291.2 | P | 506.4 | P | --- |
| 267331_at | 108.7 | P | 137.1 | P | 136.2 | P | 134.3 | P | --- |
| 267332_at | 3 | A | 1.2 | A | 1.7 | A | 1.8 | A | --- |
| 267333_at | 263.4 | P | 298 | P | 321.9 | P | 289.7 | P | --- |
| 267334_at | 0.8 | A | 1.1 | A | 11 | A | 2.2 | A | --- |
| 267335_s_at | 166.9 | P | 330.5 | P | 244.3 | P | 207.4 | P | --- |
| 267336_at | 1307 | P | 1278.7 | P | 1297.6 | P | 1423.6 | P | --- |
| 267337_at | 171.2 | P | 349.7 | P | 187.7 | P | 263.2 | P | --- |
| 267338_at | 2375.5 | P | 2090.8 | P | 2221.1 | P | 1881.8 | P | EIF2 |
| 267339_at | 166.6 | P | 281.5 | P | 305.5 | P | 334.6 | P | --- |
| 267340_at | 1622.8 | P | 1459 | P | 1840.6 | P | 1587.6 | P | ACD11 |
| 267341_at | 354.2 | P | 320.9 | P | 376.7 | P | 288.3 | P | --- |
| 267342_at | 826.7 | P | 741 | P | 927.7 | P | 718.4 | P | COX10 |
| 267343_at | 47.2 | P | 32.5 | P | 39.5 | P | 53.5 | P | --- |
| 267344_at | 190.8 | P | 138.1 | P | 197.8 | P | 156.3 | P | --- |
| 267345_at | 1.9 | A | 2.7 | A | 11.9 | A | 4.9 | A | --- |
| 267346_at | 487.3 | P | 566.7 | P | 414.7 | P | 514.2 | P | COI1 |
| 267347_at | 262.4 | P | 230.9 | P | 199.1 | P | 242.3 | P | --- |
| 267348_at | 849.8 | P | 759.7 | P | 788 | P | 1171.7 | P | --- |
| 267349_at | 379.9 | P | 265.2 | P | 409.9 | P | 300.7 | P | --- |
| 267350_at | 14.3 | A | 12.7 | A | 22.6 | A | 31.1 | P | NRPD1B |

| | | | | | | | | | |
|---|---|---|---|---|---|---|---|---|---|
| 267351_at | 95.6 | P | 111.8 | P | 89.8 | P | 175.5 | P | NRPD1B |
| 267352_at | 11.8 | A | 13.2 | A | 4.8 | A | 4.5 | A | --- |
| 267353_at | 285.4 | P | 369.8 | P | 371.7 | P | 452.7 | P | --- |
| 267354_at | 54.5 | A | 134.7 | P | 80 | P | 134.5 | P | MYB25 |
| 267355_at | 925.6 | P | 1264.6 | P | 929.5 | P | 1321.1 | P | --- |
| 267356_at | 571.3 | P | 784 | P | 617.6 | P | 757.5 | P | ISA1 |
| 267357_at | 255.8 | P | 266.8 | P | 159.9 | P | 211.8 | P | HSPRO2 |
| 267358_at | 388.2 | P | 593.4 | P | 365.6 | P | 537.3 | P | PROT1 |
| 267359_at | 99.4 | P | 116.8 | P | 176.8 | P | 173.7 | P | --- |
| 267360_at | 934 | P | 1066.4 | P | 1020.5 | P | 1124.3 | P | --- |
| 267361_at | 236.3 | P | 394.5 | P | 216.9 | P | 320.1 | P | --- |
| 267362_at | 210.7 | P | 165 | P | 218 | P | 178 | P | --- |
| 267363_at | 567.8 | P | 808.7 | P | 513.8 | P | 664.3 | P | --- |
| 267364_at | 2835.4 | P | 2166.6 | P | 1962.5 | P | 2549.2 | P | ELF4 |
| 267365_at | 771.9 | P | 726.4 | P | 655.8 | P | 742 | P | --- |
| 267366_at | 3570.1 | P | 2792.2 | P | 2878.7 | P | 3087.1 | P | --- |
| 267367_at | 2283.7 | P | 2129.5 | P | 1928.4 | P | 2131.2 | P | --- |
| 267368_at | 3981.9 | P | 2851.1 | P | 4024.9 | P | 3214.6 | P | ATCS |
| 267369_at | 119.3 | P | 113.5 | P | 128.5 | P | 128.1 | P | --- |
| 267370_at | 908 | P | 888.7 | P | 915.8 | P | 805.9 | P | --- |
| 267371_at | 191.2 | P | 151.5 | P | 234.4 | P | 137 | P | --- |
| 267372_at | 14.7 | A | 11.1 | A | 20.6 | A | 4.9 | A | ARSK1 |
| 267373_at | 487.4 | P | 539.7 | P | 480.2 | P | 563.4 | P | CID7 |
| 267374_at | 1518.9 | P | 1186.4 | P | 1409.9 | P | 1361.2 | P | --- |
| 267375_at | 1044.4 | P | 964.4 | P | 823.1 | P | 849.8 | P | GP ALPHA 1 |
| 267376_at | 745.7 | P | 1018.6 | P | 1013.2 | P | 1016.1 | P | ER |
| 267377_at | 2184.2 | P | 2389.1 | P | 2393.7 | P | 2670.8 | P | KCS10 |
| 267378_at | 405 | P | 450.2 | P | 360.3 | P | 431.8 | P | --- |
| 267379_at | 1192.3 | P | 1441.8 | P | 1165.6 | P | 1274.7 | P | --- |
| 267380_at | 434.6 | P | 364.7 | P | 486.5 | P | 453.2 | P | CYP711A1 |
| 267381_at | 505 | P | 478.1 | P | 599.8 | P | 424.3 | P | --- |
| 267382_at | 157.1 | P | 254.1 | P | 195.5 | P | 275.4 | P | --- |
| 267383_at | 657 | P | 656.4 | P | 547.5 | P | 632 | P | --- |
| 267384_at | 16.7 | A | 4.6 | A | 6.2 | A | 2.2 | A | --- |
| 267385_at | 25.7 | A | 47.3 | P | 24 | A | 45.4 | A | --- |
| 267386_at | 80.9 | P | 45 | P | 55.2 | A | 61.9 | M | --- |
| 267387_at | 119.2 | P | 206.8 | P | 163.5 | P | 185.7 | P | --- |
| 267388_at | 210.5 | P | 248.6 | P | 180.1 | P | 231.4 | P | BGLU15 |
| 267389_at | 83 | P | 32.6 | A | 294.2 | P | 36 | A | BGLU28 |
| 267390_at | 15.7 | A | 23 | A | 35.7 | P | 21.8 | A | BGLU29 |
| 267391_at | 26.9 | A | 32.7 | A | 3.8 | A | 1.8 | A | BGLU17 |
| 267392_at | 2808 | P | 3245.8 | P | 2789.2 | P | 3179.6 | P | PEN2 |
| 267393_at | 229.2 | P | 204 | P | 196.5 | P | 159 | P | --- |
| 267394_s_at | 0.6 | A | 1.4 | A | 1.4 | A | 0.8 | A | AtGH9B9 |
| 267395_at | 1.4 | A | 19.1 | A | 8.5 | A | 2.4 | A | --- |
| 267396_at | 422.4 | P | 386 | P | 388.1 | P | 334.2 | P | --- |
| 267397_at | 156.7 | P | 127.8 | P | 143.4 | P | 126.9 | P | --- |
| 267398_at | 18.1 | A | 1.3 | A | 12.9 | A | 0.4 | A | AtGH9B11 |
| 267399_at | 1.3 | A | 1 | A | 1.4 | A | 1.5 | A | --- |
| 267400_at | 1112.4 | P | 1257 | P | 1089.6 | P | 1167.7 | P | --- |
| 267401_at | 158.3 | P | 173.5 | P | 150.3 | P | 193.6 | P | --- |
| 267402_at | 117.9 | P | 197.2 | P | 106.3 | P | 196.8 | P | IQD6 |
| 267403_at | 21.8 | P | 2.1 | A | 2.5 | A | 0.5 | A | AGL33 |
| 267404_at | 26.4 | A | 56.2 | A | 48.7 | A | 37.8 | A | atnudt22 |
| 267405_at | 1413.4 | P | 1340.4 | P | 1027.4 | P | 1214.3 | P | CUTA |
| 267406_at | 173.4 | P | 208 | P | 237.4 | P | 191 | P | MEE22 |
| 267407_at | 4.5 | A | 48.9 | A | 13.8 | A | 34.4 | A | MEE27 |
| 267408_at | 3.1 | A | 4.2 | A | 8.9 | A | 4.3 | A | --- |
| 267409_at | 11.8 | A | 13 | A | 4.1 | A | 6.4 | A | --- |
| 267410_at | 99.3 | P | 82.5 | P | 107.6 | P | 115.2 | P | EDA18 |
| 267411_at | 688.9 | P | 357.5 | P | 440.4 | P | 314.4 | P | --- |
| 267412_at | 49.6 | A | 87 | P | 54.4 | A | 73.4 | A | --- |
| 267413_at | 140.6 | P | 124.3 | P | 172.6 | P | 101.8 | P | CAT5 |
| 267414_at | 37.3 | A | 75.3 | P | 55.2 | A | 41.9 | P | MEE23 |
| 267415_at | 585.2 | P | 516.1 | P | 540.8 | P | 511.5 | P | --- |
| 267416_at | 35.4 | M | 43.8 | A | 27.5 | A | 29.9 | M | SETH1 |
| 267417_at | 31 | A | 17.3 | A | 59.4 | A | 43.9 | A | --- |
| 267418_at | 41.8 | M | 38.3 | A | 31.1 | A | 19.2 | A | --- |
| 267419_at | 687.1 | P | 798.4 | P | 702.8 | P | 684.5 | P | ATO1 |
| 267420_at | 95.6 | P | 100.9 | P | 123.8 | P | 65.8 | P | --- |
| 267421_at | 1764.9 | P | 1135.7 | P | 1976.1 | P | 1351.8 | P | --- |
| 267422_at | 1197.6 | P | 1313.8 | P | 1017.6 | P | 1320.2 | P | --- |
| 267423_at | 533 | P | 475.2 | P | 451.1 | P | 527.2 | P | KUP11 |
| 267424_at | 4 | A | 2.9 | A | 4.3 | A | 0.6 | A | --- |
| 267425_at | 876.1 | P | 442.2 | P | 677.7 | P | 488.9 | P | --- |
| 267426_at | 4.1 | A | 4.2 | A | 8.5 | A | 4 | A | --- |
| 267427_at | 23.3 | A | 19.1 | A | 16.4 | A | 26.7 | A | WRKY35 |
| 267428_at | 217.6 | P | 250.4 | P | 164.4 | P | 293.2 | P | --- |
| 267429_at | 33.7 | A | 39.9 | A | 34 | A | 65.6 | A | MEE25 |
| 267430_at | 4925.8 | P | 3793.5 | P | 3956.5 | P | 3622 | P | EDA3 |
| 267431_at | 8.6 | A | 5.2 | A | 41.4 | A | 13.9 | A | MEE26 |
| 267432_at | 440.9 | P | 975.4 | P | 577.6 | P | 925.8 | P | --- |
| 267433_at | 308.9 | P | 309.7 | P | 321.8 | P | 258.1 | P | PEX10 |
| 267434_at | 163.3 | P | 253.9 | P | 218.6 | P | 317.8 | P | AT3BETAHSD/D2 |
| 267435_at | 6373.4 | P | 6595.6 | P | 5687.6 | P | 6633.2 | P | --- |
| 267436_at | 32.3 | A | 129.8 | P | 47.5 | A | 47.5 | A | FRK1 |
| 267437_at | 20 | A | 6 | A | 16.9 | A | 19.7 | A | --- |
| 267438_at | 19.7 | A | 8.6 | A | 28.6 | A | 23.8 | A | --- |
| 267439_at | 23.7 | A | 2.3 | A | 6 | A | 7.1 | A | --- |
| 267440_at | 5.8 | A | 2.3 | A | 2.1 | A | 22.2 | A | SHT |
| 267441_at | 3.3 | A | 2.8 | A | 4.2 | A | 1.9 | A | --- |
| 267442_at | 521 | P | 510.3 | P | 718 | P | 647 | P | --- |
| 267443_at | 5.6 | A | 3 | A | 20.9 | A | 7.3 | A | --- |
| 267444_at | 299.2 | P | 353.6 | P | 296.5 | P | 348.4 | P | --- |
| 267445_at | 91.7 | P | 55.7 | P | 57 | A | 90.3 | P | --- |
| 267446_s_at | 644.2 | P | 568.4 | P | 620 | P | 588.2 | P | --- |
| 267447_at | 5.2 | A | 5.5 | A | 2.9 | A | 17.9 | A | ArRABA1h |
| 267448_at | 165.8 | P | 185.3 | P | 212.1 | P | 218.1 | P | --- |
| 267449_at | 1.4 | A | 8.1 | A | 3.5 | A | 1.1 | A | --- |
| 267450_at | 30.8 | A | 7.7 | A | 3.8 | A | 20.5 | A | --- |
| 267451_at | 97.4 | P | 56.7 | P | 131.8 | P | 73.8 | P | --- |
| 267452_at | 274.9 | P | 309.8 | P | 372.2 | P | 398.4 | P | ETT |
| 267453_at | 45.5 | P | 22.7 | P | 61.9 | P | 43.5 | P | HB-3 |
| 267454_at | 660.9 | P | 872.6 | P | 708 | P | 823 | P | --- |
| 267455_at | 69.1 | P | 80.1 | P | 114.3 | P | 91.5 | P | --- |
| 267456_at | 303.7 | P | 953.8 | P | 678.1 | P | 1228 | P | PHO2 |
| 267457_at | 30.2 | A | 33.3 | A | 15 | A | 31.6 | A | AGP30 |
| 267458_at | 9.5 | A | 42.8 | P | 57.1 | M | 26.2 | A | MLO5 |
| 267459_at | 896.5 | P | 1195 | P | 846 | P | 1771.9 | P | --- |

| | | | | | | | | | |
|---|---|---|---|---|---|---|---|---|---|
| 267460_at | 81.3 | P | 93.3 | P | 106 | P | 80.9 | P | SPL3 |
| 267461_at | 1644.4 | P | 1346.2 | P | 1396.6 | P | 1144.1 | P | --- |
| 267462_at | 94.9 | P | 146.9 | P | 131.1 | P | 116.9 | P | --- |
| 267463_at | 630.4 | P | 726.4 | P | 614.4 | P | 713.7 | P | --- |
| 267464_at | 27.6 | P | 33.4 | A | 33.6 | A | 19.2 | A | --- |
| 267465_at | 160.3 | P | 228.7 | P | 232.8 | P | 265 | P | SLP3 |
| 267466_at | 3.2 | A | 16.4 | A | 31.2 | A | 5.3 | A | --- |
| 267467_at | 117.9 | P | 123.4 | P | 146.2 | P | 123.2 | P | DRIP2 |
| 267468_at | 441.6 | P | 428.5 | P | 397 | P | 395.4 | P | --- |
| 267469_at | 324.5 | P | 397.9 | P | 374.3 | P | 358.5 | P | WRKY21 |
| 267470_at | 2460.5 | P | 2102.3 | P | 2621.5 | P | 2394.6 | P | C4H |
| 267471_at | 1267.1 | P | 1491.9 | P | 1437.7 | P | 1503.3 | P | FC2 |
| 267472_at | 886.1 | P | 390.6 | P | 527.7 | P | 548.3 | P | ARPN |
| 267473_at | 93.3 | P | 77.1 | P | 94.9 | P | 84.1 | P | --- |
| 267474_at | 279.2 | P | 415.7 | P | 364.7 | P | 420.6 | P | WHY3 |
| 267475_at | 200.9 | P | 190.2 | P | 157 | P | 201.4 | P | --- |
| 267476_at | 13.5 | A | 25.3 | A | 13.4 | A | 12.9 | A | --- |
| 267477_at | 1764 | P | 918.7 | P | 1145.2 | P | 878.2 | P | PLPB |
| 267478_at | 6.1 | A | 7.1 | A | 4.7 | A | 27.2 | A | --- |
| 267479_at | 4.6 | A | 1.1 | A | 5.6 | A | 7.5 | A | --- |
| 267480_at | 2.1 | A | 3 | A | 24.8 | A | 11.9 | A | --- |
| 267481_at | 229.3 | P | 263.5 | P | 213.3 | P | 296.2 | P | --- |
| 267482_s_at | 119.5 | P | 104.7 | P | 129 | P | 110.1 | P | --- |
| 267483_at | 398.1 | P | 501.1 | P | 412.1 | P | 348.6 | P | UTR1 |
| 267484_at | 7011.2 | P | 6524 | P | 4437.7 | P | 6466.1 | P | ATUBC2 |
| 267485_at | 39.4 | P | 55.3 | P | 61 | P | 56 | P | MYB88 |
| 267486_at | 1090.8 | P | 1183.1 | P | 948.2 | P | 1136.7 | P | APK2B |
| 267487_at | 2.2 | A | 4.2 | A | 2.1 | A | 5.3 | A | --- |
| 267488_at | 294.5 | P | 361.6 | P | 384 | P | 354.6 | P | HMA4 |
| 267489_s_at | 367.7 | P | 419.2 | P | 295.4 | P | 413.2 | P | --- |
| 267490_at | 204.1 | P | 207.9 | P | 162.8 | P | 197.4 | P | --- |
| 267491_at | 2.9 | A | 4.2 | A | 8.4 | A | 20.7 | A | --- |
| 267492_at | 2262.4 | P | 2617.3 | P | 2392.3 | P | 2669.6 | P | --- |
| 267493_at | 21.3 | A | 31.3 | A | 25.1 | A | 11.2 | A | OFP2 |
| 267494_at | 1593.3 | P | 1666 | P | 1589.5 | P | 1657.4 | P | KIS |
| 267495_at | 41.9 | P | 48.8 | P | 69 | P | 28.5 | A | ETC2 |
| 267496_at | 883.2 | P | 363.9 | P | 301.7 | P | 361.7 | P | --- |
| 267497_at | 54.4 | P | 90.6 | P | 127.9 | P | 77 | P | --- |
| 267498_at | 452.6 | P | 495.2 | P | 277.5 | P | 417.7 | P | --- |
| 267499_at | 179.6 | P | 211.7 | P | 85.5 | P | 225.8 | P | --- |
| 267500_s_at | 210.3 | P | 350.4 | P | 228.8 | P | 308.9 | P | CYP704A1 |
| 267501_at | 401.6 | P | 439.2 | P | 373.6 | P | 379.3 | P | --- |
| 267502_at | 13.8 | A | 9.9 | A | 21.3 | A | 22.3 | A | CYP76C4 |
| 267503_at | 167.2 | P | 126.1 | P | 120.3 | P | 161.7 | P | --- |
| 267504_at | 263.8 | P | 328.1 | P | 256.5 | P | 289.4 | P | --- |
| 267505_at | 491.5 | P | 658.9 | P | 490.9 | P | 791.7 | P | CYP76C1 |
| 267506_at | 985.3 | P | 800.1 | P | 916.6 | P | 826.7 | P | --- |
| 267507_at | 1274.9 | P | 1198.1 | P | 1730.1 | P | 1380.9 | P | --- |
| 267508_at | 209.9 | P | 292.9 | P | 244.2 | P | 261.1 | P | --- |
| 267509_at | 31.9 | M | 112.2 | P | 36.1 | M | 96.2 | P | AGL20 |
| 267510_at | 1457.3 | P | 1357.4 | P | 1345.9 | P | 1139.8 | P | SAP18 |
| 267511_at | 427.6 | P | 714.8 | P | 508.5 | P | 642.5 | P | --- |
| 267512_at | 215 | P | 224.4 | P | 207.3 | P | 266.1 | P | SSE1 |
| 267513_at | 739.5 | P | 728.6 | P | 788.2 | P | 685.2 | P | --- |
| 267514_at | 159 | P | 169.8 | P | 176.4 | P | 180.5 | P | --- |
| 267515_at | 206.3 | P | 241.1 | P | 281.6 | P | 303.7 | P | --- |
| 267516_at | 270.5 | P | 900.1 | P | 365.1 | P | 609.7 | P | RPT2 |
| 267517_at | 1223.7 | P | 2215.5 | P | 1384.6 | P | 1985.3 | P | RPT2 |
| 267518_at | 465.2 | P | 636.4 | P | 244.7 | P | 549.3 | P | --- |
| 267519_at | 192.9 | P | 180.6 | P | 187.1 | P | 190.2 | P | HSI2 |
| 267520_at | 357.8 | P | 528.6 | P | 383.1 | P | 455.5 | P | --- |
| 267521_at | 25.5 | P | 43.7 | P | 44.9 | M | 60.5 | P | --- |
| 267522_at | 3.2 | A | 2.9 | A | 1.4 | A | 1.4 | A | --- |
| 267523_at | 771.5 | P | 479.2 | P | 599.2 | P | 455.1 | P | --- |
| 267524_at | 467.8 | P | 366.5 | P | 393.2 | P | 324.8 | P | --- |
| 267525_at | 36.7 | P | 13.8 | A | 24.2 | A | 44.6 | A | --- |
| 267526_at | 8963.7 | P | 9006.8 | P | 7448.2 | P | 8550.2 | P | PSBW |
| 267527_at | 4.5 | A | 13.8 | A | 22.5 | A | 15.8 | A | --- |
| 267528_at | 8.8 | A | 0.6 | A | 4.7 | A | 3.9 | A | AGL6 |
| 267529_at | 62.4 | A | 78.5 | P | 61.4 | A | 71 | P | AtAUR3 |
| 267530_at | 214.6 | P | 301.1 | P | 279.8 | P | 285.5 | P | --- |
| 267531_at | 8.2 | A | 3.8 | A | 11.9 | A | 15.4 | A | CPK14 |
| 267532_at | 175.5 | P | 179.7 | P | 129.4 | P | 207.5 | P | --- |
| 267533_at | 591.9 | P | 594.7 | P | 678.3 | P | 594.2 | P | ATNUDX23 |
| 267534_at | 667.7 | P | 656.8 | P | 695.5 | P | 699.9 | P | --- |
| 267535_at | 377.1 | P | 511.7 | P | 498.7 | P | 507.1 | P | ZFP8 |
| 267536_at | 296.4 | P | 343.7 | P | 237.1 | P | 297.6 | P | PLDBETA1 |
| 267537_at | 173.3 | P | 252 | P | 174.3 | P | 205.3 | P | GK-1 |
| 267538_at | 193.3 | P | 173.9 | P | 127.4 | P | 189.1 | P | --- |
| 267539_at | 428.7 | P | 522.4 | P | 608.3 | P | 539 | P | --- |
| 267540_at | 237.8 | P | 234 | P | 201.5 | P | 216.3 | P | --- |
| 267541_at | 13.6 | A | 20.6 | P | 34.5 | A | 21.4 | A | --- |
| 267542_at | 1 | A | 0.7 | A | 2.1 | A | 6.2 | A | --- |
| 267543_at | 759.3 | P | 846.2 | P | 870.5 | P | 860.2 | P | --- |
| 267544_at | 372.3 | P | 438.4 | P | 397.4 | P | 485.2 | P | CB5-B |
| 267545_at | 2782 | P | 2046 | P | 2038.7 | P | 2454.3 | P | GRP23 |
| 267546_at | 348.3 | P | 730.3 | P | 420.4 | P | 479.3 | P | AtRLP23 |
| 267547_at | 35.9 | A | 57 | P | 50.9 | A | 48.7 | A | ATVAMP725 |
| 267548_at | 6.3 | A | 21.8 | P | 28.8 | A | 23.2 | P | AtRLP22 |
| 267549_at | 159.6 | P | 262.9 | P | 206.5 | P | 286.5 | P | --- |
| 267550_at | 190.4 | P | 226 | P | 133.1 | P | 190.6 | P | AP4.3A |
| 267551_at | 45.8 | A | 24.2 | A | 34.7 | A | 33.2 | A | UBP1 |
| 267552_at | 30.5 | A | 18.7 | A | 7.6 | A | 20 | A | PAP13 |
| 267553_s_at | 2491.9 | P | 2745.5 | P | 2468 | P | 2803.4 | P | --- |
| 267554_at | 14.9 | A | 21.4 | A | 16 | A | 5.2 | A | --- |
| 267555_at | 273 | P | 281.2 | P | 454.7 | P | 388.7 | P | SUMO5 |
| 267556_at | 859.6 | P | 1209.3 | P | 830.4 | P | 1159.2 | P | BGAL9 |
| 267557_at | 149.3 | P | 156.9 | P | 160 | P | 182.5 | P | KRP4 |
| 267558_at | 1468.4 | P | 1341.1 | P | 1240.7 | P | 1299.1 | P | LUH |
| 267559_at | 322.1 | P | 75 | P | 135.8 | P | 176.5 | P | CYP76C2 |
| 267560_at | 63.6 | A | 52 | A | 58 | A | 16.3 | A | CYP76C3 |
| 267561_at | 315.6 | P | 351.5 | P | 315.7 | P | 372.9 | P | --- |
| 267562_at | 1324 | P | 1427.9 | P | 1826.6 | P | 1444.4 | P | --- |
| 267563_at | 6.5 | A | 15.6 | A | 13.9 | A | 14.4 | A | --- |
| 267564_at | 176.5 | P | 205.2 | P | 216.8 | P | 225.4 | P | --- |
| 267565_at | 46.7 | P | 18.4 | A | 31.3 | A | 7.3 | A | CYP71A12 |
| 267566_at | 5 | A | 36 | A | 4.5 | A | 10.7 | A | --- |
| 267567_at | 89.8 | P | 72.9 | P | 29.4 | A | 42.9 | P | CYP71A13 |
| 267568_at | 74.1 | P | 140.3 | P | 153.2 | P | 107.3 | P | --- |

| | | | | | | | | | |
|---|---|---|---|---|---|---|---|---|---|
| 267569_at | 144 | P | 142.2 | P | 121.8 | P | 112.1 | P | PSBP-2 |
| 267570_at | 177.1 | P | 155.7 | P | 153.4 | P | 147.1 | P | HVT1 |
| 267571_at | 9.5 | A | 25.2 | A | 14.9 | A | 2.3 | A | --- |
| 267572_at | 49.8 | P | 40.1 | A | 59.2 | P | 71.2 | P | --- |
| 267573_at | 47.7 | P | 63.7 | P | 51.9 | P | 68.4 | P | --- |
| 267574_at | 20.9 | A | 40 | P | 16.4 | A | 21.9 | A | --- |
| 267575_at | 49.4 | P | 35.9 | P | 53.3 | A | 28.7 | A | --- |
| 267576_at | 37.5 | A | 58.5 | A | 4.9 | A | 39.6 | P | --- |
| 267577_at | 460.4 | P | 552.3 | P | 445 | P | 509.1 | P | --- |
| 267578_at | 462.3 | P | 541.1 | P | 435.1 | P | 506.1 | P | --- |
| 267579_at | 22.3 | A | 8.6 | A | 11 | A | 3.6 | A | --- |
| 267580_at | 98.9 | P | 133.2 | P | 90.8 | P | 129.7 | P | --- |
| 267581_at | 354.7 | P | 357.5 | P | 306.2 | P | 372.6 | P | --- |
| 267582_at | 9.8 | A | 11.7 | A | 31.1 | A | 57.5 | A | --- |
| 267583_at | 478.5 | P | 492.7 | P | 506.6 | P | 556.8 | P | --- |
| 267584_at | 14.2 | A | 37.6 | A | 7.5 | A | 8 | A | --- |
| 267585_s_at | 4.2 | A | 7.8 | A | 3.8 | A | 15.7 | A | ACT9 |
| 267586_at | 465.3 | P | 1019.3 | P | 751.6 | P | 829.3 | P | --- |
| 267587_at | 269.1 | P | 308.2 | P | 350.4 | P | 283.2 | P | --- |
| 267588_at | 16.1 | A | 17.8 | A | 59.2 | A | 5.5 | A | --- |
| 267589_at | 5.4 | A | 3.8 | A | 1 | A | 1 | A | --- |
| 267590_at | 896.2 | P | 1060.2 | P | 901.6 | P | 1070.6 | P | ATEXPA4 |
| 267591_at | 292 | P | 174.3 | P | 192.3 | P | 251.6 | P | RTFL8 |
| 267592_at | 543.4 | P | 222.3 | P | 417.3 | P | 213.3 | P | --- |
| 267593_at | 11 | A | 12 | A | 18.4 | A | 0.6 | A | AtRLP25 |
| 267594_at | 23.9 | A | 33.1 | A | 53.8 | A | 3.9 | A | --- |
| 267595_at | 127 | P | 346.9 | P | 170.4 | P | 440.9 | P | AtGH9B8 |
| 267596_s_at | 373.4 | P | 512 | P | 365.6 | P | 462.5 | P | AtRLP26 |
| 267597_at | 176.4 | P | 48.6 | P | 91.5 | P | 52.3 | P | AtRLP24 |
| 267598_at | 6 | A | 7.1 | A | 2.9 | A | 3 | A | --- |
| 267599_at | 402.7 | P | 362.8 | P | 298.3 | P | 325.9 | P | --- |
| 267600_at | 6578.1 | P | 6005.5 | P | 7309.9 | P | 6248.1 | P | --- |
| 267601_at | 281.2 | P | 349.8 | P | 320.4 | P | 351.7 | P | --- |
| 267602_at | 232.4 | P | 264.4 | P | 247.5 | P | 252.3 | P | --- |
| 267603_at | 208.9 | P | 270.9 | P | 214.1 | P | 245.4 | P | ATZW10 |
| 267604_at | 158.5 | P | 172.4 | P | 194.3 | P | 188.8 | P | ZFN2 |
| 267605_at | 1711.4 | P | 1837.2 | P | 1531.8 | P | 1961 | P | ATPDIL2-3 |
| 267606_at | 354.2 | P | 378.4 | P | 427.1 | P | 433.6 | P | KCS11 |
| 267607_s_at | 3780 | P | 3709.1 | P | 3793.3 | P | 3705.8 | P | ATSEH |
| 267608_at | 666.2 | P | 786.3 | P | 684 | P | 845.5 | P | --- |
| 267609_at | 502.7 | P | 487.6 | P | 454.2 | P | 512.3 | P | --- |
| 267610_at | 196.1 | P | 171.1 | P | 103.2 | A | 199.6 | P | AKT1 |
| 267611_at | 447.8 | P | 597.1 | P | 517.9 | P | 538.6 | P | SPX2 |
| 267612_at | 909.4 | P | 1228.8 | P | 980.1 | P | 1257.5 | P | --- |
| 267613_at | 16 | A | 13.7 | A | 34 | A | 33 | P | PID2 |
| 267614_at | 43.8 | A | 47.4 | A | 70.7 | A | 37.8 | A | BAS1 |
| 267615_at | 1.4 | A | 1 | A | 1.8 | A | 1 | A | --- |
| 267616_at | 342.9 | P | 327.9 | P | 293.3 | P | 288.3 | P | --- |
| 267617_at | 2606.7 | P | 2918.4 | P | 2348.9 | P | 2483.6 | P | TED4 |
| 267618_at | 102.2 | A | 149.8 | P | 96.6 | A | 141.2 | P | CYCB1;4 |
| 267619_at | 477 | P | 602.9 | P | 543.5 | P | 599 | P | --- |
| 267620_at | 16.3 | A | 30.4 | A | 38.2 | A | 23.1 | A | --- |
| 267621_at | 180.5 | P | 61.2 | P | 112.1 | P | 88 | P | --- |
| 267622_at | 86.4 | P | 141.1 | P | 90.8 | P | 111 | P | --- |
| 267623_at | 167.1 | P | 158.9 | P | 171 | P | 151.5 | P | --- |
| 267624_at | 440.3 | P | 652.4 | P | 459.9 | P | 617 | P | BIK1 |
| 267625_at | 96 | P | 216.9 | P | 186.1 | P | 184.5 | P | --- |
| 267626_at | 48.2 | A | 32.1 | A | 20.8 | A | 36.4 | A | CYP712A1 |
| 267627_at | 174.7 | P | 114.6 | P | 105.2 | P | 126.2 | P | --- |
| 267628_at | 364 | P | 319 | P | 403.6 | P | 363 | P | --- |
| 267629_at | 166.1 | P | 170.1 | P | 203.3 | P | 207.2 | P | POLD2 |
| 267630_at | 828.7 | P | 1054.7 | P | 1058.5 | P | 1145.7 | P | --- |
| 267631_at | 35.3 | P | 20.5 | P | 45.4 | A | 24.7 | A | --- |
| 267632_at | 218.3 | P | 212.4 | P | 176.3 | P | 188.8 | P | --- |
| 267633_at | 3 | A | 3.2 | A | 6.3 | A | 19.6 | A | --- |
| 267634_at | 18.9 | A | 3.5 | A | 26.5 | A | 4.4 | A | --- |
| 267635_at | 2226.2 | P | 2603.1 | P | 2179.3 | P | 2415.6 | P | --- |
| 267636_at | 103.7 | P | 160.7 | P | 129.2 | P | 153.8 | P | --- |
| 267637_at | 267.4 | P | 322 | P | 232.2 | P | 331.4 | P | --- |
| 267638_at | 4147.4 | P | 4109.9 | P | 3653.6 | P | 3853.8 | P | OEP16-3 |
| 267639_at | 154.6 | P | 154.2 | P | 188 | P | 204.5 | P | SPL9 |
| 267640_at | 737.6 | P | 747.1 | P | 762.1 | P | 744.3 | P | COP1 |
| 267641_at | 32.1 | A | 27.8 | A | 19.6 | A | 41.7 | A | AGO6 |
| 267642_at | 439.7 | P | 497.4 | P | 532.4 | P | 522.8 | P | --- |
| 267643_at | 32.1 | A | 37.7 | A | 58.2 | A | 3.6 | A | RALFL17 |
| 267644_s_at | 169.4 | P | 140.5 | P | 179.8 | P | 159.6 | P | --- |
| 267645_at | 254.5 | P | 428.4 | P | 444.8 | P | 471.1 | P | BGLU33 |
| 267646_at | 108 | P | 65.9 | P | 83.8 | P | 36.5 | A | PHT5 |
| AFFX-Athal-25SrRNA_s_at | 9924.7 | P | 8239.9 | P | 10603.6 | P | 7421.1 | P | --- |
| AFFX-Athal-5SrRNA_at | 20.6 | P | 8.6 | A | 23.3 | A | 12.2 | A | --- |
| AFFX-Athal-Actin_3_f_at | 5363.3 | P | 6134.8 | P | 4481.5 | P | 5581.6 | P | ACT7 |
| AFFX-Athal-Actin_5_r_at | 544.9 | P | 687.8 | P | 698.2 | P | 586.2 | P | --- |
| AFFX-Athal-Actin_M_at | 4281.9 | P | 4955.5 | P | 4298.9 | P | 4938.8 | P | --- |
| AFFX-Athal-GAPDH_3_s_at | 7075 | P | 6658.8 | P | 6433.2 | P | 6834.8 | P | GAPC1 |
| AFFX-Athal-GAPDH_5_s_at | 6191.8 | P | 6025.2 | P | 5650.7 | P | 5683.3 | P | GAPC1 |
| AFFX-Athal-GAPDH_M_s_at | 9856.1 | P | 9381.4 | P | 10181 | P | 9977.9 | P | GAPC1 |
| AFFX-Athal-Ubq_3_f_at | 18487.5 | P | 17081 | P | 13713.9 | P | 18410.6 | P | --- |
| AFFX-Athal-Ubq_5_f_at | 6254 | P | 6831.6 | P | 5614.5 | P | 6494.5 | P | --- |
| AFFX-Athal-Ubq_M_f_at | 5782.9 | P | 6345.1 | P | 5110.4 | P | 6419.8 | P | --- |
| AFFX-BioB-3_at | 332.2 | P | 349.4 | P | 424.6 | P | 361.3 | P | --- |
| AFFX-BioB-5_at | 306.2 | P | 253.4 | P | 294.2 | P | 277.6 | P | --- |
| AFFX-BioB-M_at | 466.4 | P | 398.9 | P | 423.3 | P | 512.2 | P | --- |
| AFFX-BioC-3_at | 776.8 | P | 768.5 | P | 969.3 | P | 869.7 | P | --- |
| AFFX-BioC-5_at | 1028.4 | P | 903.7 | P | 1109 | P | 1040.3 | P | --- |
| AFFX-BioDn-3_at | 3093.1 | P | 2856.2 | P | 3526.4 | P | 3240.7 | P | --- |
| AFFX-BioDn-5_at | 1985.4 | P | 1739.8 | P | 2228.7 | P | 1944.8 | P | --- |
| AFFX-CreX-3_at | 12546.1 | P | 11820.6 | P | 15002.8 | P | 12700.9 | P | --- |
| AFFX-CreX-5_at | 11206.9 | P | 9162.2 | P | 10697.1 | P | 10342.9 | P | --- |
| AFFX-DapX-3_at | 1951.1 | P | 993.3 | P | 1205.2 | P | 1053 | P | --- |
| AFFX-DapX-5_at | 599.7 | P | 303.7 | P | 386.3 | P | 280.4 | P | --- |
| AFFX-DapX-M_at | 1203.3 | P | 575.8 | P | 749.9 | P | 657.7 | P | --- |
| AFFX-LysX-3_at | 240.2 | P | 127.1 | P | 122.4 | P | 104.1 | P | --- |
| AFFX-LysX-5_at | 58.8 | P | 57.9 | P | 57.1 | P | 40 | P | --- |
| AFFX-LysX-M_at | 106.3 | P | 84.4 | P | 129.6 | P | 57.9 | P | --- |
| AFFX-PheX-3_at | 258.2 | P | 155.8 | P | 187.5 | P | 124.4 | P | --- |
| AFFX-PheX-5_at | 143.2 | P | 77.2 | P | 90.9 | P | 81.2 | P | --- |
| AFFX-PheX-M_at | 174.8 | P | 92.5 | P | 120.4 | P | 84.2 | P | --- |
| AFFX-r2-At-Actin-3_s_at | 9653.8 | P | 10654.6 | P | 10782.8 | P | 11177.6 | P | --- |
| AFFX-r2-At-Actin-5_s_at | 5041.9 | P | 6655.2 | P | 5766.8 | P | 6724.5 | P | --- |

| | | | | | | | | | |
|---|---|---|---|---|---|---|---|---|---|
| AFFX-r2-At-Actin-M_s_at | 7083.2 | P | 8472.6 | P | 8076.9 | P | 8788.7 | P | --- |
| AFFX-r2-At-GAPDH-3_s_at | 12919 | P | 12176.9 | P | 14224.5 | P | 12683.2 | P | --- |
| AFFX-r2-At-GAPDH-5_s_at | 14832.2 | P | 13556.6 | P | 13590 | P | 14038.2 | P | --- |
| AFFX-r2-At-U12639_at | 4.5 | A | 16.9 | A | 28.1 | A | 2.4 | A | --- |
| AFFX-r2-At-Ubq-3_x_at | 15086.4 | P | 14047.2 | P | 13227.2 | P | 14958.2 | P | --- |
| AFFX-r2-At-Ubq-5_x_at | 12396.1 | P | 12630.1 | P | 12255.5 | P | 13293.1 | P | --- |
| AFFX-r2-Bs-dap-3_at | 1315.3 | P | 672.4 | P | 783.2 | P | 619.8 | P | --- |
| AFFX-r2-Bs-dap-5_at | 798.1 | P | 293.1 | P | 443.3 | P | 317.4 | P | --- |
| AFFX-r2-Bs-dap-M_at | 1069.9 | P | 566.3 | P | 630.8 | P | 556.4 | P | --- |
| AFFX-r2-Bs-lys-3_at | 155.3 | P | 99 | P | 126.1 | P | 113.1 | P | --- |
| AFFX-r2-Bs-lys-5_at | 102.6 | P | 59.5 | P | 68.5 | P | 59.7 | P | --- |
| AFFX-r2-Bs-lys-M_at | 72.8 | P | 60 | P | 63.7 | P | 50.1 | P | --- |
| AFFX-r2-Bs-phe-3_at | 227 | P | 132.9 | P | 190.2 | P | 133.8 | P | --- |
| AFFX-r2-Bs-phe-5_at | 190.1 | P | 95.8 | P | 111.3 | P | 86.3 | P | --- |
| AFFX-r2-Bs-phe-M_at | 199.4 | P | 103.3 | P | 141.9 | P | 111.8 | P | --- |
| AFFX-r2-Bs-thr-3_s_at | 667.2 | P | 325 | P | 491.6 | P | 374.2 | P | --- |
| AFFX-r2-Bs-thr-5_s_at | 255.9 | P | 129.1 | P | 151 | P | 119.8 | P | --- |
| AFFX-r2-Bs-thr-M_s_at | 557.6 | P | 284.5 | P | 338.7 | P | 307.4 | P | --- |
| AFFX-r2-Ec-bioB-3_at | 455 | P | 445.2 | P | 497.1 | P | 473.8 | P | --- |
| AFFX-r2-Ec-bioB-5_at | 458.9 | P | 426 | P | 279.9 | P | 449.3 | P | --- |
| AFFX-r2-Ec-bioB-M_at | 535.7 | P | 494.7 | P | 516.5 | P | 511.1 | P | --- |
| AFFX-r2-Ec-bioC-3_at | 1819.6 | P | 1309.9 | P | 1468.6 | P | 1343.3 | P | --- |
| AFFX-r2-Ec-bioC-5_at | 1318.6 | P | 1137.5 | P | 1300.7 | P | 1323.5 | P | --- |
| AFFX-r2-Ec-bioD-3_at | 5048.6 | P | 4820.3 | P | 4903.9 | P | 4968.4 | P | --- |
| AFFX-r2-Ec-bioD-5_at | 3798.5 | P | 3582.8 | P | 3501.4 | P | 3708 | P | --- |
| AFFX-r2-P1-cre-3_at | 19260.4 | P | 17572.4 | P | 16842.7 | P | 18376.6 | P | --- |
| AFFX-r2-P1-cre-5_at | 15549.7 | P | 13072.2 | P | 14978.4 | P | 15371.2 | P | --- |
| AFFX-ThrX-3_at | 585.6 | P | 290.4 | P | 360.7 | P | 305.7 | P | --- |
| AFFX-ThrX-5_at | 179.8 | P | 64.1 | P | 111.8 | P | 79.6 | P | --- |
| AFFX-ThrX-M_at | 300 | P | 164.8 | P | 220.8 | P | 155.2 | P | --- |
| AFFX-TrpnX-3_at | 3.9 | A | 8.4 | A | 2.4 | A | 3.4 | A | --- |
| AFFX-TrpnX-5_at | 23.4 | A | 3.6 | A | 7.4 | A | 20.5 | A | --- |
| AFFX-TrpnX-M_at | 0.8 | A | 0.5 | A | 0.8 | A | 0.6 | A | --- |
